# Local translation in perisynaptic astrocytic processes is specific and regulated by fear conditioning

**DOI:** 10.1101/2020.01.21.913970

**Authors:** Noémie Mazaré, Marc Oudart, Julien Moulard, Giselle Cheung, Romain Tortuyaux, Philippe Mailly, David Mazaud, Alexis-Pierre Bemelmans, Anne-Cécile Boulay, Corinne Blugeon, Laurent Jourdren, Stéphane Le Crom, Nathalie Rouach, Martine Cohen-Salmon

**Author notes:** The authors declare no conflicts of interest.

## Abstract

Local translation is a conserved molecular mechanism conferring cells the ability to quickly respond to local stimuli. It not only permits cells with complex morphology to bypass somatic protein synthesis and transport, but also contributes locally to the establishment of molecular and functional polarity. In the brain, local translation has been extensively studied in neurons and has only been recently reported in astrocytes, whose fine processes contact both blood vessels and synapses. Yet the specificity and regulation of astrocyte local translation remain unknown. Here, we studied hippocampal perisynaptic astrocytic processes (PAPs) and show that they contain all the machinery for translation. Using our recently refined polysome immunoprecipitation technique, we then characterized the pool of polysomal mRNAs in PAPs, referred to as the PAPome, and compared it to the one found in the whole astrocyte. We found that the PAPome encoded an unexpected molecular repertoire, mostly composed of cytoplasmic proteins and of proteins involved in iron homeostasis, translation, cell cycle and cytoskeleton. Among them, ezrin (Ezr), ferritin heavy chain 1 (Fth1) and 60S acidic ribosomal protein1 (Rplp1) were enriched in PAPs compared to perivascular astrocytic processes, indicating that local translation differs at these two interfaces. Remarkably, PAPs were also enriched in transcripts coding for proteins involved in learning and memory, such as ferritin (Ftl1 and Fth1), G1/S-specific cyclin-D2 (Ccnd2), E3 ubiquitin-protein ligase (Mdm2), Receptor of activated protein C kinase 1 (Gnb2l1) and Elongation factor 1-alpha 1 (Eef1a1). To address their regulation in a physiological context, we assessed their local translation after fear conditioning. We found alterations in their density and/or distribution in astrocytes as well as a drop in their translation specifically in PAPs. In all, our results reveal an unexpected molecular repertoire of hippocampal PAPs, which is regulated by local translation during learning and memory processes.

## Introduction

Astrocytes constitute the most abundant population of glial cells in the mammalian brain. They are morphologically complex cells, with many ramifications extending towards both blood vessels and neurons (1). Specific domains called endfeet contact the blood vessels and thus enable astrocytes to modulate important vascular functions, such as blood-brain barrier integrity, immunity (2) and cerebral blood flow (3). The perisynaptic astrocytic processes (PAPs) interact with both synapses and dendrites, and regulate synaptic transmission (4). Furthermore, PAPs can sense changes in the composition of the perisynaptic extracellular space produced by active neurotransmission (5). The PAPs prevent prolonged neuronal activation and excitotoxicity by clearing ions and neurotransmitters released by the synapse. Astroglial processes are indeed equipped with transporters and channels, such as glutamate transporters and inward rectifying potassium channels, which tightly control the level of perisynaptic glutamate (6) and potassium respectively (7). Perisynaptic astrocytic processes also release neuroactive factors such as ATP and D-serine through various pathways including connexin hemichannels (8) or vesicles (9) . Astrocytes also influence synaptic function by dynamically modulating their synaptic coverage (10, 11). Lastly, these cells eliminate weak synapses and actively refine neuronal circuits during development, and orchestrate synaptogenesis in the mature brain (12–14). Understanding how astrocytes control this wide variety of synaptic functions is crucial because aberrant communication between neurons and astrocytes is known to contribute to several brain diseases. Characterization of the underlying molecular mechanisms may not only reveal the brain’s fundamental workings but also facilitate the development of new therapeutic tools.

Specialized cell polarity is a hallmark of astrocytes, but the underlying mechanisms are still unknown. The evolutionarily conserved cellular strategies involved in functional polarization notably include compartmentalization of mRNAs in distal regions of the cytoplasm, and local translation for spatiotemporal targeting of protein delivery (15). In the brain, both processes have been described in oligodendrocytes (16) and have been extensively studied in neurons - the cytoplasmic processes of which may be more than 1000 times longer than the cell body (17) (18). It has been shown that local translation provides synapses with a rapid access to new proteins, and has a crucial role in synaptic function and plasticity (19, 20). Importantly, the impairment of local translation in neurons has been implicated in several neuropathological diseases, such as fragile-X syndrome (21), amyotrophic lateral sclerosis (ALS) (22, 23) and spinal muscular atrophy (24, 25).

We recently showed that local translation occurs in perivascular astrocyte processes (PvAP), and characterized the locally translated molecular repertoire (26). Two other studies have also highlighted out local translation in radial glia during brain development (27) and in PAPs from the adult cortex (28). It has been suggested that impaired local translation in astrocytes and oligodendrocytes is involved in ALS (29).

In the present study, we characterized local translation in PAPs in the dorsal hippocampus, a region of the brain involved in memory and learning. We first checked for the presence in PAPs of mRNAs, ribosomes, local translation, the endoplasmic reticulum (ER)-Golgi intermediate compartment (ERGIC), and elements of the Golgi apparatus that might be crucial for local translation and protein maturation. We then used our recently refined translating ribosome affinity purification (TRAP) protocol (30) to characterize polysomal mRNAs in PAPs, and notably the mRNAs present at higher levels in the processes than in the astrocyte as a whole. Lastly, we probed the physiological relevance of local translation in PAPs by characterizing changes in levels of memory-related, PAP-enriched polysomal mRNAs after contextual memory acquisition.

## Results

### Hippocampal PAPs contain protein synthesis and maturation organelles

To test for the existence of local translation in PAPs from the hippocampus, we first characterized the mRNA distribution in astrocytes. We performed a FISH experiment on adult mouse CA1 hippocampal sections in order to detect Slc1a2 mRNA, a transcript coding for the astrocyte-specific glutamate transporter GLT1. The samples were co-immunostained for the astrocyte-specific intermediate filament glial fibrillary acidic protein (GFAP) (31). As shown in **Fig. 1A**, Slc1a2 mRNA FISH dots appeared to be distributed throughout the astrocyte. We used our recently developed *AstroDot* ImageJ plugin to characterize the mRNA distribution by counting the number of FISH dots localized on GFAP-immunolabeled intermediate filament in the somata and the large and fine processes (31). Since Slc1a2 is specifically expressed by astrocytes in the CA1 region, we considered that mRNA FISH dots localized outside GFAP-immunolabeled processes could be attributed to PAPs, where GFAP is known to be less present (32). The estimated mean ± standard error (SEM) proportion of Slc1a2 mRNAs was 16.1 ± 0.7% in the astrocyte somata, 20.5 ± 1.0% in GFAP-positive large processes, 63.3±1.0% in GFAP-positive fine processes, and 19.8 ± 0.9% in GFAP-negative areas (n=68 cells) (**Fig. 1B**). Overall, these results indicate that Slc1a2 mRNAs are preferentially distributed in distal areas of astrocytes that probably include PAPs. We next sought to localize the ribosomes in astrocytes by studying the Aldh1l1:L10a-eGFP transgenic mouse line, which expresses the eGFP-tagged ribosomal protein Rpl10a specifically in astrocytes (33) (**Fig. 1C**), by looking for eGFP-labeled ribosomes in mouse hippocampal sections. To visualize synapses, the presynaptic VGluT1 protein and the postsynaptic Homer1 protein were co-immunolabeled. Astrocyte ribosomes were present in somata and distal processes, including PAPs (**Fig. 1C**). In parallel, we detected local translation events (ribopuromycylation of nascent chains) in hippocampal CA1 PAPs (34). Puromycin (PMY) is an aminoglycoside antibiotic that mimics charged tRNATyr and is incorporated into the ribosome A site. It induces premature translation termination by ribosome-catalyzed covalent incorporation into the nascent carboxy-terminal chain (35). Freshly prepared hippocampal sections from Aldh1l1:L10a-eGFP mice were incubated in the presence or absence (negative control, not shown) of PMY. The PMY was detected by immunofluorescence (**Fig. 1C**); it co-localized with eGFP, and was found to be proximal to VGluT1/Homer1 immunolabeling - indicating the presence of local translation in PAPs.

**Figure 1:**
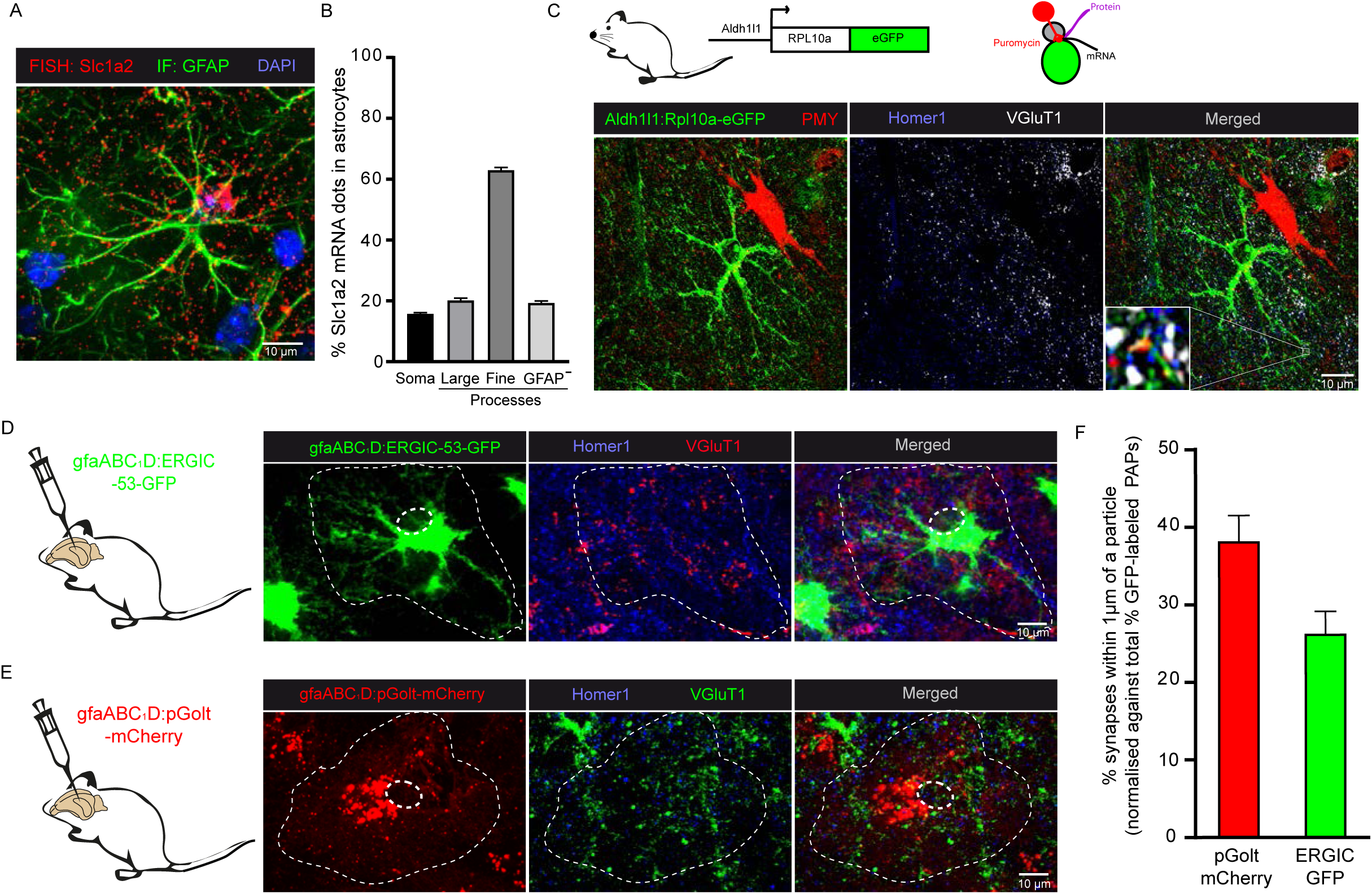
The perisynaptic processes of hippocampal astrocytes synthesize proteins and contain protein maturation organelles. **A.** Representative confocal microscopy image of the FISH detection of Slc1a2 mRNAs (red dots) in a dorsal hippocampal CA1 astrocyte immunolabeled for GFAP (green). Nuclei were stained with DAPI (blue). **B.** *AstroDot* analysis of the distribution of Slc1a2 mRNA in somata, large (>0.3 μm) and fine (<0.3 μm) GFAP-immunolabeled processes, and GFAP-negative processes (n=64) of CA1 astrocytes. **C.** Detection of CA1 astrocyte ribosomes and local translation events in PAPs by puromycylation (+PMY, with puromycin) in Aldh1l1:L10a-eGFP transgenic mice. Astrocyte ribosomes were immunolabeled for GFP (green) and PMY (red). Synapses were co-immunolabeled for VGluT1 (pre-synapse) (grey) and Homer1 (post-synapse) (blue). The white square is a magnified view (x10) of an astrocyte translation event near synapses. **D.** Visualization of ERGIC in CA1 astrocytes infected with a gfaABC_1_D-ERGIC-GFP-expressing AAV (green). Synapses were immunolabeled for VGluT1 (red) and Homer1 (blue) **E.** Visualization of Golgi particles in CA1 astrocytes infected with a gfaABC_1_D-pGolt-Cherry-expressing AAV (red). Synapses were immunolabeled for VGluT1 (green) and Homer1 (blue). In (**D**) and (**E**), the astrocyte domain and its nucleus are indicated by a dotted line. **F.** Quantification in (**D**) and (**E**) of the percentage of synapses within 1 μm of ERGIC or pGolt particles. n=20 cells from three mice per condition. The values were normalized against the total number of cytosolic eGFP-labeled PAPs within 1 μm of a synapse in mice infected with a gfaABC_1_D-eGFP expressing AAV in control experiments (see **Fig. S1**).

Perisynaptic astrocytic processes are extremely thin structures (<50 nm) (36), the subcellular organization of which has not been substantially addressed. Nevertheless, were a membrane protein such as GLT1 to be translated inside a PAP, it would have to pass through the ER and Golgi to be properly folded and functional at the plasma membrane. Here, we addressed the presence of these organelles in PAPs. We used an adeno-associated virus (AAV) bearing the gfaABC_1_D synthetic promoter (derived from *Gfap* (37)) to drive the expression of the Golgi tracker pGolt (38) and ERGIC-53 (an integral membrane protein localized in the ERGIC) (39). These proteins were tagged with mCherry and GFP, respectively. The AAVs were injected separately into the CA1 region of the dorsal hippocampus of adult mice (**Fig. 1D, E**). VGluT1 and Homer1 proteins were co-immunolabeled to visualize the synapses. As a control, adult mice were injected with an AAV driving the expression of eGFP alone in astrocytes (gfaABC_1_D-eGFP) (**Fig. S1A**). Co-immunofluorescent detection of GFAP with gfaABC_1_D-eGFP demonstrated the astrocytic specificity of infection and transgene expression (**Fig. S1B**). This control experiment also enabled us to quantify the number of VGluT1/Homer-immunolabeled synapses within 1 μm of a GFP-positive site, corresponding to the total amount of synapses contacted by PAPs. We then quantified VGlut1/Homer1-immunolabeled synapses within 1 μm of ERGIC-53-GFP and pGolt-mCherry-positive PAPs. ERGIC-53-GFP was detected homogeneously throughout astrocytes - suggesting the presence of a continuous ERGIC network - and was present in 27 ± 3% of PAPs (**Fig. 1D, F**). In contrast, pGolt-mCherry fluorescence was discontinuous, and positive puncta were detected in 38 ± 3% of the PAPs (**Fig. 1E, F**). Thus, it appeared that some PAPs contain ERGIC and elements of the Golgi apparatus. Taken as a whole, these results indicate that local translation and possibly post-translational modifications occur in hippocampal PAPs.

### Identification of the pool of polysomal mRNAs in PAPs (the PAPome) in the dorsal hippocampus

To further study local translation in PAPs, we next characterized the pool of polysomal mRNAs inside these processes. It has been shown that synaptogliosome preparations, consisting of apposed pre- and post-synaptic membranes, contain PAPs that remain attached to synaptic neuronal membranes (40). Here, we purified synaptogliosomes from the dorsal hippocampus of Aldh1l1:L10a-eGFP mice and characterized the various fractions of our preparation by Western blots (**Fig. 2A, B**). Compared with the first supernatant obtain from hippocampal homogenate (S1, whole astrocytes), the synaptogliosome fraction (P3) contained low levels of GFAP and high levels of the postsynaptic protein PSD95 and the cytosolic PAP protein Ezrin (41, 42). Rpl10a-eGFP was also detected in this fraction (**Fig. 2B**). These results indicated that the P3 fraction comprised ribosome-containing PAPs. We also checked for the presence of astrocyte ribosomes in the P3 fraction via the immunofluorescent detection of eGFP, VGluT1 and Homer1 (**Fig. 2C**). As we previously observed Slc1a2 mRNA in fine astrocyte processes (**Fig. 1A**), we used TRAP and qRT-PCR (see below) to test for the presence of Slc1a2 polysomal mRNAs in PAPs in the P3 fraction (**Fig. 2D**). Taken as a whole, these results indicate that PAPs co-purify with hippocampal synapses and contain polysomal mRNAs.

**Figure 2:**
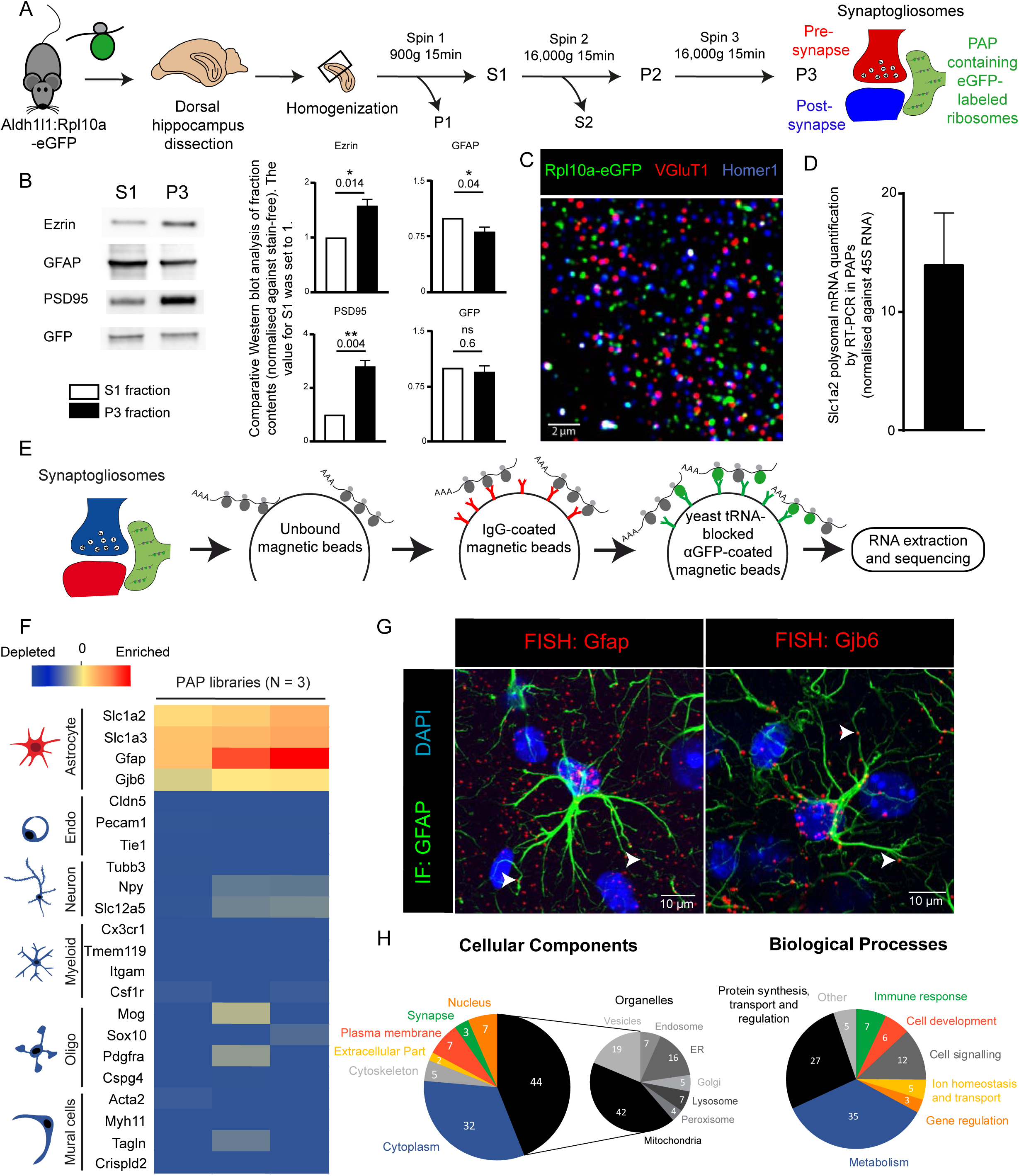
Identification of the PAPome, the pool of ribosome-bound mRNAs in PAPs from the dorsal hippocampus. **A.** Flowchart for the preparation of synaptogliosomes from the dorsal hippocampus of Aldh1l1:L10a-eGFP transgenic mice. S, supernatant; P, pellet. **B.** Western blot comparison of S1 and P3 fractions. *, p ≤ 0.05; **, p ≤ 0.01, and ns: not significant in a two-tailed, paired t-test. The data are presented as the mean ± SEM (n=4) **C.** A confocal immunofluorescence microscopy image of synaptogliosomes from an Aldh1l1:L10a-eGFP mouse. The ribosomes in PAPs were immunolabeled with eGFP (green). Pre- and post-synaptic areas were immunolabeled for VGluT1 (red) and Homer1 (blue), respectively. **D.** RT-PCR detection of Slc1a2 polysomal mRNAs in synaptogliosomes. The levels were normalized against 45S RNA. **E.** Flowchart for TRAP and analysis of PAP polysomal mRNAs extracted from dorsal hippocampal synaptogliosomes. The raw sequencing data are given in **Table S2**. **F.** Purity heat map of the RNA-seq data for a selection of mRNAs specific for each type of brain cell. The null level corresponds to mRNAs with 500 reads and 70% exon coverage (our selected detection threshold). Each column represents an independent cDNA library (n=3). **G.** Confocal microscopy images (FISH detection) of Gfap and Gjb6 mRNAs (red dots) in dorsal hippocampal astrocytes immunolabeled for GFAP (green). Nuclei were stained with DAPI. White arrowheads indicate FISH dots in fine astrocytic processes. **H.** Gene Ontology analysis of the PAPome.

We next aimed at purifying and characterizing polysomal mRNAs present in PAPs. To preclude the copurification of mRNAs from brain cells other than astrocytes (which would result from non-specific affinity of mRNAs to the TRAP column), we added pre-cleaning and blocking steps to the TRAP protocol (30) (**Fig. 2E**) and extracted GFP-tagged ribosomes from PAPs sampled from the dorsal hippocampus of adult Aldh1l1:L10a-eGFP mice. After reverse transcription, cDNA libraries were sequenced and the reads were aligned against the *Mus musculus* genome, in order to identify the corresponding genes. We first checked that our libraries were enriched in astrocyte-specific genes and depleted in genes specific to neurons, myeloid cells, oligodendrocytes, mural cells (vascular smooth muscle cells and pericytes), and endothelial cells (**Fig. 2F**). The raw list was then refined by filtering for mRNAs with ≥500 reads and an exon coverage ≥70%; this selected the most highly represented, complete mRNAs corresponding to potential translational events (for an exhaustive list, see **Table S2**). A total of 1118 mRNAs were detected in PAPs. This set included most of the known astrocyte-specific and astrocyte-enriched mRNAs (e.g. Slc1a2, Slc1a3, Mlc1, Hepacam, Gfap, Ezr, Gjb6, Gja1 and Gpr37ll1) (**Table S2**) (43). These results were validated by FISH experiments evidencing the distal distribution of some of these mRNAs (Gfap and Gjb6) (**Fig. 2G**). Interestingly, a number of astrocyte-specific mRNAs (Sox9, Aldh1l1, Slc25a18 and Ptprz1) were not found - suggesting that mRNAs are selectively distributed in PAPs (**Table S2**). A GO analysis indicated that most of the dorsal hippocampal PAP mRNAs encode cytosolic (32%) and intracellular organelle (44%) proteins. Large proportions are involved in protein synthesis (27%), metabolism (35%), and cell signaling (12%) (**Fig. 2H**). Consistently, some of the most abundant mRNAs in PAPs coded for ribosomal proteins (such as Rpl4 and Rplp1) or proteins involved in mRNA stability and translation (such as the elongation factors eEF1A1 and eEF2 and the Poly(A) binding protein PABPC1) (**Table S2**). This indicates that the molecular machinery for translation itself might be translated in PAPs. Lastly, mRNAs encoding the two ferritin isoforms Ftl1 and Fth1 were also highly represented, suggesting that local translation has a role in local iron storage and reduction (44).

After characterizing the pool of polysomal mRNAs present in dorsal hippocampal PAPs, we decided to refer to it as the “PAPome”, by analogy to our recent description of the “endfeetome”, the polysomal transcriptome found in perivascular astrocyte endfeet (26).

### Polysomal mRNAs are differentially distributed in hippocampal PAPs

After having characterized the PAPome, we determined which of its mRNA components were enriched in the PAP relative to the astrocyte as a whole. After homogenizing dorsal hippocampus tissue from Aldh1l1:L10a-eGFP mice, we used TRAP to purify all the polysomal mRNAs in astrocytes (**Fig. 3A**). As described above, three independent cDNA libraries were synthesized and sequenced. Astrocyte-specific mRNAs were highly enriched, whereas other brain cell-specific RNAs were depleted (**Fig. 3B**). Application of the criteria used in the previous experiment (≥500 reads and ≥70% exon coverage) to the raw list gave a refined list of 2,546 mRNAs, including those coding for most of the known astrocyte-specific or-enriched proteins such as Sox9, Aldh1l1, Slc25a18 or Ptprz1, which were not present in the PAPome (**Table S2**). The distribution of polysomal mRNAs within the astrocyte was then analyzed by comparing the PAPome with the whole-astrocyte repertoire (**Table S2**). We found that 557 mRNAs were significantly enriched in whole astrocytes (p value ≤ 0.05; log2 fold-change ≥1), 82 were enriched in PAPs (p value ≤ 0.05; log2 fold-change ≤ −1) and 678 mRNAs were similarly present in whole astrocytes and PAPs (-1≤log2 fold-change≤1). The astrocyte-specific transcripts Gfap, Slc1a2, Slc1a3, Mlc1, Hepacam, Ezr, Gjb6, Gja1 and Gpr37ll1 were all equally detected in PAPs and whole astrocytes, indicating that they are probably homogeneously translated in hippocampal astrocytes. The PAP-enriched mRNAs had an unexpected profile; a GO analysis indicated that most of them encoded cytosolic proteins involved in metabolism (24%), cell signaling (23%), development (15%) or protein synthesis (13%) (**Fig. 3C**). Twenty-seven of the transcripts coded for ribosomal proteins, such as Rplp1or Rpl4 (**Fig. 3D**). We also noted the presence of the two ferritin-subunit-encoding mRNAs Fth1 and Ftl1, as well as mRNAs related to cell cycle mechanisms (e.g. Ccnd2 and Mdm2) (**Fig. 3D**). The PAPs were also rich in mRNAs coding for cytoskeletal proteins, including not only microtubule-related proteins (Tubb5 and Mapre1), but also the ezrin and radixin proteins involved in linking the cell membrane to the actin cytoskeleton (**Fig. 3D**). The results of qPCRs for a selection of the most abundant PAP-enriched mRNAs validated these findings (**Table S3**). Overall, our data show that the polysomal mRNA distribution has a highly specific molecular polarity in PAPs and that those mRNAs are involved in specific functions related to protein synthesis, cell cycle regulation, iron homeostasis, and cytoskeleton dynamics.

**Figure 3:**
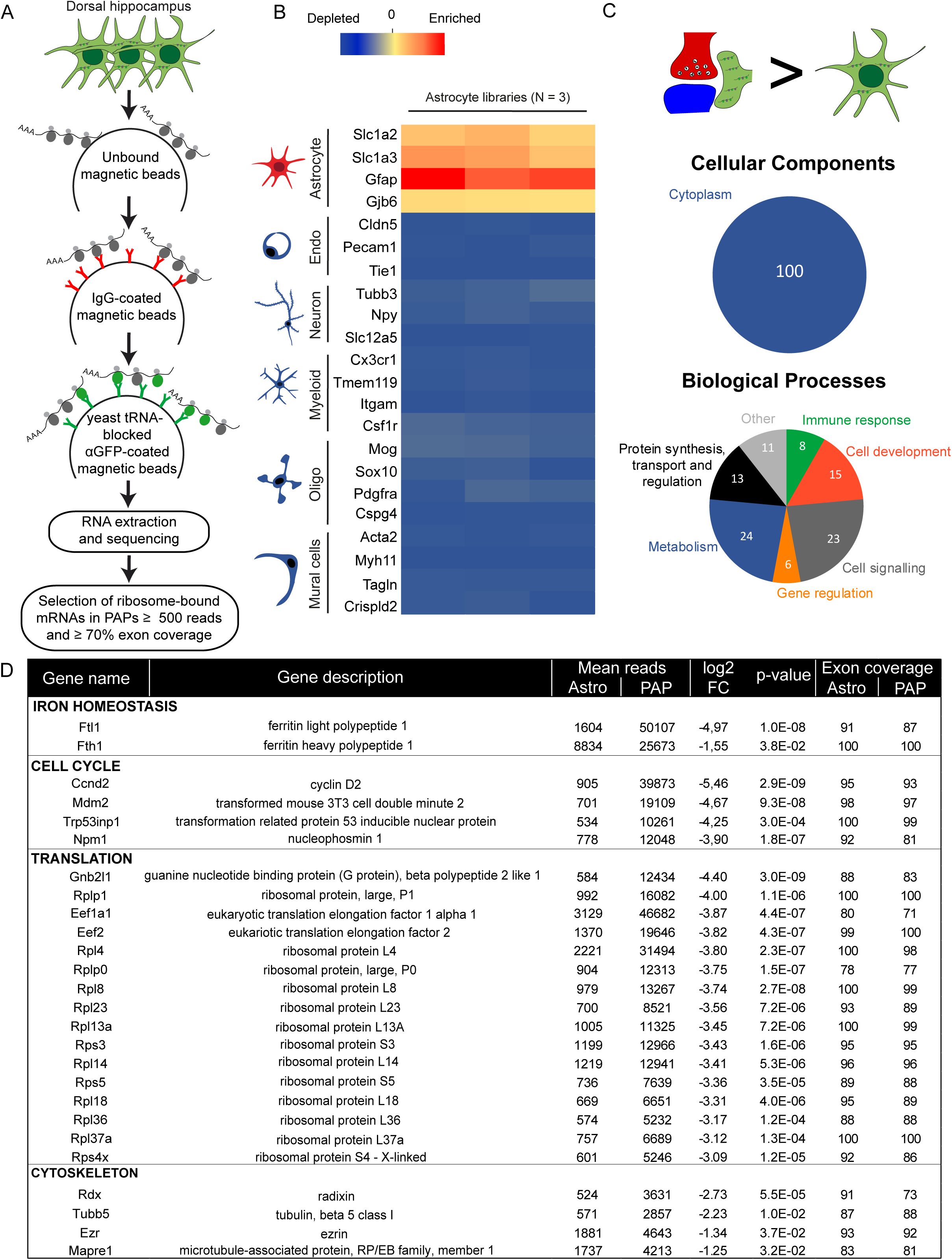
Identification of polysomal mRNAs enriched in PAPs from the dorsal hippocampus. **A.** Flowchart for the TRAP and analysis of polysomal mRNAs in astrocytes from the dorsal hippocampus. Sequencing raw data are in **Table S2**. **B.** Purity heat map of the RNA-seq data for a selection of mRNAs specific for each type of brain cell. The null level corresponds to mRNAs with 500 reads and 70% exon coverage. Each column represents an independent cDNA library. **C.** Gene ontology analysis of mRNAs enriched in PAPs, relative to whole astrocytes (p value ≤ 0.05; log2 fold-change ≤ −1). **D.** Raw sequencing data for a selection of the most PAP-enriched mRNAs (p-value ≤ 0.05; log2 fold-change ≤ −1). FC, fold-change.

### Differences in polysomal mRNA distribution between PAPs and perivascular astrocyte processes (PvAPs)

We have previously reported that local translation occurs in PvAPs (26). Although PvAPs contact synapses and therefore also qualify as PAPs (26), most PAPs are far from blood vessels and thus are likely to be dedicated to specific neuroglial interactions. We hypothesized that if local translation sustains astrocyte functional polarity, it would differ between PvAPs and PAPs. To address this question, we used qPCR assay to compare the expression of a selection of PAP-enriched polysomal mRNAs in hippocampal PAPs and PvAPs. Polysomal mRNAs were extracted from adult Aldh1l1:Rpl10a-eGFP whole hippocampus, using TRAP. For PAPs, we isolated synaptogliosomes (as described above). For PvAPs, we purified gliovascular units (30) (**Fig. 4A**). Interestingly, Ezr, Fth1, and Rplp1 mRNAs were more enriched in PAPs than in PvAPs, while all other mRNAs were equally detected in PAPs and PvAPs (**Fig. 4B**, **Table S4**). These results demonstrate that local translation differs in PAPs and PvAPs, and may thus sustain distinct molecular identities at each of these astrocytic interfaces.

**Figure 4:**
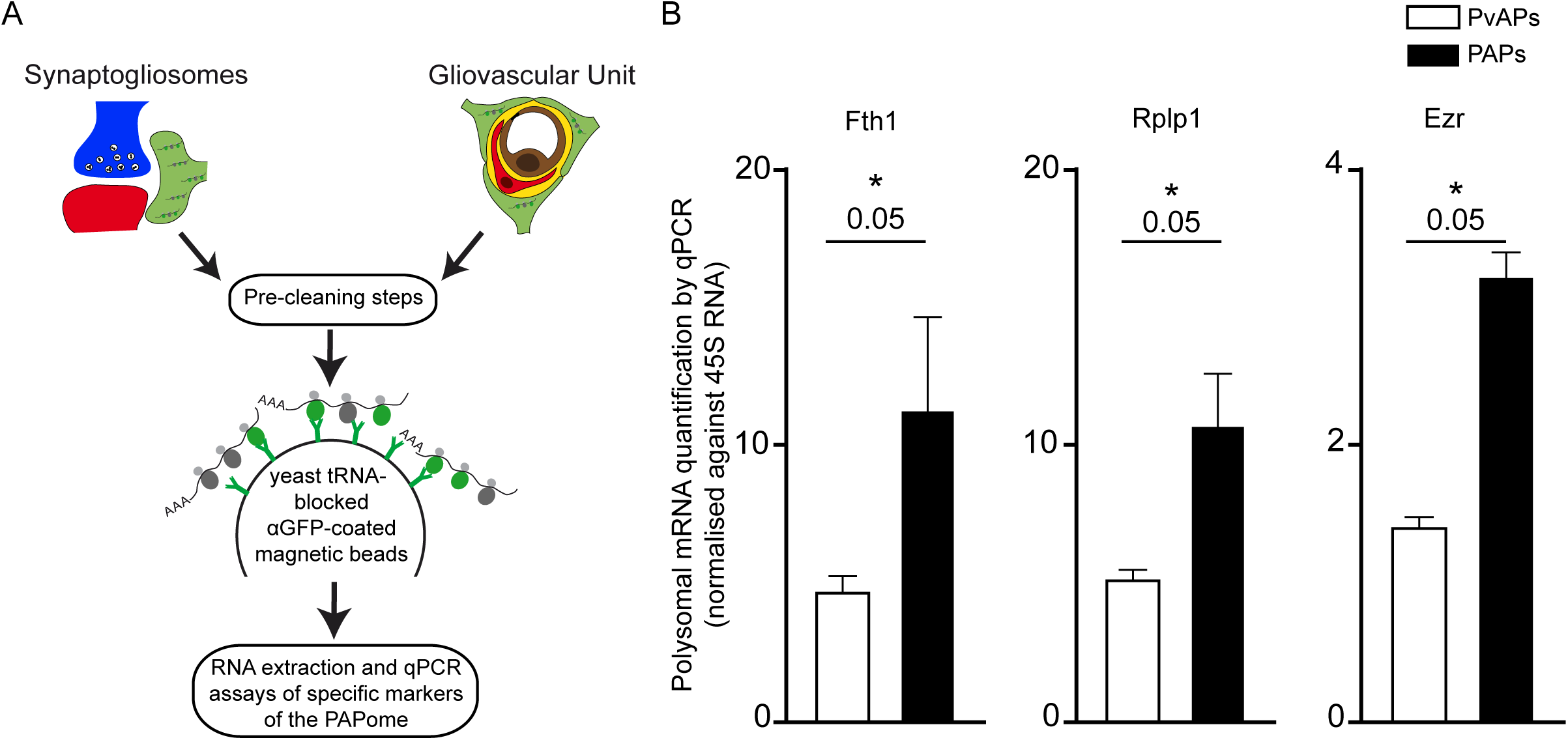
Comparison of enriched mRNA levels in PAPs vs. PvAPs. **A.** Flowchart for the comparison of a selection of enriched mRNAs in PAPs vs. PvAPs. Polysomal mRNAs were isolated by TRAP from Aldh1l1:L10a-eGFP whole hippocampal synaptogliosomes or purified gliovascular units. The raw qPCR data are given in **Table S4**. **B.** qPCR results for Fth1, Rplp1 and Ezr, demonstrating their enrichment in PAPs vs. PvAPs. *, p ≤ 0.05 in a one-tailed Mann-Whitney test. The data are quoted as the mean ± SEM. n=3.

### Fear conditioning influences local translation in PAPs

Recent studies have shown that astrocyte directly regulate memory formation through its contribution to local synaptic transmission and plasticity via exchange of regulatory signals with synapses (45–47). Although mRNA distribution and local translation in neurons have already been explored in the context of synaptic plasticity (48, 49) and memory formation (50–52), these mechanisms have not yet been evaluated in astrocytes. Strikingly, several of the polysomal mRNAs found to be enriched in PAPs (including Flt1, Fth1 (44, 53), Ccnd2 (54–56), Mdm2 (57–61), Gnb2l1 (62–64) and Eef1a1 (65–67)) have been linked to learning, memory or cognitive decline. We therefore investigated whether the local translation of these mRNAs in PAPs might be regulated by learning and memory processes. Hence, we exposed mice to a simple fear conditioning task (**Fig. 5A**): on Day 1, they were placed in a specific context (a new cage with white noise (5000 Hz, 60 dB) and an acetic acid (1%) smell). They were left to explore for three minutes before being exposed to four consecutive foot-shocks. The mice thus learnt to associate the cage with the unpleasant stimulus. On Day 2, the mice were placed in the same cage but did not receive foot-shocks. Re-exposure of the animal to the same aversive context allowed us to check the efficacy of the conditioning-induction, as estimated by the duration of freezing (> 70 s). Before comparing mRNAs in normal vs. conditioned mice, we first checked that the distance between the PAP and the synapse had not been affected by the conditioning protocol (**Fig. S2**). Indeed, it has been reported that the consolidation of fear conditioning induces morphological rearrangements in astrocytes from the lateral amygdala, notably with the emergence of synapses lacking PAPs, (68). Here, using super-resolution STED microscopy, we first measured individual PAP-synapse distances (ranging from 0 to 1500 nm) in the CA1 region of the dorsal hippocampus from hGfap:eGFP transgenic mice (in which eGFP fills the astrocyte cytoplasm) under control and fear conditions (69). STED imaging and co-immunostaining of eGFP with the pre- and post-synaptic markers VGluT1 and Homer1 generated data on the cumulative frequency of the distances between eGFP and VGluT1/Homer1 co-labeled synapses (**Fig. S2A, B**). There were no significant difference between control and fear-conditioned mice, indicating that the PAP-synapses distance was unchanged 24 h after exposure to contextual fear conditioning. Next, we performed Western blot analysis on the S1 (whole astrocyte) and P3 (synaptogliosome) fractions (**see** **Fig 2A**) from hGfap:eGFP mice. We did not observe any difference in eGFP content in either fraction, which suggests that the same amount of PAPs was purified from our synaptogliosomes (**Fig. S2C, D**). Lastly, we analyzed Aldh1l1:L10a-eGFP synaptogliosomes, and again did not observe any intercondition differences in the ribosomal content of PAPs (**Fig. S2E, F**). Overall, these results indicate that PAPs from the dorsal hippocampus did not retract, and retained their cytosolic content 24 h after fear conditioning.

**Figure 5:**
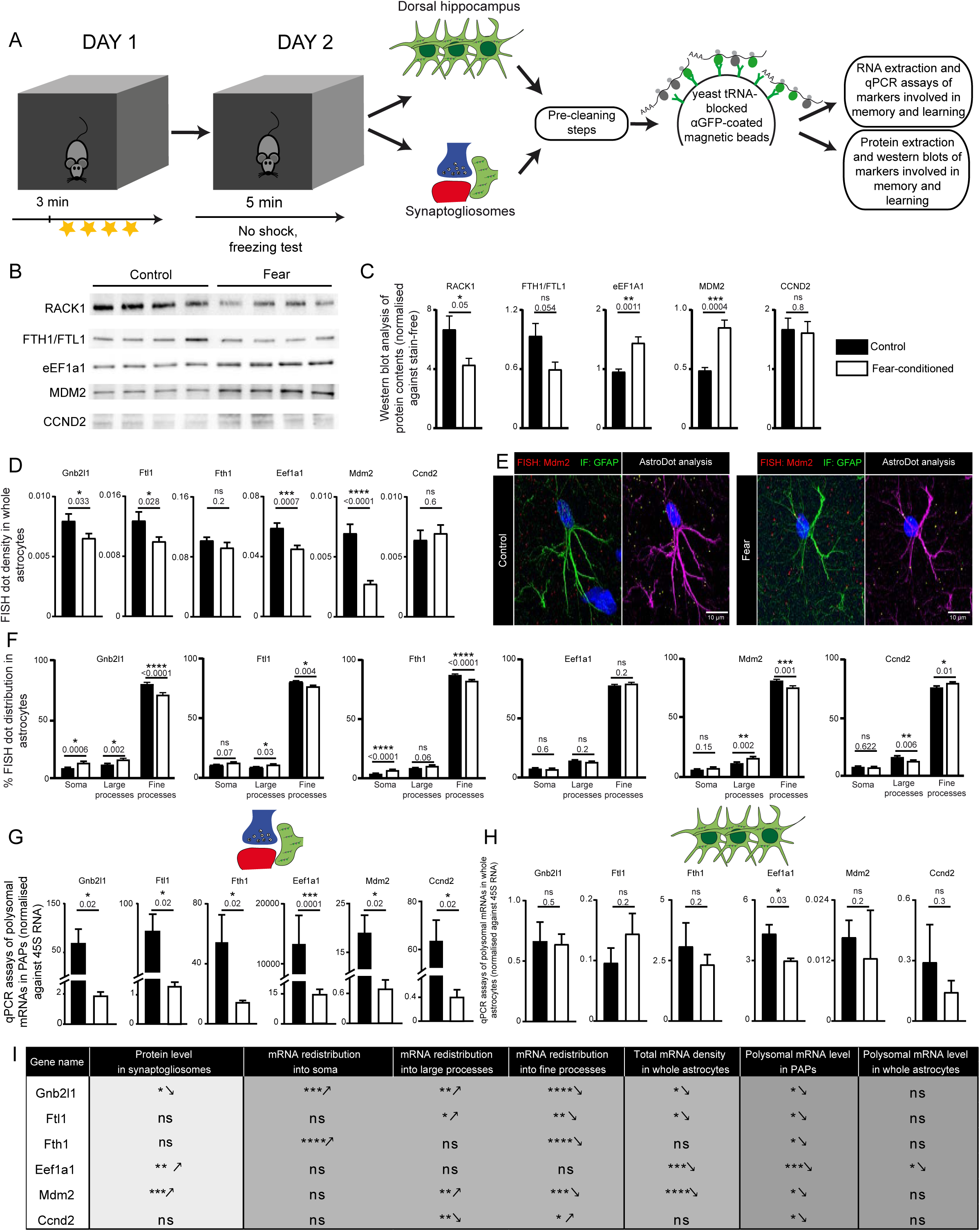
Fear conditioning regulates local translation in PAPs. **A.** Flowchart for the fear-conditioning protocol, followed by analyses of proteins and mRNAs in astrocytes and PAPs from the dorsal hippocampus. **B.** Western blots of RACK1, FTH1/FTL1, CCND2, MDM2 and eEF1A1 in dorsal hippocampal synaptogliosomes prepared from control mice and fear-conditioned mice. **C.** Analysis of the Western blots in **B**; signals were normalized against stain-free membranes. *, p ≤ 0.05, **, p ≤ 0.01, ***, p ≤ 0.001, and ns: not significant in a two-tailed, paired t-test. The data are presented as mean ± SEM (n=4). The raw data are given in **Table S5**. **D.** *AstroDot* analysis of the density of PAP-enriched mRNAs encoding RACK1 (Gnb2l1), FTL1, FTH1, eEF1A1 MDM2 and CCND2 in GFAP-immunolabeled astrocytes in the dorsal hippocampus of control mice and fear-conditioned mice. *, p<0.05; ***, p<0.001; ****p<0.0001 and ns: not significant in an unpaired, two-tailed t-test; n ≥35 cells from three mice per condition. The data are presented as the mean ± SEM, and the raw data are given in **Table S6**. **E. left panel,** confocal microscopy image of the FISH detection of Mdm2 mRNAs (red dots) in a dorsal hippocampal astrocyte immunolabeled for GFAP (green) in a control mouse and a fear-conditioned mouse. Nuclei are stained with DAPI (blue); **right panel,** *AstroDot* analysis of the left-panel image for each condition. The green dots were located in the soma or in GFAP-immunolabeled large processes; the yellow dots were located in GFAP-immunolabeled fine processes. GFAP immunostaining is shown in magenta. **F.** *AstroDot* analysis of the distribution of PAP-enriched mRNAs encoding RACK1 (Gnb2l1), FTL1, FTH1, eEF1A1 MDM2 and CCND2 in somata, large (> 0.3 μm) and fine (< 0.3 μm) GFAP-immunolabeled astrocyte processes from the dorsal hippocampus of control and fear-conditioned mice. *, p ≤ 0.05; ***, p ≤ 0.001; ****p ≤ 0.0001, and ns: not significant in a two-tailed, unpaired t-test; n ≥35 cells from three mice per condition. Data are presented as the mean ± SEM, and the raw data are given in Table S6. **G, H.** qPCR quantification of polysomal mRNAs in dorsal hippocampal PAPs (**G**) and whole astrocytes (**H**) from control mice and fear-conditioned mice. Signals are normalized against 45S RNA. *, p ≤ 0.05; ***, p ≤ 0.0001, and ns: not significant in a one-tailed Mann-Whitney test. The data are presented as the mean ± SEM (n=3 to 5 samples per condition), and the raw data are given in **Table S7**. **I.** A summary of the results.

To assess the impact of learning on our selected markers, we first analyzed the overall amount of each protein (using Western blots) in fear-conditioned and control synaptogliosomes extracted from the dorsal hippocampus (**Fig 5B, C**, **Table S5**). Levels of RACK1 (encoded by Gnb2l1) and ferritins (although not significantly) were lower in the fear-conditioned group, levels of MDM2 and eEF1A1were higher in the fear-conditioned group, and levels of CCND2 were similar in the fear-conditioned and control groups. These results indicated that fear conditioning influences the levels of most of these proteins at the neuroglial interface. We then looked for astrocyte-specific changes. Firstly, we looked at whether the density and distribution of these PAP-enriched mRNAs were modified by fear learning (**Fig. 5D, E**, **Table S6**). To this end, we used our *AstroDot* technique to analyze the distribution of fluorescently labeled mRNAs on GFAP-immunolabeled intermediate filaments in control and conditioned mice (31). With the exception of Fth1 and Ccnd2, the amounts of astrocyte mRNA were lower in the fear-conditioned group (**Fig. 5D, E**, **Table S6**). The distribution differed for almost all the mRNAs, with higher levels in large processes and lower levels in fine processes; the only exceptions were Eef1a1 (no difference in distribution) and Ccnd2 (lower levels in large processes and higher levels in fine processes) (**Fig. 5F**, **Table S6**). These results suggest that fear conditioning induces changes in the density and/or distribution of these mRNAs in astrocytes. Lastly, we determined whether the translation of these mRNAs in PAPs was modified upon fear conditioning. We used TRAP to extract polysomal mRNAs from dorsal hippocampus astrocytes and synaptogliosomes sampled from control and fear-conditioned Aldh1l1:L10a-eGFP mice, and then used qPCR to compare our panel of markers in whole astrocytes and PAPs (**Fig. 5G**, **Table S7**). In whole astrocytes, levels of all markers were similar with the exception of Eef1a1, which appeared to be slightly downregulated. In marked contrast, the PAP levels of all markers were lower in the fear-conditioned group (**Fig. 5H**, **Table S7**). In summary, levels of PAP-enriched transcripts may be regulated in different ways, i.e. mRNA transcription, mRNA redistribution, and ribosome binding for local translation (**Fig. 5I**). Upon fear conditioning, Eef1a1 (for example) might be less extensively transcribed in whole astrocytes and less extensively translated in PAPs. The opposite process might take place in neurons, as suggested by the higher level of protein expression in synaptogliosomes. Taken as a whole, these results suggest for the first time that learning and memory acquisition affects local translation in PAPs from the dorsal hippocampus.

## Discussion

The role of mRNA distribution and local translation in establishing cell functional polarity is widely established (15). In the central nervous system, these mechanisms contribute to neuronal development and synaptic transmission (18) and their potential involvement in the control of distal synaptic and vascular functions in astrocytes (a highly ramified type of glial cell) has recently been suggested (26, 28). In the present study, we focused on the perisynaptic astrocytic interface in the dorsal hippocampus, a region of the brain involved in memory and learning. We identified the pool of polysomal mRNAs in PAPs and showed that it is locally modified after a behavioral task related to learning and memory. To study local translation in PAPs, we used TRAP to purify PAP polysomes from synaptogliosomes. A modification of the TRAP protocol developed in our laboratory (the inclusion of cytosolic extract pre-cleaning and column-blocking steps) enabled us to eliminate non-specific mRNA binding events (30). As a result, we were able to purify a wide range of astrocytic mRNAs, without using their cell-specificity or enrichment as a selection criteria. In addition to intrinsic transcriptomic differences between astrocytes in different brain areas (70, 71), the use of this new procedure probably explains the drastic differences observed in our PAP-enriched polysomal transcriptome compared to the one described previously in the cortex, which was mainly composed of astrocyte-enriched or astrocyte-specific mRNAs (28).

The nature of enriched polysomal mRNAs in dorsal hippocampus PAPs suggests a role for local translation in specific biological processes: (i) The **structural plasticity of PAPs**: radixin and ezrin are actin-binding proteins that have already been involved in PAP morphology and motility (40–42). The cytoskeleton-associated protein category contained *β*-tubulin (encoded by Tubb5 mRNA, and which was recently described as being essential for neuronal differentiation and dendritic spine formation *in vivo* (72)) and EB1 (encoded by Mapre1, and which binds to the plus-end of microtubules and regulates their dynamics (73)). *In vitro,* microtubules have been implicated in the regulation of intermediate filament polarization in astrocytes (74). Although, microtubules have not been observed in PAPs *in vivo*, microtubule dynamics in these structures might also regulate the polarity. (ii) **Iron homeostasis**: Ferritin sequesters iron ions and regulates their toxicity through its ferroxidase activity. The local synthesis of ferritin in PAPs might therefore allow astrocytes to locally buffer iron and protect the synaptic environment from oxidative stress (44, 75). (iii) **Translation**: A large proportion of PAP-enriched mRNAs is thought to be involved in protein synthesis; the mRNAs code for ribosomal proteins, elongation factors, MDM2 (a ubiquitin E3 ligase that binds to ribosomal proteins (76) and acts as a translational repressor (61)) and NPM1 (a nucleolar phosphoprotein that regulates ribosome biogenesis (77)). Similar observations were recently made in neuronal processes (65, 78–80). Moreover, it was recently reported that newly translated ribosomal proteins in axons of primary cultured neurons incorporate into axonal ribosomes and sustain local translation (81). These convergent results strongly suggest that in astrocytes, as in neurons, local translation is sustained by the local synthesis of translation machinery, with either the *de novo* assembly of translational complexes or the replacement of damaged proteins. It is also possible that these proteins display non-canonical functions in PAPs, as has been described for eEF1A1 (82, 83).

In the present study, we did not look for alternative mRNA isoforms. In fact, our Illumina sequencing was based on the reconstruction of 75 bp sequences, which is not well suited to isoform analysis. Nevertheless, by comparing the results in PAPs vs. whole astrocytes, we noted that the C-terminal exons of Gfap *α* and *δ* (84) were differentially distributed: Gfap *α* was more abundant in PAPs, and GFAP *δ* was more abundant in the cell soma (**Table S2**). These findings corroborate our recent observation (based on FISH) than GFAP *δ* mRNA is more abundant that GFAP *α* mRNA in the soma (31), and suggest that alternative splicing may contribute to the PAPs’ molecular identity.

To establish whether the astrocyte’s functional polarity is mediated by local translation, we compared the levels of PAP-enriched polysomal mRNAs in PAPs and in PvAPs. Interestingly, some mRNAs appeared to be enriched in PAPs compared to PvAPs, indicating that the distribution and local translation of mRNAs differ at these two interfaces. The enrichment of Rplp1 mRNA in PAPs suggests that PvAPs and PAPs have different repertoires of ribosomal proteins and thus may follow different translational rules. The enrichment of Fth1 mRNA in PAPs suggests that iron reduction rather than storage prevails in PAPs (44). Another striking difference between PAPs and PvAPs relates to their ultrastructure. Although both processes display a continuous ER network, PvAPs (which are much larger than PAPs) can contain a full Golgi apparatus; in contrast, only pGolt-containing Golgi vesicles were observed in PAPs. Moreover, most of the mRNAs in the previously identified PvAP-enriched translatome code for secreted proteins and membrane proteins, whereas those in the PAP code for cytoplasmic proteins. These results indicate that post-translational protein modifications might be quite different in the two compartments and might again confer different functions to locally translated proteins.

Some of the mRNAs that are most highly enriched in PAPs code for proteins involved in learning and memory, namely FTH1, FTL1, eEF1a1, CCND2, MDM2 and RACK1. We therefore investigated the influence of fear conditioning on the translation of the corresponding mRNAs. We found that the level of polysomal mRNAs dropped specifically in PAPs 24 h after fear conditioning, indicating that their local translation might be altered. Interestingly, not all mRNAs behaved in the same way. Some showed a transcriptional effect and in some cases redistribution in astrocytes; this suggests that mRNA transport in astrocyte processes was also involved. The lower level of polysomal mRNAs in PAPs after fear conditioning was not always accompanied by a lower level of protein in the synaptogliosomes, which indicates the possible presence of opposing mechanisms in synapses. This was particularly striking for Eef1a1, with higher protein levels and lower levels of both total mRNA and polysomal mRNA in the synaptogliosomes after fear conditioning. Remarkably, Eef1a1 mRNA has been detected in axons, where it might be locally translated to maintain synaptic plasticity (65) and long-term potentiation (66). Its decrease at the synaptic level has been related to *α*-synucleinopathy (67). Since long-term potentiation and fear conditioning are intimately linked (85), opposite changes might occur in neurons and astrocytes (i.e. an increase in Eef1a1 levels on the neuronal side and a decrease on the astrocytic side).

Overall, our findings provide novel, unexpected insights into the PAP’s molecular identity - particularly the preferential translation of mRNAs related to iron homeostasis, cytoskeletal dynamics and the translation machinery in these compartments. Our findings also constitute the first *in vivo* evidence of local, neuronal-activity-induced translation changes in PAPs – changes that might be linked to the regulation of synaptic and circuit functions underlying complex behaviors.

## Material and Methods

### Mice

Tg(Aldh1l1-eGFP/Rpl10a) JD130Htz (MGI: 5496674) (Aldh1l1:L10a-eGFP) mice were obtained from Nathaniel Heintz’s laboratory (Rockefeller University, New York City, NY) and kept under pathogen-free conditions (86). The genotyping protocol is described on the bacTRAP project’s web site (www.bactrap.org). hGfap-eGFP mice were obtained from the Max Delbrück Center for Molecular Medicine (Berlin-Buch, Germany) (69). *C57BL6* mice were purchased from Janvier Labs (France) and kept in pathogen-free conditions.

### Ethical approval

All animal experiments were carried out in compliance with the European Directive 2010/63/EU on the protection of animals used for scientific purposes and the guidelines issued by the French National Animal Care and Use Committee (reference: 2013/118). The project was also approved by the French Ministry for Research and Higher Education’s institutional review board (reference 2018051809274047).

### High-resolution fluorescent in situ hybridization and GFAP co-immunofluorescent detection and analysis

Fluorescent in situ hybridization (FISH) was performed on floating brain sections obtained from mice perfused with 4% PFA/PBS, according to the v2 Multiplex RNAscope technique (Advanced Cell Diagnostics, Inc., Newark, CA, USA). After the FISH procedure, GFAP was immunofluorescently detected. Astrocyte-specific FISH dots were recognized on the base of their localization on the GFAP immunolabeling, using the *AstroDot* ImageJ plug-in. This method has been described in detail elsewhere (31). The Bacillus subtilis dihydrodipicolinate reductase (dapB) gene was used as a negative control. The probes are listed in Supplementary **Table S1**. Independent experiments were performed on three animals and two brain slices per animal.

### Gene ontology (GO) analyses

We used the text-mining Pathway Studio ResNet database (Ariadne Genomics, Rockville, MD) and the gene set enrichment analysis (GSEA) tool (87) in Pathway Studio (version 12.1.0.9) (88) to identify signaling pathways and biological processes that were overrepresented in our list of differentially expressed genes. To determine pathways for PAPs and whole astrocytes, we used the GO tool (categories: Biological Processes and Cellular Components; p-value threshold: 0.01; percentage overlap: >15%). To compare the lists from PAPs vs. whole astrocytes, we selected the Kolmogorov-Smirnov test in the GSEA tool (p-value threshold: 0.05).

### Preparation of synaptogliosomes

Synaptogliosomes were prepared from the dorsal hippocampi of 2-month-old mice. All steps were performed at 4°C. Dorsal hippocampi were dissected and homogenized with a tight glass homogenizer (20 strokes) in buffer solution (0.32 M sucrose and 10 mM HEPES in diethyl-dicarbonate-treated water, with 0.5 mM DTT, Protease Inhibitors (Complete-EDTA free) 1 mini tablet/10 mL, RNasin1µL/mL, CHX 100µg/mL added extratemporaneously). The homogenate was then centrifuged at 900g for 15 min. The pellet P1 was discarded, and the supernatant S1 was centrifuged at 16,000g for 15 min. The supernatant S2 was discarded, and the pellet P2 (containing synaptogliosomes) was diluted in 600 µl of buffer solution and centrifuged again at 16,000g for 15 min. The final pellet P3 contained the synaptogliosomes.

### Aldh1l1:L10a-eGFP TRAP, and RNA sequencing and analysis

The TRAP experiments were performed on male mice only since hippocampal astrocytes display sex-difference (89). Two dorsal hippocampi were used for whole astrocyte polysome extraction (*n*=3 libraries). For PAP polysome extraction, synaptogliosomes were isolated from dorsal hippocampi pooled from two mice (*n*=3 libraries). Polysomes were extracted using our recently published TRAP protocol (30). In order to avoid non-specific binding of the mRNAs to the beads or to IgG, and in contrast to the initial TRAP protocol (86), the cytosolic extracts were pre-cleaned more extensively prior to immunoprecipitation. Lysates were first placed on an antibody-free pre-cleaning column and then on a column with bound non-specific mouse IgGs. Non-specific sites on the immunoprecipitation column were blocked with yeast tRNA and BSA (30).

Polysomal mRNAs were purified using the RNeasy Lipid tissue kit (Qiagen). Ten nanograms of total RNA were amplified and converted into cDNA using a SMART-Seq v4 Ultra Low Input RNA kit (Clontech). Next, an average of 150 pg of amplified cDNA was used per library, with a Nextera XT DNA kit (Illumina). Libraries were multiplexed on two high-output flow cells and sequenced (75 bp reads) on a NextSeq 500 device (Illumina). The mean ± standard deviation number of reads per sample meeting the Illumina quality criterion was 23 ± 6 million.

The data were analyzed (read filtering, mapping, alignment filtering, read quantification, normalization and differential analysis) using the Eoulsan pipeline (90). Before mapping, adapters were removed (using TrimGalore (91) version 0.4.1), poly N read tails were trimmed, reads ≤ 40 bases were removed, and reads with a mean quality score ≤ 30 were discarded. The reads were then aligned against the *Mus musculus* genome (Ensembl version 84) using STAR (version 2.5.2b) (92). Alignments from reads matching more than once on the reference genome were removed using a Java version of samtools (93). The annotated *Mus musculus* gene transfer format file (Ensembl version 84) was used to compute gene expression. The regions in which alignments and referenced exons overlapped were counted using HTSeq-count (version 0.5.3) (93).

The RNA-seq gene expression data and raw fastq files are available on the GEO repository (www.ncbi.nlm.nih.gov/geo/) under the accession number GSE143531. The sample counts were normalized using DESeq2 (version 1.8.1) (94). Statistical processing and differential analyses were also performed using DESeq2 (version 1.8.1). The presence of an mRNA in whole astrocytes or in PAPs was defined by a read count ≥ 500 and ≥ 70% exon coverage. Enrichment in PAPs was defined as a log2 fold-change ≤ −1 and a p-value ≤ 0.05, whereas depletion was defined as a log2 fold-change ≥1 and a p-value ≤ 0.05.

### Quantitative RT-PCR

RNA was extracted using the Rneasy Lipid tissue mini kit (Qiagen, Hilden, Germany). cDNA was then generated using the Superscript™ III Reverse Transcriptase kit and pre-amplified with a mixture of the tested TaqMan® probes (**Table S1**) using the SsoAdvanced™ PreAmp Supermix (Biorad). Differential levels of cDNA expression were measured using the droplet digital PCR (ddPCR) system (Biorad) and TaqMan® copy number assay probes. Briefly, cDNA and 6-carboxyfluorescein (FAM) probes and primers were distributed into approximately 10,000 to 20,000 droplets. The nucleic acids were then PCR-amplified in a thermal cycler and read (as the number of positive and negative droplets) with a QX200 Droplet Digital PCR System. The ratio for each tested gene was normalized against the total number of positive droplets against 45S RNA.

### Western blots

Synaptogliosome pellets were sonicated three times for 10 s at 20 Hz (Vibra cell VCX130) and boiled in Laemmli loading buffer. Protein content was measured using the Pierce 660 nm protein assay reagent (Thermo Scientific, Waltham, MA, USA). Equal amounts of proteins were separated by denaturing electrophoresis in Mini-Protean TGX stain-free gels (Biorad) and then electrotransferred to nitrocellulose membranes using the Trans-blot Turbo Transfer System (Biorad). Membranes were hybridized as described previously (95). The antibodies used in this study are listed in **Table S1**. Horseradish peroxidase activity was visualized by enhanced chemiluminescence in a Western Lightning Plus system (Perkin Elmer, Waltham, MA, USA). Chemiluminescent imaging was performed on a LAS4000 system (Fujifilm, Minato-ku, Tokyo, Japan). At least four independent samples were analyzed in each experiment. The level of chemiluminescence for each antibody was normalized against stain-free signal on membranes.

### Viral vectors and stereotaxic injection

Two independent transgenes (composed of ERGIC-53 and pGolt cDNAs fused with GFP or mCherry respectively) were inserted under the control of the gfaABC1D synthetic promoter (37) (to drive their expression in astrocytes) in an AAV shuttle plasmid containing the inverted terminal repeats of AAV2. Pseudotyped serotype 9 AAV particles were produced by transient co-transfection of HEK-293T cells, as described previously (96). Viral titles were determined by quantitative PCR amplification of the inverted terminal repeats on DNase-resistant particles, and expressed in vector genomes (vg) per ml.

Two-to five-month-old mice were anesthetized with a mixture of ketamine (95 mg/kg; Merial) and xylazine (10 mg/kg; Bayer) in 0.9% NaCl and placed on a stereotaxic frame with constant body temperature monitoring. Adeno-associated viruses were diluted in PBS with 0.01% Pluronic F-68 at a concentration of 9x10^12^ vg/ml and 1 µl of virus was injected unilaterally into the right hippocampus at a rate of 0.1 µl/min, using a 29-gauge blunt-tip needle linked to a 2 µl Hamilton syringe (Phymep). The stereotaxic coordinates relative to the bregma were: anteroposterior, −2 mm; mediolateral: +1.5 mm; and dorsoventral, −1.5 mm. The needle was left in place for 5 min and was then slowly removed. The skin was glued back in place, and the animals’ recovery was checked regularly for the next 24 h. After 3 weeks, the mice were sacrificed and processed for immunostaining (see below). To determine the number of astrocytic pGolt and ERGIC-53 particles within 1µm of synapses, we developed an ImageJ plugin. We first defined a synapse as a VGluT1 particle sharing at least one common voxel with a Homer1 particle. The synapse location was calculated as the half distance between the VGluT1 and Homer1 particle centers. The inverted distance map of the ERGIC-53-GFP or pGolt-mCherry channel allowed to measure the distance from a synapse to the astrocyte compartment. We define a range between 0 and 1 µm to define a PAP.

### Immunohistolabeling, puromycylation and confocal imaging

Brain sections were fixed in PBS/PFA 4% for 15 min, rinsed in PBS, incubated for 1 h at room temperature in blocking solution (2% goat serum, 0.2% Triton X-100 in PBS). Next, the sections were incubated with primary antibodies diluted in the blocking solution for 12 h at 4 °C, rinsed for 5 min in PBS three times, incubated with secondary antibodies diluted in blocking solution for 2h at room temperature, rinsed for 5 min in PBS three times, and mounted in Fluoromount (Southern Biotech, Birmingham, AL). Brain sections were imaged on X1 and W1 spinning-disk confocal microscopes (Yokogawa). Images were acquired with a 63X oil immersion objective (Zeiss). Synaptogliosomes were immobilized on a glass slide coated with Cell-Tak (Corning) and immunostained as described previously. In order to measure the distance from the PAP to the synapse, we performed super-resolution stimulated emission depletion (STED) microscopy on a home-built, time-gated system. The latter was based on a commercial point scanning microscope (RESOLFT, Abberior Instruments) comprising a microscope frame (IX83, Olympus), galvanometer mirrors (Quad scanner, Abberior Instruments), and a detection unit consisting of two avalanche photodiodes (SPCM-AQRH, Excelitas Technologies). Images were acquired with a 100X/1.4 NA oil immersion objective lens (UPLSAPO 100X, Olympus). Confocal images were then deconvolved using the Huygens software and combined with the STED images in one file for analysis. For each astrocyte, 4 regions of interest of about 600 µm² were imaged. Homer1 and VGluT1 stainings were acquired in both confocal and STED resolution. The analysis was performed using an in-house developed plugin on ImageJ to analyze the distance from an identified synapse to the closest astrocyte process. In brief, maxima intensities were identified in STED images and compared to the deconvolved confocal images to avoid false positive punctae. A synapse was defined as two punctae from Homer1 and VGluT1 found within 300 µm of each other. Distances were calculated from the center of the synapse to the closest eGFP-labeled astrocyte process. For puromycylation, transverse slices of 150µm from the hippocampi of a P17 mouse were prepared using a vibrating blade microtome (Leica) in cold (4°C) Artificial cerebrospinal fluid (aCSF) containing (in mM): 119 NaCl, 2.5 KCl, 2.5 CaCl_2_, 1.3 MgSO_4_, 1 NaH_2_PO_4_, 26.2 NaHCO_3_ and 11 glucose saturated with 95% O_2_ and 5% CO_2_. Slices were left to rest in storage chamber for 1h at room temperature, then incubated at 32°C for 1h. Half of the slices were placed in a chamber with 91µM puromycin (Sigma) at 32°C for 10min. The other half was left untouched and used as negative control. Slices were then washed out for 2min in fresh aCSF, fixed in PBS/PFA 4% overnight at 4°C, rinsed for 3min in PBS three times and permeabilized and blocked in 4% goat serum, 0.5% Triton X-100 in PBS overnight at 4°C. Next, slices were incubated overnight at 4°C with primary antibodies in the blocking solution without Triton, rinsed 3 times in PBS for 20 min, and incubated with secondary antibodies for 3h at room temperature. Slices were rinsed again with PBS and mounted in Fluoromount (Southern Biotech, Birmingham, AL) on glass cover slips. Images were acquired using Zeiss LSM 980 Confocal – Airyscan 2, with a 63X oil immersion objective (Zeiss). Antibodies and applications are listed in Supplementary **Table S1**.

### Fear conditioning

Behavioral tests were performed on two-month-old male Aldh1l1:L10a-eGFP, hGFAP-GFP and wild-type mice. On day 1, the fear-conditioning group was placed in a conditioning chamber (characterized by white noise (5000 Hz, 60 dB) and a smell of acetic acid (1%)) and left to explore for 3 min. The mice were exposed to four consecutive 1 s scrambled foot shocks (current: 0.6 mA; inter-shock interval: ∼60 s) and then returned to their home cage. On day 2, the mice were put back in the conditioning chamber to assess contextual conditioning. The time spent freezing during a 5 min session was recorded automatically using a camera Sony effio-e 700TVL and the infrared tracking device. It was quantified using the PolyFear Software. The mice were sacrificed shortly after the freezing test. Home-caged animals were used as controls.

## Acknowledgments

This work was funded by the Fondation pour la Recherche Médicale to M. Cohen-Salmon (AJE20171039094) and N. Mazaré (FDT201904008077), the ED3C doctoral school to N. Mazaré, the European Research Council (Consolidator grant #683154) and European Union’s Horizon 2020 research and innovation program (Marie Sklodowska-Curie Innovative Training Networks, grant #722053, EU-GliaPhD) to N. Rouach, the FP7-PEOPLE Marie Curie Intra-European Fellowship for career development (grant #622289) to G. Cheung, and the “Journées de Neurologie de Langue Française” to R. Tortuyaux . The creation of the Center for Interdisciplinary Research in Biology (CIRB) was funded by the “Fondation Bettencourt Schueller”. We thank Carole Escartin and Océane Guillemaud for helpful discussions and all members of the Orion imaging facility for their high-quality technical support. The École Normale Supérieure genomic core facility was supported by the France Génomique national infrastructure, funded as part of the “Investissements d’Avenir” program managed by the Agence Nationale de la Recherche (contract ANR-10-INBS-09).

**Supplementary figure legends**

**Figure S1: Quantification of PAPs in the dorsal hippocampus.**

**A.** gfaABC_1_D-eGFP-AAV-infected hippocampal astrocytes immunolabeled with Homer1 (blue) and VGluT1 (red). **B.** The number of GFP-positive sites within 1 μm of VgluT1/Homer1. Quantification in (**A**) of the percentage of eGFP staining within 1 μm of a synapse, determined as the percentage of PAPs. N=16 cells in three mice. Immunolabeled synapses, representing the total number of PAPs detected by this technique. **C.** Specificity of the AAV gfaABC_1_D-eGFP infection in astrocytes. Hippocampal astrocytes are immunostained with GFAP (red).

**Figure S2: Characterization of the PAP-synapse distance upon fear conditioning.**

**A.** Deconvoluted confocal microscopy image of a single plane containing an astrocyte and synapses in the dorsal hippocampus of a control hGfap-eGFP mouse. The astrocyte is immunolabeled for eGFP. Pre- and post-synapses are immunolabeled for VGluT1 (blue) and Homer1 (red), respectively. The magnified area shows the STED image for VGluT1 and Homer1 merged with deconvoluted confocal image for eGFP **B.** Cumulative frequency of eGFP as a function of the distance to the synapse (from 0 to 800 nm). Solid lines represent the calculated means, and dotted lines represent the SEM for astrocytes from control mice (green) and fear-conditioned (red) mice. n≥270 synapses per cell, three cells per mouse, and three mice per condition. Two-way ANOVA, interaction value, p=0.5 (not significant). **C.** Western blot analysis of S1 (whole astrocyte extracts) and P3 (synaptogliosomes) (see Fig. 2) from hGfap:eGFP control mice or fear-conditioned mice. **D.** Analysis of the results in **C**, indicating that the amount of eGFP in astrocytes and PAPs was stable upon fear conditioning. Signals were normalized against stain-free membranes. The data are presented as the mean ± SEM (n=4); ns: not significant in an unpaired, two-tailed t-test. **E.** Western blot analysis of S1 (whole astrocyte extracts) and P3 (synaptogliosomes) from Aldh1l1:L10a-eGFP control mice and fear-conditioned mice. **F.** Analysis of the results in **E**, indicating that the amount of eGFP-tagged ribosomes was stable in astrocytes and PAPs. The signals were normalized against stain-free membrane. The data are presented as the mean ± SEM (n=4); ns: not significant in an unpaired, two-tailed t-test.

**Table S1: List of reagents**

**Table S2: Raw RNA-seq data for polysomal mRNAs from dorsal hippocampal PAPs and whole astrocytes.** This table contains 7 pages: **1.** RNA-seq raw data of polysomal mRNAs extracted from whole astrocytes and PAPs; **2.** PAPome: list of polysomal transcripts in PAPs with ≥500 reads and exon coverage ≥70%; **3.** Whole astrocytes: list of polysomal transcripts in whole astrocytes with ≥500 reads and exon coverage ≥70%; **4.** PAP-enriched: list of polysomal transcripts enriched in PAPs: p value ≤ 0.05; log2 fold-change ≤ −1; **5.** Astrocyte-enriched: list of polysomal transcripts enriched in whole astrocytes: p value ≤ 0.05; log2 fold-change ≥1; **6.** Equally present: list of polysomal transcripts equally present in PAPs and Whole astrocytes; −1≤log2 fold-change≤1. 7. GFAP exon study: comparison of whole astrocytes/PAPs for each GFAP exon. Id: identity number in *Ensembl.org*; BaseMean Read, mean of the reads in all libraries; padj, Adjusted p-value.

**Table S3: qPCR analysis of polysomal mRNAs in whole astrocytes and PAPs from the dorsal hippocampus.** SEM: standard error of the mean; *, p ≤ 0.05; n=3; one-tailed Mann-Whitney test.

**Table S4: qPCR analysis of polysomal mRNAs in PAPs and PvAPs from the hippocampus.** SEM: standard error of the mean; *, p ≤ 0.05, ns: not significant; n=3; one-tailed Mann-Whitney test.

**Table S5: Western blot analysis of proteins encoded by PAP-enriched mRNAs in synaptogliosomes purified from the dorsal hippocampus in control mice and fear-conditioned mice.** SEM: standard error of the mean; *, p ≤ 0.05, **, p ≤ 0.01, *** p ≤ 0.001, ns: not significant in an unpaired two-tailed t-test; n=4 to 5 per condition.

**Table S6: Analysis of the density and distribution of PAP-enriched mRNAs in control mice and fear-conditioned mice.** AstroDot raw data: *AstroDot* data for the density and distribution of mRNAs Ccnd2, Gnb2l1, Eef1a1, Mdm2, Fth1, and Ftl1 mRNAs. Descriptive statistics: summary of the results for each marker in control mice and fear-conditioned mice. SEM: standard error of the mean; n ≥35 cells for all markers, three mice per condition. Unpaired two-tailed t-test.

**Table S7: qPCR analysis of dorsal hippocampus polysomal mRNAs in whole astrocytes and PAPs in control and fear-conditioned mice.** SEM: standard error of the mean; *, p ≤ 0.05; ***, p ≤ 0.0001, ns: not significant in a one-tailed Mann-Whitney test; n=3 to 5.

