## Supplementary files for "Local translation in perisynaptic astrocytic processes is specific and regulated by fear conditioning"

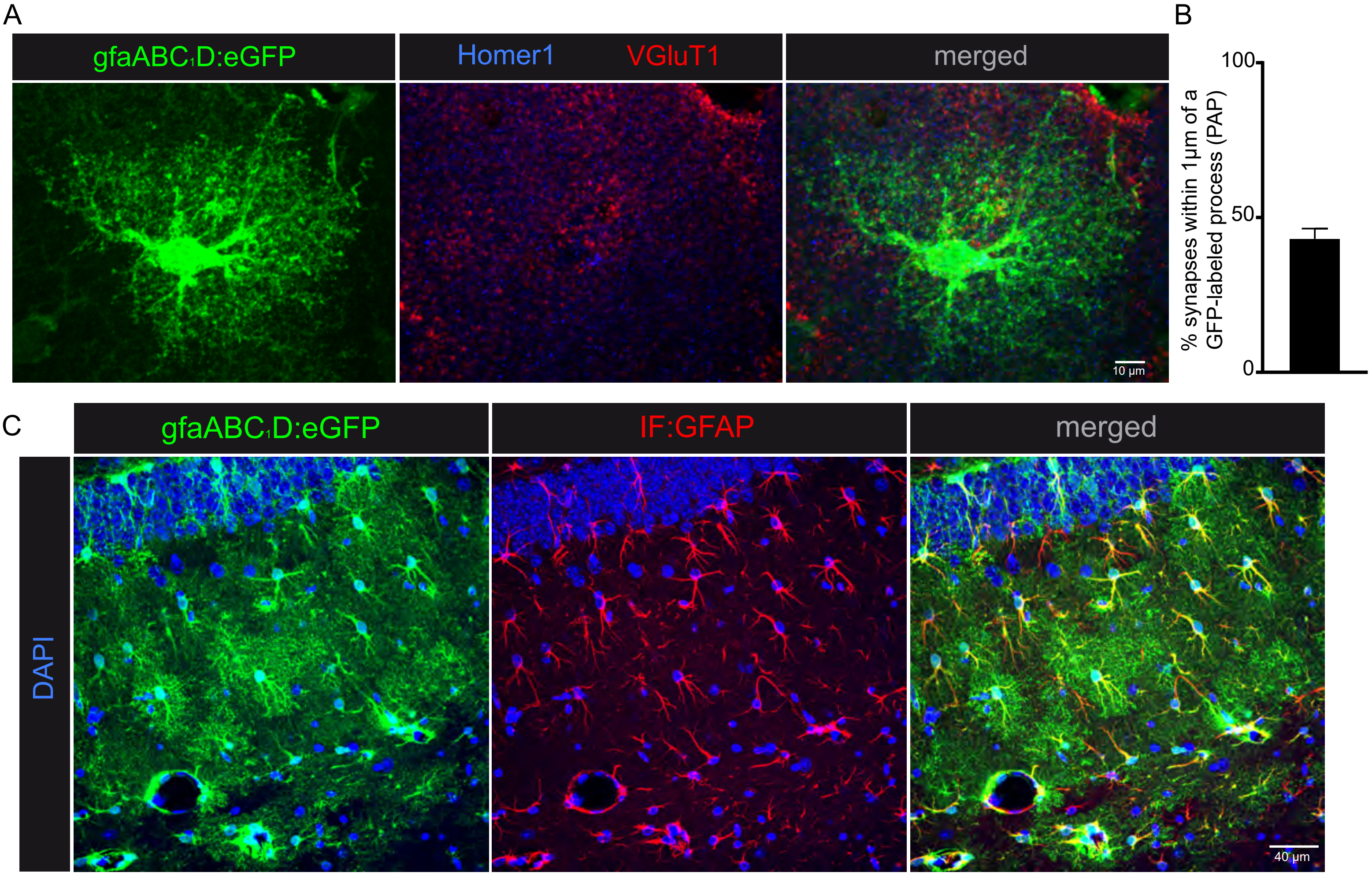
A
gfaABC1D:eGFP
Homer1
VGluT1
merged
10 µm
B
% synapses within 1µm of a GFP-labeled process (PAP)
100
50
0
C
gfaABC1D:eGFP
IF:GFAP
merged
DAPI
40 µm

Figure S1

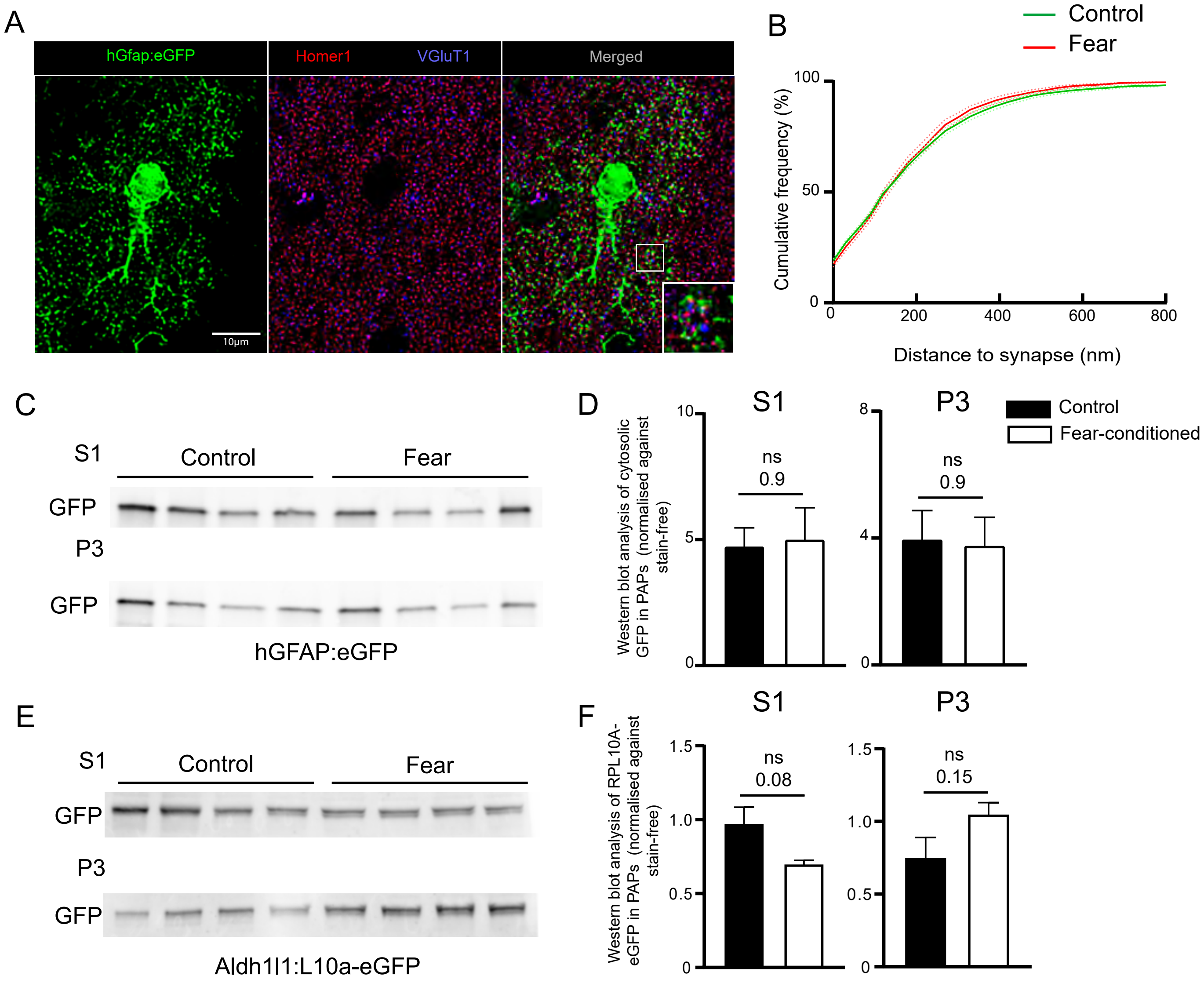

A
hGfap:eGFP
Homer1
VGluT1
Merged
10µm
B
Control
Fear
Cumulative frequency (%)
100
50
0
0
200
400
600
800
Distance to synapse (nm)
C
S1
Control
Fear
GFP
P3
GFP
hGFAP:eGFP
D
S1
P3
Control
Fear-conditioned
Western blot analysis of cytosolic GFP in PAPs (normalised against stain-free)
10
5
0
8
4
0
ns
0.9
ns
0.9
E
S1
Control
Fear
GFP
P3
GFP
Aldh1l1:L10a-eGFP
F
S1
P3
Western blot analysis of RPL10A-eGFP in PAPs (normalised against stain-free)
1.5
1.0
0.5
0
ns
0.08
ns
0.15

Table S1

| | Gene name | Accession Number | Catalog number Advanced Cell Diagnostic | Target Region | Channel |
|---|---|---|---|---|---|
| FISH probes | Fth | NM_010239.2 | 511561 | 3 - 926 | C1 |
| | Ftl1 | NM_010240.2 | 314771 | 6 - 908 | C1 |
| | Gfap | NM_001131020.1 | 313211 | 2 - 1761 | C1 |
| | Gjb6 | NM_001010937.2 | 458811 | 251 - 1383 | C1 |
| | Slc1a2 | NM_001077514.3 | 441341 | 1001 - 2251 | C1 |
| | Gnb2l1 | NM_008143.3 | 443621 | 152 - 439 | C1 |
| | Mdm2 | NM_010786.4 | 447641 | 832 - 1789 | C1 |
| | Eef1a1 | NM_010106.2 | 564581 | 2 - 1412 | C1 |
| | Ccnd2 | NM_009829.3 | 433211 | 546 - 1783 | C1 |

| | Protein name | Species | Company | Type | Dilution | |
|---|---|---|---|---|---|---|
| | | | | | Immunofluorescence | Western blot |
| Primary Antibodies | CCND2 | mouse | Clinisciences | Primary | 1/500 | 1/500 |
| | EEF1A1 | rabbit | Abcam | Primary | 1/500 | 1/500 |
| | Ezrin | mouse | Abcam | Primary | | 1/500 |
| | FTH1 | rabbit | Cell Signalling | Primary | 1/500 | 1/500 |
| | GFAP | rabbit | Sigma | Primary | 1/500 | 1/1000 |
| | GFP | chicken | Aves | Primary | 1/500 | 1/1000 |
| | Homer1 | rabbit | Synaptic Systems | Primary | 1/500 | |
| | MDM2 | mouse | Abcam | Primary | 1/500 | 1/1000 |
| | PSD95 | mouse | BD Biosciences | Primary | | 1/500 |
| | RACK1 | mouse | Clinisciences | Primary | 1/500 | 1/500 |
| | VGlut1 | mouse | Synaptic Systems | Primary | 1/500 | |
| | PMY-647 | mouse | Sigma | Conjugated | 1/5000 | |

| | Targeted species | Fluorophore/ HRP | Host Species | Company | Use |
|---|---|---|---|---|---|
| Secondary Antibodies | mouse | 488 | goat | Invitrogen | Immunostaining |
| | mouse | 555 | goat | Invitrogen | Immunostaining |
| | chicken | 488 | goat | Invitrogen | Immunostaining |
| | rabbit | 488 | goat | Invitrogen | Immunostaining |
| | rabbit | 647 | goat | Invitrogen | Immunostaining |
| | mouse | 647 | goat | Invitrogen | Immunostaining |
| | rabbit | 405 | goat | Invitrogen | Immunostaining |
| | chicken | HRP | donkey | Jackson | Western blot |
| | rabbit IgG | HRP | goat | Cohesion | Western blot |
| | mouse IgG | HRP | goat | Cohesion | Western blot |

| | Gene Name | Reference |
|---|---|---|
| Taqman probes | Ccnd2 | Mm00438070_m1 |
| | Ezr | Mm00447761_m1 |
| | Mdm2 | Mm00487656_m1 |
| | Rdx | Mm00501337_m1 |
| | Rps3 | Mm00656272_m1 |
| | Rpl8 | Mm00657299_g1 |
| | Hsp90ab1 | Mm00833431_g1 |
| | Rps14 | Mm00849906_g1 |
| | Fth1 | Mm00850707_g1 |
| | Slc1a2 | Mm01275814_m1 |
| | Gnb2l1 | Mm01291968_g1 |
| | Eef1a1 | Mm01973893_g1 |
| | Npm1 | Mm02391781_g1 |
| | Rplp1 | Mm02601846_g1 |
| | Ftl1 | Mm03030144_g1 |
| | Eef2 | Mm05700170_g1 |
| | Rpl4 | Mm05781370_g1 |

Table S2.1

| Id | baseMean Read | Mean read Astrocyte | Mean read PAP | log2FoldChange | padj | Associated Gene Name | Description | Mean Astrocyte Exon Coverage | Mean PAP exon Coverage |
|---|---|---|---|---|---|---|---|---|---|
| ENSMUSG00000089728 | 63,69588585 | 0,0 | 96,4 | -12,87289164 | 0,000213026 | Clec2f | C-type lectin domain family 2, member f [Source:MGI Symbol;Acc:MGI:3522133] | #DIV/0! | #DIV/0! |
| ENSMUSG00000079009 | 13,05408068 | 0,0 | 63,1 | -12,07776434 | 0,007465481 | Gm14139 | predicted gene 14139 [Source:MGI Symbol;Acc:MGI:3836251] | 0,00 | 25,2 |
| ENSMUSG00000020143 | 54,6345833 | 0,0 | 105,6 | -12,03668627 | 0,000597131 | Dock2 | dedicator of cyto-kinesis 2 [Source:MGI Symbol;Acc:MGI:2149010] | 0,00 | 11,1 |
| ENSMUSG00000023959 | 30,30353339 | 0,0 | 158,8 | -11,8366452 | 0,000273852 | Clic5 | chloride intracellular channel 5 [Source:MGI Symbol;Acc:MGI:1917912] | 0,00 | 12,0 |
| ENSMUSG00000089365 | 14,23002582 | 0,0 | 45,5 | -11,79215419 | 0,001619671 | Gm23823 | predicted gene, 23823 [Source:MGI Symbol;Acc:MGI:5453600] | #DIV/0! | #DIV/0! |
| ENSMUSG00000037922 | 21,91489306 | 0,0 | 79,7 | -11,46988569 | 0,002063127 | Bank1 | B cell scaffold protein with ankyrin repeats 1 [Source:MGI Symbol;Acc:MGI:2442120] | 0,00 | 19,5 |
| ENSMUSG00000042386 | 33,55065835 | 0,0 | 98,6 | -11,13464667 | 0,000449249 | Tex13 | testis expressed gene 13 [Source:MGI Symbol;Acc:MGI:1890544] | 0,00 | 29,8 |
| ENSMUSG00000076586 | 17,97228005 | 0,0 | 67,9 | -11,12575992 | 0,003510352 | Igkv8-21 | immunoglobulin kappa variable 8-21 [Source:MGI Symbol;Acc:MGI:1330840] | 0,00 | 72,0 |
| ENSMUSG00000085651 | 23,65921826 | 0,0 | 35,4 | -11,10461875 | 0,002344062 | Gm11695 | predicted gene 11695 [Source:MGI Symbol;Acc:MGI:3649841] | 0,00 | 62,8 |
| ENSMUSG00000106599 | 10,02015475 | 0,0 | 27,2 | -10,79187826 | 0,004094298 | Ighv1-73 | immunoglobulin heavy variable 1-73 [Source:MGI Symbol;Acc:MGI:3645131] | 0,00 | 15,1 |
| ENSMUSG00000021569 | 72,27199538 | 0,0 | 90,5 | -10,70240002 | 0,005806239 | Trip13 | thyroid hormone receptor interactor 13 [Source:MGI Symbol;Acc:MGI:1916966] | 0,00 | 16,7 |
| ENSMUSG00000087547 | 51,99076796 | 0,1 | 143,4 | -10,6738417 | 4,10056E-05 | Platr27 | pluripotency associated transcript 27 [Source:MGI Symbol;Acc:MGI:1925566] | #DIV/0! | #DIV/0! |
| ENSMUSG00000034116 | 118,615124 | 0,8 | 387,3 | -10,54848171 | 0,000323446 | Vav1 | vav 1 oncogene [Source:MGI Symbol;Acc:MGI:98923] | 3,10 | 62,6 |
| ENSMUSG00000021886 | 22,84737478 | 0,0 | 70,1 | -10,35913963 | 0,004482858 | Gpr65 | G-protein coupled receptor 65 [Source:MGI Symbol;Acc:MGI:108031] | 0,00 | 21,0 |
| ENSMUSG00000105788 | 6,220607739 | 0,0 | 17,6 | -10,29899315 | 0,006117719 | AC099934.1 | | #DIV/0! | #DIV/0! |
| ENSMUSG00000026288 | 32,2488908 | 3,3 | 172,2 | -10,11613474 | 0,000237317 | Inpp5d | inositol polyphosphate-5-phosphatase D [Source:MGI Symbol;Acc:MGI:107357] | 9,83 | 36,4 |
| ENSMUSG00000090673 | 44,28698776 | 0,3 | 204,9 | -10,07898001 | 0,000528972 | Gm340 | predicted gene 340 [Source:MGI Symbol;Acc:MGI:2685186] | 2,01 | 31,2 |
| ENSMUSG00000107788 | 9,33162789 | 0,0 | 13,9 | -10,06504661 | 0,006824289 | Gm6266 | predicted gene 6266 [Source:MGI Symbol;Acc:MGI:3646310] | #DIV/0! | #DIV/0! |
| ENSMUSG00000105181 | 7,305873248 | 0,0 | 18,7 | -10,03977562 | 0,007237627 | Gm19620 | predicted gene, 19620 [Source:MGI Symbol;Acc:MGI:5011805] | #DIV/0! | #DIV/0! |
| ENSMUSG00000044303 | 14,54187233 | 0,0 | 43,2 | -9,817664391 | 0,019798401 | Cdkn2a | cyclin-dependent kinase inhibitor 2A [Source:MGI Symbol;Acc:MGI:104738] | 0,00 | 27,6 |
| ENSMUSG00000034329 | 14,01322817 | 1,7 | 78,9 | -9,808908396 | 0,025444664 | Brip1 | BRCA1 interacting protein C-terminal helicase 1 [Source:MGI Symbol;Acc:MGI:2442836] | 3,32 | 4,8 |
| ENSMUSG00000066382 | 34,63085056 | 0,1 | 83,2 | -9,7075985 | 0,000100784 | Iqcf5 | IQ motif containing F5 [Source:MGI Symbol;Acc:MGI:1922720] | 4,48 | 74,8 |
| ENSMUSG00000042129 | 105,2486947 | 7,3 | 199,1 | -9,681211597 | 0,000950496 | Rassf4 | Ras association (RalGDS/AF-6) domain family member 4 [Source:MGI Symbol;Acc:MGI:2386853] | 14,40 | 21,9 |
| ENSMUSG00000031285 | 34,75633825 | 0,1 | 195,8 | -9,661371965 | 0,000910617 | Dcx | doublecortin [Source:MGI Symbol;Acc:MGI:1277171] | 0,23 | 5,9 |
| ENSMUSG00000058838 | 4,284714235 | 0,0 | 12,8 | -9,6521321 | 0,013801968 | Rps27a-ps2 | ribosomal protein S27A, pseudogene 2 [Source:MGI Symbol;Acc:MGI:3643614] | #DIV/0! | #DIV/0! |
| ENSMUSG00000035395 | 62,01028707 | 0,6 | 173,0 | -9,61462326 | 1,21102E-05 | Pet2 | plasmacytoma expressed transcript 2 [Source:MGI Symbol;Acc:MGI:101758] | 2,67 | 22,7 |
| ENSMUSG00000031383 | 23,78496601 | 0,0 | 30,8 | -9,585920641 | 0,010337501 | Dusp9 | dual specificity phosphatase 9 [Source:MGI Symbol;Acc:MGI:2387107] | 0,00 | 10,3 |
| ENSMUSG00000058216 | 12,94014738 | 0,0 | 43,1 | -9,521656443 | 0,038333083 | BC021614 | cDNA sequence BC021614 [Source:MGI Symbol;Acc:MGI:2385078] | 0,00 | 38,5 |
| ENSMUSG00000022831 | 21,47214401 | 0,0 | 15,8 | -9,429970147 | 0,010158021 | Hcls1 | hematopoietic cell specific Lyn substrate 1 [Source:MGI Symbol;Acc:MGI:104568] | 0,00 | 10,1 |
| ENSMUSG00000030157 | 11,97072508 | 0,8 | 58,9 | -9,404277097 | 0,018865545 | Clec2d | C-type lectin domain family 2, member d [Source:MGI Symbol;Acc:MGI:2135589] | 7,31 | 31,6 |
| ENSMUSG00000093208 | 4,505817378 | 0,0 | 11,0 | -9,367589484 | 0,01719502 | Gm27325 | predicted gene, 27325 [Source:MGI Symbol;Acc:MGI:5530707] | #DIV/0! | #DIV/0! |
| ENSMUSG00000040498 | 81,84824142 | 0,5 | 151,9 | -9,359013924 | 8,49619E-07 | Igsf23 | immunoglobulin superfamily, member 23 [Source:MGI Symbol;Acc:MGI:1917330] | 3,21 | 54,4 |
| ENSMUSG00000044201 | 97,81935734 | 0,9 | 390,3 | -9,356289186 | 1,54644E-06 | Cdc25c | cell division cycle 25C [Source:MGI Symbol;Acc:MGI:88350] | 8,17 | 94,8 |
| ENSMUSG00000084131 | 4,427746047 | 0,0 | 13,7 | -9,320183685 | 0,01146856 | Rpl3-ps2 | ribosomal protein L3, pseudogene 2 [Source:MGI Symbol;Acc:MGI:3645345] | #DIV/0! | #DIV/0! |
| ENSMUSG00000019836 | 8,518769402 | 0,0 | 32,3 | -9,240879045 | 0,004888202 | Amd-ps4 | S-adenosylmethionine decarboxylase, pseudogene 4 [Source:MGI Symbol;Acc:MGI:3650439] | #DIV/0! | #DIV/0! |
| ENSMUSG00000100512 | 140,2052037 | 0,6 | 258,5 | -9,209809258 | 1,46039E-09 | Ovol3 | OVO homolog-like 3 (Drosophila) [Source:MGI Symbol;Acc:MGI:2388075] | 14,14 | 90,2 |
| ENSMUSG00000048126 | 51,24498625 | 0,1 | 284,7 | -9,171289296 | 0,000239392 | Col6a3 | collagen, type VI, alpha 3 [Source:MGI Symbol;Acc:MGI:88461] | 0,23 | 22,8 |
| ENSMUSG00000027102 | 40,38916636 | 0,6 | 151,6 | -9,171248343 | 0,001220733 | Hoxd8 | homeobox D8 [Source:MGI Symbol;Acc:MGI:96209] | 2,88 | 31,2 |
| ENSMUSG00000020185 | 68,75840317 | 0,2 | 79,4 | -9,162364264 | 0,000939547 | E2f7 | E2F transcription factor 7 [Source:MGI Symbol;Acc:MGI:1289147] | 0,62 | 9,1 |
| ENSMUSG00000099767 | 20,27467324 | 0,9 | 52,0 | -9,134550807 | 0,00680684 | Gm28884 | predicted gene 28884 [Source:MGI Symbol;Acc:MGI:5579590] | 3,15 | 8,5 |
| ENSMUSG00000047347 | 3,276815649 | 0,0 | 7,0 | -8,99571349 | 0,043523277 | Tdg-ps | thymine DNA glycosylase, pseudogene [Source:MGI Symbol;Acc:MGI:3645587] | #DIV/0! | #DIV/0! |
| ENSMUSG00000024795 | 55,90097825 | 0,7 | 168,4 | -8,988957817 | 0,000950496 | Kif20b | kinesin family member 20B [Source:MGI Symbol;Acc:MGI:2444576] | 1,53 | 9,1 |
| ENSMUSG00000033036 | 7,370154038 | 0,0 | 6,6 | -8,988087848 | 0,021434724 | Gm7879 | predicted pseudogene 7879 [Source:MGI Symbol;Acc:MGI:3645078] | 0,00 | 11,6 |
| ENSMUSG00000040705 | 7,715212036 | 0,0 | 17,4 | -8,97121214 | 0,020897978 | A930016O22Rik | RIKEN cDNA A930016O22 gene [Source:MGI Symbol;Acc:MGI:3605623] | 0,00 | 4,9 |
| ENSMUSG00000104044 | 3,580510947 | 0,0 | 6,9 | -8,950009051 | 0,028435689 | Gm37566 | predicted gene, 37566 [Source:MGI Symbol;Acc:MGI:5610794] | 0,00 | 0,3 |
| ENSMUSG00000090104 | 3,857123291 | 0,0 | 9,1 | -8,916835476 | 0,015957979 | Slmapos2 | sarcolemma associated protein, opposite strand transcript 2 [Source:MGI Symbol;Acc:MGI:3801907] | #DIV/0! | #DIV/0! |
| ENSMUSG00000020330 | 145,7263491 | 1,3 | 435,6 | -8,902300654 | 1,82175E-05 | Hmmr | hyaluronan mediated motility receptor (RHAMM) [Source:MGI Symbol;Acc:MGI:104667] | 3,50 | 46,7 |
| ENSMUSG00000096177 | 3,351715577 | 0,0 | 6,3 | -8,832556827 | 0,024203883 | Gm5070 | predicted gene 5070 [Source:MGI Symbol;Acc:MGI:3643219] | #DIV/0! | #DIV/0! |
| ENSMUSG00000012705 | 7,814481945 | 0,0 | 36,1 | -8,818439028 | 0,022766165 | Retn | resistin [Source:MGI Symbol;Acc:MGI:1888506] | 0,00 | 14,6 |
| ENSMUSG00000056328 | 3,365355899 | 0,0 | 14,2 | -8,816345615 | 0,026467138 | Myh1 | myosin, heavy polypeptide 1, skeletal muscle, adult [Source:MGI Symbol;Acc:MGI:1339711] | 0,00 | 1,8 |
| ENSMUSG00000030399 | 7,412725361 | 0,0 | 14,5 | -8,798942336 | 0,02650583 | Ckm | creatine kinase, muscle [Source:MGI Symbol;Acc:MGI:88413] | 0,00 | 3,9 |
| ENSMUSG00000061469 | 23,02373565 | 0,1 | 72,9 | -8,760846398 | 0,000449249 | Gm12569 | predicted gene 12569 [Source:MGI Symbol;Acc:MGI:3650740] | 2,76 | 56,8 |
| ENSMUSG00000024989 | 57,11933486 | 4,9 | 276,6 | -8,730660564 | 0,00278201 | Cep55 | centrosomal protein 55 [Source:MGI Symbol;Acc:MGI:1921357] | 17,19 | 46,4 |
| ENSMUSG00000031026 | 3,07164826 | 0,0 | 6,4 | -8,64480854 | 0,028643912 | Trim66 | tripartite motif-containing 66 [Source:MGI Symbol;Acc:MGI:2152406] | 0,00 | 0,9 |
| ENSMUSG00000064493 | 7,131894516 | 0,0 | 10,7 | -8,6391343 | 0,036236029 | Snora28 | small nucleolar RNA, H/ACA box 28 [Source:MGI Symbol;Acc:MGI:3819496] | #DIV/0! | #DIV/0! |
| ENSMUSG00000029844 | 5,419046606 | 0,0 | 27,1 | -8,58439953 | 0,03977239 | Hoxa1 | homeobox A1 [Source:MGI Symbol;Acc:MGI:96170] | 0,00 | 11,6 |
| ENSMUSG00000011267 | 45,64812321 | 2,5 | 50,1 | -8,542302084 | 0,002611645 | Zfp296 | zinc finger protein 296 [Source:MGI Symbol;Acc:MGI:1926956] | 17,23 | 23,5 |
| ENSMUSG00000015314 | 63,69385277 | 0,3 | 162,7 | -8,469483382 | 0,001995574 | Slamf6 | SLAM family member 6 [Source:MGI Symbol;Acc:MGI:1353620] | 0,74 | 11,9 |
| ENSMUSG00000062393 | 59,50003078 | 1,3 | 99,3 | -8,462374048 | 0,00036757 | Dgkk | diacylglycerol kinase kappa [Source:MGI Symbol;Acc:MGI:3580254] | 2,22 | 10,6 |
| ENSMUSG00000078954 | 23,39496117 | 0,0 | 33,3 | -8,420144396 | 0,003282085 | Arhgap8 | Rho GTPase activating protein 8 [Source:MGI Symbol;Acc:MGI:1920417] | 0,00 | 23,7 |
| ENSMUSG00000044103 | 3,734263354 | 0,0 | 9,8 | -8,385590606 | 0,048726664 | Il1f9 | interleukin 1 family, member 9 [Source:MGI Symbol;Acc:MGI:2449929] | 0,00 | 0,8 |
| ENSMUSG00000038007 | 26,04126624 | 2,0 | 81,0 | -8,333543797 | 0,002117304 | Acer2 | alkaline ceramidase 2 [Source:MGI Symbol;Acc:MGI:1920932] | 3,85 | 8,4 |
| ENSMUSG00000062687 | 4,808459005 | 0,0 | 9,7 | -8,327281165 | 0,031168192 | Cks1brt | CDC28 protein kinase 1b, retrogene [Source:MGI Symbol;Acc:MGI:3643620] | 0,00 | 2,5 |
| ENSMUSG00000035711 | 40,9369764 | 0,9 | 152,5 | -8,184416325 | 7,89907E-07 | Dok3 | docking protein 3 [Source:MGI Symbol;Acc:MGI:1351490] | 10,67 | 71,2 |
| ENSMUSG00000074385 | 29,52901476 | 0,3 | 86,0 | -8,178886756 | 0,001014933 | Gm10684 | predicted gene 10684 [Source:MGI Symbol;Acc:MGI:3641851] | 1,86 | 12,3 |
| ENSMUSG00000107304 | 35,80808299 | 0,6 | 124,4 | -8,160039282 | 7,21588E-06 | Gm43775 | predicted gene 43775 [Source:MGI Symbol;Acc:MGI:5663912] | 2,75 | 14,0 |
| ENSMUSG00000089716 | 9,401797268 | 0,1 | 30,3 | -8,15209067 | 0,001750815 | Gm6264 | predicted gene 6264 [Source:MGI Symbol;Acc:MGI:3779580] | #DIV/0! | #DIV/0! |
| ENSMUSG00000043592 | 10,01290961 | 0,0 | 11,4 | -8,147178892 | 0,041791906 | Unc5cl | unc-5 homolog C (C. elegans)-like [Source:MGI Symbol;Acc:MGI:1923839] | 0,00 | 4,9 |
| ENSMUSG00000036223 | 13,56640756 | 1,2 | 19,8 | -8,112115247 | 0,031824215 | Ska1 | spindle and kinetochore associated complex subunit 1 [Source:MGI Symbol;Acc:MGI:1913718] | 2,82 | 7,9 |
| ENSMUSG00000024542 | 5,651477667 | 0,0 | 3,7 | -8,107932624 | 0,032792261 | Cep192 | centrosomal protein 192 [Source:MGI Symbol;Acc:MGI:1918049] | 0,00 | 0,9 |
| ENSMUSG00000020573 | 35,82887198 | 0,2 | 126,2 | -8,098369794 | 0,007303788 | Pik3cg | phosphoinositide-3-kinase, catalytic, gamma polypeptide [Source:MGI Symbol;Acc:MGI:1353576] | 0,80 | 15,3 |
| ENSMUSG00000069910 | 11,71823776 | 0,0 | 15,6 | -8,083827649 | 0,02338691 | Spdl1 | spindle apparatus coiled-coil protein 1 [Source:MGI Symbol;Acc:MGI:1917635] | 0,00 | 4,5 |
| ENSMUSG00000008193 | 22,69433712 | 0,6 | 60,7 | -8,069767193 | 0,00438382 | Spib | Spi-B transcription factor (Spi-1/PU.1 related) [Source:MGI Symbol;Acc:MGI:892986] | 3,58 | 20,5 |
| ENSMUSG00000018654 | 193,3881116 | 2,5 | 588,5 | -7,948338481 | 8,79878E-06 | Ikzf1 | IKAROS family zinc finger 1 [Source:MGI Symbol;Acc:MGI:1342540] | 7,21 | 49,5 |
| ENSMUSG00000087107 | 585,7003812 | 7,2 | 1528,1 | -7,948218774 | 4,26581E-12 | AI662270 | expressed sequence AI662270 [Source:MGI Symbol;Acc:MGI:2144254] | #DIV/0! | #DIV/0! |
| ENSMUSG00000049916 | 43,27887829 | 1,1 | 171,4 | -7,935128467 | 0,000629254 | 2610318N02Rik | RIKEN cDNA 2610318N02 gene [Source:MGI Symbol;Acc:MGI:1917708] | 12,65 | 49,8 |
| ENSMUSG00000045775 | 70,21720643 | 0,8 | 163,3 | -7,906462853 | 2,32992E-06 | Slc16a5 | lute carrier family 16 (monocarboxylic acid transporters), member 5 [Source:MGI Symbol;Acc:MGI:244351 | 3,32 | 29,5 |
| ENSMUSG00000033952 | 24,83338668 | 0,1 | 29,6 | -7,903255757 | 0,013582641 | Aspm | asp (abnormal spindle)-like, microcephaly associated (Drosophila) [Source:MGI Symbol;Acc:MGI:1334448 | 0,25 | 1,9 |
| ENSMUSG00000097519 | 29,59669299 | 0,5 | 173,6 | -7,811728773 | 0,001720238 | 4930558J18Rik | RIKEN cDNA 4930558J18 gene [Source:MGI Symbol;Acc:MGI:1922573] | #DIV/0! | #DIV/0! |
| ENSMUSG00000038253 | 11,20789663 | 0,1 | 61,8 | -7,80669151 | 0,011857656 | Hoxa5 | homeobox A5 [Source:MGI Symbol;Acc:MGI:96177] | 1,33 | 11,9 |
| ENSMUSG00000016758 | 16,76282178 | 1,0 | 47,0 | -7,780399868 | 0,00123279 | Bik | BCL2-interacting killer [Source:MGI Symbol;Acc:MGI:1206591] | 20,57 | 55,2 |
| ENSMUSG00000096326 | 10,87481718 | 0,1 | 21,3 | -7,762354244 | 0,003008247 | Ighv1-78 | immunoglobulin heavy variable 1-78 [Source:MGI Symbol;Acc:MGI:4439736] | 6,65 | 21,2 |
| ENSMUSG00000031343 | 3,895801195 | 0,0 | 8,4 | -7,698003304 | 0,039413959 | Gabra3 | gamma-aminobutyric acid (GABA) A receptor, subunit alpha 3 [Source:MGI Symbol;Acc:MGI:95615] | 0,00 | 1,9 |
| ENSMUSG00000068604 | 4,686099018 | 0,0 | 7,9 | -7,638759719 | 0,049495484 | Gm8225 | predicted gene 8225 [Source:MGI Symbol;Acc:MGI:3643826] | 0,00 | 5,7 |
| ENSMUSG00000104475 | 9,137199968 | 0,0 | 44,4 | -7,638254823 | 0,031656379 | D630036G22Rik | RIKEN cDNA D630036G22 gene [Source:MGI Symbol;Acc:MGI:3045265] | 0,00 | 19,8 |
| ENSMUSG00000097536 | 60,07850063 | 1,9 | 230,8 | -7,588622413 | 0,000916738 | 2610037D02Rik | RIKEN cDNA 2610037D02 gene [Source:MGI Symbol;Acc:MGI:1917290] | #DIV/0! | #DIV/0! |
| ENSMUSG00000102798 | 11,22517809 | 0,5 | 34,5 | -7,584067353 | 0,001659214 | Gm37271 | predicted gene, 37271 [Source:MGI Symbol;Acc:MGI:5610499] | 2,17 | 2,7 |
| ENSMUSG00000024673 | 133,4753033 | 5,4 | 464,5 | -7,533814608 | 0,001936366 | Ms4a1 | membrane-spanning 4-domains, subfamily A, member 1 [Source:MGI Symbol;Acc:MGI:88321] | 22,69 | 58,1 |
| ENSMUSG00000045730 | 16,62982648 | 0,7 | 95,7 | -7,509584668 | 0,002292486 | Adrb2 | adrenergic receptor, beta 2 [Source:MGI Symbol;Acc:MGI:87938] | 6,91 | 30,8 |
| ENSMUSG00000068105 | 38,95369603 | 1,1 | 128,4 | -7,488249194 | 0,009917731 | Tnfrsf13c | tumor necrosis factor receptor superfamily, member 13c [Source:MGI Symbol;Acc:MGI:1919299] | 8,96 | 44,1 |
| ENSMUSG00000045328 | 161,6178587 | 2,1 | 487,8 | -7,484265116 | 0,003152677 | Cenpe | centromere protein E [Source:MGI Symbol;Acc:MGI:1098230] | 3,61 | 21,2 |
| ENSMUSG00000090243 | 7,945814876 | 0,1 | 15,4 | -7,474697474 | 0,005859943 | Gm16103 | predicted gene 16103 [Source:MGI Symbol;Acc:MGI:3802086] | #DIV/0! | #DIV/0! |
| ENSMUSG00000054702 | 55,74106765 | 0,8 | 164,8 | -7,471394718 | 0,000999635 | Ap1s3 | adaptor-related protein complex AP-1, sigma 3 [Source:MGI Symbol;Acc:MGI:1891304] | 4,96 | 19,7 |
| ENSMUSG00000094940 | 16,91798048 | 0,3 | 48,2 | -7,452413345 | 0,001256291 | Ighv1-84 | immunoglobulin heavy variable 1-84 [Source:MGI Symbol;Acc:MGI:3644235] | 15,00 | 19,9 |
| ENSMUSG00000098713 | 6,462149692 | 0,1 | 17,3 | -7,450499762 | 0,004783257 | Rps2-ps5 | ribosomal protein S2, pseudogene 5 [Source:MGI Symbol;Acc:MGI:3647821] | #DIV/0! | #DIV/0! |
| ENSMUSG00000083382 | 3,728683775 | 0,1 | 15,2 | -7,411310902 | 0,016730362 | Gm6433 | predicted gene 6433 [Source:MGI Symbol;Acc:MGI:3645394] | #DIV/0! | #DIV/0! |
| ENSMUSG00000052613 | 36,43465038 | 1,7 | 184,2 | -7,411203435 | 0,003152677 | Pcdh15 | protocadherin 15 [Source:MGI Symbol;Acc:MGI:1891428] | 1,02 | 7,9 |
| ENSMUSG00000099021 | 10,50090853 | 0,2 | 24,1 | -7,409424385 | 0,002705858 | Rn7s1 | 7S RNA 1 [Source:MGI Symbol;Acc:MGI:97950] | 9,22 | 42,4 |
| ENSMUSG00000024211 | 26,48858467 | 0,1 | 27,4 | -7,393688269 | 0,017919652 | Grm8 | glutamate receptor, metabotropic 8 [Source:MGI Symbol;Acc:MGI:1351345] | 0,60 | 4,8 |
| ENSMUSG00000017697 | 108,8273181 | 5,5 | 225,4 | -7,38610165 | 1,73687E-05 | Ada | adenosine deaminase [Source:MGI Symbol;Acc:MGI:87916] | 27,90 | 72,9 |
| ENSMUSG00000091199 | 9,598537724 | 0,2 | 38,2 | -7,344214912 | 0,014130848 | Gm2619 | predicted gene 2619 [Source:MGI Symbol;Acc:MGI:3780787] | #DIV/0! | #DIV/0! |
| ENSMUSG00000032254 | 78,4379728 | 2,5 | 277,8 | -7,334475236 | 0,00125629 | Kif23 | kinesin family member 23 [Source:MGI Symbol;Acc:MGI:1919069] | 9,39 | 41,3 |
| ENSMUSG00000105601 | 68,85828253 | 1,3 | 145,5 | -7,332609959 | 2,49425E-07 | Gm42725 | predicted gene 42725 [Source:MGI Symbol;Acc:MGI:5662862] | 1,30 | 2,4 |
| ENSMUSG00000020914 | 360,2020894 | 2,9 | 404,0 | -7,301088796 | 1,42091E-07 | Top2a | topoisomerase (DNA) II alpha [Source:MGI Symbol;Acc:MGI:98790] | 9,90 | 33,7 |
| ENSMUSG00000097415 | 349,3295467 | 3,7 | 855,0 | -7,266807657 | 1,56781E-07 | AU020206 | expressed sequence AU020206 [Source:MGI Symbol;Acc:MGI:2142134] | #DIV/0! | #DIV/0! |
| ENSMUSG00000025154 | 359,6688039 | 6,5 | 571,9 | -7,261202535 | 2,34175E-08 | Arhgap19 | Rho GTPase activating protein 19 [Source:MGI Symbol;Acc:MGI:1918335] | 13,72 | 48,8 |
| ENSMUSG00000038179 | 47,0091889 | 0,9 | 135,6 | -7,240342231 | 0,008269402 | Slamf7 | SLAM family member 7 [Source:MGI Symbol;Acc:MGI:1922595] | 2,79 | 15,9 |
| ENSMUSG00000046179 | 79,32851451 | 0,5 | 86,3 | -7,22922709 | 0,022528419 | E2f8 | E2F transcription factor 8 [Source:MGI Symbol;Acc:MGI:1922038] | 2,16 | 14,9 |
| ENSMUSG00000102330 | 6,330848223 | 0,1 | 22,5 | -7,226563328 | 0,035785983 | Gm37234 | predicted gene, 37234 [Source:MGI Symbol;Acc:MGI:5610462] | 1,73 | 6,5 |
| ENSMUSG00000032089 | 5,063576894 | 0,0 | 24,0 | -7,218775319 | 0,049693667 | Il10ra | interleukin 10 receptor, alpha [Source:MGI Symbol;Acc:MGI:96538] | 0,00 | 9,0 |
| ENSMUSG00000108608 | 46,83660634 | 1,0 | 107,7 | -7,181669974 | 8,65077E-07 | RP23-412J12.1 | | #DIV/0! | #DIV/0! |
| ENSMUSG00000018919 | 8,661294203 | 0,2 | 36,6 | -7,17422362 | 0,043042394 | Tm4sf5 | transmembrane 4 superfamily member 5 [Source:MGI Symbol;Acc:MGI:1922854] | 4,33 | 30,2 |
| ENSMUSG00000051457 | 8,733673695 | 0,1 | 27,3 | -7,170831086 | 0,009571041 | Spn | sialophorin [Source:MGI Symbol;Acc:MGI:98384] | 0,10 | 6,3 |
| ENSMUSG00000089764 | 5,635308551 | 0,1 | 12,5 | -7,143251641 | 0,005428678 | Gm16580 | predicted gene 16580 [Source:MGI Symbol;Acc:MGI:4415000] | #DIV/0! | #DIV/0! |
| ENSMUSG00000059493 | 20,01080679 | 0,2 | 109,3 | -7,142445052 | 0,02763554 | Nhs | Nance-Horan syndrome (human) [Source:MGI Symbol;Acc:MGI:2684894] | 0,56 | 8,9 |
| ENSMUSG00000085786 | 106,8661729 | 0,7 | 269,9 | -7,126011434 | 5,74761E-05 | Gm15987 | predicted gene 15987 [Source:MGI Symbol;Acc:MGI:3801849] | 5,71 | 37,1 |
| ENSMUSG00000020897 | 120,831736 | 4,0 | 125,7 | -7,113227923 | 0,002117304 | Aurkb | aurora kinase B [Source:MGI Symbol;Acc:MGI:107168] | 23,68 | 49,1 |
| ENSMUSG00000001497 | 35,71653773 | 2,2 | 66,4 | -7,107305424 | 0,011515238 | Pax9 | paired box 9 [Source:MGI Symbol;Acc:MGI:97493] | 5,80 | 8,6 |
| ENSMUSG00000073274 | 9,663111117 | 0,2 | 56,1 | -7,07425568 | 0,028885757 | Gm14636 | predicted gene 14636 [Source:MGI Symbol;Acc:MGI:3641976] | #DIV/0! | #DIV/0! |
| ENSMUSG00000039304 | 13,26270009 | 0,5 | 68,4 | -7,063272678 | 0,008403466 | Tnfsf10 | tumor necrosis factor (ligand) superfamily, member 10 [Source:MGI Symbol;Acc:MGI:107414] | 0,76 | 9,1 |
| ENSMUSG00000025324 | 18,88595138 | 0,8 | 91,3 | -7,050765828 | 0,019280959 | Atp10a | ATPase, class V, type 10A [Source:MGI Symbol;Acc:MGI:1330809] | 2,65 | 15,3 |
| ENSMUSG00000098359 | 10,21697291 | 0,1 | 22,4 | -7,049812106 | 0,006314703 | Mir7649 | microRNA 7649 [Source:MGI Symbol;Acc:MGI:5562774] | #DIV/0! | #DIV/0! |
| ENSMUSG00000080775 | 16,12330285 | 0,3 | 39,8 | -7,028458361 | 2,3349E-05 | Gm6368 | predicted gene 6368 [Source:MGI Symbol;Acc:MGI:3644537] | #DIV/0! | #DIV/0! |
| ENSMUSG00000040345 | 82,13830164 | 1,0 | 155,3 | -7,011451629 | 0,000203727 | Arhgap9 | Rho GTPase activating protein 9 [Source:MGI Symbol;Acc:MGI:2143764] | 4,14 | 60,3 |
| ENSMUSG00000058147 | 27,74430755 | 3,2 | 38,6 | -6,981883943 | 0,013133221 | Xlr3c | X-linked lymphocyte-regulated 3C [Source:MGI Symbol;Acc:MGI:3047103] | 12,14 | 17,5 |
| ENSMUSG00000097669 | 4,828659516 | 0,1 | 11,0 | -6,956752385 | 0,009535772 | Gm8649 | predicted gene 8649 [Source:MGI Symbol;Acc:MGI:3644257] | #DIV/0! | #DIV/0! |
| ENSMUSG00000072109 | 150,7486416 | 2,5 | 308,2 | -6,941047588 | 1,54136E-08 | A530040E14Rik | RIKEN cDNA A530040E14 gene [Source:MGI Symbol;Acc:MGI:3612703] | 9,60 | 27,7 |
| ENSMUSG00000070332 | 12,73035661 | 0,5 | 45,1 | -6,936027605 | 0,003507645 | Trim80 | tripartite motif-containing 80 [Source:MGI Symbol;Acc:MGI:3588186] | 4,62 | 22,3 |
| ENSMUSG00000101625 | 46,33402846 | 0,7 | 84,2 | -6,929852385 | 1,51429E-06 | Gm29371 | predicted gene 29371 [Source:MGI Symbol;Acc:MGI:5580077] | #DIV/0! | #DIV/0! |
| ENSMUSG00000021998 | 209,9036953 | 5,3 | 361,1 | -6,920010044 | 0,000101356 | Lcp1 | lymphocyte cytosolic protein 1 [Source:MGI Symbol;Acc:MGI:104808] | 17,62 | 31,0 |
| ENSMUSG00000104211 | 6,017997671 | 0,2 | 18,9 | -6,919729166 | 0,002815277 | Gm37985 | predicted gene, 37985 [Source:MGI Symbol;Acc:MGI:5611213] | 0,06 | 0,2 |
| ENSMUSG00000010142 | 182,2274644 | 7,1 | 411,3 | -6,894711139 | 0,000210376 | Tnfrsf13b | tumor necrosis factor receptor superfamily, member 13b [Source:MGI Symbol;Acc:MGI:1889411] | 29,22 | 61,4 |
| ENSMUSG00000032818 | 3,205301106 | 0,1 | 8,4 | -6,890058853 | 0,032184538 | Loxhd1 | lipoxygenase homology domains 1 [Source:MGI Symbol;Acc:MGI:1914609] | 0,23 | 1,8 |
| ENSMUSG00000050550 | 76,18266643 | 1,7 | 186,7 | -6,82361864 | 9,39576E-10 | Gm11868 | predicted gene 11868 [Source:MGI Symbol;Acc:MGI:3650507] | #DIV/0! | #DIV/0! |
| ENSMUSG00000097352 | 21,05745571 | 0,2 | 38,7 | -6,818052548 | 0,030390259 | C920009B18Rik | RIKEN cDNA C920009B18 gene [Source:MGI Symbol;Acc:MGI:3583961] | #DIV/0! | #DIV/0! |
| ENSMUSG00000048534 | 145,6639565 | 3,1 | 385,8 | -6,748505923 | 0,000644921 | Amica1 | adhesion molecule, interacts with CXADR antigen 1 [Source:MGI Symbol;Acc:MGI:2685484] | 20,58 | 54,7 |
| ENSMUSG00000030867 | 94,76256401 | 3,1 | 242,9 | -6,74668107 | 1,51429E-06 | Plk1 | polo-like kinase 1 [Source:MGI Symbol;Acc:MGI:97621] | 22,69 | 78,7 |
| ENSMUSG00000025983 | 11,77875179 | 0,5 | 22,6 | -6,715105375 | 0,005444277 | Ccdc150 | coiled-coil domain containing 150 [Source:MGI Symbol;Acc:MGI:1925266] | 1,94 | 0,0 |
| ENSMUSG00000035722 | 38,89490063 | 5,7 | 105,5 | -6,712683562 | 0,015129033 | Abca7 | ATP-binding cassette, sub-family A (ABC1), member 7 [Source:MGI Symbol;Acc:MGI:1351646] | 10,35 | 19,4 |
| ENSMUSG00000094497 | 24,52475599 | 0,5 | 47,5 | -6,70382126 | 7,28907E-07 | Gm8210 | predicted pseudogene 8210 [Source:MGI Symbol;Acc:MGI:3643593] | #DIV/0! | #DIV/0! |
| ENSMUSG00000106304 | 6,947051589 | 0,1 | 39,3 | -6,66506543 | 0,049273686 | Gm42518 | predicted gene 42518 [Source:MGI Symbol;Acc:MGI:5662655] | #DIV/0! | #DIV/0! |
| ENSMUSG00000089844 | 118,5812465 | 2,3 | 178,7 | -6,659922181 | 9,78467E-08 | A530032D15Rik | RIKEN cDNA A530032D15Rik gene [Source:MGI Symbol;Acc:MGI:3037746] | 17,05 | 47,0 |
| ENSMUSG00000104277 | 3,643692635 | 0,2 | 11,7 | -6,644970664 | 0,022615306 | Gm38299 | predicted gene, 38299 [Source:MGI Symbol;Acc:MGI:5611527] | 0,81 | 3,0 |
| ENSMUSG00000082762 | 10,31145897 | 0,3 | 27,5 | -6,611657553 | 0,00113626 | Gm12366 | predicted gene 12366 [Source:MGI Symbol;Acc:MGI:3649557] | #DIV/0! | #DIV/0! |
| ENSMUSG00000095170 | 8,140924562 | 0,1 | 13,4 | -6,591882262 | 0,007556503 | Ighv8-11 | immunoglobulin heavy variable V8-11 [Source:MGI Symbol;Acc:MGI:3644925] | 6,98 | 17,1 |
| ENSMUSG00000039529 | 6,057406844 | 0,7 | 13,4 | -6,581538247 | 0,03263599 | Atp8b1 | ATPase, class I, type 8B, member 1 [Source:MGI Symbol;Acc:MGI:1859665] | 2,08 | 1,1 |
| ENSMUSG00000017204 | 21,66353433 | 0,6 | 55,4 | -6,577918916 | 0,0349724 | Gsdma | gasdermin A [Source:MGI Symbol;Acc:MGI:1889509] | 3,67 | 15,6 |
| ENSMUSG00000008496 | 116,6334102 | 1,7 | 234,9 | -6,576715932 | 9,69722E-06 | Pou2f2 | POU domain, class 2, transcription factor 2 [Source:MGI Symbol;Acc:MGI:101897] | 3,92 | 20,5 |
| ENSMUSG00000080059 | 8,227149439 | 0,6 | 38,9 | -6,574197138 | 0,000283891 | Rps19-ps3 | ribosomal protein S19, pseudogene 3 [Source:MGI Symbol;Acc:MGI:3701125] | #DIV/0! | #DIV/0! |
| ENSMUSG00000098489 | 187,162573 | 4,1 | 382,3 | -6,565290768 | 1,64725E-07 | Mir7678 | microRNA 7678 [Source:MGI Symbol;Acc:MGI:5530770] | #DIV/0! | #DIV/0! |
| ENSMUSG00000034652 | 11,49166272 | 1,0 | 59,5 | -6,564820981 | 0,013801968 | Cd300a | CD300A antigen [Source:MGI Symbol;Acc:MGI:2443411] | 4,02 | 8,7 |
| ENSMUSG00000076613 | 728,8173151 | 20,2 | 1461,6 | -6,560550826 | 2,84158E-07 | Ighg2b | immunoglobulin heavy constant gamma 2B [Source:MGI Symbol;Acc:MGI:96445] | #DIV/0! | #DIV/0! |
| ENSMUSG00000080790 | 27,3993957 | 0,7 | 64,4 | -6,555966481 | 0,000409724 | Gm7638 | predicted gene 7638 [Source:MGI Symbol;Acc:MGI:3643470] | #DIV/0! | #DIV/0! |
| ENSMUSG00000037337 | 91,70735992 | 2,0 | 137,8 | -6,533314751 | 0,007372654 | Map4k1 | mitogen-activated protein kinase kinase kinase kinase 1 [Source:MGI Symbol;Acc:MGI:1346882] | 11,72 | 33,8 |
| ENSMUSG00000052477 | 145,4851703 | 3,2 | 263,2 | -6,515575484 | 9,78467E-08 | C130026I21Rik | RIKEN cDNA C130026I21 gene [Source:MGI Symbol;Acc:MGI:3612702] | 11,46 | 20,7 |
| ENSMUSG00000104399 | 6,870791704 | 0,2 | 23,5 | -6,482433649 | 0,017886688 | Gm37963 | predicted gene, 37963 [Source:MGI Symbol;Acc:MGI:5611191] | 7,45 | 17,7 |
| ENSMUSG00000104074 | 5,32548379 | 0,2 | 21,1 | -6,475270256 | 0,005026681 | Gm37025 | predicted gene, 37025 [Source:MGI Symbol;Acc:MGI:5610253] | #DIV/0! | #DIV/0! |
| ENSMUSG00000098318 | 856,8351677 | 21,3 | 1873,9 | -6,463129956 | 4,18023E-08 | Lockd | lncRNA downstream of Cdkn1b [Source:MGI Symbol;Acc:MGI:1915081] | #DIV/0! | #DIV/0! |
| ENSMUSG00000081434 | 152,1421003 | 3,5 | 284,2 | -6,461170332 | 4,7481E-08 | Gm14165 | predicted gene 14165 [Source:MGI Symbol;Acc:MGI:3650537] | #DIV/0! | #DIV/0! |
| ENSMUSG00000067149 | 4645,357636 | 98,2 | 8893,5 | -6,454970554 | 3,01216E-10 | Jchain | immunoglobulin joining chain [Source:MGI Symbol;Acc:MGI:96493] | 94,19 | 96,1 |
| ENSMUSG00000008932 | 12,24099588 | 0,5 | 31,9 | -6,454774522 | 0,000928541 | Slc1a7 | solute carrier family 1 (glutamate transporter), member 7 [Source:MGI Symbol;Acc:MGI:2444087] | 1,56 | 2,8 |
| ENSMUSG00000039853 | 14,51328993 | 0,8 | 47,7 | -6,448006461 | 0,006045628 | Trim14 | tripartite motif-containing 14 [Source:MGI Symbol;Acc:MGI:1921985] | 6,18 | 23,9 |
| ENSMUSG00000076614 | 2651,869628 | 59,9 | 5577,7 | -6,440568215 | 2,31178E-09 | Ighg1 | immunoglobulin heavy constant gamma 1 (G1m marker) [Source:MGI Symbol;Acc:MGI:96446] | #DIV/0! | #DIV/0! |
| ENSMUSG00000040204 | 346,3719742 | 9,1 | 782,7 | -6,437569102 | 1,76712E-08 | 2810417H13Rik | RIKEN cDNA 2810417H13 gene [Source:MGI Symbol;Acc:MGI:1915276] | 32,50 | 34,0 |
| ENSMUSG00000098537 | 8,546820884 | 0,1 | 10,4 | -6,414211719 | 0,049283007 | Gm27867 | predicted gene, 27867 [Source:MGI Symbol;Acc:MGI:5531249] | 7,20 | 43,7 |
| ENSMUSG00000096150 | 57,68876008 | 1,2 | 144,0 | -6,413816296 | 7,86835E-06 | Ighv1-85 | immunoglobulin heavy variable 1-85 [Source:MGI Symbol;Acc:MGI:3645723] | 34,57 | 49,7 |
| ENSMUSG00000069117 | 1243,719348 | 32,1 | 2564,7 | -6,395888864 | 1,46039E-09 | Gm10260 | predicted gene 10260 [Source:MGI Symbol;Acc:MGI:3642298] | 36,38 | 37,3 |
| ENSMUSG00000097346 | 1243,821567 | 32,1 | 2564,7 | -6,395888799 | 1,46039E-09 | Gm26619 | predicted gene, 26619 [Source:MGI Symbol;Acc:MGI:5477113] | #DIV/0! | #DIV/0! |
| ENSMUSG00000090054 | 5,478523857 | 0,3 | 15,4 | -6,392471953 | 0,004328655 | Gm15629 | predicted gene 15629 [Source:MGI Symbol;Acc:MGI:3783073] | 1,68 | 8,8 |

| | | | | | | | | | |
|---|---|---|---|---|---|---|---|---|---|
| ENSMUSG00000105263 | 69,75091135 | 1,7 | 142,8 | -6,389002995 | 1,34963E-07 | Gm42427 | predicted gene 42427 [Source:MGI Symbol;Acc:MGI:5662564] | 4,45 | 23,4 |
| ENSMUSG00000026480 | 18,92249924 | 3,1 | 49,9 | -6,386873319 | 0,028922607 | Ncf2 | neutrophil cytosolic factor 2 [Source:MGI Symbol;Acc:MGI:97284] | 9,83 | 11,9 |
| ENSMUSG00000001228 | 87,1737505 | 6,8 | 301,6 | -6,380149849 | 0,010750274 | Uhrf1 | ubiquitin-like, containing PHD and RING finger domains, 1 [Source:MGI Symbol;Acc:MGI:1338889] | 23,31 | 47,0 |
| ENSMUSG00000009145 | 103,6815214 | 1,6 | 134,6 | -6,378872635 | 1,89182E-07 | Dqx1 | DEAQ RNA-dependent ATPase [Source:MGI Symbol;Acc:MGI:2136388] | 4,52 | 5,9 |
| ENSMUSG00000022033 | 81,46308349 | 7,8 | 371,2 | -6,37595782 | 9,35276E-05 | Pbk | PDZ binding kinase [Source:MGI Symbol;Acc:MGI:1289156] | 38,23 | 71,8 |
| ENSMUSG00000076620 | 59,2109534 | 1,1 | 98,5 | -6,374168972 | 4,7481E-08 | Ighj2 | immunoglobulin heavy joining 2 [Source:MGI Symbol;Acc:MGI:4439803] | 87,50 | 98,6 |
| ENSMUSG00000076604 | 236,4426664 | 6,7 | 528,1 | -6,366250992 | 5,22092E-08 | Igkj1 | immunoglobulin kappa joining 1 [Source:MGI Symbol;Acc:MGI:1316689] | 100,00 | 100,0 |
| ENSMUSG00000078249 | 81,36182666 | 1,4 | 102,9 | -6,361050312 | 4,88823E-06 | Hmga1-rs1 | high mobility group AT-hook I, related sequence 1 [Source:MGI Symbol;Acc:MGI:96161] | 10,31 | 26,1 |
| ENSMUSG00000081640 | 70,71666038 | 1,6 | 116,7 | -6,343553428 | 6,21322E-06 | Rplp0-ps1 | ribosomal protein, large, P0, pseudogene 1 [Source:MGI Symbol;Acc:MGI:3648599] | #DIV/0! | #DIV/0! |
| ENSMUSG00000100922 | 13,65742887 | 0,4 | 32,7 | -6,343362268 | 4,55509E-05 | Gm8520 | predicted gene 8520 [Source:MGI Symbol;Acc:MGI:3645038] | #DIV/0! | #DIV/0! |
| ENSMUSG00000062353 | 1924,960682 | 49,3 | 3887,1 | -6,342087063 | 5,6048E-09 | Gm15772 | predicted gene 15772 [Source:MGI Symbol;Acc:MGI:3805541] | #DIV/0! | #DIV/0! |
| ENSMUSG00000062365 | 3,332642291 | 0,1 | 6,4 | -6,340590824 | 0,041551111 | Gm4968 | predicted gene 4968 [Source:MGI Symbol;Acc:MGI:3647516] | #DIV/0! | #DIV/0! |
| ENSMUSG00000032496 | 1370,848603 | 37,5 | 3016,6 | -6,332260807 | 5,6048E-09 | Ltf | lactotransferrin [Source:MGI Symbol;Acc:MGI:96837] | 26,85 | 27,4 |
| ENSMUSG00000052760 | 52,30959565 | 1,0 | 62,5 | -6,31671842 | 0,008531258 | A630001G21Rik | RIKEN cDNA A630001G21 gene [Source:MGI Symbol;Acc:MGI:2443131] | 6,78 | 15,5 |
| ENSMUSG00000107383 | 394,0129962 | 9,7 | 744,8 | -6,303601211 | 4,88775E-09 | Gm4366 | predicted gene 4366 [Source:MGI Symbol;Acc:MGI:3782551] | #DIV/0! | #DIV/0! |
| ENSMUSG00000057244 | 3,439718934 | 0,1 | 6,3 | -6,296364547 | 0,032148443 | Gm6139 | predicted gene 6139 [Source:MGI Symbol;Acc:MGI:3643279] | #DIV/0! | #DIV/0! |
| ENSMUSG00000017002 | 1640,604775 | 43,2 | 3467,8 | -6,292994457 | 1,28744E-08 | Slpi | secretory leukocyte peptidase inhibitor [Source:MGI Symbol;Acc:MGI:109297] | 62,06 | 67,7 |
| ENSMUSG00000104452 | 29,3970038 | 0,7 | 55,4 | -6,292346895 | 0,000303492 | Ighv8-8 | immunoglobulin heavy variable 8-8 [Source:MGI Symbol;Acc:MGI:3815333] | 24,21 | 39,3 |
| ENSMUSG00000068151 | 10,90004523 | 0,5 | 61,0 | -6,289268568 | 0,01756774 | A230006K03Rik | RIKEN cDNA A230006K03 gene [Source:MGI Symbol;Acc:MGI:3644205] | #DIV/0! | #DIV/0! |
| ENSMUSG00000019737 | 75,78875169 | 3,4 | 151,8 | -6,287759095 | 0,000240449 | Syne4 | spectrin repeat containing, nuclear envelope family member 4 [Source:MGI Symbol;Acc:MGI:2141950] | 35,60 | 63,9 |
| ENSMUSG00000065701 | 3,804881418 | 0,1 | 5,9 | -6,250361352 | 0,033465884 | Rny1 | RNA, Y1 small cytoplasmic, Ro-associated [Source:MGI Symbol;Acc:MGI:97995] | 18,15 | 72,9 |
| ENSMUSG00000096979 | 5,804015823 | 0,2 | 10,8 | -6,240328756 | 0,015746369 | Gm26880 | predicted gene, 26880 [Source:MGI Symbol;Acc:MGI:5477374] | #DIV/0! | #DIV/0! |
| ENSMUSG00000005320 | 119,8003179 | 2,9 | 226,3 | -6,232624855 | 7,26463E-10 | Fgfr4 | fibroblast growth factor receptor 4 [Source:MGI Symbol;Acc:MGI:95525] | 0,28 | 1,0 |
| ENSMUSG00000094117 | 447,6668683 | 11,2 | 883,0 | -6,229705193 | 1,54136E-08 | Igkv3-12 | immunoglobulin kappa variable 3-12 [Source:MGI Symbol;Acc:MGI:1330815] | 98,61 | 98,6 |
| ENSMUSG00000093894 | 28,13106711 | 0,4 | 40,8 | -6,226218544 | 0,000101356 | Ighv1-53 | immunoglobulin heavy variable 1-53 [Source:MGI Symbol;Acc:MGI:3576502] | 21,65 | 43,9 |
| ENSMUSG00000040592 | 31,56696221 | 1,9 | 105,4 | -6,223401248 | 0,006502921 | Cd79b | CD79B antigen [Source:MGI Symbol;Acc:MGI:96431] | 18,15 | 42,8 |
| ENSMUSG00000097694 | 88,39497134 | 1,7 | 35,5 | -6,215683959 | 0,00764456 | G730013B05Rik | RIKEN cDNA G730013B05 gene [Source:MGI Symbol;Acc:MGI:3588276] | #DIV/0! | #DIV/0! |
| ENSMUSG00000095007 | 594,4280617 | 15,1 | 1242,9 | -6,196879145 | 2,32518E-09 | Igkv12-41 | immunoglobulin kappa chain variable 12-41 [Source:MGI Symbol;Acc:MGI:4439772] | 97,98 | 98,0 |
| ENSMUSG00000061132 | 881,9158537 | 23,7 | 1635,8 | -6,183567196 | 1,16098E-07 | Blnk | B cell linker [Source:MGI Symbol;Acc:MGI:96878] | 67,29 | 85,4 |
| ENSMUSG00000105233 | 155,4271588 | 3,8 | 271,2 | -6,181611833 | 1,49817E-07 | Gm20568 | predicted gene, 20568 [Source:MGI Symbol;Acc:MGI:5295675] | #DIV/0! | #DIV/0! |
| ENSMUSG00000023439 | 39,41384388 | 3,8 | 214,5 | -6,170293395 | 1,02311E-06 | Gnb3 | guanine nucleotide binding protein (G protein), beta 3 [Source:MGI Symbol;Acc:MGI:95785] | 24,61 | 52,3 |
| ENSMUSG00000097698 | 71,90655763 | 2,0 | 156,8 | -6,164286264 | 1,0658E-09 | Gm26906 | predicted gene, 26906 [Source:MGI Symbol;Acc:MGI:5477400] | #DIV/0! | #DIV/0! |
| ENSMUSG00000106567 | 131,6466548 | 3,1 | 243,3 | -6,155325747 | 4,30787E-06 | 2010309G21Rik | RIKEN cDNA 2010309G21 gene [Source:MGI Symbol;Acc:MGI:1917268] | 19,43 | 20,3 |
| ENSMUSG00000033220 | 583,8489667 | 17,9 | 1128,8 | -6,142470952 | 7,64774E-09 | Rac2 | RAS-related C3 botulinum substrate 2 [Source:MGI Symbol;Acc:MGI:97846] | 38,31 | 73,1 |
| ENSMUSG00000107730 | 299,1757151 | 9,0 | 629,3 | -6,130657396 | 8,96005E-07 | Gm44238 | predicted gene, 44238 [Source:MGI Symbol;Acc:MGI:5690630] | #DIV/0! | #DIV/0! |
| ENSMUSG00000076609 | 4968,129246 | 144,7 | 10402,2 | -6,115606718 | 7,26463E-10 | Igkc | immunoglobulin kappa constant [Source:MGI Symbol;Acc:MGI:96495] | #DIV/0! | #DIV/0! |
| ENSMUSG00000105231 | 33,332692 | 0,7 | 52,2 | -6,10809446 | 1,19892E-06 | Iglj3 | immunoglobulin lambda joining 3 [Source:MGI Symbol;Acc:MGI:5293407] | 92,11 | 96,5 |
| ENSMUSG00000105353 | 7,614658035 | 0,2 | 8,5 | -6,076731848 | 0,015962962 | Gm42428 | predicted gene 42428 [Source:MGI Symbol;Acc:MGI:5662565] | #DIV/0! | #DIV/0! |
| ENSMUSG00000086454 | 8,696577107 | 1,5 | 48,6 | -6,035105292 | 0,027265477 | Platr14 | pluripotency associated transcript 14 [Source:MGI Symbol;Acc:MGI:1922209] | #DIV/0! | #DIV/0! |
| ENSMUSG00000097906 | 489,0636633 | 14,5 | 937,6 | -6,030105184 | 3,15634E-11 | Gm9625 | predicted gene 9625 [Source:MGI Symbol;Acc:MGI:3780033] | #DIV/0! | #DIV/0! |
| ENSMUSG00000101609 | 39,04327497 | 2,3 | 170,6 | -6,026776764 | 0,000560297 | Kcnq1ot1 | KCNQ1 overlapping transcript 1 [Source:MGI Symbol;Acc:MGI:1926855] | 0,36 | 1,3 |
| ENSMUSG00000076576 | 623,686119 | 21,1 | 1314,9 | -6,019470699 | 3,02708E-07 | Igkv6-32 | immunoglobulin kappa variable 6-32 [Source:MGI Symbol;Acc:MGI:3641634] | 97,54 | 99,2 |
| ENSMUSG00000024014 | 1595,135403 | 44,5 | 2725,6 | -6,017534084 | 2,06622E-08 | Pim1 | proviral integration site 1 [Source:MGI Symbol;Acc:MGI:97584] | 82,63 | 95,0 |
| ENSMUSG00000032053 | 2262,980425 | 76,1 | 4579,7 | -5,99665777 | 1,35675E-07 | Pou2af1 | POU domain, class 2, associating factor 1 [Source:MGI Symbol;Acc:MGI:105086] | 90,09 | 100,0 |
| ENSMUSG00000086320 | 10,46073938 | 0,4 | 22,4 | -5,994527269 | 0,000414415 | Gm12840 | predicted gene 12840 [Source:MGI Symbol;Acc:MGI:3650917] | #DIV/0! | #DIV/0! |
| ENSMUSG00000063696 | 669,7730701 | 20,1 | 1271,9 | -5,988463517 | 2,32518E-09 | Gm8730 | predicted pseudogene 8730 [Source:MGI Symbol;Acc:MGI:3644565] | 25,02 | 33,4 |
| ENSMUSG00000012819 | 8,498122226 | 0,6 | 37,5 | -5,975308478 | 0,006571721 | Cdh23 | cadherin 23 (otocadherin) [Source:MGI Symbol;Acc:MGI:1890219] | 0,53 | 2,4 |
| ENSMUSG00000078161 | 109,7129435 | 6,1 | 205,5 | -5,969168808 | 0,003186092 | Erich3 | glutamate rich 3 [Source:MGI Symbol;Acc:MGI:1919095] | 11,75 | 17,4 |
| ENSMUSG00000083773 | 244,196087 | 5,6 | 373,7 | -5,966596 | 1,11176E-05 | Gm13394 | predicted gene 13394 [Source:MGI Symbol;Acc:MGI:3651848] | #DIV/0! | #DIV/0! |
| ENSMUSG00000085666 | 391,7156593 | 14,2 | 759,4 | -5,954775241 | 5,58598E-06 | Gm9855 | predicted pseudogene 9855 [Source:MGI Symbol;Acc:MGI:3704357] | #DIV/0! | #DIV/0! |
| ENSMUSG00000098183 | 12,25859872 | 0,2 | 7,4 | -5,95299946 | 0,030440412 | Gm27010 | predicted gene, 27010 [Source:MGI Symbol;Acc:MGI:5504125] | #DIV/0! | #DIV/0! |
| ENSMUSG00000063816 | 5,594166646 | 0,5 | 12,9 | -5,946721809 | 0,012541575 | Gm4987 | predicted gene 4987 [Source:MGI Symbol;Acc:MGI:3647751] | #DIV/0! | #DIV/0! |
| ENSMUSG00000065145 | 4,623504159 | 0,2 | 12,4 | -5,946060263 | 0,014677393 | Vaultrc5 | vault RNA component 5 [Source:MGI Symbol;Acc:MGI:2673990] | 30,30 | 79,0 |
| ENSMUSG00000105547 | 431,8053344 | 12,3 | 788,1 | -5,945031656 | 3,02708E-07 | Iglc3 | immunoglobulin lambda constant 3 [Source:MGI Symbol;Acc:MGI:99886] | #DIV/0! | #DIV/0! |
| ENSMUSG00000017716 | 387,9696244 | 11,9 | 741,3 | -5,941913664 | 6,42969E-11 | Birc5 | baculoviral IAP repeat-containing 5 [Source:MGI Symbol;Acc:MGI:1203517] | 22,25 | 26,5 |
| ENSMUSG00000024353 | 1289,981283 | 44,1 | 2498,4 | -5,938617439 | 6,36686E-07 | Mzb1 | marginal zone B and B1 cell-specific protein 1 [Source:MGI Symbol;Acc:MGI:1917066] | 99,78 | 100,0 |
| ENSMUSG00000044349 | 2213,765193 | 111,4 | 5430,4 | -5,935979557 | 3,39498E-15 | Snhg11 | small nucleolar RNA host gene 11 [Source:MGI Symbol;Acc:MGI:2441845] | 69,40 | 94,0 |
| ENSMUSG00000107470 | 73,01920442 | 2,3 | 137,1 | -5,934936417 | 8,80139E-08 | Gm3375 | predicted gene 3375 [Source:MGI Symbol;Acc:MGI:3781553] | #DIV/0! | #DIV/0! |
| ENSMUSG00000102117 | 3,832012889 | 0,2 | 12,5 | -5,93346134 | 0,006367725 | Rpsa-ps1 | ribosomal protein SA, pseudogene 1 [Source:MGI Symbol;Acc:MGI:3644886] | #DIV/0! | #DIV/0! |
| ENSMUSG00000017861 | 61,71543548 | 1,5 | 35,6 | -5,918737011 | 0,001952593 | Mybl2 | myeloblastosis oncogene-like 2 [Source:MGI Symbol;Acc:MGI:101785] | 6,13 | 13,0 |
| ENSMUSG00000100037 | 86,97611107 | 3,0 | 171,4 | -5,914353993 | 3,30532E-06 | Gm29103 | predicted gene 29103 [Source:MGI Symbol;Acc:MGI:5579809] | #DIV/0! | #DIV/0! |
| ENSMUSG00000000628 | 114,4873591 | 5,1 | 353,1 | -5,908049066 | 3,20288E-05 | Hk2 | hexokinase 2 [Source:MGI Symbol;Acc:MGI:1315197] | 9,98 | 22,3 |
| ENSMUSG00000030798 | 95,45559866 | 5,0 | 295,5 | -5,897947528 | 1,29366E-05 | Cd37 | CD37 antigen [Source:MGI Symbol;Acc:MGI:88330] | 38,55 | 88,0 |
| ENSMUSG00000107379 | 2474,5358 | 78,1 | 4706,2 | -5,893145649 | 3,47477E-09 | Gm43126 | predicted gene 43126 [Source:MGI Symbol;Acc:MGI:5663263] | 86,71 | 86,7 |
| ENSMUSG00000032332 | 4,46011723 | 0,5 | 8,3 | -5,885580975 | 0,042239085 | Col12a1 | collagen, type XII, alpha 1 [Source:MGI Symbol;Acc:MGI:88448] | 0,00 | 1,3 |
| ENSMUSG00000095123 | 28,16863709 | 1,3 | 101,7 | -5,876963217 | 0,010820707 | Gm21781 | predicted gene, 21781 [Source:MGI Symbol;Acc:MGI:5433945] | #DIV/0! | #DIV/0! |
| ENSMUSG00000048865 | 438,4891033 | 19,4 | 1103,5 | -5,870050698 | 3,81875E-07 | Arhgap30 | Rho GTPase activating protein 30 [Source:MGI Symbol;Acc:MGI:2684948] | 43,93 | 81,3 |
| ENSMUSG00000104975 | 37,47205434 | 1,0 | 80,5 | -5,862523853 | 9,35276E-05 | Iglj2 | immunoglobulin lambda joining 2 [Source:MGI Symbol;Acc:MGI:5293409] | 45,61 | 79,8 |
| ENSMUSG00000026548 | 188,0421509 | 10,1 | 567,5 | -5,8564339 | 0,001311604 | Slamf9 | SLAM family member 9 [Source:MGI Symbol;Acc:MGI:1923692] | 55,39 | 91,4 |
| ENSMUSG00000109208 | 77,49835453 | 4,5 | 247,1 | -5,840201777 | 4,81571E-06 | Cd37 | CD37 antigen [Source:MGI Symbol;Acc:MGI:88330] | 43,31 | 86,0 |
| ENSMUSG00000078956 | 30,77470349 | 1,1 | 66,5 | -5,839781253 | 3,31881E-08 | Gm14221 | predicted gene 14221 [Source:MGI Symbol;Acc:MGI:3650696] | 32,55 | 41,2 |
| ENSMUSG00000074802 | 206,0412526 | 5,4 | 203,8 | -5,820035366 | 0,00025459 | Gas2l3 | growth arrest-specific 2 like 3 [Source:MGI Symbol;Acc:MGI:1918780] | 10,76 | 23,6 |
| ENSMUSG00000092674 | 21,01549508 | 0,8 | 39,3 | -5,81797658 | 0,000247061 | Gm24105 | predicted gene, 24105 [Source:MGI Symbol;Acc:MGI:5453882] | 22,18 | 45,6 |
| ENSMUSG00000087658 | 18,46817933 | 0,5 | 19,3 | -5,814813313 | 0,012060102 | Hotairm1 | Hoxa transcript antisense RNA, myeloid-specific 1 [Source:MGI Symbol;Acc:MGI:3705155] | 7,09 | 19,3 |
| ENSMUSG00000006585 | 103,1069119 | 8,2 | 377,9 | -5,809684302 | 0,001402484 | Cdt1 | chromatin licensing and DNA replication factor 1 [Source:MGI Symbol;Acc:MGI:1914427] | 41,57 | 67,2 |
| ENSMUSG00000055413 | 7,512001782 | 0,3 | 24,2 | -5,807642106 | 0,018594281 | H2-Q5 | histocompatibility 2, Q region locus 5 [Source:MGI Symbol;Acc:MGI:95934] | #DIV/0! | #DIV/0! |
| ENSMUSG00000018983 | 2916,310894 | 99,7 | 5206,7 | -5,790292071 | 1,80466E-07 | E2f2 | E2F transcription factor 2 [Source:MGI Symbol;Acc:MGI:1096341] | 81,85 | 85,7 |
| ENSMUSG00000073628 | 25,28750506 | 0,7 | 38,4 | -5,787046751 | 5,47741E-05 | Gm10552 | predicted gene 10552 [Source:MGI Symbol;Acc:MGI:3642083] | 3,35 | 7,8 |
| ENSMUSG00000018166 | 9,846779722 | 0,9 | 7,0 | -5,777290363 | 0,040706087 | Erbb3 | erb-b2 receptor tyrosine kinase 3 [Source:MGI Symbol;Acc:MGI:95411] | 2,68 | 1,1 |
| ENSMUSG00000041431 | 150,6392189 | 7,0 | 194,0 | -5,767224936 | 1,57793E-05 | Ccnb1 | cyclin B1 [Source:MGI Symbol;Acc:MGI:88302] | 28,62 | 46,4 |
| ENSMUSG00000076939 | 22,45651384 | 1,0 | 57,3 | -5,766878846 | 0,001631465 | Iglv3 | immunoglobulin lambda variable 3 [Source:MGI Symbol;Acc:MGI:3645509] | 35,70 | 52,5 |
| ENSMUSG00000076605 | 108,7654962 | 3,6 | 194,4 | -5,759682415 | 7,80062E-07 | Igkj2 | immunoglobulin kappa joining 2 [Source:MGI Symbol;Acc:MGI:1316690] | 100,00 | 100,0 |
| ENSMUSG00000070031 | 263,3126634 | 10,2 | 589,0 | -5,758885811 | 7,74547E-07 | Sp140 | Sp140 nuclear body protein [Source:MGI Symbol;Acc:MGI:3702467] | 30,64 | 58,0 |
| ENSMUSG00000096452 | 10,61140679 | 0,5 | 22,5 | -5,756572052 | 0,003390135 | Ighv1-77 | immunoglobulin heavy variable 1-77 [Source:MGI Symbol;Acc:MGI:4439670] | 12,32 | 27,8 |
| ENSMUSG00000097384 | 129,2717924 | 5,0 | 273,1 | -5,756342145 | 7,28907E-07 | Gm26815 | predicted gene, 26815 [Source:MGI Symbol;Acc:MGI:5477309] | #DIV/0! | #DIV/0! |
| ENSMUSG00000032218 | 152,748582 | 4,9 | 78,0 | -5,742788193 | 0,004908134 | Ccnb2 | cyclin B2 [Source:MGI Symbol;Acc:MGI:88311] | 29,50 | 42,5 |
| ENSMUSG00000053158 | 74,27548128 | 4,1 | 215,4 | -5,741454004 | 0,001506164 | Fes | feline sarcoma oncogene [Source:MGI Symbol;Acc:MGI:95514] | 13,15 | 25,7 |
| ENSMUSG00000020475 | 1233,276676 | 38,3 | 2082,6 | -5,737620063 | 8,139E-09 | Pgam2 | phosphoglycerate mutase 2 [Source:MGI Symbol;Acc:MGI:1933118] | 97,82 | 98,5 |
| ENSMUSG00000095597 | 206,7010198 | 8,1 | 435,6 | -5,731316322 | 3,50653E-09 | Gm6472 | predicted pseudogene 6472 [Source:MGI Symbol;Acc:MGI:3648396] | #DIV/0! | #DIV/0! |
| ENSMUSG00000037638 | 35,94762579 | 1,0 | 29,4 | -5,730295742 | 0,024493876 | Zbtb42 | zinc finger and BTB domain containing 42 [Source:MGI Symbol;Acc:MGI:3644133] | 3,96 | 12,1 |
| ENSMUSG00000053604 | 148,0877698 | 6,5 | 351,9 | -5,728685898 | 8,20261E-06 | Rpia | ribose 5-phosphate isomerase A [Source:MGI Symbol;Acc:MGI:103254] | 31,20 | 39,9 |
| ENSMUSG00000076937 | 142,1108727 | 2,8 | 246,4 | -5,716544931 | 0,000339105 | Iglc2 | immunoglobulin lambda constant 2 [Source:MGI Symbol;Acc:MGI:99547] | 35,20 | 80,1 |
| ENSMUSG00000054418 | 31,17823409 | 2,2 | 103,2 | -5,716095542 | 0,000674007 | 2900041M22Rik | RIKEN cDNA 2900041M22 gene [Source:MGI Symbol;Acc:MGI:1925653] | #DIV/0! | #DIV/0! |
| ENSMUSG00000038352 | 894,2178775 | 34,0 | 1847,5 | -5,712408411 | 4,73487E-08 | Arl5c | ADP-ribosylation factor-like 5C [Source:MGI Symbol;Acc:MGI:3028577] | 52,66 | 51,8 |
| ENSMUSG00000072770 | 8,471025971 | 0,5 | 45,9 | -5,706151512 | 0,04289021 | Acrbp | proacrosin binding protein [Source:MGI Symbol;Acc:MGI:1859515] | 3,85 | 8,6 |
| ENSMUSG00000004891 | 167,8283401 | 10,7 | 360,2 | -5,688457656 | 0,001792169 | Nes | nestin [Source:MGI Symbol;Acc:MGI:101784] | 24,83 | 39,3 |
| ENSMUSG00000023505 | 158,8414388 | 10,7 | 350,3 | -5,680066308 | 0,000506281 | Cdca3 | cell division cycle associated 3 [Source:MGI Symbol;Acc:MGI:1315198] | 43,72 | 40,3 |
| ENSMUSG00000103880 | 95,30779947 | 4,0 | 209,6 | -5,673383531 | 2,08003E-06 | Gm37009 | predicted gene, 37009 [Source:MGI Symbol;Acc:MGI:5610237] | #DIV/0! | #DIV/0! |
| ENSMUSG00000102700 | 173,4656237 | 6,7 | 330,0 | -5,671101933 | 1,00938E-06 | Gm38312 | predicted gene, 38312 [Source:MGI Symbol;Acc:MGI:5611540] | 6,66 | 8,4 |
| ENSMUSG00000048087 | 24,55938153 | 0,9 | 53,2 | -5,655375969 | 0,000740471 | Gm4737 | predicted gene 4737 [Source:MGI Symbol;Acc:MGI:3643647] | 6,33 | 8,9 |
| ENSMUSG00000097125 | 381,8850143 | 12,6 | 594,1 | -5,654600937 | 2,03568E-07 | Gm26885 | predicted gene, 26885 [Source:MGI Symbol;Acc:MGI:5477379] | #DIV/0! | #DIV/0! |
| ENSMUSG00000002297 | 848,0934016 | 35,2 | 1734,5 | -5,643042967 | 1,35675E-07 | Dbf4 | DBF4 zinc finger [Source:MGI Symbol;Acc:MGI:1351328] | 75,15 | 83,3 |
| ENSMUSG00000050332 | 444,5329074 | 19,7 | 988,1 | -5,642635584 | 7,30123E-12 | Amer1 | APC membrane recruitment 1 [Source:MGI Symbol;Acc:MGI:1919595] | 24,52 | 32,0 |
| ENSMUSG00000105827 | 9,700197019 | 0,7 | 42,9 | -5,584592842 | 0,014533827 | Hist2h2bb | histone cluster 2, H2bb [Source:MGI Symbol;Acc:MGI:2448413] | 16,62 | 41,2 |
| ENSMUSG00000093283 | 25,84562218 | 1,3 | 56,7 | -5,583137902 | 0,000492468 | Gm22567 | predicted gene, 22567 [Source:MGI Symbol;Acc:MGI:5452344] | 22,12 | 38,0 |
| ENSMUSG00000081603 | 82,0868823 | 3,4 | 161,4 | -5,57981788 | 1,42091E-07 | Gm14681 | predicted gene 14681 [Source:MGI Symbol;Acc:MGI:3705734] | #DIV/0! | #DIV/0! |
| ENSMUSG00000099250 | 6,577145913 | 0,2 | 12,2 | -5,571137052 | 0,006282973 | Rn7s2 | 7S RNA 2 [Source:MGI Symbol;Acc:MGI:97953] | 5,89 | 39,0 |
| ENSMUSG00000001517 | 98,50452213 | 4,2 | 198,9 | -5,564269707 | 1,13441E-06 | Foxm1 | forkhead box M1 [Source:MGI Symbol;Acc:MGI:1347487] | 16,59 | 40,9 |
| ENSMUSG00000026126 | 209,4792757 | 9,3 | 378,6 | -5,563374668 | 9,92483E-12 | Ptpn18 | protein tyrosine phosphatase, non-receptor type 18 [Source:MGI Symbol;Acc:MGI:108410] | 34,54 | 32,2 |
| ENSMUSG00000071713 | 139,7185834 | 2,9 | 435,3 | -5,56130907 | 0,001437604 | Csf2rb | stimulating factor 2 receptor, beta, low-affinity (granulocyte-macrophage) [Source:MGI Symbol;Acc:MGI:13 | 9,59 | 33,4 |
| ENSMUSG00000100801 | 398,8861067 | 15,2 | 742,5 | -5,560928033 | 7,01353E-09 | Gm15459 | predicted gene 15459 [Source:MGI Symbol;Acc:MGI:3705702] | #DIV/0! | #DIV/0! |
| ENSMUSG00000082145 | 24,07253664 | 1,0 | 52,6 | -5,545080425 | 6,29267E-05 | Gm12312 | predicted gene 12312 [Source:MGI Symbol;Acc:MGI:3649321] | #DIV/0! | #DIV/0! |
| ENSMUSG00000105646 | 65,84611698 | 1,8 | 133,0 | -5,534784717 | 0,002149466 | Gm43291 | predicted gene 43291 [Source:MGI Symbol;Acc:MGI:5663428] | #DIV/0! | #DIV/0! |
| ENSMUSG00000001403 | 123,4131261 | 5,8 | 290,9 | -5,525039436 | 0,000216121 | Ube2c | ubiquitin-conjugating enzyme E2C [Source:MGI Symbol;Acc:MGI:1915862] | 23,78 | 33,7 |
| ENSMUSG00000022584 | 49,27001936 | 1,8 | 145,7 | -5,524447361 | 0,009933686 | Ly6c2 | lymphocyte antigen 6 complex, locus C2 [Source:MGI Symbol;Acc:MGI:3712069] | 26,46 | 53,8 |
| ENSMUSG00000106874 | 1350,483045 | 55,7 | 2556,4 | -5,497546316 | 7,01353E-09 | Gm20186 | predicted gene, 20186 [Source:MGI Symbol;Acc:MGI:5012371] | #DIV/0! | #DIV/0! |
| ENSMUSG00000026950 | 9,129082538 | 0,5 | 9,4 | -5,473935557 | 0,009087284 | Neb | nebulin [Source:MGI Symbol;Acc:MGI:97292] | 0,15 | 0,9 |
| ENSMUSG00000100280 | 93,71739176 | 4,8 | 180,4 | -5,471510486 | 9,80902E-13 | Gm28417 | predicted gene 28417 [Source:MGI Symbol;Acc:MGI:5579123] | 20,87 | 23,9 |
| ENSMUSG00000061393 | 400,5856393 | 21,1 | 1020,8 | -5,467700396 | 2,96566E-09 | Acvr2b | activin receptor IIB [Source:MGI Symbol;Acc:MGI:87912] | 89,16 | 94,8 |
| ENSMUSG00000000184 | 21813,72757 | 905,0 | 39873,2 | -5,456549541 | 2,9364E-09 | Ccnd2 | cyclin D2 [Source:MGI Symbol;Acc:MGI:88314] | 94,55 | 93,2 |
| ENSMUSG00000104280 | 5,751314644 | 0,3 | 17,5 | -5,455297503 | 0,011917528 | Rps2-ps11 | ribosomal protein S2, pseudogene 11 [Source:MGI Symbol;Acc:MGI:3648819] | #DIV/0! | #DIV/0! |
| ENSMUSG00000026447 | 33,23263492 | 1,7 | 70,8 | -5,447931287 | 4,50341E-05 | Pik3c2b | phosphoinositide-3-kinase, class 2, beta polypeptide [Source:MGI Symbol;Acc:MGI:2685045] | 3,06 | 7,5 |
| ENSMUSG00000021356 | 1646,083425 | 51,2 | 2385,5 | -5,441477243 | 2,76801E-06 | Irf4 | interferon regulatory factor 4 [Source:MGI Symbol;Acc:MGI:1096873] | 73,98 | 71,6 |
| ENSMUSG00000044468 | 2005,40503 | 80,3 | 3776,5 | -5,432856356 | 1,46039E-09 | Fam46c | family with sequence similarity 46, member C [Source:MGI Symbol;Acc:MGI:1921895] | 78,69 | 80,7 |
| ENSMUSG00000033871 | 133,6181269 | 6,2 | 317,2 | -5,421258339 | 0,000210376 | Ppargc1b | roxisome proliferative activated receptor, gamma, coactivator 1 beta [Source:MGI Symbol;Acc:MGI:24449 | 24,81 | 52,0 |
| ENSMUSG00000079033 | 288,495623 | 9,9 | 418,0 | -5,411542861 | 3,63141E-08 | Mef2b | myocyte enhancer factor 2B [Source:MGI Symbol;Acc:MGI:104526] | 67,19 | 83,3 |
| ENSMUSG00000092746 | 41,31289728 | 1,8 | 79,4 | -5,406511728 | 0,000253433 | Rn7s6 | 7S RNA 6 [Source:MGI Symbol;Acc:MGI:97956] | 26,12 | 33,7 |
| ENSMUSG00000026605 | 224,9755947 | 10,8 | 380,2 | -5,40117495 | 8,80139E-08 | Cenpf | centromere protein F [Source:MGI Symbol;Acc:MGI:1313302] | 9,50 | 22,8 |
| ENSMUSG00000108688 | 6,444120645 | 0,5 | 14,4 | -5,394431034 | 0,01628919 | RP23-136G1.3 | | 5,31 | 11,1 |
| ENSMUSG00000088025 | 155,7962019 | 5,6 | 237,3 | -5,390915518 | 5,93596E-07 | Rprl3 | ribonuclease P RNA-like 3 [Source:MGI Symbol;Acc:MGI:105103] | 35,52 | 53,0 |
| ENSMUSG00000003283 | 274,7728559 | 12,4 | 628,2 | -5,385997626 | 2,6287E-05 | Hck | hemopoietic cell kinase [Source:MGI Symbol;Acc:MGI:96052] | 50,13 | 74,0 |
| ENSMUSG00000092837 | 580,9036049 | 22,4 | 930,6 | -5,380453495 | 3,08491E-06 | Rpph1 | ribonuclease P RNA component H1 [Source:MGI Symbol;Acc:MGI:1934664] | 89,24 | 98,3 |
| ENSMUSG00000024456 | 1009,772227 | 41,3 | 1900,7 | -5,365631128 | 6,45349E-08 | Diaph1 | diaphanous related formin 1 [Source:MGI Symbol;Acc:MGI:1194490] | 56,90 | 62,2 |
| ENSMUSG00000097815 | 1453,918419 | 63,0 | 2715,6 | -5,361196632 | 9,46005E-09 | Gm26809 | predicted gene, 26809 [Source:MGI Symbol;Acc:MGI:5477303] | #DIV/0! | #DIV/0! |
| ENSMUSG00000091957 | 128,0053198 | 6,4 | 269,8 | -5,356982094 | 1,55505E-06 | Rps2-ps10 | ribosomal protein S2, pseudogene 10 [Source:MGI Symbol;Acc:MGI:3645604] | #DIV/0! | #DIV/0! |
| ENSMUSG00000027868 | 39,42344337 | 2,4 | 223,6 | -5,352596236 | 0,021038504 | Tbx15 | T-box 15 [Source:MGI Symbol;Acc:MGI:1277234] | 8,54 | 25,5 |
| ENSMUSG00000034634 | 32,46709091 | 1,8 | 92,6 | -5,349724657 | 0,011702426 | Ly6d | lymphocyte antigen 6 complex, locus D [Source:MGI Symbol;Acc:MGI:96881] | 28,81 | 61,7 |
| ENSMUSG00000027326 | 29,83191581 | 1,0 | 159,8 | -5,343644213 | 0,032353047 | Casc5 | cancer susceptibility candidate 5 [Source:MGI Symbol;Acc:MGI:1923714] | 0,96 | 6,7 |
| ENSMUSG00000051510 | 7844,165703 | 367,0 | 14556,2 | -5,34102334 | 3,52039E-09 | Mafg | af musculoaponeurotic fibrosarcoma oncogene family, protein G (avian) [Source:MGI Symbol;Acc:MGI:96 | 93,16 | 92,5 |
| ENSMUSG00000036197 | 757,8304356 | 42,0 | 1775,3 | -5,330438599 | 1,21468E-05 | Gxylt1 | glucoside xylosyltransferase 1 [Source:MGI Symbol;Acc:MGI:2684933] | 57,37 | 70,5 |
| ENSMUSG00000057036 | 12,31500647 | 0,9 | 25,0 | -5,326297935 | 0,003685841 | Gm7536 | predicted gene 7536 [Source:MGI Symbol;Acc:MGI:3645137] | #DIV/0! | #DIV/0! |
| ENSMUSG00000084535 | 32,79631168 | 1,5 | 58,5 | -5,326191289 | 2,05301E-07 | Mir1906-1 | microRNA 1906-1 [Source:MGI Symbol;Acc:MGI:3811417] | #DIV/0! | #DIV/0! |
| ENSMUSG00000076612 | 93,59374214 | 3,9 | 191,5 | -5,312309544 | 0,000284004 | Ighg2c | immunoglobulin heavy constant gamma 2C [Source:MGI Symbol;Acc:MGI:2686979] | 32,87 | 56,6 |
| ENSMUSG00000061330 | 3,759028747 | 0,3 | 10,5 | -5,274259448 | 0,032205242 | Gm11361 | predicted pseudogene 11361 [Source:MGI Symbol;Acc:MGI:3649931] | #DIV/0! | #DIV/0! |
| ENSMUSG00000041361 | 73,4133108 | 4,2 | 252,0 | -5,273066611 | 0,005670529 | Myzap | myocardial zonula adherens protein [Source:MGI Symbol;Acc:MGI:2142908] | 19,75 | 47,7 |
| ENSMUSG00000041779 | 108,195717 | 5,0 | 317,5 | -5,264558775 | 0,001055726 | Tram2 | translocating chain-associating membrane protein 2 [Source:MGI Symbol;Acc:MGI:1924817] | 11,87 | 30,0 |
| ENSMUSG00000033644 | 34,81970358 | 0,5 | 26,8 | -5,26158087 | 0,045510333 | Piwil2 | piwi-like RNA-mediated gene silencing 2 [Source:MGI Symbol;Acc:MGI:1930036] | 1,52 | 2,0 |
| ENSMUSG00000049511 | 29,48140725 | 3,2 | 162,8 | -5,259291168 | 0,022275942 | Htr1b | 5-hydroxytryptamine (serotonin) receptor 1B [Source:MGI Symbol;Acc:MGI:96274] | 8,71 | 27,7 |
| ENSMUSG00000079627 | 48,16550414 | 0,7 | 67,0 | -5,24350706 | 0,027465973 | Rhox2h | reproductive homeobox 2H [Source:MGI Symbol;Acc:MGI:3713490] | 9,87 | 38,6 |
| ENSMUSG00000052557 | 493,2670664 | 24,0 | 869,6 | -5,237603324 | 9,71962E-06 | Gan | giant axonal neuropathy [Source:MGI Symbol;Acc:MGI:1890619] | 24,02 | 21,9 |
| ENSMUSG00000078497 | 7,137539188 | 0,5 | 22,8 | -5,235399208 | 0,016951686 | Gm13145 | predicted gene 13145 [Source:MGI Symbol;Acc:MGI:3701123] | 2,90 | 8,9 |
| ENSMUSG00000026193 | 54,62713511 | 5,6 | 202,2 | -5,226138273 | 0,006894788 | Fn1 | fibronectin 1 [Source:MGI Symbol;Acc:MGI:95566] | 7,86 | 11,7 |
| ENSMUSG00000005410 | 83,82610691 | 1,6 | 137,5 | -5,211940054 | 0,012980249 | Mcm5 | romosome maintenance deficient 5, cell division cycle 46 (S. cerevisiae) [Source:MGI Symbol;Acc:MGI:10 | 8,11 | 28,2 |
| ENSMUSG00000020423 | 5881,089427 | 269,7 | 9732,1 | -5,209200662 | 4,66881E-08 | Btg2 | B cell translocation gene 2, anti-proliferative [Source:MGI Symbol;Acc:MGI:108384] | 95,56 | 95,4 |
| ENSMUSG00000093327 | 24,45128137 | 2,2 | 64,0 | -5,196997769 | 6,04359E-05 | Mir5107 | microRNA 5107 [Source:MGI Symbol;Acc:MGI:4950427] | #DIV/0! | #DIV/0! |
| ENSMUSG00000029188 | 3,952070332 | 0,2 | 7,3 | -5,1859067 | 0,026538475 | Slc34a2 | solute carrier family 34 (sodium phosphate), member 2 [Source:MGI Symbol;Acc:MGI:1342284] | 0,00 | 0,0 |
| ENSMUSG00000092805 | 118,2902493 | 5,7 | 209,1 | -5,180330483 | 2,45431E-05 | Gm26461 | predicted gene, 26461 [Source:MGI Symbol;Acc:MGI:5456238] | 28,97 | 44,1 |
| ENSMUSG00000097505 | 78,5870172 | 5,3 | 172,0 | -5,176896341 | 2,15802E-05 | Gm26540 | predicted gene, 26540 [Source:MGI Symbol;Acc:MGI:5477034] | #DIV/0! | #DIV/0! |
| ENSMUSG00000034832 | 803,0776777 | 37,0 | 1283,4 | -5,172645273 | 3,20413E-06 | Tet3 | tet methylcytosine dioxygenase 3 [Source:MGI Symbol;Acc:MGI:2446229] | 34,74 | 40,0 |
| ENSMUSG00000022283 | 8139,156487 | 422,9 | 15751,5 | -5,170753872 | 1,0805E-09 | Pabpc1 | poly(A) binding protein, cytoplasmic 1 [Source:MGI Symbol;Acc:MGI:1349722] | 94,48 | 97,6 |
| ENSMUSG00000044617 | 40,62590232 | 4,6 | 115,6 | -5,163738451 | 0,003324027 | Zbtb39 | zinc finger and BTB domain containing 39 [Source:MGI Symbol;Acc:MGI:2443316] | 11,46 | 19,0 |
| ENSMUSG00000083087 | 30,5384067 | 1,8 | 67,0 | -5,162567978 | 1,58743E-06 | Gm11249 | predicted gene 11249 [Source:MGI Symbol;Acc:MGI:3650834] | #DIV/0! | #DIV/0! |
| ENSMUSG00000026436 | 2946,80653 | 136,7 | 5085,2 | -5,146051251 | 3,91128E-07 | Elk4 | ELK4, member of ETS oncogene family [Source:MGI Symbol;Acc:MGI:102853] | 68,78 | 64,6 |
| ENSMUSG00000023015 | 276,7501578 | 15,9 | 597,9 | -5,115173952 | 0,007404599 | Racgap1 | Rac GTPase-activating protein 1 [Source:MGI Symbol;Acc:MGI:1349423] | 46,47 | 57,4 |
| ENSMUSG00000058126 | 136,70626 | 8,6 | 288,4 | -5,112507846 | 4,28329E-06 | Tpm3-rs7 | tropomyosin 3, related sequence 7 [Source:MGI Symbol;Acc:MGI:99705] | 33,57 | 44,6 |
| ENSMUSG00000064109 | 23,30066315 | 1,1 | 44,3 | -5,108753022 | 0,020740744 | Hcst | hematopoietic cell signal transducer [Source:MGI Symbol;Acc:MGI:1344360] | 28,58 | 53,9 |
| ENSMUSG00000066842 | 4,168205297 | 0,3 | 15,0 | -5,082751501 | 0,039984724 | Hmcn1 | hemicentin 1 [Source:MGI Symbol;Acc:MGI:2685047] | 0,24 | 0,9 |
| ENSMUSG00000026622 | 375,6310548 | 17,3 | 1049,5 | -5,074674143 | 0,000376467 | Nek2 | NIMA (never in mitosis gene a)-related expressed kinase 2 [Source:MGI Symbol;Acc:MGI:109359] | 47,22 | 63,9 |
| ENSMUSG00000074873 | 144,1723334 | 6,7 | 255,1 | -5,074427841 | 7,1321E-05 | AI606181 | expressed sequence AI606181 [Source:MGI Symbol;Acc:MGI:2147586] | 29,57 | 54,1 |
| ENSMUSG00000074887 | 10,35731214 | 0,7 | 16,7 | -5,055208627 | 0,019947179 | EU599041 | expressed sequence EU599041 [Source:MGI Symbol;Acc:MGI:3796554] | #DIV/0! | #DIV/0! |

| | | | | | | | | | |
|---|---|---|---|---|---|---|---|---|---|
| ENSMUSG00000038872 | 60,03632949 | 6,1 | 278,3 | -5,045464779 | 0,000747081 | Zfhx3 | zinc finger homeobox 3 [Source:MGI Symbol;Acc:MGI:99948] | 5,52 | 8,5 |
| ENSMUSG00000062611 | 25,87721819 | 1,4 | 48,3 | -5,045361571 | 1,73585E-05 | Rps3a2 | ribosomal protein S3A2 [Source:MGI Symbol;Acc:MGI:3642853] | #DIV/0! | #DIV/0! |
| ENSMUSG00000029204 | 53,12678572 | 2,8 | 123,6 | -5,02978603 | 0,049415649 | Rhoh | ras homolog gene family, member H [Source:MGI Symbol;Acc:MGI:1921984] | 5,87 | 18,3 |
| ENSMUSG00000076731 | 34,46174002 | 1,0 | 54,5 | -5,024449895 | 0,003193267 | Ighv8-12 | immunoglobulin heavy variable V8-12 [Source:MGI Symbol;Acc:MGI:3642873] | 28,12 | 31,9 |
| ENSMUSG00000028068 | 18,19067824 | 3,5 | 87,9 | -5,021181479 | 0,003504465 | Iqgap3 | IQ motif containing GTPase activating protein 3 [Source:MGI Symbol;Acc:MGI:3028642] | 9,67 | 14,4 |
| ENSMUSG00000076036 | 9598,997426 | 577,3 | 18610,4 | -5,01309555 | 1,28744E-08 | Gm22133 | predicted gene, 22133 [Source:MGI Symbol;Acc:MGI:5451910] | #DIV/0! | #DIV/0! |
| ENSMUSG00000038587 | 14,96613971 | 3,0 | 44,5 | -5,007863324 | 0,005499467 | Akap12 | A kinase (PRKA) anchor protein (gravin) 12 [Source:MGI Symbol;Acc:MGI:1932576] | 8,00 | 7,8 |
| ENSMUSG00000023982 | 21,16738532 | 1,9 | 53,8 | -5,007163205 | 0,009115328 | Guca1a | guanylate cyclase activator 1a (retina) [Source:MGI Symbol;Acc:MGI:102770] | 22,85 | 29,5 |
| ENSMUSG00000081406 | 8,222231566 | 0,8 | 24,4 | -5,002859204 | 0,00229615 | Rps6-ps4 | ribosomal protein S6, pseudogene 4 [Source:MGI Symbol;Acc:MGI:3650907] | #DIV/0! | #DIV/0! |
| ENSMUSG00000082536 | 12,63717516 | 0,7 | 19,9 | -5,001793536 | 0,000141741 | Gm13456 | predicted gene 13456 [Source:MGI Symbol;Acc:MGI:3651389] | #DIV/0! | #DIV/0! |
| ENSMUSG00000027947 | 44,89258547 | 0,7 | 27,4 | -4,997225542 | 0,016318124 | Il6ra | interleukin 6 receptor, alpha [Source:MGI Symbol;Acc:MGI:105304] | 1,55 | 2,9 |
| ENSMUSG00000006360 | 2525,547188 | 149,8 | 4999,5 | -4,979942091 | 1,36322E-09 | Crip1 | cysteine-rich protein 1 (intestinal) [Source:MGI Symbol;Acc:MGI:88501] | 62,10 | 60,2 |
| ENSMUSG00000021268 | 1369,310439 | 92,4 | 2838,6 | -4,974512821 | 5,41933E-10 | Meg3 | maternally expressed 3 [Source:MGI Symbol;Acc:MGI:1202886] | #DIV/0! | #DIV/0! |
| ENSMUSG00000082465 | 7,401982211 | 0,7 | 17,9 | -4,972579396 | 0,00511186 | Rps6-ps3 | ribosomal protein S6, pseudogene 3 [Source:MGI Symbol;Acc:MGI:3643565] | #DIV/0! | #DIV/0! |
| ENSMUSG00000050708 | 25890,21632 | 1603,8 | 50106,7 | -4,972222268 | 9,99202E-09 | Ftl1 | ferritin light polypeptide 1 [Source:MGI Symbol;Acc:MGI:95589] | 90,71 | 87,1 |
| ENSMUSG00000026873 | 83,98606612 | 7,3 | 235,2 | -4,950168787 | 0,000303492 | Phf19 | PHD finger protein 19 [Source:MGI Symbol;Acc:MGI:1921266] | 25,46 | 59,0 |
| ENSMUSG00000092702 | 8,115709604 | 0,5 | 16,9 | -4,940938037 | 0,009369036 | Gm24514 | predicted gene, 24514 [Source:MGI Symbol;Acc:MGI:5454291] | 23,16 | 24,6 |
| ENSMUSG00000093064 | 4,267071813 | 0,3 | 8,1 | -4,932861599 | 0,028021437 | Gm23153 | predicted gene, 23153 [Source:MGI Symbol;Acc:MGI:5452930] | 11,40 | 26,3 |
| ENSMUSG00000062382 | 17,13486472 | 1,2 | 32,6 | -4,92864971 | 9,68725E-05 | Gm10116 | predicted pseudogene 10116 [Source:MGI Symbol;Acc:MGI:3779109] | 6,69 | 15,5 |
| ENSMUSG00000003355 | 146,5051251 | 9,0 | 306,6 | -4,923640503 | 1,16153E-07 | Fkbp11 | FK506 binding protein 11 [Source:MGI Symbol;Acc:MGI:1913370] | 77,98 | 93,7 |
| ENSMUSG00000079455 | 14,13836585 | 1,2 | 29,6 | -4,917440234 | 0,003253813 | Gm16026 | predicted pseudogene 16026 [Source:MGI Symbol;Acc:MGI:3783243] | #DIV/0! | #DIV/0! |
| ENSMUSG00000098112 | 168,9948644 | 17,0 | 350,5 | -4,906680235 | 0,02437757 | Bin2 | bridging integrator 2 [Source:MGI Symbol;Acc:MGI:3611448] | 58,01 | 60,6 |
| ENSMUSG00000065037 | 3,939076534 | 0,4 | 16,8 | -4,905913905 | 0,01657662 | Rn7sk | RNA, 7SK, nuclear [Source:MGI Symbol;Acc:MGI:103186] | 19,54 | 30,0 |
| ENSMUSG00000033327 | 17,43443412 | 1,3 | 99,4 | -4,904904773 | 0,026852488 | Tnxb | tenascin XB [Source:MGI Symbol;Acc:MGI:1932137] | 2,15 | 6,4 |
| ENSMUSG00000037991 | 43,14646572 | 2,9 | 83,3 | -4,902984895 | 0,000753061 | Rmi2 | MI2, RecQ mediated genome instability 2, homolog (S. cerevisiae) [Source:MGI Symbol;Acc:MGI:268538 | 10,88 | 18,8 |
| ENSMUSG00000057098 | 20,68036841 | 1,4 | 39,2 | -4,876523467 | 0,007221778 | Ebf1 | early B cell factor 1 [Source:MGI Symbol;Acc:MGI:95275] | 2,90 | 7,7 |
| ENSMUSG00000097554 | 481,5209941 | 29,2 | 910,6 | -4,875154704 | 3,72827E-07 | Gm26825 | predicted gene, 26825 [Source:MGI Symbol;Acc:MGI:5477319] | #DIV/0! | #DIV/0! |
| ENSMUSG00000024610 | 272,8664983 | 21,6 | 627,7 | -4,859462332 | 2,32518E-09 | Cd74 | ıvariant polypeptide of major histocompatibility complex, class II antigen-associated) [Source:MGI Symbol; | 79,34 | 85,5 |
| ENSMUSG00000091383 | 8,619674137 | 0,7 | 18,9 | -4,845162565 | 0,003764373 | Hist1h2al | histone cluster 1, H2al [Source:MGI Symbol;Acc:MGI:3646032] | 23,46 | 26,9 |
| ENSMUSG00000025407 | 244,2628535 | 12,6 | 389,8 | -4,840081239 | 3,3386E-06 | Gli1 | GLI-Kruppel family member GLI1 [Source:MGI Symbol;Acc:MGI:95727] | 41,80 | 61,2 |
| ENSMUSG00000096942 | 42,59128759 | 3,9 | 95,9 | -4,840012161 | 5,40052E-06 | Rps19-ps6 | ribosomal protein S19, pseudogene 6 [Source:MGI Symbol;Acc:MGI:5011818] | #DIV/0! | #DIV/0! |
| ENSMUSG00000053559 | 177,6213597 | 7,8 | 265,9 | -4,834515485 | 1,98013E-08 | Smagp | small cell adhesion glycoprotein [Source:MGI Symbol;Acc:MGI:2448476] | 58,41 | 78,5 |
| ENSMUSG00000059422 | 31,37201464 | 1,2 | 48,9 | -4,834149297 | 0,006792606 | Gm8116 | predicted gene 8116 [Source:MGI Symbol;Acc:MGI:3648797] | #DIV/0! | #DIV/0! |
| ENSMUSG00000090246 | 53,38780729 | 3,5 | 116,2 | -4,828728462 | 0,001811264 | Gm17017 | predicted gene 17017 [Source:MGI Symbol;Acc:MGI:4820558] | 6,00 | 11,4 |
| ENSMUSG00000024937 | 329,3262644 | 27,7 | 768,2 | -4,816542234 | 2,30444E-07 | Ehbp1l1 | EH domain binding protein 1-like 1 [Source:MGI Symbol;Acc:MGI:3612340] | 38,58 | 45,1 |
| ENSMUSG00000070034 | 56,65684299 | 1,7 | 57,6 | -4,816255791 | 9,46583E-06 | Sp110 | Sp110 nuclear body protein [Source:MGI Symbol;Acc:MGI:1923364] | 12,55 | 15,0 |
| ENSMUSG00000105790 | 11,06159584 | 0,7 | 17,2 | -4,812926278 | 0,0032678 | AC122546.1 | | #DIV/0! | #DIV/0! |
| ENSMUSG00000054116 | 148,4771359 | 8,0 | 222,6 | -4,806781937 | 1,22483E-07 | E130116L18Rik | RIKEN cDNA E130116L18 gene [Source:MGI Symbol;Acc:MGI:1925120] | 87,15 | 84,4 |
| ENSMUSG00000097736 | 25,50417217 | 6,4 | 127,4 | -4,801118114 | 0,027404618 | 9530059O14Rik | RIKEN cDNA 9530059O14 gene [Source:MGI Symbol;Acc:MGI:2442421] | #DIV/0! | #DIV/0! |
| ENSMUSG00000020592 | 151,9597165 | 7,8 | 203,6 | -4,794106525 | 0,001615582 | Sdc1 | syndecan 1 [Source:MGI Symbol;Acc:MGI:1349162] | 35,54 | 48,4 |
| ENSMUSG00000045694 | 10,91272542 | 0,7 | 21,6 | -4,794096324 | 0,00415514 | Gm4997 | predicted gene 4997 [Source:MGI Symbol;Acc:MGI:3643651] | #DIV/0! | #DIV/0! |
| ENSMUSG00000030104 | 2125,006824 | 145,7 | 4430,7 | -4,785184421 | 1,0847E-09 | Edem1 | ER degradation enhancer, mannosidase alpha-like 1 [Source:MGI Symbol;Acc:MGI:2180139] | 96,45 | 95,8 |
| ENSMUSG00000099597 | 697,943171 | 52,9 | 1463,6 | -4,774156142 | 3,26295E-09 | Gm28933 | predicted gene 28933 [Source:MGI Symbol;Acc:MGI:5579639] | #DIV/0! | #DIV/0! |
| ENSMUSG00000084020 | 25,84664892 | 1,9 | 59,9 | -4,765930546 | 1,00287E-05 | Wsb2-ps | WD repeat and SOCS box-containing 2, pseudogene [Source:MGI Symbol;Acc:MGI:3649891] | #DIV/0! | #DIV/0! |
| ENSMUSG00000019942 | 39,577135 | 8,5 | 155,5 | -4,760758434 | 0,020593229 | Cdk1 | cyclin-dependent kinase 1 [Source:MGI Symbol;Acc:MGI:88351] | 20,49 | 30,2 |
| ENSMUSG00000057836 | 46,97716758 | 3,4 | 75,3 | -4,758959286 | 0,001239038 | Xlr3a | X-linked lymphocyte-regulated 3A [Source:MGI Symbol;Acc:MGI:109506] | 26,43 | 32,1 |
| ENSMUSG00000083483 | 4,775678862 | 0,3 | 9,1 | -4,753449493 | 0,040204319 | Gm14044 | predicted gene 14044 [Source:MGI Symbol;Acc:MGI:3650452] | #DIV/0! | #DIV/0! |
| ENSMUSG00000042644 | 24,53671083 | 5,1 | 78,8 | -4,752797929 | 0,009535772 | Itpr3 | inositol 1,4,5-triphosphate receptor 3 [Source:MGI Symbol;Acc:MGI:96624] | 10,07 | 5,7 |
| ENSMUSG00000030785 | 138,8463258 | 7,6 | 216,4 | -4,750874417 | 2,75023E-05 | Cox6a2 | cytochrome c oxidase subunit VIa polypeptide 2 [Source:MGI Symbol;Acc:MGI:104649] | 69,26 | 71,2 |
| ENSMUSG00000005667 | 927,088201 | 72,1 | 1920,4 | -4,733736365 | 1,38937E-07 | Mthfd2 | ofolate dehydrogenase (NAD+ dependent), methenyltetrahydrofolate cyclohydrolase [Source:MGI Symbol; | 96,26 | 89,0 |
| ENSMUSG00000064945 | 6,333533816 | 0,7 | 18,2 | -4,725482773 | 0,004061773 | Rny3 | RNA, Y3 small cytoplasmic (associated with Ro protein) [Source:MGI Symbol;Acc:MGI:107304] | 68,95 | 87,3 |
| ENSMUSG00000021514 | 389,37353 | 27,4 | 726,3 | -4,724685918 | 3,00981E-05 | Zfp369 | zinc finger protein 369 [Source:MGI Symbol;Acc:MGI:2176229] | 52,74 | 12,4 |
| ENSMUSG00000082935 | 19,56744953 | 1,5 | 38,4 | -4,723312279 | 0,002222804 | Gm7658 | predicted gene 7658 [Source:MGI Symbol;Acc:MGI:3648029] | #DIV/0! | #DIV/0! |
| ENSMUSG00000105987 | 157,8728486 | 12,5 | 334,7 | -4,714039861 | 6,28475E-14 | AI506816 | expressed sequence AI506816 [Source:MGI Symbol;Acc:MGI:2140929] | #DIV/0! | #DIV/0! |
| ENSMUSG00000094344 | 10,48350132 | 1,2 | 27,4 | -4,70550334 | 0,000652266 | Gm11942 | predicted gene 11942 [Source:MGI Symbol;Acc:MGI:3650608] | #DIV/0! | #DIV/0! |
| ENSMUSG00000042225 | 345,4726374 | 22,4 | 644,7 | -4,690672035 | 9,83737E-05 | Ammecr1 | ıental retardation, midface hypoplasia and elliptocytosis chromosomal region gene 1 [Source:MGI Symbol; | 49,11 | 33,4 |
| ENSMUSG00000029177 | 487,6594281 | 28,4 | 898,8 | -4,687941773 | 1,28384E-05 | Cenpa | centromere protein A [Source:MGI Symbol;Acc:MGI:88375] | 78,31 | 63,3 |
| ENSMUSG00000094088 | 59,39384965 | 2,9 | 90,4 | -4,677673179 | 0,001689804 | Ighv1-64 | immunoglobulin heavy variable 1-64 [Source:MGI Symbol;Acc:MGI:4439789] | 50,70 | 60,7 |
| ENSMUSG00000084349 | 203,9446293 | 14,1 | 378,4 | -4,676456751 | 8,84247E-08 | Rpl3-ps1 | ribosomal protein L3, pseudogene 1 [Source:MGI Symbol;Acc:MGI:3644217] | #DIV/0! | #DIV/0! |
| ENSMUSG00000020184 | 10069,65634 | 700,9 | 19108,8 | -4,670599456 | 9,25997E-08 | Mdm2 | transformed mouse 3T3 cell double minute 2 [Source:MGI Symbol;Acc:MGI:96952] | 97,72 | 97,5 |
| ENSMUSG00000109509 | 11,66462918 | 0,9 | 20,3 | -4,668607214 | 0,006618112 | RP24-471H15.4 | | #DIV/0! | #DIV/0! |
| ENSMUSG00000025480 | 168,6660051 | 11,9 | 414,8 | -4,63799009 | 0,006385371 | Syce1 | synaptonemal complex central element protein 1 [Source:MGI Symbol;Acc:MGI:1921325] | 53,60 | 65,7 |
| ENSMUSG00000096188 | 955,8155426 | 72,4 | 1786,7 | -4,637033398 | 6,66747E-09 | Cmtm4 | CKLF-like MARVEL transmembrane domain containing 4 [Source:MGI Symbol;Acc:MGI:2142888] | 48,30 | 51,8 |
| ENSMUSG00000080893 | 71,78321434 | 6,1 | 150,3 | -4,588814774 | 3,76204E-06 | Gm15920 | predicted gene 15920 [Source:MGI Symbol;Acc:MGI:3801934] | #DIV/0! | #DIV/0! |
| ENSMUSG00000047676 | 12,37291759 | 1,1 | 27,9 | -4,586543396 | 0,000196744 | Rpsa-ps10 | ribosomal protein SA, pseudogene 10 [Source:MGI Symbol;Acc:MGI:3704228] | #DIV/0! | #DIV/0! |
| ENSMUSG00000026832 | 237,1753175 | 9,9 | 391,6 | -4,56053694 | 0,034693621 | Cytip | cytohesin 1 interacting protein [Source:MGI Symbol;Acc:MGI:2183535] | 17,94 | 12,8 |
| ENSMUSG00000058443 | 72,24922445 | 5,2 | 132,5 | -4,553436876 | 2,21133E-05 | Rpl10-ps3 | ribosomal protein L10, pseudogene 3 [Source:MGI Symbol;Acc:MGI:3704336] | 11,01 | 16,1 |
| ENSMUSG00000098661 | 7,4933425 | 1,1 | 26,0 | -4,548562753 | 0,017048315 | Mir7052 | microRNA 7052 [Source:MGI Symbol;Acc:MGI:5530965] | #DIV/0! | #DIV/0! |
| ENSMUSG00000105461 | 7,498083575 | 0,6 | 16,4 | -4,545126155 | 0,019798401 | Gm4742 | predicted gene 4742 [Source:MGI Symbol;Acc:MGI:3645661] | #DIV/0! | #DIV/0! |
| ENSMUSG00000106895 | 10,49234338 | 0,8 | 18,6 | -4,534800013 | 0,00331406 | Gm4754 | predicted gene 4754 [Source:MGI Symbol;Acc:MGI:3645925] | #DIV/0! | #DIV/0! |
| ENSMUSG00000040860 | 130,3699322 | 27,2 | 599,5 | -4,529584108 | 0,000822862 | Crocc | ciliary rootlet coiled-coil, rootletin [Source:MGI Symbol;Acc:MGI:3529431] | 47,28 | 72,7 |
| ENSMUSG00000024399 | 30,39305133 | 2,9 | 85,5 | -4,512544884 | 0,0080152 | Ltb | lymphotoxin B [Source:MGI Symbol;Acc:MGI:104796] | 29,65 | 51,9 |
| ENSMUSG00000097994 | 90,25303762 | 4,8 | 148,2 | -4,506797012 | 0,018209447 | Gm26982 | predicted gene, 26982 [Source:MGI Symbol;Acc:MGI:5504097] | 43,47 | 48,9 |
| ENSMUSG00000070305 | 314,179915 | 24,9 | 568,4 | -4,506129174 | 0,002016411 | Mpzl3 | myelin protein zero-like 3 [Source:MGI Symbol;Acc:MGI:2442647] | 68,65 | 69,0 |
| ENSMUSG00000044533 | 7786,727512 | 678,8 | 15416,8 | -4,492492705 | 3,47815E-08 | Rps2 | ribosomal protein S2 [Source:MGI Symbol;Acc:MGI:105110] | 68,76 | 58,4 |
| ENSMUSG00000034333 | 28,67066197 | 2,1 | 70,0 | -4,489047024 | 0,005334587 | Zbed4 | zinc finger, BED type containing 4 [Source:MGI Symbol;Acc:MGI:2682302] | 5,82 | 5,6 |
| ENSMUSG00000091845 | 5,181156846 | 0,6 | 8,2 | -4,488212713 | 0,02468674 | Gm4604 | predicted gene 4604 [Source:MGI Symbol;Acc:MGI:3782787] | #DIV/0! | #DIV/0! |
| ENSMUSG00000011960 | 2304,394541 | 188,4 | 4239,8 | -4,487709234 | 7,75904E-08 | Ccnt1 | cyclin T1 [Source:MGI Symbol;Acc:MGI:1328363] | 68,03 | 82,0 |
| ENSMUSG00000020183 | 407,3225969 | 34,1 | 715,6 | -4,480081415 | 5,555E-08 | Cpm | carboxypeptidase M [Source:MGI Symbol;Acc:MGI:1917824] | 48,60 | 27,7 |
| ENSMUSG00000025795 | 463,7039266 | 45,3 | 1021,2 | -4,473333379 | 5,555E-08 | Rassf3 | Ras association (RalGDS/AF-6) domain family member 3 [Source:MGI Symbol;Acc:MGI:2179722] | 85,76 | 97,1 |
| ENSMUSG00000020680 | 30,40316582 | 2,6 | 55,8 | -4,472152096 | 0,003823378 | Taf15 | ; RNA polymerase II, TATA box binding protein (TBP)-associated factor [Source:MGI Symbol;Acc:MGI:191 | 19,44 | 31,3 |
| ENSMUSG00000024769 | 76,82222904 | 6,8 | 124,7 | -4,465422943 | 0,001838255 | Cdc42bpg | CDC42 binding protein kinase gamma (DMPK-like) [Source:MGI Symbol;Acc:MGI:2652845] | 17,79 | 21,3 |
| ENSMUSG00000064193 | 4,374698552 | 0,8 | 12,2 | -4,456621722 | 0,019433244 | Gm4735 | predicted gene 4735 [Source:MGI Symbol;Acc:MGI:3645521] | #DIV/0! | #DIV/0! |
| ENSMUSG00000084416 | 66,21855928 | 5,6 | 125,1 | -4,455502996 | 7,08271E-06 | Rpl10a-ps1 | ribosomal protein L10A, pseudogene 1 [Source:MGI Symbol;Acc:MGI:3705885] | #DIV/0! | #DIV/0! |
| ENSMUSG00000040084 | 92,91837585 | 11,2 | 327,1 | -4,433065351 | 0,012135925 | Bub1b | BUB1B, mitotic checkpoint serine/threonine kinase [Source:MGI Symbol;Acc:MGI:1333889] | 44,59 | 41,0 |
| ENSMUSG00000024472 | 335,2266456 | 29,5 | 690,8 | -4,423770647 | 3,1628E-09 | Dcp2 | decapping mRNA 2 [Source:MGI Symbol;Acc:MGI:1917890] | 27,38 | 21,5 |
| ENSMUSG00000051747 | 44,87235462 | 4,2 | 80,0 | -4,410894958 | 0,001308733 | Ttn | titin [Source:MGI Symbol;Acc:MGI:98864] | 0,50 | 1,2 |
| ENSMUSG00000036391 | 271,3747666 | 35,1 | 727,0 | -4,406522514 | 5,84859E-05 | Sec24a | Sec24 related gene family, member A (S. cerevisiae) [Source:MGI Symbol;Acc:MGI:1924621] | 47,58 | 32,7 |
| ENSMUSG00000041258 | 577,5177457 | 59,6 | 1141,7 | -4,401796357 | 1,17878E-06 | Zfp236 | zinc finger protein 236 [Source:MGI Symbol;Acc:MGI:1926950] | 41,85 | 23,4 |
| ENSMUSG00000020372 | 6053,641255 | 583,7 | 12434,4 | -4,398034559 | 2,96566E-09 | Gnb2l1 | anine nucleotide binding protein (G protein), beta polypeptide 2 like 1 [Source:MGI Symbol;Acc:MGI:1018 | 87,70 | 82,7 |
| ENSMUSG00000020715 | 106,3550586 | 18,5 | 274,6 | -4,397347554 | 0,003159434 | Ern1 | endoplasmic reticulum (ER) to nucleus signalling 1 [Source:MGI Symbol;Acc:MGI:1930134] | 29,90 | 17,2 |
| ENSMUSG00000040195 | 266,5486904 | 37,1 | 843,5 | -4,395261308 | 0,002832565 | Tmem194 | transmembrane protein 194 [Source:MGI Symbol;Acc:MGI:2446113] | 29,92 | 51,2 |
| ENSMUSG00000069862 | 37,07597893 | 3,3 | 80,6 | -4,392250227 | 0,000894573 | Rps12-ps9 | ribosomal protein S12, pseudogene 9 [Source:MGI Symbol;Acc:MGI:3704324] | #DIV/0! | #DIV/0! |
| ENSMUSG00000097545 | 48,27013572 | 2,6 | 138,1 | -4,386402883 | 0,023083401 | Mir124a-1hg | Mir124-1 host gene (non-protein coding) [Source:MGI Symbol;Acc:MGI:2442197] | #DIV/0! | #DIV/0! |
| ENSMUSG00000097327 | 21,74783253 | 1,8 | 35,9 | -4,369914806 | 0,031222156 | E030030I06Rik | RIKEN cDNA E030030I06 gene [Source:MGI Symbol;Acc:MGI:2442914] | 7,95 | 9,2 |
| ENSMUSG00000059751 | 4,966493075 | 0,7 | 8,3 | -4,365533303 | 0,033025772 | Rps3a3 | ribosomal protein S3A3 [Source:MGI Symbol;Acc:MGI:3643406] | #DIV/0! | #DIV/0! |
| ENSMUSG00000038437 | 266,659877 | 33,9 | 694,7 | -4,364499481 | 0,000295215 | Mllt6 | oid or mixed-lineage leukemia (trithorax homolog, Drosophila); translocated to, 6 [Source:MGI Symbol;Acc: | 42,57 | 42,2 |
| ENSMUSG00000096403 | 35,03209266 | 4,0 | 90,2 | -4,351943607 | 0,000690653 | Gm9825 | predicted gene 9825 [Source:MGI Symbol;Acc:MGI:3708729] | #DIV/0! | #DIV/0! |
| ENSMUSG00000025330 | 19,28063523 | 2,0 | 83,5 | -4,349912793 | 0,022461671 | Padi4 | peptidyl arginine deiminase, type IV [Source:MGI Symbol;Acc:MGI:1338898] | 14,35 | 24,0 |
| ENSMUSG00000030091 | 53,32401952 | 10,2 | 139,6 | -4,346614887 | 0,018853713 | Nup210 | nucleoporin 210 [Source:MGI Symbol;Acc:MGI:1859555] | 19,22 | 15,0 |
| ENSMUSG00000027514 | 78,93826311 | 4,2 | 99,1 | -4,321275165 | 4,6433E-05 | Zbp1 | Z-DNA binding protein 1 [Source:MGI Symbol;Acc:MGI:1927449] | 22,84 | 23,9 |
| ENSMUSG00000036698 | 2153,187514 | 180,0 | 3504,3 | -4,317333362 | 2,26612E-06 | Ago2 | argonaute RISC catalytic subunit 2 [Source:MGI Symbol;Acc:MGI:2446632] | 69,96 | 59,8 |
| ENSMUSG00000024378 | 599,7785597 | 59,9 | 1335,5 | -4,314897475 | 0,000361488 | Stard4 | StAR-related lipid transfer (START) domain containing 4 [Source:MGI Symbol;Acc:MGI:2156764] | 77,98 | 79,7 |
| ENSMUSG00000099492 | 30,37989066 | 4,1 | 78,7 | -4,311355755 | 7,65615E-05 | Gm5525 | predicted gene 5525 [Source:MGI Symbol;Acc:MGI:3646042] | #DIV/0! | #DIV/0! |
| ENSMUSG00000080242 | 22,12533393 | 0,9 | 20,4 | -4,311147444 | 0,00054709 | Gm15487 | predicted gene 15487 [Source:MGI Symbol;Acc:MGI:3709610] | #DIV/0! | #DIV/0! |
| ENSMUSG00000028649 | 513,0763115 | 55,7 | 1151,3 | -4,298070241 | 0,000213007 | Macf1 | microtubule-actin crosslinking factor 1 [Source:MGI Symbol;Acc:MGI:108559] | 22,90 | 27,6 |
| ENSMUSG00000008682 | 4532,812726 | 447,3 | 8938,8 | -4,287965668 | 7,01353E-09 | Rpl10 | ribosomal protein L10 [Source:MGI Symbol;Acc:MGI:105943] | 81,73 | 79,8 |
| ENSMUSG00000073409 | 9,082459149 | 1,8 | 17,4 | -4,277208321 | 0,035618532 | H2-Q6 | histocompatibility 2, Q region locus 6 [Source:MGI Symbol;Acc:MGI:95935] | 11,05 | 9,5 |
| ENSMUSG00000055184 | 74,94025788 | 11,7 | 231,3 | -4,263336736 | 0,007768705 | Fam72a | family with sequence similarity 72, member A [Source:MGI Symbol;Acc:MGI:1919669] | 32,60 | 49,7 |
| ENSMUSG00000087412 | 32,84010461 | 3,6 | 74,2 | -4,25311081 | 5,74761E-05 | Gm15501 | predicted pseudogene 15501 [Source:MGI Symbol;Acc:MGI:3704296] | #DIV/0! | #DIV/0! |
| ENSMUSG00000102289 | 27,09544473 | 2,8 | 50,4 | -4,251857976 | 0,000544672 | Gm31258 | predicted gene, 31258 [Source:MGI Symbol;Acc:MGI:5590417] | 4,55 | 3,4 |
| ENSMUSG00000055093 | 43,07000989 | 4,1 | 93,5 | -4,248407703 | 0,000358526 | Gm8430 | predicted pseudogene 8430 [Source:MGI Symbol;Acc:MGI:3645406] | #DIV/0! | #DIV/0! |
| ENSMUSG00000028211 | 6452,519215 | 534,2 | 10260,5 | -4,248306982 | 0,000303492 | Trp53inp1 | transformation related protein 53 inducible nuclear protein 1 [Source:MGI Symbol;Acc:MGI:1926609] | 99,65 | 98,6 |
| ENSMUSG00000094685 | 36,21143755 | 3,2 | 58,5 | -4,23251306 | 0,008077579 | Gm5900 | predicted pseudogene 5900 [Source:MGI Symbol;Acc:MGI:3643907] | 27,67 | 30,7 |
| ENSMUSG00000094862 | 9,179062856 | 0,7 | 13,3 | -4,226494248 | 0,008081216 | Ighv1-56 | immunoglobulin heavy variable 1-56 [Source:MGI Symbol;Acc:MGI:4439715] | 21,27 | 24,5 |
| ENSMUSG00000032035 | 293,2521258 | 18,9 | 579,3 | -4,223356828 | 0,005908159 | Ets1 | E26 avian leukemia oncogene 1, 5' domain [Source:MGI Symbol;Acc:MGI:95455] | 42,25 | 51,6 |
| ENSMUSG00000041313 | 129,4135367 | 16,0 | 295,9 | -4,197762888 | 7,97277E-08 | Slc7a1 | e carrier family 7 (cationic amino acid transporter, y+ system), member 1 [Source:MGI Symbol;Acc:MGI:88 | 25,65 | 30,1 |
| ENSMUSG00000031732 | 78,69659104 | 8,7 | 210,8 | -4,196937381 | 0,027465973 | Phlpp2 | PH domain and leucine rich repeat protein phosphatase 2 [Source:MGI Symbol;Acc:MGI:2444928] | 14,84 | 15,5 |
| ENSMUSG00000097767 | 151,8900347 | 20,4 | 437,7 | -4,196294191 | 0,030503172 | Miat | myocardial infarction associated transcript (non-protein coding) [Source:MGI Symbol;Acc:MGI:2444886] | #DIV/0! | #DIV/0! |
| ENSMUSG00000043091 | 259,8076472 | 32,1 | 574,7 | -4,14761083 | 4,02324E-09 | Tuba1c | tubulin, alpha 1C [Source:MGI Symbol;Acc:MGI:1095409] | 37,58 | 33,4 |
| ENSMUSG00000032264 | 385,9192577 | 67,0 | 1176,8 | -4,146784539 | 6,90135E-11 | Zw10 | zw10 kinetochore protein [Source:MGI Symbol;Acc:MGI:1349478] | 92,09 | 87,2 |
| ENSMUSG00000027381 | 469,6618265 | 24,2 | 663,6 | -4,143286474 | 0,000372966 | Bcl2l11 | BCL2-like 11 (apoptosis facilitator) [Source:MGI Symbol;Acc:MGI:1197519] | 38,86 | 33,1 |
| ENSMUSG00000019564 | 322,4820306 | 10,6 | 338,0 | -4,13851078 | 0,00236833 | Arid3a | AT rich interactive domain 3A (BRIGHT-like) [Source:MGI Symbol;Acc:MGI:1328360] | 26,87 | 41,2 |
| ENSMUSG00000107092 | 13,37237055 | 0,8 | 13,8 | -4,137982914 | 0,003510998 | Gm7993 | predicted gene 7993 [Source:MGI Symbol;Acc:MGI:3647399] | #DIV/0! | #DIV/0! |
| ENSMUSG00000028044 | 669,223202 | 72,6 | 1329,4 | -4,120573845 | 1,49817E-07 | Cks1b | CDC28 protein kinase 1b [Source:MGI Symbol;Acc:MGI:1889208] | 71,91 | 72,4 |
| ENSMUSG00000105471 | 48,18701596 | 5,0 | 85,3 | -4,118648348 | 6,21322E-06 | A430073D23Rik | RIKEN cDNA A430073D23 gene [Source:MGI Symbol;Acc:MGI:4361019] | #DIV/0! | #DIV/0! |
| ENSMUSG00000031637 | 50,71539744 | 7,3 | 119,4 | -4,118021818 | 0,001516172 | Lrp2bp | Lrp2 binding protein [Source:MGI Symbol;Acc:MGI:1914870] | 19,61 | 10,7 |
| ENSMUSG00000105031 | 315,9133488 | 35,6 | 614,4 | -4,111023739 | 7,17312E-07 | Gm3511 | predicted gene 3511 [Source:MGI Symbol;Acc:MGI:3781688] | #DIV/0! | #DIV/0! |
| ENSMUSG00000020788 | 184,733299 | 13,7 | 317,2 | -4,101128134 | 0,017742249 | Atp2a3 | ATPase, Ca++ transporting, ubiquitous [Source:MGI Symbol;Acc:MGI:1194503] | 39,96 | 37,9 |
| ENSMUSG00000060019 | 91,3995725 | 8,1 | 144,3 | -4,100343518 | 0,000693649 | Gm10073 | predicted pseudogene 10073 [Source:MGI Symbol;Acc:MGI:3641908] | 28,76 | 31,8 |
| ENSMUSG00000070343 | 43,10326785 | 5,5 | 97,4 | -4,10023716 | 0,000146715 | Gm10288 | predicted gene 10288 [Source:MGI Symbol;Acc:MGI:3704227] | #DIV/0! | #DIV/0! |
| ENSMUSG00000040952 | 7701,172479 | 837,7 | 14529,2 | -4,098015794 | 5,65874E-07 | Rps19 | ribosomal protein S19 [Source:MGI Symbol;Acc:MGI:1333780] | 44,43 | 41,6 |
| ENSMUSG00000068396 | 4,173624578 | 0,6 | 9,6 | -4,089297945 | 0,012522787 | Rpl34-ps1 | ribosomal protein L34, pseudogene 1 [Source:MGI Symbol;Acc:MGI:3704270] | #DIV/0! | #DIV/0! |
| ENSMUSG00000030189 | 3465,697206 | 364,6 | 6332,0 | -4,08006836 | 3,81817E-08 | Ybx3 | Y box protein 3 [Source:MGI Symbol;Acc:MGI:2137670] | 98,33 | 99,5 |
| ENSMUSG00000091408 | 26,45183063 | 1,4 | 26,6 | -4,075688584 | 0,011104678 | Gm6728 | predicted gene 6728 [Source:MGI Symbol;Acc:MGI:3643418] | #DIV/0! | #DIV/0! |
| ENSMUSG00000000486 | 120,3371151 | 12,0 | 211,0 | -4,070742028 | 0,000339105 | Sept1 | septin 1 [Source:MGI Symbol;Acc:MGI:1858916] | 42,75 | 73,4 |
| ENSMUSG00000071644 | 4158,937501 | 468,0 | 7855,1 | -4,064461657 | 1,93102E-07 | Eef1g | eukaryotic translation elongation factor 1 gamma [Source:MGI Symbol;Acc:MGI:1914410] | 75,11 | 64,7 |
| ENSMUSG00000027490 | 512,9507869 | 48,1 | 816,1 | -4,062898767 | 2,32992E-06 | E2f1 | E2F transcription factor 1 [Source:MGI Symbol;Acc:MGI:101941] | 74,15 | 73,1 |
| ENSMUSG00000029752 | 1096,2716 | 121,9 | 2202,8 | -4,06141559 | 3,76013E-05 | Asns | asparagine synthetase [Source:MGI Symbol;Acc:MGI:1350929] | 80,40 | 77,7 |
| ENSMUSG00000027469 | 146,5127368 | 21,1 | 395,2 | -4,049418883 | 0,003833396 | Tpx2 | TPX2, microtubule-associated protein homolog (Xenopus laevis) [Source:MGI Symbol;Acc:MGI:1919369] | 46,78 | 42,0 |
| ENSMUSG00000036908 | 326,8737756 | 43,7 | 801,5 | -4,03503258 | 9,4607E-06 | Unc93b1 | unc-93 homolog B1 (C. elegans) [Source:MGI Symbol;Acc:MGI:1859307] | 87,36 | 91,8 |
| ENSMUSG00000108366 | 8,549054331 | 0,4 | 9,1 | -4,031723376 | 0,015142907 | RP24-490A22.9 | | #DIV/0! | #DIV/0! |
| ENSMUSG00000041133 | 1510,404606 | 210,4 | 3303,9 | -4,01990808 | 2,21133E-05 | Smc1a | structural maintenance of chromosomes 1A [Source:MGI Symbol;Acc:MGI:1344345] | 85,00 | 82,8 |
| ENSMUSG00000076138 | 3734,809382 | 424,6 | 7232,3 | -4,01978897 | 7,86835E-06 | Gm26377 | predicted gene, 26377 [Source:MGI Symbol;Acc:MGI:5456154] | #DIV/0! | #DIV/0! |
| ENSMUSG00000032518 | 3651,633269 | 412,1 | 6751,8 | -4,003594763 | 5,555E-08 | Rpsa | ribosomal protein SA [Source:MGI Symbol;Acc:MGI:105381] | 65,52 | 58,1 |
| ENSMUSG00000022311 | 19,35408013 | 5,5 | 37,7 | -4,002682907 | 0,021890207 | Csmd3 | CUB and Sushi multiple domains 3 [Source:MGI Symbol;Acc:MGI:2386403] | 5,29 | 3,8 |
| ENSMUSG00000028333 | 3577,280412 | 456,4 | 7480,6 | -3,999312201 | 1,25858E-08 | Anp32b | acidic (leucine-rich) nuclear phosphoprotein 32 family, member B [Source:MGI Symbol;Acc:MGI:1914878] | 85,86 | 86,0 |
| ENSMUSG00000007892 | 9021,236542 | 992,1 | 16082,4 | -3,996641756 | 1,10834E-06 | Rplp1 | ribosomal protein, large, P1 [Source:MGI Symbol;Acc:MGI:1927099] | 100,00 | 100,0 |
| ENSMUSG00000058655 | 3499,878575 | 443,0 | 7109,0 | -3,995784791 | 5,94025E-09 | Eif4b | eukaryotic translation initiation factor 4B [Source:MGI Symbol;Acc:MGI:95304] | 97,71 | 94,5 |
| ENSMUSG00000039985 | 53,65734564 | 5,1 | 88,0 | -3,994156589 | 0,001827916 | Fam60a | family with sequence similarity 60, member A [Source:MGI Symbol;Acc:MGI:1929091] | 11,66 | 10,1 |
| ENSMUSG00000081684 | 4,622433735 | 0,6 | 9,0 | -3,989245985 | 0,033174792 | Rps2-ps13 | ribosomal protein S2, pseudogene 13 [Source:MGI Symbol;Acc:MGI:3705640] | #DIV/0! | #DIV/0! |
| ENSMUSG00000075470 | 189,7378882 | 20,2 | 337,9 | -3,989114482 | 5,74761E-05 | Alg10b | sparagine-linked glycosylation 10B (alpha-1,2-glucosyltransferase) [Source:MGI Symbol;Acc:MGI:214615 | 55,85 | 51,3 |
| ENSMUSG00000037466 | 209,971086 | 11,6 | 422,0 | -3,98716658 | 0,002133857 | 4930427A07Rik | RIKEN cDNA 4930427A07 gene [Source:MGI Symbol;Acc:MGI:2144738] | 50,21 | 59,4 |
| ENSMUSG00000050914 | 146,2090012 | 19,1 | 343,5 | -3,984991858 | 2,22705E-05 | Ankrd37 | ankyrin repeat domain 37 [Source:MGI Symbol;Acc:MGI:3603344] | 82,27 | 93,2 |
| ENSMUSG00000060657 | 69,30377246 | 8,5 | 233,1 | -3,98458743 | 0,01949513 | Marf1 | meiosis arrest female 1 [Source:MGI Symbol;Acc:MGI:2444505] | 12,86 | 26,3 |
| ENSMUSG00000026234 | 7227,918776 | 863,8 | 14210,7 | -3,98442721 | 1,76545E-07 | Ncl | nucleolin [Source:MGI Symbol;Acc:MGI:97286] | 76,48 | 65,5 |
| ENSMUSG00000043279 | 349,5535636 | 36,0 | 582,0 | -3,980658766 | 4,50336E-06 | Trim56 | tripartite motif-containing 56 [Source:MGI Symbol;Acc:MGI:2685298] | 40,25 | 35,7 |
| ENSMUSG00000058006 | 32,94798709 | 4,5 | 110,1 | -3,965066912 | 0,019744889 | Mdn1 | midasin homolog (yeast) [Source:MGI Symbol;Acc:MGI:1926159] | 2,78 | 4,4 |
| ENSMUSG00000026558 | 867,8330169 | 81,8 | 1362,3 | -3,963490017 | 2,11226E-06 | Uck2 | uridine-cytidine kinase 2 [Source:MGI Symbol;Acc:MGI:1931744] | 82,78 | 76,6 |
| ENSMUSG00000044847 | 241,8096786 | 25,4 | 403,7 | -3,9621268 | 2,35035E-06 | Lsm11 | U7 snRNP-specific Sm-like protein LSM11 [Source:MGI Symbol;Acc:MGI:1919540] | 45,42 | 37,7 |
| ENSMUSG00000028234 | 5007,303709 | 606,1 | 9492,3 | -3,955322411 | 4,37541E-07 | Rps20 | ribosomal protein S20 [Source:MGI Symbol;Acc:MGI:1914677] | 51,69 | 14,6 |
| ENSMUSG00000056209 | 52,41440675 | 8,6 | 113,9 | -3,949915017 | 0,000261448 | Npm3 | nucleoplasmin 3 [Source:MGI Symbol;Acc:MGI:894653] | 64,83 | 69,6 |
| ENSMUSG00000034674 | 632,448457 | 81,1 | 1242,1 | -3,948251308 | 7,87586E-06 | Tdg | thymine DNA glycosylase [Source:MGI Symbol;Acc:MGI:108247] | 38,11 | 40,2 |
| ENSMUSG00000080877 | 4,26348706 | 0,6 | 9,9 | -3,947384727 | 0,026680113 | Rpl22-ps1 | ribosomal protein L22, pseudogene 1 [Source:MGI Symbol;Acc:MGI:3644276] | #DIV/0! | #DIV/0! |
| ENSMUSG00000071041 | 5,387808748 | 0,7 | 10,9 | -3,934092664 | 0,021890207 | Gm15210 | predicted gene 15210 [Source:MGI Symbol;Acc:MGI:3705743] | #DIV/0! | #DIV/0! |
| ENSMUSG00000037795 | 101,1349663 | 16,3 | 149,3 | -3,929664069 | 0,040542253 | N4bp2 | NEDD4 binding protein 2 [Source:MGI Symbol;Acc:MGI:2684414] | 21,63 | 16,3 |
| ENSMUSG00000062296 | 89,17993815 | 12,0 | 227,7 | -3,928923868 | 0,003267939 | Trank1 | tetratricopeptide repeat and ankyrin repeat containing 1 [Source:MGI Symbol;Acc:MGI:1341834] | 15,90 | 20,3 |
| ENSMUSG00000033589 | 92,5586337 | 13,0 | 225,5 | -3,9208126 | 0,000206152 | Reep4 | receptor accessory protein 4 [Source:MGI Symbol;Acc:MGI:1919799] | 71,36 | 72,5 |
| ENSMUSG00000028873 | 127,1544822 | 14,4 | 238,2 | -3,908605113 | 0,003585785 | Cdca8 | cell division cycle associated 8 [Source:MGI Symbol;Acc:MGI:1196274] | 44,24 | 55,6 |
| ENSMUSG00000075266 | 339,8185315 | 36,2 | 568,1 | -3,906093396 | 3,109E-06 | Cenpw | centromere protein W [Source:MGI Symbol;Acc:MGI:1913561] | 87,28 | 27,7 |

| | | | | | | | | | |
|---|---|---|---|---|---|---|---|---|---|
| ENSMUSG00000057113 | 6483,911835 | 778,3 | 12048,4 | -3,895958317 | 1,76545E-07 | Npm1 | nucleophosmin 1 [Source:MGI Symbol;Acc:MGI:106184] | 92,12 | 80,7 |
| ENSMUSG00000033237 | 623,7472615 | 69,2 | 1036,6 | -3,893886703 | 2,10928E-07 | Arid2 | AT rich interactive domain 2 (ARID, RFX-like) [Source:MGI Symbol;Acc:MGI:1924294] | 56,18 | 30,0 |
| ENSMUSG00000097971 | 86,0807512 | 10,0 | 179,9 | -3,890048882 | 0,001850015 | Gm26917 | predicted gene, 26917 [Source:MGI Symbol;Acc:MGI:5504032] | #DIV/0! | #DIV/0! |
| ENSMUSG00000043987 | 48,42641378 | 10,7 | 110,0 | -3,880379426 | 0,001363825 | Cep164 | centrosomal protein 164 [Source:MGI Symbol;Acc:MGI:2384878] | 17,45 | 14,4 |
| ENSMUSG00000037742 | 24476,69821 | 3129,0 | 46682,3 | -3,873586275 | 4,43069E-07 | Eef1a1 | eukaryotic translation elongation factor 1 alpha 1 [Source:MGI Symbol;Acc:MGI:1096881] | 80,10 | 71,1 |
| ENSMUSG00000107176 | 120,7309483 | 14,6 | 228,5 | -3,865410418 | 7,08271E-06 | Gm9794 | predicted pseudogene 9794 [Source:MGI Symbol;Acc:MGI:3642653] | #DIV/0! | #DIV/0! |
| ENSMUSG00000047281 | 9,166071967 | 3,0 | 46,2 | -3,864521709 | 0,037914345 | Sfn | stratifin [Source:MGI Symbol;Acc:MGI:1891831] | 26,08 | 28,1 |
| ENSMUSG00000020460 | 4762,165298 | 623,7 | 9248,6 | -3,860575047 | 5,47199E-07 | Rps27a | ribosomal protein S27A [Source:MGI Symbol;Acc:MGI:1925544] | 72,54 | 71,7 |
| ENSMUSG00000105814 | 249,7029612 | 29,7 | 470,7 | -3,858615671 | 4,28999E-05 | Mir703 | microRNA 703 [Source:MGI Symbol;Acc:MGI:3629657] | #DIV/0! | #DIV/0! |
| ENSMUSG00000105388 | 13,40811053 | 1,9 | 27,4 | -3,857978878 | 0,003107701 | Rpl36a-ps2 | ribosomal protein L36A, pseudogene 2 [Source:MGI Symbol;Acc:MGI:3780887] | #DIV/0! | #DIV/0! |
| ENSMUSG00000060419 | 64,33230317 | 8,5 | 126,8 | -3,844541663 | 1,24537E-05 | Rps16-ps2 | ribosomal protein S16, pseudogene 2 [Source:MGI Symbol;Acc:MGI:3704259] | #DIV/0! | #DIV/0! |
| ENSMUSG00000058603 | 27,38239285 | 4,3 | 62,6 | -3,832877115 | 0,000913521 | Rpl28-ps1 | ribosomal protein L28, pseudogene 1 [Source:MGI Symbol;Acc:MGI:3705349] | #DIV/0! | #DIV/0! |
| ENSMUSG00000000682 | 157,4691545 | 9,1 | 300,7 | -3,832062558 | 0,006918725 | Cd52 | CD52 antigen [Source:MGI Symbol;Acc:MGI:1346088] | 91,89 | 65,6 |
| ENSMUSG00000106037 | 80,16321896 | 10,5 | 153,0 | -3,830583815 | 0,000175807 | Gm4332 | predicted gene 4332 [Source:MGI Symbol;Acc:MGI:3782515] | #DIV/0! | #DIV/0! |
| ENSMUSG00000072692 | 361,6777276 | 51,1 | 742,7 | -3,830143593 | 2,07836E-05 | Rpl37rt | ribosomal protein L37, retrotransposed [Source:MGI Symbol;Acc:MGI:3651519] | #DIV/0! | #DIV/0! |
| ENSMUSG00000034994 | 10952,36374 | 1370,5 | 19645,5 | -3,820031026 | 4,31989E-07 | Eef2 | eukaryotic translation elongation factor 2 [Source:MGI Symbol;Acc:MGI:95288] | 99,49 | 99,5 |
| ENSMUSG00000021733 | 143,9230226 | 24,1 | 599,9 | -3,819467209 | 0,016824826 | Slc4a7 | olute carrier family 4, sodium bicarbonate cotransporter, member 7 [Source:MGI Symbol;Acc:MGI:244387 | 37,56 | 36,6 |
| ENSMUSG00000041841 | 5777,786689 | 825,4 | 11719,5 | -3,806037508 | 3,52178E-06 | Rpl37 | ribosomal protein L37 [Source:MGI Symbol;Acc:MGI:1914531] | 59,09 | 45,8 |
| ENSMUSG00000028081 | 7035,223989 | 918,0 | 12993,1 | -3,801840478 | 2,37047E-07 | Rps3a1 | ribosomal protein S3A1 [Source:MGI Symbol;Acc:MGI:1202063] | 66,80 | 66,1 |
| ENSMUSG00000083337 | 43,40530525 | 5,4 | 80,5 | -3,798951452 | 0,000171406 | Gm11539 | predicted gene 11539 [Source:MGI Symbol;Acc:MGI:3651911] | #DIV/0! | #DIV/0! |
| ENSMUSG00000032399 | 16817,97492 | 2220,7 | 31493,7 | -3,796835035 | 2,29172E-07 | Rpl4 | ribosomal protein L4 [Source:MGI Symbol;Acc:MGI:1915141] | 100,00 | 98,3 |
| ENSMUSG00000089883 | 4,570287887 | 1,2 | 13,1 | -3,796538734 | 0,013075767 | Gm3355 | predicted gene 3355 [Source:MGI Symbol;Acc:MGI:3781533] | #DIV/0! | #DIV/0! |
| ENSMUSG00000001741 | 151,8625254 | 26,8 | 445,7 | -3,793042181 | 0,007266466 | Il16 | interleukin 16 [Source:MGI Symbol;Acc:MGI:1270855] | 47,88 | 34,1 |
| ENSMUSG00000083621 | 15,6856226 | 2,5 | 34,8 | -3,792534101 | 0,000416631 | Gm14586 | predicted gene 14586 [Source:MGI Symbol;Acc:MGI:3705507] | #DIV/0! | #DIV/0! |
| ENSMUSG00000027663 | 1207,558491 | 178,3 | 2516,4 | -3,787085639 | 1,90966E-07 | Zmat3 | zinc finger matrin type 3 [Source:MGI Symbol;Acc:MGI:1195270] | 87,75 | 53,4 |
| ENSMUSG00000090761 | 10,47378759 | 0,9 | 24,7 | -3,782798503 | 0,044715456 | Gm17201 | predicted gene 17201 [Source:MGI Symbol;Acc:MGI:4938028] | 19,15 | 48,0 |
| ENSMUSG00000022346 | 372,2263221 | 55,9 | 811,1 | -3,775260879 | 6,12101E-06 | Myc | myelocytomatosis oncogene [Source:MGI Symbol;Acc:MGI:97250] | 76,91 | 77,8 |
| ENSMUSG00000040212 | 186,6026729 | 22,6 | 309,6 | -3,771737362 | 0,000210376 | Emp3 | epithelial membrane protein 3 [Source:MGI Symbol;Acc:MGI:1098729] | 66,56 | 74,4 |
| ENSMUSG00000104649 | 16,85727186 | 3,1 | 44,7 | -3,768037767 | 0,000131934 | Gm43712 | predicted gene 43712 [Source:MGI Symbol;Acc:MGI:5663849] | #DIV/0! | #DIV/0! |
| ENSMUSG00000032803 | 1710,801444 | 211,5 | 2931,4 | -3,766093967 | 5,03624E-08 | Cdv3 | carnitine deficiency-associated gene expressed in ventricle 3 [Source:MGI Symbol;Acc:MGI:2448759] | 66,15 | 65,6 |
| ENSMUSG00000072640 | 551,3246775 | 105,7 | 1377,9 | -3,760435465 | 5,19236E-08 | Lyrm9 | LYR motif containing 9 [Source:MGI Symbol;Acc:MGI:1913524] | 46,65 | 26,4 |
| ENSMUSG00000083992 | 97,4696099 | 13,1 | 179,0 | -3,75576643 | 9,6913E-05 | Gm11478 | predicted gene 11478 [Source:MGI Symbol;Acc:MGI:3650888] | #DIV/0! | #DIV/0! |
| ENSMUSG00000067274 | 6635,247618 | 904,4 | 12313,4 | -3,752169344 | 1,50113E-07 | Rplp0 | ribosomal protein, large, P0 [Source:MGI Symbol;Acc:MGI:1927636] | 77,99 | 77,0 |
| ENSMUSG00000066626 | 5,736775791 | 0,9 | 13,4 | -3,751816566 | 0,020396796 | Gm5121 | predicted gene 5121 [Source:MGI Symbol;Acc:MGI:3647258] | #DIV/0! | #DIV/0! |
| ENSMUSG00000038900 | 2177,725545 | 307,5 | 4157,2 | -3,751349999 | 1,46512E-06 | Rpl12 | ribosomal protein L12 [Source:MGI Symbol;Acc:MGI:98002] | 68,63 | 74,1 |
| ENSMUSG00000083019 | 5,469753275 | 1,0 | 11,2 | -3,748037963 | 0,01262077 | Gm6222 | predicted pseudogene 6222 [Source:MGI Symbol;Acc:MGI:3648722] | #DIV/0! | #DIV/0! |
| ENSMUSG00000003970 | 7106,387782 | 979,0 | 13267,5 | -3,742325153 | 2,68524E-08 | Rpl8 | ribosomal protein L8 [Source:MGI Symbol;Acc:MGI:1350927] | 99,54 | 99,0 |
| ENSMUSG00000041219 | 320,5050964 | 28,4 | 382,2 | -3,728028144 | 0,000121115 | Arhgap11a | Rho GTPase activating protein 11A [Source:MGI Symbol;Acc:MGI:2444300] | 54,49 | 32,5 |
| ENSMUSG00000023027 | 901,7039206 | 118,4 | 1778,9 | -3,721847738 | 4,82546E-06 | Atf1 | activating transcription factor 1 [Source:MGI Symbol;Acc:MGI:1298366] | 77,33 | 79,1 |
| ENSMUSG00000037563 | 2004,079942 | 295,7 | 3922,5 | -3,71905362 | 7,12389E-07 | Rps16 | ribosomal protein S16 [Source:MGI Symbol;Acc:MGI:98118] | 44,30 | 41,1 |
| ENSMUSG00000101795 | 12,62989148 | 1,0 | 12,5 | -3,712250347 | 0,005174925 | Gm5835 | predicted gene 5835 [Source:MGI Symbol;Acc:MGI:3643957] | #DIV/0! | #DIV/0! |
| ENSMUSG00000020091 | 1276,168788 | 123,8 | 1779,6 | -3,706565136 | 0,000213007 | Eif4ebp2 | eukaryotic translation initiation factor 4E binding protein 2 [Source:MGI Symbol;Acc:MGI:109198] | 96,75 | 98,0 |
| ENSMUSG00000004661 | 80,09080059 | 18,6 | 211,1 | -3,702583254 | 0,02327851 | Arid3b | AT rich interactive domain 3B (BRIGHT-like) [Source:MGI Symbol;Acc:MGI:1930768] | 46,50 | 38,4 |
| ENSMUSG00000033713 | 2698,101568 | 398,1 | 5242,0 | -3,702205025 | 6,54077E-06 | Foxn3 | forkhead box N3 [Source:MGI Symbol;Acc:MGI:1918625] | 88,73 | 85,4 |
| ENSMUSG00000061983 | 1534,723109 | 214,5 | 2800,8 | -3,6844869 | 1,27617E-05 | Rps12 | ribosomal protein S12 [Source:MGI Symbol;Acc:MGI:98105] | 100,00 | 99,9 |
| ENSMUSG00000035284 | 217,9055455 | 21,2 | 236,7 | -3,682373461 | 0,003190773 | Vps13c | vacuolar protein sorting 13C (yeast) [Source:MGI Symbol;Acc:MGI:2444207] | 23,42 | 12,7 |
| ENSMUSG00000047215 | 6304,164006 | 893,5 | 11573,9 | -3,68164856 | 6,36686E-07 | Rpl9 | ribosomal protein L9 [Source:MGI Symbol;Acc:MGI:1298373] | 38,83 | 30,3 |
| ENSMUSG00000086841 | 417,3319593 | 56,5 | 743,2 | -3,67841003 | 7,89907E-07 | 2410006H16Rik | RIKEN cDNA 2410006H16 gene [Source:MGI Symbol;Acc:MGI:1916471] | #DIV/0! | #DIV/0! |
| ENSMUSG00000050299 | 148,2907605 | 21,4 | 285,7 | -3,668632862 | 1,21468E-05 | Gm9843 | predicted gene 9843 [Source:MGI Symbol;Acc:MGI:3708621] | 59,99 | 68,9 |
| ENSMUSG00000034786 | 25,37032233 | 4,4 | 66,5 | -3,661371549 | 0,000850563 | Gpsm3 | G-protein signalling modulator 3 (AGS3-like, C. elegans) [Source:MGI Symbol;Acc:MGI:2146785] | 35,12 | 42,3 |
| ENSMUSG00000049764 | 283,8061899 | 53,6 | 793,8 | -3,660118551 | 0,001079602 | Zfp280b | zinc finger protein 280B [Source:MGI Symbol;Acc:MGI:1927865] | 68,48 | 61,2 |
| ENSMUSG00000029622 | 1407,222163 | 207,9 | 2617,3 | -3,656803851 | 1,76712E-08 | Arpc1b | actin related protein 2/3 complex, subunit 1B [Source:MGI Symbol;Acc:MGI:1343142] | 64,01 | 44,1 |
| ENSMUSG00000069682 | 34,12562902 | 4,7 | 59,3 | -3,649604565 | 3,44747E-05 | Gm10275 | predicted pseudogene 10275 [Source:MGI Symbol;Acc:MGI:3704354] | #DIV/0! | #DIV/0! |
| ENSMUSG00000046364 | 2040,729934 | 299,3 | 3811,3 | -3,646183018 | 7,17095E-06 | Rpl27a | ribosomal protein L27A [Source:MGI Symbol;Acc:MGI:1347076] | 91,11 | 91,7 |
| ENSMUSG00000081051 | 44,09246092 | 6,0 | 79,7 | -3,64504446 | 0,000374782 | Gm15427 | predicted pseudogene 15427 [Source:MGI Symbol;Acc:MGI:3642341] | #DIV/0! | #DIV/0! |
| ENSMUSG00000081382 | 77,34487854 | 9,8 | 122,0 | -3,639372049 | 2,15802E-05 | Rpl18-ps1 | ribosomal protein L18, pseudogene 1 [Source:MGI Symbol;Acc:MGI:98018] | #DIV/0! | #DIV/0! |
| ENSMUSG00000057841 | 6551,687344 | 990,5 | 12270,3 | -3,617877175 | 1,16724E-06 | Rpl32 | ribosomal protein L32 [Source:MGI Symbol;Acc:MGI:98038] | 62,24 | 53,7 |
| ENSMUSG00000030536 | 457,7118066 | 58,6 | 842,4 | -3,609214331 | 2,15802E-05 | Iqgap1 | IQ motif containing GTPase activating protein 1 [Source:MGI Symbol;Acc:MGI:1352757] | 66,13 | 46,6 |
| ENSMUSG00000108520 | 42,66040505 | 7,5 | 107,7 | -3,603023474 | 0,026392096 | RP23-311C19.5 | | #DIV/0! | #DIV/0! |
| ENSMUSG00000046721 | 53,76513282 | 8,8 | 112,6 | -3,599220123 | 0,000486956 | Rpl14-ps1 | ribosomal protein L14, pseudogene 1 [Source:MGI Symbol;Acc:MGI:3710579] | #DIV/0! | #DIV/0! |
| ENSMUSG00000020661 | 470,3765152 | 81,2 | 989,7 | -3,598226389 | 3,52178E-06 | Dnmt3a | DNA methyltransferase 3A [Source:MGI Symbol;Acc:MGI:1261827] | 54,75 | 43,7 |
| ENSMUSG00000051650 | 175,9070434 | 42,6 | 534,1 | -3,597692795 | 0,01463197 | B3gnt2 | JDP-GlcNAc:betaGal beta-1,3-N-acetylglucosaminyltransferase 2 [Source:MGI Symbol;Acc:MGI:1889505 | 76,47 | 73,9 |
| ENSMUSG00000005447 | 242,0601082 | 31,1 | 395,9 | -3,594785934 | 1,48473E-05 | Pafah1b3 | platelet-activating factor acetylhydrolase, isoform 1b, subunit 3 [Source:MGI Symbol;Acc:MGI:108414] | 65,11 | 60,2 |
| ENSMUSG00000034342 | 518,3373836 | 62,0 | 854,8 | -3,594728189 | 0,000130703 | Cbl | Casitas B-lineage lymphoma [Source:MGI Symbol;Acc:MGI:88279] | 47,87 | 32,4 |
| ENSMUSG00000068966 | 65,20256288 | 14,0 | 108,4 | -3,588046902 | 0,031449128 | Zbtb34 | zinc finger and BTB domain containing 34 [Source:MGI Symbol;Acc:MGI:2685195] | 26,11 | 11,5 |
| ENSMUSG00000042453 | 56,25785456 | 5,5 | 89,2 | -3,585297829 | 0,004679087 | Reln | reelin [Source:MGI Symbol;Acc:MGI:103022] | 6,60 | 7,7 |
| ENSMUSG00000006333 | 6972,777405 | 1068,9 | 12835,1 | -3,580947178 | 1,51429E-06 | Rps9 | ribosomal protein S9 [Source:MGI Symbol;Acc:MGI:1924096] | 63,04 | 48,6 |
| ENSMUSG00000028018 | 39,57810117 | 6,1 | 159,3 | -3,57792821 | 0,016066419 | Gstcd | glutathione S-transferase, C-terminal domain containing [Source:MGI Symbol;Acc:MGI:1914803] | 19,64 | 29,6 |
| ENSMUSG00000102349 | 5422,44528 | 811,0 | 10992,7 | -3,577752685 | 0,000238304 | Gm37376 | predicted gene, 37376 [Source:MGI Symbol;Acc:MGI:5610604] | 100,00 | 100,0 |
| ENSMUSG00000067367 | 843,6789848 | 137,8 | 1649,6 | -3,574227904 | 1,73585E-05 | Lyar | Ly1 antibody reactive clone [Source:MGI Symbol;Acc:MGI:107470] | 85,39 | 86,7 |
| ENSMUSG00000092341 | 9007,697267 | 1378,8 | 18338,3 | -3,567126202 | 0,000226397 | Malat1 | stasis associated lung adenocarcinoma transcript 1 (non-coding RNA) [Source:MGI Symbol;Acc:MGI:191 | #DIV/0! | #DIV/0! |
| ENSMUSG00000026565 | 347,6187342 | 49,4 | 567,8 | -3,562424122 | 0,000377046 | Pou2f1 | POU domain, class 2, transcription factor 1 [Source:MGI Symbol;Acc:MGI:101898] | 26,89 | 21,5 |
| ENSMUSG00000071415 | 4645,282686 | 699,7 | 8521,2 | -3,560964177 | 7,22388E-06 | Rpl23 | ribosomal protein L23 [Source:MGI Symbol;Acc:MGI:1929455] | 93,26 | 88,6 |
| ENSMUSG00000028906 | 213,2978305 | 38,6 | 605,5 | -3,556560103 | 0,017409758 | Epb41 | erythrocyte membrane protein band 4.1 [Source:MGI Symbol;Acc:MGI:95401] | 51,77 | 39,7 |
| ENSMUSG00000062456 | 31,48860693 | 5,1 | 63,7 | -3,552932085 | 0,000253433 | Rpl9-ps6 | ribosomal protein L9, pseudogene 6 [Source:MGI Symbol;Acc:MGI:3642682] | 22,29 | 16,4 |
| ENSMUSG00000028438 | 49,3646373 | 8,6 | 105,2 | -3,552723183 | 0,009975811 | Kif24 | kinesin family member 24 [Source:MGI Symbol;Acc:MGI:1918345] | 11,46 | 10,2 |
| ENSMUSG00000062867 | 719,6069675 | 117,7 | 1343,3 | -3,55188122 | 6,89676E-07 | Impdh2 | inosine 5'-phosphate dehydrogenase 2 [Source:MGI Symbol;Acc:MGI:109367] | 83,47 | 73,3 |
| ENSMUSG00000098292 | 27,33103626 | 5,2 | 59,9 | -3,543193923 | 0,000131252 | Gm27194 | predicted gene 27194 [Source:MGI Symbol;Acc:MGI:5521037] | #DIV/0! | #DIV/0! |
| ENSMUSG00000099764 | 42,58025071 | 6,6 | 82,2 | -3,531607464 | 0,000641627 | Rps10-ps2 | ribosomal protein S10, pseudogene 2 [Source:MGI Symbol;Acc:MGI:3645627] | #DIV/0! | #DIV/0! |
| ENSMUSG00000038485 | 314,0211195 | 45,7 | 560,6 | -3,528496516 | 0,002949302 | Socs7 | suppressor of cytokine signaling 7 [Source:MGI Symbol;Acc:MGI:2651588] | 52,15 | 39,9 |
| ENSMUSG00000027331 | 206,0239098 | 16,2 | 548,2 | -3,522262929 | 0,020886638 | Knstrn | kinetochore-localized astrin/SPAG5 binding [Source:MGI Symbol;Acc:MGI:1289298] | 24,04 | 28,6 |
| ENSMUSG00000083757 | 17,80593412 | 2,7 | 36,5 | -3,517858532 | 0,008751033 | Rps15a-ps4 | ribosomal protein S15A, pseudogene 4 [Source:MGI Symbol;Acc:MGI:3652187] | #DIV/0! | #DIV/0! |
| ENSMUSG00000027932 | 41,7437033 | 9,0 | 138,6 | -3,512233037 | 0,022294132 | Slc27a3 | solute carrier family 27 (fatty acid transporter), member 3 [Source:MGI Symbol;Acc:MGI:1347358] | 46,11 | 28,1 |
| ENSMUSG00000076615 | 10,86635991 | 0,7 | 12,9 | -3,510843494 | 0,041507394 | Ighg3 | Immunoglobulin heavy constant gamma 3 [Source:MGI Symbol;Acc:MGI:2144790] | #DIV/0! | #DIV/0! |
| ENSMUSG00000048371 | 334,105496 | 50,6 | 597,1 | -3,510293501 | 6,00758E-05 | Pdp2 | pyruvate dehyrogenase phosphatase catalytic subunit 2 [Source:MGI Symbol;Acc:MGI:1918878] | 55,22 | 53,2 |
| ENSMUSG00000094935 | 8,65318337 | 1,1 | 17,7 | -3,501975503 | 0,019428823 | Gm9726 | predicted gene 9726 [Source:MGI Symbol;Acc:MGI:3648149] | 5,27 | 2,0 |
| ENSMUSG00000079435 | 2521,256704 | 399,7 | 4619,8 | -3,496320334 | 4,40626E-06 | Rpl36a | ribosomal protein L36A [Source:MGI Symbol;Acc:MGI:1201789] | 100,00 | 93,0 |
| ENSMUSG00000097141 | 644,9940887 | 85,7 | 961,8 | -3,492491159 | 5,06949E-05 | Gm10524 | predicted gene 10524 [Source:MGI Symbol;Acc:MGI:3642791] | 48,67 | 54,8 |
| ENSMUSG00000001435 | 94,39417384 | 18,3 | 174,2 | -3,491312031 | 0,012906012 | Col18a1 | collagen, type XVIII, alpha 1 [Source:MGI Symbol;Acc:MGI:88451] | 26,74 | 15,3 |
| ENSMUSG00000104581 | 11,216128 | 2,2 | 24,8 | -3,476393078 | 0,001962054 | Gm42497 | predicted gene 42497 [Source:MGI Symbol;Acc:MGI:5662634] | 1,40 | 1,2 |
| ENSMUSG00000031838 | 72,91954179 | 14,0 | 180,1 | -3,472446456 | 0,000154767 | Ifi30 | interferon gamma inducible protein 30 [Source:MGI Symbol;Acc:MGI:2137648] | 93,23 | 93,2 |
| ENSMUSG00000104960 | 269,8508187 | 35,9 | 397,1 | -3,469020727 | 0,002179882 | Snhg8 | small nucleolar RNA host gene 8 [Source:MGI Symbol;Acc:MGI:1917145] | #DIV/0! | #DIV/0! |
| ENSMUSG00000041560 | 1693,031917 | 332,7 | 3738,4 | -3,466105288 | 2,21133E-05 | Gltscr2 | glioma tumor suppressor candidate region gene 2 [Source:MGI Symbol;Acc:MGI:2154441] | 49,74 | 46,3 |
| ENSMUSG00000043424 | 28,66205651 | 3,1 | 36,2 | -3,458634709 | 0,016628071 | Eif3j2 | eukaryotic translation initiation factor 3, subunit J2 [Source:MGI Symbol;Acc:MGI:3704486] | 14,81 | 11,4 |
| ENSMUSG00000032298 | 176,4176712 | 42,0 | 473,7 | -3,455703549 | 0,000106962 | Neil1 | nei endonuclease VIII-like 1 (E. coli) [Source:MGI Symbol;Acc:MGI:1920024] | 96,72 | 85,6 |
| ENSMUSG00000047749 | 52,54159956 | 15,9 | 180,3 | -3,454878792 | 0,02395112 | Zc3hav1l | zinc finger CCCH-type, antiviral 1-like [Source:MGI Symbol;Acc:MGI:2443387] | 29,50 | 30,1 |
| ENSMUSG00000074129 | 6108,031172 | 1005,4 | 11325,3 | -3,45260639 | 7,17095E-06 | Rpl13a | ribosomal protein L13A [Source:MGI Symbol;Acc:MGI:1351455] | 100,00 | 99,4 |
| ENSMUSG00000029638 | 327,3509124 | 43,3 | 471,7 | -3,44982347 | 0,00133698 | Glcci1 | glucocorticoid induced transcript 1 [Source:MGI Symbol;Acc:MGI:2179717] | 49,33 | 27,7 |
| ENSMUSG00000062083 | 9,815216689 | 1,5 | 22,7 | -3,448919238 | 0,033116718 | Rpl13a-ps1 | ribosomal protein 13A, pseudogene 1 [Source:MGI Symbol;Acc:MGI:3648883] | 10,89 | 8,6 |
| ENSMUSG00000051413 | 126,8851439 | 24,0 | 261,1 | -3,443350655 | 5,81422E-05 | Plagl2 | pleiomorphic adenoma gene-like 2 [Source:MGI Symbol;Acc:MGI:1933165] | 48,60 | 19,4 |
| ENSMUSG00000083326 | 27,23004985 | 3,9 | 44,8 | -3,44056539 | 0,00300717 | Rpl38-ps1 | ribosomal protein L38, pseudogene 1 [Source:MGI Symbol;Acc:MGI:3649580] | #DIV/0! | #DIV/0! |
| ENSMUSG00000025192 | 75,73700187 | 17,4 | 196,5 | -3,436179393 | 0,003911784 | Entpd7 | ectonucleoside triphosphate diphosphohydrolase 7 [Source:MGI Symbol;Acc:MGI:2135885] | 35,18 | 20,4 |
| ENSMUSG00000030319 | 1049,719614 | 179,8 | 1984,1 | -3,435243159 | 4,50336E-06 | Cand2 | cullin-associated and neddylation-dissociated 2 (putative) [Source:MGI Symbol;Acc:MGI:1914338] | 72,99 | 11,2 |
| ENSMUSG00000056394 | 109,4886107 | 21,0 | 226,4 | -3,432897558 | 0,003510352 | Lig1 | ligase I, DNA, ATP-dependent [Source:MGI Symbol;Acc:MGI:101789] | 50,02 | 40,3 |
| ENSMUSG00000030067 | 702,13773 | 110,6 | 1275,2 | -3,432084886 | 0,000494516 | Foxp1 | forkhead box P1 [Source:MGI Symbol;Acc:MGI:1914004] | 67,15 | 49,9 |
| ENSMUSG00000030744 | 7066,868887 | 1198,7 | 12966,5 | -3,429732311 | 1,55505E-06 | Rps3 | ribosomal protein S3 [Source:MGI Symbol;Acc:MGI:1350917] | 95,25 | 95,2 |
| ENSMUSG00000074269 | 82,75741668 | 22,6 | 238,1 | -3,42877691 | 0,001083239 | Rec114 | REC114 meiotic recombination protein [Source:MGI Symbol;Acc:MGI:1920923] | 28,02 | 79,0 |
| ENSMUSG00000040618 | 524,74267 | 76,7 | 853,9 | -3,427295436 | 0,000104218 | Pck2 | phosphoenolpyruvate carboxykinase 2 (mitochondrial) [Source:MGI Symbol;Acc:MGI:1860456] | 85,25 | 73,8 |
| ENSMUSG00000060438 | 15,59265175 | 2,2 | 26,2 | -3,414899401 | 0,004371466 | Rps10-ps1 | ribosomal protein S10, pseudogene 1 [Source:MGI Symbol;Acc:MGI:3644167] | #DIV/0! | #DIV/0! |
| ENSMUSG00000025794 | 6977,535508 | 1218,9 | 12940,7 | -3,407211413 | 5,25356E-06 | Rpl14 | ribosomal protein L14 [Source:MGI Symbol;Acc:MGI:1914365] | 96,36 | 96,5 |
| ENSMUSG00000067288 | 1968,127397 | 336,2 | 3589,9 | -3,405684737 | 5,8902E-05 | Rps28 | ribosomal protein S28 [Source:MGI Symbol;Acc:MGI:1859516] | 68,22 | 52,6 |
| ENSMUSG00000038811 | 38,93501664 | 3,8 | 78,2 | -3,401926816 | 0,01843514 | Gngt2 | cleotide binding protein (G protein), gamma transducing activity polypeptide 2 [Source:MGI Symbol;Acc:M | 23,18 | 17,8 |
| ENSMUSG00000045128 | 6173,115318 | 998,3 | 10807,6 | -3,399256041 | 1,10796E-05 | Rpl18a | ribosomal protein L18A [Source:MGI Symbol;Acc:MGI:1924058] | 75,99 | 43,5 |
| ENSMUSG00000020649 | 554,0967423 | 78,0 | 991,1 | -3,388868726 | 0,003112756 | Rrm2 | ribonucleotide reductase M2 [Source:MGI Symbol;Acc:MGI:98181] | 89,36 | 83,6 |
| ENSMUSG00000058064 | 9,210480919 | 1,7 | 18,5 | -3,385342067 | 0,008502043 | Gm10036 | predicted gene 10036 [Source:MGI Symbol;Acc:MGI:3642334] | 14,90 | 19,6 |
| ENSMUSG00000082741 | 10,59940889 | 1,2 | 13,1 | -3,380721613 | 0,021626772 | Gm9703 | predicted gene 9703 [Source:MGI Symbol;Acc:MGI:3780110] | #DIV/0! | #DIV/0! |
| ENSMUSG00000025967 | 3304,879794 | 647,0 | 6702,1 | -3,366685645 | 4,01052E-07 | Eef1b2 | eukaryotic translation elongation factor 1 beta 2 [Source:MGI Symbol;Acc:MGI:1929520] | 81,06 | 59,6 |
| ENSMUSG00000059291 | 3162,814437 | 591,0 | 6143,7 | -3,360512327 | 3,65109E-06 | Rpl11 | ribosomal protein L11 [Source:MGI Symbol;Acc:MGI:1914275] | 92,14 | 69,3 |
| ENSMUSG00000012848 | 4655,01696 | 736,4 | 7638,9 | -3,35613699 | 3,46276E-05 | Rps5 | ribosomal protein S5 [Source:MGI Symbol;Acc:MGI:1097682] | 88,60 | 87,9 |
| ENSMUSG00000005225 | 361,4018895 | 65,3 | 670,0 | -3,354338805 | 1,64923E-05 | Plekha8 | ology domain containing, family A (phosphoinositide binding specific) member 8 [Source:MGI Symbol;Acc | 71,23 | 49,2 |
| ENSMUSG00000026407 | 102,1213589 | 16,4 | 196,0 | -3,347781455 | 0,011776046 | Cacna1s | calcium channel, voltage-dependent, L type, alpha 1S subunit [Source:MGI Symbol;Acc:MGI:88294] | 13,30 | 13,7 |
| ENSMUSG00000056612 | 2017,581495 | 358,0 | 3595,9 | -3,337369546 | 1,70388E-05 | Ppp1r14b | protein phosphatase 1, regulatory (inhibitor) subunit 14B [Source:MGI Symbol;Acc:MGI:107682] | 75,59 | 70,3 |
| ENSMUSG00000035476 | 93,73076867 | 35,1 | 400,8 | -3,334920705 | 0,03739492 | Tab3 | TGF-beta activated kinase 1/MAP3K7 binding protein 3 [Source:MGI Symbol;Acc:MGI:1913974] | 51,59 | 24,9 |
| ENSMUSG00000068101 | 202,0412335 | 22,1 | 234,8 | -3,333374404 | 0,003257241 | Cenpm | centromere protein M [Source:MGI Symbol;Acc:MGI:1913820] | 82,96 | 89,2 |
| ENSMUSG00000038970 | 452,1432298 | 96,3 | 1091,4 | -3,330980596 | 0,003302984 | Lmtk2 | lemur tyrosine kinase 2 [Source:MGI Symbol;Acc:MGI:3036247] | 81,01 | 44,6 |
| ENSMUSG00000038539 | 161,8936452 | 32,0 | 344,0 | -3,330578917 | 0,001526084 | Atf5 | activating transcription factor 5 [Source:MGI Symbol;Acc:MGI:2141857] | 69,73 | 44,5 |
| ENSMUSG00000059159 | 7,900374132 | 1,7 | 18,5 | -3,330256297 | 0,018196783 | Gm8129 | predicted pseudogene 8129 [Source:MGI Symbol;Acc:MGI:3646863] | #DIV/0! | #DIV/0! |
| ENSMUSG00000028495 | 3249,747196 | 573,7 | 5774,7 | -3,328382007 | 2,03568E-07 | Rps6 | ribosomal protein S6 [Source:MGI Symbol;Acc:MGI:98159] | 67,43 | 63,4 |
| ENSMUSG00000019066 | 264,5409044 | 42,0 | 450,0 | -3,327889744 | 0,002563239 | Rab3d | RAB3D, member RAS oncogene family [Source:MGI Symbol;Acc:MGI:97844] | 77,75 | 66,0 |
| ENSMUSG00000037926 | 283,0174312 | 45,4 | 564,0 | -3,325983881 | 0,004192399 | Ssh2 | slingshot homolog 2 (Drosophila) [Source:MGI Symbol;Acc:MGI:2679255] | 42,98 | 23,9 |
| ENSMUSG00000001280 | 382,9962677 | 92,8 | 913,1 | -3,325168779 | 0,000890876 | Sp1 | trans-acting transcription factor 1 [Source:MGI Symbol;Acc:MGI:98372] | 62,07 | 41,4 |
| ENSMUSG00000074578 | 424,7733538 | 59,4 | 652,3 | -3,322207516 | 0,000652266 | Zfos1 | zinc finger, NFX1-type containing 1, opposite strand RNA 1 [Source:MGI Symbol;Acc:MGI:1916199] | 87,99 | 64,4 |
| ENSMUSG00000011257 | 218,1655256 | 34,4 | 381,7 | -3,32164772 | 2,15802E-05 | Pabpc4 | poly(A) binding protein, cytoplasmic 4 [Source:MGI Symbol;Acc:MGI:2385206] | 57,21 | 40,5 |
| ENSMUSG00000037243 | 55,47062756 | 22,7 | 231,3 | -3,315017658 | 0,034990446 | Zfp692 | zinc finger protein 692 [Source:MGI Symbol;Acc:MGI:2144276] | 86,19 | 82,3 |
| ENSMUSG00000060143 | 268,806121 | 50,5 | 503,4 | -3,314169608 | 1,73695E-05 | Gm10076 | predicted gene 10076 [Source:MGI Symbol;Acc:MGI:3704451] | 86,28 | 50,3 |
| ENSMUSG00000007029 | 600,6031922 | 105,5 | 1089,3 | -3,312533447 | 8,56428E-05 | Vars | valyl-tRNA synthetase [Source:MGI Symbol;Acc:MGI:90675] | 86,22 | 74,8 |
| ENSMUSG00000038538 | 250,0557326 | 75,8 | 763,6 | -3,309764138 | 0,000218895 | Ubn2 | ubinuclein 2 [Source:MGI Symbol;Acc:MGI:2444236] | 39,42 | 22,5 |
| ENSMUSG00000059070 | 3805,434615 | 669,5 | 6651,3 | -3,309531537 | 4,04787E-06 | Rpl18 | ribosomal protein L18 [Source:MGI Symbol;Acc:MGI:98003] | 95,09 | 88,8 |
| ENSMUSG00000073640 | 14,22493101 | 2,7 | 27,1 | -3,293445693 | 0,004084001 | Rpl27-ps3 | ribosomal protein L27, pseudogene 3 [Source:MGI Symbol;Acc:MGI:3646174] | 10,04 | 10,5 |
| ENSMUSG00000037754 | 1047,116301 | 155,0 | 1619,6 | -3,288181686 | 0,001474981 | Ppp1r16b | protein phosphatase 1, regulatory (inhibitor) subunit 16B [Source:MGI Symbol;Acc:MGI:2151841] | 70,20 | 69,6 |
| ENSMUSG00000052698 | 160,1926656 | 42,6 | 335,0 | -3,283135385 | 0,02579482 | Tln2 | talin 2 [Source:MGI Symbol;Acc:MGI:1917799] | 34,97 | 12,3 |
| ENSMUSG00000010608 | 1222,711141 | 247,7 | 2643,9 | -3,278827823 | 0,000392863 | Rbm25 | RNA binding motif protein 25 [Source:MGI Symbol;Acc:MGI:1914289] | 87,35 | 70,7 |
| ENSMUSG00000081926 | 15,79534113 | 3,1 | 30,4 | -3,278069478 | 0,011576497 | Gm15536 | predicted gene 15536 [Source:MGI Symbol;Acc:MGI:3782984] | #DIV/0! | #DIV/0! |
| ENSMUSG00000024073 | 143,9999794 | 53,1 | 491,7 | -3,273442936 | 0,009302039 | Birc6 | baculoviral IAP repeat-containing 6 [Source:MGI Symbol;Acc:MGI:1276108] | 35,11 | 13,7 |
| ENSMUSG00000029992 | 585,3385595 | 126,4 | 1472,0 | -3,2713081 | 3,99098E-06 | Gfpt1 | glutamine fructose-6-phosphate transaminase 1 [Source:MGI Symbol;Acc:MGI:95698] | 80,96 | 50,6 |
| ENSMUSG00000018697 | 210,5129821 | 48,2 | 486,6 | -3,264442908 | 0,020396796 | Aatf | apoptosis antagonizing transcription factor [Source:MGI Symbol;Acc:MGI:1929608] | 96,21 | 95,4 |
| ENSMUSG00000005087 | 243,2337171 | 36,1 | 299,7 | -3,261027176 | 0,040488644 | Cd44 | CD44 antigen [Source:MGI Symbol;Acc:MGI:88338] | 49,78 | 19,3 |
| ENSMUSG00000020140 | 9,140367986 | 1,8 | 19,4 | -3,258956764 | 0,009864246 | Lgr5 | leucine rich repeat containing G protein coupled receptor 5 [Source:MGI Symbol;Acc:MGI:1341817] | 0,00 | 0,0 |
| ENSMUSG00000024991 | 2605,732091 | 392,1 | 4058,8 | -3,255963934 | 2,80086E-05 | Eif3a | eukaryotic translation initiation factor 3, subunit A [Source:MGI Symbol;Acc:MGI:95301] | 94,94 | 92,4 |
| ENSMUSG00000074800 | 128,7849952 | 24,8 | 238,5 | -3,255688014 | 0,000219136 | Gm4149 | predicted pseudogene 4149 [Source:MGI Symbol;Acc:MGI:3782325] | #DIV/0! | #DIV/0! |
| ENSMUSG00000060600 | 252,5216942 | 33,3 | 473,1 | -3,245750194 | 0,00903868 | Eno3 | enolase 3, beta muscle [Source:MGI Symbol;Acc:MGI:95395] | 82,02 | 65,0 |
| ENSMUSG00000073705 | 125,2782286 | 20,5 | 196,7 | -3,244913223 | 0,001619671 | Apitd1 | apoptosis-inducing, TAF9-like domain 1 [Source:MGI Symbol;Acc:MGI:1917178] | 62,91 | 49,6 |
| ENSMUSG00000102717 | 102,0782048 | 17,2 | 195,1 | -3,243412891 | 0,019280959 | Gm37759 | predicted gene, 37759 [Source:MGI Symbol;Acc:MGI:5610987] | 34,46 | 29,0 |
| ENSMUSG00000107724 | 84,06872107 | 9,5 | 89,0 | -3,239458662 | 0,030793208 | Gm16042 | predicted gene 16042 [Source:MGI Symbol;Acc:MGI:3801935] | #DIV/0! | #DIV/0! |
| ENSMUSG00000047045 | 1117,259559 | 204,1 | 2032,3 | -3,228611462 | 0,001753896 | Tmem164 | transmembrane protein 164 [Source:MGI Symbol;Acc:MGI:2148020] | 80,42 | 67,4 |
| ENSMUSG00000064337 | 411,642785 | 123,4 | 767,6 | -3,224981132 | 0,021097502 | mt-Rnr1 | mitochondrially encoded 12S rRNA [Source:MGI Symbol;Acc:MGI:102493] | 95,92 | 91,6 |
| ENSMUSG00000020075 | 359,3850571 | 74,6 | 613,1 | -3,212669437 | 0,000323542 | Ddx21 | DEAD (Asp-Glu-Ala-Asp) box polypeptide 21 [Source:MGI Symbol;Acc:MGI:1860494] | 87,60 | 54,1 |
| ENSMUSG00000062328 | 1807,984231 | 372,5 | 3448,9 | -3,202744013 | 2,295E-05 | Rpl17 | ribosomal protein L17 [Source:MGI Symbol;Acc:MGI:2448270] | 93,46 | 87,5 |
| ENSMUSG00000087590 | 104,5311206 | 27,7 | 250,3 | -3,197471125 | 0,000495092 | Epb41l4aos | erythrocyte membrane protein band 4.1 like 4a, opposite strand [Source:MGI Symbol;Acc:MGI:1916999] | #DIV/0! | #DIV/0! |
| ENSMUSG00000022565 | 269,6751429 | 21,4 | 280,4 | -3,196823338 | 0,00355232 | Plec | plectin [Source:MGI Symbol;Acc:MGI:1277961] | 14,60 | 10,7 |
| ENSMUSG00000043131 | 2915,651384 | 594,8 | 5588,2 | -3,18293955 | 3,02051E-06 | Mob1a | MOB kinase activator 1A [Source:MGI Symbol;Acc:MGI:2442631] | 98,10 | 96,1 |
| ENSMUSG00000004099 | 234,4502629 | 49,1 | 624,9 | -3,18084392 | 0,005385528 | Dnmt1 | DNA methyltransferase (cytosine-5) 1 [Source:MGI Symbol;Acc:MGI:94912] | 62,08 | 32,7 |
| ENSMUSG00000022177 | 108,9294988 | 25,6 | 224,9 | -3,178916145 | 0,025470758 | Haus4 | HAUS augmin-like complex, subunit 4 [Source:MGI Symbol;Acc:MGI:1261794] | 90,40 | 85,6 |
| ENSMUSG00000022856 | 121,8912262 | 33,0 | 304,3 | -3,175004494 | 0,019063996 | Tmem41a | transmembrane protein 41a [Source:MGI Symbol;Acc:MGI:1913914] | 93,33 | 88,1 |
| ENSMUSG00000105913 | 492,834296 | 93,7 | 860,4 | -3,17270974 | 8,64599E-06 | Mir5121 | microRNA 5121 [Source:MGI Symbol;Acc:MGI:4950446] | #DIV/0! | #DIV/0! |
| ENSMUSG00000057863 | 2897,925885 | 574,3 | 5232,3 | -3,170735729 | 0,000119227 | Rpl36 | ribosomal protein L36 [Source:MGI Symbol;Acc:MGI:1860603] | 87,87 | 88,0 |
| ENSMUSG00000093656 | 15,1237205 | 6,5 | 51,8 | -3,164966885 | 0,02528919 | Gm20628 | predicted gene 20628 [Source:MGI Symbol;Acc:MGI:5313075] | 20,35 | 7,9 |
| ENSMUSG00000036450 | 514,8784644 | 111,0 | 1095,8 | -3,164768438 | 0,001697391 | Hif1an | hypoxia-inducible factor 1, alpha subunit inhibitor [Source:MGI Symbol;Acc:MGI:2442345] | 94,51 | 77,3 |
| ENSMUSG00000083716 | 19,08863494 | 3,6 | 31,9 | -3,161762465 | 0,002376866 | Gm13436 | predicted gene 13436 [Source:MGI Symbol;Acc:MGI:3651005] | #DIV/0! | #DIV/0! |
| ENSMUSG00000032300 | 367,454057 | 86,7 | 783,5 | -3,158278741 | 0,023215974 | 1700017B05Rik | RIKEN cDNA 1700017B05 gene [Source:MGI Symbol;Acc:MGI:1921461] | 79,87 | 65,7 |

| | | | | | | | | | |
|---|---|---|---|---|---|---|---|---|---|
| ENSMUSG00000052146 | 1370,401729 | 267,6 | 2547,4 | -3,155389914 | 1,99491E-05 | Rps10 | ribosomal protein S10 [Source:MGI Symbol;Acc:MGI:1914347] | 81,89 | 68,8 |
| ENSMUSG00000008668 | 1926,011458 | 385,4 | 3441,1 | -3,152511531 | 4,78721E-05 | Rps18 | ribosomal protein S18 [Source:MGI Symbol;Acc:MGI:98146] | 76,44 | 71,8 |
| ENSMUSG00000064330 | 64,46916735 | 9,5 | 113,6 | -3,15181897 | 0,038466782 | Pde6h | phosphodiesterase 6H, cGMP-specific, cone, gamma [Source:MGI Symbol;Acc:MGI:1925850] | 3,64 | 1,9 |
| ENSMUSG00000105879 | 25,11052938 | 4,4 | 41,1 | -3,149029244 | 0,003553886 | Gm6204 | predicted gene 6204 [Source:MGI Symbol;Acc:MGI:3649038] | #DIV/0! | #DIV/0! |
| ENSMUSG00000000384 | 124,8485201 | 28,0 | 251,0 | -3,148996328 | 0,035985021 | Tbrg4 | transforming growth factor beta regulated gene 4 [Source:MGI Symbol;Acc:MGI:1100868] | 81,05 | 65,8 |
| ENSMUSG00000021391 | 44,65553291 | 21,6 | 181,2 | -3,147672512 | 0,02501549 | Cenpp | centromere protein P [Source:MGI Symbol;Acc:MGI:1913586] | 89,83 | 83,5 |
| ENSMUSG00000025362 | 1737,54051 | 349,2 | 3194,6 | -3,146905053 | 0,000212251 | Rps26 | ribosomal protein S26 [Source:MGI Symbol;Acc:MGI:1351628] | 97,17 | 97,7 |
| ENSMUSG00000074886 | 921,8778752 | 170,8 | 1545,8 | -3,138806577 | 5,63806E-05 | Grk6 | G protein-coupled receptor kinase 6 [Source:MGI Symbol;Acc:MGI:1347078] | 94,42 | 92,1 |
| ENSMUSG00000046330 | 3650,847536 | 757,0 | 6688,8 | -3,123765101 | 0,000132751 | Rpl37a | ribosomal protein L37a [Source:MGI Symbol;Acc:MGI:98068] | 100,00 | 99,6 |
| ENSMUSG00000066595 | 64,09964564 | 28,7 | 292,2 | -3,121538811 | 0,010341486 | Mfsd7b | major facilitator superfamily domain containing 7B [Source:MGI Symbol;Acc:MGI:2444881] | 64,03 | 45,7 |
| ENSMUSG00000058446 | 904,739529 | 166,4 | 1499,0 | -3,113952849 | 0,002300741 | Znrf2 | zinc and ring finger 2 [Source:MGI Symbol;Acc:MGI:1196246] | 85,05 | 77,8 |
| ENSMUSG00000106431 | 18,46018895 | 1,4 | 15,2 | -3,105033452 | 0,037087563 | Gm44492 | predicted gene, 44492 [Source:MGI Symbol;Acc:MGI:5690884] | #DIV/0! | #DIV/0! |
| ENSMUSG00000028382 | 590,4359488 | 80,5 | 749,8 | -3,103625328 | 0,008933079 | Ptbp3 | polypyrimidine tract binding protein 3 [Source:MGI Symbol;Acc:MGI:1923334] | 71,93 | 39,4 |
| ENSMUSG00000060671 | 295,0377479 | 74,7 | 785,8 | -3,097491525 | 0,008966028 | Atp8b2 | ATPase, class I, type 8B, member 2 [Source:MGI Symbol;Acc:MGI:1859660] | 80,35 | 61,6 |
| ENSMUSG00000031320 | 2902,037563 | 601,0 | 5246,0 | -3,092556917 | 1,21468E-05 | Rps4x | ribosomal protein S4, X-linked [Source:MGI Symbol;Acc:MGI:98158] | 91,73 | 86,3 |
| ENSMUSG00000107969 | 18,02640291 | 3,5 | 30,3 | -3,089177144 | 0,026732351 | Gm44085 | predicted gene, 44085 [Source:MGI Symbol;Acc:MGI:5690477] | 30,65 | 30,8 |
| ENSMUSG00000031245 | 735,3372975 | 162,5 | 1412,5 | -3,087579698 | 9,58593E-08 | Hmgn5 | high-mobility group nucleosome binding domain 5 [Source:MGI Symbol;Acc:MGI:1355295] | 96,26 | 94,4 |
| ENSMUSG00000067212 | 153,0823687 | 36,0 | 293,5 | -3,086179602 | 0,001469408 | H2-T23 | histocompatibility 2, T region locus 23 [Source:MGI Symbol;Acc:MGI:95957] | 78,44 | 76,5 |
| ENSMUSG00000024013 | 69,35504593 | 6,6 | 120,4 | -3,080070978 | 0,044019342 | Fgd2 | FYVE, RhoGEF and PH domain containing 2 [Source:MGI Symbol;Acc:MGI:1347084] | 38,13 | 35,8 |
| ENSMUSG00000093133 | 26,70464886 | 5,7 | 54,8 | -3,074197598 | 0,040204319 | Gm25994 | predicted gene, 25994 [Source:MGI Symbol;Acc:MGI:5455771] | #DIV/0! | #DIV/0! |
| ENSMUSG00000038708 | 938,6032226 | 263,6 | 2532,5 | -3,074105313 | 0,000140268 | Golga4 | golgi autoantigen, golgin subfamily a, 4 [Source:MGI Symbol;Acc:MGI:1859646] | 93,54 | 71,6 |
| ENSMUSG00000061313 | 242,7763165 | 55,2 | 513,2 | -3,065912862 | 0,010552384 | Ddhd2 | DDHD domain containing 2 [Source:MGI Symbol;Acc:MGI:1919358] | 84,03 | 67,6 |
| ENSMUSG00000026110 | 178,4534534 | 25,3 | 201,1 | -3,061612161 | 0,021430598 | Mgat4a | mannoside acetylglucosaminyltransferase 4, isoenzyme A [Source:MGI Symbol;Acc:MGI:2662992] | 36,50 | 16,6 |
| ENSMUSG00000064369 | 25,26997504 | 5,8 | 64,5 | -3,059046004 | 0,046874479 | mt-Te | mitochondrially encoded tRNA glutamic acid [Source:MGI Symbol;Acc:MGI:102488] | 86,96 | 79,7 |
| ENSMUSG00000040327 | 151,4968176 | 57,4 | 579,4 | -3,052067967 | 0,032646765 | Cul9 | cullin 9 [Source:MGI Symbol;Acc:MGI:1925559] | 62,67 | 59,7 |
| ENSMUSG00000042462 | 750,9273599 | 144,9 | 1209,7 | -3,051248399 | 0,000275443 | Dctpp1 | dCTP pyrophosphatase 1 [Source:MGI Symbol;Acc:MGI:1913672] | 100,00 | 98,0 |
| ENSMUSG00000032735 | 39,00064172 | 15,9 | 187,3 | -3,05049618 | 0,019207141 | Ablim3 | actin binding LIM protein family, member 3 [Source:MGI Symbol;Acc:MGI:2442582] | 43,44 | 48,4 |
| ENSMUSG00000067344 | 8,001528693 | 2,0 | 18,2 | -3,048951421 | 0,039878766 | Rps25-ps1 | ribosomal protein S25, pseudogene 1 [Source:MGI Symbol;Acc:MGI:3642902] | #DIV/0! | #DIV/0! |
| ENSMUSG00000038274 | 2436,760178 | 522,7 | 4377,7 | -3,04399687 | 3,74958E-05 | Fau | -Reilly murine sarcoma virus (FBR-MuSV) ubiquitously expressed (fox derived) [Source:MGI Symbol;Acc:M | 83,35 | 85,6 |
| ENSMUSG00000021215 | 348,1213893 | 63,8 | 551,7 | -3,042923636 | 0,014453882 | Net1 | neuroepithelial cell transforming gene 1 [Source:MGI Symbol;Acc:MGI:1927138] | 76,84 | 57,3 |
| ENSMUSG00000058761 | 134,484793 | 20,4 | 133,2 | -3,03686007 | 0,04335039 | Rnf169 | ring finger protein 169 [Source:MGI Symbol;Acc:MGI:1920257] | 29,43 | 17,4 |
| ENSMUSG00000034445 | 397,6392228 | 73,9 | 579,0 | -3,035888716 | 0,000286746 | Cyb561a3 | cytochrome b561 family, member A3 [Source:MGI Symbol;Acc:MGI:2686925] | 98,48 | 75,4 |
| ENSMUSG00000057322 | 2234,13364 | 477,9 | 3961,9 | -3,031042209 | 0,000105789 | Rpl38 | ribosomal protein L38 [Source:MGI Symbol;Acc:MGI:1914921] | 78,47 | 81,2 |
| ENSMUSG00000016487 | 292,8906568 | 67,0 | 587,7 | -3,027865965 | 0,008993454 | Ppfibp1 | PTPRF interacting protein, binding protein 1 (liprin beta 1) [Source:MGI Symbol;Acc:MGI:1914783] | 80,54 | 56,4 |
| ENSMUSG00000047675 | 2653,753854 | 569,4 | 4826,7 | -3,023021225 | 3,67112E-05 | Rps8 | ribosomal protein S8 [Source:MGI Symbol;Acc:MGI:98166] | 91,74 | 80,5 |
| ENSMUSG00000097908 | 887,9399922 | 111,1 | 981,9 | -3,022233329 | 0,000291915 | 4933404O12Rik | RIKEN cDNA 4933404O12 gene [Source:MGI Symbol;Acc:MGI:1914002] | #DIV/0! | #DIV/0! |
| ENSMUSG00000037544 | 35,68872767 | 3,2 | 24,0 | -3,005141893 | 0,037750973 | Dlgap5 | discs, large (Drosophila) homolog-associated protein 5 [Source:MGI Symbol;Acc:MGI:2183453] | 13,36 | 8,4 |
| ENSMUSG00000026020 | 486,0090303 | 98,6 | 761,6 | -3,00132364 | 0,003324528 | Nop58 | NOP58 ribonucleoprotein [Source:MGI Symbol;Acc:MGI:1933184] | 21,90 | 19,0 |
| ENSMUSG00000057406 | 345,4135896 | 93,7 | 916,4 | -3,00071339 | 0,009623952 | Whsc1 | Wolf-Hirschhorn syndrome candidate 1 (human) [Source:MGI Symbol;Acc:MGI:1276574] | 52,43 | 35,6 |
| ENSMUSG00000039987 | 163,3893365 | 44,1 | 410,7 | -3,000282823 | 0,007521921 | Phtf2 | putative homeodomain transcription factor 2 [Source:MGI Symbol;Acc:MGI:1916020] | 73,57 | 31,7 |
| ENSMUSG00000018583 | 2353,155654 | 531,2 | 4112,1 | -2,993425709 | 1,23964E-05 | G3bp1 | GTPase activating protein (SH3 domain) binding protein 1 [Source:MGI Symbol;Acc:MGI:1351465] | 86,70 | 80,1 |
| ENSMUSG00000100865 | 24,50930833 | 5,1 | 44,9 | -2,98113078 | 0,015786213 | Gm9320 | predicted gene 9320 [Source:MGI Symbol;Acc:MGI:3645000] | #DIV/0! | #DIV/0! |
| ENSMUSG00000040661 | 111,4025839 | 22,3 | 164,9 | -2,980283588 | 0,008569622 | Rad54l2 | RAD54 like 2 (S. cerevisiae) [Source:MGI Symbol;Acc:MGI:1933196] | 34,95 | 14,5 |
| ENSMUSG00000061477 | 2094,103572 | 446,9 | 3537,1 | -2,973645355 | 2,46932E-05 | Rps7 | ribosomal protein S7 [Source:MGI Symbol;Acc:MGI:1333818] | 67,36 | 60,8 |
| ENSMUSG00000025949 | 219,6153616 | 87,1 | 577,8 | -2,960322702 | 0,009970368 | Pikfyve | phosphoinositide kinase, FYVE finger containing [Source:MGI Symbol;Acc:MGI:1335106] | 51,34 | 27,5 |
| ENSMUSG00000046841 | 444,0738937 | 74,9 | 557,5 | -2,957825928 | 1,57793E-05 | Ckap4 | cytoskeleton-associated protein 4 [Source:MGI Symbol;Acc:MGI:2444926] | 96,67 | 81,0 |
| ENSMUSG00000053094 | 2005,455958 | 473,1 | 3860,8 | -2,956031741 | 0,000190848 | Tmem248 | transmembrane protein 248 [Source:MGI Symbol;Acc:MGI:1918917] | 88,42 | 87,1 |
| ENSMUSG00000081049 | 27,71143101 | 7,5 | 60,4 | -2,954555794 | 0,002960247 | Rps24-ps3 | ribosomal protein S24, pseudogene 3 [Source:MGI Symbol;Acc:MGI:3644407] | #DIV/0! | #DIV/0! |
| ENSMUSG00000025290 | 1789,098516 | 434,5 | 3408,9 | -2,953903531 | 1,57793E-05 | Rps24 | ribosomal protein S24 [Source:MGI Symbol;Acc:MGI:98147] | 76,07 | 88,2 |
| ENSMUSG00000094955 | 83,14454442 | 17,4 | 133,9 | -2,951792722 | 0,001031677 | Gm3699 | predicted gene 3699 [Source:MGI Symbol;Acc:MGI:3781875] | #DIV/0! | #DIV/0! |
| ENSMUSG00000020482 | 830,8046766 | 177,1 | 1569,0 | -2,945844463 | 0,001256291 | Ccdc117 | coiled-coil domain containing 117 [Source:MGI Symbol;Acc:MGI:2144383] | 99,32 | 94,0 |
| ENSMUSG00000054280 | 376,0929349 | 70,1 | 686,5 | -2,93215562 | 0,008795457 | Prr14l | proline rich 14-like [Source:MGI Symbol;Acc:MGI:2443658] | 53,65 | 36,6 |
| ENSMUSG00000030168 | 431,4546671 | 85,6 | 693,9 | -2,926325168 | 0,001886339 | Adipor2 | adiponectin receptor 2 [Source:MGI Symbol;Acc:MGI:93830] | 68,94 | 51,9 |
| ENSMUSG00000092274 | 367,5100363 | 81,8 | 651,1 | -2,925076944 | 0,001115225 | Neat1 | nuclear paraspeckle assembly transcript 1 (non-protein coding) [Source:MGI Symbol;Acc:MGI:1914211] | #DIV/0! | #DIV/0! |
| ENSMUSG00000101431 | 26,74581568 | 5,8 | 42,7 | -2,912513939 | 0,003281236 | Gm7901 | predicted gene 7901 [Source:MGI Symbol;Acc:MGI:3645047] | #DIV/0! | #DIV/0! |
| ENSMUSG00000081306 | 220,6377036 | 50,6 | 404,2 | -2,912296765 | 0,000361488 | Gm15266 | predicted gene 15266 [Source:MGI Symbol;Acc:MGI:3705867] | #DIV/0! | #DIV/0! |
| ENSMUSG00000025403 | 634,8699007 | 130,6 | 1050,2 | -2,903111251 | 7,37339E-07 | Shmt2 | serine hydroxymethyltransferase 2 (mitochondrial) [Source:MGI Symbol;Acc:MGI:1277989] | 95,33 | 93,3 |
| ENSMUSG00000028385 | 196,5338353 | 32,9 | 302,7 | -2,902036289 | 0,009208575 | Snx30 | sorting nexin family member 30 [Source:MGI Symbol;Acc:MGI:2443882] | 43,84 | 32,0 |
| ENSMUSG00000023025 | 723,1557319 | 162,1 | 1313,3 | -2,901646587 | 0,027311701 | Larp4 | La ribonucleoprotein domain family, member 4 [Source:MGI Symbol;Acc:MGI:2443114] | 82,54 | 52,9 |
| ENSMUSG00000004446 | 276,5869663 | 77,9 | 550,4 | -2,896316842 | 0,003324027 | Bid | BH3 interacting domain death agonist [Source:MGI Symbol;Acc:MGI:108093] | 77,00 | 36,8 |
| ENSMUSG00000008683 | 1768,797134 | 409,0 | 3145,7 | -2,88139042 | 0,000347895 | Rps15a | ribosomal protein S15A [Source:MGI Symbol;Acc:MGI:2389091] | 35,46 | 8,3 |
| ENSMUSG00000043923 | 630,8023349 | 149,0 | 1109,9 | -2,874303824 | 0,000106962 | Ccdc84 | coiled-coil domain containing 84 [Source:MGI Symbol;Acc:MGI:2685960] | 84,46 | 31,9 |
| ENSMUSG00000030313 | 196,3825221 | 52,3 | 399,2 | -2,873843588 | 0,003056312 | Dennd5b | DENN/MADD domain containing 5B [Source:MGI Symbol;Acc:MGI:2444273] | 44,41 | 21,6 |
| ENSMUSG00000040054 | 206,8267227 | 64,1 | 407,8 | -2,865799136 | 0,004175259 | Baz2a | bromodomain adjacent to zinc finger domain, 2A [Source:MGI Symbol;Acc:MGI:2151152] | 62,33 | 37,9 |
| ENSMUSG00000038612 | 2908,13207 | 660,4 | 4907,4 | -2,865627693 | 7,88054E-05 | Mcl1 | myeloid cell leukemia sequence 1 [Source:MGI Symbol;Acc:MGI:101769] | 99,96 | 99,0 |
| ENSMUSG00000030245 | 574,5348372 | 120,9 | 937,5 | -2,863382285 | 0,000617566 | Golt1b | golgi transport 1B [Source:MGI Symbol;Acc:MGI:1914214] | 99,46 | 78,7 |
| ENSMUSG00000053173 | 11,19119834 | 2,9 | 22,4 | -2,857400798 | 0,041726586 | Rpl18-ps2 | ribosomal protein L18, pseudogene 2 [Source:MGI Symbol;Acc:MGI:3646997] | #DIV/0! | #DIV/0! |
| ENSMUSG00000040945 | 1866,809958 | 405,6 | 3091,0 | -2,848984919 | 2,9727E-06 | Rcc2 | regulator of chromosome condensation 2 [Source:MGI Symbol;Acc:MGI:1919784] | 93,46 | 92,7 |
| ENSMUSG00000017404 | 3831,906176 | 933,3 | 6821,4 | -2,846447961 | 0,000164224 | Rpl19 | ribosomal protein L19 [Source:MGI Symbol;Acc:MGI:98020] | 94,36 | 85,4 |
| ENSMUSG00000098992 | 1236,982094 | 304,3 | 2219,1 | -2,842901996 | 0,000190079 | Gm27597 | predicted gene, 27597 [Source:MGI Symbol;Acc:MGI:5530979] | #DIV/0! | #DIV/0! |
| ENSMUSG00000085156 | 111,1370062 | 30,3 | 221,3 | -2,83654006 | 0,020521944 | Gm11974 | predicted gene 11974 [Source:MGI Symbol;Acc:MGI:3650059] | #DIV/0! | #DIV/0! |
| ENSMUSG00000107143 | 89,62268646 | 21,5 | 159,3 | -2,833356252 | 0,000211436 | Gm42469 | predicted gene 42469 [Source:MGI Symbol;Acc:MGI:5662606] | 18,90 | 8,6 |
| ENSMUSG00000060036 | 4534,716872 | 1055,9 | 7601,6 | -2,833146281 | 5,58598E-06 | Rpl3 | ribosomal protein L3 [Source:MGI Symbol;Acc:MGI:1351605] | 81,75 | 69,6 |
| ENSMUSG00000015363 | 813,261801 | 222,0 | 1617,6 | -2,827313344 | 0,002311456 | Trabd | TraB domain containing [Source:MGI Symbol;Acc:MGI:1915226] | 84,33 | 87,3 |
| ENSMUSG00000046434 | 914,4753717 | 227,6 | 1604,6 | -2,82698984 | 9,92487E-06 | Hnrnpa1 | heterogeneous nuclear ribonucleoprotein A1 [Source:MGI Symbol;Acc:MGI:104820] | 69,66 | 67,0 |
| ENSMUSG00000037295 | 137,5070369 | 30,1 | 255,1 | -2,82527468 | 0,009341226 | Ldlrap1 | low density lipoprotein receptor adaptor protein 1 [Source:MGI Symbol;Acc:MGI:2140175] | 35,22 | 27,1 |
| ENSMUSG00000038095 | 1132,580894 | 276,0 | 1945,5 | -2,821786591 | 1,18866E-05 | Sbno1 | sno, strawberry notch homolog 1 (Drosophila) [Source:MGI Symbol;Acc:MGI:2384298] | 81,47 | 61,5 |
| ENSMUSG00000034243 | 308,1333181 | 93,8 | 646,8 | -2,808783531 | 0,003447469 | Golgb1 | golgi autoantigen, golgin subfamily b, macrogolgin 1 [Source:MGI Symbol;Acc:MGI:1099447] | 52,11 | 34,2 |
| ENSMUSG00000104623 | 12,46596985 | 3,3 | 26,4 | -2,806610019 | 0,030683297 | Gm6394 | predicted gene 6394 [Source:MGI Symbol;Acc:MGI:3648319] | #DIV/0! | #DIV/0! |
| ENSMUSG00000065126 | 36,08367722 | 6,7 | 60,4 | -2,800469989 | 0,030229016 | Snord104 | small nucleolar RNA, C/D box 104 [Source:MGI Symbol;Acc:MGI:3819515] | #DIV/0! | #DIV/0! |
| ENSMUSG00000030019 | 330,6070839 | 70,3 | 489,9 | -2,796960929 | 0,008348916 | Fbxl14 | F-box and leucine-rich repeat protein 14 [Source:MGI Symbol;Acc:MGI:2141676] | 93,22 | 76,2 |
| ENSMUSG00000049036 | 137,999756 | 32,5 | 230,6 | -2,79466487 | 0,007327727 | Tmem121 | transmembrane protein 121 [Source:MGI Symbol;Acc:MGI:1916445] | 91,27 | 88,0 |
| ENSMUSG00000064345 | 491,8191797 | 198,0 | 779,5 | -2,793452306 | 0,031924952 | mt-Nd2 | mitochondrially encoded NADH dehydrogenase 2 [Source:MGI Symbol;Acc:MGI:102500] | 97,85 | 90,8 |
| ENSMUSG00000048109 | 98,42497924 | 23,3 | 152,8 | -2,793245264 | 0,019020269 | Rbm15 | RNA binding motif protein 15 [Source:MGI Symbol;Acc:MGI:2443205] | 32,52 | 12,6 |
| ENSMUSG00000029640 | 441,481566 | 142,3 | 988,3 | -2,788353292 | 0,002221384 | Usp12 | ubiquitin specific peptidase 12 [Source:MGI Symbol;Acc:MGI:1270128] | 79,79 | 77,5 |
| ENSMUSG00000024594 | 822,0327966 | 144,0 | 1051,8 | -2,787352227 | 0,001676218 | Prrc1 | proline-rich coiled-coil 1 [Source:MGI Symbol;Acc:MGI:1916106] | 96,28 | 70,2 |
| ENSMUSG00000041235 | 209,8849234 | 82,3 | 542,8 | -2,782751787 | 0,0171364 | Chd7 | chromodomain helicase DNA binding protein 7 [Source:MGI Symbol;Acc:MGI:2444748] | 48,28 | 28,0 |
| ENSMUSG00000086583 | 20,98233939 | 3,2 | 23,8 | -2,772675693 | 0,006985873 | Gm15500 | predicted pseudogene 15500 [Source:MGI Symbol;Acc:MGI:3782947] | #DIV/0! | #DIV/0! |
| ENSMUSG00000043716 | 4494,274326 | 1146,3 | 7868,7 | -2,77156225 | 4,78721E-05 | Rpl7 | ribosomal protein L7 [Source:MGI Symbol;Acc:MGI:98073] | 71,27 | 67,7 |
| ENSMUSG00000011254 | 73,30132653 | 18,0 | 150,2 | -2,771278267 | 0,012288115 | Thg1l | tRNA-histidine guanylyltransferase 1-like (S. cerevisiae) [Source:MGI Symbol;Acc:MGI:1913878] | 57,60 | 33,4 |
| ENSMUSG00000038893 | 200,9517047 | 62,6 | 478,3 | -2,769634065 | 0,011671606 | Fam117a | family with sequence similarity 117, member A [Source:MGI Symbol;Acc:MGI:2144564] | 91,25 | 78,9 |
| ENSMUSG00000022336 | 1520,257428 | 421,9 | 2994,5 | -2,765767287 | 0,002175921 | Eif3e | eukaryotic translation initiation factor 3, subunit E [Source:MGI Symbol;Acc:MGI:99257] | 99,37 | 97,4 |
| ENSMUSG00000062480 | 1006,75555 | 256,0 | 1735,3 | -2,761748725 | 0,000471215 | Acat3 | acetyl-Coenzyme A acetyltransferase 3 [Source:MGI Symbol;Acc:MGI:109182] | 71,13 | 16,7 |
| ENSMUSG00000061787 | 2309,568286 | 601,5 | 4096,3 | -2,757215888 | 0,000565985 | Rps17 | ribosomal protein S17 [Source:MGI Symbol;Acc:MGI:1309526] | 96,29 | 91,0 |
| ENSMUSG00000008730 | 1514,744392 | 397,4 | 2798,2 | -2,746839072 | 0,00119092 | Hipk1 | homeodomain interacting protein kinase 1 [Source:MGI Symbol;Acc:MGI:1314873] | 94,47 | 69,5 |
| ENSMUSG00000002997 | 283,5997213 | 98,4 | 831,9 | -2,744656385 | 0,025074159 | Prkar2b | protein kinase, cAMP dependent regulatory, type II beta [Source:MGI Symbol;Acc:MGI:97760] | 88,78 | 80,6 |
| ENSMUSG00000030432 | 1973,771962 | 522,0 | 3498,0 | -2,738638165 | 0,000123894 | Rpl28 | ribosomal protein L28 [Source:MGI Symbol;Acc:MGI:101839] | 71,64 | 52,7 |
| ENSMUSG00000078240 | 7,326310268 | 1,7 | 11,6 | -2,738261749 | 0,041751163 | Gm3550 | predicted gene 3550 [Source:MGI Symbol;Acc:MGI:3781727] | 2,58 | 3,8 |
| ENSMUSG00000018362 | 135,2730879 | 38,5 | 253,7 | -2,736829804 | 0,001227127 | Kpna2 | karyopherin (importin) alpha 2 [Source:MGI Symbol;Acc:MGI:103561] | 44,82 | 37,2 |
| ENSMUSG00000032050 | 2231,358465 | 523,8 | 3631,2 | -2,728851539 | 5,51281E-05 | Rdx | radixin [Source:MGI Symbol;Acc:MGI:97887] | 91,37 | 72,8 |
| ENSMUSG00000031939 | 143,1401122 | 34,1 | 279,0 | -2,726529971 | 0,003486296 | Taf1d | 'A box binding protein (Tbp)-associated factor, RNA polymerase I, D [Source:MGI Symbol;Acc:MGI:1922 | 88,56 | 60,4 |
| ENSMUSG00000035891 | 1437,96107 | 373,4 | 2674,7 | -2,72550567 | 0,000749905 | Cerk | ceramide kinase [Source:MGI Symbol;Acc:MGI:2386052] | 91,64 | 86,3 |
| ENSMUSG00000032604 | 437,9654717 | 132,2 | 921,1 | -2,723048395 | 0,005003395 | Qars | glutaminyl-tRNA synthetase [Source:MGI Symbol;Acc:MGI:1915851] | 83,66 | 66,7 |
| ENSMUSG00000093674 | 5878,878701 | 1538,3 | 10211,9 | -2,721072825 | 0,000652685 | Rpl41 | ribosomal protein L41 [Source:MGI Symbol;Acc:MGI:1915195] | 91,04 | 65,2 |
| ENSMUSG00000007050 | 430,6888982 | 107,6 | 803,9 | -2,720896986 | 0,001074931 | Lsm2 | LSM2 homolog, U6 small nuclear RNA associated (S. cerevisiae) [Source:MGI Symbol;Acc:MGI:90676] | 89,32 | 93,9 |
| ENSMUSG00000054008 | 806,8124191 | 250,3 | 1663,5 | -2,716423749 | 0,001237421 | Ndst1 | N-deacetylase/N-sulfotransferase (heparan glucosaminyl) 1 [Source:MGI Symbol;Acc:MGI:104719] | 99,37 | 75,0 |
| ENSMUSG00000049517 | 2022,520846 | 531,2 | 3493,2 | -2,712308579 | 0,000154358 | Rps23 | ribosomal protein S23 [Source:MGI Symbol;Acc:MGI:1913725] | 90,08 | 86,6 |
| ENSMUSG00000009876 | 87,96656764 | 26,9 | 186,4 | -2,692757673 | 0,000987959 | Cox4i2 | cytochrome c oxidase subunit IV isoform 2 [Source:MGI Symbol;Acc:MGI:2135755] | 88,03 | 65,0 |
| ENSMUSG00000061536 | 264,4414251 | 111,2 | 790,4 | -2,689859437 | 0,033174792 | Sec22c | SEC22 homolog C, vesicle trafficking protein [Source:MGI Symbol;Acc:MGI:2447871] | 62,02 | 53,6 |
| ENSMUSG00000026811 | 1305,285515 | 343,5 | 2253,5 | -2,683830401 | 0,001227127 | St6galnac6 | uraminyl-2,3-beta-galactosyl-1,3)-N-acetylgalactosaminide alpha-2,6-sialyltransferase 6 [Source:MGI Sym | 86,15 | 73,2 |
| ENSMUSG00000026131 | 225,4032055 | 66,5 | 427,5 | -2,682222749 | 0,006314703 | Dst | dystonin [Source:MGI Symbol;Acc:MGI:104627] | 17,99 | 7,0 |
| ENSMUSG00000060730 | 17,79673081 | 4,0 | 27,4 | -2,682124559 | 0,010393496 | Gm10086 | predicted pseudogene 10086 [Source:MGI Symbol;Acc:MGI:3642342] | #DIV/0! | #DIV/0! |
| ENSMUSG00000066232 | 544,3752766 | 160,3 | 1270,2 | -2,68126633 | 0,003757381 | Ipo7 | importin 7 [Source:MGI Symbol;Acc:MGI:2152414] | 97,72 | 78,3 |
| ENSMUSG00000006784 | 66,49025808 | 17,0 | 106,0 | -2,68003105 | 0,005687287 | Ttc25 | tetratricopeptide repeat domain 25 [Source:MGI Symbol;Acc:MGI:1921657] | 51,12 | 0,8 |
| ENSMUSG00000064326 | 402,2526838 | 104,7 | 678,3 | -2,661526642 | 0,001864447 | Siva1 | SIVA1, apoptosis-inducing factor [Source:MGI Symbol;Acc:MGI:1353606] | 94,71 | 88,6 |
| ENSMUSG00000025085 | 1060,428357 | 384,2 | 2561,8 | -2,647197094 | 0,00478346 | Ablim1 | actin-binding LIM protein 1 [Source:MGI Symbol;Acc:MGI:1194500] | 72,39 | 56,1 |
| ENSMUSG00000025138 | 356,0432829 | 96,3 | 671,5 | -2,64577053 | 0,001506164 | Sirt7 | sirtuin 7 [Source:MGI Symbol;Acc:MGI:2385849] | 93,21 | 59,5 |
| ENSMUSG00000029246 | 210,0401493 | 60,2 | 360,2 | -2,644384606 | 0,020349883 | Ppat | phosphoribosyl pyrophosphate amidotransferase [Source:MGI Symbol;Acc:MGI:2387203] | 86,76 | 44,2 |
| ENSMUSG00000009927 | 2954,88596 | 814,2 | 5100,5 | -2,63845456 | 0,000492468 | Rps25 | ribosomal protein S25 [Source:MGI Symbol;Acc:MGI:1922867] | 89,07 | 80,3 |
| ENSMUSG00000074280 | 93,64727601 | 24,9 | 176,7 | -2,636953287 | 0,00696183 | Gm6166 | predicted gene 6166 [Source:MGI Symbol;Acc:MGI:3645893] | #DIV/0! | #DIV/0! |
| ENSMUSG00000099708 | 31,67569911 | 10,0 | 64,0 | -2,634165972 | 0,015786213 | Gm29361 | predicted gene 29361 [Source:MGI Symbol;Acc:MGI:5580067] | 14,80 | 8,8 |
| ENSMUSG00000030045 | 406,3823924 | 144,3 | 939,5 | -2,632157598 | 0,000144914 | Mrpl19 | mitochondrial ribosomal protein L19 [Source:MGI Symbol;Acc:MGI:1926274] | 84,61 | 46,6 |
| ENSMUSG00000062006 | 1892,574734 | 510,4 | 3214,6 | -2,631688869 | 0,000666709 | Rpl34 | ribosomal protein L34 [Source:MGI Symbol;Acc:MGI:1915686] | 48,48 | 43,6 |
| ENSMUSG00000039844 | 702,5078494 | 178,6 | 1166,1 | -2,623522432 | 0,006792606 | Rapgef1 | Rap guanine nucleotide exchange factor (GEF) 1 [Source:MGI Symbol;Acc:MGI:104580] | 87,29 | 60,4 |
| ENSMUSG00000037646 | 257,2926401 | 53,3 | 340,3 | -2,621939338 | 0,001571784 | Vps13b | vacuolar protein sorting 13B (yeast) [Source:MGI Symbol;Acc:MGI:1916380] | 41,46 | 10,0 |
| ENSMUSG00000083022 | 8,430718471 | 2,2 | 13,0 | -2,621096397 | 0,03902333 | Rps15a-ps6 | ribosomal protein S15A, pseudogene 6 [Source:MGI Symbol;Acc:MGI:3650298] | #DIV/0! | #DIV/0! |
| ENSMUSG00000010048 | 160,3885364 | 45,6 | 308,6 | -2,617678616 | 0,005219865 | Ifrd2 | interferon-related developmental regulator 2 [Source:MGI Symbol;Acc:MGI:1316708] | 86,54 | 75,2 |
| ENSMUSG00000021149 | 601,0396389 | 187,3 | 1123,0 | -2,61626096 | 0,000344206 | Gtpbp4 | GTP binding protein 4 [Source:MGI Symbol;Acc:MGI:1916487] | 93,39 | 76,2 |
| ENSMUSG00000037805 | 1570,262274 | 486,4 | 3009,6 | -2,614638708 | 0,001115212 | Rpl10a | ribosomal protein L10A [Source:MGI Symbol;Acc:MGI:1343877] | 61,28 | 51,7 |
| ENSMUSG00000030275 | 1966,378523 | 521,8 | 3229,3 | -2,611311353 | 2,15802E-05 | Etnk1 | ethanolamine kinase 1 [Source:MGI Symbol;Acc:MGI:1922570] | 73,95 | 61,3 |
| ENSMUSG00000029616 | 1031,07195 | 290,5 | 1697,0 | -2,607654137 | 0,000132368 | Erp29 | endoplasmic reticulum protein 29 [Source:MGI Symbol;Acc:MGI:1914647] | 47,54 | 31,5 |
| ENSMUSG00000000282 | 80,27145264 | 36,3 | 210,0 | -2,60640123 | 0,027123079 | Mnt | max binding protein [Source:MGI Symbol;Acc:MGI:109150] | 62,26 | 41,3 |
| ENSMUSG00000042419 | 82,5116676 | 36,5 | 219,1 | -2,605187313 | 0,038269539 | Nfkbil1 | factor of kappa light polypeptide gene enhancer in B cells inhibitor like 1 [Source:MGI Symbol;Acc:MGI:13 | 89,15 | 82,1 |
| ENSMUSG00000093071 | 73,45940029 | 19,6 | 117,6 | -2,598701446 | 0,000652685 | Gm25951 | predicted gene, 25951 [Source:MGI Symbol;Acc:MGI:5455728] | #DIV/0! | #DIV/0! |
| ENSMUSG00000040658 | 136,1987126 | 47,5 | 285,3 | -2,594621322 | 0,017171551 | Dnph1 | 2'-deoxynucleoside 5'-phosphate N-hydrolase 1 [Source:MGI Symbol;Acc:MGI:3039376] | 99,49 | 97,4 |
| ENSMUSG00000080921 | 23,81036705 | 7,5 | 45,5 | -2,583300579 | 0,009540698 | Rpl38-ps2 | ribosomal protein L38, pseudogene 2 [Source:MGI Symbol;Acc:MGI:3646625] | #DIV/0! | #DIV/0! |
| ENSMUSG00000079260 | 139,0273002 | 26,3 | 172,3 | -2,582577279 | 0,007628162 | Tmppe | transmembrane protein with metallophosphoesterase domain [Source:MGI Symbol;Acc:MGI:5317335] | 39,44 | 15,7 |
| ENSMUSG00000023272 | 759,2696235 | 202,5 | 1117,8 | -2,572202125 | 0,000506281 | Creld2 | cysteine-rich with EGF-like domains 2 [Source:MGI Symbol;Acc:MGI:1923987] | 99,73 | 99,9 |
| ENSMUSG00000069972 | 25,48959665 | 6,9 | 41,1 | -2,569792546 | 0,003423993 | Rps13-ps2 | ribosomal protein S13, pseudogene 2 [Source:MGI Symbol;Acc:MGI:3704295] | #DIV/0! | #DIV/0! |
| ENSMUSG00000078193 | 77,33340911 | 25,8 | 153,8 | -2,563210722 | 0,010028845 | Gm2000 | predicted gene 2000 [Source:MGI Symbol;Acc:MGI:3780170] | 27,27 | 24,2 |
| ENSMUSG00000031388 | 766,1552288 | 268,5 | 1595,0 | -2,560988243 | 0,000351418 | Naa10 | N(alpha)-acetyltransferase 10, NatA catalytic subunit [Source:MGI Symbol;Acc:MGI:1915255] | 76,58 | 77,6 |
| ENSMUSG00000029415 | 195,2859722 | 31,5 | 207,0 | -2,558071039 | 0,027388901 | Sdad1 | SDA1 domain containing 1 [Source:MGI Symbol;Acc:MGI:2140779] | 56,52 | 29,7 |
| ENSMUSG00000003429 | 4102,407704 | 1154,4 | 6890,2 | -2,554113086 | 0,000910617 | Rps11 | ribosomal protein S11 [Source:MGI Symbol;Acc:MGI:1351329] | 83,39 | 83,4 |
| ENSMUSG00000053565 | 1779,189802 | 561,1 | 3282,0 | -2,546552653 | 0,000362015 | Eif3k | eukaryotic translation initiation factor 3, subunit K [Source:MGI Symbol;Acc:MGI:1921080] | 84,40 | 71,6 |
| ENSMUSG00000005583 | 957,0372554 | 175,2 | 1124,4 | -2,537581156 | 0,013341264 | Mef2c | myocyte enhancer factor 2C [Source:MGI Symbol;Acc:MGI:99458] | 76,41 | 45,6 |
| ENSMUSG00000042744 | 86,67409309 | 53,9 | 316,7 | -2,532818374 | 0,048051986 | Gm15800 | predicted gene 15800 [Source:MGI Symbol;Acc:MGI:3647820] | 37,19 | 14,3 |
| ENSMUSG00000068039 | 3732,511154 | 1040,4 | 5987,1 | -2,530588177 | 0,001128767 | Tcp1 | t-complex protein 1 [Source:MGI Symbol;Acc:MGI:98535] | 93,60 | 82,8 |
| ENSMUSG00000018160 | 559,1595285 | 164,8 | 988,8 | -2,521787533 | 0,001313881 | Med1 | mediator complex subunit 1 [Source:MGI Symbol;Acc:MGI:1100846] | 74,96 | 40,6 |
| ENSMUSG00000030870 | 1200,242068 | 314,0 | 1796,0 | -2,521615946 | 0,004682892 | Ubfd1 | ubiquitin family domain containing 1 [Source:MGI Symbol;Acc:MGI:107301] | 98,89 | 92,0 |
| ENSMUSG00000081344 | 46,65470233 | 14,8 | 88,6 | -2,52017325 | 0,009879242 | Gm14303 | predicted gene 14303 [Source:MGI Symbol;Acc:MGI:3651379] | #DIV/0! | #DIV/0! |
| ENSMUSG00000038642 | 448,9083965 | 51,0 | 588,0 | -2,51497332 | 0,04729708 | Ctss | cathepsin S [Source:MGI Symbol;Acc:MGI:107341] | 92,86 | 71,2 |
| ENSMUSG00000052040 | 1004,398834 | 272,7 | 1537,6 | -2,505311116 | 0,000753834 | Klf13 | Kruppel-like factor 13 [Source:MGI Symbol;Acc:MGI:1354948] | 44,30 | 39,5 |
| ENSMUSG00000029625 | 283,7644398 | 77,1 | 444,6 | -2,499971034 | 0,004107521 | Cpsf4 | cleavage and polyadenylation specific factor 4 [Source:MGI Symbol;Acc:MGI:1861602] | 93,99 | 85,7 |
| ENSMUSG00000030844 | 191,0170325 | 44,4 | 256,1 | -2,496775513 | 0,029679181 | Rgs10 | regulator of G-protein signalling 10 [Source:MGI Symbol;Acc:MGI:1915115] | 72,48 | 63,1 |
| ENSMUSG00000029014 | 731,0573432 | 268,1 | 1521,9 | -2,487190482 | 1,25744E-06 | Dnajc2 | DnaJ heat shock protein family (Hsp40) member C2 [Source:MGI Symbol;Acc:MGI:99470] | 72,26 | 73,9 |
| ENSMUSG00000034892 | 4204,406769 | 1244,4 | 7113,4 | -2,481884087 | 0,002985684 | Rps29 | ribosomal protein S29 [Source:MGI Symbol;Acc:MGI:107681] | 93,50 | 79,7 |
| ENSMUSG00000009575 | 2514,133514 | 750,8 | 4187,8 | -2,479551265 | 0,000146715 | Cbx5 | chromobox 5 [Source:MGI Symbol;Acc:MGI:109372] | 98,91 | 66,7 |
| ENSMUSG00000092827 | 9,750195646 | 3,4 | 25,1 | -2,470974588 | 0,049495484 | Mir3091 | microRNA 3091 [Source:MGI Symbol;Acc:MGI:4834305] | #DIV/0! | #DIV/0! |
| ENSMUSG00000049327 | 875,5310667 | 268,9 | 1533,5 | -2,469541301 | 0,001541667 | Setd8 | SET domain containing (lysine methyltransferase) 8 [Source:MGI Symbol;Acc:MGI:1915206] | 96,69 | 88,9 |
| ENSMUSG00000068036 | 213,1978179 | 89,4 | 523,7 | -2,468116745 | 0,001710679 | Mllt4 | oid or mixed-lineage leukemia (trithorax homolog, Drosophila); translocated to, 4 [Source:MGI Symbol;Acc: | 62,16 | 28,3 |
| ENSMUSG00000062762 | 3127,65182 | 841,4 | 4645,9 | -2,467298274 | 0,000907964 | Ei24 | etoposide induced 2.4 mRNA [Source:MGI Symbol;Acc:MGI:108090] | 95,22 | 94,5 |
| ENSMUSG00000059040 | 51,7660495 | 19,4 | 108,3 | -2,4648904 | 0,005174925 | Eno1b | enolase 1B, retrotransposed [Source:MGI Symbol;Acc:MGI:3648653] | 9,96 | 9,8 |
| ENSMUSG00000090000 | 1819,692041 | 611,1 | 3334,1 | -2,463551176 | 0,005455322 | Ier3ip1 | immediate early response 3 interacting protein 1 [Source:MGI Symbol;Acc:MGI:1913441] | 99,88 | 98,4 |
| ENSMUSG00000063556 | 118,599341 | 35,4 | 195,0 | -2,454933176 | 0,001775185 | Gm10132 | predicted gene 10132 [Source:MGI Symbol;Acc:MGI:3704449] | #DIV/0! | #DIV/0! |
| ENSMUSG00000007827 | 351,4357535 | 55,5 | 364,8 | -2,446287548 | 0,007837213 | Ankrd26 | ankyrin repeat domain 26 [Source:MGI Symbol;Acc:MGI:1917887] | 49,03 | 28,8 |
| ENSMUSG00000059474 | 209,3574337 | 51,5 | 306,5 | -2,440655436 | 0,010465506 | Mbtd1 | mbt domain containing 1 [Source:MGI Symbol;Acc:MGI:2143977] | 54,45 | 30,6 |
| ENSMUSG00000000740 | 4239,369353 | 1340,0 | 7298,1 | -2,439813905 | 0,000641048 | Rpl13 | ribosomal protein L13 [Source:MGI Symbol;Acc:MGI:105922] | 84,53 | 79,1 |
| ENSMUSG00000078515 | 401,8888895 | 108,0 | 605,7 | -2,432028863 | 0,013593364 | Ddi2 | DNA-damage inducible protein 2 [Source:MGI Symbol;Acc:MGI:1917244] | 71,38 | 33,4 |
| ENSMUSG00000032212 | 495,7833755 | 157,4 | 917,6 | -2,420230869 | 0,003629297 | Sltm | SAFB-like, transcription modulator [Source:MGI Symbol;Acc:MGI:1913910] | 97,07 | 72,5 |
| ENSMUSG00000027715 | 327,562454 | 61,1 | 389,8 | -2,419862036 | 0,003911784 | Ccna2 | cyclin A2 [Source:MGI Symbol;Acc:MGI:108069] | 43,17 | 27,9 |

| | | | | | | | | | |
|---|---|---|---|---|---|---|---|---|---|
| ENSMUSG00000001056 | 1199,455521 | 383,2 | 2195,4 | -2,41807779 | 0,001230025 | Nhp2 | NHP2 ribonucleoprotein [Source:MGI Symbol;Acc:MGI:1098547] | 93,86 | 76,9 |
| ENSMUSG00000027630 | 1354,388075 | 452,9 | 2697,4 | -2,41049349 | 0,023575759 | Tbl1xr1 | transducin (beta)-like 1X-linked receptor 1 [Source:MGI Symbol;Acc:MGI:2441730] | 89,30 | 78,4 |
| ENSMUSG00000047888 | 806,7169175 | 208,3 | 1171,8 | -2,407193861 | 0,023604417 | Tnrc6b | trinucleotide repeat containing 6b [Source:MGI Symbol;Acc:MGI:2443730] | 65,92 | 36,4 |
| ENSMUSG00000039001 | 2401,635816 | 808,1 | 4487,3 | -2,406922041 | 0,005578315 | Rps21 | ribosomal protein S21 [Source:MGI Symbol;Acc:MGI:1913731] | 99,75 | 99,2 |
| ENSMUSG00000064370 | 1269,584883 | 570,3 | 2333,8 | -2,405933538 | 0,042667317 | mt-Cytb | mitochondrially encoded cytochrome b [Source:MGI Symbol;Acc:MGI:102501] | 100,00 | 99,9 |
| ENSMUSG00000029147 | 488,9638481 | 176,0 | 1051,6 | -2,395224827 | 0,00602753 | Ppm1g | phosphatase 1G (formerly 2C), magnesium-dependent, gamma isoform [Source:MGI Symbol;Acc:MGI:1( | 94,32 | 78,6 |
| ENSMUSG00000024608 | 6040,472539 | 1867,0 | 9884,1 | -2,387567444 | 0,002347168 | Rps14 | ribosomal protein S14 [Source:MGI Symbol;Acc:MGI:98107] | 78,66 | 61,7 |
| ENSMUSG00000049354 | 2124,930454 | 717,1 | 4331,6 | -2,387158893 | 0,004736532 | Dcaf7 | DDB1 and CUL4 associated factor 7 [Source:MGI Symbol;Acc:MGI:1919083] | 99,02 | 86,6 |
| ENSMUSG00000067148 | 362,3540177 | 126,1 | 688,9 | -2,385495727 | 0,021531594 | Polr1c | polymerase (RNA) I polypeptide C [Source:MGI Symbol;Acc:MGI:103288] | 92,54 | 82,3 |
| ENSMUSG00000067870 | 45,58528201 | 13,8 | 77,9 | -2,379630422 | 0,026765281 | Rpl31-ps8 | ribosomal protein L31, pseudogene 8 [Source:MGI Symbol;Acc:MGI:3647726] | #DIV/0! | #DIV/0! |
| ENSMUSG00000026434 | 1736,777739 | 744,6 | 3922,2 | -2,375009144 | 0,00123279 | Nucks1 | nuclear casein kinase and cyclin-dependent kinase substrate 1 [Source:MGI Symbol;Acc:MGI:1934811] | 99,45 | 90,0 |
| ENSMUSG00000029687 | 140,707529 | 38,3 | 227,9 | -2,36906857 | 0,030153207 | Ezh2 | enhancer of zeste 2 polycomb repressive complex 2 subunit [Source:MGI Symbol;Acc:MGI:107940] | 64,82 | 31,9 |
| ENSMUSG00000037674 | 258,2206199 | 122,1 | 610,5 | -2,367477699 | 0,009417009 | Rfx7 | regulatory factor X, 7 [Source:MGI Symbol;Acc:MGI:2442675] | 74,85 | 43,6 |
| ENSMUSG00000031146 | 117,6098367 | 25,1 | 145,3 | -2,367063972 | 0,004301928 | Plp2 | proteolipid protein 2 [Source:MGI Symbol;Acc:MGI:1298382] | 64,30 | 55,4 |
| ENSMUSG00000037905 | 478,0635886 | 160,5 | 794,0 | -2,353726292 | 0,000480627 | Bri3bp | Bri3 binding protein [Source:MGI Symbol;Acc:MGI:1924059] | 74,47 | 32,5 |
| ENSMUSG00000070738 | 619,3128231 | 151,3 | 858,3 | -2,352600113 | 0,004061773 | Dgkd | diacylglycerol kinase, delta [Source:MGI Symbol;Acc:MGI:2138334] | 91,79 | 38,7 |
| ENSMUSG00000031389 | 405,9180891 | 159,0 | 817,2 | -2,336411069 | 0,002288581 | Arhgap4 | Rho GTPase activating protein 4 [Source:MGI Symbol;Acc:MGI:2159577] | 0,00 | 0,0 |
| ENSMUSG00000063316 | 1155,601922 | 374,5 | 1874,8 | -2,335783268 | 0,00133698 | Rpl27 | ribosomal protein L27 [Source:MGI Symbol;Acc:MGI:98036] | 72,72 | 72,9 |
| ENSMUSG00000033499 | 1547,787965 | 461,9 | 2422,9 | -2,330138126 | 0,00438382 | Larp4b | La ribonucleoprotein domain family, member 4B [Source:MGI Symbol;Acc:MGI:106330] | 97,07 | 89,0 |
| ENSMUSG00000109194 | 187,227815 | 60,5 | 307,8 | -2,32737694 | 0,00146498 | RP24-225H21.4 | | 14,22 | 14,2 |
| ENSMUSG00000020362 | 1855,787994 | 612,7 | 3361,0 | -2,324602013 | 0,005326329 | Cnot6 | CCR4-NOT transcription complex, subunit 6 [Source:MGI Symbol;Acc:MGI:2144529] | 98,61 | 87,5 |
| ENSMUSG00000091478 | 121,8944312 | 43,7 | 219,5 | -2,316106683 | 0,001031677 | Gm10039 | predicted pseudogene 10039 [Source:MGI Symbol;Acc:MGI:3704382] | #DIV/0! | #DIV/0! |
| ENSMUSG00000038738 | 263,6457897 | 96,7 | 604,9 | -2,313543807 | 0,002540682 | Shank1 | SH3/ankyrin domain gene 1 [Source:MGI Symbol;Acc:MGI:3613677] | 28,28 | 26,7 |
| ENSMUSG00000020315 | 573,7367697 | 156,8 | 1206,5 | -2,309152189 | 0,023617627 | Sptbn1 | spectrin beta, non-erythrocytic 1 [Source:MGI Symbol;Acc:MGI:98388] | 68,39 | 38,6 |
| ENSMUSG00000082755 | 19,94855407 | 6,7 | 37,7 | -2,307019431 | 0,041122242 | Gm8692 | predicted gene 8692 [Source:MGI Symbol;Acc:MGI:3646818] | #DIV/0! | #DIV/0! |
| ENSMUSG00000063457 | 2861,403994 | 959,3 | 4788,3 | -2,306708321 | 0,002039489 | Rps15 | ribosomal protein S15 [Source:MGI Symbol;Acc:MGI:98117] | 90,16 | 86,2 |
| ENSMUSG00000027330 | 269,6176326 | 89,6 | 419,4 | -2,301179811 | 0,016861921 | Cdc25b | cell division cycle 25B [Source:MGI Symbol;Acc:MGI:99701] | 90,07 | 67,1 |
| ENSMUSG00000079677 | 591,0955612 | 201,1 | 973,4 | -2,282050351 | 0,001811364 | Fdx1l | ferredoxin 1-like [Source:MGI Symbol;Acc:MGI:1915415] | 100,00 | 100,0 |
| ENSMUSG00000061665 | 226,5105165 | 58,0 | 415,6 | -2,281375688 | 0,032713879 | Cd2ap | CD2-associated protein [Source:MGI Symbol;Acc:MGI:1330281] | 79,41 | 40,4 |
| ENSMUSG00000035268 | 2064,61097 | 652,1 | 3203,0 | -2,278002248 | 0,007481539 | Pkig | protein kinase inhibitor, gamma [Source:MGI Symbol;Acc:MGI:1343086] | 86,98 | 83,1 |
| ENSMUSG00000044783 | 319,570986 | 132,2 | 715,8 | -2,276941819 | 0,028127231 | Hjurp | Holliday junction recognition protein [Source:MGI Symbol;Acc:MGI:2685821] | 84,57 | 53,5 |
| ENSMUSG00000028639 | 1856,535722 | 590,0 | 2884,8 | -2,276330921 | 0,000164224 | Ybx1 | Y box protein 1 [Source:MGI Symbol;Acc:MGI:99146] | 93,25 | 89,1 |
| ENSMUSG00000026028 | 1163,713224 | 507,8 | 2473,6 | -2,271898162 | 0,001111771 | Trak2 | trafficking protein, kinesin binding 2 [Source:MGI Symbol;Acc:MGI:1918077] | 99,22 | 81,7 |
| ENSMUSG00000032239 | 625,898957 | 225,7 | 1130,4 | -2,269543816 | 0,001506164 | Rp9 | retinitis pigmentosa 9 (human) [Source:MGI Symbol;Acc:MGI:2157166] | 99,85 | 87,5 |
| ENSMUSG00000020994 | 473,1706799 | 143,2 | 852,2 | -2,265732667 | 0,021389235 | Pnn | pinin [Source:MGI Symbol;Acc:MGI:1100514] | 93,60 | 65,3 |
| ENSMUSG00000049516 | 221,3971297 | 87,7 | 471,7 | -2,26453863 | 0,020593229 | Spty2d1 | SPT2, Suppressor of Ty, domain containing 1 (S. cerevisiae) [Source:MGI Symbol;Acc:MGI:2142062] | 85,88 | 44,5 |
| ENSMUSG00000032946 | 242,9910491 | 96,1 | 461,1 | -2,262466446 | 0,001809308 | Rasgrp2 | RAS, guanyl releasing protein 2 [Source:MGI Symbol;Acc:MGI:1333849] | 30,30 | 23,0 |
| ENSMUSG00000069135 | 337,5592519 | 155,7 | 769,3 | -2,258928619 | 0,01162181 | Fgfr1op | Fgfr1 oncogene partner [Source:MGI Symbol;Acc:MGI:1922546] | 83,24 | 66,5 |
| ENSMUSG00000009112 | 802,6421692 | 296,3 | 1673,8 | -2,258874715 | 0,016961595 | Bcl2l13 | BCL2-like 13 (apoptosis facilitator) [Source:MGI Symbol;Acc:MGI:2136959] | 86,55 | 74,6 |
| ENSMUSG00000073702 | 1395,639297 | 465,5 | 2252,3 | -2,250641689 | 0,003250126 | Rpl31 | ribosomal protein L31 [Source:MGI Symbol;Acc:MGI:2149632] | 77,84 | 55,1 |
| ENSMUSG00000064368 | 232,691076 | 99,4 | 397,7 | -2,24698004 | 0,048536982 | mt-Nd6 | mitochondrially encoded NADH dehydrogenase 6 [Source:MGI Symbol;Acc:MGI:102495] | 100,00 | 100,0 |
| ENSMUSG00000068735 | 280,6907696 | 70,5 | 349,6 | -2,245817887 | 0,013755682 | Trp53i11 | transformation related protein 53 inducible protein 11 [Source:MGI Symbol;Acc:MGI:2670995] | 92,17 | 77,2 |
| ENSMUSG00000001542 | 504,4877419 | 175,9 | 850,9 | -2,240738963 | 0,003911784 | Ell2 | elongation factor RNA polymerase II 2 [Source:MGI Symbol;Acc:MGI:2183438] | 97,89 | 86,9 |
| ENSMUSG00000108414 | 121,1299623 | 46,4 | 307,3 | -2,238206468 | 0,040351622 | Snhg1 | small nucleolar RNA host gene 1 [Source:MGI Symbol;Acc:MGI:3763743] | #DIV/0! | #DIV/0! |
| ENSMUSG00000023004 | 1193,171513 | 370,8 | 1803,8 | -2,23789101 | 0,000795283 | Tuba1b | tubulin, alpha 1B [Source:MGI Symbol;Acc:MGI:107804] | 37,81 | 31,8 |
| ENSMUSG00000005481 | 292,7414324 | 127,7 | 660,6 | -2,237284839 | 0,017922902 | Ddx39 | DEAD (Asp-Glu-Ala-Asp) box polypeptide 39 [Source:MGI Symbol;Acc:MGI:1915528] | 55,84 | 32,9 |
| ENSMUSG00000001525 | 1768,974525 | 570,7 | 2857,5 | -2,233341064 | 0,010254873 | Tubb5 | tubulin, beta 5 class I [Source:MGI Symbol;Acc:MGI:107812] | 87,07 | 87,7 |
| ENSMUSG00000027752 | 250,3694276 | 114,3 | 600,6 | -2,230441753 | 0,042287198 | Exosc8 | exosome component 8 [Source:MGI Symbol;Acc:MGI:1916889] | 83,51 | 75,4 |
| ENSMUSG00000026103 | 618,2451719 | 215,3 | 1012,1 | -2,223604703 | 0,007232084 | Gls | glutaminase [Source:MGI Symbol;Acc:MGI:95752] | 78,64 | 39,7 |
| ENSMUSG00000002222 | 901,2350117 | 287,7 | 1361,0 | -2,214521869 | 0,012884608 | Rmnd5a | required for meiotic nuclear division 5 homolog A (S. cerevisiae) [Source:MGI Symbol;Acc:MGI:1915727] | 94,26 | 71,5 |
| ENSMUSG00000023944 | 5424,421096 | 1938,0 | 8787,9 | -2,209574748 | 0,004084001 | Hsp90ab1 | heat shock protein 90 alpha (cytosolic), class B member 1 [Source:MGI Symbol;Acc:MGI:96247] | 98,17 | 95,8 |
| ENSMUSG00000040888 | 374,8900931 | 146,8 | 664,6 | -2,207729956 | 0,01543911 | Gfer | owth factor, erv1 (S. cerevisiae)-like (augmenter of liver regeneration) [Source:MGI Symbol;Acc:MGI:1077! | 82,70 | 66,3 |
| ENSMUSG00000031985 | 2087,854959 | 745,2 | 3634,2 | -2,207332522 | 0,001323936 | Gnpat | glyceronephosphate O-acyltransferase [Source:MGI Symbol;Acc:MGI:1343460] | 99,32 | 98,5 |
| ENSMUSG00000053510 | 890,0130168 | 375,9 | 1766,8 | -2,198030596 | 0,00809641 | Nrd1 | nardilysin, N-arginine dibasic convertase, NRD convertase 1 [Source:MGI Symbol;Acc:MGI:1201386] | 83,84 | 74,5 |
| ENSMUSG00000037503 | 4965,443605 | 1901,9 | 8599,9 | -2,19304745 | 0,003347776 | Fam168b | family with sequence similarity 168, member B [Source:MGI Symbol;Acc:MGI:2448487] | 99,04 | 96,4 |
| ENSMUSG00000041220 | 384,9630116 | 108,9 | 515,1 | -2,181799422 | 0,009954247 | Elovl6 | LOVL family member 6, elongation of long chain fatty acids (yeast) [Source:MGI Symbol;Acc:MGI:215652 | 59,19 | 29,9 |
| ENSMUSG00000082674 | 36,06293138 | 8,7 | 39,3 | -2,181464273 | 0,008902961 | Gm11914 | predicted gene 11914 [Source:MGI Symbol;Acc:MGI:3649889] | #DIV/0! | #DIV/0! |
| ENSMUSG00000029621 | 1691,156089 | 582,8 | 3059,2 | -2,180372259 | 0,018081101 | Arpc1a | actin related protein 2/3 complex, subunit 1A [Source:MGI Symbol;Acc:MGI:1928896] | 91,43 | 90,4 |
| ENSMUSG00000096993 | 186,0692305 | 53,9 | 246,1 | -2,176637449 | 0,005100924 | Gm26787 | predicted gene, 26787 [Source:MGI Symbol;Acc:MGI:5477281] | 25,27 | 13,5 |
| ENSMUSG00000020737 | 828,391758 | 291,2 | 1314,0 | -2,176613813 | 0,000220733 | Hn1 | hematological and neurological expressed sequence 1 [Source:MGI Symbol;Acc:MGI:1096361] | 94,98 | 93,6 |
| ENSMUSG00000006005 | 813,0551593 | 275,4 | 1244,8 | -2,176608894 | 0,000253001 | Tpr | translocated promoter region, nuclear basket protein [Source:MGI Symbol;Acc:MGI:1922066] | 81,03 | 41,5 |
| ENSMUSG00000025764 | 564,0191202 | 136,1 | 695,6 | -2,168770621 | 0,015231624 | Jade1 | jade family PHD finger 1 [Source:MGI Symbol;Acc:MGI:1925835] | 80,62 | 39,3 |
| ENSMUSG00000055760 | 175,3671282 | 78,5 | 450,5 | -2,164344211 | 0,021059027 | Gemin6 | gem (nuclear organelle) associated protein 6 [Source:MGI Symbol;Acc:MGI:1914492] | 96,36 | 77,7 |
| ENSMUSG00000058558 | 790,1055976 | 282,1 | 1320,1 | -2,148567335 | 0,00171394 | Rpl5 | ribosomal protein L5 [Source:MGI Symbol;Acc:MGI:102854] | 79,42 | 44,0 |
| ENSMUSG00000046865 | 140,585895 | 62,8 | 301,3 | -2,146521526 | 0,016481368 | Fbl | fibrillarin [Source:MGI Symbol;Acc:MGI:95486] | 80,62 | 63,5 |
| ENSMUSG00000107705 | 378,7233255 | 115,5 | 507,2 | -2,142348963 | 0,033841736 | Umad1 | UMAP1-MVP12 associated (UMA) domain containing 1 [Source:MGI Symbol;Acc:MGI:3840148] | #DIV/0! | #DIV/0! |
| ENSMUSG00000065922 | 128,0733355 | 59,4 | 255,9 | -2,139561655 | 0,045551113 | n-R5-8s1 | nuclear encoded rRNA 5.8S 1 [Source:MGI Symbol;Acc:MGI:4421735] | #DIV/0! | #DIV/0! |
| ENSMUSG00000026918 | 899,682144 | 254,8 | 1176,1 | -2,134500209 | 0,004166854 | Brd3 | bromodomain containing 3 [Source:MGI Symbol;Acc:MGI:1914632] | 90,54 | 70,0 |
| ENSMUSG00000019132 | 1033,262206 | 368,3 | 1722,1 | -2,13134336 | 0,006168003 | BC005537 | cDNA sequence BC005537 [Source:MGI Symbol;Acc:MGI:2441726] | 85,17 | 72,0 |
| ENSMUSG00000025130 | 1421,529502 | 671,0 | 2762,3 | -2,131298414 | 0,000935031 | P4hb | prolyl 4-hydroxylase, beta polypeptide [Source:MGI Symbol;Acc:MGI:97464] | 95,99 | 86,6 |
| ENSMUSG00000018446 | 1873,587826 | 755,6 | 3348,3 | -2,128023523 | 0,004054846 | C1qbp | complement component 1, q subcomponent binding protein [Source:MGI Symbol;Acc:MGI:1194505] | 100,00 | 98,1 |
| ENSMUSG00000035726 | 274,9624292 | 97,5 | 431,1 | -2,128003851 | 0,0178765 | Supt16 | suppressor of Ty 16 [Source:MGI Symbol;Acc:MGI:1890948] | 79,92 | 49,7 |
| ENSMUSG00000105703 | 90,2890643 | 30,0 | 139,5 | -2,120316197 | 0,004462196 | Gm43305 | predicted gene 43305 [Source:MGI Symbol;Acc:MGI:5663442] | 22,12 | 16,6 |
| ENSMUSG00000062997 | 1073,640105 | 423,7 | 1854,9 | -2,117641432 | 0,00393381 | Rpl35 | ribosomal protein L35 [Source:MGI Symbol;Acc:MGI:1913739] | 64,24 | 36,4 |
| ENSMUSG00000058600 | 1099,67125 | 403,7 | 1741,0 | -2,113318215 | 0,005484728 | Rpl30 | ribosomal protein L30 [Source:MGI Symbol;Acc:MGI:98037] | 88,55 | 54,3 |
| ENSMUSG00000028633 | 439,0450117 | 100,1 | 549,0 | -2,112902248 | 0,036800251 | Ctps | cytidine 5'-triphosphate synthase [Source:MGI Symbol;Acc:MGI:1858304] | 99,21 | 67,0 |
| ENSMUSG00000056692 | 770,1397863 | 280,8 | 1271,2 | -2,102133993 | 0,001188816 | D17Wsu92e | DNA segment, Chr 17, Wayne State University 92, expressed [Source:MGI Symbol;Acc:MGI:106281] | 94,62 | 70,4 |
| ENSMUSG00000094410 | 549,0351863 | 157,2 | 674,5 | -2,096201373 | 0,000820713 | Gm38394 | predicted gene, 38394 [Source:MGI Symbol;Acc:MGI:5614057] | 48,46 | 30,1 |
| ENSMUSG00000018846 | 964,7967308 | 371,4 | 1593,9 | -2,094465061 | 0,012504255 | Pank3 | pantothenate kinase 3 [Source:MGI Symbol;Acc:MGI:2387464] | 95,83 | 54,0 |
| ENSMUSG00000024617 | 562,8286501 | 309,1 | 1349,8 | -2,089623393 | 0,037127946 | Camk2a | calcium/calmodulin-dependent protein kinase II alpha [Source:MGI Symbol;Acc:MGI:88256] | 82,33 | 68,6 |
| ENSMUSG00000086839 | 150,5247406 | 55,1 | 227,3 | -2,072241909 | 0,021573976 | Gm11973 | predicted gene 11973 [Source:MGI Symbol;Acc:MGI:3650058] | 35,97 | 29,2 |
| ENSMUSG00000031490 | 589,8213547 | 274,5 | 1159,4 | -2,071670764 | 0,01046065 | Eif4ebp1 | eukaryotic translation initiation factor 4E binding protein 1 [Source:MGI Symbol;Acc:MGI:103267] | 95,75 | 84,6 |
| ENSMUSG00000025762 | 172,452711 | 82,0 | 386,3 | -2,058638291 | 0,008430306 | Larp1b | La ribonucleoprotein domain family, member 1B [Source:MGI Symbol;Acc:MGI:1914604] | 64,08 | 26,0 |
| ENSMUSG00000099083 | 352,0826511 | 181,4 | 727,8 | -2,058283227 | 0,017837685 | Atf7 | activating transcription factor 7 [Source:MGI Symbol;Acc:MGI:2443472] | 75,77 | 37,9 |
| ENSMUSG00000030403 | 128,9120175 | 49,8 | 230,7 | -2,05664981 | 0,040220583 | Vasp | vasodilator-stimulated phosphoprotein [Source:MGI Symbol;Acc:MGI:109268] | 88,16 | 72,6 |
| ENSMUSG00000085438 | 836,2955679 | 504,0 | 2014,8 | -2,056553294 | 0,025878717 | 1700020I14Rik | RIKEN cDNA 1700020I14 gene [Source:MGI Symbol;Acc:MGI:1913852] | #DIV/0! | #DIV/0! |
| ENSMUSG00000028069 | 266,254176 | 72,0 | 324,9 | -2,05296646 | 0,007393309 | Gpatch4 | G patch domain containing 4 [Source:MGI Symbol;Acc:MGI:1913864] | 74,50 | 59,4 |
| ENSMUSG00000057278 | 724,987449 | 264,2 | 1087,7 | -2,039368498 | 0,016438185 | Snrpg | small nuclear ribonucleoprotein polypeptide G [Source:MGI Symbol;Acc:MGI:1915261] | 48,09 | 32,5 |
| ENSMUSG00000049124 | 82,3274197 | 28,8 | 118,9 | -2,037757386 | 0,01149796 | Gm8186 | predicted gene 8186 [Source:MGI Symbol;Acc:MGI:3643110] | 75,99 | 42,0 |
| ENSMUSG00000018669 | 882,7082965 | 360,3 | 1501,2 | -2,036270153 | 0,020940193 | Cdk5rap3 | CDK5 regulatory subunit associated protein 3 [Source:MGI Symbol;Acc:MGI:1933126] | 88,88 | 77,7 |
| ENSMUSG00000083910 | 22,31957134 | 8,2 | 34,1 | -2,030167556 | 0,034734788 | Gm13604 | predicted gene 13604 [Source:MGI Symbol;Acc:MGI:3652203] | #DIV/0! | #DIV/0! |
| ENSMUSG00000102961 | 77,09375425 | 34,8 | 151,3 | -2,030111024 | 0,037127946 | Gm19918 | predicted gene, 19918 [Source:MGI Symbol;Acc:MGI:5012103] | 98,75 | 91,1 |
| ENSMUSG00000032312 | 237,4422593 | 59,8 | 442,6 | -2,016944357 | 0,04938358 | Csk | c-src tyrosine kinase [Source:MGI Symbol;Acc:MGI:88537] | 86,58 | 63,5 |
| ENSMUSG00000007739 | 1569,08876 | 588,9 | 2302,2 | -2,011873071 | 0,004307395 | Cct4 | chaperonin containing Tcp1, subunit 4 (delta) [Source:MGI Symbol;Acc:MGI:104689] | 95,53 | 77,7 |
| ENSMUSG00000001642 | 245,5242353 | 85,5 | 348,1 | -2,011059171 | 0,043026894 | Akr1b3 | aldo-keto reductase family 1, member B3 (aldose reductase) [Source:MGI Symbol;Acc:MGI:1353494] | 65,73 | 59,1 |
| ENSMUSG00000032601 | 1404,717129 | 573,5 | 2315,5 | -1,999030518 | 0,024619628 | Prkar2a | protein kinase, cAMP dependent regulatory, type II alpha [Source:MGI Symbol;Acc:MGI:108025] | 95,75 | 82,8 |
| ENSMUSG00000048668 | 313,684383 | 124,7 | 503,7 | -1,991254677 | 0,032996306 | Rhno1 | RAD9-HUS1-RAD1 interacting nuclear orphan 1 [Source:MGI Symbol;Acc:MGI:1915315] | 39,76 | 33,0 |
| ENSMUSG00000021427 | 1717,994465 | 668,7 | 3047,3 | -1,988867523 | 0,015473364 | Ssr1 | signal sequence receptor, alpha [Source:MGI Symbol;Acc:MGI:105082] | 88,57 | 47,3 |
| ENSMUSG00000028029 | 428,5752292 | 160,5 | 638,3 | -1,986625903 | 0,029520617 | Aimp1 | inoacyl tRNA synthetase complex-interacting multifunctional protein 1 [Source:MGI Symbol;Acc:MGI:1027 | 78,53 | 71,6 |
| ENSMUSG00000035171 | 800,5749716 | 404,8 | 1682,5 | -1,984905447 | 0,028180393 | 1110059E24Rik | RIKEN cDNA 1110059E24 gene [Source:MGI Symbol;Acc:MGI:1913456] | 67,38 | 58,1 |
| ENSMUSG00000036371 | 1912,041009 | 899,7 | 3565,5 | -1,970990974 | 0,004170187 | Serbp1 | serpine1 mRNA binding protein 1 [Source:MGI Symbol;Acc:MGI:1914120] | 56,26 | 24,2 |
| ENSMUSG00000043733 | 820,4975876 | 332,7 | 1364,9 | -1,970238151 | 0,040344484 | Ptpn11 | protein tyrosine phosphatase, non-receptor type 11 [Source:MGI Symbol;Acc:MGI:99511] | 80,08 | 58,3 |
| ENSMUSG00000029313 | 548,1441227 | 166,4 | 724,1 | -1,969034265 | 0,044261601 | Aff1 | AF4/FMR2 family, member 1 [Source:MGI Symbol;Acc:MGI:1100819] | 92,38 | 42,9 |
| ENSMUSG00000020863 | 662,6139839 | 281,5 | 1372,5 | -1,96817103 | 0,030961686 | Luc7l3 | LUC7-like 3 (S. cerevisiae) [Source:MGI Symbol;Acc:MGI:1914934] | 52,74 | 38,0 |
| ENSMUSG00000034636 | 396,1643748 | 230,9 | 862,6 | -1,957757351 | 0,017066302 | Zyg11b | zyg-ll family member B, cell cycle regulator [Source:MGI Symbol;Acc:MGI:2685277] | 94,55 | 59,6 |
| ENSMUSG00000031029 | 1098,986367 | 433,1 | 1731,9 | -1,956088402 | 0,008067252 | Eif3f | eukaryotic translation initiation factor 3, subunit F [Source:MGI Symbol;Acc:MGI:1913335] | 70,47 | 49,8 |
| ENSMUSG00000030431 | 200,027288 | 77,3 | 281,6 | -1,952906841 | 0,010931263 | Tmem238 | transmembrane protein 238 [Source:MGI Symbol;Acc:MGI:1922935] | 89,60 | 74,3 |
| ENSMUSG00000025813 | 496,3649634 | 196,8 | 768,5 | -1,952582019 | 0,006419354 | Homer2 | homer scaffolding protein 2 [Source:MGI Symbol;Acc:MGI:1347354] | 62,63 | 17,8 |
| ENSMUSG00000020677 | 405,525774 | 206,7 | 770,1 | -1,945426659 | 0,046765233 | Ddx52 | DEAD (Asp-Glu-Ala-Asp) box polypeptide 52 [Source:MGI Symbol;Acc:MGI:1925644] | 94,33 | 63,7 |
| ENSMUSG00000033987 | 189,3644065 | 81,4 | 314,4 | -1,944821472 | 0,008019388 | Dnah17 | dynein, axonemal, heavy chain 17 [Source:MGI Symbol;Acc:MGI:1917176] | 1,49 | 2,0 |
| ENSMUSG00000055239 | 1263,304131 | 562,2 | 2172,5 | -1,941717294 | 0,005720701 | Kcmf1 | potassium channel modulatory factor 1 [Source:MGI Symbol;Acc:MGI:1921537] | 92,72 | 64,8 |
| ENSMUSG00000090862 | 1361,808743 | 511,9 | 2011,3 | -1,938828312 | 0,005751825 | Rps13 | ribosomal protein S13 [Source:MGI Symbol;Acc:MGI:1915302] | 39,43 | 35,8 |
| ENSMUSG00000022241 | 391,1838439 | 158,3 | 662,4 | -1,935808758 | 0,035197959 | Tars | threonyl-tRNA synthetase [Source:MGI Symbol;Acc:MGI:106314] | 99,30 | 78,3 |
| ENSMUSG00000037331 | 850,6386719 | 294,5 | 1213,1 | -1,930015664 | 0,017570204 | Larp1 | La ribonucleoprotein domain family, member 1 [Source:MGI Symbol;Acc:MGI:1890165] | 90,39 | 73,1 |
| ENSMUSG00000029249 | 290,3959263 | 91,6 | 358,2 | -1,929577013 | 0,048790899 | Rest | RE1-silencing transcription factor [Source:MGI Symbol;Acc:MGI:104897] | 54,85 | 28,0 |
| ENSMUSG00000090553 | 1617,753716 | 608,9 | 2541,6 | -1,927611008 | 0,022233865 | Snrpe | small nuclear ribonucleoprotein E [Source:MGI Symbol;Acc:MGI:98346] | 52,89 | 51,0 |
| ENSMUSG00000102976 | 544,9747238 | 167,4 | 650,3 | -1,924166173 | 0,002214351 | Zc3h11a | zinc finger CCCH type containing 11A [Source:MGI Symbol;Acc:MGI:1917829] | 96,31 | 60,0 |
| ENSMUSG00000020857 | 2408,26337 | 999,5 | 3835,6 | -1,923658836 | 0,004092103 | Nme2 | NME/NM23 nucleoside diphosphate kinase 2 [Source:MGI Symbol;Acc:MGI:97356] | 83,80 | 72,3 |
| ENSMUSG00000001440 | 690,3493514 | 343,1 | 1389,6 | -1,912108426 | 0,012422812 | Kpnb1 | karyopherin (importin) beta 1 [Source:MGI Symbol;Acc:MGI:107532] | 93,13 | 66,6 |
| ENSMUSG00000034118 | 615,7568357 | 241,9 | 907,1 | -1,911919191 | 0,017387155 | Tpst1 | protein-tyrosine sulfotransferase 1 [Source:MGI Symbol;Acc:MGI:1298231] | 87,55 | 85,4 |
| ENSMUSG00000058799 | 1239,899022 | 427,5 | 1612,1 | -1,909626843 | 0,005971788 | Nap1l1 | nucleosome assembly protein 1-like 1 [Source:MGI Symbol;Acc:MGI:1855693] | 79,17 | 58,0 |
| ENSMUSG00000039382 | 950,3942333 | 364,2 | 1428,7 | -1,909143992 | 0,017459821 | Wdr45 | WD repeat domain 45 [Source:MGI Symbol;Acc:MGI:1859606] | 84,48 | 57,9 |
| ENSMUSG00000017639 | 1754,738925 | 766,2 | 2901,8 | -1,908235507 | 0,01949513 | Rab11fip4 | RAB11 family interacting protein 4 (class II) [Source:MGI Symbol;Acc:MGI:2442920] | 97,99 | 88,4 |
| ENSMUSG00000022587 | 478,1543835 | 192,8 | 840,4 | -1,905398835 | 0,006291444 | Ly6e | lymphocyte antigen 6 complex, locus E [Source:MGI Symbol;Acc:MGI:106651] | 68,69 | 46,4 |
| ENSMUSG00000105001 | 47,12090601 | 19,3 | 73,5 | -1,902319928 | 0,024534508 | AC117232.1 | | #DIV/0! | #DIV/0! |
| ENSMUSG00000022234 | 1538,296136 | 652,0 | 2621,3 | -1,899305899 | 0,021568781 | Cct5 | chaperonin containing Tcp1, subunit 5 (epsilon) [Source:MGI Symbol;Acc:MGI:107185] | 95,81 | 87,6 |
| ENSMUSG00000034024 | 1584,521891 | 577,2 | 2666,9 | -1,896572479 | 0,041917199 | Cct2 | chaperonin containing Tcp1, subunit 2 (beta) [Source:MGI Symbol;Acc:MGI:107186] | 99,90 | 94,6 |
| ENSMUSG00000026974 | 176,5012002 | 71,1 | 319,3 | -1,889322167 | 0,036907111 | Zmynd19 | zinc finger, MYND domain containing 19 [Source:MGI Symbol;Acc:MGI:1914437] | 62,52 | 40,8 |
| ENSMUSG00000090733 | 1872,028442 | 755,8 | 2792,5 | -1,885613528 | 0,015750201 | Rps27 | ribosomal protein S27 [Source:MGI Symbol;Acc:MGI:1888676] | 91,16 | 74,9 |
| ENSMUSG00000026615 | 1083,440134 | 505,0 | 2064,5 | -1,876413275 | 0,035858194 | Eprs | glutamyl-prolyl-tRNA synthetase [Source:MGI Symbol;Acc:MGI:97838] | 99,67 | 87,3 |
| ENSMUSG00000013076 | 271,7865757 | 91,2 | 331,2 | -1,864373374 | 0,014423451 | Amotl1 | angiomotin-like 1 [Source:MGI Symbol;Acc:MGI:1922973] | 73,19 | 18,2 |
| ENSMUSG00000046062 | 549,0193699 | 193,7 | 862,3 | -1,859596021 | 0,014556756 | Ppp1r15b | protein phosphatase 1, regulatory (inhibitor) subunit 15b [Source:MGI Symbol;Acc:MGI:2444211] | 92,15 | 55,5 |
| ENSMUSG00000024188 | 703,1997326 | 217,8 | 790,9 | -1,857877137 | 0,020923564 | Luc7l | Luc7 homolog (S. cerevisiae)-like [Source:MGI Symbol;Acc:MGI:1914228] | 89,97 | 51,2 |
| ENSMUSG00000056851 | 1144,737468 | 602,7 | 2190,3 | -1,853477619 | 0,003181242 | Pcbp2 | poly(rC) binding protein 2 [Source:MGI Symbol;Acc:MGI:108202] | 95,23 | 74,0 |
| ENSMUSG00000038072 | 830,4075458 | 289,5 | 1137,6 | -1,85183649 | 0,04621204 | Galnt11 | tyl-alpha-D-galactosamine:polypeptide N-acetylgalactosaminyltransferase 11 [Source:MGI Symbol;Acc:MG | 84,73 | 25,8 |
| ENSMUSG00000020766 | 377,117681 | 150,3 | 555,3 | -1,846176419 | 0,021158696 | Galk1 | galactokinase 1 [Source:MGI Symbol;Acc:MGI:95730] | 96,63 | 95,1 |
| ENSMUSG00000024516 | 1456,187784 | 624,3 | 2272,9 | -1,844074025 | 0,006566597 | Sec11c | SEC11 homolog C, signal peptidase complex subunit [Source:MGI Symbol;Acc:MGI:1913536] | 98,15 | 96,2 |
| ENSMUSG00000058546 | 1370,693598 | 567,7 | 2062,4 | -1,84353665 | 0,007716469 | Rpl23a | ribosomal protein L23A [Source:MGI Symbol;Acc:MGI:3040672] | 75,95 | 57,5 |
| ENSMUSG00000035295 | 283,5798677 | 130,5 | 472,2 | -1,842209054 | 0,01149796 | Wdr38 | WD repeat domain 38 [Source:MGI Symbol;Acc:MGI:1923896] | 28,62 | 0,0 |
| ENSMUSG00000021840 | 475,0928785 | 164,5 | 753,9 | -1,841935696 | 0,040356401 | Mapk1ip1l | mitogen-activated protein kinase 1 interacting protein 1-like [Source:MGI Symbol;Acc:MGI:2444022] | 71,92 | 49,2 |
| ENSMUSG00000027852 | 1382,003522 | 559,6 | 2061,9 | -1,841175577 | 0,020740575 | Nras | neuroblastoma ras oncogene [Source:MGI Symbol;Acc:MGI:97376] | 92,71 | 74,4 |
| ENSMUSG00000022216 | 889,8763999 | 413,4 | 1502,1 | -1,840571489 | 0,010769986 | Psme1 | roteasome (prosome, macropain) activator subunit 1 (PA28 alpha) [Source:MGI Symbol;Acc:MGI:109636 | 88,56 | 79,2 |
| ENSMUSG00000032525 | 365,9033534 | 65,8 | 263,7 | -1,829506592 | 0,024671043 | Nktr | natural killer tumor recognition sequence [Source:MGI Symbol;Acc:MGI:97346] | 55,56 | 28,5 |
| ENSMUSG00000022194 | 660,9788631 | 299,5 | 1199,0 | -1,828538832 | 0,009853147 | Pabpn1 | poly(A) binding protein, nuclear 1 [Source:MGI Symbol;Acc:MGI:1859158] | 63,48 | 64,1 |
| ENSMUSG00000030282 | 697,3176725 | 361,9 | 1290,2 | -1,826995653 | 0,009025989 | Cmas | cytidine monophospho-N-acetylneuraminic acid synthetase [Source:MGI Symbol;Acc:MGI:1337124] | 99,53 | 97,3 |
| ENSMUSG00000094483 | 1786,818222 | 786,3 | 2772,0 | -1,82330192 | 0,027142231 | Purb | purine rich element binding protein B [Source:MGI Symbol;Acc:MGI:1338779] | 99,11 | 69,6 |
| ENSMUSG00000079197 | 1045,94362 | 496,2 | 1758,9 | -1,813615852 | 0,004671396 | Psme2 | roteasome (prosome, macropain) activator subunit 2 (PA28 beta) [Source:MGI Symbol;Acc:MGI:1096365 | 89,20 | 85,2 |
| ENSMUSG00000039910 | 226,0789083 | 84,8 | 312,1 | -1,811044446 | 0,018671178 | Cited2 | 0-interacting transactivator, with Glu/Asp-rich carboxy-terminal domain, 2 [Source:MGI Symbol;Acc:MGI:1 | 96,27 | 73,9 |
| ENSMUSG00000105072 | 36,73334284 | 12,5 | 45,8 | -1,800186134 | 0,029520617 | Gm44335 | predicted gene, 44335 [Source:MGI Symbol;Acc:MGI:5690727] | #DIV/0! | #DIV/0! |
| ENSMUSG00000057522 | 1919,007695 | 774,9 | 2760,1 | -1,796794735 | 0,031022712 | Spop | speckle-type POZ protein [Source:MGI Symbol;Acc:MGI:1343085] | 98,17 | 97,7 |
| ENSMUSG00000024845 | 1071,246469 | 521,8 | 1857,9 | -1,786124754 | 0,011579966 | Tmem134 | transmembrane protein 134 [Source:MGI Symbol;Acc:MGI:1914240] | #DIV/0! | #DIV/0! |
| ENSMUSG00000020307 | 669,2382995 | 274,8 | 964,5 | -1,778496638 | 0,005739649 | Cdc34 | cell division cycle 34 [Source:MGI Symbol;Acc:MGI:102657] | 100,00 | 97,3 |
| ENSMUSG00000049932 | 398,6852773 | 135,0 | 485,2 | -1,771272264 | 0,043292904 | H2afx | H2A histone family, member X [Source:MGI Symbol;Acc:MGI:102688] | 99,49 | 97,1 |
| ENSMUSG00000032376 | 258,093774 | 99,3 | 380,6 | -1,755862809 | 0,048346724 | Usp3 | ubiquitin specific peptidase 3 [Source:MGI Symbol;Acc:MGI:2152450] | 69,04 | 38,2 |
| ENSMUSG00000029614 | 991,3116259 | 399,1 | 1357,0 | -1,751340101 | 0,012772283 | Rpl6 | ribosomal protein L6 [Source:MGI Symbol;Acc:MGI:108057] | 60,08 | 52,1 |
| ENSMUSG00000098274 | 1007,109937 | 433,6 | 1467,0 | -1,750677284 | 0,011677328 | Rpl24 | ribosomal protein L24 [Source:MGI Symbol;Acc:MGI:1915443] | 67,90 | 54,9 |
| ENSMUSG00000098371 | 1006,397036 | 433,3 | 1465,4 | -1,750398272 | 0,011843997 | Gm28037 | predicted gene, 28037 [Source:MGI Symbol;Acc:MGI:5547773] | #DIV/0! | #DIV/0! |
| ENSMUSG00000068328 | 960,0880752 | 517,4 | 1709,7 | -1,735767076 | 0,003898346 | Aup1 | ancient ubiquitous protein 1 [Source:MGI Symbol;Acc:MGI:107789] | 92,65 | 81,9 |
| ENSMUSG00000046562 | 1082,955718 | 668,7 | 2246,3 | -1,725947919 | 0,015376826 | Unc119b | unc-119 homolog B (C. elegans) [Source:MGI Symbol;Acc:MGI:2147162] | 98,84 | 97,8 |
| ENSMUSG00000053332 | 1880,499266 | 865,1 | 2827,7 | -1,719835539 | 0,01149796 | Gas5 | growth arrest specific 5 [Source:MGI Symbol;Acc:MGI:95659] | #DIV/0! | #DIV/0! |
| ENSMUSG00000086859 | 101,3227445 | 52,2 | 178,7 | -1,717448678 | 0,049666351 | Snhg20 | small nucleolar RNA host gene 20 [Source:MGI Symbol;Acc:MGI:1924222] | #DIV/0! | #DIV/0! |
| ENSMUSG00000040848 | 1696,488964 | 722,8 | 2607,2 | -1,709341604 | 0,009576467 | Sft2d2 | SFT2 domain containing 2 [Source:MGI Symbol;Acc:MGI:1917362] | 98,52 | 94,5 |
| ENSMUSG00000058013 | 553,5558121 | 326,6 | 1115,2 | -1,709018979 | 0,015765903 | Sept11 | septin 11 [Source:MGI Symbol;Acc:MGI:1277214] | 89,52 | 57,7 |
| ENSMUSG00000057541 | 236,2922534 | 79,6 | 230,0 | -1,700766593 | 0,021504289 | Pus7 | pseudouridylate synthase 7 homolog (S. cerevisiae) [Source:MGI Symbol;Acc:MGI:1925947] | 74,56 | 23,5 |
| ENSMUSG00000097234 | 62,66436412 | 37,0 | 119,9 | -1,694055171 | 0,032701694 | Gm26518 | predicted gene, 26518 [Source:MGI Symbol;Acc:MGI:5477012] | #DIV/0! | #DIV/0! |
| ENSMUSG00000031155 | 510,1901712 | 193,7 | 627,7 | -1,692583174 | 0,026907503 | Pim2 | proviral integration site 2 [Source:MGI Symbol;Acc:MGI:97587] | 91,90 | 88,6 |
| ENSMUSG00000021115 | 199,9917513 | 71,8 | 235,9 | -1,690594973 | 0,016602103 | Vrk1 | vaccinia related kinase 1 [Source:MGI Symbol;Acc:MGI:1261847] | 64,70 | 27,3 |
| ENSMUSG00000062647 | 1176,277331 | 539,1 | 1737,7 | -1,687696787 | 0,012275369 | Rpl7a | ribosomal protein L7A [Source:MGI Symbol;Acc:MGI:1353472] | 93,07 | 88,1 |
| ENSMUSG00000049751 | 712,9077747 | 347,9 | 1124,6 | -1,661880627 | 0,014785083 | Rpl36al | ribosomal protein L36A-like [Source:MGI Symbol;Acc:MGI:1913733] | 42,47 | 31,8 |
| ENSMUSG00000028433 | 363,9246458 | 153,1 | 476,2 | -1,651253355 | 0,023538411 | Ubap2 | ubiquitin-associated protein 2 [Source:MGI Symbol;Acc:MGI:1916176] | 95,16 | 59,4 |

| | | | | | | | | | |
|---|---|---|---|---|---|---|---|---|---|
| ENSMUSG00000029538 | 1927,565427 | 894,9 | 2802,4 | -1,650396913 | 0,019292726 | Srsf9 | serine/arginine-rich splicing factor 9 [Source:MGI Symbol;Acc:MGI:104896] | 100,00 | 99,8 |
| ENSMUSG00000039852 | 754,6646683 | 341,8 | 1069,8 | -1,643871172 | 0,034990446 | Rere | arginine glutamic acid dipeptide (RE) repeats [Source:MGI Symbol;Acc:MGI:2683486] | 89,15 | 45,0 |
| ENSMUSG00000028936 | 1429,951303 | 587,6 | 1858,5 | -1,643344124 | 0,022414935 | Rpl22 | ribosomal protein L22 [Source:MGI Symbol;Acc:MGI:99262] | 99,95 | 65,6 |
| ENSMUSG00000027999 | 749,584816 | 359,8 | 1106,9 | -1,629152547 | 0,006758871 | Pla2g12a | phospholipase A2, group XIIA [Source:MGI Symbol;Acc:MGI:1913600] | 92,49 | 80,9 |
| ENSMUSG00000022255 | 813,8068874 | 283,1 | 917,8 | -1,624084454 | 0,016226964 | Mtdh | metadherin [Source:MGI Symbol;Acc:MGI:1914404] | 98,53 | 53,4 |
| ENSMUSG00000041736 | 439,6890229 | 207,8 | 666,6 | -1,616775928 | 0,015702473 | Tspo | translocator protein [Source:MGI Symbol;Acc:MGI:88222] | 85,79 | 78,9 |
| ENSMUSG00000047721 | 1037,607185 | 489,8 | 1620,9 | -1,611282312 | 0,046896001 | Bola2 | bolA-like 2 (E. coli) [Source:MGI Symbol;Acc:MGI:1913412] | 83,90 | 46,8 |
| ENSMUSG00000020962 | 507,4896588 | 236,1 | 880,3 | -1,610614027 | 0,047671394 | Gtf2a1 | general transcription factor II A, 1 [Source:MGI Symbol;Acc:MGI:1933277] | 78,10 | 46,4 |
| ENSMUSG00000020922 | 653,7261249 | 305,5 | 955,8 | -1,604222897 | 0,036691306 | Lsm12 | LSM12 homolog (S. cerevisiae) [Source:MGI Symbol;Acc:MGI:1919592] | 97,33 | 94,9 |
| ENSMUSG00000020840 | 608,3958001 | 314,3 | 965,8 | -1,600952855 | 0,028127231 | Blmh | bleomycin hydrolase [Source:MGI Symbol;Acc:MGI:1345186] | 85,37 | 71,5 |
| ENSMUSG00000059213 | 255,4616999 | 177,1 | 488,0 | -1,599845381 | 0,026882294 | Ddn | dendrin [Source:MGI Symbol;Acc:MGI:108101] | 81,25 | 57,9 |
| ENSMUSG00000025364 | 449,663769 | 213,2 | 653,2 | -1,595263283 | 0,020865585 | Pa2g4 | proliferation-associated 2G4 [Source:MGI Symbol;Acc:MGI:894684] | 91,90 | 66,8 |
| ENSMUSG00000020484 | 1529,384883 | 504,8 | 1725,9 | -1,590412666 | 0,048661456 | Xbp1 | X-box binding protein 1 [Source:MGI Symbol;Acc:MGI:98970] | 76,30 | 81,4 |
| ENSMUSG00000026095 | 1023,117591 | 540,1 | 1634,3 | -1,585685964 | 0,042232315 | Asnsd1 | asparagine synthetase domain containing 1 [Source:MGI Symbol;Acc:MGI:1917646] | 85,81 | 83,8 |
| ENSMUSG00000025508 | 1915,392064 | 975,0 | 3035,0 | -1,585309932 | 0,02909233 | Rplp2 | ribosomal protein, large P2 [Source:MGI Symbol;Acc:MGI:1914436] | 62,74 | 42,4 |
| ENSMUSG00000031167 | 344,3472873 | 165,5 | 517,0 | -1,582978003 | 0,014910111 | Rbm3 | RNA binding motif protein 3 [Source:MGI Symbol;Acc:MGI:1099460] | 22,83 | 20,8 |
| ENSMUSG00000033685 | 1558,882554 | 616,2 | 1758,1 | -1,582213154 | 0,024345705 | Ucp2 | uncoupling protein 2 (mitochondrial, proton carrier) [Source:MGI Symbol;Acc:MGI:109354] | 41,39 | 35,9 |
| ENSMUSG00000092232 | 547,9166041 | 276,7 | 909,6 | -1,579036137 | 0,02097435 | Gm20521 | predicted gene 20521 [Source:MGI Symbol;Acc:MGI:5141986] | 78,58 | 52,3 |
| ENSMUSG00000024248 | 1001,61802 | 530,5 | 1586,2 | -1,577515157 | 0,008448967 | Cox7a2l | cytochrome c oxidase subunit VIIa polypeptide 2-like [Source:MGI Symbol;Acc:MGI:106015] | 82,33 | 71,0 |
| ENSMUSG00000040701 | 93,31832649 | 64,6 | 202,5 | -1,57410998 | 0,033916681 | Ap1g2 | adaptor protein complex AP-1, gamma 2 subunit [Source:MGI Symbol;Acc:MGI:1328307] | 40,71 | 37,1 |
| ENSMUSG00000024661 | 17122,44557 | 8833,9 | 25673,0 | -1,548844492 | 0,038318939 | Fth1 | ferritin heavy polypeptide 1 [Source:MGI Symbol;Acc:MGI:95588] | 100,00 | 100,0 |
| ENSMUSG00000018559 | 283,6212283 | 169,8 | 525,9 | -1,541357657 | 0,032234575 | Ctdnep1 | CTD nuclear envelope phosphatase 1 [Source:MGI Symbol;Acc:MGI:1914431] | 93,48 | 84,9 |
| ENSMUSG00000026087 | 1351,124825 | 751,2 | 2157,5 | -1,519205091 | 0,014287526 | Mrpl30 | mitochondrial ribosomal protein L30 [Source:MGI Symbol;Acc:MGI:1333820] | 57,57 | 30,0 |
| ENSMUSG00000037894 | 887,6720257 | 441,2 | 1262,8 | -1,508568653 | 0,014274168 | H2afz | H2A histone family, member Z [Source:MGI Symbol;Acc:MGI:1888388] | 72,50 | 66,0 |
| ENSMUSG00000036781 | 2500,904047 | 1245,3 | 3537,6 | -1,503845315 | 0,020374405 | Rps27l | ribosomal protein S27-like [Source:MGI Symbol;Acc:MGI:1915191] | 100,00 | 94,5 |
| ENSMUSG00000023050 | 793,909324 | 516,2 | 1488,5 | -1,493898588 | 0,041785654 | Map3k12 | mitogen-activated protein kinase kinase kinase 12 [Source:MGI Symbol;Acc:MGI:1346881] | 86,03 | 39,3 |
| ENSMUSG00000060636 | 487,9981432 | 254,9 | 714,6 | -1,479172483 | 0,028752676 | Rpl35a | ribosomal protein L35A [Source:MGI Symbol;Acc:MGI:1928894] | 70,92 | 55,0 |
| ENSMUSG00000042305 | 1401,817417 | 922,6 | 2495,1 | -1,450965537 | 0,020610821 | Tmem183a | transmembrane protein 183A [Source:MGI Symbol;Acc:MGI:1914729] | 97,97 | 96,5 |
| ENSMUSG00000025381 | 1174,393307 | 724,0 | 1981,3 | -1,444115694 | 0,012772283 | Cnpy2 | canopy FGF signaling regulator 2 [Source:MGI Symbol;Acc:MGI:1928477] | 97,51 | 96,1 |
| ENSMUSG00000042747 | 1247,90687 | 623,0 | 1705,6 | -1,440700269 | 0,029889368 | Krtcap2 | keratinocyte associated protein 2 [Source:MGI Symbol;Acc:MGI:1913309] | 74,64 | 47,8 |
| ENSMUSG00000068749 | 780,9165316 | 444,9 | 1295,5 | -1,43694269 | 0,049533768 | Psma5 | proteasome (prosome, macropain) subunit, alpha type 5 [Source:MGI Symbol;Acc:MGI:1347009] | 76,30 | 70,4 |
| ENSMUSG00000002014 | 723,4029094 | 411,7 | 1174,5 | -1,420919906 | 0,038925983 | Ssr4 | signal sequence receptor, delta [Source:MGI Symbol;Acc:MGI:1099464] | 99,92 | 93,5 |
| ENSMUSG00000012405 | 1125,718014 | 626,6 | 1682,6 | -1,3935643 | 0,03848596 | Rpl15 | ribosomal protein L15 [Source:MGI Symbol;Acc:MGI:1913730] | 81,60 | 56,3 |
| ENSMUSG00000032215 | 711,4573511 | 377,0 | 976,1 | -1,340784348 | 0,021890207 | Rsl24d1 | ribosomal L24 domain containing 1 [Source:MGI Symbol;Acc:MGI:2681840] | 77,84 | 76,1 |
| ENSMUSG00000052397 | 3201,255426 | 1881,3 | 4642,6 | -1,339596073 | 0,036909524 | Ezr | ezrin [Source:MGI Symbol;Acc:MGI:98931] | 92,84 | 91,7 |
| ENSMUSG00000027479 | 3264,489934 | 1737,3 | 4213,5 | -1,249305238 | 0,031661078 | Mapre1 | microtubule-associated protein, RP/EB family, member 1 [Source:MGI Symbol;Acc:MGI:891995] | 82,72 | 81,2 |
| ENSMUSG00000030695 | 6852,655302 | 9155,1 | 4096,1 | 1,156792513 | 0,028422025 | Aldoa | aldolase A, fructose-bisphosphate [Source:MGI Symbol;Acc:MGI:87994] | 78,40 | 54,6 |
| ENSMUSG00000078923 | 207,1349922 | 262,8 | 111,3 | 1,243365042 | 0,049402668 | Ube2v1 | ubiquitin-conjugating enzyme E2 variant 1 [Source:MGI Symbol;Acc:MGI:1913839] | 37,68 | 16,7 |
| ENSMUSG00000033751 | 369,530322 | 542,8 | 225,4 | 1,288144208 | 0,049415649 | Gadd45gip1 | wth arrest and DNA-damage-inducible, gamma interacting protein 1 [Source:MGI Symbol;Acc:MGI:19149 | 74,57 | 40,7 |
| ENSMUSG00000024099 | 2884,777939 | 4121,3 | 1704,8 | 1,302587323 | 0,038303911 | Ndufv2 | NADH dehydrogenase (ubiquinone) flavoprotein 2 [Source:MGI Symbol;Acc:MGI:1920150] | 83,33 | 56,7 |
| ENSMUSG00000011752 | 471,4363048 | 564,3 | 231,8 | 1,319376649 | 0,027864545 | Pgam1 | phosphoglycerate mutase 1 [Source:MGI Symbol;Acc:MGI:97552] | 40,71 | 25,9 |
| ENSMUSG00000072235 | 1496,440879 | 2315,2 | 941,1 | 1,34071097 | 0,03834123 | Tuba1a | tubulin, alpha 1A [Source:MGI Symbol;Acc:MGI:98869] | 63,47 | 60,9 |
| ENSMUSG00000039682 | 1068,970325 | 1447,4 | 583,2 | 1,346464514 | 0,044124433 | Lap3 | leucine aminopeptidase 3 [Source:MGI Symbol;Acc:MGI:1914238] | 99,44 | 90,6 |
| ENSMUSG00000031633 | 5994,209438 | 8533,9 | 3340,3 | 1,351773778 | 0,022655966 | Slc25a4 | r family 25 (mitochondrial carrier, adenine nucleotide translocator), member 4 [Source:MGI Symbol;Acc:MGI | 88,76 | 68,1 |
| ENSMUSG00000027533 | 1556,38122 | 2029,8 | 786,0 | 1,360619246 | 0,032399561 | Fabp5 | fatty acid binding protein 5, epidermal [Source:MGI Symbol;Acc:MGI:101790] | 91,02 | 48,7 |
| ENSMUSG00000026568 | 1400,223172 | 1961,9 | 773,2 | 1,365040645 | 0,04419823 | Mpc2 | mitochondrial pyruvate carrier 2 [Source:MGI Symbol;Acc:MGI:1917706] | 99,77 | 73,6 |
| ENSMUSG00000030057 | 5128,913023 | 6781,9 | 2662,1 | 1,384311052 | 0,012526629 | Cnbp | cellular nucleic acid binding protein [Source:MGI Symbol;Acc:MGI:88431] | 96,35 | 71,5 |
| ENSMUSG00000035242 | 1226,91357 | 1721,7 | 665,1 | 1,418093689 | 0,043293286 | Oaz1 | ornithine decarboxylase antizyme 1 [Source:MGI Symbol;Acc:MGI:109433] | 72,03 | 58,2 |
| ENSMUSG00000005505 | 433,2811363 | 539,5 | 206,2 | 1,423373462 | 0,023822826 | Kbtbd4 | kelch repeat and BTB (POZ) domain containing 4 [Source:MGI Symbol;Acc:MGI:1914386] | 99,63 | 8,8 |
| ENSMUSG00000029624 | 213,2644256 | 337,8 | 129,7 | 1,471968283 | 0,048859254 | Ptcd1 | pentatricopeptide repeat domain 1 [Source:MGI Symbol;Acc:MGI:1919049] | 90,78 | 8,1 |
| ENSMUSG00000020153 | 1894,682385 | 2725,9 | 1043,4 | 1,479969006 | 0,02491849 | Ndufs7 | NADH dehydrogenase (ubiquinone) Fe-S protein 7 [Source:MGI Symbol;Acc:MGI:1922656] | 100,00 | 77,9 |
| ENSMUSG00000053768 | 1343,433037 | 1761,2 | 672,7 | 1,484451715 | 0,049495484 | Chchd3 | coiled-coil-helix-coiled-coil-helix domain containing 3 [Source:MGI Symbol;Acc:MGI:1913325] | 91,00 | 65,5 |
| ENSMUSG00000016427 | 1216,394086 | 1566,6 | 546,8 | 1,491259401 | 0,035329739 | Ndufa1 | NADH dehydrogenase (ubiquinone) 1 alpha subcomplex, 1 [Source:MGI Symbol;Acc:MGI:1929511] | 100,00 | 96,4 |
| ENSMUSG00000003299 | 651,390733 | 877,1 | 355,5 | 1,492016094 | 0,049460093 | Mrpl4 | mitochondrial ribosomal protein L4 [Source:MGI Symbol;Acc:MGI:2137210] | 99,66 | 82,2 |
| ENSMUSG00000029213 | 1254,891883 | 1619,4 | 599,5 | 1,493828888 | 0,035667203 | Commd8 | COMM domain containing 8 [Source:MGI Symbol;Acc:MGI:1343485] | 99,74 | 60,9 |
| ENSMUSG00000014610 | 1377,465746 | 1750,1 | 678,4 | 1,494929256 | 0,040198466 | Etfb | electron transferring flavoprotein, beta polypeptide [Source:MGI Symbol;Acc:MGI:106098] | 98,10 | 93,6 |
| ENSMUSG00000018567 | 2797,049007 | 4116,9 | 1521,0 | 1,496759153 | 0,02894751 | Gabarap | gamma-aminobutyric acid receptor associated protein [Source:MGI Symbol;Acc:MGI:1861742] | 86,43 | 63,7 |
| ENSMUSG00000001175 | 10205,77799 | 15676,0 | 5669,9 | 1,497333076 | 0,02058664 | Calm1 | calmodulin 1 [Source:MGI Symbol;Acc:MGI:88251] | 95,53 | 78,5 |
| ENSMUSG00000015092 | 1483,773919 | 2160,2 | 836,7 | 1,503425603 | 0,033078575 | Edf1 | endothelial differentiation-related factor 1 [Source:MGI Symbol;Acc:MGI:1891227] | 100,00 | 97,4 |
| ENSMUSG00000019494 | 1089,297468 | 1449,2 | 546,9 | 1,506204476 | 0,03434132 | Cops6 | COP9 signalosome subunit 6 [Source:MGI Symbol;Acc:MGI:1349439] | 96,27 | 57,4 |
| ENSMUSG00000028843 | 556,9498559 | 734,5 | 273,2 | 1,510393074 | 0,033172832 | Sh3bgrl3 | SH3 domain binding glutamic acid-rich protein-like 3 [Source:MGI Symbol;Acc:MGI:1920973] | 99,60 | 90,8 |
| ENSMUSG00000046110 | 546,0272381 | 699,0 | 253,8 | 1,515361401 | 0,04454386 | Serinc4 | serine incorporator 4 [Source:MGI Symbol;Acc:MGI:2441842] | 44,17 | 33,1 |
| ENSMUSG00000020107 | 1016,63675 | 1488,8 | 585,6 | 1,516180585 | 0,042818634 | Anapc16 | anaphase promoting complex subunit 16 [Source:MGI Symbol;Acc:MGI:1289325] | 93,03 | 53,0 |
| ENSMUSG00000108996 | 313,5773891 | 427,9 | 163,9 | 1,523677726 | 0,045311419 | l7Rn6 | lethal, Chr 7, Rinchik 6 [Source:MGI Symbol;Acc:MGI:96738] | 100,00 | 90,8 |
| ENSMUSG00000021771 | 1487,728861 | 2251,0 | 846,1 | 1,530492292 | 0,032785753 | Vdac2 | voltage-dependent anion channel 2 [Source:MGI Symbol;Acc:MGI:106915] | 74,71 | 64,5 |
| ENSMUSG00000019923 | 2037,42836 | 3006,5 | 1072,7 | 1,535743182 | 0,012214782 | Zwint | ZW10 interactor [Source:MGI Symbol;Acc:MGI:1289227] | 70,55 | 46,6 |
| ENSMUSG00000006699 | 3239,697166 | 5008,4 | 1725,7 | 1,543907669 | 0,018427617 | Cdc42 | cell division cycle 42 [Source:MGI Symbol;Acc:MGI:106211] | 99,92 | 71,9 |
| ENSMUSG00000072772 | 1445,586207 | 2108,5 | 769,2 | 1,548932818 | 0,040451804 | Grcc10 | gene rich cluster, C10 gene [Source:MGI Symbol;Acc:MGI:1315201] | 91,85 | 63,9 |
| ENSMUSG00000017221 | 831,2640235 | 1237,3 | 451,4 | 1,551167653 | 0,035445955 | Psmd3 | proteasome (prosome, macropain) 26S subunit, non-ATPase, 3 [Source:MGI Symbol;Acc:MGI:98858] | 96,64 | 89,1 |
| ENSMUSG00000058267 | 1767,32929 | 2481,6 | 877,7 | 1,565661973 | 0,018346333 | Mrps14 | mitochondrial ribosomal protein S14 [Source:MGI Symbol;Acc:MGI:1928141] | 90,41 | 26,5 |
| ENSMUSG00000032470 | 1160,094472 | 1652,4 | 651,7 | 1,568896614 | 0,038333083 | Mras | muscle and microspikes RAS [Source:MGI Symbol;Acc:MGI:1100856] | 95,61 | 54,5 |
| ENSMUSG00000056201 | 4225,503326 | 6077,7 | 2100,2 | 1,572990399 | 0,011616912 | Cfl1 | cofilin 1, non-muscle [Source:MGI Symbol;Acc:MGI:101757] | 100,00 | 99,5 |
| ENSMUSG00000065990 | 913,5002483 | 1346,1 | 456,5 | 1,588464021 | 0,042232315 | Aurkaip1 | aurora kinase A interacting protein 1 [Source:MGI Symbol;Acc:MGI:1913327] | 93,06 | 53,9 |
| ENSMUSG00000031819 | 1725,682957 | 2664,0 | 918,1 | 1,589262123 | 0,031449128 | Emc8 | ER membrane protein complex subunit 8 [Source:MGI Symbol;Acc:MGI:1343095] | 99,96 | 45,6 |
| ENSMUSG00000062376 | 227,0766245 | 324,7 | 141,9 | 1,589904907 | 0,048349535 | Borcs7 | BLOC-1 related complex subunit 7 [Source:MGI Symbol;Acc:MGI:1913689] | 98,64 | 23,5 |
| ENSMUSG00000020349 | 2449,356034 | 3792,9 | 1279,7 | 1,590142862 | 0,016448139 | Ppp2ca | otein phosphatase 2 (formerly 2A), catalytic subunit, alpha isoform [Source:MGI Symbol;Acc:MGI:132115 | 58,58 | 37,9 |
| ENSMUSG00000062270 | 579,3338625 | 805,0 | 272,2 | 1,596270678 | 0,027395054 | Morf4l1 | mortality factor 4 like 1 [Source:MGI Symbol;Acc:MGI:1096551] | 60,24 | 35,9 |
| ENSMUSG00000104208 | 416,0432738 | 575,3 | 202,3 | 1,605226905 | 0,029520617 | Gm37401 | predicted gene, 37401 [Source:MGI Symbol;Acc:MGI:5610629] | 89,53 | 79,0 |
| ENSMUSG00000021024 | 1069,402194 | 1508,9 | 517,8 | 1,60913497 | 0,012234787 | Psma6 | proteasome (prosome, macropain) subunit, alpha type 6 [Source:MGI Symbol;Acc:MGI:1347006] | 89,26 | 84,0 |
| ENSMUSG00000062797 | 737,9690946 | 1075,0 | 380,6 | 1,617310689 | 0,01942823 | l7Rn6 | lethal, Chr 7, Rinchik 6 [Source:MGI Symbol;Acc:MGI:96738] | 81,91 | 44,9 |
| ENSMUSG00000096210 | 1685,333745 | 2419,1 | 958,6 | 1,622162446 | 0,048969442 | H1f0 | H1 histone family, member 0 [Source:MGI Symbol;Acc:MGI:95893] | 95,98 | 83,6 |
| ENSMUSG00000021417 | 1275,863532 | 1873,8 | 668,6 | 1,63798989 | 0,037737088 | Eci2 | enoyl-Coenzyme A delta isomerase 2 [Source:MGI Symbol;Acc:MGI:1346064] | 93,12 | 78,0 |
| ENSMUSG00000032314 | 1559,219018 | 2222,0 | 783,2 | 1,651294618 | 0,026876988 | Etfa | electron transferring flavoprotein, alpha polypeptide [Source:MGI Symbol;Acc:MGI:106092] | 98,37 | 87,3 |
| ENSMUSG00000007338 | 448,2562701 | 629,6 | 202,4 | 1,657096996 | 0,019933848 | Mrpl49 | mitochondrial ribosomal protein L49 [Source:MGI Symbol;Acc:MGI:108180] | 100,00 | 83,4 |
| ENSMUSG00000014956 | 2019,202506 | 2697,5 | 868,3 | 1,660803412 | 0,045604239 | Ppp1cb | protein phosphatase 1, catalytic subunit, beta isoform [Source:MGI Symbol;Acc:MGI:104871] | 68,20 | 42,8 |
| ENSMUSG00000021877 | 2483,707402 | 3635,1 | 1122,5 | 1,688334094 | 0,026989554 | Arf4 | ADP-ribosylation factor 4 [Source:MGI Symbol;Acc:MGI:99433] | 99,51 | 84,6 |
| ENSMUSG00000022023 | 335,7990111 | 448,3 | 136,8 | 1,698612265 | 0,021157074 | Wbp4 | WW domain binding protein 4 [Source:MGI Symbol;Acc:MGI:109568] | 98,07 | 25,0 |
| ENSMUSG00000040048 | 2137,904751 | 3248,3 | 1034,9 | 1,702389882 | 0,017208103 | Ndufb10 | NADH dehydrogenase (ubiquinone) 1 beta subcomplex, 10 [Source:MGI Symbol;Acc:MGI:1915592] | 98,76 | 87,5 |
| ENSMUSG00000029106 | 981,31401 | 1548,2 | 491,2 | 1,707967149 | 0,025833693 | Add1 | adducin 1 (alpha) [Source:MGI Symbol;Acc:MGI:87918] | 84,75 | 56,7 |
| ENSMUSG00000033059 | 3110,054318 | 4740,9 | 1500,3 | 1,714160302 | 0,025988092 | Pygb | brain glycogen phosphorylase [Source:MGI Symbol;Acc:MGI:97828] | 100,00 | 97,3 |
| ENSMUSG00000031847 | 3898,788056 | 6064,7 | 1874,0 | 1,714536245 | 0,035066767 | Rac1 | RAS-related C3 botulinum substrate 1 [Source:MGI Symbol;Acc:MGI:97845] | 76,23 | 53,6 |
| ENSMUSG00000015575 | 1233,883383 | 1900,5 | 591,8 | 1,715137181 | 0,034237557 | Atp6v0e | ATPase, H+ transporting, lysosomal V0 subunit E [Source:MGI Symbol;Acc:MGI:1328318] | 34,94 | 23,6 |
| ENSMUSG00000021039 | 560,4026852 | 646,8 | 195,9 | 1,717055624 | 0,024493876 | Snw1 | SNW domain containing 1 [Source:MGI Symbol;Acc:MGI:1913604] | 100,00 | 55,2 |
| ENSMUSG00000028745 | 1860,041406 | 2618,2 | 858,4 | 1,72263377 | 0,008385673 | Capzb | capping protein (actin filament) muscle Z-line, beta [Source:MGI Symbol;Acc:MGI:104652] | 79,73 | 72,5 |
| ENSMUSG00000039183 | 544,0310683 | 792,2 | 246,4 | 1,725802066 | 0,010921483 | Nubp2 | nucleotide binding protein 2 [Source:MGI Symbol;Acc:MGI:1347072] | 99,77 | 92,2 |
| ENSMUSG00000063694 | 818,6256562 | 1130,5 | 367,3 | 1,734909065 | 0,02753792 | Cycs | cytochrome c, somatic [Source:MGI Symbol;Acc:MGI:88578] | 74,03 | 12,5 |
| ENSMUSG00000024646 | 2228,291289 | 3590,4 | 1172,4 | 1,74365462 | 0,032220951 | Cyb5a | cytochrome b5 type A (microsomal) [Source:MGI Symbol;Acc:MGI:1926952] | 72,23 | 57,7 |
| ENSMUSG00000022477 | 2393,048746 | 3240,0 | 1001,4 | 1,750042761 | 0,016615332 | Aco2 | aconitase 2, mitochondrial [Source:MGI Symbol;Acc:MGI:87880] | 97,36 | 89,9 |
| ENSMUSG00000009863 | 2311,688329 | 3260,2 | 982,6 | 1,750697322 | 0,010559107 | Sdhb | succinate dehydrogenase complex, subunit B, iron sulfur (Ip) [Source:MGI Symbol;Acc:MGI:1914930] | 99,42 | 87,3 |
| ENSMUSG00000059734 | 1316,029065 | 1838,4 | 559,7 | 1,758414445 | 0,01404616 | Ndufs8 | NADH dehydrogenase (ubiquinone) Fe-S protein 8 [Source:MGI Symbol;Acc:MGI:2385079] | 100,00 | 83,0 |
| ENSMUSG00000061232 | 330,6851525 | 404,9 | 124,8 | 1,763490591 | 0,018209447 | H2-K1 | histocompatibility 2, K1, K region [Source:MGI Symbol;Acc:MGI:95904] | 97,82 | 38,4 |
| ENSMUSG00000007815 | 1527,43217 | 2229,5 | 683,7 | 1,766259746 | 0,011984883 | Rhoa | ras homolog gene family, member A [Source:MGI Symbol;Acc:MGI:1096342] | 89,57 | 68,6 |
| ENSMUSG00000031021 | 1417,297086 | 1984,0 | 604,9 | 1,768879675 | 0,043819032 | Tmem9b | TMEM9 domain family, member B [Source:MGI Symbol;Acc:MGI:1915254] | 99,87 | 72,3 |
| ENSMUSG00000032519 | 374,6063052 | 405,0 | 127,6 | 1,771648651 | 0,020897978 | Slc25a38 | solute carrier family 25, member 38 [Source:MGI Symbol;Acc:MGI:2384782] | 84,45 | 38,6 |
| ENSMUSG00000030647 | 3334,965764 | 4999,8 | 1664,9 | 1,781381849 | 0,026582956 | Ndufc2 | NADH dehydrogenase (ubiquinone) 1, subcomplex unknown, 2 [Source:MGI Symbol;Acc:MGI:1344370] | 93,63 | 86,1 |
| ENSMUSG00000026782 | 824,0320584 | 1030,6 | 305,3 | 1,782752341 | 0,009850501 | Abi2 | abl-interactor 2 [Source:MGI Symbol;Acc:MGI:106913] | 99,36 | 34,3 |
| ENSMUSG00000029559 | 334,6757925 | 465,4 | 146,4 | 1,786210078 | 0,012728172 | 2210016L21Rik | RIKEN cDNA 2210016L21 gene [Source:MGI Symbol;Acc:MGI:1919607] | 71,41 | 38,0 |
| ENSMUSG00000027406 | 1230,697481 | 1815,6 | 541,8 | 1,786623633 | 0,036470737 | Idh3b | isocitrate dehydrogenase 3 (NAD+) beta [Source:MGI Symbol;Acc:MGI:2158650] | 98,66 | 92,0 |
| ENSMUSG00000062691 | 515,852525 | 775,5 | 233,5 | 1,78859764 | 0,024124442 | Cebpzos | CCAAT/enhancer binding protein (C/EBP), zeta, opposite strand [Source:MGI Symbol;Acc:MGI:1915804] | 99,20 | 54,6 |
| ENSMUSG00000022476 | 1439,630778 | 2319,6 | 676,8 | 1,792890454 | 0,008890511 | Polr3h | polymerase (RNA) III (DNA directed) polypeptide H [Source:MGI Symbol;Acc:MGI:1926179] | 100,00 | 77,4 |
| ENSMUSG00000039206 | 1025,17827 | 1682,0 | 487,7 | 1,795300187 | 0,02076277 | Daglb | diacylglycerol lipase, beta [Source:MGI Symbol;Acc:MGI:2442032] | 79,67 | 13,3 |
| ENSMUSG00000021520 | 2041,329657 | 2911,2 | 933,2 | 1,805019809 | 0,044783053 | Uqcrb | ubiquinol-cytochrome c reductase binding protein [Source:MGI Symbol;Acc:MGI:1914780] | 99,19 | 90,7 |
| ENSMUSG00000015837 | 1934,48964 | 3246,5 | 934,4 | 1,807555197 | 0,0291501 | Sqstm1 | sequestosome 1 [Source:MGI Symbol;Acc:MGI:107931] | 92,89 | 75,4 |
| ENSMUSG00000046432 | 786,057818 | 1117,8 | 337,9 | 1,814063785 | 0,00624442 | Ngfrap1 | nerve growth factor receptor (TNFRSF16) associated protein 1 [Source:MGI Symbol;Acc:MGI:1338016] | 91,71 | 78,9 |
| ENSMUSG00000022472 | 946,9749004 | 1395,0 | 415,3 | 1,815806113 | 0,023276764 | Desi1 | desumoylating isopeptidase 1 [Source:MGI Symbol;Acc:MGI:106313] | 99,95 | 74,8 |
| ENSMUSG00000020949 | 1264,009972 | 1939,9 | 621,9 | 1,816015528 | 0,015863155 | Fkbp3 | FK506 binding protein 3 [Source:MGI Symbol;Acc:MGI:1353460] | 100,00 | 99,1 |
| ENSMUSG00000025410 | 889,7881771 | 1092,8 | 327,8 | 1,818068269 | 0,009373845 | Dctn2 | dynactin 2 [Source:MGI Symbol;Acc:MGI:107733] | 99,46 | 65,9 |
| ENSMUSG00000029918 | 367,3767333 | 563,9 | 192,8 | 1,823727662 | 0,042278602 | Mrps33 | mitochondrial ribosomal protein S33 [Source:MGI Symbol;Acc:MGI:1338046] | 56,46 | 20,1 |
| ENSMUSG00000022540 | 457,5144483 | 643,4 | 213,2 | 1,825661497 | 0,037073718 | Rogdi | rogdi homolog (Drosophila) [Source:MGI Symbol;Acc:MGI:1913299] | 91,33 | 67,0 |
| ENSMUSG00000060743 | 620,5077361 | 966,1 | 269,8 | 1,82984422 | 0,01071547 | H3f3a | H3 histone, family 3A [Source:MGI Symbol;Acc:MGI:1097686] | 53,42 | 11,2 |
| ENSMUSG00000024875 | 471,9737984 | 610,7 | 195,3 | 1,832398626 | 0,035785983 | Yif1a | Yip1 interacting factor homolog A (S. cerevisiae) [Source:MGI Symbol;Acc:MGI:1915340] | 100,00 | 65,4 |
| ENSMUSG00000084950 | 280,3719106 | 433,3 | 157,7 | 1,832728696 | 0,0277816 | Gm5577 | predicted gene 5577 [Source:MGI Symbol;Acc:MGI:3648213] | 51,73 | 30,1 |
| ENSMUSG00000038008 | 640,8541555 | 983,5 | 294,7 | 1,842227032 | 0,004225179 | Hmgn2 | high mobility group nucleosomal binding domain 2 [Source:MGI Symbol;Acc:MGI:96136] | 46,05 | 33,5 |
| ENSMUSG00000036309 | 5370,101594 | 8601,9 | 2451,8 | 1,842311099 | 0,013575227 | Skp1a | S-phase kinase-associated protein 1A [Source:MGI Symbol;Acc:MGI:103575] | 93,96 | 85,5 |
| ENSMUSG00000020149 | 2176,358434 | 3214,8 | 910,7 | 1,8555957 | 0,02655426 | Rab1a | RAB1A, member RAS oncogene family [Source:MGI Symbol;Acc:MGI:97842] | 98,51 | 76,1 |
| ENSMUSG00000026223 | 1930,130961 | 2790,6 | 854,1 | 1,863351175 | 0,041869735 | Itm2c | integral membrane protein 2C [Source:MGI Symbol;Acc:MGI:1927594] | 91,24 | 86,4 |
| ENSMUSG00000059323 | 1019,890899 | 1473,5 | 420,4 | 1,871603195 | 0,008244491 | Tonsl | tonsoku-like, DNA repair protein [Source:MGI Symbol;Acc:MGI:1919999] | 41,44 | 16,6 |
| ENSMUSG00000039105 | 782,1106069 | 1180,9 | 351,8 | 1,874845356 | 0,038989198 | Atp6v1g1 | ATPase, H+ transporting, lysosomal V1 subunit G1 [Source:MGI Symbol;Acc:MGI:1913540] | 88,60 | 71,9 |
| ENSMUSG00000022210 | 310,5321102 | 489,9 | 135,0 | 1,878250077 | 0,031022712 | Dhrs4 | dehydrogenase/reductase (SDR family) member 4 [Source:MGI Symbol;Acc:MGI:90169] | 98,87 | 69,6 |
| ENSMUSG00000079477 | 2551,660479 | 3970,3 | 1164,3 | 1,882824329 | 0,004893845 | Rab7 | RAB7, member RAS oncogene family [Source:MGI Symbol;Acc:MGI:105068] | 83,23 | 76,2 |
| ENSMUSG00000019804 | 3374,039649 | 5299,6 | 1889,2 | 1,884236659 | 0,045031522 | Snx3 | sorting nexin 3 [Source:MGI Symbol;Acc:MGI:1860188] | 99,53 | 88,1 |
| ENSMUSG00000028691 | 5370,474218 | 8949,8 | 2330,1 | 1,890915356 | 0,009128616 | Prdx1 | peroxiredoxin 1 [Source:MGI Symbol;Acc:MGI:99523] | 94,53 | 45,4 |
| ENSMUSG00000023904 | 748,862289 | 1004,4 | 394,1 | 1,909496949 | 0,041609376 | Hcfc1r1 | host cell factor C1 regulator 1 (XPO1-dependent) [Source:MGI Symbol;Acc:MGI:2663619] | 100,00 | 74,2 |
| ENSMUSG00000017631 | 1724,899954 | 2803,6 | 792,9 | 1,913469508 | 0,032006231 | Abr | active BCR-related gene [Source:MGI Symbol;Acc:MGI:107771] | 78,13 | 51,3 |
| ENSMUSG00000057342 | 487,1653518 | 723,3 | 207,2 | 1,918092421 | 0,039812398 | Sphk2 | sphingosine kinase 2 [Source:MGI Symbol;Acc:MGI:1861380] | 93,05 | 27,0 |
| ENSMUSG00000012126 | 273,1724985 | 340,9 | 97,2 | 1,918838447 | 0,011280576 | Ubxn11 | UBX domain protein 11 [Source:MGI Symbol;Acc:MGI:1914836] | 70,07 | 8,8 |
| ENSMUSG00000087165 | 136,5171366 | 174,2 | 54,0 | 1,919015145 | 0,04724066 | 2010001A14Rik | RIKEN cDNA 2010001A14 gene [Source:MGI Symbol;Acc:MGI:1923766] | #DIV/0! | #DIV/0! |
| ENSMUSG00000030161 | 1781,509257 | 2811,1 | 835,5 | 1,91998543 | 0,037496059 | Gabarapl1 | mma-aminobutyric acid (GABA) A receptor-associated protein-like 1 [Source:MGI Symbol;Acc:MGI:19149 | 98,97 | 90,6 |
| ENSMUSG00000018965 | 820,592509 | 1141,5 | 331,0 | 1,920194769 | 0,008430306 | Ywhah | onooxygenase/tryptophan 5-monooxygenase activation protein, eta polypeptide [Source:MGI Symbol;Acc:I | 77,72 | 63,5 |
| ENSMUSG00000021127 | 287,8428393 | 414,9 | 141,3 | 1,923411669 | 0,030338824 | Zfp36l1 | zinc finger protein 36, C3H type-like 1 [Source:MGI Symbol;Acc:MGI:107946] | 97,07 | 31,9 |
| ENSMUSG00000052738 | 1952,195315 | 3010,1 | 854,5 | 1,930174958 | 0,019361857 | Suclg1 | succinate-CoA ligase, GDP-forming, alpha subunit [Source:MGI Symbol;Acc:MGI:1927234] | 89,20 | 80,1 |
| ENSMUSG00000022884 | 918,567066 | 1099,6 | 336,3 | 1,935333157 | 0,029323948 | Eif4a2 | eukaryotic translation initiation factor 4A2 [Source:MGI Symbol;Acc:MGI:106906] | 77,36 | 29,2 |
| ENSMUSG00000031762 | 7759,039747 | 14105,0 | 4348,8 | 1,935607756 | 0,028442082 | Mt2 | metallothionein 2 [Source:MGI Symbol;Acc:MGI:97172] | 100,00 | 89,0 |
| ENSMUSG00000029223 | 204,3742354 | 273,1 | 71,7 | 1,935894291 | 0,042931635 | Uchl1 | ubiquitin carboxy-terminal hydrolase L1 [Source:MGI Symbol;Acc:MGI:103149] | 90,00 | 61,7 |
| ENSMUSG00000018287 | 512,1549412 | 764,1 | 228,5 | 1,938276547 | 0,01416993 | Spag7 | sperm associated antigen 7 [Source:MGI Symbol;Acc:MGI:107380] | 98,62 | 79,6 |
| ENSMUSG00000101503 | 316,8212895 | 474,5 | 134,9 | 1,94001277 | 0,031573046 | Gm28364 | predicted gene 28364 [Source:MGI Symbol;Acc:MGI:5579070] | 40,42 | 50,7 |
| ENSMUSG00000024091 | 1811,973348 | 2899,8 | 751,0 | 1,944708908 | 0,011034089 | Vapa | vesicle-associated membrane protein, associated protein A [Source:MGI Symbol;Acc:MGI:1353561] | 100,00 | 92,2 |
| ENSMUSG00000042613 | 550,3691547 | 862,0 | 222,0 | 1,957736177 | 0,049958565 | Pbxip1 | pre B cell leukemia transcription factor interacting protein 1 [Source:MGI Symbol;Acc:MGI:2441670] | 99,89 | 39,6 |
| ENSMUSG00000058076 | 2257,431826 | 3494,8 | 973,0 | 1,978017414 | 0,024107075 | Sdhc | inate dehydrogenase complex, subunit C, integral membrane protein [Source:MGI Symbol;Acc:MGI:1913 | 42,93 | 36,2 |
| ENSMUSG00000035133 | 2617,959338 | 3949,3 | 1097,4 | 1,978087273 | 0,043707608 | Arhgap5 | Rho GTPase activating protein 5 [Source:MGI Symbol;Acc:MGI:1332637] | 100,00 | 74,7 |
| ENSMUSG00000053453 | 566,9933803 | 838,6 | 238,6 | 1,978428361 | 0,016274719 | Thoc7 | THO complex 7 homolog (Drosophila) [Source:MGI Symbol;Acc:MGI:1913481] | 100,00 | 74,6 |
| ENSMUSG00000070493 | 2358,197659 | 3578,3 | 968,6 | 1,98810357 | 0,009471772 | Chchd2 | coiled-coil-helix-coiled-coil-helix domain containing 2 [Source:MGI Symbol;Acc:MGI:1261428] | 60,62 | 51,1 |
| ENSMUSG00000037361 | 973,4242926 | 1448,2 | 381,2 | 2,000299186 | 0,005071598 | Sf3b6 | splicing factor 3B, subunit 6 [Source:MGI Symbol;Acc:MGI:1913305] | 99,76 | 64,2 |
| ENSMUSG00000035770 | 551,5439822 | 608,9 | 165,9 | 2,004989817 | 0,032713879 | Dync1li2 | dynein, cytoplasmic 1 light intermediate chain 2 [Source:MGI Symbol;Acc:MGI:107738] | 97,68 | 31,8 |
| ENSMUSG00000002129 | 349,6511744 | 406,7 | 133,0 | 2,019412443 | 0,033928676 | Sf3a1 | splicing factor 3a, subunit 1 [Source:MGI Symbol;Acc:MGI:1914715] | 99,12 | 21,2 |
| ENSMUSG00000035559 | 522,1958701 | 728,7 | 179,6 | 2,021587406 | 0,013631482 | Mpv17l2 | MPV17 mitochondrial membrane protein-like 2 [Source:MGI Symbol;Acc:MGI:2681846] | 100,00 | 93,0 |
| ENSMUSG00000022680 | 997,0763851 | 1240,3 | 450,0 | 2,029272382 | 0,04176854 | Pdxdc1 | pyridoxal-dependent decarboxylase domain containing 1 [Source:MGI Symbol;Acc:MGI:1920909] | 74,52 | 29,3 |
| ENSMUSG00000027199 | 1102,213686 | 1518,1 | 384,6 | 2,029583876 | 0,009341226 | Gatm | lycine amidinotransferase (L-arginine:glycine amidinotransferase) [Source:MGI Symbol;Acc:MGI:1914342 | 99,80 | 87,0 |
| ENSMUSG00000020923 | 360,9146729 | 518,2 | 167,8 | 2,033148227 | 0,02958134 | Ubtf | upstream binding transcription factor, RNA polymerase I [Source:MGI Symbol;Acc:MGI:98512] | 61,76 | 18,9 |
| ENSMUSG00000019689 | 943,0729735 | 1388,5 | 435,7 | 2,039557532 | 0,025713947 | 1110001J03Rik | RIKEN cDNA 1110001J03 gene [Source:MGI Symbol;Acc:MGI:1913367] | 100,00 | 94,4 |
| ENSMUSG00000029446 | 950,342606 | 1402,7 | 434,0 | 2,05864391 | 0,014153278 | Psph | phosphoserine phosphatase [Source:MGI Symbol;Acc:MGI:97788] | 96,60 | 66,1 |
| ENSMUSG00000059447 | 898,8476499 | 1393,1 | 338,6 | 2,060632758 | 0,041531226 | Hadhb | ase/3-ketoacyl-Coenzyme A thiolase/enoyl-Coenzyme A hydratase (trifunctional protein), beta subunit [So | 71,71 | 60,1 |
| ENSMUSG00000024654 | 5574,076828 | 8843,8 | 2613,1 | 2,065770252 | 0,031924952 | Asrgl1 | asparaginase like 1 [Source:MGI Symbol;Acc:MGI:1913764] | 99,91 | 81,3 |
| ENSMUSG00000048644 | 347,2326346 | 514,3 | 191,7 | 2,07021208 | 0,041767397 | Ctxn1 | cortexin 1 [Source:MGI Symbol;Acc:MGI:88566] | 97,47 | 72,4 |
| ENSMUSG00000026032 | 1083,248404 | 1636,9 | 441,1 | 2,083151555 | 0,005912613 | Ndufb3 | NADH dehydrogenase (ubiquinone) 1 beta subcomplex 3 [Source:MGI Symbol;Acc:MGI:1913745] | 99,48 | 65,7 |
| ENSMUSG00000077990 | 276,6858173 | 485,7 | 143,5 | 2,084841496 | 0,04557986 | Gm23134 | predicted gene, 23134 [Source:MGI Symbol;Acc:MGI:5452911] | #DIV/0! | #DIV/0! |
| ENSMUSG00000038387 | 419,0321684 | 700,7 | 200,7 | 2,087108059 | 0,044827322 | Rras | Harvey rat sarcoma oncogene, subgroup R [Source:MGI Symbol;Acc:MGI:98179] | 99,86 | 86,1 |
| ENSMUSG00000033174 | 2261,334726 | 3294,5 | 821,7 | 2,087457622 | 0,022729402 | Mgll | monoglyceride lipase [Source:MGI Symbol;Acc:MGI:1346042] | 88,24 | 48,8 |
| ENSMUSG00000026895 | 1575,596634 | 2427,0 | 605,8 | 2,094147255 | 0,008019388 | Ndufa8 | NADH dehydrogenase (ubiquinone) 1 alpha subcomplex, 8 [Source:MGI Symbol;Acc:MGI:1915625] | 99,54 | 89,4 |
| ENSMUSG00000027602 | 1044,470816 | 1477,6 | 476,5 | 2,097219576 | 0,039651736 | Map1lc3a | microtubule-associated protein 1 light chain 3 alpha [Source:MGI Symbol;Acc:MGI:1915661] | 98,86 | 85,0 |

| | | | | | | | | | |
|---|---|---|---|---|---|---|---|---|---|
| ENSMUSG00000024423 | 664,8487652 | 831,8 | 319,0 | 2,103693997 | 0,0282791 | Impact | impact, RWD domain protein [Source:MGI Symbol;Acc:MGI:1098233] | 99,98 | 43,9 |
| ENSMUSG00000000346 | 6265,36545 | 10160,4 | 2772,7 | 2,114820033 | 0,020067293 | Dazap2 | DAZ associated protein 2 [Source:MGI Symbol;Acc:MGI:1344344] | 100,00 | 99,8 |
| ENSMUSG00000000159 | 258,1003633 | 298,5 | 125,1 | 2,128084169 | 0,048578581 | Igsf5 | immunoglobulin superfamily, member 5 [Source:MGI Symbol;Acc:MGI:1919308] | 57,42 | 9,2 |
| ENSMUSG00000039533 | 7834,091637 | 12560,3 | 3170,0 | 2,130947325 | 0,025444664 | Mmd2 | monocyte to macrophage differentiation-associated 2 [Source:MGI Symbol;Acc:MGI:1922354] | 99,95 | 65,7 |
| ENSMUSG00000019428 | 1202,659648 | 1689,2 | 417,0 | 2,1358753 | 0,020672134 | Fkbp8 | FK506 binding protein 8 [Source:MGI Symbol;Acc:MGI:1341070] | 87,80 | 61,9 |
| ENSMUSG00000075706 | 2789,889853 | 4102,1 | 1129,6 | 2,142257017 | 0,024379257 | Gpx4 | glutathione peroxidase 4 [Source:MGI Symbol;Acc:MGI:104767] | 89,05 | 50,5 |
| ENSMUSG00000014303 | 49,93656692 | 62,9 | 19,6 | 2,143088115 | 0,045695789 | Glis2 | GLIS family zinc finger 2 [Source:MGI Symbol;Acc:MGI:1932535] | 81,15 | 3,6 |
| ENSMUSG00000000552 | 173,281182 | 252,8 | 68,0 | 2,158023177 | 0,024653333 | Zfp385a | zinc finger protein 385A [Source:MGI Symbol;Acc:MGI:1352495] | 94,70 | 37,3 |
| ENSMUSG00000027835 | 302,4268936 | 473,0 | 112,7 | 2,158948212 | 0,030939401 | Pdcd10 | programmed cell death 10 [Source:MGI Symbol;Acc:MGI:1928396] | 82,74 | 23,6 |
| ENSMUSG00000031916 | 668,286609 | 874,8 | 241,9 | 2,160611846 | 0,016739916 | Cog8 | component of oligomeric golgi complex 8 [Source:MGI Symbol;Acc:MGI:2142885] | 85,39 | 27,5 |
| ENSMUSG00000006299 | 1719,96549 | 2712,9 | 619,4 | 2,162223385 | 0,009115328 | Aamp | angio-associated migratory protein [Source:MGI Symbol;Acc:MGI:107809] | 88,33 | 84,9 |
| ENSMUSG00000028517 | 5294,244796 | 8309,9 | 1852,3 | 2,163435082 | 0,038475475 | Plpp3 | phospholipid phosphatase 3 [Source:MGI Symbol;Acc:MGI:1915166] | 100,00 | 96,4 |
| ENSMUSG00000004071 | 2050,891936 | 3245,1 | 730,9 | 2,164359026 | 0,000211054 | Cdip1 | cell death inducing Trp53 target 1 [Source:MGI Symbol;Acc:MGI:1913876] | 97,33 | 65,7 |
| ENSMUSG00000077450 | 921,3516496 | 1435,8 | 351,6 | 2,167389697 | 0,003898346 | Rab11b | RAB11B, member RAS oncogene family [Source:MGI Symbol;Acc:MGI:99425] | 88,21 | 23,0 |
| ENSMUSG00000055912 | 61,41438738 | 85,3 | 18,8 | 2,17355235 | 0,017459821 | Tmem150a | transmembrane protein 150A [Source:MGI Symbol;Acc:MGI:2385244] | 88,10 | 2,6 |
| ENSMUSG00000009585 | 72,55334509 | 103,2 | 23,5 | 2,173646604 | 0,017011651 | Apobec3 | apolipoprotein B mRNA editing enzyme, catalytic polypeptide 3 [Source:MGI Symbol;Acc:MGI:1933111] | 65,76 | 2,1 |
| ENSMUSG00000052456 | 376,9891458 | 555,5 | 132,8 | 2,175549671 | 0,00331406 | Asna1 | arsA arsenite transporter, ATP-binding, homolog 1 (bacterial) [Source:MGI Symbol;Acc:MGI:1928379] | 99,15 | 76,3 |
| ENSMUSG00000086938 | 121,5658047 | 191,4 | 47,8 | 2,177613127 | 0,009223407 | 4930481A15Rik | RIKEN cDNA 4930481A15 gene [Source:MGI Symbol;Acc:MGI:1922181] | 80,51 | 12,5 |
| ENSMUSG00000106967 | 65,73865771 | 93,6 | 23,7 | 2,177874044 | 0,04176854 | Gm42477 | predicted gene 42477 [Source:MGI Symbol;Acc:MGI:5662614] | 51,60 | 30,6 |
| ENSMUSG00000004558 | 24355,62011 | 37500,0 | 10834,1 | 2,181282147 | 0,042457815 | Ndrg2 | N-myc downstream regulated gene 2 [Source:MGI Symbol;Acc:MGI:1352498] | 100,00 | 95,8 |
| ENSMUSG00000024826 | 442,1781409 | 541,3 | 201,7 | 2,185263437 | 0,038936575 | Dpf2 | D4, zinc and double PHD fingers family 2 [Source:MGI Symbol;Acc:MGI:109529] | 81,65 | 37,5 |
| ENSMUSG00000031517 | 10067,10059 | 15521,9 | 3318,6 | 2,185972251 | 0,038235961 | Gpm6a | glycoprotein m6a [Source:MGI Symbol;Acc:MGI:107671] | 99,47 | 83,6 |
| ENSMUSG00000001700 | 1565,983308 | 2022,6 | 691,6 | 2,200625467 | 0,034742336 | Gramd3 | GRAM domain containing 3 [Source:MGI Symbol;Acc:MGI:1914815] | 99,99 | 70,1 |
| ENSMUSG00000076432 | 837,7218682 | 1359,0 | 306,8 | 2,201301859 | 0,014040727 | Ywhaq | nooxygenase/tryptophan 5-monooxygenase activation protein, theta polypeptide [Source:MGI Symbol;Acc | 63,22 | 35,7 |
| ENSMUSG00000022678 | 1718,410152 | 3006,1 | 727,3 | 2,202016842 | 0,007525357 | Nde1 | nuclear distribution gene E homolog 1 (A nidulans) [Source:MGI Symbol;Acc:MGI:1914453] | 100,00 | 85,4 |
| ENSMUSG00000036565 | 453,2412347 | 668,1 | 155,0 | 2,210138495 | 0,039710916 | Ttyh3 | tweety homolog 3 (Drosophila) [Source:MGI Symbol;Acc:MGI:1925589] | 97,39 | 38,8 |
| ENSMUSG00000030120 | 5066,190039 | 7983,9 | 2097,3 | 2,213138383 | 0,011883954 | Mlf2 | myeloid leukemia factor 2 [Source:MGI Symbol;Acc:MGI:1353554] | 98,76 | 88,4 |
| ENSMUSG00000091512 | 428,8963462 | 636,2 | 185,5 | 2,21681615 | 0,036439871 | Lamtor3 | late endosomal/lysosomal adaptor, MAPK and MTOR activator 3 [Source:MGI Symbol;Acc:MGI:1929467] | 97,40 | 52,4 |
| ENSMUSG00000002524 | 787,3787945 | 1164,1 | 303,8 | 2,221990018 | 0,009219475 | Puf60 | poly-U binding splicing factor 60 [Source:MGI Symbol;Acc:MGI:1915209] | 99,74 | 71,7 |
| ENSMUSG00000019433 | 470,6447524 | 725,8 | 180,1 | 2,224493651 | 0,025104874 | Gipc1 | GIPC PDZ domain containing family, member 1 [Source:MGI Symbol;Acc:MGI:1926252] | 100,00 | 83,3 |
| ENSMUSG00000043445 | 1063,991897 | 1573,1 | 421,1 | 2,228539158 | 0,003934342 | Pgp | phosphoglycolate phosphatase [Source:MGI Symbol;Acc:MGI:1914328] | 100,00 | 93,1 |
| ENSMUSG00000079478 | 512,834649 | 613,9 | 150,0 | 2,232782898 | 0,009806038 | Sssca1 | Sjogren's syndrome/scleroderma autoantigen 1 homolog (human) [Source:MGI Symbol;Acc:MGI:1913482 | 92,57 | 76,8 |
| ENSMUSG00000079108 | 71,74870845 | 91,7 | 21,4 | 2,244618728 | 0,041339457 | Srp54b | signal recognition particle 54B [Source:MGI Symbol;Acc:MGI:3714357] | 11,22 | 1,6 |
| ENSMUSG00000035198 | 302,3791999 | 363,1 | 131,4 | 2,246358942 | 0,049215399 | Tubg1 | tubulin, gamma 1 [Source:MGI Symbol;Acc:MGI:101834] | 95,99 | 53,3 |
| ENSMUSG00000036026 | 223,1351592 | 245,4 | 68,8 | 2,247821928 | 0,034264344 | Tmem63b | transmembrane protein 63b [Source:MGI Symbol;Acc:MGI:2387609] | 84,52 | 14,0 |
| ENSMUSG00000041570 | 278,7195371 | 445,8 | 182,2 | 2,260018823 | 0,047316917 | Camsap2 | almodulin regulated spectrin-associated protein family, member 2 [Source:MGI Symbol;Acc:MGI:1922434 | 97,05 | 23,3 |
| ENSMUSG00000027805 | 2694,994484 | 3989,0 | 1130,1 | 2,262200441 | 0,022794374 | Pfn2 | profilin 2 [Source:MGI Symbol;Acc:MGI:97550] | 71,96 | 49,6 |
| ENSMUSG00000039197 | 2391,377095 | 4193,7 | 879,4 | 2,265107272 | 0,028021437 | Adk | adenosine kinase [Source:MGI Symbol;Acc:MGI:87930] | 99,68 | 95,8 |
| ENSMUSG00000004849 | 546,1437265 | 842,5 | 239,8 | 2,268581824 | 0,009302039 | Ap1s1 | adaptor protein complex AP-1, sigma 1 [Source:MGI Symbol;Acc:MGI:1098244] | 73,74 | 53,2 |
| ENSMUSG00000029030 | 3162,941696 | 4771,5 | 1449,4 | 2,269373971 | 0,013831112 | Tprgl | transformation related protein 63 regulated like [Source:MGI Symbol;Acc:MGI:1915058] | 100,00 | 99,8 |
| ENSMUSG00000002205 | 522,0306241 | 781,4 | 215,3 | 2,28667077 | 0,009976334 | Vrk3 | vaccinia related kinase 3 [Source:MGI Symbol;Acc:MGI:2182465] | 77,32 | 43,7 |
| ENSMUSG00000032388 | 582,5763096 | 809,8 | 167,0 | 2,289529332 | 0,008088217 | Spg21 | spastic paraplegia 21 homolog (human) [Source:MGI Symbol;Acc:MGI:106403] | 87,04 | 48,4 |
| ENSMUSG00000034343 | 544,5646541 | 674,0 | 144,5 | 2,290284923 | 0,000674007 | Ube2f | ubiquitin-conjugating enzyme E2F (putative) [Source:MGI Symbol;Acc:MGI:1915171] | 64,12 | 13,7 |
| ENSMUSG00000032051 | 284,5157124 | 462,4 | 96,6 | 2,294964673 | 0,013312483 | Fdx1 | ferredoxin 1 [Source:MGI Symbol;Acc:MGI:103224] | 100,00 | 58,8 |
| ENSMUSG00000036564 | 1703,765749 | 2659,6 | 561,0 | 2,301540314 | 0,037604751 | Ndrg4 | N-myc downstream regulated gene 4 [Source:MGI Symbol;Acc:MGI:2384590] | 86,34 | 67,0 |
| ENSMUSG00000005034 | 649,1593359 | 1026,6 | 374,8 | 2,304170887 | 0,01781242 | Prkacb | protein kinase, cAMP dependent, catalytic, beta [Source:MGI Symbol;Acc:MGI:97594] | 87,87 | 32,1 |
| ENSMUSG00000019210 | 1460,060984 | 2443,7 | 554,1 | 2,306610391 | 0,043042394 | Atp6v1e1 | ATPase, H+ transporting, lysosomal V1 subunit E1 [Source:MGI Symbol;Acc:MGI:894326] | 95,31 | 77,5 |
| ENSMUSG00000008348 | 697,4738932 | 1087,5 | 256,5 | 2,313645436 | 0,011568682 | Ubc | ubiquitin C [Source:MGI Symbol;Acc:MGI:98889] | 36,65 | 21,7 |
| ENSMUSG00000032615 | 420,9453983 | 605,7 | 207,3 | 2,315520156 | 0,035956226 | Nt5m | 5',3'-nucleotidase, mitochondrial [Source:MGI Symbol;Acc:MGI:1917127] | 100,00 | 52,1 |
| ENSMUSG00000041846 | 352,5240599 | 473,9 | 226,0 | 2,322683723 | 0,043929689 | Smek1 | SMEK homolog 1, suppressor of mek1 (Dictyostelium) [Source:MGI Symbol;Acc:MGI:1915984] | 98,53 | 36,5 |
| ENSMUSG00000041769 | 279,95048 | 392,7 | 89,4 | 2,324413944 | 0,020289918 | Ppp2r2d | protein phosphatase 2, regulatory subunit B, delta [Source:MGI Symbol;Acc:MGI:1289252] | 98,08 | 38,8 |
| ENSMUSG00000031812 | 4078,713088 | 6913,7 | 1436,0 | 2,358111527 | 0,004016285 | Map1lc3b | microtubule-associated protein 1 light chain 3 beta [Source:MGI Symbol;Acc:MGI:1914693] | 62,27 | 54,7 |
| ENSMUSG00000045438 | 255,4207087 | 375,8 | 86,0 | 2,35897683 | 0,006235071 | Cox19 | cytochrome c oxidase assembly protein 19 [Source:MGI Symbol;Acc:MGI:1915283] | 70,36 | 24,3 |
| ENSMUSG00000045092 | 6861,182325 | 10766,9 | 2331,6 | 2,36893765 | 0,02830088 | S1pr1 | sphingosine-1-phosphate receptor 1 [Source:MGI Symbol;Acc:MGI:1096355] | 99,84 | 95,0 |
| ENSMUSG00000032553 | 290,159152 | 279,0 | 71,2 | 2,371386443 | 0,032190185 | Srprb | signal recognition particle receptor, B subunit [Source:MGI Symbol;Acc:MGI:102964] | 98,88 | 23,5 |
| ENSMUSG00000024870 | 1448,181509 | 2402,2 | 548,7 | 2,376310822 | 0,012503927 | Rab1b | RAB1B, member RAS oncogene family [Source:MGI Symbol;Acc:MGI:1923558] | 100,00 | 94,2 |
| ENSMUSG00000022800 | 541,9182751 | 714,5 | 153,5 | 2,376581851 | 0,018401236 | Fyttd1 | forty-two-three domain containing 1 [Source:MGI Symbol;Acc:MGI:1917955] | 84,53 | 23,9 |
| ENSMUSG00000015478 | 1566,798617 | 2428,2 | 621,6 | 2,378579114 | 0,01670605 | Rnf5 | ring finger protein 5 [Source:MGI Symbol;Acc:MGI:1860076] | 97,71 | 84,1 |
| ENSMUSG00000078931 | 553,1619615 | 752,5 | 191,6 | 2,37991509 | 0,010159129 | Pdf | peptide deformylase (mitochondrial) [Source:MGI Symbol;Acc:MGI:1915273] | 100,00 | 49,3 |
| ENSMUSG00000036790 | 70,5844788 | 113,8 | 30,2 | 2,38367787 | 0,032019326 | Slitrk2 | SLIT and NTRK-like family, member 2 [Source:MGI Symbol;Acc:MGI:2679449] | 62,56 | 2,6 |
| ENSMUSG00000024414 | 488,0545719 | 673,9 | 150,8 | 2,383952208 | 0,006314703 | Mrpl27 | mitochondrial ribosomal protein L27 [Source:MGI Symbol;Acc:MGI:2137224] | 98,59 | 74,8 |
| ENSMUSG00000025153 | 534,1794263 | 778,9 | 148,0 | 2,387598616 | 0,00303887 | Fasn | fatty acid synthase [Source:MGI Symbol;Acc:MGI:95485] | 84,19 | 10,3 |
| ENSMUSG00000054364 | 686,0014878 | 1222,2 | 312,3 | 2,388504401 | 0,021531594 | Rhob | ras homolog gene family, member B [Source:MGI Symbol;Acc:MGI:107949] | 99,97 | 72,6 |
| ENSMUSG00000059409 | 539,0084069 | 729,1 | 174,5 | 2,398224107 | 0,009132349 | Ppp2r5d | protein phosphatase 2, regulatory subunit B', delta [Source:MGI Symbol;Acc:MGI:2388481] | 100,00 | 32,9 |
| ENSMUSG00000034066 | 108,5605814 | 168,8 | 36,3 | 2,398277757 | 0,006659046 | Farp2 | FERM, RhoGEF and pleckstrin domain protein 2 [Source:MGI Symbol;Acc:MGI:2385126] | 69,24 | 1,5 |
| ENSMUSG00000047187 | 2986,571069 | 4964,9 | 950,8 | 2,406560618 | 0,008355771 | Rab2a | RAB2A, member RAS oncogene family [Source:MGI Symbol;Acc:MGI:1928750] | 97,97 | 92,0 |
| ENSMUSG00000021076 | 708,7366497 | 1019,7 | 326,4 | 2,407450913 | 0,047498192 | Actr10 | ARP10 actin-related protein 10 [Source:MGI Symbol;Acc:MGI:1891654] | 93,32 | 38,8 |
| ENSMUSG00000024454 | 794,5743063 | 1190,8 | 346,3 | 2,415219963 | 0,009892318 | Hdac3 | histone deacetylase 3 [Source:MGI Symbol;Acc:MGI:1343091] | 99,00 | 67,4 |
| ENSMUSG00000028603 | 1832,008607 | 2754,7 | 795,8 | 2,4328615 | 0,039405259 | Scp2 | sterol carrier protein 2, liver [Source:MGI Symbol;Acc:MGI:98254] | 78,74 | 29,7 |
| ENSMUSG00000034341 | 1925,996557 | 2945,6 | 565,2 | 2,436461757 | 0,011411264 | Wbp2 | WW domain binding protein 2 [Source:MGI Symbol;Acc:MGI:104709] | 100,00 | 93,3 |
| ENSMUSG00000017390 | 27993,96595 | 45498,3 | 9857,7 | 2,442382009 | 0,039994072 | Aldoc | aldolase C, fructose-bisphosphate [Source:MGI Symbol;Acc:MGI:101863] | 100,00 | 82,3 |
| ENSMUSG00000016534 | 1071,441088 | 1494,1 | 258,7 | 2,443334079 | 0,003836524 | Lamp2 | lysosomal-associated membrane protein 2 [Source:MGI Symbol;Acc:MGI:96748] | 97,51 | 27,1 |
| ENSMUSG00000001891 | 3376,807116 | 5626,4 | 1082,7 | 2,463630929 | 0,00551827 | Ugp2 | UDP-glucose pyrophosphorylase 2 [Source:MGI Symbol;Acc:MGI:2183447] | 100,00 | 89,3 |
| ENSMUSG00000009291 | 1239,941908 | 1784,4 | 325,4 | 2,467305244 | 0,0047196 | Pttg1ip | pituitary tumor-transforming 1 interacting protein [Source:MGI Symbol;Acc:MGI:2652132] | 92,41 | 79,7 |
| ENSMUSG00000097675 | 312,5648649 | 535,6 | 111,8 | 2,478936215 | 0,009892318 | 1700101I11Rik | RIKEN cDNA 1700101I11 gene [Source:MGI Symbol;Acc:MGI:1915180] | 71,18 | 22,9 |
| ENSMUSG00000031924 | 2188,954268 | 2963,3 | 959,0 | 2,480193552 | 0,02185231 | Cyb5b | cytochrome b5 type B [Source:MGI Symbol;Acc:MGI:1913677] | 99,98 | 56,2 |
| ENSMUSG00000058135 | 15538,58151 | 27393,6 | 5406,5 | 2,481091227 | 0,014154423 | Gstm1 | glutathione S-transferase, mu 1 [Source:MGI Symbol;Acc:MGI:95860] | 100,00 | 86,7 |
| ENSMUSG00000092395 | 783,6529183 | 1229,1 | 300,9 | 2,481219493 | 0,019372396 | Gm20463 | predicted gene 20463 [Source:MGI Symbol;Acc:MGI:5141928] | 95,72 | 86,4 |
| ENSMUSG00000022658 | 2871,127137 | 4602,4 | 1041,8 | 2,486328788 | 0,022252444 | Tagln3 | transgelin 3 [Source:MGI Symbol;Acc:MGI:1926784] | 100,00 | 82,5 |
| ENSMUSG00000031748 | 1880,105392 | 3210,9 | 662,7 | 2,490622748 | 0,01949513 | Gnao1 | guanine nucleotide binding protein, alpha O [Source:MGI Symbol;Acc:MGI:95775] | 98,13 | 58,3 |
| ENSMUSG00000085793 | 977,1521822 | 1476,1 | 325,4 | 2,490694723 | 0,026653364 | Lin52 | lin-52 homolog (C. elegans) [Source:MGI Symbol;Acc:MGI:3045391] | 92,65 | 68,9 |
| ENSMUSG00000093006 | 111,6094179 | 185,2 | 36,8 | 2,494200497 | 0,006385371 | Gm24157 | predicted gene, 24157 [Source:MGI Symbol;Acc:MGI:5453934] | #DIV/0! | #DIV/0! |
| ENSMUSG00000026154 | 617,7917886 | 1014,5 | 265,8 | 2,4942935 | 0,038235961 | Sdhaf4 | succinate dehydrogenase complex assembly factor 4 [Source:MGI Symbol;Acc:MGI:1915252] | 99,66 | 87,8 |
| ENSMUSG00000097347 | 241,1597428 | 421,4 | 73,8 | 2,497891103 | 0,002312946 | Gm17275 | predicted gene, 17275 [Source:MGI Symbol;Acc:MGI:4936909] | #DIV/0! | #DIV/0! |
| ENSMUSG00000050565 | 559,7682589 | 835,6 | 202,0 | 2,499472278 | 0,006726105 | Tor1aip2 | torsin A interacting protein 2 [Source:MGI Symbol;Acc:MGI:3582695] | 85,30 | 13,2 |
| ENSMUSG00000041058 | 337,8283682 | 452,9 | 90,5 | 2,500075271 | 0,031656379 | Wwp1 | WW domain containing E3 ubiquitin protein ligase 1 [Source:MGI Symbol;Acc:MGI:1861728] | 98,96 | 9,9 |
| ENSMUSG00000022474 | 2290,444281 | 3622,5 | 643,3 | 2,501701865 | 0,001903062 | Pmm1 | phosphomannomutase 1 [Source:MGI Symbol;Acc:MGI:1353418] | 96,82 | 94,8 |
| ENSMUSG00000002808 | 1109,202615 | 1724,2 | 366,7 | 2,510182537 | 0,004023292 | Epdr1 | ependymin related protein 1 (zebrafish) [Source:MGI Symbol;Acc:MGI:2145369] | 99,61 | 60,1 |
| ENSMUSG00000041556 | 2597,615743 | 4362,4 | 923,1 | 2,510377292 | 0,027415843 | Fbxo2 | F-box protein 2 [Source:MGI Symbol;Acc:MGI:2446216] | 100,00 | 98,9 |
| ENSMUSG00000072612 | 68,36751632 | 109,4 | 20,1 | 2,5110845 | 0,011984883 | Gm10382 | predicted gene 10382 [Source:MGI Symbol;Acc:MGI:3647829] | 54,23 | 4,7 |
| ENSMUSG00000021607 | 500,8761745 | 699,9 | 170,4 | 2,520429145 | 0,017920081 | Mrpl36 | mitochondrial ribosomal protein L36 [Source:MGI Symbol;Acc:MGI:2137228] | 100,00 | 64,0 |
| ENSMUSG00000026701 | 25807,46035 | 44118,1 | 8096,1 | 2,52666555 | 0,005912613 | Prdx6 | peroxiredoxin 6 [Source:MGI Symbol;Acc:MGI:894320] | 99,72 | 63,2 |
| ENSMUSG00000005103 | 963,5555563 | 1381,2 | 338,4 | 2,527830655 | 0,031192751 | Wdr1 | WD repeat domain 1 [Source:MGI Symbol;Acc:MGI:1337100] | 99,40 | 50,3 |
| ENSMUSG00000102495 | 70,15144988 | 111,7 | 21,9 | 2,530922422 | 0,036691306 | Gm26524 | predicted gene, 26524 [Source:MGI Symbol;Acc:MGI:5477018] | 28,46 | 14,0 |
| ENSMUSG00000003269 | 510,3468141 | 776,6 | 195,6 | 2,533360847 | 0,012958678 | Cyth2 | cytohesin 2 [Source:MGI Symbol;Acc:MGI:1334255] | 78,92 | 40,2 |
| ENSMUSG00000027273 | 2212,826448 | 3297,9 | 637,5 | 2,542920331 | 0,001885975 | Snap25 | synaptosomal-associated protein 25 [Source:MGI Symbol;Acc:MGI:98331] | 99,91 | 90,0 |
| ENSMUSG00000041408 | 377,8709587 | 490,2 | 313,1 | 2,551438418 | 0,049369644 | Wapl | WAPL cohesin release factor [Source:MGI Symbol;Acc:MGI:2675859] | 99,57 | 12,8 |
| ENSMUSG00000097417 | 167,0771903 | 240,6 | 60,5 | 2,551598197 | 0,016066419 | Gm26669 | predicted gene, 26669 [Source:MGI Symbol;Acc:MGI:5477163] | #DIV/0! | #DIV/0! |
| ENSMUSG00000022214 | 1110,993832 | 1667,5 | 329,0 | 2,552926558 | 0,00303887 | Dcaf11 | DDB1 and CUL4 associated factor 11 [Source:MGI Symbol;Acc:MGI:90168] | 99,77 | 65,7 |
| ENSMUSG00000049422 | 3675,660824 | 5927,8 | 1328,1 | 2,559704302 | 0,034635525 | Chchd10 | coiled-coil-helix-coiled-coil-helix domain containing 10 [Source:MGI Symbol;Acc:MGI:2143558] | 85,59 | 68,5 |
| ENSMUSG00000024614 | 269,9832691 | 268,9 | 44,5 | 2,564175329 | 0,016496871 | Tmx3 | thioredoxin-related transmembrane protein 3 [Source:MGI Symbol;Acc:MGI:2442418] | 96,52 | 10,7 |
| ENSMUSG00000030246 | 6025,373907 | 10219,7 | 1961,4 | 2,566471378 | 0,018209447 | Ldhb | lactate dehydrogenase B [Source:MGI Symbol;Acc:MGI:96763] | 89,81 | 80,7 |
| ENSMUSG00000031840 | 402,1136673 | 637,9 | 105,9 | 2,567483766 | 0,000212251 | Rab3a | RAB3A, member RAS oncogene family [Source:MGI Symbol;Acc:MGI:97843] | 83,20 | 57,9 |
| ENSMUSG00000028191 | 384,1904572 | 512,8 | 175,0 | 2,582263178 | 0,043968139 | Bcl10 | B cell leukemia/lymphoma 10 [Source:MGI Symbol;Acc:MGI:1337994] | 99,06 | 41,5 |
| ENSMUSG00000027562 | 5040,418104 | 7637,9 | 1579,5 | 2,587544728 | 0,033825651 | Car2 | carbonic anhydrase 2 [Source:MGI Symbol;Acc:MGI:88269] | 100,00 | 96,6 |
| ENSMUSG00000030605 | 2679,837537 | 3595,0 | 906,0 | 2,587940381 | 0,039595924 | Mfge8 | milk fat globule-EGF factor 8 protein [Source:MGI Symbol;Acc:MGI:102768] | 97,62 | 81,3 |
| ENSMUSG00000000399 | 1097,394795 | 1541,2 | 539,7 | 2,591507616 | 0,033632765 | Ndufa9 | NADH dehydrogenase (ubiquinone) 1 alpha subcomplex, 9 [Source:MGI Symbol;Acc:MGI:1913358] | 89,25 | 56,8 |
| ENSMUSG00000041716 | 216,8198054 | 359,1 | 70,2 | 2,610035429 | 0,010931263 | Gm20604 | predicted gene 20604 [Source:MGI Symbol;Acc:MGI:5305180] | 91,53 | 10,4 |
| ENSMUSG00000005312 | 1109,756903 | 1498,5 | 413,8 | 2,617859019 | 0,009401723 | Ubqln1 | ubiquilin 1 [Source:MGI Symbol;Acc:MGI:1860276] | 99,90 | 49,7 |
| ENSMUSG00000064254 | 213,0298991 | 286,9 | 85,9 | 2,618037709 | 0,035956226 | Ethe1 | ethylmalonic encephalopathy 1 [Source:MGI Symbol;Acc:MGI:1913321] | 70,89 | 36,4 |
| ENSMUSG00000059824 | 884,9900427 | 1378,4 | 280,0 | 2,618336911 | 0,010339674 | Dbp | D site albumin promoter binding protein [Source:MGI Symbol;Acc:MGI:94866] | 99,46 | 73,6 |
| ENSMUSG00000025351 | 2573,716586 | 3643,7 | 807,8 | 2,620052873 | 0,046266669 | Cd63 | CD63 antigen [Source:MGI Symbol;Acc:MGI:99529] | 99,81 | 84,7 |
| ENSMUSG00000033768 | 188,805667 | 273,0 | 50,5 | 2,627997514 | 0,026614038 | Nrxn2 | neurexin II [Source:MGI Symbol;Acc:MGI:1096362] | 75,11 | 9,1 |
| ENSMUSG00000050157 | 186,8281102 | 322,3 | 61,5 | 2,63673946 | 0,036115609 | Gm867 | predicted gene 867 [Source:MGI Symbol;Acc:MGI:2685713] | 46,76 | 2,0 |
| ENSMUSG00000030652 | 510,4126897 | 769,4 | 211,9 | 2,641599137 | 0,023617627 | Coq7 | demethyl-Q 7 [Source:MGI Symbol;Acc:MGI:107207] | 63,26 | 37,5 |
| ENSMUSG00000025766 | 523,609447 | 616,3 | 160,8 | 2,65247345 | 0,026344109 | D3Ertd751e | DNA segment, Chr 3, ERATO Doi 751, expressed [Source:MGI Symbol;Acc:MGI:1289213] | 82,24 | 26,1 |
| ENSMUSG00000001270 | 15673,42835 | 25914,7 | 4548,4 | 2,653357783 | 0,006924761 | Ckb | creatine kinase, brain [Source:MGI Symbol;Acc:MGI:88407] | 98,92 | 95,7 |
| ENSMUSG00000040659 | 1865,924429 | 2832,3 | 535,9 | 2,654244529 | 0,036094827 | Efhd2 | EF hand domain containing 2 [Source:MGI Symbol;Acc:MGI:106504] | 99,83 | 86,3 |
| ENSMUSG00000052837 | 151,8474423 | 115,8 | 34,8 | 2,671915718 | 0,017221499 | Junb | jun B proto-oncogene [Source:MGI Symbol;Acc:MGI:96647] | 98,68 | 24,3 |
| ENSMUSG00000040025 | 181,5515184 | 176,2 | 84,4 | 2,671996655 | 0,024493876 | Ythdf2 | YTH domain family 2 [Source:MGI Symbol;Acc:MGI:2444233] | 77,11 | 16,0 |
| ENSMUSG00000028207 | 778,6014344 | 949,7 | 350,0 | 2,675242486 | 0,040270912 | Asph | aspartate-beta-hydroxylase [Source:MGI Symbol;Acc:MGI:1914186] | 87,73 | 19,3 |
| ENSMUSG00000016637 | 624,8752474 | 1026,2 | 233,7 | 2,688511809 | 0,007880777 | Ift27 | intraflagellar transport 27 [Source:MGI Symbol;Acc:MGI:1914292] | 100,00 | 73,2 |
| ENSMUSG00000044927 | 54,37183594 | 90,4 | 17,2 | 2,695746362 | 0,031229742 | H1fx | H1 histone family, member X [Source:MGI Symbol;Acc:MGI:2685307] | 90,52 | 20,8 |
| ENSMUSG00000085587 | 607,5720966 | 1048,5 | 172,8 | 2,69998563 | 0,043140291 | Gm14493 | predicted gene 14493 [Source:MGI Symbol;Acc:MGI:3705313] | 100,00 | 97,2 |
| ENSMUSG00000056515 | 2959,495762 | 4392,7 | 1427,3 | 2,702277605 | 0,032958225 | Rab31 | RAB31, member RAS oncogene family [Source:MGI Symbol;Acc:MGI:1914603] | 99,44 | 69,4 |
| ENSMUSG00000057236 | 917,2594864 | 1202,3 | 341,8 | 2,712466555 | 0,020333974 | Rbbp4 | retinoblastoma binding protein 4 [Source:MGI Symbol;Acc:MGI:1194912] | 96,42 | 26,1 |
| ENSMUSG00000058672 | 1059,397743 | 1899,0 | 298,0 | 2,712962672 | 0,001000199 | Tubb2a | tubulin, beta 2A class IIA [Source:MGI Symbol;Acc:MGI:107861] | 77,27 | 64,9 |
| ENSMUSG00000028180 | 869,7005151 | 916,7 | 367,1 | 2,717079903 | 0,037067021 | Zranb2 | zinc finger, RAN-binding domain containing 2 [Source:MGI Symbol;Acc:MGI:1858211] | 95,26 | 40,7 |
| ENSMUSG00000057880 | 2713,525491 | 3680,1 | 1936,4 | 2,725275362 | 0,049616998 | Abat | 4-aminobutyrate aminotransferase [Source:MGI Symbol;Acc:MGI:2443582] | 95,36 | 57,0 |
| ENSMUSG00000040759 | 2169,567804 | 3259,6 | 712,5 | 2,734264431 | 0,024671043 | Cmtm5 | CKLF-like MARVEL transmembrane domain containing 5 [Source:MGI Symbol;Acc:MGI:2447164] | 100,00 | 97,0 |
| ENSMUSG00000078566 | 588,5516072 | 1008,8 | 174,5 | 2,752086181 | 0,012127993 | Bnip3 | BCL2/adenovirus E1B interacting protein 3 [Source:MGI Symbol;Acc:MGI:109326] | 88,00 | 51,2 |
| ENSMUSG00000030685 | 292,6804875 | 422,7 | 124,6 | 2,753385848 | 0,042045862 | Kctd13 | potassium channel tetramerisation domain containing 13 [Source:MGI Symbol;Acc:MGI:1923739] | 99,72 | 51,1 |
| ENSMUSG00000052752 | 385,3506829 | 494,4 | 128,8 | 2,75342884 | 0,018671178 | Traf7 | TNF receptor-associated factor 7 [Source:MGI Symbol;Acc:MGI:3042141] | 90,72 | 32,6 |
| ENSMUSG00000027430 | 363,3638555 | 453,8 | 147,5 | 2,773593072 | 0,046826836 | Dtd1 | D-tyrosyl-tRNA deacylase 1 [Source:MGI Symbol;Acc:MGI:1913294] | 95,75 | 47,0 |
| ENSMUSG00000020538 | 325,2731552 | 379,3 | 61,4 | 2,775467097 | 0,029679181 | Srebf1 | sterol regulatory element binding transcription factor 1 [Source:MGI Symbol;Acc:MGI:107606] | 97,62 | 17,7 |
| ENSMUSG00000060601 | 521,663964 | 666,5 | 162,3 | 2,778218622 | 0,003206771 | Nr1h2 | nuclear receptor subfamily 1, group H, member 2 [Source:MGI Symbol;Acc:MGI:1352463] | 79,57 | 41,0 |
| ENSMUSG00000096944 | 273,5257525 | 359,1 | 103,8 | 2,791018842 | 0,038003043 | Gm26722 | predicted gene, 26722 [Source:MGI Symbol;Acc:MGI:5477216] | #DIV/0! | #DIV/0! |
| ENSMUSG00000021750 | 7821,690492 | 11038,8 | 3003,6 | 2,793243523 | 0,041551111 | Fam107a | family with sequence similarity 107, member A [Source:MGI Symbol;Acc:MGI:3041256] | 91,94 | 77,4 |
| ENSMUSG00000066735 | 94,05432835 | 134,5 | 25,3 | 2,794356958 | 0,017614538 | Vkorc1l1 | vitamin K epoxide reductase complex, subunit 1-like 1 [Source:MGI Symbol;Acc:MGI:1916818] | 79,33 | 3,3 |
| ENSMUSG00000079357 | 179,493245 | 266,1 | 62,6 | 2,794846233 | 0,018673563 | Gm11100 | predicted gene 11100 [Source:MGI Symbol;Acc:MGI:3779338] | 23,49 | 9,6 |
| ENSMUSG00000085403 | 485,9552079 | 896,2 | 134,4 | 2,796005365 | 0,005241385 | Gm13068 | predicted gene 13068 [Source:MGI Symbol;Acc:MGI:3651967] | 32,08 | 31,4 |
| ENSMUSG00000083679 | 30,49493778 | 48,2 | 8,2 | 2,80048593 | 0,028325605 | Gm12892 | predicted gene 12892 [Source:MGI Symbol;Acc:MGI:3649649] | #DIV/0! | #DIV/0! |
| ENSMUSG00000029036 | 220,6344229 | 236,2 | 69,3 | 2,816077793 | 0,037509548 | Atad3a | ATPase family, AAA domain containing 3A [Source:MGI Symbol;Acc:MGI:1919214] | 99,77 | 32,2 |
| ENSMUSG00000024767 | 399,3464628 | 653,0 | 138,7 | 2,821819904 | 0,0135501 | Otub1 | OTU domain, ubiquitin aldehyde binding 1 [Source:MGI Symbol;Acc:MGI:2147616] | 88,17 | 49,7 |
| ENSMUSG00000057219 | 58,98886609 | 79,6 | 11,9 | 2,82686624 | 0,016761689 | Armc7 | armadillo repeat containing 7 [Source:MGI Symbol;Acc:MGI:2679719] | 98,07 | 2,8 |
| ENSMUSG00000039347 | 1178,61283 | 1761,1 | 273,6 | 2,830326952 | 0,002086313 | Atp6v0e2 | ATPase, H+ transporting, lysosomal V0 subunit E2 [Source:MGI Symbol;Acc:MGI:1923502] | 99,98 | 76,6 |
| ENSMUSG00000018340 | 554,1598538 | 572,1 | 149,4 | 2,835073367 | 0,006043248 | Anxa6 | annexin A6 [Source:MGI Symbol;Acc:MGI:88255] | 100,00 | 30,1 |
| ENSMUSG00000021371 | 1243,973363 | 2060,9 | 350,3 | 2,838174058 | 0,046938122 | Mcur1 | mitochondrial calcium uniporter regulator 1 [Source:MGI Symbol;Acc:MGI:1923387] | 99,98 | 43,4 |
| ENSMUSG00000093930 | 1515,438221 | 2470,9 | 447,6 | 2,838680654 | 0,005058806 | Hmgcs1 | 3-hydroxy-3-methylglutaryl-Coenzyme A synthase 1 [Source:MGI Symbol;Acc:MGI:107592] | 95,81 | 63,0 |
| ENSMUSG00000020882 | 125,2377263 | 134,6 | 42,8 | 2,845638915 | 0,030710999 | Cacnb1 | calcium channel, voltage-dependent, beta 1 subunit [Source:MGI Symbol;Acc:MGI:102522] | 86,81 | 8,6 |
| ENSMUSG00000028669 | 971,5740285 | 1366,3 | 340,9 | 2,847730765 | 0,032095287 | Pithd1 | ninal proteasome-interacting domain of thioredoxin-like) domain containing 1 [Source:MGI Symbol;Acc:MC | 92,52 | 55,1 |
| ENSMUSG00000018451 | 2184,911665 | 4675,8 | 767,3 | 2,851845982 | 0,033594061 | 6330403K07Rik | RIKEN cDNA 6330403K07 gene [Source:MGI Symbol;Acc:MGI:1918001] | #DIV/0! | #DIV/0! |
| ENSMUSG00000086544 | 241,8365834 | 382,8 | 88,0 | 2,854936157 | 0,007785817 | Chn1os3 | chimerin 1, opposite strand 3 [Source:MGI Symbol;Acc:MGI:3705094] | 10,80 | 4,9 |
| ENSMUSG00000008035 | 2010,53251 | 3547,8 | 525,3 | 2,861309636 | 0,040314285 | Mid1ip1 | id1 interacting protein 1 (gastrulation specific G12-like (zebrafish)) [Source:MGI Symbol;Acc:MGI:191529 | 99,82 | 92,7 |
| ENSMUSG00000030986 | 366,1728408 | 546,1 | 94,2 | 2,86289479 | 0,020710182 | Dhx32 | DEAH (Asp-Glu-Ala-His) box polypeptide 32 [Source:MGI Symbol;Acc:MGI:2141813] | 84,68 | 6,1 |
| ENSMUSG00000026473 | 5271,908081 | 8817,2 | 1752,7 | 2,864783904 | 0,018591859 | Glul | glutamate-ammonia ligase (glutamine synthetase) [Source:MGI Symbol;Acc:MGI:95739] | 99,68 | 70,1 |
| ENSMUSG00000001687 | 2450,620557 | 3740,5 | 600,0 | 2,872811188 | 0,001753896 | Ubl3 | ubiquitin-like 3 [Source:MGI Symbol;Acc:MGI:1344373] | 99,92 | 87,9 |
| ENSMUSG00000033430 | 273,5304782 | 282,1 | 40,9 | 2,873222679 | 0,024429058 | Terf2ip | telomeric repeat binding factor 2, interacting protein [Source:MGI Symbol;Acc:MGI:1929871] | 98,20 | 4,5 |
| ENSMUSG00000028982 | 1716,50645 | 3218,5 | 454,7 | 2,87780834 | 0,003532165 | Slc25a33 | solute carrier family 25, member 33 [Source:MGI Symbol;Acc:MGI:1917806] | 100,00 | 80,0 |
| ENSMUSG00000004032 | 13402,09956 | 22563,9 | 4169,9 | 2,878087256 | 0,009805025 | Gstm5 | glutathione S-transferase, mu 5 [Source:MGI Symbol;Acc:MGI:1309466] | 95,99 | 81,5 |
| ENSMUSG00000042506 | 415,6496038 | 566,7 | 78,2 | 2,88243317 | 0,001135361 | Usp22 | ubiquitin specific peptidase 22 [Source:MGI Symbol;Acc:MGI:2144157] | 99,54 | 16,8 |
| ENSMUSG00000040843 | 536,3132902 | 716,1 | 181,9 | 2,890402737 | 0,047131037 | Tiprl | TIP41, TOR signalling pathway regulator-like (S. cerevisiae) [Source:MGI Symbol;Acc:MGI:1915087] | 46,09 | 23,2 |
| ENSMUSG00000024413 | 161,971305 | 213,9 | 35,8 | 2,892256077 | 0,043946234 | Npc1 | Niemann-Pick type C1 [Source:MGI Symbol;Acc:MGI:1097712] | 94,38 | 6,8 |
| ENSMUSG00000099227 | 482,4546937 | 805,4 | 195,5 | 2,893275811 | 0,022115108 | Mir8114 | microRNA 8114 [Source:MGI Symbol;Acc:MGI:5530764] | #DIV/0! | #DIV/0! |
| ENSMUSG00000026851 | 381,7554733 | 582,3 | 90,1 | 2,894196241 | 0,003820711 | BC005624 | cDNA sequence BC005624 [Source:MGI Symbol;Acc:MGI:2385132] | 95,03 | 33,0 |
| ENSMUSG00000089922 | 144,4232131 | 195,9 | 38,3 | 2,897416342 | 0,026132051 | 3110039M20Rik | RIKEN cDNA 3110039M20 gene [Source:MGI Symbol;Acc:MGI:1914543] | 32,48 | 8,7 |
| ENSMUSG00000040997 | 4139,711407 | 7373,1 | 1172,5 | 2,898001054 | 0,010278719 | Abhd4 | abhydrolase domain containing 4 [Source:MGI Symbol;Acc:MGI:1915938] | 92,71 | 78,9 |
| ENSMUSG00000010095 | 805,4621326 | 1150,7 | 316,1 | 2,898416317 | 0,016311869 | Slc3a2 | ier family 3 (activators of dibasic and neutral amino acid transport), member 2 [Source:MGI Symbol;Acc:M | 72,16 | 36,0 |
| ENSMUSG00000022051 | 744,5990077 | 1090,4 | 171,4 | 2,912796724 | 0,00065863 | Bnip3l | BCL2/adenovirus E1B interacting protein 3-like [Source:MGI Symbol;Acc:MGI:1332659] | 89,08 | 24,7 |
| ENSMUSG00000031787 | 209,1142639 | 303,5 | 68,8 | 2,91405214 | 0,044187226 | Katnb1 | katanin p80 (WD40-containing) subunit B 1 [Source:MGI Symbol;Acc:MGI:1921437] | 89,30 | 11,3 |

| | | | | | | | | | |
|---|---|---|---|---|---|---|---|---|---|
| ENSMUSG00000021010 | 303,8941622 | 544,9 | 183,1 | 2,914481234 | 0,028723909 | Npas3 | neuronal PAS domain protein 3 [Source:MGI Symbol;Acc:MGI:1351610] | 99,79 | 22,9 |
| ENSMUSG00000030034 | 70,31907009 | 99,1 | 23,1 | 2,916719723 | 0,017886688 | Ino80b | INO80 complex subunit B [Source:MGI Symbol;Acc:MGI:1917270] | 97,87 | 24,4 |
| ENSMUSG00000027012 | 245,8638753 | 310,7 | 74,3 | 2,921247247 | 0,002485914 | Dync1i2 | dynein cytoplasmic 1 intermediate chain 2 [Source:MGI Symbol;Acc:MGI:107750] | 93,47 | 19,0 |
| ENSMUSG00000023452 | 786,3439518 | 1226,5 | 253,8 | 2,92791599 | 0,031441632 | Pisd | phosphatidylserine decarboxylase [Source:MGI Symbol;Acc:MGI:2445114] | 93,41 | 42,4 |
| ENSMUSG00000019998 | 1127,205005 | 1811,1 | 251,7 | 2,936334293 | 0,036029848 | Stx7 | syntaxin 7 [Source:MGI Symbol;Acc:MGI:1858210] | 92,43 | 63,8 |
| ENSMUSG00000078350 | 282,1195485 | 300,3 | 95,5 | 2,958210383 | 0,020710182 | Smim1 | small integral membrane protein 1 [Source:MGI Symbol;Acc:MGI:1916109] | 66,15 | 8,2 |
| ENSMUSG00000018378 | 247,3012585 | 330,9 | 194,4 | 2,960598931 | 0,04454386 | Cuedc1 | CUE domain containing 1 [Source:MGI Symbol;Acc:MGI:2144281] | 96,86 | 23,0 |
| ENSMUSG00000036940 | 198,4391547 | 206,5 | 30,5 | 2,961336534 | 0,025618029 | Kdm1a | lysine (K)-specific demethylase 1A [Source:MGI Symbol;Acc:MGI:1196256] | 95,10 | 11,2 |
| ENSMUSG00000022466 | 173,9725922 | 180,8 | 25,5 | 2,964241864 | 0,001283323 | Rpap3 | RNA polymerase II associated protein 3 [Source:MGI Symbol;Acc:MGI:1277218] | 100,00 | 13,4 |
| ENSMUSG00000022323 | 4453,483543 | 9257,7 | 1530,0 | 2,965120876 | 0,015512448 | Hrsp12 | heat-responsive protein 12 [Source:MGI Symbol;Acc:MGI:1095401] | 99,87 | 98,9 |
| ENSMUSG00000020024 | 281,0295318 | 193,9 | 55,3 | 2,965132862 | 0,032701694 | Cep83 | centrosomal protein 83 [Source:MGI Symbol;Acc:MGI:1924298] | 97,68 | 15,2 |
| ENSMUSG00000020894 | 1871,688246 | 3441,5 | 652,1 | 2,968334165 | 0,037808165 | Vamp2 | vesicle-associated membrane protein 2 [Source:MGI Symbol;Acc:MGI:1313277] | 93,67 | 66,3 |
| ENSMUSG00000021069 | 25,86964131 | 35,0 | 5,4 | 2,971181507 | 0,045543859 | Pygl | liver glycogen phosphorylase [Source:MGI Symbol;Acc:MGI:97829] | 70,36 | 1,7 |
| ENSMUSG00000029474 | 440,4184696 | 727,2 | 129,1 | 2,97388898 | 0,009176851 | Rnf34 | ring finger protein 34 [Source:MGI Symbol;Acc:MGI:2153340] | 75,82 | 18,2 |
| ENSMUSG00000030924 | 18,67156068 | 43,5 | 5,4 | 2,98281901 | 0,040821386 | 2610020H08Rik | RIKEN cDNA 2610020H08 gene [Source:MGI Symbol;Acc:MGI:1919402] | 76,32 | 3,3 |
| ENSMUSG00000091498 | 255,1289384 | 403,7 | 80,1 | 2,984917965 | 0,014731525 | Mpc1-ps | mitochondrial pyruvate carrier 1, pseudogene [Source:MGI Symbol;Acc:MGI:3781628] | #DIV/0! | #DIV/0! |
| ENSMUSG00000024422 | 139,0899203 | 188,4 | 38,4 | 2,986906121 | 0,032019326 | Dhx16 | DEAH (Asp-Glu-Ala-His) box polypeptide 16 [Source:MGI Symbol;Acc:MGI:1916442] | 99,96 | 12,2 |
| ENSMUSG00000031314 | 127,5496708 | 164,7 | 20,7 | 2,988062786 | 0,01226813 | Taf1 | RNA polymerase II, TATA box binding protein (TBP)-associated factor [Source:MGI Symbol;Acc:MGI:133 | 83,01 | 3,9 |
| ENSMUSG00000027712 | 1247,022224 | 2006,0 | 366,1 | 2,988737599 | 0,00133698 | Anxa5 | annexin A5 [Source:MGI Symbol;Acc:MGI:106008] | 99,11 | 79,2 |
| ENSMUSG00000035754 | 235,8555527 | 353,4 | 78,2 | 3,000452494 | 0,016973184 | Wdr18 | WD repeat domain 18 [Source:MGI Symbol;Acc:MGI:2158400] | 95,45 | 26,5 |
| ENSMUSG00000030310 | 2505,336743 | 4233,9 | 607,6 | 3,002877566 | 0,007479235 | Slc6a1 | olute carrier family 6 (neurotransmitter transporter, GABA), member 1 [Source:MGI Symbol;Acc:MGI:9562 | 99,84 | 74,0 |
| ENSMUSG00000020745 | 364,1114375 | 494,3 | 135,1 | 3,005482796 | 0,028160503 | Pafah1b1 | platelet-activating factor acetylhydrolase, isoform 1b, subunit 1 [Source:MGI Symbol;Acc:MGI:109520] | 90,23 | 15,7 |
| ENSMUSG00000030447 | 725,8870621 | 970,2 | 243,9 | 3,008836977 | 0,027751101 | Cyfip1 | cytoplasmic FMR1 interacting protein 1 [Source:MGI Symbol;Acc:MGI:1338801] | 90,55 | 16,2 |
| ENSMUSG00000002459 | 1068,107793 | 1815,1 | 262,5 | 3,011057567 | 0,036785298 | Rgs20 | regulator of G-protein signaling 20 [Source:MGI Symbol;Acc:MGI:1929866] | 95,53 | 54,1 |
| ENSMUSG00000022684 | 227,074382 | 223,8 | 109,5 | 3,020546593 | 0,030009825 | Bfar | bifunctional apoptosis regulator [Source:MGI Symbol;Acc:MGI:1914368] | 96,45 | 23,1 |
| ENSMUSG00000021286 | 791,0110138 | 1296,5 | 179,5 | 3,02189402 | 0,027719879 | Zfyve21 | zinc finger, FYVE domain containing 21 [Source:MGI Symbol;Acc:MGI:1915770] | 99,57 | 77,6 |
| ENSMUSG00000063801 | 319,5421567 | 450,7 | 177,5 | 3,034221469 | 0,048781064 | Ap3s2 | adaptor-related protein complex 3, sigma 2 subunit [Source:MGI Symbol;Acc:MGI:1337060] | 93,92 | 22,4 |
| ENSMUSG00000039680 | 640,3152604 | 973,7 | 140,6 | 3,034833904 | 0,001260716 | Mrps6 | mitochondrial ribosomal protein S6 [Source:MGI Symbol;Acc:MGI:2153111] | 97,87 | 75,1 |
| ENSMUSG00000039770 | 753,2811402 | 1217,8 | 178,1 | 3,036069369 | 0,023777367 | Ypel5 | yippee-like 5 (Drosophila) [Source:MGI Symbol;Acc:MGI:1916937] | 91,51 | 29,0 |
| ENSMUSG00000027650 | 51,19238715 | 78,1 | 22,8 | 3,036504298 | 0,04708563 | Tti1 | TELO2 interacting protein 1 [Source:MGI Symbol;Acc:MGI:1922675] | 87,37 | 6,2 |
| ENSMUSG00000024018 | 173,2807022 | 238,6 | 34,3 | 3,039972727 | 0,036067348 | Ccdc167 | coiled-coil domain containing 167 [Source:MGI Symbol;Acc:MGI:1915847] | 97,75 | 2,1 |
| ENSMUSG00000073639 | 2312,973779 | 4276,3 | 697,2 | 3,048622156 | 0,005008091 | Rab18 | RAB18, member RAS oncogene family [Source:MGI Symbol;Acc:MGI:102790] | 100,00 | 74,5 |
| ENSMUSG00000004056 | 2029,523432 | 3419,9 | 497,4 | 3,056415922 | 0,002796786 | Akt2 | thymoma viral proto-oncogene 2 [Source:MGI Symbol;Acc:MGI:104874] | 86,94 | 35,6 |
| ENSMUSG00000078786 | 83,35208476 | 123,5 | 14,9 | 3,061420519 | 0,02830088 | BC024978 | cDNA sequence BC024978 [Source:MGI Symbol;Acc:MGI:3041247] | 75,92 | 1,6 |
| ENSMUSG00000051403 | 224,4250429 | 323,9 | 56,3 | 3,063417471 | 0,043040966 | Ppp1r37 | protein phosphatase 1, regulatory subunit 37 [Source:MGI Symbol;Acc:MGI:2687042] | 82,62 | 23,6 |
| ENSMUSG00000026421 | 9539,536171 | 15542,5 | 2631,6 | 3,069311308 | 0,005598936 | Csrp1 | cysteine and glycine-rich protein 1 [Source:MGI Symbol;Acc:MGI:88549] | 96,06 | 87,4 |
| ENSMUSG00000036402 | 2064,768853 | 3228,4 | 742,3 | 3,071097663 | 0,032017121 | Gng12 | guanine nucleotide binding protein (G protein), gamma 12 [Source:MGI Symbol;Acc:MGI:1336171] | 98,48 | 41,0 |
| ENSMUSG00000003573 | 221,8494175 | 279,8 | 47,0 | 3,087258531 | 0,007327727 | Homer3 | homer scaffolding protein 3 [Source:MGI Symbol;Acc:MGI:1347359] | 90,43 | 9,5 |
| ENSMUSG00000044018 | 632,9854342 | 869,4 | 216,6 | 3,088304034 | 0,021130539 | Mrpl50 | mitochondrial ribosomal protein L50 [Source:MGI Symbol;Acc:MGI:107329] | 99,95 | 41,3 |
| ENSMUSG00000103651 | 48,22664751 | 60,4 | 33,0 | 3,09194301 | 0,041751163 | Gm37206 | predicted gene, 37206 [Source:MGI Symbol;Acc:MGI:5610434] | 46,06 | 5,6 |
| ENSMUSG00000020257 | 306,8106217 | 354,0 | 231,4 | 3,09441374 | 0,0347406 | Wdr82 | WD repeat domain containing 82 [Source:MGI Symbol;Acc:MGI:1924555] | 99,19 | 25,6 |
| ENSMUSG00000029613 | 713,2738151 | 966,9 | 242,2 | 3,096960796 | 0,005587554 | Eif2ak1 | eukaryotic translation initiation factor 2 alpha kinase 1 [Source:MGI Symbol;Acc:MGI:1353448] | 98,98 | 29,8 |
| ENSMUSG00000042707 | 48,03087316 | 75,4 | 16,7 | 3,101814633 | 0,046009917 | Dnali1 | dynein, axonemal, light intermediate polypeptide 1 [Source:MGI Symbol;Acc:MGI:1922813] | 97,23 | 1,0 |
| ENSMUSG00000030127 | 975,4883976 | 1630,7 | 266,1 | 3,103750097 | 0,014694066 | Cops7a | COP9 signalosome subunit 7A [Source:MGI Symbol;Acc:MGI:1349400] | 98,88 | 60,2 |
| ENSMUSG00000028737 | 334,4139007 | 483,7 | 103,1 | 3,104552607 | 0,032701694 | Aldh4a1 | aldehyde dehydrogenase 4 family, member A1 [Source:MGI Symbol;Acc:MGI:2443883] | 99,26 | 28,4 |
| ENSMUSG00000053931 | 2555,568949 | 4571,4 | 569,3 | 3,113843633 | 0,01918874 | Cnn3 | calponin 3, acidic [Source:MGI Symbol;Acc:MGI:1919244] | 91,83 | 82,0 |
| ENSMUSG00000007411 | 832,2872984 | 1410,2 | 264,6 | 3,114436716 | 0,04449741 | Mark3 | MAP/microtubule affinity regulating kinase 3 [Source:MGI Symbol;Acc:MGI:1341865] | 99,91 | 47,8 |
| ENSMUSG00000030682 | 509,5327067 | 658,9 | 139,8 | 3,117524926 | 0,031082957 | Cdipt | ylglycerol--inositol 3-phosphatidyltransferase (phosphatidylinositol synthase) [Source:MGI Symbol;Acc:MG | 83,19 | 28,2 |
| ENSMUSG00000006782 | 1391,115634 | 2135,2 | 336,9 | 3,121961733 | 0,000397712 | Cnp | 2',3'-cyclic nucleotide 3' phosphodiesterase [Source:MGI Symbol;Acc:MGI:88437] | 98,85 | 49,9 |
| ENSMUSG00000029212 | 343,7372114 | 496,2 | 111,0 | 3,129099103 | 0,025520772 | Gabrb1 | gamma-aminobutyric acid (GABA) A receptor, subunit beta 1 [Source:MGI Symbol;Acc:MGI:95619] | 74,42 | 5,5 |
| ENSMUSG00000039485 | 496,5078394 | 669,5 | 137,5 | 3,146433223 | 0,043422761 | Tspyl4 | TSPY-like 4 [Source:MGI Symbol;Acc:MGI:106393] | 99,86 | 30,4 |
| ENSMUSG00000026113 | 28,32611506 | 53,7 | 8,7 | 3,147302522 | 0,031057321 | Inpp4a | inositol polyphosphate-4-phosphatase, type I [Source:MGI Symbol;Acc:MGI:1931123] | 53,70 | 1,8 |
| ENSMUSG00000049672 | 132,2760608 | 80,7 | 11,9 | 3,149486817 | 0,029745314 | Zbtb14 | zinc finger and BTB domain containing 14 [Source:MGI Symbol;Acc:MGI:1195345] | 60,16 | 4,7 |
| ENSMUSG00000000028 | 63,48952688 | 43,1 | 6,1 | 3,155246044 | 0,025353928 | Cdc45 | cell division cycle 45 [Source:MGI Symbol;Acc:MGI:1338073] | 87,94 | 4,3 |
| ENSMUSG00000040021 | 174,551282 | 213,9 | 39,8 | 3,157721061 | 0,013799563 | Lats1 | large tumor suppressor [Source:MGI Symbol;Acc:MGI:1333883] | 93,86 | 7,9 |
| ENSMUSG00000007872 | 2402,980681 | 4238,2 | 567,3 | 3,163569043 | 0,035908736 | Id3 | inhibitor of DNA binding 3 [Source:MGI Symbol;Acc:MGI:96398] | 84,74 | 75,2 |
| ENSMUSG00000054604 | 907,272714 | 1535,8 | 271,8 | 3,164185179 | 0,047974717 | Cggbp1 | CGG triplet repeat binding protein 1 [Source:MGI Symbol;Acc:MGI:2146370] | 99,65 | 30,3 |
| ENSMUSG00000054863 | 512,7219966 | 787,3 | 132,0 | 3,165286878 | 0,023150736 | Fam19a5 | family with sequence similarity 19, member A5 [Source:MGI Symbol;Acc:MGI:2146182] | 100,00 | 33,4 |
| ENSMUSG00000106561 | 33,64361696 | 46,5 | 6,7 | 3,165608233 | 0,035534808 | AC151967.1 | | #DIV/0! | #DIV/0! |
| ENSMUSG00000022558 | 73,02931392 | 92,3 | 9,8 | 3,166574222 | 0,04454386 | Mroh1 | maestro heat-like repeat family member 1 [Source:MGI Symbol;Acc:MGI:2442558] | 67,05 | 1,1 |
| ENSMUSG00000054423 | 122,2329628 | 221,6 | 41,5 | 3,166714873 | 0,048183751 | Cadps | Ca2+-dependent secretion activator [Source:MGI Symbol;Acc:MGI:1350922] | 93,02 | 11,1 |
| ENSMUSG00000023861 | 2038,66445 | 3171,1 | 660,9 | 3,168173127 | 0,013481882 | Mpc1 | mitochondrial pyruvate carrier 1 [Source:MGI Symbol;Acc:MGI:1915240] | 61,85 | 33,7 |
| ENSMUSG00000021684 | 650,5198139 | 1189,0 | 149,1 | 3,180344554 | 0,003685841 | Pde8b | phosphodiesterase 8B [Source:MGI Symbol;Acc:MGI:2443999] | 92,48 | 26,8 |
| ENSMUSG00000032010 | 531,6443062 | 775,0 | 190,3 | 3,183145006 | 0,004192399 | Usp2 | ubiquitin specific peptidase 2 [Source:MGI Symbol;Acc:MGI:1858178] | 95,43 | 25,6 |
| ENSMUSG00000027797 | 1488,855498 | 1881,1 | 643,4 | 3,185303448 | 0,030148615 | Dclk1 | doublecortin-like kinase 1 [Source:MGI Symbol;Acc:MGI:1330861] | 94,18 | 21,7 |
| ENSMUSG00000039449 | 280,069302 | 312,1 | 42,6 | 3,186897272 | 0,01696342 | Prpf18 | PRP18 pre-mRNA processing factor 18 homolog (yeast) [Source:MGI Symbol;Acc:MGI:1914479] | 96,68 | 7,1 |
| ENSMUSG00000001627 | 849,9692241 | 1160,2 | 344,7 | 3,19325936 | 0,039810883 | Ifrd1 | interferon-related developmental regulator 1 [Source:MGI Symbol;Acc:MGI:1316717] | 69,77 | 28,3 |
| ENSMUSG00000018322 | 227,2938795 | 401,7 | 84,9 | 3,200654621 | 0,028550438 | Tomm34 | translocase of outer mitochondrial membrane 34 [Source:MGI Symbol;Acc:MGI:1914395] | 90,71 | 27,1 |
| ENSMUSG00000029787 | 153,9892533 | 342,7 | 48,4 | 3,212270714 | 0,048346724 | Avl9 | AVL9 homolog (S. cerevisiase) [Source:MGI Symbol;Acc:MGI:1926187] | 96,54 | 4,9 |
| ENSMUSG00000019143 | 238,9793146 | 223,4 | 45,9 | 3,213258431 | 0,025306615 | Hars2 | histidyl-tRNA synthetase 2, mitochondrial (putative) [Source:MGI Symbol;Acc:MGI:1918041] | 93,19 | 6,2 |
| ENSMUSG00000032656 | 21,36808577 | 30,9 | 4,0 | 3,220142014 | 0,048122098 | March3 | membrane-associated ring finger (C3HC4) 3 [Source:MGI Symbol;Acc:MGI:2443667] | 55,89 | 1,2 |
| ENSMUSG00000029673 | 82,55609106 | 105,6 | 15,0 | 3,221356323 | 0,04729708 | Auts2 | autism susceptibility candidate 2 [Source:MGI Symbol;Acc:MGI:1919847] | 63,88 | 0,3 |
| ENSMUSG00000032059 | 126,979887 | 203,1 | 33,0 | 3,221451309 | 0,014853753 | Alg9 | ısparagine-linked glycosylation 9 (alpha 1,2 mannosyltransferase) [Source:MGI Symbol;Acc:MGI:1924753 | 95,66 | 7,9 |
| ENSMUSG00000025477 | 454,3526982 | 502,8 | 160,8 | 3,221558502 | 0,023648568 | Inpp5a | inositol polyphosphate-5-phosphatase A [Source:MGI Symbol;Acc:MGI:2686961] | 91,04 | 36,8 |
| ENSMUSG00000006676 | 238,867168 | 293,0 | 156,9 | 3,222593877 | 0,04897189 | Usp19 | ubiquitin specific peptidase 19 [Source:MGI Symbol;Acc:MGI:1918722] | 95,07 | 23,0 |
| ENSMUSG00000074457 | 3310,028881 | 5645,4 | 874,7 | 3,223754098 | 0,014747957 | S100a16 | S100 calcium binding protein A16 [Source:MGI Symbol;Acc:MGI:1915110] | 98,95 | 71,5 |
| ENSMUSG00000074781 | 531,3799325 | 910,3 | 122,4 | 3,231326716 | 6,21322E-06 | Ube2n | ubiquitin-conjugating enzyme E2N [Source:MGI Symbol;Acc:MGI:1934835] | 86,52 | 13,1 |
| ENSMUSG00000085888 | 373,7509483 | 584,9 | 62,3 | 3,23258482 | 0,003908753 | Gm12224 | predicted gene 12224 [Source:MGI Symbol;Acc:MGI:3651369] | 24,99 | 17,0 |
| ENSMUSG00000033720 | 924,4757416 | 1618,3 | 223,0 | 3,234648542 | 0,005708695 | Sfxn5 | sideroflexin 5 [Source:MGI Symbol;Acc:MGI:2137681] | 96,41 | 25,3 |
| ENSMUSG00000037712 | 922,2854553 | 1445,4 | 250,7 | 3,248149635 | 0,049283007 | Fermt2 | fermitin family member 2 [Source:MGI Symbol;Acc:MGI:2385001] | 91,84 | 38,2 |
| ENSMUSG00000025373 | 448,8758499 | 494,1 | 51,5 | 3,259588935 | 0,002180801 | Rnf41 | ring finger protein 41 [Source:MGI Symbol;Acc:MGI:1914838] | 86,12 | 12,0 |
| ENSMUSG00000029651 | 54,9531284 | 44,7 | 8,9 | 3,262324162 | 0,048114522 | Mtus2 | microtubule associated tumor suppressor candidate 2 [Source:MGI Symbol;Acc:MGI:1915388] | 44,41 | 1,1 |
| ENSMUSG00000045136 | 226,0365755 | 420,6 | 76,5 | 3,263020637 | 0,010481298 | Tubb2b | tubulin, beta 2B class IIB [Source:MGI Symbol;Acc:MGI:1920960] | 78,01 | 30,3 |
| ENSMUSG00000031703 | 977,2546134 | 1306,3 | 217,1 | 3,265502326 | 0,010660604 | Itfg1 | integrin alpha FG-GAP repeat containing 1 [Source:MGI Symbol;Acc:MGI:106419] | 99,72 | 44,0 |
| ENSMUSG00000072949 | 131,7286611 | 262,7 | 49,1 | 3,266235424 | 0,049273767 | Acot1 | acyl-CoA thioesterase 1 [Source:MGI Symbol;Acc:MGI:1349396] | 80,09 | 22,7 |
| ENSMUSG00000024827 | 810,761106 | 1545,1 | 321,5 | 3,269587192 | 0,021874999 | Gldc | glycine decarboxylase [Source:MGI Symbol;Acc:MGI:1341155] | 99,84 | 58,3 |
| ENSMUSG00000025862 | 333,125678 | 486,2 | 66,2 | 3,269898927 | 0,015905714 | Stag2 | stromal antigen 2 [Source:MGI Symbol;Acc:MGI:1098583] | 92,10 | 9,2 |
| ENSMUSG00000000308 | 64,17972285 | 75,0 | 7,3 | 3,274620813 | 0,048486407 | Ckmt1 | creatine kinase, mitochondrial 1, ubiquitous [Source:MGI Symbol;Acc:MGI:99441] | 82,17 | 3,5 |
| ENSMUSG00000029458 | 301,7093414 | 297,0 | 236,2 | 3,27998516 | 0,013609945 | Brap | BRCA1 associated protein [Source:MGI Symbol;Acc:MGI:1919649] | 89,03 | 29,8 |
| ENSMUSG00000024442 | 78,81698025 | 128,8 | 21,9 | 3,283355835 | 0,034957448 | 0610009O20Rik | RIKEN cDNA 0610009O20 gene [Source:MGI Symbol;Acc:MGI:1914089] | 98,66 | 8,4 |
| ENSMUSG00000018102 | 3090,352678 | 5158,8 | 742,9 | 3,284319456 | 0,007337311 | Hist1h2bc | histone cluster 1, H2bc [Source:MGI Symbol;Acc:MGI:1915274] | 71,78 | 65,9 |
| ENSMUSG00000039414 | 26,99572854 | 50,5 | 10,5 | 3,284483058 | 0,034106967 | Heatr5b | HEAT repeat containing 5B [Source:MGI Symbol;Acc:MGI:2444098] | 70,06 | 2,4 |
| ENSMUSG00000094439 | 167,2995977 | 248,3 | 52,9 | 3,286795944 | 0,012702191 | Gm21969 | predicted gene 21969 [Source:MGI Symbol;Acc:MGI:5439438] | #DIV/0! | #DIV/0! |
| ENSMUSG00000048482 | 62,08507092 | 71,2 | 12,4 | 3,291810598 | 0,041952726 | Bdnf | brain derived neurotrophic factor [Source:MGI Symbol;Acc:MGI:88145] | 48,86 | 1,5 |
| ENSMUSG00000033569 | 75,06730325 | 99,6 | 17,4 | 3,30339372 | 0,041984702 | Adgrb3 | adhesion G protein-coupled receptor B3 [Source:MGI Symbol;Acc:MGI:2441837] | 81,80 | 3,2 |
| ENSMUSG00000018398 | 566,6545399 | 983,7 | 108,2 | 3,309033335 | 0,014694066 | Sept8 | septin 8 [Source:MGI Symbol;Acc:MGI:894310] | 96,65 | 16,9 |
| ENSMUSG00000022010 | 1215,684324 | 2723,5 | 359,5 | 3,309580977 | 0,004779948 | Tsc22d1 | TSC22 domain family, member 1 [Source:MGI Symbol;Acc:MGI:109127] | 70,62 | 24,3 |
| ENSMUSG00000026235 | 109,3263608 | 119,0 | 18,6 | 3,31704811 | 0,02528919 | Epha4 | Eph receptor A4 [Source:MGI Symbol;Acc:MGI:98277] | 85,16 | 3,4 |
| ENSMUSG00000034912 | 96,30963524 | 170,3 | 22,6 | 3,326926626 | 0,022313912 | Mdga2 | MAM domain containing glycosylphosphatidylinositol anchor 2 [Source:MGI Symbol;Acc:MGI:2444706] | 94,50 | 4,2 |
| ENSMUSG00000043463 | 33,39006717 | 79,8 | 9,7 | 3,333637841 | 0,039272304 | Rab9b | RAB9B, member RAS oncogene family [Source:MGI Symbol;Acc:MGI:2442454] | 86,52 | 2,3 |
| ENSMUSG00000059173 | 49,93463427 | 98,4 | 28,8 | 3,335209701 | 0,031924952 | Pde1a | phosphodiesterase 1A, calmodulin-dependent [Source:MGI Symbol;Acc:MGI:1201792] | 74,04 | 4,5 |
| ENSMUSG00000042046 | 108,8214261 | 190,5 | 30,3 | 3,338121516 | 0,015371493 | Dstyk | dual serine/threonine and tyrosine protein kinase [Source:MGI Symbol;Acc:MGI:1925064] | 93,46 | 5,2 |
| ENSMUSG00000031516 | 474,0064812 | 782,6 | 180,1 | 3,349060354 | 0,02300487 | Dctn6 | dynactin 6 [Source:MGI Symbol;Acc:MGI:1343154] | 68,69 | 37,2 |
| ENSMUSG00000049225 | 157,9984151 | 292,5 | 80,7 | 3,354122744 | 0,024052207 | Pdp1 | pyruvate dehyrogenase phosphatase catalytic subunit 1 [Source:MGI Symbol;Acc:MGI:2685870] | 94,39 | 18,1 |
| ENSMUSG00000039067 | 674,9342418 | 1076,0 | 137,4 | 3,363206497 | 0,021130539 | Psmd7 | proteasome (prosome, macropain) 26S subunit, non-ATPase, 7 [Source:MGI Symbol;Acc:MGI:1351511] | 98,98 | 60,1 |
| ENSMUSG00000040003 | 318,1989736 | 510,1 | 115,4 | 3,367948324 | 0,016973227 | Magi2 | rane associated guanylate kinase, WW and PDZ domain containing 2 [Source:MGI Symbol;Acc:MGI:135 | 84,07 | 14,7 |
| ENSMUSG00000004902 | 1954,037323 | 3148,9 | 339,5 | 3,368045064 | 0,001512458 | Slc25a18 | solute carrier family 25 (mitochondrial carrier), member 18 [Source:MGI Symbol;Acc:MGI:1919053] | 100,00 | 80,6 |
| ENSMUSG00000028028 | 16,96251309 | 28,2 | 2,9 | 3,376284578 | 0,048449631 | Alpk1 | alpha-kinase 1 [Source:MGI Symbol;Acc:MGI:1918731] | 50,35 | 1,0 |
| ENSMUSG00000028857 | 564,1920553 | 733,2 | 218,2 | 3,381106392 | 0,006684206 | Tmem222 | transmembrane protein 222 [Source:MGI Symbol;Acc:MGI:1098568] | 100,00 | 44,4 |
| ENSMUSG00000020629 | 856,0173071 | 1254,7 | 314,0 | 3,383464105 | 0,020940193 | Adi1 | acireductone dioxygenase 1 [Source:MGI Symbol;Acc:MGI:2144929] | 98,58 | 58,1 |
| ENSMUSG00000025856 | 75,03204322 | 118,3 | 20,3 | 3,386027472 | 0,042818634 | Pdgfa | platelet derived growth factor, alpha [Source:MGI Symbol;Acc:MGI:97527] | 77,27 | 9,9 |
| ENSMUSG00000030965 | 162,8941222 | 226,0 | 31,5 | 3,388343926 | 0,004094298 | Fam175b | family with sequence similarity 175, member B [Source:MGI Symbol;Acc:MGI:1926116] | 86,49 | 9,3 |
| ENSMUSG00000024308 | 387,7575166 | 468,7 | 113,9 | 3,391079478 | 0,014423451 | Tapbp | TAP binding protein [Source:MGI Symbol;Acc:MGI:1201689] | 97,75 | 10,3 |
| ENSMUSG00000002981 | 667,7155322 | 981,9 | 185,5 | 3,391958464 | 0,025913375 | Clptm1 | cleft lip and palate associated transmembrane protein 1 [Source:MGI Symbol;Acc:MGI:1927155] | 88,07 | 29,9 |
| ENSMUSG00000033799 | 55,26048851 | 89,4 | 9,9 | 3,392394149 | 0,047976404 | Fam208b | family with sequence similarity 208, member B [Source:MGI Symbol;Acc:MGI:2145274] | 64,42 | 1,2 |
| ENSMUSG00000020889 | 844,2585255 | 1585,4 | 180,7 | 3,392786163 | 0,008552163 | Nr1d1 | nuclear receptor subfamily 1, group D, member 1 [Source:MGI Symbol;Acc:MGI:2444210] | 98,84 | 48,6 |
| ENSMUSG00000042302 | 47,13714326 | 63,7 | 8,8 | 3,397201549 | 0,037002047 | Ehbp1 | EH domain binding protein 1 [Source:MGI Symbol;Acc:MGI:2667252] | 65,98 | 1,6 |
| ENSMUSG00000026781 | 1045,220997 | 1693,4 | 208,4 | 3,398455386 | 0,032118278 | Acbd5 | acyl-Coenzyme A binding domain containing 5 [Source:MGI Symbol;Acc:MGI:1921409] | 94,94 | 25,0 |
| ENSMUSG00000042439 | 54,42815741 | 101,3 | 11,9 | 3,406150812 | 0,012416894 | Zfp532 | zinc finger protein 532 [Source:MGI Symbol;Acc:MGI:3036282] | 81,55 | 2,3 |
| ENSMUSG00000058444 | 346,0784542 | 302,5 | 124,0 | 3,40667923 | 0,044965954 | Map2k5 | mitogen-activated protein kinase kinase 5 [Source:MGI Symbol;Acc:MGI:1346345] | 96,84 | 34,5 |
| ENSMUSG00000017009 | 1296,644605 | 2148,1 | 429,7 | 3,406763036 | 0,033778987 | Sdc4 | syndecan 4 [Source:MGI Symbol;Acc:MGI:1349164] | 100,00 | 54,4 |
| ENSMUSG00000040147 | 928,3758273 | 1700,1 | 366,9 | 3,407038748 | 0,045877678 | Maob | monoamine oxidase B [Source:MGI Symbol;Acc:MGI:96916] | 100,00 | 58,1 |
| ENSMUSG00000055707 | 234,7846275 | 270,1 | 44,6 | 3,410183729 | 0,031383924 | Klhl26 | kelch-like 26 [Source:MGI Symbol;Acc:MGI:2443079] | 99,66 | 10,7 |
| ENSMUSG00000030929 | 111,3099373 | 83,9 | 30,8 | 3,410980834 | 0,023332026 | Eri2 | exoribonuclease 2 [Source:MGI Symbol;Acc:MGI:1918401] | 81,00 | 1,5 |
| ENSMUSG00000031913 | 847,8854329 | 1208,0 | 257,4 | 3,414817368 | 0,022180393 | Vps4a | vacuolar protein sorting 4a (yeast) [Source:MGI Symbol;Acc:MGI:1890520] | 98,61 | 57,5 |
| ENSMUSG00000020741 | 324,8738499 | 371,8 | 101,9 | 3,41993437 | 0,041726586 | Cluh | clustered mitochondria (cluA/CLU1) homolog [Source:MGI Symbol;Acc:MGI:1921398] | 99,69 | 21,6 |
| ENSMUSG00000038594 | 83,87573121 | 95,0 | 108,7 | 3,42032798 | 0,038574977 | Cep85l | centrosomal protein 85-like [Source:MGI Symbol;Acc:MGI:3642684] | 92,33 | 20,9 |
| ENSMUSG00000040415 | 611,5031199 | 1000,5 | 127,0 | 3,421926085 | 0,008196587 | Dtx3 | deltex 3, E3 ubiquitin ligase [Source:MGI Symbol;Acc:MGI:2135752] | 97,43 | 50,6 |
| ENSMUSG00000097121 | 100,0501703 | 88,8 | 16,7 | 3,424463763 | 0,045265578 | D130020L05Rik | RIKEN cDNA D130020L05 gene [Source:MGI Symbol;Acc:MGI:2450270] | #DIV/0! | #DIV/0! |
| ENSMUSG00000058594 | 403,4253431 | 564,5 | 126,4 | 3,4323884 | 0,030993462 | Fbxo18 | F-box protein 18 [Source:MGI Symbol;Acc:MGI:1354699] | 98,94 | 36,2 |
| ENSMUSG00000019837 | 345,7396928 | 552,2 | 91,3 | 3,433964336 | 0,04497998 | Gtf3c6 | general transcription factor IIIC, polypeptide 6, alpha [Source:MGI Symbol;Acc:MGI:1914621] | 100,00 | 55,6 |
| ENSMUSG00000023971 | 152,8701049 | 211,6 | 40,5 | 3,434846723 | 0,049937405 | Rrp36 | ribosomal RNA processing 36 homolog (S. cerevisiae) [Source:MGI Symbol;Acc:MGI:2385053] | 100,00 | 33,9 |
| ENSMUSG00000040260 | 230,2791277 | 423,9 | 47,0 | 3,436493918 | 0,024128905 | Daam2 | dishevelled associated activator of morphogenesis 2 [Source:MGI Symbol;Acc:MGI:1923691] | 95,91 | 6,1 |
| ENSMUSG00000028128 | 1180,710347 | 2050,6 | 286,4 | 3,439879082 | 0,029459488 | F3 | coagulation factor III [Source:MGI Symbol;Acc:MGI:88381] | 100,00 | 63,3 |
| ENSMUSG00000017670 | 990,9450104 | 1557,2 | 299,2 | 3,440091995 | 0,025410497 | Elmo2 | engulfment and cell motility 2 [Source:MGI Symbol;Acc:MGI:2153045] | 81,45 | 37,6 |
| ENSMUSG00000022914 | 515,8408387 | 733,6 | 121,5 | 3,442341337 | 0,008263525 | Brwd1 | bromodomain and WD repeat domain containing 1 [Source:MGI Symbol;Acc:MGI:1890651] | 39,76 | 7,2 |
| ENSMUSG00000071573 | 39,99624536 | 83,5 | 11,7 | 3,444043105 | 0,038574663 | Rnls | renalase, FAD-dependent amine oxidase [Source:MGI Symbol;Acc:MGI:1915045] | 97,50 | 6,8 |
| ENSMUSG00000014776 | 187,9599187 | 245,8 | 55,0 | 3,445175837 | 0,021867771 | Nol3 | nucleolar protein 3 (apoptosis repressor with CARD domain) [Source:MGI Symbol;Acc:MGI:1925938] | 95,43 | 14,1 |
| ENSMUSG00000023923 | 40,42727894 | 69,9 | 8,5 | 3,449190814 | 0,040220583 | Tbc1d5 | TBC1 domain family, member 5 [Source:MGI Symbol;Acc:MGI:1919488] | 74,93 | 2,7 |
| ENSMUSG00000041297 | 91,0405164 | 130,9 | 37,6 | 3,449603702 | 0,025023 | Cdk13 | cyclin-dependent kinase 13 [Source:MGI Symbol;Acc:MGI:1916812] | 59,77 | 6,0 |
| ENSMUSG00000030766 | 114,8764374 | 76,9 | 111,9 | 3,450216305 | 0,045524527 | Arhgap17 | Rho GTPase activating protein 17 [Source:MGI Symbol;Acc:MGI:1917747] | 89,51 | 24,0 |
| ENSMUSG00000076441 | 76,48602976 | 101,2 | 13,7 | 3,452897069 | 0,00331406 | Ass1 | argininosuccinate synthetase 1 [Source:MGI Symbol;Acc:MGI:88090] | 67,75 | 10,3 |
| ENSMUSG00000015354 | 133,6135775 | 150,1 | 51,7 | 3,461209929 | 0,032176357 | Pcolce2 | procollagen C-endopeptidase enhancer 2 [Source:MGI Symbol;Acc:MGI:1923727] | 48,51 | 5,8 |
| ENSMUSG00000020668 | 216,3795323 | 413,4 | 64,3 | 3,46155252 | 0,023869497 | Kif3c | kinesin family member 3C [Source:MGI Symbol;Acc:MGI:107979] | 94,94 | 9,5 |
| ENSMUSG00000033862 | 139,6260772 | 140,7 | 42,3 | 3,462361757 | 0,028006187 | Cdk10 | cyclin-dependent kinase 10 [Source:MGI Symbol;Acc:MGI:2448549] | 92,62 | 24,5 |
| ENSMUSG00000097257 | 99,97951419 | 156,4 | 31,6 | 3,466214376 | 0,021445849 | Gm26768 | predicted gene, 26768 [Source:MGI Symbol;Acc:MGI:5477262] | #DIV/0! | #DIV/0! |
| ENSMUSG00000022272 | 611,5575088 | 829,7 | 111,0 | 3,470886686 | 0,019063996 | Myo10 | myosin X [Source:MGI Symbol;Acc:MGI:107716] | 71,29 | 10,1 |
| ENSMUSG00000049739 | 106,4435845 | 52,2 | 7,7 | 3,477382398 | 0,045272001 | Zfp646 | zinc finger protein 646 [Source:MGI Symbol;Acc:MGI:3665412] | 49,91 | 0,3 |
| ENSMUSG00000050315 | 36,72830022 | 56,5 | 5,1 | 3,479439404 | 0,0277816 | Synpo2 | synaptopodin 2 [Source:MGI Symbol;Acc:MGI:2153070] | 41,94 | 0,9 |
| ENSMUSG00000043969 | 369,7379423 | 579,7 | 215,1 | 3,485824536 | 0,037609424 | Emx2 | empty spiracles homeobox 2 [Source:MGI Symbol;Acc:MGI:95388] | 97,60 | 30,2 |
| ENSMUSG00000020333 | 2718,540867 | 4355,3 | 398,2 | 3,48703406 | 0,00138233 | Acsl6 | acyl-CoA synthetase long-chain family member 6 [Source:MGI Symbol;Acc:MGI:894291] | 94,11 | 22,7 |
| ENSMUSG00000061410 | 42,66722954 | 80,6 | 12,1 | 3,490188477 | 0,033811349 | Zcchc14 | zinc finger, CCHC domain containing 14 [Source:MGI Symbol;Acc:MGI:2159407] | 57,85 | 2,1 |
| ENSMUSG00000001833 | 3636,853996 | 6315,9 | 938,9 | 3,492282414 | 0,001270433 | Sept7 | septin 7 [Source:MGI Symbol;Acc:MGI:1335094] | 98,20 | 73,2 |
| ENSMUSG00000087478 | 902,4672165 | 1382,4 | 229,3 | 3,495268015 | 0,006182165 | 4930506C21Rik | RIKEN cDNA 4930506C21 gene [Source:MGI Symbol;Acc:MGI:1922310] | #DIV/0! | #DIV/0! |
| ENSMUSG00000060475 | 150,3798329 | 183,2 | 28,7 | 3,496781181 | 0,007221778 | Wtap | Wilms tumour 1-associating protein [Source:MGI Symbol;Acc:MGI:1926395] | 55,78 | 5,2 |
| ENSMUSG00000024191 | 126,5661557 | 142,6 | 33,8 | 3,502752689 | 0,04192962 | Bnip1 | BCL2/adenovirus E1B interacting protein 1 [Source:MGI Symbol;Acc:MGI:109328] | 99,78 | 30,7 |
| ENSMUSG00000049939 | 44,99699398 | 54,4 | 6,5 | 3,503693514 | 0,031879257 | Lrrc4 | leucine rich repeat containing 4 [Source:MGI Symbol;Acc:MGI:2182081] | 66,63 | 1,9 |
| ENSMUSG00000031609 | 160,3764878 | 233,6 | 43,9 | 3,504196436 | 0,048905248 | Sap30 | sin3 associated polypeptide [Source:MGI Symbol;Acc:MGI:1929129] | 100,00 | 34,2 |
| ENSMUSG00000020886 | 52,69926329 | 79,2 | 32,4 | 3,508228787 | 0,027806939 | Dlg4 | discs, large homolog 4 (Drosophila) [Source:MGI Symbol;Acc:MGI:1277959] | 61,17 | 7,0 |
| ENSMUSG00000024410 | 161,102125 | 190,0 | 16,7 | 3,510545018 | 0,0231486 | 3110002H16Rik | RIKEN cDNA 3110002H16 gene [Source:MGI Symbol;Acc:MGI:1916528] | 98,99 | 3,8 |
| ENSMUSG00000025451 | 97,67012017 | 120,5 | 17,4 | 3,5118043 | 0,00461509 | Paip1 | polyadenylate binding protein-interacting protein 1 [Source:MGI Symbol;Acc:MGI:2384993] | 91,39 | 7,3 |
| ENSMUSG00000020137 | 67,52950928 | 85,8 | 21,8 | 3,513074811 | 0,018164583 | Thap2 | THAP domain containing, apoptosis associated protein 2 [Source:MGI Symbol;Acc:MGI:1914066] | 94,57 | 5,4 |
| ENSMUSG00000041132 | 325,9403107 | 254,3 | 234,0 | 3,513624827 | 0,041059844 | N4bp2l1 | NEDD4 binding protein 2-like 1 [Source:MGI Symbol;Acc:MGI:2140872] | 85,74 | 30,5 |
| ENSMUSG00000039697 | 25,09474982 | 57,6 | 5,1 | 3,514111253 | 0,024112472 | Ncoa7 | nuclear receptor coactivator 7 [Source:MGI Symbol;Acc:MGI:2444847] | 72,43 | 1,6 |
| ENSMUSG00000020776 | 317,3486792 | 402,6 | 71,2 | 3,514914767 | 0,029839664 | Fbf1 | Fas (TNFRSF6) binding factor 1 [Source:MGI Symbol;Acc:MGI:1922033] | 99,50 | 12,5 |
| ENSMUSG00000039220 | 131,8512804 | 152,4 | 20,9 | 3,516084374 | 0,017048315 | Ppp1r10 | protein phosphatase 1, regulatory subunit 10 [Source:MGI Symbol;Acc:MGI:1289273] | 90,98 | 3,2 |
| ENSMUSG00000027777 | 625,9823286 | 1113,4 | 173,2 | 3,516976598 | 0,010101794 | Schip1 | schwannomin interacting protein 1 [Source:MGI Symbol;Acc:MGI:1353557] | 82,97 | 28,8 |
| ENSMUSG00000002550 | 370,5328053 | 638,2 | 126,3 | 3,519702036 | 0,039939641 | Uck1 | uridine-cytidine kinase 1 [Source:MGI Symbol;Acc:MGI:98904] | 74,75 | 32,9 |
| ENSMUSG00000037075 | 193,1818851 | 274,1 | 111,9 | 3,525400344 | 0,038574252 | Rnf139 | ring finger protein 139 [Source:MGI Symbol;Acc:MGI:1923091] | 70,70 | 7,8 |
| ENSMUSG00000022139 | 735,7644587 | 1291,7 | 313,9 | 3,527580559 | 0,04509201 | Mbnl2 | muscleblind-like 2 [Source:MGI Symbol;Acc:MGI:2145597] | 99,96 | 41,2 |

| | | | | | | | | | |
|---|---|---|---|---|---|---|---|---|---|
| ENSMUSG00000045095 | 148,6739786 | 153,6 | 297,8 | 3,530236454 | 0,035076824 | Magi1 | rane associated guanylate kinase, WW and PDZ domain containing 1 [Source:MGI Symbol;Acc:MGI:120 | 66,36 | 18,8 |
| ENSMUSG00000037111 | 170,96791 | 132,8 | 37,6 | 3,536005206 | 0,00551827 | Setd7 | SET domain containing (lysine methyltransferase) 7 [Source:MGI Symbol;Acc:MGI:1920501] | 84,94 | 1,8 |
| ENSMUSG00000031309 | 23,84938945 | 50,3 | 8,9 | 3,540478522 | 0,032783163 | Rps6ka3 | ribosomal protein S6 kinase polypeptide 3 [Source:MGI Symbol;Acc:MGI:104557] | 57,75 | 1,0 |
| ENSMUSG00000056486 | 1631,697437 | 2658,4 | 569,2 | 3,549330761 | 0,001198022 | Chn1 | chimerin 1 [Source:MGI Symbol;Acc:MGI:1915674] | 90,92 | 29,7 |
| ENSMUSG00000024498 | 170,4421972 | 136,8 | 42,3 | 3,553034773 | 0,038027541 | Tcerg1 | transcription elongation regulator 1 (CA150) [Source:MGI Symbol;Acc:MGI:1926421] | 99,41 | 6,2 |
| ENSMUSG00000027425 | 151,0840805 | 140,3 | 91,3 | 3,562221999 | 0,036830707 | Csrp2bp | cysteine and glycine-rich protein 2 binding protein [Source:MGI Symbol;Acc:MGI:1917264] | 93,71 | 23,2 |
| ENSMUSG00000026657 | 133,2382488 | 265,9 | 53,7 | 3,563295126 | 0,012957623 | Frmd4a | FERM domain containing 4A [Source:MGI Symbol;Acc:MGI:1919850] | 70,84 | 3,9 |
| ENSMUSG00000062627 | 107,4331136 | 103,4 | 56,7 | 3,572480863 | 0,01858131 | Mysm1 | myb-like, SWIRM and MPN domains 1 [Source:MGI Symbol;Acc:MGI:2444584] | 77,83 | 5,9 |
| ENSMUSG00000030822 | 218,3533569 | 364,8 | 118,0 | 3,572704198 | 0,014386881 | Prr14 | proline rich 14 [Source:MGI Symbol;Acc:MGI:2384565] | 97,04 | 33,9 |
| ENSMUSG00000035983 | 62,42524382 | 91,7 | 27,3 | 3,576352505 | 0,048873194 | Gm7008 | predicted gene 7008 [Source:MGI Symbol;Acc:MGI:3647211] | 83,46 | 11,1 |
| ENSMUSG00000024603 | 380,7942734 | 547,3 | 83,7 | 3,577209313 | 0,008733864 | Dctn4 | dynactin 4 [Source:MGI Symbol;Acc:MGI:1914915] | 99,94 | 13,5 |
| ENSMUSG00000098912 | 183,6044868 | 296,4 | 41,1 | 3,582319403 | 0,018472201 | 1500004A13Rik | RIKEN cDNA 1500004A13 gene [Source:MGI Symbol;Acc:MGI:2442808] | #DIV/0! | #DIV/0! |
| ENSMUSG00000064215 | 556,6591641 | 666,7 | 94,9 | 3,584757005 | 0,014423451 | Ifi27 | interferon, alpha-inducible protein 27 [Source:MGI Symbol;Acc:MGI:1277180] | 99,74 | 58,5 |
| ENSMUSG00000100252 | 143,6473948 | 193,9 | 25,7 | 3,588117464 | 0,006489994 | Mir124-2hg | Mir124-2 host gene (non-protein coding) [Source:MGI Symbol;Acc:MGI:1917691] | #DIV/0! | #DIV/0! |
| ENSMUSG00000039530 | 507,7168092 | 742,6 | 109,7 | 3,590002148 | 0,019481169 | Tusc3 | tumor suppressor candidate 3 [Source:MGI Symbol;Acc:MGI:1933134] | 100,00 | 54,2 |
| ENSMUSG00000038070 | 55,79204293 | 89,2 | 33,8 | 3,590702695 | 0,04449741 | Cntln | centlein, centrosomal protein [Source:MGI Symbol;Acc:MGI:2443104] | 67,92 | 2,4 |
| ENSMUSG00000086994 | 22,14694389 | 26,7 | 5,5 | 3,598769645 | 0,032701694 | Gm13402 | predicted gene 13402 [Source:MGI Symbol;Acc:MGI:3649787] | 34,20 | 9,1 |
| ENSMUSG00000090861 | 28,93264677 | 48,0 | 4,9 | 3,599039335 | 0,027588436 | Arf4os | ADP-ribosylation factor 4, opposite strand [Source:MGI Symbol;Acc:MGI:3642411] | 88,10 | 5,8 |
| ENSMUSG00000055373 | 179,5221789 | 309,1 | 105,9 | 3,604194472 | 0,034908947 | Fut9 | fucosyltransferase 9 [Source:MGI Symbol;Acc:MGI:1330859] | 82,19 | 5,0 |
| ENSMUSG00000098004 | 29,63529798 | 35,8 | 30,5 | 3,605516538 | 0,036038224 | Gm27027 | predicted gene, 27027 [Source:MGI Symbol;Acc:MGI:5504142] | 64,23 | 10,5 |
| ENSMUSG00000039217 | 558,9442091 | 823,3 | 123,9 | 3,60657922 | 0,00921148 | Il18 | interleukin 18 [Source:MGI Symbol;Acc:MGI:107936] | 98,88 | 37,8 |
| ENSMUSG00000006456 | 116,2245521 | 153,3 | 41,2 | 3,607802516 | 0,032425801 | Rbm14 | RNA binding motif protein 14 [Source:MGI Symbol;Acc:MGI:1929092] | 79,68 | 14,6 |
| ENSMUSG00000078517 | 264,6808663 | 422,1 | 37,0 | 3,610278296 | 0,004999214 | Emc1 | ER membrane protein complex subunit 1 [Source:MGI Symbol;Acc:MGI:2443696] | 90,95 | 5,6 |
| ENSMUSG00000041762 | 125,0930194 | 213,8 | 55,6 | 3,614580226 | 0,008882968 | Gpr155 | G protein-coupled receptor 155 [Source:MGI Symbol;Acc:MGI:1915776] | 95,42 | 10,5 |
| ENSMUSG00000021453 | 738,739459 | 1412,0 | 213,2 | 3,617881543 | 0,000663947 | Gadd45g | growth arrest and DNA-damage-inducible 45 gamma [Source:MGI Symbol;Acc:MGI:1346325] | 100,00 | 58,3 |
| ENSMUSG00000032311 | 94,52740317 | 129,3 | 46,4 | 3,619158227 | 0,045477031 | Nrg4 | neuregulin 4 [Source:MGI Symbol;Acc:MGI:1933833] | 11,29 | 2,4 |
| ENSMUSG00000102422 | 585,8950719 | 1038,6 | 161,2 | 3,619618972 | 0,015518472 | Schip1 | schwannomin interacting protein 1 [Source:MGI Symbol;Acc:MGI:1353557] | 82,95 | 33,3 |
| ENSMUSG00000006356 | 1712,865211 | 3009,7 | 281,2 | 3,625101782 | 0,012282057 | Crip2 | cysteine rich protein 2 [Source:MGI Symbol;Acc:MGI:1915587] | 93,73 | 67,8 |
| ENSMUSG00000027998 | 212,6432245 | 246,1 | 80,8 | 3,626640023 | 0,014975476 | Plrg1 | pleiotropic regulator 1, PRL1 homolog (Arabidopsis) [Source:MGI Symbol;Acc:MGI:1858197] | 98,18 | 28,0 |
| ENSMUSG00000049807 | 110,2215032 | 136,2 | 11,4 | 3,628422762 | 0,021479852 | Arhgap23 | Rho GTPase activating protein 23 [Source:MGI Symbol;Acc:MGI:3697726] | 91,61 | 2,3 |
| ENSMUSG00000048264 | 53,39601886 | 111,6 | 15,8 | 3,637795335 | 0,020250365 | Dip2c | disco interacting protein 2 homolog C [Source:MGI Symbol;Acc:MGI:1920179] | 73,81 | 2,4 |
| ENSMUSG00000030397 | 91,12434732 | 111,8 | 13,2 | 3,637873349 | 0,030076884 | Mark4 | MAP/microtubule affinity regulating kinase 4 [Source:MGI Symbol;Acc:MGI:1920955] | 85,63 | 3,0 |
| ENSMUSG00000055421 | 184,7660644 | 205,1 | 109,3 | 3,637911389 | 0,01410677 | Pcdh9 | protocadherin 9 [Source:MGI Symbol;Acc:MGI:1306801] | 92,07 | 13,9 |
| ENSMUSG00000040466 | 112,6044086 | 143,7 | 21,4 | 3,644574713 | 0,025878717 | Blvrb | biliverdin reductase B (flavin reductase (NADPH)) [Source:MGI Symbol;Acc:MGI:2385271] | 64,36 | 14,9 |
| ENSMUSG00000032640 | 117,17113 | 113,1 | 21,8 | 3,645530692 | 0,007496462 | Chsy1 | chondroitin sulfate synthase 1 [Source:MGI Symbol;Acc:MGI:2681120] | 89,68 | 7,0 |
| ENSMUSG00000033885 | 146,4013779 | 250,4 | 35,9 | 3,647733963 | 0,007556503 | Pxk | PX domain containing serine/threonine kinase [Source:MGI Symbol;Acc:MGI:1289230] | 98,38 | 10,7 |
| ENSMUSG00000037286 | 203,3390975 | 258,2 | 22,0 | 3,651156371 | 0,014210371 | Stag1 | stromal antigen 1 [Source:MGI Symbol;Acc:MGI:1098658] | 91,84 | 2,9 |
| ENSMUSG00000002280 | 280,4419306 | 452,0 | 44,2 | 3,656029978 | 0,003764373 | Narfl | nuclear prelamin A recognition factor-like [Source:MGI Symbol;Acc:MGI:1914813] | 97,12 | 13,3 |
| ENSMUSG00000020644 | 7067,298447 | 13113,0 | 1346,6 | 3,658053668 | 0,032295618 | Id2 | inhibitor of DNA binding 2 [Source:MGI Symbol;Acc:MGI:96397] | 100,00 | 98,5 |
| ENSMUSG00000020366 | 670,6595536 | 1051,5 | 228,7 | 3,660470531 | 0,016261287 | Mapk9 | mitogen-activated protein kinase 9 [Source:MGI Symbol;Acc:MGI:1346862] | 100,00 | 30,3 |
| ENSMUSG00000051497 | 327,5861593 | 372,6 | 59,3 | 3,667681699 | 0,045334553 | Kcnj16 | potassium inwardly-rectifying channel, subfamily J, member 16 [Source:MGI Symbol;Acc:MGI:1314842] | 93,13 | 17,8 |
| ENSMUSG00000037573 | 278,9323155 | 471,2 | 76,4 | 3,667932262 | 0,021163503 | Tob1 | transducer of ErbB-2.1 [Source:MGI Symbol;Acc:MGI:1349721] | 98,27 | 38,0 |
| ENSMUSG00000021579 | 70,24339767 | 71,5 | 15,4 | 3,668045115 | 0,01922771 | Lrrc14b | leucine rich repeat containing 14B [Source:MGI Symbol;Acc:MGI:2145269] | 78,63 | 1,8 |
| ENSMUSG00000032118 | 1340,551878 | 2134,5 | 499,7 | 3,669510186 | 0,008615342 | Fez1 | fasciculation and elongation protein zeta 1 (zygin I) [Source:MGI Symbol;Acc:MGI:2670976] | 69,36 | 30,9 |
| ENSMUSG00000027016 | 60,30798616 | 107,7 | 50,4 | 3,672437652 | 0,046627412 | Zfp385b | zinc finger protein 385B [Source:MGI Symbol;Acc:MGI:2444734] | 59,58 | 7,4 |
| ENSMUSG00000004567 | 143,69211 | 141,0 | 74,7 | 3,673240663 | 0,049615669 | Mcoln1 | mucolipin 1 [Source:MGI Symbol;Acc:MGI:1890498] | 92,80 | 20,0 |
| ENSMUSG00000052248 | 19,57505306 | 35,2 | 10,3 | 3,676317033 | 0,03782395 | Zeb2os | zinc finger E-box binding homeobox 2, opposite strand [Source:MGI Symbol;Acc:MGI:3652108] | 85,66 | 16,3 |
| ENSMUSG00000068479 | 62,30541346 | 91,1 | 11,4 | 3,677191361 | 0,010956967 | Mfap1a | microfibrillar-associated protein 1A [Source:MGI Symbol;Acc:MGI:1914782] | 48,50 | 2,2 |
| ENSMUSG00000021661 | 264,6948191 | 478,4 | 84,1 | 3,678488465 | 0,011341232 | Ankra2 | ankyrin repeat, family A (RFXANK-like), 2 [Source:MGI Symbol;Acc:MGI:1915808] | 80,81 | 19,4 |
| ENSMUSG00000033392 | 201,0146086 | 339,7 | 25,8 | 3,681933809 | 0,003090135 | Clasp2 | CLIP associating protein 2 [Source:MGI Symbol;Acc:MGI:1923749] | 96,21 | 4,6 |
| ENSMUSG00000053560 | 71,00273577 | 40,7 | 48,1 | 3,683699442 | 0,049587815 | Ier2 | immediate early response 2 [Source:MGI Symbol;Acc:MGI:104815] | 97,13 | 25,9 |
| ENSMUSG00000028049 | 218,7360055 | 287,1 | 69,1 | 3,686258285 | 0,005859943 | Scamp3 | secretory carrier membrane protein 3 [Source:MGI Symbol;Acc:MGI:1346346] | 100,00 | 35,8 |
| ENSMUSG00000041229 | 31,83563861 | 67,3 | 5,6 | 3,686948308 | 0,019554562 | Phf8 | PHD finger protein 8 [Source:MGI Symbol;Acc:MGI:2444341] | 46,74 | 0,8 |
| ENSMUSG00000025885 | 58,39824854 | 71,0 | 7,7 | 3,690290538 | 0,027588436 | Myo5b | myosin VB [Source:MGI Symbol;Acc:MGI:106598] | 58,52 | 1,6 |
| ENSMUSG00000020654 | 53,81191878 | 93,3 | 12,3 | 3,691873495 | 0,043467208 | Adcy3 | adenylate cyclase 3 [Source:MGI Symbol;Acc:MGI:99675] | 90,02 | 0,7 |
| ENSMUSG00000020273 | 33,3696064 | 50,1 | 4,5 | 3,694569356 | 0,0300223 | Papolg | poly(A) polymerase gamma [Source:MGI Symbol;Acc:MGI:2442119] | 60,83 | 1,7 |
| ENSMUSG00000041650 | 317,1521434 | 496,1 | 42,3 | 3,695676001 | 0,000412708 | Pcca | propionyl-Coenzyme A carboxylase, alpha polypeptide [Source:MGI Symbol;Acc:MGI:97499] | 96,24 | 3,9 |
| ENSMUSG00000028556 | 63,91875745 | 80,9 | 6,2 | 3,695768328 | 0,025126393 | Dock7 | dedicator of cytokinesis 7 [Source:MGI Symbol;Acc:MGI:1914549] | 77,01 | 1,4 |
| ENSMUSG00000049489 | 251,0571837 | 333,2 | 165,8 | 3,700653909 | 0,04736372 | Fam58b | family with sequence similarity 58, member B [Source:MGI Symbol;Acc:MGI:1916359] | 99,62 | 39,7 |
| ENSMUSG00000030589 | 53,88081255 | 43,4 | 7,3 | 3,700869005 | 0,01149796 | Rasgrp4 | RAS guanyl releasing protein 4 [Source:MGI Symbol;Acc:MGI:2386851] | 19,76 | 3,2 |
| ENSMUSG00000021760 | 346,8294753 | 535,4 | 77,0 | 3,702796559 | 0,036798784 | Gpx8 | glutathione peroxidase 8 (putative) [Source:MGI Symbol;Acc:MGI:1916840] | 100,00 | 52,6 |
| ENSMUSG00000079555 | 198,6484629 | 235,4 | 31,0 | 3,710282158 | 0,015403999 | Haus3 | HAUS augmin-like complex, subunit 3 [Source:MGI Symbol;Acc:MGI:2387633] | 89,39 | 6,4 |
| ENSMUSG00000041961 | 39,68552177 | 67,1 | 7,2 | 3,71032361 | 0,049892314 | Znrf3 | zinc and ring finger 3 [Source:MGI Symbol;Acc:MGI:3039616] | 52,13 | 1,0 |
| ENSMUSG00000036114 | 238,7692083 | 285,6 | 35,6 | 3,710916892 | 0,002147266 | Rpp25l | ribonuclease P/MRP 25 subunit-like [Source:MGI Symbol;Acc:MGI:1917211] | 100,00 | 48,8 |
| ENSMUSG00000031788 | 545,5937213 | 894,7 | 140,1 | 3,711332977 | 0,044692 | Kifc3 | kinesin family member C3 [Source:MGI Symbol;Acc:MGI:109202] | 96,06 | 38,0 |
| ENSMUSG00000066026 | 253,8342968 | 385,9 | 62,4 | 3,713594185 | 0,029845441 | Dhrs3 | dehydrogenase/reductase (SDR family) member 3 [Source:MGI Symbol;Acc:MGI:1315215] | 84,04 | 25,4 |
| ENSMUSG00000093574 | 399,708487 | 607,9 | 94,6 | 3,717471292 | 0,01949513 | Gm20671 | predicted gene 20671 [Source:MGI Symbol;Acc:MGI:5313118] | 80,53 | 39,8 |
| ENSMUSG00000007777 | 487,7046392 | 803,2 | 119,2 | 3,719021449 | 0,031598818 | 0610009B22Rik | RIKEN cDNA 0610009B22 gene [Source:MGI Symbol;Acc:MGI:1913300] | 88,51 | 50,9 |
| ENSMUSG00000078451 | 89,74106996 | 141,2 | 16,1 | 3,719687927 | 0,019371516 | Ppil6 | peptidylprolyl isomerase (cyclophilin)-like 6 [Source:MGI Symbol;Acc:MGI:1920325] | 98,31 | 9,8 |
| ENSMUSG00000031402 | 169,7588564 | 160,0 | 132,7 | 3,721241177 | 0,013799563 | Mpp1 | membrane protein, palmitoylated [Source:MGI Symbol;Acc:MGI:105941] | 76,80 | 25,1 |
| ENSMUSG00000024381 | 209,1334197 | 268,1 | 109,3 | 3,721285541 | 0,040594928 | Bin1 | bridging integrator 1 [Source:MGI Symbol;Acc:MGI:108092] | 98,33 | 31,5 |
| ENSMUSG00000065954 | 397,6527289 | 462,7 | 81,8 | 3,726760251 | 0,011883954 | Tacc1 | transforming, acidic coiled-coil containing protein 1 [Source:MGI Symbol;Acc:MGI:2443510] | 93,01 | 10,5 |
| ENSMUSG00000020647 | 594,7663381 | 1023,7 | 80,2 | 3,733771615 | 0,020359596 | Ncoa1 | nuclear receptor coactivator 1 [Source:MGI Symbol;Acc:MGI:1276523] | 99,40 | 11,3 |
| ENSMUSG00000058258 | 170,620036 | 299,6 | 23,9 | 3,733838715 | 9,38647E-06 | Idi1 | isopentenyl-diphosphate delta isomerase [Source:MGI Symbol;Acc:MGI:2442264] | 39,52 | 6,6 |
| ENSMUSG00000020792 | 308,1239909 | 310,8 | 142,1 | 3,734944507 | 0,00883569 | Exoc7 | exocyst complex component 7 [Source:MGI Symbol;Acc:MGI:1859270] | 89,37 | 27,5 |
| ENSMUSG00000002109 | 170,3332477 | 94,6 | 158,1 | 3,735647492 | 0,0223201 | Ddb2 | damage specific DNA binding protein 2 [Source:MGI Symbol;Acc:MGI:1355314] | 69,46 | 24,5 |
| ENSMUSG00000029913 | 56,2265729 | 108,0 | 52,6 | 3,735676391 | 0,035777858 | Prdm5 | PR domain containing 5 [Source:MGI Symbol;Acc:MGI:1918029] | 98,87 | 22,9 |
| ENSMUSG00000029174 | 96,54084278 | 82,9 | 81,9 | 3,742111655 | 0,030699902 | Tbc1d1 | TBC1 domain family, member 1 [Source:MGI Symbol;Acc:MGI:1889508] | 68,89 | 15,9 |
| ENSMUSG00000034472 | 119,1314322 | 144,1 | 76,0 | 3,743430117 | 0,048071924 | Rasd2 | RASD family, member 2 [Source:MGI Symbol;Acc:MGI:1922391] | 86,93 | 24,1 |
| ENSMUSG00000071064 | 26,01468623 | 41,9 | 4,9 | 3,743627036 | 0,018089692 | Zfp827 | zinc finger protein 827 [Source:MGI Symbol;Acc:MGI:2444807] | 46,84 | 0,9 |
| ENSMUSG00000007989 | 75,04118678 | 102,3 | 61,5 | 3,744406088 | 0,040128039 | Fzd3 | frizzled class receptor 3 [Source:MGI Symbol;Acc:MGI:108476] | 94,35 | 11,0 |
| ENSMUSG00000035459 | 14,90017508 | 39,0 | 4,0 | 3,744561024 | 0,049283007 | Stab2 | stabilin 2 [Source:MGI Symbol;Acc:MGI:2178743] | 31,51 | 0,9 |
| ENSMUSG00000030519 | 285,9838615 | 414,8 | 73,6 | 3,746208795 | 0,008196587 | Apba2 | amyloid beta (A4) precursor protein-binding, family A, member 2 [Source:MGI Symbol;Acc:MGI:1261791] | 96,36 | 25,3 |
| ENSMUSG00000047284 | 78,29055506 | 45,6 | 3,8 | 3,746591679 | 0,026398355 | Neurl4 | neuralized E3 ubiquitin protein ligase 4 [Source:MGI Symbol;Acc:MGI:1921092] | 69,58 | 1,0 |
| ENSMUSG00000024642 | 355,2898146 | 510,8 | 230,7 | 3,750901666 | 0,046232076 | Tle4 | transducin-like enhancer of split 4, homolog of Drosophila E(spl) [Source:MGI Symbol;Acc:MGI:104633] | 97,78 | 29,7 |
| ENSMUSG00000030750 | 316,3438701 | 491,8 | 66,6 | 3,7522945 | 0,02586384 | Nsmce1 | non-SMC element 1 homolog (S. cerevisiae) [Source:MGI Symbol;Acc:MGI:1914961] | 88,18 | 43,3 |
| ENSMUSG00000022546 | 119,0404642 | 108,1 | 84,7 | 3,753060497 | 0,019472254 | Gpt | glutamic pyruvic transaminase, soluble [Source:MGI Symbol;Acc:MGI:95802] | 99,62 | 18,7 |
| ENSMUSG00000106247 | 110,1221183 | 156,0 | 44,5 | 3,761004317 | 0,015462113 | Gm43720 | predicted gene 43720 [Source:MGI Symbol;Acc:MGI:5663857] | #DIV/0! | #DIV/0! |
| ENSMUSG00000067860 | 79,50752458 | 52,1 | 4,4 | 3,761339512 | 0,026466872 | Zic3 | zinc finger protein of the cerebellum 3 [Source:MGI Symbol;Acc:MGI:106676] | 71,74 | 1,6 |
| ENSMUSG00000033671 | 63,29564118 | 54,0 | 5,2 | 3,763044997 | 0,036438839 | Cep350 | centrosomal protein 350 [Source:MGI Symbol;Acc:MGI:1921331] | 38,59 | 0,8 |
| ENSMUSG00000105647 | 52,28534846 | 82,2 | 13,1 | 3,763883922 | 0,028022415 | Gm43695 | predicted gene 43695 [Source:MGI Symbol;Acc:MGI:5663832] | #DIV/0! | #DIV/0! |
| ENSMUSG00000032120 | 205,5496121 | 222,9 | 53,9 | 3,766147962 | 0,048323693 | C2cd2l | C2 calcium-dependent domain containing 2-like [Source:MGI Symbol;Acc:MGI:1919014] | 81,41 | 17,3 |
| ENSMUSG00000000538 | 633,3079565 | 1132,1 | 104,0 | 3,767832524 | 0,018030954 | Tom1l2 | target of myb1-like 2 (chicken) [Source:MGI Symbol;Acc:MGI:2443306] | 84,22 | 17,0 |
| ENSMUSG00000024878 | 87,73607049 | 168,3 | 73,5 | 3,780200545 | 0,019371516 | Cbwd1 | COBW domain containing 1 [Source:MGI Symbol;Acc:MGI:2385089] | 98,44 | 18,0 |
| ENSMUSG00000004562 | 96,14030275 | 165,9 | 17,1 | 3,780654746 | 0,004430539 | Arhgef40 | Rho guanine nucleotide exchange factor (GEF) 40 [Source:MGI Symbol;Acc:MGI:2685515] | 86,50 | 2,1 |
| ENSMUSG00000049488 | 22,82154049 | 46,8 | 4,0 | 3,780786952 | 0,024766487 | Tmem67 | transmembrane protein 67 [Source:MGI Symbol;Acc:MGI:1923928] | 66,46 | 1,3 |
| ENSMUSG00000021451 | 111,5982748 | 180,9 | 116,2 | 3,782833515 | 0,032017121 | Sema4d | lin domain (Ig), transmembrane domain (TM) and short cytoplasmic domain, (semaphorin) 4D [Source:MC | 90,44 | 19,0 |
| ENSMUSG00000022197 | 28,78076966 | 41,6 | 3,8 | 3,785791133 | 0,012660573 | Pdzd2 | PDZ domain containing 2 [Source:MGI Symbol;Acc:MGI:1922394] | 39,36 | 0,1 |
| ENSMUSG00000055932 | 529,8185053 | 728,0 | 458,3 | 3,790825004 | 0,016316251 | Fto | fat mass and obesity associated [Source:MGI Symbol;Acc:MGI:1347093] | 88,01 | 32,5 |
| ENSMUSG00000009621 | 40,64414331 | 42,7 | 3,7 | 3,792682033 | 0,044173727 | Vav2 | vav 2 oncogene [Source:MGI Symbol;Acc:MGI:102718] | 70,07 | 0,8 |
| ENSMUSG00000014932 | 238,2456344 | 305,3 | 106,9 | 3,798175744 | 0,041328052 | Yes1 | Yamaguchi sarcoma viral (v-yes) oncogene homolog 1 [Source:MGI Symbol;Acc:MGI:99147] | 82,04 | 11,3 |
| ENSMUSG00000039477 | 47,82053387 | 82,0 | 18,8 | 3,798630918 | 0,016560862 | Tnrc18 | trinucleotide repeat containing 18 [Source:MGI Symbol;Acc:MGI:3648294] | 59,99 | 3,6 |
| ENSMUSG00000041935 | 596,8820503 | 898,3 | 157,6 | 3,800317794 | 0,008682778 | AW549877 | expressed sequence AW549877 [Source:MGI Symbol;Acc:MGI:2146232] | 99,85 | 25,9 |
| ENSMUSG00000046707 | 71,27532716 | 71,8 | 22,6 | 3,800616183 | 0,014677393 | Csnk2a2 | casein kinase 2, alpha prime polypeptide [Source:MGI Symbol;Acc:MGI:88547] | 44,28 | 3,3 |
| ENSMUSG00000040711 | 72,44174502 | 142,0 | 27,2 | 3,803563964 | 0,021434724 | Sh3pxd2b | SH3 and PX domains 2B [Source:MGI Symbol;Acc:MGI:2442062] | 89,68 | 3,4 |
| ENSMUSG00000030935 | 57,23367678 | 38,1 | 3,9 | 3,804734862 | 0,018627369 | Acsm3 | acyl-CoA synthetase medium-chain family member 3 [Source:MGI Symbol;Acc:MGI:99538] | 29,96 | 1,6 |
| ENSMUSG00000014837 | 64,1137573 | 98,3 | 7,8 | 3,812396848 | 0,008993454 | 4931428F04Rik | RIKEN cDNA 4931428F04 gene [Source:MGI Symbol;Acc:MGI:1921606] | 93,59 | 2,9 |
| ENSMUSG00000000266 | 173,3926895 | 291,7 | 88,6 | 3,825446844 | 0,039826269 | Mid2 | midline 2 [Source:MGI Symbol;Acc:MGI:1344333] | 95,31 | 9,5 |
| ENSMUSG00000033306 | 58,50232479 | 115,4 | 11,6 | 3,8274545 | 0,026471687 | Lpp | LIM domain containing preferred translocation partner in lipoma [Source:MGI Symbol;Acc:MGI:2441849] | 50,86 | 0,8 |
| ENSMUSG00000028923 | 922,4617878 | 1432,4 | 184,4 | 3,83310751 | 0,046935289 | Necap2 | NECAP endocytosis associated 2 [Source:MGI Symbol;Acc:MGI:1913397] | 99,91 | 56,5 |
| ENSMUSG00000040537 | 95,03582868 | 134,8 | 76,9 | 3,839543979 | 0,039272304 | Adam22 | a disintegrin and metallopeptidase domain 22 [Source:MGI Symbol;Acc:MGI:1340046] | 72,74 | 4,6 |
| ENSMUSG00000060275 | 18,87343638 | 39,2 | 3,3 | 3,839765812 | 0,031958344 | Nrg2 | neuregulin 2 [Source:MGI Symbol;Acc:MGI:1098246] | 77,84 | 1,0 |
| ENSMUSG00000040536 | 57,86893742 | 103,9 | 34,8 | 3,840718895 | 0,044863632 | Necab1 | N-terminal EF-hand calcium binding protein 1 [Source:MGI Symbol;Acc:MGI:1916602] | 45,63 | 3,3 |
| ENSMUSG00000000876 | 379,8415621 | 662,2 | 62,0 | 3,842006462 | 0,005979142 | Pxmp4 | peroxisomal membrane protein 4 [Source:MGI Symbol;Acc:MGI:1891701] | 95,07 | 21,1 |
| ENSMUSG00000100826 | 31,60683023 | 48,1 | 7,9 | 3,846090801 | 0,018666759 | Snhg14 | small nucleolar RNA host gene 14 [Source:MGI Symbol;Acc:MGI:1289201] | 15,71 | 0,3 |
| ENSMUSG00000022332 | 653,3312119 | 991,3 | 169,2 | 3,84943369 | 0,026680113 | Khdrbs3 | I domain containing, RNA binding, signal transduction associated 3 [Source:MGI Symbol;Acc:MGI:13133 | 98,60 | 41,7 |
| ENSMUSG00000022718 | 72,91518764 | 96,1 | 96,2 | 3,855496346 | 0,034693621 | Dgcr8 | DiGeorge syndrome critical region gene 8 [Source:MGI Symbol;Acc:MGI:2151114] | 86,11 | 7,3 |
| ENSMUSG00000055320 | 44,78034646 | 84,0 | 7,6 | 3,85823013 | 0,029297578 | Tead1 | TEA domain family member 1 [Source:MGI Symbol;Acc:MGI:101876] | 55,72 | 0,9 |
| ENSMUSG00000032590 | 115,0950807 | 57,0 | 7,5 | 3,858661842 | 0,016135407 | Apeh | acylpeptide hydrolase [Source:MGI Symbol;Acc:MGI:88041] | 90,76 | 4,2 |
| ENSMUSG00000031295 | 24,73218591 | 31,1 | 2,9 | 3,860863327 | 0,039939641 | Phka2 | phosphorylase kinase alpha 2 [Source:MGI Symbol;Acc:MGI:97577] | 60,83 | 0,7 |
| ENSMUSG00000031519 | 10,02778679 | 22,3 | 1,6 | 3,866249136 | 0,046020341 | Asb5 | ankyrin repeat and SOCs box-containing 5 [Source:MGI Symbol;Acc:MGI:1923544] | 79,82 | 2,6 |
| ENSMUSG00000046085 | 15,0793196 | 27,3 | 5,5 | 3,871091627 | 0,043929689 | 4931422A03Rik | RIKEN cDNA 4931422A03 gene [Source:MGI Symbol;Acc:MGI:1918236] | 16,20 | 2,6 |
| ENSMUSG00000048170 | 151,709613 | 151,0 | 109,0 | 3,873472597 | 0,02830088 | Mcmbp | MCM (minichromosome maintenance deficient) binding protein [Source:MGI Symbol;Acc:MGI:1920977] | 94,01 | 23,3 |
| ENSMUSG00000070315 | 65,63716885 | 89,6 | 8,0 | 3,876051265 | 0,006895432 | 4930581F22Rik | RIKEN cDNA 4930581F22 gene [Source:MGI Symbol;Acc:MGI:1926184] | #DIV/0! | #DIV/0! |
| ENSMUSG00000079671 | 226,9426518 | 370,8 | 81,9 | 3,882875105 | 0,00764456 | 2610203C22Rik | RIKEN cDNA 2610203C22 gene [Source:MGI Symbol;Acc:MGI:1919731] | 11,99 | 0,7 |
| ENSMUSG00000022562 | 344,1642148 | 531,4 | 34,8 | 3,890070123 | 0,000263846 | Oplah | 5-oxoprolinase (ATP-hydrolysing) [Source:MGI Symbol;Acc:MGI:1922725] | 91,11 | 6,3 |
| ENSMUSG00000031568 | 375,1032714 | 606,7 | 122,3 | 3,896733059 | 0,03063669 | Rwdd4a | RWD domain containing 4A [Source:MGI Symbol;Acc:MGI:2681000] | 99,94 | 29,6 |
| ENSMUSG00000089774 | 89,24858197 | 123,4 | 11,4 | 3,897488597 | 0,007785233 | Slc5a3 | solute carrier family 5 (inositol transporters), member 3 [Source:MGI Symbol;Acc:MGI:1858226] | 34,05 | 1,3 |
| ENSMUSG00000027327 | 196,9707613 | 277,5 | 124,4 | 3,898970038 | 0,039083391 | 1700037H04Rik | RIKEN cDNA 1700037H04 gene [Source:MGI Symbol;Acc:MGI:1914576] | 90,20 | 34,0 |
| ENSMUSG00000026489 | 108,1256577 | 144,0 | 33,2 | 3,899838012 | 0,006712627 | Adck3 | aarF domain containing kinase 3 [Source:MGI Symbol;Acc:MGI:1914676] | 67,01 | 9,1 |
| ENSMUSG00000025666 | 5058,207393 | 8454,8 | 1975,7 | 3,902021586 | 0,034693621 | Tmem47 | transmembrane protein 47 [Source:MGI Symbol;Acc:MGI:2177570] | 100,00 | 66,3 |
| ENSMUSG00000025466 | 372,0473977 | 597,2 | 90,5 | 3,906383826 | 0,026432656 | Fuom | fucose mutarotase [Source:MGI Symbol;Acc:MGI:1916314] | 69,36 | 22,2 |
| ENSMUSG00000074797 | 186,9406056 | 266,6 | 28,8 | 3,907044121 | 0,020886638 | Itpa | inosine triphosphatase (nucleoside triphosphate pyrophosphatase) [Source:MGI Symbol;Acc:MGI:96622] | 85,08 | 10,6 |
| ENSMUSG00000027168 | 292,511282 | 493,2 | 78,7 | 3,909875429 | 0,004750039 | Pax6 | paired box 6 [Source:MGI Symbol;Acc:MGI:97490] | 83,51 | 8,1 |
| ENSMUSG00000041949 | 33,74170409 | 53,9 | 4,8 | 3,915641453 | 0,042667317 | Tango6 | transport and golgi organization 6 [Source:MGI Symbol;Acc:MGI:2142786] | 82,36 | 1,2 |
| ENSMUSG00000019338 | 103,8036103 | 87,7 | 49,1 | 3,916058767 | 0,040378847 | Zfp687 | zinc finger protein 687 [Source:MGI Symbol;Acc:MGI:1925516] | 88,62 | 5,8 |
| ENSMUSG00000031782 | 739,7624998 | 1224,2 | 215,4 | 3,917082161 | 0,048045881 | Coq9 | coenzyme Q9 [Source:MGI Symbol;Acc:MGI:1915164] | 98,36 | 57,4 |
| ENSMUSG00000018209 | 245,4916135 | 260,7 | 170,6 | 3,918178646 | 0,012135124 | Stk4 | serine/threonine kinase 4 [Source:MGI Symbol;Acc:MGI:1929004] | 92,09 | 15,4 |
| ENSMUSG00000031879 | 316,3364487 | 368,3 | 70,0 | 3,920793747 | 0,005839703 | Fam96b | family with sequence similarity 96, member B [Source:MGI Symbol;Acc:MGI:1915773] | 77,39 | 40,5 |
| ENSMUSG00000035585 | 400,6338273 | 682,1 | 145,6 | 3,92325707 | 0,026432771 | Tsen34 | tRNA splicing endonuclease 34 homolog (S. cerevisiae) [Source:MGI Symbol;Acc:MGI:1913328] | 89,44 | 33,2 |
| ENSMUSG00000021222 | 147,8748953 | 218,0 | 84,4 | 3,923937106 | 0,041732256 | Dcaf4 | DDB1 and CUL4 associated factor 4 [Source:MGI Symbol;Acc:MGI:1921078] | 99,76 | 33,1 |
| ENSMUSG00000041540 | 132,7356007 | 193,2 | 57,2 | 3,924175044 | 0,038686613 | Sox5 | SRY (sex determining region Y)-box 5 [Source:MGI Symbol;Acc:MGI:98367] | 65,80 | 8,7 |
| ENSMUSG00000051375 | 25,48545181 | 42,0 | 3,6 | 3,92596757 | 0,011318875 | Pcdh1 | protocadherin 1 [Source:MGI Symbol;Acc:MGI:104692] | 42,30 | 0,9 |
| ENSMUSG00000106943 | 60,64564276 | 30,4 | 8,3 | 3,930386793 | 0,049462071 | Dancr | differentiation antagonizing non-protein coding RNA [Source:MGI Symbol;Acc:MGI:1917286] | #DIV/0! | #DIV/0! |
| ENSMUSG00000032355 | 52,24281948 | 29,7 | 3,6 | 3,931235311 | 0,043042394 | Mlip | muscular LMNA-interacting protein [Source:MGI Symbol;Acc:MGI:1916892] | 58,21 | 0,7 |
| ENSMUSG00000053656 | 60,60898698 | 30,4 | 8,3 | 3,931764622 | 0,049495484 | Dancr | differentiation antagonizing non-protein coding RNA [Source:MGI Symbol;Acc:MGI:1917286] | #DIV/0! | #DIV/0! |
| ENSMUSG00000017734 | 711,9491356 | 1079,1 | 150,5 | 3,936562623 | 0,036438839 | Dbndd2 | dysbindin (dystrobrevin binding protein 1) domain containing 2 [Source:MGI Symbol;Acc:MGI:106562] | 80,00 | 18,0 |
| ENSMUSG00000022055 | 342,6120833 | 507,1 | 263,9 | 3,943360753 | 0,020000617 | Nefl | neurofilament, light polypeptide [Source:MGI Symbol;Acc:MGI:97313] | 99,14 | 32,0 |
| ENSMUSG00000033610 | 312,383273 | 549,2 | 173,0 | 3,94370195 | 0,032722538 | Pank1 | pantothenate kinase 1 [Source:MGI Symbol;Acc:MGI:1922985] | 98,25 | 19,7 |
| ENSMUSG00000006649 | 22,43855829 | 31,3 | 2,5 | 3,945054499 | 0,043707608 | Nphs1 | nephrosis 1, nephrin [Source:MGI Symbol;Acc:MGI:1859637] | 42,56 | 0,1 |
| ENSMUSG00000056476 | 26,20895416 | 61,3 | 3,5 | 3,950083346 | 0,015498573 | Med12l | mediator complex subunit 12-like [Source:MGI Symbol;Acc:MGI:2139916] | 44,66 | 0,4 |
| ENSMUSG00000002393 | 281,9106134 | 343,1 | 47,9 | 3,951170531 | 0,012570572 | Nr2f6 | nuclear receptor subfamily 2, group F, member 6 [Source:MGI Symbol;Acc:MGI:1352453] | 96,61 | 22,7 |
| ENSMUSG00000086976 | 19,01169611 | 31,0 | 4,6 | 3,952783939 | 0,021369444 | Gm12316 | predicted gene 12316 [Source:MGI Symbol;Acc:MGI:3651760] | 54,52 | 15,3 |
| ENSMUSG00000020166 | 335,4892113 | 478,1 | 118,6 | 3,954795592 | 0,010788277 | Cnot2 | CCR4-NOT transcription complex, subunit 2 [Source:MGI Symbol;Acc:MGI:1919318] | 83,57 | 24,8 |
| ENSMUSG00000026842 | 127,3940675 | 95,8 | 191,0 | 3,957841149 | 0,032150585 | Abl1 | c-abl oncogene 1, non-receptor tyrosine kinase [Source:MGI Symbol;Acc:MGI:87859] | 77,32 | 11,1 |
| ENSMUSG00000052139 | 129,9975742 | 198,6 | 14,3 | 3,960405933 | 0,016615332 | Bre | brain and reproductive organ-expressed protein [Source:MGI Symbol;Acc:MGI:1333875] | 76,45 | 8,6 |
| ENSMUSG00000020794 | 671,2508399 | 916,0 | 262,2 | 3,963150889 | 0,019734779 | Ube2g1 | ubiquitin-conjugating enzyme E2G 1 [Source:MGI Symbol;Acc:MGI:1914378] | 99,58 | 21,2 |
| ENSMUSG00000021022 | 180,3476162 | 286,0 | 54,0 | 3,964871021 | 0,01192108 | Ppp2r3c | protein phosphatase 2, regulatory subunit B", gamma [Source:MGI Symbol;Acc:MGI:1930009] | 89,39 | 16,1 |
| ENSMUSG00000041923 | 148,7504854 | 228,1 | 43,3 | 3,96933919 | 0,030440978 | Nol4 | nucleolar protein 4 [Source:MGI Symbol;Acc:MGI:2441684] | 93,74 | 7,6 |
| ENSMUSG00000086237 | 53,16480177 | 75,3 | 7,8 | 3,971469621 | 0,017525943 | Gm15591 | predicted gene 15591 [Source:MGI Symbol;Acc:MGI:3783038] | 22,28 | 10,7 |
| ENSMUSG00000019803 | 625,0499056 | 997,3 | 362,4 | 3,973994495 | 0,027719879 | Nr2e1 | nuclear receptor subfamily 2, group E, member 1 [Source:MGI Symbol;Acc:MGI:1100526] | 91,57 | 36,9 |
| ENSMUSG00000048787 | 62,39737413 | 65,7 | 4,4 | 3,974982834 | 0,005598936 | Dcun1d3 | L, defective in cullin neddylation 1, domain containing 3 (S. cerevisiae) [Source:MGI Symbol;Acc:MGI:267 | 59,01 | 0,9 |
| ENSMUSG00000035354 | 147,2481282 | 161,9 | 46,6 | 3,975409852 | 0,044124433 | Uvrag | UV radiation resistance associated gene [Source:MGI Symbol;Acc:MGI:1925860] | 80,91 | 5,7 |
| ENSMUSG00000066415 | 249,2505243 | 201,0 | 21,7 | 3,978779487 | 0,044517252 | Msl2 | male-specific lethal 2 homolog (Drosophila) [Source:MGI Symbol;Acc:MGI:1925103] | 87,55 | 4,3 |
| ENSMUSG00000032413 | 501,8604574 | 763,4 | 108,3 | 3,981208163 | 0,025391383 | Rasa2 | RAS p21 protein activator 2 [Source:MGI Symbol;Acc:MGI:2149960] | 99,32 | 9,3 |
| ENSMUSG00000020733 | 2595,653112 | 4739,3 | 385,4 | 3,981956606 | 3,84838E-05 | Slc9a3r1 | e carrier family 9 (sodium/hydrogen exchanger), member 3 regulator 1 [Source:MGI Symbol;Acc:MGI:134 | 100,00 | 85,4 |
| ENSMUSG00000053297 | 51,11578555 | 97,5 | 55,1 | 3,987610145 | 0,04116795 | AI854703 | expressed sequence AI854703 [Source:MGI Symbol;Acc:MGI:2141510] | 72,76 | 11,2 |
| ENSMUSG00000002396 | 202,7651734 | 238,6 | 33,3 | 3,99434985 | 0,016756908 | Ocel1 | occludin/ELL domain containing 1 [Source:MGI Symbol;Acc:MGI:1924340] | 52,40 | 4,4 |
| ENSMUSG00000052593 | 159,1396954 | 195,6 | 25,0 | 3,995027027 | 0,030177101 | Adam17 | a disintegrin and metallopeptidase domain 17 [Source:MGI Symbol;Acc:MGI:1096335] | 97,59 | 4,2 |
| ENSMUSG00000028576 | 350,1474769 | 575,5 | 72,1 | 3,995197818 | 0,019258586 | Ift74 | intraflagellar transport 74 [Source:MGI Symbol;Acc:MGI:1914944] | 82,16 | 22,7 |

| | | | | | | | | | |
|---|---|---|---|---|---|---|---|---|---|
| ENSMUSG00000049044 | 702,8718037 | 288,0 | 138,7 | 4,002724635 | 0,040678949 | Rapgef4 | Rap guanine nucleotide exchange factor (GEF) 4 [Source:MGI Symbol;Acc:MGI:1917723] | 92,90 | 26,6 |
| ENSMUSG00000038010 | 53,26930972 | 99,4 | 29,4 | 4,003615504 | 0,023150736 | Ccdc138 | coiled-coil domain containing 138 [Source:MGI Symbol;Acc:MGI:1923388] | 98,93 | 14,0 |
| ENSMUSG00000048834 | 77,35089889 | 124,8 | 17,1 | 4,005205764 | 0,01862641 | Vstm2a | V-set and transmembrane domain containing 2A [Source:MGI Symbol;Acc:MGI:2384826] | 72,86 | 4,1 |
| ENSMUSG00000022711 | 173,0522702 | 263,1 | 114,0 | 4,005246849 | 0,039966556 | Pmm2 | phosphomannomutase 2 [Source:MGI Symbol;Acc:MGI:1859214] | 98,59 | 31,5 |
| ENSMUSG00000062908 | 807,8869061 | 1151,0 | 144,5 | 4,007039645 | 0,016615332 | Acadm | acyl-Coenzyme A dehydrogenase, medium chain [Source:MGI Symbol;Acc:MGI:87867] | 94,62 | 37,5 |
| ENSMUSG00000022353 | 38,79792658 | 50,7 | 3,1 | 4,009515884 | 0,048601587 | Mtss1 | metastasis suppressor 1 [Source:MGI Symbol;Acc:MGI:2384818] | 68,47 | 1,2 |
| ENSMUSG00000027652 | 109,5308204 | 161,4 | 13,0 | 4,010912444 | 0,0095525 | Ralgapb | Ral GTPase activating protein, beta subunit (non-catalytic) [Source:MGI Symbol;Acc:MGI:2444531] | 86,76 | 1,1 |
| ENSMUSG00000026173 | 401,6669988 | 651,1 | 332,7 | 4,011660005 | 0,038651503 | Plcd4 | phospholipase C, delta 4 [Source:MGI Symbol;Acc:MGI:107469] | 87,92 | 20,9 |
| ENSMUSG00000040276 | 130,5648103 | 147,8 | 12,6 | 4,012394075 | 0,005087538 | Pacsin1 | protein kinase C and casein kinase substrate in neurons 1 [Source:MGI Symbol;Acc:MGI:1345181] | 59,48 | 5,3 |
| ENSMUSG00000040177 | 97,87108307 | 90,0 | 130,0 | 4,014109665 | 0,040959203 | 2310057M21Rik | RIKEN cDNA 2310057M21 gene [Source:MGI Symbol;Acc:MGI:1915527] | 37,86 | 14,4 |
| ENSMUSG00000050213 | 99,34007873 | 95,2 | 27,1 | 4,014857838 | 0,018168706 | Snip1 | Smad nuclear interacting protein 1 [Source:MGI Symbol;Acc:MGI:2156003] | 98,96 | 10,6 |
| ENSMUSG00000025892 | 13,99967378 | 19,3 | 2,0 | 4,014991836 | 0,043233 | Gria4 | glutamate receptor, ionotropic, AMPA4 (alpha 4) [Source:MGI Symbol;Acc:MGI:95811] | 28,80 | 0,4 |
| ENSMUSG00000089940 | 21,20443276 | 46,8 | 4,9 | 4,015992266 | 0,027123079 | Gm4117 | predicted gene 4117 [Source:MGI Symbol;Acc:MGI:3782293] | #DIV/0! | #DIV/0! |
| ENSMUSG00000028034 | 280,5593255 | 298,6 | 334,3 | 4,017260509 | 0,037371421 | Fubp1 | far upstream element (FUSE) binding protein 1 [Source:MGI Symbol;Acc:MGI:1196294] | 76,88 | 16,0 |
| ENSMUSG00000036503 | 1121,304896 | 1778,8 | 230,9 | 4,018600589 | 0,000755551 | Rnf13 | ring finger protein 13 [Source:MGI Symbol;Acc:MGI:1346341] | 76,85 | 25,9 |
| ENSMUSG00000008489 | 18,04602718 | 32,0 | 2,2 | 4,021335745 | 0,031168192 | Elavl2 | / (embryonic lethal, abnormal vision, Drosophila)-like 2 (Hu antigen B) [Source:MGI Symbol;Acc:MGI:110( | 50,84 | 0,8 |
| ENSMUSG00000025899 | 128,7021941 | 175,0 | 46,6 | 4,02205221 | 0,036806912 | Alkbh8 | alkB homolog 8, tRNA methyltransferase [Source:MGI Symbol;Acc:MGI:1914917] | 95,85 | 7,7 |
| ENSMUSG00000031565 | 520,5452656 | 905,4 | 68,6 | 4,023382579 | 0,005279836 | Fgfr1 | fibroblast growth factor receptor 1 [Source:MGI Symbol;Acc:MGI:95522] | 90,65 | 17,5 |
| ENSMUSG00000029610 | 206,9262596 | 250,3 | 59,7 | 4,027206288 | 0,020740744 | Aimp2 | noacyl tRNA synthetase complex-interacting multifunctional protein 2 [Source:MGI Symbol;Acc:MGI:2385 | 99,39 | 31,0 |
| ENSMUSG00000009739 | 150,5928701 | 158,2 | 57,7 | 4,028844622 | 0,037355635 | Pou6f1 | POU domain, class 6, transcription factor 1 [Source:MGI Symbol;Acc:MGI:102935] | 77,53 | 8,0 |
| ENSMUSG00000003452 | 122,458854 | 166,2 | 39,0 | 4,029135368 | 0,015934582 | Bicd1 | bicaudal D homolog 1 (Drosophila) [Source:MGI Symbol;Acc:MGI:1101760] | 78,80 | 4,7 |
| ENSMUSG00000034401 | 126,0176841 | 99,7 | 147,5 | 4,034850379 | 0,047028333 | Spata6 | spermatogenesis associated 6 [Source:MGI Symbol;Acc:MGI:1915196] | 69,37 | 23,3 |
| ENSMUSG00000029178 | 272,3631596 | 451,8 | 119,7 | 4,036126148 | 0,016333387 | Klf3 | Kruppel-like factor 3 (basic) [Source:MGI Symbol;Acc:MGI:1342773] | 94,33 | 16,5 |
| ENSMUSG00000021796 | 829,1429023 | 959,6 | 92,6 | 4,037759421 | 0,0347406 | Bmpr1a | bone morphogenetic protein receptor, type 1A [Source:MGI Symbol;Acc:MGI:1338938] | 92,38 | 6,8 |
| ENSMUSG00000005882 | 376,0408137 | 587,3 | 54,8 | 4,038164113 | 0,009961272 | Uqcc1 | ubiquinol-cytochrome c reductase complex assembly factor 1 [Source:MGI Symbol;Acc:MGI:1929472] | 91,25 | 20,0 |
| ENSMUSG00000009394 | 367,3098017 | 542,3 | 100,6 | 4,040948312 | 0,018744531 | Syn2 | synapsin II [Source:MGI Symbol;Acc:MGI:103020] | 98,84 | 16,6 |
| ENSMUSG00000051133 | 24,23837934 | 37,5 | 6,1 | 4,041166625 | 0,029414886 | Gm6293 | predicted pseudogene 6293 [Source:MGI Symbol;Acc:MGI:3649011] | #DIV/0! | #DIV/0! |
| ENSMUSG00000057716 | 23,50350324 | 42,4 | 2,9 | 4,04683272 | 0,017837685 | Tmem178b | transmembrane protein 178B [Source:MGI Symbol;Acc:MGI:3647581] | 24,75 | 0,7 |
| ENSMUSG00000035403 | 30,80772684 | 43,2 | 2,8 | 4,050044806 | 0,01036577 | Crb2 | crumbs family member 2 [Source:MGI Symbol;Acc:MGI:2679260] | 44,79 | 0,5 |
| ENSMUSG00000107846 | 34,52174809 | 70,6 | 6,4 | 4,05317257 | 0,016320434 | Gm43963 | predicted gene, 43963 [Source:MGI Symbol;Acc:MGI:5690355] | 8,94 | 2,1 |
| ENSMUSG00000024897 | 33,87233665 | 60,7 | 6,6 | 4,061001858 | 0,005912613 | Apba1 | amyloid beta (A4) precursor protein binding, family A, member 1 [Source:MGI Symbol;Acc:MGI:1860297] | 77,29 | 0,7 |
| ENSMUSG00000044628 | 182,7693058 | 347,4 | 59,2 | 4,062884812 | 0,031115635 | Rnf208 | ring finger protein 208 [Source:MGI Symbol;Acc:MGI:1916096] | 99,43 | 29,8 |
| ENSMUSG00000007812 | 171,7507879 | 239,0 | 83,8 | 4,06508176 | 0,030939401 | Zfp655 | zinc finger protein 655 [Source:MGI Symbol;Acc:MGI:1919861] | 96,21 | 10,2 |
| ENSMUSG00000028282 | 106,8868326 | 73,1 | 66,2 | 4,066817283 | 0,042914633 | Casp8ap2 | caspase 8 associated protein 2 [Source:MGI Symbol;Acc:MGI:1349399] | 59,83 | 7,5 |
| ENSMUSG00000031485 | 455,0666049 | 658,8 | 230,9 | 4,069753848 | 0,032949239 | Prosc | proline synthetase co-transcribed [Source:MGI Symbol;Acc:MGI:1891207] | 70,34 | 16,2 |
| ENSMUSG00000002428 | 68,57242991 | 48,0 | 3,3 | 4,071170901 | 0,035625893 | Hltf | helicase-like transcription factor [Source:MGI Symbol;Acc:MGI:1196437] | 76,62 | 0,8 |
| ENSMUSG00000022947 | 180,5449187 | 322,5 | 42,9 | 4,072593191 | 0,047549521 | Cbr3 | carbonyl reductase 3 [Source:MGI Symbol;Acc:MGI:1309992] | 99,91 | 35,1 |
| ENSMUSG00000021448 | 80,30215214 | 105,5 | 83,9 | 4,07391137 | 0,048162859 | Shc3 | src homology 2 domain-containing transforming protein C3 [Source:MGI Symbol;Acc:MGI:106179] | 100,00 | 31,4 |
| ENSMUSG00000000711 | 248,0582706 | 373,8 | 111,4 | 4,075028561 | 0,003553886 | Rab5b | RAB5B, member RAS oncogene family [Source:MGI Symbol;Acc:MGI:105938] | 80,68 | 22,5 |
| ENSMUSG00000052656 | 197,6166394 | 334,4 | 41,5 | 4,076564995 | 0,037147528 | Rnf103 | ring finger protein 103 [Source:MGI Symbol;Acc:MGI:109483] | 94,70 | 8,2 |
| ENSMUSG00000027665 | 418,2734575 | 674,2 | 72,6 | 4,076634085 | 0,012371023 | Pik3ca | phosphatidylinositol 3-kinase, catalytic, alpha polypeptide [Source:MGI Symbol;Acc:MGI:1206581] | 78,78 | 4,2 |
| ENSMUSG00000048406 | 40,4109158 | 70,6 | 13,0 | 4,077488831 | 0,033897948 | B330016D10Rik | RIKEN cDNA B330016D10 gene [Source:MGI Symbol;Acc:MGI:2444063] | #DIV/0! | #DIV/0! |
| ENSMUSG00000035181 | 19,89893749 | 41,3 | 2,9 | 4,077543885 | 0,022284032 | Heatr5a | HEAT repeat containing 5A [Source:MGI Symbol;Acc:MGI:2444133] | 50,81 | 0,8 |
| ENSMUSG00000106951 | 21,40273568 | 34,8 | 3,6 | 4,078770816 | 0,022541113 | 5930430L01Rik | RIKEN cDNA 5930430L01 gene [Source:MGI Symbol;Acc:MGI:2443110] | #DIV/0! | #DIV/0! |
| ENSMUSG00000031347 | 691,0435734 | 1071,1 | 156,9 | 4,081195167 | 0,009302039 | Cetn2 | centrin 2 [Source:MGI Symbol;Acc:MGI:1347085] | 86,51 | 27,3 |
| ENSMUSG00000036949 | 845,8175333 | 1315,0 | 237,1 | 4,083614209 | 0,00706941 | Slc39a12 | solute carrier family 39 (zinc transporter), member 12 [Source:MGI Symbol;Acc:MGI:2139274] | 96,08 | 32,8 |
| ENSMUSG00000051149 | 129,5356422 | 138,0 | 42,3 | 4,0861624 | 0,017398599 | Adnp | activity-dependent neuroprotective protein [Source:MGI Symbol;Acc:MGI:1338758] | 85,02 | 7,7 |
| ENSMUSG00000095139 | 142,6115686 | 183,9 | 22,3 | 4,090083648 | 0,010172918 | Pou3f2 | POU domain, class 3, transcription factor 2 [Source:MGI Symbol;Acc:MGI:101895] | 88,53 | 2,6 |
| ENSMUSG00000001518 | 74,02844482 | 84,3 | 6,7 | 4,090441287 | 0,01023543 | Itfg2 | integrin alpha FG-GAP repeat containing 2 [Source:MGI Symbol;Acc:MGI:1915450] | 69,34 | 2,7 |
| ENSMUSG00000092006 | 73,36719451 | 72,7 | 20,6 | 4,091580054 | 0,021996124 | Gm17139 | predicted gene 17139 [Source:MGI Symbol;Acc:MGI:4937966] | 99,82 | 20,6 |
| ENSMUSG00000032017 | 27,50349511 | 34,9 | 2,1 | 4,099987595 | 0,040198466 | Grik4 | glutamate receptor, ionotropic, kainate 4 [Source:MGI Symbol;Acc:MGI:95817] | 45,04 | 0,8 |
| ENSMUSG00000021635 | 185,3811666 | 213,9 | 157,3 | 4,100122784 | 0,0202613 | Rad17 | RAD17 homolog (S. pombe) [Source:MGI Symbol;Acc:MGI:1333807] | 95,35 | 30,2 |
| ENSMUSG00000097280 | 760,6201532 | 1126,7 | 221,0 | 4,102923804 | 0,030440412 | AI849053 | expressed sequence AI849053 [Source:MGI Symbol;Acc:MGI:2139818] | #DIV/0! | #DIV/0! |
| ENSMUSG00000007987 | 391,3506562 | 605,1 | 60,6 | 4,104220322 | 0,015569151 | Ift22 | intraflagellar transport 22 [Source:MGI Symbol;Acc:MGI:1914536] | 98,30 | 15,1 |
| ENSMUSG00000093577 | 25,32141264 | 63,5 | 4,6 | 4,106351713 | 0,01066796 | Gm20632 | predicted gene 20632 [Source:MGI Symbol;Acc:MGI:5313079] | 69,84 | 3,3 |
| ENSMUSG00000032673 | 226,5688612 | 294,1 | 64,2 | 4,106545779 | 0,02387903 | Prorsd1 | prolyl-tRNA synthetase domain containing 1 [Source:MGI Symbol;Acc:MGI:1915189] | 50,81 | 16,5 |
| ENSMUSG00000024074 | 73,26554395 | 97,8 | 6,4 | 4,108186361 | 0,016261287 | Crim1 | cysteine rich transmembrane BMP regulator 1 (chordin like) [Source:MGI Symbol;Acc:MGI:1354756] | 87,52 | 1,8 |
| ENSMUSG00000051721 | 16,7220697 | 34,4 | 2,1 | 4,11531226 | 0,035764165 | BC068281 | cDNA sequence BC068281 [Source:MGI Symbol;Acc:MGI:3040699] | 51,55 | 1,1 |
| ENSMUSG00000043019 | 123,2651492 | 119,5 | 374,4 | 4,118514912 | 0,031441632 | Edem3 | ER degradation enhancer, mannosidase alpha-like 3 [Source:MGI Symbol;Acc:MGI:1914217] | 90,96 | 28,4 |
| ENSMUSG00000038393 | 242,1687534 | 291,1 | 26,8 | 4,119987141 | 0,00354734 | Txnip | thioredoxin interacting protein [Source:MGI Symbol;Acc:MGI:1889549] | 99,29 | 10,1 |
| ENSMUSG00000039100 | 164,7129271 | 153,7 | 9,0 | 4,122537157 | 0,005393236 | March6 | membrane-associated ring finger (C3HC4) 6 [Source:MGI Symbol;Acc:MGI:2442773] | 85,92 | 2,7 |
| ENSMUSG00000071646 | 298,8513334 | 381,8 | 151,8 | 4,12357802 | 0,022551392 | Mta2 | metastasis-associated gene family, member 2 [Source:MGI Symbol;Acc:MGI:1346340] | 99,80 | 32,8 |
| ENSMUSG00000026826 | 83,89106726 | 49,0 | 2,5 | 4,126757908 | 0,018872392 | Nr4a2 | nuclear receptor subfamily 4, group A, member 2 [Source:MGI Symbol;Acc:MGI:1352456] | 76,89 | 1,9 |
| ENSMUSG00000062785 | 51,98116225 | 37,5 | 2,9 | 4,127045735 | 0,048657589 | Kcnc3 | otassium voltage gated channel, Shaw-related subfamily, member 3 [Source:MGI Symbol;Acc:MGI:9666 | 52,53 | 1,1 |
| ENSMUSG00000022240 | 966,1032321 | 1685,6 | 341,3 | 4,127390369 | 0,013193418 | Ctnnd2 | catenin (cadherin associated protein), delta 2 [Source:MGI Symbol;Acc:MGI:1195966] | 98,79 | 31,3 |
| ENSMUSG00000096941 | 15,96590469 | 31,6 | 4,2 | 4,129041492 | 0,042763369 | Gm26720 | predicted gene, 26720 [Source:MGI Symbol;Acc:MGI:5477214] | #DIV/0! | #DIV/0! |
| ENSMUSG00000057469 | 855,7155963 | 968,0 | 407,9 | 4,131249221 | 0,002708293 | E2f6 | E2F transcription factor 6 [Source:MGI Symbol;Acc:MGI:1354159] | 100,00 | 37,6 |
| ENSMUSG00000049288 | 70,07510273 | 103,5 | 29,4 | 4,133262848 | 0,011875427 | Lix1l | Lix1-like [Source:MGI Symbol;Acc:MGI:3036267] | 85,12 | 7,0 |
| ENSMUSG00000034021 | 93,98217926 | 134,1 | 8,8 | 4,134756659 | 0,000247052 | Pds5b | DS5, regulator of cohesion maintenance, homolog B (S. cerevisiae) [Source:MGI Symbol;Acc:MGI:214094 | 85,36 | 1,7 |
| ENSMUSG00000032997 | 119,1017171 | 192,1 | 17,5 | 4,139694513 | 0,015131294 | Chpf | chondroitin polymerizing factor [Source:MGI Symbol;Acc:MGI:106576] | 97,70 | 5,1 |
| ENSMUSG00000029763 | 203,4293075 | 328,5 | 34,7 | 4,140255123 | 0,010922467 | Exoc4 | exocyst complex component 4 [Source:MGI Symbol;Acc:MGI:1096376] | 86,70 | 4,2 |
| ENSMUSG00000040410 | 100,3793718 | 187,0 | 19,5 | 4,140560141 | 0,030254323 | Fbxl4 | F-box and leucine-rich repeat protein 4 [Source:MGI Symbol;Acc:MGI:2140367] | 98,02 | 7,9 |
| ENSMUSG00000037369 | 47,99878569 | 96,1 | 6,2 | 4,141374071 | 0,005518094 | Kdm6a | lysine (K)-specific demethylase 6A [Source:MGI Symbol;Acc:MGI:1095419] | 82,52 | 1,9 |
| ENSMUSG00000085819 | 20,41764825 | 38,1 | 14,8 | 4,142878953 | 0,048596989 | Ube4bos1 | ubiquitination factor E4B, opposite strand 1 [Source:MGI Symbol;Acc:MGI:1925072] | 15,86 | 5,1 |
| ENSMUSG00000021113 | 95,43475342 | 190,6 | 21,3 | 4,143623024 | 0,003009476 | Snapc1 | small nuclear RNA activating complex, polypeptide 1 [Source:MGI Symbol;Acc:MGI:1922877] | 96,33 | 7,7 |
| ENSMUSG00000105203 | 86,53307393 | 147,3 | 31,5 | 4,144302257 | 0,012282057 | 4933428P19Rik | RIKEN cDNA 4933428P19 gene [Source:MGI Symbol;Acc:MGI:1918479] | 41,47 | 14,6 |
| ENSMUSG00000028139 | 102,0289699 | 108,0 | 59,6 | 4,147118366 | 0,024429058 | Riiad1 | ulatory subunit of type II PKA R-subunit (RIIa) domain containing 1 [Source:MGI Symbol;Acc:MGI:1913( | 73,23 | 23,8 |
| ENSMUSG00000057335 | 118,6438728 | 108,2 | 8,0 | 4,148111927 | 0,005058806 | Cep170 | centrosomal protein 170 [Source:MGI Symbol;Acc:MGI:1918348] | 63,06 | 1,8 |
| ENSMUSG00000022208 | 174,0122803 | 246,6 | 66,6 | 4,149300078 | 0,011576497 | Jph4 | junctophilin 4 [Source:MGI Symbol;Acc:MGI:2443113] | 86,42 | 13,1 |
| ENSMUSG00000023961 | 218,8711441 | 129,1 | 139,6 | 4,149827658 | 0,028133658 | Enpp4 | ectonucleotide pyrophosphatase/phosphodiesterase 4 [Source:MGI Symbol;Acc:MGI:2682634] | 89,84 | 11,8 |
| ENSMUSG00000106224 | 21,86294683 | 33,0 | 4,2 | 4,156936303 | 0,040594928 | Gm43823 | predicted gene 43823 [Source:MGI Symbol;Acc:MGI:5663960] | 69,84 | 1,2 |
| ENSMUSG00000032932 | 91,46257174 | 109,2 | 15,9 | 4,158597845 | 0,048189025 | Hspa13 | heat shock protein 70 family, member 13 [Source:MGI Symbol;Acc:MGI:1309463] | 94,64 | 2,7 |
| ENSMUSG00000038451 | 25,04492681 | 39,9 | 2,9 | 4,160173077 | 0,01521912 | Spsb2 | splA/ryanodine receptor domain and SOCS box containing 2 [Source:MGI Symbol;Acc:MGI:1315199] | 75,57 | 0,3 |
| ENSMUSG00000106978 | 9,162248695 | 24,6 | 3,0 | 4,163334933 | 0,044231766 | N4bp2os | NEDD4 binding protein 2, opposite strand [Source:MGI Symbol;Acc:MGI:1921484] | 14,31 | 3,1 |
| ENSMUSG00000002103 | 265,8450618 | 279,0 | 273,9 | 4,163431562 | 0,010179082 | Acp2 | acid phosphatase 2, lysosomal [Source:MGI Symbol;Acc:MGI:87882] | 98,76 | 28,6 |
| ENSMUSG00000026787 | 76,85719518 | 93,2 | 11,9 | 4,163731723 | 0,03214507 | Gad2 | glutamic acid decarboxylase 2 [Source:MGI Symbol;Acc:MGI:95634] | 82,14 | 1,9 |
| ENSMUSG00000019173 | 133,9052165 | 189,3 | 21,6 | 4,164181449 | 0,003008727 | Rab5c | RAB5C, member RAS oncogene family [Source:MGI Symbol;Acc:MGI:105306] | 84,91 | 11,0 |
| ENSMUSG00000013150 | 96,86083371 | 167,9 | 14,9 | 4,165475184 | 0,015929831 | Gfod2 | glucose-fructose oxidoreductase domain containing 2 [Source:MGI Symbol;Acc:MGI:1917825] | 65,92 | 4,2 |
| ENSMUSG00000019856 | 57,8471268 | 103,6 | 5,9 | 4,167952014 | 0,008966028 | Fam184a | family with sequence similarity 184, member A [Source:MGI Symbol;Acc:MGI:1923156] | 94,65 | 1,8 |
| ENSMUSG00000022428 | 504,7432095 | 836,1 | 83,7 | 4,169409615 | 0,006499377 | Cby1 | chibby homolog 1 (Drosophila) [Source:MGI Symbol;Acc:MGI:1920989] | 100,00 | 60,9 |
| ENSMUSG00000049106 | 206,3483556 | 143,3 | 143,5 | 4,172948078 | 0,032271384 | Dcaf5 | DDB1 and CUL4 associated factor 5 [Source:MGI Symbol;Acc:MGI:2444785] | 93,09 | 19,5 |
| ENSMUSG00000038564 | 52,2178901 | 61,2 | 3,1 | 4,178560839 | 0,010577991 | Ift172 | intraflagellar transport 172 [Source:MGI Symbol;Acc:MGI:2682064] | 75,86 | 1,3 |
| ENSMUSG00000047409 | 152,4006535 | 276,2 | 20,6 | 4,181975697 | 0,000458034 | Ctdspl | y-terminal domain, RNA polymerase II, polypeptide A) small phosphatase-like [Source:MGI Symbol;Acc:M | 97,56 | 3,5 |
| ENSMUSG00000000056 | 187,3061894 | 313,7 | 24,3 | 4,184766996 | 0,017985246 | Narf | nuclear prelamin A recognition factor [Source:MGI Symbol;Acc:MGI:1914858] | 82,91 | 3,6 |
| ENSMUSG00000034748 | 26,6546817 | 34,2 | 6,5 | 4,191752501 | 0,032042223 | Sirt6 | sirtuin 6 [Source:MGI Symbol;Acc:MGI:1354161] | 67,27 | 2,7 |
| ENSMUSG00000024140 | 517,6015488 | 832,8 | 168,4 | 4,192429595 | 0,014602 | Epas1 | endothelial PAS domain protein 1 [Source:MGI Symbol;Acc:MGI:109169] | 99,14 | 27,1 |
| ENSMUSG00000035769 | 10,33216399 | 26,1 | 2,4 | 4,195763738 | 0,048006925 | Xylb | xylulokinase homolog (H. influenzae) [Source:MGI Symbol;Acc:MGI:2142985] | 62,95 | 1,1 |
| ENSMUSG00000075289 | 23,83085932 | 40,8 | 27,7 | 4,196329769 | 0,0341018 | Carns1 | carnosine synthase 1 [Source:MGI Symbol;Acc:MGI:2147595] | 66,50 | 6,3 |
| ENSMUSG00000003585 | 383,2636972 | 589,5 | 79,0 | 4,197100972 | 0,031195337 | Sec14l2 | SEC14-like lipid binding 2 [Source:MGI Symbol;Acc:MGI:1915065] | 98,75 | 28,5 |
| ENSMUSG00000045210 | 172,859437 | 217,0 | 185,1 | 4,203532304 | 0,030384029 | Vcpip1 | valosin containing protein (p97)/p47 complex interacting protein 1 [Source:MGI Symbol;Acc:MGI:1917925 | 73,88 | 9,5 |
| ENSMUSG00000086341 | 90,49052333 | 140,8 | 11,7 | 4,206944745 | 0,000900205 | Gm15932 | predicted gene 15932 [Source:MGI Symbol;Acc:MGI:3802106] | 34,90 | 18,6 |
| ENSMUSG00000050229 | 101,671466 | 61,8 | 5,6 | 4,210536727 | 0,024302188 | Pigm | phosphatidylinositol glycan anchor biosynthesis, class M [Source:MGI Symbol;Acc:MGI:1914806] | 41,98 | 0,6 |
| ENSMUSG00000020986 | 335,218255 | 460,4 | 250,2 | 4,211285888 | 0,02712276 | Sec23a | SEC23 homolog A, COPII coat complex component [Source:MGI Symbol;Acc:MGI:1349635] | 99,80 | 28,0 |
| ENSMUSG00000021182 | 64,75195983 | 36,6 | 2,2 | 4,211803813 | 0,049293996 | Ccdc88c | coiled-coil domain containing 88C [Source:MGI Symbol;Acc:MGI:1915589] | 53,83 | 0,5 |
| ENSMUSG00000038712 | 234,341901 | 363,9 | 84,2 | 4,212642431 | 0,019798401 | Fam63a | family with sequence similarity 63, member A [Source:MGI Symbol;Acc:MGI:1922257] | 92,86 | 5,3 |
| ENSMUSG00000038848 | 227,0041405 | 427,0 | 74,4 | 4,213475439 | 0,013627914 | Ythdf1 | YTH domain family 1 [Source:MGI Symbol;Acc:MGI:1917431] | 96,00 | 13,5 |
| ENSMUSG00000056812 | 107,8630504 | 60,1 | 60,1 | 4,215885487 | 0,038925983 | St8sia3 | ST8 alpha-N-acetyl-neuraminide alpha-2,8-sialyltransferase 3 [Source:MGI Symbol;Acc:MGI:106019] | 50,09 | 9,9 |
| ENSMUSG00000030443 | 20,99251975 | 40,9 | 17,0 | 4,218052486 | 0,035915426 | Zfp583 | zinc finger protein 583 [Source:MGI Symbol;Acc:MGI:2682297] | 81,84 | 3,1 |
| ENSMUSG00000039607 | 23,5242956 | 36,6 | 2,1 | 4,218246829 | 0,025347578 | Rbms3 | RNA binding motif, single stranded interacting protein [Source:MGI Symbol;Acc:MGI:2444477] | 48,73 | 0,6 |
| ENSMUSG00000030516 | 247,6972171 | 395,5 | 65,7 | 4,219537847 | 0,027397662 | Tjp1 | tight junction protein 1 [Source:MGI Symbol;Acc:MGI:98759] | 91,22 | 9,5 |
| ENSMUSG00000020599 | 16,71432196 | 30,7 | 1,5 | 4,22019162 | 0,04773446 | Rgs9 | regulator of G-protein signaling 9 [Source:MGI Symbol;Acc:MGI:1338824] | 43,77 | 0,6 |
| ENSMUSG00000014418 | 103,5416763 | 127,8 | 6,8 | 4,224052794 | 0,023952235 | Hps5 | Hermansky-Pudlak syndrome 5 [Source:MGI Symbol;Acc:MGI:2180307] | 94,75 | 2,2 |
| ENSMUSG00000038324 | 593,4117314 | 825,2 | 298,0 | 4,228417732 | 0,015231624 | Trpc4ap | eceptor potential cation channel, subfamily C, member 4 associated protein [Source:MGI Symbol;Acc:MGI | 98,92 | 37,0 |
| ENSMUSG00000032905 | 643,4864475 | 806,8 | 96,4 | 4,232918111 | 0,008088217 | Atg12 | autophagy related 12 [Source:MGI Symbol;Acc:MGI:1914776] | 99,76 | 30,4 |
| ENSMUSG00000041688 | 56,37087411 | 87,4 | 92,1 | 4,233481099 | 0,027533811 | Amot | angiomotin [Source:MGI Symbol;Acc:MGI:108440] | 72,34 | 10,5 |
| ENSMUSG00000030347 | 112,8191064 | 157,2 | 19,4 | 4,235083218 | 0,012526629 | D6Wsu163e | DNA segment, Chr 6, Wayne State University 163, expressed [Source:MGI Symbol;Acc:MGI:107893] | 90,35 | 4,0 |
| ENSMUSG00000039033 | 122,2268191 | 171,9 | 78,8 | 4,237336891 | 0,029137742 | Tasp1 | taspase, threonine aspartase 1 [Source:MGI Symbol;Acc:MGI:1923062] | 96,54 | 21,4 |
| ENSMUSG00000020495 | 29,3086367 | 59,0 | 3,2 | 4,237646675 | 0,01751344 | Smg8 | g-8 homolog, nonsense mediated mRNA decay factor (C. elegans) [Source:MGI Symbol;Acc:MGI:19213 | 88,37 | 2,3 |
| ENSMUSG00000055485 | 22,49849621 | 47,3 | 2,8 | 4,245970333 | 0,014224668 | Soga1 | suppressor of glucose, autophagy associated 1 [Source:MGI Symbol;Acc:MGI:2444575] | 62,39 | 0,6 |
| ENSMUSG00000036606 | 265,1375922 | 313,1 | 240,6 | 4,248644327 | 0,031245874 | Plxnb2 | plexin B2 [Source:MGI Symbol;Acc:MGI:2154239] | 95,94 | 22,4 |
| ENSMUSG00000028776 | 38,37143496 | 46,8 | 2,9 | 4,248677113 | 0,033603209 | Tinagl1 | tubulointerstitial nephritis antigen-like 1 [Source:MGI Symbol;Acc:MGI:2137617] | 71,72 | 0,1 |
| ENSMUSG00000071711 | 26,56064541 | 33,5 | 5,5 | 4,250318085 | 0,031837467 | Mpst | mercaptopyruvate sulfurtransferase [Source:MGI Symbol;Acc:MGI:2179733] | 92,51 | 5,7 |
| ENSMUSG00000045318 | 14,24573202 | 27,5 | 4,0 | 4,251593702 | 0,038574663 | Adra2c | adrenergic receptor, alpha 2c [Source:MGI Symbol;Acc:MGI:87936] | 62,07 | 2,1 |
| ENSMUSG00000087406 | 52,60783421 | 92,4 | 7,3 | 4,2524711 | 0,015957979 | E130215H24Rik | RIKEN cDNA E130215H24 gene [Source:MGI Symbol;Acc:MGI:1926091] | #DIV/0! | #DIV/0! |
| ENSMUSG00000070056 | 154,5526056 | 196,3 | 165,4 | 4,254292296 | 0,035427617 | Mfhas1 | malignant fibrous histiocytoma amplified sequence 1 [Source:MGI Symbol;Acc:MGI:1098644] | 94,78 | 16,2 |
| ENSMUSG00000032898 | 292,1814783 | 461,6 | 180,4 | 4,256127263 | 0,035777858 | Fbxo21 | F-box protein 21 [Source:MGI Symbol;Acc:MGI:1924223] | 99,06 | 28,0 |
| ENSMUSG00000025145 | 171,6057958 | 71,2 | 88,7 | 4,25718375 | 0,012275369 | Lrrc45 | leucine rich repeat containing 45 [Source:MGI Symbol;Acc:MGI:2387183] | 93,47 | 21,7 |
| ENSMUSG00000075514 | 23,78217058 | 46,1 | 2,6 | 4,259233788 | 0,024203883 | Gm13375 | predicted gene 13375 [Source:MGI Symbol;Acc:MGI:3649913] | 100,00 | 6,6 |
| ENSMUSG00000034480 | 46,13521822 | 80,1 | 38,8 | 4,260487837 | 0,033841736 | Diaph2 | diaphanous related formin 2 [Source:MGI Symbol;Acc:MGI:1858500] | 59,39 | 2,2 |
| ENSMUSG00000028444 | 319,3522002 | 520,3 | 76,0 | 4,261566469 | 0,009023159 | Cntfr | ciliary neurotrophic factor receptor [Source:MGI Symbol;Acc:MGI:99605] | 83,98 | 27,7 |
| ENSMUSG00000070476 | 102,6569913 | 146,1 | 17,1 | 4,262626777 | 0,012288115 | Fam217b | family with sequence similarity 217, member B [Source:MGI Symbol;Acc:MGI:1918782] | 72,83 | 5,1 |
| ENSMUSG00000021773 | 95,09258779 | 106,5 | 7,7 | 4,268897801 | 0,003758957 | Comtd1 | catechol-O-methyltransferase domain containing 1 [Source:MGI Symbol;Acc:MGI:1916406] | 63,80 | 0,2 |
| ENSMUSG00000062590 | 41,41546468 | 58,9 | 3,9 | 4,272015253 | 0,010179082 | Armc9 | armadillo repeat containing 9 [Source:MGI Symbol;Acc:MGI:1926045] | 67,75 | 0,7 |
| ENSMUSG00000027905 | 30,34004869 | 73,3 | 4,3 | 4,275565465 | 0,006773063 | Ddx20 | DEAD (Asp-Glu-Ala-Asp) box polypeptide 20 [Source:MGI Symbol;Acc:MGI:1858415] | 93,51 | 2,6 |
| ENSMUSG00000055200 | 57,18328812 | 45,8 | 17,6 | 4,278832566 | 0,032982686 | Sertad3 | SERTA domain containing 3 [Source:MGI Symbol;Acc:MGI:2180697] | 95,81 | 17,9 |
| ENSMUSG00000040651 | 63,7219993 | 108,4 | 8,1 | 4,285301713 | 0,005438495 | Fam208a | family with sequence similarity 208, member A [Source:MGI Symbol;Acc:MGI:1921694] | 79,20 | 1,7 |
| ENSMUSG00000021136 | 93,94726969 | 87,0 | 13,9 | 4,289859021 | 0,031441632 | Smoc1 | SPARC related modular calcium binding 1 [Source:MGI Symbol;Acc:MGI:1929878] | 89,75 | 4,7 |
| ENSMUSG00000046679 | 147,3538846 | 263,7 | 59,4 | 4,292184295 | 0,034003397 | C87436 | expressed sequence C87436 [Source:MGI Symbol;Acc:MGI:2141787] | 98,90 | 11,9 |
| ENSMUSG00000029534 | 85,00653332 | 114,9 | 12,7 | 4,295905474 | 0,029745314 | St7 | suppression of tumorigenicity 7 [Source:MGI Symbol;Acc:MGI:1927450] | 84,55 | 6,7 |
| ENSMUSG00000089824 | 146,0235147 | 159,9 | 17,1 | 4,296664917 | 0,018091557 | Rbm12 | RNA binding motif protein 12 [Source:MGI Symbol;Acc:MGI:1922960] | 60,53 | 2,2 |
| ENSMUSG00000022270 | 106,4614547 | 139,6 | 54,7 | 4,298361635 | 0,037512084 | Fam134b | family with sequence similarity 134, member B [Source:MGI Symbol;Acc:MGI:1913520] | 91,99 | 9,0 |
| ENSMUSG00000027806 | 73,36450219 | 114,1 | 6,1 | 4,299828513 | 0,004712445 | Tsc22d2 | TSC22 domain family, member 2 [Source:MGI Symbol;Acc:MGI:1919283] | 55,01 | 1,2 |
| ENSMUSG00000027187 | 794,9331423 | 1374,1 | 212,9 | 4,302970593 | 0,014342208 | Cat | catalase [Source:MGI Symbol;Acc:MGI:88271] | 95,99 | 26,5 |
| ENSMUSG00000029263 | 36,64757423 | 60,9 | 2,9 | 4,303558119 | 0,029730157 | Pigg | phosphatidylinositol glycan anchor biosynthesis, class G [Source:MGI Symbol;Acc:MGI:3576484] | 60,77 | 0,9 |
| ENSMUSG00000022607 | 314,2304779 | 517,9 | 38,9 | 4,306253034 | 0,002238453 | Ptk2 | PTK2 protein tyrosine kinase 2 [Source:MGI Symbol;Acc:MGI:95481] | 90,15 | 8,9 |
| ENSMUSG00000033684 | 37,97559939 | 71,7 | 4,0 | 4,308159326 | 0,007755903 | Qsox1 | quiescin Q6 sulfhydryl oxidase 1 [Source:MGI Symbol;Acc:MGI:1330818] | 84,73 | 0,0 |
| ENSMUSG00000036435 | 71,4124267 | 103,0 | 98,7 | 4,314319299 | 0,032967308 | Exoc1 | exocyst complex component 1 [Source:MGI Symbol;Acc:MGI:2445020] | 96,30 | 0,5 |
| ENSMUSG00000051451 | 259,7729984 | 187,1 | 265,4 | 4,31753803 | 0,016481368 | Crebzf | CREB/ATF bZIP transcription factor [Source:MGI Symbol;Acc:MGI:2675296] | 93,47 | 28,4 |
| ENSMUSG00000029782 | 112,9428522 | 206,2 | 72,6 | 4,318075534 | 0,009976306 | Tmem209 | transmembrane protein 209 [Source:MGI Symbol;Acc:MGI:1919899] | 96,14 | 4,8 |
| ENSMUSG00000035382 | 33,83832146 | 67,1 | 13,6 | 4,322697058 | 0,028556088 | Pcsk7 | proprotein convertase subtilisin/kexin type 7 [Source:MGI Symbol;Acc:MGI:107421] | 92,56 | 1,7 |
| ENSMUSG00000040720 | 50,46049091 | 59,3 | 4,7 | 4,326666955 | 0,028086723 | 1110037F02Rik | RIKEN cDNA 1110037F02 gene [Source:MGI Symbol;Acc:MGI:1913435] | 68,67 | 1,3 |
| ENSMUSG00000054252 | 1421,456376 | 2177,6 | 334,5 | 4,327974932 | 0,032460068 | Fgfr3 | fibroblast growth factor receptor 3 [Source:MGI Symbol;Acc:MGI:95524] | 95,18 | 34,0 |
| ENSMUSG00000027668 | 1082,5317 | 1643,4 | 208,5 | 4,331102778 | 8,21045E-05 | Mfn1 | mitofusin 1 [Source:MGI Symbol;Acc:MGI:1914664] | 83,49 | 20,0 |
| ENSMUSG00000025077 | 78,56970481 | 126,6 | 28,8 | 4,333609841 | 0,046024278 | Dclre1a | DNA cross-link repair 1A [Source:MGI Symbol;Acc:MGI:1930042] | 96,92 | 5,7 |
| ENSMUSG00000060371 | 46,46922351 | 88,5 | 4,4 | 4,335907693 | 0,006336387 | Caln1 | calneuron 1 [Source:MGI Symbol;Acc:MGI:2155987] | 33,19 | 0,8 |
| ENSMUSG00000030590 | 118,490918 | 91,0 | 24,8 | 4,343449377 | 0,003911784 | Fam98c | family with sequence similarity 98, member C [Source:MGI Symbol;Acc:MGI:1921083] | 95,02 | 25,2 |
| ENSMUSG00000022306 | 51,20912509 | 105,6 | 8,5 | 4,343849853 | 0,009373845 | Zfpm2 | zinc finger protein, multitype 2 [Source:MGI Symbol;Acc:MGI:1334444] | 91,32 | 2,2 |
| ENSMUSG00000042686 | 22,05967506 | 25,4 | 2,0 | 4,346550307 | 0,028969349 | Jph1 | junctophilin 1 [Source:MGI Symbol;Acc:MGI:1891495] | 58,16 | 0,8 |
| ENSMUSG00000047446 | 165,8675297 | 232,3 | 72,9 | 4,348010759 | 0,015231624 | Arl4a | ADP-ribosylation factor-like 4A [Source:MGI Symbol;Acc:MGI:99437] | 96,34 | 9,9 |
| ENSMUSG00000023072 | 100,8322801 | 50,6 | 86,9 | 4,349251517 | 0,030151954 | Cep89 | centrosomal protein 89 [Source:MGI Symbol;Acc:MGI:1919390] | 95,76 | 29,6 |
| ENSMUSG00000036810 | 395,1012214 | 616,4 | 107,3 | 4,351050999 | 0,014165371 | Cnep1r1 | CTD nuclear envelope phosphatase 1 regulatory subunit 1 [Source:MGI Symbol;Acc:MGI:1921981] | 69,91 | 16,6 |
| ENSMUSG00000034891 | 78,91130761 | 150,1 | 10,4 | 4,351150395 | 0,009091942 | Sncb | synuclein, beta [Source:MGI Symbol;Acc:MGI:1889011] | 86,03 | 12,3 |
| ENSMUSG00000107476 | 22,41935957 | 31,2 | 5,0 | 4,354149735 | 0,033106856 | Zfp862-ps | zinc finger protein 862, pseudogene [Source:MGI Symbol;Acc:MGI:1889827] | 48,06 | 0,0 |
| ENSMUSG00000045658 | 464,6370821 | 804,0 | 65,1 | 4,356277217 | 0,021434724 | Pid1 | phosphotyrosine interaction domain containing 1 [Source:MGI Symbol;Acc:MGI:2138391] | 43,81 | 2,2 |
| ENSMUSG00000022636 | 146,8240396 | 160,6 | 62,6 | 4,358134577 | 0,020597957 | Alcam | activated leukocyte cell adhesion molecule [Source:MGI Symbol;Acc:MGI:1313266] | 69,77 | 6,3 |
| ENSMUSG00000028613 | 257,7945443 | 257,4 | 21,5 | 4,35936185 | 0,013210643 | Lrp8 | lensity lipoprotein receptor-related protein 8, apolipoprotein e receptor [Source:MGI Symbol;Acc:MGI:134( | 94,34 | 2,4 |
| ENSMUSG00000074909 | 101,9394174 | 132,1 | 46,9 | 4,37107049 | 0,036404933 | Ranbp6 | RAN binding protein 6 [Source:MGI Symbol;Acc:MGI:2683212] | 94,96 | 7,3 |
| ENSMUSG00000008429 | 482,923573 | 846,3 | 135,1 | 4,371340023 | 0,015819883 | Herpud2 | HERPUD family member 2 [Source:MGI Symbol;Acc:MGI:1915393] | 100,00 | 18,8 |
| ENSMUSG00000029790 | 44,41474884 | 82,7 | 12,6 | 4,372164923 | 0,030723512 | Cep41 | centrosomal protein 41 [Source:MGI Symbol;Acc:MGI:1891414] | 85,21 | 3,4 |
| ENSMUSG00000048280 | 47,15341492 | 80,5 | 4,5 | 4,374157042 | 0,014619677 | Zfp738 | zinc finger protein 738 [Source:MGI Symbol;Acc:MGI:3040706] | 73,16 | 1,3 |

| | | | | | | | | | |
|---|---|---|---|---|---|---|---|---|---|
| ENSMUSG00000014763 | 192,773075 | 274,8 | 28,3 | 4,375894989 | 0,015013494 | Fam120b | family with sequence similarity 120, member B [Source:MGI Symbol;Acc:MGI:1914794] | 98,96 | 3,6 |
| ENSMUSG00000028675 | 1496,90851 | 2993,6 | 372,1 | 4,37713914 | 0,03340453 | Pnrc2 | proline-rich nuclear receptor coactivator 2 [Source:MGI Symbol;Acc:MGI:106512] | 100,00 | 54,5 |
| ENSMUSG00000043421 | 64,27930342 | 76,8 | 21,3 | 4,380160679 | 0,007999802 | Hilpda | hypoxia inducible lipid droplet associated [Source:MGI Symbol;Acc:MGI:1916823] | 87,94 | 19,8 |
| ENSMUSG00000022512 | 121,1420806 | 130,0 | 213,5 | 4,380423831 | 0,046113091 | Cldn1 | claudin 1 [Source:MGI Symbol;Acc:MGI:1276109] | 89,84 | 28,7 |
| ENSMUSG00000031153 | 209,0692533 | 280,4 | 21,2 | 4,382184567 | 0,007785817 | Gripap1 | GRIP1 associated protein 1 [Source:MGI Symbol;Acc:MGI:1859616] | 91,73 | 7,0 |
| ENSMUSG00000097622 | 523,9481019 | 1006,2 | 55,5 | 4,382783607 | 0,001689804 | A330033J07Rik | RIKEN cDNA A330033J07 gene [Source:MGI Symbol;Acc:MGI:2444369] | #DIV/0! | #DIV/0! |
| ENSMUSG00000006423 | 576,2405048 | 918,9 | 67,4 | 4,38351691 | 0,014738218 | C330007P06Rik | RIKEN cDNA C330007P06 gene [Source:MGI Symbol;Acc:MGI:1924894] | 75,09 | 10,5 |
| ENSMUSG00000059981 | 77,83364244 | 131,5 | 48,2 | 4,385528031 | 0,025831491 | Taok2 | TAO kinase 2 [Source:MGI Symbol;Acc:MGI:1915919] | 87,36 | 9,1 |
| ENSMUSG00000033166 | 169,9764968 | 107,7 | 24,0 | 4,386710832 | 0,029520617 | Dis3 | S3 homolog, exosome endoribonuclease and 3'-5' exoribonuclease [Source:MGI Symbol;Acc:MGI:191991 | 93,59 | 4,1 |
| ENSMUSG00000033159 | 170,5652976 | 160,8 | 73,3 | 4,388263079 | 0,009076555 | Cnppd1 | cyclin Pas1/PHO80 domain containing 1 [Source:MGI Symbol;Acc:MGI:1916421] | 68,47 | 17,6 |
| ENSMUSG00000037465 | 48,56015419 | 106,3 | 5,7 | 4,390596281 | 0,017689038 | Klf10 | Kruppel-like factor 10 [Source:MGI Symbol;Acc:MGI:1101353] | 97,25 | 2,0 |
| ENSMUSG00000049184 | 35,62343979 | 32,6 | 6,0 | 4,392101665 | 0,046107493 | Purg | purine-rich element binding protein G [Source:MGI Symbol;Acc:MGI:1922279] | 85,53 | 2,4 |
| ENSMUSG00000024797 | 155,9607074 | 210,0 | 35,4 | 4,396778807 | 0,006175831 | Vps51 | vacuolar protein sorting 51 homolog (S. cerevisiae) [Source:MGI Symbol;Acc:MGI:1915755] | 86,96 | 7,8 |
| ENSMUSG00000056367 | 294,0385074 | 455,7 | 92,5 | 4,397818737 | 0,017191018 | Actr3b | ARP3 actin-related protein 3B [Source:MGI Symbol;Acc:MGI:2661120] | 99,94 | 32,9 |
| ENSMUSG00000042121 | 46,09646236 | 52,6 | 2,7 | 4,399881223 | 0,010799495 | Ssh1 | slingshot homolog 1 (Drosophila) [Source:MGI Symbol;Acc:MGI:2686240] | 53,28 | 0,8 |
| ENSMUSG00000027709 | 257,6523312 | 286,7 | 180,4 | 4,400994823 | 0,025878717 | Mccc1 | methylcrotonoyl-Coenzyme A carboxylase 1 (alpha) [Source:MGI Symbol;Acc:MGI:1919289] | 99,96 | 33,0 |
| ENSMUSG00000106662 | 13,70964279 | 32,1 | 6,0 | 4,401501206 | 0,047690705 | Gm43034 | predicted gene 43034 [Source:MGI Symbol;Acc:MGI:5663171] | 93,77 | 4,5 |
| ENSMUSG00000026977 | 251,8650477 | 231,9 | 32,2 | 4,40222987 | 0,015863155 | March7 | membrane-associated ring finger (C3HC4) 7 [Source:MGI Symbol;Acc:MGI:1931053] | 96,59 | 4,0 |
| ENSMUSG00000019970 | 466,5309259 | 556,9 | 202,2 | 4,404179458 | 0,038497541 | Sgk1 | serum/glucocorticoid regulated kinase 1 [Source:MGI Symbol;Acc:MGI:1340062] | 78,21 | 20,9 |
| ENSMUSG00000026259 | 844,6142754 | 1235,4 | 261,6 | 4,405705429 | 0,043053311 | Ngef | neuronal guanine nucleotide exchange factor [Source:MGI Symbol;Acc:MGI:1858414] | 82,91 | 34,5 |
| ENSMUSG00000026797 | 370,9472849 | 488,6 | 130,3 | 4,405768771 | 0,024215367 | Stxbp1 | syntaxin binding protein 1 [Source:MGI Symbol;Acc:MGI:107363] | 96,69 | 5,9 |
| ENSMUSG00000024883 | 57,91601157 | 43,3 | 2,5 | 4,408107476 | 0,033965089 | Rin1 | Ras and Rab interactor 1 [Source:MGI Symbol;Acc:MGI:2385695] | 77,54 | 1,6 |
| ENSMUSG00000030094 | 110,3483212 | 120,4 | 9,9 | 4,409042688 | 0,009975811 | Xpc | xeroderma pigmentosum, complementation group C [Source:MGI Symbol;Acc:MGI:103557] | 98,44 | 3,7 |
| ENSMUSG00000032340 | 157,4937967 | 314,6 | 86,3 | 4,411800101 | 0,010769986 | Neo1 | neogenin [Source:MGI Symbol;Acc:MGI:1097159] | 98,49 | 5,6 |
| ENSMUSG00000038170 | 280,8077427 | 526,8 | 80,8 | 4,414207482 | 0,005219865 | Pde4dip | phosphodiesterase 4D interacting protein (myomegalin) [Source:MGI Symbol;Acc:MGI:1891434] | 78,82 | 4,5 |
| ENSMUSG00000028082 | 97,51467294 | 117,7 | 6,2 | 4,415593094 | 0,010341616 | Sh3d19 | SH3 domain protein D19 [Source:MGI Symbol;Acc:MGI:1350923] | 72,34 | 0,5 |
| ENSMUSG00000079508 | 189,8192271 | 290,7 | 45,5 | 4,42086111 | 0,005913701 | Apoo | apolipoprotein O [Source:MGI Symbol;Acc:MGI:1915566] | 66,51 | 20,2 |
| ENSMUSG00000085913 | 106,0343708 | 130,8 | 80,8 | 4,422606413 | 0,033353365 | Gm15601 | predicted gene 15601 [Source:MGI Symbol;Acc:MGI:3783048] | 21,34 | 5,6 |
| ENSMUSG00000042874 | 21,68733043 | 37,2 | 3,6 | 4,424188122 | 0,030699902 | D930007J09Rik | RIKEN cDNA D930007J09 gene [Source:MGI Symbol;Acc:MGI:3686989] | 100,00 | 7,5 |
| ENSMUSG00000012187 | 28,47446254 | 37,6 | 17,0 | 4,424324005 | 0,041784204 | Mogat1 | monoacylglycerol O-acyltransferase 1 [Source:MGI Symbol;Acc:MGI:1915643] | 17,97 | 5,7 |
| ENSMUSG00000001768 | 931,1079024 | 1541,2 | 250,2 | 4,426095014 | 0,00220446 | Rin2 | Ras and Rab interactor 2 [Source:MGI Symbol;Acc:MGI:1921280] | 93,09 | 23,2 |
| ENSMUSG00000017144 | 48,03057789 | 68,3 | 4,0 | 4,427540892 | 0,013896003 | Rnd3 | Rho family GTPase 3 [Source:MGI Symbol;Acc:MGI:1921444] | 62,27 | 1,0 |
| ENSMUSG00000015748 | 73,9905747 | 67,6 | 3,8 | 4,428151761 | 0,013120608 | Prpf3 | PRP3 pre-mRNA processing factor 3 homolog (yeast) [Source:MGI Symbol;Acc:MGI:1918017] | 69,21 | 1,4 |
| ENSMUSG00000051786 | 144,8915124 | 99,9 | 22,3 | 4,428516843 | 0,025533191 | Tubgcp6 | tubulin, gamma complex associated protein 6 [Source:MGI Symbol;Acc:MGI:2146071] | 59,39 | 3,3 |
| ENSMUSG00000032198 | 32,7236607 | 54,7 | 3,1 | 4,430182307 | 0,031958344 | Dock6 | dedicator of cytokinesis 6 [Source:MGI Symbol;Acc:MGI:1914789] | 29,35 | 0,6 |
| ENSMUSG00000048148 | 645,6487335 | 1013,9 | 91,9 | 4,430547884 | 0,047499556 | Nwd1 | NACHT and WD repeat domain containing 1 [Source:MGI Symbol;Acc:MGI:2442268] | 92,39 | 10,7 |
| ENSMUSG00000043881 | 68,93567898 | 115,9 | 23,7 | 4,431066815 | 0,013799306 | Kbtbd7 | kelch repeat and BTB (POZ) domain containing 7 [Source:MGI Symbol;Acc:MGI:2685141] | 80,30 | 3,3 |
| ENSMUSG00000043929 | 46,23799884 | 62,1 | 3,4 | 4,431281108 | 0,019003596 | Klhl15 | kelch-like 15 [Source:MGI Symbol;Acc:MGI:1923400] | 39,78 | 0,3 |
| ENSMUSG00000029544 | 28,79022695 | 45,8 | 25,9 | 4,432992005 | 0,032701694 | Cabp1 | calcium binding protein 1 [Source:MGI Symbol;Acc:MGI:1352750] | 69,51 | 9,9 |
| ENSMUSG00000025911 | 1586,347819 | 2611,2 | 470,9 | 4,433672572 | 0,018661386 | Adhfe1 | alcohol dehydrogenase, iron containing, 1 [Source:MGI Symbol;Acc:MGI:1923437] | 86,12 | 38,3 |
| ENSMUSG00000029726 | 108,136768 | 157,7 | 93,3 | 4,433871253 | 0,049788304 | Mepce | methylphosphate capping enzyme [Source:MGI Symbol;Acc:MGI:106477] | 87,42 | 19,9 |
| ENSMUSG00000028799 | 64,35187595 | 112,4 | 29,9 | 4,435608599 | 0,010956967 | Zfp362 | zinc finger protein 362 [Source:MGI Symbol;Acc:MGI:2652839] | 98,94 | 11,1 |
| ENSMUSG00000038828 | 95,87124249 | 143,4 | 66,9 | 4,435701714 | 0,013798776 | Tmem214 | transmembrane protein 214 [Source:MGI Symbol;Acc:MGI:1916046] | 48,18 | 9,7 |
| ENSMUSG00000061751 | 178,9419624 | 211,9 | 29,9 | 4,436519142 | 0,002746664 | Kalrn | kalirin, RhoGEF kinase [Source:MGI Symbol;Acc:MGI:2685385] | 59,17 | 1,5 |
| ENSMUSG00000021379 | 4031,702164 | 7929,5 | 465,2 | 4,43826935 | 0,000420757 | Id4 | inhibitor of DNA binding 4 [Source:MGI Symbol;Acc:MGI:99414] | 100,00 | 77,3 |
| ENSMUSG00000024926 | 184,2669243 | 267,0 | 24,5 | 4,438460982 | 0,010671413 | Kat5 | K(lysine) acetyltransferase 5 [Source:MGI Symbol;Acc:MGI:1932051] | 98,12 | 12,7 |
| ENSMUSG00000042745 | 366,7859004 | 635,1 | 110,7 | 4,441513155 | 0,033753697 | Id1 | inhibitor of DNA binding 1 [Source:MGI Symbol;Acc:MGI:96396] | 81,01 | 37,8 |
| ENSMUSG00000043602 | 92,09943251 | 151,4 | 22,8 | 4,442290306 | 0,036993453 | Zfp3 | zinc finger protein 3 [Source:MGI Symbol;Acc:MGI:99177] | 100,00 | 7,6 |
| ENSMUSG00000030934 | 1689,644553 | 2751,1 | 252,7 | 4,443483032 | 0,004375869 | Oat | ornithine aminotransferase [Source:MGI Symbol;Acc:MGI:97394] | 99,35 | 60,4 |
| ENSMUSG00000057440 | 74,05738021 | 83,8 | 27,5 | 4,445120452 | 0,04799095 | Mpp7 | mbrane protein, palmitoylated 7 (MAGUK p55 subfamily member 7) [Source:MGI Symbol;Acc:MGI:19229 | 88,64 | 3,2 |
| ENSMUSG00000095325 | 158,6586513 | 190,9 | 92,3 | 4,446566095 | 0,007209328 | Zfp870 | zinc finger protein 870 [Source:MGI Symbol;Acc:MGI:3029586] | 87,73 | 13,7 |
| ENSMUSG00000042834 | 334,8825172 | 398,7 | 440,1 | 4,453949839 | 0,014821484 | Nrep | neuronal regeneration related protein [Source:MGI Symbol;Acc:MGI:99444] | 88,55 | 34,3 |
| ENSMUSG00000047030 | 249,2397156 | 238,9 | 156,3 | 4,45469994 | 0,022541113 | Spata2 | spermatogenesis associated 2 [Source:MGI Symbol;Acc:MGI:2146885] | 91,36 | 26,7 |
| ENSMUSG00000090534 | 90,05900272 | 122,6 | 125,7 | 4,455105067 | 0,046488178 | Gm4675 | predicted gene 4675 [Source:MGI Symbol;Acc:MGI:3782856] | #DIV/0! | #DIV/0! |
| ENSMUSG00000040268 | 276,0937791 | 407,6 | 40,7 | 4,456871225 | 0,024779473 | Plekha1 | ology domain containing, family A (phosphoinositide binding specific) member 1 [Source:MGI Symbol;Acc | 74,79 | 7,3 |
| ENSMUSG00000100627 | 26,03247412 | 51,5 | 13,3 | 4,458422035 | 0,045559406 | A830008E24Rik | RIKEN cDNA A830008E24 gene [Source:MGI Symbol;Acc:MGI:3041183] | #DIV/0! | #DIV/0! |
| ENSMUSG00000027605 | 413,427811 | 618,6 | 33,9 | 4,458938397 | 0,009417009 | Acss2 | acyl-CoA synthetase short-chain family member 2 [Source:MGI Symbol;Acc:MGI:1890410] | 75,99 | 3,3 |
| ENSMUSG00000046201 | 67,91969352 | 118,4 | 5,3 | 4,461186302 | 0,014697605 | Scaf8 | SR-related CTD-associated factor 8 [Source:MGI Symbol;Acc:MGI:1925212] | 89,35 | 1,8 |
| ENSMUSG00000037098 | 153,7623496 | 255,5 | 49,6 | 4,461736419 | 0,041106794 | Rab11fip3 | RAB11 family interacting protein 3 (class II) [Source:MGI Symbol;Acc:MGI:2444431] | 92,14 | 8,4 |
| ENSMUSG00000031930 | 125,3442013 | 234,7 | 31,3 | 4,464361676 | 0,019292726 | Wwp2 | WW domain containing E3 ubiquitin protein ligase 2 [Source:MGI Symbol;Acc:MGI:1914144] | 99,90 | 7,3 |
| ENSMUSG00000031792 | 94,34268917 | 137,4 | 44,6 | 4,467662221 | 0,04891304 | Usb1 | U6 snRNA biogenesis 1 [Source:MGI Symbol;Acc:MGI:2142454] | 71,44 | 21,4 |
| ENSMUSG00000029376 | 345,7495603 | 491,0 | 36,3 | 4,470467474 | 0,035996626 | Mthfd2l | ethylenetetrahydrofolate dehydrogenase (NADP+ dependent) 2-like [Source:MGI Symbol;Acc:MGI:191587 | 94,08 | 12,5 |
| ENSMUSG00000003746 | 88,90652378 | 104,9 | 6,8 | 4,472361373 | 0,021890207 | Man1a | mannosidase 1, alpha [Source:MGI Symbol;Acc:MGI:104677] | 73,36 | 2,0 |
| ENSMUSG00000030613 | 730,0011595 | 1155,9 | 99,1 | 4,474191675 | 0,032419045 | Ccdc90b | coiled-coil domain containing 90B [Source:MGI Symbol;Acc:MGI:1913615] | 99,83 | 41,0 |
| ENSMUSG00000034663 | 148,5578321 | 191,6 | 92,2 | 4,47439889 | 0,016481368 | Bmp2k | BMP2 inducible kinase [Source:MGI Symbol;Acc:MGI:2155456] | 68,96 | 6,7 |
| ENSMUSG00000035392 | 216,8939791 | 340,6 | 43,5 | 4,474440736 | 0,004212449 | Dennd1a | DENN/MADD domain containing 1A [Source:MGI Symbol;Acc:MGI:2442794] | 98,72 | 14,0 |
| ENSMUSG00000039706 | 66,22090797 | 80,2 | 6,8 | 4,477497075 | 0,049693667 | Ldb2 | LIM domain binding 2 [Source:MGI Symbol;Acc:MGI:894670] | 93,07 | 1,7 |
| ENSMUSG00000038366 | 589,2746621 | 818,7 | 146,1 | 4,479469101 | 0,003764373 | Lasp1 | LIM and SH3 protein 1 [Source:MGI Symbol;Acc:MGI:109656] | 64,63 | 21,6 |
| ENSMUSG00000028696 | 57,05545708 | 122,8 | 12,3 | 4,481574403 | 0,005345699 | Ipp | IAP promoted placental gene [Source:MGI Symbol;Acc:MGI:96581] | 97,35 | 6,0 |
| ENSMUSG00000027220 | 66,248224 | 94,4 | 22,3 | 4,482188961 | 0,045609433 | Syt13 | synaptotagmin XIII [Source:MGI Symbol;Acc:MGI:1933945] | 93,74 | 6,4 |
| ENSMUSG00000057715 | 62,83053065 | 124,3 | 8,1 | 4,488763127 | 0,003034051 | A830018L16Rik | RIKEN cDNA A830018L16 gene [Source:MGI Symbol;Acc:MGI:2444149] | 73,81 | 1,8 |
| ENSMUSG00000103707 | 17,63713046 | 32,8 | 1,8 | 4,489977143 | 0,016437872 | Pcdha6 | protocadherin alpha 6 [Source:MGI Symbol;Acc:MGI:1298367] | 41,77 | 0,4 |
| ENSMUSG00000041343 | 232,8954012 | 317,0 | 23,9 | 4,49235128 | 0,010561845 | Ankrd42 | ankyrin repeat domain 42 [Source:MGI Symbol;Acc:MGI:1921095] | 62,93 | 2,5 |
| ENSMUSG00000024382 | 80,32880183 | 91,2 | 24,6 | 4,496842571 | 0,035357864 | Ercc3 | epair cross-complementing rodent repair deficiency, complementation group 3 [Source:MGI Symbol;Acc:M | 97,30 | 11,4 |
| ENSMUSG00000020564 | 333,7090826 | 453,7 | 229,8 | 4,497144431 | 0,015024331 | Atxn7l1 | ataxin 7-like 1 [Source:MGI Symbol;Acc:MGI:3584458] | 45,00 | 6,2 |
| ENSMUSG00000028967 | 109,3823751 | 127,5 | 241,2 | 4,497833409 | 0,041869735 | Errfi1 | ERBB receptor feedback inhibitor 1 [Source:MGI Symbol;Acc:MGI:1921405] | 93,08 | 30,0 |
| ENSMUSG00000066621 | 241,9389207 | 273,6 | 241,5 | 4,500063581 | 0,010756829 | Tecpr1 | tectonin beta-propeller repeat containing 1 [Source:MGI Symbol;Acc:MGI:1917631] | 98,79 | 26,9 |
| ENSMUSG00000031093 | 60,70371405 | 51,3 | 7,9 | 4,500127334 | 0,022976457 | Dock11 | dedicator of cytokinesis 11 [Source:MGI Symbol;Acc:MGI:1923224] | 65,18 | 1,6 |
| ENSMUSG00000043298 | 16,20074462 | 33,7 | 10,2 | 4,50148114 | 0,048578581 | Smco3 | single-pass membrane protein with coiled-coil domains 3 [Source:MGI Symbol;Acc:MGI:2443451] | 80,43 | 2,9 |
| ENSMUSG00000003923 | 296,8956941 | 456,7 | 207,5 | 4,501487668 | 0,01036577 | Tfam | transcription factor A, mitochondrial [Source:MGI Symbol;Acc:MGI:107810] | 89,48 | 12,0 |
| ENSMUSG00000039154 | 141,6260314 | 214,9 | 13,6 | 4,502966916 | 0,011486179 | Shd | src homology 2 domain-containing transforming protein D [Source:MGI Symbol;Acc:MGI:1099461] | 98,17 | 12,5 |
| ENSMUSG00000007216 | 114,8735366 | 144,9 | 123,9 | 4,506784568 | 0,047974717 | Zfp775 | zinc finger protein 775 [Source:MGI Symbol;Acc:MGI:2683557] | 63,64 | 19,2 |
| ENSMUSG00000028527 | 331,5419051 | 515,7 | 97,8 | 4,507246676 | 0,017796356 | Ak4 | adenylate kinase 4 [Source:MGI Symbol;Acc:MGI:87979] | 90,31 | 21,0 |
| ENSMUSG00000003581 | 202,9056783 | 349,4 | 135,4 | 4,507508648 | 0,043069248 | Rnf215 | ring finger protein 215 [Source:MGI Symbol;Acc:MGI:1918923] | 100,00 | 32,1 |
| ENSMUSG00000032359 | 21,11423121 | 34,9 | 1,6 | 4,510759238 | 0,01933552 | Ctsh | cathepsin H [Source:MGI Symbol;Acc:MGI:107285] | 72,50 | 2,8 |
| ENSMUSG00000010609 | 62,84793149 | 69,8 | 49,9 | 4,510845808 | 0,010179275 | Psen2 | presenilin 2 [Source:MGI Symbol;Acc:MGI:109284] | 81,47 | 20,0 |
| ENSMUSG00000029233 | 98,81630131 | 112,4 | 10,2 | 4,511845239 | 0,033841736 | Srd5a3 | steroid 5 alpha-reductase 3 [Source:MGI Symbol;Acc:MGI:1930252] | 81,75 | 3,3 |
| ENSMUSG00000038860 | 72,14399068 | 107,9 | 6,7 | 4,51450181 | 0,021867771 | Garnl3 | GTPase activating RANGAP domain-like 3 [Source:MGI Symbol;Acc:MGI:2139309] | 80,81 | 0,6 |
| ENSMUSG00000048376 | 32,19769374 | 56,5 | 15,7 | 4,519175667 | 0,024963366 | F2r | coagulation factor II (thrombin) receptor [Source:MGI Symbol;Acc:MGI:101802] | 92,72 | 3,3 |
| ENSMUSG00000020953 | 39,24989612 | 31,6 | 1,8 | 4,521000616 | 0,026876988 | Coch | cochlin [Source:MGI Symbol;Acc:MGI:1278313] | 63,25 | 0,9 |
| ENSMUSG00000045817 | 302,8993588 | 584,1 | 50,6 | 4,521790191 | 0,008002308 | Zfp36l2 | zinc finger protein 36, C3H type-like 2 [Source:MGI Symbol;Acc:MGI:107945] | 82,67 | 17,5 |
| ENSMUSG00000026185 | 515,4174616 | 729,2 | 105,9 | 4,521962538 | 0,013986814 | Igfbp5 | insulin-like growth factor binding protein 5 [Source:MGI Symbol;Acc:MGI:96440] | 99,58 | 11,9 |
| ENSMUSG00000021666 | 191,4420271 | 241,6 | 52,1 | 4,526160808 | 0,00624442 | Gfm2 | G elongation factor, mitochondrial 2 [Source:MGI Symbol;Acc:MGI:2444783] | 64,26 | 5,3 |
| ENSMUSG00000031095 | 335,4371376 | 346,8 | 41,5 | 4,526176831 | 0,041339457 | Cul4b | cullin 4B [Source:MGI Symbol;Acc:MGI:1919834] | 88,17 | 5,2 |
| ENSMUSG00000033088 | 180,2377966 | 245,7 | 49,8 | 4,526738986 | 0,011405516 | Triobp | TRIO and F-actin binding protein [Source:MGI Symbol;Acc:MGI:1349410] | 59,37 | 4,7 |
| ENSMUSG00000036552 | 22,97127266 | 41,4 | 15,7 | 4,529792721 | 0,038574663 | Ermard | ER membrane associated RNA degradation [Source:MGI Symbol;Acc:MGI:1917317] | 32,49 | 1,4 |
| ENSMUSG00000034471 | 102,3549975 | 174,9 | 22,9 | 4,53292714 | 0,028589358 | Caskin2 | CASK-interacting protein 2 [Source:MGI Symbol;Acc:MGI:2157062] | 93,83 | 5,8 |
| ENSMUSG00000045252 | 117,7521461 | 175,4 | 7,9 | 4,533004098 | 0,00047235 | Zfp574 | zinc finger protein 574 [Source:MGI Symbol;Acc:MGI:2442951] | 80,22 | 1,2 |
| ENSMUSG00000037112 | 27,56792897 | 56,5 | 7,9 | 4,535555412 | 0,022729402 | Sik2 | salt inducible kinase 2 [Source:MGI Symbol;Acc:MGI:2445031] | 67,32 | 0,5 |
| ENSMUSG00000049176 | 98,09717817 | 75,5 | 172,6 | 4,536418935 | 0,023575759 | Frmpd4 | FERM and PDZ domain containing 4 [Source:MGI Symbol;Acc:MGI:3042378] | 60,56 | 12,2 |
| ENSMUSG00000025597 | 11,18193407 | 25,4 | 2,0 | 4,542423348 | 0,041984702 | Klhl4 | kelch-like 4 [Source:MGI Symbol;Acc:MGI:2442829] | 68,95 | 0,5 |
| ENSMUSG00000050288 | 70,10187858 | 129,9 | 7,4 | 4,543536486 | 0,001749003 | Fzd2 | frizzled class receptor 2 [Source:MGI Symbol;Acc:MGI:1888513] | 87,10 | 2,9 |
| ENSMUSG00000038024 | 16,92170323 | 29,4 | 1,8 | 4,544431939 | 0,016213872 | Dennd4c | DENN/MADD domain containing 4C [Source:MGI Symbol;Acc:MGI:1914769] | 37,17 | 0,4 |
| ENSMUSG00000028800 | 158,1255068 | 180,4 | 108,6 | 4,548723482 | 0,006045628 | Hdac1 | histone deacetylase 1 [Source:MGI Symbol;Acc:MGI:108086] | 83,95 | 22,7 |
| ENSMUSG00000020034 | 275,0200625 | 367,0 | 107,1 | 4,549435465 | 0,04083678 | Tcp11l2 | t-complex 11 (mouse) like 2 [Source:MGI Symbol;Acc:MGI:2444679] | 91,89 | 28,4 |
| ENSMUSG00000042451 | 67,52801142 | 64,8 | 11,5 | 4,55578485 | 0,011411264 | Mybph | myosin binding protein H [Source:MGI Symbol;Acc:MGI:1858196] | 55,42 | 10,5 |
| ENSMUSG00000056919 | 69,13723412 | 82,7 | 3,5 | 4,558656226 | 0,005018328 | Cep162 | centrosomal protein 162 [Source:MGI Symbol;Acc:MGI:1925343] | 60,10 | 0,6 |
| ENSMUSG00000027615 | 24,16622385 | 38,9 | 4,0 | 4,561546775 | 0,023510692 | Hps3 | Hermansky-Pudlak syndrome 3 [Source:MGI Symbol;Acc:MGI:2153839] | 73,18 | 0,6 |
| ENSMUSG00000036403 | 15,14936888 | 24,4 | 1,8 | 4,565230328 | 0,046896001 | Cep135 | centrosomal protein 135 [Source:MGI Symbol;Acc:MGI:2681869] | 44,59 | 0,1 |
| ENSMUSG00000013155 | 41,41476917 | 45,3 | 33,3 | 4,565898264 | 0,039052016 | Enkd1 | enkurin domain containing 1 [Source:MGI Symbol;Acc:MGI:2142593] | 98,77 | 26,9 |
| ENSMUSG00000079598 | 57,92032764 | 57,2 | 65,4 | 4,566959243 | 0,037914345 | Clec2l | C-type lectin domain family 2, member L [Source:MGI Symbol;Acc:MGI:2141402] | 94,21 | 25,5 |
| ENSMUSG00000028991 | 82,76721191 | 76,7 | 3,5 | 4,567155295 | 0,009076555 | Mtor | mechanistic target of rapamycin (serine/threonine kinase) [Source:MGI Symbol;Acc:MGI:1928394] | 71,28 | 0,9 |
| ENSMUSG00000000402 | 41,68635167 | 52,9 | 64,8 | 4,568377089 | 0,039591111 | Egfl6 | EGF-like-domain, multiple 6 [Source:MGI Symbol;Acc:MGI:1858599] | 94,59 | 14,1 |
| ENSMUSG00000021870 | 412,3571269 | 495,9 | 32,8 | 4,570869649 | 0,000131252 | Slmap | sarcolemma associated protein [Source:MGI Symbol;Acc:MGI:1933549] | 94,01 | 6,0 |
| ENSMUSG00000028532 | 148,7389208 | 196,2 | 155,6 | 4,572902208 | 0,031059339 | Cachd1 | cache domain containing 1 [Source:MGI Symbol;Acc:MGI:2444177] | 92,21 | 24,9 |
| ENSMUSG00000028974 | 40,41654683 | 60,8 | 49,6 | 4,573065608 | 0,036911715 | Dffa | DNA fragmentation factor, alpha subunit [Source:MGI Symbol;Acc:MGI:1196227] | 87,76 | 11,6 |
| ENSMUSG00000003161 | 1520,814897 | 2380,4 | 298,3 | 4,575573785 | 0,003406984 | Sri | sorcin [Source:MGI Symbol;Acc:MGI:98419] | 90,88 | 41,8 |
| ENSMUSG00000086693 | 12,88083422 | 24,9 | 1,8 | 4,576499611 | 0,037067021 | A730081D07Rik | RIKEN cDNA A730081D07 gene [Source:MGI Symbol;Acc:MGI:2441825] | #DIV/0! | #DIV/0! |
| ENSMUSG00000051890 | 100,5233029 | 187,0 | 14,1 | 4,577153459 | 0,011852623 | Klhdc1 | kelch domain containing 1 [Source:MGI Symbol;Acc:MGI:2672853] | 98,88 | 2,8 |
| ENSMUSG00000015882 | 127,3117463 | 141,8 | 7,4 | 4,57726754 | 0,001337014 | Lcorl | ligand dependent nuclear receptor corepressor-like [Source:MGI Symbol;Acc:MGI:2651932] | 74,49 | 0,0 |
| ENSMUSG00000025609 | 343,2243945 | 562,0 | 57,8 | 4,578406134 | 0,01521624 | Mkln1 | muskelin 1, intracellular mediator containing kelch motifs [Source:MGI Symbol;Acc:MGI:1351638] | 72,41 | 3,5 |
| ENSMUSG00000024078 | 34,5607441 | 24,7 | 1,8 | 4,578438088 | 0,042639055 | Ttc27 | tetratricopeptide repeat domain 27 [Source:MGI Symbol;Acc:MGI:1921446] | 69,69 | 1,6 |
| ENSMUSG00000034429 | 34,83071353 | 56,1 | 28,7 | 4,578724577 | 0,030430428 | Zfp707 | zinc finger protein 707 [Source:MGI Symbol;Acc:MGI:1916270] | 92,46 | 18,1 |
| ENSMUSG00000086013 | 78,48770456 | 47,9 | 107,8 | 4,578889478 | 0,032295618 | Gm15706 | predicted gene 15706 [Source:MGI Symbol;Acc:MGI:3783146] | #DIV/0! | #DIV/0! |
| ENSMUSG00000025816 | 84,71017506 | 78,8 | 5,6 | 4,580561339 | 0,005198465 | Sec61a2 | Sec61, alpha subunit 2 (S. cerevisiae) [Source:MGI Symbol;Acc:MGI:1931071] | 95,06 | 2,8 |
| ENSMUSG00000024999 | 66,654606 | 51,3 | 121,0 | 4,584900003 | 0,035299784 | Noc3l | nucleolar complex associated 3 homolog (S. cerevisiae) [Source:MGI Symbol;Acc:MGI:1932610] | 84,41 | 19,1 |
| ENSMUSG00000038679 | 195,9069407 | 239,7 | 17,0 | 4,587419181 | 0,005837578 | Trps1 | trichorhinophalangeal syndrome I (human) [Source:MGI Symbol;Acc:MGI:1927616] | 70,63 | 1,3 |
| ENSMUSG00000027365 | 158,9519474 | 183,0 | 41,9 | 4,588756726 | 0,012503368 | Trpm7 | ansient receptor potential cation channel, subfamily M, member 7 [Source:MGI Symbol;Acc:MGI:1929996 | 93,62 | 2,5 |
| ENSMUSG00000030718 | 239,7260087 | 363,9 | 38,5 | 4,59019171 | 0,011357974 | Ppme1 | protein phosphatase methylesterase 1 [Source:MGI Symbol;Acc:MGI:1919840] | 87,15 | 13,0 |
| ENSMUSG00000060038 | 189,8835733 | 274,4 | 26,2 | 4,59407822 | 0,001241521 | Dhps | deoxyhypusine synthase [Source:MGI Symbol;Acc:MGI:2683592] | 100,00 | 10,8 |
| ENSMUSG00000097825 | 29,02539889 | 47,5 | 27,2 | 4,594116187 | 0,039738048 | 9630001P10Rik | RIKEN cDNA 9630001P10 gene [Source:MGI Symbol;Acc:MGI:2441755] | 66,04 | 4,6 |
| ENSMUSG00000041685 | 216,6592956 | 297,2 | 14,2 | 4,594729627 | 0,018265505 | Fcho2 | FCH domain only 2 [Source:MGI Symbol;Acc:MGI:3505790] | 96,45 | 3,8 |
| ENSMUSG00000097863 | 25,47462106 | 46,0 | 37,5 | 4,594829961 | 0,046773879 | 1010001B22Rik | RIKEN cDNA 1010001B22 gene [Source:MGI Symbol;Acc:MGI:1922653] | #DIV/0! | #DIV/0! |
| ENSMUSG00000038570 | 83,05862569 | 111,7 | 19,7 | 4,594931664 | 0,03178474 | Saxo2 | stablizer of axonemal microtubules 2 [Source:MGI Symbol;Acc:MGI:1914618] | 96,45 | 3,2 |
| ENSMUSG00000097081 | 13,62080793 | 26,2 | 2,0 | 4,597775359 | 0,040076558 | Gm10425 | predicted gene 10425 [Source:MGI Symbol;Acc:MGI:3642471] | #DIV/0! | #DIV/0! |
| ENSMUSG00000018412 | 136,9787431 | 188,7 | 28,8 | 4,599018624 | 0,011384864 | Kansl1 | KAT8 regulatory NSL complex subunit 1 [Source:MGI Symbol;Acc:MGI:1923969] | 75,64 | 4,0 |
| ENSMUSG00000022807 | 84,02539915 | 130,7 | 93,9 | 4,600971267 | 0,039697754 | Osbpl11 | oxysterol binding protein-like 11 [Source:MGI Symbol;Acc:MGI:2146553] | 98,06 | 21,6 |
| ENSMUSG00000024940 | 168,6583035 | 265,5 | 17,4 | 4,601148793 | 0,002985684 | Ltbp3 | latent transforming growth factor beta binding protein 3 [Source:MGI Symbol;Acc:MGI:1101355] | 87,95 | 1,3 |
| ENSMUSG00000025504 | 36,22186964 | 39,0 | 23,1 | 4,601379107 | 0,036159363 | Eps8l2 | EPS8-like 2 [Source:MGI Symbol;Acc:MGI:2138828] | 82,85 | 7,0 |
| ENSMUSG00000028546 | 36,03252211 | 68,8 | 3,6 | 4,602102893 | 0,005119497 | Elavl4 | V (embryonic lethal, abnormal vision, Drosophila)-like 4 (Hu antigen D) [Source:MGI Symbol;Acc:MGI:107 | 56,98 | 1,0 |
| ENSMUSG00000001227 | 47,48625443 | 50,2 | 105,9 | 4,60215299 | 0,04559403 | Sema6b | in, transmembrane domain (TM), and cytoplasmic domain, (semaphorin) 6B [Source:MGI Symbol;Acc:MG | 83,50 | 26,4 |
| ENSMUSG00000040282 | 46,20472856 | 90,7 | 7,5 | 4,603431894 | 0,005825188 | BC052040 | cDNA sequence BC052040 [Source:MGI Symbol;Acc:MGI:3026886] | 92,86 | 2,7 |
| ENSMUSG00000030747 | 106,1439928 | 155,3 | 92,4 | 4,604181096 | 0,036395652 | Dgat2 | diacylglycerol O-acyltransferase 2 [Source:MGI Symbol;Acc:MGI:1915050] | 96,38 | 29,7 |
| ENSMUSG00000089840 | 33,54769858 | 52,8 | 66,0 | 4,605924463 | 0,040866069 | Gm4491 | predicted gene 4491 [Source:MGI Symbol;Acc:MGI:3782676] | 97,05 | 32,1 |
| ENSMUSG00000021245 | 151,0109234 | 173,9 | 19,6 | 4,606306229 | 0,01419944 | Mlh3 | mutL homolog 3 (E coli) [Source:MGI Symbol;Acc:MGI:1353455] | 97,98 | 3,1 |
| ENSMUSG00000086401 | 18,50553771 | 41,5 | 7,4 | 4,61519307 | 0,03083772 | Gm15559 | predicted gene 15559 [Source:MGI Symbol;Acc:MGI:3783008] | 81,70 | 2,0 |
| ENSMUSG00000039239 | 250,6644308 | 224,7 | 144,5 | 4,621132892 | 0,00988044 | Tgfb2 | transforming growth factor, beta 2 [Source:MGI Symbol;Acc:MGI:98726] | 95,96 | 16,4 |
| ENSMUSG00000069920 | 20,70589962 | 42,1 | 7,4 | 4,621365496 | 0,026869909 | B3gnt9 | JDP-GlcNAc:betaGal beta-1,3-N-acetylglucosaminyltransferase 9 [Source:MGI Symbol;Acc:MGI:2142841 | 28,44 | 3,1 |
| ENSMUSG00000020248 | 266,5290303 | 373,7 | 48,0 | 4,622961237 | 0,000190848 | Nfyb | nuclear transcription factor-Y beta [Source:MGI Symbol;Acc:MGI:97317] | 98,69 | 14,9 |
| ENSMUSG00000032238 | 432,0270189 | 815,2 | 67,3 | 4,623851816 | 0,006626268 | Rora | RAR-related orphan receptor alpha [Source:MGI Symbol;Acc:MGI:104661] | 86,52 | 5,0 |
| ENSMUSG00000028300 | 86,14146224 | 125,2 | 26,4 | 4,625087345 | 0,024067236 | 3110043O21Rik | RIKEN cDNA 3110043O21 gene [Source:MGI Symbol;Acc:MGI:1920455] | 90,10 | 8,2 |
| ENSMUSG00000025733 | 186,2667002 | 213,3 | 12,4 | 4,625331465 | 0,003911784 | Rhot2 | ras homolog gene family, member T2 [Source:MGI Symbol;Acc:MGI:2384892] | 91,78 | 6,9 |
| ENSMUSG00000097517 | 31,466022 | 60,8 | 33,9 | 4,626035469 | 0,048346724 | Gm26695 | predicted gene, 26695 [Source:MGI Symbol;Acc:MGI:5477189] | #DIV/0! | #DIV/0! |
| ENSMUSG00000074865 | 23,90915005 | 45,1 | 28,9 | 4,626988529 | 0,042593161 | Zfp934 | zinc finger protein 934 [Source:MGI Symbol;Acc:MGI:1924367] | 83,49 | 12,1 |
| ENSMUSG00000029560 | 129,7609262 | 184,2 | 34,8 | 4,62807416 | 0,007465481 | Snx8 | sorting nexin 8 [Source:MGI Symbol;Acc:MGI:2443816] | 90,63 | 7,9 |
| ENSMUSG00000028573 | 144,1514733 | 250,1 | 14,8 | 4,628858805 | 0,010213022 | Fggy | FGGY carbohydrate kinase domain containing [Source:MGI Symbol;Acc:MGI:1922828] | 81,28 | 4,9 |
| ENSMUSG00000031555 | 230,0046984 | 228,9 | 53,3 | 4,629030878 | 0,039272304 | Adam9 | a disintegrin and metallopeptidase domain 9 (meltrin gamma) [Source:MGI Symbol;Acc:MGI:105376] | 98,14 | 9,2 |
| ENSMUSG00000097461 | 37,54796553 | 67,9 | 11,5 | 4,629397255 | 0,012198906 | Gm26735 | predicted gene, 26735 [Source:MGI Symbol;Acc:MGI:5477229] | #DIV/0! | #DIV/0! |
| ENSMUSG00000031980 | 458,2439055 | 628,4 | 59,6 | 4,630712684 | 0,011193495 | Agt | angiotensinogen (serpin peptidase inhibitor, clade A, member 8) [Source:MGI Symbol;Acc:MGI:87963] | 97,21 | 33,8 |
| ENSMUSG00000039960 | 338,4913592 | 682,1 | 58,5 | 4,631808303 | 0,014677393 | Rhou | ras homolog gene family, member U [Source:MGI Symbol;Acc:MGI:1916831] | 99,37 | 18,3 |
| ENSMUSG00000021978 | 231,4163435 | 300,7 | 68,6 | 4,632755803 | 0,010069978 | Extl3 | exostoses (multiple)-like 3 [Source:MGI Symbol;Acc:MGI:1860765] | 99,32 | 8,2 |
| ENSMUSG00000049658 | 100,420567 | 96,3 | 11,7 | 4,633980119 | 0,006870532 | Bdp1 | e prime 1, subunit of RNA polymerase III transcription initiation factor IIIB [Source:MGI Symbol;Acc:MGI:1: | 62,55 | 1,5 |
| ENSMUSG00000035649 | 62,60351381 | 72,3 | 24,2 | 4,634263576 | 0,010895155 | Zcchc7 | zinc finger, CCHC domain containing 7 [Source:MGI Symbol;Acc:MGI:2442912] | 45,22 | 3,6 |
| ENSMUSG00000000223 | 282,8210578 | 510,9 | 46,9 | 4,635150507 | 0,009687502 | Drp2 | dystrophin related protein 2 [Source:MGI Symbol;Acc:MGI:107432] | 96,54 | 4,8 |
| ENSMUSG00000024666 | 69,66822067 | 116,3 | 54,8 | 4,636525826 | 0,042608321 | Tmem138 | transmembrane protein 138 [Source:MGI Symbol;Acc:MGI:1920232] | 97,57 | 25,6 |
| ENSMUSG00000034101 | 119,6414665 | 115,0 | 297,3 | 4,637627666 | 0,040279897 | Ctnnd1 | catenin (cadherin associated protein), delta 1 [Source:MGI Symbol;Acc:MGI:105100] | 75,91 | 24,0 |
| ENSMUSG00000031536 | 435,0680435 | 792,2 | 137,7 | 4,637662953 | 0,009723044 | Polb | polymerase (DNA directed), beta [Source:MGI Symbol;Acc:MGI:97740] | 99,76 | 32,3 |
| ENSMUSG00000037965 | 119,3330584 | 106,8 | 6,8 | 4,637665903 | 0,007999802 | Zc3h7a | zinc finger CCCH type containing 7 A [Source:MGI Symbol;Acc:MGI:2445044] | 85,25 | 1,7 |
| ENSMUSG00000054517 | 22,59495043 | 35,3 | 2,8 | 4,637912459 | 0,021087735 | Trim65 | tripartite motif-containing 65 [Source:MGI Symbol;Acc:MGI:2442815] | 57,26 | 0,9 |
| ENSMUSG00000032232 | 31,83450041 | 58,2 | 5,0 | 4,638603639 | 0,014140921 | Cgnl1 | cingulin-like 1 [Source:MGI Symbol;Acc:MGI:1915428] | 57,68 | 0,4 |

| | | | | | | | | | |
|---|---|---|---|---|---|---|---|---|---|
| ENSMUSG00000097727 | 64,97480594 | 66,5 | 205,2 | 4,638671036 | 0,046775818 | F630040K05Rik | RIKEN cDNA F630040K05 gene [Source:MGI Symbol;Acc:MGI:4437734] | #DIV/0! | #DIV/0! |
| ENSMUSG00000054074 | 92,63784834 | 128,1 | 6,7 | 4,641256276 | 0,010732531 | Skida1 | SKI/DACH domain containing 1 [Source:MGI Symbol;Acc:MGI:1919918] | 88,30 | 1,6 |
| ENSMUSG00000022019 | 133,8409879 | 148,8 | 81,4 | 4,642587682 | 0,008739075 | Tdrd3 | tudor domain containing 3 [Source:MGI Symbol;Acc:MGI:2444023] | 76,97 | 12,9 |
| ENSMUSG00000028771 | 98,58650392 | 82,6 | 4,2 | 4,642819249 | 0,017459821 | Ptpn12 | protein tyrosine phosphatase, non-receptor type 12 [Source:MGI Symbol;Acc:MGI:104673] | 89,90 | 2,0 |
| ENSMUSG00000101952 | 32,55485166 | 36,0 | 5,5 | 4,644398841 | 0,046454798 | Gm10550 | predicted gene 10550 [Source:MGI Symbol;Acc:MGI:3704193] | 70,06 | 2,5 |
| ENSMUSG00000038545 | 23,83033077 | 47,1 | 12,1 | 4,645712683 | 0,032632391 | Cul7 | cullin 7 [Source:MGI Symbol;Acc:MGI:1913765] | 76,23 | 2,3 |
| ENSMUSG00000025133 | 177,0168677 | 133,7 | 6,2 | 4,648829913 | 0,000737458 | Ints4 | integrator complex subunit 4 [Source:MGI Symbol;Acc:MGI:1917164] | 98,07 | 2,6 |
| ENSMUSG00000031290 | 66,06140536 | 42,7 | 4,6 | 4,649081984 | 0,023617627 | Lrch2 | cine-rich repeats and calponin homology (CH) domain containing 2 [Source:MGI Symbol;Acc:MGI:21478 | 60,99 | 0,7 |
| ENSMUSG00000021257 | 17,10072855 | 40,5 | 1,6 | 4,649353189 | 0,014129347 | Angel1 | angel homolog 1 (Drosophila) [Source:MGI Symbol;Acc:MGI:1915987] | 80,35 | 1,3 |
| ENSMUSG00000029469 | 88,92441317 | 161,8 | 12,9 | 4,650102689 | 0,010337447 | Ift81 | intraflagellar transport 81 [Source:MGI Symbol;Acc:MGI:1098597] | 92,23 | 3,0 |
| ENSMUSG00000043557 | 22,2428005 | 28,0 | 1,8 | 4,654439595 | 0,049215399 | Mdga1 | MAM domain containing glycosylphosphatidylinositol anchor 1 [Source:MGI Symbol;Acc:MGI:1922012] | 36,31 | 0,3 |
| ENSMUSG00000024896 | 132,8335194 | 169,1 | 56,7 | 4,654822709 | 0,043662164 | Minpp1 | multiple inositol polyphosphate histidine phosphatase 1 [Source:MGI Symbol;Acc:MGI:1336159] | 94,07 | 20,6 |
| ENSMUSG00000035704 | 63,33657144 | 52,2 | 2,9 | 4,655768246 | 0,021173437 | Alg8 | asparagine-linked glycosylation 8 (alpha-1,3-glucosyltransferase) [Source:MGI Symbol;Acc:MGI:2141959] | 89,78 | 2,1 |
| ENSMUSG00000027361 | 107,853549 | 87,3 | 48,1 | 4,661314901 | 0,005859943 | Gabpb1 | GA repeat binding protein, beta 1 [Source:MGI Symbol;Acc:MGI:95611] | 36,19 | 5,0 |
| ENSMUSG00000028344 | 46,39844418 | 56,8 | 82,3 | 4,663116005 | 0,035026938 | Invs | inversin [Source:MGI Symbol;Acc:MGI:1335082] | 73,79 | 9,9 |
| ENSMUSG00000004791 | 40,48229919 | 68,5 | 27,8 | 4,663310743 | 0,023269957 | Pgf | placental growth factor [Source:MGI Symbol;Acc:MGI:105095] | 69,76 | 15,6 |
| ENSMUSG00000020827 | 191,5122669 | 299,9 | 45,9 | 4,66384238 | 0,000425299 | Mink1 | misshapen-like kinase 1 (zebrafish) [Source:MGI Symbol;Acc:MGI:1355329] | 95,07 | 14,4 |
| ENSMUSG00000041272 | 32,58748637 | 50,6 | 5,0 | 4,665184585 | 0,010339674 | Tox | thymocyte selection-associated high mobility group box [Source:MGI Symbol;Acc:MGI:2181659] | 67,29 | 0,6 |
| ENSMUSG00000024319 | 249,7732712 | 352,0 | 21,7 | 4,66858622 | 0,006156797 | Vps52 | vacuolar protein sorting 52 (yeast) [Source:MGI Symbol;Acc:MGI:1330304] | 97,65 | 10,9 |
| ENSMUSG00000035228 | 81,21064024 | 72,4 | 56,4 | 4,671708403 | 0,013163485 | Ccdc106 | coiled-coil domain containing 106 [Source:MGI Symbol;Acc:MGI:2385900] | 78,12 | 21,7 |
| ENSMUSG00000032571 | 33,49535448 | 54,9 | 16,6 | 4,671969762 | 0,014750469 | Pik3r4 | osphatidylinositol 3 kinase, regulatory subunit, polypeptide 4, p150 [Source:MGI Symbol;Acc:MGI:192291 | 74,75 | 5,2 |
| ENSMUSG00000044950 | 159,0067555 | 169,6 | 12,6 | 4,675718968 | 0,009740385 | Pwwp2a | PWWP domain containing 2A [Source:MGI Symbol;Acc:MGI:1918052] | 85,91 | 1,4 |
| ENSMUSG00000050310 | 100,8762041 | 104,9 | 8,3 | 4,676262875 | 0,031228558 | Rictor | RPTOR independent companion of MTOR, complex 2 [Source:MGI Symbol;Acc:MGI:1926007] | 65,02 | 0,7 |
| ENSMUSG00000028322 | 237,8592256 | 357,3 | 110,8 | 4,678068018 | 0,012702191 | Exosc3 | exosome component 3 [Source:MGI Symbol;Acc:MGI:1913612] | 99,53 | 28,6 |
| ENSMUSG00000020805 | 33,39366895 | 54,9 | 26,9 | 4,681227305 | 0,033488223 | Slc13a5 | :e carrier family 13 (sodium-dependent citrate transporter), member 5 [Source:MGI Symbol;Acc:MGI:3037 | 63,72 | 4,4 |
| ENSMUSG00000003438 | 229,5835021 | 331,7 | 53,2 | 4,681624445 | 0,006652102 | Timm50 | translocase of inner mitochondrial membrane 50 [Source:MGI Symbol;Acc:MGI:1913775] | 99,52 | 23,7 |
| ENSMUSG00000027132 | 291,2757029 | 360,8 | 197,6 | 4,682265615 | 0,008290685 | Katnbl1 | katanin p80 subunit B like 1 [Source:MGI Symbol;Acc:MGI:1919675] | 94,96 | 13,0 |
| ENSMUSG00000039959 | 278,1492707 | 459,2 | 171,3 | 4,685101254 | 0,006773063 | Hip1 | huntingtin interacting protein 1 [Source:MGI Symbol;Acc:MGI:1099804] | 98,86 | 20,4 |
| ENSMUSG00000036932 | 155,4954475 | 179,8 | 13,3 | 4,686432079 | 0,013268445 | Aifm1 | apoptosis-inducing factor, mitochondrion-associated 1 [Source:MGI Symbol;Acc:MGI:1349419] | 91,10 | 5,9 |
| ENSMUSG00000034453 | 74,05691176 | 76,4 | 41,8 | 4,694211674 | 0,020710182 | Polr3b | polymerase (RNA) III (DNA directed) polypeptide B [Source:MGI Symbol;Acc:MGI:1917678] | 89,83 | 5,4 |
| ENSMUSG00000001555 | 164,6411786 | 288,1 | 19,5 | 4,694689284 | 0,019770307 | Fkbp10 | FK506 binding protein 10 [Source:MGI Symbol;Acc:MGI:104769] | 99,57 | 6,7 |
| ENSMUSG00000098950 | 390,4451128 | 576,3 | 33,0 | 4,697026559 | 1,21917E-05 | Gm28036 | predicted gene, 28036 [Source:MGI Symbol;Acc:MGI:5547772] | 100,00 | 0,0 |
| ENSMUSG00000024998 | 75,63624224 | 125,2 | 42,6 | 4,701809167 | 0,010254873 | Plce1 | phospholipase C, epsilon 1 [Source:MGI Symbol;Acc:MGI:1921305] | 75,93 | 2,3 |
| ENSMUSG00000027646 | 160,0146985 | 262,3 | 14,2 | 4,703208792 | 0,005639541 | Src | Rous sarcoma oncogene [Source:MGI Symbol;Acc:MGI:98397] | 96,49 | 6,8 |
| ENSMUSG00000040649 | 128,0305031 | 215,6 | 59,4 | 4,704204759 | 0,008268466 | Rimklb | ribosomal modification protein rimK-like family member B [Source:MGI Symbol;Acc:MGI:1918325] | 85,55 | 12,5 |
| ENSMUSG00000031889 | 122,8444584 | 224,1 | 55,6 | 4,705349554 | 0,00919271 | D230025D16Rik | RIKEN cDNA D230025D16 gene [Source:MGI Symbol;Acc:MGI:2443049] | 93,05 | 13,7 |
| ENSMUSG00000038827 | 129,0255053 | 153,6 | 9,0 | 4,705476027 | 0,01657662 | Fam206a | family with sequence similarity 206, member A [Source:MGI Symbol;Acc:MGI:2677850] | 97,68 | 2,3 |
| ENSMUSG00000087331 | 43,66708448 | 61,2 | 23,6 | 4,705748702 | 0,021479852 | 1810021B22Rik | RIKEN cDNA 1810021B22 gene [Source:MGI Symbol;Acc:MGI:1916370] | 99,70 | 24,5 |
| ENSMUSG00000017146 | 37,09024938 | 37,9 | 16,0 | 4,70694332 | 0,047318583 | Brca1 | breast cancer 1, early onset [Source:MGI Symbol;Acc:MGI:104537] | 21,83 | 1,0 |
| ENSMUSG00000028617 | 95,08898071 | 141,1 | 23,4 | 4,709950446 | 0,028169901 | Lrrc42 | leucine rich repeat containing 42 [Source:MGI Symbol;Acc:MGI:1925059] | 89,25 | 21,2 |
| ENSMUSG00000025915 | 254,0569374 | 288,6 | 55,4 | 4,710124037 | 0,011173068 | Sgk3 | serum/glucocorticoid regulated kinase 3 [Source:MGI Symbol;Acc:MGI:2182368] | 93,72 | 8,9 |
| ENSMUSG00000074635 | 27,80660476 | 49,3 | 16,6 | 4,711749754 | 0,028391752 | 3110070M22Rik | RIKEN cDNA 3110070M22 gene [Source:MGI Symbol;Acc:MGI:1914554] | 99,17 | 16,2 |
| ENSMUSG00000034484 | 555,1475291 | 864,5 | 147,7 | 4,712895578 | 0,017066504 | Snx2 | sorting nexin 2 [Source:MGI Symbol;Acc:MGI:1915054] | 98,38 | 34,2 |
| ENSMUSG00000068740 | 80,99493423 | 98,6 | 7,9 | 4,716178771 | 0,00713294 | Celsr2 | cadherin, EGF LAG seven-pass G-type receptor 2 [Source:MGI Symbol;Acc:MGI:1858235] | 66,85 | 1,0 |
| ENSMUSG00000022704 | 53,51655247 | 55,6 | 20,6 | 4,717928551 | 0,040051414 | Qtrtd1 | queuine tRNA-ribosyltransferase domain containing 1 [Source:MGI Symbol;Acc:MGI:1922194] | 95,80 | 3,1 |
| ENSMUSG00000031431 | 3733,547352 | 8167,4 | 579,5 | 4,720736378 | 2,59185E-05 | Tsc22d3 | TSC22 domain family, member 3 [Source:MGI Symbol;Acc:MGI:1196284] | 90,35 | 57,3 |
| ENSMUSG00000028293 | 103,3835522 | 184,3 | 42,7 | 4,722479346 | 0,006824289 | Slc35a1 | solute carrier family 35 (CMP-sialic acid transporter), member 1 [Source:MGI Symbol;Acc:MGI:1345622] | 96,85 | 19,5 |
| ENSMUSG00000031613 | 21,54833113 | 37,5 | 4,0 | 4,72638777 | 0,024470355 | Hpgd | hydroxyprostaglandin dehydrogenase 15 (NAD) [Source:MGI Symbol;Acc:MGI:108085] | 92,09 | 1,4 |
| ENSMUSG00000030235 | 584,8064651 | 779,4 | 227,8 | 4,729652828 | 0,004694199 | Slco1c1 | solute carrier organic anion transporter family, member 1c1 [Source:MGI Symbol;Acc:MGI:1889679] | 94,38 | 25,7 |
| ENSMUSG00000030538 | 656,373653 | 1129,0 | 169,2 | 4,730889407 | 0,012435143 | Cib1 | calcium and integrin binding 1 (calmyrin) [Source:MGI Symbol;Acc:MGI:1344418] | 88,77 | 44,4 |
| ENSMUSG00000021902 | 41,02945384 | 40,9 | 17,6 | 4,732155761 | 0,044212092 | Phf7 | PHD finger protein 7 [Source:MGI Symbol;Acc:MGI:1919088] | 94,39 | 10,6 |
| ENSMUSG00000024502 | 124,2872161 | 219,1 | 43,6 | 4,732747055 | 0,006566597 | Jakmip2 | janus kinase and microtubule interacting protein 2 [Source:MGI Symbol;Acc:MGI:1923467] | 98,96 | 11,6 |
| ENSMUSG00000042404 | 58,78082944 | 60,9 | 81,3 | 4,733583826 | 0,031804477 | Dennd4b | DENN/MADD domain containing 4B [Source:MGI Symbol;Acc:MGI:2446201] | 73,03 | 17,6 |
| ENSMUSG00000052751 | 342,57704 | 590,6 | 157,5 | 4,734354113 | 0,025306745 | Repin1 | replication initiator 1 [Source:MGI Symbol;Acc:MGI:1889817] | 99,63 | 31,6 |
| ENSMUSG00000024426 | 250,443586 | 451,0 | 22,4 | 4,73557451 | 0,011862764 | Atat1 | alpha tubulin acetyltransferase 1 [Source:MGI Symbol;Acc:MGI:1913869] | 95,50 | 11,8 |
| ENSMUSG00000025555 | 570,6495437 | 784,2 | 47,0 | 4,73606724 | 3,85469E-05 | Farp1 | RhoGEF (Arhgef) and pleckstrin domain protein 1 (chondrocyte-derived) [Source:MGI Symbol;Acc:MGI:24 | 96,21 | 10,8 |
| ENSMUSG00000054400 | 48,672898 | 51,9 | 16,6 | 4,742560664 | 0,018852943 | Cklf | chemokine-like factor [Source:MGI Symbol;Acc:MGI:1922708] | 41,88 | 5,4 |
| ENSMUSG00000042010 | 28,07530369 | 38,3 | 1,5 | 4,744646642 | 0,013609945 | Acacb | acetyl-Coenzyme A carboxylase beta [Source:MGI Symbol;Acc:MGI:2140940] | 42,43 | 0,3 |
| ENSMUSG00000020205 | 57,35932426 | 91,8 | 7,5 | 4,74519153 | 0,017418341 | Phlda1 | pleckstrin homology-like domain, family A, member 1 [Source:MGI Symbol;Acc:MGI:1096880] | 82,66 | 7,0 |
| ENSMUSG00000017713 | 32,8082667 | 47,0 | 71,4 | 4,745452131 | 0,04176244 | Tha1 | threonine aldolase 1 [Source:MGI Symbol;Acc:MGI:1919026] | 92,52 | 25,2 |
| ENSMUSG00000037579 | 38,17329115 | 52,4 | 54,5 | 4,746333523 | 0,030353701 | Kcnh3 | assium voltage-gated channel, subfamily H (eag-related), member 3 [Source:MGI Symbol;Acc:MGI:13417 | 89,12 | 20,7 |
| ENSMUSG00000053600 | 31,79356799 | 40,9 | 5,5 | 4,74847666 | 0,038976922 | Zfp472 | zinc finger protein 472 [Source:MGI Symbol;Acc:MGI:2385049] | 88,44 | 1,4 |
| ENSMUSG00000027560 | 182,7017517 | 252,6 | 58,3 | 4,750269679 | 0,024787833 | Dok5 | docking protein 5 [Source:MGI Symbol;Acc:MGI:1924079] | 94,12 | 23,5 |
| ENSMUSG00000006019 | 63,59121193 | 129,8 | 10,2 | 4,751599454 | 0,001979734 | Dhx34 | DEAH (Asp-Glu-Ala-His) box polypeptide 34 [Source:MGI Symbol;Acc:MGI:1918973] | 90,58 | 3,9 |
| ENSMUSG00000033510 | 89,30461357 | 175,4 | 54,6 | 4,753126698 | 0,012135925 | Otud7a | OTU domain containing 7A [Source:MGI Symbol;Acc:MGI:2158505] | 98,72 | 19,7 |
| ENSMUSG00000004642 | 244,5282021 | 193,5 | 206,2 | 4,754639891 | 0,004007706 | Slbp | stem-loop binding protein [Source:MGI Symbol;Acc:MGI:108402] | 55,47 | 16,4 |
| ENSMUSG00000035948 | 19,24946209 | 33,0 | 2,8 | 4,755418485 | 0,041007987 | Acss3 | acyl-CoA synthetase short-chain family member 3 [Source:MGI Symbol;Acc:MGI:2685720] | 55,50 | 1,6 |
| ENSMUSG00000049717 | 17,08395724 | 40,4 | 5,5 | 4,75599756 | 0,025459641 | Lig4 | ligase IV, DNA, ATP-dependent [Source:MGI Symbol;Acc:MGI:1335098] | 63,21 | 0,9 |
| ENSMUSG00000039585 | 95,8239928 | 147,5 | 13,6 | 4,756811661 | 0,003911996 | Myo9a | myosin IXa [Source:MGI Symbol;Acc:MGI:107735] | 72,92 | 1,2 |
| ENSMUSG00000035212 | 178,7861882 | 275,5 | 70,2 | 4,757026711 | 0,014613111 | Leprot | leptin receptor overlapping transcript [Source:MGI Symbol;Acc:MGI:2687005] | 98,51 | 29,0 |
| ENSMUSG00000019261 | 110,0927586 | 139,8 | 44,5 | 4,757252525 | 0,036425207 | Map1s | microtubule-associated protein 1S [Source:MGI Symbol;Acc:MGI:2443304] | 98,80 | 5,6 |
| ENSMUSG00000027770 | 247,179692 | 374,7 | 104,1 | 4,757252659 | 0,008475401 | Dhx36 | DEAH (Asp-Glu-Ala-His) box polypeptide 36 [Source:MGI Symbol;Acc:MGI:1919412] | 95,91 | 5,2 |
| ENSMUSG00000037035 | 14,64821353 | 29,9 | 2,0 | 4,757480515 | 0,027230265 | Inhbb | inhibin beta-B [Source:MGI Symbol;Acc:MGI:96571] | 55,87 | 0,6 |
| ENSMUSG00000086149 | 15,2260891 | 26,8 | 1,8 | 4,760735746 | 0,04126323 | Gm15598 | predicted gene 15598 [Source:MGI Symbol;Acc:MGI:3783045] | 39,71 | 3,3 |
| ENSMUSG00000039431 | 24,70149624 | 56,6 | 2,6 | 4,761119901 | 0,005499898 | Mtmr7 | myotubularin related protein 7 [Source:MGI Symbol;Acc:MGI:1891693] | 58,59 | 1,4 |
| ENSMUSG00000023905 | 20,02653569 | 31,4 | 1,8 | 4,761692771 | 0,036800251 | Tnfrsf12a | tumor necrosis factor receptor superfamily, member 12a [Source:MGI Symbol;Acc:MGI:1351484] | 93,77 | 2,0 |
| ENSMUSG00000014668 | 162,5940311 | 201,5 | 9,6 | 4,762015405 | 0,021458515 | Chfr | checkpoint with forkhead and ring finger domains [Source:MGI Symbol;Acc:MGI:2444898] | 87,52 | 1,6 |
| ENSMUSG00000085331 | 23,43262646 | 56,2 | 20,0 | 4,76325075 | 0,030153207 | Gm11274 | predicted gene 11274 [Source:MGI Symbol;Acc:MGI:3649225] | 100,00 | 32,2 |
| ENSMUSG00000009207 | 105,3080468 | 73,3 | 88,4 | 4,763497866 | 0,023012649 | Lnp | limb and neural patterns [Source:MGI Symbol;Acc:MGI:1918115] | 81,51 | 3,7 |
| ENSMUSG00000032952 | 119,194883 | 140,7 | 11,1 | 4,765988348 | 0,036395652 | Ap4b1 | adaptor-related protein complex AP-4, beta 1 [Source:MGI Symbol;Acc:MGI:1337130] | 96,80 | 3,3 |
| ENSMUSG00000055676 | 68,81559705 | 104,8 | 10,9 | 4,766146876 | 0,014560625 | Gm5069 | predicted pseudogene 5069 [Source:MGI Symbol;Acc:MGI:3644516] | #DIV/0! | #DIV/0! |
| ENSMUSG00000084781 | 35,31787644 | 54,5 | 9,1 | 4,766610465 | 0,028301947 | D930015M05Rik | RIKEN cDNA D930015M05 gene [Source:MGI Symbol;Acc:MGI:3612068] | #DIV/0! | #DIV/0! |
| ENSMUSG00000031561 | 112,8095973 | 159,5 | 107,3 | 4,768956883 | 0,014627184 | Tenm3 | teneurin transmembrane protein 3 [Source:MGI Symbol;Acc:MGI:1345183] | 72,69 | 12,2 |
| ENSMUSG00000057286 | 117,592055 | 81,0 | 2,6 | 4,769182944 | 0,012683404 | St6galnac2 | euraminyl-2,3-beta-galactosyl-1,3)-N-acetylgalactosaminide alpha-2,6-sialyltransferase 2 [Source:MGI Sym | 58,94 | 1,3 |
| ENSMUSG00000105489 | 63,83867437 | 131,0 | 42,7 | 4,76980269 | 0,027651309 | Gm43477 | predicted gene 43477 [Source:MGI Symbol;Acc:MGI:5663614] | 100,00 | 30,7 |
| ENSMUSG00000067150 | 54,22611203 | 65,3 | 119,8 | 4,770172319 | 0,038783839 | Xpo5 | exportin 5 [Source:MGI Symbol;Acc:MGI:1913789] | 92,08 | 26,5 |
| ENSMUSG00000020841 | 53,05466632 | 66,9 | 2,7 | 4,770525686 | 0,010849987 | Cpd | carboxypeptidase D [Source:MGI Symbol;Acc:MGI:107265] | 73,72 | 0,6 |
| ENSMUSG00000022790 | 208,1475751 | 376,0 | 27,2 | 4,770600136 | 0,000455775 | Igsf11 | immunoglobulin superfamily, member 11 [Source:MGI Symbol;Acc:MGI:2388477] | 94,84 | 8,7 |
| ENSMUSG00000038637 | 27,94168394 | 47,6 | 22,2 | 4,770895836 | 0,030232994 | Lrrc56 | leucine rich repeat containing 56 [Source:MGI Symbol;Acc:MGI:1917802] | 89,86 | 12,4 |
| ENSMUSG00000039801 | 26,21335406 | 37,7 | 1,8 | 4,771322952 | 0,010484536 | 2410089E03Rik | RIKEN cDNA 2410089E03 gene [Source:MGI Symbol;Acc:MGI:1920942] | 37,40 | 0,5 |
| ENSMUSG00000024135 | 37,47678412 | 81,3 | 11,5 | 4,776066145 | 0,013414871 | Srbd1 | S1 RNA binding domain 1 [Source:MGI Symbol;Acc:MGI:1925836] | 92,43 | 2,9 |
| ENSMUSG00000037029 | 22,73781397 | 36,2 | 2,4 | 4,778378172 | 0,034635525 | Zfp146 | zinc finger protein 146 [Source:MGI Symbol;Acc:MGI:1347092] | 92,08 | 1,2 |
| ENSMUSG00000026279 | 100,8991203 | 117,7 | 7,7 | 4,77968807 | 0,009440287 | Thap4 | THAP domain containing 4 [Source:MGI Symbol;Acc:MGI:1914276] | 35,80 | 1,4 |
| ENSMUSG00000020868 | 45,13221291 | 72,3 | 3,8 | 4,780729942 | 0,006538382 | Xylt2 | xylosyltransferase II [Source:MGI Symbol;Acc:MGI:2444797] | 92,09 | 2,4 |
| ENSMUSG00000034973 | 137,0519753 | 82,4 | 145,2 | 4,780966038 | 0,027485495 | Dopey1 | dopey family member 1 [Source:MGI Symbol;Acc:MGI:1289294] | 42,58 | 7,4 |
| ENSMUSG00000039813 | 37,55449416 | 44,9 | 1,9 | 4,781495607 | 0,008037903 | Tbc1d2 | TBC1 domain family, member 2 [Source:MGI Symbol;Acc:MGI:2652885] | 65,32 | 0,9 |
| ENSMUSG00000021680 | 11,78407087 | 27,4 | 1,2 | 4,781514916 | 0,027635847 | Crhbp | corticotropin releasing hormone binding protein [Source:MGI Symbol;Acc:MGI:88497] | 93,63 | 2,5 |
| ENSMUSG00000029607 | 55,68106201 | 76,5 | 32,1 | 4,78454503 | 0,013916721 | Ankrd61 | ankyrin repeat domain 61 [Source:MGI Symbol;Acc:MGI:1913979] | 7,38 | 2,5 |
| ENSMUSG00000068747 | 350,1364497 | 561,3 | 92,7 | 4,785509058 | 0,002883434 | Sort1 | sortilin 1 [Source:MGI Symbol;Acc:MGI:1338015] | 99,11 | 14,5 |
| ENSMUSG00000042626 | 75,57267793 | 140,2 | 42,2 | 4,785610304 | 0,0058277 | Shc1 | src homology 2 domain-containing transforming protein C1 [Source:MGI Symbol;Acc:MGI:98296] | 88,23 | 10,3 |
| ENSMUSG00000028434 | 15,35331794 | 19,9 | 0,9 | 4,786400248 | 0,043946234 | Epb41l4b | erythrocyte membrane protein band 4.1 like 4b [Source:MGI Symbol;Acc:MGI:1859149] | 29,76 | 0,3 |
| ENSMUSG00000002897 | 43,60265644 | 57,0 | 2,9 | 4,78839932 | 0,024997762 | Il17ra | interleukin 17 receptor A [Source:MGI Symbol;Acc:MGI:107399] | 44,49 | 0,6 |
| ENSMUSG00000021281 | 243,032566 | 313,2 | 37,6 | 4,791981566 | 0,009025989 | Tnfaip2 | tumor necrosis factor, alpha-induced protein 2 [Source:MGI Symbol;Acc:MGI:104960] | 47,41 | 1,4 |
| ENSMUSG00000044968 | 64,58155283 | 40,6 | 1,6 | 4,792037616 | 0,035299784 | Napepld | N-acyl phosphatidylethanolamine phospholipase D [Source:MGI Symbol;Acc:MGI:2140885] | 81,07 | 0,9 |
| ENSMUSG00000076435 | 108,9525048 | 159,2 | 8,7 | 4,792039176 | 0,027153105 | Acsf2 | acyl-CoA synthetase family member 2 [Source:MGI Symbol;Acc:MGI:2388287] | 89,35 | 4,7 |
| ENSMUSG00000029641 | 38,5120643 | 65,7 | 20,0 | 4,793203701 | 0,021218563 | Rasl11a | RAS-like, family 11, member A [Source:MGI Symbol;Acc:MGI:1916145] | 98,79 | 12,6 |
| ENSMUSG00000049353 | 44,33791038 | 51,8 | 2,6 | 4,794660148 | 0,031961072 | Rd3 | retinal degeneration 3 [Source:MGI Symbol;Acc:MGI:1921273] | 50,45 | 1,2 |
| ENSMUSG00000030551 | 44,74065321 | 63,6 | 2,5 | 4,796708726 | 0,013932249 | Nr2f2 | nuclear receptor subfamily 2, group F, member 2 [Source:MGI Symbol;Acc:MGI:1352452] | 48,31 | 1,0 |
| ENSMUSG00000008855 | 335,7879027 | 394,7 | 73,9 | 4,797742529 | 0,005255914 | Hdac5 | histone deacetylase 5 [Source:MGI Symbol;Acc:MGI:1333784] | 72,42 | 16,5 |
| ENSMUSG00000022449 | 90,28604269 | 68,8 | 79,3 | 4,797993345 | 0,028041904 | Adamts20 | <e and metallopeptidase (reprolysin type) with thrombospondin type 1 motif, 20 [Source:MGI Symbol;Acc:I | 46,54 | 6,5 |
| ENSMUSG00000042750 | 395,8143897 | 720,8 | 53,1 | 4,805836205 | 0,000190848 | Bex2 | brain expressed X-linked 2 [Source:MGI Symbol;Acc:MGI:1338017] | 90,37 | 33,1 |
| ENSMUSG00000029344 | 126,4188961 | 160,7 | 149,8 | 4,80784873 | 0,017981001 | Tpst2 | protein-tyrosine sulfotransferase 2 [Source:MGI Symbol;Acc:MGI:1309516] | 96,07 | 33,1 |
| ENSMUSG00000021067 | 61,4726531 | 37,4 | 4,6 | 4,808433175 | 0,03394 | Sav1 | salvador homolog 1 (Drosophila) [Source:MGI Symbol;Acc:MGI:1927144] | 86,99 | 4,2 |
| ENSMUSG00000026475 | 42,75255688 | 44,0 | 120,7 | 4,809501269 | 0,049965332 | Rgs16 | regulator of G-protein signaling 16 [Source:MGI Symbol;Acc:MGI:108407] | 94,20 | 29,0 |
| ENSMUSG00000098022 | 23,32704871 | 30,9 | 2,8 | 4,810544551 | 0,039826269 | Zfp82 | zinc finger protein 82 [Source:MGI Symbol;Acc:MGI:1890753] | 55,99 | 1,1 |
| ENSMUSG00000031077 | 38,12906979 | 51,9 | 24,0 | 4,810835501 | 0,04043499 | Fadd | Fas (TNFRSF6)-associated via death domain [Source:MGI Symbol;Acc:MGI:109324] | 70,07 | 5,7 |
| ENSMUSG00000079144 | 70,30593445 | 129,5 | 38,8 | 4,81353852 | 0,022140116 | A130010J15Rik | RIKEN cDNA A130010J15 gene [Source:MGI Symbol;Acc:MGI:2441776] | 76,65 | 5,6 |
| ENSMUSG00000074637 | 798,7731557 | 1447,9 | 52,9 | 4,814030343 | 0,010125358 | Sox2 | SRY (sex determining region Y)-box 2 [Source:MGI Symbol;Acc:MGI:98364] | 100,00 | 28,8 |
| ENSMUSG00000043572 | 37,89905162 | 57,3 | 64,9 | 4,814110345 | 0,032713879 | Pars2 | prolyl-tRNA synthetase (mitochondrial)(putative) [Source:MGI Symbol;Acc:MGI:2386296] | 83,95 | 10,8 |
| ENSMUSG00000021009 | 82,2633511 | 62,0 | 2,8 | 4,814326986 | 0,005892223 | Ptpn21 | protein tyrosine phosphatase, non-receptor type 21 [Source:MGI Symbol;Acc:MGI:1344406] | 85,19 | 0,9 |
| ENSMUSG00000016984 | 91,36886126 | 98,9 | 18,2 | 4,814496906 | 0,031961072 | Etaa1 | Ewing tumor-associated antigen 1 [Source:MGI Symbol;Acc:MGI:1915395] | 80,78 | 4,4 |
| ENSMUSG00000025898 | 44,79253028 | 65,2 | 35,1 | 4,814500645 | 0,015702473 | Cwf19l2 | CWF19-like 2, cell cycle control (S. pombe) [Source:MGI Symbol;Acc:MGI:1918023] | 89,13 | 9,4 |
| ENSMUSG00000033739 | 32,57872299 | 49,4 | 62,9 | 4,815000697 | 0,048796347 | Fkbpl | FK506 binding protein-like [Source:MGI Symbol;Acc:MGI:1932127] | 94,74 | 24,3 |
| ENSMUSG00000030956 | 129,2329731 | 157,7 | 86,3 | 4,815030953 | 0,014620403 | Fam53b | family with sequence similarity 53, member B [Source:MGI Symbol;Acc:MGI:1925188] | 82,94 | 13,4 |
| ENSMUSG00000108530 | 37,80656112 | 56,0 | 18,2 | 4,816254097 | 0,034525326 | RP23-259N3.1 | | #DIV/0! | #DIV/0! |
| ENSMUSG00000086718 | 73,01772451 | 95,2 | 15,7 | 4,816859101 | 0,001750815 | Gm16069 | predicted gene 16069 [Source:MGI Symbol;Acc:MGI:3801823] | 56,33 | 15,4 |
| ENSMUSG00000031539 | 190,6715303 | 356,2 | 19,6 | 4,818118606 | 0,007327727 | Ap3m2 | adaptor-related protein complex 3, mu 2 subunit [Source:MGI Symbol;Acc:MGI:1929214] | 98,71 | 8,7 |
| ENSMUSG00000036983 | 41,06670512 | 37,3 | 1,8 | 4,81956352 | 0,031402449 | Tfb1m | transcription factor B1, mitochondrial [Source:MGI Symbol;Acc:MGI:2146851] | 96,78 | 1,6 |
| ENSMUSG00000043895 | 85,69221867 | 48,2 | 37,9 | 4,820367416 | 0,040927497 | S1pr2 | sphingosine-1-phosphate receptor 2 [Source:MGI Symbol;Acc:MGI:99569] | 93,13 | 10,5 |
| ENSMUSG00000019558 | 165,2963259 | 219,1 | 159,3 | 4,821493132 | 0,016994889 | Slc6a8 | te carrier family 6 (neurotransmitter transporter, creatine), member 8 [Source:MGI Symbol;Acc:MGI:2147 | 91,35 | 26,8 |
| ENSMUSG00000032698 | 25,57570601 | 33,1 | 1,2 | 4,829751491 | 0,026661258 | Lmo2 | LIM domain only 2 [Source:MGI Symbol;Acc:MGI:102811] | 68,08 | 1,2 |
| ENSMUSG00000003762 | 94,4740266 | 92,0 | 72,0 | 4,829769636 | 0,009925804 | Adck4 | aarF domain containing kinase 4 [Source:MGI Symbol;Acc:MGI:1924139] | 90,15 | 23,9 |
| ENSMUSG00000039476 | 44,56196306 | 47,4 | 21,8 | 4,831099259 | 0,045695789 | Prrx2 | paired related homeobox 2 [Source:MGI Symbol;Acc:MGI:98218] | 64,87 | 16,6 |
| ENSMUSG00000027004 | 44,28709451 | 62,1 | 72,0 | 4,831920814 | 0,035465616 | Frzb | frizzled-related protein [Source:MGI Symbol;Acc:MGI:892032] | 75,03 | 15,2 |
| ENSMUSG00000025059 | 325,5065604 | 517,7 | 184,5 | 4,834950967 | 0,011804736 | Gk | glycerol kinase [Source:MGI Symbol;Acc:MGI:106594] | 81,49 | 23,5 |
| ENSMUSG00000053333 | 36,37945013 | 49,9 | 33,3 | 4,837826895 | 0,02374044 | Dis3l2 | DIS3 like 3'-5' exoribonuclease 2 [Source:MGI Symbol;Acc:MGI:2442555] | 88,74 | 10,8 |
| ENSMUSG00000046688 | 447,3657828 | 802,4 | 107,8 | 4,838028648 | 0,003602277 | Tifa | TRAF-interacting protein with forkhead-associated domain [Source:MGI Symbol;Acc:MGI:2182965] | 96,98 | 19,7 |
| ENSMUSG00000003198 | 42,51886153 | 55,9 | 72,1 | 4,839761043 | 0,037450038 | Zfp959 | zinc finger protein 959 [Source:MGI Symbol;Acc:MGI:2385058] | 92,08 | 24,2 |
| ENSMUSG00000104850 | 28,24577998 | 52,1 | 18,2 | 4,843729191 | 0,045522395 | Gm42901 | predicted gene 42901 [Source:MGI Symbol;Acc:MGI:5663038] | 37,98 | 2,2 |
| ENSMUSG00000029360 | 109,5091425 | 145,3 | 93,7 | 4,844248234 | 0,017489351 | Gm9754 | predicted gene 9754 [Source:MGI Symbol;Acc:MGI:3704256] | 53,65 | 7,1 |
| ENSMUSG00000020027 | 20,90521473 | 39,2 | 7,0 | 4,844267382 | 0,042457815 | Socs2 | suppressor of cytokine signaling 2 [Source:MGI Symbol;Acc:MGI:1201787] | 61,48 | 2,0 |
| ENSMUSG00000025078 | 108,9587071 | 84,5 | 21,8 | 4,845033832 | 0,017886688 | Nhlrc2 | NHL repeat containing 2 [Source:MGI Symbol;Acc:MGI:1914116] | 91,27 | 8,0 |
| ENSMUSG00000043388 | 37,61762802 | 35,7 | 3,7 | 4,847660405 | 0,038989198 | Tmem130 | transmembrane protein 130 [Source:MGI Symbol;Acc:MGI:3607706] | 87,97 | 1,8 |
| ENSMUSG00000021360 | 22,53494315 | 41,3 | 6,0 | 4,848471713 | 0,032353047 | Gcnt2 | glucosaminyl (N-acetyl) transferase 2, I-branching enzyme [Source:MGI Symbol;Acc:MGI:1100870] | 41,14 | 0,7 |
| ENSMUSG00000020639 | 193,3671845 | 366,4 | 58,5 | 4,849575216 | 0,025272718 | Pfn4 | profilin family, member 4 [Source:MGI Symbol;Acc:MGI:1920121] | 100,00 | 25,8 |
| ENSMUSG00000033653 | 64,80087205 | 95,6 | 5,2 | 4,850887157 | 0,011734301 | Vps8 | vacuolar protein sorting 8 homolog (S. cerevisiae) [Source:MGI Symbol;Acc:MGI:2146407] | 57,65 | 0,6 |
| ENSMUSG00000020718 | 26,57840872 | 36,8 | 2,8 | 4,851087188 | 0,025643645 | Polg2 | polymerase (DNA directed), gamma 2, accessory subunit [Source:MGI Symbol;Acc:MGI:1354947] | 94,49 | 1,3 |
| ENSMUSG00000042359 | 38,61173489 | 53,1 | 6,5 | 4,851489827 | 0,01527973 | Osbpl6 | oxysterol binding protein-like 6 [Source:MGI Symbol;Acc:MGI:2139014] | 46,22 | 1,3 |
| ENSMUSG00000036478 | 509,9247894 | 916,2 | 87,3 | 4,853231675 | 0,014524967 | Btg1 | B cell translocation gene 1, anti-proliferative [Source:MGI Symbol;Acc:MGI:88215] | 39,74 | 8,3 |
| ENSMUSG00000039168 | 95,32695394 | 123,7 | 36,6 | 4,853357432 | 0,018089692 | Dap | death-associated protein [Source:MGI Symbol;Acc:MGI:1918190] | 76,03 | 19,2 |
| ENSMUSG00000095385 | 31,50601914 | 56,4 | 18,2 | 4,85419438 | 0,031059339 | D630033O11Rik | RIKEN cDNA D630033O11 gene [Source:MGI Symbol;Acc:MGI:2685434] | 64,23 | 17,0 |
| ENSMUSG00000029022 | 77,31857671 | 48,7 | 49,0 | 4,856252116 | 0,03920992 | Miip | migration and invasion inhibitory protein [Source:MGI Symbol;Acc:MGI:106506] | 77,58 | 19,2 |
| ENSMUSG00000027811 | 57,60521391 | 46,4 | 77,6 | 4,856330975 | 0,048183751 | 4930579G24Rik | RIKEN cDNA 4930579G24 gene [Source:MGI Symbol;Acc:MGI:1923189] | 90,91 | 28,2 |
| ENSMUSG00000106847 | 220,2693479 | 406,4 | 173,3 | 4,856808854 | 0,016951686 | Peg13 | paternally expressed 13 [Source:MGI Symbol;Acc:MGI:2663476] | 96,00 | 27,4 |
| ENSMUSG00000103715 | 33,20309296 | 49,0 | 38,8 | 4,861128127 | 0,038989198 | 4933431K14Rik | RIKEN cDNA 4933431K14 gene [Source:MGI Symbol;Acc:MGI:1918545] | 96,94 | 24,9 |
| ENSMUSG00000020660 | 24,03750582 | 41,3 | 7,4 | 4,8645799 | 0,037324514 | Pomc | pro-opiomelanocortin-alpha [Source:MGI Symbol;Acc:MGI:97742] | 59,81 | 9,8 |
| ENSMUSG00000036792 | 66,85145329 | 94,0 | 3,8 | 4,866170379 | 0,003641976 | Mbd5 | methyl-CpG binding domain protein 5 [Source:MGI Symbol;Acc:MGI:2138934] | 53,78 | 0,7 |
| ENSMUSG00000038534 | 35,23490095 | 48,1 | 20,3 | 4,866641319 | 0,024997762 | Osbpl7 | oxysterol binding protein-like 7 [Source:MGI Symbol;Acc:MGI:1918490] | 84,81 | 7,1 |
| ENSMUSG00000034854 | 72,04151752 | 46,6 | 7,9 | 4,868595182 | 0,03703421 | Mfsd12 | major facilitator superfamily domain containing 12 [Source:MGI Symbol;Acc:MGI:3604804] | 83,42 | 2,3 |
| ENSMUSG00000024063 | 43,38210378 | 74,2 | 9,1 | 4,872869797 | 0,008220062 | Lbh | limb-bud and heart [Source:MGI Symbol;Acc:MGI:1925139] | 90,12 | 5,6 |
| ENSMUSG00000056234 | 72,9064558 | 119,7 | 16,3 | 4,874178421 | 0,008966028 | Ncoa4 | nuclear receptor coactivator 4 [Source:MGI Symbol;Acc:MGI:1350932] | 39,05 | 4,8 |
| ENSMUSG00000029669 | 803,3551436 | 1211,7 | 308,3 | 4,874627621 | 0,037161083 | Tspan12 | tetraspanin 12 [Source:MGI Symbol;Acc:MGI:1889818] | 88,14 | 26,5 |
| ENSMUSG00000051435 | 68,18062553 | 101,8 | 30,9 | 4,874877296 | 0,009281644 | Fhad1 | forkhead-associated (FHA) phosphopeptide binding domain 1 [Source:MGI Symbol;Acc:MGI:1920323] | 68,62 | 3,9 |
| ENSMUSG00000022898 | 612,9403905 | 947,3 | 148,5 | 4,875269003 | 0,012119144 | Dscr3 | Down syndrome critical region gene 3 [Source:MGI Symbol;Acc:MGI:1206040] | 96,81 | 31,6 |
| ENSMUSG00000107689 | 37,69699604 | 51,0 | 45,4 | 4,876842891 | 0,0447485 | Gm44386 | predicted gene, 44386 [Source:MGI Symbol;Acc:MGI:5690778] | #DIV/0! | #DIV/0! |
| ENSMUSG00000018999 | 203,0693152 | 284,0 | 187,4 | 4,878654905 | 0,017476584 | Slc35b4 | solute carrier family 35, member B4 [Source:MGI Symbol;Acc:MGI:1931249] | 64,37 | 15,2 |
| ENSMUSG00000024222 | 104,3662389 | 172,4 | 126,7 | 4,878754621 | 0,015956754 | Fkbp5 | FK506 binding protein 5 [Source:MGI Symbol;Acc:MGI:104670] | 81,96 | 21,6 |
| ENSMUSG00000090173 | 8,688404686 | 19,9 | 1,0 | 4,879733204 | 0,046236448 | Fbxw10 | F-box and WD-40 domain protein 10 [Source:MGI Symbol;Acc:MGI:3052463] | 66,07 | 0,4 |
| ENSMUSG00000046822 | 102,7710769 | 170,0 | 93,7 | 4,881198736 | 0,010077387 | Slc39a3 | solute carrier family 39 (zinc transporter), member 3 [Source:MGI Symbol;Acc:MGI:2147269] | 96,95 | 21,9 |
| ENSMUSG00000004151 | 92,34141156 | 143,3 | 58,8 | 4,883520133 | 0,017970476 | Etv1 | ets variant 1 [Source:MGI Symbol;Acc:MGI:99254] | 47,62 | 7,5 |

| | | | | | | | | | |
|---|---|---|---|---|---|---|---|---|---|
| ENSMUSG00000039298 | 88,13935781 | 173,8 | 26,2 | 4,883904128 | 0,004276605 | Cdk5rap2 | CDK5 regulatory subunit associated protein 2 [Source:MGI Symbol;Acc:MGI:2384875] | 45,88 | 2,2 |
| ENSMUSG00000042476 | 13,5914192 | 21,9 | 0,6 | 4,884636565 | 0,032399561 | Abcb4 | ATP-binding cassette, sub-family B (MDR/TAP), member 4 [Source:MGI Symbol;Acc:MGI:97569] | 53,39 | 0,6 |
| ENSMUSG00000027078 | 40,15969712 | 56,0 | 46,6 | 4,886464598 | 0,033653786 | Ube2l6 | ubiquitin-conjugating enzyme E2L 6 [Source:MGI Symbol;Acc:MGI:1914500] | 79,35 | 23,2 |
| ENSMUSG00000047022 | 89,9348692 | 126,5 | 6,6 | 4,887622692 | 0,005911767 | Mipol1 | mirror-image polydactyly gene 1 homolog (human) [Source:MGI Symbol;Acc:MGI:1920740] | 71,00 | 3,7 |
| ENSMUSG00000040473 | 58,54553391 | 60,4 | 2,2 | 4,890280401 | 0,006715922 | Cfap69 | cilia and flagella associated protein 69 [Source:MGI Symbol;Acc:MGI:2443778] | 62,55 | 0,8 |
| ENSMUSG00000036528 | 99,54759744 | 105,6 | 7,5 | 4,891929646 | 0,015421831 | Ppfibp2 | PTPRF interacting protein, binding protein 2 (liprin beta 2) [Source:MGI Symbol;Acc:MGI:894649] | 66,04 | 2,0 |
| ENSMUSG00000003421 | 221,006492 | 296,5 | 30,5 | 4,892094569 | 5,2044E-05 | Nosip | nitric oxide synthase interacting protein [Source:MGI Symbol;Acc:MGI:1913644] | 93,42 | 15,8 |
| ENSMUSG00000034168 | 325,1908621 | 623,2 | 39,1 | 4,893084073 | 0,000387247 | Irf2bpl | interferon regulatory factor 2 binding protein-like [Source:MGI Symbol;Acc:MGI:2442463] | 98,37 | 12,1 |
| ENSMUSG00000094372 | 34,99630809 | 51,9 | 17,6 | 4,893893275 | 0,04134294 | Gm21839 | predicted gene, 21839 [Source:MGI Symbol;Acc:MGI:5434003] | #DIV/0! | #DIV/0! |
| ENSMUSG00000013921 | 988,2428338 | 1809,4 | 121,3 | 4,89527626 | 0,02958134 | Clip3 | CAP-GLY domain containing linker protein 3 [Source:MGI Symbol;Acc:MGI:1923936] | 99,90 | 34,4 |
| ENSMUSG00000042099 | 100,3023263 | 56,7 | 186,4 | 4,897944294 | 0,045237134 | Kank3 | KN motif and ankyrin repeat domains 3 [Source:MGI Symbol;Acc:MGI:1098615] | 68,89 | 22,4 |
| ENSMUSG00000021470 | 61,21747951 | 78,4 | 3,8 | 4,898896442 | 0,00713294 | Ercc6l2 | cross-complementing rodent repair deficiency, complementation group 6 like 2 [Source:MGI Symbol;Acc: | 71,54 | 0,5 |
| ENSMUSG00000021461 | 45,60294138 | 69,2 | 73,0 | 4,89975586 | 0,027754602 | Fancc | Fanconi anemia, complementation group C [Source:MGI Symbol;Acc:MGI:95480] | 55,24 | 11,6 |
| ENSMUSG00000060681 | 149,7774848 | 198,0 | 178,8 | 4,900457798 | 0,022989747 | Slc9a6 | solute carrier family 9 (sodium/hydrogen exchanger), member 6 [Source:MGI Symbol;Acc:MGI:2443511] | 93,67 | 6,5 |
| ENSMUSG00000034793 | 189,3021808 | 238,3 | 37,6 | 4,900839951 | 0,008415371 | G6pc3 | glucose 6 phosphatase, catalytic, 3 [Source:MGI Symbol;Acc:MGI:1915651] | 91,78 | 24,2 |
| ENSMUSG00000085338 | 40,23969719 | 65,6 | 29,7 | 4,902100042 | 0,045237975 | 2410004I01Rik | RIKEN cDNA 2410004I01 gene [Source:MGI Symbol;Acc:MGI:1916989] | #DIV/0! | #DIV/0! |
| ENSMUSG00000092118 | 31,57187697 | 55,4 | 37,0 | 4,902740376 | 0,024493876 | Fancf | Fanconi anemia, complementation group F [Source:MGI Symbol;Acc:MGI:3689889] | 97,79 | 23,6 |
| ENSMUSG00000003123 | 73,99507809 | 100,9 | 3,5 | 4,903768418 | 0,000146715 | Lipe | lipase, hormone sensitive [Source:MGI Symbol;Acc:MGI:96790] | 56,06 | 0,6 |
| ENSMUSG00000032495 | 75,68440241 | 141,8 | 19,7 | 4,904441255 | 0,013322569 | Lrrc2 | leucine rich repeat containing 2 [Source:MGI Symbol;Acc:MGI:1921499] | 69,41 | 6,6 |
| ENSMUSG00000109129 | 67,10863022 | 42,8 | 3,7 | 4,904445738 | 0,019947179 | RP23-414I24.8 | | 97,61 | 3,3 |
| ENSMUSG00000036006 | 70,09033361 | 138,5 | 9,2 | 4,906045532 | 0,003673533 | Fam65b | family with sequence similarity 65, member B [Source:MGI Symbol;Acc:MGI:2444879] | 57,92 | 1,5 |
| ENSMUSG00000031709 | 110,7314027 | 89,1 | 3,1 | 4,908151504 | 0,018490227 | Tbc1d9 | TBC1 domain family, member 9 [Source:MGI Symbol;Acc:MGI:1918560] | 87,52 | 1,6 |
| ENSMUSG00000029136 | 46,71857762 | 51,1 | 10,3 | 4,908361587 | 0,033841736 | Rbks | ribokinase [Source:MGI Symbol;Acc:MGI:1918586] | 98,14 | 7,7 |
| ENSMUSG00000050428 | 45,94051888 | 44,5 | 5,0 | 4,909224935 | 0,01983382 | Fbxo46 | F-box protein 46 [Source:MGI Symbol;Acc:MGI:2444918] | 67,03 | 0,9 |
| ENSMUSG00000035067 | 38,376673 | 54,4 | 2,6 | 4,909599474 | 0,038476201 | Xkr6 | X Kell blood group precursor related family member 6 homolog [Source:MGI Symbol;Acc:MGI:2447765] | 25,17 | 1,5 |
| ENSMUSG00000027809 | 464,0805089 | 759,5 | 270,7 | 4,910085638 | 0,022041571 | Etfdh | electron transferring flavoprotein, dehydrogenase [Source:MGI Symbol;Acc:MGI:106100] | 95,50 | 11,3 |
| ENSMUSG00000109128 | 59,80978544 | 80,2 | 21,2 | 4,910376796 | 0,010820437 | RP23-106P7.5 | | 94,01 | 28,0 |
| ENSMUSG00000027573 | 125,3902639 | 186,6 | 19,8 | 4,911932318 | 0,006045628 | Gid8 | GID complex subunit 8 [Source:MGI Symbol;Acc:MGI:1923675] | 65,67 | 8,6 |
| ENSMUSG00000097574 | 72,47273985 | 69,0 | 58,7 | 4,912249723 | 0,041784204 | C920006O11Rik | RIKEN cDNA C920006O11 gene [Source:MGI Symbol;Acc:MGI:2443759] | #DIV/0! | #DIV/0! |
| ENSMUSG00000086601 | 28,46246265 | 49,0 | 10,2 | 4,912707216 | 0,037909203 | Rapgef3os1 | ap guanine nucleotide exchange factor (GEF) 3, opposite strand 1 [Source:MGI Symbol;Acc:MGI:382658 | 64,87 | 13,4 |
| ENSMUSG00000097597 | 16,70135269 | 34,2 | 5,0 | 4,913020796 | 0,048733912 | Gm26674 | predicted gene, 26674 [Source:MGI Symbol;Acc:MGI:5477168] | #DIV/0! | #DIV/0! |
| ENSMUSG00000031627 | 134,6013803 | 167,4 | 63,8 | 4,913265115 | 0,010559107 | Irf2 | interferon regulatory factor 2 [Source:MGI Symbol;Acc:MGI:96591] | 86,82 | 18,8 |
| ENSMUSG00000040111 | 70,82360735 | 106,5 | 17,2 | 4,91517434 | 0,019275359 | Gramd1b | GRAM domain containing 1B [Source:MGI Symbol;Acc:MGI:1925037] | 61,56 | 3,2 |
| ENSMUSG00000052105 | 47,46703455 | 99,9 | 4,7 | 4,915610134 | 0,00102309 | Mtcl1 | microtubule crosslinking factor 1 [Source:MGI Symbol;Acc:MGI:1915867] | 80,81 | 0,0 |
| ENSMUSG00000109274 | 47,93638994 | 51,7 | 28,7 | 4,917220884 | 0,031954791 | RP23-419K14.8 | | 69,31 | 12,3 |
| ENSMUSG00000025880 | 32,92870094 | 51,6 | 6,0 | 4,919745694 | 0,013375274 | Smad7 | SMAD family member 7 [Source:MGI Symbol;Acc:MGI:1100518] | 74,21 | 1,6 |
| ENSMUSG00000073102 | 22,68666248 | 42,4 | 3,7 | 4,923340731 | 0,018929695 | Drc1 | dynein regulatory complex subunit 1 [Source:MGI Symbol;Acc:MGI:2685906] | 90,13 | 1,7 |
| ENSMUSG00000037134 | 43,31394229 | 46,0 | 3,0 | 4,925455255 | 0,019153024 | Prmt10 | protein arginine methyltransferase 10 (putative) [Source:MGI Symbol;Acc:MGI:2142651] | 83,18 | 0,9 |
| ENSMUSG00000075271 | 15,72888164 | 31,4 | 1,2 | 4,925755666 | 0,023671899 | Ttc30a1 | tetratricopeptide repeat domain 30A1 [Source:MGI Symbol;Acc:MGI:1926052] | 44,13 | 0,4 |
| ENSMUSG00000022906 | 89,37743659 | 77,0 | 64,7 | 4,927264489 | 0,014785672 | Parp9 | poly (ADP-ribose) polymerase family, member 9 [Source:MGI Symbol;Acc:MGI:1933117] | 79,48 | 16,8 |
| ENSMUSG00000063458 | 21,74250602 | 36,8 | 3,0 | 4,928387787 | 0,031998123 | 1700112E06Rik | RIKEN cDNA 1700112E06 gene [Source:MGI Symbol;Acc:MGI:1923883] | 72,03 | 2,1 |
| ENSMUSG00000060568 | 18,67292776 | 40,4 | 3,0 | 4,929299422 | 0,01751344 | Fam78b | family with sequence similarity 78, member B [Source:MGI Symbol;Acc:MGI:2443050] | 54,58 | 0,4 |
| ENSMUSG00000030518 | 24,61861859 | 37,5 | 6,0 | 4,929334486 | 0,04158631 | Fam189a1 | family with sequence similarity 189, member A1 [Source:MGI Symbol;Acc:MGI:1917888] | 74,77 | 1,5 |
| ENSMUSG00000059316 | 79,88529778 | 83,1 | 17,6 | 4,929335482 | 0,005792904 | Slc27a4 | solute carrier family 27 (fatty acid transporter), member 4 [Source:MGI Symbol;Acc:MGI:1347347] | 78,38 | 5,5 |
| ENSMUSG00000087006 | 48,52324928 | 94,2 | 6,9 | 4,934151364 | 0,01835823 | Gm13889 | predicted gene 13889 [Source:MGI Symbol;Acc:MGI:3652053] | 90,23 | 6,9 |
| ENSMUSG00000022159 | 246,9786917 | 380,5 | 62,7 | 4,939734914 | 0,032017121 | Rab2b | RAB2B, member RAS oncogene family [Source:MGI Symbol;Acc:MGI:1923588] | 96,10 | 15,1 |
| ENSMUSG00000023079 | 51,49050057 | 67,8 | 13,0 | 4,940479666 | 0,008348916 | Gtf2ird1 | general transcription factor II I repeat domain-containing 1 [Source:MGI Symbol;Acc:MGI:1861942] | 79,51 | 2,3 |
| ENSMUSG00000014850 | 41,20467746 | 70,2 | 14,8 | 4,943082595 | 0,006635377 | Msh3 | mutS homolog 3 (E. coli) [Source:MGI Symbol;Acc:MGI:109519] | 89,54 | 3,9 |
| ENSMUSG00000074794 | 240,8956876 | 432,1 | 98,0 | 4,943730217 | 0,014768969 | Arrdc3 | arrestin domain containing 3 [Source:MGI Symbol;Acc:MGI:2145242] | 98,65 | 26,6 |
| ENSMUSG00000056536 | 23,42028434 | 65,8 | 2,5 | 4,943974902 | 0,00433647 | Pign | phosphatidylinositol glycan anchor biosynthesis, class N [Source:MGI Symbol;Acc:MGI:1351629] | 60,50 | 0,4 |
| ENSMUSG00000015970 | 20,09566221 | 33,7 | 1,8 | 4,944513481 | 0,024031707 | Chdh | choline dehydrogenase [Source:MGI Symbol;Acc:MGI:1860776] | 60,42 | 0,3 |
| ENSMUSG00000045078 | 151,3126052 | 221,7 | 29,7 | 4,944773933 | 0,026765281 | Rnf216 | ring finger protein 216 [Source:MGI Symbol;Acc:MGI:1344349] | 98,99 | 7,0 |
| ENSMUSG00000029627 | 58,4613517 | 67,3 | 90,8 | 4,944808543 | 0,04083678 | Zkscan14 | zinc finger with KRAB and SCAN domains 14 [Source:MGI Symbol;Acc:MGI:1914485] | 60,97 | 16,6 |
| ENSMUSG00000027340 | 299,1670621 | 314,8 | 125,4 | 4,945897321 | 0,009398001 | Slc23a2 | solute carrier family 23 (nucleobase transporters), member 2 [Source:MGI Symbol;Acc:MGI:1859682] | 98,12 | 17,7 |
| ENSMUSG00000070287 | 40,98270364 | 65,8 | 48,9 | 4,947298639 | 0,027588436 | Slc35g2 | solute carrier family 35, member G2 [Source:MGI Symbol;Acc:MGI:2685365] | 99,22 | 18,8 |
| ENSMUSG00000038241 | 72,108833 | 82,2 | 43,4 | 4,948907748 | 0,014604568 | Cep250 | centrosomal protein 250 [Source:MGI Symbol;Acc:MGI:108084] | 57,46 | 4,2 |
| ENSMUSG00000022528 | 194,0401597 | 372,5 | 47,2 | 4,95043057 | 0,016481368 | Hes1 | hairy and enhancer of split 1 (Drosophila) [Source:MGI Symbol;Acc:MGI:104853] | 99,91 | 30,0 |
| ENSMUSG00000054679 | 34,41789789 | 30,7 | 2,0 | 4,952491395 | 0,037914345 | Srsf12 | serine/arginine-rich splicing factor 12 [Source:MGI Symbol;Acc:MGI:2661424] | 72,15 | 0,4 |
| ENSMUSG00000053004 | 52,61330539 | 67,5 | 41,8 | 4,954133752 | 0,02650583 | Hrh1 | histamine receptor H1 [Source:MGI Symbol;Acc:MGI:107619] | 64,28 | 6,0 |
| ENSMUSG00000039456 | 123,1236521 | 93,9 | 74,4 | 4,954194088 | 0,014210371 | Morc3 | microrchidia 3 [Source:MGI Symbol;Acc:MGI:2136841] | 93,39 | 9,7 |
| ENSMUSG00000059810 | 178,492042 | 301,7 | 20,6 | 4,954423598 | 0,008924977 | Rgs3 | regulator of G-protein signaling 3 [Source:MGI Symbol;Acc:MGI:1354734] | 60,13 | 2,5 |
| ENSMUSG00000050530 | 178,0493471 | 258,3 | 18,5 | 4,955284781 | 0,007374158 | Fam171a1 | family with sequence similarity 171, member A1 [Source:MGI Symbol;Acc:MGI:2442917] | 91,68 | 3,4 |
| ENSMUSG00000020876 | 222,1817665 | 259,9 | 49,1 | 4,958123416 | 0,009540698 | Snx11 | sorting nexin 11 [Source:MGI Symbol;Acc:MGI:1921729] | 96,95 | 20,9 |
| ENSMUSG00000097047 | 42,48949855 | 61,3 | 22,4 | 4,959319323 | 0,031057321 | 1110020A21Rik | RIKEN cDNA 1110020A21 gene [Source:MGI Symbol;Acc:MGI:1915781] | #DIV/0! | #DIV/0! |
| ENSMUSG00000054162 | 27,61178821 | 47,4 | 1,6 | 4,96149722 | 0,004936822 | Spock3 | sparc/osteonectin, cwcv and kazal-like domains proteoglycan 3 [Source:MGI Symbol;Acc:MGI:1920152] | 81,76 | 1,1 |
| ENSMUSG00000025759 | 130,5473789 | 74,5 | 138,6 | 4,962616774 | 0,046525682 | Mfsd8 | major facilitator superfamily domain containing 8 [Source:MGI Symbol;Acc:MGI:1919425] | 86,05 | 26,1 |
| ENSMUSG00000086472 | 33,87190488 | 36,9 | 1,8 | 4,964703038 | 0,014866324 | Gm16172 | predicted gene 16172 [Source:MGI Symbol;Acc:MGI:3801735] | 42,28 | 3,9 |
| ENSMUSG00000041112 | 43,1485144 | 49,5 | 62,8 | 4,967623196 | 0,04102085 | Elmo1 | engulfment and cell motility 1 [Source:MGI Symbol;Acc:MGI:2153044] | 75,49 | 14,8 |
| ENSMUSG00000036339 | 102,8205898 | 71,7 | 3,4 | 4,969139405 | 0,026547142 | Tmem260 | transmembrane protein 260 [Source:MGI Symbol;Acc:MGI:2443219] | 77,50 | 0,4 |
| ENSMUSG00000045333 | 210,111791 | 336,5 | 71,4 | 4,969949251 | 0,020710182 | Zfp423 | zinc finger protein 423 [Source:MGI Symbol;Acc:MGI:1891217] | 90,50 | 19,0 |
| ENSMUSG00000056258 | 94,93711197 | 125,9 | 134,3 | 4,9699793 | 0,026469842 | Kcnq3 | potassium voltage-gated channel, subfamily Q, member 3 [Source:MGI Symbol;Acc:MGI:1336181] | 67,16 | 4,2 |
| ENSMUSG00000047759 | 40,38200081 | 63,1 | 23,0 | 4,972215917 | 0,032493928 | Hs3st3a1 | heparan sulfate (glucosamine) 3-O-sulfotransferase 3A1 [Source:MGI Symbol;Acc:MGI:1333861] | 84,19 | 7,9 |
| ENSMUSG00000034751 | 79,72122035 | 141,7 | 34,9 | 4,97705242 | 0,0080152 | Mast4 | microtubule associated serine/threonine kinase family member 4 [Source:MGI Symbol;Acc:MGI:1918885] | 66,39 | 1,6 |
| ENSMUSG00000074398 | 54,4675328 | 63,7 | 24,2 | 4,977385784 | 0,04070987 | Gm15441 | predicted gene 15441 [Source:MGI Symbol;Acc:MGI:3641753] | 62,29 | 8,7 |
| ENSMUSG00000086003 | 39,05000195 | 61,5 | 13,9 | 4,977499935 | 0,034670823 | B230206L02Rik | RIKEN cDNA B230206L02 gene [Source:MGI Symbol;Acc:MGI:1924598] | #DIV/0! | #DIV/0! |
| ENSMUSG00000024420 | 232,9175356 | 406,0 | 32,3 | 4,977592939 | 0,000415246 | Zfp521 | zinc finger protein 521 [Source:MGI Symbol;Acc:MGI:95459] | 94,09 | 2,7 |
| ENSMUSG00000036580 | 273,7744354 | 348,4 | 339,1 | 4,985410606 | 0,010213022 | Spg20 | pastic paraplegia 20, spartin (Troyer syndrome) homolog (human) [Source:MGI Symbol;Acc:MGI:213980 | 73,18 | 24,1 |
| ENSMUSG00000022369 | 14,23540789 | 22,3 | 0,9 | 4,986173557 | 0,030135494 | Mtbp | Mdm2, transformed 3T3 cell double minute p53 binding protein [Source:MGI Symbol;Acc:MGI:2146005] | 44,97 | 0,4 |
| ENSMUSG00000029162 | 103,2393487 | 118,1 | 61,9 | 4,986591259 | 0,004455351 | Khk | ketohexokinase [Source:MGI Symbol;Acc:MGI:1096353] | 85,18 | 24,3 |
| ENSMUSG00000046572 | 11,62630714 | 22,6 | 0,9 | 4,99066407 | 0,046236448 | Zfp518b | zinc finger protein 518B [Source:MGI Symbol;Acc:MGI:2140750] | 43,80 | 0,3 |
| ENSMUSG00000106892 | 40,90224288 | 48,1 | 3,6 | 4,992224353 | 0,02470018 | Gm42791 | predicted gene 42791 [Source:MGI Symbol;Acc:MGI:5662928] | 100,00 | 1,4 |
| ENSMUSG00000040253 | 42,57379924 | 58,4 | 72,1 | 4,994307213 | 0,030961686 | Gbp7 | guanylate binding protein 7 [Source:MGI Symbol;Acc:MGI:2444421] | 68,72 | 10,7 |
| ENSMUSG00000030727 | 74,97109682 | 60,9 | 121,7 | 4,994808909 | 0,020784121 | Rabep2 | rabaptin, RAB GTPase binding effector protein 2 [Source:MGI Symbol;Acc:MGI:1917564] | 93,57 | 23,2 |
| ENSMUSG00000033295 | 223,0234843 | 189,3 | 99,0 | 4,995368536 | 0,022538154 | Ptprf | protein tyrosine phosphatase, receptor type, F [Source:MGI Symbol;Acc:MGI:102695] | 95,78 | 8,9 |
| ENSMUSG00000027194 | 27,65794307 | 46,0 | 16,0 | 4,996591341 | 0,038475475 | Ttc17 | tetratricopeptide repeat domain 17 [Source:MGI Symbol;Acc:MGI:1921819] | 76,66 | 1,9 |
| ENSMUSG00000063632 | 42,30726288 | 42,0 | 6,5 | 4,998233873 | 0,03496998 | Sox11 | SRY (sex determining region Y)-box 11 [Source:MGI Symbol;Acc:MGI:98359] | 44,04 | 0,3 |
| ENSMUSG00000062031 | 149,2267552 | 218,5 | 19,6 | 4,99950808 | 0,022294132 | Athl1 | ATH1, acid trehalase-like 1 (yeast) [Source:MGI Symbol;Acc:MGI:2444047] | 99,45 | 4,6 |
| ENSMUSG00000020961 | 57,23685893 | 61,0 | 84,1 | 5,000696893 | 0,044220312 | Ston2 | stonin 2 [Source:MGI Symbol;Acc:MGI:1918272] | 69,33 | 11,7 |
| ENSMUSG00000004508 | 76,92668821 | 63,9 | 243,3 | 5,000994331 | 0,049495484 | Gab2 | growth factor receptor bound protein 2-associated protein 2 [Source:MGI Symbol;Acc:MGI:1333854] | 76,04 | 26,1 |
| ENSMUSG00000022469 | 351,4399669 | 629,8 | 133,4 | 5,005476843 | 0,019553887 | Rapgef3 | Rap guanine nucleotide exchange factor (GEF) 3 [Source:MGI Symbol;Acc:MGI:2441741] | 95,18 | 25,5 |
| ENSMUSG00000025226 | 77,54824858 | 131,3 | 31,9 | 5,005793633 | 0,038936575 | Fbxl15 | F-box and leucine-rich repeat protein 15 [Source:MGI Symbol;Acc:MGI:1915681] | 99,60 | 25,9 |
| ENSMUSG00000024909 | 206,6881228 | 300,8 | 45,4 | 5,007123806 | 0,016761689 | Efemp2 | ermal growth factor-containing fibulin-like extracellular matrix protein 2 [Source:MGI Symbol;Acc:MGI:1891 | 81,94 | 17,8 |
| ENSMUSG00000031333 | 196,3292623 | 119,3 | 33,9 | 5,011115761 | 0,027862291 | Abcb7 | ATP-binding cassette, sub-family B (MDR/TAP), member 7 [Source:MGI Symbol;Acc:MGI:109533] | 91,32 | 6,0 |
| ENSMUSG00000085037 | 55,11168843 | 62,9 | 110,0 | 5,011559939 | 0,038847525 | 4933421O10Rik | RIKEN cDNA 4933421O10 gene [Source:MGI Symbol;Acc:MGI:1918323] | #DIV/0! | #DIV/0! |
| ENSMUSG00000062991 | 19,0402979 | 52,5 | 1,6 | 5,014128157 | 0,007251624 | Nrg1 | neuregulin 1 [Source:MGI Symbol;Acc:MGI:96083] | 50,22 | 0,3 |
| ENSMUSG00000101225 | 75,64287255 | 58,6 | 56,9 | 5,014511255 | 0,046809582 | 1700008J07Rik | RIKEN cDNA 1700008J07 gene [Source:MGI Symbol;Acc:MGI:1916610] | 77,82 | 9,0 |
| ENSMUSG00000020329 | 45,57717099 | 54,8 | 49,6 | 5,015206035 | 0,046454798 | Polrmt | polymerase (RNA) mitochondrial (DNA directed) [Source:MGI Symbol;Acc:MGI:1915843] | 92,30 | 20,0 |
| ENSMUSG00000041936 | 107,0486017 | 54,7 | 107,8 | 5,016397331 | 0,038235961 | Agrn | agrin [Source:MGI Symbol;Acc:MGI:87961] | 61,01 | 11,2 |
| ENSMUSG00000031143 | 40,17764565 | 45,6 | 3,7 | 5,018339316 | 0,018715896 | Ccdc22 | coiled-coil domain containing 22 [Source:MGI Symbol;Acc:MGI:1859608] | 90,14 | 3,2 |
| ENSMUSG00000038240 | 48,49623327 | 65,0 | 59,3 | 5,019251592 | 0,032204168 | Pdss2 | prenyl (solanesyl) diphosphate synthase, subunit 2 [Source:MGI Symbol;Acc:MGI:1918615] | 87,98 | 19,2 |
| ENSMUSG00000020246 | 140,8149238 | 215,2 | 6,9 | 5,019895336 | 0,00570154 | Hcfc2 | host cell factor C2 [Source:MGI Symbol;Acc:MGI:1915183] | 95,57 | 1,9 |
| ENSMUSG00000039904 | 188,9128993 | 283,5 | 102,5 | 5,022993061 | 0,028969349 | Gpr37 | G protein-coupled receptor 37 [Source:MGI Symbol;Acc:MGI:1313297] | 71,74 | 17,9 |
| ENSMUSG00000050556 | 86,82348715 | 54,7 | 131,7 | 5,023227544 | 0,04126323 | Kcnb1 | otassium voltage gated channel, Shab-related subfamily, member 1 [Source:MGI Symbol;Acc:MGI:9666 | 49,27 | 8,6 |
| ENSMUSG00000017747 | 43,10746748 | 65,6 | 41,2 | 5,024973368 | 0,029860531 | Ghdc | GH3 domain containing [Source:MGI Symbol;Acc:MGI:1931556] | 95,79 | 11,2 |
| ENSMUSG00000026585 | 271,7138312 | 259,8 | 96,3 | 5,026945278 | 0,013959093 | Kifap3 | kinesin-associated protein 3 [Source:MGI Symbol;Acc:MGI:107566] | 97,41 | 7,3 |
| ENSMUSG00000035517 | 202,7815108 | 269,4 | 93,1 | 5,029906551 | 0,042593161 | Tdrd7 | tudor domain containing 7 [Source:MGI Symbol;Acc:MGI:2140279] | 94,17 | 23,0 |
| ENSMUSG00000023009 | 12,92333989 | 22,1 | 1,0 | 5,03021211 | 0,043929689 | Nckap5l | NCK-associated protein 5-like [Source:MGI Symbol;Acc:MGI:3609653] | 52,40 | 0,4 |
| ENSMUSG00000085242 | 48,13919247 | 41,5 | 4,6 | 5,030996913 | 0,038574663 | Gm11537 | predicted gene 11537 [Source:MGI Symbol;Acc:MGI:3705160] | 86,51 | 9,9 |
| ENSMUSG00000030316 | 84,40379784 | 78,3 | 41,6 | 5,032473323 | 0,012980249 | Tamm41 | ondrial translocator assembly and maintenance protein, homolog (S. cerevisiae) [Source:MGI Symbol;Acc | 99,46 | 28,8 |
| ENSMUSG00000030768 | 49,81540085 | 47,8 | 32,3 | 5,033021371 | 0,037377105 | Disp1 | dispatched homolog 1 (Drosophila) [Source:MGI Symbol;Acc:MGI:1916147] | 74,71 | 7,2 |
| ENSMUSG00000026577 | 202,6525291 | 160,7 | 225,2 | 5,033231322 | 0,01696342 | Blzf1 | basic leucine zipper nuclear factor 1 [Source:MGI Symbol;Acc:MGI:1201607] | 73,76 | 19,5 |
| ENSMUSG00000084917 | 25,33907406 | 33,0 | 1,8 | 5,034473341 | 0,026488139 | Gm17477 | predicted gene, 17477 [Source:MGI Symbol;Acc:MGI:4937111] | #DIV/0! | #DIV/0! |
| ENSMUSG00000025086 | 60,95195757 | 72,0 | 49,9 | 5,036715015 | 0,011455016 | Trub1 | TruB pseudouridine (psi) synthase homolog 1 (E. coli) [Source:MGI Symbol;Acc:MGI:1919383] | 29,01 | 3,1 |
| ENSMUSG00000005907 | 46,5577745 | 60,7 | 32,7 | 5,036969934 | 0,032425801 | Pex1 | peroxisomal biogenesis factor 1 [Source:MGI Symbol;Acc:MGI:1918632] | 83,92 | 2,2 |
| ENSMUSG00000040302 | 64,90473698 | 75,5 | 23,0 | 5,038366664 | 0,024990021 | Rbm48 | RNA binding motif protein 48 [Source:MGI Symbol;Acc:MGI:2442653] | 87,89 | 9,2 |
| ENSMUSG00000086392 | 25,89424891 | 54,3 | 8,0 | 5,039055713 | 0,020595116 | Mccc1os | ethylcrotonoyl-Coenzyme A carboxylase 1 (alpha), opposite strand [Source:MGI Symbol;Acc:MGI:359066 | 47,68 | 0,6 |
| ENSMUSG00000068742 | 588,4488923 | 915,0 | 148,2 | 5,039207022 | 0,004170187 | Cry2 | cryptochrome 2 (photolyase-like) [Source:MGI Symbol;Acc:MGI:1270859] | 91,55 | 26,6 |
| ENSMUSG00000004044 | 48,37787297 | 82,8 | 3,1 | 5,039285971 | 0,013593364 | Ptrf | polymerase I and transcript release factor [Source:MGI Symbol;Acc:MGI:1277968] | 87,06 | 2,4 |
| ENSMUSG00000097080 | 50,96992471 | 75,0 | 27,8 | 5,039427272 | 0,020740575 | 1700086O06Rik | RIKEN cDNA 1700086O06 gene [Source:MGI Symbol;Acc:MGI:1920766] | #DIV/0! | #DIV/0! |
| ENSMUSG00000030672 | 10,66054049 | 22,9 | 0,9 | 5,04020716 | 0,039810883 | Mylpf | myosin light chain, phosphorylatable, fast skeletal muscle [Source:MGI Symbol;Acc:MGI:97273] | 80,80 | 0,0 |
| ENSMUSG00000097145 | 55,96603475 | 42,7 | 71,2 | 5,04031907 | 0,047974717 | 9230114K14Rik | RIKEN cDNA 9230114K14 gene [Source:MGI Symbol;Acc:MGI:3041212] | #DIV/0! | #DIV/0! |
| ENSMUSG00000102615 | 55,13426038 | 74,9 | 58,7 | 5,044377197 | 0,026466872 | Gm37844 | predicted gene, 37844 [Source:MGI Symbol;Acc:MGI:5611072] | 38,55 | 3,6 |
| ENSMUSG00000026339 | 52,03710051 | 66,9 | 64,2 | 5,045137831 | 0,031656379 | Ccdc93 | coiled-coil domain containing 93 [Source:MGI Symbol;Acc:MGI:1918079] | 77,95 | 7,5 |
| ENSMUSG00000052302 | 43,32331249 | 43,6 | 114,8 | 5,045388468 | 0,048935638 | Tbc1d30 | TBC1 domain family, member 30 [Source:MGI Symbol;Acc:MGI:1921944] | 57,28 | 17,8 |
| ENSMUSG00000039738 | 37,06418687 | 62,9 | 53,9 | 5,045541244 | 0,022959296 | Slx4 | .X4 structure-specific endonuclease subunit homolog (S. cerevisiae) [Source:MGI Symbol;Acc:MGI:1062 | 80,34 | 3,9 |
| ENSMUSG00000033769 | 224,4878107 | 326,5 | 263,0 | 5,046048909 | 0,005408155 | Exoc6b | exocyst complex component 6B [Source:MGI Symbol;Acc:MGI:1923164] | 99,54 | 23,3 |
| ENSMUSG00000022773 | 69,16600778 | 114,2 | 24,2 | 5,046174145 | 0,007588935 | Ypel1 | yippee-like 1 (Drosophila) [Source:MGI Symbol;Acc:MGI:1913303] | 73,52 | 6,0 |
| ENSMUSG00000024241 | 135,6877925 | 128,2 | 22,8 | 5,046260693 | 0,007238478 | Sos1 | son of sevenless homolog 1 (Drosophila) [Source:MGI Symbol;Acc:MGI:98354] | 75,53 | 2,6 |
| ENSMUSG00000031901 | 108,8956621 | 92,1 | 97,4 | 5,048404431 | 0,010332307 | Dus2 | dihydrouridine synthase 2 [Source:MGI Symbol;Acc:MGI:1913619] | 86,14 | 26,7 |
| ENSMUSG00000025175 | 69,88026476 | 87,8 | 78,1 | 5,051333077 | 0,020000617 | Fn3k | fructosamine 3 kinase [Source:MGI Symbol;Acc:MGI:1926834] | 23,25 | 7,2 |
| ENSMUSG00000106261 | 28,01506469 | 46,8 | 8,0 | 5,051896546 | 0,027404618 | Gm34086 | predicted gene, 34086 [Source:MGI Symbol;Acc:MGI:5593245] | #DIV/0! | #DIV/0! |
| ENSMUSG00000000325 | 62,99202597 | 100,5 | 5,6 | 5,054680477 | 0,011601357 | Arvcf | armadillo repeat gene deleted in velo-cardio-facial syndrome [Source:MGI Symbol;Acc:MGI:109620] | 80,94 | 1,2 |
| ENSMUSG00000050721 | 716,7367821 | 830,7 | 116,9 | 5,055084201 | 0,000674007 | Plekho2 | pleckstrin homology domain containing, family O member 2 [Source:MGI Symbol;Acc:MGI:2143132] | 100,00 | 28,2 |
| ENSMUSG00000042726 | 84,08609304 | 143,1 | 40,3 | 5,055324358 | 0,01805405 | Trafd1 | TRAF type zinc finger domain containing 1 [Source:MGI Symbol;Acc:MGI:1923551] | 95,51 | 14,5 |
| ENSMUSG00000058454 | 147,5758741 | 192,0 | 137,6 | 5,05618872 | 0,006538382 | Dhcr7 | 7-dehydrocholesterol reductase [Source:MGI Symbol;Acc:MGI:1298378] | 77,17 | 21,6 |
| ENSMUSG00000027079 | 127,9999913 | 72,6 | 92,6 | 5,056339414 | 0,022828578 | Clp1 | CLP1, cleavage and polyadenylation factor I subunit [Source:MGI Symbol;Acc:MGI:2138968] | 70,13 | 21,6 |
| ENSMUSG00000037649 | 33,22673354 | 20,9 | 0,6 | 5,061172661 | 0,026432656 | H2-DMa | histocompatibility 2, class II, locus DMa [Source:MGI Symbol;Acc:MGI:95921] | 88,49 | 1,5 |
| ENSMUSG00000051000 | 17,59924062 | 23,5 | 1,0 | 5,061856533 | 0,034679175 | Fam160a1 | family with sequence similarity 160, member A1 [Source:MGI Symbol;Acc:MGI:2444746] | 49,96 | 0,5 |
| ENSMUSG00000086247 | 76,581896 | 92,8 | 26,0 | 5,062744487 | 0,014196298 | Gm15787 | predicted gene 15787 [Source:MGI Symbol;Acc:MGI:3783229] | 61,60 | 4,4 |
| ENSMUSG00000060121 | 66,77524957 | 74,2 | 82,3 | 5,06285925 | 0,024379257 | Gemin2 | gem (nuclear organelle) associated protein 2 [Source:MGI Symbol;Acc:MGI:1913853] | 88,75 | 28,9 |
| ENSMUSG00000035024 | 54,93971072 | 53,2 | 171,9 | 5,065730457 | 0,041755486 | Ncapd3 | non-SMC condensin II complex, subunit D3 [Source:MGI Symbol;Acc:MGI:2142989] | 79,84 | 25,6 |
| ENSMUSG00000047423 | 115,9168336 | 189,8 | 33,1 | 5,066068865 | 0,015924147 | AI837181 | expressed sequence AI837181 [Source:MGI Symbol;Acc:MGI:2147598] | 95,67 | 20,2 |
| ENSMUSG00000025213 | 11,65235518 | 19,9 | 0,6 | 5,068264172 | 0,045725793 | Kazald1 | Kazal-type serine peptidase inhibitor domain 1 [Source:MGI Symbol;Acc:MGI:2147606] | 69,11 | 0,9 |
| ENSMUSG00000038530 | 74,77793906 | 151,3 | 5,2 | 5,068860959 | 0,00354517 | Rgs4 | regulator of G-protein signaling 4 [Source:MGI Symbol;Acc:MGI:108409] | 98,40 | 2,5 |
| ENSMUSG00000023806 | 26,7326911 | 33,7 | 2,8 | 5,068891305 | 0,042696781 | Rsph3b | radial spoke 3B homolog (Chlamydomonas) [Source:MGI Symbol;Acc:MGI:3630308] | 34,47 | 2,4 |
| ENSMUSG00000078762 | 34,89670855 | 24,3 | 0,9 | 5,069037845 | 0,028969349 | Haus5 | HAUS augmin-like complex, subunit 5 [Source:MGI Symbol;Acc:MGI:1919159] | 79,44 | 0,2 |
| ENSMUSG00000092124 | 33,89701234 | 50,3 | 2,4 | 5,069557818 | 0,01361908 | B930094E09Rik | RIKEN cDNA B930094E09 gene [Source:MGI Symbol;Acc:MGI:3603750] | 84,27 | 5,9 |
| ENSMUSG00000029228 | 58,43193067 | 67,6 | 104,1 | 5,07360122 | 0,032701694 | Lnx1 | ligand of numb-protein X 1 [Source:MGI Symbol;Acc:MGI:1278335] | 57,94 | 17,5 |
| ENSMUSG00000040606 | 54,12360352 | 62,3 | 47,9 | 5,075427671 | 0,020150316 | Kazn | kazrin, periplakin interacting protein [Source:MGI Symbol;Acc:MGI:1918779] | 64,10 | 3,8 |
| ENSMUSG00000002814 | 28,7550759 | 33,1 | 1,0 | 5,076534496 | 0,036080939 | Top3a | topoisomerase (DNA) III alpha [Source:MGI Symbol;Acc:MGI:1197527] | 56,07 | 0,6 |
| ENSMUSG00000048232 | 39,17283555 | 64,0 | 2,2 | 5,077051287 | 0,022785134 | Fbxo10 | F-box protein 10 [Source:MGI Symbol;Acc:MGI:2686937] | 85,93 | 0,9 |
| ENSMUSG00000028278 | 585,1836176 | 783,8 | 68,0 | 5,077811641 | 0,014778862 | Rragd | Ras-related GTP binding D [Source:MGI Symbol;Acc:MGI:1098604] | 93,98 | 9,7 |
| ENSMUSG00000022335 | 10,91047632 | 32,2 | 1,2 | 5,079376603 | 0,033169014 | Zfat | zinc finger and AT hook domain containing [Source:MGI Symbol;Acc:MGI:2681865] | 69,74 | 1,1 |
| ENSMUSG00000026109 | 160,1201728 | 74,6 | 74,9 | 5,079909856 | 0,018385995 | Tmeff2 | ısmembrane protein with EGF-like and two follistatin-like domains 2 [Source:MGI Symbol;Acc:MGI:1861 | 47,04 | 11,8 |
| ENSMUSG00000026872 | 192,7242825 | 314,1 | 15,1 | 5,080231924 | 4,12477E-05 | Zeb2 | zinc finger E-box binding homeobox 2 [Source:MGI Symbol;Acc:MGI:1344407] | 71,60 | 2,1 |
| ENSMUSG00000037395 | 86,27771055 | 142,0 | 25,6 | 5,081810849 | 0,003913071 | Rcor3 | REST corepressor 3 [Source:MGI Symbol;Acc:MGI:2441920] | 86,23 | 3,7 |
| ENSMUSG00000040557 | 53,64549912 | 51,7 | 4,6 | 5,081847244 | 0,013941177 | Wbscr27 | Williams Beuren syndrome chromosome region 27 (human) [Source:MGI Symbol;Acc:MGI:1933146] | 78,29 | 2,5 |
| ENSMUSG00000023826 | 23,45450313 | 31,7 | 0,6 | 5,082617526 | 0,02712276 | Park2 | Parkinson disease (autosomal recessive, juvenile) 2, parkin [Source:MGI Symbol;Acc:MGI:1355296] | 47,52 | 0,6 |
| ENSMUSG00000034871 | 323,5928665 | 488,7 | 152,5 | 5,085575855 | 0,022538154 | Fam151a | family with sequence simliarity 151, member A [Source:MGI Symbol;Acc:MGI:2657115] | 65,68 | 25,4 |
| ENSMUSG00000033790 | 54,48193508 | 86,3 | 2,5 | 5,086461163 | 0,011335402 | Tubgcp5 | tubulin, gamma complex associated protein 5 [Source:MGI Symbol;Acc:MGI:2178836] | 90,91 | 1,7 |
| ENSMUSG00000019577 | 310,7903641 | 477,4 | 208,3 | 5,088761865 | 0,016698171 | Pdk4 | pyruvate dehydrogenase kinase, isoenzyme 4 [Source:MGI Symbol;Acc:MGI:1351481] | 99,96 | 31,7 |
| ENSMUSG00000063087 | 39,50356686 | 64,8 | 27,2 | 5,090398904 | 0,032353047 | Gm10125 | predicted gene 10125 [Source:MGI Symbol;Acc:MGI:3642044] | 60,44 | 5,7 |
| ENSMUSG00000028441 | 68,34975571 | 74,9 | 15,1 | 5,091632541 | 0,017418341 | 1110017D15Rik | RIKEN cDNA 1110017D15 gene [Source:MGI Symbol;Acc:MGI:1920971] | 51,39 | 15,5 |
| ENSMUSG00000022342 | 17,19854407 | 30,5 | 0,9 | 5,092703292 | 0,033293558 | Kcnv1 | potassium channel, subfamily V, member 1 [Source:MGI Symbol;Acc:MGI:1914748] | 56,21 | 0,5 |
| ENSMUSG00000032285 | 515,0352402 | 705,8 | 96,5 | 5,099225382 | 0,012689677 | Dnaja4 | DnaJ heat shock protein family (Hsp40) member A4 [Source:MGI Symbol;Acc:MGI:1927638] | 85,26 | 24,7 |
| ENSMUSG00000036968 | 51,43796095 | 70,0 | 53,9 | 5,099818775 | 0,028922607 | Cnpy4 | canopy FGF signaling regulator 4 [Source:MGI Symbol;Acc:MGI:1913705] | 73,23 | 20,9 |
| ENSMUSG00000038369 | 107,7090238 | 193,9 | 73,1 | 5,101549262 | 0,024052207 | Ncoa6 | nuclear receptor coactivator 6 [Source:MGI Symbol;Acc:MGI:1929915] | 85,04 | 4,0 |
| ENSMUSG00000053985 | 32,53419349 | 76,2 | 2,9 | 5,10225116 | 0,00303887 | Zfp14 | zinc finger protein 14 [Source:MGI Symbol;Acc:MGI:99160] | 94,06 | 1,6 |
| ENSMUSG00000035372 | 80,51237198 | 110,1 | 4,8 | 5,103188324 | 0,030781503 | 1810055G02Rik | RIKEN cDNA 1810055G02 gene [Source:MGI Symbol;Acc:MGI:1919306] | 99,11 | 3,0 |
| ENSMUSG00000056737 | 66,81488419 | 53,2 | 49,9 | 5,104436483 | 0,046024278 | Capg | capping protein (actin filament), gelsolin-like [Source:MGI Symbol;Acc:MGI:1098259] | 57,99 | 18,5 |

| | | | | | | | | | |
|---|---|---|---|---|---|---|---|---|---|
| ENSMUSG00000029502 | 104,9344604 | 149,3 | 5,2 | 5,104895667 | 0,003836524 | Golga3 | golgi autoantigen, golgin subfamily a, 3 [Source:MGI Symbol;Acc:MGI:96958] | 89,17 | 0,7 |
| ENSMUSG00000047407 | 80,26799436 | 67,1 | 10,2 | 5,105194037 | 0,016761518 | Tgif1 | TGFB-induced factor homeobox 1 [Source:MGI Symbol;Acc:MGI:1194497] | 47,12 | 4,9 |
| ENSMUSG00000036002 | 185,8385032 | 303,2 | 100,0 | 5,10634373 | 0,011485191 | Fam214b | family with sequence similarity 214, member B [Source:MGI Symbol;Acc:MGI:2441854] | 81,78 | 23,5 |
| ENSMUSG00000020656 | 91,46543011 | 91,8 | 3,2 | 5,107200482 | 0,014627184 | Grhl1 | grainyhead-like 1 (Drosophila) [Source:MGI Symbol;Acc:MGI:2182540] | 92,47 | 1,5 |
| ENSMUSG00000022468 | 36,70930584 | 56,5 | 4,6 | 5,110084892 | 0,013798776 | Endou | endonuclease, polyU-specific [Source:MGI Symbol;Acc:MGI:97746] | 67,07 | 4,0 |
| ENSMUSG00000029545 | 124,6233019 | 178,7 | 7,0 | 5,112383116 | 0,016949068 | Acads | acyl-Coenzyme A dehydrogenase, short chain [Source:MGI Symbol;Acc:MGI:87868] | 98,15 | 4,8 |
| ENSMUSG00000019820 | 58,9107904 | 55,5 | 1,9 | 5,112449295 | 0,011914035 | Utrn | utrophin [Source:MGI Symbol;Acc:MGI:104631] | 38,26 | 0,3 |
| ENSMUSG00000006576 | 99,9885202 | 122,6 | 4,8 | 5,113192255 | 0,011568682 | Slc4a3 | solute carrier family 4 (anion exchanger), member 3 [Source:MGI Symbol;Acc:MGI:109350] | 57,55 | 0,2 |
| ENSMUSG00000030103 | 725,4548401 | 1192,8 | 98,4 | 5,113210667 | 0,037822042 | Bhlhe40 | basic helix-loop-helix family, member e40 [Source:MGI Symbol;Acc:MGI:1097714] | 96,08 | 25,3 |
| ENSMUSG00000008763 | 175,3644089 | 131,2 | 78,1 | 5,114005111 | 0,003947394 | Man1a2 | mannosidase, alpha, class 1A, member 2 [Source:MGI Symbol;Acc:MGI:104676] | 79,56 | 5,4 |
| ENSMUSG00000018733 | 79,14785544 | 159,6 | 5,8 | 5,116927725 | 0,002402222 | Pex12 | peroxisomal biogenesis factor 12 [Source:MGI Symbol;Acc:MGI:2144177] | 76,60 | 2,5 |
| ENSMUSG00000023022 | 25,72976245 | 39,0 | 2,0 | 5,117625913 | 0,015403999 | Lima1 | LIM domain and actin binding 1 [Source:MGI Symbol;Acc:MGI:1920992] | 75,47 | 0,0 |
| ENSMUSG00000040820 | 68,8201688 | 74,6 | 29,6 | 5,117786187 | 0,014040727 | Hlcs | synthetase (biotin- [propriony-Coenzyme A-carboxylase (ATP-hydrolysing)] ligase) [Source:MGI Symbol;A | 88,12 | 4,8 |
| ENSMUSG00000085795 | 71,22618403 | 126,2 | 8,0 | 5,121607969 | 0,003025102 | Zfp703 | zinc finger protein 703 [Source:MGI Symbol;Acc:MGI:2662729] | 96,22 | 4,7 |
| ENSMUSG00000038668 | 40,83155936 | 57,6 | 16,0 | 5,12552131 | 0,021568781 | Lpar1 | lysophosphatidic acid receptor 1 [Source:MGI Symbol;Acc:MGI:108429] | 75,15 | 2,7 |
| ENSMUSG00000097684 | 10,63876612 | 25,7 | 0,9 | 5,126429992 | 0,037108704 | Gm26645 | predicted gene, 26645 [Source:MGI Symbol;Acc:MGI:5477139] | 93,79 | 1,9 |
| ENSMUSG00000018900 | 58,09831823 | 86,2 | 82,3 | 5,131343503 | 0,024220695 | Slc22a5 | solute carrier family 22 (organic cation transporter), member 5 [Source:MGI Symbol;Acc:MGI:1329012] | 88,47 | 16,6 |
| ENSMUSG00000097063 | 47,74266413 | 60,4 | 36,3 | 5,132142976 | 0,04126323 | Pantr2 | domain, class 3, transcription factor 3 adjacent noncoding transcript 2 [Source:MGI Symbol;Acc:MGI:191 | 52,43 | 5,7 |
| ENSMUSG00000036622 | 163,3048584 | 173,4 | 10,8 | 5,133631323 | 0,028808928 | Atp13a2 | ATPase type 13A2 [Source:MGI Symbol;Acc:MGI:1922022] | 93,17 | 3,7 |
| ENSMUSG00000027935 | 167,9503901 | 237,1 | 46,6 | 5,133870032 | 0,032855246 | Rab13 | RAB13, member RAS oncogene family [Source:MGI Symbol;Acc:MGI:1927232] | 96,89 | 22,2 |
| ENSMUSG00000048277 | 93,7020071 | 119,4 | 3,5 | 5,134870551 | 0,004712445 | Syngr2 | synaptogyrin 2 [Source:MGI Symbol;Acc:MGI:1328324] | 87,84 | 3,0 |
| ENSMUSG00000025170 | 68,62411638 | 81,7 | 2,8 | 5,135486 | 0,031057321 | Rab40b | Rab40B, member RAS oncogene family [Source:MGI Symbol;Acc:MGI:2183451] | 96,22 | 3,1 |
| ENSMUSG00000026546 | 23,16391482 | 30,4 | 0,9 | 5,135716894 | 0,032353047 | Cfap45 | cilia and flagella associated protein 45 [Source:MGI Symbol;Acc:MGI:1919120] | 58,33 | 1,0 |
| ENSMUSG00000036208 | 37,5741341 | 67,5 | 22,2 | 5,136212065 | 0,020466978 | Nepro | nucleolus and neural progenitor protein [Source:MGI Symbol;Acc:MGI:2384836] | 81,80 | 5,5 |
| ENSMUSG00000040339 | 197,4358207 | 179,2 | 389,2 | 5,136343902 | 0,016628071 | Fam102b | family with sequence similarity 102, member B [Source:MGI Symbol;Acc:MGI:3036259] | 97,39 | 32,0 |
| ENSMUSG00000071037 | 46,40392245 | 60,4 | 57,9 | 5,138614354 | 0,024345705 | Camkmt | calmodulin-lysine N-methyltransferase [Source:MGI Symbol;Acc:MGI:1920832] | 99,93 | 25,3 |
| ENSMUSG00000027575 | 156,8536001 | 185,9 | 89,3 | 5,139589565 | 0,007237627 | Arfgap1 | ADP-ribosylation factor GTPase activating protein 1 [Source:MGI Symbol;Acc:MGI:2183559] | 93,20 | 21,5 |
| ENSMUSG00000019854 | 393,4623558 | 433,3 | 148,5 | 5,1398102 | 0,001301333 | Reps1 | RalBP1 associated Eps domain containing protein [Source:MGI Symbol;Acc:MGI:1196373] | 99,48 | 30,3 |
| ENSMUSG00000039990 | 54,8296414 | 42,8 | 5,0 | 5,140449922 | 0,031057321 | Edrf1 | erythroid differentiation regulatory factor 1 [Source:MGI Symbol;Acc:MGI:1919831] | 68,58 | 1,2 |
| ENSMUSG00000018334 | 12,58638431 | 31,8 | 1,0 | 5,142303444 | 0,034820752 | Ksr1 | kinase suppressor of ras 1 [Source:MGI Symbol;Acc:MGI:105051] | 39,36 | 0,3 |
| ENSMUSG00000033965 | 254,0796763 | 336,1 | 80,7 | 5,142448216 | 0,014653362 | Slc16a2 | lute carrier family 16 (monocarboxylic acid transporters), member 2 [Source:MGI Symbol;Acc:MGI:12037 | 98,39 | 14,8 |
| ENSMUSG00000030350 | 60,09181989 | 70,8 | 105,8 | 5,142526198 | 0,021479852 | Prmt8 | protein arginine N-methyltransferase 8 [Source:MGI Symbol;Acc:MGI:3043083] | 80,04 | 10,5 |
| ENSMUSG00000058997 | 132,0446396 | 195,5 | 128,7 | 5,144736239 | 0,028301947 | Vwa8 | von Willebrand factor A domain containing 8 [Source:MGI Symbol;Acc:MGI:1919008] | 97,27 | 12,3 |
| ENSMUSG00000027296 | 172,0723836 | 229,4 | 70,5 | 5,14796119 | 0,040594928 | Itpka | inositol 1,4,5-trisphosphate 3-kinase A [Source:MGI Symbol;Acc:MGI:1333822] | 99,65 | 30,9 |
| ENSMUSG00000001751 | 42,20736951 | 38,8 | 3,7 | 5,148258831 | 0,041851674 | Naglu | alpha-N-acetylglucosaminidase (Sanfilippo disease IIIB) [Source:MGI Symbol;Acc:MGI:1351641] | 86,78 | 1,2 |
| ENSMUSG00000048442 | 33,37312524 | 39,2 | 3,0 | 5,148963514 | 0,0349724 | Smim5 | small integral membrane protein 5 [Source:MGI Symbol;Acc:MGI:1913778] | 67,90 | 0,4 |
| ENSMUSG00000072875 | 66,7234689 | 55,3 | 31,4 | 5,149043617 | 0,048566449 | Gpr27 | G protein-coupled receptor 27 [Source:MGI Symbol;Acc:MGI:1202299] | 93,63 | 15,6 |
| ENSMUSG00000042647 | 46,53236282 | 66,0 | 45,4 | 5,149509721 | 0,034824858 | Acad12 | acyl-Coenzyme A dehydrogenase family, member 12 [Source:MGI Symbol;Acc:MGI:2443320] | 50,00 | 5,2 |
| ENSMUSG00000090093 | 36,333196 | 62,1 | 43,9 | 5,149850138 | 0,032701694 | Gm14399 | predicted gene 14399 [Source:MGI Symbol;Acc:MGI:3650082] | 51,07 | 5,3 |
| ENSMUSG00000044231 | 134,591697 | 212,4 | 66,4 | 5,153502571 | 0,024711206 | Nhlrc1 | NHL repeat containing 1 [Source:MGI Symbol;Acc:MGI:2145264] | 98,85 | 27,3 |
| ENSMUSG00000040446 | 243,6997894 | 311,6 | 69,1 | 5,154260053 | 0,016949068 | Rprd1a | regulation of nuclear pre-mRNA domain containing 1A [Source:MGI Symbol;Acc:MGI:2385066] | 53,25 | 15,0 |
| ENSMUSG00000104415 | 47,68715312 | 71,2 | 49,0 | 5,154455844 | 0,023293298 | Gm37069 | predicted gene, 37069 [Source:MGI Symbol;Acc:MGI:5610297] | 87,90 | 6,7 |
| ENSMUSG00000055652 | 39,12922482 | 73,3 | 14,8 | 5,154674026 | 0,012713796 | Klhl25 | kelch-like 25 [Source:MGI Symbol;Acc:MGI:2668031] | 84,15 | 4,3 |
| ENSMUSG00000029207 | 71,68205716 | 128,6 | 27,5 | 5,156739904 | 0,018852943 | Apbb2 | amyloid beta (A4) precursor protein-binding, family B, member 2 [Source:MGI Symbol;Acc:MGI:108405] | 82,53 | 2,9 |
| ENSMUSG00000071379 | 44,35233608 | 39,9 | 8,3 | 5,157155079 | 0,047394949 | Hpcal1 | hippocalcin-like 1 [Source:MGI Symbol;Acc:MGI:1855689] | 95,00 | 6,1 |
| ENSMUSG00000040250 | 285,5054749 | 266,1 | 136,5 | 5,15727548 | 0,016213872 | Asun | asunder, spermatogenesis regulator [Source:MGI Symbol;Acc:MGI:1918427] | 96,40 | 24,3 |
| ENSMUSG00000021418 | 48,77326851 | 76,1 | 52,7 | 5,157689022 | 0,019292726 | Rpp40 | ribonuclease P 40 subunit [Source:MGI Symbol;Acc:MGI:1346084] | 39,02 | 8,0 |
| ENSMUSG00000047881 | 76,30678071 | 52,6 | 20,3 | 5,158783526 | 0,035836561 | Rell1 | RELT-like 1 [Source:MGI Symbol;Acc:MGI:2140767] | 86,81 | 3,8 |
| ENSMUSG00000085316 | 41,55075697 | 54,3 | 58,9 | 5,159333655 | 0,035624713 | D330050G23Rik | RIKEN cDNA D330050G23 gene [Source:MGI Symbol;Acc:MGI:2445155] | #DIV/0! | #DIV/0! |
| ENSMUSG00000040918 | 154,8408441 | 140,6 | 15,8 | 5,16351798 | 0,013906882 | Slc19a2 | solute carrier family 19 (thiamine transporter), member 2 [Source:MGI Symbol;Acc:MGI:1928761] | 96,60 | 7,0 |
| ENSMUSG00000036699 | 41,01963145 | 46,2 | 128,7 | 5,164770679 | 0,041059844 | Zcchc12 | zinc finger, CCHC domain containing 12 [Source:MGI Symbol;Acc:MGI:1919943] | 89,69 | 27,4 |
| ENSMUSG00000024948 | 136,5097488 | 118,7 | 121,0 | 5,165252085 | 0,006675325 | Map4k2 | mitogen-activated protein kinase kinase kinase kinase 2 [Source:MGI Symbol;Acc:MGI:1346883] | 91,43 | 17,8 |
| ENSMUSG00000033542 | 14,98811942 | 28,1 | 0,9 | 5,165545033 | 0,036159363 | Arhgef5 | Rho guanine nucleotide exchange factor (GEF) 5 [Source:MGI Symbol;Acc:MGI:1858952] | 53,14 | 0,3 |
| ENSMUSG00000025425 | 85,18650562 | 130,8 | 13,5 | 5,165745227 | 0,016615332 | St8sia5 | ST8 alpha-N-acetyl-neuraminide alpha-2,8-sialyltransferase 5 [Source:MGI Symbol;Acc:MGI:109243] | 99,36 | 9,1 |
| ENSMUSG00000037845 | 39,3298292 | 55,2 | 8,5 | 5,166546077 | 0,038260068 | Fdxacb1 | ferredoxin-fold anticodon binding domain containing 1 [Source:MGI Symbol;Acc:MGI:3584513] | 84,84 | 2,5 |
| ENSMUSG00000031684 | 34,64193907 | 44,4 | 3,7 | 5,170434793 | 0,02435978 | Slc10a7 | te carrier family 10 (sodium/bile acid cotransporter family), member 7 [Source:MGI Symbol;Acc:MGI:1924 | 83,76 | 0,5 |
| ENSMUSG00000028948 | 101,2409005 | 60,2 | 54,9 | 5,172692714 | 0,035504063 | Nol9 | nucleolar protein 9 [Source:MGI Symbol;Acc:MGI:1921285] | 85,29 | 5,7 |
| ENSMUSG00000041096 | 171,1173756 | 217,6 | 130,5 | 5,174904244 | 0,0277816 | Tspyl2 | TSPY-like 2 [Source:MGI Symbol;Acc:MGI:106244] | 97,23 | 28,7 |
| ENSMUSG00000020396 | 22,39075056 | 29,3 | 1,0 | 5,17694258 | 0,039554987 | Nefh | neurofilament, heavy polypeptide [Source:MGI Symbol;Acc:MGI:97309] | 55,12 | 0,6 |
| ENSMUSG00000079550 | 11,78583971 | 24,2 | 1,0 | 5,177999276 | 0,035949079 | Mpp4 | mbrane protein, palmitoylated 4 (MAGUK p55 subfamily member 4) [Source:MGI Symbol;Acc:MGI:23866 | 39,16 | 0,0 |
| ENSMUSG00000015942 | 43,53180055 | 26,6 | 0,9 | 5,179000225 | 0,045752938 | Gtf2ird2 | GTF2I repeat domain containing 2 [Source:MGI Symbol;Acc:MGI:2149780] | 64,39 | 0,6 |
| ENSMUSG00000000148 | 46,2921206 | 72,4 | 21,3 | 5,180793187 | 0,028391752 | Brat1 | BRCA1-associated ATM activator 1 [Source:MGI Symbol;Acc:MGI:1891679] | 78,05 | 6,8 |
| ENSMUSG00000071723 | 82,87963755 | 65,2 | 2,1 | 5,182128496 | 0,048897618 | Gspt2 | G1 to S phase transition 2 [Source:MGI Symbol;Acc:MGI:1316727] | 91,62 | 2,7 |
| ENSMUSG00000040118 | 48,39799425 | 37,5 | 1,2 | 5,182562846 | 0,036115609 | Cacna2d1 | calcium channel, voltage-dependent, alpha2/delta subunit 1 [Source:MGI Symbol;Acc:MGI:88295] | 37,37 | 0,3 |
| ENSMUSG00000042770 | 34,30390378 | 60,9 | 6,1 | 5,182874815 | 0,022180393 | Hebp1 | heme binding protein 1 [Source:MGI Symbol;Acc:MGI:1333880] | 96,60 | 16,3 |
| ENSMUSG00000018470 | 51,28050178 | 51,1 | 22,2 | 5,183095088 | 0,036792124 | Kcnab3 | sium voltage-gated channel, shaker-related subfamily, beta member 3 [Source:MGI Symbol;Acc:MGI:133 | 82,16 | 9,8 |
| ENSMUSG00000073176 | 65,08291579 | 76,6 | 156,2 | 5,184407795 | 0,025360525 | Zfp449 | zinc finger protein 449 [Source:MGI Symbol;Acc:MGI:1925869] | 94,81 | 26,5 |
| ENSMUSG00000105518 | 45,1103132 | 51,5 | 4,2 | 5,186033469 | 0,034536859 | Gm42674 | predicted gene 42674 [Source:MGI Symbol;Acc:MGI:5662811] | #DIV/0! | #DIV/0! |
| ENSMUSG00000037606 | 78,19439619 | 100,0 | 67,2 | 5,18687639 | 0,026432656 | Osbpl5 | oxysterol binding protein-like 5 [Source:MGI Symbol;Acc:MGI:1930265] | 86,79 | 14,7 |
| ENSMUSG00000094786 | 55,19959402 | 72,1 | 20,6 | 5,187379713 | 0,025109118 | Gm14403 | predicted gene 14403 [Source:MGI Symbol;Acc:MGI:3649813] | 97,38 | 4,0 |
| ENSMUSG00000054737 | 27,1093844 | 47,3 | 3,0 | 5,188376341 | 0,01508771 | Zfp182 | zinc finger protein 182 [Source:MGI Symbol;Acc:MGI:2442220] | 79,75 | 0,8 |
| ENSMUSG00000018395 | 215,134701 | 213,3 | 194,2 | 5,190944745 | 0,010895155 | Kif3a | kinesin family member 3A [Source:MGI Symbol;Acc:MGI:107689] | 92,78 | 18,6 |
| ENSMUSG00000045967 | 76,50220896 | 57,5 | 305,3 | 5,194870638 | 0,034971199 | Gpr158 | G protein-coupled receptor 158 [Source:MGI Symbol;Acc:MGI:2441697] | 66,49 | 24,4 |
| ENSMUSG00000025893 | 91,89131739 | 132,2 | 5,8 | 5,197206098 | 0,003911996 | Kbtbd3 | kelch repeat and BTB (POZ) domain containing 3 [Source:MGI Symbol;Acc:MGI:1916399] | 89,12 | 3,0 |
| ENSMUSG00000026187 | 161,968736 | 186,0 | 105,0 | 5,197501258 | 0,0223201 | Xrcc5 | ray repair complementing defective repair in Chinese hamster cells 5 [Source:MGI Symbol;Acc:MGI:1045 | 99,80 | 28,3 |
| ENSMUSG00000035776 | 249,2498894 | 236,4 | 53,6 | 5,197548872 | 0,0027533 | Cd99l2 | CD99 antigen-like 2 [Source:MGI Symbol;Acc:MGI:2177151] | 95,55 | 11,5 |
| ENSMUSG00000034059 | 45,47766007 | 55,6 | 50,8 | 5,202565675 | 0,036731538 | Ypel4 | yippee-like 4 (Drosophila) [Source:MGI Symbol;Acc:MGI:3605071] | 93,12 | 17,5 |
| ENSMUSG00000078490 | 74,98911136 | 70,2 | 117,4 | 5,206655281 | 0,023446158 | Cfap74 | cilia and flagella associated protein 74 [Source:MGI Symbol;Acc:MGI:1917130] | 37,11 | 9,3 |
| ENSMUSG00000048794 | 41,12567431 | 50,6 | 92,8 | 5,206741081 | 0,043183971 | Cfap100 | cilia and flagella associated protein 100 [Source:MGI Symbol;Acc:MGI:2141635] | 73,66 | 13,9 |
| ENSMUSG00000051343 | 59,5609676 | 81,3 | 24,8 | 5,207089375 | 0,023617627 | Rab11fip5 | RAB11 family interacting protein 5 (class I) [Source:MGI Symbol;Acc:MGI:1098586] | 64,16 | 5,0 |
| ENSMUSG00000004698 | 38,9637144 | 63,4 | 3,0 | 5,2104274 | 0,008060872 | Hdac9 | histone deacetylase 9 [Source:MGI Symbol;Acc:MGI:1931221] | 86,97 | 0,6 |
| ENSMUSG00000038059 | 1023,446181 | 2280,3 | 177,3 | 5,210595532 | 0,016782598 | Smim3 | small integral membrane protein 3 [Source:MGI Symbol;Acc:MGI:1917088] | 97,63 | 41,4 |
| ENSMUSG00000071847 | 13,68566577 | 27,7 | 1,0 | 5,215469487 | 0,03214507 | Apcdd1 | adenomatosis polyposis coli down-regulated 1 [Source:MGI Symbol;Acc:MGI:3513977] | 74,07 | 0,8 |
| ENSMUSG00000025533 | 102,3547323 | 109,2 | 80,5 | 5,2167861 | 0,012772283 | Asl | argininosuccinate lyase [Source:MGI Symbol;Acc:MGI:88084] | 54,29 | 15,7 |
| ENSMUSG00000053475 | 32,56073551 | 53,5 | 5,0 | 5,2174576 | 0,036734909 | Tnfaip6 | tumor necrosis factor alpha induced protein 6 [Source:MGI Symbol;Acc:MGI:1195266] | 94,39 | 2,6 |
| ENSMUSG00000020196 | 112,6500307 | 139,0 | 3,7 | 5,219663403 | 0,000243456 | Cabin1 | calcineurin binding protein 1 [Source:MGI Symbol;Acc:MGI:1298375] | 94,20 | 1,0 |
| ENSMUSG00000019996 | 136,6125369 | 204,4 | 89,7 | 5,220539806 | 0,012850723 | Map7 | microtubule-associated protein 7 [Source:MGI Symbol;Acc:MGI:1328328] | 89,38 | 18,0 |
| ENSMUSG00000002108 | 140,6689186 | 166,8 | 108,7 | 5,221626189 | 0,030035439 | Nr1h3 | nuclear receptor subfamily 1, group H, member 3 [Source:MGI Symbol;Acc:MGI:1352462] | 91,27 | 25,3 |
| ENSMUSG00000091102 | 50,29022456 | 71,6 | 33,3 | 5,224662732 | 0,023290812 | 5830462I19Rik | RIKEN cDNA 5830462I19 gene [Source:MGI Symbol;Acc:MGI:3642105] | 23,25 | 6,9 |
| ENSMUSG00000024330 | 62,72775304 | 96,9 | 35,1 | 5,226704925 | 0,016670975 | Col11a2 | collagen, type XI, alpha 2 [Source:MGI Symbol;Acc:MGI:88447] | 83,56 | 4,8 |
| ENSMUSG00000026792 | 80,14356034 | 131,7 | 4,6 | 5,22673954 | 0,016560862 | Lrsam1 | leucine rich repeat and sterile alpha motif containing 1 [Source:MGI Symbol;Acc:MGI:2684789] | 92,04 | 0,9 |
| ENSMUSG00000041268 | 172,6345725 | 97,3 | 3,8 | 5,227538859 | 0,007327727 | Dmxl2 | Dmx-like 2 [Source:MGI Symbol;Acc:MGI:2444630] | 61,31 | 0,6 |
| ENSMUSG00000022321 | 118,9404582 | 182,0 | 74,4 | 5,227680184 | 0,014564133 | Cdh10 | cadherin 10 [Source:MGI Symbol;Acc:MGI:107436] | 91,19 | 9,2 |
| ENSMUSG00000026018 | 70,53183861 | 101,2 | 64,2 | 5,233191259 | 0,02010831 | Ica1l | islet cell autoantigen 1-like [Source:MGI Symbol;Acc:MGI:1917625] | 88,44 | 2,8 |
| ENSMUSG00000050423 | 526,5832015 | 1043,3 | 49,7 | 5,234971213 | 0,012259683 | Ppp1r3g | protein phosphatase 1, regulatory (inhibitor) subunit 3G [Source:MGI Symbol;Acc:MGI:1923737] | 99,76 | 22,3 |
| ENSMUSG00000038954 | 58,46530433 | 58,0 | 12,0 | 5,235247988 | 0,026432771 | Supt3 | suppressor of Ty 3 [Source:MGI Symbol;Acc:MGI:1923723] | 88,12 | 8,7 |
| ENSMUSG00000021171 | 125,7756086 | 110,1 | 2,6 | 5,237514876 | 0,00246054 | Esyt2 | extended synaptotagmin-like protein 2 [Source:MGI Symbol;Acc:MGI:1261845] | 79,20 | 0,9 |
| ENSMUSG00000040616 | 63,1237672 | 70,5 | 43,4 | 5,238313636 | 0,037169362 | Tmem51 | transmembrane protein 51 [Source:MGI Symbol;Acc:MGI:2384874] | 95,04 | 24,1 |
| ENSMUSG00000028709 | 28,93443034 | 32,0 | 1,2 | 5,238962524 | 0,049937405 | Mob3c | MOB kinase activator 3C [Source:MGI Symbol;Acc:MGI:2140623] | 76,44 | 0,6 |
| ENSMUSG00000050910 | 97,16432805 | 106,4 | 119,8 | 5,240897785 | 0,019111794 | Cdr2l | cerebellar degeneration-related protein 2-like [Source:MGI Symbol;Acc:MGI:2684867] | 96,84 | 28,9 |
| ENSMUSG00000000838 | 479,110113 | 623,1 | 85,7 | 5,242950626 | 0,005424796 | Fmr1 | fragile X mental retardation syndrome 1 [Source:MGI Symbol;Acc:MGI:95564] | 98,27 | 13,3 |
| ENSMUSG00000098789 | 52,12895607 | 68,2 | 3,0 | 5,243212915 | 0,001753896 | Jmjd7 | jumonji domain containing 7 [Source:MGI Symbol;Acc:MGI:3845785] | 97,98 | 1,1 |
| ENSMUSG00000030619 | 208,3610213 | 226,8 | 93,9 | 5,245047611 | 0,001565802 | Eed | embryonic ectoderm development [Source:MGI Symbol;Acc:MGI:95286] | 97,52 | 27,9 |
| ENSMUSG00000025645 | 39,73814018 | 65,3 | 25,4 | 5,245131093 | 0,043006015 | Ccdc51 | coiled-coil domain containing 51 [Source:MGI Symbol;Acc:MGI:1913908] | 99,61 | 20,1 |
| ENSMUSG00000036613 | 125,2551244 | 197,0 | 24,1 | 5,247759007 | 0,008268466 | Tssc1 | tumor suppressing subtransferable candidate 1 [Source:MGI Symbol;Acc:MGI:1289332] | 100,00 | 18,6 |
| ENSMUSG00000028469 | 164,5205442 | 160,8 | 84,5 | 5,247891918 | 0,027862291 | Npr2 | natriuretic peptide receptor 2 [Source:MGI Symbol;Acc:MGI:97372] | 97,51 | 24,9 |
| ENSMUSG00000020562 | 37,35457237 | 44,4 | 8,3 | 5,247985019 | 0,039742097 | Efcab10 | EF-hand calcium binding domain 10 [Source:MGI Symbol;Acc:MGI:1922290] | 69,68 | 11,6 |
| ENSMUSG00000024170 | 92,05062156 | 83,9 | 18,5 | 5,253655778 | 0,014277996 | Telo2 | TEL2, telomere maintenance 2, homolog (S. cerevisiae) [Source:MGI Symbol;Acc:MGI:1918968] | 94,58 | 2,8 |
| ENSMUSG00000022788 | 168,186099 | 253,7 | 142,6 | 5,258146162 | 0,025335912 | Fgd4 | FYVE, RhoGEF and PH domain containing 4 [Source:MGI Symbol;Acc:MGI:2183747] | 86,01 | 10,6 |
| ENSMUSG00000035351 | 71,32076864 | 71,6 | 90,6 | 5,258715089 | 0,030847065 | Nup37 | nucleoporin 37 [Source:MGI Symbol;Acc:MGI:1919964] | 91,61 | 29,1 |
| ENSMUSG00000021109 | 352,2828596 | 439,0 | 57,1 | 5,261088759 | 0,012408832 | Hif1a | hypoxia inducible factor 1, alpha subunit [Source:MGI Symbol;Acc:MGI:106918] | 85,24 | 9,4 |
| ENSMUSG00000022383 | 55,13099199 | 76,8 | 24,2 | 5,262028328 | 0,038545922 | Ppara | peroxisome proliferator activated receptor alpha [Source:MGI Symbol;Acc:MGI:104740] | 70,73 | 3,9 |
| ENSMUSG00000021745 | 181,7760138 | 321,9 | 45,4 | 5,263130608 | 0,005424796 | Ptprg | protein tyrosine phosphatase, receptor type, G [Source:MGI Symbol;Acc:MGI:97814] | 85,32 | 4,2 |
| ENSMUSG00000035310 | 146,2652254 | 124,5 | 76,3 | 5,263874972 | 0,012405249 | Lin54 | lin-54 homolog (C. elegans) [Source:MGI Symbol;Acc:MGI:2140902] | 87,28 | 6,6 |
| ENSMUSG00000070699 | 57,15718595 | 75,8 | 21,0 | 5,264814251 | 0,01071547 | Sars2 | seryl-aminoacyl-tRNA synthetase 2 [Source:MGI Symbol;Acc:MGI:1919234] | 99,38 | 9,8 |
| ENSMUSG00000029338 | 14,15824376 | 26,5 | 1,0 | 5,266115519 | 0,028809138 | Antxr2 | anthrax toxin receptor 2 [Source:MGI Symbol;Acc:MGI:1919164] | 35,67 | 0,4 |
| ENSMUSG00000017765 | 251,320559 | 317,7 | 19,7 | 5,267422178 | 0,017459821 | Slc12a4 | solute carrier family 12, member 4 [Source:MGI Symbol;Acc:MGI:1309465] | 99,06 | 6,1 |
| ENSMUSG00000023571 | 74,67930317 | 65,1 | 15,7 | 5,270191893 | 0,031804477 | Fam132a | family with sequence similarity 132, member A [Source:MGI Symbol;Acc:MGI:1914639] | 94,65 | 10,6 |
| ENSMUSG00000086740 | 54,19213845 | 87,2 | 19,0 | 5,271352673 | 0,007191641 | Gm17029 | predicted gene 17029 [Source:MGI Symbol;Acc:MGI:4937856] | 12,46 | 2,6 |
| ENSMUSG00000073293 | 94,8924683 | 171,9 | 5,5 | 5,272062185 | 0,003096333 | Nudt10 | nudix (nucleoside diphosphate linked moiety X)-type motif 10 [Source:MGI Symbol;Acc:MGI:2147931] | 79,34 | 8,8 |
| ENSMUSG00000031585 | 173,3698052 | 224,2 | 40,3 | 5,272494012 | 0,001565802 | Gtf2e2 | general transcription factor II E, polypeptide 2 (beta subunit) [Source:MGI Symbol;Acc:MGI:1915403] | 79,24 | 16,8 |
| ENSMUSG00000041491 | 42,25465883 | 44,3 | 2,4 | 5,277521432 | 0,04304535 | Cep78 | centrosomal protein 78 [Source:MGI Symbol;Acc:MGI:1924386] | 86,64 | 1,2 |
| ENSMUSG00000041261 | 102,8341143 | 66,4 | 113,8 | 5,278876552 | 0,040165668 | Car8 | carbonic anhydrase 8 [Source:MGI Symbol;Acc:MGI:88253] | 53,63 | 16,1 |
| ENSMUSG00000018372 | 22,28199949 | 43,6 | 13,0 | 5,281547281 | 0,032010435 | Cep95 | centrosomal protein 95 [Source:MGI Symbol;Acc:MGI:2443502] | 62,90 | 2,4 |
| ENSMUSG00000017057 | 130,973264 | 103,1 | 276,0 | 5,281983738 | 0,017011651 | Il13ra1 | interleukin 13 receptor, alpha 1 [Source:MGI Symbol;Acc:MGI:105052] | 95,95 | 31,8 |
| ENSMUSG00000043153 | 51,08900882 | 74,1 | 16,3 | 5,282480909 | 0,022999178 | Ispd | isoprenoid synthase domain containing [Source:MGI Symbol;Acc:MGI:1923097] | 96,06 | 4,9 |
| ENSMUSG00000045348 | 16,81897033 | 38,7 | 1,2 | 5,285719316 | 0,016730362 | Nyap1 | uronal tyrosine-phosphorylated phosphoinositide 3-kinase adaptor 1 [Source:MGI Symbol;Acc:MGI:24438 | 52,90 | 0,4 |
| ENSMUSG00000022180 | 50,1120043 | 81,3 | 84,8 | 5,285963029 | 0,019785874 | Slc7a8 | carrier family 7 (cationic amino acid transporter, y+ system), member 8 [Source:MGI Symbol;Acc:MGI:135 | 95,16 | 12,3 |
| ENSMUSG00000037686 | 45,45385385 | 77,1 | 52,7 | 5,286819644 | 0,031650087 | Aspg | asparaginase [Source:MGI Symbol;Acc:MGI:2144822] | 95,11 | 23,0 |
| ENSMUSG00000022987 | 24,9785729 | 48,8 | 4,0 | 5,28748505 | 0,025618029 | Zfp641 | zinc finger protein 641 [Source:MGI Symbol;Acc:MGI:2442788] | 82,52 | 0,5 |
| ENSMUSG00000067071 | 259,6583772 | 400,9 | 114,2 | 5,287667554 | 0,02223371 | Hes6 | hairy and enhancer of split 6 [Source:MGI Symbol;Acc:MGI:1859852] | 99,50 | 36,6 |
| ENSMUSG00000030623 | 46,62235245 | 55,0 | 19,4 | 5,289382281 | 0,045477031 | Prss23os | protease, serine 23, opposite strand [Source:MGI Symbol;Acc:MGI:1916654] | 54,27 | 13,0 |
| ENSMUSG00000032890 | 92,15679441 | 81,6 | 130,7 | 5,289511275 | 0,029889368 | Rims3 | regulating synaptic membrane exocytosis 3 [Source:MGI Symbol;Acc:MGI:2443331] | 83,72 | 15,5 |
| ENSMUSG00000103730 | 51,30008819 | 68,8 | 10,2 | 5,292429899 | 0,014411479 | Gm38316 | predicted gene, 38316 [Source:MGI Symbol;Acc:MGI:5611544] | 41,93 | 11,9 |
| ENSMUSG00000040612 | 156,5224669 | 186,6 | 14,5 | 5,294515534 | 0,002277685 | Ildr2 | immunoglobulin-like domain containing receptor 2 [Source:MGI Symbol;Acc:MGI:1196370] | 94,63 | 3,1 |
| ENSMUSG00000029708 | 143,9752282 | 155,3 | 296,0 | 5,294715486 | 0,024619628 | Gcc1 | golgi coiled coil 1 [Source:MGI Symbol;Acc:MGI:1921625] | 79,16 | 21,9 |
| ENSMUSG00000020781 | 72,58744155 | 73,2 | 27,7 | 5,295398845 | 0,017796356 | Tsen54 | tRNA splicing endonuclease 54 homolog (S. cerevisiae) [Source:MGI Symbol;Acc:MGI:1923515] | 91,76 | 19,7 |
| ENSMUSG00000021668 | 23,70887606 | 53,5 | 1,8 | 5,298131448 | 0,009703371 | Polk | polymerase (DNA directed), kappa [Source:MGI Symbol;Acc:MGI:1349767] | 87,01 | 0,9 |
| ENSMUSG00000050074 | 26,98156033 | 51,0 | 4,2 | 5,299078256 | 0,037768849 | Spink8 | serine peptidase inhibitor, Kazal type 8 [Source:MGI Symbol;Acc:MGI:1925959] | 83,15 | 4,7 |
| ENSMUSG00000070939 | 130,8598531 | 235,5 | 8,0 | 5,300426249 | 0,00513118 | Tgfbrap1 | transforming growth factor, beta receptor associated protein 1 [Source:MGI Symbol;Acc:MGI:2447427] | 89,11 | 1,3 |
| ENSMUSG00000032512 | 216,838719 | 416,1 | 40,9 | 5,300443285 | 0,004774503 | Wdr48 | WD repeat domain 48 [Source:MGI Symbol;Acc:MGI:1914811] | 100,00 | 11,4 |
| ENSMUSG00000053046 | 97,92990596 | 147,3 | 11,6 | 5,302329873 | 0,005334587 | Brsk2 | BR serine/threonine kinase 2 [Source:MGI Symbol;Acc:MGI:1923020] | 82,21 | 2,6 |
| ENSMUSG00000049892 | 55,27677322 | 87,8 | 84,1 | 5,302836738 | 0,047725025 | Rasd1 | RAS, dexamethasone-induced 1 [Source:MGI Symbol;Acc:MGI:1270848] | 98,37 | 31,8 |
| ENSMUSG00000076439 | 74,36787024 | 65,4 | 104,8 | 5,303981479 | 0,032701694 | Mog | myelin oligodendrocyte glycoprotein [Source:MGI Symbol;Acc:MGI:97435] | 94,66 | 30,1 |
| ENSMUSG00000030966 | 26,76427984 | 54,5 | 7,0 | 5,305051581 | 0,015702473 | Trim21 | tripartite motif-containing 21 [Source:MGI Symbol;Acc:MGI:106657] | 90,49 | 1,0 |
| ENSMUSG00000026380 | 97,03732417 | 167,5 | 72,8 | 5,305170534 | 0,008849967 | Tfcp2l1 | transcription factor CP2-like 1 [Source:MGI Symbol;Acc:MGI:2444691] | 86,63 | 2,9 |
| ENSMUSG00000037697 | 425,0357441 | 548,6 | 280,6 | 5,305866348 | 0,005026681 | Ddhd1 | DDHD domain containing 1 [Source:MGI Symbol;Acc:MGI:2150302] | 98,94 | 21,2 |
| ENSMUSG00000033854 | 16,53829052 | 42,2 | 1,8 | 5,305872935 | 0,013278452 | Kcnk10 | potassium channel, subfamily K, member 10 [Source:MGI Symbol;Acc:MGI:1919508] | 91,68 | 1,9 |
| ENSMUSG00000068335 | 41,22067993 | 34,9 | 1,0 | 5,306622988 | 0,019104214 | Dok1 | docking protein 1 [Source:MGI Symbol;Acc:MGI:893587] | 86,88 | 1,3 |
| ENSMUSG00000028629 | 45,46105018 | 51,6 | 64,9 | 5,307635719 | 0,028391752 | Exo5 | exonuclease 5 [Source:MGI Symbol;Acc:MGI:1920422] | 93,16 | 16,7 |
| ENSMUSG00000061411 | 48,0942254 | 54,2 | 16,0 | 5,308071187 | 0,017640758 | Nol4l | nucleolar protein 4-like [Source:MGI Symbol;Acc:MGI:1918765] | 63,50 | 2,9 |
| ENSMUSG00000038368 | 29,40591137 | 49,7 | 2,8 | 5,308877425 | 0,013688075 | Focad | focadhesin [Source:MGI Symbol;Acc:MGI:2676921] | 69,49 | 0,8 |
| ENSMUSG00000041396 | 83,10643547 | 58,7 | 235,5 | 5,309481581 | 0,037067021 | Mettl18 | methyltransferase like 18 [Source:MGI Symbol;Acc:MGI:1917212] | 97,47 | 30,9 |
| ENSMUSG00000078716 | 88,9054473 | 103,5 | 73,2 | 5,313355069 | 0,017796356 | Tmem8b | transmembrane protein 8B [Source:MGI Symbol;Acc:MGI:2441680] | 90,56 | 10,4 |
| ENSMUSG00000024538 | 14,30345409 | 27,5 | 1,0 | 5,314602044 | 0,03541018 | Ppic | peptidylprolyl isomerase C [Source:MGI Symbol;Acc:MGI:97751] | 86,37 | 1,9 |
| ENSMUSG00000031200 | 78,075114 | 90,9 | 42,5 | 5,317364924 | 0,009905094 | Mtcp1 | mature T cell proliferation 1 [Source:MGI Symbol;Acc:MGI:102699] | 82,39 | 10,0 |
| ENSMUSG00000026723 | 50,45976155 | 79,6 | 93,2 | 5,317625551 | 0,032701694 | Trdmt1 | tRNA aspartic acid methyltransferase 1 [Source:MGI Symbol;Acc:MGI:1274787] | 76,70 | 13,4 |
| ENSMUSG00000086600 | 60,11069409 | 69,6 | 11,5 | 5,318411779 | 0,022313912 | C030005K06Rik | RIKEN cDNA C030005K06 gene [Source:MGI Symbol;Acc:MGI:1925945] | #DIV/0! | #DIV/0! |
| ENSMUSG00000024978 | 515,0201062 | 947,9 | 242,2 | 5,319261838 | 0,019063996 | Gpam | glycerol-3-phosphate acyltransferase, mitochondrial [Source:MGI Symbol;Acc:MGI:109162] | 99,36 | 37,8 |
| ENSMUSG00000042228 | 91,25272355 | 57,2 | 241,5 | 5,319893284 | 0,038936575 | Lyn | Yamaguchi sarcoma viral (v-yes-1) oncogene homolog [Source:MGI Symbol;Acc:MGI:96892] | 73,62 | 23,3 |
| ENSMUSG00000036452 | 136,3107696 | 220,4 | 19,1 | 5,32112829 | 0,003583224 | Arhgap26 | Rho GTPase activating protein 26 [Source:MGI Symbol;Acc:MGI:1918552] | 80,95 | 1,2 |
| ENSMUSG00000051034 | 53,43368676 | 70,4 | 95,2 | 5,322580363 | 0,022976457 | Zfp11 | zinc finger protein 11 [Source:MGI Symbol;Acc:MGI:99156] | 89,57 | 17,6 |
| ENSMUSG00000023938 | 39,55392988 | 58,0 | 6,5 | 5,322866798 | 0,023850871 | Aars2 | alanyl-tRNA synthetase 2, mitochondrial (putative) [Source:MGI Symbol;Acc:MGI:2681839] | 92,58 | 1,3 |
| ENSMUSG00000000753 | 105,5511797 | 177,6 | 48,7 | 5,325917151 | 0,020800123 | Serpinf1 | serine (or cysteine) peptidase inhibitor, clade F, member 1 [Source:MGI Symbol;Acc:MGI:108080] | 68,27 | 18,2 |
| ENSMUSG00000028114 | 115,5370844 | 188,0 | 61,6 | 5,32770028 | 0,009115328 | Mettl14 | methyltransferase like 14 [Source:MGI Symbol;Acc:MGI:2442926] | 39,25 | 7,8 |
| ENSMUSG00000035173 | 300,2619313 | 119,3 | 60,1 | 5,328545696 | 0,004405578 | Ccdc186 | coiled-coil domain containing 186 [Source:MGI Symbol;Acc:MGI:2445022] | 88,09 | 8,5 |
| ENSMUSG00000062542 | 51,96400263 | 70,6 | 1,6 | 5,328687816 | 0,00379295 | Syt9 | synaptotagmin IX [Source:MGI Symbol;Acc:MGI:1926373] | 81,90 | 1,0 |
| ENSMUSG00000018651 | 104,666547 | 96,9 | 25,0 | 5,328889116 | 0,006905151 | Tada2a | transcriptional adaptor 2A [Source:MGI Symbol;Acc:MGI:2144471] | 91,02 | 9,1 |
| ENSMUSG00000038349 | 117,957673 | 244,9 | 8,6 | 5,329921159 | 0,001689804 | Plcl1 | phospholipase C-like 1 [Source:MGI Symbol;Acc:MGI:3036262] | 90,35 | 2,6 |
| ENSMUSG00000028879 | 706,3689385 | 1098,7 | 188,1 | 5,333893475 | 0,004681539 | Stx12 | syntaxin 12 [Source:MGI Symbol;Acc:MGI:1931027] | 99,76 | 29,2 |

| | | | | | | | | | |
|---|---|---|---|---|---|---|---|---|---|
| ENSMUSG00000108368 | 51,32917785 | 80,0 | 12,9 | 5,334107442 | 0,018938033 | RP23-335C3.10 | | 12,81 | 3,4 |
| ENSMUSG00000045103 | 376,6631293 | 642,4 | 15,8 | 5,334634444 | 0,000931105 | Dmd | dystrophin, muscular dystrophy [Source:MGI Symbol;Acc:MGI:94909] | 46,78 | 1,7 |
| ENSMUSG00000042363 | 942,6844532 | 1364,8 | 86,3 | 5,334901017 | 3,44087E-05 | Lgalsl | lectin, galactoside binding-like [Source:MGI Symbol;Acc:MGI:1916114] | 99,99 | 21,4 |
| ENSMUSG00000031483 | 308,8706624 | 439,4 | 52,9 | 5,335631167 | 0,01751344 | Erlin2 | ER lipid raft associated 2 [Source:MGI Symbol;Acc:MGI:2387215] | 99,83 | 7,1 |
| ENSMUSG00000024193 | 147,4829131 | 218,3 | 16,1 | 5,336780818 | 0,003826312 | Phf1 | PHD finger protein 1 [Source:MGI Symbol;Acc:MGI:98647] | 97,42 | 8,0 |
| ENSMUSG00000043671 | 113,698011 | 200,8 | 59,8 | 5,336808586 | 0,006918725 | Dpy19l3 | dpy-19-like 3 (C. elegans) [Source:MGI Symbol;Acc:MGI:2443952] | 90,85 | 4,7 |
| ENSMUSG00000107944 | 74,45649667 | 78,1 | 15,7 | 5,338610246 | 0,010671474 | Gm44280 | predicted gene, 44280 [Source:MGI Symbol;Acc:MGI:5690672] | 27,59 | 5,1 |
| ENSMUSG00000022103 | 14,01036707 | 26,1 | 0,6 | 5,343521557 | 0,027094605 | Gfra2 | glial cell line derived neurotrophic factor family receptor alpha 2 [Source:MGI Symbol;Acc:MGI:1195462] | 66,17 | 0,7 |
| ENSMUSG00000023800 | 57,08861396 | 90,8 | 2,8 | 5,343735812 | 0,005219865 | Tiam2 | T cell lymphoma invasion and metastasis 2 [Source:MGI Symbol;Acc:MGI:1344338] | 82,98 | 0,6 |
| ENSMUSG00000025583 | 122,025605 | 122,8 | 6,6 | 5,343772712 | 0,00624442 | Rptor | regulatory associated protein of MTOR, complex 1 [Source:MGI Symbol;Acc:MGI:1921620] | 92,48 | 1,9 |
| ENSMUSG00000013160 | 676,2240055 | 1149,6 | 39,7 | 5,344879598 | 2,80125E-06 | Atp6v0d1 | ATPase, H+ transporting, lysosomal V0 subunit D1 [Source:MGI Symbol;Acc:MGI:1201778] | 99,61 | 35,3 |
| ENSMUSG00000024079 | 59,01838911 | 66,4 | 91,4 | 5,34494194 | 0,040314285 | Eif2ak2 | eukaryotic translation initiation factor 2-alpha kinase 2 [Source:MGI Symbol;Acc:MGI:1353449] | 83,23 | 11,9 |
| ENSMUSG00000041879 | 306,2951113 | 367,6 | 66,3 | 5,348336794 | 0,000448202 | Ipo9 | importin 9 [Source:MGI Symbol;Acc:MGI:1918944] | 97,09 | 11,7 |
| ENSMUSG00000026544 | 42,0599491 | 59,7 | 8,0 | 5,348813411 | 0,024810203 | Dusp23 | dual specificity phosphatase 23 [Source:MGI Symbol;Acc:MGI:1915690] | 92,84 | 6,7 |
| ENSMUSG00000024151 | 128,5641571 | 87,2 | 59,1 | 5,350553024 | 0,013044142 | Msh2 | mutS homolog 2 (E. coli) [Source:MGI Symbol;Acc:MGI:101816] | 91,28 | 13,7 |
| ENSMUSG00000014329 | 29,02301004 | 40,0 | 2,8 | 5,350829525 | 0,035908736 | Bicc1 | BicC family RNA binding protein 1 [Source:MGI Symbol;Acc:MGI:1933388] | 60,64 | 0,9 |
| ENSMUSG00000038463 | 18,44373765 | 29,9 | 1,0 | 5,351212924 | 0,025461744 | Olfml2b | olfactomedin-like 2B [Source:MGI Symbol;Acc:MGI:2443310] | 72,21 | 0,6 |
| ENSMUSG00000052812 | 63,7793688 | 49,2 | 3,0 | 5,352528395 | 0,017219893 | Atad2b | ATPase family, AAA domain containing 2B [Source:MGI Symbol;Acc:MGI:2444798] | 72,50 | 0,5 |
| ENSMUSG00000026956 | 138,0467381 | 179,0 | 65,5 | 5,355933789 | 0,006717397 | Uap1l1 | UDP-N-acteylglucosamine pyrophosphorylase 1-like 1 [Source:MGI Symbol;Acc:MGI:2443318] | 66,45 | 14,1 |
| ENSMUSG00000055125 | 31,06459717 | 33,6 | 1,2 | 5,358167568 | 0,039869743 | M5C1000I18Rik | RIKEN cDNA M5C1000I18 gene [Source:MGI Symbol;Acc:MGI:3642038] | #DIV/0! | #DIV/0! |
| ENSMUSG00000040387 | 64,40104852 | 121,4 | 4,0 | 5,360070029 | 0,005859943 | Klhl32 | kelch-like 32 [Source:MGI Symbol;Acc:MGI:3612790] | 59,71 | 0,7 |
| ENSMUSG00000030725 | 68,71713118 | 80,3 | 30,9 | 5,362181236 | 0,024173494 | Lipt2 | lipoyl(octanoyl) transferase 2 (putative) [Source:MGI Symbol;Acc:MGI:1914414] | 87,23 | 15,5 |
| ENSMUSG00000026163 | 116,4823877 | 178,4 | 211,3 | 5,36330127 | 0,009960995 | Sphkap | SPHK1 interactor, AKAP domain containing [Source:MGI Symbol;Acc:MGI:1924879] | 78,15 | 17,2 |
| ENSMUSG00000097756 | 167,5086382 | 237,9 | 85,0 | 5,363340084 | 0,015968705 | A730056A06Rik | RIKEN cDNA A730056A06 gene [Source:MGI Symbol;Acc:MGI:2442718] | 99,14 | 18,9 |
| ENSMUSG00000042213 | 24,65490625 | 31,9 | 1,0 | 5,363825098 | 0,032804617 | Zfand4 | zinc finger, AN1-type domain 4 [Source:MGI Symbol;Acc:MGI:1914742] | 61,43 | 0,8 |
| ENSMUSG00000039886 | 152,1831028 | 149,5 | 49,9 | 5,364334413 | 0,001008328 | Tmem120a | transmembrane protein 120A [Source:MGI Symbol;Acc:MGI:2686991] | 87,56 | 26,9 |
| ENSMUSG00000037211 | 39,55720162 | 51,5 | 7,4 | 5,364618423 | 0,029289201 | Spry1 | sprouty homolog 1 (Drosophila) [Source:MGI Symbol;Acc:MGI:1345139] | 78,72 | 3,7 |
| ENSMUSG00000048720 | 132,6627671 | 212,3 | 51,8 | 5,365289613 | 0,009369036 | Tbc1d12 | TBC1D12: TBC1 domain family, member 12 [Source:MGI Symbol;Acc:MGI:2384803] | 92,30 | 11,3 |
| ENSMUSG00000062380 | 35,77708669 | 54,3 | 1,8 | 5,36749714 | 0,006335422 | Tubb3 | tubulin, beta 3 class III [Source:MGI Symbol;Acc:MGI:107813] | 91,70 | 1,2 |
| ENSMUSG00000019785 | 29,53577634 | 66,5 | 6,0 | 5,367978355 | 0,009076555 | Clvs2 | clavesin 2 [Source:MGI Symbol;Acc:MGI:2443223] | 54,18 | 1,0 |
| ENSMUSG00000073478 | 64,62469132 | 72,2 | 65,6 | 5,367983223 | 0,018731642 | D730003I15Rik | RIKEN cDNA D730003I15 gene [Source:MGI Symbol;Acc:MGI:1925971] | #DIV/0! | #DIV/0! |
| ENSMUSG00000022978 | 48,4605873 | 67,8 | 25,9 | 5,369376606 | 0,031528835 | Mis18a | MIS18 kinetochore protein homolog A (S. pombe) [Source:MGI Symbol;Acc:MGI:1913828] | 96,35 | 24,4 |
| ENSMUSG00000039086 | 150,9007672 | 177,6 | 163,3 | 5,371487835 | 0,018653255 | Ss18l1 | synovial sarcoma translocation gene on chromosome 18-like 1 [Source:MGI Symbol;Acc:MGI:2444061] | 98,01 | 27,1 |
| ENSMUSG00000018861 | 108,0207192 | 152,8 | 4,8 | 5,375411165 | 0,000191526 | Fdxr | ferredoxin reductase [Source:MGI Symbol;Acc:MGI:104724] | 95,55 | 2,2 |
| ENSMUSG00000041577 | 579,4489536 | 914,7 | 142,5 | 5,376070811 | 0,00404364 | Prelp | proline arginine-rich end leucine-rich repeat [Source:MGI Symbol;Acc:MGI:2151110] | 100,00 | 28,0 |
| ENSMUSG00000000600 | 270,4631676 | 317,4 | 335,8 | 5,378656261 | 0,015059 | Krit1 | KRIT1, ankyrin repeat containing [Source:MGI Symbol;Acc:MGI:1930618] | 62,61 | 16,3 |
| ENSMUSG00000020898 | 120,4410345 | 82,1 | 131,9 | 5,381373104 | 0,049797078 | Ctc1 | CTS telomere maintenance complex component 1 [Source:MGI Symbol;Acc:MGI:1916214] | 92,68 | 27,8 |
| ENSMUSG00000039509 | 140,5733908 | 56,6 | 5,5 | 5,381805667 | 0,032377354 | Nup133 | nucleoporin 133 [Source:MGI Symbol;Acc:MGI:2442620] | 55,10 | 1,5 |
| ENSMUSG00000034312 | 151,9868041 | 180,8 | 131,3 | 5,384164833 | 0,010323287 | Iqsec1 | IQ motif and Sec7 domain 1 [Source:MGI Symbol;Acc:MGI:1196356] | 53,63 | 5,9 |
| ENSMUSG00000064090 | 46,96482293 | 65,2 | 37,9 | 5,384258358 | 0,021514342 | Vrk2 | vaccinia related kinase 2 [Source:MGI Symbol;Acc:MGI:1917172] | 86,33 | 12,9 |
| ENSMUSG00000041975 | 40,56230471 | 56,5 | 7,4 | 5,387246311 | 0,033653786 | Mettl8 | methyltransferase like 8 [Source:MGI Symbol;Acc:MGI:2385142] | 58,01 | 2,8 |
| ENSMUSG00000060771 | 101,0212935 | 148,6 | 83,4 | 5,387307067 | 0,026697736 | Tsga10 | testis specific 10 [Source:MGI Symbol;Acc:MGI:2685063] | 73,16 | 4,7 |
| ENSMUSG00000028007 | 46,21254702 | 82,1 | 2,1 | 5,390231844 | 0,004490891 | Snx7 | sorting nexin 7 [Source:MGI Symbol;Acc:MGI:1923811] | 92,29 | 1,8 |
| ENSMUSG00000028195 | 39,13615415 | 71,7 | 1,6 | 5,390777609 | 0,002856502 | Cyr61 | cysteine rich protein 61 [Source:MGI Symbol;Acc:MGI:88613] | 96,91 | 2,2 |
| ENSMUSG00000104861 | 40,7106243 | 67,9 | 14,0 | 5,391619143 | 0,01933552 | 3110039M20Rik | RIKEN cDNA 3110039M20 gene [Source:MGI Symbol;Acc:MGI:1914543] | #DIV/0! | #DIV/0! |
| ENSMUSG00000042659 | 37,5039311 | 63,0 | 7,4 | 5,392106663 | 0,027651309 | Arrdc4 | arrestin domain containing 4 [Source:MGI Symbol;Acc:MGI:1913662] | 84,55 | 0,8 |
| ENSMUSG00000089990 | 67,48818822 | 88,4 | 18,8 | 5,392649505 | 0,016213872 | Gm15852 | predicted gene 15852 [Source:MGI Symbol;Acc:MGI:3801733] | #DIV/0! | #DIV/0! |
| ENSMUSG00000042104 | 58,98321749 | 77,8 | 16,3 | 5,393246414 | 0,016940586 | Uggt2 | UDP-glucose glycoprotein glucosyltransferase 2 [Source:MGI Symbol;Acc:MGI:1913685] | 57,08 | 1,0 |
| ENSMUSG00000061838 | 376,5989317 | 503,0 | 18,1 | 5,39460277 | 0,00015953 | Suclg2 | succinate-Coenzyme A ligase, GDP-forming, beta subunit [Source:MGI Symbol;Acc:MGI:1306824] | 72,70 | 4,4 |
| ENSMUSG00000021071 | 567,5835207 | 1024,7 | 73,1 | 5,395387432 | 0,010167511 | Trim9 | tripartite motif-containing 9 [Source:MGI Symbol;Acc:MGI:2137354] | 84,34 | 10,8 |
| ENSMUSG00000097520 | 33,07914642 | 65,5 | 24,0 | 5,396795377 | 0,016738321 | Gm26843 | predicted gene, 26843 [Source:MGI Symbol;Acc:MGI:5477337] | #DIV/0! | #DIV/0! |
| ENSMUSG00000047230 | 181,9973396 | 215,6 | 155,7 | 5,399020202 | 0,017275672 | Cldn2 | claudin 2 [Source:MGI Symbol;Acc:MGI:1276110] | 86,69 | 16,7 |
| ENSMUSG00000021884 | 183,1440219 | 273,1 | 174,5 | 5,399763483 | 0,014697605 | Hacl1 | 2-hydroxyacyl-CoA lyase 1 [Source:MGI Symbol;Acc:MGI:1929657] | 76,05 | 24,4 |
| ENSMUSG00000024773 | 39,24763512 | 52,8 | 56,9 | 5,403967702 | 0,023471074 | Atg2a | autophagy related 2A [Source:MGI Symbol;Acc:MGI:1916291] | 70,05 | 13,6 |
| ENSMUSG00000034371 | 38,39545666 | 31,8 | 1,0 | 5,404911065 | 0,028559261 | Tkfc | triokinase, FMN cyclase [Source:MGI Symbol;Acc:MGI:2385084] | 89,64 | 0,3 |
| ENSMUSG00000025916 | 18,81016384 | 27,9 | 0,6 | 5,405477357 | 0,021387317 | Ppp1r42 | protein phosphatase 1, regulatory subunit 42 [Source:MGI Symbol;Acc:MGI:1921138] | 88,33 | 0,0 |
| ENSMUSG00000028224 | 82,63974826 | 135,1 | 7,6 | 5,405989744 | 0,005718965 | Nbn | nibrin [Source:MGI Symbol;Acc:MGI:1351625] | 97,08 | 2,9 |
| ENSMUSG00000021221 | 43,85037615 | 91,8 | 2,6 | 5,409368336 | 0,001775381 | Dpf3 | D4, zinc and double PHD fingers, family 3 [Source:MGI Symbol;Acc:MGI:1917377] | 90,34 | 0,4 |
| ENSMUSG00000022637 | 54,87807667 | 45,0 | 1,8 | 5,410635808 | 0,027062154 | Cblb | Casitas B-lineage lymphoma b [Source:MGI Symbol;Acc:MGI:2146430] | 68,69 | 0,7 |
| ENSMUSG00000007610 | 50,16217777 | 54,0 | 64,7 | 5,412057977 | 0,037457717 | Gtpbp3 | GTP binding protein 3 [Source:MGI Symbol;Acc:MGI:1917609] | 78,78 | 17,9 |
| ENSMUSG00000032583 | 72,35131425 | 139,0 | 5,6 | 5,413876008 | 0,00213042 | Mon1a | MON1 homolog A (yeast) [Source:MGI Symbol;Acc:MGI:1920075] | 87,34 | 1,9 |
| ENSMUSG00000057409 | 34,03042942 | 76,3 | 15,7 | 5,41420056 | 0,012870305 | Zfp53 | zinc finger protein 53 [Source:MGI Symbol;Acc:MGI:99200] | 99,29 | 4,3 |
| ENSMUSG00000078317 | 121,2263267 | 133,6 | 18,5 | 5,417449629 | 0,001184981 | F8a | factor 8-associated gene A [Source:MGI Symbol;Acc:MGI:95474] | 61,56 | 9,0 |
| ENSMUSG00000020553 | 134,1894017 | 93,3 | 120,1 | 5,418195651 | 0,01696342 | Pctp | phosphatidylcholine transfer protein [Source:MGI Symbol;Acc:MGI:107375] | 88,35 | 22,6 |
| ENSMUSG00000026227 | 1185,538334 | 2028,4 | 361,1 | 5,42136497 | 0,019428823 | 2810459M11Rik | RIKEN cDNA 2810459M11 gene [Source:MGI Symbol;Acc:MGI:1920042] | 99,77 | 38,4 |
| ENSMUSG00000063550 | 17,57041015 | 33,0 | 1,0 | 5,421663272 | 0,017919652 | Nup98 | nucleoporin 98 [Source:MGI Symbol;Acc:MGI:109404] | 71,51 | 0,5 |
| ENSMUSG00000027247 | 223,2686465 | 422,0 | 83,3 | 5,421809809 | 0,000176485 | Arhgap1 | Rho GTPase activating protein 1 [Source:MGI Symbol;Acc:MGI:2445003] | 83,90 | 22,5 |
| ENSMUSG00000022747 | 48,40318017 | 53,3 | 3,0 | 5,421918667 | 0,03458377 | St3gal6 | ST3 beta-galactoside alpha-2,3-sialyltransferase 6 [Source:MGI Symbol;Acc:MGI:1888707] | 56,40 | 1,0 |
| ENSMUSG00000023330 | 70,60012342 | 73,0 | 23,1 | 5,422420526 | 0,014298916 | Dtwd1 | DTW domain containing 1 [Source:MGI Symbol;Acc:MGI:1916435] | 96,27 | 20,8 |
| ENSMUSG00000036501 | 178,3975537 | 304,6 | 65,7 | 5,422433258 | 0,00229615 | Fam13b | family with sequence similarity 13, member B [Source:MGI Symbol;Acc:MGI:2447834] | 98,22 | 12,6 |
| ENSMUSG00000036022 | 166,2470161 | 66,1 | 31,4 | 5,423469961 | 0,025618029 | Fam122b | family with sequence similarity 122, member B [Source:MGI Symbol;Acc:MGI:1926005] | 83,38 | 2,2 |
| ENSMUSG00000033852 | 65,0058097 | 66,4 | 4,0 | 5,424666575 | 0,008933079 | Gm28042 | predicted gene, 28042 [Source:MGI Symbol;Acc:MGI:5547778] | 23,79 | 0,4 |
| ENSMUSG00000031533 | 156,2092955 | 149,5 | 24,0 | 5,425521371 | 0,000822862 | Mrps31 | mitochondrial ribosomal protein S31 [Source:MGI Symbol;Acc:MGI:1913153] | 95,92 | 9,6 |
| ENSMUSG00000028274 | 118,8792497 | 160,5 | 5,8 | 5,426022613 | 0,002290724 | Rngtt | RNA guanylyltransferase and 5'-phosphatase [Source:MGI Symbol;Acc:MGI:1329041] | 98,75 | 1,5 |
| ENSMUSG00000037824 | 299,1013482 | 254,3 | 58,3 | 5,426956355 | 0,008651939 | Tspan14 | tetraspanin 14 [Source:MGI Symbol;Acc:MGI:1196325] | 99,38 | 28,9 |
| ENSMUSG00000042284 | 18,409499 | 30,1 | 1,0 | 5,42732321 | 0,03834123 | Itga1 | integrin alpha 1 [Source:MGI Symbol;Acc:MGI:96599] | 46,70 | 0,2 |
| ENSMUSG00000038094 | 154,2412151 | 291,3 | 159,6 | 5,427775672 | 0,003374021 | Atp13a4 | ATPase type 13A4 [Source:MGI Symbol;Acc:MGI:1924456] | 92,40 | 26,9 |
| ENSMUSG00000051910 | 155,8529992 | 207,5 | 84,4 | 5,428418274 | 0,005026681 | Sox6 | SRY (sex determining region Y)-box 6 [Source:MGI Symbol;Acc:MGI:98368] | 83,28 | 2,5 |
| ENSMUSG00000045777 | 37,49171827 | 51,6 | 41,9 | 5,428513667 | 0,045031522 | Ifitm10 | interferon induced transmembrane protein 10 [Source:MGI Symbol;Acc:MGI:2444776] | 29,33 | 6,7 |
| ENSMUSG00000046417 | 150,3168692 | 215,8 | 47,6 | 5,429191293 | 0,014146058 | Lrrc75a | leucine rich repeat containing 75A [Source:MGI Symbol;Acc:MGI:2682293] | 98,49 | 26,7 |
| ENSMUSG00000086767 | 14,75358795 | 31,4 | 0,9 | 5,429549593 | 0,018089692 | Gm13070 | predicted gene 13070 [Source:MGI Symbol;Acc:MGI:3651970] | 13,41 | 0,1 |
| ENSMUSG00000102101 | 80,18919053 | 106,3 | 26,6 | 5,430308039 | 0,009242871 | Zbtb11os1 | zinc finger and BTB domain containing 11, opposite strand 1 [Source:MGI Symbol;Acc:MGI:1913641] | 98,02 | 18,4 |
| ENSMUSG00000028538 | 106,9967469 | 109,9 | 3,8 | 5,431876732 | 0,008128452 | St3gal3 | ST3 beta-galactoside alpha-2,3-sialyltransferase 3 [Source:MGI Symbol;Acc:MGI:1316659] | 90,53 | 2,5 |
| ENSMUSG00000100113 | 37,86784586 | 57,4 | 3,7 | 5,43226209 | 0,015495909 | Gm28592 | predicted gene 28592 [Source:MGI Symbol;Acc:MGI:5579298] | #DIV/0! | #DIV/0! |
| ENSMUSG00000053263 | 51,14497294 | 78,9 | 12,0 | 5,433111366 | 0,007343792 | Gm12592 | predicted gene 12592 [Source:MGI Symbol;Acc:MGI:3649892] | #DIV/0! | #DIV/0! |
| ENSMUSG00000028019 | 54,4550751 | 85,4 | 2,2 | 5,434216369 | 0,004628223 | Pdgfc | platelet-derived growth factor, C polypeptide [Source:MGI Symbol;Acc:MGI:1859631] | 80,46 | 1,4 |
| ENSMUSG00000028518 | 39,76295043 | 90,4 | 2,5 | 5,435437872 | 0,002076273 | Prkaa2 | protein kinase, AMP-activated, alpha 2 catalytic subunit [Source:MGI Symbol;Acc:MGI:1336173] | 76,64 | 0,6 |
| ENSMUSG00000030271 | 71,85631577 | 90,0 | 56,4 | 5,436052982 | 0,009369036 | Ogg1 | 8-oxoguanine DNA-glycosylase 1 [Source:MGI Symbol;Acc:MGI:1097693] | 98,17 | 28,7 |
| ENSMUSG00000033308 | 32,37722038 | 36,6 | 1,8 | 5,437748268 | 0,038466782 | Dpyd | dihydropyrimidine dehydrogenase [Source:MGI Symbol;Acc:MGI:2139667] | 70,33 | 0,5 |
| ENSMUSG00000045287 | 145,7676093 | 77,5 | 122,9 | 5,440199896 | 0,046955815 | Rtn4rl1 | reticulon 4 receptor-like 1 [Source:MGI Symbol;Acc:MGI:2661375] | 90,98 | 22,9 |
| ENSMUSG00000038894 | 207,7813292 | 217,7 | 57,0 | 5,440341617 | 0,000749905 | Irs2 | insulin receptor substrate 2 [Source:MGI Symbol;Acc:MGI:109334] | 97,76 | 13,8 |
| ENSMUSG00000039501 | 62,36387885 | 107,2 | 2,5 | 5,442586985 | 0,000521979 | Znfx1 | zinc finger, NFX1-type containing 1 [Source:MGI Symbol;Acc:MGI:2138982] | 85,88 | 1,0 |
| ENSMUSG00000025576 | 148,3088486 | 205,8 | 124,3 | 5,443540579 | 0,006792606 | Rbfox3 | RNA binding protein, fox-1 homolog (C. elegans) 3 [Source:MGI Symbol;Acc:MGI:106368] | 75,02 | 15,3 |
| ENSMUSG00000025200 | 32,7377802 | 43,3 | 2,0 | 5,444513316 | 0,021217726 | Cwf19l1 | CWF19-like 1, cell cycle control (S. pombe) [Source:MGI Symbol;Acc:MGI:1919752] | 70,42 | 1,1 |
| ENSMUSG00000086810 | 23,94340971 | 50,6 | 1,8 | 5,445055849 | 0,012114415 | Gm13110 | predicted gene 13110 [Source:MGI Symbol;Acc:MGI:3651716] | #DIV/0! | #DIV/0! |
| ENSMUSG00000074782 | 60,28583552 | 74,2 | 73,8 | 5,445248882 | 0,04104837 | 4833422C13Rik | RIKEN cDNA 4833422C13 gene [Source:MGI Symbol;Acc:MGI:3603826] | 64,31 | 10,6 |
| ENSMUSG00000070047 | 34,4352838 | 48,2 | 3,0 | 5,446624196 | 0,02327851 | Fat1 | FAT atypical cadherin 1 [Source:MGI Symbol;Acc:MGI:109168] | 28,48 | 0,3 |
| ENSMUSG00000024246 | 79,45270201 | 73,2 | 88,7 | 5,448159046 | 0,032811127 | Thumpd2 | THUMP domain containing 2 [Source:MGI Symbol;Acc:MGI:1919417] | 98,29 | 29,4 |
| ENSMUSG00000037982 | 21,8383313 | 26,3 | 0,9 | 5,450777879 | 0,047550795 | Gm9725 | predicted gene 9725 [Source:MGI Symbol;Acc:MGI:3646552] | #DIV/0! | #DIV/0! |
| ENSMUSG00000024526 | 22,02562061 | 40,3 | 1,2 | 5,451190048 | 0,030803761 | Cidea | leath-inducing DNA fragmentation factor, alpha subunit-like effector A [Source:MGI Symbol;Acc:MGI:1270 | 89,95 | 2,1 |
| ENSMUSG00000055733 | 58,62561767 | 78,8 | 102,6 | 5,451644536 | 0,017005951 | Nap1l3 | nucleosome assembly protein 1-like 3 [Source:MGI Symbol;Acc:MGI:1859565] | 93,90 | 26,9 |
| ENSMUSG00000046861 | 88,79878157 | 142,5 | 17,6 | 5,452049142 | 0,014295743 | Hectd3 | HECT domain containing 3 [Source:MGI Symbol;Acc:MGI:1923858] | 89,29 | 3,8 |
| ENSMUSG00000029718 | 103,6138276 | 103,0 | 59,9 | 5,454467177 | 0,048578581 | Pcolce | procollagen C-endopeptidase enhancer protein [Source:MGI Symbol;Acc:MGI:105099] | 80,66 | 21,6 |
| ENSMUSG00000049556 | 64,4953343 | 101,6 | 58,1 | 5,454532372 | 0,012408832 | Lingo1 | leucine rich repeat and Ig domain containing 1 [Source:MGI Symbol;Acc:MGI:1915522] | 85,57 | 19,2 |
| ENSMUSG00000086773 | 33,37767068 | 54,4 | 23,9 | 5,45457637 | 0,035504063 | Gm16192 | predicted gene 16192 [Source:MGI Symbol;Acc:MGI:3802024] | 48,88 | 11,6 |
| ENSMUSG00000036273 | 33,33597089 | 51,6 | 1,8 | 5,457645951 | 0,005948324 | Lrrk2 | leucine-rich repeat kinase 2 [Source:MGI Symbol;Acc:MGI:1913975] | 54,70 | 0,5 |
| ENSMUSG00000022305 | 69,20044791 | 42,5 | 1,2 | 5,4595199 | 0,027238648 | Lrp12 | low density lipoprotein-related protein 12 [Source:MGI Symbol;Acc:MGI:2443132] | 73,67 | 0,5 |
| ENSMUSG00000050144 | 429,4937676 | 684,1 | 65,9 | 5,460875972 | 0,002750301 | Slc25a44 | solute carrier family 25, member 44 [Source:MGI Symbol;Acc:MGI:2444391] | 99,92 | 21,4 |
| ENSMUSG00000031166 | 355,5496308 | 475,1 | 135,7 | 5,461371592 | 0,007343792 | Wdr13 | WD repeat domain 13 [Source:MGI Symbol;Acc:MGI:1914661] | 95,32 | 25,6 |
| ENSMUSG00000009356 | 12,35457085 | 30,6 | 1,0 | 5,461423287 | 0,024247603 | Lpo | lactoperoxidase [Source:MGI Symbol;Acc:MGI:1923363] | 75,14 | 0,8 |
| ENSMUSG00000042804 | 49,53305503 | 71,1 | 70,2 | 5,462139142 | 0,023114372 | Gpr153 | G protein-coupled receptor 153 [Source:MGI Symbol;Acc:MGI:1916157] | 87,79 | 19,4 |
| ENSMUSG00000055116 | 243,827577 | 404,0 | 76,5 | 5,464144449 | 0,010814203 | Arntl | aryl hydrocarbon receptor nuclear translocator-like [Source:MGI Symbol;Acc:MGI:1096381] | 98,52 | 25,9 |
| ENSMUSG00000021938 | 67,9335764 | 119,2 | 4,6 | 5,464844225 | 0,006156797 | Pspc1 | paraspeckle protein 1 [Source:MGI Symbol;Acc:MGI:1913895] | 63,14 | 1,9 |
| ENSMUSG00000038418 | 113,2011372 | 159,7 | 135,3 | 5,465962401 | 0,0135501 | Egr1 | early growth response 1 [Source:MGI Symbol;Acc:MGI:95295] | 66,41 | 18,7 |
| ENSMUSG00000021007 | 38,78456999 | 86,1 | 13,0 | 5,466084924 | 0,008345158 | Spata7 | spermatogenesis associated 7 [Source:MGI Symbol;Acc:MGI:2144877] | 98,41 | 4,2 |
| ENSMUSG00000070814 | 96,47165274 | 68,4 | 57,3 | 5,466422349 | 0,015050976 | 6330408A02Rik | RIKEN cDNA 6330408A02 gene [Source:MGI Symbol;Acc:MGI:2447816] | 77,16 | 15,1 |
| ENSMUSG00000027188 | 40,70215992 | 70,7 | 4,0 | 5,4674552 | 0,005495494 | Pamr1 | eptidase domain containing associated with muscle regeneration 1 [Source:MGI Symbol;Acc:MGI:244508 | 94,09 | 1,4 |
| ENSMUSG00000092356 | 59,75951135 | 74,2 | 25,9 | 5,467555125 | 0,018333315 | Gm20532 | predicted gene 20532 [Source:MGI Symbol;Acc:MGI:5141997] | #DIV/0! | #DIV/0! |
| ENSMUSG00000049744 | 24,06799077 | 26,8 | 0,9 | 5,468863907 | 0,04883748 | Arhgap15 | Rho GTPase activating protein 15 [Source:MGI Symbol;Acc:MGI:1923367] | 35,31 | 0,0 |
| ENSMUSG00000028597 | 51,71043834 | 72,4 | 57,5 | 5,471482187 | 0,041940594 | Gpx7 | glutathione peroxidase 7 [Source:MGI Symbol;Acc:MGI:1914555] | 82,62 | 12,7 |
| ENSMUSG00000022089 | 50,03840808 | 69,8 | 30,5 | 5,473758464 | 0,0231486 | Bin3 | bridging integrator 3 [Source:MGI Symbol;Acc:MGI:1929883] | 88,34 | 25,6 |
| ENSMUSG00000085609 | 53,36159942 | 49,7 | 60,9 | 5,476778111 | 0,043136566 | 1700016P03Rik | RIKEN cDNA 1700016P03 gene [Source:MGI Symbol;Acc:MGI:1922792] | #DIV/0! | #DIV/0! |
| ENSMUSG00000100838 | 52,44255131 | 78,6 | 20,0 | 5,477197813 | 0,021744159 | Gm29094 | predicted gene 29094 [Source:MGI Symbol;Acc:MGI:5579800] | 47,84 | 12,1 |
| ENSMUSG00000038271 | 84,98672492 | 65,7 | 134,9 | 5,478148902 | 0,031245874 | Iffo1 | intermediate filament family orphan 1 [Source:MGI Symbol;Acc:MGI:2444516] | 94,51 | 29,1 |
| ENSMUSG00000034263 | 206,2981395 | 217,9 | 126,3 | 5,48060667 | 0,00903868 | Vwa9 | von Willebrand factor A domain containing 9 [Source:MGI Symbol;Acc:MGI:1917132] | 88,93 | 24,5 |
| ENSMUSG00000032322 | 14,12803959 | 29,8 | 0,9 | 5,483934128 | 0,02720012 | Pstpip1 | proline-serine-threonine phosphatase-interacting protein 1 [Source:MGI Symbol;Acc:MGI:1321396] | 76,90 | 1,2 |
| ENSMUSG00000041380 | 80,33814285 | 75,5 | 7,3 | 5,48416591 | 0,021697227 | Htr2c | 5-hydroxytryptamine (serotonin) receptor 2C [Source:MGI Symbol;Acc:MGI:96281] | 79,74 | 2,2 |
| ENSMUSG00000021303 | 62,43571458 | 81,7 | 105,4 | 5,484607301 | 0,01751344 | Gng4 | guanine nucleotide binding protein (G protein), gamma 4 [Source:MGI Symbol;Acc:MGI:102703] | 97,18 | 27,9 |
| ENSMUSG00000040263 | 158,3597852 | 109,4 | 138,6 | 5,484828272 | 0,008561703 | Klhdc4 | kelch domain containing 4 [Source:MGI Symbol;Acc:MGI:2384569] | 98,59 | 31,3 |
| ENSMUSG00000025089 | 178,8900884 | 248,7 | 152,9 | 5,48563103 | 0,017150991 | Gfra1 | glial cell line derived neurotrophic factor family receptor alpha 1 [Source:MGI Symbol;Acc:MGI:1100842] | 89,18 | 21,6 |
| ENSMUSG00000086350 | 33,36059231 | 67,7 | 7,0 | 5,486464988 | 0,010552384 | B230369F24Rik | RIKEN cDNA B230369F24 gene [Source:MGI Symbol;Acc:MGI:2443392] | 87,27 | 1,3 |
| ENSMUSG00000036167 | 285,7756848 | 370,8 | 51,0 | 5,487344802 | 0,004444364 | Pphln1 | periphilin 1 [Source:MGI Symbol;Acc:MGI:1917029] | 90,43 | 7,1 |
| ENSMUSG00000038046 | 35,77233774 | 71,2 | 41,9 | 5,48796357 | 0,014833244 | Rnmtl1 | RNA methyltransferase like 1 [Source:MGI Symbol;Acc:MGI:1914640] | 97,09 | 9,8 |
| ENSMUSG00000051469 | 572,4937631 | 925,6 | 96,4 | 5,489464119 | 0,001912223 | Zfp24 | zinc finger protein 24 [Source:MGI Symbol;Acc:MGI:1929704] | 98,68 | 9,4 |
| ENSMUSG00000020921 | 145,411372 | 86,7 | 73,0 | 5,489991973 | 0,013831112 | Tmem101 | transmembrane protein 101 [Source:MGI Symbol;Acc:MGI:1923797] | 98,77 | 27,8 |
| ENSMUSG00000074622 | 21,67604582 | 44,3 | 1,2 | 5,491683607 | 0,010956967 | Mafb | af musculoaponeurotic fibrosarcoma oncogene family, protein B (avian) [Source:MGI Symbol;Acc:MGI:104 | 85,45 | 0,2 |
| ENSMUSG00000040564 | 50,37638397 | 65,7 | 15,7 | 5,492329903 | 0,036425207 | Apoc1 | apolipoprotein C-I [Source:MGI Symbol;Acc:MGI:88053] | 82,97 | 20,9 |
| ENSMUSG00000024735 | 479,8537994 | 551,1 | 106,4 | 5,493694629 | 0,00025412 | Prpf19 | RP19/PSO4 pre-mRNA processing factor 19 homolog (S. cerevisiae) [Source:MGI Symbol;Acc:MGI:10624 | 41,02 | 10,6 |
| ENSMUSG00000021224 | 90,8230472 | 180,4 | 4,8 | 5,493740888 | 0,000581979 | Numb | numb gene homolog (Drosophila) [Source:MGI Symbol;Acc:MGI:107423] | 94,24 | 1,3 |
| ENSMUSG00000074892 | 57,73824113 | 92,7 | 23,9 | 5,493871364 | 0,012151295 | B3galt5 | DP-Gal:betaGlcNAc beta 1,3-galactosyltransferase, polypeptide 5 [Source:MGI Symbol;Acc:MGI:213687 | 78,97 | 3,7 |
| ENSMUSG00000044150 | 57,49905106 | 68,1 | 163,7 | 5,497613632 | 0,026592475 | A830080D01Rik | RIKEN cDNA A830080D01 gene [Source:MGI Symbol;Acc:MGI:2685992] | 77,79 | 11,3 |
| ENSMUSG00000038520 | 159,7322521 | 216,9 | 23,2 | 5,499180762 | 0,012277477 | Tbc1d17 | TBC1 domain family, member 17 [Source:MGI Symbol;Acc:MGI:2449973] | 29,86 | 3,6 |
| ENSMUSG00000001995 | 44,67862284 | 60,4 | 1,5 | 5,50050935 | 0,020289918 | Sipa1l2 | signal-induced proliferation-associated 1 like 2 [Source:MGI Symbol;Acc:MGI:2676970] | 77,18 | 0,8 |
| ENSMUSG00000038855 | 114,4023676 | 168,9 | 31,5 | 5,500882469 | 0,001220733 | Itpkb | inositol 1,4,5-trisphosphate 3-kinase B [Source:MGI Symbol;Acc:MGI:109235] | 91,73 | 4,4 |
| ENSMUSG00000027778 | 162,0402961 | 184,0 | 29,3 | 5,501239143 | 0,015806369 | Ift80 | intraflagellar transport 80 [Source:MGI Symbol;Acc:MGI:1915509] | 94,75 | 5,8 |
| ENSMUSG00000074282 | 39,50492851 | 74,7 | 9,0 | 5,503679161 | 0,01011622 | Zfp94 | zinc finger protein 94 [Source:MGI Symbol;Acc:MGI:107610] | 87,11 | 1,4 |
| ENSMUSG00000101429 | 64,45144504 | 66,5 | 66,6 | 5,505512403 | 0,044191388 | BC055402 | cDNA sequence BC055402 [Source:MGI Symbol;Acc:MGI:3039597] | #DIV/0! | #DIV/0! |
| ENSMUSG00000055980 | 37,88099509 | 50,1 | 2,0 | 5,506756412 | 0,011331665 | Irs1 | insulin receptor substrate 1 [Source:MGI Symbol;Acc:MGI:99454] | 44,04 | 0,3 |
| ENSMUSG00000033377 | 51,87570969 | 79,6 | 26,9 | 5,507562068 | 0,01780352 | Palmd | palmdelphin [Source:MGI Symbol;Acc:MGI:2148896] | 57,97 | 7,1 |
| ENSMUSG00000051335 | 40,48626259 | 29,3 | 1,0 | 5,509027159 | 0,040932701 | Gfod1 | glucose-fructose oxidoreductase domain containing 1 [Source:MGI Symbol;Acc:MGI:2145304] | 38,30 | 0,3 |
| ENSMUSG00000025323 | 74,88730598 | 98,1 | 2,6 | 5,510816279 | 0,00685741 | Sp4 | trans-acting transcription factor 4 [Source:MGI Symbol;Acc:MGI:107595] | 82,63 | 0,7 |
| ENSMUSG00000021556 | 132,2540098 | 85,0 | 408,5 | 5,511402028 | 0,030375698 | Golm1 | golgi membrane protein 1 [Source:MGI Symbol;Acc:MGI:1917329] | 95,12 | 31,9 |
| ENSMUSG00000020770 | 156,9876053 | 172,4 | 24,6 | 5,511688562 | 0,022819949 | Unk | unkempt homolog (Drosophila) [Source:MGI Symbol;Acc:MGI:2442456] | 99,40 | 8,1 |
| ENSMUSG00000002105 | 181,9347311 | 231,6 | 150,5 | 5,513110654 | 0,01265198 | Slc39a13 | solute carrier family 39 (metal ion transporter), member 13 [Source:MGI Symbol;Acc:MGI:1915677] | 90,26 | 21,2 |
| ENSMUSG00000038602 | 51,59381564 | 51,2 | 6,0 | 5,515291057 | 0,029991054 | Slc35f1 | solute carrier family 35, member F1 [Source:MGI Symbol;Acc:MGI:2139810] | 81,03 | 0,8 |
| ENSMUSG00000001020 | 54,25113349 | 63,1 | 4,2 | 5,517358651 | 0,027695266 | S100a4 | S100 calcium binding protein A4 [Source:MGI Symbol;Acc:MGI:1330282] | 84,47 | 12,3 |
| ENSMUSG00000027360 | 32,94416843 | 52,1 | 2,0 | 5,519504539 | 0,00739225 | Hdc | histidine decarboxylase [Source:MGI Symbol;Acc:MGI:96062] | 62,50 | 0,3 |
| ENSMUSG00000034853 | 1112,892301 | 1710,1 | 530,8 | 5,520561062 | 0,023510692 | Acot11 | acyl-CoA thioesterase 11 [Source:MGI Symbol;Acc:MGI:1913736] | 95,22 | 33,5 |
| ENSMUSG00000108255 | 57,80541327 | 91,9 | 7,3 | 5,521966151 | 0,007671013 | Gm16499 | predicted gene 16499 [Source:MGI Symbol;Acc:MGI:3704267] | #DIV/0! | #DIV/0! |
| ENSMUSG00000024683 | 243,4901099 | 311,7 | 131,3 | 5,522496998 | 0,005912613 | Mrpl16 | mitochondrial ribosomal protein L16 [Source:MGI Symbol;Acc:MGI:2137219] | 100,00 | 34,9 |
| ENSMUSG00000097431 | 57,78633704 | 85,0 | 6,5 | 5,524714354 | 0,004795666 | Gm26782 | predicted gene, 26782 [Source:MGI Symbol;Acc:MGI:5477276] | #DIV/0! | #DIV/0! |
| ENSMUSG00000031730 | 148,1374365 | 179,9 | 89,7 | 5,526031809 | 0,02720012 | Dhodh | dihydroorotate dehydrogenase [Source:MGI Symbol;Acc:MGI:1928378] | 88,38 | 20,9 |
| ENSMUSG00000050334 | 101,669339 | 151,4 | 4,0 | 5,527502629 | 0,000753834 | C130071C03Rik | RIKEN cDNA C130071C03 gene [Source:MGI Symbol;Acc:MGI:2443574] | #DIV/0! | #DIV/0! |
| ENSMUSG00000017774 | 93,0836724 | 32,9 | 1,0 | 5,529029356 | 0,035624713 | Myo1c | myosin IC [Source:MGI Symbol;Acc:MGI:106612] | 55,17 | 0,4 |
| ENSMUSG00000032727 | 79,94173684 | 106,9 | 3,9 | 5,53126916 | 0,017842158 | Mier3 | mesoderm induction early response 1, family member 3 [Source:MGI Symbol;Acc:MGI:2442317] | 85,08 | 0,9 |
| ENSMUSG00000043439 | 62,55814527 | 56,9 | 47,1 | 5,53479386 | 0,04416784 | E130012A19Rik | RIKEN cDNA E130012A19 gene [Source:MGI Symbol;Acc:MGI:2143991] | 98,27 | 21,9 |
| ENSMUSG00000042680 | 315,935694 | 453,4 | 198,1 | 5,536107997 | 0,004724652 | Garem | GRB2 associated, regulator of MAPK1 [Source:MGI Symbol;Acc:MGI:2685790] | 94,62 | 24,4 |
| ENSMUSG00000022827 | 83,1310516 | 79,8 | 50,8 | 5,536114851 | 0,023659521 | Rabl3 | RAB, member RAS oncogene family-like 3 [Source:MGI Symbol;Acc:MGI:1914907] | 98,26 | 25,2 |
| ENSMUSG00000021738 | 114,5674002 | 85,6 | 84,1 | 5,536439812 | 0,015236033 | Atxn7 | ataxin 7 [Source:MGI Symbol;Acc:MGI:2179277] | 81,02 | 7,6 |
| ENSMUSG00000006763 | 36,98850486 | 61,8 | 2,8 | 5,537396039 | 0,007941197 | Saal1 | serum amyloid A-like 1 [Source:MGI Symbol;Acc:MGI:1926185] | 59,86 | 1,0 |

| | | | | | | | | | |
|---|---|---|---|---|---|---|---|---|---|
| ENSMUSG00000014158 | 77,42367943 | 67,1 | 3,0 | 5,538894868 | 0,020710182 | Trpv4 | ransient receptor potential cation channel, subfamily V, member 4 [Source:MGI Symbol;Acc:MGI:1926945 | 83,33 | 0,7 |
| ENSMUSG00000024993 | 651,6130822 | 817,1 | 128,1 | 5,539054613 | 0,000154767 | Fam45a | family with sequence similarity 45, member A [Source:MGI Symbol;Acc:MGI:1915144] | 99,92 | 32,9 |
| ENSMUSG00000075410 | 74,21621458 | 95,9 | 9,7 | 5,540661278 | 0,019433512 | Gm11744 | predicted gene 11744 [Source:MGI Symbol;Acc:MGI:3649529] | 77,72 | 7,2 |
| ENSMUSG00000040270 | 22,08751978 | 31,7 | 1,0 | 5,542777655 | 0,023617627 | Bach2 | 3TB and CNC homology, basic leucine zipper transcription factor 2 [Source:MGI Symbol;Acc:MGI:894679 | 33,52 | 0,3 |
| ENSMUSG00000035151 | 54,56850385 | 74,5 | 31,9 | 5,543002193 | 0,01738593 | Elmod2 | ELMO/CED-12 domain containing 2 [Source:MGI Symbol;Acc:MGI:2445165] | 84,70 | 5,5 |
| ENSMUSG00000061778 | 49,65672687 | 86,2 | 7,0 | 5,543354872 | 0,004292292 | Mospd2 | motile sperm domain containing 2 [Source:MGI Symbol;Acc:MGI:1924013] | 92,23 | 1,4 |
| ENSMUSG00000048440 | 47,22793923 | 67,5 | 12,1 | 5,544408484 | 0,035076824 | Cyp4f16 | cytochrome P450, family 4, subfamily f, polypeptide 16 [Source:MGI Symbol;Acc:MGI:1917351] | 92,45 | 7,4 |
| ENSMUSG00000029279 | 51,76128074 | 71,9 | 8,0 | 5,545616211 | 0,011428385 | Brdt | bromodomain, testis-specific [Source:MGI Symbol;Acc:MGI:1891374] | 51,91 | 1,0 |
| ENSMUSG00000036282 | 246,6654373 | 457,1 | 42,7 | 5,546922191 | 0,004314422 | Naa30 | N(alpha)-acetyltransferase 30, NatC catalytic subunit [Source:MGI Symbol;Acc:MGI:1922259] | 94,61 | 4,8 |
| ENSMUSG00000038383 | 120,7547729 | 132,0 | 5,6 | 5,547303256 | 0,015641335 | Pigu | phosphatidylinositol glycan anchor biosynthesis, class U [Source:MGI Symbol;Acc:MGI:3039607] | 98,04 | 2,8 |
| ENSMUSG00000040506 | 53,65000798 | 93,6 | 36,0 | 5,548794857 | 0,007527607 | Ambra1 | autophagy/beclin 1 regulator 1 [Source:MGI Symbol;Acc:MGI:2443564] | 88,75 | 11,0 |
| ENSMUSG00000034055 | 283,1935741 | 477,2 | 17,5 | 5,548897256 | 0,00671267 | Phka1 | phosphorylase kinase alpha 1 [Source:MGI Symbol;Acc:MGI:97576] | 98,77 | 4,2 |
| ENSMUSG00000036672 | 45,84155389 | 70,6 | 37,9 | 5,549177874 | 0,030259366 | Cenpt | centromere protein T [Source:MGI Symbol;Acc:MGI:2443939] | 98,79 | 23,9 |
| ENSMUSG00000038546 | 189,2746541 | 351,6 | 10,8 | 5,549187909 | 0,001092312 | Ranbp9 | RAN binding protein 9 [Source:MGI Symbol;Acc:MGI:1928741] | 92,89 | 3,9 |
| ENSMUSG00000046658 | 31,36105176 | 50,7 | 1,8 | 5,554146541 | 0,009373845 | Zfp316 | zinc finger protein 316 [Source:MGI Symbol;Acc:MGI:1860402] | 61,60 | 0,7 |
| ENSMUSG00000035910 | 73,38351358 | 93,3 | 128,3 | 5,556519588 | 0,018488206 | Dcdc2a | doublecortin domain containing 2a [Source:MGI Symbol;Acc:MGI:2652818] | 46,16 | 16,1 |
| ENSMUSG00000020389 | 150,2954291 | 172,7 | 8,3 | 5,556811981 | 0,010341616 | Cdkl3 | cyclin-dependent kinase-like 3 [Source:MGI Symbol;Acc:MGI:2388268] | 73,67 | 1,4 |
| ENSMUSG00000028698 | 63,71575595 | 72,5 | 5,0 | 5,559295429 | 0,010825187 | Pik3r3 | hosphatidylinositol 3 kinase, regulatory subunit, polypeptide 3 (p55) [Source:MGI Symbol;Acc:MGI:109277 | 84,99 | 0,5 |
| ENSMUSG00000050714 | 77,05585443 | 101,8 | 17,6 | 5,56058995 | 0,013171957 | Zbtb26 | zinc finger and BTB domain containing 26 [Source:MGI Symbol;Acc:MGI:2444402] | 90,29 | 3,1 |
| ENSMUSG00000034121 | 22,3583427 | 52,2 | 2,0 | 5,561429157 | 0,00992454 | Mks1 | Meckel syndrome, type 1 [Source:MGI Symbol;Acc:MGI:3584243] | 97,15 | 1,3 |
| ENSMUSG00000086815 | 69,35698682 | 107,2 | 22,4 | 5,562658717 | 0,007797436 | 3110082J24Rik | RIKEN cDNA 3110082J24 gene [Source:MGI Symbol;Acc:MGI:3641994] | 97,87 | 25,0 |
| ENSMUSG00000022064 | 51,93468382 | 96,6 | 3,6 | 5,564004202 | 0,001860059 | Pibf1 | progesterone immunomodulatory binding factor 1 [Source:MGI Symbol;Acc:MGI:1261910] | 94,52 | 2,2 |
| ENSMUSG00000054556 | 53,77889183 | 69,8 | 15,1 | 5,564054716 | 0,031790417 | Gm4876 | predicted gene 4876 [Source:MGI Symbol;Acc:MGI:3647654] | 62,44 | 3,8 |
| ENSMUSG00000048581 | 55,95451855 | 79,9 | 36,0 | 5,57203895 | 0,016304962 | E130311K13Rik | RIKEN cDNA E130311K13 gene [Source:MGI Symbol;Acc:MGI:3607716] | 92,99 | 21,8 |
| ENSMUSG00000063808 | 152,5973659 | 228,5 | 87,7 | 5,573107316 | 0,016226964 | Gpatch1 | G patch domain containing 1 [Source:MGI Symbol;Acc:MGI:1914721] | 99,30 | 16,9 |
| ENSMUSG00000054931 | 26,77141163 | 47,1 | 2,0 | 5,576049046 | 0,033965089 | Zkscan4 | zinc finger with KRAB and SCAN domains 4 [Source:MGI Symbol;Acc:MGI:3649412] | 81,49 | 1,0 |
| ENSMUSG00000056673 | 35,41686128 | 82,5 | 4,0 | 5,57686756 | 0,00268435 | Kdm5d | lysine (K)-specific demethylase 5D [Source:MGI Symbol;Acc:MGI:99780] | 59,09 | 0,3 |
| ENSMUSG00000040097 | 389,7278472 | 723,2 | 97,6 | 5,577298774 | 0,002167561 | Flywch1 | FLYWCH-type zinc finger 1 [Source:MGI Symbol;Acc:MGI:2442638] | 85,94 | 24,6 |
| ENSMUSG00000037525 | 53,00814465 | 59,5 | 39,9 | 5,579077234 | 0,035624713 | Bcdin3d | BCDIN3 domain containing [Source:MGI Symbol;Acc:MGI:1922534] | 93,26 | 21,9 |
| ENSMUSG00000034177 | 12,37957314 | 28,3 | 0,6 | 5,580272051 | 0,037825201 | Rnf43 | ring finger protein 43 [Source:MGI Symbol;Acc:MGI:2442609] | 53,58 | 0,5 |
| ENSMUSG00000039756 | 280,2523614 | 168,1 | 146,9 | 5,581442215 | 0,002379889 | Dnttip2 | deoxynucleotidyltransferase, terminal, interacting protein 2 [Source:MGI Symbol;Acc:MGI:1923173] | 99,55 | 27,8 |
| ENSMUSG00000029311 | 36,27034236 | 72,3 | 14,8 | 5,581927425 | 0,021181533 | Hsd17b11 | hydroxysteroid (17-beta) dehydrogenase 11 [Source:MGI Symbol;Acc:MGI:2149821] | 36,05 | 3,0 |
| ENSMUSG00000043773 | 42,32899405 | 71,7 | 7,0 | 5,581980284 | 0,023617627 | 1700048O20Rik | RIKEN cDNA 1700048O20 gene [Source:MGI Symbol;Acc:MGI:1920637] | 97,04 | 4,3 |
| ENSMUSG00000074358 | 52,07140152 | 96,3 | 13,0 | 5,586075736 | 0,004888202 | Ccdc61 | coiled-coil domain containing 61 [Source:MGI Symbol;Acc:MGI:2685005] | 99,55 | 4,1 |
| ENSMUSG00000109061 | 107,3596597 | 111,8 | 51,8 | 5,58670949 | 0,008670473 | Map2k7 | mitogen-activated protein kinase kinase 7 [Source:MGI Symbol;Acc:MGI:1346871] | #DIV/0! | #DIV/0! |
| ENSMUSG00000039108 | 22,52747076 | 41,1 | 1,0 | 5,588067137 | 0,017399585 | Lsm14b | LSM14 homolog B (SCD6, S. cerevisiae) [Source:MGI Symbol;Acc:MGI:3040677] | 72,51 | 0,9 |
| ENSMUSG00000042570 | 59,3083196 | 81,0 | 102,8 | 5,589590242 | 0,022807088 | Mier2 | mesoderm induction early response 1, family member 2 [Source:MGI Symbol;Acc:MGI:1917677] | 74,93 | 17,8 |
| ENSMUSG00000006241 | 42,91855993 | 42,4 | 1,2 | 5,589653446 | 0,023308834 | Ccdc159 | coiled-coil domain containing 159 [Source:MGI Symbol;Acc:MGI:1914369] | 97,97 | 1,4 |
| ENSMUSG00000031367 | 444,1664251 | 647,2 | 189,4 | 5,589735535 | 0,0135501 | Ap1s2 | adaptor-related protein complex 1, sigma 2 subunit [Source:MGI Symbol;Acc:MGI:1889383] | 89,78 | 8,4 |
| ENSMUSG00000066687 | 71,23717652 | 101,9 | 53,9 | 5,594636675 | 0,012687612 | Zbtb16 | zinc finger and BTB domain containing 16 [Source:MGI Symbol;Acc:MGI:103222] | 90,19 | 10,1 |
| ENSMUSG00000078908 | 169,3694307 | 266,3 | 106,4 | 5,595303576 | 0,01092896 | Mon1b | MON1 homolog b (yeast) [Source:MGI Symbol;Acc:MGI:1923231] | 99,52 | 16,1 |
| ENSMUSG00000031337 | 81,13068869 | 116,2 | 3,4 | 5,597241888 | 0,002675426 | Mtm1 | X-linked myotubular myopathy gene 1 [Source:MGI Symbol;Acc:MGI:1099452] | 83,77 | 1,0 |
| ENSMUSG00000004677 | 58,9922027 | 65,8 | 136,7 | 5,597788871 | 0,028127231 | Myo9b | myosin IXb [Source:MGI Symbol;Acc:MGI:106624] | 77,76 | 9,0 |
| ENSMUSG00000052942 | 95,17308912 | 157,4 | 4,7 | 5,598432247 | 0,002180599 | Glis3 | GLIS family zinc finger 3 [Source:MGI Symbol;Acc:MGI:2444289] | 81,51 | 0,6 |
| ENSMUSG00000009647 | 154,6682692 | 292,2 | 14,0 | 5,599120417 | 0,003911784 | Mcu | mitochondrial calcium uniporter [Source:MGI Symbol;Acc:MGI:3026965] | 88,86 | 4,9 |
| ENSMUSG00000087341 | 37,01715415 | 61,2 | 5,0 | 5,59957148 | 0,014382513 | 0610040F04Rik | RIKEN cDNA 0610040F04 gene [Source:MGI Symbol;Acc:MGI:1922644] | 35,13 | 0,5 |
| ENSMUSG00000043621 | 54,60024646 | 79,6 | 7,4 | 5,600117041 | 0,010465506 | Ubxn10 | UBX domain protein 10 [Source:MGI Symbol;Acc:MGI:2443123] | 79,93 | 0,0 |
| ENSMUSG00000066438 | 14,39796355 | 32,5 | 0,6 | 5,602708835 | 0,017063717 | Plekhd1 | homology domain containing, family D (with coiled-coil domains) member 1 [Source:MGI Symbol;Acc:MGI | 74,58 | 0,1 |
| ENSMUSG00000044134 | 42,76077821 | 87,9 | 12,9 | 5,60362645 | 0,011734301 | Fam109a | family with sequence similarity 109, member A [Source:MGI Symbol;Acc:MGI:2442708] | 96,59 | 8,7 |
| ENSMUSG00000032231 | 55,7467367 | 99,6 | 8,5 | 5,604467336 | 0,011601357 | Anxa2 | annexin A2 [Source:MGI Symbol;Acc:MGI:88246] | 79,33 | 5,8 |
| ENSMUSG00000030774 | 356,4816932 | 628,2 | 163,6 | 5,6054888 | 0,000581979 | Pak1 | p21 protein (Cdc42/Rac)-activated kinase 1 [Source:MGI Symbol;Acc:MGI:1339975] | 94,44 | 27,3 |
| ENSMUSG00000029524 | 56,06843913 | 91,3 | 35,1 | 5,606705109 | 0,014423451 | Sirt4 | sirtuin 4 [Source:MGI Symbol;Acc:MGI:1922637] | 72,90 | 17,9 |
| ENSMUSG00000071256 | 62,13039924 | 113,2 | 79,9 | 5,60957578 | 0,021452754 | Zfp213 | zinc finger protein 213 [Source:MGI Symbol;Acc:MGI:3053094] | 97,26 | 27,5 |
| ENSMUSG00000031595 | 48,42183671 | 68,5 | 93,8 | 5,609613782 | 0,032298015 | Pdgfrl | platelet-derived growth factor receptor-like [Source:MGI Symbol;Acc:MGI:1916047] | 98,90 | 8,0 |
| ENSMUSG00000049092 | 31,6562816 | 51,4 | 2,0 | 5,610860979 | 0,01159044 | Gpr137c | G protein-coupled receptor 137C [Source:MGI Symbol;Acc:MGI:1917963] | 96,90 | 1,7 |
| ENSMUSG00000030226 | 172,8193055 | 393,2 | 23,4 | 5,613392196 | 0,002096022 | Lmo3 | LIM domain only 3 [Source:MGI Symbol;Acc:MGI:102810] | 84,51 | 1,6 |
| ENSMUSG00000034334 | 150,9984516 | 246,2 | 39,7 | 5,613464819 | 0,017418341 | Fam151b | family with sequence similarity 151, member B [Source:MGI Symbol;Acc:MGI:1921192] | 100,00 | 25,7 |
| ENSMUSG00000029104 | 20,33277461 | 42,7 | 0,9 | 5,615037591 | 0,00655659 | Htt | huntingtin [Source:MGI Symbol;Acc:MGI:96067] | 35,52 | 0,2 |
| ENSMUSG00000055799 | 95,13074546 | 137,2 | 3,8 | 5,617846137 | 0,006860894 | Tcf7l1 | transcription factor 7 like 1 (T cell specific, HMG box) [Source:MGI Symbol;Acc:MGI:1202876] | 95,67 | 2,5 |
| ENSMUSG00000002059 | 291,3869475 | 407,8 | 107,6 | 5,619591245 | 0,008771538 | Rab34 | RAB34, member RAS oncogene family [Source:MGI Symbol;Acc:MGI:104606] | 99,77 | 26,9 |
| ENSMUSG00000061950 | 86,15775939 | 79,2 | 1,9 | 5,624421695 | 0,006715922 | Ppp4r1 | protein phosphatase 4, regulatory subunit 1 [Source:MGI Symbol;Acc:MGI:1917601] | 85,37 | 1,0 |
| ENSMUSG00000041842 | 24,66957188 | 44,6 | 1,2 | 5,625838826 | 0,027763802 | Fhdc1 | FH2 domain containing 1 [Source:MGI Symbol;Acc:MGI:2684972] | 57,01 | 0,3 |
| ENSMUSG00000025510 | 54,8930753 | 71,5 | 93,8 | 5,631675432 | 0,038235961 | Cd151 | CD151 antigen [Source:MGI Symbol;Acc:MGI:1096360] | 92,74 | 30,5 |
| ENSMUSG00000030839 | 73,89289324 | 61,9 | 109,1 | 5,631697149 | 0,041952726 | Sergef | secretion regulating guanine nucleotide exchange factor [Source:MGI Symbol;Acc:MGI:1351630] | 81,73 | 24,8 |
| ENSMUSG00000034035 | 42,56709648 | 63,4 | 35,1 | 5,635383734 | 0,036170803 | Ccdc17 | coiled-coil domain containing 17 [Source:MGI Symbol;Acc:MGI:1915667] | 90,90 | 15,3 |
| ENSMUSG00000062190 | 101,6147427 | 162,5 | 8,0 | 5,642888885 | 0,00418273 | Lancl2 | LanC (bacterial lantibiotic synthetase component C)-like 2 [Source:MGI Symbol;Acc:MGI:1919085] | 99,24 | 2,1 |
| ENSMUSG00000033578 | 101,0593016 | 97,0 | 41,6 | 5,643522987 | 0,017981001 | Tmem35 | transmembrane protein 35 [Source:MGI Symbol;Acc:MGI:1914814] | 91,08 | 23,8 |
| ENSMUSG00000078349 | 22,78080469 | 42,3 | 1,8 | 5,649282175 | 0,024134552 | AW011738 | expressed sequence AW011738 [Source:MGI Symbol;Acc:MGI:2140540] | 95,18 | 0,7 |
| ENSMUSG00000027309 | 718,0173823 | 1021,0 | 592,9 | 5,650332698 | 0,008345158 | 4930402H24Rik | RIKEN cDNA 4930402H24 gene [Source:MGI Symbol;Acc:MGI:1923029] | 98,02 | 26,0 |
| ENSMUSG00000030089 | 24,80938876 | 31,0 | 1,0 | 5,651467344 | 0,044022476 | Slc41a3 | solute carrier family 41, member 3 [Source:MGI Symbol;Acc:MGI:1918949] | 61,75 | 0,8 |
| ENSMUSG00000020109 | 104,334819 | 107,0 | 17,6 | 5,651801984 | 0,012054878 | Dnajb12 | DnaJ heat shock protein family (Hsp40) member B12 [Source:MGI Symbol;Acc:MGI:1931881] | 77,14 | 4,3 |
| ENSMUSG00000030761 | 29,96157661 | 58,2 | 1,8 | 5,652070814 | 0,005639129 | Myo7a | myosin VIIA [Source:MGI Symbol;Acc:MGI:104510] | 66,07 | 0,4 |
| ENSMUSG00000003135 | 152,3656001 | 145,2 | 57,3 | 5,652531361 | 0,007930236 | Cnot11 | CCR4-NOT transcription complex, subunit 11 [Source:MGI Symbol;Acc:MGI:106580] | 89,06 | 19,0 |
| ENSMUSG00000020085 | 183,4257765 | 247,4 | 178,3 | 5,653738814 | 0,020350663 | Aifm2 | apoptosis-inducing factor, mitochondrion-associated 2 [Source:MGI Symbol;Acc:MGI:1918611] | 96,66 | 16,6 |
| ENSMUSG00000049164 | 39,24738011 | 76,0 | 10,0 | 5,654116552 | 0,011601357 | Zfp518a | zinc finger protein 518A [Source:MGI Symbol;Acc:MGI:1919922] | 76,04 | 1,5 |
| ENSMUSG00000045625 | 18,91234057 | 34,0 | 0,9 | 5,654221748 | 0,033653786 | Pigz | phosphatidylinositol glycan anchor biosynthesis, class Z [Source:MGI Symbol;Acc:MGI:2443822] | 68,94 | 0,0 |
| ENSMUSG00000020604 | 209,3703271 | 188,9 | 43,9 | 5,657575776 | 0,004301928 | Arsg | arylsulfatase G [Source:MGI Symbol;Acc:MGI:1921258] | 90,37 | 5,6 |
| ENSMUSG00000074682 | 66,99855953 | 129,6 | 47,2 | 5,659837403 | 0,010247119 | Zcchc3 | zinc finger, CCHC domain containing 3 [Source:MGI Symbol;Acc:MGI:1915167] | 96,80 | 17,5 |
| ENSMUSG00000038022 | 28,45528222 | 37,7 | 1,0 | 5,665337036 | 0,014593862 | Fam188b | family with sequence similarity 188, member B [Source:MGI Symbol;Acc:MGI:3583959] | 55,15 | 0,3 |
| ENSMUSG00000107741 | 59,72092685 | 86,1 | 9,1 | 5,665960763 | 0,012198906 | Gm2011 | predicted gene 2011 [Source:MGI Symbol;Acc:MGI:3780180] | #DIV/0! | #DIV/0! |
| ENSMUSG00000060798 | 95,2000057 | 104,8 | 65,9 | 5,667719825 | 0,005465327 | Intu | inturned planar cell polarity protein [Source:MGI Symbol;Acc:MGI:2443752] | 81,45 | 4,9 |
| ENSMUSG00000026004 | 114,8655585 | 197,5 | 9,9 | 5,668040416 | 0,000935031 | Kansl1l | KAT8 regulatory NSL complex subunit 1-like [Source:MGI Symbol;Acc:MGI:1915941] | 94,09 | 2,2 |
| ENSMUSG00000097350 | 40,91070558 | 82,9 | 27,7 | 5,66832738 | 0,021424278 | 4732491K20Rik | RIKEN cDNA 4732491K20 gene [Source:MGI Symbol;Acc:MGI:3026975] | #DIV/0! | #DIV/0! |
| ENSMUSG00000034800 | 53,60531558 | 65,9 | 7,0 | 5,668420249 | 0,018777336 | Zfp661 | zinc finger protein 661 [Source:MGI Symbol;Acc:MGI:1919430] | 60,58 | 0,5 |
| ENSMUSG00000027710 | 213,5382205 | 282,9 | 92,4 | 5,668792271 | 0,013322569 | Acad9 | acyl-Coenzyme A dehydrogenase family, member 9 [Source:MGI Symbol;Acc:MGI:1914272] | 78,63 | 22,1 |
| ENSMUSG00000105692 | 86,43641044 | 120,0 | 15,7 | 5,67029987 | 0,003489448 | Gm43737 | predicted gene 43737 [Source:MGI Symbol;Acc:MGI:5663874] | 48,64 | 3,3 |
| ENSMUSG00000038344 | 65,90344578 | 88,4 | 30,5 | 5,671734833 | 0,016730362 | Txlng | taxilin gamma [Source:MGI Symbol;Acc:MGI:3590652] | 56,64 | 6,1 |
| ENSMUSG00000028690 | 186,625565 | 293,3 | 112,9 | 5,673825283 | 0,007016941 | Mmachc | methylmalonic aciduria cblC type, with homocystinuria [Source:MGI Symbol;Acc:MGI:1914346] | 99,76 | 31,9 |
| ENSMUSG00000039661 | 66,06354869 | 78,9 | 58,7 | 5,676481703 | 0,047499556 | Dusp26 | dual specificity phosphatase 26 (putative) [Source:MGI Symbol;Acc:MGI:1914209] | 89,65 | 24,7 |
| ENSMUSG00000036975 | 88,50680002 | 83,8 | 179,6 | 5,679913358 | 0,029703799 | Tmem177 | transmembrane protein 177 [Source:MGI Symbol;Acc:MGI:1913593] | 92,05 | 29,1 |
| ENSMUSG00000030188 | 78,4617537 | 73,8 | 1,8 | 5,680027797 | 0,004999214 | Magohb | mago homolog B, exon junction complex core component [Source:MGI Symbol;Acc:MGI:1913691] | 68,92 | 1,9 |
| ENSMUSG00000041119 | 93,97944705 | 103,7 | 74,8 | 5,680814291 | 0,005408155 | Pde9a | phosphodiesterase 9A [Source:MGI Symbol;Acc:MGI:1277179] | 91,89 | 26,4 |
| ENSMUSG00000005267 | 39,6682248 | 66,9 | 2,0 | 5,680825121 | 0,00713294 | Zfp287 | zinc finger protein 287 [Source:MGI Symbol;Acc:MGI:2176561] | 69,82 | 0,4 |
| ENSMUSG00000030816 | 191,735303 | 238,7 | 99,5 | 5,681211064 | 0,003568742 | Rnf40 | ring finger protein 40 [Source:MGI Symbol;Acc:MGI:2142048] | 82,76 | 18,8 |
| ENSMUSG00000025384 | 54,23666422 | 87,7 | 9,0 | 5,686387687 | 0,005598936 | Faap100 | Fanconi anemia core complex associated protein 100 [Source:MGI Symbol;Acc:MGI:1919135] | 96,61 | 2,3 |
| ENSMUSG00000034105 | 70,80516179 | 117,8 | 76,3 | 5,686897433 | 0,012371023 | Tldc1 | TBC/LysM associated domain containing 1 [Source:MGI Symbol;Acc:MGI:1921597] | 99,83 | 28,3 |
| ENSMUSG00000039637 | 52,15639558 | 78,9 | 56,4 | 5,687191559 | 0,028160503 | Coro7 | coronin 7 [Source:MGI Symbol;Acc:MGI:1926135] | 81,03 | 12,3 |
| ENSMUSG00000033316 | 52,82944361 | 68,4 | 45,3 | 5,687960578 | 0,026432656 | Galnt9 | etyl-alpha-D-galactosamine:polypeptide N-acetylgalactosaminyltransferase 9 [Source:MGI Symbol;Acc:MG | 93,83 | 19,8 |
| ENSMUSG00000034109 | 193,7636741 | 252,1 | 92,5 | 5,688593934 | 0,004130605 | Golim4 | golgi integral membrane protein 4 [Source:MGI Symbol;Acc:MGI:1920374] | 77,72 | 15,4 |
| ENSMUSG00000063047 | 38,76903403 | 54,3 | 1,8 | 5,689960962 | 0,02437134 | Zfp780b | zinc finger protein 780B [Source:MGI Symbol;Acc:MGI:2444764] | 54,56 | 1,0 |
| ENSMUSG00000029408 | 260,2150368 | 429,6 | 81,7 | 5,691173884 | 0,001973287 | Abcb9 | ATP-binding cassette, sub-family B (MDR/TAP), member 9 [Source:MGI Symbol;Acc:MGI:1861729] | 70,35 | 11,6 |
| ENSMUSG00000003070 | 60,87926745 | 70,6 | 56,3 | 5,691292101 | 0,036572038 | Efna2 | ephrin A2 [Source:MGI Symbol;Acc:MGI:102707] | 86,19 | 25,8 |
| ENSMUSG00000068838 | 302,9191447 | 526,2 | 79,3 | 5,691606114 | 0,001198022 | Chn1os1 | chimerin 1, opposite strand 1 [Source:MGI Symbol;Acc:MGI:3702041] | 91,98 | 34,7 |
| ENSMUSG00000037086 | 82,87666545 | 73,4 | 9,2 | 5,692845156 | 0,009954247 | Prr32 | proline rich 32 [Source:MGI Symbol;Acc:MGI:1916050] | 97,39 | 8,6 |
| ENSMUSG00000025217 | 166,4056363 | 272,3 | 30,6 | 5,693583359 | 0,010288191 | Btrc | beta-transducin repeat containing protein [Source:MGI Symbol;Acc:MGI:1338871] | 86,53 | 3,5 |
| ENSMUSG00000026307 | 44,53046521 | 64,8 | 10,0 | 5,693788745 | 0,021281295 | Scly | selenocysteine lyase [Source:MGI Symbol;Acc:MGI:1355310] | 87,20 | 6,0 |
| ENSMUSG00000027698 | 88,12176022 | 99,0 | 37,9 | 5,694298755 | 0,021451613 | Nceh1 | neutral cholesterol ester hydrolase 1 [Source:MGI Symbol;Acc:MGI:2443191] | 88,25 | 2,5 |
| ENSMUSG00000037206 | 89,25039238 | 87,4 | 74,4 | 5,694602665 | 0,02626137 | Islr | immunoglobulin superfamily containing leucine-rich repeat [Source:MGI Symbol;Acc:MGI:1349645] | 94,52 | 24,6 |
| ENSMUSG00000096764 | 91,46375853 | 98,1 | 6,5 | 5,695083791 | 0,003714772 | Gm21985 | predicted gene 21985 [Source:MGI Symbol;Acc:MGI:5439454] | #DIV/0! | #DIV/0! |
| ENSMUSG00000033031 | 38,83489055 | 36,3 | 0,6 | 5,696770336 | 0,030939401 | C330027C09Rik | RIKEN cDNA C330027C09 gene [Source:MGI Symbol;Acc:MGI:2146335] | 58,94 | 0,4 |
| ENSMUSG00000020464 | 100,1624101 | 131,7 | 2,8 | 5,700609436 | 0,006314703 | Pnpt1 | polyribonucleotide nucleotidyltransferase 1 [Source:MGI Symbol;Acc:MGI:1918951] | 94,25 | 1,3 |
| ENSMUSG00000024177 | 53,79852744 | 69,1 | 89,6 | 5,702543604 | 0,040591858 | Nme4 | NME/NM23 nucleoside diphosphate kinase 4 [Source:MGI Symbol;Acc:MGI:1931148] | 96,78 | 29,8 |
| ENSMUSG00000028957 | 80,13924467 | 69,5 | 52,9 | 5,702723618 | 0,0277816 | Per3 | period circadian clock 3 [Source:MGI Symbol;Acc:MGI:1277134] | 71,03 | 7,3 |
| ENSMUSG00000059540 | 89,59628787 | 95,7 | 54,5 | 5,703319281 | 0,016039545 | Tcea2 | transcription elongation factor A (SII), 2 [Source:MGI Symbol;Acc:MGI:107368] | 91,08 | 23,0 |
| ENSMUSG00000028521 | 202,526713 | 286,3 | 201,2 | 5,703413765 | 0,010828118 | Slc35d1 | 35 (UDP-glucuronic acid/UDP-N-acetylgalactosamine dual transporter), member D1 [Source:MGI Symbol | 95,09 | 29,5 |
| ENSMUSG00000018143 | 71,95259731 | 79,4 | 156,2 | 5,704566638 | 0,021389235 | Mafk | af musculoaponeurotic fibrosarcoma oncogene family, protein K (avian) [Source:MGI Symbol;Acc:MGI:99 | 77,12 | 23,3 |
| ENSMUSG00000034509 | 55,32128164 | 79,5 | 21,3 | 5,706351466 | 0,034203497 | Mad2l1bp | MAD2L1 binding protein [Source:MGI Symbol;Acc:MGI:1913841] | 97,24 | 14,1 |
| ENSMUSG00000056004 | 28,48072612 | 31,0 | 1,0 | 5,712146595 | 0,043427113 | 9330182L06Rik | RIKEN cDNA 9330182L06 gene [Source:MGI Symbol;Acc:MGI:2443264] | 50,16 | 0,4 |
| ENSMUSG00000041995 | 221,7272645 | 314,3 | 57,1 | 5,714649476 | 0,006022019 | Zbed3 | zinc finger, BED type containing 3 [Source:MGI Symbol;Acc:MGI:1919364] | 82,79 | 18,2 |
| ENSMUSG00000084960 | 37,7308203 | 60,9 | 16,0 | 5,71599127 | 0,030555657 | B430010I23Rik | RIKEN cDNA B430010I23 gene [Source:MGI Symbol;Acc:MGI:1926099] | #DIV/0! | #DIV/0! |
| ENSMUSG00000037363 | 31,09946311 | 62,4 | 5,0 | 5,720621063 | 0,021467434 | Letm2 | leucine zipper-EF-hand containing transmembrane protein 2 [Source:MGI Symbol;Acc:MGI:2444979] | 93,99 | 2,7 |
| ENSMUSG00000035206 | 99,05057928 | 95,5 | 194,9 | 5,727743147 | 0,024256825 | Sppl2b | signal peptide peptidase like 2B [Source:MGI Symbol;Acc:MGI:1920468] | 97,95 | 32,3 |
| ENSMUSG00000026360 | 85,01976674 | 106,7 | 52,9 | 5,728892729 | 0,006905151 | Rgs2 | regulator of G-protein signaling 2 [Source:MGI Symbol;Acc:MGI:1098271] | 63,92 | 6,8 |
| ENSMUSG00000029557 | 63,97696424 | 94,6 | 45,3 | 5,734196306 | 0,01539322 | Ftsj2 | FtsJ RNA methyltransferase homolog 2 (E. coli) [Source:MGI Symbol;Acc:MGI:1915267] | 98,52 | 29,5 |
| ENSMUSG00000031828 | 45,77676705 | 83,6 | 30,5 | 5,735307196 | 0,011298525 | Klhl36 | kelch-like 36 [Source:MGI Symbol;Acc:MGI:2385305] | 99,83 | 21,4 |
| ENSMUSG00000038305 | 66,65329083 | 88,0 | 27,9 | 5,735624985 | 0,013075767 | Spats2l | spermatogenesis associated, serine-rich 2-like [Source:MGI Symbol;Acc:MGI:1914448] | 72,99 | 1,8 |
| ENSMUSG00000022529 | 69,20120931 | 83,3 | 111,8 | 5,735844405 | 0,028301947 | Zfp263 | zinc finger protein 263 [Source:MGI Symbol;Acc:MGI:1921370] | 88,81 | 13,9 |
| ENSMUSG00000021469 | 40,44044231 | 82,2 | 1,6 | 5,737954626 | 0,007360105 | Msx2 | msh homeobox 2 [Source:MGI Symbol;Acc:MGI:97169] | 85,20 | 1,5 |
| ENSMUSG00000026627 | 25,94364841 | 34,4 | 0,9 | 5,738325221 | 0,033679154 | Tmem206 | transmembrane protein 206 [Source:MGI Symbol;Acc:MGI:1914200] | 90,47 | 1,1 |
| ENSMUSG00000036054 | 79,31727667 | 97,7 | 74,9 | 5,743159059 | 0,01738593 | Sugp2 | SURP and G patch domain containing 2 [Source:MGI Symbol;Acc:MGI:2678085] | 59,73 | 11,2 |
| ENSMUSG00000022797 | 365,4044515 | 496,8 | 41,7 | 5,747243673 | 0,000672842 | Tfrc | transferrin receptor [Source:MGI Symbol;Acc:MGI:98822] | 91,23 | 7,8 |
| ENSMUSG00000075593 | 104,9861096 | 143,4 | 41,8 | 5,74831936 | 0,005058806 | Gal3st4 | galactose-3-O-sulfotransferase 4 [Source:MGI Symbol;Acc:MGI:1916254] | 57,74 | 5,4 |
| ENSMUSG00000027366 | 171,6413323 | 211,1 | 15,6 | 5,750632543 | 0,009223407 | Sppl2a | signal peptide peptidase like 2A [Source:MGI Symbol;Acc:MGI:1913802] | 72,20 | 2,2 |
| ENSMUSG00000048039 | 84,31758019 | 101,3 | 40,7 | 5,750643987 | 0,016777724 | Isg20l2 | interferon stimulated exonuclease gene 20-like 2 [Source:MGI Symbol;Acc:MGI:2140076] | 93,42 | 16,5 |
| ENSMUSG00000022313 | 54,42501211 | 97,8 | 18,5 | 5,750944521 | 0,009853147 | Utp23 | P23, small subunit (SSU) processome component, homolog (yeast) [Source:MGI Symbol;Acc:MGI:19258 | 89,64 | 8,5 |
| ENSMUSG00000084565 | 58,72559198 | 97,3 | 9,5 | 5,753321524 | 0,013688075 | Mir1901 | microRNA 1901 [Source:MGI Symbol;Acc:MGI:3811425] | #DIV/0! | #DIV/0! |
| ENSMUSG00000022125 | 197,4513445 | 236,3 | 8,8 | 5,755764042 | 0,007603603 | Cln5 | ceroid-lipofuscinosis, neuronal 5 [Source:MGI Symbol;Acc:MGI:2442253] | 97,27 | 4,8 |
| ENSMUSG00000040189 | 30,30370336 | 63,2 | 14,0 | 5,756865742 | 0,019275359 | Ccdc114 | coiled-coil domain containing 114 [Source:MGI Symbol;Acc:MGI:2446120] | 92,99 | 8,2 |
| ENSMUSG00000023232 | 75,52541629 | 71,6 | 35,1 | 5,75962519 | 0,023444444 | Serinc2 | serine incorporator 2 [Source:MGI Symbol;Acc:MGI:1919132] | 74,74 | 16,8 |
| ENSMUSG00000023206 | 52,16355763 | 69,1 | 53,6 | 5,760677308 | 0,026031374 | Il15ra | interleukin 15 receptor, alpha chain [Source:MGI Symbol;Acc:MGI:104644] | 72,17 | 19,3 |
| ENSMUSG00000035944 | 40,31868514 | 81,5 | 43,9 | 5,761053173 | 0,008982589 | Ttc38 | tetratricopeptide repeat domain 38 [Source:MGI Symbol;Acc:MGI:2146198] | 74,30 | 5,3 |
| ENSMUSG00000020130 | 169,4529699 | 182,5 | 10,3 | 5,763942781 | 0,000290728 | Tbc1d15 | TBC1 domain family, member 15 [Source:MGI Symbol;Acc:MGI:1913937] | 85,56 | 4,6 |
| ENSMUSG00000026740 | 111,0594496 | 120,1 | 49,0 | 5,766999882 | 0,006924761 | Dnajc1 | DnaJ heat shock protein family (Hsp40) member C1 [Source:MGI Symbol;Acc:MGI:103268] | 47,65 | 4,1 |
| ENSMUSG00000023192 | 45,1083655 | 30,6 | 0,9 | 5,767653952 | 0,047650897 | Grm2 | glutamate receptor, metabotropic 2 [Source:MGI Symbol;Acc:MGI:1351339] | 51,66 | 0,5 |
| ENSMUSG00000097787 | 62,6293 | 88,5 | 26,0 | 5,767684147 | 0,028122388 | 2700046G09Rik | RIKEN cDNA 2700046G09 gene [Source:MGI Symbol;Acc:MGI:1914438] | #DIV/0! | #DIV/0! |
| ENSMUSG00000031562 | 153,7280947 | 120,4 | 68,4 | 5,768410722 | 0,005583538 | Dctd | dCMP deaminase [Source:MGI Symbol;Acc:MGI:2444529] | 90,00 | 17,7 |
| ENSMUSG00000007617 | 207,8165457 | 341,2 | 54,9 | 5,768661296 | 0,000323446 | Homer1 | homer scaffolding protein 1 [Source:MGI Symbol;Acc:MGI:1347345] | 78,00 | 1,5 |
| ENSMUSG00000021975 | 62,56007078 | 72,4 | 54,9 | 5,768707643 | 0,022180393 | Ints9 | integrator complex subunit 9 [Source:MGI Symbol;Acc:MGI:1098533] | 89,96 | 13,6 |
| ENSMUSG00000099964 | 56,03495779 | 62,3 | 2,0 | 5,769756916 | 0,005487545 | Gm29514 | predicted gene 29514 [Source:MGI Symbol;Acc:MGI:5580220] | 42,32 | 1,0 |
| ENSMUSG00000039463 | 131,6895122 | 121,1 | 36,0 | 5,769917461 | 0,012135124 | Slc9a8 | solute carrier family 9 (sodium/hydrogen exchanger), member 8 [Source:MGI Symbol;Acc:MGI:1924281] | 97,42 | 5,2 |
| ENSMUSG00000022629 | 192,2397273 | 201,0 | 116,4 | 5,771752649 | 0,015550491 | Kif21a | kinesin family member 21A [Source:MGI Symbol;Acc:MGI:109188] | 90,76 | 5,8 |
| ENSMUSG00000108778 | 47,57188036 | 73,6 | 53,9 | 5,773693885 | 0,024970514 | RP24-136L4.5 | | #DIV/0! | #DIV/0! |
| ENSMUSG00000062116 | 75,92436316 | 91,8 | 124,8 | 5,776690669 | 0,016139505 | Zfp954 | zinc finger protein 954 [Source:MGI Symbol;Acc:MGI:1917764] | 99,31 | 31,2 |
| ENSMUSG00000057554 | 242,4541234 | 431,5 | 110,9 | 5,777755429 | 0,008276051 | Lgals8 | lectin, galactose binding, soluble 8 [Source:MGI Symbol;Acc:MGI:1928481] | 90,85 | 25,0 |
| ENSMUSG00000052688 | 27,75916445 | 32,6 | 1,0 | 5,778862007 | 0,049466459 | Rab7b | RAB7B, member RAS oncogene family [Source:MGI Symbol;Acc:MGI:2442295] | 45,09 | 0,5 |
| ENSMUSG00000024732 | 118,078908 | 119,6 | 27,7 | 5,779663889 | 0,006419354 | Ccdc86 | coiled-coil domain containing 86 [Source:MGI Symbol;Acc:MGI:1277220] | 76,33 | 11,7 |
| ENSMUSG00000025008 | 65,31830732 | 63,1 | 115,8 | 5,781227804 | 0,032460068 | Tctn3 | tectonic family member 3 [Source:MGI Symbol;Acc:MGI:1914840] | 86,59 | 23,4 |
| ENSMUSG00000011148 | 164,7194631 | 97,2 | 131,2 | 5,783684368 | 0,002150316 | Adssl1 | adenylosuccinate synthetase like 1 [Source:MGI Symbol;Acc:MGI:87947] | 95,45 | 30,3 |
| ENSMUSG00000032702 | 145,5172142 | 292,0 | 34,3 | 5,783728024 | 0,008239323 | Kank1 | KN motif and ankyrin repeat domains 1 [Source:MGI Symbol;Acc:MGI:2147707] | 96,18 | 3,7 |
| ENSMUSG00000053460 | 89,69505932 | 77,3 | 131,9 | 5,783806612 | 0,027119598 | Ggcx | gamma-glutamyl carboxylase [Source:MGI Symbol;Acc:MGI:1927655] | 95,78 | 30,0 |
| ENSMUSG00000027822 | 196,3046064 | 236,8 | 5,4 | 5,786476581 | 0,004145381 | Slc33a1 | solute carrier family 33 (acetyl-CoA transporter), member 1 [Source:MGI Symbol;Acc:MGI:1332247] | 85,20 | 1,7 |
| ENSMUSG00000035142 | 94,60033181 | 75,1 | 28,7 | 5,789815634 | 0,030440978 | Nubpl | nucleotide binding protein-like [Source:MGI Symbol;Acc:MGI:1924076] | 81,68 | 7,8 |
| ENSMUSG00000039620 | 45,74593559 | 70,4 | 2,8 | 5,791478346 | 0,008591434 | 6430573F11Rik | RIKEN cDNA 6430573F11 gene [Source:MGI Symbol;Acc:MGI:2442328] | 80,70 | 1,0 |
| ENSMUSG00000027858 | 62,24713501 | 92,8 | 29,9 | 5,791666352 | 0,015558559 | Tspan2 | tetraspanin 2 [Source:MGI Symbol;Acc:MGI:1917997] | 80,71 | 3,8 |
| ENSMUSG00000024790 | 125,9980952 | 105,9 | 83,8 | 5,792027805 | 0,008088217 | Sac3d1 | SAC3 domain containing 1 [Source:MGI Symbol;Acc:MGI:1913656] | 97,26 | 31,1 |
| ENSMUSG00000097284 | 23,28084908 | 36,0 | 0,6 | 5,795019862 | 0,015786213 | 4930480K23Rik | RIKEN cDNA 4930480K23 gene [Source:MGI Symbol;Acc:MGI:1922266] | #DIV/0! | #DIV/0! |
| ENSMUSG00000071724 | 17,71551981 | 36,0 | 0,9 | 5,795659456 | 0,023784817 | Smpd5 | sphingomyelin phosphodiesterase 5 [Source:MGI Symbol;Acc:MGI:3709877] | 64,23 | 1,4 |
| ENSMUSG00000005686 | 226,2279275 | 266,2 | 80,8 | 5,798349443 | 0,012282057 | Ampd3 | adenosine monophosphate deaminase 3 [Source:MGI Symbol;Acc:MGI:1096344] | 86,08 | 5,4 |
| ENSMUSG00000021385 | 174,8548887 | 161,5 | 4,6 | 5,799271595 | 0,013245991 | Ippk | inositol 1,3,4,5,6-pentakisphosphate 2-kinase [Source:MGI Symbol;Acc:MGI:1922928] | 99,54 | 3,6 |
| ENSMUSG00000021374 | 66,40245977 | 81,0 | 2,4 | 5,802568652 | 0,005585424 | Nup153 | nucleoporin 153 [Source:MGI Symbol;Acc:MGI:2385621] | 79,56 | 0,1 |

| | | | | | | | | | |
|---|---|---|---|---|---|---|---|---|---|
| ENSMUSG00000026404 | 53,40204801 | 82,4 | 4,8 | 5,802667382 | 0,016517421 | Ddx59 | DEAD (Asp-Glu-Ala-Asp) box polypeptide 59 [Source:MGI Symbol;Acc:MGI:1915247] | 97,71 | 1,8 |
| ENSMUSG00000061603 | 47,62087503 | 69,2 | 1,8 | 5,804499611 | 0,005859943 | Akap6 | A kinase (PRKA) anchor protein 6 [Source:MGI Symbol;Acc:MGI:3050566] | 53,71 | 0,5 |
| ENSMUSG00000035234 | 51,65067756 | 74,3 | 28,7 | 5,805478971 | 0,035985021 | Fam175a | family with sequence similarity 175, member A [Source:MGI Symbol;Acc:MGI:1917931] | 98,05 | 14,8 |
| ENSMUSG00000004018 | 89,24665848 | 100,9 | 46,2 | 5,809033276 | 0,011393132 | Fancl | Fanconi anemia, complementation group L [Source:MGI Symbol;Acc:MGI:1914280] | 84,72 | 14,6 |
| ENSMUSG00000021062 | 106,7096253 | 154,9 | 142,8 | 5,810190294 | 0,014620403 | Rab15 | RAB15, member RAS oncogene family [Source:MGI Symbol;Acc:MGI:1916865] | 83,29 | 26,3 |
| ENSMUSG00000025171 | 58,8575758 | 73,7 | 3,0 | 5,812467272 | 0,00599345 | Ubtd1 | ubiquitin domain containing 1 [Source:MGI Symbol;Acc:MGI:2385092] | 99,67 | 2,9 |
| ENSMUSG00000059742 | 59,04835105 | 103,6 | 12,0 | 5,813826678 | 0,009087284 | Kcnh7 | assium voltage-gated channel, subfamily H (eag-related), member 7 [Source:MGI Symbol;Acc:MGI:21595 | 30,76 | 0,8 |
| ENSMUSG00000020135 | 84,47420287 | 116,0 | 75,8 | 5,816521922 | 0,014295743 | Apc2 | adenomatosis polyposis coli 2 [Source:MGI Symbol;Acc:MGI:1346052] | 77,31 | 10,8 |
| ENSMUSG00000043999 | 46,70399363 | 107,6 | 3,6 | 5,817622846 | 0,002057817 | Gpr75 | G protein-coupled receptor 75 [Source:MGI Symbol;Acc:MGI:2441843] | 99,07 | 1,8 |
| ENSMUSG00000108713 | 31,54437513 | 49,5 | 2,0 | 5,818178191 | 0,026466872 | RP23-223O9.2 | | #DIV/0! | #DIV/0! |
| ENSMUSG00000103770 | 19,01813194 | 34,0 | 0,6 | 5,818950466 | 0,026680113 | Pcdha9 | protocadherin alpha 9 [Source:MGI Symbol;Acc:MGI:2447322] | 46,23 | 0,5 |
| ENSMUSG00000029875 | 17,93685794 | 39,8 | 1,0 | 5,81938997 | 0,014652128 | Ccdc184 | coiled-coil domain containing 184 [Source:MGI Symbol;Acc:MGI:2146066] | 92,51 | 0,8 |
| ENSMUSG00000073424 | 160,4360378 | 246,1 | 20,4 | 5,820659583 | 0,014747957 | Cyp4f15 | cytochrome P450, family 4, subfamily f, polypeptide 15 [Source:MGI Symbol;Acc:MGI:2146921] | 99,03 | 13,3 |
| ENSMUSG00000056459 | 50,52254096 | 84,8 | 1,6 | 5,821204384 | 0,002072649 | Zbtb25 | zinc finger and BTB domain containing 25 [Source:MGI Symbol;Acc:MGI:99197] | 84,85 | 1,2 |
| ENSMUSG00000022022 | 81,45345311 | 92,5 | 47,1 | 5,822307158 | 0,020838076 | Mtrf1 | mitochondrial translational release factor 1 [Source:MGI Symbol;Acc:MGI:2384815] | 93,98 | 28,3 |
| ENSMUSG00000020086 | 60,27381274 | 121,2 | 29,7 | 5,823274416 | 0,009208575 | H2afy2 | H2A histone family, member Y2 [Source:MGI Symbol;Acc:MGI:3037658] | 98,71 | 7,6 |
| ENSMUSG00000067586 | 22,72852514 | 47,8 | 0,6 | 5,824394601 | 0,012405249 | S1pr3 | sphingosine-1-phosphate receptor 3 [Source:MGI Symbol;Acc:MGI:1339365] | 67,48 | 0,6 |
| ENSMUSG00000075000 | 58,18303621 | 86,7 | 22,2 | 5,828801069 | 0,015462113 | Nrbf2 | nuclear receptor binding factor 2 [Source:MGI Symbol;Acc:MGI:1354950] | 52,99 | 11,9 |
| ENSMUSG00000002320 | 276,6730442 | 302,7 | 103,6 | 5,82895227 | 0,004309472 | Tm9sf1 | transmembrane 9 superfamily member 1 [Source:MGI Symbol;Acc:MGI:1921390] | 86,30 | 28,7 |
| ENSMUSG00000019960 | 301,1396462 | 514,0 | 40,1 | 5,829645373 | 0,006792606 | Dusp6 | dual specificity phosphatase 6 [Source:MGI Symbol;Acc:MGI:1914853] | 99,81 | 10,7 |
| ENSMUSG00000026556 | 63,36841073 | 92,3 | 14,0 | 5,829948426 | 0,011078997 | Vangl2 | vang-like 2 (van gogh, Drosophila) [Source:MGI Symbol;Acc:MGI:2135272] | 60,36 | 1,5 |
| ENSMUSG00000024298 | 190,4100557 | 134,4 | 102,6 | 5,830252773 | 0,006792606 | Zfp871 | zinc finger protein 871 [Source:MGI Symbol;Acc:MGI:1921793] | 81,40 | 15,7 |
| ENSMUSG00000037001 | 53,5041671 | 61,6 | 2,8 | 5,83149313 | 0,027453443 | Zfp39 | zinc finger protein 39 [Source:MGI Symbol;Acc:MGI:99183] | 81,95 | 1,0 |
| ENSMUSG00000032013 | 22,98603235 | 36,5 | 0,6 | 5,832010598 | 0,027238648 | Trim29 | tripartite motif-containing 29 [Source:MGI Symbol;Acc:MGI:1919419] | 39,18 | 0,7 |
| ENSMUSG00000022376 | 38,5075378 | 33,5 | 0,9 | 5,832530143 | 0,035493043 | Adcy8 | adenylate cyclase 8 [Source:MGI Symbol;Acc:MGI:1341110] | 65,85 | 0,4 |
| ENSMUSG00000029571 | 719,5971036 | 767,4 | 20,1 | 5,832771925 | 5,39421E-06 | Tmem106b | transmembrane protein 106B [Source:MGI Symbol;Acc:MGI:1919150] | 97,90 | 2,8 |
| ENSMUSG00000042535 | 149,3091049 | 180,9 | 47,1 | 5,832784332 | 0,00080457 | Gtpbp1 | GTP binding protein 1 [Source:MGI Symbol;Acc:MGI:109443] | 99,55 | 13,0 |
| ENSMUSG00000041801 | 194,9546992 | 368,9 | 56,1 | 5,834515784 | 0,003583651 | Phlda3 | pleckstrin homology-like domain, family A, member 3 [Source:MGI Symbol;Acc:MGI:1351485] | 99,81 | 32,9 |
| ENSMUSG00000028651 | 121,0657544 | 132,6 | 62,3 | 5,836706305 | 0,008531851 | Ppie | peptidylprolyl isomerase E (cyclophilin E) [Source:MGI Symbol;Acc:MGI:1917118] | 99,89 | 28,4 |
| ENSMUSG00000026767 | 278,8400747 | 460,5 | 110,6 | 5,837331042 | 0,004170187 | Fam188a | family with sequence similarity 188, member A [Source:MGI Symbol;Acc:MGI:1914210] | 99,42 | 22,6 |
| ENSMUSG00000022508 | 103,1978582 | 160,0 | 62,9 | 5,837348391 | 0,005254334 | Bcl6 | B cell leukemia/lymphoma 6 [Source:MGI Symbol;Acc:MGI:107187] | 92,43 | 19,2 |
| ENSMUSG00000027030 | 564,9515063 | 577,3 | 9,9 | 5,841018099 | 6,8013E-05 | Stk39 | serine/threonine kinase 39 [Source:MGI Symbol;Acc:MGI:1858416] | 92,69 | 2,6 |
| ENSMUSG00000047603 | 48,54137606 | 80,5 | 2,4 | 5,843930457 | 0,006499377 | Zfp235 | zinc finger protein 235 [Source:MGI Symbol;Acc:MGI:1929117] | 90,25 | 0,6 |
| ENSMUSG00000037643 | 267,3738268 | 517,7 | 37,1 | 5,844163516 | 0,002774311 | Prkci | protein kinase C, iota [Source:MGI Symbol;Acc:MGI:99260] | 91,24 | 5,7 |
| ENSMUSG00000026799 | 132,4415911 | 147,1 | 29,6 | 5,847175746 | 0,003328442 | Med27 | mediator complex subunit 27 [Source:MGI Symbol;Acc:MGI:1916225] | 75,61 | 11,6 |
| ENSMUSG00000039634 | 64,46499211 | 114,6 | 2,1 | 5,847391284 | 0,00303887 | Zfp189 | zinc finger protein 189 [Source:MGI Symbol;Acc:MGI:2444707] | 95,39 | 1,6 |
| ENSMUSG00000056260 | 59,00594861 | 106,8 | 13,0 | 5,847618553 | 0,003276704 | Lrif1 | ligand dependent nuclear receptor interacting factor 1 [Source:MGI Symbol;Acc:MGI:2445214] | 91,19 | 1,4 |
| ENSMUSG00000021185 | 86,68108595 | 178,3 | 78,1 | 5,848512016 | 0,007180942 | 9030617O03Rik | RIKEN cDNA 9030617O03 gene [Source:MGI Symbol;Acc:MGI:2444813] | 80,79 | 12,6 |
| ENSMUSG00000038213 | 69,62366517 | 85,1 | 10,2 | 5,848519664 | 0,015197515 | Tapbpl | TAP binding protein-like [Source:MGI Symbol;Acc:MGI:2384853] | 84,71 | 4,9 |
| ENSMUSG00000029179 | 72,92624879 | 80,0 | 83,5 | 5,849233223 | 0,023617627 | Zcchc4 | zinc finger, CCHC domain containing 4 [Source:MGI Symbol;Acc:MGI:1926046] | 36,87 | 6,3 |
| ENSMUSG00000028497 | 34,52020847 | 61,0 | 1,8 | 5,849394258 | 0,018449156 | Hacd4 | 3-hydroxyacyl-CoA dehydratase 4 [Source:MGI Symbol;Acc:MGI:1914025] | 46,31 | 0,4 |
| ENSMUSG00000032839 | 29,98922061 | 39,6 | 0,6 | 5,851240613 | 0,039810883 | Trpc1 | ransient receptor potential cation channel, subfamily C, member 1 [Source:MGI Symbol;Acc:MGI:109528 | 62,05 | 0,5 |
| ENSMUSG00000039835 | 117,7485938 | 183,7 | 4,2 | 5,85125354 | 0,002624115 | Nhsl1 | NHS-like 1 [Source:MGI Symbol;Acc:MGI:106390] | 80,14 | 0,8 |
| ENSMUSG00000020212 | 37,42713272 | 51,7 | 1,0 | 5,852078725 | 0,008240475 | Mdm1 | transformed mouse 3T3 cell double minute 1 [Source:MGI Symbol;Acc:MGI:96951] | 87,66 | 0,6 |
| ENSMUSG00000015597 | 127,471537 | 59,9 | 2,0 | 5,85541183 | 0,01149796 | Zfp318 | zinc finger protein 318 [Source:MGI Symbol;Acc:MGI:1889348] | 55,10 | 0,3 |
| ENSMUSG00000059897 | 53,65890982 | 63,0 | 14,0 | 5,855453624 | 0,02579482 | Zfp930 | zinc finger protein 930 [Source:MGI Symbol;Acc:MGI:2675306] | 99,02 | 4,2 |
| ENSMUSG00000020910 | 119,5041739 | 200,5 | 34,5 | 5,855781845 | 0,020588478 | Adprm | ADP-ribose/CDP-alcohol diphosphatase, manganese dependent [Source:MGI Symbol;Acc:MGI:1913608] | 100,00 | 8,6 |
| ENSMUSG00000104548 | 126,1325979 | 76,2 | 16,6 | 5,856287042 | 0,019869101 | Gm43857 | predicted gene 43857 [Source:MGI Symbol;Acc:MGI:5663994] | 40,35 | 2,4 |
| ENSMUSG00000074415 | 263,204636 | 310,3 | 146,6 | 5,857076283 | 0,006824289 | 2610203C20Rik | RIKEN cDNA 2610203C20 gene [Source:MGI Symbol;Acc:MGI:1917705] | #DIV/0! | #DIV/0! |
| ENSMUSG00000029029 | 115,5561494 | 122,8 | 63,8 | 5,858250713 | 0,006806315 | Wrap73 | WD repeat containing, antisense to Trp73 [Source:MGI Symbol;Acc:MGI:1891749] | 98,87 | 27,9 |
| ENSMUSG00000047539 | 296,0815649 | 456,1 | 29,1 | 5,858457206 | 0,001077359 | Fbxo28 | F-box protein 28 [Source:MGI Symbol;Acc:MGI:1261890] | 99,28 | 4,5 |
| ENSMUSG00000034825 | 108,3719681 | 198,8 | 25,6 | 5,858591171 | 0,006824289 | Nrip3 | nuclear receptor interacting protein 3 [Source:MGI Symbol;Acc:MGI:1925843] | 96,86 | 6,4 |
| ENSMUSG00000026698 | 106,5671524 | 89,3 | 83,5 | 5,86015555 | 0,024970514 | Pigc | phosphatidylinositol glycan anchor biosynthesis, class C [Source:MGI Symbol;Acc:MGI:1914542] | 52,99 | 10,6 |
| ENSMUSG00000085558 | 69,50004962 | 101,6 | 50,8 | 5,860443719 | 0,024560598 | 4930412C18Rik | RIKEN cDNA 4930412C18 gene [Source:MGI Symbol;Acc:MGI:2443490] | #DIV/0! | #DIV/0! |
| ENSMUSG00000037139 | 36,67659766 | 61,0 | 2,0 | 5,861374858 | 0,008546153 | Myom3 | myomesin family, member 3 [Source:MGI Symbol;Acc:MGI:2685280] | 43,14 | 0,8 |
| ENSMUSG00000052981 | 108,9374493 | 133,1 | 45,4 | 5,861776958 | 0,006990244 | Ube2ql1 | ubiquitin-conjugating enzyme E2Q family-like 1 [Source:MGI Symbol;Acc:MGI:1924230] | 94,10 | 14,1 |
| ENSMUSG00000039357 | 21,75149466 | 41,0 | 0,6 | 5,862283348 | 0,0136442 | Fut11 | fucosyltransferase 11 [Source:MGI Symbol;Acc:MGI:1920318] | 82,57 | 0,7 |
| ENSMUSG00000012640 | 69,17505733 | 77,7 | 20,3 | 5,862385419 | 0,02468674 | Zfp715 | zinc finger protein 715 [Source:MGI Symbol;Acc:MGI:1917180] | 79,65 | 2,7 |
| ENSMUSG00000055407 | 82,74163639 | 96,9 | 222,5 | 5,867527912 | 0,016112423 | Map6 | microtubule-associated protein 6 [Source:MGI Symbol;Acc:MGI:1201690] | 63,49 | 15,9 |
| ENSMUSG00000030409 | 98,68263535 | 114,1 | 50,9 | 5,869844545 | 0,022632531 | Dmpk | dystrophia myotonica-protein kinase [Source:MGI Symbol;Acc:MGI:94906] | 97,15 | 17,2 |
| ENSMUSG00000043257 | 72,10829784 | 107,6 | 49,9 | 5,870927716 | 0,00903868 | Pigv | phosphatidylinositol glycan anchor biosynthesis, class V [Source:MGI Symbol;Acc:MGI:2442480] | 81,23 | 12,8 |
| ENSMUSG00000042632 | 128,3722264 | 137,7 | 43,4 | 5,872213297 | 0,009099596 | Pla2g6 | phospholipase A2, group VI [Source:MGI Symbol;Acc:MGI:1859152] | 79,39 | 7,5 |
| ENSMUSG00000045744 | 58,50650747 | 108,7 | 28,7 | 5,872734416 | 0,012307119 | Bricd5 | BRICHOS domain containing 5 [Source:MGI Symbol;Acc:MGI:2441766] | 100,00 | 28,0 |
| ENSMUSG00000033278 | 59,24415842 | 61,2 | 2,8 | 5,875664404 | 0,016421201 | Ptprm | protein tyrosine phosphatase, receptor type, M [Source:MGI Symbol;Acc:MGI:102694] | 71,86 | 0,7 |
| ENSMUSG00000054871 | 60,51435404 | 94,1 | 25,9 | 5,877289462 | 0,020183342 | Tmem158 | transmembrane protein 158 [Source:MGI Symbol;Acc:MGI:1919559] | 99,08 | 16,3 |
| ENSMUSG00000022206 | 38,49670964 | 56,1 | 0,6 | 5,88129034 | 0,013136736 | Npr3 | natriuretic peptide receptor 3 [Source:MGI Symbol;Acc:MGI:97373] | 54,25 | 0,2 |
| ENSMUSG00000026879 | 252,6274757 | 279,0 | 26,0 | 5,882067205 | 0,006175831 | Gsn | gelsolin [Source:MGI Symbol;Acc:MGI:95851] | 96,37 | 12,8 |
| ENSMUSG00000039579 | 72,36835161 | 115,9 | 61,9 | 5,882905019 | 0,005454688 | Grin3a | glutamate receptor ionotropic, NMDA3A [Source:MGI Symbol;Acc:MGI:1933206] | 89,32 | 2,7 |
| ENSMUSG00000089945 | 57,18688874 | 42,1 | 1,0 | 5,882925993 | 0,031501162 | Pakap | paralemmin A kinase anchor protein [Source:MGI Symbol;Acc:MGI:5141924] | 38,01 | 0,2 |
| ENSMUSG00000044647 | 21,2891301 | 44,6 | 1,0 | 5,883553022 | 0,02387903 | Csrnp3 | cysteine-serine-rich nuclear protein 3 [Source:MGI Symbol;Acc:MGI:1925021] | 38,45 | 0,2 |
| ENSMUSG00000051236 | 34,06043722 | 65,1 | 1,2 | 5,884195039 | 0,004433184 | Msrb3 | methionine sulfoxide reductase B3 [Source:MGI Symbol;Acc:MGI:2443538] | 85,56 | 0,0 |
| ENSMUSG00000105509 | 25,20303043 | 56,5 | 1,2 | 5,884365295 | 0,009372077 | C130013H08Rik | RIKEN cDNA C130013H08 gene [Source:MGI Symbol;Acc:MGI:3697343] | 57,48 | 0,7 |
| ENSMUSG00000070709 | 78,99470448 | 58,8 | 18,0 | 5,885328242 | 0,037442075 | 1700049G17Rik | RIKEN cDNA 1700049G17 gene [Source:MGI Symbol;Acc:MGI:1920680] | 70,08 | 0,0 |
| ENSMUSG00000066829 | 109,0837675 | 89,7 | 27,7 | 5,886324773 | 0,014653362 | Zfp810 | zinc finger protein 810 [Source:MGI Symbol;Acc:MGI:2384563] | 98,64 | 5,2 |
| ENSMUSG00000024687 | 134,9800268 | 155,8 | 16,6 | 5,886346376 | 0,001050231 | Osbp | oxysterol binding protein [Source:MGI Symbol;Acc:MGI:97447] | 94,23 | 2,2 |
| ENSMUSG00000035245 | 52,79653564 | 91,9 | 1,6 | 5,888041274 | 0,003702318 | Eogt | domain-specific O-linked N-acetylglucosamine (GlcNAc) transferase [Source:MGI Symbol;Acc:MGI:214 | 87,42 | 0,6 |
| ENSMUSG00000035172 | 58,05203527 | 82,2 | 61,9 | 5,891733597 | 0,025034564 | Plekhh3 | n homology domain containing, family H (with MyTH4 domain) member 3 [Source:MGI Symbol;Acc:MGI:2 | 89,87 | 14,4 |
| ENSMUSG00000049303 | 60,72626421 | 88,4 | 1,9 | 5,892312688 | 0,008268466 | Syt12 | synaptotagmin XII [Source:MGI Symbol;Acc:MGI:2159601] | 88,02 | 1,3 |
| ENSMUSG00000027298 | 79,23022735 | 87,5 | 3,0 | 5,892724544 | 0,007414379 | Tyro3 | TYRO3 protein tyrosine kinase 3 [Source:MGI Symbol;Acc:MGI:104294] | 76,56 | 1,5 |
| ENSMUSG00000016995 | 17,79374396 | 37,5 | 0,9 | 5,894828994 | 0,024792362 | Matn4 | matrilin 4 [Source:MGI Symbol;Acc:MGI:1328314] | 83,73 | 1,0 |
| ENSMUSG00000018500 | 188,947533 | 287,5 | 77,9 | 5,895417201 | 0,010973739 | Adora2b | adenosine A2b receptor [Source:MGI Symbol;Acc:MGI:99403] | 99,57 | 32,2 |
| ENSMUSG00000018217 | 289,0741749 | 434,2 | 141,9 | 5,895940862 | 0,005293924 | Pmp22 | peripheral myelin protein 22 [Source:MGI Symbol;Acc:MGI:97631] | 89,31 | 27,2 |
| ENSMUSG00000032492 | 88,73268552 | 104,0 | 128,9 | 5,897391412 | 0,022976457 | Pth1r | parathyroid hormone 1 receptor [Source:MGI Symbol;Acc:MGI:97801] | 73,41 | 22,5 |
| ENSMUSG00000032040 | 162,98729 | 171,0 | 28,7 | 5,89977588 | 0,001068526 | Dcps | decapping enzyme, scavenger [Source:MGI Symbol;Acc:MGI:1916555] | 87,55 | 21,0 |
| ENSMUSG00000003469 | 68,82201027 | 127,3 | 8,3 | 5,900709489 | 0,001376439 | Phyhip | phytanoyl-CoA hydroxylase interacting protein [Source:MGI Symbol;Acc:MGI:1860417] | 91,94 | 3,7 |
| ENSMUSG00000042616 | 206,8534273 | 357,1 | 18,5 | 5,901742598 | 9,51995E-05 | Oscp1 | organic solute carrier partner 1 [Source:MGI Symbol;Acc:MGI:1916308] | 80,73 | 2,4 |
| ENSMUSG00000106896 | 55,94972865 | 82,0 | 5,0 | 5,902061341 | 0,009390431 | G630022F23Rik | RIKEN cDNA G630022F23 gene [Source:MGI Symbol;Acc:MGI:3704248] | 50,19 | 1,5 |
| ENSMUSG00000036306 | 32,90307934 | 41,0 | 0,9 | 5,904456655 | 0,027362816 | Lzts1 | leucine zipper, putative tumor suppressor 1 [Source:MGI Symbol;Acc:MGI:2684762] | 53,66 | 0,2 |
| ENSMUSG00000009633 | 68,72899275 | 91,3 | 19,4 | 5,909185122 | 0,020183342 | G0s2 | G0/G1 switch gene 2 [Source:MGI Symbol;Acc:MGI:1316737] | 92,46 | 23,6 |
| ENSMUSG00000024718527 | 26,43718527 | 37,0 | 1,0 | 5,909898667 | 0,035624713 | Plod2 | procollagen lysine, 2-oxoglutarate 5-dioxygenase 2 [Source:MGI Symbol;Acc:MGI:1347007] | 72,50 | 0,5 |
| ENSMUSG00000033444 | 217,5616132 | 294,4 | 4,7 | 5,911665751 | 4,4765E-05 | Specc1l | erm antigen with calponin homology and coiled-coil domains 1-like [Source:MGI Symbol;Acc:MGI:19216 | 98,86 | 1,9 |
| ENSMUSG00000040033 | 100,355712 | 61,5 | 147,9 | 5,913341089 | 0,041952726 | Stat2 | signal transducer and activator of transcription 2 [Source:MGI Symbol;Acc:MGI:103039] | 76,36 | 18,2 |
| ENSMUSG00000041771 | 68,39391287 | 71,4 | 51,8 | 5,915389719 | 0,033216815 | Slc24a4 | e carrier family 24 (sodium/potassium/calcium exchanger), member 4 [Source:MGI Symbol;Acc:MGI:2447 | 68,80 | 14,2 |
| ENSMUSG00000058402 | 131,9342544 | 253,1 | 43,5 | 5,916976227 | 0,004375869 | Zfp420 | zinc finger protein 420 [Source:MGI Symbol;Acc:MGI:2444666] | 99,86 | 2,4 |
| ENSMUSG00000012429 | 135,3405106 | 209,5 | 6,5 | 5,918982982 | 0,001493938 | Mplkip | M-phase specific PLK1 intereacting protein [Source:MGI Symbol;Acc:MGI:1913558] | 96,47 | 3,5 |
| ENSMUSG00000033900 | 65,8486891 | 91,7 | 5,4 | 5,919320475 | 0,016011612 | Map9 | microtubule-associated protein 9 [Source:MGI Symbol;Acc:MGI:2442208] | 71,13 | 0,9 |
| ENSMUSG00000036086 | 41,83877814 | 78,0 | 30,9 | 5,91936484 | 0,012371023 | Zranb3 | zinc finger, RAN-binding domain containing 3 [Source:MGI Symbol;Acc:MGI:1918362] | 76,72 | 0,0 |
| ENSMUSG00000090626 | 101,7868762 | 86,4 | 12,1 | 5,920300472 | 0,024555286 | Tex9 | testis expressed gene 9 [Source:MGI Symbol;Acc:MGI:1201610] | 77,25 | 2,4 |
| ENSMUSG00000041974 | 106,0095278 | 141,2 | 3,9 | 5,925068526 | 0,015100456 | Spidr | scaffolding protein involved in DNA repair [Source:MGI Symbol;Acc:MGI:1924834] | 98,70 | 1,3 |
| ENSMUSG00000031596 | 148,5832768 | 150,4 | 25,0 | 5,925594051 | 0,015087331 | Slc7a2 | e carrier family 7 (cationic amino acid transporter, y+ system), member 2 [Source:MGI Symbol;Acc:MGI:9 | 71,04 | 2,9 |
| ENSMUSG00000066189 | 20,12870981 | 46,3 | 1,0 | 5,927072105 | 0,011265464 | Cacng3 | calcium channel, voltage-dependent, gamma subunit 3 [Source:MGI Symbol;Acc:MGI:1859165] | 69,21 | 0,8 |
| ENSMUSG00000030341 | 174,5322597 | 308,4 | 51,2 | 5,929739449 | 0,008644815 | Tnfrsf1a | tumor necrosis factor receptor superfamily, member 1a [Source:MGI Symbol;Acc:MGI:1314884] | 93,38 | 16,6 |
| ENSMUSG00000021280 | 33,219532 | 71,2 | 2,0 | 5,930398583 | 0,005058806 | Exoc3l4 | exocyst complex component 3-like 4 [Source:MGI Symbol;Acc:MGI:1921363] | 94,42 | 0,7 |
| ENSMUSG00000054894 | 343,2835097 | 596,0 | 96,2 | 5,931093464 | 0,019015056 | Atp5s | TP synthase, H+ transporting, mitochondrial F0 complex, subunit S [Source:MGI Symbol;Acc:MGI:191530 | 100,00 | 17,4 |
| ENSMUSG00000028933 | 80,22772035 | 86,7 | 25,9 | 5,931730552 | 0,026514271 | Xrcc2 | ay repair complementing defective repair in Chinese hamster cells 2 [Source:MGI Symbol;Acc:MGI:19273 | 89,93 | 7,6 |
| ENSMUSG00000020364 | 41,11103193 | 58,7 | 1,8 | 5,931827191 | 0,02435978 | Zfp354a | zinc finger protein 354A [Source:MGI Symbol;Acc:MGI:103172] | 63,76 | 0,6 |
| ENSMUSG00000043372 | 101,1551766 | 115,1 | 73,9 | 5,933299858 | 0,009526192 | Hexim2 | hexamethylene bis-acetamide inducible 2 [Source:MGI Symbol;Acc:MGI:1918309] | 90,60 | 24,5 |
| ENSMUSG00000034958 | 40,92688388 | 37,2 | 0,6 | 5,933653709 | 0,016788041 | Atcay | ataxia, cerebellar, Cayman type [Source:MGI Symbol;Acc:MGI:2448730] | 82,76 | 0,5 |
| ENSMUSG00000051169 | 118,2907729 | 119,8 | 29,6 | 5,934532936 | 0,0055907 | Rpusd3 | RNA pseudouridylate synthase domain containing 3 [Source:MGI Symbol;Acc:MGI:2141440] | 85,62 | 18,7 |
| ENSMUSG00000024201 | 108,1555996 | 124,6 | 80,5 | 5,936788722 | 0,015024331 | Kdm4b | lysine (K)-specific demethylase 4B [Source:MGI Symbol;Acc:MGI:2442355] | 96,16 | 14,2 |
| ENSMUSG00000022382 | 336,6156626 | 537,0 | 52,4 | 5,937932335 | 0,006161154 | Wnt7b | wingless-type MMTV integration site family, member 7B [Source:MGI Symbol;Acc:MGI:98962] | 81,63 | 13,1 |
| ENSMUSG00000048200 | 52,30968871 | 92,3 | 81,8 | 5,940527891 | 0,025353928 | Cracr2b | calcium release activated channel regulator 2B [Source:MGI Symbol;Acc:MGI:2446129] | 66,51 | 16,1 |
| ENSMUSG00000022682 | 227,7790498 | 270,0 | 16,3 | 5,942677331 | 0,001230025 | Rrn3 | RRN3 RNA polymerase I transcription factor homolog (yeast) [Source:MGI Symbol;Acc:MGI:1925255] | 99,43 | 5,4 |
| ENSMUSG00000068205 | 103,1982212 | 143,6 | 25,0 | 5,943252803 | 0,003119232 | Macrod2 | MACRO domain containing 2 [Source:MGI Symbol;Acc:MGI:1920149] | 56,66 | 1,9 |
| ENSMUSG00000047085 | 98,11527636 | 110,4 | 71,8 | 5,943430692 | 0,010069978 | Lrrc4b | leucine rich repeat containing 4B [Source:MGI Symbol;Acc:MGI:3027390] | 90,14 | 20,8 |
| ENSMUSG00000028999 | 183,9388731 | 113,2 | 82,3 | 5,943471732 | 0,008249347 | Rint1 | RAD50 interactor 1 [Source:MGI Symbol;Acc:MGI:1916233] | 70,68 | 11,8 |
| ENSMUSG00000075437 | 398,5926045 | 699,4 | 67,3 | 5,943527876 | 0,002565197 | Gm11681 | predicted gene 11681 [Source:MGI Symbol;Acc:MGI:3650778] | #DIV/0! | #DIV/0! |
| ENSMUSG00000031502 | 106,483887 | 32,1 | 0,6 | 5,944123227 | 0,048578581 | Col4a1 | collagen, type IV, alpha 1 [Source:MGI Symbol;Acc:MGI:88454] | 56,23 | 0,4 |
| ENSMUSG00000027230 | 50,44662589 | 51,4 | 1,2 | 5,945012952 | 0,027440825 | Creb3l1 | cAMP responsive element binding protein 3-like 1 [Source:MGI Symbol;Acc:MGI:1347062] | 82,83 | 0,8 |
| ENSMUSG00000070426 | 83,29390825 | 104,8 | 137,7 | 5,945240581 | 0,016814527 | Rnf121 | ring finger protein 121 [Source:MGI Symbol;Acc:MGI:1922462] | 55,80 | 17,7 |
| ENSMUSG00000032741 | 66,89392091 | 92,4 | 3,0 | 5,946248024 | 0,007221778 | Tpcn1 | two pore channel 1 [Source:MGI Symbol;Acc:MGI:2182472] | 68,74 | 1,2 |
| ENSMUSG00000033805 | 54,79693248 | 92,9 | 30,5 | 5,947250955 | 0,011233016 | Ephx4 | epoxide hydrolase 4 [Source:MGI Symbol;Acc:MGI:2686228] | 74,79 | 5,8 |
| ENSMUSG00000020231 | 186,3003585 | 282,4 | 19,5 | 5,947977251 | 0,002224865 | Dip2a | disco interacting protein 2 homolog A [Source:MGI Symbol;Acc:MGI:2385920] | 97,90 | 2,6 |
| ENSMUSG00000071604 | 193,625143 | 306,7 | 92,9 | 5,948485065 | 0,013848169 | Fam189a2 | family with sequence similarity 189, member A2 [Source:MGI Symbol;Acc:MGI:2685813] | 98,44 | 30,6 |
| ENSMUSG00000010017 | 37,74325755 | 36,6 | 0,9 | 5,948747278 | 0,043566196 | 2410022M11Rik | RIKEN cDNA 2410022M11 gene [Source:MGI Symbol;Acc:MGI:1917005] | 37,69 | 0,7 |
| ENSMUSG00000085105 | 44,96998386 | 70,8 | 6,5 | 5,950797001 | 0,022999178 | Gm12758 | predicted gene 12758 [Source:MGI Symbol;Acc:MGI:3702543] | 54,68 | 1,5 |
| ENSMUSG00000034917 | 29,02667657 | 38,7 | 1,0 | 5,951457171 | 0,024985178 | Tjp3 | tight junction protein 3 [Source:MGI Symbol;Acc:MGI:1351650] | 89,15 | 0,1 |
| ENSMUSG00000033342 | 24,92020479 | 46,6 | 0,9 | 5,952210472 | 0,011734301 | Plppr5 | phospholipid phosphatase related 5 [Source:MGI Symbol;Acc:MGI:1923019] | 73,63 | 0,5 |
| ENSMUSG00000051065 | 69,75575515 | 80,7 | 118,7 | 5,956778681 | 0,016182522 | Mb21d2 | Mab-21 domain containing 2 [Source:MGI Symbol;Acc:MGI:1917028] | 93,24 | 24,0 |
| ENSMUSG00000044433 | 24,31462695 | 46,0 | 1,0 | 5,957046329 | 0,017068678 | Camsap3 | almodulin regulated spectrin-associated protein family, member 3 [Source:MGI Symbol;Acc:MGI:1916947 | 67,40 | 0,2 |
| ENSMUSG00000072915 | 82,9356887 | 73,0 | 18,5 | 5,957753839 | 0,040411089 | Gm12258 | predicted gene 12258 [Source:MGI Symbol;Acc:MGI:3651534] | 86,70 | 4,6 |
| ENSMUSG00000027933 | 76,92572469 | 92,2 | 85,0 | 5,958433622 | 0,015702473 | Ints3 | integrator complex subunit 3 [Source:MGI Symbol;Acc:MGI:2140050] | 92,63 | 17,1 |
| ENSMUSG00000044881 | 64,50197691 | 88,3 | 13,9 | 5,958952893 | 0,016496288 | Coa4 | cytochrome c oxidase assembly factor 4 [Source:MGI Symbol;Acc:MGI:1915435] | 93,12 | 10,9 |
| ENSMUSG00000048920 | 175,264906 | 131,0 | 311,1 | 5,961923637 | 0,013540873 | Fkrp | fukutin related protein [Source:MGI Symbol;Acc:MGI:2447586] | 89,09 | 29,3 |
| ENSMUSG00000074785 | 113,1924285 | 168,5 | 3,6 | 5,964100197 | 0,000652685 | Plxnc1 | plexin C1 [Source:MGI Symbol;Acc:MGI:1890127] | 80,62 | 0,7 |
| ENSMUSG00000041439 | 111,4721166 | 67,5 | 4,0 | 5,964106037 | 0,027404618 | Mfsd6 | major facilitator superfamily domain containing 6 [Source:MGI Symbol;Acc:MGI:1922925] | 69,25 | 0,9 |
| ENSMUSG00000057133 | 69,9832338 | 115,9 | 55,9 | 5,96430304 | 0,00678003 | Chd6 | chromodomain helicase DNA binding protein 6 [Source:MGI Symbol;Acc:MGI:1918639] | 74,13 | 1,8 |
| ENSMUSG00000005897 | 24,89224767 | 49,6 | 1,0 | 5,966008888 | 0,011692489 | Nr2c1 | nuclear receptor subfamily 2, group C, member 1 [Source:MGI Symbol;Acc:MGI:1352465] | 66,17 | 0,4 |
| ENSMUSG00000050605 | 49,63353184 | 45,4 | 0,9 | 5,971300177 | 0,007871312 | Zfp61 | zinc finger protein 61 [Source:MGI Symbol;Acc:MGI:99663] | 95,12 | 1,0 |
| ENSMUSG00000026655 | 283,5940645 | 405,1 | 83,3 | 5,971524581 | 0,010931263 | Fam107b | family with sequence similarity 107, member B [Source:MGI Symbol;Acc:MGI:1913790] | 83,33 | 20,0 |
| ENSMUSG00000046157 | 58,29549204 | 77,3 | 14,0 | 5,971559489 | 0,023434196 | Tmem229b | transmembrane protein 229B [Source:MGI Symbol;Acc:MGI:2444389] | 78,84 | 3,6 |
| ENSMUSG00000019822 | 193,0862365 | 319,7 | 29,4 | 5,972096487 | 0,010490558 | Smpd2 | sphingomyelin phosphodiesterase 2, neutral [Source:MGI Symbol;Acc:MGI:1278330] | 99,96 | 1,6 |
| ENSMUSG00000063146 | 42,56084088 | 77,8 | 1,8 | 5,973104064 | 0,00207752 | Clip2 | CAP-GLY domain containing linker protein 2 [Source:MGI Symbol;Acc:MGI:1313136] | 87,67 | 1,0 |
| ENSMUSG00000034032 | 85,37806667 | 65,9 | 2,4 | 5,97395302 | 0,02430791 | Rpap1 | RNA polymerase II associated protein 1 [Source:MGI Symbol;Acc:MGI:1916175] | 77,81 | 0,0 |
| ENSMUSG00000102697 | 24,222883 | 42,0 | 0,6 | 5,974431771 | 0,009623952 | Pcdhac2 | protocadherin alpha subfamily C, 2 [Source:MGI Symbol;Acc:MGI:1891443] | 59,85 | 0,4 |
| ENSMUSG00000030862 | 42,31423656 | 57,5 | 1,8 | 5,975071027 | 0,014697605 | Cpxm2 | carboxypeptidase X 2 (M14 family) [Source:MGI Symbol;Acc:MGI:1926006] | 72,82 | 0,5 |
| ENSMUSG00000032565 | 111,748078 | 110,6 | 153,7 | 5,975740135 | 0,017886688 | Nudt16 | nudix (nucleoside diphosphate linked moiety X)-type motif 16 [Source:MGI Symbol;Acc:MGI:1922936] | 92,97 | 21,2 |
| ENSMUSG00000056214 | 78,57473053 | 128,7 | 62,3 | 5,976254261 | 0,007374158 | Pard6g | par-6 family cell polarity regulator gamma [Source:MGI Symbol;Acc:MGI:2135606] | 98,02 | 20,8 |
| ENSMUSG00000029276 | 62,09579728 | 72,7 | 2,8 | 5,976813022 | 0,010763771 | Glmn | glomulin, FKBP associated protein [Source:MGI Symbol;Acc:MGI:2141180] | 84,66 | 0,8 |
| ENSMUSG00000071281 | 56,76862361 | 75,4 | 2,8 | 5,977356707 | 0,007568148 | Zfp65 | zinc finger protein 65 [Source:MGI Symbol;Acc:MGI:107769] | 88,93 | 0,6 |
| ENSMUSG00000024511 | 54,90578764 | 92,0 | 64,7 | 5,980236718 | 0,012296148 | Rab27b | RAB27B, member RAS oncogene family [Source:MGI Symbol;Acc:MGI:1931295] | 55,96 | 8,6 |
| ENSMUSG00000031712 | 61,61207609 | 70,4 | 22,0 | 5,983916223 | 0,038886269 | Il15 | interleukin 15 [Source:MGI Symbol;Acc:MGI:103014] | 80,71 | 9,3 |
| ENSMUSG00000024781 | 132,1855442 | 92,3 | 7,9 | 5,983980916 | 0,015788078 | Lipa | lysosomal acid lipase A [Source:MGI Symbol;Acc:MGI:96789] | 88,73 | 3,3 |
| ENSMUSG00000031169 | 89,35324106 | 138,1 | 19,4 | 5,985594794 | 0,00447091 | Porcn | porcupine homolog (Drosophila) [Source:MGI Symbol;Acc:MGI:1890212] | 99,26 | 5,4 |
| ENSMUSG00000055862 | 97,14663321 | 129,6 | 37,9 | 5,985707808 | 0,004975445 | Izumo4 | IZUMO family member 4 [Source:MGI Symbol;Acc:MGI:1918814] | 98,69 | 17,7 |
| ENSMUSG00000030004 | 66,42463582 | 98,0 | 34,9 | 5,985783885 | 0,013401709 | Nat8 | N-acetyltransferase 8 (GCN5-related, putative) [Source:MGI Symbol;Acc:MGI:1915646] | 81,09 | 17,7 |
| ENSMUSG00000100039 | 80,13367674 | 112,3 | 49,9 | 5,988434518 | 0,010134191 | Gm28959 | predicted gene 28959 [Source:MGI Symbol;Acc:MGI:5579665] | 100,00 | 33,3 |
| ENSMUSG00000021048 | 272,0987093 | 339,4 | 77,8 | 5,98865462 | 0,002527128 | Mthfd1 | enase (NADP+ dependent), methenyltetrahydrofolate cyclohydrolase, formyltetrahydrofolate synthase [Sou | 98,77 | 8,5 |
| ENSMUSG00000051950 | 77,88776434 | 80,8 | 99,3 | 5,990010955 | 0,048276563 | B3glct | beta-3-glucosyltransferase [Source:MGI Symbol;Acc:MGI:2685903] | 87,60 | 22,4 |
| ENSMUSG00000093436 | 74,26003197 | 105,6 | 4,2 | 5,992376152 | 0,005659757 | Gm20646 | predicted gene 20646 [Source:MGI Symbol;Acc:MGI:5313093] | 99,42 | 4,7 |
| ENSMUSG00000050064 | 22,45415813 | 42,8 | 1,0 | 5,992890585 | 0,017985246 | Zfp697 | zinc finger protein 697 [Source:MGI Symbol;Acc:MGI:2139736] | 26,53 | 0,3 |
| ENSMUSG00000039914 | 152,1528264 | 255,4 | 60,5 | 5,994826421 | 0,015080081 | Coq10a | coenzyme Q10A [Source:MGI Symbol;Acc:MGI:2684847] | 88,48 | 18,8 |
| ENSMUSG00000043670 | 20,30860103 | 46,3 | 1,0 | 5,994929846 | 0,011399147 | Diras1 | DIRAS family, GTP-binding RAS-like 1 [Source:MGI Symbol;Acc:MGI:2183442] | 85,11 | 0,6 |
| ENSMUSG00000046753 | 80,90009183 | 105,3 | 32,3 | 5,999493434 | 0,016722094 | Ccdc66 | coiled-coil domain containing 66 [Source:MGI Symbol;Acc:MGI:2443639] | 99,65 | 4,7 |
| ENSMUSG00000051379 | 39,84684441 | 45,5 | 0,6 | 6,00354659 | 0,010931263 | Flrt3 | fibronectin leucine rich transmembrane protein 3 [Source:MGI Symbol;Acc:MGI:1918686] | 73,61 | 0,6 |
| ENSMUSG00000100396 | 61,3244417 | 84,6 | 33,3 | 6,005598187 | 0,03857125 | Gm29367 | predicted gene 29367 [Source:MGI Symbol;Acc:MGI:5580073] | #DIV/0! | #DIV/0! |
| ENSMUSG00000032902 | 282,8501061 | 397,9 | 268,6 | 6,007953605 | 0,005393236 | Slc16a1 | olute carrier family 16 (monocarboxylic acid transporters), member 1 [Source:MGI Symbol;Acc:MGI:10601 | 72,53 | 23,5 |
| ENSMUSG00000039676 | 106,1684052 | 105,6 | 39,3 | 6,008282281 | 0,030961686 | Capsl | calcyphosine-like [Source:MGI Symbol;Acc:MGI:1922818] | 99,86 | 29,5 |
| ENSMUSG00000020100 | 55,68532115 | 71,0 | 8,0 | 6,009053944 | 0,022313912 | Slc29a3 | solute carrier family 29 (nucleoside transporters), member 3 [Source:MGI Symbol;Acc:MGI:1918529] | 82,80 | 1,5 |
| ENSMUSG00000022025 | 32,29144892 | 36,8 | 0,6 | 6,010917514 | 0,025713947 | Lect1 | leukocyte cell derived chemotaxin 1 [Source:MGI Symbol;Acc:MGI:1341171] | 94,96 | 1,7 |

| | | | | | | | | | |
|---|---|---|---|---|---|---|---|---|---|
| ENSMUSG00000031728 | 77,27393548 | 90,6 | 11,1 | 6,011268268 | 0,014140921 | Zfp821 | zinc finger protein 821 [Source:MGI Symbol;Acc:MGI:1923121] | 94,70 | 6,9 |
| ENSMUSG00000025626 | 126,3128466 | 159,3 | 4,2 | 6,011293325 | 0,000180309 | Phf6 | PHD finger protein 6 [Source:MGI Symbol;Acc:MGI:1918248] | 61,66 | 0,4 |
| ENSMUSG00000037519 | 162,1661432 | 169,4 | 61,9 | 6,014355047 | 0,002295782 | Ppfia1 | osphatase, receptor type, f polypeptide (PTPRF), interacting protein (liprin), alpha 1 [Source:MGI Symbol;/ | 90,03 | 13,9 |
| ENSMUSG00000026211 | 29,66270588 | 39,9 | 0,9 | 6,014797719 | 0,022742322 | Obsl1 | obscurin-like 1 [Source:MGI Symbol;Acc:MGI:2138628] | 48,71 | 0,0 |
| ENSMUSG00000054196 | 58,58336158 | 105,6 | 65,9 | 6,017321586 | 0,021190548 | Cthrc1 | collagen triple helix repeat containing 1 [Source:MGI Symbol;Acc:MGI:1915838] | 99,05 | 31,0 |
| ENSMUSG00000030515 | 221,0202424 | 100,0 | 189,6 | 6,017515629 | 0,013375274 | Tarsl2 | threonyl-tRNA synthetase-like 2 [Source:MGI Symbol;Acc:MGI:2444486] | 98,68 | 27,0 |
| ENSMUSG00000101892 | 190,6344214 | 306,6 | 35,2 | 6,017580756 | 0,006448156 | 9130401M01Rik | RIKEN cDNA 9130401M01 gene [Source:MGI Symbol;Acc:MGI:1923008] | 98,99 | 30,9 |
| ENSMUSG00000021700 | 59,83367707 | 98,8 | 57,9 | 6,018525613 | 0,014415267 | Rab3c | RAB3C, member RAS oncogene family [Source:MGI Symbol;Acc:MGI:1914545] | 66,35 | 2,5 |
| ENSMUSG00000043065 | 91,01513742 | 81,0 | 24,0 | 6,018719511 | 0,023604417 | Spice1 | spindle and centriole associated protein 1 [Source:MGI Symbol;Acc:MGI:1196252] | 68,04 | 3,0 |
| ENSMUSG00000033446 | 29,92459937 | 42,4 | 0,6 | 6,019862488 | 0,018872392 | Lpar6 | lysophosphatidic acid receptor 6 [Source:MGI Symbol;Acc:MGI:1914418] | 36,48 | 0,5 |
| ENSMUSG00000035671 | 66,16299131 | 104,2 | 70,2 | 6,020063219 | 0,010157004 | Zswim4 | zinc finger SWIM-type containing 4 [Source:MGI Symbol;Acc:MGI:2443726] | 96,52 | 21,7 |
| ENSMUSG00000030780 | 110,2859821 | 116,8 | 118,0 | 6,023781957 | 0,020332104 | BC017158 | cDNA sequence BC017158 [Source:MGI Symbol;Acc:MGI:2384572] | 89,27 | 27,7 |
| ENSMUSG00000049551 | 74,86028087 | 88,0 | 83,8 | 6,025832554 | 0,022794374 | Fzd9 | frizzled class receptor 9 [Source:MGI Symbol;Acc:MGI:1313278] | 99,07 | 26,3 |
| ENSMUSG00000043155 | 33,55970804 | 42,9 | 1,0 | 6,027042316 | 0,043026894 | Hpdl | 4-hydroxyphenylpyruvate dioxygenase-like [Source:MGI Symbol;Acc:MGI:2444646] | 78,60 | 1,0 |
| ENSMUSG00000032878 | 593,3703502 | 931,1 | 228,9 | 6,027160152 | 0,003823378 | Ccdc85a | coiled-coil domain containing 85A [Source:MGI Symbol;Acc:MGI:2445069] | 98,03 | 25,1 |
| ENSMUSG00000061740 | 423,2892587 | 715,9 | 37,5 | 6,027371583 | 0,018984452 | Cyp2d22 | cytochrome P450, family 2, subfamily d, polypeptide 22 [Source:MGI Symbol;Acc:MGI:1929474] | 99,70 | 16,0 |
| ENSMUSG00000062101 | 30,81585691 | 75,9 | 1,8 | 6,030666231 | 0,0038164 | Zfp119b | zinc finger protein 119b [Source:MGI Symbol;Acc:MGI:2385323] | 82,81 | 1,1 |
| ENSMUSG00000021943 | 54,12452521 | 88,0 | 8,3 | 6,037273809 | 0,014480481 | Gdf10 | growth differentiation factor 10 [Source:MGI Symbol;Acc:MGI:95684] | 96,88 | 3,4 |
| ENSMUSG00000023074 | 307,7300938 | 428,4 | 14,8 | 6,037372233 | 0,006265298 | Mospd1 | motile sperm domain containing 1 [Source:MGI Symbol;Acc:MGI:1917630] | 99,23 | 2,4 |
| ENSMUSG00000039294 | 38,99653483 | 71,7 | 1,8 | 6,043713796 | 0,005383997 | BC017643 | cDNA sequence BC017643 [Source:MGI Symbol;Acc:MGI:2384959] | 91,04 | 0,3 |
| ENSMUSG00000042275 | 59,76809242 | 44,3 | 0,9 | 6,044982138 | 0,016335324 | Pelo | pelota homolog (Drosophila) [Source:MGI Symbol;Acc:MGI:2145154] | 87,83 | 1,5 |
| ENSMUSG00000024165 | 253,4878304 | 199,5 | 24,0 | 6,047339256 | 0,000467449 | Hn1l | hematological and neurological expressed 1-like [Source:MGI Symbol;Acc:MGI:1196260] | 99,39 | 7,3 |
| ENSMUSG00000028439 | 85,00469936 | 126,5 | 9,1 | 6,04998981 | 0,006694352 | Fam219a | family with sequence similarity 219, member A [Source:MGI Symbol;Acc:MGI:1919151] | 89,41 | 3,7 |
| ENSMUSG00000022840 | 46,47070855 | 42,1 | 1,0 | 6,050866487 | 0,022802016 | Adcy5 | adenylate cyclase 5 [Source:MGI Symbol;Acc:MGI:99673] | 49,10 | 0,2 |
| ENSMUSG00000001948 | 84,65209116 | 114,6 | 22,2 | 6,050915051 | 0,006499377 | Spa17 | sperm autoantigenic protein 17 [Source:MGI Symbol;Acc:MGI:1333778] | 100,00 | 25,3 |
| ENSMUSG00000045312 | 144,8607229 | 193,0 | 6,9 | 6,054252148 | 0,007718687 | Lhfpl2 | lipoma HMGIC fusion partner-like 2 [Source:MGI Symbol;Acc:MGI:2145236] | 85,05 | 1,5 |
| ENSMUSG00000029307 | 79,67114156 | 143,5 | 34,5 | 6,055665095 | 0,009665406 | Dmp1 | dentin matrix protein 1 [Source:MGI Symbol;Acc:MGI:94910] | 96,27 | 6,3 |
| ENSMUSG00000038831 | 147,458232 | 271,2 | 4,3 | 6,056334676 | 0,001972796 | Ralgps1 | Ral GEF with PH domain and SH3 binding motif 1 [Source:MGI Symbol;Acc:MGI:1922008] | 97,69 | 1,1 |
| ENSMUSG00000032235 | 22,39933792 | 40,6 | 0,6 | 6,056606419 | 0,013631482 | Ice2 | interactor of little elongation complex ELL subunit 2 [Source:MGI Symbol;Acc:MGI:2135947] | 64,73 | 0,5 |
| ENSMUSG00000038372 | 115,9910518 | 109,7 | 37,5 | 6,057010822 | 0,020194747 | Gmds | GDP-mannose 4, 6-dehydratase [Source:MGI Symbol;Acc:MGI:1891112] | 91,45 | 22,6 |
| ENSMUSG00000021314 | 59,75477716 | 122,1 | 1,9 | 6,058204331 | 0,003911784 | Amph | amphiphysin [Source:MGI Symbol;Acc:MGI:103574] | 98,84 | 1,4 |
| ENSMUSG00000027676 | 61,3763417 | 53,7 | 2,0 | 6,06035572 | 0,024685482 | Ccdc39 | coiled-coil domain containing 39 [Source:MGI Symbol;Acc:MGI:1289263] | 76,38 | 0,7 |
| ENSMUSG00000035437 | 262,2688794 | 341,3 | 5,6 | 6,060401377 | 1,98735E-05 | Rabgap1 | RAB GTPase activating protein 1 [Source:MGI Symbol;Acc:MGI:2385139] | 82,40 | 0,6 |
| ENSMUSG00000033624 | 73,67360841 | 110,5 | 5,5 | 6,061369381 | 0,006314703 | Pdpr | pyruvate dehydrogenase phosphatase regulatory subunit [Source:MGI Symbol;Acc:MGI:2442188] | 87,08 | 0,5 |
| ENSMUSG00000041708 | 199,4024522 | 224,9 | 4,3 | 6,062305831 | 0,010932261 | Mpped1 | metallophosphoesterase domain containing 1 [Source:MGI Symbol;Acc:MGI:106316] | 79,28 | 1,0 |
| ENSMUSG00000015652 | 164,6072052 | 158,7 | 48,4 | 6,062433062 | 0,019207141 | Steap1 | six transmembrane epithelial antigen of the prostate 1 [Source:MGI Symbol;Acc:MGI:1917608] | 87,34 | 27,3 |
| ENSMUSG00000074166 | 61,0898125 | 82,0 | 33,9 | 6,062495258 | 0,01696342 | AW146154 | expressed sequence AW146154 [Source:MGI Symbol;Acc:MGI:2142212] | 76,87 | 0,0 |
| ENSMUSG00000026694 | 49,2024193 | 77,2 | 2,0 | 6,064104746 | 0,005773343 | Mettl13 | methyltransferase like 13 [Source:MGI Symbol;Acc:MGI:1918699] | 87,23 | 0,5 |
| ENSMUSG00000070532 | 105,5739595 | 135,8 | 29,7 | 6,065685012 | 0,009541204 | Ccdc190 | coiled-coil domain containing 190 [Source:MGI Symbol;Acc:MGI:1925715] | 93,37 | 19,3 |
| ENSMUSG00000038515 | 57,26048373 | 100,2 | 28,7 | 6,066599925 | 0,015668098 | Grtp1 | GH regulated TBC protein 1 [Source:MGI Symbol;Acc:MGI:1914040] | 99,75 | 22,6 |
| ENSMUSG00000040570 | 74,90737986 | 80,0 | 3,0 | 6,067287836 | 0,016951686 | Rundc3b | RUN domain containing 3B [Source:MGI Symbol;Acc:MGI:2685286] | 89,33 | 0,3 |
| ENSMUSG00000035473 | 104,8587175 | 171,0 | 85,9 | 6,069672455 | 0,006701963 | Galm | galactose mutarotase [Source:MGI Symbol;Acc:MGI:2442420] | 90,19 | 22,8 |
| ENSMUSG00000027895 | 60,55669268 | 99,8 | 21,0 | 6,074301522 | 0,024220928 | Kcnc4 | otassium voltage gated channel, Shaw-related subfamily, member 4 [Source:MGI Symbol;Acc:MGI:9667( | 85,87 | 6,6 |
| ENSMUSG00000030720 | 76,74524698 | 97,6 | 33,3 | 6,077701576 | 0,019111794 | Cln3 | d lipofuscinosis, neuronal 3, juvenile (Batten, Spielmeyer-Vogt disease) [Source:MGI Symbol;Acc:MGI:107 | 60,95 | 10,2 |
| ENSMUSG00000029094 | 112,2254261 | 128,9 | 49,6 | 6,078881696 | 0,011516153 | Afap1 | actin filament associated protein 1 [Source:MGI Symbol;Acc:MGI:1917542] | 84,62 | 4,5 |
| ENSMUSG00000020799 | 56,93749237 | 77,7 | 3,7 | 6,079586305 | 0,011862764 | Tekt1 | tektin 1 [Source:MGI Symbol;Acc:MGI:1333819] | 87,94 | 2,1 |
| ENSMUSG00000034573 | 61,598617 | 143,0 | 2,5 | 6,082254323 | 0,000146715 | Ptpn13 | protein tyrosine phosphatase, non-receptor type 13 [Source:MGI Symbol;Acc:MGI:103293] | 94,95 | 0,5 |
| ENSMUSG00000039672 | 279,8164358 | 319,6 | 129,3 | 6,082744698 | 0,013100799 | Kcne2 | potassium voltage-gated channel, Isk-related subfamily, gene 2 [Source:MGI Symbol;Acc:MGI:1891123] | 97,34 | 31,9 |
| ENSMUSG00000055717 | 126,9560549 | 213,4 | 89,6 | 6,084180399 | 0,002803331 | Slain1 | SLAIN motif family, member 1 [Source:MGI Symbol;Acc:MGI:2145578] | 88,37 | 24,1 |
| ENSMUSG00000015016 | 59,46555298 | 84,4 | 23,0 | 6,095056248 | 0,016824826 | Acsf3 | acyl-CoA synthetase family member 3 [Source:MGI Symbol;Acc:MGI:2182591] | 97,61 | 7,4 |
| ENSMUSG00000063018 | 117,3486409 | 151,9 | 39,9 | 6,095575709 | 0,004925718 | 2010204K13Rik | RIKEN cDNA 2010204K13 gene [Source:MGI Symbol;Acc:MGI:1922859] | #DIV/0! | #DIV/0! |
| ENSMUSG00000060395 | 46,12184221 | 46,2 | 1,0 | 6,098571498 | 0,014623166 | Hyi | hydroxypyruvate isomerase (putative) [Source:MGI Symbol;Acc:MGI:1915430] | 90,99 | 0,1 |
| ENSMUSG00000050786 | 67,46092083 | 87,5 | 106,3 | 6,098904893 | 0,026432656 | Ccdc126 | coiled-coil domain containing 126 [Source:MGI Symbol;Acc:MGI:1889376] | 96,25 | 25,7 |
| ENSMUSG00000002688 | 304,6342714 | 454,9 | 204,9 | 6,100003569 | 0,017208103 | Prkd1 | protein kinase D1 [Source:MGI Symbol;Acc:MGI:99879] | 99,46 | 32,0 |
| ENSMUSG00000026718 | 310,1822641 | 453,9 | 54,8 | 6,1002677 | 0,005264923 | Stam | gnal transducing adaptor molecule (SH3 domain and ITAM motif) 1 [Source:MGI Symbol;Acc:MGI:132901 | 62,63 | 6,9 |
| ENSMUSG00000020407 | 82,15105652 | 114,4 | 12,9 | 6,101004593 | 0,014137454 | Upp1 | uridine phosphorylase 1 [Source:MGI Symbol;Acc:MGI:1097668] | 93,02 | 6,3 |
| ENSMUSG00000059142 | 64,29433081 | 97,4 | 15,7 | 6,103590573 | 0,017626395 | Zfp945 | zinc finger protein 945 [Source:MGI Symbol;Acc:MGI:2445132] | 67,69 | 2,3 |
| ENSMUSG00000026333 | 75,22896234 | 111,6 | 74,8 | 6,104525911 | 0,01192108 | Gin1 | gypsy retrotransposon integrase 1 [Source:MGI Symbol;Acc:MGI:2182036] | 67,59 | 7,7 |
| ENSMUSG00000030055 | 104,798326 | 133,1 | 104,8 | 6,10648221 | 0,007666043 | Rab43 | RAB43, member RAS oncogene family [Source:MGI Symbol;Acc:MGI:1917084] | 84,45 | 7,6 |
| ENSMUSG00000030852 | 49,80526655 | 70,8 | 17,6 | 6,107988578 | 0,022587127 | Tacc2 | transforming, acidic coiled-coil containing protein 2 [Source:MGI Symbol;Acc:MGI:1928899] | 33,39 | 1,7 |
| ENSMUSG00000036915 | 73,61677866 | 100,7 | 61,0 | 6,108127527 | 0,019015056 | Kirrel2 | kin of IRRE like 2 (Drosophila) [Source:MGI Symbol;Acc:MGI:2442334] | 94,11 | 21,4 |
| ENSMUSG00000029173 | 104,4515807 | 109,8 | 49,9 | 6,108827944 | 0,019015056 | Sepsecs | ep (O-phosphoserine) tRNA:Sec (selenocysteine) tRNA synthase [Source:MGI Symbol;Acc:MGI:109879' | 91,98 | 5,4 |
| ENSMUSG00000025537 | 468,3290363 | 760,6 | 21,8 | 6,109328353 | 2,55052E-05 | Phkg1 | phosphorylase kinase gamma 1 [Source:MGI Symbol;Acc:MGI:97579] | 87,53 | 7,1 |
| ENSMUSG00000037492 | 37,96692263 | 72,2 | 1,8 | 6,110766541 | 0,008615342 | Zmat4 | zinc finger, matrin type 4 [Source:MGI Symbol;Acc:MGI:2443497] | 82,57 | 0,5 |
| ENSMUSG00000029287 | 56,85184086 | 66,0 | 2,0 | 6,112204582 | 0,010179082 | Tgfbr3 | transforming growth factor, beta receptor III [Source:MGI Symbol;Acc:MGI:104637] | 77,34 | 0,4 |
| ENSMUSG00000087132 | 28,67496613 | 44,8 | 1,0 | 6,112684222 | 0,018390394 | A930001C03Rik | RIKEN cDNA A930001C03 gene [Source:MGI Symbol;Acc:MGI:2441838] | #DIV/0! | #DIV/0! |
| ENSMUSG00000028411 | 221,1361916 | 290,3 | 123,5 | 6,114030359 | 0,008993454 | Aptx | aprataxin [Source:MGI Symbol;Acc:MGI:1913658] | 81,76 | 21,8 |
| ENSMUSG00000024925 | 171,5537093 | 232,6 | 12,9 | 6,114945008 | 2,05142E-05 | Rnaseh2c | ribonuclease H2, subunit C [Source:MGI Symbol;Acc:MGI:1915459] | 99,10 | 12,8 |
| ENSMUSG00000041809 | 22,52081262 | 40,1 | 0,6 | 6,115310402 | 0,023659521 | Efhc1 | EF-hand domain (C-terminal) containing 1 [Source:MGI Symbol;Acc:MGI:1919127] | 91,58 | 1,1 |
| ENSMUSG00000039826 | 168,4242516 | 274,7 | 20,9 | 6,117571778 | 0,006792606 | Trub2 | TruB pseudouridine (psi) synthase homolog 2 (E. coli) [Source:MGI Symbol;Acc:MGI:2442186] | 82,98 | 6,1 |
| ENSMUSG00000028088 | 114,968655 | 157,6 | 217,3 | 6,118107274 | 0,014604568 | Fmo5 | flavin containing monooxygenase 5 [Source:MGI Symbol;Acc:MGI:1310004] | 76,76 | 29,4 |
| ENSMUSG00000031373 | 45,17785863 | 56,2 | 1,0 | 6,119258683 | 0,006806315 | Car5b | carbonic anhydrase 5b, mitochondrial [Source:MGI Symbol;Acc:MGI:1926249] | 78,81 | 0,0 |
| ENSMUSG00000051048 | 103,4419448 | 116,9 | 144,6 | 6,120108248 | 0,01553199 | P4ha3 | oline, 2-oxoglutarate 4-dioxygenase (proline 4-hydroxylase), alpha polypeptide III [Source:MGI Symbol;Acc | 99,36 | 32,1 |
| ENSMUSG00000071324 | 21,25251844 | 48,3 | 1,0 | 6,121326804 | 0,014210371 | Armc2 | armadillo repeat containing 2 [Source:MGI Symbol;Acc:MGI:1916449] | 50,73 | 0,3 |
| ENSMUSG00000058571 | 56,81141816 | 110,6 | 8,0 | 6,124390764 | 0,003898346 | Gpc6 | glypican 6 [Source:MGI Symbol;Acc:MGI:1346322] | 87,57 | 0,3 |
| ENSMUSG00000025730 | 113,774384 | 158,7 | 82,3 | 6,125792743 | 0,009806038 | Rab40c | Rab40C, member RAS oncogene family [Source:MGI Symbol;Acc:MGI:2183454] | 94,65 | 27,0 |
| ENSMUSG00000021696 | 260,0322802 | 208,9 | 26,6 | 6,126175635 | 0,00184966 | Elovl7 | LOVL family member 7, elongation of long chain fatty acids (yeast) [Source:MGI Symbol;Acc:MGI:192180 | 76,68 | 4,6 |
| ENSMUSG00000039616 | 24,62420041 | 52,5 | 0,6 | 6,127205827 | 0,007365245 | Mocos | molybdenum cofactor sulfurase [Source:MGI Symbol;Acc:MGI:1915841] | 87,16 | 0,5 |
| ENSMUSG00000021340 | 351,584957 | 386,0 | 114,2 | 6,127293137 | 0,00346243 | Gpld1 | glycosylphosphatidylinositol specific phospholipase D1 [Source:MGI Symbol;Acc:MGI:106604] | 98,70 | 28,9 |
| ENSMUSG00000054263 | 121,0235761 | 93,4 | 42,5 | 6,127602796 | 0,0252828919 | Lifr | leukemia inhibitory factor receptor [Source:MGI Symbol;Acc:MGI:96788] | 63,08 | 2,6 |
| ENSMUSG00000035615 | 139,0836045 | 279,8 | 18,2 | 6,127933406 | 0,001851868 | Frmpd1 | FERM and PDZ domain containing 1 [Source:MGI Symbol;Acc:MGI:2446274] | 90,40 | 4,8 |
| ENSMUSG00000036745 | 90,93113467 | 158,7 | 19,4 | 6,128533721 | 0,003504465 | Ttll7 | tubulin tyrosine ligase-like family, member 7 [Source:MGI Symbol;Acc:MGI:1918142] | 89,57 | 1,6 |
| ENSMUSG00000024789 | 330,8457761 | 542,2 | 9,8 | 6,128699089 | 0,000928068 | Jak2 | Janus kinase 2 [Source:MGI Symbol;Acc:MGI:96629] | 99,62 | 2,1 |
| ENSMUSG00000101856 | 64,13697507 | 108,5 | 21,3 | 6,128965784 | 0,015722844 | 1700096K18Rik | RIKEN cDNA 1700096K18 gene [Source:MGI Symbol;Acc:MGI:1920821] | #DIV/0! | #DIV/0! |
| ENSMUSG00000033594 | 120,8060071 | 94,7 | 48,1 | 6,129441212 | 0,020897978 | Spata2l | spermatogenesis associated 2-like [Source:MGI Symbol;Acc:MGI:1926029] | 98,35 | 22,7 |
| ENSMUSG00000024604 | 91,73840841 | 145,0 | 3,7 | 6,131762472 | 7,99012E-05 | Rbm22 | RNA binding motif protein 22 [Source:MGI Symbol;Acc:MGI:1914060] | 95,54 | 3,3 |
| ENSMUSG00000048058 | 40,06903377 | 57,6 | 0,6 | 6,132839277 | 0,011233016 | Ldlrad3 | low density lipoprotein receptor class A domain containing 3 [Source:MGI Symbol;Acc:MGI:2138856] | 84,23 | 0,6 |
| ENSMUSG00000004383 | 59,83199067 | 69,2 | 2,0 | 6,134044546 | 0,009078025 | Large | like-glycosyltransferase [Source:MGI Symbol;Acc:MGI:1342270] | 78,04 | 0,9 |
| ENSMUSG00000078853 | 99,15153009 | 41,0 | 0,9 | 6,134455013 | 0,029679181 | Igtp | interferon gamma induced GTPase [Source:MGI Symbol;Acc:MGI:107729] | 87,57 | 1,2 |
| ENSMUSG00000084925 | 60,0051741 | 93,1 | 35,9 | 6,134780947 | 0,013798776 | 1810062O18Rik | RIKEN cDNA 1810062O18 gene [Source:MGI Symbol;Acc:MGI:1922852] | 73,07 | 10,8 |
| ENSMUSG00000055334 | 83,29234837 | 150,6 | 13,9 | 6,136736995 | 0,001995548 | Snupn | snurportin 1 [Source:MGI Symbol;Acc:MGI:1913319] | 99,38 | 4,2 |
| ENSMUSG00000031327 | 91,90815093 | 141,2 | 200,3 | 6,137759832 | 0,013152974 | Chic1 | cysteine-rich hydrophobic domain 1 [Source:MGI Symbol;Acc:MGI:1344694] | 84,09 | 18,5 |
| ENSMUSG00000034209 | 100,8497547 | 101,8 | 40,7 | 6,138170744 | 0,012419384 | Rasl10a | RAS-like, family 10, member A [Source:MGI Symbol;Acc:MGI:1922918] | 100,00 | 26,9 |
| ENSMUSG00000055639 | 20,21650143 | 55,4 | 1,0 | 6,139050211 | 0,010230357 | Dach1 | dachshund 1 (Drosophila) [Source:MGI Symbol;Acc:MGI:1277991] | 66,19 | 0,0 |
| ENSMUSG00000026443 | 98,88342297 | 124,8 | 53,9 | 6,139400618 | 0,01025334 | Lrrn2 | leucine rich repeat protein 2, neuronal [Source:MGI Symbol;Acc:MGI:106037] | 95,94 | 5,9 |
| ENSMUSG00000045466 | 79,10363355 | 91,9 | 86,8 | 6,139933267 | 0,015968705 | Zfp956 | zinc finger protein 956 [Source:MGI Symbol;Acc:MGI:2141515] | 98,59 | 22,3 |
| ENSMUSG00000018659 | 139,7702957 | 106,9 | 24,0 | 6,143300635 | 0,016949068 | Pnpo | pyridoxine 5'-phosphate oxidase [Source:MGI Symbol;Acc:MGI:2144151] | 92,10 | 15,3 |
| ENSMUSG00000032101 | 143,4829212 | 242,0 | 8,0 | 6,143912617 | 0,003282085 | Ddx25 | DEAD (Asp-Glu-Ala-Asp) box polypeptide 25 [Source:MGI Symbol;Acc:MGI:1353582] | 34,29 | 2,3 |
| ENSMUSG00000055003 | 128,7731166 | 168,5 | 83,2 | 6,144346733 | 0,00513118 | Lrtm2 | leucine-rich repeats and transmembrane domains 2 [Source:MGI Symbol;Acc:MGI:2141485] | 83,38 | 20,5 |
| ENSMUSG00000025907 | 311,168496 | 363,6 | 80,8 | 6,146743242 | 0,004876373 | Rb1cc1 | RB1-inducible coiled-coil 1 [Source:MGI Symbol;Acc:MGI:1341850] | 84,40 | 4,0 |
| ENSMUSG00000041907 | 18,2249431 | 44,5 | 1,0 | 6,147080216 | 0,019405117 | Gpr45 | G protein-coupled receptor 45 [Source:MGI Symbol;Acc:MGI:2135882] | 9,11 | 0,0 |
| ENSMUSG00000020182 | 21,07416459 | 43,8 | 0,6 | 6,14760286 | 0,011804736 | Ddc | dopa decarboxylase [Source:MGI Symbol;Acc:MGI:94876] | 74,71 | 0,8 |
| ENSMUSG00000062488 | 67,65035353 | 109,4 | 66,6 | 6,147759979 | 0,015495909 | Ifit3b | interferon-induced protein with tetratricopeptide repeats 3B [Source:MGI Symbol;Acc:MGI:3698419] | 80,37 | 22,9 |
| ENSMUSG00000035681 | 40,60205359 | 42,9 | 1,0 | 6,147987759 | 0,03179463 | Kcnc2 | otassium voltage gated channel, Shaw-related subfamily, member 2 [Source:MGI Symbol;Acc:MGI:96668 | 62,72 | 0,2 |
| ENSMUSG00000032763 | 81,29012085 | 135,3 | 35,9 | 6,148572875 | 0,004375869 | Ilvbl | ilvB (bacterial acetolactate synthase)-like [Source:MGI Symbol;Acc:MGI:1351911] | 98,49 | 9,8 |
| ENSMUSG00000036777 | 111,4916842 | 71,2 | 56,9 | 6,149495133 | 0,033172832 | Anln | anillin, actin binding protein [Source:MGI Symbol;Acc:MGI:1920174] | 68,06 | 2,5 |
| ENSMUSG00000020988 | 190,0264338 | 297,3 | 168,6 | 6,150948289 | 0,011657155 | L2hgdh | L-2-hydroxyglutarate dehydrogenase [Source:MGI Symbol;Acc:MGI:2384968] | 97,56 | 28,7 |
| ENSMUSG00000019774 | 50,50062619 | 65,5 | 1,8 | 6,151793924 | 0,023324993 | Mtrf1l | mitochondrial translational release factor 1-like [Source:MGI Symbol;Acc:MGI:1918830] | 75,12 | 0,9 |
| ENSMUSG00000037262 | 75,65885067 | 112,1 | 117,4 | 6,152068506 | 0,017418341 | Kin | antigenic determinant of rec-A protein [Source:MGI Symbol;Acc:MGI:96676] | 92,57 | 30,0 |
| ENSMUSG00000079235 | 152,741081 | 217,3 | 10,8 | 6,155626035 | 0,007837213 | Ccdc13 | coiled-coil domain containing 13 [Source:MGI Symbol;Acc:MGI:1920144] | 94,63 | 1,9 |
| ENSMUSG00000063873 | 95,3374576 | 107,1 | 1,9 | 6,156779611 | 0,011238312 | Slc24a3 | e carrier family 24 (sodium/potassium/calcium exchanger), member 3 [Source:MGI Symbol;Acc:MGI:2137 | 93,31 | 1,1 |
| ENSMUSG00000070498 | 98,69509742 | 116,1 | 131,3 | 6,157513089 | 0,015540783 | Tmem132b | transmembrane protein 132B [Source:MGI Symbol;Acc:MGI:3609245] | 84,87 | 9,1 |
| ENSMUSG00000038510 | 204,3055041 | 171,3 | 47,1 | 6,157909944 | 0,003086144 | Rpf2 | ribosome production factor 2 homolog (S. cerevisiae) [Source:MGI Symbol;Acc:MGI:1914489] | 89,75 | 19,4 |
| ENSMUSG00000040811 | 70,95351198 | 53,2 | 0,6 | 6,158072792 | 0,009843736 | Eml2 | echinoderm microtubule associated protein like 2 [Source:MGI Symbol;Acc:MGI:1919455] | 83,65 | 0,6 |
| ENSMUSG00000050796 | 85,9585181 | 56,8 | 1,0 | 6,158154368 | 0,004915442 | B3galt6 | UDP-Gal:betaGal beta 1,3-galactosyltransferase, polypeptide 6 [Source:MGI Symbol;Acc:MGI:2152819] | 75,79 | 0,7 |
| ENSMUSG00000038250 | 143,5650753 | 187,0 | 2,5 | 6,159989992 | 0,001860059 | Usp38 | ubiquitin specific peptidase 38 [Source:MGI Symbol;Acc:MGI:1922091] | 97,19 | 1,4 |
| ENSMUSG00000108402 | 58,77646123 | 82,2 | 3,0 | 6,161968526 | 0,011717836 | RP23-325D10.4 | | 34,71 | 0,3 |
| ENSMUSG00000024059 | 229,4505321 | 390,9 | 10,5 | 6,162012008 | 0,006314703 | Clip4 | CAP-GLY domain containing linker protein family, member 4 [Source:MGI Symbol;Acc:MGI:1919100] | 97,20 | 2,6 |
| ENSMUSG00000028607 | 107,4904111 | 191,9 | 15,1 | 6,166422218 | 0,002147266 | Cpt2 | carnitine palmitoyltransferase 2 [Source:MGI Symbol;Acc:MGI:109176] | 98,74 | 4,7 |
| ENSMUSG00000038866 | 114,5999529 | 107,1 | 124,8 | 6,171534658 | 0,010881861 | Zcchc2 | zinc finger, CCHC domain containing 2 [Source:MGI Symbol;Acc:MGI:2444114] | 79,25 | 20,7 |
| ENSMUSG00000044071 | 89,48898346 | 157,0 | 27,9 | 6,172730537 | 0,003404394 | Fam19a2 | family with sequence similarity 19, member A2 [Source:MGI Symbol;Acc:MGI:2143691] | 88,40 | 2,1 |
| ENSMUSG00000048310 | 88,03023326 | 96,4 | 49,9 | 6,173290367 | 0,022161561 | Pskh1 | protein serine kinase H1 [Source:MGI Symbol;Acc:MGI:3528383] | 99,12 | 19,3 |
| ENSMUSG00000021226 | 30,12303563 | 35,7 | 0,6 | 6,174647611 | 0,045654899 | Acot2 | acyl-CoA thioesterase 2 [Source:MGI Symbol;Acc:MGI:2159605] | 75,09 | 1,1 |
| ENSMUSG00000036270 | 169,595806 | 72,1 | 378,2 | 6,179249561 | 0,035504063 | Edc4 | enhancer of mRNA decapping 4 [Source:MGI Symbol;Acc:MGI:2446249] | 82,85 | 31,8 |
| ENSMUSG00000035568 | 27,80150052 | 45,2 | 0,6 | 6,179814059 | 0,014210371 | Jazf1 | JAZF zinc finger 1 [Source:MGI Symbol;Acc:MGI:2141450] | 92,18 | 1,0 |
| ENSMUSG00000029725 | 79,83734841 | 128,2 | 16,0 | 6,181434994 | 0,004735963 | Ppp1r35 | protein phosphatase 1, regulatory subunit 35 [Source:MGI Symbol;Acc:MGI:1922853] | 95,49 | 17,3 |
| ENSMUSG00000052125 | 71,04289071 | 85,7 | 61,9 | 6,181911963 | 0,032940967 | F730043M19Rik | RIKEN cDNA F730043M19 gene [Source:MGI Symbol;Acc:MGI:2443237] | #DIV/0! | #DIV/0! |
| ENSMUSG00000029185 | 83,67334404 | 151,7 | 21,3 | 6,182358071 | 0,003510998 | Fam114a1 | family with sequence similarity 114, member A1 [Source:MGI Symbol;Acc:MGI:1915553] | 97,58 | 8,9 |
| ENSMUSG00000071350 | 73,41711394 | 97,7 | 20,0 | 6,184962179 | 0,015462113 | Setdb2 | SET domain, bifurcated 2 [Source:MGI Symbol;Acc:MGI:2685139] | 77,02 | 2,8 |
| ENSMUSG00000029312 | 93,87434398 | 106,2 | 17,0 | 6,187161872 | 0,010545033 | Klhl8 | kelch-like 8 [Source:MGI Symbol;Acc:MGI:2179430] | 95,89 | 3,7 |
| ENSMUSG00000029798 | 92,01490185 | 103,4 | 32,9 | 6,187597038 | 0,009843736 | Herc6 | hect domain and RLD 6 [Source:MGI Symbol;Acc:MGI:1914388] | 87,53 | 3,7 |
| ENSMUSG00000025142 | 273,5388965 | 456,9 | 102,1 | 6,193670104 | 0,009728372 | Aspscr1 | veolar soft part sarcoma chromosome region, candidate 1 (human) [Source:MGI Symbol;Acc:MGI:191618 | 82,22 | 3,0 |
| ENSMUSG00000015484 | 1061,184467 | 1766,4 | 299,9 | 6,194183287 | 0,011731631 | Fam163a | family with sequence similarity 163, member A [Source:MGI Symbol;Acc:MGI:3618859] | 100,00 | 34,5 |
| ENSMUSG00000010529 | 98,15040549 | 135,6 | 18,8 | 6,195172355 | 0,007479235 | Gm266 | predicted gene 266 [Source:MGI Symbol;Acc:MGI:2685112] | 99,86 | 22,1 |
| ENSMUSG00000085227 | 143,2653084 | 187,1 | 59,3 | 6,195782794 | 0,003600329 | 6330418K02Rik | RIKEN cDNA 6330418K02 gene [Source:MGI Symbol;Acc:MGI:3697416] | 98,71 | 22,5 |
| ENSMUSG00000030711 | 92,46920724 | 176,2 | 49,6 | 6,198875995 | 0,010210943 | Sult1a1 | sulfotransferase family 1A, phenol-preferring, member 1 [Source:MGI Symbol;Acc:MGI:102896] | 86,82 | 25,5 |
| ENSMUSG00000026674 | 26,23273276 | 61,1 | 0,9 | 6,200731474 | 0,005484728 | Ddr2 | discoidin domain receptor family, member 2 [Source:MGI Symbol;Acc:MGI:1345277] | 54,66 | 0,2 |
| ENSMUSG00000038914 | 225,6378687 | 126,4 | 226,9 | 6,20132066 | 0,0347406 | Dido1 | death inducer-obliterator 1 [Source:MGI Symbol;Acc:MGI:1344352] | 50,14 | 14,1 |
| ENSMUSG00000026614 | 87,91464124 | 76,5 | 2,0 | 6,202206242 | 0,00602753 | Slc30a10 | solute carrier family 30, member 10 [Source:MGI Symbol;Acc:MGI:2685058] | 58,37 | 0,3 |
| ENSMUSG00000020009 | 58,45952823 | 97,8 | 9,2 | 6,202738278 | 0,018259241 | Ifngr1 | interferon gamma receptor 1 [Source:MGI Symbol;Acc:MGI:107655] | 97,44 | 8,4 |
| ENSMUSG00000079559 | 84,36032594 | 119,5 | 77,5 | 6,203431348 | 0,024711206 | Gm684 | predicted gene 684 [Source:MGI Symbol;Acc:MGI:2685530] | 99,97 | 33,3 |
| ENSMUSG00000034931 | 149,2457008 | 139,7 | 53,6 | 6,205970198 | 0,007679962 | Dhx8 | DEAH (Asp-Glu-Ala-His) box polypeptide 8 [Source:MGI Symbol;Acc:MGI:1306823] | 97,76 | 3,2 |
| ENSMUSG00000032123 | 109,8385319 | 105,2 | 1,5 | 6,206940566 | 0,011034089 | Dpagt1 | -N-acetylglucosamine) acetylglucosaminephosphotransferase 1 (GlcNAc-1-P transferase) [Source:MGI Sy | 98,72 | 2,3 |
| ENSMUSG00000032564 | 70,55732551 | 79,7 | 62,8 | 6,210078426 | 0,022891644 | Cpne4 | copine IV [Source:MGI Symbol;Acc:MGI:1921270] | 64,19 | 5,6 |
| ENSMUSG00000098923 | 65,0416016 | 81,5 | 6,0 | 6,212352139 | 0,02371039 | Tmem185b | transmembrane protein 185B [Source:MGI Symbol;Acc:MGI:1917634] | 73,58 | 3,2 |
| ENSMUSG00000032032 | 153,6532624 | 132,2 | 188,5 | 6,21348091 | 0,016861921 | Tmem25 | transmembrane protein 25 [Source:MGI Symbol;Acc:MGI:1918937] | 94,58 | 31,7 |
| ENSMUSG00000037021 | 93,19485377 | 93,5 | 51,8 | 6,214294054 | 0,021600203 | Syngr3 | synaptogyrin 3 [Source:MGI Symbol;Acc:MGI:1341881] | 98,59 | 17,5 |
| ENSMUSG00000031790 | 32,51429655 | 56,5 | 0,6 | 6,214588534 | 0,005219865 | Mmp15 | matrix metallopeptidase 15 [Source:MGI Symbol;Acc:MGI:109320] | 83,49 | 0,4 |
| ENSMUSG00000000708 | 121,5038085 | 203,0 | 9,7 | 6,216359441 | 0,000749905 | Kat2b | K(lysine) acetyltransferase 2B [Source:MGI Symbol;Acc:MGI:1343094] | 88,48 | 3,2 |
| ENSMUSG00000072889 | 90,90425505 | 46,3 | 0,9 | 6,218015097 | 0,029451152 | Nfxl1 | nuclear transcription factor, X-box binding-like 1 [Source:MGI Symbol;Acc:MGI:1923646] | 75,91 | 0,6 |
| ENSMUSG00000029804 | 120,3779308 | 174,0 | 46,2 | 6,218747248 | 0,002532497 | Herc3 | hect domain and RLD 3 [Source:MGI Symbol;Acc:MGI:1921248] | 76,08 | 4,8 |
| ENSMUSG00000026975 | 103,1828967 | 191,4 | 55,5 | 6,220272588 | 0,003469834 | Dph7 | diphthamine biosynethesis 7 [Source:MGI Symbol;Acc:MGI:1914478] | 98,92 | 28,3 |
| ENSMUSG00000022105 | 770,8607163 | 1178,5 | 334,6 | 6,222251532 | 0,001730007 | Rb1 | retinoblastoma 1 [Source:MGI Symbol;Acc:MGI:97874] | 98,22 | 31,2 |
| ENSMUSG00000039670 | 74,99045241 | 115,4 | 45,9 | 6,222455727 | 0,008733864 | Oxld1 | oxidoreductase like domain containing 1 [Source:MGI Symbol;Acc:MGI:1913681] | 99,63 | 31,0 |
| ENSMUSG00000049038 | 107,7817275 | 184,7 | 29,6 | 6,222531563 | 0,00155985 | Mterf2 | mitochondrial transcription termination factor 2 [Source:MGI Symbol;Acc:MGI:1921488] | 99,79 | 8,8 |
| ENSMUSG00000021468 | 115,7820016 | 140,4 | 150,7 | 6,224061245 | 0,015806084 | Sptlc1 | serine palmitoyltransferase, long chain base subunit 1 [Source:MGI Symbol;Acc:MGI:1099431] | 98,08 | 29,7 |
| ENSMUSG00000044734 | 30,75839496 | 46,1 | 0,9 | 6,22445494 | 0,021660411 | Serpinb1a | serine (or cysteine) peptidase inhibitor, clade B, member 1a [Source:MGI Symbol;Acc:MGI:1913472] | 86,65 | 1,3 |
| ENSMUSG00000029189 | 47,98015372 | 63,4 | 0,6 | 6,224504101 | 0,017923708 | Sel1l3 | sel-1 suppressor of lin-12-like 3 (C. elegans) [Source:MGI Symbol;Acc:MGI:1916941] | 75,07 | 0,4 |
| ENSMUSG00000046962 | 51,76043539 | 94,5 | 3,7 | 6,227876461 | 0,008173191 | Zbtb21 | zinc finger and BTB domain containing 21 [Source:MGI Symbol;Acc:MGI:1927240] | 86,49 | 1,5 |
| ENSMUSG00000029016 | 85,28014243 | 107,0 | 56,4 | 6,228081799 | 0,015648648 | Clcn6 | chloride channel, voltage-sensitive 6 [Source:MGI Symbol;Acc:MGI:1347049] | 95,73 | 10,5 |
| ENSMUSG00000040813 | 264,5315976 | 415,0 | 41,5 | 6,229733813 | 0,000749905 | Tex264 | testis expressed gene 264 [Source:MGI Symbol;Acc:MGI:1096570] | 96,43 | 20,4 |
| ENSMUSG00000094936 | 105,645226 | 124,9 | 12,1 | 6,230942336 | 0,011710399 | Rbm4 | RNA binding motif protein 4 [Source:MGI Symbol;Acc:MGI:1100865] | 87,13 | 6,3 |
| ENSMUSG00000052373 | 38,73832975 | 49,6 | 0,9 | 6,232538399 | 0,024247603 | Mpp3 | mbrane protein, palmitoylated 3 (MAGUK p55 subfamily member 3) [Source:MGI Symbol;Acc:MGI:13283 | 53,18 | 0,0 |
| ENSMUSG00000043154 | 131,2722172 | 268,7 | 4,2 | 6,236344289 | 5,58598E-06 | Ppp2r3a | protein phosphatase 2, regulatory subunit B'', alpha [Source:MGI Symbol;Acc:MGI:2442104] | 94,31 | 1,5 |
| ENSMUSG00000030339 | 92,89098049 | 145,2 | 30,9 | 6,237666348 | 0,007933306 | Ltbr | lymphotoxin B receptor [Source:MGI Symbol;Acc:MGI:104875] | 99,35 | 11,7 |
| ENSMUSG00000022184 | 105,2100071 | 129,1 | 7,9 | 6,237803015 | 0,010324857 | Fbxo4 | F-box protein 4 [Source:MGI Symbol;Acc:MGI:2146220] | 98,23 | 8,3 |
| ENSMUSG00000044528 | 84,87057939 | 136,9 | 134,9 | 6,238565541 | 0,008206442 | Tram1l1 | translocation associated membrane protein 1-like 1 [Source:MGI Symbol;Acc:MGI:2443503] | 51,94 | 15,2 |
| ENSMUSG00000038902 | 59,08712522 | 60,6 | 0,9 | 6,24164484 | 0,00471339 | Pogz | pogo transposable element with ZNF domain [Source:MGI Symbol;Acc:MGI:2442117] | 71,81 | 0,3 |
| ENSMUSG00000059970 | 489,4639224 | 843,7 | 21,5 | 6,241837072 | 7,50734E-05 | Hspa2 | heat shock protein 2 [Source:MGI Symbol;Acc:MGI:96243] | 95,61 | 6,8 |
| ENSMUSG00000039254 | 76,03925536 | 107,4 | 92,4 | 6,241846172 | 0,023617627 | Pomt1 | protein-O-mannosyltransferase 1 [Source:MGI Symbol;Acc:MGI:2138994] | 91,41 | 27,1 |

| | | | | | | | | | |
|---|---|---|---|---|---|---|---|---|---|
| ENSMUSG00000046792 | 121,4564978 | 151,8 | 25,0 | 6,243276824 | 0,003679909 | Zfp787 | zinc finger protein 787 [Source:MGI Symbol;Acc:MGI:1914359] | 77,33 | 11,0 |
| ENSMUSG00000086949 | 135,218198 | 198,3 | 94,4 | 6,243300485 | 0,003156877 | Gm13066 | predicted gene 13066 [Source:MGI Symbol;Acc:MGI:3701131] | 81,45 | 27,1 |
| ENSMUSG00000021234 | 69,93587213 | 100,5 | 64,7 | 6,243580313 | 0,031057321 | Fam161b | family with sequence similarity 161, member B [Source:MGI Symbol;Acc:MGI:2443027] | 85,25 | 13,8 |
| ENSMUSG00000023947 | 29,1665087 | 44,6 | 0,6 | 6,244367803 | 0,024490156 | Nfkbie | actor of kappa light polypeptide gene enhancer in B cells inhibitor, epsilon [Source:MGI Symbol;Acc:MGI:1 | 89,18 | 0,4 |
| ENSMUSG00000048402 | 43,58049609 | 90,4 | 1,8 | 6,248243284 | 0,005444277 | Gli2 | GLI-Kruppel family member GLI2 [Source:MGI Symbol;Acc:MGI:95728] | 76,56 | 0,5 |
| ENSMUSG00000055319 | 144,3107318 | 128,2 | 75,8 | 6,249892826 | 0,009132349 | Sec23ip | Sec23 interacting protein [Source:MGI Symbol;Acc:MGI:2450915] | 87,83 | 6,4 |
| ENSMUSG00000023034 | 30,88823191 | 41,3 | 0,6 | 6,250846446 | 0,035404661 | Nr4a1 | nuclear receptor subfamily 4, group A, member 1 [Source:MGI Symbol;Acc:MGI:1352454] | 94,38 | 0,9 |
| ENSMUSG00000044562 | 80,75458535 | 105,6 | 14,8 | 6,251238602 | 0,012325428 | Rasip1 | Ras interacting protein 1 [Source:MGI Symbol;Acc:MGI:1917153] | 60,42 | 6,6 |
| ENSMUSG00000020723 | 48,99272378 | 71,4 | 2,8 | 6,251455977 | 0,022038199 | Cacng4 | calcium channel, voltage-dependent, gamma subunit 4 [Source:MGI Symbol;Acc:MGI:1859167] | 72,76 | 1,3 |
| ENSMUSG00000052852 | 82,86979679 | 159,0 | 2,2 | 6,251496946 | 0,000152823 | Reep1 | receptor accessory protein 1 [Source:MGI Symbol;Acc:MGI:1098827] | 92,38 | 1,3 |
| ENSMUSG00000027300 | 67,97586441 | 125,1 | 3,6 | 6,251543389 | 0,003527765 | Ubox5 | U box domain containing 5 [Source:MGI Symbol;Acc:MGI:2154658] | 82,98 | 1,4 |
| ENSMUSG00000020850 | 267,0249241 | 263,1 | 19,4 | 6,251599984 | 0,000247693 | Prpf8 | pre-mRNA processing factor 8 [Source:MGI Symbol;Acc:MGI:2179381] | 82,00 | 2,8 |
| ENSMUSG00000037640 | 211,3836124 | 152,2 | 18,5 | 6,253044703 | 0,004375869 | Zfp60 | zinc finger protein 60 [Source:MGI Symbol;Acc:MGI:99207] | 69,55 | 2,0 |
| ENSMUSG00000108175 | 88,7227155 | 164,6 | 9,7 | 6,253466179 | 0,002556717 | Gm43938 | predicted gene, 43938 [Source:MGI Symbol;Acc:MGI:5690330] | 100,00 | 4,1 |
| ENSMUSG00000044636 | 36,74526041 | 72,2 | 2,0 | 6,254159214 | 0,011233016 | Csrnp2 | cysteine-serine-rich nuclear protein 2 [Source:MGI Symbol;Acc:MGI:2386852] | 90,15 | 0,6 |
| ENSMUSG00000027272 | 73,29128312 | 81,1 | 56,9 | 6,254175345 | 0,017912342 | Ubr1 | ubiquitin protein ligase E3 component n-recognin 1 [Source:MGI Symbol;Acc:MGI:1277977] | 67,36 | 2,3 |
| ENSMUSG00000031641 | 123,6456898 | 180,9 | 78,1 | 6,255331438 | 0,006259021 | Cbr4 | carbonyl reductase 4 [Source:MGI Symbol;Acc:MGI:2384567] | 94,02 | 27,4 |
| ENSMUSG00000003865 | 190,4239802 | 327,2 | 67,1 | 6,256398761 | 0,002344814 | Gys1 | glycogen synthase 1, muscle [Source:MGI Symbol;Acc:MGI:101805] | 99,68 | 19,3 |
| ENSMUSG00000017767 | 32,13082745 | 43,5 | 1,0 | 6,257800565 | 0,034988593 | Spata25 | spermatogenesis associated 25 [Source:MGI Symbol;Acc:MGI:1922892] | 0,00 | 2,8 |
| ENSMUSG00000053646 | 376,1360993 | 553,7 | 179,9 | 6,258723918 | 0,002856502 | Plxnb1 | plexin B1 [Source:MGI Symbol;Acc:MGI:2154238] | 94,11 | 18,3 |
| ENSMUSG00000048027 | 62,06288907 | 112,6 | 2,8 | 6,259870963 | 0,001042339 | Rgmb | repulsive guidance molecule family member B [Source:MGI Symbol;Acc:MGI:1916049] | 98,57 | 1,4 |
| ENSMUSG00000103009 | 98,10090815 | 168,8 | 52,7 | 6,264370706 | 0,0047196 | Gm4430 | predicted gene 4430 [Source:MGI Symbol;Acc:MGI:3782614] | 99,11 | 24,2 |
| ENSMUSG00000085084 | 49,51076636 | 89,7 | 4,0 | 6,264674148 | 0,008771538 | 4930570G19Rik | RIKEN cDNA 4930570G19 gene [Source:MGI Symbol;Acc:MGI:1923158] | #DIV/0! | #DIV/0! |
| ENSMUSG00000035329 | 285,4041445 | 325,5 | 32,5 | 6,266487119 | 0,001581792 | Fbxo33 | F-box protein 33 [Source:MGI Symbol;Acc:MGI:1917861] | 94,93 | 6,3 |
| ENSMUSG00000025231 | 74,27291539 | 93,0 | 4,6 | 6,267431201 | 0,01198919 | Sufu | suppressor of fused homolog (Drosophila) [Source:MGI Symbol;Acc:MGI:1345643] | 64,31 | 1,3 |
| ENSMUSG00000038156 | 92,70319014 | 190,3 | 2,5 | 6,268487808 | 0,00045752 | Spon1 | spondin 1, (f-spondin) extracellular matrix protein [Source:MGI Symbol;Acc:MGI:2385287] | 78,00 | 0,8 |
| ENSMUSG00000097616 | 92,65670559 | 127,8 | 59,9 | 6,268754897 | 0,015235909 | 1110019D14Rik | RIKEN cDNA 1110019D14 gene [Source:MGI Symbol;Acc:MGI:1923561] | #DIV/0! | #DIV/0! |
| ENSMUSG00000024190 | 143,9055766 | 250,1 | 4,0 | 6,272145029 | 0,002385293 | Dusp1 | dual specificity phosphatase 1 [Source:MGI Symbol;Acc:MGI:105120] | 98,17 | 2,4 |
| ENSMUSG00000044676 | 92,2897133 | 129,6 | 121,0 | 6,273353774 | 0,019631143 | Zfp612 | zinc finger protein 612 [Source:MGI Symbol;Acc:MGI:2443465] | 100,00 | 29,8 |
| ENSMUSG00000029449 | 56,47672527 | 74,1 | 1,8 | 6,274902538 | 0,009002706 | Rhof | ras homolog gene family, member f [Source:MGI Symbol;Acc:MGI:1345629] | 53,85 | 0,6 |
| ENSMUSG00000069171 | 201,8182336 | 343,0 | 138,3 | 6,275120175 | 0,002402222 | Nr2f1 | nuclear receptor subfamily 2, group F, member 1 [Source:MGI Symbol;Acc:MGI:1352451] | 62,56 | 17,5 |
| ENSMUSG00000026566 | 130,6186307 | 147,3 | 16,6 | 6,275545846 | 0,004990486 | Mpzl1 | myelin protein zero-like 1 [Source:MGI Symbol;Acc:MGI:1915731] | 95,87 | 3,7 |
| ENSMUSG00000040729 | 45,18925765 | 85,5 | 1,2 | 6,276099626 | 0,00184252 | Cep126 | centrosomal protein 126 [Source:MGI Symbol;Acc:MGI:2680221] | 97,14 | 0,7 |
| ENSMUSG00000027546 | 218,9017078 | 180,9 | 137,7 | 6,277238857 | 0,003674644 | Atp9a | ATPase, class II, type 9A [Source:MGI Symbol;Acc:MGI:1330826] | 76,49 | 21,3 |
| ENSMUSG00000030423 | 215,149024 | 409,3 | 140,6 | 6,278836613 | 0,000764367 | Pop4 | ocessing of precursor 4, ribonuclease P/MRP family, (S. cerevisiae) [Source:MGI Symbol;Acc:MGI:191341 | 60,66 | 18,6 |
| ENSMUSG00000037003 | 55,30540621 | 130,8 | 2,4 | 6,278922672 | 0,000850563 | Tns2 | tensin 2 [Source:MGI Symbol;Acc:MGI:2387586] | 89,52 | 0,6 |
| ENSMUSG00000011427 | 46,0529675 | 79,0 | 2,0 | 6,27912471 | 0,00786331 | Zfp790 | zinc finger protein 790 [Source:MGI Symbol;Acc:MGI:1923431] | 89,32 | 0,6 |
| ENSMUSG00000045934 | 249,4715472 | 317,7 | 240,1 | 6,279157677 | 0,003438138 | Mtmr11 | myotubularin related protein 11 [Source:MGI Symbol;Acc:MGI:2652817] | 100,00 | 33,6 |
| ENSMUSG00000021665 | 179,1450825 | 209,4 | 53,6 | 6,280579574 | 0,001799574 | Hexb | hexosaminidase B [Source:MGI Symbol;Acc:MGI:96074] | 99,61 | 24,7 |
| ENSMUSG00000029028 | 290,7293172 | 237,2 | 163,6 | 6,280777755 | 0,000675647 | Lrrc47 | leucine rich repeat containing 47 [Source:MGI Symbol;Acc:MGI:1920196] | 96,82 | 28,5 |
| ENSMUSG00000034731 | 97,46681092 | 70,0 | 96,8 | 6,2821324 | 0,034238868 | Dgkh | diacylglycerol kinase, eta [Source:MGI Symbol;Acc:MGI:2444188] | 91,89 | 3,9 |
| ENSMUSG00000009216 | 285,2225561 | 546,6 | 8,8 | 6,28684634 | 1,92458E-05 | Fam163b | family with sequence similarity 163, member B [Source:MGI Symbol;Acc:MGI:1926106] | 99,22 | 3,4 |
| ENSMUSG00000029999 | 130,9160601 | 138,2 | 107,8 | 6,287688275 | 0,01361908 | Tgfa | transforming growth factor alpha [Source:MGI Symbol;Acc:MGI:98724] | 97,70 | 21,4 |
| ENSMUSG00000040520 | 231,2297545 | 398,0 | 37,0 | 6,290909911 | 0,005876089 | Manea | mannosidase, endo-alpha [Source:MGI Symbol;Acc:MGI:2444484] | 97,53 | 3,9 |
| ENSMUSG00000026672 | 103,743239 | 162,5 | 55,5 | 6,292886085 | 0,011078997 | Optn | optineurin [Source:MGI Symbol;Acc:MGI:1918898] | 99,94 | 18,8 |
| ENSMUSG00000036840 | 242,3932412 | 390,9 | 210,5 | 6,298944901 | 0,003290842 | Siah1a | seven in absentia 1A [Source:MGI Symbol;Acc:MGI:108064] | 99,21 | 30,7 |
| ENSMUSG00000034194 | 157,9286424 | 188,0 | 56,4 | 6,300971781 | 0,001817852 | R3hcc1 | R3H domain and coiled-coil containing 1 [Source:MGI Symbol;Acc:MGI:1919093] | 98,23 | 26,2 |
| ENSMUSG00000063171 | 83,84683286 | 128,3 | 127,7 | 6,301177705 | 0,008430306 | Rps4l | ribosomal protein S4-like [Source:MGI Symbol;Acc:MGI:1913434] | #DIV/0! | #DIV/0! |
| ENSMUSG00000090608 | 92,2464638 | 125,9 | 16,6 | 6,30279145 | 0,008376539 | Gm17200 | predicted gene 17200 [Source:MGI Symbol;Acc:MGI:4938027] | #DIV/0! | #DIV/0! |
| ENSMUSG00000029186 | 174,6528369 | 136,3 | 134,9 | 6,303387121 | 0,008924977 | Pi4k2b | phosphatidylinositol 4-kinase type 2 beta [Source:MGI Symbol;Acc:MGI:1914323] | 93,86 | 19,4 |
| ENSMUSG00000002679 | 145,2187253 | 200,8 | 42,4 | 6,305871904 | 0,004245988 | Med6 | mediator complex subunit 6 [Source:MGI Symbol;Acc:MGI:1917042] | 93,62 | 25,8 |
| ENSMUSG00000062822 | 174,810964 | 200,0 | 117,4 | 6,306460049 | 0,002157466 | 4833420G17Rik | RIKEN cDNA 4833420G17 gene [Source:MGI Symbol;Acc:MGI:1914642] | 99,53 | 29,2 |
| ENSMUSG00000054277 | 89,26482271 | 109,9 | 14,8 | 6,30654699 | 0,017208103 | Arfgap3 | ADP-ribosylation factor GTPase activating protein 3 [Source:MGI Symbol;Acc:MGI:1913501] | 99,05 | 2,8 |
| ENSMUSG00000051007 | 119,0556256 | 153,0 | 56,4 | 6,308217775 | 0,009471772 | Pddc1 | Parkinson disease 7 domain containing 1 [Source:MGI Symbol;Acc:MGI:2387178] | 97,48 | 31,0 |
| ENSMUSG00000069049 | 175,3140494 | 170,9 | 58,2 | 6,309019569 | 0,004721099 | Eif2s3y | ryotic translation initiation factor 2, subunit 3, structural gene Y-linked [Source:MGI Symbol;Acc:MGI:1349 | 97,13 | 8,4 |
| ENSMUSG00000029171 | 76,22639517 | 105,6 | 68,9 | 6,309550657 | 0,010465506 | Pgm1 | phosphoglucomutase 1 [Source:MGI Symbol;Acc:MGI:97564] | 97,14 | 20,6 |
| ENSMUSG00000018750 | 128,9006431 | 184,7 | 2,5 | 6,309897256 | 1,99491E-05 | Zbtb4 | zinc finger and BTB domain containing 4 [Source:MGI Symbol;Acc:MGI:1922830] | 80,27 | 0,3 |
| ENSMUSG00000028655 | 235,3819045 | 393,7 | 27,8 | 6,310746039 | 0,002714818 | Mfsd2a | major facilitator superfamily domain containing 2A [Source:MGI Symbol;Acc:MGI:1923824] | 100,00 | 14,4 |
| ENSMUSG00000022861 | 88,22215021 | 118,7 | 13,0 | 6,312284769 | 0,007630642 | Dgkg | diacylglycerol kinase, gamma [Source:MGI Symbol;Acc:MGI:105060] | 96,52 | 1,6 |
| ENSMUSG00000089669 | 58,69721084 | 96,2 | 2,4 | 6,313004998 | 0,008948721 | Tnfsf13 | tumor necrosis factor (ligand) superfamily, member 13 [Source:MGI Symbol;Acc:MGI:1916833] | 88,82 | 0,8 |
| ENSMUSG00000024935 | 72,58536504 | 39,8 | 0,6 | 6,315527096 | 0,040594928 | Slc1a1 | ily 1 (neuronal/epithelial high affinity glutamate transporter, system Xag), member 1 [Source:MGI Symbol;/ | 78,09 | 0,4 |
| ENSMUSG00000086587 | 79,53770673 | 129,0 | 54,9 | 6,316695109 | 0,007079765 | Gm11837 | predicted gene 11837 [Source:MGI Symbol;Acc:MGI:3702175] | 86,66 | 19,9 |
| ENSMUSG00000029575 | 296,2640143 | 460,1 | 81,4 | 6,317146974 | 0,001079602 | Mmab | :hylmalonic aciduria (cobalamin deficiency) type B homolog (human) [Source:MGI Symbol;Acc:MGI:19249 | 97,86 | 21,8 |
| ENSMUSG00000026571 | 57,98026597 | 69,2 | 1,8 | 6,318521045 | 0,015050976 | Dcaf6 | DDB1 and CUL4 associated factor 6 [Source:MGI Symbol;Acc:MGI:1921356] | 70,17 | 1,1 |
| ENSMUSG00000026942 | 142,6866876 | 160,5 | 164,0 | 6,318742505 | 0,01348487 | Traf2 | TNF receptor-associated factor 2 [Source:MGI Symbol;Acc:MGI:101835] | 94,30 | 28,6 |
| ENSMUSG00000028221 | 1196,441892 | 1956,7 | 221,5 | 6,318912847 | 1,68917E-06 | Tmem55a | transmembrane protein 55A [Source:MGI Symbol;Acc:MGI:1919769] | 95,77 | 34,9 |
| ENSMUSG00000058070 | 134,3440677 | 114,8 | 179,7 | 6,320894465 | 0,0277816 | Eml1 | echinoderm microtubule associated protein like 1 [Source:MGI Symbol;Acc:MGI:1915769] | 87,80 | 26,7 |
| ENSMUSG00000041438 | 170,161472 | 159,2 | 186,7 | 6,321923938 | 0,00685741 | Cirh1a | cirrhosis, autosomal recessive 1A (human) [Source:MGI Symbol;Acc:MGI:1096573] | 99,04 | 32,7 |
| ENSMUSG00000071369 | 68,73417197 | 115,4 | 3,0 | 6,323085324 | 0,001180038 | Map3k5 | mitogen-activated protein kinase kinase kinase 5 [Source:MGI Symbol;Acc:MGI:1346876] | 84,17 | 0,4 |
| ENSMUSG00000032867 | 414,2152627 | 611,5 | 365,8 | 6,323568615 | 0,002993446 | Fbxw8 | F-box and WD-40 domain protein 8 [Source:MGI Symbol;Acc:MGI:1923041] | 93,47 | 22,7 |
| ENSMUSG00000043415 | 75,31906628 | 136,3 | 14,0 | 6,325814883 | 0,002803331 | Otud1 | OTU domain containing 1 [Source:MGI Symbol;Acc:MGI:1918448] | 96,27 | 3,8 |
| ENSMUSG00000038175 | 105,6392413 | 59,7 | 1,0 | 6,328616121 | 0,004419591 | Mylip | myosin regulatory light chain interacting protein [Source:MGI Symbol;Acc:MGI:2388271] | 90,95 | 0,8 |
| ENSMUSG00000029088 | 74,62436187 | 99,8 | 3,7 | 6,328653221 | 0,007066089 | Kcnip4 | Kv channel interacting protein 4 [Source:MGI Symbol;Acc:MGI:1933131] | 76,76 | 1,5 |
| ENSMUSG00000039308 | 73,13407473 | 68,1 | 1,2 | 6,32971385 | 0,013757748 | Ndst2 | N-deacetylase/N-sulfotransferase (heparan glucosaminyl) 2 [Source:MGI Symbol;Acc:MGI:97040] | 95,53 | 0,4 |
| ENSMUSG00000026830 | 84,10265066 | 122,5 | 139,6 | 6,331099714 | 0,007094359 | Ermn | ermin, ERM-like protein [Source:MGI Symbol;Acc:MGI:1925017] | 96,39 | 28,7 |
| ENSMUSG00000021375 | 159,3929343 | 176,4 | 274,2 | 6,335040984 | 0,010465506 | Kif13a | kinesin family member 13A [Source:MGI Symbol;Acc:MGI:1098264] | 97,08 | 16,2 |
| ENSMUSG00000003549 | 130,8119573 | 186,1 | 8,3 | 6,336682595 | 0,000319142 | Ercc1 | epair cross-complementing rodent repair deficiency, complementation group 1 [Source:MGI Symbol;Acc:M | 76,76 | 2,0 |
| ENSMUSG00000000957 | 114,386606 | 156,0 | 2,1 | 6,337984309 | 0,010931263 | Mmp14 | matrix metallopeptidase 14 (membrane-inserted) [Source:MGI Symbol;Acc:MGI:101900] | 91,07 | 1,0 |
| ENSMUSG00000033809 | 109,9578659 | 143,1 | 108,3 | 6,338512892 | 0,011819715 | Alg3 | asparagine-linked glycosylation 3 (alpha-1,3-mannosyltransferase) [Source:MGI Symbol;Acc:MGI:1098592 | 99,04 | 32,4 |
| ENSMUSG00000042734 | 88,7016566 | 153,7 | 30,5 | 6,340175789 | 0,003257241 | Ttc9 | tetratricopeptide repeat domain 9 [Source:MGI Symbol;Acc:MGI:1916730] | 100,00 | 19,4 |
| ENSMUSG00000092009 | 116,3508623 | 113,1 | 105,8 | 6,343098323 | 0,019281634 | Myh15 | myosin, heavy chain 15 [Source:MGI Symbol;Acc:MGI:3643515] | 19,10 | 6,0 |
| ENSMUSG00000032114 | 198,3527989 | 156,2 | 4,8 | 6,343353099 | 0,001751225 | Slc37a4 | lute carrier family 37 (glucose-6-phosphate transporter), member 4 [Source:MGI Symbol;Acc:MGI:131665 | 95,55 | 3,7 |
| ENSMUSG00000092242 | 55,27848074 | 84,2 | 32,9 | 6,34420328 | 0,024757301 | Gm20515 | predicted gene 20515 [Source:MGI Symbol;Acc:MGI:5141980] | #DIV/0! | #DIV/0! |
| ENSMUSG00000024563 | 159,3709782 | 197,5 | 72,1 | 6,344673585 | 0,002402222 | Smad2 | SMAD family member 2 [Source:MGI Symbol;Acc:MGI:108051] | 75,33 | 16,6 |
| ENSMUSG00000029428 | 164,710144 | 184,6 | 72,6 | 6,345211334 | 0,005718965 | Stx2 | syntaxin 2 [Source:MGI Symbol;Acc:MGI:108059] | 97,95 | 28,1 |
| ENSMUSG00000049232 | 133,8290799 | 183,7 | 124,8 | 6,346116954 | 0,003758957 | Tigd2 | tigger transposable element derived 2 [Source:MGI Symbol;Acc:MGI:1915390] | 99,10 | 26,8 |
| ENSMUSG00000032714 | 56,97773131 | 121,0 | 3,0 | 6,347763847 | 0,000816967 | Syde1 | synapse defective 1, Rho GTPase, homolog 1 (C. elegans) [Source:MGI Symbol;Acc:MGI:1918959] | 99,04 | 0,6 |
| ENSMUSG00000026649 | 66,22866357 | 96,6 | 12,9 | 6,347846641 | 0,018830394 | Cfap126 | cilia and flagella associated protein 126 [Source:MGI Symbol;Acc:MGI:1922722] | 70,64 | 9,8 |
| ENSMUSG00000037808 | 46,11013768 | 105,7 | 1,5 | 6,348212079 | 0,014923234 | Fam76b | family with sequence similarity 76, member B [Source:MGI Symbol;Acc:MGI:1920076] | 93,28 | 0,8 |
| ENSMUSG00000040174 | 130,9199093 | 158,2 | 119,2 | 6,34861468 | 0,004721099 | Alkbh3 | alkB homolog 3, alpha-ketoglutarate-dependent dioxygenase [Source:MGI Symbol;Acc:MGI:1916363] | 97,99 | 29,0 |
| ENSMUSG00000031174 | 83,14511667 | 116,5 | 58,2 | 6,348915919 | 0,010732319 | Rpgr | retinitis pigmentosa GTPase regulator [Source:MGI Symbol;Acc:MGI:1344037] | 61,69 | 7,8 |
| ENSMUSG00000030880 | 155,1205751 | 150,7 | 42,5 | 6,349788445 | 0,006344489 | Polr3e | polymerase (RNA) III (DNA directed) polypeptide E [Source:MGI Symbol;Acc:MGI:1349452] | 78,26 | 12,7 |
| ENSMUSG00000029712 | 38,94000694 | 68,3 | 1,8 | 6,352198856 | 0,018980258 | Actl6b | actin-like 6B [Source:MGI Symbol;Acc:MGI:1933548] | 88,50 | 1,4 |
| ENSMUSG00000032387 | 88,48964406 | 132,7 | 42,4 | 6,35225012 | 0,017721513 | Rbpms2 | RNA binding protein with multiple splicing 2 [Source:MGI Symbol;Acc:MGI:1919223] | 91,69 | 21,6 |
| ENSMUSG00000068250 | 130,5628663 | 181,3 | 48,1 | 6,355925723 | 0,004041278 | Amn1 | antagonist of mitotic exit network 1 [Source:MGI Symbol;Acc:MGI:2442933] | 83,09 | 14,7 |
| ENSMUSG00000056228 | 83,40042951 | 93,5 | 23,1 | 6,356299292 | 0,029043621 | Cars2 | cysteinyl-tRNA synthetase 2 (mitochondrial)(putative) [Source:MGI Symbol;Acc:MGI:1919191] | 97,31 | 11,6 |
| ENSMUSG00000036052 | 150,2126409 | 195,7 | 41,6 | 6,35681088 | 0,003091172 | Dnajb5 | DnaJ heat shock protein family (Hsp40) member B5 [Source:MGI Symbol;Acc:MGI:1930018] | 79,35 | 9,2 |
| ENSMUSG00000039952 | 342,2513054 | 522,1 | 46,1 | 6,357333701 | 0,006961964 | Dag1 | dystroglycan 1 [Source:MGI Symbol;Acc:MGI:101864] | 91,94 | 5,4 |
| ENSMUSG00000005804 | 454,4563505 | 653,5 | 183,6 | 6,360406926 | 0,011506412 | Bloc1s6 | biogenesis of lysosomal organelles complex-1, subunit 6, pallidin [Source:MGI Symbol;Acc:MGI:1927580] | 77,62 | 20,5 |
| ENSMUSG00000097164 | 211,9405062 | 146,2 | 109,1 | 6,362776357 | 0,007785233 | Cep83os | centrosomal protein 83, opposite strand [Source:MGI Symbol;Acc:MGI:1914973] | #DIV/0! | #DIV/0! |
| ENSMUSG00000000202 | 403,309056 | 595,1 | 110,3 | 6,362865499 | 0,005518094 | Btbd17 | BTB (POZ) domain containing 17 [Source:MGI Symbol;Acc:MGI:1919264] | 99,94 | 29,6 |
| ENSMUSG00000021549 | 158,3945497 | 297,6 | 8,9 | 6,363589524 | 0,000718453 | Rasa1 | RAS p21 protein activator 1 [Source:MGI Symbol;Acc:MGI:97860] | 95,97 | 1,3 |
| ENSMUSG00000063698 | 67,65529732 | 121,8 | 28,9 | 6,373018319 | 0,008306247 | Sfxn4 | sideroflexin 4 [Source:MGI Symbol;Acc:MGI:2137680] | 97,12 | 8,1 |
| ENSMUSG00000026650 | 58,30780595 | 97,0 | 4,6 | 6,37338934 | 0,015421831 | Meig1 | meiosis expressed gene 1 [Source:MGI Symbol;Acc:MGI:1202878] | 60,89 | 1,8 |
| ENSMUSG00000040690 | 61,63648719 | 46,3 | 0,9 | 6,376719354 | 0,030993462 | Col16a1 | collagen, type XVI, alpha 1 [Source:MGI Symbol;Acc:MGI:1095396] | 65,52 | 0,2 |
| ENSMUSG00000028292 | 91,20073782 | 161,0 | 61,0 | 6,37799397 | 0,004936822 | Rars2 | arginyl-tRNA synthetase 2, mitochondrial [Source:MGI Symbol;Acc:MGI:1923596] | 99,52 | 24,8 |
| ENSMUSG00000031584 | 352,0166391 | 510,2 | 314,3 | 6,378094559 | 0,002535368 | Gsr | glutathione reductase [Source:MGI Symbol;Acc:MGI:95804] | 93,93 | 29,4 |
| ENSMUSG00000042579 | 54,54632328 | 103,3 | 60,9 | 6,378663278 | 0,017657042 | 4632404H12Rik | RIKEN cDNA 4632404H12 gene [Source:MGI Symbol;Acc:MGI:1921284] | #DIV/0! | #DIV/0! |
| ENSMUSG00000021711 | 424,7727081 | 755,9 | 75,5 | 6,382079098 | 0,000358526 | Trappc13 | trafficking protein particle complex 13 [Source:MGI Symbol;Acc:MGI:1914225] | 99,42 | 13,7 |
| ENSMUSG00000053581 | 430,0723381 | 746,2 | 125,2 | 6,383954851 | 0,008263317 | Zfand2a | zinc finger, AN1-type domain 2A [Source:MGI Symbol;Acc:MGI:2140729] | 98,89 | 26,6 |
| ENSMUSG00000021589 | 428,2824058 | 740,9 | 208,1 | 6,386136024 | 0,000756646 | Rhobtb3 | Rho-related BTB domain containing 3 [Source:MGI Symbol;Acc:MGI:1920546] | 98,35 | 24,9 |
| ENSMUSG00000031993 | 161,4080476 | 159,8 | 28,4 | 6,386791853 | 0,005334587 | Snx19 | sorting nexin 19 [Source:MGI Symbol;Acc:MGI:1921581] | 94,30 | 3,3 |
| ENSMUSG00000026960 | 75,41043163 | 123,6 | 3,0 | 6,388864705 | 0,003679909 | Arl6ip6 | ADP-ribosylation factor-like 6 interacting protein 6 [Source:MGI Symbol;Acc:MGI:1929507] | 85,86 | 2,0 |
| ENSMUSG00000029148 | 457,2027418 | 608,8 | 113,8 | 6,3891806 | 7,31758E-05 | Nrbp1 | nuclear receptor binding protein 1 [Source:MGI Symbol;Acc:MGI:2183436] | 99,51 | 31,2 |
| ENSMUSG00000019470 | 132,1522143 | 153,5 | 95,6 | 6,389612709 | 0,02185231 | Xab2 | XPA binding protein 2 [Source:MGI Symbol;Acc:MGI:1914689] | 99,39 | 30,0 |
| ENSMUSG00000095432 | 157,3003799 | 77,5 | 140,7 | 6,392080003 | 0,034786131 | Zfp748 | zinc finger protein 748 [Source:MGI Symbol;Acc:MGI:1916455] | 77,91 | 4,9 |
| ENSMUSG00000027339 | 309,5512202 | 593,4 | 57,1 | 6,392745356 | 0,000131252 | Rassf2 | Ras association (RalGDS/AF-6) domain family member 2 [Source:MGI Symbol;Acc:MGI:2442060] | 75,17 | 5,4 |
| ENSMUSG00000037621 | 82,56930261 | 128,5 | 1,6 | 6,394848548 | 0,000137743 | Atoh8 | atonal bHLH transcription factor 8 [Source:MGI Symbol;Acc:MGI:1918343] | 91,99 | 1,7 |
| ENSMUSG00000037907 | 59,94232988 | 54,2 | 0,9 | 6,396493955 | 0,023115987 | Ankrd13b | ankyrin repeat domain 13b [Source:MGI Symbol;Acc:MGI:2144501] | 79,20 | 0,7 |
| ENSMUSG00000039917 | 378,5573085 | 528,5 | 133,5 | 6,398213727 | 0,000756646 | Rhbdd2 | rhomboid domain containing 2 [Source:MGI Symbol;Acc:MGI:1915612] | 96,06 | 18,5 |
| ENSMUSG00000003208 | 64,26865734 | 112,0 | 11,0 | 6,399183744 | 0,009341191 | Ccdc94 | coiled-coil domain containing 94 [Source:MGI Symbol;Acc:MGI:1920136] | 99,52 | 7,7 |
| ENSMUSG00000020038 | 193,9861657 | 371,4 | 7,8 | 6,40298183 | 0,000551724 | Cry1 | cryptochrome 1 (photolyase-like) [Source:MGI Symbol;Acc:MGI:1270841] | 99,71 | 3,1 |
| ENSMUSG00000047181 | 60,82914884 | 91,3 | 1,8 | 6,403917838 | 0,008019388 | Samd14 | sterile alpha motif domain containing 14 [Source:MGI Symbol;Acc:MGI:2384945] | 86,44 | 0,7 |
| ENSMUSG00000056091 | 163,7425379 | 253,5 | 3,1 | 6,405427265 | 0,003190773 | St3gal5 | ST3 beta-galactoside alpha-2,3-sialyltransferase 5 [Source:MGI Symbol;Acc:MGI:1339963] | 87,92 | 2,5 |
| ENSMUSG00000079467 | 51,27586944 | 60,1 | 0,6 | 6,406720378 | 0,006314703 | Gm14966 | predicted gene 14966 [Source:MGI Symbol;Acc:MGI:3705223] | 81,84 | 3,1 |
| ENSMUSG00000022244 | 97,36059969 | 183,0 | 55,5 | 6,413570413 | 0,00416495 | Amacr | alpha-methylacyl-CoA racemase [Source:MGI Symbol;Acc:MGI:1098273] | 62,12 | 18,3 |
| ENSMUSG00000058979 | 206,160703 | 322,9 | 10,2 | 6,416201461 | 0,003204932 | Cecr5 | cat eye syndrome chromosome region, candidate 5 [Source:MGI Symbol;Acc:MGI:2136976] | 99,81 | 6,9 |
| ENSMUSG00000024908 | 242,7039743 | 329,5 | 4,3 | 6,41625265 | 4,30787E-06 | Ppp6r3 | protein phosphatase 6, regulatory subunit 3 [Source:MGI Symbol;Acc:MGI:1921807] | 99,72 | 2,1 |
| ENSMUSG00000052062 | 63,23958926 | 52,4 | 1,0 | 6,416274542 | 0,021600203 | Pard3b | par-3 family cell polarity regulator beta [Source:MGI Symbol;Acc:MGI:1919301] | 46,07 | 0,0 |
| ENSMUSG00000024486 | 85,17472731 | 157,1 | 12,0 | 6,416281908 | 0,004418169 | Hbegf | heparin-binding EGF-like growth factor [Source:MGI Symbol;Acc:MGI:96070] | 97,65 | 7,8 |
| ENSMUSG00000074219 | 46,96279237 | 76,8 | 1,2 | 6,418938552 | 0,010855466 | Gm10644 | predicted gene 10644 [Source:MGI Symbol;Acc:MGI:3704314] | 98,44 | 0,7 |
| ENSMUSG00000006021 | 120,0998829 | 161,3 | 36,9 | 6,422822829 | 0,009954247 | Kptn | kaptin [Source:MGI Symbol;Acc:MGI:1890380] | 99,37 | 12,0 |
| ENSMUSG00000036814 | 88,80748306 | 94,3 | 49,0 | 6,425327567 | 0,020180659 | Slc6a20a | solute carrier family 6 (neurotransmitter transporter), member 20A [Source:MGI Symbol;Acc:MGI:2143217 | 85,99 | 21,9 |
| ENSMUSG00000044730 | 89,604824 | 106,8 | 49,0 | 6,426142739 | 0,019277519 | 9930104L06Rik | RIKEN cDNA 9930104L06 gene [Source:MGI Symbol;Acc:MGI:3041172] | #DIV/0! | #DIV/0! |
| ENSMUSG00000037022 | 83,31749785 | 126,5 | 59,9 | 6,428419562 | 0,021374931 | Mmaa | methylmalonic aciduria (cobalamin deficiency) type A [Source:MGI Symbol;Acc:MGI:1923805] | 79,69 | 14,3 |
| ENSMUSG00000021892 | 142,6283159 | 153,6 | 110,9 | 6,42869821 | 0,005518094 | Sh3bp5 | SH3-domain binding protein 5 (BTK-associated) [Source:MGI Symbol;Acc:MGI:1344391] | 90,81 | 24,0 |
| ENSMUSG00000042246 | 116,1774264 | 198,4 | 118,6 | 6,429716734 | 0,00670992 | Tmc7 | transmembrane channel-like gene family 7 [Source:MGI Symbol;Acc:MGI:2443317] | 98,76 | 14,0 |
| ENSMUSG00000059288 | 87,20752146 | 136,8 | 7,0 | 6,430243756 | 0,002207462 | Cdyl | chromodomain protein, Y chromosome-like [Source:MGI Symbol;Acc:MGI:1339956] | 84,00 | 1,8 |
| ENSMUSG00000021219 | 125,2914715 | 126,5 | 1,5 | 6,43059727 | 0,000203151 | Rgs6 | regulator of G-protein signaling 6 [Source:MGI Symbol;Acc:MGI:1354730] | 58,55 | 0,2 |
| ENSMUSG00000044934 | 337,7957611 | 385,5 | 9,2 | 6,431713819 | 0,000323009 | Zfp367 | zinc finger protein 367 [Source:MGI Symbol;Acc:MGI:2442266] | 56,83 | 2,2 |
| ENSMUSG00000073700 | 118,5501923 | 115,7 | 109,8 | 6,433186317 | 0,016994889 | Klhl21 | kelch-like 21 [Source:MGI Symbol;Acc:MGI:1919288] | 94,81 | 11,7 |
| ENSMUSG00000021720 | 159,6620737 | 217,7 | 2,8 | 6,436142935 | 0,000201702 | Rnf180 | ring finger protein 180 [Source:MGI Symbol;Acc:MGI:1919066] | 91,38 | 1,6 |
| ENSMUSG00000059975 | 73,76647831 | 111,1 | 70,8 | 6,436599984 | 0,015403999 | Zfp74 | zinc finger protein 74 [Source:MGI Symbol;Acc:MGI:107784] | 73,02 | 3,5 |
| ENSMUSG00000051238 | 125,3929527 | 176,2 | 82,3 | 6,437820231 | 0,009302039 | Swsap1 | SWIM type zinc finger 7 associated protein 1 [Source:MGI Symbol;Acc:MGI:1914212] | 97,17 | 28,2 |
| ENSMUSG00000041303 | 39,03623823 | 60,6 | 0,9 | 6,439575877 | 0,005499898 | Gtf3c3 | general transcription factor IIIC, polypeptide 3 [Source:MGI Symbol;Acc:MGI:2138383] | 76,17 | 0,5 |
| ENSMUSG00000015889 | 141,3674736 | 225,5 | 68,4 | 6,441167995 | 0,001311838 | Lta4h | leukotriene A4 hydrolase [Source:MGI Symbol;Acc:MGI:96836] | 98,14 | 24,6 |
| ENSMUSG00000022664 | 354,0112876 | 484,7 | 9,5 | 6,44233274 | 0,000666709 | Slc35a5 | solute carrier family 35, member A5 [Source:MGI Symbol;Acc:MGI:1921352] | 95,73 | 1,4 |
| ENSMUSG00000036298 | 52,54808522 | 49,7 | 0,6 | 6,442936474 | 0,017048315 | Slc2a13 | solute carrier family 2 (facilitated glucose transporter), member 13 [Source:MGI Symbol;Acc:MGI:2146030 | 58,84 | 0,4 |
| ENSMUSG00000048490 | 77,9477043 | 125,2 | 58,9 | 6,443196766 | 0,005859943 | Nrip1 | nuclear receptor interacting protein 1 [Source:MGI Symbol;Acc:MGI:1315213] | 77,08 | 3,1 |
| ENSMUSG00000050840 | 88,450129 | 119,2 | 59,3 | 6,444205174 | 0,015819883 | Cdh20 | cadherin 20 [Source:MGI Symbol;Acc:MGI:1346069] | 92,31 | 6,4 |
| ENSMUSG00000018541 | 71,60472237 | 112,2 | 5,5 | 6,445520161 | 0,007785233 | Cwc25 | CWC25 spliceosome-associated protein [Source:MGI Symbol;Acc:MGI:1914730] | 65,71 | 1,7 |
| ENSMUSG00000003119 | 44,18077136 | 56,9 | 0,6 | 6,446102033 | 0,003979898 | Cdk12 | cyclin-dependent kinase 12 [Source:MGI Symbol;Acc:MGI:1098802] | 64,14 | 0,4 |
| ENSMUSG00000041471 | 43,74233368 | 71,2 | 1,0 | 6,446686708 | 0,008665422 | Fam35a | family with sequence similarity 35, member A [Source:MGI Symbol;Acc:MGI:1922948] | 85,45 | 0,7 |
| ENSMUSG00000063145 | 51,08459647 | 75,1 | 1,2 | 6,449539311 | 0,012054878 | Bbs5 | Bardet-Biedl syndrome 5 (human) [Source:MGI Symbol;Acc:MGI:1919819] | 98,50 | 1,2 |
| ENSMUSG00000024339 | 84,15582369 | 131,0 | 99,3 | 6,455178129 | 0,014620403 | Tap2 | transporter 2, ATP-binding cassette, sub-family B (MDR/TAP) [Source:MGI Symbol;Acc:MGI:98484] | 69,62 | 20,7 |
| ENSMUSG00000055371 | 127,4977259 | 160,7 | 12,1 | 6,45543775 | 0,007673692 | Stam2 | gnal transducing adaptor molecule (SH3 domain and ITAM motif) 2 [Source:MGI Symbol;Acc:MGI:192910 | 75,42 | 1,2 |
| ENSMUSG00000024424 | 61,84400698 | 99,5 | 2,0 | 6,456931034 | 0,002343294 | Ttc39c | tetratricopeptide repeat domain 39C [Source:MGI Symbol;Acc:MGI:1919997] | 81,09 | 0,1 |
| ENSMUSG00000058056 | 45,55918691 | 58,0 | 0,9 | 6,458221259 | 0,012906012 | Palld | palladin, cytoskeletal associated protein [Source:MGI Symbol;Acc:MGI:1919583] | 63,42 | 0,0 |
| ENSMUSG00000050730 | 131,2488181 | 186,3 | 2,6 | 6,458399117 | 0,001740859 | Arhgap42 | Rho GTPase activating protein 42 [Source:MGI Symbol;Acc:MGI:1918794] | 92,17 | 0,7 |
| ENSMUSG00000087497 | 223,055464 | 262,4 | 68,9 | 6,459545902 | 0,000213923 | 2810001G20Rik | RIKEN cDNA 2810001G20 gene [Source:MGI Symbol;Acc:MGI:1913706] | 86,72 | 22,7 |
| ENSMUSG00000073680 | 134,8540578 | 131,5 | 195,6 | 6,461193703 | 0,016226964 | Tmem88b | transmembrane protein 88B [Source:MGI Symbol;Acc:MGI:2444329] | 98,12 | 29,4 |
| ENSMUSG00000028005 | 70,56267963 | 104,5 | 2,4 | 6,46126746 | 0,009075917 | Gucy1b3 | guanylate cyclase 1, soluble, beta 3 [Source:MGI Symbol;Acc:MGI:1860604] | 96,34 | 0,8 |
| ENSMUSG00000042350 | 321,4016324 | 421,2 | 25,6 | 6,461955259 | 8,42939E-05 | Arel1 | apoptosis resistant E3 ubiquitin protein ligase 1 [Source:MGI Symbol;Acc:MGI:1915747] | 95,39 | 2,7 |
| ENSMUSG00000042167 | 109,4327552 | 141,4 | 97,8 | 6,464044404 | 0,008924977 | Papd4 | PAP associated domain containing 4 [Source:MGI Symbol;Acc:MGI:2140950] | 87,60 | 20,7 |
| ENSMUSG00000039830 | 101,9735353 | 125,3 | 18,8 | 6,46409405 | 0,017459821 | Olig2 | oligodendrocyte transcription factor 2 [Source:MGI Symbol;Acc:MGI:1355331] | 99,66 | 16,0 |
| ENSMUSG00000037788 | 136,6305111 | 141,4 | 187,6 | 6,464260247 | 0,018229935 | Vopp1 | vesicular, overexpressed in cancer, prosurvival protein 1 [Source:MGI Symbol;Acc:MGI:2141658] | 33,89 | 8,4 |
| ENSMUSG00000030499 | 25,0593841 | 51,6 | 0,6 | 6,464266896 | 0,013845306 | Kctd15 | potassium channel tetramerisation domain containing 15 [Source:MGI Symbol;Acc:MGI:2385276] | 80,66 | 0,0 |
| ENSMUSG00000002210 | 75,04130577 | 110,1 | 8,0 | 6,466893902 | 0,008561703 | Smg9 | g-9 homolog, nonsense mediated mRNA decay factor (C. elegans) [Source:MGI Symbol;Acc:MGI:19192 | 93,87 | 3,7 |
| ENSMUSG00000063063 | 150,1228169 | 150,9 | 9,1 | 6,468641418 | 0,009917731 | Ctnna2 | catenin (cadherin associated protein), alpha 2 [Source:MGI Symbol;Acc:MGI:88275] | 65,34 | 1,0 |
| ENSMUSG00000054459 | 178,2790421 | 333,2 | 4,8 | 6,469091133 | 0,000139785 | Vsnl1 | visinin-like 1 [Source:MGI Symbol;Acc:MGI:1349453] | 98,88 | 3,4 |

| | | | | | | | | | |
|---|---|---|---|---|---|---|---|---|---|
| ENSMUSG00000063895 | 77,44434272 | 81,2 | 3,0 | 6,46956319 | 0,014621969 | Nupl1 | nucleoporin like 1 [Source:MGI Symbol;Acc:MGI:1919094] | 95,41 | 0,2 |
| ENSMUSG00000032396 | 200,2904499 | 183,2 | 153,7 | 6,469882525 | 0,006374552 | Dis3l | DIS3 like exosome 3'-5' exoribonuclease [Source:MGI Symbol;Acc:MGI:2143272] | 78,76 | 18,9 |
| ENSMUSG00000037679 | 33,55838803 | 65,9 | 0,9 | 6,470254816 | 0,010206514 | Inf2 | inverted formin, FH2 and WH2 domain containing [Source:MGI Symbol;Acc:MGI:1917685] | 70,78 | 0,5 |
| ENSMUSG00000032449 | 105,0391754 | 127,0 | 11,0 | 6,471588517 | 0,009556603 | Slc25a36 | solute carrier family 25, member 36 [Source:MGI Symbol;Acc:MGI:1924909] | 48,26 | 1,1 |
| ENSMUSG00000022131 | 160,0867908 | 182,4 | 14,8 | 6,473490478 | 0,001279007 | Gpr180 | G protein-coupled receptor 180 [Source:MGI Symbol;Acc:MGI:1930949] | 74,27 | 4,7 |
| ENSMUSG00000028431 | 73,15663138 | 81,0 | 2,0 | 6,474513709 | 0,009369036 | Ikbkap | appa light polypeptide enhancer in B cells, kinase complex-associated protein [Source:MGI Symbol;Acc:M | 76,40 | 0,4 |
| ENSMUSG00000025169 | 104,1855628 | 116,2 | 3,0 | 6,47850936 | 0,004465947 | Ogfod3 | -oxoglutarate and iron-dependent oxygenase domain containing 3 [Source:MGI Symbol;Acc:MGI:191342 | 97,66 | 3,0 |
| ENSMUSG00000044348 | 219,11019 | 241,1 | 121,1 | 6,479245089 | 0,001506164 | Slc25a53 | solute carrier family 25, member 53 [Source:MGI Symbol;Acc:MGI:1914312] | 91,87 | 23,6 |
| ENSMUSG00000036206 | 29,67645827 | 63,3 | 0,9 | 6,48184649 | 0,009078025 | Sh3bp4 | SH3-domain binding protein 4 [Source:MGI Symbol;Acc:MGI:2138297] | 77,38 | 0,4 |
| ENSMUSG00000020238 | 123,7430904 | 214,6 | 5,5 | 6,485016916 | 2,63496E-05 | Ncln | nicalin homolog (zebrafish) [Source:MGI Symbol;Acc:MGI:1926081] | 93,01 | 2,6 |
| ENSMUSG00000022510 | 47,25895432 | 94,0 | 1,8 | 6,487723924 | 0,003647101 | Trp63 | transformation related protein 63 [Source:MGI Symbol;Acc:MGI:1330810] | 40,79 | 0,5 |
| ENSMUSG00000031409 | 47,40961619 | 56,7 | 1,0 | 6,489030648 | 0,014153278 | Tceal6 | transcription elongation factor A (SII)-like 6 [Source:MGI Symbol;Acc:MGI:1923939] | 88,03 | 1,9 |
| ENSMUSG00000026896 | 57,68411563 | 65,5 | 1,2 | 6,489969328 | 0,02528919 | Ifih1 | interferon induced with helicase C domain 1 [Source:MGI Symbol;Acc:MGI:1918836] | 78,71 | 0,5 |
| ENSMUSG00000006464 | 81,85510432 | 132,0 | 5,5 | 6,490031258 | 0,003947394 | Bbs1 | Bardet-Biedl syndrome 1 (human) [Source:MGI Symbol;Acc:MGI:1277215] | 92,65 | 1,0 |
| ENSMUSG00000042097 | 114,4764127 | 139,0 | 95,2 | 6,491216602 | 0,018591859 | Zfp239 | zinc finger protein 239 [Source:MGI Symbol;Acc:MGI:1306812] | 83,58 | 10,6 |
| ENSMUSG00000073590 | 25,61525931 | 63,6 | 0,9 | 6,492456002 | 0,006739541 | 3222401L13Rik | RIKEN cDNA 3222401L13 gene [Source:MGI Symbol;Acc:MGI:2444186] | #DIV/0! | #DIV/0! |
| ENSMUSG00000027285 | 380,2655836 | 551,0 | 65,7 | 6,492643241 | 0,000762765 | Haus2 | HAUS augmin-like complex, subunit 2 [Source:MGI Symbol;Acc:MGI:1913546] | 98,17 | 18,4 |
| ENSMUSG00000036941 | 87,76150009 | 106,1 | 2,4 | 6,49288827 | 0,008713505 | Elac1 | elaC ribonuclease Z 1 [Source:MGI Symbol;Acc:MGI:1890495] | 93,37 | 0,8 |
| ENSMUSG00000026709 | 192,5811394 | 150,4 | 270,4 | 6,497072941 | 0,00764456 | Dars2 | aspartyl-tRNA synthetase 2 (mitochondrial) [Source:MGI Symbol;Acc:MGI:2442510] | 92,21 | 25,9 |
| ENSMUSG00000034394 | 23,93695188 | 66,1 | 1,0 | 6,499300792 | 0,005768982 | Lif | leukemia inhibitory factor [Source:MGI Symbol;Acc:MGI:96787] | 86,23 | 0,6 |
| ENSMUSG00000033436 | 63,30709684 | 112,2 | 1,8 | 6,503506481 | 0,000870187 | Armcx2 | armadillo repeat containing, X-linked 2 [Source:MGI Symbol;Acc:MGI:1914666] | 85,41 | 0,8 |
| ENSMUSG00000056019 | 57,84429021 | 85,5 | 2,0 | 6,503615176 | 0,008345158 | Zfp709 | zinc finger protein 709 [Source:MGI Symbol;Acc:MGI:2384299] | 54,84 | 0,2 |
| ENSMUSG00000025551 | 29,78353431 | 64,8 | 0,9 | 6,508951921 | 0,004205779 | Fgf14 | fibroblast growth factor 14 [Source:MGI Symbol;Acc:MGI:109189] | 78,27 | 0,6 |
| ENSMUSG00000001942 | 115,3496233 | 123,1 | 12,9 | 6,509772887 | 0,015668098 | Siae | sialic acid acetylesterase [Source:MGI Symbol;Acc:MGI:104803] | 98,39 | 3,2 |
| ENSMUSG00000034807 | 63,59274615 | 129,7 | 20,3 | 6,511515432 | 0,009802604 | Colgalt1 | collagen beta(1-O)galactosyltransferase 1 [Source:MGI Symbol;Acc:MGI:1924348] | 94,81 | 7,5 |
| ENSMUSG00000028035 | 362,6037309 | 527,4 | 124,8 | 6,513107715 | 0,007303788 | Dnajb4 | DnaJ heat shock protein family (Hsp40) member B4 [Source:MGI Symbol;Acc:MGI:1914285] | 91,57 | 25,3 |
| ENSMUSG00000040688 | 98,23063736 | 159,1 | 60,1 | 6,513769304 | 0,006896685 | Tbl3 | transducin (beta)-like 3 [Source:MGI Symbol;Acc:MGI:2384863] | 72,54 | 12,5 |
| ENSMUSG00000028133 | 34,44469326 | 68,3 | 1,0 | 6,515379143 | 0,00513118 | Rwdd3 | RWD domain containing 3 [Source:MGI Symbol;Acc:MGI:1920420] | 76,29 | 1,1 |
| ENSMUSG00000010307 | 131,7464716 | 163,8 | 46,9 | 6,515535313 | 0,022056508 | Tmem86a | transmembrane protein 86A [Source:MGI Symbol;Acc:MGI:1915143] | 98,56 | 18,5 |
| ENSMUSG00000038697 | 132,2824101 | 124,0 | 151,7 | 6,517399113 | 0,015231624 | Taf5l | RNA polymerase II, p300/CBP-associated factor (PCAF)-associated factor [Source:MGI Symbol;Acc:MGI: | 82,48 | 25,9 |
| ENSMUSG00000038844 | 125,7346468 | 138,7 | 1,5 | 6,517606976 | 0,002611917 | Kif16b | kinesin family member 16B [Source:MGI Symbol;Acc:MGI:1098240] | 86,04 | 0,6 |
| ENSMUSG00000026584 | 97,17067394 | 107,6 | 101,8 | 6,520613175 | 0,018579442 | Scyl3 | SCY1-like 3 (S. cerevisiae) [Source:MGI Symbol;Acc:MGI:1921385] | 89,93 | 14,2 |
| ENSMUSG00000089872 | 134,8113007 | 106,8 | 106,8 | 6,522552635 | 0,017067502 | Rps6kc1 | ribosomal protein S6 kinase polypeptide 1 [Source:MGI Symbol;Acc:MGI:2443419] | 98,39 | 25,2 |
| ENSMUSG00000090208 | 106,9400704 | 158,3 | 24,9 | 6,522704225 | 0,006988159 | Gm15851 | predicted gene 15851 [Source:MGI Symbol;Acc:MGI:3801734] | 53,02 | 16,0 |
| ENSMUSG00000097348 | 71,22789325 | 96,2 | 4,0 | 6,524880928 | 0,010341486 | Rmst | rhabdomyosarcoma 2 associated transcript (non-coding RNA) [Source:MGI Symbol;Acc:MGI:1099806] | 100,00 | 0,0 |
| ENSMUSG00000019853 | 108,8635999 | 120,1 | 166,6 | 6,524914817 | 0,016571747 | Hebp2 | heme binding protein 2 [Source:MGI Symbol;Acc:MGI:1860084] | 98,50 | 32,1 |
| ENSMUSG00000055723 | 143,4159753 | 136,9 | 34,9 | 6,525575338 | 0,010077387 | Rras2 | related RAS viral (r-ras) oncogene homolog 2 [Source:MGI Symbol;Acc:MGI:1914172] | 99,52 | 4,2 |
| ENSMUSG00000018339 | 82,55580702 | 151,4 | 27,8 | 6,527717586 | 0,010930314 | Gpx3 | glutathione peroxidase 3 [Source:MGI Symbol;Acc:MGI:105102] | 83,47 | 22,1 |
| ENSMUSG00000041117 | 37,02614334 | 62,9 | 0,9 | 6,527854124 | 0,005859943 | Ccdc8 | coiled-coil domain containing 8 [Source:MGI Symbol;Acc:MGI:3612184] | 71,23 | 0,2 |
| ENSMUSG00000030509 | 112,2715054 | 116,9 | 9,2 | 6,529024919 | 0,015231624 | Asb7 | ankyrin repeat and SOCS box-containing 7 [Source:MGI Symbol;Acc:MGI:2152835] | 92,13 | 1,1 |
| ENSMUSG00000027167 | 226,4213286 | 394,2 | 16,7 | 6,530336827 | 0,007785817 | Elp4 | elongator acetyltransferase complex subunit 4 [Source:MGI Symbol;Acc:MGI:1925016] | 96,93 | 4,0 |
| ENSMUSG00000031360 | 79,85456512 | 93,1 | 2,0 | 6,531441578 | 0,005503191 | Ctps2 | cytidine 5'-triphosphate synthase 2 [Source:MGI Symbol;Acc:MGI:1933185] | 74,19 | 0,6 |
| ENSMUSG00000029466 | 127,8429362 | 137,4 | 40,9 | 6,531527594 | 0,007099625 | Anapc7 | anaphase promoting complex subunit 7 [Source:MGI Symbol;Acc:MGI:1929711] | 57,34 | 5,2 |
| ENSMUSG00000082229 | 44,3390591 | 67,0 | 0,9 | 6,531827687 | 0,010167511 | Nap1l2 | nucleosome assembly protein 1-like 2 [Source:MGI Symbol;Acc:MGI:106654] | 96,01 | 0,6 |
| ENSMUSG00000016253 | 239,7159207 | 351,9 | 4,9 | 6,532264848 | 0,000738207 | Nelfcd | negative elongation factor complex member C/D, Th1l [Source:MGI Symbol;Acc:MGI:1926424] | 97,61 | 0,2 |
| ENSMUSG00000029427 | 94,26289542 | 140,9 | 4,6 | 6,53230938 | 0,0022357 | Zcchc8 | zinc finger, CCHC domain containing 8 [Source:MGI Symbol;Acc:MGI:1917900] | 78,53 | 1,0 |
| ENSMUSG00000029047 | 84,29363499 | 108,8 | 200,6 | 6,533285835 | 0,013377415 | Pex10 | peroxisomal biogenesis factor 10 [Source:MGI Symbol;Acc:MGI:2684988] | 96,13 | 26,5 |
| ENSMUSG00000023892 | 149,6786736 | 120,1 | 219,5 | 6,533808917 | 0,012376795 | Zfp51 | zinc finger protein 51 [Source:MGI Symbol;Acc:MGI:99198] | 98,40 | 31,7 |
| ENSMUSG00000099034 | 35,84928058 | 55,3 | 0,6 | 6,534992662 | 0,011400919 | 2810039B14Rik | RIKEN cDNA 2810039B14 gene [Source:MGI Symbol;Acc:MGI:1919915] | #DIV/0! | #DIV/0! |
| ENSMUSG00000057265 | 102,2439062 | 162,9 | 39,9 | 6,536742628 | 0,003674644 | Bbof1 | basal body orientation factor 1 [Source:MGI Symbol;Acc:MGI:1920123] | 72,76 | 7,9 |
| ENSMUSG00000032777 | 92,93030223 | 130,4 | 5,0 | 6,539526595 | 0,003284776 | Gtf3c1 | general transcription factor III C 1 [Source:MGI Symbol;Acc:MGI:107887] | 85,68 | 0,4 |
| ENSMUSG00000040146 | 34,70096655 | 58,1 | 1,0 | 6,539740544 | 0,01696342 | Rgl3 | ral guanine nucleotide dissociation stimulator-like 3 [Source:MGI Symbol;Acc:MGI:1918996] | 93,14 | 0,9 |
| ENSMUSG00000022895 | 60,22536303 | 101,6 | 2,8 | 6,541185813 | 0,007785817 | Ets2 | E26 avian leukemia oncogene 2, 3' domain [Source:MGI Symbol;Acc:MGI:95456] | 86,75 | 0,9 |
| ENSMUSG00000029330 | 90,90042537 | 92,6 | 2,0 | 6,542657648 | 0,008349516 | Cds1 | CDP-diacylglycerol synthase 1 [Source:MGI Symbol;Acc:MGI:1921846] | 87,14 | 0,6 |
| ENSMUSG00000018433 | 182,8331204 | 132,6 | 8,5 | 6,544841052 | 0,013227625 | Nol11 | nucleolar protein 11 [Source:MGI Symbol;Acc:MGI:1916229] | 94,80 | 3,0 |
| ENSMUSG00000038042 | 102,5734355 | 111,9 | 201,6 | 6,547708689 | 0,016189098 | Ptpdc1 | protein tyrosine phosphatase domain containing 1 [Source:MGI Symbol;Acc:MGI:2145430] | 94,50 | 28,0 |
| ENSMUSG00000034467 | 94,78422504 | 135,1 | 15,0 | 6,548791728 | 0,012288115 | Dynlrb2 | dynein light chain roadblock-type 2 [Source:MGI Symbol;Acc:MGI:1922715] | 100,00 | 27,8 |
| ENSMUSG00000030471 | 105,3160008 | 87,7 | 3,0 | 6,551053428 | 0,015059 | Zdhhc13 | zinc finger, DHHC domain containing 13 [Source:MGI Symbol;Acc:MGI:1919227] | 97,63 | 1,7 |
| ENSMUSG00000022297 | 43,47488778 | 99,7 | 1,2 | 6,551993814 | 0,001366453 | Fzd6 | frizzled class receptor 6 [Source:MGI Symbol;Acc:MGI:108474] | 96,65 | 0,6 |
| ENSMUSG00000089815 | 65,46668913 | 106,7 | 13,9 | 6,553426372 | 0,017999606 | Gm5083 | predicted gene 5083 [Source:MGI Symbol;Acc:MGI:3779454] | #DIV/0! | #DIV/0! |
| ENSMUSG00000026199 | 133,6331663 | 98,4 | 105,8 | 6,55374219 | 0,026228876 | Ankzf1 | ankyrin repeat and zinc finger domain containing 1 [Source:MGI Symbol;Acc:MGI:1098746] | 92,98 | 22,1 |
| ENSMUSG00000057895 | 62,60843492 | 115,0 | 3,0 | 6,557008514 | 0,005369631 | Zfp105 | zinc finger protein 105 [Source:MGI Symbol;Acc:MGI:1277119] | 96,04 | 2,1 |
| ENSMUSG00000028578 | 116,2849716 | 181,5 | 146,5 | 6,55803516 | 0,008948721 | Caap1 | caspase activity and apoptosis inhibitor 1 [Source:MGI Symbol;Acc:MGI:1915020] | 99,81 | 32,0 |
| ENSMUSG00000016024 | 130,5380465 | 134,8 | 43,4 | 6,56112751 | 0,017398596 | Lbp | lipopolysaccharide binding protein [Source:MGI Symbol;Acc:MGI:1098776] | 57,88 | 13,1 |
| ENSMUSG00000039179 | 35,03499811 | 45,5 | 0,6 | 6,564828595 | 0,035534808 | Tekt5 | tektin 5 [Source:MGI Symbol;Acc:MGI:1917676] | 89,36 | 1,4 |
| ENSMUSG00000057147 | 167,709671 | 134,2 | 144,7 | 6,567144774 | 0,016226964 | Dph6 | diphthamine biosynthesis 6 [Source:MGI Symbol;Acc:MGI:1913882] | 96,36 | 27,4 |
| ENSMUSG00000021913 | 86,7161646 | 162,2 | 7,3 | 6,56921947 | 0,006018371 | Ogdhl | oxoglutarate dehydrogenase-like [Source:MGI Symbol;Acc:MGI:3616088] | 76,31 | 4,8 |
| ENSMUSG00000074896 | 80,38566137 | 132,6 | 59,3 | 6,569663176 | 0,01543911 | Ifit3 | interferon-induced protein with tetratricopeptide repeats 3 [Source:MGI Symbol;Acc:MGI:1101055] | 81,32 | 22,0 |
| ENSMUSG00000033931 | 121,9109512 | 156,8 | 5,4 | 6,56974714 | 0,004023292 | Rbm34 | RNA binding motif protein 34 [Source:MGI Symbol;Acc:MGI:1098653] | 86,06 | 2,7 |
| ENSMUSG00000022175 | 371,4231996 | 528,5 | 56,4 | 6,572750112 | 0,00012881 | Lrp10 | low-density lipoprotein receptor-related protein 10 [Source:MGI Symbol;Acc:MGI:1929480] | 75,30 | 15,5 |
| ENSMUSG00000030801 | 120,1463258 | 170,5 | 7,0 | 6,573557755 | 0,000907964 | Kat8 | K(lysine) acetyltransferase 8 [Source:MGI Symbol;Acc:MGI:1915023] | 79,11 | 0,0 |
| ENSMUSG00000020681 | 176,5941818 | 185,8 | 1,9 | 6,575526159 | 0,005018328 | Ace | angiotensin I converting enzyme (peptidyl-dipeptidase A) 1 [Source:MGI Symbol;Acc:MGI:87874] | 87,35 | 0,8 |
| ENSMUSG00000027222 | 107,302236 | 181,2 | 41,8 | 6,576602733 | 0,007221114 | Pex16 | peroxisomal biogenesis factor 16 [Source:MGI Symbol;Acc:MGI:1338829] | 95,09 | 24,2 |
| ENSMUSG00000054484 | 134,4832387 | 143,6 | 76,8 | 6,576932657 | 0,006235071 | Tmem62 | transmembrane protein 62 [Source:MGI Symbol;Acc:MGI:2139461] | 98,16 | 11,1 |
| ENSMUSG00000034706 | 32,76027343 | 73,2 | 1,0 | 6,579364135 | 0,004329646 | Dnaic2 | dynein, axonemal, intermediate chain 2 [Source:MGI Symbol;Acc:MGI:2685574] | 81,54 | 0,8 |
| ENSMUSG00000021097 | 122,4111086 | 198,8 | 79,9 | 6,580845414 | 0,00903868 | Clmn | calmin [Source:MGI Symbol;Acc:MGI:2136957] | 71,54 | 7,3 |
| ENSMUSG00000021482 | 148,1487213 | 200,1 | 35,1 | 6,581063285 | 0,004402332 | Aaed1 | AhpC/TSA antioxidant enzyme domain containing 1 [Source:MGI Symbol;Acc:MGI:1913379] | 100,00 | 26,1 |
| ENSMUSG00000071757 | 125,9850305 | 190,0 | 2,2 | 6,585126549 | 0,003090135 | Zhx2 | zinc fingers and homeoboxes 2 [Source:MGI Symbol;Acc:MGI:2683087] | 97,50 | 1,1 |
| ENSMUSG00000029860 | 86,87171543 | 150,2 | 66,9 | 6,592300442 | 0,006314703 | Zyx | zyxin [Source:MGI Symbol;Acc:MGI:103072] | 57,28 | 12,3 |
| ENSMUSG00000076431 | 111,6093524 | 198,8 | 11,5 | 6,592418516 | 0,002985684 | Sox4 | SRY (sex determining region Y)-box 4 [Source:MGI Symbol;Acc:MGI:98366] | 94,22 | 4,0 |
| ENSMUSG00000021338 | 258,143183 | 371,0 | 88,4 | 6,596122394 | 0,004016612 | Lrrc16a | leucine rich repeat containing 16A [Source:MGI Symbol;Acc:MGI:1915982] | 91,33 | 19,6 |
| ENSMUSG00000002980 | 71,22239424 | 108,6 | 1,8 | 6,599494359 | 0,014975476 | Bcam | basal cell adhesion molecule [Source:MGI Symbol;Acc:MGI:1929940] | 85,48 | 1,8 |
| ENSMUSG00000028992 | 82,14450854 | 117,0 | 50,9 | 6,603718719 | 0,014231804 | Nmnat1 | nicotinamide nucleotide adenylyltransferase 1 [Source:MGI Symbol;Acc:MGI:1913704] | 73,56 | 9,7 |
| ENSMUSG00000018599 | 61,8365863 | 106,9 | 6,0 | 6,607585959 | 0,009447034 | Mief2 | mitochondrial elongation factor 2 [Source:MGI Symbol;Acc:MGI:2144199] | 97,92 | 2,5 |
| ENSMUSG00000029843 | 235,0880039 | 230,3 | 102,9 | 6,609622582 | 0,007496462 | Slc13a4 | solute carrier family 13 (sodium/sulfate symporters), member 4 [Source:MGI Symbol;Acc:MGI:2442367] | 99,20 | 22,0 |
| ENSMUSG00000021559 | 149,9749339 | 114,1 | 74,8 | 6,610130131 | 0,021387317 | Dapk1 | death associated protein kinase 1 [Source:MGI Symbol;Acc:MGI:1916885] | 94,01 | 9,5 |
| ENSMUSG00000046798 | 120,4074789 | 174,7 | 43,4 | 6,61556478 | 0,013574976 | Cldn12 | claudin 12 [Source:MGI Symbol;Acc:MGI:1929288] | 91,76 | 8,0 |
| ENSMUSG00000038417 | 145,2877431 | 215,4 | 125,9 | 6,615682541 | 0,004455351 | Fig4 | FIG4 phosphoinositide 5-phosphatase [Source:MGI Symbol;Acc:MGI:2143585] | 99,98 | 30,4 |
| ENSMUSG00000018752 | 100,6013299 | 166,5 | 2,4 | 6,620339256 | 0,000393615 | Tnfsfm13 | or necrosis factor (ligand) superfamily, membrane-bound member 13 [Source:MGI Symbol;Acc:MGI:3845 | 86,86 | 0,6 |
| ENSMUSG00000001773 | 116,9998801 | 201,1 | 179,7 | 6,622176823 | 0,005100924 | Folh1 | folate hydrolase 1 [Source:MGI Symbol;Acc:MGI:1858193] | 96,83 | 28,2 |
| ENSMUSG00000029059 | 167,795611 | 184,7 | 172,6 | 6,622213007 | 0,004674463 | Fam213b | family with sequence similarity 213, member B [Source:MGI Symbol;Acc:MGI:1913719] | 96,78 | 31,5 |
| ENSMUSG00000062198 | 90,34265058 | 105,1 | 135,7 | 6,622218459 | 0,024884574 | 2700097O09Rik | RIKEN cDNA 2700097O09 gene [Source:MGI Symbol;Acc:MGI:1919908] | 99,28 | 14,5 |
| ENSMUSG00000042428 | 63,89022253 | 100,3 | 5,0 | 6,627006593 | 0,014848232 | Mgat3 | mannoside acetylglucosaminyltransferase 3 [Source:MGI Symbol;Acc:MGI:104532] | 95,02 | 0,9 |
| ENSMUSG00000022911 | 45,9432592 | 65,5 | 1,0 | 6,629023821 | 0,010372707 | Arl13b | ADP-ribosylation factor-like 13B [Source:MGI Symbol;Acc:MGI:1915396] | 86,12 | 0,4 |
| ENSMUSG00000032478 | 125,0658102 | 179,4 | 17,6 | 6,629690532 | 0,003204932 | Nme6 | NME/NM23 nucleoside diphosphate kinase 6 [Source:MGI Symbol;Acc:MGI:1861676] | 92,45 | 18,7 |
| ENSMUSG00000031521 | 105,0352559 | 145,4 | 73,9 | 6,634670791 | 0,011353576 | Aga | aspartylglucosaminidase [Source:MGI Symbol;Acc:MGI:104873] | 95,97 | 24,7 |
| ENSMUSG00000079083 | 80,33562106 | 83,0 | 1,8 | 6,635714031 | 0,01629438 | Jrkl | Jrk-like [Source:MGI Symbol;Acc:MGI:1924782] | 93,38 | 0,5 |
| ENSMUSG00000021646 | 257,9132942 | 513,6 | 75,9 | 6,635843429 | 0,00489173 | Mccc2 | methylcrotonoyl-Coenzyme A carboxylase 2 (beta) [Source:MGI Symbol;Acc:MGI:1925288] | 100,00 | 31,3 |
| ENSMUSG00000050697 | 196,1444957 | 181,9 | 78,8 | 6,638574194 | 0,001689804 | Prkaa1 | protein kinase, AMP-activated, alpha 1 catalytic subunit [Source:MGI Symbol;Acc:MGI:2145955] | 96,43 | 4,3 |
| ENSMUSG00000020019 | 95,26445737 | 131,2 | 73,8 | 6,642235078 | 0,01918874 | Ntn4 | netrin 4 [Source:MGI Symbol;Acc:MGI:1888978] | 94,75 | 22,2 |
| ENSMUSG00000055972 | 124,2881006 | 155,7 | 36,9 | 6,642657204 | 0,0136442 | 2810407A14Rik | RIKEN cDNA 2810407A14 gene [Source:MGI Symbol;Acc:MGI:1917461] | #DIV/0! | #DIV/0! |
| ENSMUSG00000033106 | 86,64838572 | 161,9 | 29,9 | 6,648658267 | 0,003911784 | Slc7a6os | solute carrier family 7, member 6 opposite strand [Source:MGI Symbol;Acc:MGI:1916951] | 98,01 | 13,9 |
| ENSMUSG00000021013 | 141,8845116 | 177,2 | 98,0 | 6,648770685 | 0,00778335 | Ttc8 | tetratricopeptide repeat domain 8 [Source:MGI Symbol;Acc:MGI:1923510] | 92,46 | 28,3 |
| ENSMUSG00000038365 | 124,3624099 | 218,8 | 2,8 | 6,65236778 | 0,000805606 | Fbxo25 | F-box protein 25 [Source:MGI Symbol;Acc:MGI:1914072] | 99,88 | 2,8 |
| ENSMUSG00000010362 | 252,3923203 | 371,1 | 14,6 | 6,654873314 | 0,000935031 | Rdm1 | RAD52 motif 1 [Source:MGI Symbol;Acc:MGI:1913849] | 99,67 | 9,2 |
| ENSMUSG00000073062 | 35,2194403 | 60,1 | 0,9 | 6,655086792 | 0,015968705 | Zxdb | zinc finger, X-linked, duplicated B [Source:MGI Symbol;Acc:MGI:3694898] | 45,54 | 0,4 |
| ENSMUSG00000031955 | 97,53844137 | 143,6 | 31,5 | 6,655868793 | 0,0223201 | Bcar1 | breast cancer anti-estrogen resistance 1 [Source:MGI Symbol;Acc:MGI:108091] | 99,10 | 10,6 |
| ENSMUSG00000040297 | 63,57924704 | 74,0 | 1,0 | 6,6574724 | 0,009206088 | Suco | SUN domain containing ossification factor [Source:MGI Symbol;Acc:MGI:2138346] | 70,73 | 0,3 |
| ENSMUSG00000025321 | 558,6375429 | 666,3 | 17,1 | 6,659339955 | 0,004625348 | Itgb8 | integrin beta 8 [Source:MGI Symbol;Acc:MGI:1338035] | 99,87 | 2,6 |
| ENSMUSG00000063646 | 233,6668735 | 409,1 | 112,4 | 6,661091545 | 0,009471772 | Jakmip1 | janus kinase and microtubule interacting protein 1 [Source:MGI Symbol;Acc:MGI:1923321] | 76,91 | 14,8 |
| ENSMUSG00000020593 | 174,5850861 | 204,4 | 33,3 | 6,66133067 | 0,0023893 | Lpin1 | lipin 1 [Source:MGI Symbol;Acc:MGI:1891340] | 93,36 | 5,5 |
| ENSMUSG00000023868 | 58,26841188 | 65,1 | 0,9 | 6,661358351 | 0,024112304 | Pde10a | phosphodiesterase 10A [Source:MGI Symbol;Acc:MGI:1345143] | 43,68 | 0,2 |
| ENSMUSG00000000776 | 96,91785751 | 154,1 | 58,2 | 6,666003061 | 0,010159129 | Polr3d | polymerase (RNA) III (DNA directed) polypeptide D [Source:MGI Symbol;Acc:MGI:1914315] | 97,50 | 24,9 |
| ENSMUSG00000022552 | 204,9800685 | 238,2 | 101,7 | 6,671377047 | 0,001966172 | Sharpin | SHANK-associated RH domain interacting protein [Source:MGI Symbol;Acc:MGI:1913331] | 98,24 | 29,1 |
| ENSMUSG00000046556 | 51,05402998 | 94,3 | 1,8 | 6,673510327 | 0,007871312 | Zfp319 | zinc finger protein 319 [Source:MGI Symbol;Acc:MGI:1890618] | 90,24 | 1,2 |
| ENSMUSG00000025777 | 90,90566322 | 106,5 | 4,0 | 6,677546427 | 0,008306247 | Gdap1 | ganglioside-induced differentiation-associated-protein 1 [Source:MGI Symbol;Acc:MGI:1338002] | 95,91 | 0,6 |
| ENSMUSG00000033883 | 111,081798 | 125,3 | 2,8 | 6,677643516 | 0,005065683 | D3Ertd254e | DNA segment, Chr 3, ERATO Doi 254, expressed [Source:MGI Symbol;Acc:MGI:1098769] | 82,99 | 0,5 |
| ENSMUSG00000028189 | 119,1932219 | 160,9 | 17,6 | 6,678251561 | 0,014295743 | Ctbs | chitobiase, di-N-acetyl- [Source:MGI Symbol;Acc:MGI:1921495] | 97,96 | 1,8 |
| ENSMUSG00000068196 | 76,68348641 | 123,9 | 0,6 | 6,680194174 | 0,004174921 | Col8a1 | collagen, type VIII, alpha 1 [Source:MGI Symbol;Acc:MGI:88463] | 74,15 | 0,5 |
| ENSMUSG00000022801 | 118,6822383 | 186,1 | 2,5 | 6,68393612 | 0,004232404 | Lrch3 | cine-rich repeats and calponin homology (CH) domain containing 3 [Source:MGI Symbol;Acc:MGI:19173 | 94,79 | 0,7 |
| ENSMUSG00000028567 | 97,87174652 | 135,8 | 48,4 | 6,684204193 | 0,015071241 | Txndc12 | thioredoxin domain containing 12 (endoplasmic reticulum) [Source:MGI Symbol;Acc:MGI:1913323] | 96,76 | 25,8 |
| ENSMUSG00000025474 | 142,444951 | 208,7 | 13,9 | 6,684776101 | 0,001277999 | Tubgcp2 | tubulin, gamma complex associated protein 2 [Source:MGI Symbol;Acc:MGI:1921487] | 92,29 | 6,6 |
| ENSMUSG00000020623 | 34,76818242 | 63,4 | 0,9 | 6,684912125 | 0,012198906 | Map2k6 | mitogen-activated protein kinase kinase 6 [Source:MGI Symbol;Acc:MGI:1346870] | 67,69 | 0,5 |
| ENSMUSG00000036867 | 38,4160038 | 64,8 | 0,9 | 6,686017548 | 0,012834265 | Smad6 | SMAD family member 6 [Source:MGI Symbol;Acc:MGI:1336883] | 91,53 | 0,7 |
| ENSMUSG00000028063 | 141,0098416 | 126,7 | 39,7 | 6,688159356 | 0,012622114 | Lmna | lamin A [Source:MGI Symbol;Acc:MGI:96794] | 84,22 | 12,8 |
| ENSMUSG00000023963 | 58,682583 | 56,9 | 0,6 | 6,692413864 | 0,023732685 | Cyp39a1 | cytochrome P450, family 39, subfamily a, polypeptide 1 [Source:MGI Symbol;Acc:MGI:1927096] | 76,17 | 0,8 |
| ENSMUSG00000024182 | 185,3167322 | 180,7 | 260,8 | 6,693714752 | 0,009954247 | Axin1 | axin 1 [Source:MGI Symbol;Acc:MGI:1096327] | 98,87 | 31,8 |
| ENSMUSG00000017802 | 40,33931823 | 77,3 | 1,0 | 6,694907244 | 0,003406984 | Fam134c | family with sequence similarity 134, member C [Source:MGI Symbol;Acc:MGI:1915248] | 88,78 | 0,7 |
| ENSMUSG00000035629 | 206,3809616 | 212,3 | 2,6 | 6,695631234 | 0,00030966 | Rubcn | omain and cysteine-rich domain containing, Beclin 1-interacting protein [Source:MGI Symbol;Acc:MGI:19 | 96,80 | 0,7 |
| ENSMUSG00000047141 | 159,5043687 | 79,9 | 1,2 | 6,699657363 | 0,02211382 | Zfp654 | zinc finger protein 654 [Source:MGI Symbol;Acc:MGI:1919270] | 79,74 | 0,3 |
| ENSMUSG00000021501 | 111,6539261 | 168,2 | 7,4 | 6,700632279 | 0,003702984 | Caml | calcium modulating ligand [Source:MGI Symbol;Acc:MGI:104728] | 99,83 | 7,6 |
| ENSMUSG00000042642 | 296,7939571 | 335,0 | 55,5 | 6,700636146 | 7,54388E-05 | Flad1 | flavin adenine dinucleotide synthetase 1 [Source:MGI Symbol;Acc:MGI:2443030] | 91,93 | 12,3 |
| ENSMUSG00000048355 | 100,2338008 | 129,1 | 74,8 | 6,702900384 | 0,014621969 | Arxes1 | adipocyte-related X-chromosome expressed sequence 1 [Source:MGI Symbol;Acc:MGI:1923469] | 67,55 | 20,5 |
| ENSMUSG00000040389 | 106,6021159 | 165,0 | 1,8 | 6,704671279 | 2,11115E-05 | Wdr47 | WD repeat domain 47 [Source:MGI Symbol;Acc:MGI:2139593] | 96,40 | 1,2 |
| ENSMUSG00000029175 | 153,2139204 | 219,8 | 80,4 | 6,70744272 | 0,005081605 | Slc35f6 | solute carrier family 35, member F6 [Source:MGI Symbol;Acc:MGI:1922169] | 86,03 | 19,8 |
| ENSMUSG00000053390 | 59,70937398 | 74,3 | 1,2 | 6,707637366 | 0,028064963 | Zfp952 | zinc finger protein 952 [Source:MGI Symbol;Acc:MGI:2441928] | 89,20 | 0,9 |
| ENSMUSG00000031665 | 89,88576916 | 158,0 | 1,5 | 6,710891832 | 0,000105076 | Sall1 | sal-like 1 (Drosophila) [Source:MGI Symbol;Acc:MGI:1889585] | 94,76 | 0,9 |
| ENSMUSG00000049929 | 87,18869358 | 127,1 | 86,8 | 6,711771619 | 0,017459821 | Lpar4 | lysophosphatidic acid receptor 4 [Source:MGI Symbol;Acc:MGI:1925384] | 85,55 | 15,5 |
| ENSMUSG00000031371 | 113,1188883 | 188,5 | 47,1 | 6,711911 | 0,005385528 | Haus7 | HAUS augmin-like complex, subunit 7 [Source:MGI Symbol;Acc:MGI:1920988] | 99,61 | 26,6 |
| ENSMUSG00000028016 | 132,3584258 | 107,0 | 71,8 | 6,712078328 | 0,026432656 | Ints12 | integrator complex subunit 12 [Source:MGI Symbol;Acc:MGI:1919043] | 94,26 | 11,7 |
| ENSMUSG00000029090 | 177,963034 | 325,4 | 4,0 | 6,712397463 | 0,000453513 | Adgra3 | adhesion G protein-coupled receptor A3 [Source:MGI Symbol;Acc:MGI:1917943] | 99,27 | 2,0 |
| ENSMUSG00000041992 | 73,21680915 | 65,3 | 0,6 | 6,716463901 | 0,003250126 | Rapgef5 | Rap guanine nucleotide exchange factor (GEF) 5 [Source:MGI Symbol;Acc:MGI:2444365] | 60,73 | 0,5 |
| ENSMUSG00000041775 | 103,3872769 | 159,5 | 46,2 | 6,71954071 | 0,008233331 | Mapk1ip1 | mitogen-activated protein kinase 1 interacting protein 1 [Source:MGI Symbol;Acc:MGI:1916796] | 84,14 | 22,4 |
| ENSMUSG00000027371 | 127,4285067 | 243,4 | 59,3 | 6,721548674 | 0,00303887 | Fahd2a | fumarylacetoacetate hydrolase domain containing 2A [Source:MGI Symbol;Acc:MGI:1915376] | 100,00 | 31,1 |
| ENSMUSG00000026831 | 88,36163539 | 150,6 | 28,9 | 6,729216162 | 0,013227625 | 1700007K13Rik | RIKEN cDNA 1700007K13 gene [Source:MGI Symbol;Acc:MGI:1916577] | 99,88 | 24,7 |
| ENSMUSG00000034227 | 47,08604399 | 80,9 | 0,9 | 6,733666415 | 0,00306943 | Foxj1 | forkhead box J1 [Source:MGI Symbol;Acc:MGI:1347474] | 93,38 | 0,6 |
| ENSMUSG00000029684 | 356,7173895 | 616,0 | 19,8 | 6,734072051 | 0,000212251 | Wasl | Wiskott-Aldrich syndrome-like (human) [Source:MGI Symbol;Acc:MGI:1920428] | 87,15 | 3,6 |
| ENSMUSG00000020087 | 128,4038377 | 188,6 | 44,4 | 6,734093126 | 0,006961964 | Tysnd1 | trypsin domain containing 1 [Source:MGI Symbol;Acc:MGI:1919017] | 99,15 | 22,1 |
| ENSMUSG00000038069 | 124,104377 | 222,4 | 145,2 | 6,736112964 | 0,004452642 | Cdkn2aip | CDKN2A interacting protein [Source:MGI Symbol;Acc:MGI:1918175] | 95,12 | 23,0 |
| ENSMUSG00000032942 | 134,0909744 | 164,5 | 86,9 | 6,738211091 | 0,014282711 | Ucp3 | uncoupling protein 3 (mitochondrial, proton carrier) [Source:MGI Symbol;Acc:MGI:1099787] | 99,65 | 29,6 |
| ENSMUSG00000025437 | 199,0034986 | 188,3 | 16,6 | 6,740985823 | 0,002167561 | Usp33 | ubiquitin specific peptidase 33 [Source:MGI Symbol;Acc:MGI:2159711] | 93,40 | 1,7 |
| ENSMUSG00000097061 | 122,3326106 | 131,4 | 156,7 | 6,74879115 | 0,023068642 | 9330151L19Rik | RIKEN cDNA 9330151L19 gene [Source:MGI Symbol;Acc:MGI:3041168] | 99,20 | 31,7 |
| ENSMUSG00000018809 | 124,7957256 | 137,9 | 258,4 | 6,753231235 | 0,013463688 | Smyd4 | SET and MYND domain containing 4 [Source:MGI Symbol;Acc:MGI:2442796] | 94,58 | 26,5 |
| ENSMUSG00000092536 | 135,8434804 | 185,5 | 33,9 | 6,756829085 | 0,011987908 | Gm20501 | predicted gene 20501 [Source:MGI Symbol;Acc:MGI:5141966] | #DIV/0! | #DIV/0! |
| ENSMUSG00000040964 | 296,0030253 | 500,2 | 153,6 | 6,766314641 | 0,002669967 | Arhgef10l | Rho guanine nucleotide exchange factor (GEF) 10-like [Source:MGI Symbol;Acc:MGI:1920004] | 82,40 | 21,5 |
| ENSMUSG00000031921 | 95,28333921 | 143,9 | 21,0 | 6,766343254 | 0,00598038 | Terf2 | telomeric repeat binding factor 2 [Source:MGI Symbol;Acc:MGI:1195972] | 72,13 | 2,7 |
| ENSMUSG00000035798 | 155,4431074 | 189,2 | 205,2 | 6,76649998 | 0,007941197 | Zdhhc17 | zinc finger, DHHC domain containing 17 [Source:MGI Symbol;Acc:MGI:2445110] | 90,99 | 26,3 |
| ENSMUSG00000020363 | 45,75286798 | 90,2 | 1,0 | 6,767889808 | 0,001998475 | Gfpt2 | glutamine fructose-6-phosphate transaminase 2 [Source:MGI Symbol;Acc:MGI:1338883] | 97,92 | 0,8 |
| ENSMUSG00000085069 | 285,7607269 | 497,4 | 97,7 | 6,769274642 | 0,010707851 | Gm13111 | predicted gene 13111 [Source:MGI Symbol;Acc:MGI:3651882] | 99,80 | 9,1 |
| ENSMUSG00000056153 | 110,3176421 | 150,4 | 159,7 | 6,769375143 | 0,010465506 | Socs6 | suppressor of cytokine signaling 6 [Source:MGI Symbol;Acc:MGI:1924885] | 80,33 | 10,0 |
| ENSMUSG00000028247 | 103,261173 | 157,2 | 22,2 | 6,770302304 | 0,01046065 | Coq3 | coenzyme Q3 methyltransferase [Source:MGI Symbol;Acc:MGI:101813] | 98,60 | 6,8 |
| ENSMUSG00000096014 | 139,0262328 | 235,0 | 19,4 | 6,770463158 | 0,000666709 | Sox1 | SRY (sex determining region Y)-box 1 [Source:MGI Symbol;Acc:MGI:98357] | 73,54 | 6,0 |
| ENSMUSG00000033880 | 164,8769387 | 235,0 | 62,9 | 6,781688728 | 0,006792606 | Lgals3bp | lectin, galactoside-binding, soluble, 3 binding protein [Source:MGI Symbol;Acc:MGI:99554] | 80,38 | 17,9 |
| ENSMUSG00000097204 | 136,7357092 | 187,8 | 101,8 | 6,781754698 | 0,002776493 | Gm17690 | predicted gene, 17690 [Source:MGI Symbol;Acc:MGI:4937324] | 52,94 | 9,8 |
| ENSMUSG00000020400 | 106,7300563 | 137,9 | 5,5 | 6,785135801 | 0,005859943 | Tnip1 | TNFAIP3 interacting protein 1 [Source:MGI Symbol;Acc:MGI:1926194] | 88,91 | 3,6 |
| ENSMUSG00000021918 | 99,35056013 | 145,4 | 6,5 | 6,785182254 | 0,00602753 | Nek4 | NIMA (never in mitosis gene a)-related expressed kinase 4 [Source:MGI Symbol;Acc:MGI:1344404] | 99,84 | 1,2 |
| ENSMUSG00000038683 | 296,5685748 | 440,5 | 87,5 | 6,785566893 | 0,008403466 | Pak1ip1 | PAK1 interacting protein 1 [Source:MGI Symbol;Acc:MGI:1915333] | 100,00 | 30,6 |
| ENSMUSG00000017667 | 64,0399971 | 147,6 | 39,9 | 6,786489615 | 0,004610597 | Zfp334 | zinc finger protein 334 [Source:MGI Symbol;Acc:MGI:2388656] | 71,96 | 5,6 |
| ENSMUSG00000019817 | 75,21033533 | 64,3 | 1,0 | 6,787483275 | 0,022313912 | Plagl1 | pleiomorphic adenoma gene-like 1 [Source:MGI Symbol;Acc:MGI:1100874] | 62,29 | 0,2 |
| ENSMUSG00000026798 | 133,7493507 | 185,1 | 32,3 | 6,789399331 | 0,004999214 | Coq4 | coenzyme Q4 [Source:MGI Symbol;Acc:MGI:1098826] | 91,06 | 16,3 |

| | | | | | | | | | |
|---|---|---|---|---|---|---|---|---|---|
| ENSMUSG00000030257 | 205,8227405 | 291,4 | 4,3 | 6,790337484 | 0,000503351 | Srgap3 | SLIT-ROBO Rho GTPase activating protein 3 [Source:MGI Symbol;Acc:MGI:2152938] | 94,03 | 1,0 |
| ENSMUSG00000038233 | 86,59517542 | 138,3 | 109,8 | 6,790765573 | 0,013120608 | Fam198a | family with sequence similarity 198, member A [Source:MGI Symbol;Acc:MGI:3041196] | 98,39 | 23,2 |
| ENSMUSG00000028222 | 110,7329448 | 187,0 | 45,3 | 6,793695588 | 0,004041278 | Calb1 | calbindin 1 [Source:MGI Symbol;Acc:MGI:88248] | 64,35 | 7,5 |
| ENSMUSG00000039531 | 32,21477091 | 75,0 | 0,9 | 6,797285867 | 0,004525086 | Zufsp | zinc finger with UFM1-specific peptidase domain [Source:MGI Symbol;Acc:MGI:1919830] | 79,92 | 1,0 |
| ENSMUSG00000025224 | 160,0730236 | 185,1 | 62,9 | 6,79808539 | 0,001908984 | Gbf1 | golgi-specific brefeldin A-resistance factor 1 [Source:MGI Symbol;Acc:MGI:1861607] | 88,90 | 4,2 |
| ENSMUSG00000026342 | 162,6680315 | 197,6 | 218,5 | 6,799556823 | 0,00727828 | Slc35f5 | solute carrier family 35, member F5 [Source:MGI Symbol;Acc:MGI:1921400] | 95,55 | 26,9 |
| ENSMUSG00000020083 | 95,07994052 | 167,5 | 85,8 | 6,801512764 | 0,005506166 | 2010107G23Rik | RIKEN cDNA 2010107G23 gene [Source:MGI Symbol;Acc:MGI:1917144] | 48,02 | 11,9 |
| ENSMUSG00000030222 | 36,53612297 | 64,8 | 0,6 | 6,805169176 | 0,014287526 | Rerg | RAS-like, estrogen-regulated, growth-inhibitor [Source:MGI Symbol;Acc:MGI:2665139] | 81,02 | 0,5 |
| ENSMUSG00000023915 | 158,0944897 | 158,6 | 196,6 | 6,805458691 | 0,00954709 | Tnfrsf21 | tumor necrosis factor receptor superfamily, member 21 [Source:MGI Symbol;Acc:MGI:2151075] | 98,16 | 27,3 |
| ENSMUSG00000022329 | 179,8485996 | 178,2 | 27,7 | 6,805703149 | 0,003720993 | Stk3 | serine/threonine kinase 3 [Source:MGI Symbol;Acc:MGI:1928487] | 96,92 | 5,0 |
| ENSMUSG00000035958 | 158,8259342 | 186,1 | 193,2 | 6,806125506 | 0,008088217 | Tdp2 | tyrosyl-DNA phosphodiesterase 2 [Source:MGI Symbol;Acc:MGI:1860486] | 99,47 | 29,7 |
| ENSMUSG00000060935 | 250,8301413 | 384,7 | 16,3 | 6,809837216 | 1,86418E-05 | Tmem263 | transmembrane protein 263 [Source:MGI Symbol;Acc:MGI:2143652] | 93,48 | 5,3 |
| ENSMUSG00000049999 | 154,1740089 | 278,4 | 170,1 | 6,809871321 | 0,003674047 | Ppp1r3d | protein phosphatase 1, regulatory subunit 3D [Source:MGI Symbol;Acc:MGI:1917664] | 98,84 | 29,9 |
| ENSMUSG00000027827 | 123,9444118 | 232,3 | 106,5 | 6,810292559 | 0,005752492 | Kcnab1 | ssium voltage-gated channel, shaker-related subfamily, beta member 1 [Source:MGI Symbol;Acc:MGI:10! | 99,91 | 17,5 |
| ENSMUSG00000022219 | 156,4374869 | 258,4 | 3,9 | 6,810511834 | 0,004472862 | Cideb | leath-inducing DNA fragmentation factor, alpha subunit-like effector B [Source:MGI Symbol;Acc:MGI:1270 | 100,00 | 0,4 |
| ENSMUSG00000051855 | 141,7151159 | 134,0 | 2,8 | 6,811273058 | 0,010820437 | Mest | mesoderm specific transcript [Source:MGI Symbol;Acc:MGI:96968] | 73,60 | 1,9 |
| ENSMUSG00000035469 | 326,3044668 | 501,8 | 8,0 | 6,815395039 | 2,70315E-08 | Rcbtb1 | romosome condensation (RCC1) and BTB (POZ) domain containing protein 1 [Source:MGI Symbol;Acc:N | 92,93 | 2,2 |
| ENSMUSG00000037957 | 135,7075823 | 220,4 | 36,0 | 6,82187441 | 0,003406984 | Wdr20 | WD repeat domain 20 [Source:MGI Symbol;Acc:MGI:1916891] | 92,91 | 13,5 |
| ENSMUSG00000029386 | 127,7264184 | 174,1 | 70,2 | 6,822747553 | 0,015360766 | Tctn2 | tectonic family member 2 [Source:MGI Symbol;Acc:MGI:1915228] | #DIV/0! | #DIV/0! |
| ENSMUSG00000057315 | 159,7518915 | 131,4 | 53,9 | 6,824475395 | 0,022180393 | Arhgap24 | Rho GTPase activating protein 24 [Source:MGI Symbol;Acc:MGI:1922647] | 66,73 | 6,0 |
| ENSMUSG00000032373 | 263,0968101 | 281,9 | 31,9 | 6,825639286 | 0,017750337 | Car12 | carbonic anyhydrase 12 [Source:MGI Symbol;Acc:MGI:1923709] | 97,55 | 3,2 |
| ENSMUSG00000026153 | 151,4505129 | 190,2 | 1,9 | 6,828046992 | 0,00337462 | Fam135a | family with sequence similarity 135, member A [Source:MGI Symbol;Acc:MGI:1915437] | 90,51 | 0,8 |
| ENSMUSG00000032422 | 151,954295 | 209,8 | 104,4 | 6,829636084 | 0,003706942 | Snx14 | sorting nexin 14 [Source:MGI Symbol;Acc:MGI:2155664] | 98,41 | 27,0 |
| ENSMUSG00000024867 | 141,0896619 | 152,7 | 38,8 | 6,829764465 | 0,003781359 | Pip5k1b | phosphatidylinositol-4-phosphate 5-kinase, type 1 beta [Source:MGI Symbol;Acc:MGI:107930] | 98,74 | 10,7 |
| ENSMUSG00000029561 | 45,64098297 | 75,1 | 1,0 | 6,831066494 | 0,018852943 | Oasl2 | 2'-5' oligoadenylate synthetase-like 2 [Source:MGI Symbol;Acc:MGI:1344390] | 71,88 | 0,0 |
| ENSMUSG00000051727 | 102,6955835 | 172,0 | 36,3 | 6,832677013 | 0,010577991 | Kctd14 | potassium channel tetramerisation domain containing 14 [Source:MGI Symbol;Acc:MGI:1289222] | 68,32 | 9,8 |
| ENSMUSG00000000001 | 715,7066754 | 846,7 | 108,0 | 6,832794982 | 5,39421E-06 | Gnai3 | guanine nucleotide binding protein (G protein), alpha inhibiting 3 [Source:MGI Symbol;Acc:MGI:95773] | 99,86 | 25,6 |
| ENSMUSG00000056158 | 123,4822779 | 123,5 | 61,0 | 6,837685888 | 0,01066796 | Car10 | carbonic anhydrase 10 [Source:MGI Symbol;Acc:MGI:1919855] | 52,17 | 9,8 |
| ENSMUSG00000028580 | 177,5340056 | 256,9 | 172,8 | 6,838553444 | 0,000753834 | Pum1 | pumilio RNA-binding family member 1 [Source:MGI Symbol;Acc:MGI:1931749] | 98,00 | 20,1 |
| ENSMUSG00000001827 | 354,2512144 | 372,3 | 110,8 | 6,839127709 | 0,01081369 | Folr1 | folate receptor 1 (adult) [Source:MGI Symbol;Acc:MGI:95568] | 59,64 | 15,3 |
| ENSMUSG00000047342 | 100,6015601 | 142,2 | 122,7 | 6,848429841 | 0,016261287 | Zfp286 | zinc finger protein 286 [Source:MGI Symbol;Acc:MGI:2384758] | 86,97 | 17,1 |
| ENSMUSG00000071862 | 44,85758413 | 84,4 | 0,6 | 6,852391967 | 0,004402332 | Lrrtm2 | leucine rich repeat transmembrane neuronal 2 [Source:MGI Symbol;Acc:MGI:2389174] | 74,96 | 0,1 |
| ENSMUSG00000074165 | 147,4966822 | 192,9 | 137,4 | 6,853376115 | 0,008966028 | Zfp788 | zinc finger protein 788 [Source:MGI Symbol;Acc:MGI:1914857] | 86,50 | 22,1 |
| ENSMUSG00000038058 | 16,3667247 | 35,2 | 0,0 | 6,853862007 | 0,038787017 | Nod1 | nucleotide-binding oligomerization domain containing 1 [Source:MGI Symbol;Acc:MGI:1341839] | 51,31 | 0,0 |
| ENSMUSG00000020021 | 166,8684536 | 211,7 | 107,7 | 6,856764125 | 0,01343637 | Fgd6 | FYVE, RhoGEF and PH domain containing 6 [Source:MGI Symbol;Acc:MGI:1261419] | 93,72 | 9,9 |
| ENSMUSG00000039405 | 118,1680959 | 148,4 | 45,3 | 6,857154626 | 0,01805405 | Prss23 | protease, serine 23 [Source:MGI Symbol;Acc:MGI:1923703] | 70,28 | 11,8 |
| ENSMUSG00000028064 | 122,2970675 | 164,5 | 87,8 | 6,861077422 | 0,012408832 | Sema4a | lin domain (Ig), transmembrane domain (TM) and short cytoplasmic domain, (semaphorin) 4A [Source:MC | 82,69 | 23,2 |
| ENSMUSG00000024228 | 70,3955046 | 125,1 | 1,2 | 6,863151585 | 0,000657962 | Nudt12 | nudix (nucleoside diphosphate linked moiety X)-type motif 12 [Source:MGI Symbol;Acc:MGI:1915243] | 83,36 | 0,6 |
| ENSMUSG00000015023 | 51,74988525 | 84,6 | 1,0 | 6,863917471 | 0,00398148 | Ddx19a | DEAD (Asp-Glu-Ala-Asp) box polypeptide 19a [Source:MGI Symbol;Acc:MGI:99526] | 76,08 | 0,8 |
| ENSMUSG00000029093 | 201,8224367 | 155,9 | 363,2 | 6,865265135 | 0,018385995 | Sorcs2 | sortilin-related VPS10 domain containing receptor 2 [Source:MGI Symbol;Acc:MGI:1932289] | 83,26 | 24,1 |
| ENSMUSG00000028654 | 54,61210535 | 73,1 | 1,0 | 6,865837985 | 0,01192108 | Mycl | nyelocytomatosis viral oncogene homolog, lung carcinoma derived (avian) [Source:MGI Symbol;Acc:MGI: | 84,34 | 0,0 |
| ENSMUSG00000045284 | 110,468774 | 111,3 | 47,1 | 6,865991266 | 0,016739916 | Dcaf12l1 | DDB1 and CUL4 associated factor 12-like 1 [Source:MGI Symbol;Acc:MGI:2444462] | 93,24 | 15,4 |
| ENSMUSG00000038984 | 120,39037 | 107,4 | 205,6 | 6,866887424 | 0,021100062 | Tspyl5 | testis-specific protein, Y-encoded-like 5 [Source:MGI Symbol;Acc:MGI:2442458] | 86,88 | 28,8 |
| ENSMUSG00000036053 | 177,9756583 | 252,5 | 96,2 | 6,867584667 | 0,003759773 | Fmnl2 | formin-like 2 [Source:MGI Symbol;Acc:MGI:1918659] | 83,68 | 9,2 |
| ENSMUSG00000026077 | 147,4134226 | 273,4 | 132,5 | 6,869354726 | 0,003163529 | Npas2 | neuronal PAS domain protein 2 [Source:MGI Symbol;Acc:MGI:109232] | 90,32 | 26,0 |
| ENSMUSG00000048930 | 92,17839945 | 143,6 | 41,6 | 6,873508336 | 0,018529292 | Tada3 | transcriptional adaptor 3 [Source:MGI Symbol;Acc:MGI:1915724] | 76,19 | 17,1 |
| ENSMUSG00000014039 | 15,9838136 | 27,2 | 0,0 | 6,873691785 | 0,031870795 | Prdm15 | PR domain containing 15 [Source:MGI Symbol;Acc:MGI:1930121] | 39,15 | 0,0 |
| ENSMUSG00000025956 | 109,2875731 | 176,3 | 57,3 | 6,873998055 | 0,012527681 | Mettl21a | methyltransferase like 21A [Source:MGI Symbol;Acc:MGI:1914349] | 76,94 | 14,3 |
| ENSMUSG00000026836 | 52,94303542 | 71,2 | 0,6 | 6,878259723 | 0,009302039 | Acvr1 | activin A receptor, type 1 [Source:MGI Symbol;Acc:MGI:87911] | 75,63 | 0,7 |
| ENSMUSG00000049321 | 144,956163 | 247,1 | 23,0 | 6,881430101 | 0,002239901 | Zfp2 | zinc finger protein 2 [Source:MGI Symbol;Acc:MGI:99167] | 99,01 | 5,6 |
| ENSMUSG00000030323 | 162,2197144 | 219,3 | 128,3 | 6,881858423 | 0,008983635 | Ift122 | intraflagellar transport 122 [Source:MGI Symbol;Acc:MGI:1932386] | 93,13 | 16,0 |
| ENSMUSG00000029600 | 104,400371 | 170,8 | 11,1 | 6,882803076 | 0,005058806 | Rita1 | RBPJ interacting and tubulin associated 1 [Source:MGI Symbol;Acc:MGI:1922021] | 93,85 | 8,8 |
| ENSMUSG00000039208 | 68,16819966 | 107,9 | 1,2 | 6,882815009 | 0,005753019 | Metrnl | meteorin, glial cell differentiation regulator-like [Source:MGI Symbol;Acc:MGI:2384806] | 87,05 | 1,6 |
| ENSMUSG00000027598 | 284,8203909 | 383,5 | 9,6 | 6,8838297 | 0,000604595 | Itch | itchy, E3 ubiquitin protein ligase [Source:MGI Symbol;Acc:MGI:1202301] | 99,39 | 1,7 |
| ENSMUSG00000057101 | 127,4399889 | 125,7 | 2,8 | 6,886212451 | 0,005326329 | Zfp180 | zinc finger protein 180 [Source:MGI Symbol;Acc:MGI:1923701] | 98,62 | 0,4 |
| ENSMUSG00000031990 | 101,929708 | 161,6 | 9,2 | 6,886342438 | 0,005845667 | Jam3 | junction adhesion molecule 3 [Source:MGI Symbol;Acc:MGI:1933825] | 79,31 | 2,5 |
| ENSMUSG00000020546 | 241,922337 | 278,5 | 139,2 | 6,895279554 | 0,006091109 | Stxbp4 | syntaxin binding protein 4 [Source:MGI Symbol;Acc:MGI:1342296] | 60,30 | 6,5 |
| ENSMUSG00000028975 | 143,7789101 | 228,5 | 48,9 | 6,897304284 | 0,002350465 | Pex14 | peroxisomal biogenesis factor 14 [Source:MGI Symbol;Acc:MGI:1927868] | 97,67 | 19,6 |
| ENSMUSG00000064068 | 194,1779701 | 302,2 | 47,1 | 6,898208319 | 0,000493699 | Mtx1 | metaxin 1 [Source:MGI Symbol;Acc:MGI:103025] | 69,67 | 15,4 |
| ENSMUSG00000020704 | 77,7839464 | 139,0 | 3,7 | 6,900715391 | 0,004681539 | Asic2 | acid-sensing (proton-gated) ion channel 2 [Source:MGI Symbol;Acc:MGI:1100867] | 76,89 | 1,6 |
| ENSMUSG00000071078 | 157,4081545 | 239,7 | 58,2 | 6,902697664 | 0,002542704 | Nr2c2ap | nuclear receptor 2C2-associated protein [Source:MGI Symbol;Acc:MGI:1922942] | 100,00 | 30,4 |
| ENSMUSG00000041199 | 148,774063 | 212,2 | 42,9 | 6,90918435 | 0,001798609 | Rpusd1 | RNA pseudouridylate synthase domain containing 1 [Source:MGI Symbol;Acc:MGI:1919186] | 99,82 | 20,9 |
| ENSMUSG00000062444 | 197,0077466 | 140,6 | 413,1 | 6,910488058 | 0,008644815 | Ap3b2 | adaptor-related protein complex 3, beta 2 subunit [Source:MGI Symbol;Acc:MGI:1100869] | 62,29 | 21,1 |
| ENSMUSG00000054648 | 113,3102267 | 182,6 | 75,0 | 6,910795933 | 0,009754334 | Zfp869 | zinc finger protein 869 [Source:MGI Symbol;Acc:MGI:1914119] | 100,00 | 28,1 |
| ENSMUSG00000038077 | 88,15037713 | 115,5 | 2,8 | 6,91120468 | 0,010956967 | Kcna6 | tassium voltage-gated channel, shaker-related, subfamily, member 6 [Source:MGI Symbol;Acc:MGI:9666 | 92,79 | 1,2 |
| ENSMUSG00000025340 | 93,98707206 | 185,1 | 7,0 | 6,91239073 | 0,001310959 | Rabgef1 | RAB guanine nucleotide exchange factor (GEF) 1 [Source:MGI Symbol;Acc:MGI:1929459] | 90,65 | 2,1 |
| ENSMUSG00000048796 | 49,41069546 | 67,6 | 1,0 | 6,914149463 | 0,020396796 | Cyb561d1 | cytochrome b-561 domain containing 1 [Source:MGI Symbol;Acc:MGI:1919273] | 80,76 | 0,5 |
| ENSMUSG00000019796 | 248,5077024 | 346,7 | 110,1 | 6,91534354 | 0,000532199 | Lrp11 | low density lipoprotein receptor-related protein 11 [Source:MGI Symbol;Acc:MGI:2442989] | 99,27 | 28,9 |
| ENSMUSG00000005268 | 271,5760312 | 278,4 | 285,7 | 6,91625703 | 0,008593034 | Prlr | prolactin receptor [Source:MGI Symbol;Acc:MGI:97763] | 54,36 | 12,9 |
| ENSMUSG00000039058 | 161,8058966 | 210,4 | 22,2 | 6,91741402 | 0,001950623 | Ak5 | adenylate kinase 5 [Source:MGI Symbol;Acc:MGI:2677491] | 91,01 | 8,4 |
| ENSMUSG00000028405 | 215,332532 | 312,2 | 32,3 | 6,919293476 | 0,0001986 | Aco1 | aconitase 1 [Source:MGI Symbol;Acc:MGI:87879] | 94,82 | 7,1 |
| ENSMUSG00000105053 | 97,65344272 | 121,3 | 1,8 | 6,924184691 | 0,007365245 | Gm43064 | predicted gene 43064 [Source:MGI Symbol;Acc:MGI:5663201] | #DIV/0! | #DIV/0! |
| ENSMUSG00000010290 | 555,863795 | 530,5 | 249,1 | 6,927258229 | 0,001631726 | AI597479 | expressed sequence AI597479 [Source:MGI Symbol;Acc:MGI:2138299] | 99,65 | 30,3 |
| ENSMUSG00000074925 | 70,2059973 | 119,6 | 2,0 | 6,92750422 | 0,004452642 | Ptar1 | protein prenyltransferase alpha subunit repeat containing 1 [Source:MGI Symbol;Acc:MGI:1921875] | 98,76 | 0,9 |
| ENSMUSG00000058325 | 344,9268633 | 623,5 | 310,3 | 6,930938041 | 0,001089664 | Dock1 | dedicator of cytokinesis 1 [Source:MGI Symbol;Acc:MGI:2429765] | 99,99 | 28,0 |
| ENSMUSG00000070880 | 185,8674592 | 191,7 | 218,5 | 6,932028231 | 0,00671309 | Gad1 | glutamate decarboxylase 1 [Source:MGI Symbol;Acc:MGI:95632] | 71,29 | 21,6 |
| ENSMUSG00000026678 | 10,08587558 | 26,9 | 0,0 | 6,93993845 | 0,039971944 | Rgs5 | regulator of G-protein signaling 5 [Source:MGI Symbol;Acc:MGI:1098434] | 50,26 | 0,0 |
| ENSMUSG00000040972 | 112,1068181 | 171,9 | 4,8 | 6,940125613 | 0,005499938 | Igsf21 | immunoglobulin superfamily, member 21 [Source:MGI Symbol;Acc:MGI:2681842] | 99,71 | 2,5 |
| ENSMUSG00000069206 | 97,63063994 | 170,3 | 42,5 | 6,940240335 | 0,009373845 | Zfp874a | zinc finger protein 874a [Source:MGI Symbol;Acc:MGI:3040703] | 65,92 | 7,3 |
| ENSMUSG00000027217 | 58,92199324 | 71,5 | 1,0 | 6,951896224 | 0,016615332 | Tspan18 | tetraspanin 18 [Source:MGI Symbol;Acc:MGI:1917186] | 88,84 | 0,4 |
| ENSMUSG00000091768 | 7,794742612 | 11,7 | 0,0 | 6,952573945 | 0,04970542 | Gm17617 | predicted gene, 17617 [Source:MGI Symbol;Acc:MGI:4937251] | 100,00 | 0,0 |
| ENSMUSG00000027086 | 68,74844362 | 87,1 | 0,9 | 6,952616435 | 0,002893214 | Fastkd1 | FAST kinase domains 1 [Source:MGI Symbol;Acc:MGI:2444596] | 97,48 | 0,9 |
| ENSMUSG00000034957 | 90,24935101 | 201,1 | 21,3 | 6,953272332 | 0,003911784 | Cebpa | CCAAT/enhancer binding protein (C/EBP), alpha [Source:MGI Symbol;Acc:MGI:99480] | 98,77 | 12,7 |
| ENSMUSG00000043190 | 124,4613171 | 183,4 | 40,5 | 6,953422809 | 0,013421166 | Rfesd | Rieske (Fe-S) domain containing [Source:MGI Symbol;Acc:MGI:2145198] | 76,54 | 8,9 |
| ENSMUSG00000046352 | 106,3429365 | 187,0 | 43,0 | 6,959187187 | 0,007871312 | Gjb2 | gap junction protein, beta 2 [Source:MGI Symbol;Acc:MGI:95720] | 98,89 | 19,4 |
| ENSMUSG00000047921 | 141,9757025 | 230,9 | 146,7 | 6,96106903 | 0,001155151 | Trappc9 | trafficking protein particle complex 9 [Source:MGI Symbol;Acc:MGI:1923760] | 61,94 | 16,3 |
| ENSMUSG00000040123 | 157,4169473 | 177,5 | 97,0 | 6,964256465 | 0,008596492 | Zmym5 | zinc finger, MYM-type 5 [Source:MGI Symbol;Acc:MGI:3041170] | 71,80 | 14,7 |
| ENSMUSG00000067995 | 136,0012683 | 158,2 | 57,3 | 6,966945455 | 0,009754334 | Gtf2f2 | general transcription factor IIF, polypeptide 2 [Source:MGI Symbol;Acc:MGI:1915955] | 99,86 | 14,7 |
| ENSMUSG00000020659 | 117,0530831 | 178,2 | 3,0 | 6,969329671 | 0,000432662 | Cbll1 | Casitas B-lineage lymphoma-like 1 [Source:MGI Symbol;Acc:MGI:2144842] | 95,28 | 0,0 |
| ENSMUSG00000021051 | 204,8896084 | 376,6 | 9,7 | 6,970536112 | 1,92946E-05 | Ppp2r5e | protein phosphatase 2, regulatory subunit B', epsilon [Source:MGI Symbol;Acc:MGI:1349473] | 95,23 | 0,9 |
| ENSMUSG00000071267 | 146,7213332 | 210,7 | 10,9 | 6,970982699 | 0,005503191 | Zfp942 | zinc finger protein 942 [Source:MGI Symbol;Acc:MGI:1920483] | 98,02 | 2,2 |
| ENSMUSG00000039145 | 121,2607831 | 187,1 | 1,6 | 6,972858294 | 0,000753834 | Camk1d | calcium/calmodulin-dependent protein kinase ID [Source:MGI Symbol;Acc:MGI:2442190] | 93,57 | 0,6 |
| ENSMUSG00000006611 | 117,1295614 | 186,7 | 35,9 | 6,973527858 | 0,007141754 | Hfe | hemochromatosis [Source:MGI Symbol;Acc:MGI:109191] | 93,29 | 9,9 |
| ENSMUSG00000084910 | 109,8194685 | 172,1 | 35,1 | 6,974776642 | 0,00954709 | C630043F03Rik | RIKEN cDNA C630043F03 gene [Source:MGI Symbol;Acc:MGI:1915535] | #DIV/0! | #DIV/0! |
| ENSMUSG00000049100 | 317,2339408 | 421,7 | 82,3 | 6,976349593 | 0,000121115 | Pcdh10 | protocadherin 10 [Source:MGI Symbol;Acc:MGI:1338042] | 73,61 | 5,0 |
| ENSMUSG00000032456 | 42,96001473 | 82,4 | 0,9 | 6,977144038 | 0,005708695 | Nmnat3 | nicotinamide nucleotide adenylyltransferase 3 [Source:MGI Symbol;Acc:MGI:1921330] | 77,14 | 0,9 |
| ENSMUSG00000041773 | 73,91205087 | 68,2 | 0,6 | 6,981857163 | 0,015050976 | Enc1 | ectodermal-neural cortex 1 [Source:MGI Symbol;Acc:MGI:109610] | 65,77 | 0,5 |
| ENSMUSG00000061306 | 140,40327 | 240,5 | 12,0 | 6,985246378 | 0,001506164 | Slc38a10 | solute carrier family 38, member 10 [Source:MGI Symbol;Acc:MGI:1919305] | 78,14 | 2,2 |
| ENSMUSG00000024076 | 38,6719934 | 96,8 | 0,6 | 6,988030056 | 0,002347168 | Vit | vitrin [Source:MGI Symbol;Acc:MGI:1921449] | 95,54 | 0,1 |
| ENSMUSG00000021044 | 90,95683868 | 175,5 | 1,8 | 6,990514257 | 0,000295215 | Adck1 | aarF domain containing kinase 1 [Source:MGI Symbol;Acc:MGI:1919363] | 98,76 | 1,1 |
| ENSMUSG00000090110 | 142,4473288 | 220,7 | 98,0 | 6,993179112 | 0,003647101 | Cmc4 | C-x(9)-C motif containing 4 [Source:MGI Symbol;Acc:MGI:5637812] | 77,18 | 21,5 |
| ENSMUSG00000053870 | 102,4870117 | 210,4 | 33,3 | 6,995390747 | 0,006712627 | Fpgt | fucose-1-phosphate guanylyltransferase [Source:MGI Symbol;Acc:MGI:1922790] | 98,89 | 3,1 |
| ENSMUSG00000048251 | 120,0147442 | 154,4 | 1,8 | 6,997883985 | 0,00138233 | Bcl11b | B cell leukemia/lymphoma 11B [Source:MGI Symbol;Acc:MGI:1929913] | 72,33 | 0,2 |
| ENSMUSG00000048218 | 144,0214897 | 72,8 | 1,0 | 6,999025347 | 0,018209447 | Amigo2 | adhesion molecule with Ig like domain 2 [Source:MGI Symbol;Acc:MGI:2145995] | 89,62 | 0,8 |
| ENSMUSG00000059263 | 210,4029216 | 294,5 | 27,9 | 7,001224329 | 9,20331E-05 | Usp47 | ubiquitin specific peptidase 47 [Source:MGI Symbol;Acc:MGI:1922246] | 98,50 | 1,6 |
| ENSMUSG00000079057 | 133,3194257 | 206,0 | 175,6 | 7,002544327 | 0,007564018 | Cyp4v3 | cytochrome P450, family 4, subfamily v, polypeptide 3 [Source:MGI Symbol;Acc:MGI:2142763] | 85,78 | 22,5 |
| ENSMUSG00000000078 | 156,8041328 | 164,9 | 327,3 | 7,004382591 | 0,01265198 | Klf6 | Kruppel-like factor 6 [Source:MGI Symbol;Acc:MGI:1346318] | 86,90 | 15,8 |
| ENSMUSG00000046546 | 37,60762554 | 91,4 | 1,0 | 7,005085479 | 0,003008247 | Fam43a | family with sequence similarity 43, member A [Source:MGI Symbol;Acc:MGI:2676309] | 93,55 | 0,3 |
| ENSMUSG00000031171 | 190,3480861 | 235,8 | 85,9 | 7,009046647 | 0,004573327 | Ftsj1 | FtsJ RNA methyltransferase homolog 1 (E. coli) [Source:MGI Symbol;Acc:MGI:1859648] | 61,68 | 15,5 |
| ENSMUSG00000034220 | 89,2540808 | 138,2 | 2,0 | 7,01010821 | 0,001483845 | Gpc1 | glypican 1 [Source:MGI Symbol;Acc:MGI:1194891] | 87,36 | 0,8 |
| ENSMUSG00000051590 | 42,62183985 | 72,4 | 0,6 | 7,010768597 | 0,009197733 | Map3k19 | mitogen-activated protein kinase kinase kinase 19 [Source:MGI Symbol;Acc:MGI:1203481] | 89,59 | 0,3 |
| ENSMUSG00000027890 | 110,5956202 | 204,3 | 14,0 | 7,013129952 | 0,001774683 | Gstm4 | glutathione S-transferase, mu 4 [Source:MGI Symbol;Acc:MGI:95862] | 98,03 | 7,0 |
| ENSMUSG00000043004 | 315,6551338 | 456,4 | 199,1 | 7,015916532 | 0,000512016 | Gng2 | guanine nucleotide binding protein (G protein), gamma 2 [Source:MGI Symbol;Acc:MGI:102705] | 83,39 | 25,9 |
| ENSMUSG00000038816 | 293,3841813 | 300,5 | 239,1 | 7,016435529 | 0,002176746 | Ctnnal1 | catenin (cadherin associated protein), alpha-like 1 [Source:MGI Symbol;Acc:MGI:1859649] | 94,76 | 31,2 |
| ENSMUSG00000008136 | 53,60675707 | 80,1 | 0,9 | 7,016677354 | 0,018585862 | Fhl2 | four and a half LIM domains 2 [Source:MGI Symbol;Acc:MGI:1338762] | 78,74 | 1,1 |
| ENSMUSG00000039958 | 54,11162021 | 114,0 | 1,8 | 7,01723504 | 0,005837578 | Mettl20 | methyltransferase like 20 [Source:MGI Symbol;Acc:MGI:2443575] | 73,46 | 1,0 |
| ENSMUSG00000046324 | 104,4836819 | 150,6 | 5,5 | 7,019639176 | 0,0080152 | Ermp1 | endoplasmic reticulum metallopeptidase 1 [Source:MGI Symbol;Acc:MGI:106250] | 71,20 | 0,8 |
| ENSMUSG00000036782 | 97,5278842 | 200,7 | 4,0 | 7,019656743 | 0,000339105 | Klhl13 | kelch-like 13 [Source:MGI Symbol;Acc:MGI:1914705] | 84,13 | 2,2 |
| ENSMUSG00000002055 | 116,1005062 | 212,8 | 21,3 | 7,020659088 | 0,003489762 | Spag5 | sperm associated antigen 5 [Source:MGI Symbol;Acc:MGI:1927470] | 52,65 | 3,7 |
| ENSMUSG00000041703 | 70,4445115 | 141,1 | 1,8 | 7,026411019 | 0,001147305 | Zic5 | zinc finger protein of the cerebellum 5 [Source:MGI Symbol;Acc:MGI:1929518] | 76,10 | 0,8 |
| ENSMUSG00000071176 | 114,9050238 | 228,1 | 75,8 | 7,02644922 | 0,003527765 | Arhgef10 | Rho guanine nucleotide exchange factor (GEF) 10 [Source:MGI Symbol;Acc:MGI:2444453] | 98,11 | 14,7 |
| ENSMUSG00000035596 | 114,3976847 | 179,1 | 12,0 | 7,028941103 | 0,005444277 | Mboat7 | membrane bound O-acyltransferase domain containing 7 [Source:MGI Symbol;Acc:MGI:1924832] | 85,70 | 2,0 |
| ENSMUSG00000020409 | 316,1051452 | 389,3 | 97,4 | 7,032618804 | 0,000753834 | Slu7 | SLU7 splicing factor homolog (S. cerevisiae) [Source:MGI Symbol;Acc:MGI:2385598] | 84,96 | 20,7 |
| ENSMUSG00000055401 | 120,2934461 | 150,7 | 2,0 | 7,034014803 | 0,00068619 | Fbxo6 | F-box protein 6 [Source:MGI Symbol;Acc:MGI:1354743] | 66,71 | 1,1 |
| ENSMUSG00000024180 | 71,08658044 | 116,4 | 1,2 | 7,036060401 | 0,005859943 | Tmem8 | transmembrane protein 8 (five membrane-spanning domains) [Source:MGI Symbol;Acc:MGI:1926283] | 91,00 | 0,6 |
| ENSMUSG00000024805 | 208,711695 | 259,8 | 93,3 | 7,037758321 | 0,001949616 | Pcgf5 | polycomb group ring finger 5 [Source:MGI Symbol;Acc:MGI:1923505] | 89,33 | 27,1 |
| ENSMUSG00000038605 | 93,80201582 | 68,8 | 0,6 | 7,039883388 | 0,01543911 | Samd10 | sterile alpha motif domain containing 10 [Source:MGI Symbol;Acc:MGI:2443872] | 97,38 | 1,1 |
| ENSMUSG00000041544 | 88,59007234 | 142,9 | 2,0 | 7,044686738 | 0,001256291 | Ptchd2 | patched domain containing 2 [Source:MGI Symbol;Acc:MGI:2444403] | 96,17 | 0,5 |
| ENSMUSG00000013495 | 246,8840115 | 301,2 | 64,8 | 7,045528016 | 0,001697391 | Tmem175 | transmembrane protein 175 [Source:MGI Symbol;Acc:MGI:1919642] | 80,21 | 11,4 |
| ENSMUSG00000021838 | 40,90892164 | 77,9 | 0,9 | 7,048301627 | 0,012259683 | Samd4 | sterile alpha motif domain containing 4 [Source:MGI Symbol;Acc:MGI:1921730] | 68,78 | 0,3 |
| ENSMUSG00000037104 | 134,1307806 | 191,0 | 4,6 | 7,052092038 | 0,00119092 | Socs5 | suppressor of cytokine signaling 5 [Source:MGI Symbol;Acc:MGI:2385459] | 96,65 | 1,6 |
| ENSMUSG00000030306 | 94,02253566 | 90,0 | 0,9 | 7,056043696 | 0,008948721 | Tmtc1 | transmembrane and tetratricopeptide repeat containing 1 [Source:MGI Symbol;Acc:MGI:3039590] | 80,75 | 0,0 |
| ENSMUSG00000027378 | 300,9383364 | 327,3 | 89,6 | 7,060244682 | 0,00171394 | Nphp1 | nephronophthisis 1 (juvenile) homolog (human) [Source:MGI Symbol;Acc:MGI:1858233] | 99,16 | 31,1 |
| ENSMUSG00000036430 | 105,2819102 | 192,4 | 12,0 | 7,064909791 | 0,004690238 | Tbcc | tubulin-specific chaperone C [Source:MGI Symbol;Acc:MGI:1919976] | 83,78 | 10,8 |
| ENSMUSG00000024142 | 141,8061079 | 258,2 | 77,6 | 7,065667941 | 0,00418273 | Mlst8 | MTOR associated protein, LST8 homolog (S. cerevisiae) [Source:MGI Symbol;Acc:MGI:1929514] | 99,29 | 24,7 |
| ENSMUSG00000056267 | 85,87777966 | 146,9 | 2,8 | 7,066554045 | 0,003568742 | Cep70 | centrosomal protein 70 [Source:MGI Symbol;Acc:MGI:1915371] | 81,85 | 1,5 |
| ENSMUSG00000072941 | 202,2197858 | 241,0 | 37,9 | 7,068397841 | 0,005254334 | Sod3 | superoxide dismutase 3, extracellular [Source:MGI Symbol;Acc:MGI:103181] | 49,90 | 8,5 |
| ENSMUSG00000038195 | 48,33409201 | 78,3 | 0,6 | 7,069588561 | 0,009049788 | Rilp | Rab interacting lysosomal protein [Source:MGI Symbol;Acc:MGI:2144271] | 94,36 | 1,7 |
| ENSMUSG00000018995 | 134,4010818 | 228,4 | 82,3 | 7,07286276 | 0,006806315 | Nars2 | asparaginyl-tRNA synthetase 2 (mitochondrial)(putative) [Source:MGI Symbol;Acc:MGI:2142075] | 75,60 | 16,3 |
| ENSMUSG00000002608 | 168,542751 | 254,0 | 70,2 | 7,076832721 | 0,001323936 | Ccdc97 | coiled-coil domain containing 97 [Source:MGI Symbol;Acc:MGI:1196455] | 98,56 | 20,5 |
| ENSMUSG00000036721 | 83,97188903 | 82,4 | 1,0 | 7,085542986 | 0,011309178 | Zscan12 | zinc finger and SCAN domain containing 12 [Source:MGI Symbol;Acc:MGI:1099444] | 87,33 | 1,1 |
| ENSMUSG00000078640 | 167,435316 | 232,1 | 89,0 | 7,088462317 | 0,009369036 | Gm11627 | predicted gene 11627 [Source:MGI Symbol;Acc:MGI:3650659] | 98,14 | 25,4 |
| ENSMUSG00000029994 | 174,2778287 | 243,2 | 167,0 | 7,088721997 | 0,005518094 | Anxa4 | annexin A4 [Source:MGI Symbol;Acc:MGI:88030] | 92,19 | 28,3 |
| ENSMUSG00000034473 | 178,125535 | 163,1 | 113,8 | 7,089478022 | 0,011768063 | Sec22a | SEC22 homolog A, vesicle trafficking protein [Source:MGI Symbol;Acc:MGI:2447876] | 50,93 | 12,8 |
| ENSMUSG00000059439 | 160,1478969 | 216,3 | 132,7 | 7,091741033 | 0,003119232 | Bcas3 | breast carcinoma amplified sequence 3 [Source:MGI Symbol;Acc:MGI:2385848] | 72,59 | 16,6 |
| ENSMUSG00000032766 | 86,41040128 | 167,7 | 12,9 | 7,092883974 | 0,009302039 | Gng11 | guanine nucleotide binding protein (G protein), gamma 11 [Source:MGI Symbol;Acc:MGI:1913316] | 97,17 | 11,7 |
| ENSMUSG00000025986 | 151,7759093 | 195,9 | 108,8 | 7,097106429 | 0,003290842 | Slc39a10 | solute carrier family 39 (zinc transporter), member 10 [Source:MGI Symbol;Acc:MGI:1914515] | 87,67 | 12,1 |
| ENSMUSG00000002578 | 9,816457916 | 12,9 | 0,0 | 7,097924953 | 0,041737816 | Ikzf4 | IKAROS family zinc finger 4 [Source:MGI Symbol;Acc:MGI:1343139] | 30,44 | 0,0 |
| ENSMUSG00000041660 | 146,3664229 | 273,0 | 145,9 | 7,105891772 | 0,007465481 | Bbox1 | (gamma), 2-oxoglutarate dioxygenase 1 (gamma-butyrobetaine hydroxylase) [Source:MGI Symbol;Acc:M | 88,67 | 13,7 |
| ENSMUSG00000024947 | 270,0726503 | 303,8 | 22,0 | 7,106169417 | 0,000261321 | Men1 | multiple endocrine neoplasia 1 [Source:MGI Symbol;Acc:MGI:1316736] | 93,36 | 6,5 |
| ENSMUSG00000004895 | 61,36725762 | 87,1 | 1,0 | 7,109709859 | 0,009369036 | Prcc | papillary renal cell carcinoma (translocation-associated) [Source:MGI Symbol;Acc:MGI:2137738] | 94,79 | 0,9 |
| ENSMUSG00000035614 | 176,8696122 | 206,9 | 1,5 | 7,111633662 | 0,000341676 | Fam179b | family with sequence similarity 179, member B [Source:MGI Symbol;Acc:MGI:2684313] | 97,67 | 0,5 |
| ENSMUSG00000021306 | 78,31130941 | 141,1 | 1,2 | 7,112168915 | 0,001310959 | Gpr137b | G protein-coupled receptor 137B [Source:MGI Symbol;Acc:MGI:1891463] | 82,68 | 0,8 |
| ENSMUSG00000032577 | 123,2908621 | 205,5 | 16,9 | 7,119579664 | 0,006895432 | Mapkapk3 | mitogen-activated protein kinase-activated protein kinase 3 [Source:MGI Symbol;Acc:MGI:2143163] | 88,72 | 6,7 |
| ENSMUSG00000093598 | 153,463001 | 194,2 | 50,9 | 7,119726395 | 0,006853154 | A730085K08Rik | RIKEN cDNA A730085K08 gene [Source:MGI Symbol;Acc:MGI:3605633] | #DIV/0! | #DIV/0! |
| ENSMUSG00000019851 | 140,4797171 | 154,1 | 25,0 | 7,125814718 | 0,011287367 | Perp | PERP, TP53 apoptosis effector [Source:MGI Symbol;Acc:MGI:1929938] | 97,35 | 6,3 |
| ENSMUSG00000031979 | 200,7525168 | 217,8 | 209,8 | 7,126624604 | 0,007327727 | Cog2 | component of oligomeric golgi complex 2 [Source:MGI Symbol;Acc:MGI:1923582] | 97,69 | 31,2 |
| ENSMUSG00000025607 | 134,3864552 | 197,2 | 6,5 | 7,127108529 | 0,002393298 | Copg2 | coatomer protein complex, subunit gamma 2 [Source:MGI Symbol;Acc:MGI:1858683] | 98,15 | 2,7 |
| ENSMUSG00000000732 | 84,05896197 | 173,9 | 70,2 | 7,128114981 | 0,014211001 | Icosl | icos ligand [Source:MGI Symbol;Acc:MGI:1354701] | 99,08 | 25,3 |
| ENSMUSG00000070806 | 9,003413325 | 12,9 | 0,0 | 7,128947368 | 0,048733912 | Zmynd12 | zinc finger, MYND domain containing 12 [Source:MGI Symbol;Acc:MGI:2140259] | 68,26 | 0,0 |
| ENSMUSG00000090758 | 52,34851844 | 95,1 | 1,0 | 7,130012737 | 0,006918725 | Gm17178 | predicted gene 17178 [Source:MGI Symbol;Acc:MGI:4938005] | 90,85 | 4,4 |
| ENSMUSG00000064120 | 46,1532764 | 93,2 | 0,9 | 7,133116939 | 0,003823775 | Mocs1 | molybdenum cofactor synthesis 1 [Source:MGI Symbol;Acc:MGI:1928904] | 89,44 | 0,7 |
| ENSMUSG00000034235 | 502,8089575 | 826,0 | 244,7 | 7,13709247 | 0,002117304 | Usp54 | ubiquitin specific peptidase 54 [Source:MGI Symbol;Acc:MGI:1926037] | 99,95 | 24,1 |
| ENSMUSG00000086677 | 8,245887186 | 15,3 | 0,0 | 7,139022398 | 0,048905248 | Tvp23bos | trans-golgi network vesicle protein 23B, opposite strand [Source:MGI Symbol;Acc:MGI:3649886] | 72,11 | 0,0 |
| ENSMUSG00000020122 | 41,40352531 | 101,4 | 1,0 | 7,139224106 | 0,002393298 | Egfr | epidermal growth factor receptor [Source:MGI Symbol;Acc:MGI:95294] | 66,50 | 0,1 |
| ENSMUSG00000024664 | 157,7333735 | 160,1 | 85,0 | 7,140111073 | 0,010533806 | Fads3 | fatty acid desaturase 3 [Source:MGI Symbol;Acc:MGI:1928740] | 95,89 | 23,4 |
| ENSMUSG00000024292 | 154,9204406 | 255,5 | 93,8 | 7,140544126 | 0,00428721 | Cyp4f14 | cytochrome P450, family 4, subfamily f, polypeptide 14 [Source:MGI Symbol;Acc:MGI:1927669] | 92,70 | 24,3 |
| ENSMUSG00000020607 | 176,0406473 | 325,9 | 72,0 | 7,146308054 | 0,0012678 | Fam84a | family with sequence similarity 84, member A [Source:MGI Symbol;Acc:MGI:2145011] | 100,00 | 4,9 |
| ENSMUSG00000037251 | 124,392056 | 189,9 | 3,7 | 7,146979094 | 0,001501406 | Pomk | protein-O-mannose kinase [Source:MGI Symbol;Acc:MGI:1921903] | 91,50 | 2,5 |
| ENSMUSG00000030545 | 223,5194201 | 362,8 | 90,8 | 7,147091397 | 0,000685443 | Pex11a | peroxisomal biogenesis factor 11 alpha [Source:MGI Symbol;Acc:MGI:1338788] | 99,81 | 22,7 |

| | | | | | | | | | |
|---|---|---|---|---|---|---|---|---|---|
| ENSMUSG00000025475 | 115,2948309 | 186,0 | 9,7 | 7,147809417 | 0,009033509 | Adgra1 | adhesion G protein-coupled receptor A1 [Source:MGI Symbol;Acc:MGI:1277167] | 94,42 | 1,8 |
| ENSMUSG00000046567 | 96,90976468 | 228,7 | 1,9 | 7,148842869 | 6,26813E-05 | 4930430F08Rik | RIKEN cDNA 4930430F08 gene [Source:MGI Symbol;Acc:MGI:1921197] | 98,97 | 1,9 |
| ENSMUSG00000073609 | 74,87446094 | 90,7 | 0,6 | 7,14976288 | 0,007449232 | D2hgdh | D-2-hydroxyglutarate dehydrogenase [Source:MGI Symbol;Acc:MGI:2138209] | 68,76 | 0,3 |
| ENSMUSG00000027569 | 166,068423 | 185,0 | 110,9 | 7,150794137 | 0,011416763 | Mrgbp | MRG/MORF4L binding protein [Source:MGI Symbol;Acc:MGI:1920497] | 81,07 | 25,2 |
| ENSMUSG00000047669 | 153,3200147 | 310,0 | 51,8 | 7,151115656 | 0,001200332 | Msl3l2 | male-specific lethal 3-like 2 (Drosophila) [Source:MGI Symbol;Acc:MGI:1920640] | 99,97 | 21,6 |
| ENSMUSG00000028555 | 10,84748395 | 13,1 | 0,0 | 7,152307774 | 0,04659058 | Ttc39a | tetratricopeptide repeat domain 39A [Source:MGI Symbol;Acc:MGI:2444350] | 31,56 | 0,0 |
| ENSMUSG00000046961 | 11,5983974 | 18,4 | 0,0 | 7,152672737 | 0,047598759 | Gpr156 | G protein-coupled receptor 156 [Source:MGI Symbol;Acc:MGI:2653880] | 36,73 | 0,0 |
| ENSMUSG00000090071 | 18,15688569 | 15,4 | 0,0 | 7,156232205 | 0,046581088 | Cdk5r2 | cyclin-dependent kinase 5, regulatory subunit 2 (p39) [Source:MGI Symbol;Acc:MGI:1330828] | 51,58 | 0,0 |
| ENSMUSG00000044229 | 117,2502416 | 190,1 | 93,8 | 7,16543901 | 0,004625348 | Nxpe4 | neurexophilin and PC-esterase domain family, member 4 [Source:MGI Symbol;Acc:MGI:1924792] | 96,33 | 12,9 |
| ENSMUSG00000073856 | 114,1939635 | 188,5 | 2,4 | 7,168212982 | 0,00161365 | Iqck | IQ motif containing K [Source:MGI Symbol;Acc:MGI:3612188] | 45,06 | 0,9 |
| ENSMUSG00000000811 | 52,17487984 | 85,3 | 0,9 | 7,168734798 | 0,010159129 | Txnrd3 | thioredoxin reductase 3 [Source:MGI Symbol;Acc:MGI:2386711] | 96,32 | 0,8 |
| ENSMUSG00000021710 | 166,5818845 | 264,3 | 139,7 | 7,168989901 | 0,004541667 | Nln | neurolysin (metallopeptidase M3 family) [Source:MGI Symbol;Acc:MGI:1923055] | 98,43 | 20,5 |
| ENSMUSG00000025648 | 57,73828708 | 87,4 | 0,9 | 7,170667432 | 0,008088217 | Pfkfb4 | 6-phosphofructo-2-kinase/fructose-2,6-biphosphatase 4 [Source:MGI Symbol;Acc:MGI:2687284] | 91,41 | 0,5 |
| ENSMUSG00000089917 | 132,6547786 | 230,3 | 79,5 | 7,172372339 | 0,005659757 | Uckl1 | uridine-cytidine kinase 1-like 1 [Source:MGI Symbol;Acc:MGI:1915806] | 79,41 | 17,9 |
| ENSMUSG00000000126 | 8,156074024 | 13,6 | 0,0 | 7,17920305 | 0,049495484 | Wnt9a | wingless-type MMTV integration site family, member 9A [Source:MGI Symbol;Acc:MGI:2446084] | 53,40 | 0,0 |
| ENSMUSG00000057816 | 8,331020388 | 14,1 | 0,0 | 7,181925451 | 0,047122655 | 1700007G11Rik | RIKEN cDNA 1700007G11 gene [Source:MGI Symbol;Acc:MGI:1916571] | 90,94 | 0,0 |
| ENSMUSG00000066151 | 74,15731332 | 87,3 | 0,9 | 7,184675887 | 0,010179082 | Fkbp15 | FK506 binding protein 15 [Source:MGI Symbol;Acc:MGI:2444782] | 74,82 | 0,4 |
| ENSMUSG00000086890 | 13,15523164 | 20,3 | 0,0 | 7,186063325 | 0,04176854 | 1700021J08Rik | RIKEN cDNA 1700021J08 gene [Source:MGI Symbol;Acc:MGI:1919485] | 48,64 | 0,0 |
| ENSMUSG00000052539 | 184,353009 | 323,9 | 3,6 | 7,189836632 | 0,001506164 | Magi3 | orane associated guanylate kinase, WW and PDZ domain containing 3 [Source:MGI Symbol;Acc:MGI:192 | 96,90 | 1,0 |
| ENSMUSG00000036411 | 42,58140436 | 95,6 | 0,6 | 7,19036478 | 0,002114832 | 9530077C05Rik | RIKEN cDNA 9530077C05 gene [Source:MGI Symbol;Acc:MGI:1915533] | 99,81 | 1,1 |
| ENSMUSG00000105265 | 150,4476426 | 198,6 | 37,9 | 7,193982794 | 0,005912613 | Sox2ot | SOX2 overlapping transcript (non-protein coding) [Source:MGI Symbol;Acc:MGI:2444112] | 44,72 | 5,9 |
| ENSMUSG00000048416 | 80,38984128 | 91,6 | 0,9 | 7,194199415 | 0,009302039 | Mlf1 | myeloid leukemia factor 1 [Source:MGI Symbol;Acc:MGI:1341819] | 94,76 | 1,7 |
| ENSMUSG00000031227 | 179,9057809 | 181,1 | 263,4 | 7,196448173 | 0,009108953 | Magee1 | melanoma antigen, family E, 1 [Source:MGI Symbol;Acc:MGI:2148149] | 99,52 | 29,7 |
| ENSMUSG00000023990 | 56,45103751 | 91,0 | 0,9 | 7,197208767 | 0,006895432 | Tfeb | transcription factor EB [Source:MGI Symbol;Acc:MGI:103270] | 44,11 | 0,5 |
| ENSMUSG00000032737 | 174,4779373 | 262,2 | 28,7 | 7,198032095 | 0,003260047 | Inppl1 | inositol polyphosphate phosphatase-like 1 [Source:MGI Symbol;Acc:MGI:1333787] | 98,11 | 10,4 |
| ENSMUSG00000025218 | 116,2949053 | 167,6 | 43,9 | 7,198837621 | 0,010820707 | Poll | polymerase (DNA directed), lambda [Source:MGI Symbol;Acc:MGI:1889000] | 99,76 | 12,3 |
| ENSMUSG00000026686 | 12,26143971 | 15,1 | 0,0 | 7,198887884 | 0,045829137 | Lmx1a | LIM homeobox transcription factor 1 alpha [Source:MGI Symbol;Acc:MGI:1888519] | 55,09 | 0,0 |
| ENSMUSG00000047635 | 136,1604579 | 202,5 | 39,7 | 7,200134778 | 0,008372435 | 2810006K23Rik | RIKEN cDNA 2810006K23 gene [Source:MGI Symbol;Acc:MGI:1919900] | 82,64 | 11,4 |
| ENSMUSG00000086741 | 7,520261938 | 15,3 | 0,0 | 7,204609316 | 0,044187226 | Gm15816 | predicted gene 15816 [Source:MGI Symbol;Acc:MGI:3802114] | 74,37 | 0,0 |
| ENSMUSG00000073367 | 7,471678088 | 13,8 | 0,0 | 7,205042164 | 0,046874479 | Gm10493 | predicted gene 10493 [Source:MGI Symbol;Acc:MGI:3644330] | #DIV/0! | #DIV/0! |
| ENSMUSG00000034709 | 165,7018421 | 286,0 | 108,9 | 7,208084916 | 0,003624055 | Ppp1r21 | protein phosphatase 1, regulatory subunit 21 [Source:MGI Symbol;Acc:MGI:1921075] | 96,60 | 26,9 |
| ENSMUSG00000097839 | 53,70835908 | 107,1 | 0,9 | 7,212821506 | 0,001437604 | Gm26909 | predicted gene, 26909 [Source:MGI Symbol;Acc:MGI:5477403] | #DIV/0! | #DIV/0! |
| ENSMUSG00000034544 | 256,8853926 | 428,4 | 4,0 | 7,215139036 | 0,001863752 | Rsrc1 | arginine/serine-rich coiled-coil 1 [Source:MGI Symbol;Acc:MGI:1914130] | 57,10 | 1,4 |
| ENSMUSG00000039084 | 6,314224766 | 13,7 | 0,0 | 7,215309462 | 0,049192431 | Chad | chondroadherin [Source:MGI Symbol;Acc:MGI:1096866] | 81,05 | 0,0 |
| ENSMUSG00000054715 | 159,2268261 | 218,9 | 264,4 | 7,216588686 | 0,004750039 | Zscan22 | zinc finger and SCAN domain containing 22 [Source:MGI Symbol;Acc:MGI:2443312] | 93,78 | 28,3 |
| ENSMUSG00000057329 | 165,8342996 | 301,8 | 2,5 | 7,217354098 | 0,000135047 | Bcl2 | B cell leukemia/lymphoma 2 [Source:MGI Symbol;Acc:MGI:88138] | 74,04 | 0,3 |
| ENSMUSG00000093485 | 6,745550523 | 14,3 | 0,0 | 7,217498798 | 0,047122655 | Gm20708 | predicted gene 20708 [Source:MGI Symbol;Acc:MGI:5313155] | 56,57 | 0,0 |
| ENSMUSG00000016552 | 47,81103738 | 105,5 | 0,9 | 7,218401247 | 0,001753896 | Foxred2 | FAD-dependent oxidoreductase domain containing 2 [Source:MGI Symbol;Acc:MGI:106315] | 94,04 | 0,4 |
| ENSMUSG00000034463 | 795,2566497 | 1272,9 | 320,9 | 7,21896488 | 0,00461289 | Scara3 | scavenger receptor class A, member 3 [Source:MGI Symbol;Acc:MGI:2444418] | 100,00 | 33,5 |
| ENSMUSG00000040856 | 9,400208673 | 14,6 | 0,0 | 7,219178446 | 0,044827322 | Dlk1 | delta-like 1 homolog (Drosophila) [Source:MGI Symbol;Acc:MGI:94900] | 22,00 | 0,0 |
| ENSMUSG00000069631 | 88,71729016 | 147,1 | 1,2 | 7,220181567 | 0,002179882 | Strada | STE20-related kinase adaptor alpha [Source:MGI Symbol;Acc:MGI:1919399] | 39,07 | 0,5 |
| ENSMUSG00000037752 | 10,33409703 | 13,9 | 0,0 | 7,226436254 | 0,04590072 | Xkr8 | X Kell blood group precursor related family member 8 homolog [Source:MGI Symbol;Acc:MGI:2685877] | 42,16 | 0,0 |
| ENSMUSG00000099094 | 6,785699481 | 16,0 | 0,0 | 7,22726116 | 0,049814539 | Gm19569 | predicted gene, 19569 [Source:MGI Symbol;Acc:MGI:5011754] | #DIV/0! | #DIV/0! |
| ENSMUSG00000094626 | 8,262707515 | 16,0 | 0,0 | 7,228059683 | 0,047939867 | Cecr6 | cat eye syndrome chromosome region, candidate 6 [Source:MGI Symbol;Acc:MGI:2136977] | 42,93 | 0,0 |
| ENSMUSG00000036120 | 264,0973177 | 452,2 | 55,5 | 7,231317165 | 4,21793E-05 | Rfxank | regulatory factor X-associated ankyrin-containing protein [Source:MGI Symbol;Acc:MGI:1333865] | 99,66 | 8,7 |
| ENSMUSG00000016382 | 314,1518169 | 329,1 | 207,6 | 7,233510555 | 0,000641627 | Pls3 | plastin 3 (T-isoform) [Source:MGI Symbol;Acc:MGI:104807] | 86,24 | 24,2 |
| ENSMUSG00000044287 | 6,879701561 | 14,4 | 0,0 | 7,233795805 | 0,049460093 | Nrn1l | neuritin 1-like [Source:MGI Symbol;Acc:MGI:2443642] | 93,52 | 0,0 |
| ENSMUSG00000003974 | 438,6457561 | 710,8 | 183,1 | 7,237895722 | 0,005428678 | Grm3 | glutamate receptor, metabotropic 3 [Source:MGI Symbol;Acc:MGI:1351340] | 100,00 | 32,9 |
| ENSMUSG00000086224 | 7,207582222 | 16,5 | 0,0 | 7,239259745 | 0,048859254 | 2700069I18Rik | RIKEN cDNA 2700069I18 gene [Source:MGI Symbol;Acc:MGI:1919858] | 64,06 | 0,0 |
| ENSMUSG00000029060 | 201,104952 | 223,3 | 101,7 | 7,240187394 | 0,008225786 | Mib2 | mindbomb homolog 2 (Drosophila) [Source:MGI Symbol;Acc:MGI:2679684] | 96,61 | 22,4 |
| ENSMUSG00000108616 | 53,50116037 | 79,2 | 0,6 | 7,242701752 | 0,016161315 | RP23-207N5.2 | | #DIV/0! | #DIV/0! |
| ENSMUSG00000026471 | 171,0911767 | 308,8 | 62,9 | 7,245552611 | 0,002221477 | Mr1 | major histocompatibility complex, class I-related [Source:MGI Symbol;Acc:MGI:1195463] | 100,00 | 21,2 |
| ENSMUSG00000097589 | 51,29317542 | 99,1 | 1,0 | 7,24805168 | 0,006792606 | Dleu2 | deleted in lymphocytic leukemia, 2 [Source:MGI Symbol;Acc:MGI:1934030] | 0,00 | 0,4 |
| ENSMUSG00000034006 | 113,2593243 | 168,3 | 49,9 | 7,252296129 | 0,011181037 | Pqlc1 | PQ loop repeat containing 1 [Source:MGI Symbol;Acc:MGI:1914193] | 64,40 | 16,8 |
| ENSMUSG00000025357 | 89,01617754 | 82,6 | 1,0 | 7,253292399 | 0,020239433 | Dgka | diacylglycerol kinase, alpha [Source:MGI Symbol;Acc:MGI:102952] | 98,65 | 0,9 |
| ENSMUSG00000035984 | 174,5081173 | 228,4 | 107,1 | 7,257223211 | 0,008002308 | Nme5 | NME/NM23 family member 5 [Source:MGI Symbol;Acc:MGI:1922783] | 99,57 | 20,9 |
| ENSMUSG00000042073 | 169,7810807 | 237,6 | 109,8 | 7,257934066 | 0,003823378 | Abhd14b | abhydrolase domain containing 14b [Source:MGI Symbol;Acc:MGI:1923741] | 92,95 | 25,0 |
| ENSMUSG00000025812 | 99,41659033 | 208,5 | 2,0 | 7,258695009 | 0,000218161 | Pard3 | par-3 family cell polarity regulator [Source:MGI Symbol;Acc:MGI:2135608] | 83,60 | 0,1 |
| ENSMUSG00000047787 | 11,48022043 | 19,0 | 0,0 | 7,259518285 | 0,044276202 | Flrt1 | fibronectin leucine rich transmembrane protein 1 [Source:MGI Symbol;Acc:MGI:3026647] | 31,96 | 0,0 |
| ENSMUSG00000056124 | 194,1006406 | 268,3 | 14,5 | 7,260490683 | 0,004068359 | B4galt6 | DP-Gal:betaGlcNAc beta 1,4-galactosyltransferase, polypeptide 6 [Source:MGI Symbol;Acc:MGI:192838( | 97,92 | 4,4 |
| ENSMUSG00000004637 | 156,204223 | 183,8 | 42,9 | 7,262605246 | 0,005575965 | Wwox | WW domain-containing oxidoreductase [Source:MGI Symbol;Acc:MGI:1931237] | 44,16 | 5,2 |
| ENSMUSG00000028099 | 164,6010573 | 204,6 | 7,4 | 7,263023866 | 0,004084001 | Polr3c | polymerase (RNA) III (DNA directed) polypeptide C [Source:MGI Symbol;Acc:MGI:1921664] | 93,35 | 2,6 |
| ENSMUSG00000108624 | 6,328215326 | 14,8 | 0,0 | 7,263971204 | 0,049415649 | RP23-435E16.3 | | 98,02 | 0,0 |
| ENSMUSG00000011589 | 50,38140163 | 101,6 | 1,0 | 7,267059105 | 0,005219865 | Fsd1 | fibronectin type 3 and SPRY domain-containing protein [Source:MGI Symbol;Acc:MGI:1934858] | 99,73 | 1,4 |
| ENSMUSG00000097274 | 152,5971215 | 218,8 | 28,7 | 7,267635514 | 0,00516674 | Gm26564 | predicted gene, 26564 [Source:MGI Symbol;Acc:MGI:5477058] | #DIV/0! | #DIV/0! |
| ENSMUSG00000047473 | 52,08887207 | 101,9 | 0,9 | 7,268140368 | 0,006720124 | Zfp30 | zinc finger protein 30 [Source:MGI Symbol;Acc:MGI:99178] | 85,05 | 0,5 |
| ENSMUSG00000027875 | 99,62830856 | 167,7 | 1,8 | 7,270867548 | 0,001037497 | Hmgcs2 | 3-hydroxy-3-methylglutaryl-Coenzyme A synthase 2 [Source:MGI Symbol;Acc:MGI:101939] | 96,19 | 0,7 |
| ENSMUSG00000024087 | 138,1711561 | 156,0 | 326,3 | 7,271142942 | 0,015131294 | Cyp1b1 | cytochrome P450, family 1, subfamily b, polypeptide 1 [Source:MGI Symbol;Acc:MGI:88590] | 94,00 | 25,6 |
| ENSMUSG00000024903 | 6,601682048 | 15,3 | 0,0 | 7,271204239 | 0,048733912 | Lao1 | L-amino acid oxidase 1 [Source:MGI Symbol;Acc:MGI:2140628] | 67,56 | 0,0 |
| ENSMUSG00000050379 | 218,3911183 | 245,0 | 7,0 | 7,271368296 | 0,00083198 | Sept6 | septin 6 [Source:MGI Symbol;Acc:MGI:1888939] | 79,74 | 0,9 |
| ENSMUSG00000061207 | 117,1263777 | 193,7 | 2,8 | 7,272111373 | 0,000806125 | Stk19 | serine/threonine kinase 19 [Source:MGI Symbol;Acc:MGI:1860085] | 95,09 | 4,7 |
| ENSMUSG00000066441 | 157,2880937 | 230,0 | 21,3 | 7,272334735 | 0,005275873 | Rdh11 | retinol dehydrogenase 11 [Source:MGI Symbol;Acc:MGI:102581] | 83,30 | 5,6 |
| ENSMUSG00000097613 | 6,740264595 | 14,7 | 0,0 | 7,272545576 | 0,048492727 | Gm17597 | predicted gene, 17597 [Source:MGI Symbol;Acc:MGI:4937231] | #DIV/0! | #DIV/0! |
| ENSMUSG00000047242 | 649,9223064 | 1020,8 | 298,1 | 7,276453746 | 0,005222124 | Taf9b | RNA polymerase II, TATA box binding protein (TBP)-associated factor [Source:MGI Symbol;Acc:MGI:30: | 83,10 | 25,2 |
| ENSMUSG00000108832 | 24,43954729 | 14,9 | 0,0 | 7,277116615 | 0,049283007 | RP24-325H17.1 | | 34,14 | 0,0 |
| ENSMUSG00000019975 | 182,9111368 | 274,7 | 134,4 | 7,27764753 | 0,004936283 | Ikbip | IKBKB interacting protein [Source:MGI Symbol;Acc:MGI:1914704] | 96,30 | 9,6 |
| ENSMUSG00000029916 | 73,38146974 | 107,6 | 1,0 | 7,284692003 | 0,004092403 | Agk | acylglycerol kinase [Source:MGI Symbol;Acc:MGI:1917173] | 68,33 | 0,5 |
| ENSMUSG00000054733 | 167,0126981 | 237,4 | 53,6 | 7,288238526 | 0,003029261 | Msra | methionine sulfoxide reductase A [Source:MGI Symbol;Acc:MGI:106916] | 71,75 | 18,6 |
| ENSMUSG00000049721 | 30,24535537 | 17,3 | 0,0 | 7,289320824 | 0,046384855 | Gal3st1 | galactose-3-O-sulfotransferase 1 [Source:MGI Symbol;Acc:MGI:1858277] | 67,22 | 0,0 |
| ENSMUSG00000068299 | 495,2325802 | 895,1 | 382,9 | 7,290950822 | 0,003685841 | 1700019G17Rik | RIKEN cDNA 1700019G17 gene [Source:MGI Symbol;Acc:MGI:1922791] | 90,63 | 30,3 |
| ENSMUSG00000021408 | 49,13578696 | 105,8 | 1,0 | 7,295103502 | 0,003629297 | Ripk1 | receptor (TNFRSF)-interacting serine-threonine kinase 1 [Source:MGI Symbol;Acc:MGI:108212] | 94,41 | 0,1 |
| ENSMUSG00000047992 | 112,3638602 | 209,9 | 38,8 | 7,300252932 | 0,007140229 | Fam69c | family with sequence similarity 69, member C [Source:MGI Symbol;Acc:MGI:3041188] | 94,87 | 16,3 |
| ENSMUSG00000024065 | 208,7990454 | 405,3 | 3,6 | 7,300573526 | 1,15387E-05 | Ehd3 | EH-domain containing 3 [Source:MGI Symbol;Acc:MGI:1928900] | 99,76 | 1,3 |
| ENSMUSG00000054676 | 229,9833745 | 335,5 | 76,8 | 7,306426235 | 0,000417252 | 1600014C10Rik | RIKEN cDNA 1600014C10 gene [Source:MGI Symbol;Acc:MGI:1919494] | 83,30 | 8,8 |
| ENSMUSG00000036019 | 8,161464182 | 15,6 | 0,0 | 7,31377196 | 0,045269835 | Tmtc2 | transmembrane and tetratricopeptide repeat containing 2 [Source:MGI Symbol;Acc:MGI:1914057] | 34,96 | 0,0 |
| ENSMUSG00000050270 | 10,85960959 | 20,0 | 0,0 | 7,315068825 | 0,042818634 | Tmem220 | transmembrane protein 220 [Source:MGI Symbol;Acc:MGI:2443691] | 70,38 | 0,0 |
| ENSMUSG00000025981 | 283,8241608 | 410,6 | 101,1 | 7,315510076 | 0,000907964 | Coq10b | coenzyme Q10B [Source:MGI Symbol;Acc:MGI:1915126] | 95,92 | 30,3 |
| ENSMUSG00000044320 | 22,37987972 | 19,3 | 0,0 | 7,31751163 | 0,044863632 | 1700001O22Rik | RIKEN cDNA 1700001O22 gene [Source:MGI Symbol;Acc:MGI:1923631] | 65,45 | 0,0 |
| ENSMUSG00000035179 | 8,580113625 | 16,1 | 0,0 | 7,317571129 | 0,044667804 | Ppp1r32 | protein phosphatase 1, regulatory subunit 32 [Source:MGI Symbol;Acc:MGI:1915002] | 82,03 | 0,0 |
| ENSMUSG00000021065 | 502,168909 | 829,5 | 73,1 | 7,318091804 | 0,000378972 | Fut8 | fucosyltransferase 8 [Source:MGI Symbol;Acc:MGI:1858901] | 94,83 | 20,1 |
| ENSMUSG00000002769 | 20,19820001 | 15,5 | 0,0 | 7,318781303 | 0,03769351 | Gnmt | glycine N-methyltransferase [Source:MGI Symbol;Acc:MGI:1202304] | 88,86 | 0,0 |
| ENSMUSG00000042015 | 290,7407715 | 456,5 | 114,4 | 7,318866852 | 0,000903303 | Wdr41 | WD repeat domain 41 [Source:MGI Symbol;Acc:MGI:2445123] | 60,47 | 17,6 |
| ENSMUSG00000045639 | 112,2942356 | 141,9 | 1,8 | 7,324313244 | 0,004555798 | Zfp629 | zinc finger protein 629 [Source:MGI Symbol;Acc:MGI:2444524] | 82,62 | 0,7 |
| ENSMUSG00000085030 | 6,04046577 | 16,6 | 0,0 | 7,324702532 | 0,048089003 | 2810455O05Rik | RIKEN cDNA 2810455O05 gene [Source:MGI Symbol;Acc:MGI:1920072] | 49,42 | 0,0 |
| ENSMUSG00000032220 | 75,46547651 | 111,2 | 1,0 | 7,326072479 | 0,00351257 | Myo1e | myosin IE [Source:MGI Symbol;Acc:MGI:106621] | 93,03 | 0,4 |
| ENSMUSG00000074406 | 15,12630922 | 14,9 | 0,0 | 7,327577292 | 0,049215399 | Zfp628 | zinc finger protein 628 [Source:MGI Symbol;Acc:MGI:2665174] | 51,22 | 0,0 |
| ENSMUSG00000091636 | 11,75228735 | 15,7 | 0,0 | 7,328226397 | 0,04176854 | A330050F15Rik | RIKEN cDNA A330050F15 gene [Source:MGI Symbol;Acc:MGI:2444600] | 65,17 | 0,0 |
| ENSMUSG00000019590 | 421,3197419 | 486,2 | 199,6 | 7,329185384 | 0,007671013 | Cyb561 | cytochrome b-561 [Source:MGI Symbol;Acc:MGI:103253] | 85,24 | 24,2 |
| ENSMUSG00000074607 | 217,9921559 | 394,0 | 237,2 | 7,329629342 | 0,00328299 | Tox2 | TOX high mobility group box family member 2 [Source:MGI Symbol;Acc:MGI:3611233] | 89,04 | 28,7 |
| ENSMUSG00000073968 | 8,91105735 | 17,2 | 0,0 | 7,331964136 | 0,043851134 | Trim68 | tripartite motif-containing 68 [Source:MGI Symbol;Acc:MGI:2142077] | 65,56 | 0,0 |
| ENSMUSG00000040219 | 16,15517356 | 24,6 | 0,0 | 7,333232713 | 0,048552305 | Ttc12 | tetratricopeptide repeat domain 12 [Source:MGI Symbol;Acc:MGI:2444588] | 51,56 | 0,0 |
| ENSMUSG00000104684 | 7,27201969 | 15,0 | 0,0 | 7,333989077 | 0,045654899 | 5430427N15Rik | RIKEN cDNA 5430427N15 gene [Source:MGI Symbol;Acc:MGI:2441699] | #DIV/0! | #DIV/0! |
| ENSMUSG00000040482 | 70,25451192 | 93,3 | 0,6 | 7,335605223 | 0,006782761 | Dxo | decapping exoribonuclease [Source:MGI Symbol;Acc:MGI:1890444] | 85,36 | 1,3 |
| ENSMUSG00000030555 | 104,0126643 | 199,0 | 2,4 | 7,335676304 | 0,002001696 | Ttc23 | tetratricopeptide repeat domain 23 [Source:MGI Symbol;Acc:MGI:1914259] | 86,53 | 0,0 |
| ENSMUSG00000061510 | 8,259179024 | 16,8 | 0,0 | 7,337019935 | 0,049283007 | Gm10101 | predicted gene 10101 [Source:MGI Symbol;Acc:MGI:3641713] | #DIV/0! | #DIV/0! |
| ENSMUSG00000021130 | 327,3942485 | 316,1 | 565,9 | 7,337592244 | 0,003307477 | Galnt16 | tyl-alpha-D-galactosamine:polypeptide N-acetylgalactosaminyltransferase 16 [Source:MGI Symbol;Acc:MG | 98,66 | 32,8 |
| ENSMUSG00000017491 | 141,003315 | 227,4 | 1,5 | 7,337969433 | 0,001015712 | Rarb | retinoic acid receptor, beta [Source:MGI Symbol;Acc:MGI:97857] | 99,97 | 1,3 |
| ENSMUSG00000050103 | 159,8025986 | 224,2 | 52,7 | 7,339056853 | 0,008924977 | Agmo | alkylglycerol monooxygenase [Source:MGI Symbol;Acc:MGI:2442495] | 63,84 | 8,4 |
| ENSMUSG00000029122 | 126,6342511 | 158,7 | 59,1 | 7,342361888 | 0,016189098 | Evc | Ellis van Creveld gene syndrome [Source:MGI Symbol;Acc:MGI:1890596] | 81,46 | 14,8 |
| ENSMUSG00000068962 | 8,106556821 | 18,7 | 0,0 | 7,342909411 | 0,049691937 | Zfp114 | zinc finger protein 114 [Source:MGI Symbol;Acc:MGI:3037815] | 32,70 | 0,0 |
| ENSMUSG00000025083 | 65,41423078 | 89,5 | 0,6 | 7,347438608 | 0,007785817 | Afap1l2 | actin filament associated protein 1-like 2 [Source:MGI Symbol;Acc:MGI:2147658] | 78,48 | 0,5 |
| ENSMUSG00000082163 | 7,109249308 | 15,5 | 0,0 | 7,347464211 | 0,045516692 | Gm14276 | predicted gene 14276 [Source:MGI Symbol;Acc:MGI:3701955] | #DIV/0! | #DIV/0! |
| ENSMUSG00000031242 | 184,0392126 | 207,3 | 2,0 | 7,352090115 | 9,23286E-05 | 2610002M06Rik | RIKEN cDNA 2610002M06 gene [Source:MGI Symbol;Acc:MGI:1914278] | 91,57 | 0,4 |
| ENSMUSG00000037762 | 76,98903707 | 102,0 | 1,0 | 7,359978463 | 0,010865456 | Slc16a9 | lute carrier family 16 (monocarboxylic acid transporters), member 9 [Source:MGI Symbol;Acc:MGI:19141( | 89,96 | 0,4 |
| ENSMUSG00000030095 | 240,6386413 | 190,9 | 210,6 | 7,367348671 | 0,014040727 | Tmem43 | transmembrane protein 43 [Source:MGI Symbol;Acc:MGI:1921372] | 99,51 | 30,4 |
| ENSMUSG00000085396 | 178,2141696 | 19,7 | 0,0 | 7,367579044 | 0,045791107 | Firre | functional intergenic repeating RNA element [Source:MGI Symbol;Acc:MGI:2147989] | #DIV/0! | #DIV/0! |
| ENSMUSG00000097187 | 9,363720876 | 15,6 | 0,0 | 7,368469022 | 0,041791906 | Gm19426 | predicted gene, 19426 [Source:MGI Symbol;Acc:MGI:5011611] | 93,43 | 0,0 |
| ENSMUSG00000069520 | 221,578552 | 287,4 | 2,8 | 7,368687744 | 4,88823E-06 | Tmem19 | transmembrane protein 19 [Source:MGI Symbol;Acc:MGI:1914476] | 99,50 | 2,4 |
| ENSMUSG00000019055 | 132,5098236 | 268,4 | 1,6 | 7,369361319 | 0,000247693 | Plod1 | procollagen-lysine, 2-oxoglutarate 5-dioxygenase 1 [Source:MGI Symbol;Acc:MGI:99907] | 92,89 | 1,1 |
| ENSMUSG00000030314 | 301,8112078 | 475,6 | 111,4 | 7,370043266 | 0,000782874 | Atg7 | autophagy related 7 [Source:MGI Symbol;Acc:MGI:1921494] | 82,23 | 17,6 |
| ENSMUSG00000074829 | 222,9480823 | 350,6 | 133,1 | 7,371730295 | 0,002135809 | 2010315B03Rik | RIKEN cDNA 2010315B03 gene [Source:MGI Symbol;Acc:MGI:1919321] | 99,60 | 29,4 |
| ENSMUSG00000059027 | 9,442884365 | 15,8 | 0,0 | 7,374570641 | 0,041952726 | 9630013D21Rik | RIKEN cDNA 9630013D21 gene [Source:MGI Symbol;Acc:MGI:2442649] | #DIV/0! | #DIV/0! |
| ENSMUSG00000068270 | 11,18691649 | 17,0 | 0,0 | 7,377578231 | 0,049526667 | Shroom4 | shroom family member 4 [Source:MGI Symbol;Acc:MGI:2685570] | 25,46 | 0,0 |
| ENSMUSG00000085615 | 8,228122177 | 15,8 | 0,0 | 7,378764644 | 0,043075759 | A330035P11Rik | RIKEN cDNA A330035P11 gene [Source:MGI Symbol;Acc:MGI:2442610] | #DIV/0! | #DIV/0! |
| ENSMUSG00000044835 | 155,8894084 | 240,3 | 68,9 | 7,379290148 | 0,00451739 | Ankrd45 | ankyrin repeat domain 45 [Source:MGI Symbol;Acc:MGI:1921094] | 99,32 | 17,1 |
| ENSMUSG00000042289 | 151,4847587 | 217,3 | 7,3 | 7,381290717 | 0,00551827 | Hsd3b7 | y-delta-5-steroid dehydrogenase, 3 beta- and steroid delta-isomerase 7 [Source:MGI Symbol;Acc:MGI:21 | 88,05 | 4,2 |
| ENSMUSG00000017943 | 122,6826742 | 222,2 | 83,8 | 7,383472233 | 0,005219865 | Gdap1l1 | ganglioside-induced differentiation-associated protein 1-like 1 [Source:MGI Symbol;Acc:MGI:2385163] | 91,95 | 20,8 |
| ENSMUSG00000078963 | 8,138298117 | 18,8 | 0,0 | 7,384653929 | 0,043040966 | Hsbp1l1 | heat shock factor binding protein 1-like 1 [Source:MGI Symbol;Acc:MGI:1913505] | 60,48 | 0,0 |
| ENSMUSG00000033721 | 57,52264494 | 114,8 | 0,6 | 7,384773735 | 0,00151093 | Vav3 | vav 3 oncogene [Source:MGI Symbol;Acc:MGI:1888518] | 90,42 | 0,4 |
| ENSMUSG00000020844 | 185,9710834 | 268,1 | 51,8 | 7,387368803 | 0,003081179 | Nxn | nucleoredoxin [Source:MGI Symbol;Acc:MGI:109331] | 95,35 | 18,8 |
| ENSMUSG00000047604 | 9,85832913 | 16,0 | 0,0 | 7,388580904 | 0,04128043 | Frat2 | frequently rearranged in advanced T cell lymphomas 2 [Source:MGI Symbol;Acc:MGI:2673967] | 68,51 | 0,0 |
| ENSMUSG00000074876 | 9,171311263 | 16,1 | 0,0 | 7,390225112 | 0,045877678 | Spata5l1 | spermatogenesis associated 5-like 1 [Source:MGI Symbol;Acc:MGI:3036261] | #DIV/0! | #DIV/0! |
| ENSMUSG00000020878 | 10,64832496 | 15,9 | 0,0 | 7,390348598 | 0,03857125 | Lrrc46 | leucine rich repeat containing 46 [Source:MGI Symbol;Acc:MGI:1916547] | 89,70 | 0,0 |
| ENSMUSG00000034850 | 195,9435706 | 267,5 | 87,8 | 7,392320778 | 0,008502063 | Tmem127 | transmembrane protein 127 [Source:MGI Symbol;Acc:MGI:1916720] | 98,04 | 10,9 |
| ENSMUSG00000021365 | 108,7698879 | 126,4 | 1,2 | 7,392789993 | 0,009395344 | Nedd9 | ural precursor cell expressed, developmentally down-regulated gene 9 [Source:MGI Symbol;Acc:MGI:973( | 90,97 | 0,5 |
| ENSMUSG00000108621 | 58,44986687 | 115,2 | 0,9 | 7,395068451 | 0,001729152 | Gm37494 | predicted gene, 37494 [Source:MGI Symbol;Acc:MGI:5610722] | #DIV/0! | #DIV/0! |
| ENSMUSG00000033454 | 87,19168701 | 110,4 | 0,6 | 7,397089638 | 0,005224013 | Zbtb1 | zinc finger and BTB domain containing 1 [Source:MGI Symbol;Acc:MGI:2442326] | 40,50 | 0,2 |
| ENSMUSG00000050587 | 196,0084094 | 230,7 | 7,4 | 7,399467078 | 0,003286802 | Lrrc4c | leucine rich repeat containing 4C [Source:MGI Symbol;Acc:MGI:2442636] | 92,25 | 2,7 |
| ENSMUSG00000097049 | 11,05049583 | 16,5 | 0,0 | 7,400734826 | 0,044692 | 6530411M01Rik | RIKEN cDNA 6530411M01 gene [Source:MGI Symbol;Acc:MGI:1915041] | #DIV/0! | #DIV/0! |
| ENSMUSG00000042540 | 7,712410515 | 15,5 | 0,0 | 7,401403961 | 0,04296762 | Acot5 | acyl-CoA thioesterase 5 [Source:MGI Symbol;Acc:MGI:2384969] | 72,13 | 0,0 |
| ENSMUSG00000075020 | 12,46532893 | 16,0 | 0,0 | 7,402986991 | 0,045847926 | E530001K10Rik | RIKEN cDNA E530001K10 gene [Source:MGI Symbol;Acc:MGI:3041234] | #DIV/0! | #DIV/0! |
| ENSMUSG00000024477 | 80,45728729 | 135,2 | 1,2 | 7,405142327 | 0,006284954 | Pggt1b | protein geranylgeranyltransferase type I, beta subunit [Source:MGI Symbol;Acc:MGI:1917514] | 98,67 | 0,9 |
| ENSMUSG00000027087 | 229,6991072 | 168,6 | 1,8 | 7,405548434 | 0,005764095 | Itgav | integrin alpha V [Source:MGI Symbol;Acc:MGI:96608] | 84,37 | 0,3 |
| ENSMUSG00000091329 | 10,09566287 | 15,9 | 0,0 | 7,407271592 | 0,045673564 | 1700112D23Rik | RIKEN cDNA 1700112D23 gene [Source:MGI Symbol;Acc:MGI:1920861] | 79,85 | 0,0 |
| ENSMUSG00000022799 | 110,0886274 | 161,9 | 2,0 | 7,409247302 | 0,002292486 | Arhgap31 | Rho GTPase activating protein 31 [Source:MGI Symbol;Acc:MGI:1333857] | 85,79 | 0,0 |
| ENSMUSG00000029176 | 193,9639553 | 308,3 | 64,7 | 7,409984488 | 0,000907964 | Anapc4 | anaphase promoting complex subunit 4 [Source:MGI Symbol;Acc:MGI:1098673] | 87,24 | 9,0 |
| ENSMUSG00000067879 | 146,2915458 | 266,1 | 143,7 | 7,411079175 | 0,003119232 | 3110035E14Rik | RIKEN cDNA 3110035E14 gene [Source:MGI Symbol;Acc:MGI:1924232] | 99,79 | 18,3 |
| ENSMUSG00000087413 | 129,8488803 | 183,1 | 126,7 | 7,412374955 | 0,013575227 | Gm11266 | predicted gene 11266 [Source:MGI Symbol;Acc:MGI:3649705] | 100,00 | 32,9 |
| ENSMUSG00000107605 | 5,701276696 | 16,6 | 0,0 | 7,412538462 | 0,045981415 | Gm44117 | predicted gene, 44117 [Source:MGI Symbol;Acc:MGI:5690509] | #DIV/0! | #DIV/0! |
| ENSMUSG00000036766 | 199,8795026 | 280,9 | 98,9 | 7,416752776 | 0,00427333 | Dner | delta/notch-like EGF repeat containing [Source:MGI Symbol;Acc:MGI:2152889] | 84,29 | 22,0 |
| ENSMUSG00000104729 | 9,907591873 | 16,0 | 0,0 | 7,417278706 | 0,048189025 | Gm44471 | predicted gene, 44471 [Source:MGI Symbol;Acc:MGI:5690863] | #DIV/0! | #DIV/0! |
| ENSMUSG00000030281 | 133,1022751 | 195,1 | 140,7 | 7,420720482 | 0,01090299 | Il17rc | interleukin 17 receptor C [Source:MGI Symbol;Acc:MGI:2159336] | 97,77 | 29,7 |
| ENSMUSG00000030823 | 20,3643633 | 18,3 | 0,0 | 7,422055487 | 0,04083678 | 9130019O22Rik | RIKEN cDNA 9130019O22 gene [Source:MGI Symbol;Acc:MGI:1926171] | 48,51 | 0,0 |
| ENSMUSG00000003458 | 238,0363719 | 319,5 | 26,6 | 7,422625367 | 0,001089664 | Ncstn | nicastrin [Source:MGI Symbol;Acc:MGI:1891700] | 99,02 | 11,0 |
| ENSMUSG00000087620 | 9,794819896 | 15,8 | 0,0 | 7,422975353 | 0,038128913 | 5330434G04Rik | RIKEN cDNA 5330434G04 gene [Source:MGI Symbol;Acc:MGI:1925519] | #DIV/0! | #DIV/0! |
| ENSMUSG00000041654 | 158,6932478 | 176,7 | 2,8 | 7,422997603 | 0,004736532 | Slc39a11 | solute carrier family 39 (metal ion transporter), member 11 [Source:MGI Symbol;Acc:MGI:1917056] | 86,11 | 2,2 |
| ENSMUSG00000027832 | 18,55406233 | 18,3 | 0,0 | 7,425416267 | 0,045609433 | Ptx3 | pentraxin related gene [Source:MGI Symbol;Acc:MGI:104641] | 45,31 | 0,0 |
| ENSMUSG00000085058 | 7,964701987 | 16,5 | 0,0 | 7,426388505 | 0,04189693 | 8030453O22Rik | RIKEN cDNA 8030453O22 gene [Source:MGI Symbol;Acc:MGI:1924459] | 78,17 | 0,0 |
| ENSMUSG00000005917 | 110,7771498 | 212,6 | 2,0 | 7,429230727 | 0,000123894 | Otx1 | orthodenticle homolog 1 [Source:MGI Symbol;Acc:MGI:97450] | 96,40 | 0,8 |
| ENSMUSG00000023919 | 172,6775639 | 312,5 | 2,5 | 7,429288051 | 0,00161365 | Cenpq | centromere protein Q [Source:MGI Symbol;Acc:MGI:1933744] | 95,73 | 1,3 |
| ENSMUSG00000027305 | 136,5215355 | 244,4 | 3,0 | 7,43189153 | 0,000991795 | Ndufaf1 | H dehydrogenase (ubiquinone) 1 alpha subcomplex, assembly factor 1 [Source:MGI Symbol;Acc:MGI:191( | 99,95 | 3,2 |
| ENSMUSG00000055435 | 22,18027375 | 21,2 | 0,0 | 7,431920551 | 0,041940594 | Maf | an musculoaponeurotic fibrosarcoma (v-maf) AS42 oncogene homolog [Source:MGI Symbol;Acc:MGI:969 | 22,67 | 0,0 |
| ENSMUSG00000097836 | 7,420760533 | 16,1 | 0,0 | 7,432365105 | 0,042379612 | Gm26903 | predicted gene, 26903 [Source:MGI Symbol;Acc:MGI:5477397] | 17,72 | 0,0 |
| ENSMUSG00000035692 | 15,00045794 | 18,2 | 0,0 | 7,432660231 | 0,047394949 | Isg15 | ISG15 ubiquitin-like modifier [Source:MGI Symbol;Acc:MGI:1855694] | 78,09 | 0,0 |
| ENSMUSG00000094002 | 7,988442153 | 17,3 | 0,0 | 7,433655878 | 0,041742988 | Gm9866 | predicted gene 9866 [Source:MGI Symbol;Acc:MGI:3643118] | #DIV/0! | #DIV/0! |
| ENSMUSG00000024963 | 115,8643231 | 154,2 | 1,8 | 7,435636466 | 0,003706942 | Dnajc4 | DnaJ heat shock protein family (Hsp40) member C4 [Source:MGI Symbol;Acc:MGI:1927346] | 97,70 | 2,0 |
| ENSMUSG00000021671 | 162,3531416 | 245,2 | 74,9 | 7,437404502 | 0,008920937 | Poc5 | POC5 centriolar protein homolog (Chlamydomonas) [Source:MGI Symbol;Acc:MGI:1914713] | 99,88 | 23,9 |
| ENSMUSG00000041012 | 22,56404045 | 22,2 | 0,0 | 7,443933767 | 0,04304535 | Cmtm8 | CKLF-like MARVEL transmembrane domain containing 8 [Source:MGI Symbol;Acc:MGI:2447167] | 89,85 | 0,0 |
| ENSMUSG00000104822 | 12,72264751 | 16,3 | 0,0 | 7,445562251 | 0,04865636 | Gm42967 | predicted gene 42967 [Source:MGI Symbol;Acc:MGI:5663104] | 11,03 | 0,0 |
| ENSMUSG00000028476 | 12,57217383 | 18,3 | 0,0 | 7,446736934 | 0,033679154 | Reck | reversion-inducing-cysteine-rich protein with kazal motifs [Source:MGI Symbol;Acc:MGI:1855698] | 46,56 | 0,0 |

| | | | | | | | | | |
|---|---|---|---|---|---|---|---|---|---|
| ENSMUSG00000060336 | 21,60952521 | 16,7 | 0,0 | 7,447478873 | 0,046938122 | Zfp937 | zinc finger protein 937 [Source:MGI Symbol;Acc:MGI:3644476] | 57,60 | 0,0 |
| ENSMUSG00000034758 | 38,1054684 | 16,9 | 0,0 | 7,448765782 | 0,042808065 | Tle6 | transducin-like enhancer of split 6, homolog of Drosophila E(spl) [Source:MGI Symbol;Acc:MGI:2149593] | 63,14 | 0,0 |
| ENSMUSG00000048108 | 113,1685758 | 119,9 | 0,6 | 7,450781823 | 0,014935367 | Tmem72 | transmembrane protein 72 [Source:MGI Symbol;Acc:MGI:2442707] | 31,41 | 0,1 |
| ENSMUSG00000097102 | 154,8415306 | 241,2 | 128,9 | 7,452573165 | 0,012357059 | 2310069G16Rik | RIKEN cDNA 2310069G16 gene [Source:MGI Symbol;Acc:MGI:1916909] | #DIV/0! | #DIV/0! |
| ENSMUSG00000091971 | 6,605373678 | 16,7 | 0,0 | 7,453683346 | 0,04338551 | Hspa1a | heat shock protein 1A [Source:MGI Symbol;Acc:MGI:96244] | 38,73 | 0,0 |
| ENSMUSG00000104283 | 15,83016006 | 17,9 | 0,0 | 7,45513435 | 0,04043499 | Gm37459 | predicted gene, 37459 [Source:MGI Symbol;Acc:MGI:5610687] | 11,26 | 0,0 |
| ENSMUSG00000044340 | 455,3316389 | 681,4 | 36,6 | 7,456646219 | 0,001647634 | Phlpp1 | PH domain and leucine rich repeat protein phosphatase 1 [Source:MGI Symbol;Acc:MGI:2138327] | 98,28 | 5,7 |
| ENSMUSG00000094320 | 10,56736726 | 16,6 | 0,0 | 7,456691221 | 0,04124584 | Gm13202 | predicted gene 13202 [Source:MGI Symbol;Acc:MGI:3649942] | #DIV/0! | #DIV/0! |
| ENSMUSG00000036916 | 13,11851859 | 25,7 | 0,0 | 7,45701398 | 0,028969349 | Zfp280c | zinc finger protein 280C [Source:MGI Symbol;Acc:MGI:2387585] | 49,85 | 0,0 |
| ENSMUSG00000022358 | 18,81862008 | 20,5 | 0,0 | 7,457540542 | 0,038037761 | Fbxo32 | F-box protein 32 [Source:MGI Symbol;Acc:MGI:1914981] | 28,87 | 0,0 |
| ENSMUSG00000039891 | 13,19124629 | 20,2 | 0,0 | 7,458965925 | 0,044827322 | Txlnb | taxilin beta [Source:MGI Symbol;Acc:MGI:2671945] | 44,63 | 0,0 |
| ENSMUSG00000026688 | 130,8020299 | 233,2 | 18,0 | 7,46264555 | 0,004602677 | Mgst3 | microsomal glutathione S-transferase 3 [Source:MGI Symbol;Acc:MGI:1913697] | 95,93 | 17,0 |
| ENSMUSG00000103366 | 8,339195916 | 16,7 | 0,0 | 7,462993116 | 0,044965954 | C630004L07Rik | RIKEN cDNA C630004L07 gene [Source:MGI Symbol;Acc:MGI:3641643] | 51,27 | 0,0 |
| ENSMUSG00000054381 | 13,29729777 | 17,6 | 0,0 | 7,464064788 | 0,038356903 | Zfp747 | zinc finger protein 747 [Source:MGI Symbol;Acc:MGI:2443581] | 65,15 | 0,0 |
| ENSMUSG00000029596 | 183,534315 | 321,2 | 98,0 | 7,467149537 | 0,004357665 | Sdsl | serine dehydratase-like [Source:MGI Symbol;Acc:MGI:2182607] | 83,50 | 27,5 |
| ENSMUSG00000031796 | 260,1657842 | 416,2 | 30,5 | 7,468126225 | 0,000118074 | Cfap20 | cilia and flagella associated protein 20 [Source:MGI Symbol;Acc:MGI:107428] | 98,45 | 27,0 |
| ENSMUSG00000042249 | 259,567523 | 337,6 | 223,9 | 7,469512948 | 0,005408155 | Adrbk2 | adrenergic receptor kinase, beta 2 [Source:MGI Symbol;Acc:MGI:87941] | 84,21 | 23,5 |
| ENSMUSG00000039043 | 216,2644681 | 397,8 | 204,6 | 7,469764249 | 0,001337014 | Arpin | actin-related protein 2/3 complex inhibitor [Source:MGI Symbol;Acc:MGI:1917670] | 97,69 | 25,9 |
| ENSMUSG00000021725 | 72,50816597 | 108,9 | 0,6 | 7,470229815 | 0,00330193 | Parp8 | poly (ADP-ribose) polymerase family, member 8 [Source:MGI Symbol;Acc:MGI:1098713] | 95,13 | 0,8 |
| ENSMUSG00000042826 | 10,88753478 | 20,0 | 0,0 | 7,474879521 | 0,035299784 | Fgf11 | fibroblast growth factor 11 [Source:MGI Symbol;Acc:MGI:109167] | 60,36 | 0,0 |
| ENSMUSG00000097929 | 8,240230256 | 16,4 | 0,0 | 7,476597788 | 0,04024368 | Tunar | Tcl1 upstream neural differentiation associated RNA [Source:MGI Symbol;Acc:MGI:1917202] | #DIV/0! | #DIV/0! |
| ENSMUSG00000035773 | 13,36979439 | 16,6 | 0,0 | 7,480303036 | 0,040811789 | Kiss1r | KISS1 receptor [Source:MGI Symbol;Acc:MGI:2148793] | 59,54 | 0,0 |
| ENSMUSG00000037738 | 16,23968693 | 18,6 | 0,0 | 7,48074026 | 0,035858194 | Nek5 | NIMA (never in mitosis gene a)-related expressed kinase 5 [Source:MGI Symbol;Acc:MGI:2142824] | 61,54 | 0,0 |
| ENSMUSG00000030643 | 200,514088 | 335,9 | 50,8 | 7,483164598 | 0,002914962 | Rab30 | RAB30, member RAS oncogene family [Source:MGI Symbol;Acc:MGI:1923235] | 79,28 | 4,2 |
| ENSMUSG00000006143 | 7,259478347 | 17,8 | 0,0 | 7,484235252 | 0,041157959 | Upk3bl | uroplakin 3B-like [Source:MGI Symbol;Acc:MGI:1916915] | 56,05 | 0,0 |
| ENSMUSG00000097339 | 9,28759814 | 16,9 | 0,0 | 7,485213815 | 0,039016968 | Gm26671 | predicted gene, 26671 [Source:MGI Symbol;Acc:MGI:5477165] | #DIV/0! | #DIV/0! |
| ENSMUSG00000034110 | 109,0418081 | 185,3 | 4,8 | 7,485564346 | 0,01149796 | Kctd7 | potassium channel tetramerisation domain containing 7 [Source:MGI Symbol;Acc:MGI:2442265] | 82,99 | 1,2 |
| ENSMUSG00000067455 | 7,991423955 | 17,1 | 0,0 | 7,487670424 | 0,040217345 | Hist1h4j | histone cluster 1, H4j [Source:MGI Symbol;Acc:MGI:2448436] | 74,72 | 0,0 |
| ENSMUSG00000042348 | 133,5133463 | 166,8 | 1,8 | 7,490452209 | 0,002487422 | Arl15 | ADP-ribosylation factor-like 15 [Source:MGI Symbol;Acc:MGI:2442308] | 88,48 | 0,7 |
| ENSMUSG00000048458 | 72,98443387 | 82,4 | 0,6 | 7,491353739 | 0,024345705 | Fam212b | family with sequence similarity 212, member B [Source:MGI Symbol;Acc:MGI:1923497] | 73,11 | 0,5 |
| ENSMUSG00000026791 | 132,6400046 | 227,7 | 31,4 | 7,492173093 | 0,006551757 | Slc2a8 | solute carrier family 2, (facilitated glucose transporter), member 8 [Source:MGI Symbol;Acc:MGI:1860103] | 99,23 | 21,2 |
| ENSMUSG00000042185 | 16,66240903 | 16,9 | 0,0 | 7,492831648 | 0,040839827 | Nfrkb | nuclear factor related to kappa B binding protein [Source:MGI Symbol;Acc:MGI:2442410] | 42,62 | 0,0 |
| ENSMUSG00000054855 | 19,67151853 | 18,9 | 0,0 | 7,49601963 | 0,036562028 | Rnd1 | Rho family GTPase 1 [Source:MGI Symbol;Acc:MGI:2444878] | 44,35 | 0,0 |
| ENSMUSG00000109336 | 327,4332195 | 539,9 | 131,9 | 7,499087225 | 0,000218895 | Samd4b | sterile alpha motif domain containing 4B [Source:MGI Symbol;Acc:MGI:2448542] | 98,14 | 23,7 |
| ENSMUSG00000027978 | 9,974367252 | 17,8 | 0,0 | 7,501590434 | 0,038161439 | Prss12 | protease, serine 12 neurotrypsin (motopsin) [Source:MGI Symbol;Acc:MGI:1100881] | 63,95 | 0,0 |
| ENSMUSG00000091183 | 19,39253153 | 16,8 | 0,0 | 7,503925505 | 0,034511518 | Gm5141 | predicted gene 5141 [Source:MGI Symbol;Acc:MGI:3779466] | 60,24 | 0,0 |
| ENSMUSG00000027088 | 172,4951828 | 331,4 | 7,4 | 7,504048376 | 0,000339105 | Phospho2 | phosphatase, orphan 2 [Source:MGI Symbol;Acc:MGI:1920623] | 61,40 | 1,8 |
| ENSMUSG00000037260 | 186,9146411 | 331,7 | 77,6 | 7,504647174 | 0,001927375 | Hgsnat | heparan-alpha-glucosaminide N-acetyltransferase [Source:MGI Symbol;Acc:MGI:1196297] | 99,60 | 28,9 |
| ENSMUSG00000064023 | 7,583745573 | 18,0 | 0,0 | 7,504684956 | 0,040206203 | Klk8 | kallikrein related-peptidase 8 [Source:MGI Symbol;Acc:MGI:1343327] | 75,19 | 0,0 |
| ENSMUSG00000073144 | 12,34819921 | 17,0 | 0,0 | 7,505012911 | 0,042818634 | 4930599N23Rik | RIKEN cDNA 4930599N23 gene [Source:MGI Symbol;Acc:MGI:1922629] | 82,08 | 0,0 |
| ENSMUSG00000055737 | 137,4887599 | 224,6 | 46,0 | 7,505059952 | 0,012135124 | Ghr | growth hormone receptor [Source:MGI Symbol;Acc:MGI:95708] | 87,20 | 6,7 |
| ENSMUSG00000072999 | 10,95449065 | 17,1 | 0,0 | 7,50707442 | 0,036413023 | Gm15401 | predicted gene 15401 [Source:MGI Symbol;Acc:MGI:3642201] | 17,56 | 0,0 |
| ENSMUSG00000028039 | 16,31891617 | 18,8 | 0,0 | 7,50719925 | 0,032858453 | Efna3 | ephrin A3 [Source:MGI Symbol;Acc:MGI:106644] | 62,52 | 0,0 |
| ENSMUSG00000042115 | 8,868469279 | 17,4 | 0,0 | 7,507264054 | 0,043523277 | Klhdc8a | kelch domain containing 8A [Source:MGI Symbol;Acc:MGI:2442630] | 35,99 | 0,0 |
| ENSMUSG00000053334 | 100,7475728 | 158,2 | 2,0 | 7,509337709 | 0,00408373 | Ficd | FIC domain containing [Source:MGI Symbol;Acc:MGI:1098550] | 98,05 | 1,0 |
| ENSMUSG00000050035 | 9,621071232 | 20,6 | 0,0 | 7,510955798 | 0,038215945 | Fhl4 | four and a half LIM domains 4 [Source:MGI Symbol;Acc:MGI:1338765] | 60,10 | 0,0 |
| ENSMUSG00000036095 | 450,561104 | 687,5 | 35,5 | 7,512888667 | 0,000296708 | Dgkb | diacylglycerol kinase, beta [Source:MGI Symbol;Acc:MGI:2442474] | 100,00 | 4,5 |
| ENSMUSG00000049686 | 76,0054238 | 112,1 | 1,0 | 7,517502611 | 0,007630642 | Orai1 | ORAI calcium release-activated calcium modulator 1 [Source:MGI Symbol;Acc:MGI:1925542] | 92,46 | 1,3 |
| ENSMUSG00000105868 | 193,5639811 | 297,8 | 38,1 | 7,519376672 | 0,006473549 | Gm43766 | predicted gene 43766 [Source:MGI Symbol;Acc:MGI:5663903] | 38,07 | 4,0 |
| ENSMUSG00000031090 | 12,86591071 | 26,0 | 0,0 | 7,524249008 | 0,035907432 | Nadsyn1 | NAD synthetase 1 [Source:MGI Symbol;Acc:MGI:1926164] | 62,48 | 0,0 |
| ENSMUSG00000042211 | 182,8930402 | 207,8 | 1,8 | 7,524493776 | 0,000228804 | Fbxo38 | F-box protein 38 [Source:MGI Symbol;Acc:MGI:2444639] | 98,66 | 0,7 |
| ENSMUSG00000037826 | 147,3388526 | 272,7 | 1,5 | 7,530088001 | 0,000432263 | Ppm1k | protein phosphatase 1K (PP2C domain containing) [Source:MGI Symbol;Acc:MGI:2442111] | 91,56 | 0,7 |
| ENSMUSG00000036446 | 22,32406531 | 20,6 | 0,0 | 7,533786079 | 0,032117629 | Lum | lumican [Source:MGI Symbol;Acc:MGI:109347] | 66,91 | 0,0 |
| ENSMUSG00000015189 | 201,371139 | 320,2 | 10,9 | 7,533847054 | 0,001537657 | Casd1 | CAS1 domain containing 1 [Source:MGI Symbol;Acc:MGI:2384865] | 95,08 | 1,7 |
| ENSMUSG00000030030 | 62,42944533 | 125,0 | 1,0 | 7,534705262 | 0,003415823 | 1700003E16Rik | RIKEN cDNA 1700003E16 gene [Source:MGI Symbol;Acc:MGI:1919087] | 99,80 | 1,1 |
| ENSMUSG00000064540 | 11,51397444 | 17,5 | 0,0 | 7,535316925 | 0,037291229 | Snord42a | small nucleolar RNA, C/D box 42A [Source:MGI Symbol;Acc:MGI:3819532] | #DIV/0! | #DIV/0! |
| ENSMUSG00000015850 | 9,335526233 | 18,0 | 0,0 | 7,535922191 | 0,037783459 | Adamtsl4 | ADAMTS-like 4 [Source:MGI Symbol;Acc:MGI:2389008] | 47,10 | 0,0 |
| ENSMUSG00000007867 | 172,5831449 | 277,8 | 28,9 | 7,536222615 | 0,002574765 | Ift43 | intraflagellar transport 43 [Source:MGI Symbol;Acc:MGI:1923661] | 100,00 | 26,8 |
| ENSMUSG00000084866 | 7,514034175 | 17,4 | 0,0 | 7,536685323 | 0,039413959 | A930006K02Rik | RIKEN cDNA A930006K02 gene [Source:MGI Symbol;Acc:MGI:1925683] | #DIV/0! | #DIV/0! |
| ENSMUSG00000021650 | 255,2296232 | 351,7 | 209,6 | 7,538535697 | 0,00119092 | Ptcd2 | pentatricopeptide repeat domain 2 [Source:MGI Symbol;Acc:MGI:1916177] | 99,98 | 32,1 |
| ENSMUSG00000005057 | 11,98022052 | 17,1 | 0,0 | 7,543222372 | 0,035855891 | Sh2b2 | SH2B adaptor protein 2 [Source:MGI Symbol;Acc:MGI:1345171] | 64,31 | 0,0 |
| ENSMUSG00000015093 | 19,61793931 | 17,7 | 0,0 | 7,546229363 | 0,033353365 | Clic3 | chloride intracellular channel 3 [Source:MGI Symbol;Acc:MGI:1916704] | 92,66 | 0,0 |
| ENSMUSG00000046321 | 7,614420678 | 17,6 | 0,0 | 7,548222309 | 0,038936575 | Hs3st2 | heparan sulfate (glucosamine) 3-O-sulfotransferase 2 [Source:MGI Symbol;Acc:MGI:1333802] | 52,85 | 0,0 |
| ENSMUSG00000061013 | 14,51137268 | 19,2 | 0,0 | 7,550913995 | 0,048850675 | Mkx | mohawk homeobox [Source:MGI Symbol;Acc:MGI:2687286] | 61,58 | 0,0 |
| ENSMUSG00000039686 | 282,9111432 | 518,6 | 22,2 | 7,552399183 | 1,683E-05 | Zer1 | zyg-11 related, cell cycle regulator [Source:MGI Symbol;Acc:MGI:2442511] | 95,56 | 7,6 |
| ENSMUSG00000102969 | 11,05075946 | 17,7 | 0,0 | 7,552973461 | 0,046024278 | Gm33366 | predicted gene, 33366 [Source:MGI Symbol;Acc:MGI:5592525] | 19,62 | 0,0 |
| ENSMUSG00000085133 | 278,6806959 | 416,5 | 74,4 | 7,553019155 | 0,00292659 | B930095G15Rik | RIKEN cDNA B930095G15 gene [Source:MGI Symbol;Acc:MGI:2443701] | 99,25 | 14,2 |
| ENSMUSG00000058331 | 14,44583731 | 18,5 | 0,0 | 7,553576425 | 0,038411401 | Zfp85 | zinc finger protein 85 [Source:MGI Symbol;Acc:MGI:107767] | 68,04 | 0,0 |
| ENSMUSG00000030255 | 237,4717237 | 257,7 | 168,6 | 7,554071282 | 0,008999244 | Sspn | sarcospan [Source:MGI Symbol;Acc:MGI:1353511] | 88,13 | 13,7 |
| ENSMUSG00000097626 | 10,84792566 | 17,3 | 0,0 | 7,554266934 | 0,0349724 | 4921504A21Rik | RIKEN cDNA 4921504A21 gene [Source:MGI Symbol;Acc:MGI:1918082] | 55,88 | 0,0 |
| ENSMUSG00000071637 | 143,4938117 | 276,1 | 90,6 | 7,554457974 | 0,006473692 | Cebpd | CCAAT/enhancer binding protein (C/EBP), delta [Source:MGI Symbol;Acc:MGI:103573] | #DIV/0! | #DIV/0! |
| ENSMUSG00000003527 | 82,38882004 | 107,6 | 0,6 | 7,555319546 | 0,00779471 | Dgcr14 | DiGeorge syndrome critical region gene 14 [Source:MGI Symbol;Acc:MGI:107854] | 98,50 | 0,8 |
| ENSMUSG00000020641 | 7,415601024 | 18,5 | 0,0 | 7,559559464 | 0,048006925 | Rsad2 | radical S-adenosyl methionine domain containing 2 [Source:MGI Symbol;Acc:MGI:1929628] | 45,28 | 0,0 |
| ENSMUSG00000022889 | 294,0502576 | 478,9 | 84,7 | 7,560023883 | 0,000411596 | Mrpl39 | mitochondrial ribosomal protein L39 [Source:MGI Symbol;Acc:MGI:1351620] | 64,09 | 14,8 |
| ENSMUSG00000064280 | 16,0275854 | 22,1 | 0,0 | 7,564044167 | 0,033841736 | Ccdc146 | coiled-coil domain containing 146 [Source:MGI Symbol;Acc:MGI:1922422] | 36,86 | 0,0 |
| ENSMUSG00000041426 | 186,0789718 | 366,3 | 17,0 | 7,567334975 | 0,000121115 | Hibch | 3-hydroxyisobutyryl-Coenzyme A hydrolase [Source:MGI Symbol;Acc:MGI:1923792] | 98,35 | 1,6 |
| ENSMUSG00000087028 | 11,06399062 | 17,9 | 0,0 | 7,568427715 | 0,035726554 | Gm13387 | predicted gene 13387 [Source:MGI Symbol;Acc:MGI:3651242] | #DIV/0! | #DIV/0! |
| ENSMUSG00000031681 | 212,0795205 | 333,5 | 51,8 | 7,56923087 | 0,001988309 | Smad1 | SMAD family member 1 [Source:MGI Symbol;Acc:MGI:109452] | 52,63 | 12,6 |
| ENSMUSG00000072845 | 16,87389977 | 18,6 | 0,0 | 7,57212238 | 0,045454473 | Tmprss11a | transmembrane protease, serine 11a [Source:MGI Symbol;Acc:MGI:2684853] | 70,15 | 0,0 |
| ENSMUSG00000036817 | 104,6060224 | 177,8 | 2,8 | 7,572763021 | 0,005254334 | Sun1 | Sad1 and UNC84 domain containing 1 [Source:MGI Symbol;Acc:MGI:1924303] | 83,51 | 0,8 |
| ENSMUSG00000010751 | 17,36343006 | 19,2 | 0,0 | 7,577110442 | 0,048260699 | Tnfrsf22 | tumor necrosis factor receptor superfamily, member 22 [Source:MGI Symbol;Acc:MGI:1930270] | 50,77 | 0,0 |
| ENSMUSG00000021003 | 84,58344258 | 110,2 | 0,6 | 7,577843129 | 0,007522213 | Galc | galactosylceramidase [Source:MGI Symbol;Acc:MGI:95636] | 95,94 | 0,3 |
| ENSMUSG00000037993 | 129,7015732 | 126,9 | 0,9 | 7,578363885 | 0,003260047 | Dhx38 | DEAH (Asp-Glu-Ala-His) box polypeptide 38 [Source:MGI Symbol;Acc:MGI:1927617] | 99,57 | 0,6 |
| ENSMUSG00000048899 | 15,35338667 | 17,6 | 0,0 | 7,578391538 | 0,032017121 | Rimkla | ribosomal modification protein rimK-like family member A [Source:MGI Symbol;Acc:MGI:3040686] | 55,44 | 0,0 |
| ENSMUSG00000018459 | 201,0827795 | 287,4 | 1,6 | 7,579439193 | 0,002611198 | Slc13a3 | arrier family 13 (sodium-dependent dicarboxylate transporter), member 3 [Source:MGI Symbol;Acc:MGI:2 | 91,50 | 1,2 |
| ENSMUSG00000104117 | 161,4121746 | 253,9 | 132,7 | 7,580026702 | 0,005923855 | Gm20743 | predicted gene, 20743 [Source:MGI Symbol;Acc:MGI:5434099] | 45,58 | 2,8 |
| ENSMUSG00000109228 | 15,48121677 | 19,2 | 0,0 | 7,580357146 | 0,033603209 | Fam81b | family with sequence similarity 81, member B [Source:MGI Symbol;Acc:MGI:2685122] | 45,17 | 0,0 |
| ENSMUSG00000057060 | 78,00223499 | 106,2 | 0,6 | 7,580935626 | 0,009540698 | Slc35f3 | solute carrier family 35, member F3 [Source:MGI Symbol;Acc:MGI:2444426] | 98,12 | 1,1 |
| ENSMUSG00000006191 | 139,6866432 | 227,4 | 3,7 | 7,585538307 | 0,002104533 | Cdkal1 | CDK5 regulatory subunit associated protein 1-like 1 [Source:MGI Symbol;Acc:MGI:1921765] | 50,15 | 0,9 |
| ENSMUSG00000087013 | 13,8503673 | 18,1 | 0,0 | 7,585786197 | 0,039448787 | 2610027K06Rik | RIKEN cDNA 2610027K06 gene [Source:MGI Symbol;Acc:MGI:1917159] | #DIV/0! | #DIV/0! |
| ENSMUSG00000089798 | 13,56218397 | 20,9 | 0,0 | 7,586479976 | 0,036025061 | 1700028K03Rik | RIKEN cDNA 1700028K03 gene [Source:MGI Symbol;Acc:MGI:1923671] | 16,50 | 0,0 |
| ENSMUSG00000054939 | 13,19326768 | 19,8 | 0,0 | 7,588672751 | 0,034237557 | Zfp174 | zinc finger protein 174 [Source:MGI Symbol;Acc:MGI:2686600] | 45,49 | 0,0 |
| ENSMUSG00000029155 | 11,19726492 | 19,1 | 0,0 | 7,589958871 | 0,032492661 | Spata18 | spermatogenesis associated 18 [Source:MGI Symbol;Acc:MGI:1920722] | 57,87 | 0,0 |
| ENSMUSG00000031981 | 9,152079367 | 20,5 | 0,0 | 7,593213827 | 0,035667203 | Capn9 | calpain 9 [Source:MGI Symbol;Acc:MGI:1920897] | 30,46 | 0,0 |
| ENSMUSG00000039629 | 187,9443462 | 144,4 | 0,9 | 7,596056089 | 0,008873387 | Strip2 | striatin interacting protein 2 [Source:MGI Symbol;Acc:MGI:2444363] | 71,50 | 0,3 |
| ENSMUSG00000022237 | 10,40382893 | 18,0 | 0,0 | 7,596956338 | 0,034990446 | Ankrd33b | ankyrin repeat domain 33B [Source:MGI Symbol;Acc:MGI:1917904] | 28,31 | 0,0 |
| ENSMUSG00000045573 | 28,43613685 | 19,2 | 0,0 | 7,599137787 | 0,026299636 | Penk | preproenkephalin [Source:MGI Symbol;Acc:MGI:104629] | 68,99 | 0,0 |
| ENSMUSG00000054967 | 8,291677203 | 18,6 | 0,0 | 7,608318424 | 0,04126323 | Zfp647 | zinc finger protein 647 [Source:MGI Symbol;Acc:MGI:3052806] | 81,51 | 0,0 |
| ENSMUSG00000026188 | 10,09809925 | 22,3 | 0,0 | 7,610176935 | 0,035534808 | Tmem169 | transmembrane protein 169 [Source:MGI Symbol;Acc:MGI:2442781] | 64,17 | 0,0 |
| ENSMUSG00000085541 | 9,245943137 | 18,6 | 0,0 | 7,613250435 | 0,035799496 | Gm16010 | predicted gene 16010 [Source:MGI Symbol;Acc:MGI:3802126] | 22,82 | 0,0 |
| ENSMUSG00000025492 | 219,3089602 | 279,6 | 13,3 | 7,614351098 | 0,003912703 | Ifitm3 | interferon induced transmembrane protein 3 [Source:MGI Symbol;Acc:MGI:1913391] | 99,46 | 25,0 |
| ENSMUSG00000086119 | 11,77586781 | 19,5 | 0,0 | 7,616066355 | 0,041940594 | Gm2415 | predicted gene 2415 [Source:MGI Symbol;Acc:MGI:3780582] | 40,35 | 0,0 |
| ENSMUSG00000028085 | 80,85657151 | 117,6 | 0,9 | 7,61805312 | 0,006659046 | Gatb | glutamyl-tRNA(Gln) amidotransferase, subunit B [Source:MGI Symbol;Acc:MGI:2442496] | 54,41 | 0,6 |
| ENSMUSG00000079450 | 12,02914666 | 19,0 | 0,0 | 7,618207993 | 0,040458196 | Cldn34c1 | claudin 34C1 [Source:MGI Symbol;Acc:MGI:1920311] | 22,59 | 0,0 |
| ENSMUSG00000109371 | 8,255134299 | 19,8 | 0,0 | 7,618379829 | 0,037747822 | RP23-274K7.1 | | #DIV/0! | #DIV/0! |
| ENSMUSG00000020142 | 258,4972404 | 358,7 | 180,6 | 7,622594777 | 0,001850845 | Slc1a4 | e carrier family 1 (glutamate/neutral amino acid transporter), member 4 [Source:MGI Symbol;Acc:MGI:213 | 99,58 | 28,1 |
| ENSMUSG00000027737 | 126,029779 | 109,2 | 0,6 | 7,625282231 | 0,006618112 | Slc7a11 | carrier family 7 (cationic amino acid transporter, y+ system), member 11 [Source:MGI Symbol;Acc:MGI:13 | 74,73 | 0,3 |
| ENSMUSG00000036586 | 16,53460401 | 18,6 | 0,0 | 7,627405424 | 0,039413959 | Grifin | galectin-related inter-fiber protein [Source:MGI Symbol;Acc:MGI:1925248] | 97,88 | 0,0 |
| ENSMUSG00000023886 | 16,79588093 | 20,9 | 0,0 | 7,628500481 | 0,030390259 | Smoc2 | SPARC related modular calcium binding 2 [Source:MGI Symbol;Acc:MGI:1929881] | 69,07 | 0,0 |
| ENSMUSG00000056023 | 9,264552684 | 18,8 | 0,0 | 7,62914514 | 0,041007846 | Gm9989 | predicted gene 9989 [Source:MGI Symbol;Acc:MGI:3642361] | #DIV/0! | #DIV/0! |
| ENSMUSG00000002147 | 32,08356397 | 24,3 | 0,0 | 7,631509203 | 0,041755486 | Stat6 | signal transducer and activator of transcription 6 [Source:MGI Symbol;Acc:MGI:103034] | 54,00 | 0,0 |
| ENSMUSG00000027997 | 144,4239531 | 275,5 | 76,8 | 7,632492279 | 0,003112756 | Casp6 | caspase 6 [Source:MGI Symbol;Acc:MGI:1312921] | 96,59 | 26,3 |
| ENSMUSG00000098093 | 13,71756141 | 23,5 | 0,0 | 7,634443049 | 0,032933683 | Gm8707 | predicted gene 8707 [Source:MGI Symbol;Acc:MGI:3643201] | #DIV/0! | #DIV/0! |
| ENSMUSG00000017764 | 143,5361575 | 202,1 | 7,4 | 7,636756011 | 0,00903868 | Zswim1 | zinc finger SWIM-type containing 1 [Source:MGI Symbol;Acc:MGI:1919221] | 86,35 | 2,9 |
| ENSMUSG00000001034 | 73,92602402 | 127,3 | 0,6 | 7,639519957 | 0,002720628 | Mapk7 | mitogen-activated protein kinase 7 [Source:MGI Symbol;Acc:MGI:1346347] | 93,20 | 0,5 |
| ENSMUSG00000108207 | 31,13822051 | 19,9 | 0,0 | 7,642515379 | 0,042551917 | 1810059H22Rik | RIKEN cDNA 1810059H22 gene [Source:MGI Symbol;Acc:MGI:1917070] | #DIV/0! | #DIV/0! |
| ENSMUSG00000097515 | 14,28704117 | 19,3 | 0,0 | 7,64323334 | 0,047775952 | 1700040D17Rik | RIKEN cDNA 1700040D17 gene [Source:MGI Symbol;Acc:MGI:1923852] | #DIV/0! | #DIV/0! |
| ENSMUSG00000032006 | 167,3124439 | 242,6 | 31,4 | 7,644253486 | 0,011838927 | Pdgfd | platelet-derived growth factor, D polypeptide [Source:MGI Symbol;Acc:MGI:1919035] | 100,00 | 7,2 |
| ENSMUSG00000108955 | 11,54477963 | 18,8 | 0,0 | 7,644696503 | 0,036159363 | RP23-442J18.3 | | 74,24 | 0,0 |
| ENSMUSG00000021318 | 76,05365841 | 148,2 | 0,9 | 7,646135882 | 0,001014306 | Gli3 | GLI-Kruppel family member GLI3 [Source:MGI Symbol;Acc:MGI:95729] | 89,35 | 0,3 |
| ENSMUSG00000108935 | 8,828630682 | 20,0 | 0,0 | 7,649124136 | 0,03529281 | RP23-17C12.5 | | 36,23 | 0,0 |
| ENSMUSG00000013936 | 7,136166677 | 18,8 | 0,0 | 7,649221308 | 0,036915526 | Myl2 | myosin, light polypeptide 2, regulatory, cardiac, slow [Source:MGI Symbol;Acc:MGI:97272] | 33,25 | 0,0 |
| ENSMUSG00000089760 | 8,839659804 | 18,7 | 0,0 | 7,653109112 | 0,037027415 | D030046N08Rik | RIKEN cDNA D030046N08 gene [Source:MGI Symbol;Acc:MGI:3645234] | #DIV/0! | #DIV/0! |
| ENSMUSG00000052525 | 67,3895844 | 125,4 | 1,0 | 7,653671128 | 0,005863782 | Spdya | speedy/RINGO cell cycle regulator family, member A [Source:MGI Symbol;Acc:MGI:1918141] | 57,54 | 1,1 |
| ENSMUSG00000040624 | 73,81579986 | 131,6 | 1,0 | 7,653796609 | 0,004151407 | Plekhg1 | n homology domain containing, family G (with RhoGef domain) member 1 [Source:MGI Symbol;Acc:MGI:2 | 89,39 | 0,3 |
| ENSMUSG00000078185 | 13,54022902 | 20,1 | 0,0 | 7,656241617 | 0,032493928 | Chml | choroideremia-like [Source:MGI Symbol;Acc:MGI:101913] | 44,05 | 0,0 |
| ENSMUSG00000105651 | 10,06137996 | 18,7 | 0,0 | 7,6569689 | 0,049169375 | Gm43003 | predicted gene 43003 [Source:MGI Symbol;Acc:MGI:5663140] | #DIV/0! | #DIV/0! |
| ENSMUSG00000027886 | 16,40668859 | 21,0 | 0,0 | 7,65725099 | 0,036403657 | 1700013F07Rik | RIKEN cDNA 1700013F07 gene [Source:MGI Symbol;Acc:MGI:1922754] | 94,96 | 0,0 |
| ENSMUSG00000030450 | 25,84167988 | 25,7 | 0,0 | 7,658085233 | 0,036115609 | Oca2 | oculocutaneous albinism II [Source:MGI Symbol;Acc:MGI:97454] | 63,46 | 0,0 |
| ENSMUSG00000040666 | 21,61551985 | 23,1 | 0,0 | 7,659053707 | 0,037032158 | Sh3bgr | SH3-binding domain glutamic acid-rich protein [Source:MGI Symbol;Acc:MGI:1354740] | 76,33 | 0,0 |
| ENSMUSG00000074064 | 221,5158924 | 382,9 | 36,0 | 7,661066004 | 0,000803302 | Mlycd | malonyl-CoA decarboxylase [Source:MGI Symbol;Acc:MGI:1928485] | 99,62 | 19,7 |
| ENSMUSG00000044566 | 6,244430668 | 22,6 | 0,0 | 7,662482314 | 0,036733794 | Cage1 | cancer antigen 1 [Source:MGI Symbol;Acc:MGI:1918463] | 43,58 | 0,0 |
| ENSMUSG00000060530 | 11,78894957 | 21,9 | 0,0 | 7,664058677 | 0,033009037 | A930017M01Rik | RIKEN cDNA A930017M01 gene [Source:MGI Symbol;Acc:MGI:2685151] | #DIV/0! | #DIV/0! |
| ENSMUSG00000103217 | 830,4869917 | 1421,6 | 316,8 | 7,667103101 | 1,3278E-05 | Gm38287 | predicted gene, 38287 [Source:MGI Symbol;Acc:MGI:5611515] | 100,00 | 34,4 |
| ENSMUSG00000052415 | 8,78694969 | 19,9 | 0,0 | 7,66849942 | 0,037127946 | Tchh | trichohyalin [Source:MGI Symbol;Acc:MGI:2177944] | 37,85 | 0,0 |
| ENSMUSG00000022816 | 177,675013 | 300,7 | 1,5 | 7,670254407 | 0,001109028 | Fstl1 | follistatin-like 1 [Source:MGI Symbol;Acc:MGI:102793] | 95,30 | 0,7 |
| ENSMUSG00000059498 | 30,12841498 | 19,4 | 0,0 | 7,672128565 | 0,024707886 | Fcgr3 | Fc receptor, IgG, low affinity III [Source:MGI Symbol;Acc:MGI:95500] | 71,31 | 0,0 |
| ENSMUSG00000029468 | 13,93005404 | 23,9 | 0,0 | 7,672803602 | 0,031241937 | P2rx7 | purinergic receptor P2X, ligand-gated ion channel, 7 [Source:MGI Symbol;Acc:MGI:1339957] | 42,10 | 0,0 |
| ENSMUSG00000024990 | 8,206186037 | 19,4 | 0,0 | 7,673670907 | 0,035197959 | Rbp4 | retinol binding protein 4, plasma [Source:MGI Symbol;Acc:MGI:97879] | 67,39 | 0,0 |
| ENSMUSG00000025723 | 14,17323688 | 20,3 | 0,0 | 7,67458546 | 0,047131668 | Nmb | neuromedin B [Source:MGI Symbol;Acc:MGI:1915289] | 95,93 | 0,0 |
| ENSMUSG00000020176 | 17,27773248 | 27,0 | 0,0 | 7,675594573 | 0,03703421 | Grb10 | growth factor receptor bound protein 10 [Source:MGI Symbol;Acc:MGI:103232] | 39,44 | 0,0 |
| ENSMUSG00000028294 | 16,38472649 | 21,3 | 0,0 | 7,676145183 | 0,031354119 | Cfap206 | cilia and flagella associated protein 206 [Source:MGI Symbol;Acc:MGI:1916579] | 69,63 | 0,0 |
| ENSMUSG00000028173 | 254,0454335 | 284,0 | 14,0 | 7,680036772 | 0,003324528 | Wls | wntless homolog (Drosophila) [Source:MGI Symbol;Acc:MGI:1915401] | 94,31 | 2,3 |
| ENSMUSG00000107314 | 6,983303701 | 19,2 | 0,0 | 7,680108216 | 0,036310348 | Gm20488 | predicted gene 20488 [Source:MGI Symbol;Acc:MGI:5141953] | #DIV/0! | #DIV/0! |
| ENSMUSG00000074867 | 14,85603288 | 24,9 | 0,0 | 7,682572019 | 0,038487558 | Zfp808 | zinc finger protein 80 [Source:MGI Symbol;Acc:MGI:3704127] | 59,03 | 0,0 |
| ENSMUSG00000029026 | 16,66667562 | 20,2 | 0,0 | 7,683543958 | 0,049283007 | Trp73 | transformation related protein 73 [Source:MGI Symbol;Acc:MGI:1336991] | 44,82 | 0,0 |
| ENSMUSG00000087633 | 15,33074752 | 19,2 | 0,0 | 7,684138964 | 0,044490222 | Gm14455 | predicted gene 14455 [Source:MGI Symbol;Acc:MGI:3711286] | #DIV/0! | #DIV/0! |
| ENSMUSG00000055963 | 218,8176185 | 301,0 | 83,5 | 7,684139351 | 0,007337311 | Triqk | triple QxxK/R motif containing [Source:MGI Symbol;Acc:MGI:3650048] | 93,98 | 13,5 |
| ENSMUSG00000044576 | 333,1030866 | 494,5 | 93,2 | 7,686428092 | 0,000416716 | Gareml | GRB2 associated, regulator of MAPK1-like [Source:MGI Symbol;Acc:MGI:2685290] | 100,00 | 28,5 |
| ENSMUSG00000063802 | 326,2606295 | 540,7 | 56,3 | 7,686561654 | 0,000339105 | Hspbp1 | PA (heat shock 70kDa) binding protein, cytoplasmic cochaperone 1 [Source:MGI Symbol;Acc:MGI:19134 | 93,57 | 23,4 |
| ENSMUSG00000035493 | 13,87414127 | 20,7 | 0,0 | 7,687632161 | 0,027511569 | Tgfbi | transforming growth factor, beta induced [Source:MGI Symbol;Acc:MGI:99959] | 68,14 | 0,0 |
| ENSMUSG00000042763 | 81,46041066 | 139,1 | 0,9 | 7,687660096 | 0,004110442 | Maneal | mannosidase, endo-alpha-like [Source:MGI Symbol;Acc:MGI:2684896] | 97,84 | 0,1 |
| ENSMUSG00000081219 | 9,597275374 | 18,9 | 0,0 | 7,68803968 | 0,033679589 | Bambi-ps1 | and activin membrane-bound inhibitor, pseudogene (Xenopus laevis) [Source:MGI Symbol;Acc:MGI:1932 | #DIV/0! | #DIV/0! |
| ENSMUSG00000020098 | 13,32208002 | 19,2 | 0,0 | 7,688107886 | 0,036067348 | Pcbd1 | olamine dehydratase/dimerization cofactor of hepatocyte nuclear factor 1 alpha (TCF1) 1 [Source:MGI Sym | 94,64 | 0,0 |
| ENSMUSG00000025854 | 162,8983447 | 249,3 | 7,0 | 7,688889107 | 0,004671396 | Fam20c | family with sequence similarity 20, member C [Source:MGI Symbol;Acc:MGI:2136853] | 98,67 | 2,8 |
| ENSMUSG00000045062 | 22,25418639 | 22,3 | 0,0 | 7,689313349 | 0,041726586 | Pcdhb7 | protocadherin beta 7 [Source:MGI Symbol;Acc:MGI:2136741] | 60,51 | 0,0 |
| ENSMUSG00000046269 | 10,13095039 | 19,9 | 0,0 | 7,690327919 | 0,040475764 | Usp27x | ubiquitin specific peptidase 27, X chromosome [Source:MGI Symbol;Acc:MGI:1859645] | 43,93 | 0,0 |
| ENSMUSG00000060962 | 9,308651973 | 19,6 | 0,0 | 7,692999178 | 0,032686055 | Dmkn | dermokine [Source:MGI Symbol;Acc:MGI:1920962] | 46,35 | 0,0 |
| ENSMUSG00000026094 | 224,5243041 | 311,5 | 168,2 | 7,70041786 | 0,005518259 | Stk17b | serine/threonine kinase 17b (apoptosis-inducing) [Source:MGI Symbol;Acc:MGI:2138162] | 99,32 | 15,7 |
| ENSMUSG00000023122 | 23,23939842 | 22,7 | 0,0 | 7,701648311 | 0,032492661 | Sult1c2 | sulfotransferase family, cytosolic, 1C, member 2 [Source:MGI Symbol;Acc:MGI:1916333] | 71,70 | 0,0 |
| ENSMUSG00000042045 | 10,40886686 | 19,4 | 0,0 | 7,702305231 | 0,032855246 | Sln | sarcolipin [Source:MGI Symbol;Acc:MGI:1913652] | 100,00 | 0,0 |
| ENSMUSG00000029127 | 21,40383694 | 20,5 | 0,0 | 7,703676725 | 0,046538228 | Zbtb49 | zinc finger and BTB domain containing 49 [Source:MGI Symbol;Acc:MGI:1922329] | 41,08 | 0,0 |
| ENSMUSG00000033488 | 160,4155674 | 268,2 | 11,1 | 7,70373859 | 0,005004394 | BC026585 | cDNA sequence BC026585 [Source:MGI Symbol;Acc:MGI:2448516] | 99,92 | 14,3 |
| ENSMUSG00000086886 | 9,132925215 | 19,8 | 0,0 | 7,706599849 | 0,033603209 | Gm15135 | predicted gene 15135 [Source:MGI Symbol;Acc:MGI:3705309] | 99,45 | 0,0 |
| ENSMUSG00000020930 | 13,10665437 | 19,1 | 0,0 | 7,707605706 | 0,031573046 | Ccdc103 | coiled-coil domain containing 103 [Source:MGI Symbol;Acc:MGI:1920543] | 71,36 | 0,0 |
| ENSMUSG00000104093 | 9,863437433 | 20,1 | 0,0 | 7,708351748 | 0,03848596 | A330015K06Rik | RIKEN cDNA A330015K06 gene [Source:MGI Symbol;Acc:MGI:2443553] | 17,21 | 0,0 |
| ENSMUSG00000029392 | 198,2886925 | 317,0 | 105,8 | 7,708848682 | 0,002876863 | Rilpl1 | Rab interacting lysosomal protein-like 1 [Source:MGI Symbol;Acc:MGI:1922945] | 94,86 | 26,9 |
| ENSMUSG00000031974 | 145,6264102 | 217,7 | 1,2 | 7,710315495 | 0,00086999 | Abcb10 | ATP-binding cassette, sub-family B (MDR/TAP), member 10 [Source:MGI Symbol;Acc:MGI:1860508] | 94,33 | 0,3 |
| ENSMUSG00000105541 | 16,89755564 | 19,8 | 0,0 | 7,710798833 | 0,036453483 | Gm43136 | predicted gene 43136 [Source:MGI Symbol;Acc:MGI:5663273] | 100,00 | 0,0 |
| ENSMUSG00000045319 | 16,70415612 | 26,1 | 0,0 | 7,711340718 | 0,040711104 | Proser2 | proline and serine rich 2 [Source:MGI Symbol;Acc:MGI:2442238] | 42,72 | 0,0 |

| | | | | | | | | | |
|---|---|---|---|---|---|---|---|---|---|
| ENSMUSG00000003477 | 25,67471407 | 19,6 | 0,0 | 7,712787917 | 0,03855583 | Inmt | indolethylamine N-methyltransferase [Source:MGI Symbol;Acc:MGI:102963] | 92,63 | 0,0 |
| ENSMUSG00000049300 | 85,25288609 | 185,1 | 1,2 | 7,712927181 | 0,001856511 | Prmt6 | protein arginine N-methyltransferase 6 [Source:MGI Symbol;Acc:MGI:2139971] | 98,95 | 0,6 |
| ENSMUSG00000003344 | 232,662838 | 404,2 | 14,5 | 7,714706898 | 0,000814172 | Btbd2 | BTB (POZ) domain containing 2 [Source:MGI Symbol;Acc:MGI:1933831] | 80,83 | 5,5 |
| ENSMUSG00000031576 | 8,676867443 | 20,5 | 0,0 | 7,715174497 | 0,033753697 | Kcnu1 | potassium channel, subfamily U, member 1 [Source:MGI Symbol;Acc:MGI:1202300] | 36,13 | 0,0 |
| ENSMUSG00000043164 | 9,630176049 | 20,2 | 0,0 | 7,716012735 | 0,032996306 | Tmem212 | transmembrane protein 212 [Source:MGI Symbol;Acc:MGI:2685410] | 90,46 | 0,0 |
| ENSMUSG00000047671 | 11,96770805 | 20,6 | 0,0 | 7,716876172 | 0,032201517 | Tctex1d4 | Tctex1 domain containing 4 [Source:MGI Symbol;Acc:MGI:3045358] | 87,10 | 0,0 |
| ENSMUSG00000028256 | 125,5666713 | 119,1 | 1,0 | 7,719817796 | 0,011576497 | Odf2l | outer dense fiber of sperm tails 2-like [Source:MGI Symbol;Acc:MGI:1098600] | 83,65 | 0,8 |
| ENSMUSG00000074024 | 19,70559623 | 23,2 | 0,0 | 7,720749365 | 0,0453227 | 4632427E13Rik | RIKEN cDNA 4632427E13 gene [Source:MGI Symbol;Acc:MGI:1915436] | 14,14 | 0,0 |
| ENSMUSG00000015659 | 41,08575871 | 22,3 | 0,0 | 7,722509153 | 0,044856546 | Serac1 | serine active site containing 1 [Source:MGI Symbol;Acc:MGI:2447813] | 59,32 | 0,0 |
| ENSMUSG00000056329 | 12,17180462 | 19,7 | 0,0 | 7,72289234 | 0,036800251 | 1810010K12Rik | RIKEN cDNA 1810010K12 gene [Source:MGI Symbol;Acc:MGI:1921419] | #DIV/0! | #DIV/0! |
| ENSMUSG00000055202 | 7,060951215 | 20,1 | 0,0 | 7,723126778 | 0,039697754 | Zfp811 | zinc finger protein 811 [Source:MGI Symbol;Acc:MGI:2682944] | 52,90 | 0,0 |
| ENSMUSG00000015843 | 13,49097773 | 20,9 | 0,0 | 7,727647277 | 0,036067348 | Rxrg | retinoid X receptor gamma [Source:MGI Symbol;Acc:MGI:98216] | 67,48 | 0,0 |
| ENSMUSG00000003354 | 12,2400468 | 20,4 | 0,0 | 7,728032452 | 0,031057321 | Ccdc65 | coiled-coil domain containing 65 [Source:MGI Symbol;Acc:MGI:2146001] | 84,29 | 0,0 |
| ENSMUSG00000005102 | 25,72389415 | 20,2 | 0,0 | 7,72852138 | 0,032017121 | Eif2ak4 | eukaryotic translation initiation factor 2 alpha kinase 4 [Source:MGI Symbol;Acc:MGI:1353427] | 34,86 | 0,0 |
| ENSMUSG00000057614 | 270,9134659 | 421,1 | 33,3 | 7,729785529 | 0,003119232 | Gnai1 | guanine nucleotide binding protein (G protein), alpha inhibiting 1 [Source:MGI Symbol;Acc:MGI:95771] | 99,81 | 9,4 |
| ENSMUSG00000105006 | 21,36455473 | 19,9 | 0,0 | 7,729894783 | 0,046522378 | Gm9484 | predicted gene 9484 [Source:MGI Symbol;Acc:MGI:3779894] | #DIV/0! | #DIV/0! |
| ENSMUSG00000078580 | 15,42926596 | 24,9 | 0,0 | 7,73075641 | 0,033301917 | E430018J23Rik | RIKEN cDNA E430018J23 gene [Source:MGI Symbol;Acc:MGI:2141981] | 49,04 | 0,0 |
| ENSMUSG00000026807 | 31,69795977 | 22,8 | 0,0 | 7,734068496 | 0,04553704 | Ak8 | adenylate kinase 8 [Source:MGI Symbol;Acc:MGI:1916120] | 83,02 | 0,0 |
| ENSMUSG00000066643 | 131,9852534 | 158,3 | 0,9 | 7,734265644 | 0,001276418 | Wdr35 | WD repeat domain 35 [Source:MGI Symbol;Acc:MGI:1921932] | 95,70 | 0,4 |
| ENSMUSG00000107040 | 9,889318873 | 19,9 | 0,0 | 7,735618003 | 0,034974929 | Gm43791 | predicted gene 43791 [Source:MGI Symbol;Acc:MGI:5663928] | 99,72 | 0,0 |
| ENSMUSG00000021414 | 10,61230793 | 24,3 | 0,0 | 7,736416264 | 0,031195337 | Fam217a | family with sequence similarity 217, member A [Source:MGI Symbol;Acc:MGI:1919114] | 62,88 | 0,0 |
| ENSMUSG00000028687 | 15,5211355 | 20,0 | 0,0 | 7,737983615 | 0,03834123 | Mutyh | mutY homolog (E. coli) [Source:MGI Symbol;Acc:MGI:1917853] | 84,93 | 0,0 |
| ENSMUSG00000021592 | 98,83882546 | 128,0 | 0,9 | 7,739000567 | 0,010341486 | Arsk | arylsulfatase K [Source:MGI Symbol;Acc:MGI:1924291] | 87,28 | 0,2 |
| ENSMUSG00000040836 | 10,07201769 | 20,8 | 0,0 | 7,739269923 | 0,034849401 | Gpr161 | G protein-coupled receptor 161 [Source:MGI Symbol;Acc:MGI:2685054] | 42,28 | 0,0 |
| ENSMUSG00000006134 | 258,8858604 | 427,1 | 45,3 | 7,741807044 | 0,000291776 | Crkl | v-crk avian sarcoma virus CT10 oncogene homolog-like [Source:MGI Symbol;Acc:MGI:104686] | 100,00 | 12,7 |
| ENSMUSG00000086915 | 15,55411631 | 20,4 | 0,0 | 7,742048777 | 0,033243415 | Gm16365 | predicted gene 16365 [Source:MGI Symbol;Acc:MGI:3840143] | 66,15 | 0,0 |
| ENSMUSG00000026024 | 110,5420412 | 138,0 | 0,9 | 7,742052342 | 0,003758957 | Als2 | amyotrophic lateral sclerosis 2 (juvenile) [Source:MGI Symbol;Acc:MGI:1921268] | 92,56 | 0,4 |
| ENSMUSG00000021752 | 267,2244267 | 363,0 | 89,8 | 7,74431236 | 0,00143212 | Kctd6 | potassium channel tetramerisation domain containing 6 [Source:MGI Symbol;Acc:MGI:1918643] | 77,09 | 13,9 |
| ENSMUSG00000091864 | 15,71157501 | 19,9 | 0,0 | 7,744407054 | 0,047416673 | Gm17102 | predicted gene 17102 [Source:MGI Symbol;Acc:MGI:4937929] | #DIV/0! | #DIV/0! |
| ENSMUSG00000029371 | 140,5377617 | 264,4 | 38,7 | 7,746058658 | 0,005392139 | Cxcl5 | chemokine (C-X-C motif) ligand 5 [Source:MGI Symbol;Acc:MGI:1096868] | 98,75 | 17,3 |
| ENSMUSG00000021432 | 24,31470921 | 24,6 | 0,0 | 7,747745895 | 0,032573357 | Slc35b3 | solute carrier family 35, member B3 [Source:MGI Symbol;Acc:MGI:1913978] | 69,21 | 0,0 |
| ENSMUSG00000073207 | 8,368556096 | 20,7 | 0,0 | 7,747856711 | 0,037512084 | Ccdc160 | coiled-coil domain containing 160 [Source:MGI Symbol;Acc:MGI:3588225] | 86,20 | 0,0 |
| ENSMUSG00000022489 | 84,66905138 | 138,8 | 1,0 | 7,748420831 | 0,003820711 | Pde1b | phosphodiesterase 1B, Ca2+-calmodulin dependent [Source:MGI Symbol;Acc:MGI:97523] | 99,53 | 0,2 |
| ENSMUSG00000039782 | 92,64590061 | 115,2 | 0,6 | 7,752283505 | 0,007146733 | Cpeb2 | cytoplasmic polyadenylation element binding protein 2 [Source:MGI Symbol;Acc:MGI:2442640] | 75,52 | 0,4 |
| ENSMUSG00000039328 | 12,39624884 | 22,1 | 0,0 | 7,75239446 | 0,02940813 | Rnf122 | ring finger protein 122 [Source:MGI Symbol;Acc:MGI:1916117] | 44,99 | 0,0 |
| ENSMUSG00000043857 | 10,44253111 | 21,3 | 0,0 | 7,752605822 | 0,030326102 | Mgat5b | mannoside acetylglucosaminyltransferase 5, isoenzyme B [Source:MGI Symbol;Acc:MGI:3606200] | 53,81 | 0,0 |
| ENSMUSG00000079737 | 12,08992599 | 21,4 | 0,0 | 7,756078591 | 0,030970165 | 3110001I22Rik | RIKEN cDNA 3110001I22 gene [Source:MGI Symbol;Acc:MGI:1913848] | 67,50 | 0,0 |
| ENSMUSG00000106990 | 10,08195667 | 21,1 | 0,0 | 7,756779858 | 0,032425801 | Gm42547 | predicted gene 42547 [Source:MGI Symbol;Acc:MGI:5662684] | 57,71 | 0,0 |
| ENSMUSG00000086996 | 20,04043534 | 24,5 | 0,0 | 7,760601088 | 0,043183971 | 4933416E14Rik | RIKEN cDNA 4933416E14 gene [Source:MGI Symbol;Acc:MGI:1918430] | #DIV/0! | #DIV/0! |
| ENSMUSG00000026096 | 308,3190504 | 418,7 | 147,9 | 7,761937503 | 0,000860505 | Osgepl1 | O-sialoglycoprotein endopeptidase-like 1 [Source:MGI Symbol;Acc:MGI:1919335] | 72,79 | 19,9 |
| ENSMUSG00000036469 | 52,41101865 | 24,5 | 0,0 | 7,764069531 | 0,034990446 | March1 | membrane-associated ring finger (C3HC4) 1 [Source:MGI Symbol;Acc:MGI:1920175] | 40,82 | 0,0 |
| ENSMUSG00000020890 | 84,36411815 | 130,8 | 0,6 | 7,764848675 | 0,004118196 | Gucy2e | guanylate cyclase 2e [Source:MGI Symbol;Acc:MGI:105123] | 61,53 | 0,2 |
| ENSMUSG00000043913 | 20,52274197 | 20,4 | 0,0 | 7,765136199 | 0,04126323 | Ccdc60 | coiled-coil domain containing 60 [Source:MGI Symbol;Acc:MGI:2141043] | 44,65 | 0,0 |
| ENSMUSG00000097193 | 13,94904297 | 21,2 | 0,0 | 7,76649188 | 0,047650897 | Gm26664 | predicted gene, 26664 [Source:MGI Symbol;Acc:MGI:5477158] | #DIV/0! | #DIV/0! |
| ENSMUSG00000036376 | 121,7373827 | 130,4 | 0,9 | 7,766656425 | 0,006385371 | Abt1 | activator of basal transcription 1 [Source:MGI Symbol;Acc:MGI:1353636] | 99,30 | 2,0 |
| ENSMUSG00000057173 | 7,351937021 | 21,5 | 0,0 | 7,767540196 | 0,033825651 | Rfx8 | regulatory factor X 8 [Source:MGI Symbol;Acc:MGI:3588206] | 80,06 | 0,0 |
| ENSMUSG00000040586 | 17,52029515 | 23,6 | 0,0 | 7,768858976 | 0,030374361 | Ofd1 | oral-facial-digital syndrome 1 gene homolog (human) [Source:MGI Symbol;Acc:MGI:1350328] | 44,82 | 0,0 |
| ENSMUSG00000078622 | 273,8173118 | 380,6 | 33,3 | 7,769803218 | 0,001705667 | Ccdc47 | coiled-coil domain containing 47 [Source:MGI Symbol;Acc:MGI:1914413] | 70,14 | 4,0 |
| ENSMUSG00000010057 | 77,80016556 | 145,0 | 0,9 | 7,770921681 | 0,002096022 | Nprl2 | nitrogen permease regulator-like 2 [Source:MGI Symbol;Acc:MGI:1914482] | 99,81 | 1,7 |
| ENSMUSG00000077625 | 17,34379235 | 20,8 | 0,0 | 7,771018232 | 0,032460068 | Snord4a | small nucleolar RNA, C/D box 4A [Source:MGI Symbol;Acc:MGI:3819540] | #DIV/0! | #DIV/0! |
| ENSMUSG00000039384 | 8,989528162 | 23,4 | 0,0 | 7,772828401 | 0,030485218 | Dusp10 | dual specificity phosphatase 10 [Source:MGI Symbol;Acc:MGI:1927070] | 61,39 | 0,0 |
| ENSMUSG00000031673 | 224,3583654 | 231,5 | 119,7 | 7,773183656 | 0,00877641 | Cdh11 | cadherin 11 [Source:MGI Symbol;Acc:MGI:99217] | 91,83 | 9,6 |
| ENSMUSG00000078779 | 14,253632 | 23,3 | 0,0 | 7,775954962 | 0,039622276 | Zfp59 | zinc finger protein 59 [Source:MGI Symbol;Acc:MGI:99206] | 55,54 | 0,0 |
| ENSMUSG00000030522 | 345,2786717 | 446,1 | 281,4 | 7,775999385 | 0,003686734 | Mtmr10 | myotubularin related protein 10 [Source:MGI Symbol;Acc:MGI:2142292] | 99,78 | 23,0 |
| ENSMUSG00000033053 | 15,53614271 | 23,8 | 0,0 | 7,776035239 | 0,029326086 | 1700028P14Rik | RIKEN cDNA 1700028P14 gene [Source:MGI Symbol;Acc:MGI:1914733] | 88,13 | 0,0 |
| ENSMUSG00000024427 | 67,56245997 | 128,8 | 0,9 | 7,776123918 | 0,006198296 | Spry4 | sprouty homolog 4 (Drosophila) [Source:MGI Symbol;Acc:MGI:1345144] | 96,73 | 0,5 |
| ENSMUSG00000032940 | 8,260118194 | 25,4 | 0,0 | 7,776352398 | 0,030348234 | Rbm11 | RNA binding motif protein 11 [Source:MGI Symbol;Acc:MGI:2447622] | 63,73 | 0,0 |
| ENSMUSG00000032551 | 119,1508959 | 237,1 | 2,0 | 7,776920625 | 0,000408627 | 1110059G10Rik | RIKEN cDNA 1110059G10 gene [Source:MGI Symbol;Acc:MGI:1913452] | 33,86 | 0,7 |
| ENSMUSG00000106887 | 8,921279129 | 20,9 | 0,0 | 7,777210153 | 0,04741438 | A930005G22Rik | RIKEN cDNA A930005G22 gene [Source:MGI Symbol;Acc:MGI:3528178] | 30,22 | 0,0 |
| ENSMUSG00000059811 | 416,3495456 | 565,4 | 138,6 | 7,779855946 | 0,000285361 | Atl2 | atlastin GTPase 2 [Source:MGI Symbol;Acc:MGI:1929492] | 99,10 | 17,5 |
| ENSMUSG00000056870 | 157,6726917 | 293,2 | 3,0 | 7,781173634 | 0,000135047 | Gulp1 | GULP, engulfment adaptor PTB domain containing 1 [Source:MGI Symbol;Acc:MGI:1920407] | 80,73 | 0,1 |
| ENSMUSG00000031970 | 22,73136148 | 21,6 | 0,0 | 7,784296413 | 0,046342945 | Dbndd1 | dysbindin (dystrobrevin binding protein 1) domain containing 1 [Source:MGI Symbol;Acc:MGI:1919435] | 52,46 | 0,0 |
| ENSMUSG00000099686 | 10,78704598 | 20,9 | 0,0 | 7,784749562 | 0,030961686 | Gm29460 | predicted gene 29460 [Source:MGI Symbol;Acc:MGI:5580166] | #DIV/0! | #DIV/0! |
| ENSMUSG00000090122 | 157,0654586 | 246,9 | 68,4 | 7,784972435 | 0,010973739 | Kcne1l | m voltage-gated channel, Isk-related family, member 1-like, pseudogene [Source:MGI Symbol;Acc:MGI:1 | 95,98 | 29,0 |
| ENSMUSG00000070923 | 234,4848441 | 418,4 | 16,9 | 7,787230388 | 0,001568134 | Klhl9 | kelch-like 9 [Source:MGI Symbol;Acc:MGI:2180122] | 99,82 | 4,9 |
| ENSMUSG00000046949 | 200,9334263 | 380,8 | 29,0 | 7,788353705 | 0,003206771 | Nqo2 | NAD(P)H dehydrogenase, quinone 2 [Source:MGI Symbol;Acc:MGI:104513] | 98,00 | 6,9 |
| ENSMUSG00000027956 | 109,4563842 | 173,8 | 1,8 | 7,79075144 | 0,008723079 | Tmem144 | transmembrane protein 144 [Source:MGI Symbol;Acc:MGI:1917902] | 87,39 | 0,6 |
| ENSMUSG00000011837 | 183,7623448 | 288,8 | 7,4 | 7,790995973 | 0,003406984 | Snapc2 | small nuclear RNA activating complex, polypeptide 2 [Source:MGI Symbol;Acc:MGI:1914861] | 92,76 | 5,8 |
| ENSMUSG00000074158 | 63,73357497 | 22,8 | 0,0 | 7,791938683 | 0,047000046 | 9830147E19Rik | RIKEN cDNA 9830147E19 gene [Source:MGI Symbol;Acc:MGI:3036263] | 47,89 | 0,0 |
| ENSMUSG00000029521 | 13,01982388 | 23,4 | 0,0 | 7,794978819 | 0,029889368 | Chek2 | checkpoint kinase 2 [Source:MGI Symbol;Acc:MGI:1355321] | 64,58 | 0,0 |
| ENSMUSG00000096458 | 113,6320022 | 209,3 | 1,8 | 7,796612423 | 0,00125629 | Moap1 | modulator of apoptosis 1 [Source:MGI Symbol;Acc:MGI:1915555] | 99,60 | 0,3 |
| ENSMUSG00000036819 | 74,80814806 | 114,5 | 0,9 | 7,79704405 | 0,013090322 | Jmjd4 | jumonji domain containing 4 [Source:MGI Symbol;Acc:MGI:2144404] | 74,37 | 0,4 |
| ENSMUSG00000031217 | 11,74392702 | 21,1 | 0,0 | 7,798105123 | 0,030390259 | Efnb1 | ephrin B1 [Source:MGI Symbol;Acc:MGI:102708] | 65,60 | 0,0 |
| ENSMUSG00000079045 | 10,43141061 | 20,8 | 0,0 | 7,799473751 | 0,031639983 | Prox1os | prospero homeobox 1, opposite strand [Source:MGI Symbol;Acc:MGI:4937200] | 98,79 | 0,0 |
| ENSMUSG00000053111 | 11,39850929 | 24,2 | 0,0 | 7,803561549 | 0,030384029 | Fank1 | fibronectin type 3 and ankyrin repeat domains 1 [Source:MGI Symbol;Acc:MGI:1914180] | 72,73 | 0,0 |
| ENSMUSG00000016918 | 199,2980946 | 169,7 | 0,9 | 7,804117961 | 0,00303887 | Sulf1 | sulfatase 1 [Source:MGI Symbol;Acc:MGI:2138563] | 66,07 | 0,0 |
| ENSMUSG00000091890 | 11,13883367 | 20,9 | 0,0 | 7,805235883 | 0,030868759 | A830073O21Rik | RIKEN cDNA A830073O21 gene [Source:MGI Symbol;Acc:MGI:2443692] | 22,30 | 0,0 |
| ENSMUSG00000052658 | 8,952032489 | 21,3 | 0,0 | 7,807895764 | 0,031553476 | 5830454E08Rik | RIKEN cDNA 5830454E08 gene [Source:MGI Symbol;Acc:MGI:1923350] | 94,22 | 0,0 |
| ENSMUSG00000029095 | 110,2326013 | 143,6 | 0,6 | 7,808563242 | 0,000848438 | Ablim2 | actin-binding LIM protein 2 [Source:MGI Symbol;Acc:MGI:2385758] | 87,01 | 0,4 |
| ENSMUSG00000045404 | 10,58515607 | 21,2 | 0,0 | 7,811895279 | 0,030475503 | Kcnk13 | potassium channel, subfamily K, member 13 [Source:MGI Symbol;Acc:MGI:2384976] | 61,31 | 0,0 |
| ENSMUSG00000100760 | 14,50390353 | 22,5 | 0,0 | 7,813691874 | 0,045607135 | Gm4035 | predicted gene 4035 [Source:MGI Symbol;Acc:MGI:3782209] | #DIV/0! | #DIV/0! |
| ENSMUSG00000020268 | 124,6761098 | 228,5 | 1,8 | 7,813929663 | 0,000632718 | Lyrm7 | LYR motif containing 7 [Source:MGI Symbol;Acc:MGI:1922780] | 98,92 | 1,8 |
| ENSMUSG00000039474 | 124,3588209 | 184,1 | 1,8 | 7,817076268 | 0,005425973 | Wfs1 | Wolfram syndrome 1 homolog (human) [Source:MGI Symbol;Acc:MGI:1328355] | 91,87 | 0,6 |
| ENSMUSG00000026809 | 25,66098527 | 22,2 | 0,0 | 7,818051068 | 0,034966623 | 1700026L06Rik | RIKEN cDNA 1700026L06 gene [Source:MGI Symbol;Acc:MGI:1917237] | 94,93 | 0,0 |
| ENSMUSG00000030029 | 140,1688123 | 199,8 | 2,0 | 7,820363013 | 0,002730442 | Lrig1 | leucine-rich repeats and immunoglobulin-like domains 1 [Source:MGI Symbol;Acc:MGI:107935] | 99,21 | 0,4 |
| ENSMUSG00000068264 | 245,2908829 | 381,2 | 64,8 | 7,82186049 | 0,003373155 | Ap5s1 | adaptor-related protein 5 complex, sigma 1 subunit [Source:MGI Symbol;Acc:MGI:1916846] | 96,02 | 25,0 |
| ENSMUSG00000055745 | 150,6534002 | 212,0 | 80,8 | 7,822041633 | 0,006824289 | Ldoc1l | leucine zipper, down-regulated in cancer 1-like [Source:MGI Symbol;Acc:MGI:2675858] | 97,29 | 16,4 |
| ENSMUSG00000045031 | 145,0138844 | 226,9 | 2,0 | 7,824925763 | 0,000820713 | Cetn4 | centrin 4 [Source:MGI Symbol;Acc:MGI:2677454] | 43,68 | 0,7 |
| ENSMUSG00000038506 | 210,6356697 | 360,3 | 200,6 | 7,826072336 | 0,002264897 | Dcun1d2 | l, defective in cullin neddylation 1, domain containing 2 (S. cerevisiae) [Source:MGI Symbol;Acc:MGI:214 | 65,32 | 19,2 |
| ENSMUSG00000096753 | 19,85349231 | 21,8 | 0,0 | 7,828081077 | 0,034412661 | Fam181a | family with sequence similarity 181, member A [Source:MGI Symbol;Acc:MGI:3647570] | 79,01 | 0,0 |
| ENSMUSG00000060572 | 17,05200872 | 24,5 | 0,0 | 7,829046154 | 0,039741539 | Mfap2 | microfibrillar-associated protein 2 [Source:MGI Symbol;Acc:MGI:99559] | 84,21 | 0,0 |
| ENSMUSG00000021871 | 255,7133129 | 371,9 | 26,8 | 7,830895884 | 0,001697391 | Pnp | purine-nucleoside phosphorylase [Source:MGI Symbol;Acc:MGI:97365] | 51,95 | 9,2 |
| ENSMUSG00000104876 | 18,47099908 | 22,0 | 0,0 | 7,835384484 | 0,038356903 | Trdc | T cell receptor delta, constant region [Source:MGI Symbol;Acc:MGI:98612] | #DIV/0! | #DIV/0! |
| ENSMUSG00000100155 | 14,02875584 | 23,7 | 0,0 | 7,836651884 | 0,037903724 | Gm2109 | predicted gene 2109 [Source:MGI Symbol;Acc:MGI:3780277] | #DIV/0! | #DIV/0! |
| ENSMUSG00000056306 | 8,453449761 | 21,5 | 0,0 | 7,83929201 | 0,031234123 | Sertm1 | serine rich and transmembrane domain containing 1 [Source:MGI Symbol;Acc:MGI:3607715] | 64,70 | 0,0 |
| ENSMUSG00000045671 | 199,2421156 | 389,9 | 10,9 | 7,840486898 | 0,001860059 | Spred2 | sprouty-related, EVH1 domain containing 2 [Source:MGI Symbol;Acc:MGI:2150019] | 92,63 | 6,4 |
| ENSMUSG00000073650 | 17,96455001 | 22,9 | 0,0 | 7,842620407 | 0,039016968 | Catip | ciliogenesis associated TTC17 interacting protein [Source:MGI Symbol;Acc:MGI:2685062] | 72,90 | 0,0 |
| ENSMUSG00000007817 | 232,1516433 | 374,4 | 19,0 | 7,844585873 | 0,000384973 | Zmiz1 | zinc finger, MIZ-type containing 1 [Source:MGI Symbol;Acc:MGI:3040693] | 97,81 | 1,2 |
| ENSMUSG00000024259 | 522,4021203 | 777,6 | 185,2 | 7,846447205 | 7,83545E-05 | Slc25a46 | solute carrier family 25, member 46 [Source:MGI Symbol;Acc:MGI:1914703] | 85,66 | 18,1 |
| ENSMUSG00000058486 | 125,9627989 | 160,8 | 1,0 | 7,84820822 | 0,002172517 | Wdr91 | WD repeat domain 91 [Source:MGI Symbol;Acc:MGI:2141558] | 92,69 | 0,6 |
| ENSMUSG00000018865 | 199,1840594 | 286,5 | 25,0 | 7,85360462 | 0,002450082 | Sult4a1 | sulfotransferase family 4A, member 1 [Source:MGI Symbol;Acc:MGI:1888971] | 95,90 | 12,4 |
| ENSMUSG00000030796 | 22,02911355 | 25,9 | 0,0 | 7,854349501 | 0,043183971 | Tead2 | TEA domain family member 2 [Source:MGI Symbol;Acc:MGI:104904] | 48,44 | 0,0 |
| ENSMUSG00000029352 | 10,1156734 | 22,0 | 0,0 | 7,854517264 | 0,037119338 | Crybb3 | crystallin, beta B3 [Source:MGI Symbol;Acc:MGI:102717] | 64,31 | 0,0 |
| ENSMUSG00000054659 | 9,134223935 | 21,9 | 0,0 | 7,860684729 | 0,028775952 | Pm20d2 | peptidase M20 domain containing 2 [Source:MGI Symbol;Acc:MGI:2685270] | 69,87 | 0,0 |
| ENSMUSG00000087596 | 29,84110715 | 22,3 | 0,0 | 7,86167057 | 0,041041021 | Gm11715 | predicted gene 11715 [Source:MGI Symbol;Acc:MGI:3702026] | 56,93 | 0,0 |
| ENSMUSG00000021866 | 23,50591597 | 22,9 | 0,0 | 7,862037205 | 0,038936575 | Anxa11 | annexin A11 [Source:MGI Symbol;Acc:MGI:108481] | 36,93 | 0,0 |
| ENSMUSG00000021143 | 236,8248485 | 368,2 | 4,6 | 7,862375974 | 9,06777E-05 | Pacs2 | phosphofurin acidic cluster sorting protein 2 [Source:MGI Symbol;Acc:MGI:1924399] | 99,38 | 1,3 |
| ENSMUSG00000105843 | 99,9393531 | 241,2 | 1,8 | 7,864069518 | 0,001838255 | Gm42644 | predicted gene 42644 [Source:MGI Symbol;Acc:MGI:5662781] | 56,59 | 1,3 |
| ENSMUSG00000081525 | 13,82518383 | 22,0 | 0,0 | 7,867633609 | 0,027862291 | Gm13325 | predicted gene 13325 [Source:MGI Symbol;Acc:MGI:3650421] | #DIV/0! | #DIV/0! |
| ENSMUSG00000027459 | 21,67580966 | 23,1 | 0,0 | 7,86764192 | 0,039812398 | Fam110a | family with sequence similarity 110, member A [Source:MGI Symbol;Acc:MGI:1921097] | 69,34 | 0,0 |
| ENSMUSG00000027244 | 310,5918455 | 451,0 | 32,3 | 7,868122912 | 0,000432263 | Atg13 | autophagy related 13 [Source:MGI Symbol;Acc:MGI:1196429] | 99,65 | 12,6 |
| ENSMUSG00000050619 | 114,7134943 | 153,7 | 1,0 | 7,868561887 | 0,002764018 | Zscan29 | zinc finger SCAN domains 29 [Source:MGI Symbol;Acc:MGI:2139317] | 72,19 | 0,2 |
| ENSMUSG00000054072 | 11,43223295 | 23,4 | 0,0 | 7,869106792 | 0,030939401 | Iigp1 | interferon inducible GTPase 1 [Source:MGI Symbol;Acc:MGI:1926259] | 61,81 | 0,0 |
| ENSMUSG00000024843 | 351,6647078 | 541,6 | 98,0 | 7,869335012 | 7,23739E-05 | Chka | choline kinase alpha [Source:MGI Symbol;Acc:MGI:107760] | 95,64 | 23,7 |
| ENSMUSG00000108350 | 21,35689116 | 22,1 | 0,0 | 7,870352828 | 0,034351736 | RP24-174G2.1 | | 69,93 | 0,0 |
| ENSMUSG00000021747 | 12,35590439 | 22,4 | 0,0 | 7,870851213 | 0,027283073 | 4930452B06Rik | RIKEN cDNA 4930452B06 gene [Source:MGI Symbol;Acc:MGI:1926154] | 73,18 | 0,0 |
| ENSMUSG00000078202 | 962,0327184 | 1648,6 | 359,8 | 7,871820206 | 8,51191E-06 | Nrarp | Notch-regulated ankyrin repeat protein [Source:MGI Symbol;Acc:MGI:1914372] | 100,00 | 34,2 |
| ENSMUSG00000007594 | 93,97757499 | 32,6 | 0,0 | 7,872939398 | 0,033811244 | Hapln4 | hyaluronan and proteoglycan link protein 4 [Source:MGI Symbol;Acc:MGI:2679531] | 62,89 | 0,0 |
| ENSMUSG00000038523 | 10,86344176 | 22,0 | 0,0 | 7,873554589 | 0,028684582 | 1700003F12Rik | RIKEN cDNA 1700003F12 gene [Source:MGI Symbol;Acc:MGI:1922730] | 79,14 | 0,0 |
| ENSMUSG00000037455 | 305,7729406 | 341,3 | 110,8 | 7,875988155 | 0,004681539 | Slc18b1 | solute carrier family 18, subfamily B, member 1 [Source:MGI Symbol;Acc:MGI:1923556] | 96,13 | 26,6 |
| ENSMUSG00000019971 | 123,8628981 | 248,4 | 1,8 | 7,87599349 | 0,000351418 | Cep290 | centrosomal protein 290 [Source:MGI Symbol;Acc:MGI:2384917] | 85,06 | 0,6 |
| ENSMUSG00000015214 | 270,9799039 | 388,0 | 125,7 | 7,876743488 | 0,002083055 | Mtmr1 | myotubularin related protein 1 [Source:MGI Symbol;Acc:MGI:1858271] | 98,22 | 19,2 |
| ENSMUSG00000027938 | 8,04853044 | 21,6 | 0,0 | 7,877077682 | 0,030892326 | Creb3l4 | cAMP responsive element binding protein 3-like 4 [Source:MGI Symbol;Acc:MGI:1916603] | 83,96 | 0,0 |
| ENSMUSG00000087601 | 14,45780947 | 22,9 | 0,0 | 7,880940712 | 0,027588436 | Uchl1os | ubiquitin carboxy-terminal hydrolase L1, opposite strand [Source:MGI Symbol;Acc:MGI:5446771] | 6,41 | 0,0 |
| ENSMUSG00000075419 | 134,8760062 | 218,4 | 3,0 | 7,880998261 | 0,004329646 | Dolk | dolichol kinase [Source:MGI Symbol;Acc:MGI:2677836] | 99,70 | 2,3 |
| ENSMUSG00000039450 | 179,9088061 | 341,2 | 69,9 | 7,887403746 | 0,001311604 | Dcxr | dicarbonyl L-xylulose reductase [Source:MGI Symbol;Acc:MGI:1915130] | 100,00 | 28,9 |
| ENSMUSG00000099407 | 12,21378355 | 22,6 | 0,0 | 7,889986063 | 0,024110602 | 1700060O08Rik | RIKEN cDNA 1700060O08 gene [Source:MGI Symbol;Acc:MGI:1914585] | 29,54 | 0,0 |
| ENSMUSG00000024846 | 10,45990613 | 22,5 | 0,0 | 7,890666354 | 0,035197577 | Cst6 | cystatin E/M [Source:MGI Symbol;Acc:MGI:1920970] | 10,51 | 0,0 |
| ENSMUSG00000049804 | 13,79728477 | 26,4 | 0,0 | 7,891151851 | 0,030961686 | Armcx4 | armadillo repeat containing, X-linked 4 [Source:MGI Symbol;Acc:MGI:2147887] | 28,11 | 0,0 |
| ENSMUSG00000057778 | 257,5157836 | 355,6 | 208,6 | 7,895494248 | 0,001808729 | Cyb5d2 | cytochrome b5 domain containing 2 [Source:MGI Symbol;Acc:MGI:2684848] | 98,00 | 30,8 |
| ENSMUSG00000087259 | 250,620041 | 469,2 | 22,4 | 7,896702163 | 0,000699848 | 2610035D17Rik | RIKEN cDNA 2610035D17 gene [Source:MGI Symbol;Acc:MGI:1919636] | #DIV/0! | #DIV/0! |
| ENSMUSG00000052928 | 21,5043255 | 24,7 | 0,0 | 7,899858939 | 0,038524442 | Ctif | CBP80/20-dependent translation initiation factor [Source:MGI Symbol;Acc:MGI:2685518] | 40,09 | 0,0 |
| ENSMUSG00000051022 | 21,05238857 | 26,7 | 0,0 | 7,900913612 | 0,025713947 | Hs3st1 | heparan sulfate (glucosamine) 3-O-sulfotransferase 1 [Source:MGI Symbol;Acc:MGI:1201606] | 62,06 | 0,0 |
| ENSMUSG00000069372 | 14,36386777 | 22,2 | 0,0 | 7,903914138 | 0,027055875 | Ctxn3 | cortexin 3 [Source:MGI Symbol;Acc:MGI:3642816] | 84,67 | 0,0 |
| ENSMUSG00000037362 | 13,22628076 | 22,8 | 0,0 | 7,904212472 | 0,026799326 | Nov | nephroblastoma overexpressed gene [Source:MGI Symbol;Acc:MGI:109185] | 59,84 | 0,0 |
| ENSMUSG00000039934 | 265,790389 | 465,9 | 148,9 | 7,904715631 | 0,001906853 | Gsap | gamma-secretase activating protein [Source:MGI Symbol;Acc:MGI:2442259] | 95,79 | 12,9 |
| ENSMUSG00000011154 | 14,83420625 | 23,4 | 0,0 | 7,905197718 | 0,028127231 | Cfap161 | cilia and flagella associated protein 161 [Source:MGI Symbol;Acc:MGI:1922806] | 39,44 | 0,0 |
| ENSMUSG00000024457 | 113,2727764 | 155,2 | 1,0 | 7,906925492 | 0,003568742 | Trim26 | tripartite motif-containing 26 [Source:MGI Symbol;Acc:MGI:1337056] | 97,55 | 0,8 |
| ENSMUSG00000028801 | 10,50537925 | 24,9 | 0,0 | 7,908526133 | 0,028391752 | Stpg1 | sperm tail PG rich repeat containing 1 [Source:MGI Symbol;Acc:MGI:1926056] | 66,96 | 0,0 |
| ENSMUSG00000039496 | 17,85802327 | 26,7 | 0,0 | 7,909492896 | 0,034269882 | Cdnf | cerebral dopamine neurotrophic factor [Source:MGI Symbol;Acc:MGI:3606576] | 68,69 | 0,0 |
| ENSMUSG00000053552 | 17,5251068 | 22,9 | 0,0 | 7,909908321 | 0,035299784 | Ebf4 | early B cell factor 4 [Source:MGI Symbol;Acc:MGI:2385972] | 66,38 | 0,0 |
| ENSMUSG00000023017 | 16,04699362 | 22,9 | 0,0 | 7,91016756 | 0,026466872 | Asic1 | acid-sensing (proton-gated) ion channel 1 [Source:MGI Symbol;Acc:MGI:1194915] | 62,58 | 0,0 |
| ENSMUSG00000030917 | 45,2433187 | 30,4 | 0,0 | 7,911376263 | 0,042287198 | Tmem159 | transmembrane protein 159 [Source:MGI Symbol;Acc:MGI:1925752] | 77,52 | 0,0 |
| ENSMUSG00000028389 | 159,4756196 | 273,1 | 3,0 | 7,918791781 | 0,000639077 | Zfp37 | zinc finger protein 37 [Source:MGI Symbol;Acc:MGI:99181] | 99,61 | 1,4 |
| ENSMUSG00000072258 | 18,91027904 | 22,5 | 0,0 | 7,919106445 | 0,031924952 | Taf1a | TA box binding protein (Tbp)-associated factor, RNA polymerase I, A [Source:MGI Symbol;Acc:MGI:1095 | 77,07 | 0,0 |
| ENSMUSG00000004843 | 723,2696758 | 1143,8 | 36,1 | 7,919181402 | 0,000851624 | Chmp2b | charged multivesicular body protein 2B [Source:MGI Symbol;Acc:MGI:1916192] | 100,00 | 20,0 |
| ENSMUSG00000032024 | 23,07335394 | 25,9 | 0,0 | 7,923369201 | 0,030285345 | Clmp | CXADR-like membrane protein [Source:MGI Symbol;Acc:MGI:1918816] | 59,44 | 0,0 |
| ENSMUSG00000034258 | 24,06810004 | 31,8 | 0,0 | 7,924009681 | 0,02491829 | Mfsd7c | major facilitator superfamily domain containing 7C [Source:MGI Symbol;Acc:MGI:2384974] | 56,38 | 0,0 |
| ENSMUSG00000031066 | 142,0316358 | 191,7 | 1,2 | 7,92573482 | 0,003764373 | Usp11 | ubiquitin specific peptidase 11 [Source:MGI Symbol;Acc:MGI:2384312] | 98,25 | 0,7 |
| ENSMUSG00000089665 | 13,39636274 | 22,6 | 0,0 | 7,926880884 | 0,030265477 | Fcor | Foxo1 corepressor [Source:MGI Symbol;Acc:MGI:1915484] | 98,04 | 0,0 |
| ENSMUSG00000040359 | 110,2470982 | 144,8 | 0,6 | 7,929771891 | 0,008372435 | Ufl1 | UFM1 specific ligase 1 [Source:MGI Symbol;Acc:MGI:1914740] | 86,86 | 0,5 |
| ENSMUSG00000028150 | 105,4677997 | 148,2 | 0,6 | 7,93028164 | 0,005719419 | Rorc | RAR-related orphan receptor gamma [Source:MGI Symbol;Acc:MGI:104856] | 82,20 | 0,9 |
| ENSMUSG00000032724 | 146,9512672 | 272,7 | 2,0 | 7,930510255 | 0,000210376 | Abtb2 | ankyrin repeat and BTB (POZ) domain containing 2 [Source:MGI Symbol;Acc:MGI:2139365] | 99,77 | 1,0 |
| ENSMUSG00000015647 | 10,67429528 | 24,0 | 0,0 | 7,930603287 | 0,026531842 | Lama5 | laminin, alpha 5 [Source:MGI Symbol;Acc:MGI:105382] | 23,92 | 0,0 |
| ENSMUSG00000044772 | 20,0690003 | 22,8 | 0,0 | 7,930905534 | 0,039826269 | Sntn | sentan, cilia apical structure protein [Source:MGI Symbol;Acc:MGI:3045373] | 95,13 | 0,0 |
| ENSMUSG00000055044 | 11,22704038 | 22,9 | 0,0 | 7,93144729 | 0,0277816 | Pdlim1 | PDZ and LIM domain 1 (elfin) [Source:MGI Symbol;Acc:MGI:1860611] | 46,34 | 0,0 |
| ENSMUSG00000086316 | 261,7068821 | 407,6 | 3,0 | 7,934107247 | 5,70284E-05 | 2210013O21Rik | RIKEN cDNA 2210013O21 gene [Source:MGI Symbol;Acc:MGI:1917373] | 63,93 | 2,0 |
| ENSMUSG00000022759 | 15,72827723 | 22,5 | 0,0 | 7,937785145 | 0,026023887 | Lrrc74b | leucine rich repeat containing 74B [Source:MGI Symbol;Acc:MGI:1921935] | 65,09 | 0,0 |
| ENSMUSG00000024507 | 376,0846306 | 645,3 | 171,9 | 7,946786063 | 0,000507497 | Hsd17b4 | hydroxysteroid (17-beta) dehydrogenase 4 [Source:MGI Symbol;Acc:MGI:105089] | 98,05 | 31,2 |
| ENSMUSG00000020520 | 18,29555907 | 22,7 | 0,0 | 7,947634517 | 0,025459641 | Galnt10 | tyl-alpha-D-galactosamine:polypeptide N-acetylgalactosaminyltransferase 10 [Source:MGI Symbol;Acc:MG | 44,99 | 0,0 |
| ENSMUSG00000017314 | 81,79907016 | 128,8 | 0,6 | 7,951917096 | 0,005859943 | Mpp2 | mbrane protein, palmitoylated 2 (MAGUK p55 subfamily member 2) [Source:MGI Symbol;Acc:MGI:18582 | 93,00 | 0,5 |
| ENSMUSG00000093553 | 12,76802414 | 27,5 | 0,0 | 7,9530054 | 0,024997762 | Gm20633 | predicted gene 20633 [Source:MGI Symbol;Acc:MGI:5313080] | 52,98 | 0,0 |
| ENSMUSG00000038982 | 290,6362239 | 461,2 | 60,1 | 7,956283308 | 0,000701769 | Bloc1s5 | biogenesis of lysosomal organelles complex-1, subunit 5, muted [Source:MGI Symbol;Acc:MGI:2178598] | 99,75 | 28,4 |
| ENSMUSG00000048807 | 189,2988899 | 312,0 | 147,7 | 7,956389177 | 0,004482858 | Slc35e4 | solute carrier family 35, member E4 [Source:MGI Symbol;Acc:MGI:2144150] | 86,56 | 22,1 |
| ENSMUSG00000097944 | 18,17296878 | 24,6 | 0,0 | 7,960495154 | 0,035884097 | A130014A01Rik | RIKEN cDNA A130014A01 gene [Source:MGI Symbol;Acc:MGI:3028043] | #DIV/0! | #DIV/0! |
| ENSMUSG00000027291 | 202,4246786 | 314,9 | 168,6 | 7,962564892 | 0,002528819 | Vps39 | vacuolar protein sorting 39 (yeast) [Source:MGI Symbol;Acc:MGI:2443189] | 96,98 | 23,1 |
| ENSMUSG00000030424 | 11,18003858 | 25,8 | 0,0 | 7,964678557 | 0,027111686 | Zfp939 | zinc finger protein 939 [Source:MGI Symbol;Acc:MGI:3036240] | #DIV/0! | #DIV/0! |
| ENSMUSG00000031870 | 10,32327717 | 23,6 | 0,0 | 7,965163783 | 0,027432837 | Pgr | progesterone receptor [Source:MGI Symbol;Acc:MGI:97567] | 31,34 | 0,0 |
| ENSMUSG00000097835 | 22,81717369 | 23,9 | 0,0 | 7,966004087 | 0,035569459 | Gm26910 | predicted gene, 26910 [Source:MGI Symbol;Acc:MGI:5477404] | #DIV/0! | #DIV/0! |
| ENSMUSG00000000085 | 98,86390825 | 199,9 | 1,0 | 7,967435963 | 0,000114954 | Scmh1 | sex comb on midleg homolog 1 [Source:MGI Symbol;Acc:MGI:1352762] | 98,98 | 0,4 |

| | | | | | | | | | |
|---|---|---|---|---|---|---|---|---|---|
| ENSMUSG00000044279 | 31,83192764 | 35,6 | 0,0 | 7,973061705 | 0,032022599 | Crb3 | crumbs family member 3 [Source:MGI Symbol;Acc:MGI:2670904] | 65,69 | 0,0 |
| ENSMUSG00000022751 | 188,016312 | 262,9 | 88,8 | 7,973132143 | 0,006117719 | Nit2 | nitrilase family, member 2 [Source:MGI Symbol;Acc:MGI:1261838] | 100,00 | 24,7 |
| ENSMUSG00000024219 | 261,7067503 | 448,8 | 86,9 | 7,973819515 | 0,001995574 | Anks1 | ankyrin repeat and SAM domain containing 1 [Source:MGI Symbol;Acc:MGI:2446180] | 73,31 | 14,1 |
| ENSMUSG00000029265 | 417,9888038 | 449,2 | 282,8 | 7,97801178 | 0,00133698 | Dr1 | down-regulator of transcription 1 [Source:MGI Symbol;Acc:MGI:1100515] | 99,76 | 31,0 |
| ENSMUSG00000107121 | 12,1201792 | 23,5 | 0,0 | 7,979797806 | 0,032476003 | 1110006O24Rik | RIKEN cDNA 1110006O24 gene [Source:MGI Symbol;Acc:MGI:1913373] | #DIV/0! | #DIV/0! |
| ENSMUSG00000038206 | 613,6710587 | 1105,8 | 116,7 | 7,979856908 | 0,000686115 | Fbxo8 | F-box protein 8 [Source:MGI Symbol;Acc:MGI:1354696] | 99,82 | 29,3 |
| ENSMUSG00000078994 | 12,0114398 | 24,2 | 0,0 | 7,981415669 | 0,026919049 | Zfp429 | zinc finger protein 429 [Source:MGI Symbol;Acc:MGI:1920057] | 73,18 | 0,0 |
| ENSMUSG00000074646 | 17,86437364 | 25,4 | 0,0 | 7,981632966 | 0,036767496 | 6430550D23Rik | RIKEN cDNA 6430550D23 gene [Source:MGI Symbol;Acc:MGI:2443361] | 62,11 | 0,0 |
| ENSMUSG00000031934 | 17,63408448 | 27,5 | 0,0 | 7,981830059 | 0,030375698 | Panx1 | pannexin 1 [Source:MGI Symbol;Acc:MGI:1860055] | 64,04 | 0,0 |
| ENSMUSG00000051262 | 12,75564395 | 23,8 | 0,0 | 7,982333826 | 0,036897733 | Cml3 | camello-like 3 [Source:MGI Symbol;Acc:MGI:2136449] | 26,20 | 0,0 |
| ENSMUSG00000046814 | 11,65722702 | 23,6 | 0,0 | 7,987781425 | 0,026425594 | Gchfr | GTP cyclohydrolase I feedback regulator [Source:MGI Symbol;Acc:MGI:2443977] | 96,95 | 0,0 |
| ENSMUSG00000022112 | 760,5132483 | 1295,0 | 49,9 | 7,990192687 | 0,000121059 | Gpc5 | glypican 5 [Source:MGI Symbol;Acc:MGI:1194894] | 98,97 | 8,4 |
| ENSMUSG00000026683 | 58,04357727 | 24,1 | 0,0 | 7,991803362 | 0,0300223 | Nuf2 | 2, NDC80 kinetochore complex component, homolog (S. cerevisiae) [Source:MGI Symbol;Acc:MGI:1914 | 40,50 | 0,0 |
| ENSMUSG00000030340 | 13,16045081 | 23,5 | 0,0 | 7,991982747 | 0,025713947 | Scnn1a | sodium channel, nonvoltage-gated 1 alpha [Source:MGI Symbol;Acc:MGI:101782] | 49,68 | 0,0 |
| ENSMUSG00000028167 | 204,9181882 | 417,7 | 55,7 | 7,992113587 | 0,003523103 | Bdh2 | 3-hydroxybutyrate dehydrogenase, type 2 [Source:MGI Symbol;Acc:MGI:1917022] | 99,86 | 29,8 |
| ENSMUSG00000021791 | 26,0540982 | 24,8 | 0,0 | 7,994685063 | 0,031656379 | Dydc2 | DPY30 domain containing 2 [Source:MGI Symbol;Acc:MGI:1918450] | 87,78 | 0,0 |
| ENSMUSG00000028926 | 322,2874965 | 500,7 | 3,0 | 7,994910712 | 1,10834E-06 | Cdk14 | cyclin-dependent kinase 14 [Source:MGI Symbol;Acc:MGI:894318] | 89,83 | 1,6 |
| ENSMUSG00000029333 | 12,94818684 | 27,0 | 0,0 | 7,999582014 | 0,026981852 | Rasgef1b | RasGEF domain family, member 1B [Source:MGI Symbol;Acc:MGI:2443755] | 66,28 | 0,0 |
| ENSMUSG00000046192 | 20,79069952 | 25,5 | 0,0 | 8,001819867 | 0,033353365 | Iqub | IQ motif and ubiquitin domain containing [Source:MGI Symbol;Acc:MGI:3041159] | 58,93 | 0,0 |
| ENSMUSG00000022044 | 135,5596779 | 167,4 | 0,6 | 8,00309104 | 0,001850845 | Stmn4 | stathmin-like 4 [Source:MGI Symbol;Acc:MGI:1931224] | 93,32 | 1,1 |
| ENSMUSG00000023092 | 672,7176838 | 1125,9 | 23,7 | 8,003535773 | 0,000142552 | Fhl1 | four and a half LIM domains 1 [Source:MGI Symbol;Acc:MGI:1298387] | 78,02 | 5,3 |
| ENSMUSG00000044337 | 15,32408268 | 27,2 | 0,0 | 8,006331903 | 0,031756332 | Ackr3 | atypical chemokine receptor 3 [Source:MGI Symbol;Acc:MGI:109562] | 51,42 | 0,0 |
| ENSMUSG00000026893 | 278,080922 | 649,1 | 30,5 | 8,007145236 | 5,29242E-05 | Gca | grancalcin [Source:MGI Symbol;Acc:MGI:1918521] | 99,74 | 7,9 |
| ENSMUSG00000019861 | 99,27529959 | 137,3 | 0,6 | 8,007413336 | 0,006895432 | Gopc | golgi associated PDZ and coiled-coil motif containing [Source:MGI Symbol;Acc:MGI:2149946] | 95,15 | 0,6 |
| ENSMUSG00000054474 | 139,9721154 | 203,2 | 1,8 | 8,009367623 | 0,004575845 | Thnsl2 | threonine synthase-like 2 (bacterial) [Source:MGI Symbol;Acc:MGI:3041254] | 85,64 | 0,9 |
| ENSMUSG00000038026 | 21,22321489 | 28,7 | 0,0 | 8,012875911 | 0,025245139 | Kcnj9 | potassium inwardly-rectifying channel, subfamily J, member 9 [Source:MGI Symbol;Acc:MGI:108007] | 69,48 | 0,0 |
| ENSMUSG00000021619 | 137,6613286 | 250,0 | 1,2 | 8,01319312 | 0,000533879 | Atg10 | autophagy related 10 [Source:MGI Symbol;Acc:MGI:1914045] | 99,36 | 2,4 |
| ENSMUSG00000036977 | 236,6903653 | 355,7 | 287,4 | 8,016825838 | 0,00293155 | Anapc10 | anaphase promoting complex subunit 10 [Source:MGI Symbol;Acc:MGI:1916249] | 92,04 | 9,5 |
| ENSMUSG00000022793 | 182,5345859 | 281,2 | 35,1 | 8,019084992 | 0,008077579 | B4galt4 | DP-Gal:betaGlcNAc beta 1,4-galactosyltransferase, polypeptide 4 [Source:MGI Symbol;Acc:MGI:192838 | 89,03 | 16,4 |
| ENSMUSG00000033111 | 710,3539597 | 1073,1 | 42,4 | 8,020068689 | 4,01052E-07 | 3830406C13Rik | RIKEN cDNA 3830406C13 gene [Source:MGI Symbol;Acc:MGI:1917937] | 99,85 | 11,4 |
| ENSMUSG00000050677 | 13,94416235 | 28,7 | 0,0 | 8,021578524 | 0,023659521 | Ccdc96 | coiled-coil domain containing 96 [Source:MGI Symbol;Acc:MGI:1913967] | 63,91 | 0,0 |
| ENSMUSG00000021265 | 128,8670772 | 275,3 | 3,0 | 8,023148697 | 0,001050231 | Slc25a29 | r family 25 (mitochondrial carrier, palmitoylcarnitine transporter), member 29 [Source:MGI Symbol;Acc:MC | 99,68 | 1,9 |
| ENSMUSG00000043165 | 16,18154366 | 25,0 | 0,0 | 8,02360423 | 0,027864545 | Lor | loricrin [Source:MGI Symbol;Acc:MGI:96816] | 65,69 | 0,0 |
| ENSMUSG00000044296 | 13,99440376 | 24,0 | 0,0 | 8,02394229 | 0,024544248 | Zfp879 | zinc finger protein 879 [Source:MGI Symbol;Acc:MGI:3053099] | 82,67 | 0,0 |
| ENSMUSG00000023249 | 92,34164446 | 147,5 | 0,6 | 8,024826367 | 0,005744721 | Parp3 | poly (ADP-ribose) polymerase family, member 3 [Source:MGI Symbol;Acc:MGI:1891258] | 84,59 | 0,4 |
| ENSMUSG00000031758 | 16,2817373 | 33,4 | 0,0 | 8,025168306 | 0,024292419 | Cdyl2 | chromodomain protein, Y chromosome-like 2 [Source:MGI Symbol;Acc:MGI:1923046] | 42,15 | 0,0 |
| ENSMUSG00000061374 | 209,4553663 | 327,7 | 74,8 | 8,026241408 | 0,003510352 | Fiz1 | Flt3 interacting zinc finger protein 1 [Source:MGI Symbol;Acc:MGI:1344336] | 79,63 | 19,2 |
| ENSMUSG00000024176 | 205,3559355 | 368,8 | 1,5 | 8,028265557 | 3,48363E-05 | Sox8 | SRY (sex determining region Y)-box 8 [Source:MGI Symbol;Acc:MGI:98370] | 98,21 | 0,1 |
| ENSMUSG00000071073 | 13,95856253 | 24,5 | 0,0 | 8,035426116 | 0,023617627 | Lrrc73 | leucine rich repeat containing 73 [Source:MGI Symbol;Acc:MGI:2684934] | 88,61 | 0,0 |
| ENSMUSG00000025870 | 10,92663563 | 25,0 | 0,0 | 8,036851479 | 0,025734641 | Arl10 | ADP-ribosylation factor-like 10 [Source:MGI Symbol;Acc:MGI:1930788] | 65,30 | 0,0 |
| ENSMUSG00000098176 | 8,14456333 | 24,2 | 0,0 | 8,036871671 | 0,027404618 | Ccdc166 | coiled-coil domain containing 166 [Source:MGI Symbol;Acc:MGI:1925902] | 64,45 | 0,0 |
| ENSMUSG00000031875 | 271,1529784 | 503,5 | 35,1 | 8,037232866 | 0,000262805 | Cmtm3 | CKLF-like MARVEL transmembrane domain containing 3 [Source:MGI Symbol;Acc:MGI:2447162] | 98,16 | 21,8 |
| ENSMUSG00000037482 | 24,51578728 | 25,8 | 0,0 | 8,040900281 | 0,028611236 | Erv3 | endogenous retroviral sequence 3 [Source:MGI Symbol;Acc:MGI:1919245] | 70,60 | 0,0 |
| ENSMUSG00000093606 | 10,70952769 | 26,4 | 0,0 | 8,041238374 | 0,028249014 | B130034C11Rik | RIKEN cDNA B130034C11 gene [Source:MGI Symbol;Acc:MGI:3041186] | #DIV/0! | #DIV/0! |
| ENSMUSG00000032036 | 94,28556693 | 183,2 | 0,6 | 8,041271067 | 0,000613788 | Kirrel3 | kin of IRRE like 3 (Drosophila) [Source:MGI Symbol;Acc:MGI:1914953] | 83,05 | 0,3 |
| ENSMUSG00000085737 | 16,29655046 | 24,7 | 0,0 | 8,042555636 | 0,025023 | Gm13442 | predicted gene 13442 [Source:MGI Symbol;Acc:MGI:3650661] | 78,73 | 0,0 |
| ENSMUSG00000060090 | 260,3024397 | 309,8 | 177,6 | 8,043711098 | 0,006705673 | Rp2 | retinitis pigmentosa 2 homolog (human) [Source:MGI Symbol;Acc:MGI:1277953] | 94,13 | 21,8 |
| ENSMUSG00000091002 | 23,00333386 | 26,8 | 0,0 | 8,044088795 | 0,027485495 | Tcerg1l | transcription elongation regulator 1-like [Source:MGI Symbol;Acc:MGI:1917821] | 70,84 | 0,0 |
| ENSMUSG00000028753 | 48,32932145 | 32,8 | 0,0 | 8,045169617 | 0,032425801 | Vwa5b1 | von Willebrand factor A domain containing 5B1 [Source:MGI Symbol;Acc:MGI:1922968] | 46,22 | 0,0 |
| ENSMUSG00000049532 | 189,0746248 | 336,0 | 78,6 | 8,050950763 | 0,0058277 | Sall2 | sal-like 2 (Drosophila) [Source:MGI Symbol;Acc:MGI:1354373] | 88,03 | 18,2 |
| ENSMUSG00000050963 | 9,556085169 | 24,7 | 0,0 | 8,052594807 | 0,026876988 | Kcns2 | K+ voltage-gated channel, subfamily S, 2 [Source:MGI Symbol;Acc:MGI:1197011] | 62,45 | 0,0 |
| ENSMUSG00000047866 | 330,6547435 | 578,9 | 175,5 | 8,056079936 | 0,001907102 | Lonp2 | lon peptidase 2, peroxisomal [Source:MGI Symbol;Acc:MGI:1914137] | 97,11 | 31,7 |
| ENSMUSG00000037253 | 398,167823 | 640,5 | 21,3 | 8,056893724 | 2,07881E-05 | Mex3c | mex3 homolog C (C. elegans) [Source:MGI Symbol;Acc:MGI:2652843] | 98,61 | 5,2 |
| ENSMUSG00000038319 | 24,24658581 | 29,4 | 0,0 | 8,059492425 | 0,022540487 | Kcnh2 | assium voltage-gated channel, subfamily H (eag-related), member 2 [Source:MGI Symbol;Acc:MGI:13417 | 53,41 | 0,0 |
| ENSMUSG00000031600 | 427,4722554 | 629,8 | 39,3 | 8,059793014 | 0,000300739 | Vps37a | vacuolar protein sorting 37A (yeast) [Source:MGI Symbol;Acc:MGI:1261835] | 99,42 | 5,0 |
| ENSMUSG00000040562 | 39,17674675 | 28,1 | 0,0 | 8,061349572 | 0,036067348 | Gstm2 | glutathione S-transferase, mu 2 [Source:MGI Symbol;Acc:MGI:95861] | 83,14 | 0,0 |
| ENSMUSG00000043498 | 25,58033106 | 41,0 | 0,0 | 8,061525773 | 0,025459641 | 9330132A10Rik | RIKEN cDNA 9330132A10 gene [Source:MGI Symbol;Acc:MGI:2442380] | #DIV/0! | #DIV/0! |
| ENSMUSG00000027865 | 208,0533929 | 356,1 | 25,0 | 8,061543979 | 0,00269238 | Gdap2 | ganglioside-induced differentiation-associated-protein 2 [Source:MGI Symbol;Acc:MGI:1338001] | 86,46 | 6,5 |
| ENSMUSG00000020258 | 29,13286828 | 37,2 | 0,0 | 8,063608882 | 0,037355635 | Glyctk | glycerate kinase [Source:MGI Symbol;Acc:MGI:2444085] | 61,40 | 0,0 |
| ENSMUSG00000042589 | 15,76164941 | 25,1 | 0,0 | 8,066891728 | 0,019317301 | Cux2 | cut-like homeobox 2 [Source:MGI Symbol;Acc:MGI:107321] | 32,85 | 0,0 |
| ENSMUSG00000020937 | 22,46173097 | 25,6 | 0,0 | 8,067232644 | 0,02615526 | Plcd3 | phospholipase C, delta 3 [Source:MGI Symbol;Acc:MGI:107451] | 77,55 | 0,0 |
| ENSMUSG00000024968 | 20,96638608 | 26,0 | 0,0 | 8,06854186 | 0,028021437 | Rcor2 | REST corepressor 2 [Source:MGI Symbol;Acc:MGI:1859854] | 73,10 | 0,0 |
| ENSMUSG00000085555 | 12,06137138 | 25,1 | 0,0 | 8,069074915 | 0,024671043 | 2610035F20Rik | RIKEN cDNA 2610035F20 gene [Source:MGI Symbol;Acc:MGI:1916498] | #DIV/0! | #DIV/0! |
| ENSMUSG00000024187 | 265,3974867 | 429,7 | 154,3 | 8,072621792 | 0,002690672 | Fam234a | family with sequence similarity 234, member A [Source:MGI Symbol;Acc:MGI:2146854] | 82,11 | 23,8 |
| ENSMUSG00000022364 | 23,57015228 | 25,8 | 0,0 | 8,078087057 | 0,02818166 | Tbc1d31 | TBC1 domain family, member 31 [Source:MGI Symbol;Acc:MGI:2684931] | 69,28 | 0,0 |
| ENSMUSG00000046873 | 48,09030297 | 26,2 | 0,0 | 8,079241188 | 0,023068642 | Mbtps2 | membrane-bound transcription factor peptidase, site 2 [Source:MGI Symbol;Acc:MGI:2444506] | 30,69 | 0,0 |
| ENSMUSG00000073758 | 12,25984231 | 24,8 | 0,0 | 8,080258268 | 0,024381117 | Sh3d21 | SH3 domain containing 21 [Source:MGI Symbol;Acc:MGI:1914188] | 72,30 | 0,0 |
| ENSMUSG00000029695 | 87,62601537 | 149,4 | 0,9 | 8,080482566 | 0,007449232 | Aass | aminoadipate-semialdehyde synthase [Source:MGI Symbol;Acc:MGI:1353573] | 90,36 | 0,7 |
| ENSMUSG00000031750 | 31,49475818 | 27,3 | 0,0 | 8,08065411 | 0,035164475 | Il34 | interleukin 34 [Source:MGI Symbol;Acc:MGI:1923777] | 79,46 | 0,0 |
| ENSMUSG00000041353 | 501,294939 | 607,4 | 120,1 | 8,086509012 | 0,000170322 | Tmem29 | transmembrane protein 29 [Source:MGI Symbol;Acc:MGI:1923420] | 73,18 | 22,4 |
| ENSMUSG00000099478 | 36,14338333 | 26,4 | 0,0 | 8,086875808 | 0,041496028 | Gm28370 | predicted gene 28370 [Source:MGI Symbol;Acc:MGI:5579076] | 64,73 | 0,0 |
| ENSMUSG00000070604 | 12,40548901 | 26,5 | 0,0 | 8,087621348 | 0,024247603 | Vsig10l | V-set and immunoglobulin domain containing 10 like [Source:MGI Symbol;Acc:MGI:1922940] | 65,95 | 0,0 |
| ENSMUSG00000075427 | 16,71844382 | 25,8 | 0,0 | 8,08832245 | 0,023122546 | Olfr288 | olfactory receptor 288 [Source:MGI Symbol;Acc:MGI:3030122] | 59,23 | 0,0 |
| ENSMUSG00000032377 | 104,6408116 | 199,9 | 0,6 | 8,09425236 | 0,000898233 | Plscr4 | phospholipid scramblase 4 [Source:MGI Symbol;Acc:MGI:2143267] | 91,69 | 0,7 |
| ENSMUSG00000043587 | 62,19537119 | 26,4 | 0,0 | 8,099663086 | 0,038235961 | Pxylp1 | 2-phosphoxylose phosphatase 1 [Source:MGI Symbol;Acc:MGI:2442444] | 62,77 | 0,0 |
| ENSMUSG00000035285 | 260,6335428 | 406,5 | 202,1 | 8,100447403 | 0,004754807 | Nat14 | N-acetyltransferase 14 [Source:MGI Symbol;Acc:MGI:3039561] | 77,88 | 22,7 |
| ENSMUSG00000044081 | 10,87221474 | 25,5 | 0,0 | 8,100961812 | 0,030910399 | Zfp85os | zinc finger protein 85, opposite strand [Source:MGI Symbol;Acc:MGI:1915521] | 54,74 | 0,0 |
| ENSMUSG00000068854 | 233,905124 | 475,6 | 112,0 | 8,10323409 | 0,001705667 | Hist2h2be | histone cluster 2, H2be [Source:MGI Symbol;Acc:MGI:2448415] | 99,91 | 31,1 |
| ENSMUSG00000038224 | 10,04408408 | 25,5 | 0,0 | 8,106364708 | 0,024861835 | Serpinf2 | serine (or cysteine) peptidase inhibitor, clade F, member 2 [Source:MGI Symbol;Acc:MGI:107173] | 80,02 | 0,0 |
| ENSMUSG00000030016 | 319,9747682 | 366,9 | 46,9 | 8,106495827 | 0,001811264 | Zfp638 | zinc finger protein 638 [Source:MGI Symbol;Acc:MGI:1203484] | 74,51 | 2,4 |
| ENSMUSG00000028525 | 340,6439771 | 677,5 | 3,1 | 8,107738216 | 0,000104218 | Pde4b | phosphodiesterase 4B, cAMP specific [Source:MGI Symbol;Acc:MGI:99557] | 59,83 | 0,8 |
| ENSMUSG00000087381 | 26,22009188 | 25,8 | 0,0 | 8,107825007 | 0,041414577 | Gm16008 | predicted gene 16008 [Source:MGI Symbol;Acc:MGI:3802048] | 84,74 | 0,0 |
| ENSMUSG00000035847 | 617,3466869 | 719,8 | 106,3 | 8,109693803 | 9,58287E-06 | Ids | iduronate 2-sulfatase [Source:MGI Symbol;Acc:MGI:96417] | 96,62 | 20,7 |
| ENSMUSG00000021779 | 329,4712778 | 500,2 | 73,2 | 8,110933763 | 0,001820656 | Thrb | thyroid hormone receptor beta [Source:MGI Symbol;Acc:MGI:98743] | 91,36 | 12,5 |
| ENSMUSG00000027186 | 17,31870759 | 27,0 | 0,0 | 8,112782321 | 0,027153229 | Elf5 | E74-like factor 5 [Source:MGI Symbol;Acc:MGI:1335079] | 59,63 | 0,0 |
| ENSMUSG00000027860 | 15,81144173 | 25,7 | 0,0 | 8,113126193 | 0,025233283 | Vangl1 | vang-like 1 (van gogh, Drosophila) [Source:MGI Symbol;Acc:MGI:2159344] | 39,22 | 0,0 |
| ENSMUSG00000034591 | 15,72152147 | 26,2 | 0,0 | 8,114374487 | 0,022313912 | Slc41a2 | solute carrier family 41, member 2 [Source:MGI Symbol;Acc:MGI:2442940] | 56,68 | 0,0 |
| ENSMUSG00000043541 | 13,79764475 | 26,2 | 0,0 | 8,114932037 | 0,028808928 | Casc1 | cancer susceptibility candidate 1 [Source:MGI Symbol;Acc:MGI:2444480] | 61,10 | 0,0 |
| ENSMUSG00000067377 | 197,8650126 | 276,5 | 49,0 | 8,115280167 | 0,008240475 | Tspan6 | tetraspanin 6 [Source:MGI Symbol;Acc:MGI:1926264] | 97,20 | 17,1 |
| ENSMUSG00000038840 | 16,84971775 | 28,0 | 0,0 | 8,120492111 | 0,022540487 | Birc7 | baculoviral IAP repeat-containing 7 (livin) [Source:MGI Symbol;Acc:MGI:2676458] | 94,93 | 0,0 |
| ENSMUSG00000029182 | 23,04059176 | 27,4 | 0,0 | 8,120998774 | 0,026940557 | 1700001C02Rik | RIKEN cDNA 1700001C02 gene [Source:MGI Symbol;Acc:MGI:1922684] | 77,07 | 0,0 |
| ENSMUSG00000071660 | 217,3974317 | 271,1 | 1,2 | 8,121074849 | 0,000432662 | Ttc9c | tetratricopeptide repeat domain 9C [Source:MGI Symbol;Acc:MGI:1917637] | 100,00 | 0,6 |
| ENSMUSG00000097785 | 17,44469948 | 25,9 | 0,0 | 8,121567621 | 0,031306045 | B230217O12Rik | RIKEN cDNA B230217O12 gene [Source:MGI Symbol;Acc:MGI:2444948] | #DIV/0! | #DIV/0! |
| ENSMUSG00000022151 | 383,6558718 | 484,4 | 191,3 | 8,122492363 | 0,000606925 | Ttc33 | tetratricopeptide repeat domain 33 [Source:MGI Symbol;Acc:MGI:1914765] | 85,41 | 27,0 |
| ENSMUSG00000025491 | 11,49159293 | 26,7 | 0,0 | 8,12437602 | 0,031297359 | Ifitm1 | interferon induced transmembrane protein 1 [Source:MGI Symbol;Acc:MGI:1915963] | 50,27 | 0,0 |
| ENSMUSG00000085614 | 16,34780706 | 28,4 | 0,0 | 8,129937816 | 0,028969349 | 1700123M08Rik | RIKEN cDNA 1700123M08 gene [Source:MGI Symbol;Acc:MGI:1923915] | #DIV/0! | #DIV/0! |
| ENSMUSG00000024901 | 26,30078253 | 27,5 | 0,0 | 8,131257243 | 0,030440412 | Peli3 | pellino 3 [Source:MGI Symbol;Acc:MGI:1924963] | 73,48 | 0,0 |
| ENSMUSG00000030616 | 30,61123859 | 29,5 | 0,0 | 8,132299367 | 0,02876575 | Sytl2 | synaptotagmin-like 2 [Source:MGI Symbol;Acc:MGI:1933366] | 47,57 | 0,0 |
| ENSMUSG00000043460 | 11,28896068 | 26,0 | 0,0 | 8,132929857 | 0,023893724 | Elfn2 | leucine rich repeat and fibronectin type III, extracellular 2 [Source:MGI Symbol;Acc:MGI:3608416] | 60,49 | 0,0 |
| ENSMUSG00000037341 | 16,06247425 | 29,1 | 0,0 | 8,13370059 | 0,023963738 | Slc9a7 | solute carrier family 9 (sodium/hydrogen exchanger), member 7 [Source:MGI Symbol;Acc:MGI:2444530] | 67,97 | 0,0 |
| ENSMUSG00000030386 | 261,9631705 | 397,7 | 5,4 | 8,137070644 | 0,001505163 | Zfp606 | zinc finger protein 606 [Source:MGI Symbol;Acc:MGI:1914620] | 95,30 | 1,5 |
| ENSMUSG00000018819 | 32,76939707 | 27,3 | 0,0 | 8,139487132 | 0,0349724 | Lsp1 | lymphocyte specific 1 [Source:MGI Symbol;Acc:MGI:96832] | 60,85 | 0,0 |
| ENSMUSG00000103041 | 27,48770643 | 27,3 | 0,0 | 8,140251421 | 0,0277816 | Gm37305 | predicted gene, 37305 [Source:MGI Symbol;Acc:MGI:5610533] | 42,69 | 0,0 |
| ENSMUSG00000030259 | 18,1057927 | 26,4 | 0,0 | 8,140290189 | 0,028116857 | Rassf8 | s association (RalGDS/AF-6) domain family (N-terminal) member 8 [Source:MGI Symbol;Acc:MGI:19185 | 50,79 | 0,0 |
| ENSMUSG00000083284 | 11,43436033 | 26,4 | 0,0 | 8,143603929 | 0,023641079 | Gm13688 | predicted gene 13688 [Source:MGI Symbol;Acc:MGI:3652062] | #DIV/0! | #DIV/0! |
| ENSMUSG00000041323 | 12,07003305 | 28,2 | 0,0 | 8,143878115 | 0,023444444 | Ak7 | adenylate kinase 7 [Source:MGI Symbol;Acc:MGI:1926051] | 73,23 | 0,0 |
| ENSMUSG00000020473 | 32,11810322 | 33,5 | 0,0 | 8,147256044 | 0,028643912 | Aebp1 | AE binding protein 1 [Source:MGI Symbol;Acc:MGI:1197012] | 69,15 | 0,0 |
| ENSMUSG00000024098 | 311,727659 | 427,0 | 64,2 | 8,14908551 | 0,00338592 | Twsg1 | twisted gastrulation homolog 1 (Drosophila) [Source:MGI Symbol;Acc:MGI:2137520] | 99,30 | 5,9 |
| ENSMUSG00000084915 | 22,9526456 | 27,2 | 0,0 | 8,154035311 | 0,030860443 | C230037L18Rik | RIKEN cDNA C230037L18 gene [Source:MGI Symbol;Acc:MGI:3697701] | 61,92 | 0,0 |
| ENSMUSG00000046959 | 13,21444639 | 28,0 | 0,0 | 8,155116206 | 0,026933915 | Slc26a1 | solute carrier family 26 (sulfate transporter), member 1 [Source:MGI Symbol;Acc:MGI:2385894] | 59,31 | 0,0 |
| ENSMUSG00000045532 | 20,73106404 | 31,2 | 0,0 | 8,155931678 | 0,022549461 | C1ql1 | complement component 1, q subcomponent-like 1 [Source:MGI Symbol;Acc:MGI:1344400] | 86,63 | 0,0 |
| ENSMUSG00000032586 | 16,64995907 | 31,5 | 0,0 | 8,156635196 | 0,031573046 | Traip | TRAF-interacting protein [Source:MGI Symbol;Acc:MGI:1096377] | 64,77 | 0,0 |
| ENSMUSG00000042251 | 104,1401017 | 205,9 | 1,0 | 8,158404754 | 0,001901811 | Pm20d1 | peptidase M20 domain containing 1 [Source:MGI Symbol;Acc:MGI:2442939] | 54,89 | 0,2 |
| ENSMUSG00000038776 | 266,5702175 | 500,4 | 36,9 | 8,159807141 | 0,000392863 | Ephx1 | epoxide hydrolase 1, microsomal [Source:MGI Symbol;Acc:MGI:95405] | 87,71 | 17,5 |
| ENSMUSG00000047804 | 139,6417759 | 173,1 | 1,0 | 8,164030955 | 0,003602277 | Akap10 | A kinase (PRKA) anchor protein 10 [Source:MGI Symbol;Acc:MGI:1890218] | 60,29 | 0,3 |
| ENSMUSG00000071719 | 12,95046036 | 26,3 | 0,0 | 8,164079447 | 0,022785134 | Tmem28 | transmembrane protein 28 [Source:MGI Symbol;Acc:MGI:3648377] | 68,44 | 0,0 |
| ENSMUSG00000097078 | 15,46823241 | 28,1 | 0,0 | 8,165203346 | 0,029860531 | Gm26566 | predicted gene, 26566 [Source:MGI Symbol;Acc:MGI:5477060] | 66,28 | 0,0 |
| ENSMUSG00000032415 | 21,03531385 | 27,2 | 0,0 | 8,166581344 | 0,030176595 | Ube2cbp | ubiquitin-conjugating enzyme E2C binding protein [Source:MGI Symbol;Acc:MGI:1917598] | 88,19 | 0,0 |
| ENSMUSG00000006301 | 383,1602601 | 528,9 | 227,5 | 8,170588062 | 0,000928541 | Tmbim1 | transmembrane BAX inhibitor motif containing 1 [Source:MGI Symbol;Acc:MGI:1916910] | 22,29 | 6,2 |
| ENSMUSG00000072066 | 137,3716215 | 246,8 | 1,8 | 8,17472916 | 0,001898572 | 6720489N17Rik | RIKEN cDNA 6720489N17 gene [Source:MGI Symbol;Acc:MGI:2443901] | 98,75 | 1,1 |
| ENSMUSG00000047648 | 114,7847172 | 162,6 | 0,6 | 8,176887219 | 0,002857421 | Fbxo30 | F-box protein 30 [Source:MGI Symbol;Acc:MGI:1919115] | 92,90 | 0,5 |
| ENSMUSG00000076617 | 14,22081593 | 28,7 | 0,0 | 8,178351965 | 0,022180393 | Ighm | immunoglobulin heavy constant mu [Source:MGI Symbol;Acc:MGI:96448] | #DIV/0! | #DIV/0! |
| ENSMUSG00000036459 | 19,44312905 | 28,8 | 0,0 | 8,178597093 | 0,02563122 | Wtip | WT1-interacting protein [Source:MGI Symbol;Acc:MGI:2141920] | 81,24 | 0,0 |
| ENSMUSG00000092192 | 23,17932715 | 29,9 | 0,0 | 8,178636679 | 0,031188457 | Dyx1c1 | dyslexia susceptibility 1 candidate 1 [Source:MGI Symbol;Acc:MGI:1914935] | 83,53 | 0,0 |
| ENSMUSG00000022178 | 18,3599098 | 27,5 | 0,0 | 8,180662531 | 0,020328758 | Ajuba | ajuba LIM protein [Source:MGI Symbol;Acc:MGI:1341886] | 70,46 | 0,0 |
| ENSMUSG00000023046 | 16,84762361 | 28,0 | 0,0 | 8,181046225 | 0,030880846 | Igfbp6 | insulin-like growth factor binding protein 6 [Source:MGI Symbol;Acc:MGI:96441] | 80,91 | 0,0 |
| ENSMUSG00000031786 | 168,3336543 | 190,8 | 1,0 | 8,182795803 | 0,001239038 | Drc7 | dynein regulatory complex subunit 7 [Source:MGI Symbol;Acc:MGI:2685616] | 99,22 | 0,9 |
| ENSMUSG00000003623 | 788,2001875 | 1461,8 | 146,3 | 8,183781815 | 6,8726E-07 | Crot | carnitine O-octanoyltransferase [Source:MGI Symbol;Acc:MGI:1921364] | 96,21 | 18,7 |
| ENSMUSG00000097316 | 15,04290007 | 29,5 | 0,0 | 8,186745322 | 0,02097435 | Gm10516 | predicted gene 10516 [Source:MGI Symbol;Acc:MGI:3641979] | 70,62 | 0,0 |
| ENSMUSG00000063428 | 279,3152458 | 439,1 | 89,6 | 8,187889608 | 0,002534902 | Ddo | D-aspartate oxidase [Source:MGI Symbol;Acc:MGI:1925528] | 99,87 | 26,1 |
| ENSMUSG00000071341 | 19,02427073 | 30,1 | 0,0 | 8,188232347 | 0,021900902 | Egr4 | early growth response 4 [Source:MGI Symbol;Acc:MGI:99252] | 85,60 | 0,0 |
| ENSMUSG00000019320 | 18,69210732 | 27,3 | 0,0 | 8,189168443 | 0,021407699 | Noxo1 | NADPH oxidase organizer 1 [Source:MGI Symbol;Acc:MGI:1919143] | 82,93 | 0,0 |
| ENSMUSG00000042510 | 14,78146338 | 27,2 | 0,0 | 8,189202725 | 0,02185231 | AA986860 | expressed sequence AA986860 [Source:MGI Symbol;Acc:MGI:2138143] | 76,22 | 0,0 |
| ENSMUSG00000057156 | 148,0980761 | 183,5 | 1,0 | 8,191734104 | 0,003090135 | Homez | homeodomain leucine zipper-encoding gene [Source:MGI Symbol;Acc:MGI:2678023] | 74,28 | 0,4 |
| ENSMUSG00000085193 | 15,36840763 | 27,3 | 0,0 | 8,194666367 | 0,025966911 | Gm13322 | predicted gene 13322 [Source:MGI Symbol;Acc:MGI:3650424] | #DIV/0! | #DIV/0! |
| ENSMUSG00000026611 | 21,01454075 | 27,7 | 0,0 | 8,197006921 | 0,026443441 | Spata17 | spermatogenesis associated 17 [Source:MGI Symbol;Acc:MGI:1921967] | 58,75 | 0,0 |
| ENSMUSG00000038079 | 346,1500971 | 406,7 | 54,5 | 8,197278992 | 0,002096022 | Tmem237 | transmembrane protein 237 [Source:MGI Symbol;Acc:MGI:2138365] | 95,63 | 19,6 |
| ENSMUSG00000044252 | 1190,735998 | 1959,8 | 86,3 | 8,197390857 | 2,63496E-05 | Osbpl1a | oxysterol binding protein-like 1A [Source:MGI Symbol;Acc:MGI:1927551] | 77,30 | 13,8 |
| ENSMUSG00000022130 | 89,3212747 | 149,2 | 0,6 | 8,198915873 | 0,00759766 | Tgds | TDP-glucose 4,6-dehydratase [Source:MGI Symbol;Acc:MGI:1923605] | 79,12 | 1,1 |
| ENSMUSG00000008435 | 74,24998853 | 33,5 | 0,0 | 8,200526642 | 0,02338691 | Rdh13 | retinol dehydrogenase 13 (all-trans and 9-cis) [Source:MGI Symbol;Acc:MGI:1918732] | 67,40 | 0,0 |
| ENSMUSG00000000902 | 319,7287086 | 548,8 | 88,7 | 8,202238516 | 0,000393615 | Smarcb1 | matrix associated, actin dependent regulator of chromatin, subfamily b, member 1 [Source:MGI Symbol;A | 99,90 | 25,8 |
| ENSMUSG00000067369 | 200,2096448 | 330,3 | 146,7 | 8,202267792 | 0,004023292 | Trmt2b | TRM2 tRNA methyltransferase 2B [Source:MGI Symbol;Acc:MGI:2442530] | 96,72 | 26,1 |
| ENSMUSG00000092544 | 19,14909943 | 27,7 | 0,0 | 8,203002617 | 0,022742322 | Gm20422 | predicted gene 20422 [Source:MGI Symbol;Acc:MGI:5141887] | 92,69 | 0,0 |
| ENSMUSG00000045655 | 16,16570611 | 29,3 | 0,0 | 8,203700579 | 0,026466872 | Fam216b | family with sequence similarity 216, member B [Source:MGI Symbol;Acc:MGI:2145738] | 84,61 | 0,0 |
| ENSMUSG00000027180 | 646,0047545 | 1055,5 | 44,4 | 8,203905018 | 8,57275E-08 | Fbxo3 | F-box protein 3 [Source:MGI Symbol;Acc:MGI:1929084] | 78,74 | 5,3 |
| ENSMUSG00000049154 | 16,84719255 | 27,6 | 0,0 | 8,205256103 | 0,028582566 | Fam183b | family with sequence similarity 183, member B [Source:MGI Symbol;Acc:MGI:1922679] | 95,77 | 0,0 |
| ENSMUSG00000103755 | 11,73580767 | 27,1 | 0,0 | 8,206515238 | 0,02246819 | Gm37805 | predicted gene, 37805 [Source:MGI Symbol;Acc:MGI:5611033] | 98,17 | 0,0 |
| ENSMUSG00000034532 | 28,06704162 | 29,5 | 0,0 | 8,207116958 | 0,033174792 | Fbxo16 | F-box protein 16 [Source:MGI Symbol;Acc:MGI:1354706] | 96,53 | 0,0 |
| ENSMUSG00000103270 | 18,39196428 | 34,1 | 0,0 | 8,209178818 | 0,020808934 | Gm37915 | predicted gene, 37915 [Source:MGI Symbol;Acc:MGI:5611143] | 56,32 | 0,0 |
| ENSMUSG00000057367 | 290,7616111 | 219,9 | 1,0 | 8,21085525 | 0,000422132 | Birc2 | baculoviral IAP repeat-containing 2 [Source:MGI Symbol;Acc:MGI:1197009] | 95,06 | 0,7 |
| ENSMUSG00000030513 | 439,5542671 | 854,5 | 3,8 | 8,211432197 | 3,22281E-06 | Pcsk6 | proprotein convertase subtilisin/kexin type 6 [Source:MGI Symbol;Acc:MGI:102897] | 97,24 | 1,2 |
| ENSMUSG00000053153 | 38,60915805 | 27,5 | 0,0 | 8,211572429 | 0,0277816 | Spag16 | sperm associated antigen 16 [Source:MGI Symbol;Acc:MGI:1913972] | 56,82 | 0,0 |
| ENSMUSG00000023064 | 18,69547878 | 30,8 | 0,0 | 8,21341365 | 0,020710182 | Sncg | synuclein, gamma [Source:MGI Symbol;Acc:MGI:1298397] | 88,93 | 0,0 |
| ENSMUSG00000006154 | 18,55858761 | 44,9 | 0,0 | 8,218378144 | 0,021387317 | Eps8l1 | EPS8-like 1 [Source:MGI Symbol;Acc:MGI:1914675] | 54,27 | 0,0 |
| ENSMUSG00000068099 | 350,597587 | 533,9 | 162,2 | 8,220573823 | 0,001196977 | 1500009C09Rik | RIKEN cDNA 1500009C09 gene [Source:MGI Symbol;Acc:MGI:1923755] | 78,82 | 25,2 |
| ENSMUSG00000032081 | 19,34984546 | 29,5 | 0,0 | 8,221296228 | 0,024345705 | Apoc3 | apolipoprotein C-III [Source:MGI Symbol;Acc:MGI:88055] | 44,28 | 0,0 |
| ENSMUSG00000073791 | 22,47423172 | 28,8 | 0,0 | 8,222566183 | 0,030961686 | Efcab7 | EF-hand calcium binding domain 7 [Source:MGI Symbol;Acc:MGI:2385199] | 75,87 | 0,0 |
| ENSMUSG00000045246 | 198,3291095 | 301,2 | 2,0 | 8,225038433 | 0,000467449 | Kcng4 | potassium voltage-gated channel, subfamily G, member 4 [Source:MGI Symbol;Acc:MGI:1913983] | 98,64 | 0,0 |
| ENSMUSG00000061898 | 18,96404323 | 28,9 | 0,0 | 8,225664856 | 0,030648846 | Rbak | RB-associated KRAB zinc finger [Source:MGI Symbol;Acc:MGI:1927369] | 74,03 | 0,0 |
| ENSMUSG00000098682 | 18,11720488 | 32,1 | 0,0 | 8,228791994 | 0,019734779 | Otx2os1 | orthodenticle homolog 2 opposite strand 1 [Source:MGI Symbol;Acc:MGI:3583292] | #DIV/0! | #DIV/0! |
| ENSMUSG00000029201 | 335,0432402 | 570,3 | 286,9 | 8,230989276 | 0,001049358 | Ugdh | UDP-glucose dehydrogenase [Source:MGI Symbol;Acc:MGI:1306785] | 93,33 | 31,1 |
| ENSMUSG00000028078 | 137,4035564 | 257,8 | 2,0 | 8,23167904 | 0,001829475 | Dclk2 | doublecortin-like kinase 2 [Source:MGI Symbol;Acc:MGI:1918012] | 46,93 | 0,3 |
| ENSMUSG00000025889 | 292,2539689 | 459,6 | 333,3 | 8,232137982 | 0,001907102 | Snca | synuclein, alpha [Source:MGI Symbol;Acc:MGI:1277151] | 93,31 | 24,9 |
| ENSMUSG00000079499 | 29,0972766 | 31,5 | 0,0 | 8,232821722 | 0,024381117 | 6530402F18Rik | RIKEN cDNA 6530402F18 gene [Source:MGI Symbol;Acc:MGI:1923470] | #DIV/0! | #DIV/0! |
| ENSMUSG00000028641 | 21,82902977 | 31,9 | 0,0 | 8,233517672 | 0,020240641 | P3h1 | prolyl 3-hydroxylase 1 [Source:MGI Symbol;Acc:MGI:1888921] | 74,79 | 0,0 |
| ENSMUSG00000096972 | 17,02499868 | 28,7 | 0,0 | 8,233729311 | 0,033116718 | Gm26883 | predicted gene, 26883 [Source:MGI Symbol;Acc:MGI:5477377] | #DIV/0! | #DIV/0! |
| ENSMUSG00000036249 | 103,1258618 | 167,2 | 1,0 | 8,237304675 | 0,008002308 | Rbm43 | RNA binding motif protein 43 [Source:MGI Symbol;Acc:MGI:1918934] | 95,69 | 0,7 |
| ENSMUSG00000014791 | 18,3490713 | 28,6 | 0,0 | 8,239500444 | 0,020333974 | Elmo3 | engulfment and cell motility 3 [Source:MGI Symbol;Acc:MGI:2679007] | 72,28 | 0,0 |
| ENSMUSG00000105284 | 13,87001249 | 29,2 | 0,0 | 8,239505414 | 0,021315097 | Gm42693 | predicted gene 42693 [Source:MGI Symbol;Acc:MGI:5662830] | 97,46 | 0,0 |
| ENSMUSG00000022091 | 330,3765287 | 567,1 | 113,8 | 8,243604098 | 0,001787001 | Sorbs3 | sorbin and SH3 domain containing 3 [Source:MGI Symbol;Acc:MGI:700013] | 97,60 | 23,8 |

| | | | | | | | | | |
|---|---|---|---|---|---|---|---|---|---|
| ENSMUSG00000106894 | 17,13433041 | 28,3 | 0,0 | 8,244830553 | 0,030995326 | Gm17130 | predicted gene 17130 [Source:MGI Symbol;Acc:MGI:4937957] | 3,09 | 0,0 |
| ENSMUSG00000001750 | 53,73795745 | 30,4 | 0,0 | 8,247295695 | 0,03394 | Tcirg1 | mmune regulator 1, ATPase, H+ transporting, lysosomal V0 protein A3 [Source:MGI Symbol;Acc:MGI:13 | 66,35 | 0,0 |
| ENSMUSG00000097596 | 22,32499578 | 29,6 | 0,0 | 8,248315788 | 0,032220951 | Gm26673 | predicted gene, 26673 [Source:MGI Symbol;Acc:MGI:5477167] | 64,98 | 0,0 |
| ENSMUSG00000009378 | 34,50130348 | 48,6 | 0,0 | 8,251212779 | 0,016643822 | Slc16a12 | ute carrier family 16 (monocarboxylic acid transporters), member 12 [Source:MGI Symbol;Acc:MGI:21477 | 60,13 | 0,0 |
| ENSMUSG00000021180 | 50,41476432 | 34,6 | 0,0 | 8,253657955 | 0,033322782 | Rps6ka5 | ribosomal protein S6 kinase, polypeptide 5 [Source:MGI Symbol;Acc:MGI:1920336] | 57,92 | 0,0 |
| ENSMUSG00000100811 | 13,14675406 | 29,8 | 0,0 | 8,253802715 | 0,024577008 | Gm29083 | predicted gene 29083 [Source:MGI Symbol;Acc:MGI:5579789] | 87,68 | 0,0 |
| ENSMUSG00000031963 | 33,24201816 | 35,0 | 0,0 | 8,255306649 | 0,032492661 | Bmper | BMP-binding endothelial regulator [Source:MGI Symbol;Acc:MGI:1920480] | 63,07 | 0,0 |
| ENSMUSG00000026679 | 14,32219968 | 30,7 | 0,0 | 8,256661531 | 0,019947179 | Enkur | enkurin, TRPC channel interacting protein [Source:MGI Symbol;Acc:MGI:1918483] | 89,74 | 0,0 |
| ENSMUSG00000051107 | 18,51403992 | 28,9 | 0,0 | 8,259499247 | 0,020000617 | Gm15440 | predicted gene 15440 [Source:MGI Symbol;Acc:MGI:3642353] | 98,92 | 0,0 |
| ENSMUSG00000056854 | 17,46959509 | 34,5 | 0,0 | 8,260332444 | 0,02021691 | Pou3f4 | POU domain, class 3, transcription factor 4 [Source:MGI Symbol;Acc:MGI:101894] | 71,52 | 0,0 |
| ENSMUSG00000004951 | 15,7190402 | 29,0 | 0,0 | 8,262803457 | 0,021141624 | Hspb1 | heat shock protein 1 [Source:MGI Symbol;Acc:MGI:96240] | 58,00 | 0,0 |
| ENSMUSG00000038122 | 25,31696481 | 33,5 | 0,0 | 8,268542748 | 0,027404618 | Tbc1d32 | TBC1 domain family, member 32 [Source:MGI Symbol;Acc:MGI:2442827] | 44,09 | 0,0 |
| ENSMUSG00000022003 | 46,09938993 | 28,8 | 0,0 | 8,268598299 | 0,032996306 | Slc25a30 | solute carrier family 25, member 30 [Source:MGI Symbol;Acc:MGI:1914804] | 91,99 | 0,0 |
| ENSMUSG00000031774 | 339,3745621 | 494,9 | 145,9 | 8,269617676 | 0,00254211 | Fam192a | family with sequence similarity 192, member A [Source:MGI Symbol;Acc:MGI:1919637] | 99,94 | 31,0 |
| ENSMUSG00000087002 | 16,05947822 | 28,8 | 0,0 | 8,271917805 | 0,026366395 | Gm16277 | predicted gene 16277 [Source:MGI Symbol;Acc:MGI:3826543] | 62,84 | 0,0 |
| ENSMUSG00000030272 | 314,9858309 | 487,6 | 73,9 | 8,272154123 | 0,001176273 | Camk1 | calcium/calmodulin-dependent protein kinase I [Source:MGI Symbol;Acc:MGI:1098535] | 99,87 | 29,7 |
| ENSMUSG00000001120 | 234,2169805 | 337,8 | 1,2 | 8,272930011 | 4,85698E-05 | Pcbp3 | poly(rC) binding protein 3 [Source:MGI Symbol;Acc:MGI:1890470] | 90,60 | 0,0 |
| ENSMUSG00000031927 | 24,95180264 | 29,0 | 0,0 | 8,274181065 | 0,033760829 | 1700012B09Rik | RIKEN cDNA 1700012B09 gene [Source:MGI Symbol;Acc:MGI:1916575] | 95,88 | 0,0 |
| ENSMUSG00000024899 | 750,1755373 | 1611,3 | 20,2 | 8,277173954 | 2,15802E-05 | Papss2 | 3'-phosphoadenosine 5'-phosphosulfate synthase 2 [Source:MGI Symbol;Acc:MGI:1330223] | 99,55 | 5,7 |
| ENSMUSG00000005886 | 96,98861978 | 196,4 | 0,9 | 8,279355329 | 0,001987113 | Ncoa2 | nuclear receptor coactivator 2 [Source:MGI Symbol;Acc:MGI:1276533] | 90,16 | 0,3 |
| ENSMUSG00000079495 | 14,83541048 | 29,5 | 0,0 | 8,280019691 | 0,023510692 | Gm11128 | predicted gene 11128 [Source:MGI Symbol;Acc:MGI:3779382] | 21,26 | 0,0 |
| ENSMUSG00000000296 | 263,7840235 | 406,2 | 48,1 | 8,280097408 | 0,001393062 | Tpd52l1 | tumor protein D52-like 1 [Source:MGI Symbol;Acc:MGI:1298386] | 99,45 | 26,4 |
| ENSMUSG00000033902 | 23,4285293 | 29,2 | 0,0 | 8,282403896 | 0,025907276 | Mapkbp1 | mitogen-activated protein kinase binding protein 1 [Source:MGI Symbol;Acc:MGI:1347004] | 51,83 | 0,0 |
| ENSMUSG00000031829 | 222,5449043 | 318,6 | 2,4 | 8,28273965 | 0,002875342 | Clpb | ClpB caseinolytic peptidase B [Source:MGI Symbol;Acc:MGI:1100517] | 93,58 | 0,7 |
| ENSMUSG00000026806 | 19,68172604 | 31,0 | 0,0 | 8,286374646 | 0,029043621 | Ddx31 | DEAD/H (Asp-Glu-Ala-Asp/His) box polypeptide 31 [Source:MGI Symbol;Acc:MGI:2682639] | 76,58 | 0,0 |
| ENSMUSG00000034152 | 342,9016649 | 516,6 | 9,2 | 8,287650719 | 0,000166554 | Exoc3 | exocyst complex component 3 [Source:MGI Symbol;Acc:MGI:2443972] | 99,95 | 1,9 |
| ENSMUSG00000073987 | 432,7127801 | 666,1 | 24,8 | 8,289129699 | 8,13172E-05 | Ggh | gamma-glutamyl hydrolase [Source:MGI Symbol;Acc:MGI:1329035] | 94,01 | 13,2 |
| ENSMUSG00000038370 | 466,3617072 | 581,5 | 154,7 | 8,298082965 | 0,002876863 | Pcp4l1 | Purkinje cell protein 4-like 1 [Source:MGI Symbol;Acc:MGI:1913675] | 99,34 | 28,5 |
| ENSMUSG00000090002 | 16,91540801 | 29,5 | 0,0 | 8,302509292 | 0,02602708 | Gm16006 | predicted gene 16006 [Source:MGI Symbol;Acc:MGI:3801853] | 97,82 | 0,0 |
| ENSMUSG00000107999 | 21,76708385 | 29,9 | 0,0 | 8,30308923 | 0,027341541 | Gm44123 | predicted gene, 44123 [Source:MGI Symbol;Acc:MGI:5690515] | 99,65 | 0,0 |
| ENSMUSG00000027661 | 37,80270031 | 35,2 | 0,0 | 8,303253448 | 0,028613304 | Slc2a10 | solute carrier family 2 (facilitated glucose transporter), member 10 [Source:MGI Symbol;Acc:MGI:2156687 | 74,92 | 0,0 |
| ENSMUSG00000038141 | 19,91816588 | 29,9 | 0,0 | 8,304140633 | 0,026432771 | Tmem181a | transmembrane protein 181A [Source:MGI Symbol;Acc:MGI:1924356] | 45,86 | 0,0 |
| ENSMUSG00000069814 | 15,32703885 | 30,0 | 0,0 | 8,304784424 | 0,023659521 | E130309D14Rik | RIKEN cDNA E130309D14 gene [Source:MGI Symbol;Acc:MGI:3588240] | 70,47 | 0,0 |
| ENSMUSG00000048120 | 170,5079066 | 202,6 | 0,9 | 8,305072859 | 0,001493938 | Entpd1 | ectonucleoside triphosphate diphosphohydrolase 1 [Source:MGI Symbol;Acc:MGI:102805] | 95,83 | 0,6 |
| ENSMUSG00000090175 | 16,64207805 | 34,3 | 0,0 | 8,306800728 | 0,020381708 | Ugt1a9 | UDP glucuronosyltransferase 1 family, polypeptide A9 [Source:MGI Symbol;Acc:MGI:3576092] | 56,15 | 0,0 |
| ENSMUSG00000090171 | 16,64207805 | 34,3 | 0,0 | 8,306800728 | 0,020381708 | Ugt1a2 | UDP glucuronosyltransferase 1 family, polypeptide A2 [Source:MGI Symbol;Acc:MGI:3576049] | 55,62 | 0,0 |
| ENSMUSG00000045316 | 257,3600934 | 472,9 | 24,0 | 8,307820992 | 0,000956632 | Fahd1 | fumarylacetoacetate hydrolase domain containing 1 [Source:MGI Symbol;Acc:MGI:1915886] | 100,00 | 21,6 |
| ENSMUSG00000089943 | 16,51308538 | 34,3 | 0,0 | 8,308794219 | 0,019072678 | Ugt1a5 | UDP glucuronosyltransferase 1 family, polypeptide A5 [Source:MGI Symbol;Acc:MGI:3032634] | 55,82 | 0,0 |
| ENSMUSG00000004473 | 20,2127848 | 31,5 | 0,0 | 8,31278635 | 0,029217761 | Clec11a | C-type lectin domain family 11, member a [Source:MGI Symbol;Acc:MGI:1298219] | 40,22 | 0,0 |
| ENSMUSG00000027287 | 296,8119294 | 418,6 | 17,6 | 8,315604024 | 0,002734427 | Snap23 | synaptosomal-associated protein 23 [Source:MGI Symbol;Acc:MGI:109356] | 91,98 | 3,9 |
| ENSMUSG00000089960 | 16,38409271 | 34,3 | 0,0 | 8,316936908 | 0,01949513 | Ugt1a1 | UDP glucuronosyltransferase 1 family, polypeptide A1 [Source:MGI Symbol;Acc:MGI:98898] | 55,29 | 0,0 |
| ENSMUSG00000089675 | 16,38409271 | 34,3 | 0,0 | 8,316936908 | 0,01949513 | Ugt1a8 | UDP glucuronosyltransferase 1 family, polypeptide A8 [Source:MGI Symbol;Acc:MGI:3576090] | 55,26 | 0,0 |
| ENSMUSG00000090165 | 16,38409271 | 34,3 | 0,0 | 8,316936908 | 0,01949513 | Ugt1a10 | UDP glycosyltransferase 1 family, polypeptide A10 [Source:MGI Symbol;Acc:MGI:3580642] | 54,29 | 0,0 |
| ENSMUSG00000090145 | 16,42152977 | 34,3 | 0,0 | 8,316961508 | 0,01949513 | Ugt1a6b | UDP glucuronosyltransferase 1 family, polypeptide A6B [Source:MGI Symbol;Acc:MGI:3580629] | 51,67 | 0,0 |
| ENSMUSG00000041912 | 340,8033947 | 446,5 | 185,6 | 8,318879818 | 0,000928541 | Tdrkh | tudor and KH domain containing protein [Source:MGI Symbol;Acc:MGI:1919884] | 90,49 | 6,5 |
| ENSMUSG00000023882 | 12,77111277 | 30,7 | 0,0 | 8,31975603 | 0,02031835 | Zfp54 | zinc finger protein 54 [Source:MGI Symbol;Acc:MGI:99201] | 75,06 | 0,0 |
| ENSMUSG00000051147 | 15,13181725 | 30,9 | 0,0 | 8,320209822 | 0,024238609 | Nat2 | N-acetyltransferase 2 (arylamine N-acetyltransferase) [Source:MGI Symbol;Acc:MGI:109201] | 88,45 | 0,0 |
| ENSMUSG00000035509 | 13,73067248 | 30,2 | 0,0 | 8,322680394 | 0,020000617 | Fbxl21 | F-box and leucine-rich repeat protein 21 [Source:MGI Symbol;Acc:MGI:2442921] | 88,17 | 0,0 |
| ENSMUSG00000085912 | 14,85413296 | 33,3 | 0,0 | 8,32415668 | 0,018022427 | Trp53cor1 | tumor protein p53 pathway corepressor 1 [Source:MGI Symbol;Acc:MGI:3801771] | #DIV/0! | #DIV/0! |
| ENSMUSG00000015839 | 458,830767 | 776,2 | 131,7 | 8,326764594 | 1,15903E-05 | Nfe2l2 | nuclear factor, erythroid derived 2, like 2 [Source:MGI Symbol;Acc:MGI:108420] | 98,59 | 25,2 |
| ENSMUSG00000074345 | 13,82603265 | 29,8 | 0,0 | 8,327809571 | 0,019938425 | Tnfaip8l3 | tumor necrosis factor, alpha-induced protein 8-like 3 [Source:MGI Symbol;Acc:MGI:2685363] | 90,68 | 0,0 |
| ENSMUSG00000018920 | 28,37980555 | 36,1 | 0,0 | 8,330689656 | 0,035663716 | Cxcl16 | chemokine (C-X-C motif) ligand 16 [Source:MGI Symbol;Acc:MGI:1932682] | 66,84 | 0,0 |
| ENSMUSG00000003546 | 127,3927938 | 166,0 | 0,6 | 8,333144355 | 0,005720701 | Klc4 | kinesin light chain 4 [Source:MGI Symbol;Acc:MGI:1922014] | 100,00 | 0,7 |
| ENSMUSG00000034765 | 38,29535277 | 31,1 | 0,0 | 8,333449177 | 0,0316686 | Dusp5 | dual specificity phosphatase 5 [Source:MGI Symbol;Acc:MGI:2685183] | 83,29 | 0,0 |
| ENSMUSG00000057596 | 14,153418 | 30,2 | 0,0 | 8,335195243 | 0,019706738 | Trim30d | tripartite motif-containing 30D [Source:MGI Symbol;Acc:MGI:3035181] | 69,29 | 0,0 |
| ENSMUSG00000003420 | 344,7461987 | 513,9 | 28,9 | 8,336766772 | 0,00123279 | Fcgrt | Fc receptor, IgG, alpha chain transporter [Source:MGI Symbol;Acc:MGI:103017] | 99,52 | 11,3 |
| ENSMUSG00000044707 | 19,76836398 | 34,0 | 0,0 | 8,339909223 | 0,02911605 | Ccnjl | cyclin J-like [Source:MGI Symbol;Acc:MGI:2685723] | 79,44 | 0,0 |
| ENSMUSG00000103906 | 26,8600104 | 32,5 | 0,0 | 8,339923569 | 0,026592475 | Tigd5 | tigger transposable element derived 5 [Source:MGI Symbol;Acc:MGI:2145902] | 56,73 | 0,0 |
| ENSMUSG00000097876 | 16,97599906 | 31,0 | 0,0 | 8,340581002 | 0,019104816 | Gm16892 | predicted gene, 16892 [Source:MGI Symbol;Acc:MGI:4439816] | #DIV/0! | #DIV/0! |
| ENSMUSG00000010660 | 142,6522895 | 187,0 | 1,0 | 8,343868167 | 0,005445905 | Plcd1 | phospholipase C, delta 1 [Source:MGI Symbol;Acc:MGI:97614] | 99,22 | 0,6 |
| ENSMUSG00000096215 | 33,70325563 | 45,8 | 0,0 | 8,344480206 | 0,022551392 | Smim22 | small integral membrane protein 22 [Source:MGI Symbol;Acc:MGI:3643379] | 97,93 | 0,0 |
| ENSMUSG00000042594 | 81,0293534 | 31,4 | 0,0 | 8,345424374 | 0,020595626 | Sh2b3 | SH2B adaptor protein 3 [Source:MGI Symbol;Acc:MGI:893598] | 55,81 | 0,0 |
| ENSMUSG00000022967 | 126,6120315 | 238,5 | 0,9 | 8,346957036 | 0,000207757 | Ifnar1 | interferon (alpha and beta) receptor 1 [Source:MGI Symbol;Acc:MGI:107658] | 99,59 | 0,3 |
| ENSMUSG00000102504 | 18,00740979 | 31,1 | 0,0 | 8,347886227 | 0,024415033 | Gm21955 | predicted gene, 21955 [Source:MGI Symbol;Acc:MGI:5439424] | 21,12 | 0,0 |
| ENSMUSG00000086648 | 15,42232555 | 31,8 | 0,0 | 8,352836444 | 0,019172773 | Gm15511 | predicted gene 15511 [Source:MGI Symbol;Acc:MGI:3782959] | 14,78 | 0,0 |
| ENSMUSG00000027796 | 19,39548329 | 34,8 | 0,0 | 8,354368108 | 0,023510692 | Smad9 | SMAD family member 9 [Source:MGI Symbol;Acc:MGI:1859993] | 56,76 | 0,0 |
| ENSMUSG00000086468 | 14,85205632 | 30,7 | 0,0 | 8,354661591 | 0,023018813 | Etaa1os | Ewing tumor-associated antigen 1, opposite strand [Source:MGI Symbol;Acc:MGI:1920221] | 92,21 | 0,0 |
| ENSMUSG00000021613 | 310,6159069 | 345,6 | 372,2 | 8,358191541 | 0,005100924 | Hapln1 | hyaluronan and proteoglycan link protein 1 [Source:MGI Symbol;Acc:MGI:1337006] | 93,80 | 27,0 |
| ENSMUSG00000049119 | 106,6278509 | 187,7 | 0,6 | 8,362039231 | 0,001938948 | Fam110b | family with sequence similarity 110, member B [Source:MGI Symbol;Acc:MGI:1916593] | 95,40 | 0,7 |
| ENSMUSG00000074404 | 13,46148886 | 35,3 | 0,0 | 8,363818018 | 0,019553887 | Gm1698 | predicted gene 1698 [Source:MGI Symbol;Acc:MGI:2686544] | #DIV/0! | #DIV/0! |
| ENSMUSG00000032500 | 44,05953823 | 33,0 | 0,0 | 8,364604564 | 0,031057321 | Dclk3 | doublecortin-like kinase 3 [Source:MGI Symbol;Acc:MGI:3039580] | 80,93 | 0,0 |
| ENSMUSG00000020910 | 108,3153853 | 170,9 | 0,6 | 8,365418192 | 0,004750039 | Arrdc2 | arrestin domain containing 2 [Source:MGI Symbol;Acc:MGI:1918057] | 96,22 | 1,1 |
| ENSMUSG00000048174 | 16,9096409 | 36,9 | 0,0 | 8,366560797 | 0,019131672 | Tmem81 | transmembrane protein 81 [Source:MGI Symbol;Acc:MGI:1921876] | 84,90 | 0,0 |
| ENSMUSG00000021118 | 20,93791882 | 35,5 | 0,0 | 8,368666031 | 0,018164583 | Plek2 | pleckstrin 2 [Source:MGI Symbol;Acc:MGI:1351466] | 92,61 | 0,0 |
| ENSMUSG00000055493 | 25,94745823 | 32,6 | 0,0 | 8,369016872 | 0,027876768 | Epm2a | epilepsy, progressive myoclonic epilepsy, type 2 gene alpha [Source:MGI Symbol;Acc:MGI:1341085] | 75,95 | 0,0 |
| ENSMUSG00000074811 | 16,1058158 | 31,9 | 0,0 | 8,371663826 | 0,017981001 | Hps6 | Hermansky-Pudlak syndrome 6 [Source:MGI Symbol;Acc:MGI:2181763] | 85,31 | 0,0 |
| ENSMUSG00000027765 | 20,27324593 | 33,1 | 0,0 | 8,374886871 | 0,027736526 | P2ry1 | purinergic receptor P2Y, G-protein coupled 1 [Source:MGI Symbol;Acc:MGI:105049] | 66,28 | 0,0 |
| ENSMUSG00000098050 | 27,18221619 | 35,0 | 0,0 | 8,382615486 | 0,02374044 | Gm5345 | predicted gene 5345 [Source:MGI Symbol;Acc:MGI:3644320] | #DIV/0! | #DIV/0! |
| ENSMUSG00000071036 | 10,35984287 | 34,6 | 0,0 | 8,384216756 | 0,020332104 | Gm10309 | predicted gene 10309 [Source:MGI Symbol;Acc:MGI:3641941] | 86,85 | 0,0 |
| ENSMUSG00000045761 | 17,10587508 | 36,3 | 0,0 | 8,388864708 | 0,017614538 | Fam179a | family with sequence similarity 179, member A [Source:MGI Symbol;Acc:MGI:2443498] | 73,32 | 0,0 |
| ENSMUSG00000042705 | 325,3099971 | 510,9 | 79,3 | 8,389229078 | 0,00337462 | Commd10 | COMM domain containing 10 [Source:MGI Symbol;Acc:MGI:1916706] | 99,96 | 30,8 |
| ENSMUSG00000022949 | 682,0979044 | 753,3 | 46,9 | 8,38954608 | 0,00086999 | Clic6 | chloride intracellular channel 6 [Source:MGI Symbol;Acc:MGI:2146607] | 95,85 | 3,7 |
| ENSMUSG00000035595 | 32,07507858 | 32,2 | 0,0 | 8,390254294 | 0,02368984 | 1600002K03Rik | RIKEN cDNA 1600002K03 gene [Source:MGI Symbol;Acc:MGI:1917020] | 95,67 | 0,0 |
| ENSMUSG00000085132 | 18,5858157 | 32,5 | 0,0 | 8,39119842 | 0,019277519 | Gm12265 | predicted gene 12265 [Source:MGI Symbol;Acc:MGI:3651913] | 19,59 | 0,0 |
| ENSMUSG00000022912 | 110,0525552 | 191,6 | 0,9 | 8,392580665 | 0,005254334 | Pros1 | protein S (alpha) [Source:MGI Symbol;Acc:MGI:1095733] | 99,17 | 0,6 |
| ENSMUSG00000052221 | 36,29459487 | 33,2 | 0,0 | 8,392823298 | 0,03539223 | Ppp1r36 | protein phosphatase 1, regulatory subunit 36 [Source:MGI Symbol;Acc:MGI:2684916] | 94,31 | 0,0 |
| ENSMUSG00000001247 | 26,49917929 | 42,2 | 0,0 | 8,393440017 | 0,01712795 | Lsr | lipolysis stimulated lipoprotein receptor [Source:MGI Symbol;Acc:MGI:1927471] | 85,48 | 0,0 |
| ENSMUSG00000003559 | 333,3217482 | 514,2 | 44,2 | 8,393528798 | 0,002857421 | As3mt | arsenic (+3 oxidation state) methyltransferase [Source:MGI Symbol;Acc:MGI:1929882] | 99,70 | 23,3 |
| ENSMUSG00000057137 | 24,16906274 | 32,3 | 0,0 | 8,394208492 | 0,020710182 | Tmem140 | transmembrane protein 140 [Source:MGI Symbol;Acc:MGI:1915737] | 71,63 | 0,0 |
| ENSMUSG00000029101 | 124,0126719 | 234,2 | 1,0 | 8,394376126 | 0,000552028 | Rgs12 | regulator of G-protein signaling 12 [Source:MGI Symbol;Acc:MGI:1918979] | 85,74 | 0,3 |
| ENSMUSG00000031478 | 13,58642106 | 34,9 | 0,0 | 8,395163715 | 0,020332104 | Nek3 | NIMA (never in mitosis gene a)-related expressed kinase 3 [Source:MGI Symbol;Acc:MGI:1344371] | 88,55 | 0,0 |
| ENSMUSG00000074283 | 18,31584172 | 31,1 | 0,0 | 8,395585027 | 0,01900494 | Zfp109 | zinc finger protein 109 [Source:MGI Symbol;Acc:MGI:1929099] | 80,96 | 0,0 |
| ENSMUSG00000102206 | 17,47553046 | 32,3 | 0,0 | 8,398053739 | 0,015072412 | Pcdha11 | protocadherin alpha 11 [Source:MGI Symbol;Acc:MGI:1298372] | 40,44 | 0,0 |
| ENSMUSG00000103442 | 18,11157641 | 32,4 | 0,0 | 8,398325574 | 0,016333387 | Pcdha1 | protocadherin alpha 1 [Source:MGI Symbol;Acc:MGI:2150982] | 40,93 | 0,0 |
| ENSMUSG00000086358 | 25,45219513 | 31,5 | 0,0 | 8,399475988 | 0,031059339 | Gm13270 | predicted gene 13270 [Source:MGI Symbol;Acc:MGI:3651704] | 94,15 | 0,0 |
| ENSMUSG00000102312 | 16,91511909 | 32,3 | 0,0 | 8,404168553 | 0,017067502 | Pcdha3 | protocadherin alpha 3 [Source:MGI Symbol;Acc:MGI:2447313] | 40,65 | 0,0 |
| ENSMUSG00000104148 | 16,99200291 | 32,3 | 0,0 | 8,40418756 | 0,017068678 | Pcdha2 | protocadherin alpha 2 [Source:MGI Symbol;Acc:MGI:2681880] | 40,43 | 0,0 |
| ENSMUSG00000035759 | 17,43679034 | 32,1 | 0,0 | 8,404695987 | 0,018209447 | Bbs10 | Bardet-Biedl syndrome 10 (human) [Source:MGI Symbol;Acc:MGI:1919019] | 80,05 | 0,0 |
| ENSMUSG00000028378 | 16,9551108 | 32,3 | 0,0 | 8,405403101 | 0,016333387 | Ptgr1 | prostaglandin reductase 1 [Source:MGI Symbol;Acc:MGI:1914353] | 59,45 | 0,0 |
| ENSMUSG00000059327 | 13,82983539 | 32,6 | 0,0 | 8,405689925 | 0,017570204 | Eda | ectodysplasin-A [Source:MGI Symbol;Acc:MGI:1195272] | 67,57 | 0,0 |
| ENSMUSG00000103800 | 17,00889842 | 32,4 | 0,0 | 8,406604152 | 0,017837685 | Pcdha8 | protocadherin alpha 8 [Source:MGI Symbol;Acc:MGI:2681879] | 41,07 | 0,0 |
| ENSMUSG00000029781 | 537,1075951 | 821,3 | 16,6 | 8,411235727 | 1,33767E-05 | Fkbp9 | FK506 binding protein 9 [Source:MGI Symbol;Acc:MGI:1350921] | 98,59 | 2,6 |
| ENSMUSG00000021743 | 306,4519745 | 489,2 | 54,9 | 8,412344102 | 0,000824492 | Fezf2 | Fez family zinc finger 2 [Source:MGI Symbol;Acc:MGI:1859823] | 92,59 | 13,2 |
| ENSMUSG00000086765 | 13,33720045 | 34,2 | 0,0 | 8,413940028 | 0,018901271 | Gm11827 | predicted gene 11827 [Source:MGI Symbol;Acc:MGI:3649801] | #DIV/0! | #DIV/0! |
| ENSMUSG00000049076 | 243,200996 | 504,6 | 71,2 | 8,420934652 | 0,001743956 | Acap2 | ArfGAP with coiled-coil, ankyrin repeat and PH domains 2 [Source:MGI Symbol;Acc:MGI:1925868] | 98,56 | 6,3 |
| ENSMUSG00000000305 | 60,26216826 | 32,4 | 0,0 | 8,422399934 | 0,03251979 | Cdh4 | cadherin 4 [Source:MGI Symbol;Acc:MGI:99218] | 52,25 | 0,0 |
| ENSMUSG00000097177 | 29,80859604 | 32,6 | 0,0 | 8,422447514 | 0,024944945 | 9330159M07Rik | RIKEN cDNA 9330159M07 gene [Source:MGI Symbol;Acc:MGI:2442519] | 35,42 | 0,0 |
| ENSMUSG00000042256 | 38,74451686 | 34,7 | 0,0 | 8,430447576 | 0,024028878 | Ptchd4 | patched domain containing 4 [Source:MGI Symbol;Acc:MGI:1920485] | 49,62 | 0,0 |
| ENSMUSG00000020601 | 303,0988371 | 533,7 | 129,7 | 8,431847373 | 0,000359804 | Trib2 | tribbles homolog 2 (Drosophila) [Source:MGI Symbol;Acc:MGI:2145021] | 99,95 | 15,7 |
| ENSMUSG00000032841 | 44,07455145 | 32,8 | 0,0 | 8,440133444 | 0,03394 | Prr5l | proline rich 5 like [Source:MGI Symbol;Acc:MGI:1919696] | 57,74 | 0,0 |
| ENSMUSG00000015401 | 41,86359066 | 50,0 | 0,0 | 8,440451661 | 0,019685287 | Tmem27 | transmembrane protein 27 [Source:MGI Symbol;Acc:MGI:1926234] | 75,35 | 0,0 |
| ENSMUSG00000027495 | 400,3709525 | 587,7 | 23,9 | 8,443161691 | 0,000114954 | Fam210b | family with sequence similarity 210, member B [Source:MGI Symbol;Acc:MGI:1914267] | 99,71 | 3,7 |
| ENSMUSG00000037318 | 16,56365875 | 32,6 | 0,0 | 8,444832074 | 0,018259241 | Traf3ip3 | TRAF3 interacting protein 3 [Source:MGI Symbol;Acc:MGI:2441706] | 16,49 | 0,0 |
| ENSMUSG00000075334 | 40,8272997 | 33,0 | 0,0 | 8,445859213 | 0,016304962 | Rprm | reprimo, TP53 dependent G2 arrest mediator candidate [Source:MGI Symbol;Acc:MGI:1915124] | 98,49 | 0,0 |
| ENSMUSG00000007440 | 17,04660565 | 33,1 | 0,0 | 8,452223413 | 0,016531443 | Pcdha11 | protocadherin alpha 11 [Source:MGI Symbol;Acc:MGI:1298372] | 43,23 | 0,0 |
| ENSMUSG00000060487 | 26,55410978 | 34,2 | 0,0 | 8,452657116 | 0,026466872 | Samd5 | sterile alpha motif domain containing 5 [Source:MGI Symbol;Acc:MGI:2444815] | 85,38 | 0,0 |
| ENSMUSG00000035239 | 21,27982296 | 35,7 | 0,0 | 8,452889609 | 0,018209447 | Neu3 | neuraminidase 3 [Source:MGI Symbol;Acc:MGI:1355305] | 81,36 | 0,0 |
| ENSMUSG00000021638 | 63,76865497 | 82,4 | 0,0 | 8,455650778 | 0,01281085 | Ocln | occludin [Source:MGI Symbol;Acc:MGI:106183] | 57,63 | 0,0 |
| ENSMUSG00000103310 | 18,55638807 | 33,8 | 0,0 | 8,455677692 | 0,014362471 | Pcdha12 | protocadherin alpha 12 [Source:MGI Symbol;Acc:MGI:1298370] | 44,41 | 0,0 |
| ENSMUSG00000031860 | 25,00243822 | 33,0 | 0,0 | 8,457816635 | 0,026469842 | Pbx4 | pre B cell leukemia homeobox 4 [Source:MGI Symbol;Acc:MGI:1931321] | 78,47 | 0,0 |
| ENSMUSG00000036641 | 20,84566169 | 36,6 | 0,0 | 8,458215049 | 0,018894755 | Ccdc148 | coiled-coil domain containing 148 [Source:MGI Symbol;Acc:MGI:3039583] | 67,83 | 0,0 |
| ENSMUSG00000103255 | 17,21492577 | 33,2 | 0,0 | 8,458773386 | 0,016467147 | Pcdhac1 | protocadherin alpha subfamily C, 1 [Source:MGI Symbol;Acc:MGI:1891442] | 42,83 | 0,0 |
| ENSMUSG00000022620 | 130,3894829 | 233,4 | 0,6 | 8,459445052 | 0,002194107 | Arsa | arylsulfatase A [Source:MGI Symbol;Acc:MGI:88077] | 89,01 | 0,7 |
| ENSMUSG00000066196 | 20,87422958 | 32,7 | 0,0 | 8,460332167 | 0,019869101 | Spag8 | sperm associated antigen 8 [Source:MGI Symbol;Acc:MGI:3056295] | 92,82 | 0,0 |
| ENSMUSG00000019810 | 344,1488484 | 474,2 | 98,6 | 8,46864907 | 0,002787134 | Fuca2 | fucosidase, alpha-L- 2, plasma [Source:MGI Symbol;Acc:MGI:1914098] | 67,97 | 10,8 |
| ENSMUSG00000106631 | 29,8040091 | 36,1 | 0,0 | 8,471044368 | 0,023899059 | Gm42669 | predicted gene 42669 [Source:MGI Symbol;Acc:MGI:5662806] | 22,15 | 0,0 |
| ENSMUSG00000104318 | 17,75873939 | 33,6 | 0,0 | 8,474817403 | 0,015462113 | Pcdha7 | protocadherin alpha 7 [Source:MGI Symbol;Acc:MGI:1298369] | 43,56 | 0,0 |
| ENSMUSG00000005131 | 22,95675727 | 33,1 | 0,0 | 8,475785998 | 0,023659521 | 4930550C14Rik | RIKEN cDNA 4930550C14 gene [Source:MGI Symbol;Acc:MGI:1922561] | 94,23 | 0,0 |
| ENSMUSG00000036904 | 14,56855596 | 35,1 | 0,0 | 8,480424202 | 0,017920081 | Fzd8 | frizzled class receptor 8 [Source:MGI Symbol;Acc:MGI:108460] | 74,78 | 0,0 |
| ENSMUSG00000097487 | 28,9904999 | 34,6 | 0,0 | 8,483170869 | 0,022313912 | Ptges3l | prostaglandin E synthase 3 (cytosolic)-like [Source:MGI Symbol;Acc:MGI:1916146] | 65,49 | 0,0 |
| ENSMUSG00000063354 | 39,50903907 | 51,6 | 0,0 | 8,485132456 | 0,03661633 | Slc39a4 | solute carrier family 39 (zinc transporter), member 4 [Source:MGI Symbol;Acc:MGI:1919277] | 79,72 | 0,0 |
| ENSMUSG00000025602 | 13,33626453 | 33,4 | 0,0 | 8,485510247 | 0,020710182 | Zfp202 | zinc finger protein 202 [Source:MGI Symbol;Acc:MGI:1933401] | 78,47 | 0,0 |
| ENSMUSG00000069633 | 21,78118032 | 34,6 | 0,0 | 8,486409743 | 0,019553887 | Pex11g | peroxisomal biogenesis factor 11 gamma [Source:MGI Symbol;Acc:MGI:1920905] | 71,90 | 0,0 |
| ENSMUSG00000053141 | 387,8599904 | 529,0 | 391,6 | 8,486607887 | 0,00510352 | Ptprt | protein tyrosine phosphatase, receptor type, T [Source:MGI Symbol;Acc:MGI:1321152] | 78,49 | 18,0 |
| ENSMUSG00000055866 | 55,64791066 | 34,3 | 0,0 | 8,489363521 | 0,028079855 | Per2 | period circadian clock 2 [Source:MGI Symbol;Acc:MGI:1195265] | 59,90 | 0,0 |
| ENSMUSG00000042423 | 23,66550382 | 34,1 | 0,0 | 8,491629378 | 0,014966423 | Fbrs | fibrosin [Source:MGI Symbol;Acc:MGI:104648] | 69,85 | 0,0 |
| ENSMUSG00000067825 | 15,57739057 | 35,5 | 0,0 | 8,493532565 | 0,016994889 | Pex26 | peroxisomal biogenesis factor 26 [Source:MGI Symbol;Acc:MGI:1921293] | 71,72 | 0,0 |
| ENSMUSG00000015452 | 13,39992908 | 35,5 | 0,0 | 8,497164443 | 0,017897827 | Ager | advanced glycosylation end product-specific receptor [Source:MGI Symbol;Acc:MGI:893592] | 91,97 | 0,0 |
| ENSMUSG00000044317 | 37,51642439 | 34,2 | 0,0 | 8,504776874 | 0,027533811 | Gpr4 | G protein-coupled receptor 4 [Source:MGI Symbol;Acc:MGI:2441992] | 73,37 | 0,0 |
| ENSMUSG00000030871 | 56,14956541 | 34,6 | 0,0 | 8,50519787 | 0,016698171 | Ears2 | glutamyl-tRNA synthetase 2 (mitochondrial)(putative) [Source:MGI Symbol;Acc:MGI:1914667] | 82,71 | 0,0 |
| ENSMUSG00000060149 | 45,97882729 | 34,4 | 0,0 | 8,505609087 | 0,022238384 | BC002059 | cDNA sequence BC002059 [Source:MGI Symbol;Acc:MGI:2384864] | #DIV/0! | #DIV/0! |
| ENSMUSG00000029434 | 282,7870603 | 396,2 | 211,5 | 8,507523542 | 0,00433647 | Vps33a | vacuolar protein sorting 33A (yeast) [Source:MGI Symbol;Acc:MGI:1924823] | 99,62 | 24,0 |
| ENSMUSG00000106981 | 28,25084011 | 34,4 | 0,0 | 8,508346094 | 0,027695266 | Smim20 | small integral membrane protein 20 [Source:MGI Symbol;Acc:MGI:1913528] | #DIV/0! | #DIV/0! |
| ENSMUSG00000096141 | 20,05553411 | 35,1 | 0,0 | 8,511503975 | 0,017312893 | Dnah7a | dynein, axonemal, heavy chain 7A [Source:MGI Symbol;Acc:MGI:2685838] | 1,71 | 0,0 |
| ENSMUSG00000050335 | 24,31371837 | 34,2 | 0,0 | 8,511682725 | 0,016148158 | Lgals3 | lectin, galactose binding, soluble 3 [Source:MGI Symbol;Acc:MGI:96778] | 57,19 | 0,0 |
| ENSMUSG00000075302 | 19,97018894 | 35,3 | 0,0 | 8,512640775 | 0,016508054 | Erich2 | glutamate rich 2 [Source:MGI Symbol;Acc:MGI:1913998] | 63,31 | 0,0 |
| ENSMUSG00000103092 | 17,22837529 | 34,4 | 0,0 | 8,512890787 | 0,015803049 | Pcdha5 | protocadherin alpha 5 [Source:MGI Symbol;Acc:MGI:1298371] | 47,32 | 0,0 |
| ENSMUSG00000047246 | 180,0647579 | 306,0 | 1,2 | 8,518199781 | 0,001916771 | Hist1h2be | histone cluster 1, H2be [Source:MGI Symbol;Acc:MGI:2448380] | 96,65 | 1,0 |
| ENSMUSG00000105504 | 20,93050179 | 36,3 | 0,0 | 8,519610827 | 0,015702473 | Gbp5 | guanylate binding protein 5 [Source:MGI Symbol;Acc:MGI:2429943] | 26,56 | 0,0 |
| ENSMUSG00000020101 | 32,59806669 | 39,2 | 0,0 | 8,520713194 | 0,019433244 | Vsir | V-set immunoregulatory receptor [Source:MGI Symbol;Acc:MGI:1921298] | 55,82 | 0,0 |
| ENSMUSG00000094441 | 18,47140229 | 35,3 | 0,0 | 8,521831175 | 0,023012649 | Zfp955a | zinc finger protein 955A [Source:MGI Symbol;Acc:MGI:4834570] | 72,08 | 0,0 |
| ENSMUSG00000032988 | 57,41933253 | 60,3 | 0,0 | 8,523411985 | 0,029170539 | Slc16a8 | lute carrier family 16 (monocarboxylic acid transporters), member 8 [Source:MGI Symbol;Acc:MGI:19295 | 85,73 | 0,0 |
| ENSMUSG00000045975 | 47,95954667 | 45,5 | 0,0 | 8,524914748 | 0,028064963 | C2cd2 | C2 calcium-dependent domain containing 2 [Source:MGI Symbol;Acc:MGI:1891883] | 57,20 | 0,0 |
| ENSMUSG00000026107 | 41,61161446 | 46,1 | 0,0 | 8,530276235 | 0,028160503 | Nabp1 | nucleic acid binding protein 1 [Source:MGI Symbol;Acc:MGI:1923258] | 65,10 | 0,0 |
| ENSMUSG00000022146 | 40,8306171 | 73,0 | 0,0 | 8,531422237 | 0,023510692 | Osmr | oncostatin M receptor [Source:MGI Symbol;Acc:MGI:1330819] | 61,96 | 0,0 |
| ENSMUSG00000056310 | 72,38522568 | 36,1 | 0,0 | 8,537469548 | 0,023617627 | Tyw1 | tRNA-yW synthesizing protein 1 homolog (S. cerevisiae) [Source:MGI Symbol;Acc:MGI:2141161] | 31,65 | 0,0 |
| ENSMUSG00000059482 | 365,9242273 | 621,7 | 76,3 | 8,538786434 | 0,002402222 | 2610301B20Rik | RIKEN cDNA 2610301B20 gene [Source:MGI Symbol;Acc:MGI:1914407] | 99,37 | 25,5 |
| ENSMUSG00000026676 | 169,637598 | 301,9 | 1,2 | 8,545980729 | 0,001728687 | Ccdc3 | coiled-coil domain containing 3 [Source:MGI Symbol;Acc:MGI:1921436] | 99,85 | 0,9 |
| ENSMUSG00000026388 | 22,77660626 | 35,5 | 0,0 | 8,553445861 | 0,017837685 | 3110009E18Rik | RIKEN cDNA 3110009E18 gene [Source:MGI Symbol;Acc:MGI:1920353] | 40,90 | 0,0 |
| ENSMUSG00000041696 | 12,1571146 | 36,7 | 0,0 | 8,555767577 | 0,01725089 | Rasl12 | RAS-like, family 12 [Source:MGI Symbol;Acc:MGI:1918034] | 55,00 | 0,0 |
| ENSMUSG00000021629 | 369,6186545 | 384,3 | 2,0 | 8,558084104 | 0,000262805 | Slc30a5 | solute carrier family 30 (zinc transporter), member 5 [Source:MGI Symbol;Acc:MGI:1916298] | 99,04 | 0,4 |
| ENSMUSG00000108776 | 28,71197875 | 36,0 | 0,0 | 8,559518553 | 0,020194747 | RP23-207N5.4 | | #DIV/0! | #DIV/0! |
| ENSMUSG00000063796 | 42,69942752 | 37,8 | 0,0 | 8,562797417 | 0,030653673 | Slc22a8 | solute carrier family 22 (organic anion transporter), member 8 [Source:MGI Symbol;Acc:MGI:1336187] | 64,70 | 0,0 |
| ENSMUSG00000087700 | 27,31748715 | 36,3 | 0,0 | 8,564107645 | 0,023641079 | Gm15283 | predicted gene 15283 [Source:MGI Symbol;Acc:MGI:3705161] | #DIV/0! | #DIV/0! |
| ENSMUSG00000036775 | 320,1384869 | 591,5 | 139,7 | 8,566002616 | 0,000525404 | Decr2 | 2-4-dienoyl-Coenzyme A reductase 2, peroxisomal [Source:MGI Symbol;Acc:MGI:1347059] | 99,59 | 28,6 |
| ENSMUSG00000039983 | 248,1056371 | 446,9 | 33,3 | 8,566051037 | 0,004525086 | Ccdc32 | coiled-coil domain containing 32 [Source:MGI Symbol;Acc:MGI:2685477] | 91,09 | 13,2 |
| ENSMUSG00000097743 | 16,27420185 | 35,9 | 0,0 | 8,568380491 | 0,016517421 | Gm16973 | predicted gene, 16973 [Source:MGI Symbol;Acc:MGI:4439897] | #DIV/0! | #DIV/0! |
| ENSMUSG00000039577 | 54,09676393 | 41,7 | 0,0 | 8,569824066 | 0,020940193 | Nphp4 | nephronophthisis 4 (juvenile) homolog (human) [Source:MGI Symbol;Acc:MGI:2384210] | 71,67 | 0,0 |
| ENSMUSG00000047250 | 31,21052665 | 36,3 | 0,0 | 8,572901779 | 0,02468674 | Ptgs1 | prostaglandin-endoperoxide synthase 1 [Source:MGI Symbol;Acc:MGI:97797] | 79,83 | 0,0 |
| ENSMUSG00000039178 | 257,5011591 | 420,9 | 2,0 | 8,574202642 | 4,27871E-05 | Tbc1d19 | TBC1 domain family, member 19 [Source:MGI Symbol;Acc:MGI:1914499] | 99,63 | 1,1 |

| | | | | | | | | | |
|---|---|---|---|---|---|---|---|---|---|
| ENSMUSG00000064179 | 26,91850198 | 36,2 | 0,0 | 8,574280389 | 0,03005362 | Tnnt1 | troponin T1, skeletal, slow [Source:MGI Symbol;Acc:MGI:1333868] | 74,91 | 0,0 |
| ENSMUSG00000024175 | 21,84777749 | 44,8 | 0,0 | 8,574453666 | 0,014137454 | Tekt4 | tektin 4 [Source:MGI Symbol;Acc:MGI:1919090] | 86,44 | 0,0 |
| ENSMUSG00000071291 | 26,38322163 | 35,6 | 0,0 | 8,576234385 | 0,015239583 | Zfp58 | zinc finger protein 58 [Source:MGI Symbol;Acc:MGI:99205] | 79,78 | 0,0 |
| ENSMUSG00000004500 | 41,29895866 | 36,2 | 0,0 | 8,57638575 | 0,022539815 | Zfp324 | zinc finger protein 324 [Source:MGI Symbol;Acc:MGI:2444641] | 70,50 | 0,0 |
| ENSMUSG00000040697 | 20,52621486 | 35,6 | 0,0 | 8,580858789 | 0,019280959 | Dnajc16 | DnaJ heat shock protein family (Hsp40) member C16 [Source:MGI Symbol;Acc:MGI:2442146] | 57,00 | 0,0 |
| ENSMUSG00000054720 | 96,69145858 | 252,1 | 0,6 | 8,586162195 | 0,000154767 | Lrrc8c | leucine rich repeat containing 8 family, member C [Source:MGI Symbol;Acc:MGI:2140839] | 93,86 | 0,3 |
| ENSMUSG00000066150 | 597,3780349 | 608,3 | 7,4 | 8,588241999 | 0,000133309 | Slc31a1 | solute carrier family 31, member 1 [Source:MGI Symbol;Acc:MGI:1333843] | 87,30 | 1,4 |
| ENSMUSG00000021591 | 305,501265 | 529,1 | 39,9 | 8,590388754 | 0,000652685 | Glrx | glutaredoxin [Source:MGI Symbol;Acc:MGI:2135625] | 73,80 | 18,4 |
| ENSMUSG00000104252 | 17,97010296 | 35,9 | 0,0 | 8,591702355 | 0,014832243 | Pcdha4 | protocadherin alpha 4 [Source:MGI Symbol;Acc:MGI:1298406] | 48,67 | 0,0 |
| ENSMUSG00000032540 | 492,8816675 | 643,5 | 116,8 | 8,594956832 | 0,000358526 | Abhd5 | abhydrolase domain containing 5 [Source:MGI Symbol;Acc:MGI:1914719] | 93,73 | 22,0 |
| ENSMUSG00000032802 | 509,3481607 | 746,7 | 260,8 | 8,597928345 | 0,000564638 | Srxn1 | sulfiredoxin 1 homolog (S. cerevisiae) [Source:MGI Symbol;Acc:MGI:104971] | 97,62 | 29,7 |
| ENSMUSG00000086529 | 23,93772969 | 36,6 | 0,0 | 8,599867377 | 0,022825695 | Acss2os | cyl-CoA synthetase short-chain family member 2, opposite strand [Source:MGI Symbol;Acc:MGI:1922195 | 24,27 | 0,0 |
| ENSMUSG00000052783 | 30,64009152 | 38,6 | 0,0 | 8,600522235 | 0,021424278 | Grk4 | G protein-coupled receptor kinase 4 [Source:MGI Symbol;Acc:MGI:95801] | 51,04 | 0,0 |
| ENSMUSG00000085218 | 31,17305021 | 36,8 | 0,0 | 8,601069139 | 0,01833841 | BB218582 | expressed sequence BB218582 [Source:MGI Symbol;Acc:MGI:2139448] | #DIV/0! | #DIV/0! |
| ENSMUSG00000045064 | 23,02129179 | 36,5 | 0,0 | 8,60149921 | 0,018717018 | Zc2hc1c | zinc finger, C2HC-type containing 1C [Source:MGI Symbol;Acc:MGI:1919600] | 70,00 | 0,0 |
| ENSMUSG00000070044 | 335,1652484 | 480,6 | 97,4 | 8,602167207 | 0,004736532 | Fam149a | family with sequence similarity 149, member A [Source:MGI Symbol;Acc:MGI:2387177] | 90,48 | 11,5 |
| ENSMUSG00000023935 | 21,70565663 | 38,3 | 0,0 | 8,60668261 | 0,021669249 | Spats1 | spermatogenesis associated, serine-rich 1 [Source:MGI Symbol;Acc:MGI:1918270] | 93,22 | 0,0 |
| ENSMUSG00000052769 | 19,66689088 | 36,6 | 0,0 | 8,609914056 | 0,019553887 | Gm9889 | predicted gene 9889 [Source:MGI Symbol;Acc:MGI:3642568] | #DIV/0! | #DIV/0! |
| ENSMUSG00000022591 | 23,97478679 | 36,1 | 0,0 | 8,615717914 | 0,01498487 | Gm9747 | predicted gene 9747 [Source:MGI Symbol;Acc:MGI:3710571] | 90,23 | 0,0 |
| ENSMUSG00000035845 | 37,86572094 | 37,7 | 0,0 | 8,616694085 | 0,017309393 | Alg12 | sparagine-linked glycosylation 12 (alpha-1,6-mannosyltransferase) [Source:MGI Symbol;Acc:MGI:238502 | 88,00 | 0,0 |
| ENSMUSG00000073755 | 27,87572856 | 37,8 | 0,0 | 8,618314715 | 0,020316904 | 5730409E04Rik | RIKEN cDNA 5730409E04Rik gene [Source:MGI Symbol;Acc:MGI:3609248] | 79,88 | 0,0 |
| ENSMUSG00000060147 | 475,972002 | 651,0 | 312,3 | 8,61978222 | 0,000270676 | Serpinb6a | serine (or cysteine) peptidase inhibitor, clade B, member 6a [Source:MGI Symbol;Acc:MGI:103123] | 88,61 | 18,6 |
| ENSMUSG00000071033 | 23,99225717 | 38,4 | 0,0 | 8,625899195 | 0,017750337 | Gm10308 | predicted gene 10308 [Source:MGI Symbol;Acc:MGI:3641943] | 98,56 | 0,0 |
| ENSMUSG00000097915 | 15,66848788 | 37,1 | 0,0 | 8,626753518 | 0,017614538 | A330009N23Rik | RIKEN cDNA A330009N23 gene [Source:MGI Symbol;Acc:MGI:2443491] | #DIV/0! | #DIV/0! |
| ENSMUSG00000073073 | 16,91369747 | 37,4 | 0,0 | 8,635607032 | 0,015462113 | Gm8098 | predicted gene 8098 [Source:MGI Symbol;Acc:MGI:3645749] | #DIV/0! | #DIV/0! |
| ENSMUSG00000103622 | 17,90346769 | 37,2 | 0,0 | 8,636979626 | 0,015326302 | Gm38391 | predicted gene, 38391 [Source:MGI Symbol;Acc:MGI:5613630] | 52,02 | 0,0 |
| ENSMUSG00000034459 | 26,39232671 | 42,4 | 0,0 | 8,641056628 | 0,014619677 | Ifit1 | interferon-induced protein with tetratricopeptide repeats 1 [Source:MGI Symbol;Acc:MGI:99450] | 74,67 | 0,0 |
| ENSMUSG00000024376 | 25,15230771 | 37,7 | 0,0 | 8,646178412 | 0,014620403 | Epb41l4a | erythrocyte membrane protein band 4.1 like 4a [Source:MGI Symbol;Acc:MGI:103007] | 75,41 | 0,0 |
| ENSMUSG00000048772 | 30,91094855 | 37,8 | 0,0 | 8,650815232 | 0,018594281 | Tmem53 | transmembrane protein 53 [Source:MGI Symbol;Acc:MGI:1916027] | 84,08 | 0,0 |
| ENSMUSG00000042116 | 202,0093248 | 380,3 | 2,0 | 8,659201998 | 0,001932439 | Vwa1 | von Willebrand factor A domain containing 1 [Source:MGI Symbol;Acc:MGI:2179729] | 65,78 | 0,5 |
| ENSMUSG00000081984 | 16,25927532 | 37,3 | 0,0 | 8,66088323 | 0,014422998 | Dnajb3 | DnaJ heat shock protein family (Hsp40) member B3 [Source:MGI Symbol;Acc:MGI:1306822] | 95,15 | 0,0 |
| ENSMUSG00000027408 | 15,75752287 | 37,6 | 0,0 | 8,663540773 | 0,015338279 | Cpxm1 | carboxypeptidase X 1 (M14 family) [Source:MGI Symbol;Acc:MGI:1934569] | 93,73 | 0,0 |
| ENSMUSG00000039911 | 270,6228466 | 508,7 | 59,1 | 8,664307178 | 0,001901811 | Spsb1 | splA/ryanodine receptor domain and SOCS box containing 1 [Source:MGI Symbol;Acc:MGI:1921896] | 74,01 | 14,7 |
| ENSMUSG00000026399 | 32,00363849 | 44,7 | 0,0 | 8,665820101 | 0,016737618 | Cd55 | CD55 molecule, decay accelerating factor for complement [Source:MGI Symbol;Acc:MGI:104850] | 67,25 | 0,0 |
| ENSMUSG00000097974 | 23,01083535 | 38,7 | 0,0 | 8,667601337 | 0,024052207 | Gm10605 | predicted gene 10605 [Source:MGI Symbol;Acc:MGI:3642596] | #DIV/0! | #DIV/0! |
| ENSMUSG00000021364 | 295,1918566 | 581,2 | 1,5 | 8,672692951 | 0,000104656 | Elovl2 | tion of very long chain fatty acids (FEN1/Elo2, SUR4/Elo3, yeast)-like 2 [Source:MGI Symbol;Acc:MGI:18 | 96,73 | 0,7 |
| ENSMUSG00000051037 | 16,63632493 | 37,6 | 0,0 | 8,674812524 | 0,015075944 | Zfp455 | zinc finger protein 455 [Source:MGI Symbol;Acc:MGI:3040708] | 85,90 | 0,0 |
| ENSMUSG00000041945 | 25,49429523 | 47,3 | 0,0 | 8,675233706 | 0,014287526 | Mfsd9 | major facilitator superfamily domain containing 9 [Source:MGI Symbol;Acc:MGI:2443548] | 83,89 | 0,0 |
| ENSMUSG00000038550 | 35,09027766 | 38,3 | 0,0 | 8,675917528 | 0,029157111 | Ciart | circadian associated repressor of transcription [Source:MGI Symbol;Acc:MGI:2684975] | 71,52 | 0,0 |
| ENSMUSG00000086382 | 23,77103908 | 39,6 | 0,0 | 8,683032791 | 0,022309574 | Chrna1os | ergic receptor, nicotinic, alpha polypeptide 1 (muscle), opposite strand [Source:MGI Symbol;Acc:MGI:365 | 93,68 | 0,0 |
| ENSMUSG00000057842 | 14,50160275 | 38,9 | 0,0 | 8,684049977 | 0,014499693 | Zfp595 | zinc finger protein 595 [Source:MGI Symbol;Acc:MGI:3040707] | 56,94 | 0,0 |
| ENSMUSG00000027257 | 433,8289756 | 706,9 | 84,7 | 8,689278099 | 0,001128767 | Pacsin3 | protein kinase C and casein kinase substrate in neurons 3 [Source:MGI Symbol;Acc:MGI:1891410] | 73,91 | 21,6 |
| ENSMUSG00000039982 | 141,7384033 | 273,8 | 0,6 | 8,690312285 | 0,000126938 | Dtx4 | deltex 4, E3 ubiquitin ligase [Source:MGI Symbol;Acc:MGI:2672905] | 95,73 | 0,1 |
| ENSMUSG00000020432 | 341,3501992 | 546,1 | 22,4 | 8,69109972 | 0,002117304 | Tcn2 | transcobalamin 2 [Source:MGI Symbol;Acc:MGI:98534] | 93,88 | 10,9 |
| ENSMUSG00000028182 | 12,49355465 | 38,6 | 0,0 | 8,691198158 | 0,016615332 | Lrriq3 | leucine-rich repeats and IQ motif containing 3 [Source:MGI Symbol;Acc:MGI:1921685] | 61,51 | 0,0 |
| ENSMUSG00000022120 | 30,30960689 | 38,8 | 0,0 | 8,692071668 | 0,022461671 | Rnf219 | ring finger protein 219 [Source:MGI Symbol;Acc:MGI:1919736] | 79,55 | 0,0 |
| ENSMUSG00000054057 | 20,36989392 | 38,4 | 0,0 | 8,692104611 | 0,017872104 | A930004D18Rik | RIKEN cDNA A930004D18 gene [Source:MGI Symbol;Acc:MGI:1925190] | 79,32 | 0,0 |
| ENSMUSG00000008305 | 490,37892 | 868,3 | 84,1 | 8,697570604 | 5,63806E-05 | Tle1 | transducin-like enhancer of split 1, homolog of Drosophila E(spl) [Source:MGI Symbol;Acc:MGI:104636] | 94,64 | 12,1 |
| ENSMUSG00000045875 | 19,38196705 | 42,2 | 0,0 | 8,697680027 | 0,014075851 | Adra1a | adrenergic receptor, alpha 1a [Source:MGI Symbol;Acc:MGI:104773] | 51,41 | 0,0 |
| ENSMUSG00000037989 | 160,5035102 | 302,0 | 1,0 | 8,697711012 | 0,000850539 | Wnk2 | WNK lysine deficient protein kinase 2 [Source:MGI Symbol;Acc:MGI:1922857] | 87,79 | 0,1 |
| ENSMUSG00000085181 | 32,67709738 | 40,3 | 0,0 | 8,699495058 | 0,01833008 | Gm12709 | predicted gene 12709 [Source:MGI Symbol;Acc:MGI:3651946] | 22,35 | 0,0 |
| ENSMUSG00000041075 | 29,43417305 | 40,4 | 0,0 | 8,708706966 | 0,012330984 | Fzd7 | frizzled class receptor 7 [Source:MGI Symbol;Acc:MGI:108570] | 72,65 | 0,0 |
| ENSMUSG00000025280 | 37,63206882 | 40,8 | 0,0 | 8,711389244 | 0,019187804 | Polr3a | polymerase (RNA) III (DNA directed) polypeptide A [Source:MGI Symbol;Acc:MGI:2681836] | 76,57 | 0,0 |
| ENSMUSG00000035235 | 20,81809066 | 39,5 | 0,0 | 8,713123205 | 0,019063996 | Trim13 | tripartite motif-containing 13 [Source:MGI Symbol;Acc:MGI:1913847] | 87,48 | 0,0 |
| ENSMUSG00000053783 | 26,71656564 | 41,3 | 0,0 | 8,713236343 | 0,020250365 | 1700016K19Rik | RIKEN cDNA 1700016K19 gene [Source:MGI Symbol;Acc:MGI:1921480] | 94,68 | 0,0 |
| ENSMUSG00000009566 | 51,42873598 | 39,8 | 0,0 | 8,71359153 | 0,012527681 | Fpgs | folylpolyglutamyl synthetase [Source:MGI Symbol;Acc:MGI:95576] | 81,35 | 0,0 |
| ENSMUSG00000047907 | 34,09836292 | 40,8 | 0,0 | 8,71716482 | 0,017038985 | Tshz2 | teashirt zinc finger family member 2 [Source:MGI Symbol;Acc:MGI:2153084] | 50,95 | 0,0 |
| ENSMUSG00000096140 | 15,43106587 | 39,1 | 0,0 | 8,720652104 | 0,014747957 | Ankrd66 | ankyrin repeat domain 66 [Source:MGI Symbol;Acc:MGI:1925106] | 87,56 | 0,0 |
| ENSMUSG00000037410 | 33,62083959 | 40,1 | 0,0 | 8,723099047 | 0,014620403 | Tbc1d2b | TBC1 domain family, member 2B [Source:MGI Symbol;Acc:MGI:1914266] | 49,85 | 0,0 |
| ENSMUSG00000030107 | 30,23193606 | 40,0 | 0,0 | 8,727603808 | 0,027719879 | Usp18 | ubiquitin specific peptidase 18 [Source:MGI Symbol;Acc:MGI:1344364] | 82,02 | 0,0 |
| ENSMUSG00000086075 | 19,45473824 | 39,5 | 0,0 | 8,728060861 | 0,014211001 | Gm15728 | predicted gene 15728 [Source:MGI Symbol;Acc:MGI:3783171] | #DIV/0! | #DIV/0! |
| ENSMUSG00000063108 | 79,72192496 | 45,4 | 0,0 | 8,73284668 | 0,021882131 | Zfp26 | zinc finger protein 26 [Source:MGI Symbol;Acc:MGI:99173] | 60,14 | 0,0 |
| ENSMUSG00000108112 | 34,69073639 | 44,4 | 0,0 | 8,733585422 | 0,020331835 | RP24-491O6.1 | | #DIV/0! | #DIV/0! |
| ENSMUSG00000051486 | 15,4242728 | 40,8 | 0,0 | 8,73725316 | 0,026414215 | Pcdhb11 | protocadherin beta 11 [Source:MGI Symbol;Acc:MGI:2136746] | 77,23 | 0,0 |
| ENSMUSG00000075254 | 32,44891376 | 49,0 | 0,0 | 8,747878036 | 0,01871431 | Heg1 | heart development protein with EGF-like domains 1 [Source:MGI Symbol;Acc:MGI:1924696] | 46,17 | 0,0 |
| ENSMUSG00000054545 | 21,23585479 | 43,9 | 0,0 | 8,75042418 | 0,013817474 | Ugt1a6a | UDP glucuronosyltransferase 1 family, polypeptide A6A [Source:MGI Symbol;Acc:MGI:2137698] | 73,77 | 0,0 |
| ENSMUSG00000090124 | 21,34302047 | 43,9 | 0,0 | 8,75045943 | 0,013807459 | Ugt1a7c | UDP glucuronosyltransferase 1 family, polypeptide A7C [Source:MGI Symbol;Acc:MGI:3032636] | 41,85 | 0,0 |
| ENSMUSG00000074113 | 34,1085496 | 42,2 | 0,0 | 8,753007693 | 0,026773211 | Gm10629 | predicted gene 10629 [Source:MGI Symbol;Acc:MGI:3642307] | #DIV/0! | #DIV/0! |
| ENSMUSG00000027434 | 26,95898536 | 41,2 | 0,0 | 8,756934365 | 0,01949513 | Nkx2-2 | NK2 homeobox 2 [Source:MGI Symbol;Acc:MGI:97347] | 72,15 | 0,0 |
| ENSMUSG00000056144 | 29,9587328 | 42,9 | 0,0 | 8,761039048 | 0,016754086 | Trim34a | tripartite motif-containing 34A [Source:MGI Symbol;Acc:MGI:2137359] | 45,91 | 0,0 |
| ENSMUSG00000042515 | 27,01655127 | 44,0 | 0,0 | 8,762946816 | 0,016261287 | Mum1l1 | melanoma associated antigen (mutated) 1-like 1 [Source:MGI Symbol;Acc:MGI:2445062] | 67,87 | 0,0 |
| ENSMUSG00000097565 | 28,04601562 | 41,4 | 0,0 | 8,765484288 | 0,01405216 | Gm26965 | predicted gene, 26965 [Source:MGI Symbol;Acc:MGI:5504080] | #DIV/0! | #DIV/0! |
| ENSMUSG00000025453 | 204,06696 | 302,6 | 0,9 | 8,767852711 | 0,000258944 | Nnt | nicotinamide nucleotide transhydrogenase [Source:MGI Symbol;Acc:MGI:109279] | 77,00 | 0,0 |
| ENSMUSG00000045039 | 55,19145565 | 43,8 | 0,0 | 8,769660638 | 0,015392606 | Megf8 | multiple EGF-like-domains 8 [Source:MGI Symbol;Acc:MGI:2446294] | 43,44 | 0,0 |
| ENSMUSG00000097615 | 22,40485589 | 40,4 | 0,0 | 8,770689398 | 0,01345358 | Gm2061 | predicted gene 2061 [Source:MGI Symbol;Acc:MGI:3780228] | #DIV/0! | #DIV/0! |
| ENSMUSG00000038422 | 30,74855373 | 40,7 | 0,0 | 8,773469115 | 0,014149888 | Hdhd3 | haloacid dehalogenase-like hydrolase domain containing 3 [Source:MGI Symbol;Acc:MGI:1919998] | 96,51 | 0,0 |
| ENSMUSG00000024960 | 173,1217126 | 311,1 | 1,8 | 8,77451147 | 0,003956573 | Plcb3 | phospholipase C, beta 3 [Source:MGI Symbol;Acc:MGI:104778] | 100,00 | 0,3 |
| ENSMUSG00000096606 | 28,01996336 | 45,1 | 0,0 | 8,774641328 | 0,023012649 | Tpbgl | trophoblast glycoprotein-like [Source:MGI Symbol;Acc:MGI:3646425] | 79,68 | 0,0 |
| ENSMUSG00000038132 | 37,31184371 | 41,1 | 0,0 | 8,779738709 | 0,019172773 | Rbm24 | RNA binding motif protein 24 [Source:MGI Symbol;Acc:MGI:3610364] | 79,46 | 0,0 |
| ENSMUSG00000048967 | 22,98959202 | 42,0 | 0,0 | 8,780794676 | 0,018653255 | Yjefn3 | YjeF N-terminal domain containing 3 [Source:MGI Symbol;Acc:MGI:2681845] | 97,93 | 0,0 |
| ENSMUSG00000022236 | 29,02015402 | 42,8 | 0,0 | 8,78082255 | 0,019292726 | Ropn1l | ropporin 1-like [Source:MGI Symbol;Acc:MGI:2182357] | 82,09 | 0,0 |
| ENSMUSG00000027985 | 22,87227518 | 41,7 | 0,0 | 8,781920453 | 0,013341264 | Lef1 | lymphoid enhancer binding factor 1 [Source:MGI Symbol;Acc:MGI:96770] | 80,82 | 0,0 |
| ENSMUSG00000032816 | 463,6894341 | 793,4 | 22,4 | 8,783608846 | 0,000191491 | Igdcc4 | immunoglobulin superfamily, DCC subclass, member 4 [Source:MGI Symbol;Acc:MGI:1858497] | 98,45 | 4,3 |
| ENSMUSG00000024862 | 32,28836329 | 43,4 | 0,0 | 8,786820027 | 0,011353576 | Klc2 | kinesin light chain 2 [Source:MGI Symbol;Acc:MGI:107953] | 84,32 | 0,0 |
| ENSMUSG00000037656 | 371,847478 | 649,8 | 47,2 | 8,790219913 | 0,001199733 | Slc20a2 | solute carrier family 20, member 2 [Source:MGI Symbol;Acc:MGI:97851] | 98,83 | 12,5 |
| ENSMUSG00000108934 | 47,5581199 | 50,8 | 0,0 | 8,793384955 | 0,016803379 | RP23-124B2.7 | | 98,22 | 0,0 |
| ENSMUSG00000020474 | 36,50472454 | 42,9 | 0,0 | 8,798275933 | 0,018205458 | Polm | polymerase (DNA directed), mu [Source:MGI Symbol;Acc:MGI:1860191] | 90,30 | 0,0 |
| ENSMUSG00000085772 | 35,32070629 | 43,5 | 0,0 | 8,802160776 | 0,024133717 | D630024D03Rik | RIKEN cDNA D630024D03 gene [Source:MGI Symbol;Acc:MGI:3041224] | #DIV/0! | #DIV/0! |
| ENSMUSG00000085175 | 42,11931368 | 42,4 | 0,0 | 8,802263909 | 0,017410771 | Gm11423 | predicted gene 11423 [Source:MGI Symbol;Acc:MGI:3651335] | 97,51 | 0,0 |
| ENSMUSG00000085631 | 22,40017165 | 43,2 | 0,0 | 8,803548894 | 0,01696342 | 9630028H03Rik | RIKEN cDNA 9630028H03 gene [Source:MGI Symbol;Acc:MGI:2444526] | 4,89 | 0,0 |
| ENSMUSG00000033615 | 258,7265174 | 447,9 | 1,8 | 8,804698831 | 0,000731719 | Cplx1 | complexin 1 [Source:MGI Symbol;Acc:MGI:104727] | 99,39 | 2,2 |
| ENSMUSG00000036598 | 53,99896802 | 45,9 | 0,0 | 8,805496353 | 0,025878717 | Ccdc113 | coiled-coil domain containing 113 [Source:MGI Symbol;Acc:MGI:3606076] | 98,79 | 0,0 |
| ENSMUSG00000028885 | 21,82542024 | 46,4 | 0,0 | 8,806209521 | 0,012546899 | Smpdl3b | sphingomyelin phosphodiesterase, acid-like 3B [Source:MGI Symbol;Acc:MGI:1916022] | 84,35 | 0,0 |
| ENSMUSG00000020648 | 24,10173491 | 43,7 | 0,0 | 8,807126613 | 0,015550491 | Dus4l | dihydrouridine synthase 4-like (S. cerevisiae) [Source:MGI Symbol;Acc:MGI:1919166] | 94,61 | 0,0 |
| ENSMUSG00000028164 | 34,2830609 | 41,6 | 0,0 | 8,809871408 | 0,017748179 | Manba | mannosidase, beta A, lysosomal [Source:MGI Symbol;Acc:MGI:88175] | 85,19 | 0,0 |
| ENSMUSG00000090336 | 21,43077991 | 45,3 | 0,0 | 8,812880695 | 0,012541575 | 4932443I19Rik | RIKEN cDNA 4932443I19 gene [Source:MGI Symbol;Acc:MGI:2685952] | 92,95 | 0,0 |
| ENSMUSG00000097724 | 34,087018 | 42,6 | 0,0 | 8,817531145 | 0,024602524 | Gm26850 | predicted gene, 26850 [Source:MGI Symbol;Acc:MGI:5477344] | 35,50 | 0,0 |
| ENSMUSG00000019945 | 30,26940417 | 41,6 | 0,0 | 8,824359699 | 0,012462267 | 1700040L02Rik | RIKEN cDNA 1700040L02 gene [Source:MGI Symbol;Acc:MGI:1920537] | 95,43 | 0,0 |
| ENSMUSG00000074643 | 226,2953755 | 322,2 | 0,6 | 8,824764744 | 5,88621E-05 | Cpne1 | copine I [Source:MGI Symbol;Acc:MGI:2386621] | 94,68 | 0,6 |
| ENSMUSG00000092260 | 26,91176311 | 42,6 | 0,0 | 8,826157783 | 0,012054878 | Zfp963 | zinc finger protein 963 [Source:MGI Symbol;Acc:MGI:4867078] | 88,08 | 0,0 |
| ENSMUSG00000048965 | 17,07686196 | 42,3 | 0,0 | 8,826620177 | 0,01530672 | Mrgpre | MAS-related GPR, member E [Source:MGI Symbol;Acc:MGI:2441884] | 81,33 | 0,0 |
| ENSMUSG00000030125 | 23,75455901 | 41,7 | 0,0 | 8,826630539 | 0,012785209 | Lrrc23 | leucine rich repeat containing 23 [Source:MGI Symbol;Acc:MGI:1315192] | 97,03 | 0,0 |
| ENSMUSG00000033910 | 132,4204614 | 243,1 | 0,6 | 8,828848158 | 0,00354734 | Gucy1a3 | guanylate cyclase 1, soluble, alpha 3 [Source:MGI Symbol;Acc:MGI:1926562] | 93,14 | 0,5 |
| ENSMUSG00000040296 | 33,42458706 | 44,3 | 0,0 | 8,830707429 | 0,012318121 | Ddx58 | DEAD (Asp-Glu-Ala-Asp) box polypeptide 58 [Source:MGI Symbol;Acc:MGI:2442858] | 77,34 | 0,0 |
| ENSMUSG00000019894 | 30,26637911 | 44,4 | 0,0 | 8,83474294 | 0,014697605 | Slc6a15 | solute carrier family 6 (neurotransmitter transporter), member 15 [Source:MGI Symbol;Acc:MGI:2143484] | 83,47 | 0,0 |
| ENSMUSG00000019906 | 27,04944747 | 42,7 | 0,0 | 8,837175124 | 0,015495909 | Lin7a | lin-7 homolog A (C. elegans) [Source:MGI Symbol;Acc:MGI:2135609] | 62,24 | 0,0 |
| ENSMUSG00000029710 | 19,69405162 | 42,9 | 0,0 | 8,837677793 | 0,012689677 | Ephb4 | Eph receptor B4 [Source:MGI Symbol;Acc:MGI:104757] | 76,23 | 0,0 |
| ENSMUSG00000038286 | 411,9693714 | 700,5 | 35,1 | 8,838444465 | 0,001454212 | Bphl | ydrolase-like (serine hydrolase, breast epithelial mucin-associated antigen) [Source:MGI Symbol;Acc:MGI: | 100,00 | 26,5 |
| ENSMUSG00000052957 | 323,3520274 | 537,8 | 3,7 | 8,83932191 | 0,000291495 | Gas1 | growth arrest specific 1 [Source:MGI Symbol;Acc:MGI:95655] | 99,95 | 1,9 |
| ENSMUSG00000035486 | 46,57624822 | 43,9 | 0,0 | 8,840222262 | 0,02195965 | Plk5 | polo-like kinase 5 [Source:MGI Symbol;Acc:MGI:3026984] | 93,10 | 0,0 |
| ENSMUSG00000028347 | 180,694266 | 257,3 | 1,0 | 8,840754303 | 0,003497776 | Tmeff1 | nsmembrane protein with EGF-like and two follistatin-like domains 1 [Source:MGI Symbol;Acc:MGI:19268 | 88,49 | 1,0 |
| ENSMUSG00000042694 | 25,62723058 | 42,9 | 0,0 | 8,842381456 | 0,012060102 | Obfc1 | oligonucleotide/oligosaccharide-binding fold containing 1 [Source:MGI Symbol;Acc:MGI:1915581] | 68,58 | 0,0 |
| ENSMUSG00000090394 | 27,98921215 | 42,7 | 0,0 | 8,844434741 | 0,017528757 | 4930523C07Rik | RIKEN cDNA 4930523C07 gene [Source:MGI Symbol;Acc:MGI:1914897] | 100,00 | 0,0 |
| ENSMUSG00000046519 | 339,1168622 | 558,3 | 469,0 | 8,844541281 | 0,001463867 | Golph3l | golgi phosphoprotein 3-like [Source:MGI Symbol;Acc:MGI:1917129] | 99,86 | 32,6 |
| ENSMUSG00000057396 | 24,17358333 | 50,4 | 0,0 | 8,844623466 | 0,01226813 | Zfp759 | zinc finger protein 759 [Source:MGI Symbol;Acc:MGI:2446280] | 80,26 | 0,0 |
| ENSMUSG00000023156 | 689,0810211 | 943,2 | 354,0 | 8,845991967 | 0,000235523 | Rpp14 | ribonuclease P 14 subunit [Source:MGI Symbol;Acc:MGI:1914303] | 100,00 | 32,9 |
| ENSMUSG00000024939 | 271,6692509 | 381,7 | 1,2 | 8,848107139 | 0,000989597 | Fam89b | family with sequence similarity 89, member B [Source:MGI Symbol;Acc:MGI:106595] | 100,00 | 4,0 |
| ENSMUSG00000056313 | 16,66283333 | 52,7 | 0,0 | 8,848896579 | 0,013322569 | 1810011O10Rik | RIKEN cDNA 1810011O10 gene [Source:MGI Symbol;Acc:MGI:1916318] | 90,53 | 0,0 |
| ENSMUSG00000078546 | 20,35902185 | 43,9 | 0,0 | 8,849025126 | 0,012847287 | 2210404O09Rik | RIKEN cDNA 2210404O09 gene [Source:MGI Symbol;Acc:MGI:1917331] | 83,81 | 0,0 |
| ENSMUSG00000019772 | 30,126436 | 44,9 | 0,0 | 8,850828479 | 0,020593229 | Vip | vasoactive intestinal polypeptide [Source:MGI Symbol;Acc:MGI:98933] | 99,04 | 0,0 |
| ENSMUSG00000044461 | 29,92609867 | 45,9 | 0,0 | 8,850830178 | 0,012282057 | Shisa2 | shisa family member 2 [Source:MGI Symbol;Acc:MGI:2444716] | 78,36 | 0,0 |
| ENSMUSG00000005951 | 22,79621626 | 43,5 | 0,0 | 8,851719599 | 0,013044635 | Shpk | sedoheptulokinase [Source:MGI Symbol;Acc:MGI:1921887] | 90,47 | 0,0 |
| ENSMUSG00000087385 | 38,44765227 | 44,7 | 0,0 | 8,853241728 | 0,019513488 | AA415398 | expressed sequence AA415398 [Source:MGI Symbol;Acc:MGI:3035485] | 92,11 | 0,0 |
| ENSMUSG00000045954 | 31,98734613 | 45,8 | 0,0 | 8,854270746 | 0,01215584 | Sdpr | serum deprivation response [Source:MGI Symbol;Acc:MGI:99513] | 86,38 | 0,0 |
| ENSMUSG00000032440 | 54,18441413 | 43,6 | 0,0 | 8,854856069 | 0,019318178 | Tgfbr2 | transforming growth factor, beta receptor II [Source:MGI Symbol;Acc:MGI:98729] | 77,73 | 0,0 |
| ENSMUSG00000032489 | 23,78325569 | 43,9 | 0,0 | 8,85646067 | 0,012509594 | Kif9 | kinesin family member 9 [Source:MGI Symbol;Acc:MGI:1098237] | 69,39 | 0,0 |
| ENSMUSG00000099851 | 29,75237285 | 44,6 | 0,0 | 8,863065536 | 0,014137454 | Gm29542 | predicted gene 29542 [Source:MGI Symbol;Acc:MGI:5580248] | 96,40 | 0,0 |
| ENSMUSG00000017830 | 18,12857415 | 44,0 | 0,0 | 8,871360323 | 0,012702191 | Dhx58 | DEXH (Asp-Glu-X-His) box polypeptide 58 [Source:MGI Symbol;Acc:MGI:1931560] | 61,71 | 0,0 |
| ENSMUSG00000067931 | 16,89086996 | 45,3 | 0,0 | 8,872778278 | 0,014785672 | Zfp948 | zinc finger protein 948 [Source:MGI Symbol;Acc:MGI:3040683] | 89,99 | 0,0 |
| ENSMUSG00000096910 | 24,7915554 | 44,8 | 0,0 | 8,87503479 | 0,017208103 | Zfp955b | zinc finger protein 955B [Source:MGI Symbol;Acc:MGI:4834573] | 77,19 | 0,0 |
| ENSMUSG00000034639 | 22,665608 | 47,7 | 0,0 | 8,876520083 | 0,012408832 | Setmar | SET domain without mariner transposase fusion [Source:MGI Symbol;Acc:MGI:1921979] | 94,52 | 0,0 |
| ENSMUSG00000029049 | 63,69675969 | 44,6 | 0,0 | 8,877857885 | 0,024028878 | Morn1 | MORN repeat containing 1 [Source:MGI Symbol;Acc:MGI:1924116] | 95,20 | 0,0 |
| ENSMUSG00000021958 | 65,92626985 | 45,2 | 0,0 | 8,880534871 | 0,016491976 | Pinx1 | PIN2/TERF1 interacting, telomerase inhibitor 1 [Source:MGI Symbol;Acc:MGI:1919650] | 91,49 | 0,0 |
| ENSMUSG00000026213 | 38,7328061 | 45,3 | 0,0 | 8,882582722 | 0,016994889 | Stk11ip | serine/threonine kinase 11 interacting protein [Source:MGI Symbol;Acc:MGI:1918978] | 68,88 | 0,0 |
| ENSMUSG00000033355 | 43,03623725 | 46,4 | 0,0 | 8,885610793 | 0,017393403 | Rtp4 | receptor transporter protein 4 [Source:MGI Symbol;Acc:MGI:1915025] | 74,38 | 0,0 |
| ENSMUSG00000019832 | 27,20250347 | 50,7 | 0,0 | 8,887747377 | 0,011445243 | Rab32 | RAB32, member RAS oncogene family [Source:MGI Symbol;Acc:MGI:1915094] | 87,18 | 0,0 |
| ENSMUSG00000021327 | 144,2906472 | 238,6 | 0,6 | 8,888774464 | 0,00511715 | Zkscan3 | zinc finger with KRAB and SCAN domains 3 [Source:MGI Symbol;Acc:MGI:1919989] | 41,96 | 0,3 |
| ENSMUSG00000022537 | 438,6408429 | 560,2 | 342,5 | 8,893771657 | 0,007146733 | Tmem44 | transmembrane protein 44 [Source:MGI Symbol;Acc:MGI:1924489] | 99,90 | 28,8 |
| ENSMUSG00000021411 | 25,02529542 | 44,4 | 0,0 | 8,896084487 | 0,012098309 | Pxdc1 | PX domain containing 1 [Source:MGI Symbol;Acc:MGI:1914145] | 87,23 | 0,0 |
| ENSMUSG00000027896 | 45,09791223 | 45,1 | 0,0 | 8,897820993 | 0,015981384 | Slc16a4 | lute carrier family 16 (monocarboxylic acid transporters), member 4 [Source:MGI Symbol;Acc:MGI:238518 | 84,32 | 0,0 |
| ENSMUSG00000059401 | 36,60650227 | 47,2 | 0,0 | 8,899869482 | 0,01226813 | Mamld1 | mastermind-like domain containing 1 [Source:MGI Symbol;Acc:MGI:3045303] | 81,26 | 0,0 |
| ENSMUSG00000087598 | 20,05727599 | 45,7 | 0,0 | 8,901833716 | 0,012375469 | Zfp111 | zinc finger protein 111 [Source:MGI Symbol;Acc:MGI:1929114] | 47,96 | 0,0 |
| ENSMUSG00000024169 | 40,50335459 | 48,8 | 0,0 | 8,916985453 | 0,015861685 | Ift140 | intraflagellar transport 140 [Source:MGI Symbol;Acc:MGI:2146906] | 72,83 | 0,0 |
| ENSMUSG00000031016 | 47,49711208 | 46,9 | 0,0 | 8,922155708 | 0,019656952 | Wee1 | WEE 1 homolog 1 (S. pombe) [Source:MGI Symbol;Acc:MGI:103075] | 87,24 | 0,0 |
| ENSMUSG00000060397 | 32,26963396 | 48,4 | 0,0 | 8,922250782 | 0,015231624 | Zfp128 | zinc finger protein 128 [Source:MGI Symbol;Acc:MGI:2389445] | 85,32 | 0,0 |
| ENSMUSG00000089821 | 26,48683504 | 46,6 | 0,0 | 8,927431511 | 0,018806383 | Efhd1os | EF hand domain containing 1, opposite strand [Source:MGI Symbol;Acc:MGI:4413444] | 76,51 | 0,0 |
| ENSMUSG00000032369 | 34,7131684 | 45,2 | 0,0 | 8,928002169 | 0,018213656 | Plscr1 | phospholipid scramblase 1 [Source:MGI Symbol;Acc:MGI:893575] | 70,09 | 0,0 |
| ENSMUSG00000091462 | 17,30951762 | 45,4 | 0,0 | 8,92833698 | 0,012405249 | Gm17084 | predicted gene 17084 [Source:MGI Symbol;Acc:MGI:4937911] | #DIV/0! | #DIV/0! |
| ENSMUSG00000097358 | 26,80375299 | 45,6 | 0,0 | 8,932496341 | 0,015895519 | Gm26773 | predicted gene, 26773 [Source:MGI Symbol;Acc:MGI:5477267] | #DIV/0! | #DIV/0! |
| ENSMUSG00000078173 | 49,01811772 | 46,9 | 0,0 | 8,934917313 | 0,01768787 | Lenep | lens epithelial protein [Source:MGI Symbol;Acc:MGI:1930020] | 86,45 | 0,0 |
| ENSMUSG00000035840 | 69,03597595 | 46,0 | 0,0 | 8,934974451 | 0,018007843 | Lysmd3 | LysM, putative peptidoglycan-binding, domain containing 3 [Source:MGI Symbol;Acc:MGI:1915906] | 88,22 | 0,0 |
| ENSMUSG00000067424 | 36,49245274 | 56,9 | 0,0 | 8,938831689 | 0,013894484 | Zfp563 | zinc finger protein 563 [Source:MGI Symbol;Acc:MGI:2677168] | 78,91 | 0,0 |
| ENSMUSG00000042320 | 18,65883869 | 45,7 | 0,0 | 8,940564392 | 0,011753432 | Prox2 | prospero homeobox 2 [Source:MGI Symbol;Acc:MGI:1920672] | 85,75 | 0,0 |
| ENSMUSG00000080741 | 27,15640996 | 48,7 | 0,0 | 8,941563987 | 0,020330961 | Gm11398 | predicted gene 11398 [Source:MGI Symbol;Acc:MGI:3805547] | #DIV/0! | #DIV/0! |
| ENSMUSG00000023235 | 48,53973081 | 48,4 | 0,0 | 8,944041153 | 0,016615332 | Ccl25 | chemokine (C-C motif) ligand 25 [Source:MGI Symbol;Acc:MGI:1099448] | 67,19 | 0,0 |
| ENSMUSG00000026098 | 20,14742593 | 45,6 | 0,0 | 8,945782296 | 0,013210643 | Pms1 | postmeiotic segregation increased 1 (S. cerevisiae) [Source:MGI Symbol;Acc:MGI:1202302] | 76,48 | 0,0 |
| ENSMUSG00000003752 | 30,28078566 | 48,2 | 0,0 | 8,960860205 | 0,013996418 | Itpkc | inositol 1,4,5-trisphosphate 3-kinase C [Source:MGI Symbol;Acc:MGI:2442554] | 82,01 | 0,0 |
| ENSMUSG00000037214 | 50,11953462 | 47,4 | 0,0 | 8,963921868 | 0,010814203 | Thap1 | THAP domain containing, apoptosis associated protein 1 [Source:MGI Symbol;Acc:MGI:1921004] | 97,28 | 0,0 |
| ENSMUSG00000004105 | 62,70960547 | 84,9 | 0,0 | 8,966421113 | 0,014062406 | Angptl2 | angiopoietin-like 2 [Source:MGI Symbol;Acc:MGI:1347002] | 78,63 | 0,0 |
| ENSMUSG00000020614 | 461,8008978 | 887,6 | 103,5 | 8,966837915 | 0,000490711 | Fam20a | family with sequence similarity 20, member A [Source:MGI Symbol;Acc:MGI:2388266] | 100,00 | 31,6 |
| ENSMUSG00000030770 | 587,1226938 | 848,8 | 215,5 | 8,967445563 | 0,000323009 | Parva | parvin, alpha [Source:MGI Symbol;Acc:MGI:1931144] | 93,83 | 27,7 |
| ENSMUSG00000099655 | 27,04686089 | 48,0 | 0,0 | 8,96785664 | 0,01743718 | 2310034G01Rik | RIKEN cDNA 2310034G01 gene [Source:MGI Symbol;Acc:MGI:1922829] | #DIV/0! | #DIV/0! |
| ENSMUSG00000026072 | 23,70218345 | 46,9 | 0,0 | 8,969314287 | 0,01149796 | Il1r1 | interleukin 1 receptor, type I [Source:MGI Symbol;Acc:MGI:96545] | 63,45 | 0,0 |
| ENSMUSG00000050930 | 28,67499718 | 48,5 | 0,0 | 8,970185457 | 0,011255847 | Map10 | microtubule-associated protein 10 [Source:MGI Symbol;Acc:MGI:1921643] | 81,65 | 0,0 |
| ENSMUSG00000031832 | 32,90359004 | 47,2 | 0,0 | 8,978800516 | 0,019322705 | Taf1c | TA box binding protein (Tbp)-associated factor, RNA polymerase I, C [Source:MGI Symbol;Acc:MGI:1095 | 91,86 | 0,0 |
| ENSMUSG00000082226 | 27,25112251 | 47,4 | 0,0 | 8,980834188 | 0,014621969 | Gm715 | predicted gene 715 [Source:MGI Symbol;Acc:MGI:2685561] | #DIV/0! | #DIV/0! |
| ENSMUSG00000047842 | 173,5460389 | 337,2 | 1,0 | 8,98148536 | 0,000421664 | Diras2 | DIRAS family, GTP-binding RAS-like 2 [Source:MGI Symbol;Acc:MGI:1915453] | 98,40 | 0,6 |
| ENSMUSG00000035868 | 25,12290045 | 47,4 | 0,0 | 8,985957617 | 0,011277416 | 3110052M02Rik | RIKEN cDNA 3110052M02 gene [Source:MGI Symbol;Acc:MGI:1920479] | 98,92 | 0,0 |
| ENSMUSG00000033294 | 68,69084743 | 49,9 | 0,0 | 8,991226042 | 0,014423451 | Noc4l | nucleolar complex associated 4 homolog (S. cerevisiae) [Source:MGI Symbol;Acc:MGI:2140843] | 88,13 | 0,0 |
| ENSMUSG00000039735 | 37,62492075 | 48,5 | 0,0 | 8,993662965 | 0,01036577 | Fnbp1l | formin binding protein 1-like [Source:MGI Symbol;Acc:MGI:1925642] | 64,96 | 0,0 |

| | | | | | | | | | |
|---|---|---|---|---|---|---|---|---|---|
| ENSMUSG00000086212 | 27,06670428 | 50,8 | 0,0 | 8,994013644 | 0,013508497 | Mkln1os | kelin 1, intracellular mediator containing kelch motifs, opposite strand [Source:MGI Symbol;Acc:MGI:3651 | #DIV/0! | #DIV/0! |
| ENSMUSG00000085282 | 30,31896751 | 47,9 | 0,0 | 8,996286937 | 0,019944351 | Gm15663 | predicted gene 15663 [Source:MGI Symbol;Acc:MGI:3783105] | 58,46 | 0,0 |
| ENSMUSG00000019775 | 45,62218654 | 49,8 | 0,0 | 8,998517151 | 0,017886688 | Rgs17 | regulator of G-protein signaling 17 [Source:MGI Symbol;Acc:MGI:1927469] | 39,55 | 0,0 |
| ENSMUSG00000034308 | 30,44526473 | 51,0 | 0,0 | 9,003456259 | 0,017208103 | Sdr42e1 | short chain dehydrogenase/reductase family 42E, member 1 [Source:MGI Symbol;Acc:MGI:1921282] | 87,26 | 0,0 |
| ENSMUSG00000049600 | 27,88073825 | 48,9 | 0,0 | 9,007152935 | 0,012343509 | Zbtb45 | zinc finger and BTB domain containing 45 [Source:MGI Symbol;Acc:MGI:2685003] | 78,19 | 0,0 |
| ENSMUSG00000028744 | 30,87263449 | 52,7 | 0,0 | 9,012862673 | 0,015977886 | Pqlc2 | PQ loop repeat containing 2 [Source:MGI Symbol;Acc:MGI:2384837] | 53,41 | 0,0 |
| ENSMUSG00000087436 | 41,79692076 | 53,0 | 0,0 | 9,013066976 | 0,01427933 | Gm16156 | predicted gene 16156 [Source:MGI Symbol;Acc:MGI:3801720] | 60,68 | 0,0 |
| ENSMUSG00000020037 | 412,8020798 | 648,1 | 25,9 | 9,016134343 | 0,000492468 | Rfx4 | regulatory factor X, 4 (influences HLA class II expression) [Source:MGI Symbol;Acc:MGI:1918387] | 93,61 | 5,6 |
| ENSMUSG00000079043 | 30,81915012 | 51,2 | 0,0 | 9,018967377 | 0,01024489 | Fastkd5 | FAST kinase domains 5 [Source:MGI Symbol;Acc:MGI:2139469] | 86,31 | 0,0 |
| ENSMUSG00000053414 | 25,71555857 | 51,1 | 0,0 | 9,020449572 | 0,013706157 | Hunk | hormonally upregulated Neu-associated kinase [Source:MGI Symbol;Acc:MGI:1347352] | 73,33 | 0,0 |
| ENSMUSG00000028865 | 37,4521756 | 50,9 | 0,0 | 9,020990551 | 0,016261287 | Cd164l2 | CD164 sialomucin-like 2 [Source:MGI Symbol;Acc:MGI:1916905] | 73,70 | 0,0 |
| ENSMUSG00000074736 | 34,59974835 | 50,1 | 0,0 | 9,024224277 | 0,014423451 | Syndig1 | synapse differentiation inducing 1 [Source:MGI Symbol;Acc:MGI:3702158] | 81,82 | 0,0 |
| ENSMUSG00000037813 | 33,17583595 | 48,6 | 0,0 | 9,024955266 | 0,017063717 | D630003M21Rik | RIKEN cDNA D630003M21 gene [Source:MGI Symbol;Acc:MGI:3606579] | 66,51 | 0,0 |
| ENSMUSG00000051339 | 42,98691135 | 61,9 | 0,0 | 9,029008498 | 0,012569775 | 2900026A02Rik | RIKEN cDNA 2900026A02 gene [Source:MGI Symbol;Acc:MGI:1920194] | 70,51 | 0,0 |
| ENSMUSG00000079605 | 45,83973633 | 49,8 | 0,0 | 9,032426133 | 0,01508771 | Zbtb9 | zinc finger and BTB domain containing 9 [Source:MGI Symbol;Acc:MGI:1918022] | 91,67 | 0,0 |
| ENSMUSG00000038354 | 50,67408676 | 55,0 | 0,0 | 9,032808843 | 0,014677393 | Ankrd35 | ankyrin repeat domain 35 [Source:MGI Symbol;Acc:MGI:2442590] | 69,54 | 0,0 |
| ENSMUSG00000025198 | 84,30903153 | 55,4 | 0,0 | 9,032850273 | 0,024311853 | Erlin1 | ER lipid raft associated 1 [Source:MGI Symbol;Acc:MGI:2387613] | 81,18 | 0,0 |
| ENSMUSG00000046607 | 36,04059821 | 55,3 | 0,0 | 9,034353546 | 0,015050976 | Hrk | harakiri, BCL2 interacting protein (contains only BH3 domain) [Source:MGI Symbol;Acc:MGI:1201608] | 67,69 | 0,0 |
| ENSMUSG00000031538 | 46,15896735 | 51,5 | 0,0 | 9,034669859 | 0,015786213 | Plat | plasminogen activator, tissue [Source:MGI Symbol;Acc:MGI:97610] | 91,07 | 0,0 |
| ENSMUSG00000004655 | 100,9111054 | 97,6 | 0,0 | 9,039180549 | 0,012660573 | Aqp1 | aquaporin 1 [Source:MGI Symbol;Acc:MGI:103201] | 79,94 | 0,0 |
| ENSMUSG00000032598 | 157,4395509 | 275,7 | 0,6 | 9,043134662 | 0,002831118 | Nckipsd | NCK interacting protein with SH3 domain [Source:MGI Symbol;Acc:MGI:1931834] | 94,10 | 0,4 |
| ENSMUSG00000097343 | 21,64647163 | 51,9 | 0,0 | 9,044020957 | 0,010956967 | 9030407P20Rik | RIKEN cDNA 9030407P20 gene [Source:MGI Symbol;Acc:MGI:1918818] | #DIV/0! | #DIV/0! |
| ENSMUSG00000037735 | 21,74885318 | 49,3 | 0,0 | 9,044812342 | 0,015826582 | 2810032G03Rik | RIKEN cDNA 2810032G03 gene [Source:MGI Symbol;Acc:MGI:1919919] | #DIV/0! | #DIV/0! |
| ENSMUSG00000026894 | 24,30585003 | 48,5 | 0,0 | 9,045055557 | 0,01081369 | Morn5 | MORN repeat containing 5 [Source:MGI Symbol;Acc:MGI:1922745] | 98,13 | 0,0 |
| ENSMUSG00000048450 | 425,6923305 | 605,8 | 26,8 | 9,045793952 | 0,000942143 | Msx1 | msh homeobox 1 [Source:MGI Symbol;Acc:MGI:97168] | 99,14 | 16,3 |
| ENSMUSG00000042742 | 218,3353554 | 364,5 | 1,0 | 9,04965363 | 0,000243064 | B630005N14Rik | RIKEN cDNA B630005N14 gene [Source:MGI Symbol;Acc:MGI:2141466] | 72,70 | 0,4 |
| ENSMUSG00000009670 | 29,37557944 | 61,9 | 0,0 | 9,056840583 | 0,010481298 | Tex11 | testis expressed gene 11 [Source:MGI Symbol;Acc:MGI:1933237] | 46,06 | 0,0 |
| ENSMUSG00000032850 | 34,69858583 | 49,7 | 0,0 | 9,057151631 | 0,016961595 | Rnft2 | ring finger protein, transmembrane 2 [Source:MGI Symbol;Acc:MGI:2442859] | 71,57 | 0,0 |
| ENSMUSG00000035007 | 234,1173696 | 363,7 | 0,9 | 9,062128373 | 0,000225051 | Rundc1 | RUN domain containing 1 [Source:MGI Symbol;Acc:MGI:2144506] | 99,65 | 0,7 |
| ENSMUSG00000087628 | 21,89225862 | 49,9 | 0,0 | 9,063429466 | 0,013915735 | Gm13028 | predicted gene 13028 [Source:MGI Symbol;Acc:MGI:3651504] | #DIV/0! | #DIV/0! |
| ENSMUSG00000039831 | 22,40028958 | 61,9 | 0,0 | 9,066594005 | 0,010167511 | Arhgap29 | Rho GTPase activating protein 29 [Source:MGI Symbol;Acc:MGI:2443818] | 63,09 | 0,0 |
| ENSMUSG00000028631 | 44,69042336 | 55,8 | 0,0 | 9,069400041 | 0,014359555 | Kcnq4 | potassium voltage-gated channel, subfamily Q, member 4 [Source:MGI Symbol;Acc:MGI:1926803] | 83,10 | 0,0 |
| ENSMUSG00000038235 | 48,70512072 | 76,1 | 0,0 | 9,074358309 | 0,009440287 | F11r | F11 receptor [Source:MGI Symbol;Acc:MGI:1321398] | 76,53 | 0,0 |
| ENSMUSG00000035095 | 32,40008074 | 50,4 | 0,0 | 9,077181192 | 0,011421384 | Fam167a | family with sequence similarity 167, member A [Source:MGI Symbol;Acc:MGI:3606565] | 83,83 | 0,0 |
| ENSMUSG00000043461 | 31,35514659 | 51,4 | 0,0 | 9,083212355 | 0,015966004 | Sptssb | serine palmitoyltransferase, small subunit B [Source:MGI Symbol;Acc:MGI:1913433] | 92,02 | 0,0 |
| ENSMUSG00000029480 | 59,57884124 | 53,5 | 0,0 | 9,086291277 | 0,020886638 | Dhx37 | DEAH (Asp-Glu-Ala-His) box polypeptide 37 [Source:MGI Symbol;Acc:MGI:3028576] | 79,04 | 0,0 |
| ENSMUSG00000031104 | 18,90686926 | 50,5 | 0,0 | 9,088195801 | 0,010825187 | Rab33a | RAB33A, member RAS oncogene family [Source:MGI Symbol;Acc:MGI:109493] | 90,30 | 0,0 |
| ENSMUSG00000042857 | 25,54775406 | 52,0 | 0,0 | 9,090824575 | 0,01036577 | Gm9776 | predicted gene 9776 [Source:MGI Symbol;Acc:MGI:3641792] | 66,61 | 0,0 |
| ENSMUSG00000022723 | 42,70838215 | 52,4 | 0,0 | 9,093655733 | 0,00992454 | Crybg3 | beta-gamma crystallin domain containing 3 [Source:MGI Symbol;Acc:MGI:2676311] | 28,22 | 0,0 |
| ENSMUSG00000058093 | 87,95725585 | 52,0 | 0,0 | 9,095552788 | 0,026334424 | Zfp729b | zinc finger protein 729b [Source:MGI Symbol;Acc:MGI:2145180] | 75,95 | 0,0 |
| ENSMUSG00000097573 | 18,95197444 | 52,2 | 0,0 | 9,109583723 | 0,010213022 | G730003C15Rik | RIKEN cDNA G730003C15 gene [Source:MGI Symbol;Acc:MGI:3641700] | 87,44 | 0,0 |
| ENSMUSG00000028100 | 40,27922534 | 52,5 | 0,0 | 9,113110988 | 0,01738593 | Nudt17 | nudix (nucleoside diphosphate linked moiety X)-type motif 17 [Source:MGI Symbol;Acc:MGI:1925623] | 52,78 | 0,0 |
| ENSMUSG00000070883 | 24,39634676 | 51,5 | 0,0 | 9,115905032 | 0,01181122 | Ccdc173 | coiled-coil domain containing 173 [Source:MGI Symbol;Acc:MGI:1923100] | 91,11 | 0,0 |
| ENSMUSG00000037325 | 60,69917781 | 60,4 | 0,0 | 9,11601654 | 0,014282711 | Bbs7 | Bardet-Biedl syndrome 7 (human) [Source:MGI Symbol;Acc:MGI:1918742] | 73,03 | 0,0 |
| ENSMUSG00000033207 | 25,19524956 | 53,2 | 0,0 | 9,120415127 | 0,010179082 | Mamdc2 | MAM domain containing 2 [Source:MGI Symbol;Acc:MGI:1918988] | 82,22 | 0,0 |
| ENSMUSG00000068417 | 25,97319161 | 51,8 | 0,0 | 9,120836902 | 0,010166171 | Pnp2 | purine-nucleoside phosphorylase 2 [Source:MGI Symbol;Acc:MGI:3712328] | 82,58 | 0,0 |
| ENSMUSG00000048728 | 26,18536528 | 52,7 | 0,0 | 9,12108032 | 0,009076555 | Zfp454 | zinc finger protein 454 [Source:MGI Symbol;Acc:MGI:2679253] | 92,55 | 0,0 |
| ENSMUSG00000041390 | 39,3868109 | 53,3 | 0,0 | 9,122020965 | 0,00992454 | Mdfic | MyoD family inhibitor domain containing [Source:MGI Symbol;Acc:MGI:104611] | 41,89 | 0,0 |
| ENSMUSG00000022867 | 198,384883 | 361,5 | 0,6 | 9,126358034 | 9,23286E-05 | Usp25 | ubiquitin specific peptidase 25 [Source:MGI Symbol;Acc:MGI:1353655] | 99,97 | 0,4 |
| ENSMUSG00000031548 | 63,73159125 | 70,9 | 0,0 | 9,129111621 | 0,015953227 | Sfrp1 | secreted frizzled-related protein 1 [Source:MGI Symbol;Acc:MGI:892014] | 79,35 | 0,0 |
| ENSMUSG00000071252 | 20,22923355 | 56,7 | 0,0 | 9,1294238 | 0,013831112 | 2210408I21Rik | RIKEN cDNA 2210408I21 gene [Source:MGI Symbol;Acc:MGI:1919621] | 68,02 | 0,0 |
| ENSMUSG00000061079 | 107,9033023 | 54,0 | 0,0 | 9,135380729 | 0,01949513 | Zfp143 | zinc finger protein 143 [Source:MGI Symbol;Acc:MGI:1277969] | 91,80 | 0,0 |
| ENSMUSG00000035878 | 46,99272422 | 56,3 | 0,0 | 9,137816498 | 0,017011651 | Hykk | hydroxylysine kinase 1 [Source:MGI Symbol;Acc:MGI:2443139] | 77,21 | 0,0 |
| ENSMUSG00000028796 | 453,7648352 | 839,5 | 3,7 | 9,139545142 | 2,93161E-06 | Phc2 | polyhomeotic-like 2 (Drosophila) [Source:MGI Symbol;Acc:MGI:1860454] | 67,80 | 1,9 |
| ENSMUSG00000072762 | 30,45635547 | 53,3 | 0,0 | 9,139947328 | 0,010465506 | 4930522L14Rik | RIKEN cDNA 4930522L14 gene [Source:MGI Symbol;Acc:MGI:1925270] | 80,50 | 0,0 |
| ENSMUSG00000027534 | 44,72171701 | 52,4 | 0,0 | 9,140244064 | 0,014677393 | Snx16 | sorting nexin 16 [Source:MGI Symbol;Acc:MGI:1921968] | 87,06 | 0,0 |
| ENSMUSG00000025348 | 81,31181741 | 53,6 | 0,0 | 9,141686127 | 0,020528945 | Itga7 | integrin alpha 7 [Source:MGI Symbol;Acc:MGI:102700] | 90,34 | 0,0 |
| ENSMUSG00000040749 | 47,96608671 | 53,3 | 0,0 | 9,144122151 | 0,016020315 | Siah1b | seven in absentia 1B [Source:MGI Symbol;Acc:MGI:108063] | 88,09 | 0,0 |
| ENSMUSG00000002617 | 55,52422482 | 61,0 | 0,0 | 9,144841422 | 0,016163139 | Zfp40 | zinc finger protein 40 [Source:MGI Symbol;Acc:MGI:99185] | 75,95 | 0,0 |
| ENSMUSG00000074862 | 26,91483882 | 56,4 | 0,0 | 9,145200831 | 0,012772283 | BC025920 | cDNA sequence BC025920 [Source:MGI Symbol;Acc:MGI:2670983] | 84,02 | 0,0 |
| ENSMUSG00000038302 | 255,5724832 | 486,4 | 1,2 | 9,150603593 | 0,000318756 | Lace1 | lactation elevated 1 [Source:MGI Symbol;Acc:MGI:2148801] | 95,64 | 1,0 |
| ENSMUSG00000035275 | 248,0097551 | 334,9 | 1,0 | 9,153158546 | 0,002072649 | Raver2 | ribonucleoprotein, PTB-binding 2 [Source:MGI Symbol;Acc:MGI:2443623] | 76,45 | 0,5 |
| ENSMUSG00000026354 | 23,95607432 | 64,2 | 0,0 | 9,155298805 | 0,009961272 | Lct | lactase [Source:MGI Symbol;Acc:MGI:104576] | 70,59 | 0,0 |
| ENSMUSG00000036854 | 453,7225793 | 752,7 | 66,9 | 9,159333957 | 0,000532199 | Hspb6 | heat shock protein, alpha-crystallin-related, B6 [Source:MGI Symbol;Acc:MGI:2685325] | 99,90 | 27,0 |
| ENSMUSG00000052234 | 25,26216835 | 53,2 | 0,0 | 9,159780191 | 0,015100456 | Epx | eosinophil peroxidase [Source:MGI Symbol;Acc:MGI:107569] | 64,94 | 0,0 |
| ENSMUSG00000003411 | 35,6274346 | 56,7 | 0,0 | 9,160492575 | 0,018140331 | Rab3b | RAB3B, member RAS oncogene family [Source:MGI Symbol;Acc:MGI:1917158] | 43,75 | 0,0 |
| ENSMUSG00000024544 | 26,50131954 | 54,4 | 0,0 | 9,160625145 | 0,012958678 | Ldlrad4 | low density lipoprotein receptor class A domain containing 4 [Source:MGI Symbol;Acc:MGI:1277150] | 89,53 | 0,0 |
| ENSMUSG00000026224 | 30,18694993 | 53,7 | 0,0 | 9,161430341 | 0,015622006 | 4933407L21Rik | RIKEN cDNA 4933407L21 gene [Source:MGI Symbol;Acc:MGI:1918391] | 93,07 | 0,0 |
| ENSMUSG00000021319 | 24,26599887 | 55,8 | 0,0 | 9,162193438 | 0,008475401 | Sfrp4 | secreted frizzled-related protein 4 [Source:MGI Symbol;Acc:MGI:892010] | 95,29 | 0,0 |
| ENSMUSG00000062563 | 22,90459676 | 55,6 | 0,0 | 9,16453437 | 0,009976334 | Cys1 | cystin 1 [Source:MGI Symbol;Acc:MGI:2177632] | 52,84 | 0,0 |
| ENSMUSG00000039137 | 51,15705158 | 57,4 | 0,0 | 9,165877402 | 0,013597821 | Whrn | whirlin [Source:MGI Symbol;Acc:MGI:2682003] | 72,65 | 0,0 |
| ENSMUSG00000056832 | 23,67180324 | 57,5 | 0,0 | 9,166490334 | 0,008920937 | Ttc26 | tetratricopeptide repeat domain 26 [Source:MGI Symbol;Acc:MGI:2444853] | 85,31 | 0,0 |
| ENSMUSG00000032494 | 27,01054738 | 53,8 | 0,0 | 9,169482364 | 0,015473364 | Tdgf1 | teratocarcinoma-derived growth factor 1 [Source:MGI Symbol;Acc:MGI:98658] | 67,88 | 0,0 |
| ENSMUSG00000035126 | 34,30631662 | 54,1 | 0,0 | 9,170389238 | 0,013940839 | Wdr78 | WD repeat domain 78 [Source:MGI Symbol;Acc:MGI:2385328] | 82,47 | 0,0 |
| ENSMUSG00000029735 | 40,1462837 | 54,9 | 0,0 | 9,170454146 | 0,014935367 | Tpk1 | thiamine pyrophosphokinase [Source:MGI Symbol;Acc:MGI:1352500] | 94,00 | 0,0 |
| ENSMUSG00000024534 | 56,94591822 | 54,8 | 0,0 | 9,183522876 | 0,016490229 | Sncaip | synuclein, alpha interacting protein (synphilin) [Source:MGI Symbol;Acc:MGI:1915097] | 73,39 | 0,0 |
| ENSMUSG00000062110 | 41,78427934 | 54,7 | 0,0 | 9,193745827 | 0,015343151 | Scfd2 | Sec1 family domain containing 2 [Source:MGI Symbol;Acc:MGI:2443446] | 50,92 | 0,0 |
| ENSMUSG00000060260 | 35,02679493 | 56,5 | 0,0 | 9,195109949 | 0,012457662 | Pwwp2b | PWWP domain containing 2B [Source:MGI Symbol;Acc:MGI:2142008] | 88,96 | 0,0 |
| ENSMUSG00000032431 | 75,73571432 | 64,3 | 0,0 | 9,2018597 | 0,011155709 | Crtap | cartilage associated protein [Source:MGI Symbol;Acc:MGI:1891221] | 89,71 | 0,0 |
| ENSMUSG00000066042 | 50,2104564 | 55,9 | 0,0 | 9,205914256 | 0,017005951 | Med18 | mediator complex subunit 18 [Source:MGI Symbol;Acc:MGI:1914469] | 83,20 | 0,0 |
| ENSMUSG00000022610 | 37,2279156 | 57,8 | 0,0 | 9,209278362 | 0,015231624 | Mapk12 | mitogen-activated protein kinase 12 [Source:MGI Symbol;Acc:MGI:1353438] | 96,75 | 0,0 |
| ENSMUSG00000034379 | 31,33833125 | 58,8 | 0,0 | 9,216266338 | 0,013575227 | Wdr5b | WD repeat domain 5B [Source:MGI Symbol;Acc:MGI:1916794] | 94,49 | 0,0 |
| ENSMUSG00000043487 | 26,23231264 | 58,0 | 0,0 | 9,222659717 | 0,009373845 | Acot6 | acyl-CoA thioesterase 6 [Source:MGI Symbol;Acc:MGI:1921287] | 88,25 | 0,0 |
| ENSMUSG00000021782 | 156,7301169 | 331,8 | 0,6 | 9,229042419 | 0,000820593 | Dlg5 | discs, large homolog 5 (Drosophila) [Source:MGI Symbol;Acc:MGI:1918478] | 96,13 | 0,2 |
| ENSMUSG00000044060 | 68,72616618 | 66,0 | 0,0 | 9,235845687 | 0,007571359 | A830010M20Rik | RIKEN cDNA A830010M20 gene [Source:MGI Symbol;Acc:MGI:2445097] | 54,93 | 0,0 |
| ENSMUSG00000019718 | 27,40212162 | 58,6 | 0,0 | 9,236633728 | 0,00923671 | L3hypdh | L-3-hydroxyproline dehydratase (trans-) [Source:MGI Symbol;Acc:MGI:1914467] | 91,52 | 0,0 |
| ENSMUSG00000028558 | 37,4954821 | 57,5 | 0,0 | 9,238669294 | 0,013897211 | Calr4 | calreticulin 4 [Source:MGI Symbol;Acc:MGI:2140435] | 84,07 | 0,0 |
| ENSMUSG00000022687 | 58,56893209 | 79,4 | 0,0 | 9,241231503 | 0,015239583 | Boc | ıdhesion molecule-related/down-regulated by oncogenes (Cdon) binding protein [Source:MGI Symbol;Acc | 81,42 | 0,0 |
| ENSMUSG00000002732 | 51,8586689 | 62,3 | 0,0 | 9,243859955 | 0,019770307 | Fkbp7 | FK506 binding protein 7 [Source:MGI Symbol;Acc:MGI:1336879] | 99,69 | 0,0 |
| ENSMUSG00000079415 | 31,34596111 | 57,8 | 0,0 | 9,245570592 | 0,013193418 | Cntf | ciliary neurotrophic factor [Source:MGI Symbol;Acc:MGI:88439] | 86,42 | 0,0 |
| ENSMUSG00000026727 | 374,8116654 | 647,6 | 7,0 | 9,250409403 | 0,000693649 | Rsu1 | Ras suppressor protein 1 [Source:MGI Symbol;Acc:MGI:103040] | 90,41 | 2,8 |
| ENSMUSG00000051184 | 52,00882435 | 58,0 | 0,0 | 9,252268247 | 0,011657155 | Zfp524 | zinc finger protein 524 [Source:MGI Symbol;Acc:MGI:1916740] | 78,23 | 0,0 |
| ENSMUSG00000085398 | 47,43638055 | 61,1 | 0,0 | 9,256483845 | 0,020636595 | 4931406G06Rik | RIKEN cDNA 4931406G06 gene [Source:MGI Symbol;Acc:MGI:1918228] | #DIV/0! | #DIV/0! |
| ENSMUSG00000032122 | 65,95837448 | 63,8 | 0,0 | 9,257228507 | 0,015972989 | Slc37a2 | lute carrier family 37 (glycerol-3-phosphate transporter), member 2 [Source:MGI Symbol;Acc:MGI:192969 | 76,18 | 0,0 |
| ENSMUSG00000097718 | 29,63689705 | 59,7 | 0,0 | 9,2767616 | 0,012492576 | Gm26896 | predicted gene, 26896 [Source:MGI Symbol;Acc:MGI:5477390] | #DIV/0! | #DIV/0! |
| ENSMUSG00000024227 | 51,83891718 | 63,8 | 0,0 | 9,278005455 | 0,017886688 | Pdzph1 | PDZ and pleckstrin homology domains 1 [Source:MGI Symbol;Acc:MGI:1916489] | 26,73 | 0,0 |
| ENSMUSG00000033966 | 34,41274296 | 60,0 | 0,0 | 9,281363115 | 0,011063774 | Cdkl4 | cyclin-dependent kinase-like 4 [Source:MGI Symbol;Acc:MGI:3587025] | 92,78 | 0,0 |
| ENSMUSG00000033722 | 36,33759014 | 59,0 | 0,0 | 9,282878099 | 0,008670473 | BC034090 | cDNA sequence BC034090 [Source:MGI Symbol;Acc:MGI:2672904] | 83,40 | 0,0 |
| ENSMUSG00000020635 | 35,9163229 | 58,6 | 0,0 | 9,284602073 | 0,014935367 | Fkbp1b | FK506 binding protein 1b [Source:MGI Symbol;Acc:MGI:1336205] | 97,55 | 0,0 |
| ENSMUSG00000031239 | 74,05927672 | 63,4 | 0,0 | 9,286269129 | 0,021396674 | Itm2a | integral membrane protein 2A [Source:MGI Symbol;Acc:MGI:107706] | 92,60 | 0,0 |
| ENSMUSG00000045802 | 18,88740231 | 58,9 | 0,0 | 9,28700828 | 0,009367698 | Hsf3 | heat shock transcription factor 3 [Source:MGI Symbol;Acc:MGI:3045337] | 86,67 | 0,0 |
| ENSMUSG00000063605 | 33,73607258 | 57,8 | 0,0 | 9,2876243 | 0,008249347 | Ccdc102a | coiled-coil domain containing 102A [Source:MGI Symbol;Acc:MGI:2686927] | 91,67 | 0,0 |
| ENSMUSG00000020774 | 50,30259469 | 64,6 | 0,0 | 9,29038201 | 0,010510621 | Aspa | aspartoacylase [Source:MGI Symbol;Acc:MGI:87914] | 73,14 | 0,0 |
| ENSMUSG00000020810 | 28,2985732 | 58,5 | 0,0 | 9,29099987 | 0,012574362 | Cygb | cytoglobin [Source:MGI Symbol;Acc:MGI:2149481] | 83,24 | 0,0 |
| ENSMUSG00000043614 | 34,66588377 | 59,4 | 0,0 | 9,2966055 | 0,013575227 | Vps37d | vacuolar protein sorting 37D (yeast) [Source:MGI Symbol;Acc:MGI:2159402] | 92,66 | 0,0 |
| ENSMUSG00000060429 | 29,43708707 | 63,6 | 0,0 | 9,298277499 | 0,009561077 | Sntb1 | syntrophin, basic 1 [Source:MGI Symbol;Acc:MGI:101781] | 74,65 | 0,0 |
| ENSMUSG00000075327 | 36,32707686 | 58,6 | 0,0 | 9,298815093 | 0,012503927 | Zbtb2 | zinc finger and BTB domain containing 2 [Source:MGI Symbol;Acc:MGI:2685949] | 85,60 | 0,0 |
| ENSMUSG00000032204 | 34,85207588 | 61,4 | 0,0 | 9,298985784 | 0,008594832 | Aqp9 | aquaporin 9 [Source:MGI Symbol;Acc:MGI:1891066] | 75,48 | 0,0 |
| ENSMUSG00000045248 | 34,87659444 | 63,7 | 0,0 | 9,304485073 | 0,011804736 | Med26 | mediator complex subunit 26 [Source:MGI Symbol;Acc:MGI:1917875] | 86,04 | 0,0 |
| ENSMUSG00000039063 | 38,68207398 | 60,6 | 0,0 | 9,309207403 | 0,008430306 | Echdc3 | enoyl Coenzyme A hydratase domain containing 3 [Source:MGI Symbol;Acc:MGI:1915106] | 79,65 | 0,0 |
| ENSMUSG00000043496 | 714,9791074 | 1120,4 | 114,4 | 9,311505575 | 0,000245776 | Tril | TLR4 interactor with leucine-rich repeats [Source:MGI Symbol;Acc:MGI:1914123] | 99,98 | 20,1 |
| ENSMUSG00000003929 | 39,58579643 | 61,4 | 0,0 | 9,311600078 | 0,013686241 | Zfp81 | zinc finger protein 81 [Source:MGI Symbol;Acc:MGI:1890752] | 70,65 | 0,0 |
| ENSMUSG00000022783 | 46,37629992 | 61,5 | 0,0 | 9,316369831 | 0,009373845 | Spag6l | sperm associated antigen 6-like [Source:MGI Symbol;Acc:MGI:1354388] | 96,56 | 0,0 |
| ENSMUSG00000057835 | 28,21831154 | 60,0 | 0,0 | 9,319425091 | 0,008651939 | Zfp119a | zinc finger protein 119a [Source:MGI Symbol;Acc:MGI:1345189] | 92,10 | 0,0 |
| ENSMUSG00000093497 | 37,40297644 | 62,0 | 0,0 | 9,322679345 | 0,010849987 | Gm20713 | predicted gene 20713 [Source:MGI Symbol;Acc:MGI:5313160] | #DIV/0! | #DIV/0! |
| ENSMUSG00000022853 | 48,61475081 | 66,0 | 0,0 | 9,331561949 | 0,016066445 | Ehhadh | l-Coenzyme A, hydratase/3-hydroxyacyl Coenzyme A dehydrogenase [Source:MGI Symbol;Acc:MGI:1277 | 88,23 | 0,0 |
| ENSMUSG00000060548 | 530,8751706 | 827,3 | 118,6 | 9,332821306 | 0,000957662 | Tnfrsf19 | tumor necrosis factor receptor superfamily, member 19 [Source:MGI Symbol;Acc:MGI:1352474] | 93,82 | 13,7 |
| ENSMUSG00000066191 | 23,10567717 | 59,9 | 0,0 | 9,334164371 | 0,008355771 | Anks6 | ankyrin repeat and sterile alpha motif domain containing 6 [Source:MGI Symbol;Acc:MGI:1922941] | 92,06 | 0,0 |
| ENSMUSG00000032327 | 75,76924924 | 77,9 | 0,0 | 9,337447886 | 0,014231804 | Stra6 | stimulated by retinoic acid gene 6 [Source:MGI Symbol;Acc:MGI:107742] | 56,95 | 0,0 |
| ENSMUSG00000026705 | 44,27946454 | 64,5 | 0,0 | 9,344012863 | 0,014975476 | Klhl20 | kelch-like 20 [Source:MGI Symbol;Acc:MGI:2444855] | 88,98 | 0,0 |
| ENSMUSG00000032258 | 44,8658405 | 65,1 | 0,0 | 9,346581216 | 0,008053471 | Lca5 | Leber congenital amaurosis 5 (human) [Source:MGI Symbol;Acc:MGI:1923032] | 73,34 | 0,0 |
| ENSMUSG00000073988 | 608,1843833 | 1069,2 | 126,5 | 9,347708529 | 0,001002202 | Ttpa | tocopherol (alpha) transfer protein [Source:MGI Symbol;Acc:MGI:1354168] | 95,49 | 15,4 |
| ENSMUSG00000096960 | 28,76120672 | 60,5 | 0,0 | 9,348931555 | 0,011067849 | A230028O05Rik | RIKEN cDNA A230028O05 gene [Source:MGI Symbol;Acc:MGI:2442126] | #DIV/0! | #DIV/0! |
| ENSMUSG00000015980 | 27,1795147 | 62,9 | 0,0 | 9,350143476 | 0,008475401 | Lrrc27 | leucine rich repeat containing 27 [Source:MGI Symbol;Acc:MGI:1923862] | 77,29 | 0,0 |
| ENSMUSG00000069763 | 571,1147391 | 947,0 | 121,6 | 9,350761094 | 0,001264052 | Tmem100 | transmembrane protein 100 [Source:MGI Symbol;Acc:MGI:1915138] | 84,53 | 25,8 |
| ENSMUSG00000049387 | 30,07528999 | 61,6 | 0,0 | 9,353482107 | 0,010179082 | Cox7b2 | cytochrome c oxidase subunit VIIb2 [Source:MGI Symbol;Acc:MGI:1925424] | 93,17 | 0,0 |
| ENSMUSG00000109147 | 32,90590967 | 63,5 | 0,0 | 9,358230402 | 0,012462267 | RP24-346B15.3 | | #DIV/0! | #DIV/0! |
| ENSMUSG00000046818 | 327,0055694 | 601,4 | 2,0 | 9,358582851 | 0,000213819 | Ddit4l | DNA-damage-inducible transcript 4-like [Source:MGI Symbol;Acc:MGI:1920534] | 97,57 | 0,8 |
| ENSMUSG00000083674 | 28,98759334 | 61,2 | 0,0 | 9,359213468 | 0,013271859 | Zfp133-ps | zinc finger protein 133, pseudogene [Source:MGI Symbol;Acc:MGI:2176612] | #DIV/0! | #DIV/0! |
| ENSMUSG00000028854 | 36,28770957 | 65,1 | 0,0 | 9,367226202 | 0,00992454 | Slc9a1 | solute carrier family 9 (sodium/hydrogen exchanger), member 1 [Source:MGI Symbol;Acc:MGI:102462] | 85,40 | 0,0 |
| ENSMUSG00000036368 | 41,4904679 | 62,5 | 0,0 | 9,369175619 | 0,010490558 | Rmdn2 | regulator of microtubule dynamics 2 [Source:MGI Symbol;Acc:MGI:2147043] | 95,97 | 0,0 |
| ENSMUSG00000096433 | 29,70702091 | 64,4 | 0,0 | 9,369932457 | 0,008993454 | Gm4944 | predicted gene 4944 [Source:MGI Symbol;Acc:MGI:3643318] | 66,40 | 0,0 |
| ENSMUSG00000051029 | 67,53258271 | 71,5 | 0,0 | 9,370932733 | 0,013831112 | Serpinb1b | serine (or cysteine) peptidase inhibitor, clade B, member 1b [Source:MGI Symbol;Acc:MGI:2445361] | 97,15 | 0,0 |
| ENSMUSG00000036353 | 76,50486225 | 76,2 | 0,0 | 9,37591514 | 0,01521624 | P2ry12 | purinergic receptor P2Y, G-protein coupled 12 [Source:MGI Symbol;Acc:MGI:1918089] | 87,10 | 0,0 |
| ENSMUSG00000053536 | 424,1929643 | 488,9 | 1,0 | 9,378572231 | 0,000119227 | Cstf2t | cleavage stimulation factor, 3' pre-RNA subunit 2, tau [Source:MGI Symbol;Acc:MGI:1932622] | 95,96 | 0,7 |
| ENSMUSG00000027803 | 748,0561478 | 1156,0 | 67,5 | 9,383283909 | 0,000101185 | Wwtr1 | WW domain containing transcription regulator 1 [Source:MGI Symbol;Acc:MGI:1917649] | 98,95 | 11,8 |
| ENSMUSG00000100147 | 35,26146947 | 63,4 | 0,0 | 9,386849925 | 0,014677393 | 1700047M11Rik | RIKEN cDNA 1700047M11 gene [Source:MGI Symbol;Acc:MGI:1914580] | #DIV/0! | #DIV/0! |
| ENSMUSG00000023912 | 35,347698 | 62,7 | 0,0 | 9,38708612 | 0,010585234 | Slc25a27 | solute carrier family 25, member 27 [Source:MGI Symbol;Acc:MGI:1921261] | 95,72 | 0,0 |
| ENSMUSG00000023336 | 33,98089524 | 63,4 | 0,0 | 9,392149277 | 0,017063717 | Wfdc1 | WAP four-disulfide core domain 1 [Source:MGI Symbol;Acc:MGI:1915116] | 99,31 | 0,0 |
| ENSMUSG00000027800 | 48,56969379 | 65,7 | 0,0 | 9,396866657 | 0,016447916 | Tm4sf1 | transmembrane 4 superfamily member 1 [Source:MGI Symbol;Acc:MGI:104678] | 51,00 | 0,0 |
| ENSMUSG00000047583 | 43,12131459 | 68,5 | 0,0 | 9,397838196 | 0,011804736 | Tyw3 | tRNA-yW synthesizing protein 3 homolog (S. cerevisiae) [Source:MGI Symbol;Acc:MGI:2445040] | 57,29 | 0,0 |
| ENSMUSG00000010136 | 62,63666529 | 73,4 | 0,0 | 9,407465282 | 0,00754905 | Pifo | primary cilia formation [Source:MGI Symbol;Acc:MGI:1923670] | 79,41 | 0,0 |
| ENSMUSG00000028454 | 46,72413226 | 67,6 | 0,0 | 9,41316008 | 0,012419384 | Pigo | phosphatidylinositol glycan anchor biosynthesis, class O [Source:MGI Symbol;Acc:MGI:1861452] | 86,67 | 0,0 |
| ENSMUSG00000063558 | 30,92138368 | 64,3 | 0,0 | 9,417651529 | 0,007393309 | Aox1 | aldehyde oxidase 1 [Source:MGI Symbol;Acc:MGI:88035] | 85,17 | 0,0 |
| ENSMUSG00000029416 | 41,47512671 | 66,8 | 0,0 | 9,42411526 | 0,01077691 | Slc15a4 | solute carrier family 15, member 4 [Source:MGI Symbol;Acc:MGI:2140796] | 87,69 | 0,0 |
| ENSMUSG00000096956 | 53,86666659 | 65,6 | 0,0 | 9,425317527 | 0,011576497 | Snhg18 | small nucleolar RNA host gene 18 [Source:MGI Symbol;Acc:MGI:1914085] | #DIV/0! | #DIV/0! |
| ENSMUSG00000045867 | 40,63515029 | 64,1 | 0,0 | 9,425592119 | 0,01046065 | Cradd | CASP2 and RIPK1 domain containing adaptor with death domain [Source:MGI Symbol;Acc:MGI:1336168 | 98,67 | 0,0 |
| ENSMUSG00000031119 | 49,44126392 | 67,1 | 0,0 | 9,42672631 | 0,010472682 | Gpc4 | glypican 4 [Source:MGI Symbol;Acc:MGI:104902] | 86,38 | 0,0 |
| ENSMUSG00000074384 | 30,85251709 | 65,5 | 0,0 | 9,431517978 | 0,013056501 | AI429214 | expressed sequence AI429214 [Source:MGI Symbol;Acc:MGI:2142538] | 98,36 | 0,0 |
| ENSMUSG00000086029 | 30,99451626 | 64,4 | 0,0 | 9,435215238 | 0,011862827 | Pax6os1 | paired box 6 opposite strand 1 [Source:MGI Symbol;Acc:MGI:3028033] | 91,15 | 0,0 |
| ENSMUSG00000004633 | 41,95919775 | 65,8 | 0,0 | 9,447637141 | 0,013408895 | Chn2 | chimerin 2 [Source:MGI Symbol;Acc:MGI:1917243] | 63,68 | 0,0 |
| ENSMUSG00000045414 | 55,5980058 | 86,9 | 0,0 | 9,447822791 | 0,006835893 | 1190002N15Rik | RIKEN cDNA 1190002N15 gene [Source:MGI Symbol;Acc:MGI:1916111] | 81,74 | 0,0 |
| ENSMUSG00000032308 | 68,17676612 | 67,4 | 0,0 | 9,451422098 | 0,007871312 | Ulk3 | unc-51-like kinase 3 [Source:MGI Symbol;Acc:MGI:1918992] | 95,27 | 0,0 |
| ENSMUSG00000035274 | 56,43264285 | 66,3 | 0,0 | 9,452160205 | 0,017038985 | Tpbg | trophoblast glycoprotein [Source:MGI Symbol;Acc:MGI:1341264] | 71,21 | 0,0 |
| ENSMUSG00000079316 | 862,3735515 | 1549,2 | 144,6 | 9,45235827 | 3,2957E-05 | Rab9 | RAB9, member RAS oncogene family [Source:MGI Symbol;Acc:MGI:1890695] | 74,37 | 23,9 |
| ENSMUSG00000022893 | 61,2628318 | 90,0 | 0,0 | 9,45475465 | 0,010179082 | Adamts1 | like and metallopeptidase (reprolysin type) with thrombospondin type 1 motif, 1 [Source:MGI Symbol;Acc:I | 78,31 | 0,0 |
| ENSMUSG00000041845 | 38,68997581 | 65,8 | 0,0 | 9,454833196 | 0,01226015 | Rhod | ras homolog gene family, member D [Source:MGI Symbol;Acc:MGI:108446] | 89,46 | 0,0 |
| ENSMUSG00000030674 | 26,96025561 | 67,3 | 0,0 | 9,45483701 | 0,007814583 | Qprt | quinolinate phosphoribosyltransferase [Source:MGI Symbol;Acc:MGI:1914625] | 83,48 | 0,0 |
| ENSMUSG00000108701 | 44,61505134 | 68,1 | 0,0 | 9,458923849 | 0,00903868 | RP23-26J4.3 | | 23,56 | 0,0 |
| ENSMUSG00000022096 | 32,79402059 | 68,8 | 0,0 | 9,462574537 | 0,007775776 | Hr | hairless [Source:MGI Symbol;Acc:MGI:96223] | 64,52 | 0,0 |
| ENSMUSG00000040312 | 40,98680938 | 70,4 | 0,0 | 9,465968016 | 0,013706157 | Cchcr1 | coiled-coil alpha-helical rod protein 1 [Source:MGI Symbol;Acc:MGI:2385321] | 75,93 | 0,0 |
| ENSMUSG00000031487 | 32,53509108 | 65,9 | 0,0 | 9,46691299 | 0,009335512 | Brf2 | :F2, RNA polymerase III transcription initiation factor 50kDa subunit [Source:MGI Symbol;Acc:MGI:19139 | 95,30 | 0,0 |
| ENSMUSG00000085260 | 37,08949072 | 68,6 | 0,0 | 9,469883426 | 0,01106891 | Med9os | mediator complex subunit 9, opposite strand [Source:MGI Symbol;Acc:MGI:1916628] | 100,00 | 0,0 |
| ENSMUSG00000085928 | 27,92287895 | 66,2 | 0,0 | 9,481524461 | 0,007989752 | 4933427I22Rik | RIKEN cDNA 4933427I22 gene [Source:MGI Symbol;Acc:MGI:1918485] | #DIV/0! | #DIV/0! |
| ENSMUSG00000026389 | 34,70544813 | 68,8 | 0,0 | 9,489462638 | 0,011864079 | Steap3 | STEAP family member 3 [Source:MGI Symbol;Acc:MGI:1915678] | 45,79 | 0,0 |
| ENSMUSG00000054469 | 37,73817899 | 68,2 | 0,0 | 9,490287978 | 0,007341637 | Lclat1 | lysocardiolipin acyltransferase 1 [Source:MGI Symbol;Acc:MGI:2684937] | 89,52 | 0,0 |
| ENSMUSG00000029504 | 54,01048137 | 69,5 | 0,0 | 9,490659614 | 0,014697605 | Ddx51 | DEAD (Asp-Glu-Ala-Asp) box polypeptide 51 [Source:MGI Symbol;Acc:MGI:1916913] | 72,20 | 0,0 |
| ENSMUSG00000040483 | 39,4459835 | 69,3 | 0,0 | 9,493983376 | 0,010956967 | Xaf1 | XIAP associated factor 1 [Source:MGI Symbol;Acc:MGI:3772572] | 33,32 | 0,0 |
| ENSMUSG00000100009 | 27,96869634 | 67,0 | 0,0 | 9,496804945 | 0,00754095 | Gm7967 | predicted gene 7967 [Source:MGI Symbol;Acc:MGI:3644984] | #DIV/0! | #DIV/0! |
| ENSMUSG00000028391 | 48,1099931 | 69,2 | 0,0 | 9,506961121 | 0,00759766 | Wdr31 | WD repeat domain 31 [Source:MGI Symbol;Acc:MGI:1918604] | 88,30 | 0,0 |

| | | | | | | | | | |
|---|---|---|---|---|---|---|---|---|---|
| ENSMUSG00000040621 | 32,33856856 | 72,4 | 0,0 | 9,516379564 | 0,007320112 | Gemin8 | gem (nuclear organelle) associated protein 8 [Source:MGI Symbol;Acc:MGI:2384300] | 63,30 | 0,0 |
| ENSMUSG00000008384 | 60,93037038 | 71,2 | 0,0 | 9,51738395 | 0,010607885 | Sertad1 | SERTA domain containing 1 [Source:MGI Symbol;Acc:MGI:1913438] | 94,12 | 0,0 |
| ENSMUSG00000090564 | 62,77085505 | 70,0 | 0,0 | 9,521550035 | 0,007374158 | A430057M04Rik | RIKEN cDNA A430057M04 gene [Source:MGI Symbol;Acc:MGI:2442125] | #DIV/0! | #DIV/0! |
| ENSMUSG00000029056 | 40,47439723 | 69,4 | 0,0 | 9,521551306 | 0,006860894 | Pank4 | pantothenate kinase 4 [Source:MGI Symbol;Acc:MGI:2387466] | 93,37 | 0,0 |
| ENSMUSG00000022969 | 84,9586932 | 83,1 | 0,0 | 9,52244964 | 0,007871312 | Il10rb | interleukin 10 receptor, beta [Source:MGI Symbol;Acc:MGI:109380] | 94,40 | 0,0 |
| ENSMUSG00000021366 | 34,53416236 | 69,9 | 0,0 | 9,527833522 | 0,00697931 | Hivep1 | human immunodeficiency virus type I enhancer binding protein 1 [Source:MGI Symbol;Acc:MGI:96100] | 71,86 | 0,0 |
| ENSMUSG00000054006 | 38,25333567 | 69,6 | 0,0 | 9,536629314 | 0,012368053 | D630008O14Rik | RIKEN cDNA D630008O14 gene [Source:MGI Symbol;Acc:MGI:3698880] | 27,14 | 0,0 |
| ENSMUSG00000032469 | 45,03575481 | 70,4 | 0,0 | 9,538494133 | 0,012375469 | Dbr1 | debranching RNA lariats 1 [Source:MGI Symbol;Acc:MGI:1931520] | 85,48 | 0,0 |
| ENSMUSG00000022299 | 46,5706849 | 71,2 | 0,0 | 9,539003574 | 0,012135124 | Slc25a32 | solute carrier family 25, member 32 [Source:MGI Symbol;Acc:MGI:1917156] | 93,79 | 0,0 |
| ENSMUSG00000061544 | 35,41453058 | 68,9 | 0,0 | 9,540316203 | 0,006792606 | Zfp229 | zinc finger protein 229 [Source:MGI Symbol;Acc:MGI:2679295] | 85,29 | 0,0 |
| ENSMUSG00000039480 | 49,39851617 | 72,9 | 0,0 | 9,542156674 | 0,009755057 | Nt5dc1 | 5'-nucleotidase domain containing 1 [Source:MGI Symbol;Acc:MGI:2442446] | 88,70 | 0,0 |
| ENSMUSG00000033961 | 48,87754714 | 72,5 | 0,0 | 9,544337202 | 0,010536254 | Zfp446 | zinc finger protein 446 [Source:MGI Symbol;Acc:MGI:2442185] | 74,05 | 0,0 |
| ENSMUSG00000066235 | 34,6527837 | 69,6 | 0,0 | 9,551630184 | 0,009115328 | Pomgnt2 | otein O-linked mannose beta 1,4-N-acetylglucosaminyltransferase 2 [Source:MGI Symbol;Acc:MGI:21434 | 98,00 | 0,0 |
| ENSMUSG00000055305 | 35,91717237 | 72,5 | 0,0 | 9,554679191 | 0,007871312 | Zfp93 | zinc finger protein 93 [Source:MGI Symbol;Acc:MGI:107611] | 83,90 | 0,0 |
| ENSMUSG00000047844 | 45,89405162 | 71,4 | 0,0 | 9,560863242 | 0,008372602 | Bex4 | brain expressed gene 4 [Source:MGI Symbol;Acc:MGI:3606746] | 76,34 | 0,0 |
| ENSMUSG00000031504 | 64,28812708 | 71,3 | 0,0 | 9,561995622 | 0,016213872 | Rab20 | RAB20, member RAS oncogene family [Source:MGI Symbol;Acc:MGI:102789] | 96,54 | 0,0 |
| ENSMUSG00000097503 | 37,37259307 | 70,9 | 0,0 | 9,569176598 | 0,008268466 | 3110045C21Rik | RIKEN cDNA 3110045C21 gene [Source:MGI Symbol;Acc:MGI:1914553] | 99,82 | 0,0 |
| ENSMUSG00000062040 | 47,64329891 | 75,9 | 0,0 | 9,570148209 | 0,006519732 | Zfp27 | zinc finger protein 27 [Source:MGI Symbol;Acc:MGI:99174] | 82,58 | 0,0 |
| ENSMUSG00000035713 | 29,64232563 | 77,0 | 0,0 | 9,577543283 | 0,007625355 | Usp35 | ubiquitin specific peptidase 35 [Source:MGI Symbol;Acc:MGI:2685339] | 89,77 | 0,0 |
| ENSMUSG00000034487 | 64,86833931 | 81,5 | 0,0 | 9,58074281 | 0,010180252 | Kdelc2 | KDEL (Lys-Asp-Glu-Leu) containing 2 [Source:MGI Symbol;Acc:MGI:1923765] | 90,50 | 0,0 |
| ENSMUSG00000019826 | 39,4740166 | 80,8 | 0,0 | 9,582973513 | 0,010611249 | Zbtb24 | zinc finger and BTB domain containing 24 [Source:MGI Symbol;Acc:MGI:3039618] | 92,76 | 0,0 |
| ENSMUSG00000019461 | 85,88919058 | 75,1 | 0,0 | 9,584564114 | 0,009059372 | Plscr3 | phospholipid scramblase 3 [Source:MGI Symbol;Acc:MGI:1917560] | 67,03 | 0,0 |
| ENSMUSG00000031616 | 56,11096077 | 74,2 | 0,0 | 9,58717311 | 0,014423451 | Ednra | endothelin receptor type A [Source:MGI Symbol;Acc:MGI:105923] | 97,77 | 0,0 |
| ENSMUSG00000039164 | 72,63480866 | 74,7 | 0,0 | 9,587339599 | 0,010028792 | Naif1 | nuclear apoptosis inducing factor 1 [Source:MGI Symbol;Acc:MGI:1918504] | 66,67 | 0,0 |
| ENSMUSG00000022525 | 42,61537844 | 72,1 | 0,0 | 9,59214142 | 0,007303788 | Hrasls | HRAS-like suppressor [Source:MGI Symbol;Acc:MGI:1351473] | 92,54 | 0,0 |
| ENSMUSG00000044522 | 38,08966333 | 75,0 | 0,0 | 9,596347703 | 0,009976306 | A730020M07Rik | RIKEN cDNA A730020M07 gene [Source:MGI Symbol;Acc:MGI:2442643] | #DIV/0! | #DIV/0! |
| ENSMUSG00000060675 | 876,6093181 | 1380,5 | 244,9 | 9,599586504 | 6,00758E-05 | Pla2g16 | phospholipase A2, group XVI [Source:MGI Symbol;Acc:MGI:2179715] | 60,85 | 17,3 |
| ENSMUSG00000048332 | 467,280596 | 870,4 | 207,6 | 9,606714316 | 0,003107701 | Lhfp | lipoma HMGIC fusion partner [Source:MGI Symbol;Acc:MGI:1920048] | 99,70 | 33,0 |
| ENSMUSG00000003812 | 126,6835068 | 79,6 | 0,0 | 9,60975179 | 0,007874674 | Dnase2a | deoxyribonuclease II alpha [Source:MGI Symbol;Acc:MGI:1329019] | 76,08 | 0,0 |
| ENSMUSG00000046380 | 60,18866368 | 82,0 | 0,0 | 9,619333885 | 0,013799306 | Jrk | jerky [Source:MGI Symbol;Acc:MGI:106214] | 61,47 | 0,0 |
| ENSMUSG00000018849 | 999,007905 | 1602,2 | 347,4 | 9,62886232 | 0,000138865 | Wwc1 | WW, C2 and coiled-coil domain containing 1 [Source:MGI Symbol;Acc:MGI:2388637] | 99,93 | 31,2 |
| ENSMUSG00000031283 | 55,68076966 | 78,7 | 0,0 | 9,633598249 | 0,010197083 | Chrdl1 | chordin-like 1 [Source:MGI Symbol;Acc:MGI:1933172] | 65,84 | 0,0 |
| ENSMUSG00000066838 | 40,50308081 | 73,4 | 0,0 | 9,643020367 | 0,006489994 | Zfp772 | zinc finger protein 772 [Source:MGI Symbol;Acc:MGI:2385265] | 94,81 | 0,0 |
| ENSMUSG00000036636 | 108,7994725 | 86,1 | 0,0 | 9,64823221 | 0,011016486 | Clcn7 | chloride channel, voltage-sensitive 7 [Source:MGI Symbol;Acc:MGI:1347048] | 87,48 | 0,0 |
| ENSMUSG00000042684 | 41,39590679 | 78,2 | 0,0 | 9,64937078 | 0,006314703 | Npl | N-acetylneuraminate pyruvate lyase [Source:MGI Symbol;Acc:MGI:1921341] | 80,06 | 0,0 |
| ENSMUSG00000060579 | 44,22594549 | 75,3 | 0,0 | 9,651598729 | 0,010125358 | Fhit | fragile histidine triad gene [Source:MGI Symbol;Acc:MGI:1277947] | 85,34 | 0,0 |
| ENSMUSG00000021785 | 271,3159906 | 454,0 | 0,9 | 9,652548212 | 0,001146727 | Ngly1 | N-glycanase 1 [Source:MGI Symbol;Acc:MGI:1913276] | 97,81 | 0,9 |
| ENSMUSG00000026778 | 89,43611421 | 80,9 | 0,0 | 9,652620172 | 0,010179082 | Prkcq | protein kinase C, theta [Source:MGI Symbol;Acc:MGI:97601] | 91,07 | 0,0 |
| ENSMUSG00000031639 | 289,5384572 | 523,7 | 0,6 | 9,652978759 | 0,000253141 | Tlr3 | toll-like receptor 3 [Source:MGI Symbol;Acc:MGI:2156367] | 98,10 | 0,5 |
| ENSMUSG00000058192 | 37,47725329 | 75,5 | 0,0 | 9,654983458 | 0,010611344 | Zfp846 | zinc finger protein 846 [Source:MGI Symbol;Acc:MGI:1924012] | 93,24 | 0,0 |
| ENSMUSG00000030111 | 64,22031119 | 106,3 | 0,0 | 9,662753018 | 0,005912613 | A2m | alpha-2-macroglobulin [Source:MGI Symbol;Acc:MGI:2449119] | 76,20 | 0,0 |
| ENSMUSG00000036907 | 39,14094552 | 76,4 | 0,0 | 9,665147875 | 0,006838397 | C1ql2 | complement component 1, q subcomponent-like 2 [Source:MGI Symbol;Acc:MGI:3032521] | 99,17 | 0,0 |
| ENSMUSG00000102775 | 43,24682498 | 78,4 | 0,0 | 9,669409391 | 0,007374158 | Gm38160 | predicted gene, 38160 [Source:MGI Symbol;Acc:MGI:5611388] | 23,76 | 0,0 |
| ENSMUSG00000041684 | 69,49057685 | 77,5 | 0,0 | 9,675960087 | 0,008249347 | Bivm | basic, immunoglobulin-like variable motif containing [Source:MGI Symbol;Acc:MGI:2179809] | 85,02 | 0,0 |
| ENSMUSG00000056598 | 78,39788753 | 77,7 | 0,0 | 9,677921066 | 0,009615105 | Lrrc48 | leucine rich repeat containing 48 [Source:MGI Symbol;Acc:MGI:1921915] | 49,63 | 0,0 |
| ENSMUSG00000054708 | 43,5980235 | 77,8 | 0,0 | 9,683530042 | 0,008707404 | Ankrd24 | ankyrin repeat domain 24 [Source:MGI Symbol;Acc:MGI:1890394] | 40,71 | 0,0 |
| ENSMUSG00000038332 | 562,4530342 | 1168,5 | 3,0 | 9,6868209 | 8,36606E-06 | Sesn1 | sestrin 1 [Source:MGI Symbol;Acc:MGI:2155278] | 98,97 | 1,5 |
| ENSMUSG00000040653 | 41,4424363 | 80,0 | 0,0 | 9,68704759 | 0,010360576 | Ppp1r14c | protein phosphatase 1, regulatory (inhibitor) subunit 14c [Source:MGI Symbol;Acc:MGI:1923392] | 79,94 | 0,0 |
| ENSMUSG00000016128 | 75,23419321 | 83,8 | 0,0 | 9,689132633 | 0,010153223 | Stard13 | StAR-related lipid transfer (START) domain containing 13 [Source:MGI Symbol;Acc:MGI:2385331] | 82,67 | 0,0 |
| ENSMUSG00000071398 | 75,3783076 | 82,5 | 0,0 | 9,689648627 | 0,012689677 | 2410004P03Rik | RIKEN cDNA 2410004P03 gene [Source:MGI Symbol;Acc:MGI:1920917] | 52,08 | 0,0 |
| ENSMUSG00000062743 | 36,47670945 | 83,4 | 0,0 | 9,693916251 | 0,006284954 | Zfp677 | zinc finger protein 677 [Source:MGI Symbol;Acc:MGI:3053207] | 94,49 | 0,0 |
| ENSMUSG00000039851 | 60,50369481 | 82,7 | 0,0 | 9,698769433 | 0,015371493 | 4932438H23Rik | RIKEN cDNA 4932438H23 gene [Source:MGI Symbol;Acc:MGI:1921637] | 68,75 | 0,0 |
| ENSMUSG00000048856 | 42,38882517 | 81,6 | 0,0 | 9,700402057 | 0,010288191 | Slc25a47 | solute carrier family 25, member 47 [Source:MGI Symbol;Acc:MGI:2144766] | 94,36 | 0,0 |
| ENSMUSG00000057054 | 36,7503523 | 77,1 | 0,0 | 9,703365286 | 0,006235071 | Inca1 | inhibitor of CDK, cyclin A1 interacting protein 1 [Source:MGI Symbol;Acc:MGI:2144284] | 94,53 | 0,0 |
| ENSMUSG00000029130 | 77,14039211 | 79,8 | 0,0 | 9,704094437 | 0,01251595 | Rnf32 | ring finger protein 32 [Source:MGI Symbol;Acc:MGI:1861747] | 95,66 | 0,0 |
| ENSMUSG00000029833 | 44,85177975 | 79,2 | 0,0 | 9,709434925 | 0,005500683 | Trim24 | tripartite motif-containing 24 [Source:MGI Symbol;Acc:MGI:109275] | 82,39 | 0,0 |
| ENSMUSG00000063060 | 32,92850682 | 82,4 | 0,0 | 9,716810074 | 0,006237028 | Sox7 | SRY (sex determining region Y)-box 7 [Source:MGI Symbol;Acc:MGI:98369] | 94,06 | 0,0 |
| ENSMUSG00000032425 | 51,85056958 | 78,3 | 0,0 | 9,724893082 | 0,006824289 | Zfp949 | zinc finger protein 949 [Source:MGI Symbol;Acc:MGI:1918890] | 86,06 | 0,0 |
| ENSMUSG00000050447 | 407,1818864 | 450,9 | 1,0 | 9,728784254 | 0,00212029 | Lypd6 | LY6/PLAUR domain containing 6 [Source:MGI Symbol;Acc:MGI:2443848] | 93,22 | 0,7 |
| ENSMUSG00000045790 | 55,27308823 | 81,0 | 0,0 | 9,729917203 | 0,007527607 | Ccdc149 | coiled-coil domain containing 149 [Source:MGI Symbol;Acc:MGI:2685293] | 97,74 | 0,0 |
| ENSMUSG00000070002 | 73,17792668 | 86,3 | 0,0 | 9,730220062 | 0,014784801 | Ell | elongation factor RNA polymerase II [Source:MGI Symbol;Acc:MGI:109377] | 94,47 | 0,0 |
| ENSMUSG00000070306 | 44,76948118 | 80,5 | 0,0 | 9,732219553 | 0,00903868 | Ccdc153 | coiled-coil domain containing 153 [Source:MGI Symbol;Acc:MGI:2448587] | 99,39 | 0,0 |
| ENSMUSG00000044037 | 56,58493726 | 90,3 | 0,0 | 9,7333376 | 0,010948548 | Als2cl | ALS2 C-terminal like [Source:MGI Symbol;Acc:MGI:2447532] | 86,05 | 0,0 |
| ENSMUSG00000015202 | 39,12910196 | 79,6 | 0,0 | 9,736593899 | 0,006007756 | Cnksr3 | Cnksr family member 3 [Source:MGI Symbol;Acc:MGI:2674130] | 87,03 | 0,0 |
| ENSMUSG00000050132 | 49,80203599 | 80,7 | 0,0 | 9,739635313 | 0,007234206 | Sarm1 | sterile alpha and HEAT/Armadillo motif containing 1 [Source:MGI Symbol;Acc:MGI:2136419] | 80,91 | 0,0 |
| ENSMUSG00000036111 | 65,67970899 | 80,5 | 0,0 | 9,747631893 | 0,007839518 | Lmo1 | LIM domain only 1 [Source:MGI Symbol;Acc:MGI:102812] | 78,54 | 0,0 |
| ENSMUSG00000046731 | 52,54040713 | 82,9 | 0,0 | 9,755430386 | 0,008933079 | Kctd11 | potassium channel tetramerisation domain containing 11 [Source:MGI Symbol;Acc:MGI:2448712] | 85,04 | 0,0 |
| ENSMUSG00000019232 | 698,7722339 | 1033,1 | 105,8 | 9,755746558 | 0,000377106 | Etnppl | ethanolamine phosphate phospholyase [Source:MGI Symbol;Acc:MGI:1919010] | 97,24 | 4,7 |
| ENSMUSG00000021823 | 229,8978689 | 505,0 | 0,6 | 9,759854 | 0,0001986 | Vcl | vinculin [Source:MGI Symbol;Acc:MGI:98927] | 99,56 | 0,3 |
| ENSMUSG00000037108 | 35,34114575 | 80,6 | 0,0 | 9,763257343 | 0,005803266 | Zcwpw1 | zinc finger, CW type with PWWP domain 1 [Source:MGI Symbol;Acc:MGI:2685899] | 98,71 | 0,0 |
| ENSMUSG00000052676 | 68,55648873 | 87,6 | 0,0 | 9,765073682 | 0,007768705 | Zmat1 | zinc finger, matrin type 1 [Source:MGI Symbol;Acc:MGI:2442284] | 90,81 | 0,0 |
| ENSMUSG00000023224 | 73,53170958 | 85,2 | 0,0 | 9,77034077 | 0,008966028 | Serping1 | serine (or cysteine) peptidase inhibitor, clade G, member 1 [Source:MGI Symbol;Acc:MGI:894696] | 86,08 | 0,0 |
| ENSMUSG00000037259 | 59,57366149 | 88,9 | 0,0 | 9,773710424 | 0,009076555 | Dzank1 | double zinc ribbon and ankyrin repeat domains 1 [Source:MGI Symbol;Acc:MGI:2139080] | 77,57 | 0,0 |
| ENSMUSG00000008167 | 101,5716721 | 83,0 | 0,0 | 9,775443367 | 0,013518196 | Fbxw9 | F-box and WD-40 domain protein 9 [Source:MGI Symbol;Acc:MGI:1915878] | 96,57 | 0,0 |
| ENSMUSG00000097340 | 67,72755621 | 87,6 | 0,0 | 9,775642635 | 0,014695662 | Gm26617 | predicted gene, 26617 [Source:MGI Symbol;Acc:MGI:5477111] | #DIV/0! | #DIV/0! |
| ENSMUSG00000023143 | 58,6666804 | 84,2 | 0,0 | 9,778798232 | 0,007004937 | Nagpa | acetylglucosamine-1-phosphodiester alpha-N-acetylglucosaminidase [Source:MGI Symbol;Acc:MGI:13515 | 98,84 | 0,0 |
| ENSMUSG00000016239 | 42,5852998 | 84,6 | 0,0 | 9,780744686 | 0,010159129 | Lonrf3 | LON peptidase N-terminal domain and ring finger 3 [Source:MGI Symbol;Acc:MGI:1921615] | 76,58 | 0,0 |
| ENSMUSG00000020491 | 63,57213036 | 84,4 | 0,0 | 9,783812346 | 0,008366913 | 2810021J22Rik | RIKEN cDNA 2810021J22 gene [Source:MGI Symbol;Acc:MGI:1917194] | 92,29 | 0,0 |
| ENSMUSG00000043885 | 101,5958245 | 85,5 | 0,0 | 9,783943431 | 0,008541677 | Slc36a4 | solute carrier family 36 (proton/amino acid symporter), member 4 [Source:MGI Symbol;Acc:MGI:2442595] | 90,52 | 0,0 |
| ENSMUSG00000019789 | 62,45200316 | 82,8 | 0,0 | 9,788218536 | 0,010179082 | Hey2 | hairy/enhancer-of-split related with YRPW motif 2 [Source:MGI Symbol;Acc:MGI:1341884] | 95,25 | 0,0 |
| ENSMUSG00000021594 | 48,48358988 | 86,0 | 0,0 | 9,790885141 | 0,006498751 | Srd5a1 | steroid 5 alpha-reductase 1 [Source:MGI Symbol;Acc:MGI:98400] | 97,28 | 0,0 |
| ENSMUSG00000020669 | 38,79771288 | 82,4 | 0,0 | 9,795738085 | 0,005628623 | Sh3yl1 | Sh3 domain YSC-like 1 [Source:MGI Symbol;Acc:MGI:1346118] | 98,90 | 0,0 |
| ENSMUSG00000027887 | 53,46386905 | 83,8 | 0,0 | 9,804271807 | 0,00759766 | Sypl2 | synaptophysin-like 2 [Source:MGI Symbol;Acc:MGI:1328311] | 75,19 | 0,0 |
| ENSMUSG00000034575 | 55,68702093 | 84,0 | 0,0 | 9,809410445 | 0,008348916 | Papd7 | PAP associated domain containing 7 [Source:MGI Symbol;Acc:MGI:2682295] | 84,39 | 0,0 |
| ENSMUSG00000032372 | 43,76623119 | 84,2 | 0,0 | 9,812015887 | 0,00562029 | Plscr2 | phospholipid scramblase 2 [Source:MGI Symbol;Acc:MGI:1270860] | 68,36 | 0,0 |
| ENSMUSG00000026514 | 53,49430551 | 85,0 | 0,0 | 9,818609055 | 0,008345158 | Cnih3 | cornichon family AMPA receptor auxiliary protein 3 [Source:MGI Symbol;Acc:MGI:1920228] | 95,73 | 0,0 |
| ENSMUSG00000024970 | 42,99733494 | 85,8 | 0,0 | 9,825011336 | 0,008082239 | AI846148 | expressed sequence AI846148 [Source:MGI Symbol;Acc:MGI:2147611] | 80,13 | 0,0 |
| ENSMUSG00000029757 | 57,07597682 | 89,2 | 0,0 | 9,825078559 | 0,006499377 | Dync1i1 | dynein cytoplasmic 1 intermediate chain 1 [Source:MGI Symbol;Acc:MGI:107743] | 76,24 | 0,0 |
| ENSMUSG00000062127 | 53,06391775 | 93,3 | 0,0 | 9,835219027 | 0,005911767 | Cttnbp2nl | CTTNBP2 N-terminal like [Source:MGI Symbol;Acc:MGI:1933137] | 86,87 | 0,0 |
| ENSMUSG00000068522 | 56,57169587 | 86,3 | 0,0 | 9,842814029 | 0,010973739 | Aard | alanine and arginine rich domain containing protein [Source:MGI Symbol;Acc:MGI:2181621] | 95,94 | 0,0 |
| ENSMUSG00000062944 | 70,34936569 | 87,3 | 0,0 | 9,845623967 | 0,007384703 | 9130023H24Rik | RIKEN cDNA 9130023H24 gene [Source:MGI Symbol;Acc:MGI:2442738] | 98,47 | 0,0 |
| ENSMUSG00000002012 | 63,35591049 | 87,5 | 0,0 | 9,846528037 | 0,010159129 | Pnck | pregnancy upregulated non-ubiquitously expressed CaM kinase [Source:MGI Symbol;Acc:MGI:1347357] | 84,73 | 0,0 |
| ENSMUSG00000036972 | 60,90168887 | 93,3 | 0,0 | 9,854342706 | 0,006629959 | Zic4 | zinc finger protein of the cerebellum 4 [Source:MGI Symbol;Acc:MGI:107201] | 69,22 | 0,0 |
| ENSMUSG00000046204 | 50,90742898 | 87,3 | 0,0 | 9,85491284 | 0,008403466 | Pnma2 | paraneoplastic antigen MA2 [Source:MGI Symbol;Acc:MGI:2444129] | 89,02 | 0,0 |
| ENSMUSG00000029569 | 55,45871543 | 90,2 | 0,0 | 9,861028193 | 0,007768705 | Tmem168 | transmembrane protein 168 [Source:MGI Symbol;Acc:MGI:1921794] | 87,80 | 0,0 |
| ENSMUSG00000027074 | 29,00593419 | 86,8 | 0,0 | 9,861433408 | 0,005639364 | Slc43a3 | solute carrier family 43, member 3 [Source:MGI Symbol;Acc:MGI:1931054] | 93,84 | 0,0 |
| ENSMUSG00000047996 | 35,43293766 | 91,9 | 0,0 | 9,868026668 | 0,005487001 | Prrg1 | proline rich Gla (G-carboxyglutamic acid) 1 [Source:MGI Symbol;Acc:MGI:1917364] | 91,08 | 0,0 |
| ENSMUSG00000031379 | 58,97710632 | 93,6 | 0,0 | 9,872185442 | 0,008345158 | Pir | pirin [Source:MGI Symbol;Acc:MGI:1916906] | 85,55 | 0,0 |
| ENSMUSG00000027635 | 87,22983943 | 93,8 | 0,0 | 9,872220298 | 0,005963779 | Dsn1 | DSN1 homolog, MIS12 kinetochore complex component [Source:MGI Symbol;Acc:MGI:1914184] | 87,48 | 0,0 |
| ENSMUSG00000069208 | 59,42488976 | 93,8 | 0,0 | 9,878544621 | 0,008798314 | Zfp825 | zinc finger protein 825 [Source:MGI Symbol;Acc:MGI:2385315] | 97,49 | 0,0 |
| ENSMUSG00000014361 | 362,5514549 | 536,3 | 0,9 | 9,884057267 | 0,000939386 | Mertk | c-mer proto-oncogene tyrosine kinase [Source:MGI Symbol;Acc:MGI:96965] | 100,00 | 0,5 |
| ENSMUSG00000024528 | 87,81437272 | 91,3 | 0,0 | 9,885825608 | 0,009025989 | Srfbp1 | serum response factor binding protein 1 [Source:MGI Symbol;Acc:MGI:1914472] | 100,00 | 0,0 |
| ENSMUSG00000031161 | 70,74519847 | 89,6 | 0,0 | 9,888841653 | 0,00992454 | Hdac6 | histone deacetylase 6 [Source:MGI Symbol;Acc:MGI:1333752] | 86,60 | 0,0 |
| ENSMUSG00000045932 | 50,13855168 | 98,5 | 0,0 | 9,897560865 | 0,005195556 | Ifit2 | interferon-induced protein with tetratricopeptide repeats 2 [Source:MGI Symbol;Acc:MGI:99449] | 92,96 | 0,0 |
| ENSMUSG00000036062 | 44,79374807 | 95,2 | 0,0 | 9,898324111 | 0,007872182 | Phf24 | PHD finger protein 24 [Source:MGI Symbol;Acc:MGI:2140712] | 75,48 | 0,0 |
| ENSMUSG00000049493 | 99,28046865 | 91,6 | 0,0 | 9,908088439 | 0,009281644 | Pls1 | plastin 1 (I-isoform) [Source:MGI Symbol;Acc:MGI:104809] | 93,12 | 0,0 |
| ENSMUSG00000069227 | 42,03207213 | 89,9 | 0,0 | 9,934883703 | 0,00511186 | Gprin1 | G protein-regulated inducer of neurite outgrowth 1 [Source:MGI Symbol;Acc:MGI:1349455] | 91,57 | 0,0 |
| ENSMUSG00000019917 | 96,54836261 | 93,3 | 0,0 | 9,936564867 | 0,009843736 | Sept10 | septin 10 [Source:MGI Symbol;Acc:MGI:1918110] | 87,40 | 0,0 |
| ENSMUSG00000048439 | 61,96849802 | 94,1 | 0,0 | 9,937038927 | 0,010334871 | Nupl2 | nucleoporin like 2 [Source:MGI Symbol;Acc:MGI:2387631] | 96,41 | 0,0 |
| ENSMUSG00000040978 | 52,00850031 | 94,6 | 0,0 | 9,944291786 | 0,005428678 | Gm11992 | predicted gene 11992 [Source:MGI Symbol;Acc:MGI:3651127] | 96,11 | 0,0 |
| ENSMUSG00000041205 | 122,7418793 | 109,6 | 0,0 | 9,949536739 | 0,009369036 | Map6d1 | MAP6 domain containing 1 [Source:MGI Symbol;Acc:MGI:3607784] | 94,81 | 0,0 |
| ENSMUSG00000038080 | 64,90672003 | 93,5 | 0,0 | 9,951800206 | 0,006792606 | Kdm1b | lysine (K)-specific demethylase 1B [Source:MGI Symbol;Acc:MGI:2145261] | 93,74 | 0,0 |
| ENSMUSG00000051373 | 78,69454864 | 110,5 | 0,0 | 9,954594951 | 0,009369036 | Plpp7 | phospholipid phosphatase 7 (inactive) [Source:MGI Symbol;Acc:MGI:2445183] | 93,01 | 0,0 |
| ENSMUSG00000092416 | 33,86767254 | 98,3 | 0,0 | 9,955530123 | 0,006658752 | Zfp141 | zinc finger protein 141 [Source:MGI Symbol;Acc:MGI:3584269] | 91,85 | 0,0 |
| ENSMUSG00000027489 | 49,18954966 | 92,2 | 0,0 | 9,957738882 | 0,009049788 | Necab3 | N-terminal EF-hand calcium binding protein 3 [Source:MGI Symbol;Acc:MGI:1861721] | 98,98 | 0,0 |
| ENSMUSG00000021596 | 36,90983958 | 98,4 | 0,0 | 9,962237074 | 0,006409393 | Mctp1 | multiple C2 domains, transmembrane 1 [Source:MGI Symbol;Acc:MGI:1926021] | 64,03 | 0,0 |
| ENSMUSG00000008999 | 87,76926704 | 105,5 | 0,0 | 9,963324928 | 0,009970368 | Bmp7 | bone morphogenetic protein 7 [Source:MGI Symbol;Acc:MGI:103302] | 90,74 | 0,0 |
| ENSMUSG00000097728 | 58,0494777 | 108,1 | 0,0 | 9,966686834 | 0,009208575 | Gm26840 | predicted gene, 26840 [Source:MGI Symbol;Acc:MGI:5477334] | #DIV/0! | #DIV/0! |
| ENSMUSG00000040113 | 41,449095 | 93,4 | 0,0 | 9,967078601 | 0,004999214 | Mettl11b | methyltransferase like 11B [Source:MGI Symbol;Acc:MGI:2685053] | 99,74 | 0,0 |
| ENSMUSG00000045180 | 90,17323566 | 108,3 | 0,0 | 9,970243099 | 0,008082239 | Shroom2 | shroom family member 2 [Source:MGI Symbol;Acc:MGI:107194] | 64,96 | 0,0 |
| ENSMUSG00000010175 | 45,91650256 | 102,2 | 0,0 | 9,977613803 | 0,005844961 | Prox1 | prospero homeobox 1 [Source:MGI Symbol;Acc:MGI:97772] | 76,56 | 0,0 |
| ENSMUSG00000036686 | 84,40957119 | 100,5 | 0,0 | 9,977954771 | 0,005975581 | Cc2d1a | coiled-coil and C2 domain containing 1A [Source:MGI Symbol;Acc:MGI:2384831] | 94,07 | 0,0 |
| ENSMUSG00000030657 | 74,07899969 | 105,7 | 0,0 | 9,987148488 | 0,006918725 | Xylt1 | xylosyltransferase 1 [Source:MGI Symbol;Acc:MGI:2451073] | 77,33 | 0,0 |
| ENSMUSG00000035547 | 49,91343674 | 103,6 | 0,0 | 9,993722521 | 0,006314703 | Capn5 | calpain 5 [Source:MGI Symbol;Acc:MGI:1100859] | 84,23 | 0,0 |
| ENSMUSG00000018648 | 78,0896456 | 102,2 | 0,0 | 9,99376423 | 0,008133646 | Dusp14 | dual specificity phosphatase 14 [Source:MGI Symbol;Acc:MGI:1927168] | 75,05 | 0,0 |
| ENSMUSG00000055660 | 73,37262297 | 98,0 | 0,0 | 10,00130162 | 0,00542125 | Mettl4 | methyltransferase like 4 [Source:MGI Symbol;Acc:MGI:1924031] | 99,86 | 0,0 |
| ENSMUSG00000106019 | 65,32138644 | 98,0 | 0,0 | 10,00468652 | 0,009805025 | Gm43672 | predicted gene 43672 [Source:MGI Symbol;Acc:MGI:5663809] | #DIV/0! | #DIV/0! |
| ENSMUSG00000068457 | 59,12176729 | 97,1 | 0,0 | 10,00878251 | 0,008334437 | Uty | ubiquitously transcribed tetratricopeptide repeat gene, Y chromosome [Source:MGI Symbol;Acc:MGI:89481 | 87,11 | 0,0 |
| ENSMUSG00000054519 | 74,81074874 | 101,4 | 0,0 | 10,01057256 | 0,007374158 | Zfp867 | zinc finger protein 867 [Source:MGI Symbol;Acc:MGI:2681848] | 88,80 | 0,0 |
| ENSMUSG00000051316 | 60,29514877 | 98,4 | 0,0 | 10,01533723 | 0,005369631 | Taf7 | RNA polymerase II, TATA box binding protein (TBP)-associated factor [Source:MGI Symbol;Acc:MGI:134 | 60,94 | 0,0 |
| ENSMUSG00000073910 | 95,00546086 | 101,8 | 0,0 | 10,02258369 | 0,007640988 | Mob3b | MOB kinase activator 3B [Source:MGI Symbol;Acc:MGI:2664539] | 88,47 | 0,0 |
| ENSMUSG00000054021 | 68,80408108 | 98,4 | 0,0 | 10,02318183 | 0,007603603 | Sirt5 | sirtuin 5 [Source:MGI Symbol;Acc:MGI:1915596] | 99,78 | 0,0 |
| ENSMUSG00000032417 | 46,94413323 | 101,2 | 0,0 | 10,02469936 | 0,004763065 | Rwdd2a | RWD domain containing 2A [Source:MGI Symbol;Acc:MGI:1916769] | 97,44 | 0,0 |
| ENSMUSG00000042340 | 79,1339446 | 98,0 | 0,0 | 10,0276447 | 0,00685741 | Ctf1 | cardiotrophin 1 [Source:MGI Symbol;Acc:MGI:105115] | 94,94 | 0,0 |
| ENSMUSG00000070643 | 49,44550277 | 99,2 | 0,0 | 10,02798444 | 0,004133532 | Sox13 | SRY (sex determining region Y)-box 13 [Source:MGI Symbol;Acc:MGI:98361] | 94,48 | 0,0 |
| ENSMUSG00000097440 | 47,14416026 | 99,9 | 0,0 | 10,03595295 | 0,006918725 | Gm6277 | predicted gene 6277 [Source:MGI Symbol;Acc:MGI:3779583] | #DIV/0! | #DIV/0! |
| ENSMUSG00000025857 | 75,4215224 | 102,4 | 0,0 | 10,03756053 | 0,010268946 | Dnaaf5 | dynein, axonemal assembly factor 5 [Source:MGI Symbol;Acc:MGI:3616079] | 96,04 | 0,0 |
| ENSMUSG00000055541 | 51,83540632 | 99,7 | 0,0 | 10,04404476 | 0,007516444 | Lair1 | leukocyte-associated Ig-like receptor 1 [Source:MGI Symbol;Acc:MGI:105492] | 46,35 | 0,0 |
| ENSMUSG00000034845 | 60,54544292 | 100,4 | 0,0 | 10,04833922 | 0,007671013 | Plvap | plasmalemma vesicle associated protein [Source:MGI Symbol;Acc:MGI:1890497] | 60,51 | 0,0 |
| ENSMUSG00000045980 | 51,75768278 | 100,3 | 0,0 | 10,05020438 | 0,006899501 | Tmem104 | transmembrane protein 104 [Source:MGI Symbol;Acc:MGI:2444222] | 93,54 | 0,0 |
| ENSMUSG00000086629 | 57,23288953 | 101,6 | 0,0 | 10,05185619 | 0,008541452 | 2810403D21Rik | RIKEN cDNA 2810403D21 gene [Source:MGI Symbol;Acc:MGI:1917214] | #DIV/0! | #DIV/0! |
| ENSMUSG00000055235 | 88,87298027 | 114,0 | 0,0 | 10,05762703 | 0,006624688 | Wdr86 | WD repeat domain 86 [Source:MGI Symbol;Acc:MGI:1915466] | 93,41 | 0,0 |
| ENSMUSG00000029086 | 72,11215536 | 101,7 | 0,0 | 10,0649796 | 0,006668962 | Prom1 | prominin 1 [Source:MGI Symbol;Acc:MGI:1100886] | 72,23 | 0,0 |
| ENSMUSG00000002908 | 61,96731425 | 103,8 | 0,0 | 10,06560052 | 0,009417009 | Kcnn1 | rmediate/small conductance calcium-activated channel, subfamily N, member 1 [Source:MGI Symbol;Acc: | 71,13 | 0,0 |
| ENSMUSG00000032531 | 71,18164933 | 101,5 | 0,0 | 10,07153727 | 0,007070948 | Amotl2 | angiomotin-like 2 [Source:MGI Symbol;Acc:MGI:1929286] | 78,23 | 0,0 |
| ENSMUSG00000085314 | 46,88091995 | 108,0 | 0,0 | 10,07373044 | 0,006961964 | Gm14866 | predicted gene 14866 [Source:MGI Symbol;Acc:MGI:3704514] | #DIV/0! | #DIV/0! |
| ENSMUSG00000044033 | 70,16767037 | 110,3 | 0,0 | 10,07746892 | 0,004252527 | Ccdc141 | coiled-coil domain containing 141 [Source:MGI Symbol;Acc:MGI:1919735] | 61,13 | 0,0 |
| ENSMUSG00000035277 | 60,99750543 | 104,1 | 0,0 | 10,08092994 | 0,007666043 | Arx | aristaless related homeobox [Source:MGI Symbol;Acc:MGI:1097716] | 95,36 | 0,0 |
| ENSMUSG00000029816 | 64,85831357 | 108,3 | 0,0 | 10,08415913 | 0,011322754 | Gpnmb | glycoprotein (transmembrane) nmb [Source:MGI Symbol;Acc:MGI:1934765] | 58,99 | 0,0 |
| ENSMUSG00000097333 | 72,58569514 | 105,3 | 0,0 | 10,09575992 | 0,006884094 | Zfp87 | zinc finger protein 87 [Source:MGI Symbol;Acc:MGI:107768] | 95,75 | 0,0 |
| ENSMUSG00000027655 | 110,3833075 | 109,6 | 0,0 | 10,10321168 | 0,006282973 | Dhx35 | DEAH (Asp-Glu-Ala-His) box polypeptide 35 [Source:MGI Symbol;Acc:MGI:1918965] | 95,65 | 0,0 |
| ENSMUSG00000030087 | 785,9506496 | 1410,8 | 239,5 | 10,13344809 | 0,000145766 | Klf15 | Kruppel-like factor 15 [Source:MGI Symbol;Acc:MGI:1929988] | 91,39 | 26,9 |
| ENSMUSG00000060639 | 48,85874485 | 108,1 | 0,0 | 10,13801141 | 0,005859943 | Hist1h4i | histone cluster 1, H4i [Source:MGI Symbol;Acc:MGI:2448432] | 89,68 | 0,0 |
| ENSMUSG00000049811 | 69,53995829 | 112,7 | 0,0 | 10,13955929 | 0,005837578 | Fam161a | family with sequence similarity 161, member A [Source:MGI Symbol;Acc:MGI:1921123] | 81,89 | 0,0 |
| ENSMUSG00000048222 | 67,40432471 | 107,3 | 0,0 | 10,14010249 | 0,006434401 | Mfap1b | microfibrillar-associated protein 1B [Source:MGI Symbol;Acc:MGI:3694697] | 41,48 | 0,0 |
| ENSMUSG00000040990 | 97,31957335 | 113,6 | 0,0 | 10,1418499 | 0,004277829 | Sh3kbp1 | SH3-domain kinase binding protein 1 [Source:MGI Symbol;Acc:MGI:1889583] | 53,07 | 0,0 |
| ENSMUSG00000009828 | 78,37434864 | 108,8 | 0,0 | 10,14335506 | 0,006374552 | Ick | intestinal cell kinase [Source:MGI Symbol;Acc:MGI:1934157] | 43,19 | 0,0 |
| ENSMUSG00000041000 | 57,73383001 | 108,8 | 0,0 | 10,14687682 | 0,003904395 | Trim62 | tripartite motif-containing 62 [Source:MGI Symbol;Acc:MGI:1914775] | 96,02 | 0,0 |
| ENSMUSG00000028383 | 984,5809727 | 1462,6 | 421,8 | 10,15159043 | 0,001354712 | Hsdl2 | hydroxysteroid dehydrogenase like 2 [Source:MGI Symbol;Acc:MGI:1919729] | 93,73 | 29,9 |
| ENSMUSG00000027015 | 104,9487311 | 108,1 | 0,0 | 10,15360879 | 0,01046065 | Cybrd1 | cytochrome b reductase 1 [Source:MGI Symbol;Acc:MGI:2654575] | 91,33 | 0,0 |
| ENSMUSG00000017978 | 56,06712493 | 120,1 | 0,0 | 10,15715691 | 0,006156797 | Cadps2 | Ca2+-dependent activator protein for secretion 2 [Source:MGI Symbol;Acc:MGI:2443963] | 67,65 | 0,0 |
| ENSMUSG00000022724 | 88,20825906 | 114,9 | 0,0 | 10,16156497 | 0,004271538 | Mina | myc induced nuclear antigen [Source:MGI Symbol;Acc:MGI:1914264] | 51,93 | 0,0 |
| ENSMUSG00000022749 | 70,80326936 | 107,6 | 0,0 | 10,16223722 | 0,005058806 | Tbc1d23 | TBC1 domain family, member 23 [Source:MGI Symbol;Acc:MGI:1914831] | 97,43 | 0,0 |
| ENSMUSG00000035069 | 99,02518597 | 110,1 | 0,0 | 10,17128916 | 0,006895432 | Oma1 | OMA1 homolog, zinc metallopeptidase (S. cerevisiae) [Source:MGI Symbol;Acc:MGI:1914263] | 98,59 | 0,0 |
| ENSMUSG00000034430 | 103,8702729 | 114,5 | 0,0 | 10,17188166 | 0,007209328 | Zxdc | ZXD family zinc finger C [Source:MGI Symbol;Acc:MGI:1933108] | 79,03 | 0,0 |
| ENSMUSG00000046312 | 407,8900163 | 735,0 | 0,9 | 10,17979973 | 8,88537E-05 | AI464131 | expressed sequence AI464131 [Source:MGI Symbol;Acc:MGI:2140300] | 93,83 | 0,5 |
| ENSMUSG00000020467 | 72,1111541 | 125,7 | 0,0 | 10,18344188 | 0,006198296 | Efemp1 | ermal growth factor-containing fibulin-like extracellular matrix protein 1 [Source:MGI Symbol;Acc:MGI:1339 | 97,73 | 0,0 |
| ENSMUSG00000105957 | 70,30956366 | 112,0 | 0,0 | 10,18388582 | 0,010763569 | Gm43254 | predicted gene 43254 [Source:MGI Symbol;Acc:MGI:5663391] | #DIV/0! | #DIV/0! |
| ENSMUSG00000047216 | 128,4012368 | 110,0 | 0,0 | 10,18854984 | 0,010334871 | Cdh19 | cadherin 19, type 2 [Source:MGI Symbol;Acc:MGI:3588198] | 97,83 | 0,0 |
| ENSMUSG00000072294 | 57,12156505 | 108,3 | 0,0 | 10,19322298 | 0,004023292 | Klf12 | Kruppel-like factor 12 [Source:MGI Symbol;Acc:MGI:1333796] | 99,84 | 0,0 |

| | | | | | | | | | |
|---|---|---|---|---|---|---|---|---|---|
| ENSMUSG00000023348 | 63,55504746 | 109,2 | 0,0 | 10,20281205 | 0,003947394 | Trip6 | thyroid hormone receptor interactor 6 [Source:MGI Symbol;Acc:MGI:1343458] | 81,61 | 0,0 |
| ENSMUSG00000020525 | 181,0750682 | 113,1 | 0,0 | 10,20404878 | 0,004932028 | Ppm1d | protein phosphatase 1D magnesium-dependent, delta isoform [Source:MGI Symbol;Acc:MGI:1858214] | 88,74 | 0,0 |
| ENSMUSG00000048706 | 63,71186421 | 119,1 | 0,0 | 10,20654001 | 0,007251624 | Lurap1l | leucine rich adaptor protein 1-like [Source:MGI Symbol;Acc:MGI:106510] | 75,80 | 0,0 |
| ENSMUSG00000057234 | 56,68300831 | 113,2 | 0,0 | 10,20896221 | 0,004763065 | Mettl15 | methyltransferase like 15 [Source:MGI Symbol;Acc:MGI:1924144] | 95,49 | 0,0 |
| ENSMUSG00000039004 | 142,6137836 | 127,7 | 0,0 | 10,22384241 | 0,011589432 | Bmp6 | bone morphogenetic protein 6 [Source:MGI Symbol;Acc:MGI:88182] | 97,51 | 0,0 |
| ENSMUSG00000050875 | 72,71906228 | 113,2 | 0,0 | 10,22617 | 0,008035114 | A730017C20Rik | RIKEN cDNA A730017C20 gene [Source:MGI Symbol;Acc:MGI:2442934] | 90,12 | 0,0 |
| ENSMUSG00000034216 | 51,32932058 | 117,1 | 0,0 | 10,22852846 | 0,004166854 | Vps18 | vacuolar protein sorting 18 (yeast) [Source:MGI Symbol;Acc:MGI:2443626] | 94,91 | 0,0 |
| ENSMUSG00000078234 | 59,23167039 | 111,3 | 0,0 | 10,23213418 | 0,005170315 | Klhdc7a | kelch domain containing 7A [Source:MGI Symbol;Acc:MGI:2444612] | 75,94 | 0,0 |
| ENSMUSG00000024766 | 57,4485338 | 111,7 | 0,0 | 10,23722586 | 0,004139713 | Lipo1 | lipase, member O1 [Source:MGI Symbol;Acc:MGI:2147592] | 68,51 | 0,0 |
| ENSMUSG00000043391 | 581,8867587 | 876,7 | 1,0 | 10,24725866 | 1,96332E-05 | 2510009E07Rik | RIKEN cDNA 2510009E07 gene [Source:MGI Symbol;Acc:MGI:1919440] | 99,61 | 0,5 |
| ENSMUSG00000026102 | 145,0354343 | 124,4 | 0,0 | 10,250809 | 0,009728372 | Inpp1 | inositol polyphosphate-1-phosphatase [Source:MGI Symbol;Acc:MGI:104848] | 64,30 | 0,0 |
| ENSMUSG00000020707 | 87,15664376 | 120,7 | 0,0 | 10,25154473 | 0,010956967 | Rnf135 | ring finger protein 135 [Source:MGI Symbol;Acc:MGI:1919206] | 98,62 | 0,0 |
| ENSMUSG00000028789 | 83,14902913 | 116,4 | 0,0 | 10,26063339 | 0,005326329 | Azin2 | antizyme inhibitor 2 [Source:MGI Symbol;Acc:MGI:2442093] | 77,20 | 0,0 |
| ENSMUSG00000042389 | 92,54633419 | 117,9 | 0,0 | 10,26359112 | 0,004505742 | Tsen2 | tRNA splicing endonuclease 2 homolog (S. cerevisiae) [Source:MGI Symbol;Acc:MGI:2141599] | 83,06 | 0,0 |
| ENSMUSG00000067430 | 73,14182744 | 118,2 | 0,0 | 10,27199823 | 0,00759766 | Zfp763 | zinc finger protein 763 [Source:MGI Symbol;Acc:MGI:1920701] | 98,76 | 0,0 |
| ENSMUSG00000022008 | 79,02808841 | 119,9 | 0,0 | 10,2755704 | 0,005499898 | Gpalpp1 | GPALPP motifs containing 1 [Source:MGI Symbol;Acc:MGI:1914717] | 95,20 | 0,0 |
| ENSMUSG00000027111 | 85,44243428 | 126,1 | 0,0 | 10,27741959 | 0,006566597 | Itga6 | integrin alpha 6 [Source:MGI Symbol;Acc:MGI:96605] | 67,57 | 0,0 |
| ENSMUSG00000039765 | 56,78980491 | 118,8 | 0,0 | 10,27823866 | 0,004999214 | Cc2d2a | coiled-coil and C2 domain containing 2A [Source:MGI Symbol;Acc:MGI:1924487] | 91,32 | 0,0 |
| ENSMUSG00000037709 | 59,82068035 | 125,4 | 0,0 | 10,28282959 | 0,003602277 | Fam13a | family with sequence similarity 13, member A [Source:MGI Symbol;Acc:MGI:1889842] | 78,36 | 0,0 |
| ENSMUSG00000031441 | 79,9978675 | 120,2 | 0,0 | 10,28726523 | 0,005886206 | Atp11a | ATPase, class VI, type 11A [Source:MGI Symbol;Acc:MGI:1354735] | 81,40 | 0,0 |
| ENSMUSG00000037664 | 141,4608258 | 152,5 | 0,0 | 10,30435683 | 0,006237028 | Cdkn1c | cyclin-dependent kinase inhibitor 1C (P57) [Source:MGI Symbol;Acc:MGI:104564] | 94,24 | 0,0 |
| ENSMUSG00000040181 | 79,85283924 | 119,7 | 0,0 | 10,30832574 | 0,007234206 | Fmo1 | flavin containing monooxygenase 1 [Source:MGI Symbol;Acc:MGI:1310002] | 65,59 | 0,0 |
| ENSMUSG00000004815 | 90,27726 | 121,7 | 0,0 | 10,31832422 | 0,003586342 | Dgkq | diacylglycerol kinase, theta [Source:MGI Symbol;Acc:MGI:102918] | 88,80 | 0,0 |
| ENSMUSG00000037085 | 73,98386336 | 120,5 | 0,0 | 10,31915868 | 0,00713294 | Trmt12 | tRNA methyltranferase 12 [Source:MGI Symbol;Acc:MGI:1915510] | 58,69 | 0,0 |
| ENSMUSG00000030093 | 92,47687164 | 126,2 | 0,0 | 10,32145133 | 0,008531851 | Wnt7a | wingless-type MMTV integration site family, member 7A [Source:MGI Symbol;Acc:MGI:98961] | 98,43 | 0,0 |
| ENSMUSG00000006764 | 55,67559973 | 122,6 | 0,0 | 10,32920195 | 0,00578879 | Tph2 | tryptophan hydroxylase 2 [Source:MGI Symbol;Acc:MGI:2651811] | 71,87 | 0,0 |
| ENSMUSG00000026586 | 59,38053993 | 121,9 | 0,0 | 10,33535043 | 0,003720993 | Prrx1 | paired related homeobox 1 [Source:MGI Symbol;Acc:MGI:97712] | 64,36 | 0,0 |
| ENSMUSG00000041633 | 57,9771706 | 123,3 | 0,0 | 10,33574414 | 0,003568742 | Kctd12b | potassium channel tetramerisation domain containing 12b [Source:MGI Symbol;Acc:MGI:2444667] | 93,20 | 0,0 |
| ENSMUSG00000027784 | 59,00019006 | 124,2 | 0,0 | 10,34876592 | 0,004696232 | Ppm1l | protein phosphatase 1 (formerly 2C)-like [Source:MGI Symbol;Acc:MGI:2139740] | 77,13 | 0,0 |
| ENSMUSG00000022309 | 116,522214 | 142,7 | 0,0 | 10,34953412 | 0,004671396 | Angpt1 | angiopoietin 1 [Source:MGI Symbol;Acc:MGI:108448] | 93,50 | 0,0 |
| ENSMUSG00000053226 | 93,8916662 | 125,2 | 0,0 | 10,35043161 | 0,008002308 | Dand5 | DAN domain family member 5, BMP antagonist [Source:MGI Symbol;Acc:MGI:1344365] | 71,25 | 0,0 |
| ENSMUSG00000024620 | 54,76941512 | 134,5 | 0,0 | 10,35433327 | 0,003438138 | Pdgfrb | platelet derived growth factor receptor, beta polypeptide [Source:MGI Symbol;Acc:MGI:97531] | 94,51 | 0,0 |
| ENSMUSG00000049404 | 79,62848463 | 121,8 | 0,0 | 10,35531403 | 0,007521921 | Rarres1 | retinoic acid receptor responder (tazarotene induced) 1 [Source:MGI Symbol;Acc:MGI:1924461] | 98,57 | 0,0 |
| ENSMUSG00000022040 | 381,0775271 | 653,6 | 0,6 | 10,35718761 | 0,001096359 | Ephx2 | epoxide hydrolase 2, cytoplasmic [Source:MGI Symbol;Acc:MGI:99500] | 100,00 | 1,2 |
| ENSMUSG00000052384 | 96,38695289 | 125,3 | 0,0 | 10,35872695 | 0,006353994 | Nrros | negative regulator of reactive oxygen species [Source:MGI Symbol;Acc:MGI:2445095] | 64,85 | 0,0 |
| ENSMUSG00000038296 | 60,18524918 | 124,3 | 0,0 | 10,37407303 | 0,005157407 | Galnt18 | tyl-alpha-D-galactosamine:polypeptide N-acetylgalactosaminyltransferase 18 [Source:MGI Symbol;Acc:MG | 97,00 | 0,0 |
| ENSMUSG00000027669 | 64,44667949 | 124,3 | 0,0 | 10,37423539 | 0,0047196 | Gnb4 | guanine nucleotide binding protein (G protein), beta 4 [Source:MGI Symbol;Acc:MGI:104581] | 70,12 | 0,0 |
| ENSMUSG00000046997 | 66,22370172 | 122,4 | 0,0 | 10,3809556 | 0,003413357 | Spsb4 | splA/ryanodine receptor domain and SOCS box containing 4 [Source:MGI Symbol;Acc:MGI:2183445] | 96,34 | 0,0 |
| ENSMUSG00000048550 | 140,8056751 | 136,0 | 0,0 | 10,38114297 | 0,005177485 | Thnsl1 | threonine synthase-like 1 (bacterial) [Source:MGI Symbol;Acc:MGI:2139347] | 88,33 | 0,0 |
| ENSMUSG00000045251 | 76,39178675 | 128,4 | 0,0 | 10,38891142 | 0,007337311 | Zfp688 | zinc finger protein 688 [Source:MGI Symbol;Acc:MGI:1916484] | 99,77 | 0,0 |
| ENSMUSG00000097328 | 78,38639435 | 127,7 | 0,0 | 10,39300516 | 0,003239323 | Tnfsf12 | tumor necrosis factor (ligand) superfamily, member 12 [Source:MGI Symbol;Acc:MGI:1196259] | 84,05 | 0,0 |
| ENSMUSG00000053706 | 71,56445536 | 126,0 | 0,0 | 10,39472781 | 0,006007756 | B430305J03Rik | RIKEN cDNA B430305J03 gene [Source:MGI Symbol;Acc:MGI:3697707] | 52,05 | 0,0 |
| ENSMUSG00000000759 | 90,03436169 | 131,9 | 0,0 | 10,3953507 | 0,004779948 | Tubgcp3 | tubulin, gamma complex associated protein 3 [Source:MGI Symbol;Acc:MGI:2183752] | 94,53 | 0,0 |
| ENSMUSG00000024270 | 98,21258179 | 150,5 | 0,0 | 10,41346811 | 0,006111698 | Slc39a6 | solute carrier family 39 (metal ion transporter), member 6 [Source:MGI Symbol;Acc:MGI:2147279] | 90,65 | 0,0 |
| ENSMUSG00000023980 | 96,6479701 | 129,5 | 0,0 | 10,41516197 | 0,004984913 | Taf8 | RNA polymerase II, TATA box binding protein (TBP)-associated factorq [Source:MGI Symbol;Acc:MGI:192 | 93,92 | 0,0 |
| ENSMUSG00000030630 | 78,90335328 | 132,3 | 0,0 | 10,41655598 | 0,004041278 | Fah | fumarylacetoacetate hydrolase [Source:MGI Symbol;Acc:MGI:95482] | 89,06 | 0,0 |
| ENSMUSG00000044807 | 94,57720501 | 140,6 | 0,0 | 10,41798759 | 0,004754807 | Zfp354c | zinc finger protein 354C [Source:MGI Symbol;Acc:MGI:1353621] | 94,96 | 0,0 |
| ENSMUSG00000010721 | 68,40821879 | 129,7 | 0,0 | 10,42002455 | 0,004520755 | Lmbr1 | limb region 1 [Source:MGI Symbol;Acc:MGI:1861746] | 55,34 | 0,0 |
| ENSMUSG00000063760 | 100,3498145 | 131,5 | 0,0 | 10,43170343 | 0,004940922 | Rnf217 | ring finger protein 217 [Source:MGI Symbol;Acc:MGI:3610311] | 45,61 | 0,0 |
| ENSMUSG00000034936 | 52,62972803 | 142,4 | 0,0 | 10,44598143 | 0,005425973 | Arl4d | ADP-ribosylation factor-like 4D [Source:MGI Symbol;Acc:MGI:1933155] | 99,45 | 0,0 |
| ENSMUSG00000037017 | 114,6247145 | 136,3 | 0,0 | 10,44717935 | 0,004170187 | Zscan21 | zinc finger and SCAN domain containing 21 [Source:MGI Symbol;Acc:MGI:99182] | 71,11 | 0,0 |
| ENSMUSG00000085024 | 77,50828062 | 139,6 | 0,0 | 10,45251161 | 0,004441908 | C230035I16Rik | RIKEN cDNA C230035I16 gene [Source:MGI Symbol;Acc:MGI:2444850] | #DIV/0! | #DIV/0! |
| ENSMUSG00000037784 | 83,79227332 | 135,6 | 0,0 | 10,466 | 0,007043709 | Dzip1l | DAZ interacting protein 1-like [Source:MGI Symbol;Acc:MGI:1919757] | 93,51 | 0,0 |
| ENSMUSG00000035357 | 63,65193006 | 133,2 | 0,0 | 10,4678875 | 0,003679909 | Pdzrn3 | PDZ domain containing RING finger 3 [Source:MGI Symbol;Acc:MGI:1933157] | 96,51 | 0,0 |
| ENSMUSG00000031755 | 87,51830955 | 136,9 | 0,0 | 10,4777855 | 0,005659757 | Bbs2 | Bardet-Biedl syndrome 2 (human) [Source:MGI Symbol;Acc:MGI:2135267] | 99,17 | 0,0 |
| ENSMUSG00000079494 | 60,71761407 | 134,5 | 0,0 | 10,48145501 | 0,006453689 | Cml5 | camello-like 5 [Source:MGI Symbol;Acc:MGI:1916299] | 100,00 | 0,0 |
| ENSMUSG00000075273 | 86,24488234 | 138,0 | 0,0 | 10,49430835 | 0,004879078 | Ttc30b | tetratricopeptide repeat domain 30B [Source:MGI Symbol;Acc:MGI:1919671] | 82,17 | 0,0 |
| ENSMUSG00000044122 | 113,9665348 | 137,6 | 0,0 | 10,49513556 | 0,006574102 | Proca1 | protein interacting with cyclin A1 [Source:MGI Symbol;Acc:MGI:1918274] | 74,96 | 0,0 |
| ENSMUSG00000005413 | 67,67046042 | 137,7 | 0,0 | 10,49720195 | 0,003796029 | Hmox1 | heme oxygenase 1 [Source:MGI Symbol;Acc:MGI:96163] | 97,79 | 0,0 |
| ENSMUSG00000002279 | 83,32316862 | 138,0 | 0,0 | 10,49864905 | 0,005786317 | Lmf1 | lipase maturation factor 1 [Source:MGI Symbol;Acc:MGI:1923733] | 76,20 | 0,0 |
| ENSMUSG00000014599 | 75,60122103 | 147,7 | 0,0 | 10,50084986 | 0,004625348 | Csf1 | colony stimulating factor 1 (macrophage) [Source:MGI Symbol;Acc:MGI:1339753] | 84,25 | 0,0 |
| ENSMUSG00000040464 | 79,03823985 | 138,6 | 0,0 | 10,50540926 | 0,00594822 | Gtpbp10 | GTP-binding protein 10 (putative) [Source:MGI Symbol;Acc:MGI:2385599] | 65,30 | 0,0 |
| ENSMUSG00000025193 | 76,99237327 | 137,8 | 0,0 | 10,50570328 | 0,007709848 | Cutc | cutC copper transporter [Source:MGI Symbol;Acc:MGI:1913638] | 97,60 | 0,0 |
| ENSMUSG00000067928 | 89,23008921 | 138,5 | 0,0 | 10,50860692 | 0,004450017 | Zfp760 | zinc finger protein 760 [Source:MGI Symbol;Acc:MGI:2679257] | 98,95 | 0,0 |
| ENSMUSG00000050666 | 136,1733063 | 142,0 | 0,0 | 10,51713183 | 0,006117719 | Vstm4 | V-set and transmembrane domain containing 4 [Source:MGI Symbol;Acc:MGI:2444633] | 87,28 | 0,0 |
| ENSMUSG00000024565 | 107,9932299 | 147,6 | 0,0 | 10,5215579 | 0,003995155 | Sall3 | sal-like 3 (Drosophila) [Source:MGI Symbol;Acc:MGI:109295] | 79,76 | 0,0 |
| ENSMUSG00000021483 | 84,02899269 | 143,8 | 0,0 | 10,52869082 | 0,004712445 | Cdk20 | cyclin-dependent kinase 20 [Source:MGI Symbol;Acc:MGI:2145349] | 99,80 | 0,0 |
| ENSMUSG00000021047 | 95,75064025 | 141,5 | 0,0 | 10,53273332 | 0,003469834 | Nova1 | neuro-oncological ventral antigen 1 [Source:MGI Symbol;Acc:MGI:104297] | 52,68 | 0,0 |
| ENSMUSG00000001376 | 122,1659997 | 144,9 | 0,0 | 10,53694333 | 0,002117304 | Vps50 | VPS50 EARP/GARPII complex subunit [Source:MGI Symbol;Acc:MGI:1920538] | 60,15 | 0,0 |
| ENSMUSG00000051351 | 69,85275425 | 140,9 | 0,0 | 10,54064731 | 0,004754807 | Zfp46 | zinc finger protein 46 [Source:MGI Symbol;Acc:MGI:99192] | 92,95 | 0,0 |
| ENSMUSG00000034826 | 115,5328078 | 144,4 | 0,0 | 10,54718549 | 0,002339017 | Nup54 | nucleoporin 54 [Source:MGI Symbol;Acc:MGI:1920460] | 80,02 | 0,0 |
| ENSMUSG00000050737 | 84,973452 | 156,7 | 0,0 | 10,54923739 | 0,006868128 | Ptges | prostaglandin E synthase [Source:MGI Symbol;Acc:MGI:1927593] | 91,18 | 0,0 |
| ENSMUSG00000074170 | 94,69571225 | 142,3 | 0,0 | 10,55396385 | 0,008345158 | Plekhf1 | in homology domain containing, family F (with FYVE domain) member 1 [Source:MGI Symbol;Acc:MGI:1 | 31,60 | 0,0 |
| ENSMUSG00000050052 | 67,84683525 | 141,9 | 0,0 | 10,55897389 | 0,00513118 | Tdrp | testis development related protein [Source:MGI Symbol;Acc:MGI:1919398] | 96,32 | 0,0 |
| ENSMUSG00000026142 | 82,43168476 | 154,8 | 0,0 | 10,56759349 | 0,004779333 | Rhbdd1 | rhomboid domain containing 1 [Source:MGI Symbol;Acc:MGI:1924117] | 88,87 | 0,0 |
| ENSMUSG00000032262 | 133,647526 | 148,5 | 0,0 | 10,57063209 | 0,007303788 | Elovl4 | tion of very long chain fatty acids (FEN1/Elo2, SUR4/Elo3, yeast)-like 4 [Source:MGI Symbol;Acc:MGI:19 | 97,92 | 0,0 |
| ENSMUSG00000049791 | 91,65017404 | 157,5 | 0,0 | 10,57479028 | 0,007311599 | Fzd4 | frizzled class receptor 4 [Source:MGI Symbol;Acc:MGI:108520] | 94,64 | 0,0 |
| ENSMUSG00000026816 | 106,0002472 | 149,7 | 0,0 | 10,5822928 | 0,006735863 | Gtf3c5 | general transcription factor IIIC, polypeptide 5 [Source:MGI Symbol;Acc:MGI:1917489] | 97,00 | 0,0 |
| ENSMUSG00000029366 | 92,04701498 | 142,3 | 0,0 | 10,58363687 | 0,005971788 | Dck | deoxycytidine kinase [Source:MGI Symbol;Acc:MGI:102726] | 97,31 | 0,0 |
| ENSMUSG00000034930 | 79,86475213 | 149,7 | 0,0 | 10,59379682 | 0,003479574 | Rtkn | rhotekin [Source:MGI Symbol;Acc:MGI:107371] | 75,36 | 0,0 |
| ENSMUSG00000030032 | 75,29836321 | 146,6 | 0,0 | 10,59512511 | 0,005948324 | Wdr54 | WD repeat domain 54 [Source:MGI Symbol;Acc:MGI:1922909] | 57,50 | 0,0 |
| ENSMUSG00000036611 | 83,55551127 | 145,4 | 0,0 | 10,59975892 | 0,003911784 | Eepd1 | ndonuclease/exonuclease/phosphatase family domain containing 1 [Source:MGI Symbol;Acc:MGI:191473 | 99,11 | 0,0 |
| ENSMUSG00000007655 | 101,2015448 | 153,0 | 0,0 | 10,62275391 | 0,006198296 | Cav1 | caveolin 1, caveolae protein [Source:MGI Symbol;Acc:MGI:102709] | 81,26 | 0,0 |
| ENSMUSG00000048970 | 151,6295051 | 151,2 | 0,0 | 10,64202611 | 0,005058806 | C1galt1c1 | C1GALT1-specific chaperone 1 [Source:MGI Symbol;Acc:MGI:1913493] | 99,52 | 0,0 |
| ENSMUSG00000039246 | 75,0956255 | 149,6 | 0,0 | 10,64518832 | 0,00404034 | Lyplal1 | lysophospholipase-like 1 [Source:MGI Symbol;Acc:MGI:2385115] | 97,47 | 0,0 |
| ENSMUSG00000039410 | 160,4482855 | 154,8 | 0,0 | 10,6468605 | 0,003934342 | Prdm16 | PR domain containing 16 [Source:MGI Symbol;Acc:MGI:1917923] | 86,30 | 0,0 |
| ENSMUSG00000038102 | 97,74048265 | 161,4 | 0,0 | 10,66708892 | 0,005923855 | Trappc11 | trafficking protein particle complex 11 [Source:MGI Symbol;Acc:MGI:2444585] | 73,94 | 0,0 |
| ENSMUSG00000010080 | 126,2157797 | 191,9 | 0,0 | 10,72953282 | 0,002138615 | Epn3 | epsin 3 [Source:MGI Symbol;Acc:MGI:1919139] | 94,91 | 0,0 |
| ENSMUSG00000036898 | 68,00807751 | 157,7 | 0,0 | 10,73070449 | 0,002449971 | Zfp157 | zinc finger protein 157 [Source:MGI Symbol;Acc:MGI:1919404] | 85,25 | 0,0 |
| ENSMUSG00000025525 | 113,3049389 | 163,8 | 0,0 | 10,73241134 | 0,003510998 | Apool | apolipoprotein O-like [Source:MGI Symbol;Acc:MGI:1915367] | 97,96 | 0,0 |
| ENSMUSG00000053070 | 110,3540435 | 161,3 | 0,0 | 10,74539795 | 0,005961182 | 9230110C19Rik | RIKEN cDNA 9230110C19 gene [Source:MGI Symbol;Acc:MGI:3045346] | 99,98 | 0,0 |
| ENSMUSG00000097881 | 95,07394693 | 162,1 | 0,0 | 10,75129435 | 0,004610597 | Celrr | cerebellum expressed regulatory RNA [Source:MGI Symbol;Acc:MGI:2685664] | #DIV/0! | #DIV/0! |
| ENSMUSG00000046862 | 112,1154353 | 172,7 | 0,0 | 10,75311113 | 0,00404364 | Pramef8 | PRAME family member 8 [Source:MGI Symbol;Acc:MGI:2140473] | 89,68 | 0,0 |
| ENSMUSG00000043510 | 134,5364984 | 165,8 | 0,0 | 10,75526683 | 0,002778958 | Hscb | HscB iron-sulfur cluster co-chaperone [Source:MGI Symbol;Acc:MGI:2141135] | 99,67 | 0,0 |
| ENSMUSG00000020334 | 91,75102906 | 161,2 | 0,0 | 10,75529934 | 0,004940922 | Slc22a4 | solute carrier family 22 (organic cation transporter), member 4 [Source:MGI Symbol;Acc:MGI:1353479] | 99,51 | 0,0 |
| ENSMUSG00000024593 | 171,9658849 | 192,9 | 0,0 | 10,75663661 | 0,005739649 | Megf10 | multiple EGF-like-domains 10 [Source:MGI Symbol;Acc:MGI:2685177] | 80,14 | 0,0 |
| ENSMUSG00000024663 | 89,50020698 | 162,3 | 0,0 | 10,76014108 | 0,003587282 | Rab3il1 | RAB3A interacting protein (rabin3)-like 1 [Source:MGI Symbol;Acc:MGI:1922010] | 64,16 | 0,0 |
| ENSMUSG00000070366 | 92,33289675 | 164,6 | 0,0 | 10,76842596 | 0,005071544 | Plpp4 | phospholipid phosphatase 4 [Source:MGI Symbol;Acc:MGI:2685936] | 100,00 | 0,0 |
| ENSMUSG00000028268 | 134,143328 | 172,2 | 0,0 | 10,77116862 | 0,002154621 | Gbp3 | guanylate binding protein 3 [Source:MGI Symbol;Acc:MGI:1926263] | 88,50 | 0,0 |
| ENSMUSG00000015653 | 169,8342264 | 178,2 | 0,0 | 10,77132804 | 0,005369631 | Steap2 | six transmembrane epithelial antigen of prostate 2 [Source:MGI Symbol;Acc:MGI:1921301] | 63,43 | 0,0 |
| ENSMUSG00000045733 | 130,2396947 | 167,5 | 0,0 | 10,77215722 | 0,003086144 | Sprn | shadow of prion protein [Source:MGI Symbol;Acc:MGI:3582583] | 93,60 | 0,0 |
| ENSMUSG00000035078 | 93,76571232 | 172,4 | 0,0 | 10,77250228 | 0,005739649 | Mtmr9 | myotubularin related protein 9 [Source:MGI Symbol;Acc:MGI:2442842] | 98,29 | 0,0 |
| ENSMUSG00000026796 | 127,6184889 | 164,6 | 0,0 | 10,77379447 | 0,003679909 | Fam129b | family with sequence similarity 129, member B [Source:MGI Symbol;Acc:MGI:2442910] | 95,36 | 0,0 |
| ENSMUSG00000040124 | 93,81417809 | 165,3 | 0,0 | 10,77729376 | 0,003374021 | Gorab | golgin, RAB6-interacting [Source:MGI Symbol;Acc:MGI:2138271] | 99,87 | 0,0 |
| ENSMUSG00000078670 | 107,9336815 | 164,8 | 0,0 | 10,78255832 | 0,002157466 | Fam174b | family with sequence similarity 174, member B [Source:MGI Symbol;Acc:MGI:3698178] | 97,01 | 0,0 |
| ENSMUSG00000022150 | 113,3323266 | 172,7 | 0,0 | 10,79111037 | 0,004467035 | Dab2 | disabled 2, mitogen-responsive phosphoprotein [Source:MGI Symbol;Acc:MGI:109175] | 78,77 | 0,0 |
| ENSMUSG00000105352 | 121,3658299 | 169,8 | 0,0 | 10,80682722 | 0,00354734 | C030018K13Rik | RIKEN cDNA C030018K13 gene [Source:MGI Symbol;Acc:MGI:1924632] | #DIV/0! | #DIV/0! |
| ENSMUSG00000029650 | 111,7976043 | 169,1 | 0,0 | 10,81324471 | 0,00418273 | Slc46a3 | solute carrier family 46, member 3 [Source:MGI Symbol;Acc:MGI:1918956] | 99,36 | 0,0 |
| ENSMUSG00000024500 | 742,2971983 | 1270,2 | 0,9 | 10,81461772 | 1,46512E-06 | Ppp2r2b | protein phosphatase 2, regulatory subunit B, beta [Source:MGI Symbol;Acc:MGI:1920180] | 95,27 | 0,7 |
| ENSMUSG00000020189 | 144,2212838 | 172,3 | 0,0 | 10,81802839 | 0,004068359 | Osbpl8 | oxysterol binding protein-like 8 [Source:MGI Symbol;Acc:MGI:2443807] | 94,07 | 0,0 |
| ENSMUSG00000028494 | 99,96900079 | 171,5 | 0,0 | 10,82268824 | 0,005255281 | Plin2 | perilipin 2 [Source:MGI Symbol;Acc:MGI:87920] | 85,58 | 0,0 |
| ENSMUSG00000043822 | 131,2717278 | 178,0 | 0,0 | 10,82380646 | 0,004455351 | Adamtsl5 | ADAMTS-like 5 [Source:MGI Symbol;Acc:MGI:1913798] | 99,33 | 0,0 |
| ENSMUSG00000009640 | 107,4547484 | 174,1 | 0,0 | 10,82990321 | 0,002761475 | Dmap1 | DNA methyltransferase 1-associated protein 1 [Source:MGI Symbol;Acc:MGI:1913483] | 99,15 | 0,0 |
| ENSMUSG00000055633 | 94,64722421 | 173,3 | 0,0 | 10,83030994 | 0,00302911 | Zfp580 | zinc finger protein 580 [Source:MGI Symbol;Acc:MGI:1916242] | 87,94 | 0,0 |
| ENSMUSG00000082519 | 151,7563574 | 178,1 | 0,0 | 10,84266411 | 0,005018328 | Vamp7-ps | vesicle-associated membrane protein 7, pseudogene [Source:MGI Symbol;Acc:MGI:3652239] | #DIV/0! | #DIV/0! |
| ENSMUSG00000036291 | 113,4805548 | 184,1 | 0,0 | 10,85083332 | 0,004166827 | Ap5m1 | adaptor-related protein complex 5, mu 1 subunit [Source:MGI Symbol;Acc:MGI:1921635] | 98,67 | 0,0 |
| ENSMUSG00000068617 | 123,2919314 | 186,5 | 0,0 | 10,85310843 | 0,002102103 | Efcab1 | EF hand calcium binding domain 1 [Source:MGI Symbol;Acc:MGI:1914043] | 97,73 | 0,0 |
| ENSMUSG00000037344 | 104,4035612 | 187,8 | 0,0 | 10,86688244 | 0,004288554 | Slc12a9 | olute carrier family 12 (potassium/chloride transporters), member 9 [Source:MGI Symbol;Acc:MGI:193353 | 95,32 | 0,0 |
| ENSMUSG00000039648 | 98,95953495 | 176,7 | 0,0 | 10,86888264 | 0,0049693 | Ccbl1 | cysteine conjugate-beta lyase 1 [Source:MGI Symbol;Acc:MGI:1917516] | 86,18 | 0,0 |
| ENSMUSG00000033717 | 92,20069892 | 174,8 | 0,0 | 10,87585447 | 0,00314896 | Adra2a | adrenergic receptor, alpha 2a [Source:MGI Symbol;Acc:MGI:87934] | 96,63 | 0,0 |
| ENSMUSG00000027739 | 718,3458159 | 1241,5 | 0,9 | 10,88246443 | 7,08271E-06 | Rab33b | RAB33B, member RAS oncogene family [Source:MGI Symbol;Acc:MGI:1330805] | 91,17 | 0,5 |
| ENSMUSG00000001552 | 141,7307927 | 186,3 | 0,0 | 10,89032892 | 0,002947323 | Jup | junction plakoglobin [Source:MGI Symbol;Acc:MGI:96650] | 99,36 | 0,0 |
| ENSMUSG00000019359 | 81,37095437 | 190,7 | 0,0 | 10,90092217 | 0,003396312 | Gdpd2 | glycerophosphodiester phosphodiesterase domain containing 2 [Source:MGI Symbol;Acc:MGI:1918834] | 96,07 | 0,0 |
| ENSMUSG00000052914 | 139,5549214 | 192,8 | 0,0 | 10,94575235 | 0,007028225 | Cyp2j6 | cytochrome P450, family 2, subfamily j, polypeptide 6 [Source:MGI Symbol;Acc:MGI:1270148] | 93,24 | 0,0 |
| ENSMUSG00000031209 | 110,1907986 | 189,8 | 0,0 | 10,94699197 | 0,002985684 | Heph | hephaestin [Source:MGI Symbol;Acc:MGI:1332240] | 79,87 | 0,0 |
| ENSMUSG00000049411 | 98,18829718 | 184,4 | 0,0 | 10,95360564 | 0,002725637 | Tmem241 | transmembrane protein 241 [Source:MGI Symbol;Acc:MGI:2442435] | 97,33 | 0,0 |
| ENSMUSG00000022235 | 98,35857891 | 187,8 | 0,0 | 10,95557844 | 0,003391373 | Cmbl | carboxymethylenebutenolidase-like (Pseudomonas) [Source:MGI Symbol;Acc:MGI:1916824] | 95,82 | 0,0 |
| ENSMUSG00000025780 | 136,4329129 | 188,0 | 0,0 | 10,95810393 | 0,006640362 | Itih5 | inter-alpha (globulin) inhibitor H5 [Source:MGI Symbol;Acc:MGI:1925751] | 90,61 | 0,0 |
| ENSMUSG00000027351 | 2235,347083 | 3418,9 | 746,8 | 10,96034909 | 6,54566E-05 | Spred1 | sprouty protein with EVH-1 domain 1, related sequence [Source:MGI Symbol;Acc:MGI:2150016] | 98,93 | 28,2 |
| ENSMUSG00000093769 | 91,95381179 | 191,5 | 0,0 | 10,97242968 | 0,004455351 | Hist2h3c1 | histone cluster 2, H3c1 [Source:MGI Symbol;Acc:MGI:2448355] | 66,43 | 0,0 |
| ENSMUSG00000022109 | 140,4836616 | 191,4 | 0,0 | 10,98332693 | 0,001705667 | Med4 | mediator complex subunit 4 [Source:MGI Symbol;Acc:MGI:1914631] | 100,00 | 0,0 |
| ENSMUSG00000037594 | 100,588727 | 189,5 | 0,0 | 10,99150005 | 0,003820711 | BC022687 | cDNA sequence BC022687 [Source:MGI Symbol;Acc:MGI:2443738] | 99,48 | 0,0 |
| ENSMUSG00000021848 | 244,4538723 | 207,7 | 0,0 | 11,00305469 | 0,003728942 | Otx2 | orthodenticle homolog 2 [Source:MGI Symbol;Acc:MGI:97451] | 80,84 | 0,0 |
| ENSMUSG00000050989 | 122,6434401 | 201,2 | 0,0 | 11,03627604 | 0,004783257 | Sepn1 | selenoprotein N, 1 [Source:MGI Symbol;Acc:MGI:2151208] | 99,62 | 0,0 |
| ENSMUSG00000070392 | 96,87764324 | 201,2 | 0,0 | 11,03977778 | 0,003679909 | Gm20634 | predicted gene 20634 [Source:MGI Symbol;Acc:MGI:5313081] | #DIV/0! | #DIV/0! |
| ENSMUSG00000020393 | 99,26998617 | 218,1 | 0,0 | 11,0648064 | 0,004120523 | Kremen1 | kringle containing transmembrane protein 1 [Source:MGI Symbol;Acc:MGI:1933988] | 64,59 | 0,0 |
| ENSMUSG00000041143 | 99,42041028 | 200,3 | 0,0 | 11,07308784 | 0,003904395 | Tmco4 | transmembrane and coiled-coil domains 4 [Source:MGI Symbol;Acc:MGI:1924306] | 93,73 | 0,0 |
| ENSMUSG00000048424 | 159,4491095 | 207,6 | 0,0 | 11,0792235 | 0,005825188 | Ranbp3l | RAN binding protein 3-like [Source:MGI Symbol;Acc:MGI:2444654] | 98,21 | 0,0 |
| ENSMUSG00000045237 | 127,7311273 | 208,8 | 0,0 | 11,09687984 | 0,003198527 | 1110012L19Rik | RIKEN cDNA 1110012L19 gene [Source:MGI Symbol;Acc:MGI:1915868] | 94,49 | 0,0 |
| ENSMUSG00000042155 | 204,4413743 | 212,0 | 0,0 | 11,10909438 | 0,003257241 | Klhl23 | kelch-like 23 [Source:MGI Symbol;Acc:MGI:2683536] | 84,89 | 0,0 |
| ENSMUSG00000029096 | 112,7001275 | 210,4 | 0,0 | 11,11310869 | 0,004735963 | Htra3 | HtrA serine peptidase 3 [Source:MGI Symbol;Acc:MGI:1925808] | 81,00 | 0,0 |
| ENSMUSG00000035062 | 111,2723984 | 210,5 | 0,0 | 11,1245409 | 0,001661464 | Zc4h2 | zinc finger, C4H2 domain containing [Source:MGI Symbol;Acc:MGI:2679294] | 96,75 | 0,0 |
| ENSMUSG00000094707 | 102,803934 | 222,2 | 0,0 | 11,13138849 | 0,004232404 | A830019P07Rik | RIKEN cDNA A830019P07 gene [Source:MGI Symbol;Acc:MGI:3028035] | #DIV/0! | #DIV/0! |
| ENSMUSG00000025588 | 117,243061 | 216,1 | 0,0 | 11,14408578 | 0,003532165 | Nat1 | N-acetyl transferase 1 [Source:MGI Symbol;Acc:MGI:97279] | 84,51 | 0,0 |
| ENSMUSG00000036894 | 122,3282611 | 216,4 | 0,0 | 11,16737638 | 0,001811264 | Rap2b | RAP2B, member of RAS oncogene family [Source:MGI Symbol;Acc:MGI:1921262] | 62,54 | 0,0 |
| ENSMUSG00000030806 | 129,3012336 | 223,0 | 0,0 | 11,1785358 | 0,002821316 | Stx1b | syntaxin 1B [Source:MGI Symbol;Acc:MGI:1930705] | 91,53 | 0,0 |
| ENSMUSG00000049521 | 136,241606 | 226,1 | 0,0 | 11,18184235 | 0,003253813 | Cdc42ep1 | CDC42 effector protein (Rho GTPase binding) 1 [Source:MGI Symbol;Acc:MGI:1929763] | 99,54 | 0,0 |
| ENSMUSG00000045827 | 98,16385389 | 219,0 | 0,0 | 11,18527574 | 0,00212029 | Serpinb9 | serine (or cysteine) peptidase inhibitor, clade B, member 9 [Source:MGI Symbol;Acc:MGI:106603] | 92,44 | 0,0 |
| ENSMUSG00000026988 | 139,1296065 | 222,6 | 0,0 | 11,19262421 | 0,003192978 | Wdsub1 | WD repeat, SAM and U-box domain containing 1 [Source:MGI Symbol;Acc:MGI:1919387] | 90,23 | 0,0 |
| ENSMUSG00000068762 | 136,0194603 | 221,6 | 0,0 | 11,20118968 | 0,004888202 | Gstm6 | glutathione S-transferase, mu 6 [Source:MGI Symbol;Acc:MGI:1309467] | 91,25 | 0,0 |
| ENSMUSG00000021238 | 1986,693012 | 3555,3 | 1,9 | 11,20646142 | 3,89286E-06 | Aldh6a1 | aldehyde dehydrogenase family 6, subfamily A1 [Source:MGI Symbol;Acc:MGI:1915077] | 100,00 | 1,1 |
| ENSMUSG00000042743 | 108,9590131 | 234,7 | 0,0 | 11,22345553 | 0,002016411 | Sgtb | nall glutamine-rich tetratricopeptide repeat (TPR)-containing, beta [Source:MGI Symbol;Acc:MGI:244461 | 98,84 | 0,0 |
| ENSMUSG00000023186 | 132,1237515 | 226,0 | 0,0 | 11,22480423 | 0,003347806 | Vwa5a | von Willebrand factor A domain containing 5A [Source:MGI Symbol;Acc:MGI:1915026] | 89,19 | 0,0 |
| ENSMUSG00000042298 | 248,0909699 | 231,5 | 0,0 | 11,23689045 | 0,002302248 | Ttc19 | tetratricopeptide repeat domain 19 [Source:MGI Symbol;Acc:MGI:1920045] | 98,90 | 0,0 |
| ENSMUSG00000085492 | 137,2320001 | 237,0 | 0,0 | 11,26612553 | 0,004058917 | Trmt61b | tRNA methyltransferase 61B [Source:MGI Symbol;Acc:MGI:1916039] | #DIV/0! | #DIV/0! |
| ENSMUSG00000006931 | 163,1537514 | 240,7 | 0,0 | 11,26784127 | 0,00337772 | P3h4 | prolyl 3-hydroxylase family member 4 (non-enzymatic) [Source:MGI Symbol;Acc:MGI:1913430] | 96,34 | 0,0 |
| ENSMUSG00000027439 | 169,3576923 | 240,3 | 0,0 | 11,27396195 | 0,00461289 | Gzf1 | GDNF-inducible zinc finger protein 1 [Source:MGI Symbol;Acc:MGI:1921783] | 93,23 | 0,0 |
| ENSMUSG00000050945 | 131,860763 | 245,3 | 0,0 | 11,27559261 | 0,004375869 | Zfp438 | zinc finger protein 438 [Source:MGI Symbol;Acc:MGI:2444919] | 99,78 | 0,0 |
| ENSMUSG00000002731 | 121,8043819 | 243,9 | 0,0 | 11,30881275 | 0,003324528 | Prkra | ı kinase, interferon inducible double stranded RNA dependent activator [Source:MGI Symbol;Acc:MGI:134 | 98,14 | 0,0 |
| ENSMUSG00000026946 | 160,3422056 | 241,6 | 0,0 | 11,30892793 | 0,003865394 | Nmi | N-myc (and STAT) interactor [Source:MGI Symbol;Acc:MGI:1928368] | 92,24 | 0,0 |
| ENSMUSG00000053117 | 133,8865532 | 239,9 | 0,0 | 11,31052718 | 0,003764373 | E330013P04Rik | RIKEN cDNA E330013P04 gene [Source:MGI Symbol;Acc:MGI:2147732] | #DIV/0! | #DIV/0! |
| ENSMUSG00000097770 | 133,4976071 | 239,0 | 0,0 | 11,31376332 | 0,004271538 | Gm26776 | predicted gene, 26776 [Source:MGI Symbol;Acc:MGI:5477270] | #DIV/0! | #DIV/0! |
| ENSMUSG00000007908 | 103,7143102 | 240,2 | 0,0 | 11,31991369 | 0,001246478 | Hmgcll1 | 3-hydroxymethyl-3-methylglutaryl-Coenzyme A lyase-like 1 [Source:MGI Symbol;Acc:MGI:2446108] | 96,83 | 0,0 |
| ENSMUSG00000035246 | 123,9795441 | 241,0 | 0,0 | 11,32471853 | 0,002972881 | Pcyt1b | phosphate cytidylyltransferase 1, choline, beta isoform [Source:MGI Symbol;Acc:MGI:2147987] | 94,26 | 0,0 |
| ENSMUSG00000021615 | 150,5023627 | 243,0 | 0,0 | 11,32576913 | 0,003798588 | Xrcc4 | ay repair complementing defective repair in Chinese hamster cells 4 [Source:MGI Symbol;Acc:MGI:13337 | 97,61 | 0,0 |
| ENSMUSG00000025531 | 153,15668 | 244,8 | 0,0 | 11,33044034 | 0,003234181 | Chm | choroidermia (RAB escort protein 1) [Source:MGI Symbol;Acc:MGI:892979] | 97,63 | 0,0 |
| ENSMUSG00000038507 | 135,284191 | 258,3 | 0,0 | 11,34079332 | 0,002938374 | Parp12 | poly (ADP-ribose) polymerase family, member 12 [Source:MGI Symbol;Acc:MGI:2143990] | 98,76 | 0,0 |
| ENSMUSG00000090083 | 213,319916 | 247,9 | 0,0 | 11,3435497 | 0,001721802 | Rnf8 | ring finger protein 8 [Source:MGI Symbol;Acc:MGI:1929069] | 94,52 | 0,0 |
| ENSMUSG00000032523 | 186,2888272 | 280,7 | 0,0 | 11,34714601 | 0,003758957 | Hhatl | hedgehog acyltransferase-like [Source:MGI Symbol;Acc:MGI:1922020] | 93,49 | 0,0 |
| ENSMUSG00000035934 | 144,6370768 | 248,2 | 0,0 | 11,35403832 | 0,003406984 | Pknox2 | Pbx/knotted 1 homeobox 2 [Source:MGI Symbol;Acc:MGI:2445415] | 88,13 | 0,0 |

| | | | | | | | | | |
|---|---|---|---|---|---|---|---|---|---|
| ENSMUSG00000036764 | 139,6733425 | 249,4 | 0,0 | 11,36207887 | 0,003169049 | Dnajc12 | DnaJ heat shock protein family (Hsp40) member C12 [Source:MGI Symbol;Acc:MGI:1353428] | 95,92 | 0,0 |
| ENSMUSG00000030352 | 160,1290492 | 273,9 | 0,0 | 11,3738846 | 0,004087681 | Tspan9 | tetraspanin 9 [Source:MGI Symbol;Acc:MGI:1924558] | 83,27 | 0,0 |
| ENSMUSG00000041298 | 196,4095457 | 259,3 | 0,0 | 11,38560943 | 0,003550342 | Katnal1 | katanin p60 subunit A-like 1 [Source:MGI Symbol;Acc:MGI:2387638] | 88,86 | 0,0 |
| ENSMUSG00000022414 | 159,5062913 | 256,9 | 0,0 | 11,39113915 | 0,002298686 | Tab1 | TGF-beta activated kinase 1/MAP3K7 binding protein 1 [Source:MGI Symbol;Acc:MGI:1913763] | 100,00 | 0,0 |
| ENSMUSG00000034910 | 176,7719202 | 270,4 | 0,0 | 11,42473058 | 0,002982748 | Pygo1 | pygopus 1 [Source:MGI Symbol;Acc:MGI:1919385] | 87,25 | 0,0 |
| ENSMUSG00000015085 | 223,9268565 | 279,8 | 0,0 | 11,42979404 | 0,00303887 | Entpd2 | ectonucleoside triphosphate diphosphohydrolase 2 [Source:MGI Symbol;Acc:MGI:1096863] | 99,40 | 0,0 |
| ENSMUSG00000055991 | 154,5482654 | 278,6 | 0,0 | 11,45831504 | 0,001510534 | Zkscan5 | zinc finger with KRAB and SCAN domains 5 [Source:MGI Symbol;Acc:MGI:107533] | 99,57 | 0,0 |
| ENSMUSG00000063281 | 256,4405292 | 271,5 | 0,0 | 11,47172907 | 0,00113495 | Zfp35 | zinc finger protein 35 [Source:MGI Symbol;Acc:MGI:99179] | 95,23 | 0,0 |
| ENSMUSG00000023966 | 183,2117524 | 274,1 | 0,0 | 11,47461148 | 0,003978272 | Rsph9 | radial spoke head 9 homolog (Chlamydomonas) [Source:MGI Symbol;Acc:MGI:1922814] | 95,62 | 0,0 |
| ENSMUSG00000001864 | 174,427741 | 279,9 | 0,0 | 11,50022515 | 0,002356428 | Aif1l | allograft inflammatory factor 1-like [Source:MGI Symbol;Acc:MGI:1919598] | 95,69 | 0,0 |
| ENSMUSG00000037419 | 260,6435186 | 287,7 | 0,0 | 11,53311186 | 0,004346481 | Endod1 | endonuclease domain containing 1 [Source:MGI Symbol;Acc:MGI:1919196] | 98,14 | 0,0 |
| ENSMUSG00000030086 | 172,6188782 | 280,5 | 0,0 | 11,53356008 | 0,001368579 | Chchd6 | coiled-coil-helix-coiled-coil-helix domain containing 6 [Source:MGI Symbol;Acc:MGI:1913348] | 98,79 | 0,0 |
| ENSMUSG00000097023 | 211,9947214 | 299,1 | 0,0 | 11,57751983 | 0,003169049 | AI854517 | expressed sequence AI854517 [Source:MGI Symbol;Acc:MGI:2142071] | #DIV/0! | #DIV/0! |
| ENSMUSG00000039519 | 178,3469779 | 302,7 | 0,0 | 11,61559518 | 0,001303089 | Cyp7b1 | cytochrome P450, family 7, subfamily b, polypeptide 1 [Source:MGI Symbol;Acc:MGI:104978] | 95,61 | 0,0 |
| ENSMUSG00000025538 | 190,924172 | 307,4 | 0,0 | 11,63495665 | 0,000610788 | Sumf2 | sulfatase modifying factor 2 [Source:MGI Symbol;Acc:MGI:1915152] | 98,13 | 0,0 |
| ENSMUSG00000040183 | 195,3080488 | 306,2 | 0,0 | 11,63598299 | 0,003805051 | Ankrd6 | ankyrin repeat domain 6 [Source:MGI Symbol;Acc:MGI:2154278] | 94,92 | 0,0 |
| ENSMUSG00000024234 | 219,0570477 | 318,0 | 0,0 | 11,71456682 | 0,003553886 | Mtpap | mitochondrial poly(A) polymerase [Source:MGI Symbol;Acc:MGI:1914690] | 99,80 | 0,0 |
| ENSMUSG00000003849 | 188,433581 | 319,0 | 0,0 | 11,73896123 | 0,002617225 | Nqo1 | NAD(P)H dehydrogenase, quinone 1 [Source:MGI Symbol;Acc:MGI:103187] | 100,00 | 0,0 |
| ENSMUSG00000020638 | 203,2678194 | 326,7 | 0,0 | 11,74466961 | 0,00269501 | Cmpk2 | cytidine monophosphate (UMP-CMP) kinase 2, mitochondrial [Source:MGI Symbol;Acc:MGI:99830] | 99,91 | 0,0 |
| ENSMUSG00000044952 | 160,691116 | 321,1 | 0,0 | 11,74894173 | 0,000797365 | Kctd21 | potassium channel tetramerisation domain containing 21 [Source:MGI Symbol;Acc:MGI:3643121] | 100,00 | 0,0 |
| ENSMUSG00000044716 | 207,6409752 | 338,4 | 0,0 | 11,75835107 | 0,002402222 | Dok7 | docking protein 7 [Source:MGI Symbol;Acc:MGI:3584043] | 95,20 | 0,0 |
| ENSMUSG00000025933 | 195,6666824 | 334,9 | 0,0 | 11,76253229 | 0,002831118 | Tmem14a | transmembrane protein 14A [Source:MGI Symbol;Acc:MGI:1922962] | 91,61 | 0,0 |
| ENSMUSG00000025742 | 363,7574555 | 342,5 | 0,0 | 11,76344034 | 0,001485551 | Prps2 | phosphoribosyl pyrophosphate synthetase 2 [Source:MGI Symbol;Acc:MGI:97776] | 97,70 | 0,0 |
| ENSMUSG00000020829 | 221,6784669 | 343,4 | 0,0 | 11,76519026 | 0,00303887 | Slc46a1 | solute carrier family 46, member 1 [Source:MGI Symbol;Acc:MGI:1098733] | 99,56 | 0,0 |
| ENSMUSG00000027628 | 236,7258836 | 331,9 | 0,0 | 11,76962821 | 0,003276905 | Aar2 | AAR2 splicing factor homolog [Source:MGI Symbol;Acc:MGI:1915545] | 85,31 | 0,0 |
| ENSMUSG00000051041 | 221,9639071 | 343,3 | 0,0 | 11,80506449 | 0,003090135 | Olfml1 | olfactomedin-like 1 [Source:MGI Symbol;Acc:MGI:2679264] | 100,00 | 0,0 |
| ENSMUSG00000024535 | 185,0802724 | 340,5 | 0,0 | 11,83428534 | 0,001323936 | Snx24 | sorting nexing 24 [Source:MGI Symbol;Acc:MGI:1916476] | 98,87 | 0,0 |
| ENSMUSG00000039007 | 250,7090292 | 363,1 | 0,0 | 11,86816906 | 0,003260047 | Cpq | carboxypeptidase Q [Source:MGI Symbol;Acc:MGI:1889205] | 97,94 | 0,0 |
| ENSMUSG00000044197 | 208,1676903 | 355,9 | 0,0 | 11,87531566 | 0,001526084 | Gpr146 | G protein-coupled receptor 146 [Source:MGI Symbol;Acc:MGI:1933113] | 75,49 | 0,0 |
| ENSMUSG00000049536 | 181,8137565 | 362,3 | 0,0 | 11,90556291 | 0,002221477 | Tceal1 | transcription elongation factor A (SII)-like 1 [Source:MGI Symbol;Acc:MGI:2385317] | 73,75 | 0,0 |
| ENSMUSG00000067567 | 238,8205249 | 366,7 | 0,0 | 11,91153157 | 0,000746847 | Hdac8 | histone deacetylase 8 [Source:MGI Symbol;Acc:MGI:1917565] | 99,23 | 0,0 |
| ENSMUSG00000031714 | 169,8060147 | 363,9 | 0,0 | 11,92886321 | 0,000903303 | Gab1 | growth factor receptor bound protein 2-associated protein 1 [Source:MGI Symbol;Acc:MGI:108088] | 96,49 | 0,0 |
| ENSMUSG00000033964 | 294,9748206 | 382,1 | 0,0 | 11,94920784 | 0,000688307 | Zbtb41 | zinc finger and BTB domain containing 41 homolog [Source:MGI Symbol;Acc:MGI:2444487] | 99,27 | 0,0 |
| ENSMUSG00000020642 | 224,6960024 | 381,4 | 0,0 | 11,97037319 | 0,001366453 | Rnf144a | ring finger protein 144A [Source:MGI Symbol;Acc:MGI:1344401] | 98,93 | 0,0 |
| ENSMUSG00000036356 | 223,0048275 | 386,9 | 0,0 | 11,99273288 | 0,003510352 | Csgalnact1 | chondroitin sulfate N-acetylgalactosaminyltransferase 1 [Source:MGI Symbol;Acc:MGI:2442354] | 92,76 | 0,0 |
| ENSMUSG00000053411 | 208,6437815 | 391,4 | 0,0 | 12,01396894 | 0,002063127 | Cbx7 | chromobox 7 [Source:MGI Symbol;Acc:MGI:1196439] | 89,48 | 0,0 |
| ENSMUSG00000025037 | 246,7675242 | 405,2 | 0,0 | 12,03789633 | 0,002135809 | Maoa | monoamine oxidase A [Source:MGI Symbol;Acc:MGI:96915] | 90,65 | 0,0 |
| ENSMUSG00000074918 | 253,9414758 | 404,2 | 0,0 | 12,05095245 | 0,001031677 | Inafm2 | InaF motif containing 2 [Source:MGI Symbol;Acc:MGI:1915354] | 98,88 | 0,0 |
| ENSMUSG00000026255 | 253,7704158 | 443,9 | 0,0 | 12,19332904 | 0,002653914 | Efhd1 | EF hand domain containing 1 [Source:MGI Symbol;Acc:MGI:1921607] | 99,95 | 0,0 |
| ENSMUSG00000036833 | 193,775732 | 476,7 | 0,0 | 12,24656202 | 0,001241521 | Pnpla7 | patatin-like phospholipase domain containing 7 [Source:MGI Symbol;Acc:MGI:2385325] | 98,51 | 0,0 |
| ENSMUSG00000041658 | 249,2756818 | 488,9 | 0,0 | 12,3011532 | 0,001223013 | Rragb | Ras-related GTP binding B [Source:MGI Symbol;Acc:MGI:3038613] | 100,00 | 0,0 |
| ENSMUSG00000039428 | 361,5622167 | 523,0 | 0,0 | 12,31848053 | 0,001506164 | Tmem135 | transmembrane protein 135 [Source:MGI Symbol;Acc:MGI:1920009] | 69,21 | 0,0 |
| ENSMUSG00000033585 | 267,1700642 | 486,7 | 0,0 | 12,32053691 | 0,00045752 | Ndn | necdin [Source:MGI Symbol;Acc:MGI:97290] | 44,23 | 0,0 |
| ENSMUSG00000070934 | 275,9281599 | 524,6 | 0,0 | 12,39166006 | 0,001239038 | Rraga | Ras-related GTP binding A [Source:MGI Symbol;Acc:MGI:1915691] | 100,00 | 0,0 |
| ENSMUSG00000031591 | 346,301213 | 588,2 | 0,0 | 12,49748319 | 0,001283295 | Asah1 | N-acylsphingosine amidohydrolase 1 [Source:MGI Symbol;Acc:MGI:1277124] | 86,65 | 0,0 |
| ENSMUSG00000047694 | 365,9985655 | 558,1 | 0,0 | 12,52148051 | 0,001908984 | Yipf6 | Yip1 domain family, member 6 [Source:MGI Symbol;Acc:MGI:1925179] | 97,21 | 0,0 |
| ENSMUSG00000033022 | 344,6975024 | 567,6 | 0,0 | 12,52759032 | 0,0003463 | Cdo1 | cysteine dioxygenase 1, cytosolic [Source:MGI Symbol;Acc:MGI:105925] | 100,00 | 0,0 |
| ENSMUSG00000028159 | 351,9704584 | 602,7 | 0,0 | 12,58330101 | 0,000575064 | Dapp1 | dual adaptor for phosphotyrosine and 3-phosphoinositides 1 [Source:MGI Symbol;Acc:MGI:1347063] | 99,68 | 0,0 |
| ENSMUSG00000047205 | 361,6723135 | 600,2 | 0,0 | 12,62255679 | 0,001385088 | Dusp18 | dual specificity phosphatase 18 [Source:MGI Symbol;Acc:MGI:1922469] | 99,86 | 0,0 |
| ENSMUSG00000038497 | 338,1019725 | 660,7 | 0,0 | 12,69399095 | 0,000910617 | Tmco3 | transmembrane and coiled-coil domains 3 [Source:MGI Symbol;Acc:MGI:2444946] | 82,67 | 0,0 |
| ENSMUSG00000028443 | 399,1859145 | 658,0 | 0,0 | 12,7445966 | 0,000323542 | Nudt2 | nudix (nucleoside diphosphate linked moiety X)-type motif 2 [Source:MGI Symbol;Acc:MGI:1913651] | 100,00 | 0,0 |
| ENSMUSG00000065979 | 399,0341558 | 693,2 | 0,0 | 12,82164289 | 0,000276871 | Cpped1 | calcineurin-like phosphoesterase domain containing 1 [Source:MGI Symbol;Acc:MGI:2443300] | 96,72 | 0,0 |
| ENSMUSG00000025656 | 574,5060606 | 702,5 | 0,0 | 12,82833714 | 0,000613008 | Arhgef9 | CDC42 guanine nucleotide exchange factor (GEF) 9 [Source:MGI Symbol;Acc:MGI:2442233] | 65,71 | 0,0 |
| ENSMUSG00000027488 | 541,5952148 | 910,2 | 0,0 | 13,21805002 | 0,000652685 | Snta1 | syntrophin, acidic 1 [Source:MGI Symbol;Acc:MGI:101772] | 99,24 | 0,0 |
| ENSMUSG00000022261 | 569,4634829 | 1012,6 | 0,0 | 13,33260589 | 0,000625913 | Sdc2 | syndecan 2 [Source:MGI Symbol;Acc:MGI:1349165] | 99,08 | 0,0 |
| ENSMUSG00000025509 | 810,3082676 | 1856,8 | 0,0 | 14,21350216 | 1,76206E-05 | Pnpla2 | patatin-like phospholipase domain containing 2 [Source:MGI Symbol;Acc:MGI:1914103] | 85,24 | 0,0 |

Table S2.2

| Id | Associated Gene Name | Description | Mean read PAP | Mean PAP exon Coverage |
|---|---|---|---|---|
| ENSMUSG00000050708 | Ftl1 | ferritin light polypeptide 1 [Source:MGI Symbol;Acc:MGI:95589] | 50106,7 | 87,1 |
| ENSMUSG00000037742 | Eef1a1 | eukaryotic translation elongation factor 1 alpha 1 [Source:MGI Symbol;Acc:MGI:1096881] | 46682,3 | 71,1 |
| ENSMUSG00000000184 | Ccnd2 | cyclin D2 [Source:MGI Symbol;Acc:MGI:88314] | 39873,2 | 93,2 |
| ENSMUSG00000037852 | Cpe | carboxypeptidase E [Source:MGI Symbol;Acc:MGI:101932] | 36749,6 | 100,0 |
| ENSMUSG00000032399 | Rpl4 | ribosomal protein L4 [Source:MGI Symbol;Acc:MGI:1915141] | 31493,7 | 98,3 |
| ENSMUSG00000024661 | Fth1 | ferritin heavy polypeptide 1 [Source:MGI Symbol;Acc:MGI:95588] | 25673,0 | 100,0 |
| ENSMUSG00000061808 | Ttr | transthyretin [Source:MGI Symbol;Acc:MGI:98865] | 25063,3 | 95,5 |
| ENSMUSG00000027447 | Cst3 | cystatin C [Source:MGI Symbol;Acc:MGI:102519] | 24711,3 | 85,1 |
| ENSMUSG00000031765 | Mt1 | metallothionein 1 [Source:MGI Symbol;Acc:MGI:97171] | 23512,3 | 100,0 |
| ENSMUSG00000034994 | Eef2 | eukaryotic translation elongation factor 2 [Source:MGI Symbol;Acc:MGI:95288] | 19645,5 | 99,5 |
| ENSMUSG00000020184 | Mdm2 | transformed mouse 3T3 cell double minute 2 [Source:MGI Symbol;Acc:MGI:96952] | 19108,8 | 97,5 |
| ENSMUSG00000007892 | Rplp1 | ribosomal protein, large, P1 [Source:MGI Symbol;Acc:MGI:1927099] | 16082,4 | 100,0 |
| ENSMUSG00000022283 | Pabpc1 | poly(A) binding protein, cytoplasmic 1 [Source:MGI Symbol;Acc:MGI:1349722] | 15751,5 | 97,6 |
| ENSMUSG00000020932 | Gfap | glial fibrillary acidic protein [Source:MGI Symbol;Acc:MGI:95697] | 15244,8 | 86,7 |
| ENSMUSG00000051510 | Mafg | v-maf musculoaponeurotic fibrosarcoma oncogene family, protein G (avian) [Source:MGI Symbol;Acc:MGI:96911] | 14556,2 | 92,5 |
| ENSMUSG00000003970 | Rpl8 | ribosomal protein L8 [Source:MGI Symbol;Acc:MGI:1350927] | 13267,5 | 99,0 |
| ENSMUSG00000030744 | Rps3 | ribosomal protein S3 [Source:MGI Symbol;Acc:MGI:1350917] | 12966,5 | 95,2 |
| ENSMUSG00000025794 | Rpl14 | ribosomal protein L14 [Source:MGI Symbol;Acc:MGI:1914365] | 12940,7 | 96,5 |
| ENSMUSG00000020372 | Gnb2l1 | guanine nucleotide binding protein (G protein), beta polypeptide 2 like 1 [Source:MGI Symbol;Acc:MGI:101849] | 12434,4 | 82,7 |
| ENSMUSG00000067274 | Rplp0 | ribosomal protein, large, P0 [Source:MGI Symbol;Acc:MGI:1927636] | 12313,4 | 77,0 |
| ENSMUSG00000057113 | Npm1 | nucleophosmin 1 [Source:MGI Symbol;Acc:MGI:106184] | 12048,4 | 80,7 |
| ENSMUSG00000025428 | Atp5a1 | ATP synthase, H+ transporting, mitochondrial F1 complex, alpha subunit 1 [Source:MGI Symbol;Acc:MGI:88115] | 11614,7 | 84,6 |
| ENSMUSG00000074129 | Rpl13a | ribosomal protein L13A [Source:MGI Symbol;Acc:MGI:1351455] | 11325,3 | 99,4 |
| ENSMUSG00000031342 | Gpm6b | glycoprotein m6b [Source:MGI Symbol;Acc:MGI:107672] | 11195,8 | 97,9 |
| ENSMUSG00000102349 | Gm37376 | predicted gene, 37376 [Source:MGI Symbol;Acc:MGI:5610604] | 10992,7 | 100,0 |
| ENSMUSG00000004558 | Ndrg2 | N-myc downstream regulated gene 2 [Source:MGI Symbol;Acc:MGI:1352498] | 10834,1 | 95,8 |
| ENSMUSG00000028211 | Trp53inp1 | transformation related protein 53 inducible nuclear protein 1 [Source:MGI Symbol;Acc:MGI:1926609] | 10260,5 | 98,6 |
| ENSMUSG00000017390 | Aldoc | aldolase C, fructose-bisphosphate [Source:MGI Symbol;Acc:MGI:101863] | 9857,7 | 82,3 |
| ENSMUSG00000020423 | Btg2 | B cell translocation gene 2, anti-proliferative [Source:MGI Symbol;Acc:MGI:108384] | 9732,1 | 95,4 |
| ENSMUSG00000020460 | Rps27a | ribosomal protein S27A [Source:MGI Symbol;Acc:MGI:1925544] | 9248,6 | 71,7 |
| ENSMUSG00000025203 | Scd2 | stearoyl-Coenzyme A desaturase 2 [Source:MGI Symbol;Acc:MGI:98240] | 9057,2 | 92,1 |
| ENSMUSG00000008682 | Rpl10 | ribosomal protein L10 [Source:MGI Symbol;Acc:MGI:105943] | 8938,8 | 79,8 |
| ENSMUSG00000067149 | Jchain | immunoglobulin joining chain [Source:MGI Symbol;Acc:MGI:96493] | 8893,5 | 96,1 |
| ENSMUSG00000023944 | Hsp90ab1 | heat shock protein 90 alpha (cytosolic), class B member 1 [Source:MGI Symbol;Acc:MGI:96247] | 8787,9 | 95,8 |
| ENSMUSG00000037503 | Fam168b | family with sequence similarity 168, member B [Source:MGI Symbol;Acc:MGI:2448487] | 8599,9 | 96,4 |
| ENSMUSG00000071415 | Rpl23 | ribosomal protein L23 [Source:MGI Symbol;Acc:MGI:1929455] | 8521,2 | 88,6 |
| ENSMUSG00000032199 | Polr2m | polymerase (RNA) II (DNA directed) polypeptide M [Source:MGI Symbol;Acc:MGI:107282] | 7673,8 | 92,9 |
| ENSMUSG00000012848 | Rps5 | ribosomal protein S5 [Source:MGI Symbol;Acc:MGI:1097682] | 7638,9 | 87,9 |
| ENSMUSG00000028333 | Anp32b | acidic (leucine-rich) nuclear phosphoprotein 32 family, member B [Source:MGI Symbol;Acc:MGI:1914878] | 7480,6 | 86,0 |
| ENSMUSG00000000740 | Rpl13 | ribosomal protein L13 [Source:MGI Symbol;Acc:MGI:105922] | 7298,1 | 79,1 |
| ENSMUSG00000034892 | Rps29 | ribosomal protein S29 [Source:MGI Symbol;Acc:MGI:107681] | 7113,4 | 79,7 |
| ENSMUSG00000058655 | Eif4b | eukaryotic translation initiation factor 4B [Source:MGI Symbol;Acc:MGI:95304] | 7109,0 | 94,5 |
| ENSMUSG00000005610 | Eif4g2 | eukaryotic translation initiation factor 4, gamma 2 [Source:MGI Symbol;Acc:MGI:109207] | 7064,9 | 77,1 |
| ENSMUSG00000031760 | Mt3 | metallothionein 3 [Source:MGI Symbol;Acc:MGI:97173] | 6913,5 | 81,1 |
| ENSMUSG00000003429 | Rps11 | ribosomal protein S11 [Source:MGI Symbol;Acc:MGI:1351329] | 6890,2 | 83,4 |
| ENSMUSG00000017404 | Rpl19 | ribosomal protein L19 [Source:MGI Symbol;Acc:MGI:98020] | 6821,4 | 85,4 |
| ENSMUSG00000046330 | Rpl37a | ribosomal protein L37a [Source:MGI Symbol;Acc:MGI:98068] | 6688,8 | 99,6 |
| ENSMUSG00000059070 | Rpl18 | ribosomal protein L18 [Source:MGI Symbol;Acc:MGI:98003] | 6651,3 | 88,8 |
| ENSMUSG00000053279 | Aldh1a1 | aldehyde dehydrogenase family 1, subfamily A1 [Source:MGI Symbol;Acc:MGI:1353450] | 6567,6 | 98,3 |
| ENSMUSG00000030189 | Ybx3 | Y box protein 3 [Source:MGI Symbol;Acc:MGI:2137670] | 6332,0 | 99,5 |
| ENSMUSG00000021792 | Fam213a | family with sequence similarity 213, member A [Source:MGI Symbol;Acc:MGI:1917814] | 6162,4 | 73,7 |
| ENSMUSG00000005360 | Slc1a3 | solute carrier family 1 (glial high affinity glutamate transporter), member 3 [Source:MGI Symbol;Acc:MGI:99917] | 6135,4 | 92,6 |
| ENSMUSG00000054226 | Tprkb | Tp53rk binding protein [Source:MGI Symbol;Acc:MGI:1917036] | 6017,9 | 73,3 |
| ENSMUSG00000068039 | Tcp1 | t-complex protein 1 [Source:MGI Symbol;Acc:MGI:98535] | 5987,1 | 82,8 |
| ENSMUSG00000029309 | Sparcl1 | SPARC-like 1 [Source:MGI Symbol;Acc:MGI:108110] | 5886,9 | 98,0 |
| ENSMUSG00000033208 | S100b | S100 protein, beta polypeptide, neural [Source:MGI Symbol;Acc:MGI:98217] | 5755,8 | 100,0 |
| ENSMUSG00000078812 | Eif5a | eukaryotic translation initiation factor 5A [Source:MGI Symbol;Acc:MGI:106248] | 5701,5 | 70,3 |
| ENSMUSG00000001175 | Calm1 | calmodulin 1 [Source:MGI Symbol;Acc:MGI:88251] | 5669,9 | 78,5 |
| ENSMUSG00000035885 | Cox8a | cytochrome c oxidase subunit VIIIa [Source:MGI Symbol;Acc:MGI:105959] | 5661,2 | 87,3 |
| ENSMUSG00000043131 | Mob1a | MOB kinase activator 1A [Source:MGI Symbol;Acc:MGI:2442631] | 5588,2 | 96,1 |
| ENSMUSG00000013698 | Pea15a | phosphoprotein enriched in astrocytes 15A [Source:MGI Symbol;Acc:MGI:104799] | 5516,3 | 89,8 |
| ENSMUSG00000044349 | Snhg11 | small nucleolar RNA host gene 11 [Source:MGI Symbol;Acc:MGI:2441845] | 5430,4 | 94,0 |
| ENSMUSG00000058135 | Gstm1 | glutathione S-transferase, mu 1 [Source:MGI Symbol;Acc:MGI:95860] | 5406,5 | 86,7 |
| ENSMUSG00000025393 | Atp5b | ATP synthase, H+ transporting mitochondrial F1 complex, beta subunit [Source:MGI Symbol;Acc:MGI:107801] | 5395,0 | 92,9 |
| ENSMUSG00000031320 | Rps4x | ribosomal protein S4, X-linked [Source:MGI Symbol;Acc:MGI:98158] | 5246,0 | 86,3 |
| ENSMUSG00000033713 | Foxn3 | forkhead box N3 [Source:MGI Symbol;Acc:MGI:1918625] | 5242,0 | 85,4 |
| ENSMUSG00000057863 | Rpl36 | ribosomal protein L36 [Source:MGI Symbol;Acc:MGI:1860603] | 5232,3 | 88,0 |
| ENSMUSG00000031818 | Cox4i1 | cytochrome c oxidase subunit IV isoform 1 [Source:MGI Symbol;Acc:MGI:88473] | 5232,2 | 96,1 |
| ENSMUSG00000018983 | E2f2 | E2F transcription factor 2 [Source:MGI Symbol;Acc:MGI:1096341] | 5206,7 | 85,7 |
| ENSMUSG00000055302 | Mrfap1 | Morf4 family associated protein 1 [Source:MGI Symbol;Acc:MGI:1914818] | 5173,9 | 79,1 |
| ENSMUSG00000090625 | Gm20721 | predicted gene, 20721 [Source:MGI Symbol;Acc:MGI:5313030] | 5150,8 | 99,1 |
| ENSMUSG00000009927 | Rps25 | ribosomal protein S25 [Source:MGI Symbol;Acc:MGI:1922867] | 5100,5 | 80,3 |
| ENSMUSG00000018770 | Atp5g3 | ATP synthase, H+ transporting, mitochondrial F0 complex, subunit C3 (subunit 9) [Source:MGI Symbol;Acc:MGI:2442035 | 5031,5 | 100,0 |
| ENSMUSG00000062078 | Qk | quaking [Source:MGI Symbol;Acc:MGI:97837] | 4984,2 | 71,4 |
| ENSMUSG00000038612 | Mcl1 | myeloid cell leukemia sequence 1 [Source:MGI Symbol;Acc:MGI:101769] | 4907,4 | 99,0 |
| ENSMUSG00000047675 | Rps8 | ribosomal protein S8 [Source:MGI Symbol;Acc:MGI:98166] | 4826,7 | 80,5 |
| ENSMUSG00000064373 | Sepp1 | selenoprotein P, plasma, 1 [Source:MGI Symbol;Acc:MGI:894288] | 4823,9 | 76,7 |
| ENSMUSG00000063457 | Rps15 | ribosomal protein S15 [Source:MGI Symbol;Acc:MGI:98117] | 4788,3 | 86,2 |
| ENSMUSG00000107379 | Gm43126 | predicted gene 43126 [Source:MGI Symbol;Acc:MGI:5663263] | 4706,2 | 86,7 |
| ENSMUSG00000091228 | Gm20390 | predicted gene 20390 [Source:MGI Symbol;Acc:MGI:5141855] | 4682,5 | 97,4 |
| ENSMUSG00000062762 | Ei24 | etoposide induced 2.4 mRNA [Source:MGI Symbol;Acc:MGI:108090] | 4645,9 | 94,5 |
| ENSMUSG00000052397 | Ezr | ezrin [Source:MGI Symbol;Acc:MGI:98931] | 4642,6 | 91,7 |
| ENSMUSG00000079435 | Rpl36a | ribosomal protein L36A [Source:MGI Symbol;Acc:MGI:1201789] | 4619,8 | 93,0 |
| ENSMUSG00000032053 | Pou2af1 | POU domain, class 2, associating factor 1 [Source:MGI Symbol;Acc:MGI:105086] | 4579,7 | 100,0 |
| ENSMUSG00000027893 | Ahcyl1 | S-adenosylhomocysteine hydrolase-like 1 [Source:MGI Symbol;Acc:MGI:2385184] | 4578,2 | 93,1 |
| ENSMUSG00000001270 | Ckb | creatine kinase, brain [Source:MGI Symbol;Acc:MGI:88407] | 4548,4 | 95,7 |
| ENSMUSG00000030428 | Ttyh1 | tweety homolog 1 (Drosophila) [Source:MGI Symbol;Acc:MGI:1889007] | 4528,4 | 85,7 |
| ENSMUSG00000039001 | Rps21 | ribosomal protein S21 [Source:MGI Symbol;Acc:MGI:1913731] | 4487,3 | 99,2 |
| ENSMUSG00000022425 | Enpp2 | ectonucleotide pyrophosphatase/phosphodiesterase 2 [Source:MGI Symbol;Acc:MGI:1321390] | 4468,5 | 88,0 |
| ENSMUSG00000032294 | Pkm | pyruvate kinase, muscle [Source:MGI Symbol;Acc:MGI:97591] | 4460,3 | 89,1 |
| ENSMUSG00000030104 | Edem1 | ER degradation enhancer, mannosidase alpha-like 1 [Source:MGI Symbol;Acc:MGI:2180139] | 4430,7 | 95,8 |
| ENSMUSG00000010663 | Fads1 | fatty acid desaturase 1 [Source:MGI Symbol;Acc:MGI:1923517] | 4405,5 | 97,2 |
| ENSMUSG00000038274 | Fau | el-Biskis-Reilly murine sarcoma virus (FBR-MuSV) ubiquitously expressed (fox derived) [Source:MGI Symbol;Acc:MGI:102 | 4377,7 | 85,6 |
| ENSMUSG00000031762 | Mt2 | metallothionein 2 [Source:MGI Symbol;Acc:MGI:97172] | 4348,8 | 89,0 |
| ENSMUSG00000049354 | Dcaf7 | DDB1 and CUL4 associated factor 7 [Source:MGI Symbol;Acc:MGI:1919083] | 4331,6 | 86,6 |
| ENSMUSG00000005089 | Slc1a2 | solute carrier family 1 (glial high affinity glutamate transporter), member 2 [Source:MGI Symbol;Acc:MGI:101931] | 4289,4 | 73,5 |
| ENSMUSG00000011960 | Ccnt1 | cyclin T1 [Source:MGI Symbol;Acc:MGI:1328363] | 4239,8 | 82,0 |
| ENSMUSG00000027479 | Mapre1 | microtubule-associated protein, RP/EB family, member 1 [Source:MGI Symbol;Acc:MGI:891995] | 4213,5 | 81,2 |
| ENSMUSG00000025871 | 4833439L19Rik | RIKEN cDNA 4833439L19 gene [Source:MGI Symbol;Acc:MGI:1921162] | 4171,2 | 94,9 |
| ENSMUSG00000004032 | Gstm5 | glutathione S-transferase, mu 5 [Source:MGI Symbol;Acc:MGI:1309466] | 4169,9 | 81,5 |
| ENSMUSG00000038900 | Rpl12 | ribosomal protein L12 [Source:MGI Symbol;Acc:MGI:98002] | 4157,2 | 74,1 |
| ENSMUSG00000018583 | G3bp1 | GTPase activating protein (SH3 domain) binding protein 1 [Source:MGI Symbol;Acc:MGI:1351465] | 4112,1 | 80,1 |
| ENSMUSG00000061787 | Rps17 | ribosomal protein S17 [Source:MGI Symbol;Acc:MGI:1309526] | 4096,3 | 91,0 |
| ENSMUSG00000024991 | Eif3a | eukaryotic translation initiation factor 3, subunit A [Source:MGI Symbol;Acc:MGI:95301] | 4058,8 | 92,4 |
| ENSMUSG00000020611 | Gna13 | guanine nucleotide binding protein, alpha 13 [Source:MGI Symbol;Acc:MGI:95768] | 3986,4 | 91,8 |
| ENSMUSG00000057322 | Rpl38 | ribosomal protein L38 [Source:MGI Symbol;Acc:MGI:1914921] | 3961,9 | 81,2 |
| ENSMUSG00000022108 | Itm2b | integral membrane protein 2B [Source:MGI Symbol;Acc:MGI:1309517] | 3942,4 | 99,9 |
| ENSMUSG00000020267 | Hint1 | histidine triad nucleotide binding protein 1 [Source:MGI Symbol;Acc:MGI:1321133] | 3940,5 | 72,7 |
| ENSMUSG00000020849 | Ywhae | e 3-monooxygenase/tryptophan 5-monooxygenase activation protein, epsilon polypeptide [Source:MGI Symbol;Acc:MGI:8 | 3931,7 | 89,0 |
| ENSMUSG00000026434 | Nucks1 | nuclear casein kinase and cyclin-dependent kinase substrate 1 [Source:MGI Symbol;Acc:MGI:1934811] | 3922,2 | 90,0 |
| ENSMUSG00000022890 | Atp5j | ATP synthase, H+ transporting, mitochondrial F0 complex, subunit F [Source:MGI Symbol;Acc:MGI:107777] | 3876,8 | 71,9 |
| ENSMUSG00000053094 | Tmem248 | transmembrane protein 248 [Source:MGI Symbol;Acc:MGI:1918917] | 3860,8 | 87,1 |
| ENSMUSG00000028416 | Bag1 | BCL2-associated athanogene 1 [Source:MGI Symbol;Acc:MGI:108047] | 3853,3 | 97,5 |
| ENSMUSG00000020857 | Nme2 | NME/NM23 nucleoside diphosphate kinase 2 [Source:MGI Symbol;Acc:MGI:97356] | 3835,6 | 72,3 |
| ENSMUSG00000046364 | Rpl27a | ribosomal protein L27A [Source:MGI Symbol;Acc:MGI:1347076] | 3811,3 | 91,7 |
| ENSMUSG00000044468 | Fam46c | family with sequence similarity 46, member C [Source:MGI Symbol;Acc:MGI:1921895] | 3776,5 | 80,7 |

| | | | | |
|---|---|---|---|---|
| ENSMUSG00000031985 | Gnpat | glyceronephosphate O-acyltransferase [Source:MGI Symbol;Acc:MGI:1343460] | 3634,2 | 98,5 |
| ENSMUSG00000032050 | Rdx | radixin [Source:MGI Symbol;Acc:MGI:97887] | 3631,2 | 72,8 |
| ENSMUSG00000024213 | Nudt3 | nudix (nucleotide diphosphate linked moiety X)-type motif 3 [Source:MGI Symbol;Acc:MGI:1928484] | 3613,4 | 89,4 |
| ENSMUSG00000030224 | Strap | serine/threonine kinase receptor associated protein [Source:MGI Symbol;Acc:MGI:1329037] | 3608,6 | 96,1 |
| ENSMUSG00000056612 | Ppp1r14b | protein phosphatase 1, regulatory (inhibitor) subunit 14B [Source:MGI Symbol;Acc:MGI:107682] | 3595,9 | 70,3 |
| ENSMUSG00000059325 | Hopx | HOP homeobox [Source:MGI Symbol;Acc:MGI:1916782] | 3578,5 | 85,5 |
| ENSMUSG00000020591 | Ntsr2 | neurotensin receptor 2 [Source:MGI Symbol;Acc:MGI:108018] | 3576,6 | 97,8 |
| ENSMUSG00000036781 | Rps27l | ribosomal protein S27-like [Source:MGI Symbol;Acc:MGI:1915191] | 3537,6 | 94,5 |
| ENSMUSG00000049517 | Rps23 | ribosomal protein S23 [Source:MGI Symbol;Acc:MGI:1913725] | 3493,2 | 86,6 |
| ENSMUSG00000041571 | Sepw1 | selenoprotein W, muscle 1 [Source:MGI Symbol;Acc:MGI:1100878] | 3486,4 | 96,5 |
| ENSMUSG00000062328 | Rpl17 | ribosomal protein L17 [Source:MGI Symbol;Acc:MGI:2448270] | 3448,9 | 87,5 |
| ENSMUSG00000008668 | Rps18 | ribosomal protein S18 [Source:MGI Symbol;Acc:MGI:98146] | 3441,1 | 71,8 |
| ENSMUSG00000025290 | Rps24 | ribosomal protein S24 [Source:MGI Symbol;Acc:MGI:98147] | 3408,9 | 88,2 |
| ENSMUSG00000021282 | Eif5 | eukaryotic translation initiation factor 5 [Source:MGI Symbol;Acc:MGI:95309] | 3384,4 | 86,2 |
| ENSMUSG00000007097 | Atp1a2 | ATPase, Na+/K+ transporting, alpha 2 polypeptide [Source:MGI Symbol;Acc:MGI:88106] | 3366,1 | 86,9 |
| ENSMUSG00000020362 | Cnot6 | CCR4-NOT transcription complex, subunit 6 [Source:MGI Symbol;Acc:MGI:2144529] | 3361,0 | 87,5 |
| ENSMUSG00000018446 | C1qbp | complement component 1, q subcomponent binding protein [Source:MGI Symbol;Acc:MGI:1194505] | 3348,3 | 98,1 |
| ENSMUSG00000090000 | Ier3ip1 | immediate early response 3 interacting protein 1 [Source:MGI Symbol;Acc:MGI:1913441] | 3334,1 | 98,4 |
| ENSMUSG00000063882 | Uqcrh | ubiquinol-cytochrome c reductase hinge protein [Source:MGI Symbol;Acc:MGI:1913826] | 3328,3 | 98,4 |
| ENSMUSG00000031517 | Gpm6a | glycoprotein m6a [Source:MGI Symbol;Acc:MGI:107671] | 3318,6 | 83,6 |
| ENSMUSG00000041133 | Smc1a | structural maintenance of chromosomes 1A [Source:MGI Symbol;Acc:MGI:1344345] | 3303,9 | 82,8 |
| ENSMUSG00000053565 | Eif3k | eukaryotic translation initiation factor 3, subunit K [Source:MGI Symbol;Acc:MGI:1921080] | 3282,0 | 71,6 |
| ENSMUSG00000024782 | Ak3 | adenylate kinase 3 [Source:MGI Symbol;Acc:MGI:1860835] | 3247,4 | 97,2 |
| ENSMUSG00000035268 | Pkig | protein kinase inhibitor, gamma [Source:MGI Symbol;Acc:MGI:1343086] | 3203,0 | 83,1 |
| ENSMUSG00000025362 | Rps26 | ribosomal protein S26 [Source:MGI Symbol;Acc:MGI:1351628] | 3194,6 | 97,7 |
| ENSMUSG00000031246 | Sh3bgrl | SH3-binding domain glutamic acid-rich protein like [Source:MGI Symbol;Acc:MGI:1930849] | 3111,3 | 95,3 |
| ENSMUSG00000040945 | Rcc2 | regulator of chromosome condensation 2 [Source:MGI Symbol;Acc:MGI:1919784] | 3091,0 | 92,7 |
| ENSMUSG00000041740 | Rnf10 | ring finger protein 10 [Source:MGI Symbol;Acc:MGI:1859162] | 3067,1 | 89,5 |
| ENSMUSG00000029621 | Arpc1a | actin related protein 2/3 complex, subunit 1A [Source:MGI Symbol;Acc:MGI:1928896] | 3059,2 | 90,4 |
| ENSMUSG00000026424 | Gpr37l1 | G protein-coupled receptor 37-like 1 [Source:MGI Symbol;Acc:MGI:1928503] | 3059,1 | 94,2 |
| ENSMUSG00000032883 | Acsl3 | acyl-CoA synthetase long-chain family member 3 [Source:MGI Symbol;Acc:MGI:1921455] | 3049,8 | 95,6 |
| ENSMUSG00000026457 | Adipor1 | adiponectin receptor 1 [Source:MGI Symbol;Acc:MGI:1919924] | 3040,0 | 70,7 |
| ENSMUSG00000022186 | Oxct1 | 3-oxoacid CoA transferase 1 [Source:MGI Symbol;Acc:MGI:1914291] | 3011,9 | 95,9 |
| ENSMUSG00000021750 | Fam107a | family with sequence similarity 107, member A [Source:MGI Symbol;Acc:MGI:3041256] | 3003,6 | 77,4 |
| ENSMUSG00000022336 | Eif3e | eukaryotic translation initiation factor 3, subunit E [Source:MGI Symbol;Acc:MGI:99257] | 2994,5 | 97,4 |
| ENSMUSG00000029364 | Wsb2 | WD repeat and SOCS box-containing 2 [Source:MGI Symbol;Acc:MGI:2144041] | 2987,5 | 90,1 |
| ENSMUSG00000038388 | Mpp6 | membrane protein, palmitoylated 6 (MAGUK p55 subfamily member 6) [Source:MGI Symbol;Acc:MGI:1927340] | 2980,5 | 74,5 |
| ENSMUSG00000061904 | Slc25a3 | solute carrier family 25 (mitochondrial carrier, phosphate carrier), member 3 [Source:MGI Symbol;Acc:MGI:1353498] | 2921,1 | 79,8 |
| ENSMUSG00000017639 | Rab11fip4 | RAB11 family interacting protein 4 (class II) [Source:MGI Symbol;Acc:MGI:2442920] | 2901,8 | 88,4 |
| ENSMUSG00000050953 | Gja1 | gap junction protein, alpha 1 [Source:MGI Symbol;Acc:MGI:95713] | 2885,5 | 98,6 |
| ENSMUSG00000028639 | Ybx1 | Y box protein 1 [Source:MGI Symbol;Acc:MGI:99146] | 2884,8 | 89,1 |
| ENSMUSG00000001525 | Tubb5 | tubulin, beta 5 class I [Source:MGI Symbol;Acc:MGI:107812] | 2857,5 | 87,7 |
| ENSMUSG00000062014 | Gmfb | glia maturation factor, beta [Source:MGI Symbol;Acc:MGI:1927133] | 2857,3 | 87,8 |
| ENSMUSG00000023456 | Tpi1 | triosephosphate isomerase 1 [Source:MGI Symbol;Acc:MGI:98797] | 2830,2 | 80,2 |
| ENSMUSG00000063229 | Ldha | lactate dehydrogenase A [Source:MGI Symbol;Acc:MGI:96759] | 2819,1 | 72,8 |
| ENSMUSG00000029538 | Srsf9 | serine/arginine-rich splicing factor 9 [Source:MGI Symbol;Acc:MGI:104896] | 2802,4 | 99,8 |
| ENSMUSG00000061983 | Rps12 | ribosomal protein S12 [Source:MGI Symbol;Acc:MGI:98105] | 2800,8 | 99,9 |
| ENSMUSG00000090733 | Rps27 | ribosomal protein S27 [Source:MGI Symbol;Acc:MGI:1888676] | 2792,5 | 74,9 |
| ENSMUSG00000034566 | Atp5h | ATP synthase, H+ transporting, mitochondrial F0 complex, subunit D [Source:MGI Symbol;Acc:MGI:1918929] | 2790,1 | 84,3 |
| ENSMUSG00000028788 | Ptp4a2 | protein tyrosine phosphatase 4a2 [Source:MGI Symbol;Acc:MGI:1277117] | 2774,9 | 88,1 |
| ENSMUSG00000000346 | Dazap2 | DAZ associated protein 2 [Source:MGI Symbol;Acc:MGI:1344344] | 2772,7 | 99,8 |
| ENSMUSG00000026112 | Coa5 | cytochrome C oxidase assembly factor 5 [Source:MGI Symbol;Acc:MGI:1923428] | 2766,8 | 98,2 |
| ENSMUSG00000025130 | P4hb | prolyl 4-hydroxylase, beta polypeptide [Source:MGI Symbol;Acc:MGI:97464] | 2762,3 | 86,6 |
| ENSMUSG00000057522 | Spop | speckle-type POZ protein [Source:MGI Symbol;Acc:MGI:1343085] | 2760,1 | 97,7 |
| ENSMUSG00000005683 | Cs | citrate synthase [Source:MGI Symbol;Acc:MGI:88529] | 2729,6 | 98,6 |
| ENSMUSG00000020444 | Guk1 | guanylate kinase 1 [Source:MGI Symbol;Acc:MGI:95871] | 2726,3 | 85,6 |
| ENSMUSG00000024014 | Pim1 | proviral integration site 1 [Source:MGI Symbol;Acc:MGI:97584] | 2725,6 | 95,0 |
| ENSMUSG00000020088 | Sar1a | SAR1 gene homolog A (S. cerevisiae) [Source:MGI Symbol;Acc:MGI:98230] | 2717,1 | 99,3 |
| ENSMUSG00000027630 | Tbl1xr1 | transducin (beta)-like 1X-linked receptor 1 [Source:MGI Symbol;Acc:MGI:2441730] | 2697,4 | 78,4 |
| ENSMUSG00000037706 | Cd81 | CD81 antigen [Source:MGI Symbol;Acc:MGI:1096398] | 2678,5 | 89,8 |
| ENSMUSG00000035891 | Cerk | ceramide kinase [Source:MGI Symbol;Acc:MGI:2386052] | 2674,7 | 86,3 |
| ENSMUSG00000034024 | Cct2 | chaperonin containing Tcp1, subunit 2 (beta) [Source:MGI Symbol;Acc:MGI:107186] | 2666,9 | 94,6 |
| ENSMUSG00000030057 | Cnbp | cellular nucleic acid binding protein [Source:MGI Symbol;Acc:MGI:88431] | 2662,1 | 71,5 |
| ENSMUSG00000074093 | Svip | small VCP/p97-interacting protein [Source:MGI Symbol;Acc:MGI:1922994] | 2657,0 | 85,6 |
| ENSMUSG00000054452 | Aes | amino-terminal enhancer of split [Source:MGI Symbol;Acc:MGI:95806] | 2653,5 | 99,3 |
| ENSMUSG00000010608 | Rbm25 | RNA binding motif protein 25 [Source:MGI Symbol;Acc:MGI:1914289] | 2643,9 | 70,7 |
| ENSMUSG00000026932 | Nacc2 | nucleus accumbens associated 2, BEN and BTB (POZ) domain containing [Source:MGI Symbol;Acc:MGI:1915241] | 2636,8 | 85,0 |
| ENSMUSG00000041028 | Ghitm | growth hormone inducible transmembrane protein [Source:MGI Symbol;Acc:MGI:1913342] | 2634,5 | 85,5 |
| ENSMUSG00000026421 | Csrp1 | cysteine and glycine-rich protein 1 [Source:MGI Symbol;Acc:MGI:88549] | 2631,6 | 87,4 |
| ENSMUSG00000022234 | Cct5 | chaperonin containing Tcp1, subunit 5 (epsilon) [Source:MGI Symbol;Acc:MGI:107185] | 2621,3 | 87,6 |
| ENSMUSG00000028756 | Pink1 | PTEN induced putative kinase 1 [Source:MGI Symbol;Acc:MGI:1916193] | 2621,2 | 87,0 |
| ENSMUSG00000024654 | Asrgl1 | asparaginase like 1 [Source:MGI Symbol;Acc:MGI:1913764] | 2613,1 | 81,3 |
| ENSMUSG00000016252 | Atp5e | ATP synthase, H+ transporting, mitochondrial F1 complex, epsilon subunit [Source:MGI Symbol;Acc:MGI:1855697] | 2608,5 | 98,2 |
| ENSMUSG00000040848 | Sft2d2 | SFT2 domain containing 2 [Source:MGI Symbol;Acc:MGI:1917362] | 2607,2 | 94,5 |
| ENSMUSG00000026249 | Serpine2 | serine (or cysteine) peptidase inhibitor, clade E, member 2 [Source:MGI Symbol;Acc:MGI:101780] | 2578,7 | 87,6 |
| ENSMUSG00000020321 | Mdh1 | malate dehydrogenase 1, NAD (soluble) [Source:MGI Symbol;Acc:MGI:97051] | 2560,8 | 73,2 |
| ENSMUSG00000068823 | Csde1 | cold shock domain containing E1, RNA binding [Source:MGI Symbol;Acc:MGI:92356] | 2553,9 | 79,3 |
| ENSMUSG00000015090 | Ptgds | prostaglandin D2 synthase (brain) [Source:MGI Symbol;Acc:MGI:99261] | 2553,6 | 82,4 |
| ENSMUSG00000009630 | Ppp2cb | protein phosphatase 2 (formerly 2A), catalytic subunit, beta isoform [Source:MGI Symbol;Acc:MGI:1321161] | 2542,7 | 93,9 |
| ENSMUSG00000038708 | Golga4 | golgi autoantigen, golgin subfamily a, 4 [Source:MGI Symbol;Acc:MGI:1859646] | 2532,5 | 71,6 |
| ENSMUSG00000020368 | Canx | calnexin [Source:MGI Symbol;Acc:MGI:88261] | 2521,5 | 86,8 |
| ENSMUSG00000027423 | Snx5 | sorting nexin 5 [Source:MGI Symbol;Acc:MGI:1916428] | 2500,2 | 94,4 |
| ENSMUSG00000024353 | Mzb1 | marginal zone B and B1 cell-specific protein 1 [Source:MGI Symbol;Acc:MGI:1917066] | 2498,4 | 100,0 |
| ENSMUSG00000042305 | Tmem183a | transmembrane protein 183A [Source:MGI Symbol;Acc:MGI:1914729] | 2495,1 | 96,5 |
| ENSMUSG00000020163 | Uqcr11 | ubiquinol-cytochrome c reductase, complex III subunit XI [Source:MGI Symbol;Acc:MGI:1913844] | 2475,3 | 100,0 |
| ENSMUSG00000026028 | Trak2 | trafficking protein, kinesin binding 2 [Source:MGI Symbol;Acc:MGI:1918077] | 2473,6 | 81,7 |
| ENSMUSG00000028692 | Akr1a1 | aldo-keto reductase family 1, member A1 (aldehyde reductase) [Source:MGI Symbol;Acc:MGI:1929955] | 2469,9 | 91,3 |
| ENSMUSG00000059534 | Uqcr10 | ubiquinol-cytochrome c reductase, complex III subunit X [Source:MGI Symbol;Acc:MGI:1913402] | 2461,0 | 100,0 |
| ENSMUSG00000031839 | Hsbp1 | heat shock factor binding protein 1 [Source:MGI Symbol;Acc:MGI:1915446] | 2454,4 | 99,8 |
| ENSMUSG00000036309 | Skp1a | S-phase kinase-associated protein 1A [Source:MGI Symbol;Acc:MGI:103575] | 2451,8 | 85,5 |
| ENSMUSG00000014769 | Psmb1 | proteasome (prosome, macropain) subunit, beta type 1 [Source:MGI Symbol;Acc:MGI:104884] | 2427,9 | 99,0 |
| ENSMUSG00000036751 | Cox6b1 | cytochrome c oxidase, subunit VIb polypeptide 1 [Source:MGI Symbol;Acc:MGI:107460] | 2425,0 | 81,7 |
| ENSMUSG00000033499 | Larp4b | La ribonucleoprotein domain family, member 4B [Source:MGI Symbol;Acc:MGI:106330] | 2422,9 | 89,0 |
| ENSMUSG00000025132 | Arhgdia | Rho GDP dissociation inhibitor (GDI) alpha [Source:MGI Symbol;Acc:MGI:2178103] | 2405,9 | 91,0 |
| ENSMUSG00000026626 | Ppp2r5a | protein phosphatase 2, regulatory subunit B', alpha [Source:MGI Symbol;Acc:MGI:2388479] | 2392,4 | 94,0 |
| ENSMUSG00000037266 | Rsrp1 | arginine/serine rich protein 1 [Source:MGI Symbol;Acc:MGI:106498] | 2387,2 | 87,6 |
| ENSMUSG00000021356 | Irf4 | interferon regulatory factor 4 [Source:MGI Symbol;Acc:MGI:1096873] | 2385,5 | 71,6 |
| ENSMUSG00000032320 | Rcn2 | reticulocalbin 2 [Source:MGI Symbol;Acc:MGI:1349765] | 2360,6 | 88,9 |
| ENSMUSG00000064339 | mt-Rnr2 | mitochondrially encoded 16S rRNA [Source:MGI Symbol;Acc:MGI:102492] | 2355,1 | 99,1 |
| ENSMUSG00000064370 | mt-Cytb | mitochondrially encoded cytochrome b [Source:MGI Symbol;Acc:MGI:102501] | 2333,8 | 99,9 |
| ENSMUSG00000045092 | S1pr1 | sphingosine-1-phosphate receptor 1 [Source:MGI Symbol;Acc:MGI:1096355] | 2331,6 | 95,0 |
| ENSMUSG00000032601 | Prkar2a | protein kinase, cAMP dependent regulatory, type II alpha [Source:MGI Symbol;Acc:MGI:108025] | 2315,5 | 82,8 |
| ENSMUSG00000014294 | Ndufa2 | NADH dehydrogenase (ubiquinone) 1 alpha subcomplex, 2 [Source:MGI Symbol;Acc:MGI:1343103] | 2312,9 | 93,8 |
| ENSMUSG00000007739 | Cct4 | chaperonin containing Tcp1, subunit 4 (delta) [Source:MGI Symbol;Acc:MGI:104689] | 2302,2 | 77,7 |
| ENSMUSG00000029472 | Anapc5 | anaphase-promoting complex subunit 5 [Source:MGI Symbol;Acc:MGI:1929722] | 2287,7 | 89,4 |
| ENSMUSG00000022253 | Nadk2 | NAD kinase 2, mitochondrial [Source:MGI Symbol;Acc:MGI:1915896] | 2283,6 | 75,4 |
| ENSMUSG00000000563 | Atp5f1 | ATP synthase, H+ transporting, mitochondrial F0 complex, subunit B1 [Source:MGI Symbol;Acc:MGI:1100495] | 2279,6 | 79,0 |
| ENSMUSG00000024516 | Sec11c | SEC11 homolog C, signal peptidase complex subunit [Source:MGI Symbol;Acc:MGI:1913536] | 2272,9 | 96,2 |
| ENSMUSG00000032324 | Tspan3 | tetraspanin 3 [Source:MGI Symbol;Acc:MGI:1928098] | 2271,0 | 98,5 |
| ENSMUSG00000020326 | Ccng1 | cyclin G1 [Source:MGI Symbol;Acc:MGI:102890] | 2267,9 | 97,6 |
| ENSMUSG00000030007 | Cct7 | chaperonin containing Tcp1, subunit 7 (eta) [Source:MGI Symbol;Acc:MGI:107184] | 2254,5 | 75,8 |
| ENSMUSG00000026811 | St6galnac6 | acetyl-neuraminyl-2,3-beta-galactosyl-1,3)-N-acetylgalactosaminide alpha-2,6-sialyltransferase 6 [Source:MGI Symbol;Acc: | 2253,5 | 73,2 |
| ENSMUSG00000027367 | Stard7 | START domain containing 7 [Source:MGI Symbol;Acc:MGI:2139090] | 2249,1 | 77,9 |
| ENSMUSG00000006651 | Aplp1 | amyloid beta (A4) precursor-like protein 1 [Source:MGI Symbol;Acc:MGI:88046] | 2248,2 | 97,1 |
| ENSMUSG00000046562 | Unc119b | unc-119 homolog B (C. elegans) [Source:MGI Symbol;Acc:MGI:2147162] | 2246,3 | 97,8 |
| ENSMUSG00000037058 | Paip2 | polyadenylate-binding protein-interacting protein 2 [Source:MGI Symbol;Acc:MGI:1915119] | 2234,8 | 79,5 |
| ENSMUSG00000025702 | March8 | membrane-associated ring finger (C3HC4) 8 [Source:MGI Symbol;Acc:MGI:1919029] | 2229,6 | 82,9 |

| | | | | |
|---|---|---|---|---|
| ENSMUSG00000027828 | Ssr3 | signal sequence receptor, gamma [Source:MGI Symbol;Acc:MGI:1914687] | 2214,6 | 73,5 |
| ENSMUSG00000029752 | Asns | asparagine synthetase [Source:MGI Symbol;Acc:MGI:1350929] | 2202,8 | 77,7 |
| ENSMUSG00000032281 | Acsbg1 | acyl-CoA synthetase bubblegum family member 1 [Source:MGI Symbol;Acc:MGI:2385656] | 2199,7 | 76,9 |
| ENSMUSG00000001056 | Nhp2 | NHP2 ribonucleoprotein [Source:MGI Symbol;Acc:MGI:1098547] | 2195,4 | 76,9 |
| ENSMUSG00000056851 | Pcbp2 | poly(rC) binding protein 2 [Source:MGI Symbol;Acc:MGI:108202] | 2190,3 | 74,0 |
| ENSMUSG00000032966 | Fkbp1a | FK506 binding protein 1a [Source:MGI Symbol;Acc:MGI:95541] | 2187,3 | 88,2 |
| ENSMUSG00000022956 | Atp5o | ATP synthase, H+ transporting, mitochondrial F1 complex, O subunit [Source:MGI Symbol;Acc:MGI:106341] | 2167,0 | 73,2 |
| ENSMUSG00000029064 | Gnb1 | guanine nucleotide binding protein (G protein), beta 1 [Source:MGI Symbol;Acc:MGI:95781] | 2163,3 | 86,0 |
| ENSMUSG00000063015 | Ccni | cyclin I [Source:MGI Symbol;Acc:MGI:1341077] | 2161,4 | 73,2 |
| ENSMUSG00000041697 | Cox6a1 | cytochrome c oxidase subunit VIa polypeptide 1 [Source:MGI Symbol;Acc:MGI:103099] | 2159,6 | 96,4 |
| ENSMUSG00000011179 | Odc1 | ornithine decarboxylase, structural 1 [Source:MGI Symbol;Acc:MGI:97402] | 2157,0 | 81,9 |
| ENSMUSG00000038462 | Uqcrfs1 | ubiquinol-cytochrome c reductase, Rieske iron-sulfur polypeptide 1 [Source:MGI Symbol;Acc:MGI:1913944] | 2153,9 | 90,5 |
| ENSMUSG00000048388 | Fam171b | family with sequence similarity 171, member B [Source:MGI Symbol;Acc:MGI:2444579] | 2144,3 | 70,9 |
| ENSMUSG00000060450 | Rnf14 | ring finger protein 14 [Source:MGI Symbol;Acc:MGI:1929668] | 2142,1 | 91,2 |
| ENSMUSG00000019370 | Calm3 | calmodulin 3 [Source:MGI Symbol;Acc:MGI:103249] | 2142,0 | 82,4 |
| ENSMUSG00000035674 | Ndufa3 | NADH dehydrogenase (ubiquinone) 1 alpha subcomplex, 3 [Source:MGI Symbol;Acc:MGI:1913341] | 2128,2 | 74,3 |
| ENSMUSG00000063297 | Luzp2 | leucine zipper protein 2 [Source:MGI Symbol;Acc:MGI:1889615] | 2126,7 | 96,4 |
| ENSMUSG00000006728 | Cdk4 | cyclin-dependent kinase 4 [Source:MGI Symbol;Acc:MGI:88357] | 2121,0 | 87,7 |
| ENSMUSG00000003072 | Atp5d | ATP synthase, H+ transporting, mitochondrial F1 complex, delta subunit [Source:MGI Symbol;Acc:MGI:1913293] | 2119,6 | 72,3 |
| ENSMUSG00000079037 | Prnp | prion protein [Source:MGI Symbol;Acc:MGI:97769] | 2116,7 | 98,5 |
| ENSMUSG00000024359 | Hspa9 | heat shock protein 9 [Source:MGI Symbol;Acc:MGI:96245] | 2113,9 | 96,5 |
| ENSMUSG00000048022 | Tmem229a | transmembrane protein 229A [Source:MGI Symbol;Acc:MGI:2442812] | 2111,7 | 91,1 |
| ENSMUSG00000036046 | 5031439G07Rik | RIKEN cDNA 5031439G07 gene [Source:MGI Symbol;Acc:MGI:2444899] | 2106,2 | 83,4 |
| ENSMUSG00000056201 | Cfl1 | cofilin 1, non-muscle [Source:MGI Symbol;Acc:MGI:101757] | 2100,2 | 99,5 |
| ENSMUSG00000030120 | Mlf2 | myeloid leukemia factor 2 [Source:MGI Symbol;Acc:MGI:1353554] | 2097,3 | 88,4 |
| ENSMUSG00000032330 | Cox7a2 | cytochrome c oxidase subunit VIIa 2 [Source:MGI Symbol;Acc:MGI:1316715] | 2096,7 | 71,9 |
| ENSMUSG00000048578 | Mlec | malectin [Source:MGI Symbol;Acc:MGI:1924015] | 2086,8 | 74,5 |
| ENSMUSG00000020475 | Pgam2 | phosphoglycerate mutase 2 [Source:MGI Symbol;Acc:MGI:1933118] | 2082,6 | 98,5 |
| ENSMUSG00000035805 | Mlc1 | megalencephalic leukoencephalopathy with subcortical cysts 1 homolog (human) [Source:MGI Symbol;Acc:MGI:2157910] | 2075,6 | 89,7 |
| ENSMUSG00000005779 | Psmb4 | proteasome (prosome, macropain) subunit, beta type 4 [Source:MGI Symbol;Acc:MGI:1098257] | 2073,9 | 78,8 |
| ENSMUSG00000029632 | Ndufa4 | NADH dehydrogenase (ubiquinone) 1 alpha subcomplex, 4 [Source:MGI Symbol;Acc:MGI:107686] | 2065,1 | 86,8 |
| ENSMUSG00000026615 | Eprs | glutamyl-prolyl-tRNA synthetase [Source:MGI Symbol;Acc:MGI:97838] | 2064,5 | 87,3 |
| ENSMUSG00000027852 | Nras | neuroblastoma ras oncogene [Source:MGI Symbol;Acc:MGI:97376] | 2061,9 | 74,4 |
| ENSMUSG00000057388 | Mrpl18 | mitochondrial ribosomal protein L18 [Source:MGI Symbol;Acc:MGI:1914931] | 2055,3 | 81,5 |
| ENSMUSG00000051695 | Pcbp1 | poly(rC) binding protein 1 [Source:MGI Symbol;Acc:MGI:1345635] | 2050,3 | 84,3 |
| ENSMUSG00000018593 | Sparc | secreted acidic cysteine rich glycoprotein [Source:MGI Symbol;Acc:MGI:98373] | 2036,7 | 91,8 |
| ENSMUSG00000032181 | Scg3 | secretogranin III [Source:MGI Symbol;Acc:MGI:103032] | 2017,5 | 98,0 |
| ENSMUSG00000021124 | Vti1b | vesicle transport through interaction with t-SNAREs 1B [Source:MGI Symbol;Acc:MGI:1855688] | 1992,5 | 75,2 |
| ENSMUSG00000031667 | Aktip | thymoma viral proto-oncogene 1 interacting protein [Source:MGI Symbol;Acc:MGI:3693832] | 1991,3 | 74,2 |
| ENSMUSG00000044080 | S100a1 | S100 calcium binding protein A1 [Source:MGI Symbol;Acc:MGI:1338917] | 1986,9 | 89,2 |
| ENSMUSG00000072704 | Smim10l1 | small integral membrane protein 10 like 1 [Source:MGI Symbol;Acc:MGI:1914379] | 1982,3 | 82,7 |
| ENSMUSG00000025381 | Cnpy2 | canopy FGF signaling regulator 2 [Source:MGI Symbol;Acc:MGI:1928477] | 1981,3 | 96,1 |
| ENSMUSG00000039016 | Timm8b | translocase of inner mitochondrial membrane 8B [Source:MGI Symbol;Acc:MGI:1353424] | 1981,0 | 100,0 |
| ENSMUSG00000015749 | Anp32e | acidic (leucine-rich) nuclear phosphoprotein 32 family, member E [Source:MGI Symbol;Acc:MGI:1913721] | 1969,9 | 93,2 |
| ENSMUSG00000020719 | Ddx5 | DEAD (Asp-Glu-Ala-Asp) box polypeptide 5 [Source:MGI Symbol;Acc:MGI:105037] | 1969,8 | 85,8 |
| ENSMUSG00000030246 | Ldhb | lactate dehydrogenase B [Source:MGI Symbol;Acc:MGI:96763] | 1961,4 | 80,7 |
| ENSMUSG00000006304 | Arpc2 | actin related protein 2/3 complex, subunit 2 [Source:MGI Symbol;Acc:MGI:1923959] | 1959,7 | 72,7 |
| ENSMUSG00000026755 | Arpc5l | actin related protein 2/3 complex, subunit 5-like [Source:MGI Symbol;Acc:MGI:1921442] | 1941,7 | 97,9 |
| ENSMUSG00000023913 | Pla2g7 | phospholipase A2, group VII (platelet-activating factor acetylhydrolase, plasma) [Source:MGI Symbol;Acc:MGI:1351327] | 1935,8 | 85,4 |
| ENSMUSG00000005667 | Mthfd2 | etrahydrofolate dehydrogenase (NAD+ dependent), methenyltetrahydrofolate cyclohydrolase [Source:MGI Symbol;Acc:MG | 1920,4 | 89,0 |
| ENSMUSG00000014504 | Srp19 | signal recognition particle 19 [Source:MGI Symbol;Acc:MGI:1913634] | 1911,4 | 99,0 |
| ENSMUSG00000029580 | Actb | actin, beta [Source:MGI Symbol;Acc:MGI:87904] | 1900,6 | 85,8 |
| ENSMUSG00000021290 | 2010107E04Rik | RIKEN cDNA 2010107E04 gene [Source:MGI Symbol;Acc:MGI:1917507] | 1899,9 | 100,0 |
| ENSMUSG00000019804 | Snx3 | sorting nexin 3 [Source:MGI Symbol;Acc:MGI:1860188] | 1889,2 | 88,1 |
| ENSMUSG00000031776 | Arl2bp | ADP-ribosylation factor-like 2 binding protein [Source:MGI Symbol;Acc:MGI:1349429] | 1886,1 | 100,0 |
| ENSMUSG00000026511 | Srp9 | signal recognition particle 9 [Source:MGI Symbol;Acc:MGI:1350930] | 1881,3 | 83,4 |
| ENSMUSG00000063316 | Rpl27 | ribosomal protein L27 [Source:MGI Symbol;Acc:MGI:98036] | 1874,8 | 72,9 |
| ENSMUSG00000021474 | Sfxn1 | sideroflexin 1 [Source:MGI Symbol;Acc:MGI:2137677] | 1873,3 | 93,2 |
| ENSMUSG00000026833 | Olfm1 | olfactomedin 1 [Source:MGI Symbol;Acc:MGI:1860437] | 1867,1 | 90,2 |
| ENSMUSG00000019877 | Serinc1 | serine incorporator 1 [Source:MGI Symbol;Acc:MGI:1926228] | 1863,1 | 93,5 |
| ENSMUSG00000010277 | 2610507B11Rik | RIKEN cDNA 2610507B11 gene [Source:MGI Symbol;Acc:MGI:1919753] | 1860,3 | 71,3 |
| ENSMUSG00000024533 | Spire1 | spire homolog 1 (Drosophila) [Source:MGI Symbol;Acc:MGI:1915416] | 1860,3 | 84,1 |
| ENSMUSG00000029128 | Rab28 | RAB28, member RAS oncogene family [Source:MGI Symbol;Acc:MGI:1917285] | 1860,0 | 97,5 |
| ENSMUSG00000041126 | H2afv | H2A histone family, member V [Source:MGI Symbol;Acc:MGI:1924855] | 1853,2 | 97,8 |
| ENSMUSG00000028517 | Plpp3 | phospholipid phosphatase 3 [Source:MGI Symbol;Acc:MGI:1915166] | 1852,3 | 96,4 |
| ENSMUSG00000050856 | Atp5k | ATP synthase, H+ transporting, mitochondrial F1F0 complex, subunit E [Source:MGI Symbol;Acc:MGI:106636] | 1845,3 | 91,8 |
| ENSMUSG00000000088 | Cox5a | cytochrome c oxidase subunit Va [Source:MGI Symbol;Acc:MGI:88474] | 1844,6 | 96,9 |
| ENSMUSG00000031447 | Lamp1 | lysosomal-associated membrane protein 1 [Source:MGI Symbol;Acc:MGI:96745] | 1841,1 | 92,3 |
| ENSMUSG00000019179 | Mdh2 | malate dehydrogenase 2, NAD (mitochondrial) [Source:MGI Symbol;Acc:MGI:97050] | 1827,1 | 87,6 |
| ENSMUSG00000092137 | Gcom1 | GRINL1A complex locus [Source:MGI Symbol;Acc:MGI:5141967] | 1800,9 | 71,7 |
| ENSMUSG00000016319 | Slc25a5 | te carrier family 25 (mitochondrial carrier, adenine nucleotide translocator), member 5 [Source:MGI Symbol;Acc:MGI:1353 | 1799,0 | 97,2 |
| ENSMUSG00000030870 | Ubfd1 | ubiquitin family domain containing 1 [Source:MGI Symbol;Acc:MGI:107301] | 1796,0 | 92,0 |
| ENSMUSG00000032562 | Gnai2 | guanine nucleotide binding protein (G protein), alpha inhibiting 2 [Source:MGI Symbol;Acc:MGI:95772] | 1791,4 | 87,6 |
| ENSMUSG00000020091 | Eif4ebp2 | eukaryotic translation initiation factor 4E binding protein 2 [Source:MGI Symbol;Acc:MGI:109198] | 1779,6 | 98,0 |
| ENSMUSG00000023027 | Atf1 | activating transcription factor 1 [Source:MGI Symbol;Acc:MGI:1298366] | 1778,9 | 79,1 |
| ENSMUSG00000036197 | Gxylt1 | glucoside xylosyltransferase 1 [Source:MGI Symbol;Acc:MGI:2684933] | 1775,3 | 70,5 |
| ENSMUSG00000041329 | Atp1b2 | ATPase, Na+/K+ transporting, beta 2 polypeptide [Source:MGI Symbol;Acc:MGI:88109] | 1770,4 | 95,3 |
| ENSMUSG00000053510 | Nrd1 | nardilysin, N-arginine dibasic convertase, NRD convertase 1 [Source:MGI Symbol;Acc:MGI:1201386] | 1766,8 | 74,5 |
| ENSMUSG00000027162 | Lin7c | lin-7 homolog C (C. elegans) [Source:MGI Symbol;Acc:MGI:1330839] | 1760,0 | 77,3 |
| ENSMUSG00000079197 | Psme2 | proteasome (prosome, macropain) activator subunit 2 (PA28 beta) [Source:MGI Symbol;Acc:MGI:1096365] | 1758,9 | 85,2 |
| ENSMUSG00000006740 | Kif5b | kinesin family member 5B [Source:MGI Symbol;Acc:MGI:1098268] | 1757,4 | 72,7 |
| ENSMUSG00000026473 | Glul | glutamate-ammonia ligase (glutamine synthetase) [Source:MGI Symbol;Acc:MGI:95739] | 1752,7 | 70,1 |
| ENSMUSG00000030357 | Fkbp4 | FK506 binding protein 4 [Source:MGI Symbol;Acc:MGI:95543] | 1752,4 | 93,9 |
| ENSMUSG00000018293 | Pfn1 | profilin 1 [Source:MGI Symbol;Acc:MGI:97549] | 1749,9 | 83,3 |
| ENSMUSG00000026914 | Psmd14 | proteasome (prosome, macropain) 26S subunit, non-ATPase, 14 [Source:MGI Symbol;Acc:MGI:1913284] | 1745,3 | 89,8 |
| ENSMUSG00000004897 | Hdgf | hepatoma-derived growth factor [Source:MGI Symbol;Acc:MGI:1194494] | 1737,9 | 91,0 |
| ENSMUSG00000062647 | Rpl7a | ribosomal protein L7A [Source:MGI Symbol;Acc:MGI:1353472] | 1737,7 | 88,1 |
| ENSMUSG00000002297 | Dbf4 | DBF4 zinc finger [Source:MGI Symbol;Acc:MGI:1351328] | 1734,5 | 83,3 |
| ENSMUSG00000004980 | Hnrnpa2b1 | heterogeneous nuclear ribonucleoprotein A2/B1 [Source:MGI Symbol;Acc:MGI:104819] | 1733,2 | 83,8 |
| ENSMUSG00000004892 | Bcan | brevican [Source:MGI Symbol;Acc:MGI:1096385] | 1731,1 | 98,2 |
| ENSMUSG00000021832 | Psmc6 | proteasome (prosome, macropain) 26S subunit, ATPase, 6 [Source:MGI Symbol;Acc:MGI:1914339] | 1729,4 | 91,1 |
| ENSMUSG00000020484 | Xbp1 | X-box binding protein 1 [Source:MGI Symbol;Acc:MGI:98970] | 1725,9 | 81,4 |
| ENSMUSG00000006699 | Cdc42 | cell division cycle 42 [Source:MGI Symbol;Acc:MGI:106211] | 1725,7 | 71,9 |
| ENSMUSG00000035686 | Thrsp | thyroid hormone responsive [Source:MGI Symbol;Acc:MGI:109126] | 1724,6 | 83,6 |
| ENSMUSG00000019132 | BC005537 | cDNA sequence BC005537 [Source:MGI Symbol;Acc:MGI:2441726] | 1722,1 | 72,0 |
| ENSMUSG00000028798 | Eif3i | eukaryotic translation initiation factor 3, subunit I [Source:MGI Symbol;Acc:MGI:1860763] | 1715,6 | 92,6 |
| ENSMUSG00000024695 | Zfp91 | zinc finger protein 91 [Source:MGI Symbol;Acc:MGI:104854] | 1714,7 | 76,2 |
| ENSMUSG00000032328 | Tmem30a | transmembrane protein 30A [Source:MGI Symbol;Acc:MGI:106402] | 1713,5 | 74,4 |
| ENSMUSG00000068328 | Aup1 | ancient ubiquitous protein 1 [Source:MGI Symbol;Acc:MGI:107789] | 1709,7 | 81,9 |
| ENSMUSG00000001018 | Snapin | SNAP-associated protein [Source:MGI Symbol;Acc:MGI:1333745] | 1700,8 | 92,8 |
| ENSMUSG00000010376 | Nedd8 | neural precursor cell expressed, developmentally down-regulated gene 8 [Source:MGI Symbol;Acc:MGI:97301] | 1698,9 | 100,0 |
| ENSMUSG00000019054 | Fis1 | fission, mitochondrial 1 [Source:MGI Symbol;Acc:MGI:1913687] | 1696,9 | 99,0 |
| ENSMUSG00000040242 | Fgfr1op2 | FGFR1 oncogene partner 2 [Source:MGI Symbol;Acc:MGI:1914779] | 1696,5 | 84,4 |
| ENSMUSG00000036438 | Calm2 | calmodulin 2 [Source:MGI Symbol;Acc:MGI:103250] | 1693,7 | 94,9 |
| ENSMUSG00000009112 | Bcl2l13 | BCL2-like 13 (apoptosis facilitator) [Source:MGI Symbol;Acc:MGI:2136959] | 1673,8 | 74,6 |
| ENSMUSG00000048076 | Arf1 | ADP-ribosylation factor 1 [Source:MGI Symbol;Acc:MGI:99431] | 1667,9 | 86,2 |
| ENSMUSG00000032412 | Atp1b3 | ATPase, Na+/K+ transporting, beta 3 polypeptide [Source:MGI Symbol;Acc:MGI:107788] | 1666,3 | 89,9 |
| ENSMUSG00000024712 | Rfk | riboflavin kinase [Source:MGI Symbol;Acc:MGI:1914688] | 1665,6 | 91,9 |
| ENSMUSG00000030647 | Ndufc2 | NADH dehydrogenase (ubiquinone) 1, subcomplex unknown, 2 [Source:MGI Symbol;Acc:MGI:1344370] | 1664,9 | 86,1 |
| ENSMUSG00000054008 | Ndst1 | N-deacetylase/N-sulfotransferase (heparan glucosaminyl) 1 [Source:MGI Symbol;Acc:MGI:104719] | 1663,5 | 75,0 |
| ENSMUSG00000021484 | Lman2 | lectin, mannose-binding 2 [Source:MGI Symbol;Acc:MGI:1914140] | 1662,7 | 96,3 |
| ENSMUSG00000040731 | Eif4h | eukaryotic translation initiation factor 4H [Source:MGI Symbol;Acc:MGI:1341822] | 1660,7 | 88,3 |
| ENSMUSG00000022698 | Naa50 | N(alpha)-acetyltransferase 50, NatE catalytic subunit [Source:MGI Symbol;Acc:MGI:1919367] | 1657,0 | 78,3 |
| ENSMUSG00000031176 | Dynlt3 | dynein light chain Tctex-type 3 [Source:MGI Symbol;Acc:MGI:1914367] | 1653,2 | 98,3 |
| ENSMUSG00000067367 | Lyar | Ly1 antibody reactive clone [Source:MGI Symbol;Acc:MGI:107470] | 1649,6 | 86,7 |
| ENSMUSG00000032667 | Pon2 | paraoxonase 2 [Source:MGI Symbol;Acc:MGI:106687] | 1643,6 | 75,1 |
| ENSMUSG00000024587 | Nars | asparaginyl-tRNA synthetase [Source:MGI Symbol;Acc:MGI:1917473] | 1643,2 | 99,3 |

| | | | | |
|---|---|---|---|---|
| ENSMUSG00000025917 | Cops5 | COP9 signalosome subunit 5 [Source:MGI Symbol;Acc:MGI:1349415] | 1640,0 | 83,1 |
| ENSMUSG00000061132 | Blnk | B cell linker [Source:MGI Symbol;Acc:MGI:96878] | 1635,8 | 85,4 |
| ENSMUSG00000026095 | Asnsd1 | asparagine synthetase domain containing 1 [Source:MGI Symbol;Acc:MGI:1917646] | 1634,3 | 83,8 |
| ENSMUSG00000021242 | Npc2 | Niemann-Pick type C2 [Source:MGI Symbol;Acc:MGI:1915213] | 1633,7 | 71,4 |
| ENSMUSG00000055538 | Zcchc24 | zinc finger, CCHC domain containing 24 [Source:MGI Symbol;Acc:MGI:1919168] | 1633,5 | 73,4 |
| ENSMUSG00000023175 | Bsg | basigin [Source:MGI Symbol;Acc:MGI:88208] | 1632,8 | 76,4 |
| ENSMUSG00000004096 | Cwc15 | CWC15 spliceosome-associated protein [Source:MGI Symbol;Acc:MGI:1913320] | 1623,2 | 95,4 |
| ENSMUSG00000000149 | Gna12 | guanine nucleotide binding protein, alpha 12 [Source:MGI Symbol;Acc:MGI:95767] | 1620,5 | 92,7 |
| ENSMUSG00000030654 | Arl6ip1 | ADP-ribosylation factor-like 6 interacting protein 1 [Source:MGI Symbol;Acc:MGI:1858943] | 1619,8 | 94,0 |
| ENSMUSG00000009073 | Nf2 | neurofibromatosis 2 [Source:MGI Symbol;Acc:MGI:97307] | 1619,2 | 79,6 |
| ENSMUSG00000015363 | Trabd | TraB domain containing [Source:MGI Symbol;Acc:MGI:1915226] | 1617,6 | 87,3 |
| ENSMUSG00000032060 | Cryab | crystallin, alpha B [Source:MGI Symbol;Acc:MGI:88516] | 1615,7 | 84,1 |
| ENSMUSG00000029247 | Paics | oimidazole carboxylase, phosphoribosylaminoribosylaminoimidazole, succinocarboxamide synthetase [Source:MGI Symbo | 1615,1 | 81,7 |
| ENSMUSG00000071528 | Usmg5 | upregulated during skeletal muscle growth 5 [Source:MGI Symbol;Acc:MGI:1891435] | 1614,6 | 94,6 |
| ENSMUSG00000032381 | Fam96a | family with sequence similarity 96, member A [Source:MGI Symbol;Acc:MGI:1915500] | 1606,4 | 91,6 |
| ENSMUSG00000081534 | Slc48a1 | solute carrier family 48 (heme transporter), member 1 [Source:MGI Symbol;Acc:MGI:1914989] | 1606,3 | 96,3 |
| ENSMUSG00000091264 | Smim13 | small integral membrane protein 13 [Source:MGI Symbol;Acc:MGI:2652854] | 1602,5 | 78,0 |
| ENSMUSG00000015733 | Capza2 | capping protein (actin filament) muscle Z-line, alpha 2 [Source:MGI Symbol;Acc:MGI:106222] | 1599,4 | 87,8 |
| ENSMUSG00000031388 | Naa10 | N(alpha)-acetyltransferase 10, NatA catalytic subunit [Source:MGI Symbol;Acc:MGI:1915255] | 1595,0 | 77,6 |
| ENSMUSG00000021218 | Gdi2 | guanosine diphosphate (GDP) dissociation inhibitor 2 [Source:MGI Symbol;Acc:MGI:99845] | 1592,7 | 85,0 |
| ENSMUSG00000024248 | Cox7a2l | cytochrome c oxidase subunit VIIa polypeptide 2-like [Source:MGI Symbol;Acc:MGI:106015] | 1586,2 | 71,0 |
| ENSMUSG00000031392 | Irak1 | interleukin-1 receptor-associated kinase 1 [Source:MGI Symbol;Acc:MGI:107420] | 1581,3 | 82,8 |
| ENSMUSG00000027562 | Car2 | carbonic anhydrase 2 [Source:MGI Symbol;Acc:MGI:88269] | 1579,5 | 96,6 |
| ENSMUSG00000020048 | Hsp90b1 | heat shock protein 90, beta (Grp94), member 1 [Source:MGI Symbol;Acc:MGI:98817] | 1574,8 | 95,4 |
| ENSMUSG00000020482 | Ccdc117 | coiled-coil domain containing 117 [Source:MGI Symbol;Acc:MGI:2144383] | 1569,0 | 94,0 |
| ENSMUSG00000033475 | Tomm6 | translocase of outer mitochondrial membrane 6 homolog (yeast) [Source:MGI Symbol;Acc:MGI:1913369] | 1556,0 | 86,7 |
| ENSMUSG00000025980 | Hspd1 | heat shock protein 1 (chaperonin) [Source:MGI Symbol;Acc:MGI:96242] | 1552,5 | 77,8 |
| ENSMUSG00000023147 | Wrb | tryptophan rich basic protein [Source:MGI Symbol;Acc:MGI:2136882] | 1552,5 | 93,7 |
| ENSMUSG00000074886 | Grk6 | G protein-coupled receptor kinase 6 [Source:MGI Symbol;Acc:MGI:1347078] | 1545,8 | 92,1 |
| ENSMUSG00000020496 | Rnf187 | ring finger protein 187 [Source:MGI Symbol;Acc:MGI:1914224] | 1543,8 | 98,2 |
| ENSMUSG00000026150 | Mff | mitochondrial fission factor [Source:MGI Symbol;Acc:MGI:1922984] | 1539,2 | 82,8 |
| ENSMUSG00000006442 | Srm | spermidine synthase [Source:MGI Symbol;Acc:MGI:102690] | 1537,7 | 93,9 |
| ENSMUSG00000049327 | Setd8 | SET domain containing (lysine methyltransferase) 8 [Source:MGI Symbol;Acc:MGI:1915206] | 1533,5 | 88,9 |
| ENSMUSG00000022323 | Hrsp12 | heat-responsive protein 12 [Source:MGI Symbol;Acc:MGI:1095401] | 1530,0 | 98,9 |
| ENSMUSG00000014226 | Cacybp | calcyclin binding protein [Source:MGI Symbol;Acc:MGI:1270839] | 1529,6 | 70,4 |
| ENSMUSG00000032026 | Rexo2 | REX2, RNA exonuclease 2 homolog (S. cerevisiae) [Source:MGI Symbol;Acc:MGI:1888981] | 1525,2 | 100,0 |
| ENSMUSG00000029014 | Dnajc2 | DnaJ heat shock protein family (Hsp40) member C2 [Source:MGI Symbol;Acc:MGI:99470] | 1521,9 | 73,9 |
| ENSMUSG00000070319 | Eif3g | eukaryotic translation initiation factor 3, subunit G [Source:MGI Symbol;Acc:MGI:1858258] | 1520,1 | 100,0 |
| ENSMUSG00000003131 | Pafah1b2 | platelet-activating factor acetylhydrolase, isoform 1b, subunit 2 [Source:MGI Symbol;Acc:MGI:108415] | 1515,8 | 80,8 |
| ENSMUSG00000071655 | Ubxn1 | UBX domain protein 1 [Source:MGI Symbol;Acc:MGI:1289301] | 1508,5 | 98,7 |
| ENSMUSG00000022216 | Psme1 | proteasome (prosome, macropain) activator subunit 1 (PA28 alpha) [Source:MGI Symbol;Acc:MGI:1096367] | 1502,1 | 79,2 |
| ENSMUSG00000018669 | Cdk5rap3 | CDK5 regulatory subunit associated protein 3 [Source:MGI Symbol;Acc:MGI:1933126] | 1501,2 | 77,7 |
| ENSMUSG00000033059 | Pygb | brain glycogen phosphorylase [Source:MGI Symbol;Acc:MGI:97828] | 1500,3 | 97,3 |
| ENSMUSG00000044627 | Swi5 | SWI5 recombination repair homolog (yeast) [Source:MGI Symbol;Acc:MGI:1920181] | 1500,0 | 81,3 |
| ENSMUSG00000058446 | Znrf2 | zinc and ring finger 2 [Source:MGI Symbol;Acc:MGI:1196246] | 1499,0 | 77,8 |
| ENSMUSG00000046240 | Hepacam | hepatocyte cell adhesion molecule [Source:MGI Symbol;Acc:MGI:1920177] | 1497,0 | 83,8 |
| ENSMUSG00000020585 | Laptm4a | lysosomal-associated protein transmembrane 4A [Source:MGI Symbol;Acc:MGI:108017] | 1492,5 | 72,5 |
| ENSMUSG00000004268 | Emg1 | EMG1 N1-specific pseudouridine methyltransferase [Source:MGI Symbol;Acc:MGI:1315195] | 1490,5 | 98,4 |
| ENSMUSG00000062981 | Mrpl42 | mitochondrial ribosomal protein L42 [Source:MGI Symbol;Acc:MGI:1333774] | 1482,5 | 96,5 |
| ENSMUSG00000035597 | Prpf39 | PRP39 pre-mRNA processing factor 39 homolog (yeast) [Source:MGI Symbol;Acc:MGI:104602] | 1482,0 | 80,9 |
| ENSMUSG00000084957 | Bbip1 | BBSome interacting protein 1 [Source:MGI Symbol;Acc:MGI:1913610] | 1476,5 | 79,2 |
| ENSMUSG00000052298 | Cdc42se2 | CDC42 small effector 2 [Source:MGI Symbol;Acc:MGI:1919979] | 1473,8 | 95,9 |
| ENSMUSG00000020271 | Fbxw11 | F-box and WD-40 domain protein 11 [Source:MGI Symbol;Acc:MGI:2144023] | 1472,6 | 84,2 |
| ENSMUSG00000028419 | Chmp5 | charged multivesicular body protein 5 [Source:MGI Symbol;Acc:MGI:1924209] | 1470,5 | 90,2 |
| ENSMUSG00000022312 | Eif3h | eukaryotic translation initiation factor 3, subunit H [Source:MGI Symbol;Acc:MGI:1915385] | 1463,9 | 91,6 |
| ENSMUSG00000020664 | Dld | dihydrolipoamide dehydrogenase [Source:MGI Symbol;Acc:MGI:107450] | 1460,6 | 94,5 |
| ENSMUSG00000008690 | Ncaph2 | non-SMC condensin II complex, subunit H2 [Source:MGI Symbol;Acc:MGI:1289164] | 1457,3 | 74,6 |
| ENSMUSG00000032393 | Dpp8 | dipeptidylpeptidase 8 [Source:MGI Symbol;Acc:MGI:1921638] | 1452,5 | 88,4 |
| ENSMUSG00000029030 | Tprgl | transformation related protein 63 regulated like [Source:MGI Symbol;Acc:MGI:1915058] | 1449,4 | 99,8 |
| ENSMUSG00000031059 | Ndufb11 | NADH dehydrogenase (ubiquinone) 1 beta subcomplex, 11 [Source:MGI Symbol;Acc:MGI:1349919] | 1448,8 | 83,1 |
| ENSMUSG00000020029 | Nudt4 | nudix (nucleoside diphosphate linked moiety X)-type motif 4 [Source:MGI Symbol;Acc:MGI:1918457] | 1447,4 | 98,9 |
| ENSMUSG00000033940 | Brk1 | BRICK1, SCAR/WAVE actin-nucleating complex subunit [Source:MGI Symbol;Acc:MGI:1915406] | 1446,5 | 99,6 |
| ENSMUSG00000048429 | 1810026J23Rik | RIKEN cDNA 1810026J23 gene [Source:MGI Symbol;Acc:MGI:1917023] | 1446,1 | 90,5 |
| ENSMUSG00000028445 | Enho | energy homeostasis associated [Source:MGI Symbol;Acc:MGI:1916888] | 1441,7 | 74,2 |
| ENSMUSG00000015341 | Golga7 | golgi autoantigen, golgin subfamily a, 7 [Source:MGI Symbol;Acc:MGI:1931029] | 1431,2 | 77,9 |
| ENSMUSG00000022354 | Ndufb9 | NADH dehydrogenase (ubiquinone) 1 beta subcomplex, 9 [Source:MGI Symbol;Acc:MGI:1913468] | 1430,8 | 100,0 |
| ENSMUSG00000003380 | Rabac1 | Rab acceptor 1 (prenylated) [Source:MGI Symbol;Acc:MGI:1201692] | 1426,2 | 79,2 |
| ENSMUSG00000000171 | Sdhd | succinate dehydrogenase complex, subunit D, integral membrane protein [Source:MGI Symbol;Acc:MGI:1914175] | 1420,6 | 97,0 |
| ENSMUSG00000031245 | Hmgn5 | high-mobility group nucleosome binding domain 5 [Source:MGI Symbol;Acc:MGI:1355295] | 1412,5 | 94,4 |
| ENSMUSG00000027566 | Psma7 | proteasome (prosome, macropain) subunit, alpha type 7 [Source:MGI Symbol;Acc:MGI:1347070] | 1406,2 | 99,3 |
| ENSMUSG00000006941 | Eif1b | eukaryotic translation initiation factor 1B [Source:MGI Symbol;Acc:MGI:1916219] | 1403,8 | 99,3 |
| ENSMUSG00000028161 | Ppp3ca | protein phosphatase 3, catalytic subunit, alpha isoform [Source:MGI Symbol;Acc:MGI:107164] | 1402,5 | 78,0 |
| ENSMUSG00000014748 | Tex261 | testis expressed gene 261 [Source:MGI Symbol;Acc:MGI:1096575] | 1399,0 | 81,4 |
| ENSMUSG00000055839 | Tceb2 | transcription elongation factor B (SIII), polypeptide 2 [Source:MGI Symbol;Acc:MGI:1914923] | 1387,1 | 99,5 |
| ENSMUSG00000024962 | Vegfb | vascular endothelial growth factor B [Source:MGI Symbol;Acc:MGI:106199] | 1385,3 | 91,0 |
| ENSMUSG00000102918 | Pcdhgc3 | protocadherin gamma subfamily C, 3 [Source:MGI Symbol;Acc:MGI:1935201] | 1383,8 | 88,7 |
| ENSMUSG00000030884 | Uqcrc2 | ubiquinol cytochrome c reductase core protein 2 [Source:MGI Symbol;Acc:MGI:1914253] | 1383,3 | 87,8 |
| ENSMUSG00000006476 | Nsmf | NMDA receptor synaptonuclear signaling and neuronal migration factor [Source:MGI Symbol;Acc:MGI:1861755] | 1382,3 | 97,0 |
| ENSMUSG00000059796 | Eif4a1 | eukaryotic translation initiation factor 4A1 [Source:MGI Symbol;Acc:MGI:95303] | 1378,1 | 85,3 |
| ENSMUSG00000024844 | Banf1 | barrier to autointegration factor 1 [Source:MGI Symbol;Acc:MGI:1346330] | 1375,2 | 95,2 |
| ENSMUSG00000031996 | Aplp2 | amyloid beta (A4) precursor-like protein 2 [Source:MGI Symbol;Acc:MGI:88047] | 1365,0 | 82,3 |
| ENSMUSG00000026558 | Uck2 | uridine-cytidine kinase 2 [Source:MGI Symbol;Acc:MGI:1931744] | 1362,3 | 76,6 |
| ENSMUSG00000002222 | Rmnd5a | required for meiotic nuclear division 5 homolog A (S. cerevisiae) [Source:MGI Symbol;Acc:MGI:1915727] | 1361,0 | 71,5 |
| ENSMUSG00000034432 | Cops8 | COP9 signalosome subunit 8 [Source:MGI Symbol;Acc:MGI:1915363] | 1358,5 | 72,4 |
| ENSMUSG00000026576 | Atp1b1 | ATPase, Na+/K+ transporting, beta 1 polypeptide [Source:MGI Symbol;Acc:MGI:88108] | 1355,9 | 79,6 |
| ENSMUSG00000040385 | Ppp1ca | protein phosphatase 1, catalytic subunit, alpha isoform [Source:MGI Symbol;Acc:MGI:103016] | 1353,2 | 99,3 |
| ENSMUSG00000024411 | Aqp4 | aquaporin 4 [Source:MGI Symbol;Acc:MGI:107387] | 1352,1 | 79,0 |
| ENSMUSG00000004207 | Psap | prosaposin [Source:MGI Symbol;Acc:MGI:97783] | 1351,8 | 70,5 |
| ENSMUSG00000022450 | Ndufa6 | NADH dehydrogenase (ubiquinone) 1 alpha subcomplex, 6 (B14) [Source:MGI Symbol;Acc:MGI:1914380] | 1351,7 | 93,8 |
| ENSMUSG00000018379 | Srsf1 | serine/arginine-rich splicing factor 1 [Source:MGI Symbol;Acc:MGI:98283] | 1351,6 | 73,4 |
| ENSMUSG00000020358 | Hnrnpab | heterogeneous nuclear ribonucleoprotein A/B [Source:MGI Symbol;Acc:MGI:1330294] | 1348,1 | 78,3 |
| ENSMUSG00000020644 | Id2 | inhibitor of DNA binding 2 [Source:MGI Symbol;Acc:MGI:96397] | 1346,6 | 98,5 |
| ENSMUSG00000062867 | Impdh2 | inosine 5'-phosphate dehydrogenase 2 [Source:MGI Symbol;Acc:MGI:109367] | 1343,3 | 73,3 |
| ENSMUSG00000070394 | Tmem256 | transmembrane protein 256 [Source:MGI Symbol;Acc:MGI:1916436] | 1342,1 | 97,9 |
| ENSMUSG00000032547 | Ryk | receptor-like tyrosine kinase [Source:MGI Symbol;Acc:MGI:101766] | 1339,1 | 83,4 |
| ENSMUSG00000024378 | Stard4 | StAR-related lipid transfer (START) domain containing 4 [Source:MGI Symbol;Acc:MGI:2156764] | 1335,5 | 79,7 |
| ENSMUSG00000015759 | Cnih1 | cornichon family AMPA receptor auxiliary protein 1 [Source:MGI Symbol;Acc:MGI:1277202] | 1335,2 | 93,2 |
| ENSMUSG00000028044 | Cks1b | CDC28 protein kinase 1b [Source:MGI Symbol;Acc:MGI:1889208] | 1329,4 | 72,4 |
| ENSMUSG00000033629 | Hacd3 | 3-hydroxyacyl-CoA dehydratase 3 [Source:MGI Symbol;Acc:MGI:1889341] | 1322,7 | 98,9 |
| ENSMUSG00000008475 | Arpc5 | actin related protein 2/3 complex, subunit 5 [Source:MGI Symbol;Acc:MGI:1915021] | 1318,2 | 92,7 |
| ENSMUSG00000031065 | Cdk16 | cyclin-dependent kinase 16 [Source:MGI Symbol;Acc:MGI:97516] | 1315,0 | 75,5 |
| ENSMUSG00000076576 | Igkv6-32 | immunoglobulin kappa variable 6-32 [Source:MGI Symbol;Acc:MGI:3641634] | 1314,9 | 99,2 |
| ENSMUSG00000020737 | Hn1 | hematological and neurological expressed sequence 1 [Source:MGI Symbol;Acc:MGI:1096361] | 1314,0 | 93,6 |
| ENSMUSG00000022452 | Smdt1 | single-pass membrane protein with aspartate rich tail 1 [Source:MGI Symbol;Acc:MGI:1916279] | 1314,0 | 77,0 |
| ENSMUSG00000035864 | Syt1 | synaptotagmin I [Source:MGI Symbol;Acc:MGI:99667] | 1311,7 | 90,3 |
| ENSMUSG00000020390 | Ube2b | ubiquitin-conjugating enzyme E2B [Source:MGI Symbol;Acc:MGI:102944] | 1309,1 | 89,4 |
| ENSMUSG00000046722 | Cdc42se1 | CDC42 small effector 1 [Source:MGI Symbol;Acc:MGI:1889510] | 1308,0 | 90,9 |
| ENSMUSG00000022437 | Samm50 | sorting and assembly machinery component 50 homolog (S. cerevisiae) [Source:MGI Symbol;Acc:MGI:1915903] | 1307,5 | 98,3 |
| ENSMUSG00000025651 | Uqcrc1 | ubiquinol-cytochrome c reductase core protein 1 [Source:MGI Symbol;Acc:MGI:107876] | 1300,2 | 89,6 |
| ENSMUSG00000006205 | Htra1 | HtrA serine peptidase 1 [Source:MGI Symbol;Acc:MGI:1929076] | 1300,0 | 95,4 |
| ENSMUSG00000030265 | Kras | Kirsten rat sarcoma viral oncogene homolog [Source:MGI Symbol;Acc:MGI:96680] | 1297,0 | 75,1 |
| ENSMUSG00000068749 | Psma5 | proteasome (prosome, macropain) subunit, alpha type 5 [Source:MGI Symbol;Acc:MGI:1347009] | 1295,5 | 70,4 |
| ENSMUSG00000031299 | Pdha1 | pyruvate dehydrogenase E1 alpha 1 [Source:MGI Symbol;Acc:MGI:97532] | 1295,4 | 96,9 |
| ENSMUSG00000028194 | Ddah1 | dimethylarginine dimethylaminohydrolase 1 [Source:MGI Symbol;Acc:MGI:1916469] | 1292,0 | 92,8 |
| ENSMUSG00000030282 | Cmas | cytidine monophospho-N-acetylneuraminic acid synthetase [Source:MGI Symbol;Acc:MGI:1337124] | 1290,2 | 97,3 |
| ENSMUSG00000046671 | Mtfr1l | mitochondrial fission regulator 1-like [Source:MGI Symbol;Acc:MGI:1924074] | 1290,2 | 81,0 |
| ENSMUSG00000033933 | Vhl | von Hippel-Lindau tumor suppressor [Source:MGI Symbol;Acc:MGI:103223] | 1289,9 | 92,9 |

| | | | | |
|---|---|---|---|---|
| ENSMUSG00000047260 | Emc6 | ER membrane protein complex subunit 6 [Source:MGI Symbol;Acc:MGI:1913298] | 1288,8 | 94,9 |
| ENSMUSG00000022400 | Rbx1 | ring-box 1 [Source:MGI Symbol;Acc:MGI:1891829] | 1288,7 | 72,0 |
| ENSMUSG00000021395 | Spin1 | spindlin 1 [Source:MGI Symbol;Acc:MGI:109242] | 1285,0 | 79,8 |
| ENSMUSG00000004264 | Phb2 | prohibitin 2 [Source:MGI Symbol;Acc:MGI:102520] | 1283,0 | 98,1 |
| ENSMUSG00000023150 | Ivns1abp | influenza virus NS1A binding protein [Source:MGI Symbol;Acc:MGI:2152389] | 1279,2 | 72,5 |
| ENSMUSG00000022427 | Tomm22 | translocase of outer mitochondrial membrane 22 homolog (yeast) [Source:MGI Symbol;Acc:MGI:2450248] | 1276,7 | 99,6 |
| ENSMUSG00000056692 | D17Wsu92e | DNA segment, Chr 17, Wayne State University 92, expressed [Source:MGI Symbol;Acc:MGI:106281] | 1271,2 | 70,4 |
| ENSMUSG00000066232 | Ipo7 | importin 7 [Source:MGI Symbol;Acc:MGI:2152414] | 1270,2 | 78,3 |
| ENSMUSG00000038150 | Ormdl3 | ORM1-like 3 (S. cerevisiae) [Source:MGI Symbol;Acc:MGI:1913862] | 1269,4 | 93,4 |
| ENSMUSG00000044600 | Smim7 | small integral membrane protein 7 [Source:MGI Symbol;Acc:MGI:1914068] | 1264,3 | 70,6 |
| ENSMUSG00000034424 | Gcsh | glycine cleavage system protein H (aminomethyl carrier) [Source:MGI Symbol;Acc:MGI:1915383] | 1261,9 | 84,4 |
| ENSMUSG00000093989 | Rnasek | ribonuclease, RNase K [Source:MGI Symbol;Acc:MGI:106369] | 1261,2 | 93,4 |
| ENSMUSG00000006058 | Snf8 | SNF8, ESCRT-II complex subunit, homolog (S. cerevisiae) [Source:MGI Symbol;Acc:MGI:1343161] | 1260,7 | 88,4 |
| ENSMUSG00000024012 | Mtch1 | mitochondrial carrier homolog 1 (C. elegans) [Source:MGI Symbol;Acc:MGI:1929261] | 1255,4 | 91,1 |
| ENSMUSG00000051391 | Ywhag | e 3-monooxygenase/tryptophan 5-monooxygenase activation protein, gamma polypeptide [Source:MGI Symbol;Acc:MGI:1 | 1252,4 | 90,2 |
| ENSMUSG00000042406 | Atf4 | activating transcription factor 4 [Source:MGI Symbol;Acc:MGI:88096] | 1249,7 | 80,2 |
| ENSMUSG00000095007 | Igkv12-41 | immunoglobulin kappa chain variable 12-41 [Source:MGI Symbol;Acc:MGI:4439772] | 1242,9 | 98,0 |
| ENSMUSG00000005687 | Bcas2 | breast carcinoma amplified sequence 2 [Source:MGI Symbol;Acc:MGI:1915433] | 1240,0 | 92,0 |
| ENSMUSG00000008859 | Rala | v-ral simian leukemia viral oncogene homolog A (ras related) [Source:MGI Symbol;Acc:MGI:1927243] | 1239,2 | 92,9 |
| ENSMUSG00000039640 | Mrpl12 | mitochondrial ribosomal protein L12 [Source:MGI Symbol;Acc:MGI:1926273] | 1237,6 | 72,9 |
| ENSMUSG00000018326 | Ywhab | ne 3-monooxygenase/tryptophan 5-monooxygenase activation protein, beta polypeptide [Source:MGI Symbol;Acc:MGI:189 | 1236,3 | 75,5 |
| ENSMUSG00000064341 | mt-Nd1 | mitochondrially encoded NADH dehydrogenase 1 [Source:MGI Symbol;Acc:MGI:101787] | 1234,2 | 100,0 |
| ENSMUSG00000015994 | Fnta | farnesyltransferase, CAAX box, alpha [Source:MGI Symbol;Acc:MGI:104683] | 1232,2 | 89,6 |
| ENSMUSG00000022551 | Cyc1 | cytochrome c-1 [Source:MGI Symbol;Acc:MGI:1913695] | 1230,6 | 99,4 |
| ENSMUSG00000001100 | Poldip2 | polymerase (DNA-directed), delta interacting protein 2 [Source:MGI Symbol;Acc:MGI:1915061] | 1230,1 | 96,0 |
| ENSMUSG00000025283 | Sat1 | spermidine/spermine N1-acetyl transferase 1 [Source:MGI Symbol;Acc:MGI:98233] | 1228,3 | 77,3 |
| ENSMUSG00000033953 | Ppp3r1 | protein phosphatase 3, regulatory subunit B, alpha isoform (calcineurin B, type I) [Source:MGI Symbol;Acc:MGI:107172] | 1225,3 | 84,5 |
| ENSMUSG00000054428 | Atpif1 | ATPase inhibitory factor 1 [Source:MGI Symbol;Acc:MGI:1196457] | 1223,5 | 82,9 |
| ENSMUSG00000044408 | Sptssa | serine palmitoyltransferase, small subunit A [Source:MGI Symbol;Acc:MGI:1913399] | 1220,6 | 97,4 |
| ENSMUSG00000029063 | Nadk | NAD kinase [Source:MGI Symbol;Acc:MGI:2183149] | 1217,2 | 77,4 |
| ENSMUSG00000024853 | Sf3b2 | splicing factor 3b, subunit 2 [Source:MGI Symbol;Acc:MGI:2441856] | 1213,7 | 95,9 |
| ENSMUSG00000037331 | Larp1 | La ribonucleoprotein domain family, member 1 [Source:MGI Symbol;Acc:MGI:1890165] | 1213,1 | 73,1 |
| ENSMUSG00000018286 | Psmb6 | proteasome (prosome, macropain) subunit, beta type 6 [Source:MGI Symbol;Acc:MGI:104880] | 1211,9 | 100,0 |
| ENSMUSG00000001472 | Tcf25 | transcription factor 25 (basic helix-loop-helix) [Source:MGI Symbol;Acc:MGI:1914105] | 1210,3 | 84,8 |
| ENSMUSG00000042462 | Dctpp1 | dCTP pyrophosphatase 1 [Source:MGI Symbol;Acc:MGI:1913672] | 1209,7 | 98,0 |
| ENSMUSG00000036570 | Fxyd1 | FXYD domain-containing ion transport regulator 1 [Source:MGI Symbol;Acc:MGI:1889273] | 1209,0 | 70,2 |
| ENSMUSG00000027637 | 1110008F13Rik | RIKEN cDNA 1110008F13 gene [Source:MGI Symbol;Acc:MGI:1914638] | 1202,8 | 96,5 |
| ENSMUSG00000020869 | Lrrc59 | leucine rich repeat containing 59 [Source:MGI Symbol;Acc:MGI:2138133] | 1198,2 | 95,8 |
| ENSMUSG00000078652 | Psme3 | proteaseome (prosome, macropain) activator subunit 3 (PA28 gamma, Ki) [Source:MGI Symbol;Acc:MGI:1096366] | 1197,5 | 91,5 |
| ENSMUSG00000020650 | Bcap29 | B cell receptor associated protein 29 [Source:MGI Symbol;Acc:MGI:101917] | 1196,5 | 86,8 |
| ENSMUSG00000027002 | Nckap1 | NCK-associated protein 1 [Source:MGI Symbol;Acc:MGI:1355333] | 1193,6 | 71,6 |
| ENSMUSG00000028618 | Tmem59 | transmembrane protein 59 [Source:MGI Symbol;Acc:MGI:1929278] | 1189,7 | 83,6 |
| ENSMUSG00000038014 | Fam120a | family with sequence similarity 120, member A [Source:MGI Symbol;Acc:MGI:2446163] | 1188,6 | 76,8 |
| ENSMUSG00000046897 | Zfp740 | zinc finger protein 740 [Source:MGI Symbol;Acc:MGI:1915994] | 1183,3 | 82,6 |
| ENSMUSG00000022679 | Mpv17l | Mpv17 transgene, kidney disease mutant-like [Source:MGI Symbol;Acc:MGI:2135951] | 1182,2 | 73,6 |
| ENSMUSG00000032264 | Zw10 | zw10 kinetochore protein [Source:MGI Symbol;Acc:MGI:1349478] | 1176,8 | 87,2 |
| ENSMUSG00000026918 | Brd3 | bromodomain containing 3 [Source:MGI Symbol;Acc:MGI:1914632] | 1176,1 | 70,0 |
| ENSMUSG00000021156 | Zmynd11 | zinc finger, MYND domain containing 11 [Source:MGI Symbol;Acc:MGI:1913755] | 1175,6 | 77,9 |
| ENSMUSG00000002014 | Ssr4 | signal sequence receptor, delta [Source:MGI Symbol;Acc:MGI:1099464] | 1174,5 | 93,5 |
| ENSMUSG00000044502 | Bod1 | biorientation of chromosomes in cell division 1 [Source:MGI Symbol;Acc:MGI:1916806] | 1174,1 | 80,8 |
| ENSMUSG00000054766 | Set | SET nuclear oncogene [Source:MGI Symbol;Acc:MGI:1860267] | 1173,6 | 72,8 |
| ENSMUSG00000040997 | Abhd4 | abhydrolase domain containing 4 [Source:MGI Symbol;Acc:MGI:1915938] | 1172,5 | 78,9 |
| ENSMUSG00000005732 | Ranbp1 | RAN binding protein 1 [Source:MGI Symbol;Acc:MGI:96269] | 1170,7 | 90,6 |
| ENSMUSG00000015087 | Rabl6 | RAB, member RAS oncogene family-like 6 [Source:MGI Symbol;Acc:MGI:2442633] | 1169,1 | 91,6 |
| ENSMUSG00000079477 | Rab7 | RAB7, member RAS oncogene family [Source:MGI Symbol;Acc:MGI:105068] | 1164,3 | 76,2 |
| ENSMUSG00000029657 | Hsph1 | heat shock 105kDa/110kDa protein 1 [Source:MGI Symbol;Acc:MGI:105053] | 1162,2 | 77,1 |
| ENSMUSG00000020577 | Tspan13 | tetraspanin 13 [Source:MGI Symbol;Acc:MGI:1913359] | 1160,1 | 73,6 |
| ENSMUSG00000031490 | Eif4ebp1 | eukaryotic translation initiation factor 4E binding protein 1 [Source:MGI Symbol;Acc:MGI:103267] | 1159,4 | 84,6 |
| ENSMUSG00000046329 | Slc25a23 | solute carrier family 25 (mitochondrial carrier; phosphate carrier), member 23 [Source:MGI Symbol;Acc:MGI:1914222] | 1150,6 | 91,5 |
| ENSMUSG00000020803 | Txndc17 | thioredoxin domain containing 17 [Source:MGI Symbol;Acc:MGI:1289248] | 1147,2 | 84,3 |
| ENSMUSG00000022635 | Zcrb1 | zinc finger CCHC-type and RNA binding motif 1 [Source:MGI Symbol;Acc:MGI:1914447] | 1145,7 | 97,7 |
| ENSMUSG00000023307 | March5 | membrane-associated ring finger (C3HC4) 5 [Source:MGI Symbol;Acc:MGI:1915207] | 1144,4 | 79,0 |
| ENSMUSG00000066456 | Hmgn3 | high mobility group nucleosomal binding domain 3 [Source:MGI Symbol;Acc:MGI:2138069] | 1141,2 | 91,2 |
| ENSMUSG00000034290 | Nek9 | NIMA (never in mitosis gene a)-related expressed kinase 9 [Source:MGI Symbol;Acc:MGI:2387995] | 1138,0 | 71,1 |
| ENSMUSG00000037242 | Clic4 | chloride intracellular channel 4 (mitochondrial) [Source:MGI Symbol;Acc:MGI:1352754] | 1132,5 | 75,3 |
| ENSMUSG00000020946 | Gosr2 | golgi SNAP receptor complex member 2 [Source:MGI Symbol;Acc:MGI:1927204] | 1132,0 | 82,5 |
| ENSMUSG00000032239 | Rp9 | retinitis pigmentosa 9 (human) [Source:MGI Symbol;Acc:MGI:2157166] | 1130,4 | 87,5 |
| ENSMUSG00000047459 | Dynlrb1 | dynein light chain roadblock-type 1 [Source:MGI Symbol;Acc:MGI:1914318] | 1129,0 | 75,7 |
| ENSMUSG00000033220 | Rac2 | RAS-related C3 botulinum substrate 2 [Source:MGI Symbol;Acc:MGI:97846] | 1128,8 | 73,1 |
| ENSMUSG00000010054 | Tusc2 | tumor suppressor candidate 2 [Source:MGI Symbol;Acc:MGI:1931086] | 1123,9 | 99,2 |
| ENSMUSG00000021149 | Gtpbp4 | GTP binding protein 4 [Source:MGI Symbol;Acc:MGI:1916487] | 1123,0 | 76,2 |
| ENSMUSG00000021877 | Arf4 | ADP-ribosylation factor 4 [Source:MGI Symbol;Acc:MGI:99433] | 1122,5 | 84,6 |
| ENSMUSG00000049760 | 2410015M20Rik | RIKEN cDNA 2410015M20 gene [Source:MGI Symbol;Acc:MGI:2442174] | 1122,5 | 88,7 |
| ENSMUSG00000022110 | Sucla2 | succinate-Coenzyme A ligase, ADP-forming, beta subunit [Source:MGI Symbol;Acc:MGI:1306775] | 1122,4 | 71,2 |
| ENSMUSG00000001105 | Ift20 | intraflagellar transport 20 [Source:MGI Symbol;Acc:MGI:1915585] | 1121,7 | 92,0 |
| ENSMUSG00000003955 | Fam162a | family with sequence similarity 162, member A [Source:MGI Symbol;Acc:MGI:1917436] | 1121,2 | 99,2 |
| ENSMUSG00000023272 | Creld2 | cysteine-rich with EGF-like domains 2 [Source:MGI Symbol;Acc:MGI:1923987] | 1117,8 | 99,9 |
| ENSMUSG00000027937 | Jtb | jumping translocation breakpoint [Source:MGI Symbol;Acc:MGI:1346082] | 1116,5 | 90,4 |
| ENSMUSG00000031311 | Nono | non-POU-domain-containing, octamer binding protein [Source:MGI Symbol;Acc:MGI:1855692] | 1114,0 | 75,5 |
| ENSMUSG00000033938 | Ndufb7 | NADH dehydrogenase (ubiquinone) 1 beta subcomplex, 7 [Source:MGI Symbol;Acc:MGI:1914166] | 1113,2 | 91,0 |
| ENSMUSG00000008393 | Carhsp1 | calcium regulated heat stable protein 1 [Source:MGI Symbol;Acc:MGI:1196368] | 1107,9 | 90,7 |
| ENSMUSG00000060938 | Rpl26 | ribosomal protein L26 [Source:MGI Symbol;Acc:MGI:106022] | 1107,0 | 91,8 |
| ENSMUSG00000027999 | Pla2g12a | phospholipase A2, group XIIA [Source:MGI Symbol;Acc:MGI:1913600] | 1106,9 | 80,9 |
| ENSMUSG00000048865 | Arhgap30 | Rho GTPase activating protein 30 [Source:MGI Symbol;Acc:MGI:2684948] | 1103,5 | 81,3 |
| ENSMUSG00000016257 | Slmo2 | slowmo homolog 2 (Drosophila) [Source:MGI Symbol;Acc:MGI:1913640] | 1103,2 | 85,0 |
| ENSMUSG00000067847 | Romo1 | reactive oxygen species modulator 1 [Source:MGI Symbol;Acc:MGI:1914317] | 1100,3 | 91,4 |
| ENSMUSG00000023075 | Akirin1 | akirin 1 [Source:MGI Symbol;Acc:MGI:1915300] | 1099,9 | 75,1 |
| ENSMUSG00000034064 | Poglut1 | protein O-glucosyltransferase 1 [Source:MGI Symbol;Acc:MGI:2444232] | 1099,2 | 75,9 |
| ENSMUSG00000035133 | Arhgap5 | Rho GTPase activating protein 5 [Source:MGI Symbol;Acc:MGI:1332637] | 1097,4 | 74,7 |
| ENSMUSG00000036450 | Hif1an | hypoxia-inducible factor 1, alpha subunit inhibitor [Source:MGI Symbol;Acc:MGI:2442345] | 1095,8 | 77,3 |
| ENSMUSG00000002477 | Snrpd1 | small nuclear ribonucleoprotein D1 [Source:MGI Symbol;Acc:MGI:98344] | 1093,9 | 79,2 |
| ENSMUSG00000039515 | Ppp2r4 | protein phosphatase 2A activator, regulatory subunit B [Source:MGI Symbol;Acc:MGI:1346006] | 1092,1 | 81,4 |
| ENSMUSG00000033065 | Pfkm | phosphofructokinase, muscle [Source:MGI Symbol;Acc:MGI:97548] | 1091,3 | 94,1 |
| ENSMUSG00000007029 | Vars | valyl-tRNA synthetase [Source:MGI Symbol;Acc:MGI:90675] | 1089,3 | 74,8 |
| ENSMUSG00000021111 | Papola | poly (A) polymerase alpha [Source:MGI Symbol;Acc:MGI:109301] | 1088,5 | 76,1 |
| ENSMUSG00000025439 | Clns1a | chloride channel, nucleotide-sensitive, 1A [Source:MGI Symbol;Acc:MGI:109638] | 1086,4 | 96,5 |
| ENSMUSG00000030788 | Rnf141 | ring finger protein 141 [Source:MGI Symbol;Acc:MGI:1914400] | 1086,2 | 76,1 |
| ENSMUSG00000025728 | Pigq | phosphatidylinositol glycan anchor biosynthesis, class Q [Source:MGI Symbol;Acc:MGI:1333114] | 1086,0 | 75,7 |
| ENSMUSG00000020402 | Vdac1 | voltage-dependent anion channel 1 [Source:MGI Symbol;Acc:MGI:106919] | 1085,6 | 85,6 |
| ENSMUSG00000034868 | Myl12b | myosin, light chain 12B, regulatory [Source:MGI Symbol;Acc:MGI:107494] | 1085,4 | 78,4 |
| ENSMUSG00000026664 | Phyh | phytanoyl-CoA hydroxylase [Source:MGI Symbol;Acc:MGI:891978] | 1084,9 | 99,5 |
| ENSMUSG00000024953 | Prdx5 | peroxiredoxin 5 [Source:MGI Symbol;Acc:MGI:1859821] | 1083,2 | 78,8 |
| ENSMUSG00000001891 | Ugp2 | UDP-glucose pyrophosphorylase 2 [Source:MGI Symbol;Acc:MGI:2183447] | 1082,7 | 89,3 |
| ENSMUSG00000022295 | Atp6v1c1 | ATPase, H+ transporting, lysosomal V1 subunit C1 [Source:MGI Symbol;Acc:MGI:1913585] | 1081,6 | 96,3 |
| ENSMUSG00000053907 | Mat2a | methionine adenosyltransferase II, alpha [Source:MGI Symbol;Acc:MGI:2443731] | 1081,5 | 88,6 |
| ENSMUSG00000007891 | Ctsd | cathepsin D [Source:MGI Symbol;Acc:MGI:88562] | 1081,1 | 84,6 |
| ENSMUSG00000006315 | Tmem147 | transmembrane protein 147 [Source:MGI Symbol;Acc:MGI:1915011] | 1077,9 | 92,9 |
| ENSMUSG00000019362 | D8Ertd738e | DNA segment, Chr 8, ERATO Doi 738, expressed [Source:MGI Symbol;Acc:MGI:1289231] | 1071,6 | 93,3 |
| ENSMUSG00000026245 | Farsb | phenylalanyl-tRNA synthetase, beta subunit [Source:MGI Symbol;Acc:MGI:1346035] | 1069,7 | 84,0 |
| ENSMUSG00000027433 | Xrn2 | 5'-3' exoribonuclease 2 [Source:MGI Symbol;Acc:MGI:894687] | 1064,7 | 72,2 |
| ENSMUSG00000060992 | Copz1 | coatomer protein complex, subunit zeta 1 [Source:MGI Symbol;Acc:MGI:1929063] | 1062,8 | 99,6 |
| ENSMUSG00000033272 | Slc35a4 | solute carrier family 35, member A4 [Source:MGI Symbol;Acc:MGI:1915093] | 1061,5 | 70,8 |
| ENSMUSG00000037149 | Ddx1 | DEAD (Asp-Glu-Ala-Asp) box polypeptide 1 [Source:MGI Symbol;Acc:MGI:2144727] | 1061,2 | 74,7 |
| ENSMUSG00000036352 | Ubac1 | ubiquitin associated domain containing 1 [Source:MGI Symbol;Acc:MGI:1920995] | 1053,1 | 94,0 |
| ENSMUSG00000028861 | Mrps15 | mitochondrial ribosomal protein S15 [Source:MGI Symbol;Acc:MGI:1913657] | 1053,0 | 93,2 |
| ENSMUSG00000024594 | Prrc1 | proline-rich coiled-coil 1 [Source:MGI Symbol;Acc:MGI:1916106] | 1051,8 | 70,2 |
| ENSMUSG00000029147 | Ppm1g | protein phosphatase 1G (formerly 2C), magnesium-dependent, gamma isoform [Source:MGI Symbol;Acc:MGI:106065] | 1051,6 | 78,6 |

| | | | | |
|---|---|---|---|---|
| ENSMUSG00000024208 | Uqcc2 | ubiquinol-cytochrome c reductase complex assembly factor 2 [Source:MGI Symbol;Acc:MGI:1914517] | 1051,3 | 80,5 |
| ENSMUSG00000025403 | Shmt2 | serine hydroxymethyltransferase 2 (mitochondrial) [Source:MGI Symbol;Acc:MGI:1277989] | 1050,2 | 93,3 |
| ENSMUSG00000019471 | Cdc37 | cell division cycle 37 [Source:MGI Symbol;Acc:MGI:109531] | 1049,9 | 80,2 |
| ENSMUSG00000021102 | Glrx5 | glutaredoxin 5 [Source:MGI Symbol;Acc:MGI:1920296] | 1048,7 | 97,1 |
| ENSMUSG00000034120 | Srsf2 | serine/arginine-rich splicing factor 2 [Source:MGI Symbol;Acc:MGI:98284] | 1046,8 | 96,0 |
| ENSMUSG00000053317 | Sec61b | Sec61 beta subunit [Source:MGI Symbol;Acc:MGI:1913462] | 1045,7 | 71,3 |
| ENSMUSG00000020153 | Ndufs7 | NADH dehydrogenase (ubiquinone) Fe-S protein 7 [Source:MGI Symbol;Acc:MGI:1922656] | 1043,4 | 77,9 |
| ENSMUSG00000039470 | Zdhhc2 | zinc finger, DHHC domain containing 2 [Source:MGI Symbol;Acc:MGI:1923452] | 1042,4 | 85,5 |
| ENSMUSG00000022658 | Tagln3 | transgelin 3 [Source:MGI Symbol;Acc:MGI:1926784] | 1041,8 | 82,5 |
| ENSMUSG00000039615 | Stub1 | STIP1 homology and U-Box containing protein 1 [Source:MGI Symbol;Acc:MGI:1891731] | 1040,5 | 79,8 |
| ENSMUSG00000032002 | Dcun1d5 | DCN1, defective in cullin neddylation 1, domain containing 5 (S. cerevisiae) [Source:MGI Symbol;Acc:MGI:1924113] | 1039,6 | 98,7 |
| ENSMUSG00000024146 | Cript | cysteine-rich PDZ-binding protein [Source:MGI Symbol;Acc:MGI:1929655] | 1036,8 | 97,3 |
| ENSMUSG00000040048 | Ndufb10 | NADH dehydrogenase (ubiquinone) 1 beta subcomplex, 10 [Source:MGI Symbol;Acc:MGI:1915592] | 1034,9 | 87,5 |
| ENSMUSG00000024665 | Fads2 | fatty acid desaturase 2 [Source:MGI Symbol;Acc:MGI:1930079] | 1032,5 | 100,0 |
| ENSMUSG00000022212 | Cpne6 | copine VI [Source:MGI Symbol;Acc:MGI:1334445] | 1030,4 | 83,1 |
| ENSMUSG00000051234 | Rnf7 | ring finger protein 7 [Source:MGI Symbol;Acc:MGI:1337096] | 1027,5 | 93,3 |
| ENSMUSG00000003814 | Calr | calreticulin [Source:MGI Symbol;Acc:MGI:88252] | 1027,0 | 84,2 |
| ENSMUSG00000013593 | Ndufs2 | NADH dehydrogenase (ubiquinone) Fe-S protein 2 [Source:MGI Symbol;Acc:MGI:2385112] | 1026,1 | 89,1 |
| ENSMUSG00000063888 | Rpl7l1 | ribosomal protein L7-like 1 [Source:MGI Symbol;Acc:MGI:1913479] | 1022,3 | 87,3 |
| ENSMUSG00000024601 | Isoc1 | isochorismatase domain containing 1 [Source:MGI Symbol;Acc:MGI:1913557] | 1021,6 | 94,6 |
| ENSMUSG00000025795 | Rassf3 | Ras association (RalGDS/AF-6) domain family member 3 [Source:MGI Symbol;Acc:MGI:2179722] | 1021,2 | 97,1 |
| ENSMUSG00000061393 | Acvr2b | activin receptor IIB [Source:MGI Symbol;Acc:MGI:87912] | 1020,8 | 94,8 |
| ENSMUSG00000027067 | Ssrp1 | structure specific recognition protein 1 [Source:MGI Symbol;Acc:MGI:107912] | 1017,1 | 92,8 |
| ENSMUSG00000006024 | Napa | N-ethylmaleimide sensitive fusion protein attachment protein alpha [Source:MGI Symbol;Acc:MGI:104563] | 1016,9 | 82,2 |
| ENSMUSG00000041881 | Ndufa7 | NADH dehydrogenase (ubiquinone) 1 alpha subcomplex, 7 (B14.5a) [Source:MGI Symbol;Acc:MGI:1913666] | 1014,0 | 96,9 |
| ENSMUSG00000032383 | Ppib | peptidylprolyl isomerase B [Source:MGI Symbol;Acc:MGI:97750] | 1012,5 | 92,4 |
| ENSMUSG00000020671 | Rab10 | RAB10, member RAS oncogene family [Source:MGI Symbol;Acc:MGI:105066] | 1010,7 | 85,0 |
| ENSMUSG00000038965 | Ube2l3 | ubiquitin-conjugating enzyme E2L 3 [Source:MGI Symbol;Acc:MGI:109240] | 1010,6 | 72,1 |
| ENSMUSG00000021486 | Prelid1 | PRELI domain containing 1 [Source:MGI Symbol;Acc:MGI:1913744] | 1003,3 | 86,1 |
| ENSMUSG00000015149 | Sirt2 | sirtuin 2 [Source:MGI Symbol;Acc:MGI:1927664] | 1002,6 | 73,5 |
| ENSMUSG00000030216 | Wbp11 | WW domain binding protein 11 [Source:MGI Symbol;Acc:MGI:1891823] | 1001,5 | 91,4 |
| ENSMUSG00000022477 | Aco2 | aconitase 2, mitochondrial [Source:MGI Symbol;Acc:MGI:87880] | 1001,4 | 89,9 |
| ENSMUSG00000049612 | Omg | oligodendrocyte myelin glycoprotein [Source:MGI Symbol;Acc:MGI:106586] | 1000,5 | 79,6 |
| ENSMUSG00000009549 | Srp14 | signal recognition particle 14 [Source:MGI Symbol;Acc:MGI:107169] | 999,9 | 70,9 |
| ENSMUSG00000027620 | Rbm39 | RNA binding motif protein 39 [Source:MGI Symbol;Acc:MGI:2157953] | 998,5 | 70,5 |
| ENSMUSG00000021134 | Srsf5 | serine/arginine-rich splicing factor 5 [Source:MGI Symbol;Acc:MGI:98287] | 998,0 | 90,2 |
| ENSMUSG00000027091 | Zc3h15 | zinc finger CCCH-type containing 15 [Source:MGI Symbol;Acc:MGI:1919747] | 997,9 | 79,9 |
| ENSMUSG00000029776 | Hibadh | 3-hydroxyisobutyrate dehydrogenase [Source:MGI Symbol;Acc:MGI:1889802] | 997,6 | 92,0 |
| ENSMUSG00000067786 | Nnat | neuronatin [Source:MGI Symbol;Acc:MGI:104716] | 995,1 | 84,5 |
| ENSMUSG00000037710 | Cisd1 | CDGSH iron sulfur domain 1 [Source:MGI Symbol;Acc:MGI:1261855] | 993,5 | 80,9 |
| ENSMUSG00000020649 | Rrm2 | ribonucleotide reductase M2 [Source:MGI Symbol;Acc:MGI:98181] | 991,1 | 83,6 |
| ENSMUSG00000028622 | Mrpl37 | mitochondrial ribosomal protein L37 [Source:MGI Symbol;Acc:MGI:1926268] | 989,6 | 95,9 |
| ENSMUSG00000025613 | Cct8 | chaperonin containing Tcp1, subunit 8 (theta) [Source:MGI Symbol;Acc:MGI:107183] | 988,9 | 74,1 |
| ENSMUSG00000015671 | Psma2 | proteasome (prosome, macropain) subunit, alpha type 2 [Source:MGI Symbol;Acc:MGI:104885] | 988,7 | 96,4 |
| ENSMUSG00000029640 | Usp12 | ubiquitin specific peptidase 12 [Source:MGI Symbol;Acc:MGI:1270128] | 988,3 | 77,5 |
| ENSMUSG00000032788 | Pdxk | pyridoxal (pyridoxine, vitamin B6) kinase [Source:MGI Symbol;Acc:MGI:1351869] | 985,8 | 79,0 |
| ENSMUSG00000017843 | Ppp2r5c | protein phosphatase 2, regulatory subunit B', gamma [Source:MGI Symbol;Acc:MGI:1349475] | 985,5 | 73,5 |
| ENSMUSG00000034563 | Ccpg1 | cell cycle progression 1 [Source:MGI Symbol;Acc:MGI:1196419] | 985,1 | 76,1 |
| ENSMUSG00000002957 | Ap2a2 | adaptor-related protein complex 2, alpha 2 subunit [Source:MGI Symbol;Acc:MGI:101920] | 984,4 | 80,1 |
| ENSMUSG00000022199 | Slc22a17 | solute carrier family 22 (organic cation transporter), member 17 [Source:MGI Symbol;Acc:MGI:1926225] | 983,5 | 96,5 |
| ENSMUSG00000009863 | Sdhb | succinate dehydrogenase complex, subunit B, iron sulfur (Ip) [Source:MGI Symbol;Acc:MGI:1914930] | 982,6 | 87,3 |
| ENSMUSG00000032215 | Rsl24d1 | ribosomal L24 domain containing 1 [Source:MGI Symbol;Acc:MGI:2681840] | 976,1 | 76,1 |
| ENSMUSG00000024997 | Prdx3 | peroxiredoxin 3 [Source:MGI Symbol;Acc:MGI:88034] | 975,9 | 92,3 |
| ENSMUSG00000079677 | Fdx1l | ferredoxin 1-like [Source:MGI Symbol;Acc:MGI:1915415] | 973,4 | 100,0 |
| ENSMUSG00000021794 | Glud1 | glutamate dehydrogenase 1 [Source:MGI Symbol;Acc:MGI:95753] | 967,1 | 75,0 |
| ENSMUSG00000020840 | Blmh | bleomycin hydrolase [Source:MGI Symbol;Acc:MGI:1345186] | 965,8 | 71,5 |
| ENSMUSG00000020307 | Cdc34 | cell division cycle 34 [Source:MGI Symbol;Acc:MGI:102657] | 964,5 | 97,3 |
| ENSMUSG00000068882 | Ssb | Sjogren syndrome antigen B [Source:MGI Symbol;Acc:MGI:98423] | 963,2 | 81,9 |
| ENSMUSG00000096210 | H1f0 | H1 histone family, member 0 [Source:MGI Symbol;Acc:MGI:95893] | 958,6 | 83,6 |
| ENSMUSG00000047557 | Lxn | latexin [Source:MGI Symbol;Acc:MGI:107633] | 958,1 | 96,4 |
| ENSMUSG00000020922 | Lsm12 | LSM12 homolog (S. cerevisiae) [Source:MGI Symbol;Acc:MGI:1919592] | 955,8 | 94,9 |
| ENSMUSG00000001576 | Ergic1 | endoplasmic reticulum-golgi intermediate compartment (ERGIC) 1 [Source:MGI Symbol;Acc:MGI:1914708] | 953,0 | 76,9 |
| ENSMUSG00000032349 | Elovl5 | ELOVL family member 5, elongation of long chain fatty acids (yeast) [Source:MGI Symbol;Acc:MGI:1916051] | 951,8 | 82,8 |
| ENSMUSG00000041084 | Ostc | oligosaccharyltransferase complex subunit [Source:MGI Symbol;Acc:MGI:1913607] | 951,8 | 96,3 |
| ENSMUSG00000047187 | Rab2a | RAB2A, member RAS oncogene family [Source:MGI Symbol;Acc:MGI:1928750] | 950,8 | 92,0 |
| ENSMUSG00000015291 | Gdi1 | guanosine diphosphate (GDP) dissociation inhibitor 1 [Source:MGI Symbol;Acc:MGI:99846] | 949,7 | 85,0 |
| ENSMUSG00000036299 | BC031181 | cDNA sequence BC031181 [Source:MGI Symbol;Acc:MGI:3039614] | 949,6 | 97,6 |
| ENSMUSG00000023020 | Cox14 | cytochrome c oxidase assembly protein 14 [Source:MGI Symbol;Acc:MGI:1913629] | 949,1 | 94,2 |
| ENSMUSG00000025935 | Tram1 | translocating chain-associating membrane protein 1 [Source:MGI Symbol;Acc:MGI:1919515] | 948,7 | 90,3 |
| ENSMUSG00000052681 | Rap1b | RAS related protein 1b [Source:MGI Symbol;Acc:MGI:894315] | 947,0 | 95,0 |
| ENSMUSG00000089715 | Cbx6 | chromobox 6 [Source:MGI Symbol;Acc:MGI:3512628] | 946,5 | 71,7 |
| ENSMUSG00000024143 | Rhoq | ras homolog gene family, member Q [Source:MGI Symbol;Acc:MGI:1931553] | 943,2 | 71,3 |
| ENSMUSG00000001833 | Sept7 | septin 7 [Source:MGI Symbol;Acc:MGI:1335094] | 938,9 | 73,2 |
| ENSMUSG00000002475 | Abhd3 | abhydrolase domain containing 3 [Source:MGI Symbol;Acc:MGI:2147183] | 937,8 | 94,6 |
| ENSMUSG00000030245 | Golt1b | golgi transport 1B [Source:MGI Symbol;Acc:MGI:1914214] | 937,5 | 78,7 |
| ENSMUSG00000020483 | Dynll2 | dynein light chain LC8-type 2 [Source:MGI Symbol;Acc:MGI:1915347] | 936,4 | 85,9 |
| ENSMUSG00000071014 | Ndufb6 | NADH dehydrogenase (ubiquinone) 1 beta subcomplex, 6 [Source:MGI Symbol;Acc:MGI:2684983] | 935,5 | 86,3 |
| ENSMUSG00000060802 | B2m | beta-2 microglobulin [Source:MGI Symbol;Acc:MGI:88127] | 934,4 | 98,7 |
| ENSMUSG00000015837 | Sqstm1 | sequestosome 1 [Source:MGI Symbol;Acc:MGI:107931] | 934,4 | 75,4 |
| ENSMUSG00000021520 | Uqcrb | ubiquinol-cytochrome c reductase binding protein [Source:MGI Symbol;Acc:MGI:1914780] | 933,2 | 90,7 |
| ENSMUSG00000029066 | Mrpl20 | mitochondrial ribosomal protein L20 [Source:MGI Symbol;Acc:MGI:2137221] | 930,9 | 78,2 |
| ENSMUSG00000092837 | Rpph1 | ribonuclease P RNA component H1 [Source:MGI Symbol;Acc:MGI:1934664] | 930,6 | 98,3 |
| ENSMUSG00000039703 | Nploc4 | nuclear protein localization 4 homolog (S. cerevisiae) [Source:MGI Symbol;Acc:MGI:2679787] | 930,4 | 91,0 |
| ENSMUSG00000024181 | Mrpl28 | mitochondrial ribosomal protein L28 [Source:MGI Symbol;Acc:MGI:1915861] | 929,5 | 99,7 |
| ENSMUSG00000017715 | Pgs1 | phosphatidylglycerophosphate synthase 1 [Source:MGI Symbol;Acc:MGI:1921701] | 928,1 | 70,7 |
| ENSMUSG00000002010 | Idh3g | isocitrate dehydrogenase 3 (NAD+), gamma [Source:MGI Symbol;Acc:MGI:1099463] | 926,6 | 97,4 |
| ENSMUSG00000041556 | Fbxo2 | F-box protein 2 [Source:MGI Symbol;Acc:MGI:2446216] | 923,1 | 98,9 |
| ENSMUSG00000033307 | Mif | macrophage migration inhibitory factor [Source:MGI Symbol;Acc:MGI:96982] | 921,3 | 96,3 |
| ENSMUSG00000021737 | Psmd6 | proteasome (prosome, macropain) 26S subunit, non-ATPase, 6 [Source:MGI Symbol;Acc:MGI:1913663] | 919,4 | 93,7 |
| ENSMUSG00000024571 | Gm16286 | predicted gene 16286 [Source:MGI Symbol;Acc:MGI:3833940] | 918,9 | 90,8 |
| ENSMUSG00000032212 | Sltm | SAFB-like, transcription modulator [Source:MGI Symbol;Acc:MGI:1913910] | 917,6 | 72,5 |
| ENSMUSG00000024858 | Adrbk1 | adrenergic receptor kinase, beta 1 [Source:MGI Symbol;Acc:MGI:87940] | 917,4 | 72,9 |
| ENSMUSG00000033047 | Eif3l | eukaryotic translation initiation factor 3, subunit L [Source:MGI Symbol;Acc:MGI:2386251] | 916,1 | 98,7 |
| ENSMUSG00000078572 | 1810043H04Rik | RIKEN cDNA 1810043H04 gene [Source:MGI Symbol;Acc:MGI:1913676] | 915,9 | 86,3 |
| ENSMUSG00000059119 | Nap1l4 | nucleosome assembly protein 1-like 4 [Source:MGI Symbol;Acc:MGI:1316687] | 915,7 | 89,3 |
| ENSMUSG00000033917 | Gde1 | glycerophosphodiester phosphodiesterase 1 [Source:MGI Symbol;Acc:MGI:1891827] | 914,1 | 92,5 |
| ENSMUSG00000053898 | Ech1 | enoyl coenzyme A hydratase 1, peroxisomal [Source:MGI Symbol;Acc:MGI:1858208] | 913,6 | 92,6 |
| ENSMUSG00000066900 | Suds3 | suppressor of defective silencing 3 homolog (S. cerevisiae) [Source:MGI Symbol;Acc:MGI:1919204] | 912,5 | 85,4 |
| ENSMUSG00000032554 | Trf | transferrin [Source:MGI Symbol;Acc:MGI:98821] | 911,8 | 77,4 |
| ENSMUSG00000002274 | Metrn | meteorin, glial cell differentiation regulator [Source:MGI Symbol;Acc:MGI:1917333] | 911,6 | 90,2 |
| ENSMUSG00000027303 | Ptpra | protein tyrosine phosphatase, receptor type, A [Source:MGI Symbol;Acc:MGI:97808] | 910,8 | 81,4 |
| ENSMUSG00000020149 | Rab1a | RAB1A, member RAS oncogene family [Source:MGI Symbol;Acc:MGI:97842] | 910,7 | 76,1 |
| ENSMUSG00000029840 | Mtpn | myotrophin [Source:MGI Symbol;Acc:MGI:99445] | 910,6 | 74,4 |
| ENSMUSG00000016256 | Ctsz | cathepsin Z [Source:MGI Symbol;Acc:MGI:1891190] | 909,2 | 92,8 |
| ENSMUSG00000034118 | Tpst1 | protein-tyrosine sulfotransferase 1 [Source:MGI Symbol;Acc:MGI:1298231] | 907,1 | 85,4 |
| ENSMUSG00000030605 | Mfge8 | milk fat globule-EGF factor 8 protein [Source:MGI Symbol;Acc:MGI:102768] | 906,0 | 81,3 |
| ENSMUSG00000025978 | Rftn2 | raftlin family member 2 [Source:MGI Symbol;Acc:MGI:1921263] | 905,9 | 75,6 |
| ENSMUSG00000029777 | Gars | glycyl-tRNA synthetase [Source:MGI Symbol;Acc:MGI:2449057] | 905,6 | 82,9 |
| ENSMUSG00000022744 | Cldnd1 | claudin domain containing 1 [Source:MGI Symbol;Acc:MGI:2447860] | 901,3 | 86,5 |
| ENSMUSG00000031604 | Msmo1 | methylsterol monoxygenase 1 [Source:MGI Symbol;Acc:MGI:1913484] | 899,8 | 76,6 |
| ENSMUSG00000071662 | Polr2g | polymerase (RNA) II (DNA directed) polypeptide G [Source:MGI Symbol;Acc:MGI:1914960] | 899,2 | 99,7 |
| ENSMUSG00000090639 | Gm20425 | predicted gene 20425 [Source:MGI Symbol;Acc:MGI:5141890] | 899,2 | 79,8 |
| ENSMUSG00000027984 | Hadh | hydroxyacyl-Coenzyme A dehydrogenase [Source:MGI Symbol;Acc:MGI:96009] | 897,9 | 95,0 |
| ENSMUSG00000020440 | Arf5 | ADP-ribosylation factor 5 [Source:MGI Symbol;Acc:MGI:99434] | 895,7 | 93,0 |
| ENSMUSG00000039275 | Foxk2 | forkhead box K2 [Source:MGI Symbol;Acc:MGI:1916087] | 894,5 | 73,9 |
| ENSMUSG00000024132 | Eci1 | enoyl-Coenzyme A delta isomerase 1 [Source:MGI Symbol;Acc:MGI:94871] | 893,3 | 99,1 |
| ENSMUSG00000029146 | Snx17 | sorting nexin 17 [Source:MGI Symbol;Acc:MGI:2387801] | 892,7 | 94,0 |

| | | | | |
|---|---|---|---|---|
| ENSMUSG00000031403 | Dkc1 | dyskeratosis congenita 1, dyskerin [Source:MGI Symbol;Acc:MGI:1861727] | 890,4 | 70,7 |
| ENSMUSG00000026229 | Psmd1 | proteasome (prosome, macropain) 26S subunit, non-ATPase, 1 [Source:MGI Symbol;Acc:MGI:1917497] | 889,1 | 85,9 |
| ENSMUSG00000031617 | Tmem184c | transmembrane protein 184C [Source:MGI Symbol;Acc:MGI:2384562] | 886,8 | 86,4 |
| ENSMUSG00000056820 | Tsnax | translin-associated factor X [Source:MGI Symbol;Acc:MGI:1855672] | 883,9 | 94,0 |
| ENSMUSG00000057858 | Fam204a | family with sequence similarity 204, member A [Source:MGI Symbol;Acc:MGI:1289174] | 883,5 | 77,3 |
| ENSMUSG00000094117 | Igkv3-12 | immunoglobulin kappa variable 3-12 [Source:MGI Symbol;Acc:MGI:1330815] | 883,0 | 98,6 |
| ENSMUSG00000039197 | Adk | adenosine kinase [Source:MGI Symbol;Acc:MGI:87930] | 879,4 | 95,8 |
| ENSMUSG00000074457 | S100a16 | S100 calcium binding protein A16 [Source:MGI Symbol;Acc:MGI:1915110] | 874,7 | 71,5 |
| ENSMUSG00000024429 | Gnl1 | guanine nucleotide binding protein-like 1 [Source:MGI Symbol;Acc:MGI:95764] | 872,9 | 76,7 |
| ENSMUSG00000029432 | Gbas | glioblastoma amplified sequence [Source:MGI Symbol;Acc:MGI:1278343] | 871,9 | 75,4 |
| ENSMUSG00000026341 | Actr3 | ARP3 actin-related protein 3 [Source:MGI Symbol;Acc:MGI:1921367] | 869,6 | 78,5 |
| ENSMUSG00000035953 | Tmem55b | transmembrane protein 55b [Source:MGI Symbol;Acc:MGI:2448501] | 868,0 | 72,9 |
| ENSMUSG00000027170 | Eif3m | eukaryotic translation initiation factor 3, subunit M [Source:MGI Symbol;Acc:MGI:1351744] | 867,8 | 92,2 |
| ENSMUSG00000032397 | Tipin | timeless interacting protein [Source:MGI Symbol;Acc:MGI:1921571] | 864,7 | 95,4 |
| ENSMUSG00000007950 | Abhd8 | abhydrolase domain containing 8 [Source:MGI Symbol;Acc:MGI:1918946] | 863,0 | 99,3 |
| ENSMUSG00000040055 | Gjb6 | gap junction protein, beta 6 [Source:MGI Symbol;Acc:MGI:107588] | 862,6 | 92,2 |
| ENSMUSG00000031987 | Egln1 | egl-9 family hypoxia-inducible factor 1 [Source:MGI Symbol;Acc:MGI:1932286] | 861,6 | 86,7 |
| ENSMUSG00000038975 | Rabggtb | Rab geranylgeranyl transferase, b subunit [Source:MGI Symbol;Acc:MGI:99537] | 861,5 | 89,7 |
| ENSMUSG00000053398 | Phgdh | 3-phosphoglycerate dehydrogenase [Source:MGI Symbol;Acc:MGI:1355330] | 860,5 | 73,6 |
| ENSMUSG00000025006 | Sorbs1 | sorbin and SH3 domain containing 1 [Source:MGI Symbol;Acc:MGI:700014] | 860,5 | 78,3 |
| ENSMUSG00000066568 | Lsm14a | LSM14 homolog A (SCD6, S. cerevisiae) [Source:MGI Symbol;Acc:MGI:1914320] | 858,9 | 77,4 |
| ENSMUSG00000028745 | Capzb | capping protein (actin filament) muscle Z-line, beta [Source:MGI Symbol;Acc:MGI:104652] | 858,4 | 72,5 |
| ENSMUSG00000016946 | Kctd5 | potassium channel tetramerisation domain containing 5 [Source:MGI Symbol;Acc:MGI:1916509] | 857,7 | 94,1 |
| ENSMUSG00000032067 | Pts | 6-pyruvoyl-tetrahydropterin synthase [Source:MGI Symbol;Acc:MGI:1338783] | 854,8 | 97,4 |
| ENSMUSG00000052738 | Suclg1 | succinate-CoA ligase, GDP-forming, alpha subunit [Source:MGI Symbol;Acc:MGI:1927234] | 854,5 | 80,1 |
| ENSMUSG00000026223 | Itm2c | integral membrane protein 2C [Source:MGI Symbol;Acc:MGI:1927594] | 854,1 | 86,4 |
| ENSMUSG00000040618 | Pck2 | phosphoenolpyruvate carboxykinase 2 (mitochondrial) [Source:MGI Symbol;Acc:MGI:1860456] | 853,9 | 73,8 |
| ENSMUSG00000028822 | Tmem50a | transmembrane protein 50A [Source:MGI Symbol;Acc:MGI:1919067] | 851,0 | 96,8 |
| ENSMUSG00000001542 | Ell2 | elongation factor RNA polymerase II 2 [Source:MGI Symbol;Acc:MGI:2183438] | 850,9 | 86,9 |
| ENSMUSG00000023723 | Mrps23 | mitochondrial ribosomal protein S23 [Source:MGI Symbol;Acc:MGI:1928138] | 850,6 | 82,9 |
| ENSMUSG00000107283 | Mpv17 | MpV17 mitochondrial inner membrane protein [Source:MGI Symbol;Acc:MGI:97138] | 849,6 | 74,6 |
| ENSMUSG00000022892 | App | amyloid beta (A4) precursor protein [Source:MGI Symbol;Acc:MGI:88059] | 848,1 | 87,6 |
| ENSMUSG00000034908 | Sidt2 | SID1 transmembrane family, member 2 [Source:MGI Symbol;Acc:MGI:2446134] | 847,5 | 71,1 |
| ENSMUSG00000020152 | Actr2 | ARP2 actin-related protein 2 [Source:MGI Symbol;Acc:MGI:1913963] | 846,6 | 70,3 |
| ENSMUSG00000028837 | Psmb2 | proteasome (prosome, macropain) subunit, beta type 2 [Source:MGI Symbol;Acc:MGI:1347045] | 846,2 | 95,6 |
| ENSMUSG00000020708 | Psmc5 | protease (prosome, macropain) 26S subunit, ATPase 5 [Source:MGI Symbol;Acc:MGI:105047] | 844,5 | 99,3 |
| ENSMUSG00000015776 | Med22 | mediator complex subunit 22 [Source:MGI Symbol;Acc:MGI:98446] | 844,0 | 74,7 |
| ENSMUSG00000010406 | Mrpl52 | mitochondrial ribosomal protein L52 [Source:MGI Symbol;Acc:MGI:1916086] | 843,5 | 95,1 |
| ENSMUSG00000025190 | Got1 | glutamic-oxaloacetic transaminase 1, soluble [Source:MGI Symbol;Acc:MGI:95791] | 838,7 | 89,7 |
| ENSMUSG00000015806 | Qdpr | quinoid dihydropteridine reductase [Source:MGI Symbol;Acc:MGI:97836] | 837,6 | 75,6 |
| ENSMUSG00000015092 | Edf1 | endothelial differentiation-related factor 1 [Source:MGI Symbol;Acc:MGI:1891227] | 836,7 | 97,4 |
| ENSMUSG00000028277 | Ube2j1 | ubiquitin-conjugating enzyme E2J 1 [Source:MGI Symbol;Acc:MGI:1926245] | 836,3 | 70,2 |
| ENSMUSG00000030161 | Gabarapl1 | gamma-aminobutyric acid (GABA) A receptor-associated protein-like 1 [Source:MGI Symbol;Acc:MGI:1914980] | 835,5 | 90,6 |
| ENSMUSG00000021973 | Micu2 | mitochondrial calcium uptake 2 [Source:MGI Symbol;Acc:MGI:1915764] | 835,0 | 95,6 |
| ENSMUSG00000051853 | Arf3 | ADP-ribosylation factor 3 [Source:MGI Symbol;Acc:MGI:99432] | 834,6 | 81,7 |
| ENSMUSG00000025162 | Csnk1d | casein kinase 1, delta [Source:MGI Symbol;Acc:MGI:1355272] | 834,4 | 90,9 |
| ENSMUSG00000029048 | Rer1 | RER1 retention in endoplasmic reticulum 1 homolog (S. cerevisiae) [Source:MGI Symbol;Acc:MGI:1915080] | 834,3 | 86,8 |
| ENSMUSG00000002997 | Prkar2b | protein kinase, cAMP dependent regulatory, type II beta [Source:MGI Symbol;Acc:MGI:97760] | 831,9 | 80,6 |
| ENSMUSG00000028851 | Nudc | nuclear distribution gene C homolog (Aspergillus) [Source:MGI Symbol;Acc:MGI:106014] | 831,4 | 97,4 |
| ENSMUSG00000024914 | Drap1 | Dr1 associated protein 1 (negative cofactor 2 alpha) [Source:MGI Symbol;Acc:MGI:1913806] | 830,5 | 80,9 |
| ENSMUSG00000022553 | Maf1 | MAF1 homolog (S. cerevisiae) [Source:MGI Symbol;Acc:MGI:1916127] | 829,6 | 92,0 |
| ENSMUSG00000025151 | Maged1 | melanoma antigen, family D, 1 [Source:MGI Symbol;Acc:MGI:1930187] | 829,6 | 93,9 |
| ENSMUSG00000003345 | Csnk1g2 | casein kinase 1, gamma 2 [Source:MGI Symbol;Acc:MGI:1920014] | 823,8 | 85,2 |
| ENSMUSG00000035376 | Hacd2 | 3-hydroxyacyl-CoA dehydratase 2 [Source:MGI Symbol;Acc:MGI:1918007] | 823,7 | 70,0 |
| ENSMUSG00000002332 | Dhrs1 | dehydrogenase/reductase (SDR family) member 1 [Source:MGI Symbol;Acc:MGI:1196314] | 823,2 | 95,6 |
| ENSMUSG00000022122 | Ednrb | endothelin receptor type B [Source:MGI Symbol;Acc:MGI:102720] | 823,0 | 80,4 |
| ENSMUSG00000007721 | Ccdc124 | coiled-coil domain containing 124 [Source:MGI Symbol;Acc:MGI:1916403] | 816,8 | 99,9 |
| ENSMUSG00000027490 | E2f1 | E2F transcription factor 1 [Source:MGI Symbol;Acc:MGI:101941] | 816,1 | 73,1 |
| ENSMUSG00000045294 | Insig1 | insulin induced gene 1 [Source:MGI Symbol;Acc:MGI:1916289] | 813,8 | 84,5 |
| ENSMUSG00000037060 | Prkcdbp | protein kinase C, delta binding protein [Source:MGI Symbol;Acc:MGI:1923422] | 813,4 | 81,6 |
| ENSMUSG00000038615 | Nfe2l1 | nuclear factor, erythroid derived 2,-like 1 [Source:MGI Symbol;Acc:MGI:99421] | 812,9 | 80,4 |
| ENSMUSG00000022346 | Myc | myelocytomatosis oncogene [Source:MGI Symbol;Acc:MGI:97250] | 811,1 | 77,8 |
| ENSMUSG00000061032 | Rrp1 | ribosomal RNA processing 1 homolog (S. cerevisiae) [Source:MGI Symbol;Acc:MGI:1203500] | 808,9 | 95,0 |
| ENSMUSG00000038936 | Sccpdh | saccharopine dehydrogenase (putative) [Source:MGI Symbol;Acc:MGI:1924486] | 807,8 | 80,9 |
| ENSMUSG00000025351 | Cd63 | CD63 antigen [Source:MGI Symbol;Acc:MGI:99529] | 807,8 | 84,7 |
| ENSMUSG00000029103 | Lrpap1 | low density lipoprotein receptor-related protein associated protein 1 [Source:MGI Symbol;Acc:MGI:96829] | 804,7 | 87,8 |
| ENSMUSG00000022564 | Grina | ıate receptor, ionotropic, N-methyl D-aspartate-associated protein 1 (glutamate binding) [Source:MGI Symbol;Acc:MGI:191 | 804,4 | 94,2 |
| ENSMUSG00000053286 | Trmt1l | tRNA methyltransferase 1 like [Source:MGI Symbol;Acc:MGI:1916185] | 804,3 | 80,8 |
| ENSMUSG00000007050 | Lsm2 | LSM2 homolog, U6 small nuclear RNA associated (S. cerevisiae) [Source:MGI Symbol;Acc:MGI:90676] | 803,9 | 93,9 |
| ENSMUSG00000059336 | Slc14a1 | solute carrier family 14 (urea transporter), member 1 [Source:MGI Symbol;Acc:MGI:1351654] | 803,1 | 83,2 |
| ENSMUSG00000001666 | Ddt | D-dopachrome tautomerase [Source:MGI Symbol;Acc:MGI:1298381] | 802,9 | 81,6 |
| ENSMUSG00000038459 | Abhd17c | abhydrolase domain containing 17C [Source:MGI Symbol;Acc:MGI:1917428] | 801,8 | 90,8 |
| ENSMUSG00000036908 | Unc93b1 | unc-93 homolog B1 (C. elegans) [Source:MGI Symbol;Acc:MGI:1859307] | 801,5 | 91,8 |
| ENSMUSG00000003346 | Abhd17a | abhydrolase domain containing 17A [Source:MGI Symbol;Acc:MGI:106388] | 799,2 | 86,8 |
| ENSMUSG00000024072 | Yipf4 | Yip1 domain family, member 4 [Source:MGI Symbol;Acc:MGI:1915114] | 798,2 | 76,9 |
| ENSMUSG00000020646 | Mboat2 | membrane bound O-acyltransferase domain containing 2 [Source:MGI Symbol;Acc:MGI:1914466] | 797,0 | 88,0 |
| ENSMUSG00000024740 | Ddb1 | damage specific DNA binding protein 1 [Source:MGI Symbol;Acc:MGI:1202384] | 796,9 | 84,8 |
| ENSMUSG00000019254 | Ppp1r12c | protein phosphatase 1, regulatory (inhibitor) subunit 12C [Source:MGI Symbol;Acc:MGI:1924258] | 796,5 | 86,8 |
| ENSMUSG00000042043 | Tbca | tubulin cofactor A [Source:MGI Symbol;Acc:MGI:107549] | 796,3 | 100,0 |
| ENSMUSG00000016528 | Mapkapk2 | MAP kinase-activated protein kinase 2 [Source:MGI Symbol;Acc:MGI:109298] | 794,7 | 83,5 |
| ENSMUSG00000024780 | Cdc37l1 | cell division cycle 37-like 1 [Source:MGI Symbol;Acc:MGI:1914322] | 794,2 | 72,4 |
| ENSMUSG00000013662 | Atad1 | ATPase family, AAA domain containing 1 [Source:MGI Symbol;Acc:MGI:1915229] | 793,1 | 79,4 |
| ENSMUSG00000036106 | Prr5 | proline rich 5 (renal) [Source:MGI Symbol;Acc:MGI:1924714] | 793,0 | 89,5 |
| ENSMUSG00000074238 | Ap1ar | adaptor-related protein complex 1 associated regulatory protein [Source:MGI Symbol;Acc:MGI:2384822] | 792,6 | 89,5 |
| ENSMUSG00000015937 | H2afy | H2A histone family, member Y [Source:MGI Symbol;Acc:MGI:1349392] | 786,4 | 88,2 |
| ENSMUSG00000021508 | Cxcl14 | chemokine (C-X-C motif) ligand 14 [Source:MGI Symbol;Acc:MGI:1888514] | 785,2 | 90,4 |
| ENSMUSG00000035478 | Mbd3 | methyl-CpG binding domain protein 3 [Source:MGI Symbol;Acc:MGI:1333812] | 784,8 | 79,9 |
| ENSMUSG00000032314 | Etfa | electron transferring flavoprotein, alpha polypeptide [Source:MGI Symbol;Acc:MGI:106092] | 783,2 | 87,3 |
| ENSMUSG00000091337 | Eid1 | EP300 interacting inhibitor of differentiation 1 [Source:MGI Symbol;Acc:MGI:1889651] | 782,2 | 95,4 |
| ENSMUSG00000022500 | Litaf | LPS-induced TN factor [Source:MGI Symbol;Acc:MGI:1929512] | 779,6 | 77,5 |
| ENSMUSG00000064345 | mt-Nd2 | mitochondrially encoded NADH dehydrogenase 2 [Source:MGI Symbol;Acc:MGI:102500] | 779,5 | 90,8 |
| ENSMUSG00000038696 | Mapkap1 | mitogen-activated protein kinase associated protein 1 [Source:MGI Symbol;Acc:MGI:2444554] | 777,0 | 88,8 |
| ENSMUSG00000024537 | Psmg2 | proteasome (prosome, macropain) assembly chaperone 2 [Source:MGI Symbol;Acc:MGI:1922901] | 776,5 | 98,7 |
| ENSMUSG00000030842 | Lamtor1 | late endosomal/lysosomal adaptor, MAPK and MTOR activator 1 [Source:MGI Symbol;Acc:MGI:1913758] | 775,3 | 78,0 |
| ENSMUSG00000030890 | Ilk | integrin linked kinase [Source:MGI Symbol;Acc:MGI:1195267] | 775,1 | 74,3 |
| ENSMUSG00000028907 | Utp11l | UTP11-like, U3 small nucleolar ribonucleoprotein, (yeast) [Source:MGI Symbol;Acc:MGI:1914455] | 775,0 | 78,0 |
| ENSMUSG00000002580 | Mien1 | migration and invasion enhancer 1 [Source:MGI Symbol;Acc:MGI:1913678] | 773,7 | 97,1 |
| ENSMUSG00000019978 | Epb41l2 | erythrocyte membrane protein band 4.1 like 2 [Source:MGI Symbol;Acc:MGI:103009] | 773,7 | 79,5 |
| ENSMUSG00000068732 | Tmem167b | transmembrane protein 167B [Source:MGI Symbol;Acc:MGI:1914745] | 773,3 | 78,9 |
| ENSMUSG00000026568 | Mpc2 | mitochondrial pyruvate carrier 2 [Source:MGI Symbol;Acc:MGI:1917706] | 773,2 | 73,6 |
| ENSMUSG00000057497 | Fam136a | family with sequence similarity 136, member A [Source:MGI Symbol;Acc:MGI:1913738] | 770,8 | 95,4 |
| ENSMUSG00000011884 | Gltp | glycolipid transfer protein [Source:MGI Symbol;Acc:MGI:1929253] | 770,0 | 91,6 |
| ENSMUSG00000031848 | Lsm4 | LSM4 homolog, U6 small nuclear RNA associated (S. cerevisiae) [Source:MGI Symbol;Acc:MGI:1354692] | 768,6 | 90,5 |
| ENSMUSG00000032405 | Pias1 | protein inhibitor of activated STAT 1 [Source:MGI Symbol;Acc:MGI:1913125] | 768,3 | 70,1 |
| ENSMUSG00000064337 | mt-Rnr1 | mitochondrially encoded 12S rRNA [Source:MGI Symbol;Acc:MGI:102493] | 767,6 | 91,6 |
| ENSMUSG00000023048 | Prr13 | proline rich 13 [Source:MGI Symbol;Acc:MGI:1913401] | 766,9 | 74,3 |
| ENSMUSG00000055850 | Rnf181 | ring finger protein 181 [Source:MGI Symbol;Acc:MGI:1913760] | 765,7 | 76,4 |
| ENSMUSG00000030704 | Rab6a | RAB6A, member RAS oncogene family [Source:MGI Symbol;Acc:MGI:894313] | 764,7 | 87,5 |
| ENSMUSG00000003813 | Rad23a | RAD23a homolog (S. cerevisiae) [Source:MGI Symbol;Acc:MGI:105126] | 763,6 | 79,4 |
| ENSMUSG00000005846 | Rsl1d1 | ribosomal L1 domain containing 1 [Source:MGI Symbol;Acc:MGI:1913659] | 763,2 | 82,0 |
| ENSMUSG00000032350 | Gclc | glutamate-cysteine ligase, catalytic subunit [Source:MGI Symbol;Acc:MGI:104990] | 762,6 | 89,0 |
| ENSMUSG00000025173 | Wdr45b | WD repeat domain 45B [Source:MGI Symbol;Acc:MGI:1914090] | 762,6 | 88,8 |
| ENSMUSG00000024327 | Slc39a7 | solute carrier family 39 (zinc transporter), member 7 [Source:MGI Symbol;Acc:MGI:95909] | 762,1 | 96,5 |
| ENSMUSG00000016554 | Eif3d | eukaryotic translation initiation factor 3, subunit D [Source:MGI Symbol;Acc:MGI:1933181] | 761,8 | 73,0 |
| ENSMUSG00000015932 | Dstn | destrin [Source:MGI Symbol;Acc:MGI:1929270] | 761,7 | 72,5 |
| ENSMUSG00000024588 | Fech | ferrochelatase [Source:MGI Symbol;Acc:MGI:95513] | 759,2 | 84,0 |
| ENSMUSG00000030876 | Mettl9 | methyltransferase like 9 [Source:MGI Symbol;Acc:MGI:1914862] | 757,3 | 83,2 |

| | | | | |
|---|---|---|---|---|
| ENSMUSG00000024194 | Cuta | cutA divalent cation tolerance homolog [Source:MGI Symbol;Acc:MGI:1914925] | 753,7 | 99,4 |
| ENSMUSG00000022265 | Ank | progressive ankylosis [Source:MGI Symbol;Acc:MGI:3045421] | 751,8 | 87,4 |
| ENSMUSG00000024091 | Vapa | vesicle-associated membrane protein, associated protein A [Source:MGI Symbol;Acc:MGI:1353561] | 751,0 | 92,2 |
| ENSMUSG00000090247 | Bloc1s1 | biogenesis of lysosomal organelles complex-1, subunit 1 [Source:MGI Symbol;Acc:MGI:1195276] | 743,8 | 70,8 |
| ENSMUSG00000038412 | Higd1a | HIG1 domain family, member 1A [Source:MGI Symbol;Acc:MGI:1930666] | 743,5 | 86,5 |
| ENSMUSG00000032434 | Cmtm6 | CKLF-like MARVEL transmembrane domain containing 6 [Source:MGI Symbol;Acc:MGI:2447165] | 743,0 | 74,5 |
| ENSMUSG00000021681 | Aggf1 | angiogenic factor with G patch and FHA domains 1 [Source:MGI Symbol;Acc:MGI:1913799] | 739,3 | 94,1 |
| ENSMUSG00000109493 | RP23-109E24.10 | | 738,9 | 83,2 |
| ENSMUSG00000051154 | Commd3 | COMM domain containing 3 [Source:MGI Symbol;Acc:MGI:88218] | 738,1 | 95,0 |
| ENSMUSG00000023883 | Phf10 | PHD finger protein 10 [Source:MGI Symbol;Acc:MGI:1919307] | 737,5 | 73,9 |
| ENSMUSG00000029780 | Nt5c3 | 5'-nucleotidase, cytosolic III [Source:MGI Symbol;Acc:MGI:1927186] | 737,3 | 91,5 |
| ENSMUSG00000062963 | Ufc1 | ubiquitin-fold modifier conjugating enzyme 1 [Source:MGI Symbol;Acc:MGI:1913405] | 737,1 | 87,5 |
| ENSMUSG00000028833 | Ncdn | neurochondrin [Source:MGI Symbol;Acc:MGI:1347351] | 737,1 | 87,5 |
| ENSMUSG00000041733 | Coq5 | coenzyme Q5 methyltransferase [Source:MGI Symbol;Acc:MGI:1098643] | 737,0 | 87,2 |
| ENSMUSG00000022634 | Yaf2 | YY1 associated factor 2 [Source:MGI Symbol;Acc:MGI:1914307] | 735,3 | 94,6 |
| ENSMUSG00000030327 | Necap1 | NECAP endocytosis associated 1 [Source:MGI Symbol;Acc:MGI:1914852] | 734,2 | 90,1 |
| ENSMUSG00000037119 | D15Ertd621e | DNA segment, Chr 15, ERATO Doi 621, expressed [Source:MGI Symbol;Acc:MGI:1277178] | 734,1 | 78,0 |
| ENSMUSG00000027221 | Chst1 | carbohydrate (keratan sulfate Gal-6) sulfotransferase 1 [Source:MGI Symbol;Acc:MGI:1924219] | 733,8 | 93,9 |
| ENSMUSG00000028821 | Syf2 | SYF2 homolog, RNA splicing factor (S. cerevisiae) [Source:MGI Symbol;Acc:MGI:1915842] | 733,5 | 91,7 |
| ENSMUSG00000026496 | Parp1 | poly (ADP-ribose) polymerase family, member 1 [Source:MGI Symbol;Acc:MGI:1340806] | 731,3 | 85,8 |
| ENSMUSG00000022678 | Nde1 | nuclear distribution gene E homolog 1 (A nidulans) [Source:MGI Symbol;Acc:MGI:1914453] | 727,3 | 85,4 |
| ENSMUSG00000037916 | Ndufv1 | NADH dehydrogenase (ubiquinone) flavoprotein 1 [Source:MGI Symbol;Acc:MGI:107851] | 727,1 | 85,8 |
| ENSMUSG00000002102 | Psmc3 | proteasome (prosome, macropain) 26S subunit, ATPase 3 [Source:MGI Symbol;Acc:MGI:1098754] | 726,3 | 77,0 |
| ENSMUSG00000028953 | Abcf2 | ATP-binding cassette, sub-family F (GCN20), member 2 [Source:MGI Symbol;Acc:MGI:1351657] | 726,2 | 87,7 |
| ENSMUSG00000052934 | Fbxo31 | F-box protein 31 [Source:MGI Symbol;Acc:MGI:1354708] | 725,5 | 80,2 |
| ENSMUSG00000020076 | Ddx50 | DEAD (Asp-Glu-Ala-Asp) box polypeptide 50 [Source:MGI Symbol;Acc:MGI:2182303] | 722,1 | 97,0 |
| ENSMUSG00000031610 | Scrg1 | scrapie responsive gene 1 [Source:MGI Symbol;Acc:MGI:1328308] | 719,7 | 93,6 |
| ENSMUSG00000003062 | Stard3nl | STARD3 N-terminal like [Source:MGI Symbol;Acc:MGI:1923455] | 719,6 | 73,5 |
| ENSMUSG00000063856 | Gpx1 | glutathione peroxidase 1 [Source:MGI Symbol;Acc:MGI:104887] | 719,3 | 80,1 |
| ENSMUSG00000022789 | Dnm1l | dynamin 1-like [Source:MGI Symbol;Acc:MGI:1921256] | 718,5 | 88,8 |
| ENSMUSG00000030960 | Mettl10 | methyltransferase like 10 [Source:MGI Symbol;Acc:MGI:1919346] | 718,5 | 94,6 |
| ENSMUSG00000028393 | Alad | aminolevulinate, delta-, dehydratase [Source:MGI Symbol;Acc:MGI:96853] | 717,0 | 89,8 |
| ENSMUSG00000098650 | Gm28048 | predicted gene, 28048 [Source:MGI Symbol;Acc:MGI:5547784] | 716,4 | 96,4 |
| ENSMUSG00000000378 | Ccm2 | cerebral cavernous malformation 2 [Source:MGI Symbol;Acc:MGI:2384924] | 713,1 | 84,4 |
| ENSMUSG00000021981 | Cab39l | calcium binding protein 39-like [Source:MGI Symbol;Acc:MGI:1914081] | 712,6 | 73,9 |
| ENSMUSG00000040759 | Cmtm5 | CKLF-like MARVEL transmembrane domain containing 5 [Source:MGI Symbol;Acc:MGI:2447164] | 712,5 | 97,0 |
| ENSMUSG00000063576 | Klhdc3 | kelch domain containing 3 [Source:MGI Symbol;Acc:MGI:2651568] | 711,5 | 92,8 |
| ENSMUSG00000048978 | Nrsn1 | neurensin 1 [Source:MGI Symbol;Acc:MGI:894662] | 710,9 | 84,2 |
| ENSMUSG00000031948 | Kars | lysyl-tRNA synthetase [Source:MGI Symbol;Acc:MGI:1934754] | 709,9 | 83,4 |
| ENSMUSG00000027255 | Arfgap2 | ADP-ribosylation factor GTPase activating protein 2 [Source:MGI Symbol;Acc:MGI:1924288] | 709,3 | 80,9 |
| ENSMUSG00000040374 | Pex2 | peroxisomal biogenesis factor 2 [Source:MGI Symbol;Acc:MGI:107486] | 709,0 | 72,1 |
| ENSMUSG00000031807 | Pgls | 6-phosphogluconolactonase [Source:MGI Symbol;Acc:MGI:1913421] | 708,7 | 70,6 |
| ENSMUSG00000031532 | Saraf | store-operated calcium entry-associated regulatory factor [Source:MGI Symbol;Acc:MGI:1915137] | 707,6 | 87,4 |
| ENSMUSG00000031708 | Tecr | trans-2,3-enoyl-CoA reductase [Source:MGI Symbol;Acc:MGI:1915408] | 707,0 | 84,4 |
| ENSMUSG00000035545 | Leng8 | leukocyte receptor cluster (LRC) member 8 [Source:MGI Symbol;Acc:MGI:2142195] | 705,5 | 75,9 |
| ENSMUSG00000031197 | Vbp1 | von Hippel-Lindau binding protein 1 [Source:MGI Symbol;Acc:MGI:1333804] | 704,1 | 70,4 |
| ENSMUSG00000073639 | Rab18 | RAB18, member RAS oncogene family [Source:MGI Symbol;Acc:MGI:102790] | 697,2 | 74,5 |
| ENSMUSG00000003721 | Insig2 | insulin induced gene 2 [Source:MGI Symbol;Acc:MGI:1920249] | 697,1 | 74,6 |
| ENSMUSG00000022982 | Sod1 | superoxide dismutase 1, soluble [Source:MGI Symbol;Acc:MGI:98351] | 695,1 | 97,7 |
| ENSMUSG00000019738 | Polr2i | polymerase (RNA) II (DNA directed) polypeptide I [Source:MGI Symbol;Acc:MGI:1917170] | 692,9 | 90,1 |
| ENSMUSG00000041236 | Vps41 | vacuolar protein sorting 41 (yeast) [Source:MGI Symbol;Acc:MGI:1929215] | 692,6 | 86,2 |
| ENSMUSG00000001700 | Gramd3 | GRAM domain containing 3 [Source:MGI Symbol;Acc:MGI:1914815] | 691,6 | 70,1 |
| ENSMUSG00000028041 | Adam15 | a disintegrin and metallopeptidase domain 15 (metargidin) [Source:MGI Symbol;Acc:MGI:1333882] | 690,6 | 95,8 |
| ENSMUSG00000040010 | Slc7a5 | solute carrier family 7 (cationic amino acid transporter, y+ system), member 5 [Source:MGI Symbol;Acc:MGI:1298205] | 689,5 | 87,3 |
| ENSMUSG00000020571 | Pdia6 | protein disulfide isomerase associated 6 [Source:MGI Symbol;Acc:MGI:1919103] | 689,3 | 77,3 |
| ENSMUSG00000067148 | Polr1c | polymerase (RNA) I polypeptide C [Source:MGI Symbol;Acc:MGI:103288] | 688,9 | 82,3 |
| ENSMUSG00000106918 | Mrpl33 | mitochondrial ribosomal protein L33 [Source:MGI Symbol;Acc:MGI:2137225] | 686,9 | 93,5 |
| ENSMUSG00000021025 | Nfkbia | nuclear factor of kappa light polypeptide gene enhancer in B cells inhibitor, alpha [Source:MGI Symbol;Acc:MGI:104741] | 686,1 | 82,8 |
| ENSMUSG00000037197 | Rbm17 | RNA binding motif protein 17 [Source:MGI Symbol;Acc:MGI:1924188] | 686,0 | 82,7 |
| ENSMUSG00000090213 | Tmem189 | transmembrane protein 189 [Source:MGI Symbol;Acc:MGI:2142624] | 684,0 | 93,5 |
| ENSMUSG00000017615 | Tnfaip1 | tumor necrosis factor, alpha-induced protein 1 (endothelial) [Source:MGI Symbol;Acc:MGI:104961] | 682,4 | 74,1 |
| ENSMUSG00000016503 | Gtf3a | general transcription factor III A [Source:MGI Symbol;Acc:MGI:1913846] | 681,3 | 97,2 |
| ENSMUSG00000006736 | Tspan31 | tetraspanin 31 [Source:MGI Symbol;Acc:MGI:1914375] | 680,3 | 99,6 |
| ENSMUSG00000028797 | Tmem234 | transmembrane protein 234 [Source:MGI Symbol;Acc:MGI:1924049] | 679,1 | 84,5 |
| ENSMUSG00000023495 | Pcbp4 | poly(rC) binding protein 4 [Source:MGI Symbol;Acc:MGI:1890471] | 678,9 | 72,8 |
| ENSMUSG00000004610 | Etfb | electron transferring flavoprotein, beta polypeptide [Source:MGI Symbol;Acc:MGI:106098] | 678,4 | 93,6 |
| ENSMUSG00000064326 | Siva1 | SIVA1, apoptosis-inducing factor [Source:MGI Symbol;Acc:MGI:1353606] | 678,3 | 88,6 |
| ENSMUSG00000020375 | Rufy1 | RUN and FYVE domain containing 1 [Source:MGI Symbol;Acc:MGI:2429762] | 678,0 | 75,5 |
| ENSMUSG00000021577 | Sdha | succinate dehydrogenase complex, subunit A, flavoprotein (Fp) [Source:MGI Symbol;Acc:MGI:1914195] | 677,7 | 88,8 |
| ENSMUSG00000025260 | Hsd17b10 | hydroxysteroid (17-beta) dehydrogenase 10 [Source:MGI Symbol;Acc:MGI:1333871] | 677,4 | 92,4 |
| ENSMUSG00000024835 | Coro1b | coronin, actin binding protein 1B [Source:MGI Symbol;Acc:MGI:1345963] | 677,2 | 98,9 |
| ENSMUSG00000022476 | Polr3h | polymerase (RNA) III (DNA directed) polypeptide H [Source:MGI Symbol;Acc:MGI:1926179] | 676,8 | 77,4 |
| ENSMUSG00000030082 | Sec61a1 | Sec61 alpha 1 subunit (S. cerevisiae) [Source:MGI Symbol;Acc:MGI:1858417] | 675,5 | 89,1 |
| ENSMUSG00000020519 | Sap30l | SAP30-like [Source:MGI Symbol;Acc:MGI:1354709] | 674,8 | 97,8 |
| ENSMUSG00000064367 | mt-Nd5 | mitochondrially encoded NADH dehydrogenase 5 [Source:MGI Symbol;Acc:MGI:102496] | 674,7 | 83,1 |
| ENSMUSG00000026500 | Cox20 | COX20 Cox2 chaperone [Source:MGI Symbol;Acc:MGI:1913609] | 674,4 | 92,5 |
| ENSMUSG00000024474 | Ik | IK cytokine [Source:MGI Symbol;Acc:MGI:1345142] | 673,7 | 94,2 |
| ENSMUSG00000028772 | Zcchc17 | zinc finger, CCHC domain containing 17 [Source:MGI Symbol;Acc:MGI:1919955] | 673,2 | 72,1 |
| ENSMUSG00000021606 | Ndufs6 | NADH dehydrogenase (ubiquinone) Fe-S protein 6 [Source:MGI Symbol;Acc:MGI:107932] | 672,1 | 86,4 |
| ENSMUSG00000025465 | Echs1 | enoyl Coenzyme A hydratase, short chain, 1, mitochondrial [Source:MGI Symbol;Acc:MGI:2136460] | 668,6 | 93,8 |
| ENSMUSG00000021417 | Eci2 | enoyl-Coenzyme A delta isomerase 2 [Source:MGI Symbol;Acc:MGI:1346064] | 668,6 | 78,0 |
| ENSMUSG00000069744 | Psmb3 | proteasome (prosome, macropain) subunit, beta type 3 [Source:MGI Symbol;Acc:MGI:1347014] | 668,4 | 92,7 |
| ENSMUSG00000022257 | Laptm4b | lysosomal-associated protein transmembrane 4B [Source:MGI Symbol;Acc:MGI:1890494] | 667,0 | 83,1 |
| ENSMUSG00000032112 | Trappc4 | trafficking protein particle complex 4 [Source:MGI Symbol;Acc:MGI:1926211] | 666,8 | 73,4 |
| ENSMUSG00000035297 | Cops4 | COP9 signalosome subunit 4 [Source:MGI Symbol;Acc:MGI:1349414] | 666,6 | 81,6 |
| ENSMUSG00000041736 | Tspo | translocator protein [Source:MGI Symbol;Acc:MGI:88222] | 666,6 | 78,9 |
| ENSMUSG00000058709 | Egln2 | egl-9 family hypoxia-inducible factor 2 [Source:MGI Symbol;Acc:MGI:1932287] | 666,3 | 77,8 |
| ENSMUSG00000008036 | Ap2s1 | adaptor-related protein complex 2, sigma 1 subunit [Source:MGI Symbol;Acc:MGI:2141861] | 666,2 | 95,8 |
| ENSMUSG00000030541 | Idh2 | isocitrate dehydrogenase 2 (NADP+), mitochondrial [Source:MGI Symbol;Acc:MGI:96414] | 665,9 | 95,7 |
| ENSMUSG00000044224 | Dnajc21 | DnaJ heat shock protein family (Hsp40) member C21 [Source:MGI Symbol;Acc:MGI:1925371] | 665,5 | 73,5 |
| ENSMUSG00000013646 | Sh3bp5l | SH3 binding domain protein 5 like [Source:MGI Symbol;Acc:MGI:1933124] | 665,5 | 83,9 |
| ENSMUSG00000024981 | Acsl5 | acyl-CoA synthetase long-chain family member 5 [Source:MGI Symbol;Acc:MGI:1919129] | 664,8 | 82,5 |
| ENSMUSG00000030603 | Psmc4 | proteasome (prosome, macropain) 26S subunit, ATPase, 4 [Source:MGI Symbol;Acc:MGI:1346093] | 663,4 | 91,4 |
| ENSMUSG00000047084 | Ngrn | neugrin, neurite outgrowth associated [Source:MGI Symbol;Acc:MGI:1933212] | 662,7 | 98,2 |
| ENSMUSG00000022241 | Tars | threonyl-tRNA synthetase [Source:MGI Symbol;Acc:MGI:106314] | 662,4 | 78,3 |
| ENSMUSG00000000743 | Chmp1a | charged multivesicular body protein 1A [Source:MGI Symbol;Acc:MGI:1920159] | 662,3 | 92,6 |
| ENSMUSG00000038552 | Fndc4 | fibronectin type III domain containing 4 [Source:MGI Symbol;Acc:MGI:1917195] | 662,1 | 74,4 |
| ENSMUSG00000004394 | Tmed4 | transmembrane emp24 protein transport domain containing 4 [Source:MGI Symbol;Acc:MGI:1915070] | 662,0 | 88,8 |
| ENSMUSG00000024197 | Plin3 | perilipin 3 [Source:MGI Symbol;Acc:MGI:1914155] | 661,2 | 91,0 |
| ENSMUSG00000020744 | Slc25a19 | olute carrier family 25 (mitochondrial thiamine pyrophosphate carrier), member 19 [Source:MGI Symbol;Acc:MGI:1914533 | 659,4 | 75,9 |
| ENSMUSG00000022043 | Trim35 | tripartite motif-containing 35 [Source:MGI Symbol;Acc:MGI:1914104] | 658,8 | 74,3 |
| ENSMUSG00000040599 | Mis12 | MIS12 homolog (yeast) [Source:MGI Symbol;Acc:MGI:1914389] | 658,4 | 79,4 |
| ENSMUSG00000028757 | Ddost | dolichyl-di-phosphooligosaccharide-protein glycotransferase [Source:MGI Symbol;Acc:MGI:1194508] | 656,6 | 88,1 |
| ENSMUSG00000036850 | Mrpl41 | mitochondrial ribosomal protein L41 [Source:MGI Symbol;Acc:MGI:1333816] | 655,5 | 97,4 |
| ENSMUSG00000038909 | Kat7 | K(lysine) acetyltransferase 7 [Source:MGI Symbol;Acc:MGI:2182799] | 655,2 | 85,6 |
| ENSMUSG00000038607 | Gng10 | guanine nucleotide binding protein (G protein), gamma 10 [Source:MGI Symbol;Acc:MGI:1336169] | 654,1 | 86,0 |
| ENSMUSG00000020536 | Llgl1 | lethal giant larvae homolog 1 (Drosophila) [Source:MGI Symbol;Acc:MGI:102682] | 652,1 | 73,9 |
| ENSMUSG00000038967 | Pdk2 | pyruvate dehydrogenase kinase, isoenzyme 2 [Source:MGI Symbol;Acc:MGI:1343087] | 651,7 | 89,7 |
| ENSMUSG00000061613 | U2af1 | U2 small nuclear ribonucleoprotein auxiliary factor (U2AF) 1 [Source:MGI Symbol;Acc:MGI:98884] | 649,9 | 87,7 |
| ENSMUSG00000025208 | Mrpl43 | mitochondrial ribosomal protein L43 [Source:MGI Symbol;Acc:MGI:2137229] | 649,9 | 92,0 |
| ENSMUSG00000033530 | Ttc7b | tetratricopeptide repeat domain 7B [Source:MGI Symbol;Acc:MGI:2144724] | 648,6 | 76,1 |
| ENSMUSG00000040722 | Scamp5 | secretory carrier membrane protein 5 [Source:MGI Symbol;Acc:MGI:1928948] | 648,5 | 93,3 |
| ENSMUSG00000002345 | Borcs8 | BLOC-1 related complex subunit 8 [Source:MGI Symbol;Acc:MGI:1919618] | 647,0 | 71,2 |
| ENSMUSG00000041057 | Wdr43 | WD repeat domain 43 [Source:MGI Symbol;Acc:MGI:1919765] | 646,8 | 74,4 |
| ENSMUSG00000092345 | Gm20503 | predicted gene 20503 [Source:MGI Symbol;Acc:MGI:5141968] | 646,8 | 77,3 |
| ENSMUSG00000047514 | Tspyl1 | testis-specific protein, Y-encoded-like 1 [Source:MGI Symbol;Acc:MGI:1298395] | 646,6 | 81,9 |
| ENSMUSG00000032463 | Faim | Fas apoptotic inhibitory molecule [Source:MGI Symbol;Acc:MGI:1344387] | 645,8 | 89,5 |

| | | | | |
|---|---|---|---|---|
| ENSMUSG00000022463 | Srebf2 | sterol regulatory element binding factor 2 [Source:MGI Symbol;Acc:MGI:107585] | 644,5 | 73,9 |
| ENSMUSG00000024403 | Atp6v1g2 | ATPase, H+ transporting, lysosomal V1 subunit G2 [Source:MGI Symbol;Acc:MGI:1913487] | 643,6 | 79,5 |
| ENSMUSG00000026309 | Ilkap | integrin-linked kinase-associated serine/threonine phosphatase 2C [Source:MGI Symbol;Acc:MGI:1914694] | 643,6 | 76,6 |
| ENSMUSG00000022474 | Pmm1 | phosphomannomutase 1 [Source:MGI Symbol;Acc:MGI:1353418] | 643,3 | 94,8 |
| ENSMUSG00000003199 | Mpnd | MPN domain containing [Source:MGI Symbol;Acc:MGI:1915297] | 642,3 | 76,4 |
| ENSMUSG00000029017 | Pmpcb | peptidase (mitochondrial processing) beta [Source:MGI Symbol;Acc:MGI:1920328] | 642,3 | 76,6 |
| ENSMUSG00000000959 | Oxa1l | oxidase assembly 1-like [Source:MGI Symbol;Acc:MGI:1916339] | 640,2 | 88,7 |
| ENSMUSG00000036752 | Tubb4b | tubulin, beta 4B class IVB [Source:MGI Symbol;Acc:MGI:1915472] | 640,0 | 80,8 |
| ENSMUSG00000028029 | Aimp1 | aminoacyl tRNA synthetase complex-interacting multifunctional protein 1 [Source:MGI Symbol;Acc:MGI:102774] | 638,3 | 71,6 |
| ENSMUSG00000033863 | Klf9 | Kruppel-like factor 9 [Source:MGI Symbol;Acc:MGI:1333856] | 638,1 | 73,3 |
| ENSMUSG00000040234 | Tm7sf3 | transmembrane 7 superfamily member 3 [Source:MGI Symbol;Acc:MGI:1914873] | 637,6 | 71,5 |
| ENSMUSG00000027273 | Snap25 | synaptosomal-associated protein 25 [Source:MGI Symbol;Acc:MGI:98331] | 637,5 | 90,0 |
| ENSMUSG00000061461 | Smim20 | small integral membrane protein 20 [Source:MGI Symbol;Acc:MGI:1913528] | 637,1 | 97,0 |
| ENSMUSG00000020993 | Trappc6b | trafficking protein particle complex 6B [Source:MGI Symbol;Acc:MGI:1925482] | 636,2 | 87,5 |
| ENSMUSG00000028436 | Dcaf12 | DDB1 and CUL4 associated factor 12 [Source:MGI Symbol;Acc:MGI:1916220] | 633,1 | 79,8 |
| ENSMUSG00000025337 | Sbds | Shwachman-Bodian-Diamond syndrome homolog (human) [Source:MGI Symbol;Acc:MGI:1913961] | 633,0 | 73,6 |
| ENSMUSG00000047547 | Cltb | clathrin, light polypeptide (Lcb) [Source:MGI Symbol;Acc:MGI:1921575] | 632,6 | 86,7 |
| ENSMUSG00000019373 | Cops3 | COP9 signalosome subunit 3 [Source:MGI Symbol;Acc:MGI:1349409] | 630,0 | 92,8 |
| ENSMUSG00000024873 | Cnih2 | cornichon family AMPA receptor auxiliary protein 2 [Source:MGI Symbol;Acc:MGI:1277225] | 629,7 | 99,2 |
| ENSMUSG00000003283 | Hck | hemopoietic cell kinase [Source:MGI Symbol;Acc:MGI:96052] | 628,2 | 74,0 |
| ENSMUSG00000031155 | Pim2 | proviral integration site 2 [Source:MGI Symbol;Acc:MGI:97587] | 627,7 | 88,6 |
| ENSMUSG00000024610 | Cd74 | ntigen (invariant polypeptide of major histocompatibility complex, class II antigen-associated) [Source:MGI Symbol;Acc:MG | 627,7 | 85,5 |
| ENSMUSG00000038312 | Edem2 | ER degradation enhancer, mannosidase alpha-like 2 [Source:MGI Symbol;Acc:MGI:1915540] | 627,0 | 81,5 |
| ENSMUSG00000022962 | Gart | phosphoribosylglycinamide formyltransferase [Source:MGI Symbol;Acc:MGI:95654] | 626,5 | 72,5 |
| ENSMUSG00000052833 | Sae1 | SUMO1 activating enzyme subunit 1 [Source:MGI Symbol;Acc:MGI:1929264] | 626,0 | 98,5 |
| ENSMUSG00000041870 | Ankrd13a | ankyrin repeat domain 13a [Source:MGI Symbol;Acc:MGI:1915670] | 625,0 | 81,4 |
| ENSMUSG00000055681 | Cope | coatomer protein complex, subunit epsilon [Source:MGI Symbol;Acc:MGI:1891702] | 622,0 | 83,4 |
| ENSMUSG00000020949 | Fkbp3 | FK506 binding protein 3 [Source:MGI Symbol;Acc:MGI:1353460] | 621,9 | 99,1 |
| ENSMUSG00000001729 | Akt1 | thymoma viral proto-oncogene 1 [Source:MGI Symbol;Acc:MGI:87986] | 621,8 | 77,9 |
| ENSMUSG00000015478 | Rnf5 | ring finger protein 5 [Source:MGI Symbol;Acc:MGI:1860076] | 621,6 | 84,1 |
| ENSMUSG00000035960 | Apex1 | apurinic/apyrimidinic endonuclease 1 [Source:MGI Symbol;Acc:MGI:88042] | 620,9 | 86,3 |
| ENSMUSG00000026509 | Capn2 | calpain 2 [Source:MGI Symbol;Acc:MGI:88264] | 620,2 | 79,4 |
| ENSMUSG00000006299 | Aamp | angio-associated migratory protein [Source:MGI Symbol;Acc:MGI:107809] | 619,4 | 84,9 |
| ENSMUSG00000042380 | Smim12 | small integral membrane protein 12 [Source:MGI Symbol;Acc:MGI:1933141] | 618,9 | 98,8 |
| ENSMUSG00000028330 | Ncbp1 | nuclear cap binding protein subunit 1 [Source:MGI Symbol;Acc:MGI:1891840] | 614,1 | 72,8 |
| ENSMUSG00000061360 | Phf5a | PHD finger protein 5A [Source:MGI Symbol;Acc:MGI:2156864] | 613,9 | 97,3 |
| ENSMUSG00000036693 | Nop14 | NOP14 nucleolar protein [Source:MGI Symbol;Acc:MGI:1922666] | 613,5 | 79,2 |
| ENSMUSG00000020457 | Drg1 | developmentally regulated GTP binding protein 1 [Source:MGI Symbol;Acc:MGI:1343297] | 613,4 | 91,3 |
| ENSMUSG00000020873 | Slc35b1 | solute carrier family 35, member B1 [Source:MGI Symbol;Acc:MGI:1343133] | 612,9 | 86,9 |
| ENSMUSG00000033763 | Mtss1l | metastasis suppressor 1-like [Source:MGI Symbol;Acc:MGI:3039591] | 611,9 | 72,9 |
| ENSMUSG00000032046 | Abhd12 | abhydrolase domain containing 12 [Source:MGI Symbol;Acc:MGI:1923442] | 611,4 | 83,1 |
| ENSMUSG00000020225 | Tmbim4 | transmembrane BAX inhibitor motif containing 4 [Source:MGI Symbol;Acc:MGI:1915462] | 610,5 | 71,5 |
| ENSMUSG00000070570 | Slc17a7 | e carrier family 17 (sodium-dependent inorganic phosphate cotransporter), member 7 [Source:MGI Symbol;Acc:MGI:1920 | 610,1 | 90,8 |
| ENSMUSG00000030307 | Slc6a11 | solute carrier family 6 (neurotransmitter transporter, GABA), member 11 [Source:MGI Symbol;Acc:MGI:95630] | 609,6 | 80,4 |
| ENSMUSG00000033998 | Kcnk1 | potassium channel, subfamily K, member 1 [Source:MGI Symbol;Acc:MGI:109322] | 609,5 | 92,7 |
| ENSMUSG00000022677 | Fopnl | Fgfr1op N-terminal like [Source:MGI Symbol;Acc:MGI:1913336] | 609,1 | 85,2 |
| ENSMUSG00000029062 | Cdk11b | cyclin-dependent kinase 11B [Source:MGI Symbol;Acc:MGI:88353] | 608,8 | 74,1 |
| ENSMUSG00000087260 | Lamtor5 | late endosomal/lysosomal adaptor, MAPK and MTOR activator 5 [Source:MGI Symbol;Acc:MGI:1915826] | 607,9 | 82,1 |
| ENSMUSG00000013822 | Elof1 | ELF1 homolog, elongation factor 1 [Source:MGI Symbol;Acc:MGI:1913376] | 607,6 | 78,9 |
| ENSMUSG00000030310 | Slc6a1 | solute carrier family 6 (neurotransmitter transporter, GABA), member 1 [Source:MGI Symbol;Acc:MGI:95627] | 607,6 | 74,0 |
| ENSMUSG00000026895 | Ndufa8 | NADH dehydrogenase (ubiquinone) 1 alpha subcomplex, 8 [Source:MGI Symbol;Acc:MGI:1915625] | 605,8 | 89,4 |
| ENSMUSG00000031021 | Tmem9b | TMEM9 domain family, member B [Source:MGI Symbol;Acc:MGI:1915254] | 604,9 | 72,3 |
| ENSMUSG00000020250 | Txnrd1 | thioredoxin reductase 1 [Source:MGI Symbol;Acc:MGI:1354175] | 602,9 | 70,0 |
| ENSMUSG00000005510 | Ndufs3 | NADH dehydrogenase (ubiquinone) Fe-S protein 3 [Source:MGI Symbol;Acc:MGI:1915599] | 602,6 | 73,6 |
| ENSMUSG00000027752 | Exosc8 | exosome component 8 [Source:MGI Symbol;Acc:MGI:1916889] | 600,6 | 75,4 |
| ENSMUSG00000001687 | Ubl3 | ubiquitin-like 3 [Source:MGI Symbol;Acc:MGI:1344373] | 600,0 | 87,9 |
| ENSMUSG00000040860 | Crocc | ciliary rootlet coiled-coil, rootletin [Source:MGI Symbol;Acc:MGI:3529431] | 599,5 | 72,7 |
| ENSMUSG00000031007 | Atp6ap2 | ATPase, H+ transporting, lysosomal accessory protein 2 [Source:MGI Symbol;Acc:MGI:1917745] | 598,5 | 76,4 |
| ENSMUSG00000025868 | Higd2a | HIG1 domain family, member 2A [Source:MGI Symbol;Acc:MGI:1914294] | 592,8 | 98,6 |
| ENSMUSG00000054405 | Dnajc8 | DnaJ heat shock protein family (Hsp40) member C8 [Source:MGI Symbol;Acc:MGI:1915848] | 592,3 | 85,8 |
| ENSMUSG00000025157 | Zdhhc16 | zinc finger, DHHC domain containing 16 [Source:MGI Symbol;Acc:MGI:1921418] | 591,9 | 95,8 |
| ENSMUSG00000021178 | Psmc1 | protease (prosome, macropain) 26S subunit, ATPase 1 [Source:MGI Symbol;Acc:MGI:106054] | 590,2 | 95,5 |
| ENSMUSG00000028955 | Vamp3 | vesicle-associated membrane protein 3 [Source:MGI Symbol;Acc:MGI:1321389] | 588,9 | 89,0 |
| ENSMUSG00000026356 | Dars | aspartyl-tRNA synthetase [Source:MGI Symbol;Acc:MGI:2442544] | 588,8 | 82,7 |
| ENSMUSG00000038642 | Ctss | cathepsin S [Source:MGI Symbol;Acc:MGI:107341] | 588,0 | 71,2 |
| ENSMUSG00000025630 | Hprt | hypoxanthine guanine phosphoribosyl transferase [Source:MGI Symbol;Acc:MGI:96217] | 587,5 | 92,8 |
| ENSMUSG00000032288 | Imp3 | IMP3, U3 small nucleolar ribonucleoprotein [Source:MGI Symbol;Acc:MGI:1916119] | 584,8 | 99,0 |
| ENSMUSG00000026426 | Arl8a | ADP-ribosylation factor-like 8A [Source:MGI Symbol;Acc:MGI:1915974] | 583,5 | 91,4 |
| ENSMUSG00000039682 | Lap3 | leucine aminopeptidase 3 [Source:MGI Symbol;Acc:MGI:1914238] | 583,2 | 90,6 |
| ENSMUSG00000042396 | Rbm7 | RNA binding motif protein 7 [Source:MGI Symbol;Acc:MGI:1914260] | 582,3 | 95,6 |
| ENSMUSG00000052889 | Prkcb | protein kinase C, beta [Source:MGI Symbol;Acc:MGI:97596] | 581,2 | 81,9 |
| ENSMUSG00000001313 | Rnd2 | Rho family GTPase 2 [Source:MGI Symbol;Acc:MGI:1338755] | 581,1 | 89,6 |
| ENSMUSG00000062234 | Gak | cyclin G associated kinase [Source:MGI Symbol;Acc:MGI:2442153] | 579,9 | 78,0 |
| ENSMUSG00000025228 | Actr1a | ARP1 actin-related protein 1A, centractin alpha [Source:MGI Symbol;Acc:MGI:1858964] | 579,8 | 86,1 |
| ENSMUSG00000053838 | Nudcd3 | NudC domain containing 3 [Source:MGI Symbol;Acc:MGI:2144158] | 579,7 | 80,9 |
| ENSMUSG00000034445 | Cyb561a3 | cytochrome b561 family, member A3 [Source:MGI Symbol;Acc:MGI:2686925] | 579,0 | 75,4 |
| ENSMUSG00000035278 | Plekhj1 | pleckstrin homology domain containing, family J member 1 [Source:MGI Symbol;Acc:MGI:1925920] | 577,8 | 92,0 |
| ENSMUSG00000002395 | Use1 | unconventional SNARE in the ER 1 homolog (S. cerevisiae) [Source:MGI Symbol;Acc:MGI:1914273] | 576,4 | 73,3 |
| ENSMUSG00000079659 | Tmem243 | transmembrane protein 243, mitochondrial [Source:MGI Symbol;Acc:MGI:3606159] | 576,0 | 83,1 |
| ENSMUSG00000032299 | Commd4 | COMM domain containing 4 [Source:MGI Symbol;Acc:MGI:1913449] | 575,2 | 100,0 |
| ENSMUSG00000021241 | Isca2 | iron-sulfur cluster assembly 2 [Source:MGI Symbol;Acc:MGI:1921566] | 574,9 | 97,5 |
| ENSMUSG00000026203 | Dnajb2 | DnaJ heat shock protein family (Hsp40) member B2 [Source:MGI Symbol;Acc:MGI:1928739] | 572,2 | 71,7 |
| ENSMUSG00000027404 | Snrpb | small nuclear ribonucleoprotein B [Source:MGI Symbol;Acc:MGI:98342] | 572,2 | 93,3 |
| ENSMUSG00000003363 | Pld3 | phospholipase D family, member 3 [Source:MGI Symbol;Acc:MGI:1333782] | 571,4 | 77,0 |
| ENSMUSG00000063787 | Chchd1 | coiled-coil-helix-coiled-coil-helix domain containing 1 [Source:MGI Symbol;Acc:MGI:1913371] | 571,3 | 95,5 |
| ENSMUSG00000053931 | Cnn3 | calponin 3, acidic [Source:MGI Symbol;Acc:MGI:1919244] | 569,3 | 82,0 |
| ENSMUSG00000031729 | Ist1 | increased sodium tolerance 1 homolog (yeast) [Source:MGI Symbol;Acc:MGI:1919205] | 569,1 | 86,9 |
| ENSMUSG00000026548 | Slamf9 | SLAM family member 9 [Source:MGI Symbol;Acc:MGI:1923692] | 567,5 | 91,4 |
| ENSMUSG00000007872 | Id3 | inhibitor of DNA binding 3 [Source:MGI Symbol;Acc:MGI:96398] | 567,3 | 75,2 |
| ENSMUSG00000078656 | Vps25 | vacuolar protein sorting 25 (yeast) [Source:MGI Symbol;Acc:MGI:106354] | 566,3 | 74,8 |
| ENSMUSG00000018585 | Atox1 | antioxidant 1 copper chaperone [Source:MGI Symbol;Acc:MGI:1333855] | 566,2 | 89,2 |
| ENSMUSG00000034341 | Wbp2 | WW domain binding protein 2 [Source:MGI Symbol;Acc:MGI:104709] | 565,2 | 93,3 |
| ENSMUSG00000020132 | Rab21 | RAB21, member RAS oncogene family [Source:MGI Symbol;Acc:MGI:894308] | 563,2 | 87,6 |
| ENSMUSG00000068329 | Htra2 | HtrA serine peptidase 2 [Source:MGI Symbol;Acc:MGI:1928676] | 563,0 | 72,4 |
| ENSMUSG00000037257 | Aagab | alpha- and gamma-adaptin binding protein [Source:MGI Symbol;Acc:MGI:1914189] | 560,9 | 72,3 |
| ENSMUSG00000038722 | Bud31 | BUD31 homolog [Source:MGI Symbol;Acc:MGI:2141291] | 559,9 | 95,2 |
| ENSMUSG00000059734 | Ndufs8 | NADH dehydrogenase (ubiquinone) Fe-S protein 8 [Source:MGI Symbol;Acc:MGI:2385079] | 559,7 | 83,0 |
| ENSMUSG00000039048 | Foxred1 | FAD-dependent oxidoreductase domain containing 1 [Source:MGI Symbol;Acc:MGI:2446262] | 559,7 | 86,7 |
| ENSMUSG00000028932 | Psmc2 | proteasome (prosome, macropain) 26S subunit, ATPase 2 [Source:MGI Symbol;Acc:MGI:109555] | 557,8 | 79,8 |
| ENSMUSG00000046841 | Ckap4 | cytoskeleton-associated protein 4 [Source:MGI Symbol;Acc:MGI:2444926] | 557,5 | 81,0 |
| ENSMUSG00000042729 | Wdr74 | WD repeat domain 74 [Source:MGI Symbol;Acc:MGI:2147427] | 557,3 | 95,2 |
| ENSMUSG00000022204 | Ngdn | neuroguidin, EIF4E binding protein [Source:MGI Symbol;Acc:MGI:1916216] | 556,9 | 88,3 |
| ENSMUSG00000015095 | Fbxw5 | F-box and WD-40 domain protein 5 [Source:MGI Symbol;Acc:MGI:1354731] | 556,1 | 88,1 |
| ENSMUSG00000090841 | Myl6 | myosin, light polypeptide 6, alkali, smooth muscle and non-muscle [Source:MGI Symbol;Acc:MGI:109318] | 555,4 | 76,4 |
| ENSMUSG00000020766 | Galk1 | galactokinase 1 [Source:MGI Symbol;Acc:MGI:95730] | 555,3 | 95,1 |
| ENSMUSG00000019210 | Atp6v1e1 | ATPase, H+ transporting, lysosomal V1 subunit E1 [Source:MGI Symbol;Acc:MGI:894326] | 554,1 | 77,5 |
| ENSMUSG00000041632 | Mrps27 | mitochondrial ribosomal protein S27 [Source:MGI Symbol;Acc:MGI:1919064] | 554,0 | 95,6 |
| ENSMUSG00000022404 | Slc25a17 | e carrier family 25 (mitochondrial carrier, peroxisomal membrane protein), member 17 [Source:MGI Symbol;Acc:MGI:1342 | 551,8 | 99,0 |
| ENSMUSG00000048307 | Ankrd46 | ankyrin repeat domain 46 [Source:MGI Symbol;Acc:MGI:1916089] | 550,1 | 88,5 |
| ENSMUSG00000033020 | Polr2f | polymerase (RNA) II (DNA directed) polypeptide F [Source:MGI Symbol;Acc:MGI:1349393] | 549,6 | 99,0 |
| ENSMUSG00000024870 | Rab1b | RAB1B, member RAS oncogene family [Source:MGI Symbol;Acc:MGI:1923558] | 548,7 | 94,2 |
| ENSMUSG00000029577 | Ube3b | ubiquitin protein ligase E3B [Source:MGI Symbol;Acc:MGI:1891295] | 548,3 | 73,1 |
| ENSMUSG00000024841 | Eif1ad | eukaryotic translation initiation factor 1A domain containing [Source:MGI Symbol;Acc:MGI:1917110] | 547,9 | 75,5 |
| ENSMUSG00000016427 | Ndufa1 | NADH dehydrogenase (ubiquinone) 1 alpha subcomplex, 1 [Source:MGI Symbol;Acc:MGI:1929511] | 546,8 | 96,4 |
| ENSMUSG00000022536 | Glyr1 | glyoxylate reductase 1 homolog (Arabidopsis) [Source:MGI Symbol;Acc:MGI:1921272] | 546,6 | 85,4 |
| ENSMUSG00000062937 | Mtap | methylthioadenosine phosphorylase [Source:MGI Symbol;Acc:MGI:1914152] | 546,5 | 85,8 |
| ENSMUSG00000001774 | Chordc1 | cysteine and histidine-rich domain (CHORD)-containing, zinc-binding protein 1 [Source:MGI Symbol;Acc:MGI:1914167] | 544,4 | 75,8 |

| | | | | |
|---|---|---|---|---|
| ENSMUSG00000075528 | Aarsd1 | alanyl-tRNA synthetase domain containing 1 [Source:MGI Symbol;Acc:MGI:1916934] | 543,5 | 80,8 |
| ENSMUSG00000032305 | Fam219b | family with sequence similarity 219, member B [Source:MGI Symbol;Acc:MGI:1925573] | 542,9 | 73,5 |
| ENSMUSG00000021608 | Lpcat1 | lysophosphatidylcholine acyltransferase 1 [Source:MGI Symbol;Acc:MGI:2384812] | 541,9 | 71,0 |
| ENSMUSG00000027406 | Idh3b | isocitrate dehydrogenase 3 (NAD+) beta [Source:MGI Symbol;Acc:MGI:2158650] | 541,8 | 92,0 |
| ENSMUSG00000007564 | Ppp2r1a | protein phosphatase 2, regulatory subunit A, alpha [Source:MGI Symbol;Acc:MGI:1926334] | 541,7 | 90,4 |
| ENSMUSG00000041840 | Haus1 | HAUS augmin-like complex, subunit 1 [Source:MGI Symbol;Acc:MGI:2385076] | 541,5 | 96,5 |
| ENSMUSG00000021731 | Mrps30 | mitochondrial ribosomal protein S30 [Source:MGI Symbol;Acc:MGI:1926237] | 539,2 | 88,1 |
| ENSMUSG00000021476 | Habp4 | hyaluronic acid binding protein 4 [Source:MGI Symbol;Acc:MGI:1891713] | 538,6 | 73,8 |
| ENSMUSG00000032038 | St3gal4 | ST3 beta-galactoside alpha-2,3-sialyltransferase 4 [Source:MGI Symbol;Acc:MGI:1316743] | 538,3 | 91,4 |
| ENSMUSG00000027714 | Exosc9 | exosome component 9 [Source:MGI Symbol;Acc:MGI:1355319] | 537,4 | 82,8 |
| ENSMUSG00000049299 | Trappc1 | trafficking protein particle complex 1 [Source:MGI Symbol;Acc:MGI:1098727] | 536,7 | 89,5 |
| ENSMUSG00000040659 | Efhd2 | EF hand domain containing 2 [Source:MGI Symbol;Acc:MGI:106504] | 535,9 | 86,3 |
| ENSMUSG00000036737 | Oxsr1 | oxidative-stress responsive 1 [Source:MGI Symbol;Acc:MGI:1917378] | 535,0 | 74,6 |
| ENSMUSG00000074227 | Spint2 | serine protease inhibitor, Kunitz type 2 [Source:MGI Symbol;Acc:MGI:1338031] | 534,2 | 86,1 |
| ENSMUSG00000051650 | B3gnt2 | UDP-GlcNAc:betaGal beta-1,3-N-acetylglucosaminyltransferase 2 [Source:MGI Symbol;Acc:MGI:1889505] | 534,1 | 73,9 |
| ENSMUSG00000052296 | Ppp6r1 | protein phosphatase 6, regulatory subunit 1 [Source:MGI Symbol;Acc:MGI:2442163] | 533,5 | 83,8 |
| ENSMUSG00000027502 | Rtfdc1 | replication termination factor 2 domain containing 1 [Source:MGI Symbol;Acc:MGI:1913654] | 533,3 | 92,4 |
| ENSMUSG00000049960 | Mrps16 | mitochondrial ribosomal protein S16 [Source:MGI Symbol;Acc:MGI:1913492] | 533,3 | 99,2 |
| ENSMUSG00000031954 | Cfdp1 | craniofacial development protein 1 [Source:MGI Symbol;Acc:MGI:1344403] | 531,6 | 81,3 |
| ENSMUSG00000032526 | Deb1 | differentially expressed in B16F10 1 [Source:MGI Symbol;Acc:MGI:1349474] | 531,6 | 99,9 |
| ENSMUSG00000047067 | Dusp28 | dual specificity phosphatase 28 [Source:MGI Symbol;Acc:MGI:1914696] | 531,6 | 95,2 |
| ENSMUSG00000087687 | Pet100 | PET100 homolog (S. cerevisiae) [Source:MGI Symbol;Acc:MGI:3615306] | 530,8 | 70,4 |
| ENSMUSG00000028013 | Ppa2 | pyrophosphatase (inorganic) 2 [Source:MGI Symbol;Acc:MGI:1922026] | 530,0 | 81,9 |
| ENSMUSG00000010755 | Cars | cysteinyl-tRNA synthetase [Source:MGI Symbol;Acc:MGI:1351477] | 528,9 | 70,4 |
| ENSMUSG00000072694 | 1500011B03Rik | RIKEN cDNA 1500011B03 gene [Source:MGI Symbol;Acc:MGI:1913486] | 528,3 | 87,6 |
| ENSMUSG00000076604 | Igkj1 | immunoglobulin kappa joining 1 [Source:MGI Symbol;Acc:MGI:1316689] | 528,1 | 100,0 |
| ENSMUSG00000022757 | Tfg | Trk-fused gene [Source:MGI Symbol;Acc:MGI:1338041] | 527,8 | 77,1 |
| ENSMUSG00000032126 | Hmbs | hydroxymethylbilane synthase [Source:MGI Symbol;Acc:MGI:96112] | 527,6 | 74,9 |
| ENSMUSG00000039660 | D2Wsu81e | DNA segment, Chr 2, Wayne State University 81, expressed [Source:MGI Symbol;Acc:MGI:106544] | 527,5 | 96,8 |
| ENSMUSG00000024982 | Zdhhc6 | zinc finger, DHHC domain containing 6 [Source:MGI Symbol;Acc:MGI:1914230] | 527,2 | 75,1 |
| ENSMUSG00000050846 | Zfp623 | zinc finger protein 623 [Source:MGI Symbol;Acc:MGI:1926084] | 526,7 | 92,4 |
| ENSMUSG00000000339 | Rtca | RNA 3'-terminal phosphate cyclase [Source:MGI Symbol;Acc:MGI:1913618] | 526,7 | 87,7 |
| ENSMUSG00000022681 | Ntan1 | N-terminal Asn amidase [Source:MGI Symbol;Acc:MGI:108471] | 526,7 | 78,8 |
| ENSMUSG00000018559 | Ctdnep1 | CTD nuclear envelope phosphatase 1 [Source:MGI Symbol;Acc:MGI:1914431] | 525,9 | 84,9 |
| ENSMUSG00000008035 | Mid1ip1 | Mid1 interacting protein 1 (gastrulation specific G12-like (zebrafish)) [Source:MGI Symbol;Acc:MGI:1915291] | 525,3 | 92,7 |
| ENSMUSG00000030615 | Tmem126a | transmembrane protein 126A [Source:MGI Symbol;Acc:MGI:1913521] | 524,0 | 78,5 |
| ENSMUSG00000025035 | Arl3 | ADP-ribosylation factor-like 3 [Source:MGI Symbol;Acc:MGI:1929699] | 522,5 | 96,6 |
| ENSMUSG00000071547 | Nt5dc2 | 5'-nucleotidase domain containing 2 [Source:MGI Symbol;Acc:MGI:1917271] | 521,4 | 95,4 |
| ENSMUSG00000070003 | Ssbp4 | single stranded DNA binding protein 4 [Source:MGI Symbol;Acc:MGI:1924150] | 520,9 | 94,8 |
| ENSMUSG00000022200 | Golph3 | golgi phosphoprotein 3 [Source:MGI Symbol;Acc:MGI:1913879] | 520,6 | 89,8 |
| ENSMUSG00000020780 | Srp68 | signal recognition particle 68 [Source:MGI Symbol;Acc:MGI:1917447] | 517,9 | 79,4 |
| ENSMUSG00000021024 | Psma6 | proteasome (prosome, macropain) subunit, alpha type 6 [Source:MGI Symbol;Acc:MGI:1347006] | 517,8 | 84,0 |
| ENSMUSG00000006575 | Rundc3a | RUN domain containing 3A [Source:MGI Symbol;Acc:MGI:1858752] | 517,2 | 80,2 |
| ENSMUSG00000060679 | Mrps9 | mitochondrial ribosomal protein S9 [Source:MGI Symbol;Acc:MGI:1916777] | 516,4 | 94,5 |
| ENSMUSG00000028394 | Pole3 | polymerase (DNA directed), epsilon 3 (p17 subunit) [Source:MGI Symbol;Acc:MGI:1933378] | 514,5 | 91,4 |
| ENSMUSG00000024911 | Fibp | fibroblast growth factor (acidic) intracellular binding protein [Source:MGI Symbol;Acc:MGI:1926233] | 513,9 | 95,9 |
| ENSMUSG00000061111 | Fam195b | family with sequence similarity 195, member B [Source:MGI Symbol;Acc:MGI:2384752] | 513,3 | 96,5 |
| ENSMUSG00000068290 | Ddrgk1 | DDRGK domain containing 1 [Source:MGI Symbol;Acc:MGI:1924256] | 512,3 | 84,6 |
| ENSMUSG00000020089 | Ppa1 | pyrophosphatase (inorganic) 1 [Source:MGI Symbol;Acc:MGI:97831] | 510,5 | 83,0 |
| ENSMUSG00000051934 | Spats2 | spermatogenesis associated, serine-rich 2 [Source:MGI Symbol;Acc:MGI:1919822] | 510,3 | 73,3 |
| ENSMUSG00000031983 | 2310022B05Rik | RIKEN cDNA 2310022B05 gene [Source:MGI Symbol;Acc:MGI:1916801] | 509,4 | 87,9 |
| ENSMUSG00000002043 | Trappc6a | trafficking protein particle complex 6A [Source:MGI Symbol;Acc:MGI:1914341] | 509,2 | 98,5 |
| ENSMUSG00000021114 | Atp6v1d | ATPase, H+ transporting, lysosomal V1 subunit D [Source:MGI Symbol;Acc:MGI:1921084] | 509,2 | 91,4 |
| ENSMUSG00000058833 | 2810428I15Rik | RIKEN cDNA 2810428I15 gene [Source:MGI Symbol;Acc:MGI:1913712] | 508,4 | 92,9 |
| ENSMUSG00000002660 | Clpp | caseinolytic mitochondrial matrix peptidase proteolytic subunit [Source:MGI Symbol;Acc:MGI:1858213] | 508,4 | 78,9 |
| ENSMUSG00000005881 | Ergic3 | ERGIC and golgi 3 [Source:MGI Symbol;Acc:MGI:1913616] | 508,1 | 76,7 |
| ENSMUSG00000057649 | Brd9 | bromodomain containing 9 [Source:MGI Symbol;Acc:MGI:2145317] | 506,8 | 82,2 |
| ENSMUSG00000022951 | Rcan1 | regulator of calcineurin 1 [Source:MGI Symbol;Acc:MGI:1890564] | 505,2 | 84,9 |
| ENSMUSG00000020392 | Cdkn2aipnl | CDKN2A interacting protein N-terminal like [Source:MGI Symbol;Acc:MGI:1261797] | 505,2 | 78,7 |
| ENSMUSG00000028741 | Mrto4 | mRNA turnover 4, homolog (S. cerevisiae) [Source:MGI Symbol;Acc:MGI:1917152] | 504,0 | 77,9 |
| ENSMUSG00000025066 | Sfr1 | SWI5 dependent recombination repair 1 [Source:MGI Symbol;Acc:MGI:1915038] | 503,6 | 78,7 |
| ENSMUSG00000036023 | Parp2 | poly (ADP-ribose) polymerase family, member 2 [Source:MGI Symbol;Acc:MGI:1341112] | 502,2 | 98,4 |
| ENSMUSG00000041203 | 2310036O22Rik | RIKEN cDNA 2310036O22 gene [Source:MGI Symbol;Acc:MGI:1922833] | 501,5 | 96,7 |
| ENSMUSG00000041629 | Fam104a | family with sequence similarity 104, member A [Source:MGI Symbol;Acc:MGI:106351] | 501,1 | 73,2 |
| ENSMUSG00000009741 | Ubp1 | upstream binding protein 1 [Source:MGI Symbol;Acc:MGI:104889] | 500,9 | 79,8 |
| ENSMUSG00000014606 | Slc25a11 | solute carrier family 25 (mitochondrial carrier oxoglutarate carrier), member 11 [Source:MGI Symbol;Acc:MGI:1915113] | 500,7 | 81,5 |

Table S2.3

| Id | Associated Gene Name | Description | Mean read Astrocyte | Mean Astrocyte Coverage |
|---|---|---|---|---|
| ENSMUSG00000027447 | Cst3 | cystatin C [Source:MGI Symbol;Acc:MGI:102519] | 86625,9 | 91,68 |
| ENSMUSG00000037852 | Cpe | carboxypeptidase E [Source:MGI Symbol;Acc:MGI:101932] | 77322,8 | 100,00 |
| ENSMUSG00000031765 | Mt1 | metallothionein 1 [Source:MGI Symbol;Acc:MGI:97171] | 62434,7 | 100,00 |
| ENSMUSG00000061808 | Ttr | transthyretin [Source:MGI Symbol;Acc:MGI:98865] | 53183,3 | 100,00 |
| ENSMUSG00000002985 | Apoe | apolipoprotein E [Source:MGI Symbol;Acc:MGI:88057] | 52568,4 | 89,91 |
| ENSMUSG00000020932 | Gfap | glial fibrillary acidic protein [Source:MGI Symbol;Acc:MGI:95697] | 51709,3 | 99,40 |
| ENSMUSG00000017390 | Aldoc | aldolase C, fructose-bisphosphate [Source:MGI Symbol;Acc:MGI:101863] | 45498,3 | 100,00 |
| ENSMUSG00000026701 | Prdx6 | peroxiredoxin 6 [Source:MGI Symbol;Acc:MGI:894320] | 44118,1 | 99,72 |
| ENSMUSG00000004558 | Ndrg2 | N-myc downstream regulated gene 2 [Source:MGI Symbol;Acc:MGI:1352498] | 37500,0 | 100,00 |
| ENSMUSG00000026385 | Dbi | diazepam binding inhibitor [Source:MGI Symbol;Acc:MGI:94865] | 31118,5 | 98,12 |
| ENSMUSG00000022037 | Clu | clusterin [Source:MGI Symbol;Acc:MGI:88423] | 29982,3 | 96,67 |
| ENSMUSG00000058135 | Gstm1 | glutathione S-transferase, mu 1 [Source:MGI Symbol;Acc:MGI:95860] | 27393,6 | 100,00 |
| ENSMUSG00000001270 | Ckb | creatine kinase, brain [Source:MGI Symbol;Acc:MGI:88407] | 25914,7 | 98,92 |
| ENSMUSG00000004032 | Gstm5 | glutathione S-transferase, mu 5 [Source:MGI Symbol;Acc:MGI:1309466] | 22563,9 | 95,99 |
| ENSMUSG00000031760 | Mt3 | metallothionein 3 [Source:MGI Symbol;Acc:MGI:97173] | 21793,1 | 100,00 |
| ENSMUSG00000031342 | Gpm6b | glycoprotein m6b [Source:MGI Symbol;Acc:MGI:107672] | 19934,3 | 99,91 |
| ENSMUSG00000005360 | Slc1a3 | ;olute carrier family 1 (glial high affinity glutamate transporter), member 3 [Source:MGI Symbol;Acc:MGI:99917 | 19652,7 | 96,68 |
| ENSMUSG00000029309 | Sparcl1 | SPARC-like 1 [Source:MGI Symbol;Acc:MGI:108110] | 16389,2 | 100,00 |
| ENSMUSG00000001175 | Calm1 | calmodulin 1 [Source:MGI Symbol;Acc:MGI:88251] | 15676,0 | 95,53 |
| ENSMUSG00000026421 | Csrp1 | cysteine and glycine-rich protein 1 [Source:MGI Symbol;Acc:MGI:88549] | 15542,5 | 96,06 |
| ENSMUSG00000031517 | Gpm6a | glycoprotein m6a [Source:MGI Symbol;Acc:MGI:107671] | 15521,9 | 99,47 |
| ENSMUSG00000054226 | Tprkb | Tp53rk binding protein [Source:MGI Symbol;Acc:MGI:1917036] | 15367,4 | 99,04 |
| ENSMUSG00000053279 | Aldh1a1 | aldehyde dehydrogenase family 1, subfamily A1 [Source:MGI Symbol;Acc:MGI:1353450] | 14626,8 | 99,01 |
| ENSMUSG00000005610 | Eif4g2 | eukaryotic translation initiation factor 4, gamma 2 [Source:MGI Symbol;Acc:MGI:109207] | 14563,6 | 87,19 |
| ENSMUSG00000031762 | Mt2 | metallothionein 2 [Source:MGI Symbol;Acc:MGI:97172] | 14105,0 | 100,00 |
| ENSMUSG00000059325 | Hopx | HOP homeobox [Source:MGI Symbol;Acc:MGI:1916782] | 13770,0 | 100,00 |
| ENSMUSG00000035202 | Lars2 | leucyl-tRNA synthetase, mitochondrial [Source:MGI Symbol;Acc:MGI:2142973] | 13397,8 | 99,30 |
| ENSMUSG00000025203 | Scd2 | stearoyl-Coenzyme A desaturase 2 [Source:MGI Symbol;Acc:MGI:98240] | 13344,1 | 95,31 |
| ENSMUSG00000027893 | Ahcyl1 | S-adenosylhomocysteine hydrolase-like 1 [Source:MGI Symbol;Acc:MGI:2385184] | 13181,8 | 100,00 |
| ENSMUSG00000020644 | Id2 | inhibitor of DNA binding 2 [Source:MGI Symbol;Acc:MGI:96397] | 13113,0 | 100,00 |
| ENSMUSG00000039533 | Mmd2 | monocyte to macrophage differentiation-associated 2 [Source:MGI Symbol;Acc:MGI:1922354] | 12560,3 | 99,95 |
| ENSMUSG00000005089 | Slc1a2 | ɔlute carrier family 1 (glial high affinity glutamate transporter), member 2 [Source:MGI Symbol;Acc:MGI:10193 | 12099,8 | 99,20 |
| ENSMUSG00000022425 | Enpp2 | ectonucleotide pyrophosphatase/phosphodiesterase 2 [Source:MGI Symbol;Acc:MGI:1321390] | 12078,0 | 99,90 |
| ENSMUSG00000090625 | Gm20721 | predicted gene, 20721 [Source:MGI Symbol;Acc:MGI:5313030] | 11999,4 | 99,33 |
| ENSMUSG00000021792 | Fam213a | family with sequence similarity 213, member A [Source:MGI Symbol;Acc:MGI:1917814] | 11709,2 | 92,35 |
| ENSMUSG00000022108 | Itm2b | integral membrane protein 2B [Source:MGI Symbol;Acc:MGI:1309517] | 11491,4 | 100,00 |
| ENSMUSG00000013698 | Pea15a | phosphoprotein enriched in astrocytes 15A [Source:MGI Symbol;Acc:MGI:104799] | 11374,1 | 97,95 |
| ENSMUSG00000007097 | Atp1a2 | ATPase, Na+/K+ transporting, alpha 2 polypeptide [Source:MGI Symbol;Acc:MGI:88106] | 11364,3 | 98,03 |
| ENSMUSG00000030428 | Ttyh1 | tweety homolog 1 (Drosophila) [Source:MGI Symbol;Acc:MGI:1889007] | 11354,2 | 99,16 |
| ENSMUSG00000033208 | S100b | S100 protein, beta polypeptide, neural [Source:MGI Symbol;Acc:MGI:98217] | 11166,7 | 100,00 |
| ENSMUSG00000021750 | Fam107a | family with sequence similarity 107, member A [Source:MGI Symbol;Acc:MGI:3041256] | 11038,8 | 91,94 |
| ENSMUSG00000045092 | S1pr1 | sphingosine-1-phosphate receptor 1 [Source:MGI Symbol;Acc:MGI:1096355] | 10766,9 | 99,84 |
| ENSMUSG00000030246 | Ldhb | lactate dehydrogenase B [Source:MGI Symbol;Acc:MGI:96763] | 10219,7 | 89,81 |
| ENSMUSG00000000346 | Dazap2 | DAZ associated protein 2 [Source:MGI Symbol;Acc:MGI:1344344] | 10160,4 | 100,00 |
| ENSMUSG00000026424 | Gpr37l1 | G protein-coupled receptor 37-like 1 [Source:MGI Symbol;Acc:MGI:1928503] | 9669,7 | 99,00 |
| ENSMUSG00000037706 | Cd81 | CD81 antigen [Source:MGI Symbol;Acc:MGI:1096398] | 9604,2 | 95,12 |
| ENSMUSG00000023913 | Pla2g7 | ɔholipase A2, group VII (platelet-activating factor acetylhydrolase, plasma) [Source:MGI Symbol;Acc:MGI:135 | 9406,0 | 99,84 |
| ENSMUSG00000020849 | Ywhae | ɔnooxygenase/tryptophan 5-monooxygenase activation protein, epsilon polypeptide [Source:MGI Symbol;Acc: | 9375,3 | 99,01 |
| ENSMUSG00000022323 | Hrsp12 | heat-responsive protein 12 [Source:MGI Symbol;Acc:MGI:1095401] | 9257,7 | 99,87 |
| ENSMUSG00000030695 | Aldoa | aldolase A, fructose-bisphosphate [Source:MGI Symbol;Acc:MGI:87994] | 9155,1 | 78,40 |
| ENSMUSG00000064373 | Sepp1 | selenoprotein P, plasma, 1 [Source:MGI Symbol;Acc:MGI:894288] | 8993,2 | 89,64 |
| ENSMUSG00000028691 | Prdx1 | peroxiredoxin 1 [Source:MGI Symbol;Acc:MGI:99523] | 8949,8 | 94,53 |
| ENSMUSG00000055254 | Ntrk2 | neurotrophic tyrosine kinase, receptor, type 2 [Source:MGI Symbol;Acc:MGI:97384] | 8897,4 | 72,44 |
| ENSMUSG00000024654 | Asrgl1 | asparaginase like 1 [Source:MGI Symbol;Acc:MGI:1913764] | 8843,8 | 99,91 |
| ENSMUSG00000024661 | Fth1 | ferritin heavy polypeptide 1 [Source:MGI Symbol;Acc:MGI:95588] | 8833,9 | 100,00 |
| ENSMUSG00000020591 | Ntsr2 | neurotensin receptor 2 [Source:MGI Symbol;Acc:MGI:108018] | 8823,2 | 99,87 |
| ENSMUSG00000026473 | Glul | glutamate-ammonia ligase (glutamine synthetase) [Source:MGI Symbol;Acc:MGI:95739] | 8817,2 | 99,68 |
| ENSMUSG00000026249 | Serpine2 | serine (or cysteine) peptidase inhibitor, clade E, member 2 [Source:MGI Symbol;Acc:MGI:101780] | 8748,6 | 95,51 |
| ENSMUSG00000025393 | Atp5b | ATP synthase, H+ transporting mitochondrial F1 complex, beta subunit [Source:MGI Symbol;Acc:MGI:107801 | 8729,7 | 99,39 |
| ENSMUSG00000050953 | Gja1 | gap junction protein, alpha 1 [Source:MGI Symbol;Acc:MGI:95713] | 8700,2 | 99,99 |
| ENSMUSG00000030701 | Plekhb1 | ɔleckstrin homology domain containing, family B (evectins) member 1 [Source:MGI Symbol;Acc:MGI:1351469 | 8601,9 | 76,00 |
| ENSMUSG00000036309 | Skp1a | S-phase kinase-associated protein 1A [Source:MGI Symbol;Acc:MGI:103575] | 8601,9 | 93,96 |
| ENSMUSG00000031633 | Slc25a4 | ier family 25 (mitochondrial carrier, adenine nucleotide translocator), member 4 [Source:MGI Symbol;Acc:MGI | 8533,9 | 88,76 |
| ENSMUSG00000035686 | Thrsp | thyroid hormone responsive [Source:MGI Symbol;Acc:MGI:109126] | 8458,4 | 94,13 |
| ENSMUSG00000025666 | Tmem47 | transmembrane protein 47 [Source:MGI Symbol;Acc:MGI:2177570] | 8454,8 | 100,00 |
| ENSMUSG00000028517 | Plpp3 | phospholipid phosphatase 3 [Source:MGI Symbol;Acc:MGI:1915166] | 8309,9 | 100,00 |
| ENSMUSG00000031431 | Tsc22d3 | TSC22 domain family, member 3 [Source:MGI Symbol;Acc:MGI:1196284] | 8167,4 | 90,35 |
| ENSMUSG00000079037 | Prnp | prion protein [Source:MGI Symbol;Acc:MGI:97769] | 8124,3 | 99,69 |
| ENSMUSG00000035885 | Cox8a | cytochrome c oxidase subunit VIIIa [Source:MGI Symbol;Acc:MGI:105959] | 8039,9 | 98,72 |
| ENSMUSG00000030120 | Mlf2 | myeloid leukemia factor 2 [Source:MGI Symbol;Acc:MGI:1353554] | 7983,9 | 98,76 |
| ENSMUSG00000021379 | Id4 | inhibitor of DNA binding 4 [Source:MGI Symbol;Acc:MGI:99414] | 7929,5 | 100,00 |
| ENSMUSG00000025871 | 4833439L19Rik | RIKEN cDNA 4833439L19 gene [Source:MGI Symbol;Acc:MGI:1921162] | 7817,7 | 98,96 |
| ENSMUSG00000027562 | Car2 | carbonic anhydrase 2 [Source:MGI Symbol;Acc:MGI:88269] | 7637,9 | 100,00 |
| ENSMUSG00000018770 | Atp5g3 | ynthase, H+ transporting, mitochondrial F0 complex, subunit C3 (subunit 9) [Source:MGI Symbol;Acc:MGI:24 | 7550,4 | 100,00 |
| ENSMUSG00000040997 | Abhd4 | abhydrolase domain containing 4 [Source:MGI Symbol;Acc:MGI:1915938] | 7373,1 | 92,71 |
| ENSMUSG00000044080 | S100a1 | S100 calcium binding protein A1 [Source:MGI Symbol;Acc:MGI:1338917] | 7245,7 | 100,00 |
| ENSMUSG00000024782 | Ak3 | adenylate kinase 3 [Source:MGI Symbol;Acc:MGI:1860835] | 7240,6 | 100,00 |
| ENSMUSG00000062078 | Qk | quaking [Source:MGI Symbol;Acc:MGI:97837] | 7196,7 | 99,50 |
| ENSMUSG00000031818 | Cox4i1 | cytochrome c oxidase subunit IV isoform 1 [Source:MGI Symbol;Acc:MGI:88473] | 7165,2 | 100,00 |
| ENSMUSG00000032281 | Acsbg1 | acyl-CoA synthetase bubblegum family member 1 [Source:MGI Symbol;Acc:MGI:2385656] | 7056,1 | 84,38 |
| ENSMUSG00000032324 | Tspan3 | tetraspanin 3 [Source:MGI Symbol;Acc:MGI:1928098] | 7052,1 | 100,00 |
| ENSMUSG00000022890 | Atp5j | ATP synthase, H+ transporting, mitochondrial F0 complex, subunit F [Source:MGI Symbol;Acc:MGI:107777] | 7019,3 | 100,00 |
| ENSMUSG00000010663 | Fads1 | fatty acid desaturase 1 [Source:MGI Symbol;Acc:MGI:1923517] | 6867,2 | 99,84 |
| ENSMUSG00000037072 | Sep15 | selenoprotein 15 [Source:MGI Symbol;Acc:MGI:1927947] | 6865,4 | 78,21 |
| ENSMUSG00000030057 | Cnbp | cellular nucleic acid binding protein [Source:MGI Symbol;Acc:MGI:88431] | 6781,9 | 96,35 |
| ENSMUSG00000041329 | Atp1b2 | ATPase, Na+/K+ transporting, beta 2 polypeptide [Source:MGI Symbol;Acc:MGI:88109] | 6734,9 | 99,13 |
| ENSMUSG00000015733 | Capza2 | capping protein (actin filament) muscle Z-line, alpha 2 [Source:MGI Symbol;Acc:MGI:106222] | 6478,9 | 99,24 |
| ENSMUSG00000001833 | Sept7 | septin 7 [Source:MGI Symbol;Acc:MGI:1335094] | 6315,9 | 98,20 |
| ENSMUSG00000035805 | Mlc1 | encephalic leukoencephalopathy with subcortical cysts 1 homolog (human) [Source:MGI Symbol;Acc:MGI:21 | 6156,6 | 99,52 |
| ENSMUSG00000056201 | Cfl1 | cofilin 1, non-muscle [Source:MGI Symbol;Acc:MGI:101757] | 6077,7 | 100,00 |
| ENSMUSG00000004892 | Bcan | brevican [Source:MGI Symbol;Acc:MGI:1096385] | 6071,2 | 99,12 |
| ENSMUSG00000001847 | Rac1 | RAS-related C3 botulinum substrate 1 [Source:MGI Symbol;Acc:MGI:97845] | 6064,7 | 76,23 |
| ENSMUSG00000031447 | Lamp1 | lysosomal-associated membrane protein 1 [Source:MGI Symbol;Acc:MGI:96745] | 5978,9 | 99,79 |
| ENSMUSG00000041028 | Ghitm | growth hormone inducible transmembrane protein [Source:MGI Symbol;Acc:MGI:1913342] | 5937,9 | 100,00 |
| ENSMUSG00000049422 | Chchd10 | coiled-coil-helix-coiled-coil-helix domain containing 10 [Source:MGI Symbol;Acc:MGI:2143558] | 5927,8 | 85,59 |
| ENSMUSG00000074457 | S100a16 | S100 calcium binding protein A16 [Source:MGI Symbol;Acc:MGI:1915110] | 5645,4 | 98,95 |
| ENSMUSG00000001891 | Ugp2 | UDP-glucose pyrophosphorylase 2 [Source:MGI Symbol;Acc:MGI:2183447] | 5626,4 | 100,00 |
| ENSMUSG00000032883 | Acsl3 | acyl-CoA synthetase long-chain family member 3 [Source:MGI Symbol;Acc:MGI:1921455] | 5619,9 | 99,72 |
| ENSMUSG00000006205 | Htra1 | HtrA serine peptidase 1 [Source:MGI Symbol;Acc:MGI:1929076] | 5560,3 | 100,00 |
| ENSMUSG00000025781 | Atp5c1 | ynthase, H+ transporting, mitochondrial F1 complex, gamma polypeptide 1 [Source:MGI Symbol;Acc:MGI:12 | 5556,6 | 90,53 |
| ENSMUSG00000049775 | Tmsb4x | thymosin, beta 4, X chromosome [Source:MGI Symbol;Acc:MGI:99510] | 5480,7 | 76,64 |
| ENSMUSG00000028756 | Pink1 | PTEN induced putative kinase 1 [Source:MGI Symbol;Acc:MGI:1916193] | 5438,2 | 91,12 |
| ENSMUSG00000019804 | Snx3 | sorting nexin 3 [Source:MGI Symbol;Acc:MGI:1860188] | 5299,6 | 99,53 |
| ENSMUSG00000014313 | Cox6c | cytochrome c oxidase subunit VIc [Source:MGI Symbol;Acc:MGI:104614] | 5262,1 | 77,66 |
| ENSMUSG00000018102 | Hist1h2bc | histone cluster 1, H2bc [Source:MGI Symbol;Acc:MGI:1915274] | 5158,8 | 71,78 |
| ENSMUSG00000029580 | Actb | actin, beta [Source:MGI Symbol;Acc:MGI:87904] | 5087,8 | 92,66 |
| ENSMUSG00000023456 | Tpi1 | triosephosphate isomerase 1 [Source:MGI Symbol;Acc:MGI:98797] | 5079,6 | 92,97 |
| ENSMUSG00000020671 | Rab10 | RAB10, member RAS oncogene family [Source:MGI Symbol;Acc:MGI:105066] | 5077,5 | 96,17 |
| ENSMUSG00000020321 | Mdh1 | malate dehydrogenase 1, NAD (soluble) [Source:MGI Symbol;Acc:MGI:97051] | 5058,5 | 91,38 |
| ENSMUSG00000031839 | Hsbp1 | heat shock factor binding protein 1 [Source:MGI Symbol;Acc:MGI:1915446] | 5014,2 | 100,00 |
| ENSMUSG00000006699 | Cdc42 | cell division cycle 42 [Source:MGI Symbol;Acc:MGI:106211] | 5008,4 | 99,92 |
| ENSMUSG00000030647 | Ndufc2 | NADH dehydrogenase (ubiquinone) 1, subcomplex unknown, 2 [Source:MGI Symbol;Acc:MGI:1344370] | 4999,8 | 93,63 |
| ENSMUSG00000032562 | Gnai2 | guanine nucleotide binding protein (G protein), alpha inhibiting 2 [Source:MGI Symbol;Acc:MGI:95772] | 4993,4 | 92,18 |
| ENSMUSG00000047187 | Rab2a | RAB2A, member RAS oncogene family [Source:MGI Symbol;Acc:MGI:1928750] | 4964,9 | 97,97 |
| ENSMUSG00000026664 | Phyh | phytanoyl-CoA hydroxylase [Source:MGI Symbol;Acc:MGI:891978] | 4945,9 | 100,00 |
| ENSMUSG00000025428 | Atp5a1 | TP synthase, H+ transporting, mitochondrial F1 complex, alpha subunit 1 [Source:MGI Symbol;Acc:MGI:8811 | 4941,1 | 86,89 |

| | | | | |
|---|---|---|---|---|
| ENSMUSG00000032181 | Scg3 | secretogranin III [Source:MGI Symbol;Acc:MGI:103032] | 4924,2 | 100,00 |
| ENSMUSG00000036751 | Cox6b1 | cytochrome c oxidase, subunit VIb polypeptide 1 [Source:MGI Symbol;Acc:MGI:107460] | 4895,1 | 91,05 |
| ENSMUSG00000040055 | Gjb6 | gap junction protein, beta 6 [Source:MGI Symbol;Acc:MGI:107588] | 4828,8 | 100,00 |
| ENSMUSG00000030317 | Timp4 | tissue inhibitor of metalloproteinase 4 [Source:MGI Symbol;Acc:MGI:109125] | 4825,9 | 93,79 |
| ENSMUSG00000022132 | Cldn10 | claudin 10 [Source:MGI Symbol;Acc:MGI:1913101] | 4798,4 | 76,51 |
| ENSMUSG00000029030 | Tprgl | transformation related protein 63 regulated like [Source:MGI Symbol;Acc:MGI:1915058] | 4771,5 | 100,00 |
| ENSMUSG00000033059 | Pygb | brain glycogen phosphorylase [Source:MGI Symbol;Acc:MGI:97828] | 4740,9 | 100,00 |
| ENSMUSG00000020733 | Slc9a3r1 | ute carrier family 9 (sodium/hydrogen exchanger), member 3 regulator 1 [Source:MGI Symbol;Acc:MGI:13494 | 4739,3 | 100,00 |
| ENSMUSG00000046240 | Hepacam | hepatocyte cell adhesion molecule [Source:MGI Symbol;Acc:MGI:1920177] | 4657,1 | 100,00 |
| ENSMUSG00000032320 | Rcn2 | reticulocalbin 2 [Source:MGI Symbol;Acc:MGI:1349765] | 4624,9 | 99,52 |
| ENSMUSG00000018593 | Sparc | secreted acidic cysteine rich glycoprotein [Source:MGI Symbol;Acc:MGI:98373] | 4608,5 | 97,97 |
| ENSMUSG00000022658 | Tagln3 | transgelin 3 [Source:MGI Symbol;Acc:MGI:1926784] | 4602,4 | 100,00 |
| ENSMUSG00000030654 | Arl6ip1 | ADP-ribosylation factor-like 6 interacting protein 1 [Source:MGI Symbol;Acc:MGI:1858943] | 4595,4 | 98,43 |
| ENSMUSG00000029632 | Ndufa4 | NADH dehydrogenase (ubiquinone) 1 alpha subcomplex, 4 [Source:MGI Symbol;Acc:MGI:107686] | 4589,9 | 100,00 |
| ENSMUSG00000053931 | Cnn3 | calponin 3, acidic [Source:MGI Symbol;Acc:MGI:1919244] | 4571,4 | 91,83 |
| ENSMUSG00000047557 | Lxn | latexin [Source:MGI Symbol;Acc:MGI:107633] | 4546,5 | 100,00 |
| ENSMUSG00000032199 | Polr2m | polymerase (RNA) II (DNA directed) polypeptide M [Source:MGI Symbol;Acc:MGI:107282] | 4537,2 | 100,00 |
| ENSMUSG00000023175 | Bsg | basigin [Source:MGI Symbol;Acc:MGI:88208] | 4535,2 | 93,66 |
| ENSMUSG00000022205 | Sub1 | SUB1 homolog (S. cerevisiae) [Source:MGI Symbol;Acc:MGI:104811] | 4504,9 | 96,21 |
| ENSMUSG00000028618 | Tmem59 | transmembrane protein 59 [Source:MGI Symbol;Acc:MGI:1929278] | 4502,1 | 92,75 |
| ENSMUSG00000006651 | Aplp1 | amyloid beta (A4) precursor-like protein 1 [Source:MGI Symbol;Acc:MGI:88046] | 4500,2 | 100,00 |
| ENSMUSG00000039621 | Prex1 | osphatidylinositol-3,4,5-trisphosphate-dependent Rac exchange factor 1 [Source:MGI Symbol;Acc:MGI:30406 | 4461,3 | 92,93 |
| ENSMUSG00000028194 | Ddah1 | dimethylarginine dimethylaminohydrolase 1 [Source:MGI Symbol;Acc:MGI:1916469] | 4449,6 | 99,26 |
| ENSMUSG00000000563 | Atp5f1 | ATP synthase, H+ transporting, mitochondrial F0 complex, subunit B1 [Source:MGI Symbol;Acc:MGI:110049 | 4435,6 | 95,49 |
| ENSMUSG00000024411 | Aqp4 | aquaporin 4 [Source:MGI Symbol;Acc:MGI:107387] | 4397,6 | 99,90 |
| ENSMUSG00000020585 | Laptm4a | lysosomal-associated protein transmembrane 4A [Source:MGI Symbol;Acc:MGI:108017] | 4396,1 | 90,41 |
| ENSMUSG00000028557 | Rnf11 | ring finger protein 11 [Source:MGI Symbol;Acc:MGI:1352759] | 4395,7 | 94,82 |
| ENSMUSG00000056515 | Rab31 | RAB31, member RAS oncogene family [Source:MGI Symbol;Acc:MGI:1914603] | 4392,7 | 99,44 |
| ENSMUSG00000041697 | Cox6a1 | cytochrome c oxidase subunit VIa polypeptide 1 [Source:MGI Symbol;Acc:MGI:103099] | 4385,9 | 100,00 |
| ENSMUSG00000022956 | Atp5o | ATP synthase, H+ transporting, mitochondrial F1 complex, O subunit [Source:MGI Symbol;Acc:MGI:106341] | 4385,1 | 97,87 |
| ENSMUSG00000041556 | Fbxo2 | F-box protein 2 [Source:MGI Symbol;Acc:MGI:2446216] | 4362,4 | 100,00 |
| ENSMUSG00000020333 | Acsl6 | acyl-CoA synthetase long-chain family member 6 [Source:MGI Symbol;Acc:MGI:894291] | 4355,3 | 94,11 |
| ENSMUSG00000021939 | Ctsb | cathepsin B [Source:MGI Symbol;Acc:MGI:88561] | 4295,3 | 99,01 |
| ENSMUSG00000073639 | Rab18 | RAB18, member RAS oncogene family [Source:MGI Symbol;Acc:MGI:102790] | 4276,3 | 100,00 |
| ENSMUSG00000007872 | Id3 | inhibitor of DNA binding 3 [Source:MGI Symbol;Acc:MGI:96398] | 4238,2 | 84,74 |
| ENSMUSG00000034158 | Lrrc58 | leucine rich repeat containing 58 [Source:MGI Symbol;Acc:MGI:2443542] | 4234,6 | 91,46 |
| ENSMUSG00000030310 | Slc6a1 | solute carrier family 6 (neurotransmitter transporter, GABA), member 1 [Source:MGI Symbol;Acc:MGI:95627] | 4233,9 | 99,84 |
| ENSMUSG00000025283 | Sat1 | spermidine/spermine N1-acetyl transferase 1 [Source:MGI Symbol;Acc:MGI:98233] | 4210,7 | 91,91 |
| ENSMUSG00000039197 | Adk | adenosine kinase [Source:MGI Symbol;Acc:MGI:87930] | 4193,7 | 99,68 |
| ENSMUSG00000031176 | Dynlt3 | dynein light chain Tctex-type 3 [Source:MGI Symbol;Acc:MGI:1914367] | 4171,3 | 99,86 |
| ENSMUSG00000041571 | Sepw1 | selenoprotein W, muscle 1 [Source:MGI Symbol;Acc:MGI:1100878] | 4154,7 | 99,91 |
| ENSMUSG00000024099 | Ndufv2 | NADH dehydrogenase (ubiquinone) flavoprotein 2 [Source:MGI Symbol;Acc:MGI:1920150] | 4121,3 | 83,33 |
| ENSMUSG00000018567 | Gabarap | gamma-aminobutyric acid receptor associated protein [Source:MGI Symbol;Acc:MGI:1861742] | 4116,9 | 86,43 |
| ENSMUSG00000075706 | Gpx4 | glutathione peroxidase 4 [Source:MGI Symbol;Acc:MGI:104767] | 4102,1 | 89,05 |
| ENSMUSG00000062014 | Gmfb | glia maturation factor, beta [Source:MGI Symbol;Acc:MGI:1927133] | 4064,8 | 94,94 |
| ENSMUSG00000000247 | Lhx2 | LIM homeobox protein 2 [Source:MGI Symbol;Acc:MGI:96785] | 4042,3 | 86,54 |
| ENSMUSG00000026150 | Mff | mitochondrial fission factor [Source:MGI Symbol;Acc:MGI:1922984] | 4010,3 | 99,02 |
| ENSMUSG00000038967 | Pdk2 | pyruvate dehydrogenase kinase, isoenzyme 2 [Source:MGI Symbol;Acc:MGI:1343087] | 4005,2 | 100,00 |
| ENSMUSG00000027805 | Pfn2 | profilin 2 [Source:MGI Symbol;Acc:MGI:97550] | 3989,0 | 71,96 |
| ENSMUSG00000000088 | Cox5a | cytochrome c oxidase subunit Va [Source:MGI Symbol;Acc:MGI:88474] | 3986,5 | 100,00 |
| ENSMUSG00000079477 | Rab7 | RAB7, member RAS oncogene family [Source:MGI Symbol;Acc:MGI:105068] | 3970,3 | 83,23 |
| ENSMUSG00000035133 | Arhgap5 | Rho GTPase activating protein 5 [Source:MGI Symbol;Acc:MGI:1332637] | 3949,3 | 100,00 |
| ENSMUSG00000062397 | Zfp706 | zinc finger protein 706 [Source:MGI Symbol;Acc:MGI:1915286] | 3931,0 | 93,09 |
| ENSMUSG00000036438 | Calm2 | calmodulin 2 [Source:MGI Symbol;Acc:MGI:103250] | 3903,9 | 97,93 |
| ENSMUSG00000003072 | Atp5d | TP synthase, H+ transporting, mitochondrial F1 complex, delta subunit [Source:MGI Symbol;Acc:MGI:191329 | 3885,7 | 89,63 |
| ENSMUSG00000029063 | Nadk | NAD kinase [Source:MGI Symbol;Acc:MGI:2183149] | 3882,9 | 88,32 |
| ENSMUSG00000048022 | Tmem229a | transmembrane protein 229A [Source:MGI Symbol;Acc:MGI:2442812] | 3858,0 | 99,02 |
| ENSMUSG00000042109 | Csdc2 | cold shock domain containing C2, RNA binding [Source:MGI Symbol;Acc:MGI:2146027] | 3821,7 | 96,26 |
| ENSMUSG00000028445 | Enho | energy homeostasis associated [Source:MGI Symbol;Acc:MGI:1916888] | 3812,1 | 99,53 |
| ENSMUSG00000031246 | Sh3bgrl | SH3-binding domain glutamic acid-rich protein like [Source:MGI Symbol;Acc:MGI:1930849] | 3800,3 | 99,40 |
| ENSMUSG00000024758 | Rtn3 | reticulon 3 [Source:MGI Symbol;Acc:MGI:1339970] | 3796,0 | 82,14 |
| ENSMUSG00000029840 | Mtpn | myotrophin [Source:MGI Symbol;Acc:MGI:99445] | 3793,9 | 99,65 |
| ENSMUSG00000030704 | Rab6a | RAB6A, member RAS oncogene family [Source:MGI Symbol;Acc:MGI:894313] | 3768,5 | 97,09 |
| ENSMUSG00000028945 | Rheb | Ras homolog enriched in brain [Source:MGI Symbol;Acc:MGI:97912] | 3760,0 | 85,07 |
| ENSMUSG00000032330 | Cox7a2 | cytochrome c oxidase subunit VIIa 2 [Source:MGI Symbol;Acc:MGI:1316715] | 3740,7 | 92,48 |
| ENSMUSG00000001687 | Ubl3 | ubiquitin-like 3 [Source:MGI Symbol;Acc:MGI:1344373] | 3740,5 | 99,92 |
| ENSMUSG00000029776 | Hibadh | 3-hydroxyisobutyrate dehydrogenase [Source:MGI Symbol;Acc:MGI:1889802] | 3720,8 | 99,45 |
| ENSMUSG00000032826 | Ank2 | ankyrin 2, brain [Source:MGI Symbol;Acc:MGI:88025] | 3700,2 | 73,73 |
| ENSMUSG00000024213 | Nudt3 | nudix (nucleotide diphosphate linked moiety X)-type motif 3 [Source:MGI Symbol;Acc:MGI:1928484] | 3686,8 | 93,75 |
| ENSMUSG00000057880 | Abat | 4-aminobutyrate aminotransferase [Source:MGI Symbol;Acc:MGI:2443582] | 3680,1 | 95,36 |
| ENSMUSG00000000171 | Sdhd | ccinate dehydrogenase complex, subunit D, integral membrane protein [Source:MGI Symbol;Acc:MGI:191417 | 3649,6 | 99,82 |
| ENSMUSG00000025351 | Cd63 | CD63 antigen [Source:MGI Symbol;Acc:MGI:99529] | 3643,7 | 99,81 |
| ENSMUSG00000021877 | Arf4 | ADP-ribosylation factor 4 [Source:MGI Symbol;Acc:MGI:99433] | 3635,1 | 99,51 |
| ENSMUSG00000026626 | Ppp2r5a | protein phosphatase 2, regulatory subunit B', alpha [Source:MGI Symbol;Acc:MGI:2388479] | 3623,3 | 99,96 |
| ENSMUSG00000022474 | Pmm1 | phosphomannomutase 1 [Source:MGI Symbol;Acc:MGI:1353418] | 3622,5 | 96,82 |
| ENSMUSG00000048388 | Fam171b | family with sequence similarity 171, member B [Source:MGI Symbol;Acc:MGI:2444579] | 3611,0 | 100,00 |
| ENSMUSG00000030605 | Mfge8 | milk fat globule-EGF factor 8 protein [Source:MGI Symbol;Acc:MGI:102768] | 3595,0 | 97,62 |
| ENSMUSG00000024646 | Cyb5a | cytochrome b5 type A (microsomal) [Source:MGI Symbol;Acc:MGI:1926952] | 3590,4 | 72,23 |
| ENSMUSG00000028552 | Eps15 | epidermal growth factor receptor pathway substrate 15 [Source:MGI Symbol;Acc:MGI:104583] | 3577,1 | 99,17 |
| ENSMUSG00000031776 | Arl2bp | ADP-ribosylation factor-like 2 binding protein [Source:MGI Symbol;Acc:MGI:1349429] | 3573,5 | 100,00 |
| ENSMUSG00000053477 | Tcf4 | transcription factor 4 [Source:MGI Symbol;Acc:MGI:98506] | 3569,7 | 75,00 |
| ENSMUSG00000021238 | Aldh6a1 | aldehyde dehydrogenase family 6, subfamily A1 [Source:MGI Symbol;Acc:MGI:1915077] | 3555,3 | 100,00 |
| ENSMUSG00000032294 | Pkm | pyruvate kinase, muscle [Source:MGI Symbol;Acc:MGI:97591] | 3554,1 | 97,35 |
| ENSMUSG00000086968 | 4933431E20Rik | RIKEN cDNA 4933431E20 gene [Source:MGI Symbol;Acc:MGI:3584041] | 3550,7 | 87,57 |
| ENSMUSG00000031231 | Cox7b | cytochrome c oxidase subunit VIIb [Source:MGI Symbol;Acc:MGI:1913392] | 3550,3 | 99,87 |
| ENSMUSG00000008035 | Mid1ip1 | Mid1 interacting protein 1 (gastrulation specific G12-like (zebrafish)) [Source:MGI Symbol;Acc:MGI:1915291] | 3547,8 | 99,82 |
| ENSMUSG00000102918 | Pcdhgc3 | protocadherin gamma subfamily C, 3 [Source:MGI Symbol;Acc:MGI:1935201] | 3543,2 | 100,00 |
| ENSMUSG00000032966 | Fkbp1a | FK506 binding protein 1a [Source:MGI Symbol;Acc:MGI:95541] | 3530,9 | 99,86 |
| ENSMUSG00000001018 | Snapin | SNAP-associated protein [Source:MGI Symbol;Acc:MGI:1333745] | 3526,1 | 100,00 |
| ENSMUSG00000036570 | Fxyd1 | FXYD domain-containing ion transport regulator 1 [Source:MGI Symbol;Acc:MGI:1889273] | 3511,8 | 93,83 |
| ENSMUSG00000063882 | Uqcrh | ubiquinol-cytochrome c reductase hinge protein [Source:MGI Symbol;Acc:MGI:1913826] | 3500,0 | 100,00 |
| ENSMUSG00000036585 | Fgf1 | fibroblast growth factor 1 [Source:MGI Symbol;Acc:MGI:95515] | 3487,1 | 92,37 |
| ENSMUSG00000063297 | Luzp2 | leucine zipper protein 2 [Source:MGI Symbol;Acc:MGI:1889615] | 3485,7 | 99,99 |
| ENSMUSG00000059534 | Uqcr10 | ubiquinol-cytochrome c reductase, complex III subunit X [Source:MGI Symbol;Acc:MGI:1913402] | 3446,9 | 100,00 |
| ENSMUSG00000020894 | Vamp2 | vesicle-associated membrane protein 2 [Source:MGI Symbol;Acc:MGI:1313277] | 3441,5 | 93,67 |
| ENSMUSG00000078812 | Eif5a | eukaryotic translation initiation factor 5A [Source:MGI Symbol;Acc:MGI:106248] | 3427,9 | 73,58 |
| ENSMUSG00000033350 | Chst2 | carbohydrate sulfotransferase 2 [Source:MGI Symbol;Acc:MGI:1891160] | 3424,4 | 99,80 |
| ENSMUSG00000004056 | Akt2 | thymoma viral proto-oncogene 2 [Source:MGI Symbol;Acc:MGI:104874] | 3419,9 | 86,94 |
| ENSMUSG00000027351 | Spred1 | sprouty protein with EVH-1 domain 1, related sequence [Source:MGI Symbol;Acc:MGI:2150016] | 3418,9 | 98,93 |
| ENSMUSG00000024347 | Psd2 | pleckstrin and Sec7 domain containing 2 [Source:MGI Symbol;Acc:MGI:1921252] | 3363,0 | 95,82 |
| ENSMUSG00000034566 | Atp5h | ATP synthase, H+ transporting, mitochondrial F0 complex, subunit D [Source:MGI Symbol;Acc:MGI:1918929 | 3355,3 | 97,93 |
| ENSMUSG00000007891 | Ctsd | cathepsin D [Source:MGI Symbol;Acc:MGI:88562] | 3328,6 | 97,86 |
| ENSMUSG00000028692 | Akr1a1 | aldo-keto reductase family 1, member A1 (aldehyde reductase) [Source:MGI Symbol;Acc:MGI:1929955] | 3327,0 | 92,38 |
| ENSMUSG00000021577 | Sdha | succinate dehydrogenase complex, subunit A, flavoprotein (Fp) [Source:MGI Symbol;Acc:MGI:1914195] | 3327,0 | 99,17 |
| ENSMUSG00000005161 | Prdx2 | peroxiredoxin 2 [Source:MGI Symbol;Acc:MGI:109486] | 3321,8 | 99,05 |
| ENSMUSG00000061904 | Slc25a3 | ute carrier family 25 (mitochondrial carrier, phosphate carrier), member 3 [Source:MGI Symbol;Acc:MGI:13534 | 3316,2 | 98,75 |
| ENSMUSG00000040549 | Ckap5 | cytoskeleton associated protein 5 [Source:MGI Symbol;Acc:MGI:1923036] | 3304,8 | 94,94 |
| ENSMUSG00000027002 | Nckap1 | NCK-associated protein 1 [Source:MGI Symbol;Acc:MGI:1355333] | 3301,3 | 98,68 |
| ENSMUSG00000019370 | Calm3 | calmodulin 3 [Source:MGI Symbol;Acc:MGI:103249] | 3298,4 | 99,71 |
| ENSMUSG00000027273 | Snap25 | synaptosomal-associated protein 25 [Source:MGI Symbol;Acc:MGI:98331] | 3297,9 | 99,91 |
| ENSMUSG00000033174 | Mgll | monoglyceride lipase [Source:MGI Symbol;Acc:MGI:1346042] | 3294,5 | 88,24 |
| ENSMUSG00000033953 | Ppp3r1 | in phosphatase 3, regulatory subunit B, alpha isoform (calcineurin B, type I) [Source:MGI Symbol;Acc:MGI:10 | 3287,1 | 99,21 |
| ENSMUSG00000009863 | Sdhb | succinate dehydrogenase complex, subunit B, iron sulfur (Ip) [Source:MGI Symbol;Acc:MGI:1914930] | 3260,2 | 99,42 |
| ENSMUSG00000040759 | Cmtm5 | CKLF-like MARVEL transmembrane domain containing 5 [Source:MGI Symbol;Acc:MGI:2447164] | 3259,6 | 100,00 |
| ENSMUSG00000020612 | Prkar1a | protein kinase, cAMP dependent regulatory, type I, alpha [Source:MGI Symbol;Acc:MGI:104878] | 3249,5 | 90,12 |
| ENSMUSG00000040048 | Ndufb10 | NADH dehydrogenase (ubiquinone) 1 beta subcomplex, 10 [Source:MGI Symbol;Acc:MGI:1915592] | 3248,3 | 98,76 |
| ENSMUSG00000015837 | Sqstm1 | sequestosome 1 [Source:MGI Symbol;Acc:MGI:107931] | 3246,5 | 92,89 |

| | | | | |
|---|---|---|---|---|
| ENSMUSG00000004071 | Cdip1 | cell death inducing Trp53 target 1 [Source:MGI Symbol;Acc:MGI:1913876] | 3245,1 | 97,33 |
| ENSMUSG00000022477 | Aco2 | aconitase 2, mitochondrial [Source:MGI Symbol;Acc:MGI:87880] | 3240,0 | 97,36 |
| ENSMUSG00000036402 | Gng12 | guanine nucleotide binding protein (G protein), gamma 12 [Source:MGI Symbol;Acc:MGI:1336171] | 3228,4 | 98,48 |
| ENSMUSG00000025132 | Arhgdia | Rho GDP dissociation inhibitor (GDI) alpha [Source:MGI Symbol;Acc:MGI:2178103] | 3223,3 | 92,87 |
| ENSMUSG00000025103 | Btbd1 | BTB (POZ) domain containing 1 [Source:MGI Symbol;Acc:MGI:1933765] | 3220,1 | 89,59 |
| ENSMUSG00000028982 | Slc25a33 | solute carrier family 25, member 33 [Source:MGI Symbol;Acc:MGI:1917806] | 3218,5 | 100,00 |
| ENSMUSG00000020149 | Rab1a | RAB1A, member RAS oncogene family [Source:MGI Symbol;Acc:MGI:97842] | 3214,8 | 98,51 |
| ENSMUSG00000031748 | Gnao1 | guanine nucleotide binding protein, alpha O [Source:MGI Symbol;Acc:MGI:95775] | 3210,9 | 98,13 |
| ENSMUSG00000020611 | Gna13 | guanine nucleotide binding protein, alpha 13 [Source:MGI Symbol;Acc:MGI:95768] | 3193,9 | 99,91 |
| ENSMUSG00000021290 | 2010107E04Rik | RIKEN cDNA 2010107E04 gene [Source:MGI Symbol;Acc:MGI:1917507] | 3154,7 | 100,00 |
| ENSMUSG00000004902 | Slc25a18 | solute carrier family 25 (mitochondrial carrier), member 18 [Source:MGI Symbol;Acc:MGI:1919053] | 3148,9 | 100,00 |
| ENSMUSG00000024012 | Mtch1 | mitochondrial carrier homolog 1 (C. elegans) [Source:MGI Symbol;Acc:MGI:1929261] | 3129,5 | 96,47 |
| ENSMUSG00000037742 | Eef1a1 | eukaryotic translation elongation factor 1 alpha 1 [Source:MGI Symbol;Acc:MGI:1096881] | 3129,0 | 80,10 |
| ENSMUSG00000027333 | Smox | spermine oxidase [Source:MGI Symbol;Acc:MGI:2445356] | 3121,9 | 83,29 |
| ENSMUSG00000004207 | Psap | prosaposin [Source:MGI Symbol;Acc:MGI:97783] | 3091,9 | 95,77 |
| ENSMUSG00000102543 | Pcdhgc5 | protocadherin gamma subfamily C, 5 [Source:MGI Symbol;Acc:MGI:1935205] | 3085,7 | 99,16 |
| ENSMUSG00000067786 | Nnat | neuronatin [Source:MGI Symbol;Acc:MGI:104716] | 3078,5 | 93,02 |
| ENSMUSG00000032060 | Cryab | crystallin, alpha B [Source:MGI Symbol;Acc:MGI:88516] | 3078,2 | 95,33 |
| ENSMUSG00000038462 | Uqcrfs1 | ubiquinol-cytochrome c reductase, Rieske iron-sulfur polypeptide 1 [Source:MGI Symbol;Acc:MGI:1913944] | 3073,1 | 100,00 |
| ENSMUSG00000025931 | Paqr8 | progestin and adipoQ receptor family member VIII [Source:MGI Symbol;Acc:MGI:1921479] | 3073,0 | 97,12 |
| ENSMUSG00000020163 | Uqcr11 | ubiquinol-cytochrome c reductase, complex III subunit XI [Source:MGI Symbol;Acc:MGI:1913844] | 3052,3 | 100,00 |
| ENSMUSG00000022253 | Nadk2 | NAD kinase 2, mitochondrial [Source:MGI Symbol;Acc:MGI:1915896] | 3050,2 | 92,80 |
| ENSMUSG00000103472 | Pcdhga7 | protocadherin gamma subfamily A, 7 [Source:MGI Symbol;Acc:MGI:1935219] | 3040,1 | 86,98 |
| ENSMUSG00000103897 | Pcdhga8 | protocadherin gamma subfamily A, 8 [Source:MGI Symbol;Acc:MGI:1935221] | 3038,6 | 92,18 |
| ENSMUSG00000104063 | Pcdhgb7 | protocadherin gamma subfamily B, 7 [Source:MGI Symbol;Acc:MGI:1935199] | 3037,5 | 97,03 |
| ENSMUSG00000103088 | Pcdhgb6 | protocadherin gamma subfamily B, 6 [Source:MGI Symbol;Acc:MGI:1935197] | 3037,3 | 93,08 |
| ENSMUSG00000055302 | Mrfap1 | Morf4 family associated protein 1 [Source:MGI Symbol;Acc:MGI:1914818] | 3032,4 | 84,59 |
| ENSMUSG00000102428 | Pcdhga12 | protocadherin gamma subfamily A, 12 [Source:MGI Symbol;Acc:MGI:1935229] | 3027,4 | 80,23 |
| ENSMUSG00000102742 | Pcdhga11 | protocadherin gamma subfamily A, 11 [Source:MGI Symbol;Acc:MGI:1935228] | 3021,8 | 92,91 |
| ENSMUSG00000103793 | Pcdhga6 | protocadherin gamma subfamily A, 6 [Source:MGI Symbol;Acc:MGI:1935218] | 3018,5 | 85,90 |
| ENSMUSG00000103037 | Pcdhgb1 | protocadherin gamma subfamily B, 1 [Source:MGI Symbol;Acc:MGI:1935169] | 3015,5 | 80,78 |
| ENSMUSG00000103567 | Pcdhga5 | protocadherin gamma subfamily A, 5 [Source:MGI Symbol;Acc:MGI:1935217] | 3011,8 | 85,68 |
| ENSMUSG00000052738 | Suclg1 | succinate-CoA ligase, GDP-forming, alpha subunit [Source:MGI Symbol;Acc:MGI:1927234] | 3010,1 | 89,20 |
| ENSMUSG00000104346 | Pcdhga3 | protocadherin gamma subfamily A, 3 [Source:MGI Symbol;Acc:MGI:1935215] | 3009,9 | 85,68 |
| ENSMUSG00000006356 | Crip2 | cysteine rich protein 2 [Source:MGI Symbol;Acc:MGI:1915587] | 3009,7 | 93,73 |
| ENSMUSG00000023036 | Pcdhga8 | protocadherin gamma subfamily A, 8 [Source:MGI Symbol;Acc:MGI:1935221] | 3008,5 | 77,21 |
| ENSMUSG00000019923 | Zwint | ZW10 interactor [Source:MGI Symbol;Acc:MGI:1289227] | 3006,5 | 70,55 |
| ENSMUSG00000103677 | Pcdhga4 | protocadherin gamma subfamily A, 4 [Source:MGI Symbol;Acc:MGI:1935216] | 3006,3 | 76,57 |
| ENSMUSG00000022678 | Nde1 | nuclear distribution gene E homolog 1 (A nidulans) [Source:MGI Symbol;Acc:MGI:1914453] | 3006,1 | 100,00 |
| ENSMUSG00000005683 | Cs | citrate synthase [Source:MGI Symbol;Acc:MGI:88529] | 3002,5 | 99,99 |
| ENSMUSG00000102748 | Pcdhgb2 | protocadherin gamma subfamily B, 2 [Source:MGI Symbol;Acc:MGI:1935170] | 3001,8 | 70,34 |
| ENSMUSG00000028675 | Pnrc2 | proline-rich nuclear receptor coactivator 2 [Source:MGI Symbol;Acc:MGI:106512] | 2993,6 | 100,00 |
| ENSMUSG00000022186 | Oxct1 | 3-oxoacid CoA transferase 1 [Source:MGI Symbol;Acc:MGI:1914291] | 2988,6 | 99,57 |
| ENSMUSG00000055538 | Zcchc24 | zinc finger, CCHC domain containing 24 [Source:MGI Symbol;Acc:MGI:1919168] | 2983,0 | 99,22 |
| ENSMUSG00000024712 | Rfk | riboflavin kinase [Source:MGI Symbol;Acc:MGI:1914688] | 2977,6 | 100,00 |
| ENSMUSG00000031924 | Cyb5b | cytochrome b5 type B [Source:MGI Symbol;Acc:MGI:1913677] | 2963,3 | 99,98 |
| ENSMUSG00000009013 | Dynll1 | dynein light chain LC8-type 1 [Source:MGI Symbol;Acc:MGI:1861457] | 2951,8 | 93,11 |
| ENSMUSG00000034341 | Wbp2 | WW domain binding protein 2 [Source:MGI Symbol;Acc:MGI:104709] | 2945,6 | 100,00 |
| ENSMUSG00000015149 | Sirt2 | sirtuin 2 [Source:MGI Symbol;Acc:MGI:1927664] | 2945,4 | 83,33 |
| ENSMUSG00000025421 | Hdhd2 | haloacid dehalogenase-like hydrolase domain containing 2 [Source:MGI Symbol;Acc:MGI:1924237] | 2945,4 | 79,44 |
| ENSMUSG00000037152 | Ndufc1 | NADH dehydrogenase (ubiquinone) 1, subcomplex unknown, 1 [Source:MGI Symbol;Acc:MGI:1913627] | 2927,8 | 96,58 |
| ENSMUSG00000021520 | Uqcrb | ubiquinol-cytochrome c reductase binding protein [Source:MGI Symbol;Acc:MGI:1914780] | 2911,2 | 99,19 |
| ENSMUSG00000102252 | Snrpn | small nuclear ribonucleoprotein N [Source:MGI Symbol;Acc:MGI:98347] | 2910,3 | 94,47 |
| ENSMUSG00000024091 | Vapa | vesicle-associated membrane protein, associated protein A [Source:MGI Symbol;Acc:MGI:1353561] | 2899,8 | 100,00 |
| ENSMUSG00000038375 | Trp53inp2 | transformation related protein 53 inducible nuclear protein 2 [Source:MGI Symbol;Acc:MGI:1915978] | 2888,7 | 97,12 |
| ENSMUSG00000037058 | Paip2 | polyadenylate-binding protein-interacting protein 2 [Source:MGI Symbol;Acc:MGI:1915119] | 2877,3 | 87,25 |
| ENSMUSG00000005873 | Reep5 | receptor accessory protein 5 [Source:MGI Symbol;Acc:MGI:1270152] | 2859,5 | 99,79 |
| ENSMUSG00000027367 | Stard7 | START domain containing 7 [Source:MGI Symbol;Acc:MGI:2139090] | 2858,3 | 75,25 |
| ENSMUSG00000029064 | Gnb1 | guanine nucleotide binding protein (G protein), beta 1 [Source:MGI Symbol;Acc:MGI:95781] | 2838,0 | 90,69 |
| ENSMUSG00000040659 | Efhd2 | EF hand domain containing 2 [Source:MGI Symbol;Acc:MGI:106504] | 2832,3 | 99,83 |
| ENSMUSG00000030161 | Gabarapl1 | amma-aminobutyric acid (GABA) A receptor-associated protein-like 1 [Source:MGI Symbol;Acc:MGI:191498( | 2811,1 | 98,97 |
| ENSMUSG00000027423 | Snx5 | sorting nexin 5 [Source:MGI Symbol;Acc:MGI:1916428] | 2810,5 | 100,00 |
| ENSMUSG00000026509 | Capn2 | calpain 2 [Source:MGI Symbol;Acc:MGI:88264] | 2807,6 | 100,00 |
| ENSMUSG00000033065 | Pfkm | phosphofructokinase, muscle [Source:MGI Symbol;Acc:MGI:97548] | 2805,8 | 96,92 |
| ENSMUSG00000017631 | Abr | active BCR-related gene [Source:MGI Symbol;Acc:MGI:107771] | 2803,6 | 78,13 |
| ENSMUSG00000022452 | Smdt1 | single-pass membrane protein with aspartate rich tail 1 [Source:MGI Symbol;Acc:MGI:1916279] | 2803,2 | 87,16 |
| ENSMUSG00000026223 | Itm2c | integral membrane protein 2C [Source:MGI Symbol;Acc:MGI:1927594] | 2790,6 | 91,24 |
| ENSMUSG00000037060 | Prkcdbp | protein kinase C, delta binding protein [Source:MGI Symbol;Acc:MGI:1923422] | 2782,9 | 100,00 |
| ENSMUSG00000021282 | Eif5 | eukaryotic translation initiation factor 5 [Source:MGI Symbol;Acc:MGI:95309] | 2774,2 | 92,49 |
| ENSMUSG00000021508 | Cxcl14 | chemokine (C-X-C motif) ligand 14 [Source:MGI Symbol;Acc:MGI:1888514] | 2766,7 | 99,87 |
| ENSMUSG00000033998 | Kcnk1 | potassium channel, subfamily K, member 1 [Source:MGI Symbol;Acc:MGI:109322] | 2759,4 | 100,00 |
| ENSMUSG00000028603 | Scp2 | sterol carrier protein 2, liver [Source:MGI Symbol;Acc:MGI:98254] | 2754,7 | 78,74 |
| ENSMUSG00000061718 | Ppp1r1b | protein phosphatase 1, regulatory (inhibitor) subunit 1B [Source:MGI Symbol;Acc:MGI:94860] | 2751,4 | 84,34 |
| ENSMUSG00000030934 | Oat | ornithine aminotransferase [Source:MGI Symbol;Acc:MGI:97394] | 2751,1 | 99,35 |
| ENSMUSG00000040785 | Ttc3 | tetratricopeptide repeat domain 3 [Source:MGI Symbol;Acc:MGI:1276539] | 2750,6 | 84,35 |
| ENSMUSG00000078578 | Ube2d3 | ubiquitin-conjugating enzyme E2D 3 [Source:MGI Symbol;Acc:MGI:1913355] | 2748,2 | 92,33 |
| ENSMUSG00000034875 | Nudt19 | nudix (nucleoside diphosphate linked moiety X)-type motif 19 [Source:MGI Symbol;Acc:MGI:94203] | 2742,0 | 99,55 |
| ENSMUSG00000028416 | Bag1 | BCL2-associated athanogene 1 [Source:MGI Symbol;Acc:MGI:108047] | 2741,6 | 100,00 |
| ENSMUSG00000020153 | Ndufs7 | NADH dehydrogenase (ubiquinone) Fe-S protein 7 [Source:MGI Symbol;Acc:MGI:1922656] | 2725,9 | 100,00 |
| ENSMUSG00000022010 | Tsc22d1 | TSC22 domain family, member 1 [Source:MGI Symbol;Acc:MGI:109127] | 2723,5 | 70,62 |
| ENSMUSG00000006299 | Aamp | angio-associated migratory protein [Source:MGI Symbol;Acc:MGI:107809] | 2712,9 | 88,33 |
| ENSMUSG00000037747 | Phyhipl | phytanoyl-CoA hydroxylase interacting protein-like [Source:MGI Symbol;Acc:MGI:1918161] | 2706,7 | 84,17 |
| ENSMUSG00000002475 | Abhd3 | abhydrolase domain containing 3 [Source:MGI Symbol;Acc:MGI:2147183] | 2700,4 | 99,95 |
| ENSMUSG00000032186 | Tmod2 | tropomodulin 2 [Source:MGI Symbol;Acc:MGI:1355335] | 2699,9 | 96,62 |
| ENSMUSG00000022018 | Rgcc | regulator of cell cycle [Source:MGI Symbol;Acc:MGI:1913464] | 2693,2 | 99,82 |
| ENSMUSG00000033940 | Brk1 | BRICK1, SCAR/WAVE actin-nucleating complex subunit [Source:MGI Symbol;Acc:MGI:1915406] | 2692,6 | 100,00 |
| ENSMUSG00000025204 | Ndufb8 | NADH dehydrogenase (ubiquinone) 1 beta subcomplex 8 [Source:MGI Symbol;Acc:MGI:1914514] | 2677,8 | 95,85 |
| ENSMUSG00000047459 | Dynlrb1 | dynein light chain roadblock-type 1 [Source:MGI Symbol;Acc:MGI:1914318] | 2677,0 | 99,50 |
| ENSMUSG00000031059 | Ndufb11 | NADH dehydrogenase (ubiquinone) 1 beta subcomplex, 11 [Source:MGI Symbol;Acc:MGI:1349919] | 2674,1 | 99,96 |
| ENSMUSG00000031819 | Emc8 | ER membrane protein complex subunit 8 [Source:MGI Symbol;Acc:MGI:1343095] | 2664,0 | 99,96 |
| ENSMUSG00000036564 | Ndrg4 | N-myc downstream regulated gene 4 [Source:MGI Symbol;Acc:MGI:2384590] | 2659,6 | 86,34 |
| ENSMUSG00000056486 | Chn1 | chimerin 1 [Source:MGI Symbol;Acc:MGI:1915674] | 2658,4 | 90,92 |
| ENSMUSG00000008393 | Carhsp1 | calcium regulated heat stable protein 1 [Source:MGI Symbol;Acc:MGI:1196368] | 2646,0 | 96,86 |
| ENSMUSG00000018326 | Ywhab | onooxygenase/tryptophan 5-monooxygenase activation protein, beta polypeptide [Source:MGI Symbol;Acc:M( | 2637,9 | 86,44 |
| ENSMUSG00000015994 | Fnta | farnesyltransferase, CAAX box, alpha [Source:MGI Symbol;Acc:MGI:104683] | 2628,5 | 98,81 |
| ENSMUSG00000024953 | Prdx5 | peroxiredoxin 5 [Source:MGI Symbol;Acc:MGI:1859821] | 2623,2 | 92,86 |
| ENSMUSG00000028161 | Ppp3ca | protein phosphatase 3, catalytic subunit, alpha isoform [Source:MGI Symbol;Acc:MGI:107164] | 2618,5 | 99,98 |
| ENSMUSG00000028745 | Capzb | capping protein (actin filament) muscle Z-line, beta [Source:MGI Symbol;Acc:MGI:104652] | 2618,2 | 79,73 |
| ENSMUSG00000025911 | Adhfe1 | alcohol dehydrogenase, iron containing, 1 [Source:MGI Symbol;Acc:MGI:1923437] | 2611,2 | 86,12 |
| ENSMUSG00000016559 | H3f3b | H3 histone, family 3B [Source:MGI Symbol;Acc:MGI:1101768] | 2604,7 | 71,19 |
| ENSMUSG00000037822 | Smim14 | small integral membrane protein 14 [Source:MGI Symbol;Acc:MGI:1915802] | 2595,2 | 95,34 |
| ENSMUSG00000019179 | Mdh2 | malate dehydrogenase 2, NAD (mitochondrial) [Source:MGI Symbol;Acc:MGI:97050] | 2590,9 | 97,09 |
| ENSMUSG00000032479 | Map4 | microtubule-associated protein 4 [Source:MGI Symbol;Acc:MGI:97178] | 2585,5 | 74,96 |
| ENSMUSG00000046671 | Mtfr1l | mitochondrial fission regulator 1-like [Source:MGI Symbol;Acc:MGI:1924074] | 2581,2 | 98,05 |
| ENSMUSG00000032667 | Pon2 | paraoxonase 2 [Source:MGI Symbol;Acc:MGI:106687] | 2580,7 | 96,60 |
| ENSMUSG00000028419 | Chmp5 | charged multivesicular body protein 5 [Source:MGI Symbol;Acc:MGI:1924209] | 2579,3 | 96,49 |
| ENSMUSG00000025651 | Uqcrc1 | ubiquinol-cytochrome c reductase core protein 1 [Source:MGI Symbol;Acc:MGI:107876] | 2557,4 | 91,70 |
| ENSMUSG00000024302 | Dtna | dystrobrevin alpha [Source:MGI Symbol;Acc:MGI:106039] | 2554,0 | 99,99 |
| ENSMUSG00000037458 | Azin1 | antizyme inhibitor 1 [Source:MGI Symbol;Acc:MGI:1859169] | 2535,1 | 96,23 |
| ENSMUSG00000015932 | Dstn | destrin [Source:MGI Symbol;Acc:MGI:1929270] | 2527,6 | 82,50 |
| ENSMUSG00000048076 | Arf1 | ADP-ribosylation factor 1 [Source:MGI Symbol;Acc:MGI:99431] | 2517,5 | 97,81 |
| ENSMUSG00000033933 | Vhl | von Hippel-Lindau tumor suppressor [Source:MGI Symbol;Acc:MGI:103223] | 2512,7 | 98,94 |
| ENSMUSG00000041881 | Ndufa7 | NADH dehydrogenase (ubiquinone) 1 alpha subcomplex, 7 (B14.5a) [Source:MGI Symbol;Acc:MGI:1913666] | 2499,3 | 99,80 |
| ENSMUSG00000068923 | Syt11 | synaptotagmin XI [Source:MGI Symbol;Acc:MGI:1859547] | 2491,4 | 91,23 |
| ENSMUSG00000072946 | Ptgr2 | prostaglandin reductase 2 [Source:MGI Symbol;Acc:MGI:1916372] | 2489,3 | 80,53 |
| ENSMUSG00000016252 | Atp5e | P synthase, H+ transporting, mitochondrial F1 complex, epsilon subunit [Source:MGI Symbol;Acc:MGI:18556 | 2482,3 | 99,76 |
| ENSMUSG00000058267 | Mrps14 | mitochondrial ribosomal protein S14 [Source:MGI Symbol;Acc:MGI:1928141] | 2481,6 | 90,41 |

| | | | | |
|---|---|---|---|---|
| ENSMUSG00000019877 | Serinc1 | serine incorporator 1 [Source:MGI Symbol;Acc:MGI:1926228] | 2476,6 | 99,91 |
| ENSMUSG00000025026 | Add3 | adducin 3 (gamma) [Source:MGI Symbol;Acc:MGI:1351615] | 2473,5 | 95,12 |
| ENSMUSG00000015090 | Ptgds | prostaglandin D2 synthase (brain) [Source:MGI Symbol;Acc:MGI:99261] | 2472,2 | 99,75 |
| ENSMUSG00000054091 | 1810037I17Rik | RIKEN cDNA 1810037I17 gene [Source:MGI Symbol;Acc:MGI:1914954] | 2472,2 | 91,03 |
| ENSMUSG00000093930 | Hmgcs1 | 3-hydroxy-3-methylglutaryl-Coenzyme A synthase 1 [Source:MGI Symbol;Acc:MGI:107592] | 2470,9 | 95,81 |
| ENSMUSG00000022564 | Grina | ceptor, ionotropic, N-methyl D-aspartate-associated protein 1 (glutamate binding) [Source:MGI Symbol;Acc:M | 2465,7 | 100,00 |
| ENSMUSG00000004980 | Hnrnpa2b1 | heterogeneous nuclear ribonucleoprotein A2/B1 [Source:MGI Symbol;Acc:MGI:104819] | 2462,3 | 99,90 |
| ENSMUSG00000024665 | Fads2 | fatty acid desaturase 2 [Source:MGI Symbol;Acc:MGI:1930079] | 2454,7 | 100,00 |
| ENSMUSG00000031299 | Pdha1 | pyruvate dehydrogenase E1 alpha 1 [Source:MGI Symbol;Acc:MGI:97532] | 2454,6 | 99,94 |
| ENSMUSG00000071014 | Ndufb6 | NADH dehydrogenase (ubiquinone) 1 beta subcomplex, 6 [Source:MGI Symbol;Acc:MGI:2684983] | 2452,4 | 100,00 |
| ENSMUSG00000054920 | Klhl5 | kelch-like 5 [Source:MGI Symbol;Acc:MGI:1919028] | 2445,7 | 98,47 |
| ENSMUSG00000024533 | Spire1 | spire homolog 1 (Drosophila) [Source:MGI Symbol;Acc:MGI:1915416] | 2444,4 | 98,88 |
| ENSMUSG00000019210 | Atp6v1e1 | ATPase, H+ transporting, lysosomal V1 subunit E1 [Source:MGI Symbol;Acc:MGI:894326] | 2443,7 | 95,31 |
| ENSMUSG00000020267 | Hint1 | histidine triad nucleotide binding protein 1 [Source:MGI Symbol;Acc:MGI:1321133] | 2439,9 | 72,97 |
| ENSMUSG00000040385 | Ppp1ca | protein phosphatase 1, catalytic subunit, alpha isoform [Source:MGI Symbol;Acc:MGI:103016] | 2439,8 | 100,00 |
| ENSMUSG00000074884 | Serf2 | small EDRK-rich factor 2 [Source:MGI Symbol;Acc:MGI:1337041] | 2435,5 | 84,01 |
| ENSMUSG00000015478 | Rnf5 | ring finger protein 5 [Source:MGI Symbol;Acc:MGI:1860076] | 2428,2 | 97,71 |
| ENSMUSG00000084957 | Bbip1 | BBSome interacting protein 1 [Source:MGI Symbol;Acc:MGI:1913610] | 2427,4 | 96,29 |
| ENSMUSG00000026895 | Ndufa8 | NADH dehydrogenase (ubiquinone) 1 alpha subcomplex, 8 [Source:MGI Symbol;Acc:MGI:1915625] | 2427,0 | 99,54 |
| ENSMUSG00000031700 | Gpt2 | glutamic pyruvate transaminase (alanine aminotransferase) 2 [Source:MGI Symbol;Acc:MGI:1915391] | 2425,3 | 100,00 |
| ENSMUSG00000015222 | Map2 | microtubule-associated protein 2 [Source:MGI Symbol;Acc:MGI:97175] | 2424,0 | 86,10 |
| ENSMUSG00000096210 | H1f0 | H1 histone family, member 0 [Source:MGI Symbol;Acc:MGI:95893] | 2419,1 | 95,98 |
| ENSMUSG00000017831 | Rab5a | RAB5A, member RAS oncogene family [Source:MGI Symbol;Acc:MGI:105926] | 2415,6 | 74,11 |
| ENSMUSG00000031532 | Saraf | store-operated calcium entry-associated regulatory factor [Source:MGI Symbol;Acc:MGI:1915137] | 2414,3 | 99,58 |
| ENSMUSG00000020326 | Ccng1 | cyclin G1 [Source:MGI Symbol;Acc:MGI:102890] | 2412,0 | 99,53 |
| ENSMUSG00000014294 | Ndufa2 | NADH dehydrogenase (ubiquinone) 1 alpha subcomplex, 2 [Source:MGI Symbol;Acc:MGI:1343103] | 2406,1 | 99,18 |
| ENSMUSG00000055430 | Nap1l5 | nucleosome assembly protein 1-like 5 [Source:MGI Symbol;Acc:MGI:1923555] | 2403,0 | 100,00 |
| ENSMUSG00000024870 | Rab1b | RAB1B, member RAS oncogene family [Source:MGI Symbol;Acc:MGI:1923558] | 2402,2 | 100,00 |
| ENSMUSG00000020496 | Rnf187 | ring finger protein 187 [Source:MGI Symbol;Acc:MGI:1914224] | 2392,7 | 100,00 |
| ENSMUSG00000000149 | Gna12 | guanine nucleotide binding protein, alpha 12 [Source:MGI Symbol;Acc:MGI:95767] | 2389,9 | 100,00 |
| ENSMUSG00000017781 | Pitpna | phosphatidylinositol transfer protein, alpha [Source:MGI Symbol;Acc:MGI:99887] | 2387,3 | 99,19 |
| ENSMUSG00000003161 | Sri | sorcin [Source:MGI Symbol;Acc:MGI:98419] | 2380,4 | 90,88 |
| ENSMUSG00000025465 | Echs1 | enoyl Coenzyme A hydratase, short chain, 1, mitochondrial [Source:MGI Symbol;Acc:MGI:2136460] | 2378,4 | 99,36 |
| ENSMUSG00000022551 | Cyc1 | cytochrome c-1 [Source:MGI Symbol;Acc:MGI:1913695] | 2365,7 | 100,00 |
| ENSMUSG00000032279 | Idh3a | isocitrate dehydrogenase 3 (NAD+) alpha [Source:MGI Symbol;Acc:MGI:1915084] | 2364,8 | 100,00 |
| ENSMUSG00000022200 | Golph3 | golgi phosphoprotein 3 [Source:MGI Symbol;Acc:MGI:1913879] | 2362,7 | 100,00 |
| ENSMUSG00000041126 | H2afv | H2A histone family, member V [Source:MGI Symbol;Acc:MGI:1924855] | 2361,3 | 99,98 |
| ENSMUSG00000024640 | Psat1 | phosphoserine aminotransferase 1 [Source:MGI Symbol;Acc:MGI:2183441] | 2354,7 | 94,86 |
| ENSMUSG00000022361 | Zhx1 | zinc fingers and homeoboxes 1 [Source:MGI Symbol;Acc:MGI:109271] | 2354,6 | 98,21 |
| ENSMUSG00000033863 | Klf9 | Kruppel-like factor 9 [Source:MGI Symbol;Acc:MGI:1333856] | 2353,1 | 100,00 |
| ENSMUSG00000021794 | Glud1 | glutamate dehydrogenase 1 [Source:MGI Symbol;Acc:MGI:95753] | 2340,0 | 99,42 |
| ENSMUSG00000029462 | Vps29 | vacuolar protein sorting 29 (S. pombe) [Source:MGI Symbol;Acc:MGI:1928344] | 2333,8 | 91,66 |
| ENSMUSG00000038502 | Ptov1 | prostate tumor over expressed gene 1 [Source:MGI Symbol;Acc:MGI:1933946] | 2320,4 | 97,59 |
| ENSMUSG00000022476 | Polr3h | polymerase (RNA) III (DNA directed) polypeptide H [Source:MGI Symbol;Acc:MGI:1926179] | 2319,6 | 100,00 |
| ENSMUSG00000000567 | Sox9 | SRY (sex determining region Y)-box 9 [Source:MGI Symbol;Acc:MGI:98371] | 2319,3 | 99,78 |
| ENSMUSG00000033938 | Ndufb7 | NADH dehydrogenase (ubiquinone) 1 beta subcomplex, 7 [Source:MGI Symbol;Acc:MGI:1914166] | 2314,6 | 90,96 |
| ENSMUSG00000026878 | Rab14 | RAB14, member RAS oncogene family [Source:MGI Symbol;Acc:MGI:1915615] | 2302,5 | 71,02 |
| ENSMUSG00000072704 | Smim10l1 | small integral membrane protein 10 like 1 [Source:MGI Symbol;Acc:MGI:1914379] | 2296,4 | 89,39 |
| ENSMUSG00000021546 | Hnrnpk | heterogeneous nuclear ribonucleoprotein K [Source:MGI Symbol;Acc:MGI:99894] | 2294,4 | 82,24 |
| ENSMUSG00000001472 | Tcf25 | transcription factor 25 (basic helix-loop-helix) [Source:MGI Symbol;Acc:MGI:1914105] | 2289,5 | 97,72 |
| ENSMUSG00000081534 | Slc48a1 | solute carrier family 48 (heme transporter), member 1 [Source:MGI Symbol;Acc:MGI:1914989] | 2285,5 | 99,42 |
| ENSMUSG00000038059 | Smim3 | small integral membrane protein 3 [Source:MGI Symbol;Acc:MGI:1917088] | 2280,3 | 97,63 |
| ENSMUSG00000063229 | Ldha | lactate dehydrogenase A [Source:MGI Symbol;Acc:MGI:96759] | 2272,0 | 77,85 |
| ENSMUSG00000022354 | Ndufb9 | NADH dehydrogenase (ubiquinone) 1 beta subcomplex, 9 [Source:MGI Symbol;Acc:MGI:1913468] | 2264,9 | 100,00 |
| ENSMUSG00000050856 | Atp5k | ATP synthase, H+ transporting, mitochondrial F1F0 complex, subunit E [Source:MGI Symbol;Acc:MGI:10663 | 2264,1 | 96,32 |
| ENSMUSG00000018677 | Slc25a39 | solute carrier family 25, member 39 [Source:MGI Symbol;Acc:MGI:1196386] | 2262,5 | 87,52 |
| ENSMUSG00000051853 | Arf3 | ADP-ribosylation factor 3 [Source:MGI Symbol;Acc:MGI:99432] | 2259,2 | 94,79 |
| ENSMUSG00000026932 | Nacc2 | :leus accumbens associated 2, BEN and BTB (POZ) domain containing [Source:MGI Symbol;Acc:MGI:19152 | 2251,8 | 98,78 |
| ENSMUSG00000021771 | Vdac2 | voltage-dependent anion channel 2 [Source:MGI Symbol;Acc:MGI:106915] | 2251,0 | 74,71 |
| ENSMUSG00000022679 | Mpv17l | Mpv17 transgene, kidney disease mutant-like [Source:MGI Symbol;Acc:MGI:2135951] | 2246,5 | 93,29 |
| ENSMUSG00000038965 | Ube2l3 | ubiquitin-conjugating enzyme E2L 3 [Source:MGI Symbol;Acc:MGI:109240] | 2244,9 | 83,31 |
| ENSMUSG00000026202 | Tuba4a | tubulin, alpha 4A [Source:MGI Symbol;Acc:MGI:1095410] | 2241,0 | 86,64 |
| ENSMUSG00000034424 | Gcsh | glycine cleavage system protein H (aminomethyl carrier) [Source:MGI Symbol;Acc:MGI:1915383] | 2236,3 | 96,95 |
| ENSMUSG00000016194 | Hsd11b1 | hydroxysteroid 11-beta dehydrogenase 1 [Source:MGI Symbol;Acc:MGI:103562] | 2231,5 | 77,69 |
| ENSMUSG00000031751 | Amfr | autocrine motility factor receptor [Source:MGI Symbol;Acc:MGI:1345634] | 2231,1 | 99,09 |
| ENSMUSG00000066456 | Hmgn3 | high mobility group nucleosomal binding domain 3 [Source:MGI Symbol;Acc:MGI:2138069] | 2229,7 | 100,00 |
| ENSMUSG00000060961 | Slc4a4 | solute carrier family 4 (anion exchanger), member 4 [Source:MGI Symbol;Acc:MGI:1927555] | 2229,7 | 75,73 |
| ENSMUSG00000007815 | Rhoa | ras homolog gene family, member A [Source:MGI Symbol;Acc:MGI:1096342] | 2229,5 | 89,57 |
| ENSMUSG00000009549 | Srp14 | signal recognition particle 14 [Source:MGI Symbol;Acc:MGI:107169] | 2226,4 | 76,92 |
| ENSMUSG00000030788 | Rnf141 | ring finger protein 141 [Source:MGI Symbol;Acc:MGI:1914400] | 2223,1 | 91,46 |
| ENSMUSG00000032314 | Etfa | electron transferring flavoprotein, alpha polypeptide [Source:MGI Symbol;Acc:MGI:106092] | 2222,0 | 98,37 |
| ENSMUSG00000032399 | Rpl4 | ribosomal protein L4 [Source:MGI Symbol;Acc:MGI:1915141] | 2220,7 | 100,00 |
| ENSMUSG00000025006 | Sorbs1 | sorbin and SH3 domain containing 1 [Source:MGI Symbol;Acc:MGI:700014] | 2218,6 | 98,91 |
| ENSMUSG00000036181 | Hist1h1c | histone cluster 1, H1c [Source:MGI Symbol;Acc:MGI:1931526] | 2218,0 | 99,27 |
| ENSMUSG00000079658 | Tceb1 | transcription elongation factor B (SIII), polypeptide 1 [Source:MGI Symbol;Acc:MGI:1915173] | 2203,9 | 92,15 |
| ENSMUSG00000016319 | Slc25a5 | ier family 25 (mitochondrial carrier, adenine nucleotide translocator), member 5 [Source:MGI Symbol;Acc:MGI | 2202,9 | 99,27 |
| ENSMUSG00000024558 | Mapk4 | mitogen-activated protein kinase 4 [Source:MGI Symbol;Acc:MGI:2444559] | 2190,1 | 99,62 |
| ENSMUSG00000022400 | Rbx1 | ring-box 1 [Source:MGI Symbol;Acc:MGI:1891829] | 2181,6 | 99,17 |
| ENSMUSG00000054252 | Fgfr3 | fibroblast growth factor receptor 3 [Source:MGI Symbol;Acc:MGI:95524] | 2177,6 | 95,18 |
| ENSMUSG00000001211 | Agpat3 | 1-acylglycerol-3-phosphate O-acyltransferase 3 [Source:MGI Symbol;Acc:MGI:1336186] | 2173,4 | 88,24 |
| ENSMUSG00000060992 | Copz1 | coatomer protein complex, subunit zeta 1 [Source:MGI Symbol;Acc:MGI:1929063] | 2168,6 | 100,00 |
| ENSMUSG00000026576 | Atp1b1 | ATPase, Na+/K+ transporting, beta 1 polypeptide [Source:MGI Symbol;Acc:MGI:88108] | 2167,9 | 93,70 |
| ENSMUSG00000036257 | Pnpla8 | patatin-like phospholipase domain containing 8 [Source:MGI Symbol;Acc:MGI:1914702] | 2165,1 | 82,75 |
| ENSMUSG00000035674 | Ndufa3 | NADH dehydrogenase (ubiquinone) 1 alpha subcomplex, 3 [Source:MGI Symbol;Acc:MGI:1913341] | 2162,8 | 97,39 |
| ENSMUSG00000015092 | Edf1 | endothelial differentiation-related factor 1 [Source:MGI Symbol;Acc:MGI:1891227] | 2160,2 | 100,00 |
| ENSMUSG00000042312 | S100a13 | S100 calcium binding protein A13 [Source:MGI Symbol;Acc:MGI:109581] | 2158,7 | 100,00 |
| ENSMUSG00000020462 | Cfap36 | cilia and flagella associated protein 36 [Source:MGI Symbol;Acc:MGI:1913994] | 2157,6 | 79,59 |
| ENSMUSG00000057103 | Cml1 | camello-like 1 [Source:MGI Symbol;Acc:MGI:1913366] | 2150,2 | 95,74 |
| ENSMUSG00000017009 | Sdc4 | syndecan 4 [Source:MGI Symbol;Acc:MGI:1349164] | 2148,1 | 100,00 |
| ENSMUSG00000020265 | Sumo3 | small ubiquitin-like modifier 3 [Source:MGI Symbol;Acc:MGI:1336201] | 2140,1 | 93,68 |
| ENSMUSG00000028015 | Ctso | cathepsin O [Source:MGI Symbol;Acc:MGI:2139628] | 2137,8 | 86,01 |
| ENSMUSG00000006782 | Cnp | 2',3'-cyclic nucleotide 3' phosphodiesterase [Source:MGI Symbol;Acc:MGI:88437] | 2135,2 | 98,85 |
| ENSMUSG00000054452 | Aes | amino-terminal enhancer of split [Source:MGI Symbol;Acc:MGI:95806] | 2127,6 | 99,98 |
| ENSMUSG00000042271 | Nxt2 | nuclear transport factor 2-like export factor 2 [Source:MGI Symbol;Acc:MGI:2147914] | 2123,3 | 91,81 |
| ENSMUSG00000072772 | Grcc10 | gene rich cluster, C10 gene [Source:MGI Symbol;Acc:MGI:1315201] | 2108,5 | 91,85 |
| ENSMUSG00000061518 | Cox5b | cytochrome c oxidase subunit Vb [Source:MGI Symbol;Acc:MGI:88475] | 2101,9 | 98,15 |
| ENSMUSG00000024132 | Eci1 | enoyl-Coenzyme A delta isomerase 1 [Source:MGI Symbol;Acc:MGI:94871] | 2094,5 | 100,00 |
| ENSMUSG00000031604 | Msmo1 | methylsterol monoxygenase 1 [Source:MGI Symbol;Acc:MGI:1913484] | 2088,3 | 87,03 |
| ENSMUSG00000071528 | Usmg5 | upregulated during skeletal muscle growth 5 [Source:MGI Symbol;Acc:MGI:1891435] | 2087,8 | 100,00 |
| ENSMUSG00000002416 | Ndufb2 | NADH dehydrogenase (ubiquinone) 1 beta subcomplex, 2 [Source:MGI Symbol;Acc:MGI:1915448] | 2086,0 | 80,87 |
| ENSMUSG00000019054 | Fis1 | fission, mitochondrial 1 [Source:MGI Symbol;Acc:MGI:1913687] | 2080,4 | 100,00 |
| ENSMUSG00000074093 | Svip | small VCP/p97-interacting protein [Source:MGI Symbol;Acc:MGI:1922994] | 2078,9 | 92,39 |
| ENSMUSG00000038936 | Sccpdh | saccharopine dehydrogenase (putative) [Source:MGI Symbol;Acc:MGI:1924486] | 2078,2 | 92,71 |
| ENSMUSG00000040957 | Cables1 | CDK5 and Abl enzyme substrate 1 [Source:MGI Symbol;Acc:MGI:1927065] | 2071,3 | 100,00 |
| ENSMUSG00000027828 | Ssr3 | signal sequence receptor, gamma [Source:MGI Symbol;Acc:MGI:1914687] | 2067,5 | 97,59 |
| ENSMUSG00000021371 | Mcur1 | mitochondrial calcium uniporter regulator 1 [Source:MGI Symbol;Acc:MGI:1923387] | 2060,9 | 99,98 |
| ENSMUSG00000025350 | Rdh5 | retinol dehydrogenase 5 [Source:MGI Symbol;Acc:MGI:1201412] | 2060,1 | 96,57 |
| ENSMUSG00000020864 | Ankrd40 | ankyrin repeat domain 40 [Source:MGI Symbol;Acc:MGI:1918702] | 2054,5 | 86,86 |
| ENSMUSG00000028128 | F3 | coagulation factor III [Source:MGI Symbol;Acc:MGI:88381] | 2050,6 | 100,00 |
| ENSMUSG00000062981 | Mrpl42 | mitochondrial ribosomal protein L42 [Source:MGI Symbol;Acc:MGI:1333774] | 2047,5 | 98,85 |
| ENSMUSG00000040731 | Eif4h | eukaryotic translation initiation factor 4H [Source:MGI Symbol;Acc:MGI:1341822] | 2046,4 | 90,98 |
| ENSMUSG00000014769 | Psmb1 | proteasome (prosome, macropain) subunit, beta type 1 [Source:MGI Symbol;Acc:MGI:104884] | 2043,2 | 100,00 |
| ENSMUSG00000063358 | Mapk1 | mitogen-activated protein kinase 1 [Source:MGI Symbol;Acc:MGI:1346858] | 2034,3 | 98,74 |
| ENSMUSG00000027533 | Fabp5 | fatty acid binding protein 5, epidermal [Source:MGI Symbol;Acc:MGI:101790] | 2029,8 | 91,02 |
| ENSMUSG00000026227 | 2810459M11Rik | RIKEN cDNA 2810459M11 gene [Source:MGI Symbol;Acc:MGI:1920042] | 2028,4 | 99,77 |
| ENSMUSG00000004897 | Hdgf | hepatoma-derived growth factor [Source:MGI Symbol;Acc:MGI:1194494] | 2025,7 | 99,18 |
| ENSMUSG00000050608 | Minos1 | mitochondrial inner membrane organizing system 1 [Source:MGI Symbol;Acc:MGI:1913628] | 2024,5 | 98,80 |

| | | | | |
|---|---|---|---|---|
| ENSMUSG00000006304 | Arpc2 | actin related protein 2/3 complex, subunit 2 [Source:MGI Symbol;Acc:MGI:1923959] | 2023,1 | 81,55 |
| ENSMUSG00000001700 | Gramd3 | GRAM domain containing 3 [Source:MGI Symbol;Acc:MGI:1914815] | 2022,6 | 99,99 |
| ENSMUSG00000051223 | Bzw1 | basic leucine zipper and W2 domains 1 [Source:MGI Symbol;Acc:MGI:1914132] | 2022,5 | 90,23 |
| ENSMUSG00000060002 | Chpt1 | choline phosphotransferase 1 [Source:MGI Symbol;Acc:MGI:2384841] | 2008,5 | 92,11 |
| ENSMUSG00000027712 | Anxa5 | annexin A5 [Source:MGI Symbol;Acc:MGI:106008] | 2006,0 | 99,11 |
| ENSMUSG00000000058 | Cav2 | caveolin 2 [Source:MGI Symbol;Acc:MGI:107571] | 2004,6 | 86,19 |
| ENSMUSG00000036427 | Gpi1 | glucose phosphate isomerase 1 [Source:MGI Symbol;Acc:MGI:95797] | 1995,1 | 95,42 |
| ENSMUSG00000020390 | Ube2b | ubiquitin-conjugating enzyme E2B [Source:MGI Symbol;Acc:MGI:102944] | 1994,5 | 99,82 |
| ENSMUSG00000022450 | Ndufa6 | NADH dehydrogenase (ubiquinone) 1 alpha subcomplex, 6 (B14) [Source:MGI Symbol;Acc:MGI:1914380] | 1993,8 | 100,00 |
| ENSMUSG00000060802 | B2m | beta-2 microglobulin [Source:MGI Symbol;Acc:MGI:88127] | 1990,0 | 100,00 |
| ENSMUSG00000040681 | Hmgn1 | high mobility group nucleosomal binding domain 1 [Source:MGI Symbol;Acc:MGI:96120] | 1988,9 | 75,05 |
| ENSMUSG00000027162 | Lin7c | lin-7 homolog C (C. elegans) [Source:MGI Symbol;Acc:MGI:1330839] | 1988,0 | 99,91 |
| ENSMUSG00000043162 | Pyurf | Pigy upstream reading frame [Source:MGI Symbol;Acc:MGI:1913709] | 1987,8 | 99,92 |
| ENSMUSG00000029221 | Slc30a9 | solute carrier family 30 (zinc transporter), member 9 [Source:MGI Symbol;Acc:MGI:1923690] | 1987,6 | 86,07 |
| ENSMUSG00000044627 | Swi5 | SWI5 recombination repair homolog (yeast) [Source:MGI Symbol;Acc:MGI:1920181] | 1987,3 | 81,49 |
| ENSMUSG00000031021 | Tmem9b | TMEM9 domain family, member B [Source:MGI Symbol;Acc:MGI:1915254] | 1984,0 | 99,87 |
| ENSMUSG00000028785 | Hpca | hippocalcin [Source:MGI Symbol;Acc:MGI:1336200] | 1976,6 | 96,57 |
| ENSMUSG00000090247 | Bloc1s1 | biogenesis of lysosomal organelles complex-1, subunit 1 [Source:MGI Symbol;Acc:MGI:1195276] | 1974,3 | 88,20 |
| ENSMUSG00000091228 | Gm20390 | predicted gene 20390 [Source:MGI Symbol;Acc:MGI:5141855] | 1971,9 | 98,59 |
| ENSMUSG00000030105 | Arl8b | ADP-ribosylation factor-like 8B [Source:MGI Symbol;Acc:MGI:1914416] | 1967,1 | 85,15 |
| ENSMUSG00000025978 | Rftn2 | raftlin family member 2 [Source:MGI Symbol;Acc:MGI:1921263] | 1965,6 | 87,43 |
| ENSMUSG00000024997 | Prdx3 | peroxiredoxin 3 [Source:MGI Symbol;Acc:MGI:88034] | 1963,7 | 99,05 |
| ENSMUSG00000026568 | Mpc2 | mitochondrial pyruvate carrier 2 [Source:MGI Symbol;Acc:MGI:1917706] | 1961,9 | 99,77 |
| ENSMUSG00000044252 | Osbpl1a | oxysterol binding protein-like 1A [Source:MGI Symbol;Acc:MGI:1927551] | 1959,8 | 77,30 |
| ENSMUSG00000015291 | Gdi1 | guanosine diphosphate (GDP) dissociation inhibitor 1 [Source:MGI Symbol;Acc:MGI:99846] | 1956,7 | 91,21 |
| ENSMUSG00000028221 | Tmem55a | transmembrane protein 55A [Source:MGI Symbol;Acc:MGI:1919769] | 1956,7 | 95,77 |
| ENSMUSG00000034353 | Ramp1 | receptor (calcitonin) activity modifying protein 1 [Source:MGI Symbol;Acc:MGI:1858418] | 1948,0 | 85,97 |
| ENSMUSG00000020949 | Fkbp3 | FK506 binding protein 3 [Source:MGI Symbol;Acc:MGI:1353460] | 1939,9 | 100,00 |
| ENSMUSG00000023944 | Hsp90ab1 | heat shock protein 90 alpha (cytosolic), class B member 1 [Source:MGI Symbol;Acc:MGI:96247] | 1938,0 | 98,17 |
| ENSMUSG00000005732 | Ranbp1 | RAN binding protein 1 [Source:MGI Symbol;Acc:MGI:96269] | 1933,8 | 94,76 |
| ENSMUSG00000020777 | Acox1 | acyl-Coenzyme A oxidase 1, palmitoyl [Source:MGI Symbol;Acc:MGI:1330812] | 1928,5 | 99,13 |
| ENSMUSG00000031808 | Slc27a1 | solute carrier family 27 (fatty acid transporter), member 1 [Source:MGI Symbol;Acc:MGI:1347098] | 1927,4 | 100,00 |
| ENSMUSG00000027303 | Ptpra | protein tyrosine phosphatase, receptor type, A [Source:MGI Symbol;Acc:MGI:97808] | 1923,7 | 96,76 |
| ENSMUSG00000013593 | Ndufs2 | NADH dehydrogenase (ubiquinone) Fe-S protein 2 [Source:MGI Symbol;Acc:MGI:2385112] | 1921,5 | 93,63 |
| ENSMUSG00000043866 | Taf10 | 10 RNA polymerase II, TATA box binding protein (TBP)-associated factor [Source:MGI Symbol;Acc:MGI:1346 | 1918,8 | 94,01 |
| ENSMUSG00000006941 | Eif1b | eukaryotic translation initiation factor 1B [Source:MGI Symbol;Acc:MGI:1916219] | 1917,5 | 100,00 |
| ENSMUSG00000004096 | Cwc15 | CWC15 spliceosome-associated protein [Source:MGI Symbol;Acc:MGI:1913320] | 1917,0 | 100,00 |
| ENSMUSG00000015341 | Golga7 | golgi autoantigen, golgin subfamily a, 7 [Source:MGI Symbol;Acc:MGI:1931029] | 1914,3 | 86,52 |
| ENSMUSG00000032067 | Pts | 6-pyruvoyl-tetrahydropterin synthase [Source:MGI Symbol;Acc:MGI:1338783] | 1913,1 | 100,00 |
| ENSMUSG00000036850 | Mrpl41 | mitochondrial ribosomal protein L41 [Source:MGI Symbol;Acc:MGI:1333816] | 1910,1 | 100,00 |
| ENSMUSG00000006818 | Sod2 | superoxide dismutase 2, mitochondrial [Source:MGI Symbol;Acc:MGI:98352] | 1904,7 | 84,01 |
| ENSMUSG00000018293 | Pfn1 | profilin 1 [Source:MGI Symbol;Acc:MGI:97549] | 1904,7 | 90,16 |
| ENSMUSG00000037503 | Fam168b | family with sequence similarity 168, member B [Source:MGI Symbol;Acc:MGI:2448487] | 1901,9 | 99,04 |
| ENSMUSG00000058672 | Tubb2a | tubulin, beta 2A class IIA [Source:MGI Symbol;Acc:MGI:107861] | 1899,0 | 77,27 |
| ENSMUSG00000044408 | Sptssa | serine palmitoyltransferase, small subunit A [Source:MGI Symbol;Acc:MGI:1913399] | 1898,0 | 100,00 |
| ENSMUSG00000037062 | Sh3glb1 | SH3-domain GRB2-like B1 (endophilin) [Source:MGI Symbol;Acc:MGI:1859730] | 1897,5 | 95,37 |
| ENSMUSG00000052397 | Ezr | ezrin [Source:MGI Symbol;Acc:MGI:98931] | 1881,3 | 92,84 |
| ENSMUSG00000027797 | Dclk1 | doublecortin-like kinase 1 [Source:MGI Symbol;Acc:MGI:1330861] | 1881,1 | 94,18 |
| ENSMUSG00000031610 | Scrg1 | scrapie responsive gene 1 [Source:MGI Symbol;Acc:MGI:1328308] | 1880,2 | 100,00 |
| ENSMUSG00000031667 | Aktip | thymoma viral proto-oncogene 1 interacting protein [Source:MGI Symbol;Acc:MGI:3693832] | 1879,7 | 90,94 |
| ENSMUSG00000020263 | Appl2 | otein, phosphotyrosine interaction, PH domain and leucine zipper containing 2 [Source:MGI Symbol;Acc:MGI | 1877,4 | 98,48 |
| ENSMUSG00000021417 | Eci2 | enoyl-Coenzyme A delta isomerase 2 [Source:MGI Symbol;Acc:MGI:1346064] | 1873,8 | 93,12 |
| ENSMUSG00000037373 | Ctbp1 | C-terminal binding protein 1 [Source:MGI Symbol;Acc:MGI:1201685] | 1872,7 | 73,65 |
| ENSMUSG00000084786 | Ubl5 | ubiquitin-like 5 [Source:MGI Symbol;Acc:MGI:1913427] | 1868,5 | 75,45 |
| ENSMUSG00000024608 | Rps14 | ribosomal protein S14 [Source:MGI Symbol;Acc:MGI:98107] | 1867,0 | 78,66 |
| ENSMUSG00000055839 | Tceb2 | transcription elongation factor B (SIII), polypeptide 2 [Source:MGI Symbol;Acc:MGI:1914923] | 1866,5 | 99,93 |
| ENSMUSG00000053907 | Mat2a | methionine adenosyltransferase II, alpha [Source:MGI Symbol;Acc:MGI:2443731] | 1862,5 | 97,55 |
| ENSMUSG00000022122 | Ednrb | endothelin receptor type B [Source:MGI Symbol;Acc:MGI:102720] | 1861,1 | 100,00 |
| ENSMUSG00000025509 | Pnpla2 | patatin-like phospholipase domain containing 2 [Source:MGI Symbol;Acc:MGI:1914103] | 1856,8 | 85,24 |
| ENSMUSG00000033917 | Gde1 | glycerophosphodiester phosphodiesterase 1 [Source:MGI Symbol;Acc:MGI:1891827] | 1855,1 | 96,85 |
| ENSMUSG00000026426 | Arl8a | ADP-ribosylation factor-like 8A [Source:MGI Symbol;Acc:MGI:1915974] | 1849,8 | 99,72 |
| ENSMUSG00000060450 | Rnf14 | ring finger protein 14 [Source:MGI Symbol;Acc:MGI:1929668] | 1849,7 | 99,11 |
| ENSMUSG00000046727 | Cystm1 | cysteine-rich transmembrane module containing 1 [Source:MGI Symbol;Acc:MGI:1913310] | 1848,8 | 75,87 |
| ENSMUSG00000038607 | Gng10 | guanine nucleotide binding protein (G protein), gamma 10 [Source:MGI Symbol;Acc:MGI:1336169] | 1845,4 | 96,90 |
| ENSMUSG00000036885 | Arhgef26 | Rho guanine nucleotide exchange factor (GEF) 26 [Source:MGI Symbol;Acc:MGI:1918053] | 1841,6 | 99,21 |
| ENSMUSG00000059734 | Ndufs8 | NADH dehydrogenase (ubiquinone) Fe-S protein 8 [Source:MGI Symbol;Acc:MGI:2385079] | 1838,4 | 100,00 |
| ENSMUSG00000030591 | Psmd8 | proteasome (prosome, macropain) 26S subunit, non-ATPase, 8 [Source:MGI Symbol;Acc:MGI:1888669] | 1838,3 | 76,85 |
| ENSMUSG00000032349 | Elovl5 | ELOVL family member 5, elongation of long chain fatty acids (yeast) [Source:MGI Symbol;Acc:MGI:1916051] | 1837,7 | 94,03 |
| ENSMUSG00000067194 | Eif1ax | eukaryotic translation initiation factor 1A, X-linked [Source:MGI Symbol;Acc:MGI:1913485] | 1833,6 | 94,26 |
| ENSMUSG00000003955 | Fam162a | family with sequence similarity 162, member A [Source:MGI Symbol;Acc:MGI:1917436] | 1830,4 | 100,00 |
| ENSMUSG00000021474 | Sfxn1 | sideroflexin 1 [Source:MGI Symbol;Acc:MGI:2137677] | 1830,1 | 99,95 |
| ENSMUSG00000016946 | Kctd5 | potassium channel tetramerisation domain containing 5 [Source:MGI Symbol;Acc:MGI:1916509] | 1829,7 | 99,65 |
| ENSMUSG00000028964 | Park7 | Parkinson disease (autosomal recessive, early onset) 7 [Source:MGI Symbol;Acc:MGI:2135637] | 1827,8 | 86,10 |
| ENSMUSG00000074748 | Atxn7l3b | ataxin 7-like 3B [Source:MGI Symbol;Acc:MGI:1914971] | 1822,4 | 99,91 |
| ENSMUSG00000026003 | Acadl | acyl-Coenzyme A dehydrogenase, long-chain [Source:MGI Symbol;Acc:MGI:87866] | 1822,0 | 99,74 |
| ENSMUSG00000021576 | Pdcd6 | programmed cell death 6 [Source:MGI Symbol;Acc:MGI:109283] | 1820,4 | 90,09 |
| ENSMUSG00000028124 | Gclm | glutamate-cysteine ligase, modifier subunit [Source:MGI Symbol;Acc:MGI:104995] | 1817,9 | 88,57 |
| ENSMUSG00000023010 | Tmbim6 | transmembrane BAX inhibitor motif containing 6 [Source:MGI Symbol;Acc:MGI:99682] | 1816,0 | 71,44 |
| ENSMUSG00000027406 | Idh3b | isocitrate dehydrogenase 3 (NAD+) beta [Source:MGI Symbol;Acc:MGI:2158650] | 1815,6 | 98,66 |
| ENSMUSG00000002459 | Rgs20 | regulator of G-protein signaling 20 [Source:MGI Symbol;Acc:MGI:1929866] | 1815,1 | 95,53 |
| ENSMUSG00000019998 | Stx7 | syntaxin 7 [Source:MGI Symbol;Acc:MGI:1858210] | 1811,1 | 92,43 |
| ENSMUSG00000013921 | Clip3 | CAP-GLY domain containing linker protein 3 [Source:MGI Symbol;Acc:MGI:1923936] | 1809,4 | 99,90 |
| ENSMUSG00000030541 | Idh2 | isocitrate dehydrogenase 2 (NADP+), mitochondrial [Source:MGI Symbol;Acc:MGI:96414] | 1809,3 | 98,19 |
| ENSMUSG00000092345 | Gm20503 | predicted gene 20503 [Source:MGI Symbol;Acc:MGI:5141968] | 1798,7 | 98,48 |
| ENSMUSG00000024026 | Glo1 | glyoxalase 1 [Source:MGI Symbol;Acc:MGI:95742] | 1795,9 | 75,37 |
| ENSMUSG00000031065 | Cdk16 | cyclin-dependent kinase 16 [Source:MGI Symbol;Acc:MGI:97516] | 1792,0 | 89,22 |
| ENSMUSG00000021124 | Vti1b | vesicle transport through interaction with t-SNAREs 1B [Source:MGI Symbol;Acc:MGI:1855688] | 1790,5 | 98,81 |
| ENSMUSG00000027282 | Mtch2 | mitochondrial carrier homolog 2 (C. elegans) [Source:MGI Symbol;Acc:MGI:1929260] | 1785,0 | 95,23 |
| ENSMUSG00000009291 | Pttg1ip | pituitary tumor-transforming 1 interacting protein [Source:MGI Symbol;Acc:MGI:2652132] | 1784,4 | 92,41 |
| ENSMUSG00000026554 | Dcaf8 | DDB1 and CUL4 associated factor 8 [Source:MGI Symbol;Acc:MGI:91860] | 1780,7 | 83,30 |
| ENSMUSG00000025825 | Iscu | iron-sulfur cluster assembly enzyme [Source:MGI Symbol;Acc:MGI:1913633] | 1780,5 | 94,11 |
| ENSMUSG00000028837 | Psmb2 | proteasome (prosome, macropain) subunit, beta type 2 [Source:MGI Symbol;Acc:MGI:1347045] | 1780,2 | 100,00 |
| ENSMUSG00000036503 | Rnf13 | ring finger protein 13 [Source:MGI Symbol;Acc:MGI:1346341] | 1778,8 | 76,85 |
| ENSMUSG00000030224 | Strap | serine/threonine kinase receptor associated protein [Source:MGI Symbol;Acc:MGI:1329037] | 1771,9 | 98,96 |
| ENSMUSG00000024576 | Csnk1a1 | casein kinase 1, alpha 1 [Source:MGI Symbol;Acc:MGI:1934950] | 1767,2 | 78,89 |
| ENSMUSG00000015484 | Fam163a | family with sequence similarity 163, member A [Source:MGI Symbol;Acc:MGI:3618859] | 1766,4 | 100,00 |
| ENSMUSG00000067279 | Ppp1r3c | protein phosphatase 1, regulatory (inhibitor) subunit 3C [Source:MGI Symbol;Acc:MGI:1858229] | 1764,1 | 100,00 |
| ENSMUSG00000013275 | Slc41a1 | solute carrier family 41, member 1 [Source:MGI Symbol;Acc:MGI:2444823] | 1761,8 | 99,16 |
| ENSMUSG00000053768 | Chchd3 | coiled-coil-helix-coiled-coil-helix domain containing 3 [Source:MGI Symbol;Acc:MGI:1913325] | 1761,2 | 91,00 |
| ENSMUSG00000039347 | Atp6v0e2 | ATPase, H+ transporting, lysosomal V0 subunit E2 [Source:MGI Symbol;Acc:MGI:1923502] | 1761,1 | 99,98 |
| ENSMUSG00000004610 | Etfb | electron transferring flavoprotein, beta polypeptide [Source:MGI Symbol;Acc:MGI:106098] | 1750,1 | 98,10 |
| ENSMUSG00000026277 | Stk25 | serine/threonine kinase 25 (yeast) [Source:MGI Symbol;Acc:MGI:1891699] | 1748,3 | 83,39 |
| ENSMUSG00000024146 | Cript | cysteine-rich PDZ-binding protein [Source:MGI Symbol;Acc:MGI:1929655] | 1744,1 | 99,75 |
| ENSMUSG00000053898 | Ech1 | enoyl coenzyme A hydratase 1, peroxisomal [Source:MGI Symbol;Acc:MGI:1858208] | 1743,8 | 96,69 |
| ENSMUSG00000021218 | Gdi2 | guanosine diphosphate (GDP) dissociation inhibitor 2 [Source:MGI Symbol;Acc:MGI:99845] | 1741,7 | 97,02 |
| ENSMUSG00000027479 | Mapre1 | microtubule-associated protein, RP/EB family, member 1 [Source:MGI Symbol;Acc:MGI:891995] | 1737,3 | 82,72 |
| ENSMUSG00000021196 | Pfkp | phosphofructokinase, platelet [Source:MGI Symbol;Acc:MGI:1891833] | 1736,7 | 94,43 |
| ENSMUSG00000020271 | Fbxw11 | F-box and WD-40 domain protein 11 [Source:MGI Symbol;Acc:MGI:2144023] | 1735,2 | 98,50 |
| ENSMUSG00000025868 | Higd2a | HIG1 domain family, member 2A [Source:MGI Symbol;Acc:MGI:1914294] | 1734,5 | 99,38 |
| ENSMUSG00000015671 | Psma2 | proteasome (prosome, macropain) subunit, alpha type 2 [Source:MGI Symbol;Acc:MGI:104885] | 1733,4 | 100,00 |
| ENSMUSG00000038803 | Ost4 | oligosaccharyltransferase 4 homolog (S. cerevisiae) [Source:MGI Symbol;Acc:MGI:1914945] | 1732,6 | 96,96 |
| ENSMUSG00000027452 | Acss1 | acyl-CoA synthetase short-chain family member 1 [Source:MGI Symbol;Acc:MGI:1915988] | 1731,6 | 99,76 |
| ENSMUSG00000002808 | Epdr1 | ependymin related protein 1 (zebrafish) [Source:MGI Symbol;Acc:MGI:2145369] | 1724,2 | 99,61 |
| ENSMUSG00000035242 | Oaz1 | ornithine decarboxylase antizyme 1 [Source:MGI Symbol;Acc:MGI:109433] | 1721,7 | 72,03 |
| ENSMUSG00000020859 | Spag9 | sperm associated antigen 9 [Source:MGI Symbol;Acc:MGI:1918084] | 1721,5 | 98,38 |
| ENSMUSG00000031983 | 2310022B05Rik | RIKEN cDNA 2310022B05 gene [Source:MGI Symbol;Acc:MGI:1916801] | 1721,0 | 97,57 |
| ENSMUSG00000027566 | Psma7 | proteasome (prosome, macropain) subunit, alpha type 7 [Source:MGI Symbol;Acc:MGI:1347070] | 1712,9 | 100,00 |

| | | | | |
|---|---|---|---|---|
| ENSMUSG00000107283 | Mpv17 | MpV17 mitochondrial inner membrane protein [Source:MGI Symbol;Acc:MGI:97138] | 1711,7 | 96,04 |
| ENSMUSG00000023089 | Ndufa5 | NADH dehydrogenase (ubiquinone) 1 alpha subcomplex, 5 [Source:MGI Symbol;Acc:MGI:1915452] | 1710,7 | 81,33 |
| ENSMUSG00000034853 | Acot11 | acyl-CoA thioesterase 11 [Source:MGI Symbol;Acc:MGI:1913736] | 1710,1 | 95,22 |
| ENSMUSG00000036299 | BC031181 | cDNA sequence BC031181 [Source:MGI Symbol;Acc:MGI:3039614] | 1709,7 | 98,09 |
| ENSMUSG00000039615 | Stub1 | STIP1 homology and U-Box containing protein 1 [Source:MGI Symbol;Acc:MGI:1891731] | 1703,0 | 99,34 |
| ENSMUSG00000029247 | Paics | zole carboxylase, phosphoribosylaminoribosylaminoimidazole, succinocarboxamide synthetase [Source:MGI S | 1701,5 | 98,50 |
| ENSMUSG00000004771 | Rab11a | RAB11A, member RAS oncogene family [Source:MGI Symbol;Acc:MGI:1858202] | 1701,2 | 97,88 |
| ENSMUSG00000002580 | Mien1 | migration and invasion enhancer 1 [Source:MGI Symbol;Acc:MGI:1913678] | 1701,1 | 99,51 |
| ENSMUSG00000040147 | Maob | monoamine oxidase B [Source:MGI Symbol;Acc:MGI:96916] | 1700,1 | 100,00 |
| ENSMUSG00000025040 | Fundc1 | FUN14 domain containing 1 [Source:MGI Symbol;Acc:MGI:1919268] | 1699,4 | 94,00 |
| ENSMUSG00000028519 | Dab1 | disabled 1 [Source:MGI Symbol;Acc:MGI:108554] | 1697,0 | 95,23 |
| ENSMUSG00000017776 | Crk | v-crk avian sarcoma virus CT10 oncogene homolog [Source:MGI Symbol;Acc:MGI:88508] | 1695,3 | 92,97 |
| ENSMUSG00000026781 | Acbd5 | acyl-Coenzyme A binding domain containing 5 [Source:MGI Symbol;Acc:MGI:1921409] | 1693,4 | 94,94 |
| ENSMUSG00000014226 | Cacybp | calcyclin binding protein [Source:MGI Symbol;Acc:MGI:1270839] | 1690,6 | 93,76 |
| ENSMUSG00000019428 | Fkbp8 | FK506 binding protein 8 [Source:MGI Symbol;Acc:MGI:1341070] | 1689,2 | 87,80 |
| ENSMUSG00000022240 | Ctnnd2 | catenin (cadherin associated protein), delta 2 [Source:MGI Symbol;Acc:MGI:1195966] | 1685,6 | 98,79 |
| ENSMUSG00000031617 | Tmem184c | transmembrane protein 184C [Source:MGI Symbol;Acc:MGI:2384562] | 1683,6 | 100,00 |
| ENSMUSG00000019843 | Fyn | Fyn proto-oncogene [Source:MGI Symbol;Acc:MGI:95602] | 1683,5 | 87,18 |
| ENSMUSG00000033763 | Mtss1l | metastasis suppressor 1-like [Source:MGI Symbol;Acc:MGI:3039591] | 1682,9 | 98,73 |
| ENSMUSG00000039206 | Daglb | diacylglycerol lipase, beta [Source:MGI Symbol;Acc:MGI:2442032] | 1682,0 | 79,67 |
| ENSMUSG00000059974 | Ntm | neurotrimin [Source:MGI Symbol;Acc:MGI:2446259] | 1669,8 | 90,72 |
| ENSMUSG00000020368 | Canx | calnexin [Source:MGI Symbol;Acc:MGI:88261] | 1668,4 | 97,72 |
| ENSMUSG00000022214 | Dcaf11 | DDB1 and CUL4 associated factor 11 [Source:MGI Symbol;Acc:MGI:90168] | 1667,5 | 99,77 |
| ENSMUSG00000036504 | Phpt1 | phosphohistidine phosphatase 1 [Source:MGI Symbol;Acc:MGI:1922704] | 1665,5 | 97,94 |
| ENSMUSG00000007564 | Ppp2r1a | protein phosphatase 2, regulatory subunit A, alpha [Source:MGI Symbol;Acc:MGI:1926334] | 1664,9 | 96,67 |
| ENSMUSG00000006024 | Napa | N-ethylmaleimide sensitive fusion protein attachment protein alpha [Source:MGI Symbol;Acc:MGI:104563] | 1663,1 | 97,17 |
| ENSMUSG00000035104 | Eva1a | eva-1 homolog A (C. elegans) [Source:MGI Symbol;Acc:MGI:2385247] | 1660,5 | 88,96 |
| ENSMUSG00000032470 | Mras | muscle and microspikes RAS [Source:MGI Symbol;Acc:MGI:1100856] | 1652,4 | 95,61 |
| ENSMUSG00000020946 | Gosr2 | golgi SNAP receptor complex member 2 [Source:MGI Symbol;Acc:MGI:1927204] | 1651,2 | 92,89 |
| ENSMUSG00000078202 | Nrarp | Notch-regulated ankyrin repeat protein [Source:MGI Symbol;Acc:MGI:1914372] | 1648,6 | 100,00 |
| ENSMUSG00000027668 | Mfn1 | mitofusin 1 [Source:MGI Symbol;Acc:MGI:1914664] | 1643,4 | 83,49 |
| ENSMUSG00000007950 | Abhd8 | abhydrolase domain containing 8 [Source:MGI Symbol;Acc:MGI:1918946] | 1640,6 | 99,85 |
| ENSMUSG00000022199 | Slc22a17 | solute carrier family 22 (organic cation transporter), member 17 [Source:MGI Symbol;Acc:MGI:1926225] | 1640,2 | 99,71 |
| ENSMUSG00000020022 | Ndufa12 | NADH dehydrogenase (ubiquinone) 1 alpha subcomplex, 12 [Source:MGI Symbol;Acc:MGI:1913664] | 1638,2 | 94,95 |
| ENSMUSG00000026032 | Ndufb3 | NADH dehydrogenase (ubiquinone) 1 beta subcomplex 3 [Source:MGI Symbol;Acc:MGI:1913745] | 1636,9 | 99,48 |
| ENSMUSG00000025035 | Arl3 | ADP-ribosylation factor-like 3 [Source:MGI Symbol;Acc:MGI:1929699] | 1635,7 | 100,00 |
| ENSMUSG00000031708 | Tecr | trans-2,3-enoyl-CoA reductase [Source:MGI Symbol;Acc:MGI:1915408] | 1634,0 | 88,58 |
| ENSMUSG00000030127 | Cops7a | COP9 signalosome subunit 7A [Source:MGI Symbol;Acc:MGI:1349400] | 1630,7 | 98,88 |
| ENSMUSG00000026393 | Nek7 | NIMA (never in mitosis gene a)-related expressed kinase 7 [Source:MGI Symbol;Acc:MGI:1890645] | 1629,3 | 77,95 |
| ENSMUSG00000039016 | Timm8b | translocase of inner mitochondrial membrane 8B [Source:MGI Symbol;Acc:MGI:1353424] | 1629,2 | 100,00 |
| ENSMUSG00000015749 | Anp32e | acidic (leucine-rich) nuclear phosphoprotein 32 family, member E [Source:MGI Symbol;Acc:MGI:1913721] | 1628,6 | 95,05 |
| ENSMUSG00000024962 | Vegfb | vascular endothelial growth factor B [Source:MGI Symbol;Acc:MGI:106199] | 1627,7 | 100,00 |
| ENSMUSG00000028407 | Toporsos | topoisomerase I binding, arginine/serine-rich, opposite strand [Source:MGI Symbol;Acc:MGI:1913684] | 1626,9 | 88,98 |
| ENSMUSG00000031996 | Aplp2 | amyloid beta (A4) precursor-like protein 2 [Source:MGI Symbol;Acc:MGI:88047] | 1625,2 | 99,97 |
| ENSMUSG00000029213 | Commd8 | COMM domain containing 8 [Source:MGI Symbol;Acc:MGI:1343485] | 1619,4 | 99,74 |
| ENSMUSG00000033720 | Sfxn5 | sideroflexin 5 [Source:MGI Symbol;Acc:MGI:2137681] | 1618,3 | 96,41 |
| ENSMUSG00000035901 | Dennd5a | DENN/MADD domain containing 5A [Source:MGI Symbol;Acc:MGI:1201681] | 1616,1 | 97,42 |
| ENSMUSG00000026833 | Olfm1 | olfactomedin 1 [Source:MGI Symbol;Acc:MGI:1860437] | 1615,1 | 99,51 |
| ENSMUSG00000038876 | Rnf146 | ring finger protein 146 [Source:MGI Symbol;Acc:MGI:1915281] | 1614,8 | 90,69 |
| ENSMUSG00000024038 | Ndufv3 | NADH dehydrogenase (ubiquinone) flavoprotein 3 [Source:MGI Symbol;Acc:MGI:1890894] | 1612,0 | 96,03 |
| ENSMUSG00000024899 | Papss2 | 3'-phosphoadenosine 5'-phosphosulfate synthase 2 [Source:MGI Symbol;Acc:MGI:1330223] | 1611,3 | 99,55 |
| ENSMUSG00000026457 | Adipor1 | adiponectin receptor 1 [Source:MGI Symbol;Acc:MGI:1919924] | 1607,0 | 97,42 |
| ENSMUSG00000050708 | Ftl1 | ferritin light polypeptide 1 [Source:MGI Symbol;Acc:MGI:95589] | 1603,8 | 90,71 |
| ENSMUSG00000018849 | Wwc1 | WW, C2 and coiled-coil domain containing 1 [Source:MGI Symbol;Acc:MGI:2388637] | 1602,2 | 99,93 |
| ENSMUSG00000079426 | Arpc4 | actin related protein 2/3 complex, subunit 4 [Source:MGI Symbol;Acc:MGI:1915339] | 1601,3 | 95,25 |
| ENSMUSG00000021242 | Npc2 | Niemann-Pick type C2 [Source:MGI Symbol;Acc:MGI:1915213] | 1598,7 | 99,01 |
| ENSMUSG00000005687 | Bcas2 | breast carcinoma amplified sequence 2 [Source:MGI Symbol;Acc:MGI:1915433] | 1597,4 | 96,80 |
| ENSMUSG00000001100 | Poldip2 | polymerase (DNA-directed), delta interacting protein 2 [Source:MGI Symbol;Acc:MGI:1915061] | 1593,5 | 97,76 |
| ENSMUSG00000026260 | Ndufa10 | NADH dehydrogenase (ubiquinone) 1 alpha subcomplex 10 [Source:MGI Symbol;Acc:MGI:1914523] | 1592,8 | 81,40 |
| ENSMUSG00000070394 | Tmem256 | transmembrane protein 256 [Source:MGI Symbol;Acc:MGI:1916436] | 1590,8 | 100,00 |
| ENSMUSG00000037710 | Cisd1 | CDGSH iron sulfur domain 1 [Source:MGI Symbol;Acc:MGI:1261855] | 1590,2 | 97,02 |
| ENSMUSG00000020889 | Nr1d1 | nuclear receptor subfamily 1, group D, member 1 [Source:MGI Symbol;Acc:MGI:2444210] | 1585,4 | 98,84 |
| ENSMUSG00000021772 | Nkiras1 | NFKB inhibitor interacting Ras-like protein 1 [Source:MGI Symbol;Acc:MGI:1916971] | 1584,6 | 95,76 |
| ENSMUSG00000091337 | Eid1 | EP300 interacting inhibitor of differentiation 1 [Source:MGI Symbol;Acc:MGI:1889651] | 1576,9 | 100,00 |
| ENSMUSG00000033629 | Hacd3 | 3-hydroxyacyl-CoA dehydratase 3 [Source:MGI Symbol;Acc:MGI:1889341] | 1574,6 | 99,91 |
| ENSMUSG00000043445 | Pgp | phosphoglycolate phosphatase [Source:MGI Symbol;Acc:MGI:1914328] | 1573,1 | 100,00 |
| ENSMUSG00000054405 | Dnajc8 | DnaJ heat shock protein family (Hsp40) member C8 [Source:MGI Symbol;Acc:MGI:1915848] | 1571,1 | 92,69 |
| ENSMUSG00000068823 | Csde1 | cold shock domain containing E1, RNA binding [Source:MGI Symbol;Acc:MGI:92356] | 1570,8 | 85,78 |
| ENSMUSG00000041740 | Rnf10 | ring finger protein 10 [Source:MGI Symbol;Acc:MGI:1859162] | 1569,7 | 93,45 |
| ENSMUSG00000024462 | Gabbr1 | gamma-aminobutyric acid (GABA) B receptor, 1 [Source:MGI Symbol;Acc:MGI:1860139] | 1568,0 | 97,23 |
| ENSMUSG00000015804 | Med28 | mediator complex subunit 28 [Source:MGI Symbol;Acc:MGI:1914249] | 1567,8 | 73,18 |
| ENSMUSG00000016427 | Ndufa1 | NADH dehydrogenase (ubiquinone) 1 alpha subcomplex, 1 [Source:MGI Symbol;Acc:MGI:1929511] | 1566,6 | 100,00 |
| ENSMUSG00000030772 | Dkk3 | dickkopf WNT signaling pathway inhibitor 3 [Source:MGI Symbol;Acc:MGI:1354952] | 1558,5 | 100,00 |
| ENSMUSG00000017670 | Elmo2 | engulfment and cell motility 2 [Source:MGI Symbol;Acc:MGI:2153045] | 1557,2 | 81,45 |
| ENSMUSG00000029001 | Fbxo44 | F-box protein 44 [Source:MGI Symbol;Acc:MGI:1354744] | 1554,1 | 88,49 |
| ENSMUSG00000020088 | Sar1a | SAR1 gene homolog A (S. cerevisiae) [Source:MGI Symbol;Acc:MGI:98230] | 1553,6 | 100,00 |
| ENSMUSG00000036707 | Cab39 | calcium binding protein 39 [Source:MGI Symbol;Acc:MGI:107438] | 1549,5 | 94,48 |
| ENSMUSG00000079316 | Rab9 | RAB9, member RAS oncogene family [Source:MGI Symbol;Acc:MGI:1890695] | 1549,2 | 74,37 |
| ENSMUSG00000029106 | Add1 | adducin 1 (alpha) [Source:MGI Symbol;Acc:MGI:87918] | 1548,2 | 84,75 |
| ENSMUSG00000024827 | Gldc | glycine decarboxylase [Source:MGI Symbol;Acc:MGI:1341155] | 1545,1 | 99,84 |
| ENSMUSG00000036752 | Tubb4b | tubulin, beta 4B class IVB [Source:MGI Symbol;Acc:MGI:1915472] | 1543,9 | 86,05 |
| ENSMUSG00000000399 | Ndufa9 | NADH dehydrogenase (ubiquinone) 1 alpha subcomplex, 9 [Source:MGI Symbol;Acc:MGI:1913358] | 1541,2 | 89,25 |
| ENSMUSG00000001768 | Rin2 | Ras and Rab interactor 2 [Source:MGI Symbol;Acc:MGI:1921280] | 1541,2 | 93,09 |
| ENSMUSG00000041236 | Vps41 | vacuolar protein sorting 41 (yeast) [Source:MGI Symbol;Acc:MGI:1929215] | 1540,2 | 99,91 |
| ENSMUSG00000093674 | Rpl41 | ribosomal protein L41 [Source:MGI Symbol;Acc:MGI:1915195] | 1538,3 | 91,04 |
| ENSMUSG00000031666 | Rbl2 | retinoblastoma-like 2 [Source:MGI Symbol;Acc:MGI:105085] | 1537,8 | 100,00 |
| ENSMUSG00000041598 | Cdc42ep4 | CDC42 effector protein (Rho GTPase binding) 4 [Source:MGI Symbol;Acc:MGI:1929760] | 1537,7 | 94,94 |
| ENSMUSG00000054604 | Cggbp1 | CGG triplet repeat binding protein 1 [Source:MGI Symbol;Acc:MGI:2146370] | 1535,8 | 99,65 |
| ENSMUSG00000003531 | Dgcr6 | DiGeorge syndrome critical region gene 6 [Source:MGI Symbol;Acc:MGI:1202877] | 1533,8 | 86,44 |
| ENSMUSG00000002332 | Dhrs1 | dehydrogenase/reductase (SDR family) member 1 [Source:MGI Symbol;Acc:MGI:1196314] | 1531,3 | 97,79 |
| ENSMUSG00000029364 | Wsb2 | WD repeat and SOCS box-containing 2 [Source:MGI Symbol;Acc:MGI:2144041] | 1524,8 | 94,24 |
| ENSMUSG00000016308 | Ube2a | ubiquitin-conjugating enzyme E2A [Source:MGI Symbol;Acc:MGI:102959] | 1524,5 | 98,49 |
| ENSMUSG00000054477 | Kcnn2 | termediate/small conductance calcium-activated channel, subfamily N, member 2 [Source:MGI Symbol;Acc:M | 1524,0 | 82,33 |
| ENSMUSG00000033272 | Slc35a4 | solute carrier family 35, member A4 [Source:MGI Symbol;Acc:MGI:1915093] | 1522,8 | 94,75 |
| ENSMUSG00000026817 | Ak1 | adenylate kinase 1 [Source:MGI Symbol;Acc:MGI:87977] | 1520,2 | 93,07 |
| ENSMUSG00000021114 | Atp6v1d | ATPase, H+ transporting, lysosomal V1 subunit D [Source:MGI Symbol;Acc:MGI:1921084] | 1519,7 | 98,95 |
| ENSMUSG00000027199 | Gatm | glycine amidinotransferase (L-arginine:glycine amidinotransferase) [Source:MGI Symbol;Acc:MGI:1914342] | 1518,1 | 99,80 |
| ENSMUSG00000021024 | Psma6 | proteasome (prosome, macropain) subunit, alpha type 6 [Source:MGI Symbol;Acc:MGI:1347006] | 1508,9 | 89,26 |
| ENSMUSG00000024425 | Ndfip1 | Nedd4 family interacting protein 1 [Source:MGI Symbol;Acc:MGI:1929601] | 1508,7 | 100,00 |
| ENSMUSG00000018286 | Psmb6 | proteasome (prosome, macropain) subunit, beta type 6 [Source:MGI Symbol;Acc:MGI:104880] | 1507,7 | 100,00 |
| ENSMUSG00000079056 | Kcnip3 | Kv channel interacting protein 3, calsenilin [Source:MGI Symbol;Acc:MGI:1929258] | 1503,8 | 89,43 |
| ENSMUSG00000030342 | Cd9 | CD9 antigen [Source:MGI Symbol;Acc:MGI:88348] | 1498,7 | 99,73 |
| ENSMUSG00000005312 | Ubqln1 | ubiquilin 1 [Source:MGI Symbol;Acc:MGI:1860276] | 1498,5 | 99,90 |
| ENSMUSG00000037902 | Sirpa | signal-regulatory protein alpha [Source:MGI Symbol;Acc:MGI:108563] | 1496,9 | 94,21 |
| ENSMUSG00000031451 | Gas6 | growth arrest specific 6 [Source:MGI Symbol;Acc:MGI:95660] | 1496,1 | 99,86 |
| ENSMUSG00000002010 | Idh3g | isocitrate dehydrogenase 3 (NAD+), gamma [Source:MGI Symbol;Acc:MGI:1099463] | 1494,9 | 100,00 |
| ENSMUSG00000016534 | Lamp2 | lysosomal-associated membrane protein 2 [Source:MGI Symbol;Acc:MGI:96748] | 1494,1 | 97,51 |
| ENSMUSG00000015806 | Qdpr | quinoid dihydropteridine reductase [Source:MGI Symbol;Acc:MGI:97836] | 1494,0 | 86,48 |
| ENSMUSG00000021774 | Ube2e1 | ubiquitin-conjugating enzyme E2E 1 [Source:MGI Symbol;Acc:MGI:107411] | 1490,6 | 99,72 |
| ENSMUSG00000046722 | Cdc42se1 | CDC42 small effector 1 [Source:MGI Symbol;Acc:MGI:1889510] | 1489,5 | 93,78 |
| ENSMUSG00000020107 | Anapc16 | anaphase promoting complex subunit 16 [Source:MGI Symbol;Acc:MGI:1289325] | 1488,8 | 93,03 |
| ENSMUSG00000018196 | Glrx2 | glutaredoxin 2 (thioltransferase) [Source:MGI Symbol;Acc:MGI:1916617] | 1488,2 | 86,07 |
| ENSMUSG00000020219 | Timm13 | translocase of inner mitochondrial membrane 13 [Source:MGI Symbol;Acc:MGI:1353432] | 1487,3 | 93,03 |
| ENSMUSG00000019362 | D8Ertd738e | DNA segment, Chr 8, ERATO Doi 738, expressed [Source:MGI Symbol;Acc:MGI:1289231] | 1486,0 | 99,04 |
| ENSMUSG00000023150 | Ivns1abp | influenza virus NS1A binding protein [Source:MGI Symbol;Acc:MGI:2152389] | 1483,7 | 92,04 |
| ENSMUSG00000027428 | Rbbp9 | retinoblastoma binding protein 9 [Source:MGI Symbol;Acc:MGI:1347074] | 1483,0 | 99,98 |
| ENSMUSG00000021111 | Papola | poly (A) polymerase alpha [Source:MGI Symbol;Acc:MGI:109301] | 1481,6 | 97,56 |

| | | | | |
|---|---|---|---|---|
| ENSMUSG00000032026 | Rexo2 | REX2, RNA exonuclease 2 homolog (S. cerevisiae) [Source:MGI Symbol;Acc:MGI:1888981] | 1481,0 | 100,00 |
| ENSMUSG00000022517 | Mgrn1 | mahogunin, ring finger 1 [Source:MGI Symbol;Acc:MGI:2447670] | 1479,4 | 95,82 |
| ENSMUSG00000001666 | Ddt | D-dopachrome tautomerase [Source:MGI Symbol;Acc:MGI:1298381] | 1478,4 | 95,86 |
| ENSMUSG00000058835 | Abi1 | abl-interactor 1 [Source:MGI Symbol;Acc:MGI:104913] | 1478,4 | 98,93 |
| ENSMUSG00000027602 | Map1lc3a | microtubule-associated protein 1 light chain 3 alpha [Source:MGI Symbol;Acc:MGI:1915661] | 1477,6 | 98,86 |
| ENSMUSG00000029310 | Nudt9 | nudix (nucleoside diphosphate linked moiety X)-type motif 9 [Source:MGI Symbol;Acc:MGI:1921417] | 1477,1 | 92,21 |
| ENSMUSG00000085793 | Lin52 | lin-52 homolog (C. elegans) [Source:MGI Symbol;Acc:MGI:3045391] | 1476,1 | 92,65 |
| ENSMUSG00000010376 | Nedd8 | ıeural precursor cell expressed, developmentally down-regulated gene 8 [Source:MGI Symbol;Acc:MGI:97301 | 1475,8 | 100,00 |
| ENSMUSG00000052296 | Ppp6r1 | protein phosphatase 6, regulatory subunit 1 [Source:MGI Symbol;Acc:MGI:2442163] | 1474,2 | 99,22 |
| ENSMUSG00000041889 | Shisa4 | shisa family member 4 [Source:MGI Symbol;Acc:MGI:1924802] | 1473,6 | 99,91 |
| ENSMUSG00000049612 | Omg | oligodendrocyte myelin glycoprotein [Source:MGI Symbol;Acc:MGI:106586] | 1472,1 | 99,72 |
| ENSMUSG00000022110 | Sucla2 | succinate-Coenzyme A ligase, ADP-forming, beta subunit [Source:MGI Symbol;Acc:MGI:1306775] | 1472,0 | 87,49 |
| ENSMUSG00000024181 | Mrpl28 | mitochondrial ribosomal protein L28 [Source:MGI Symbol;Acc:MGI:1915861] | 1471,3 | 100,00 |
| ENSMUSG00000022389 | Tef | thyrotroph embryonic factor [Source:MGI Symbol;Acc:MGI:98663] | 1471,3 | 91,44 |
| ENSMUSG00000062580 | Timm17a | translocase of inner mitochondrial membrane 17a [Source:MGI Symbol;Acc:MGI:1343131] | 1468,9 | 72,92 |
| ENSMUSG00000023048 | Prr13 | proline rich 13 [Source:MGI Symbol;Acc:MGI:1913401] | 1468,2 | 96,52 |
| ENSMUSG00000027499 | Pkia | protein kinase inhibitor, alpha [Source:MGI Symbol;Acc:MGI:104747] | 1464,6 | 97,23 |
| ENSMUSG00000000743 | Chmp1a | charged multivesicular body protein 1A [Source:MGI Symbol;Acc:MGI:1920159] | 1463,6 | 100,00 |
| ENSMUSG00000028383 | Hsdl2 | hydroxysteroid dehydrogenase like 2 [Source:MGI Symbol;Acc:MGI:1919729] | 1462,6 | 93,73 |
| ENSMUSG00000021606 | Ndufs6 | NADH dehydrogenase (ubiquinone) Fe-S protein 6 [Source:MGI Symbol;Acc:MGI:107932] | 1462,5 | 97,01 |
| ENSMUSG00000003623 | Crot | carnitine O-octanoyltransferase [Source:MGI Symbol;Acc:MGI:1921364] | 1461,8 | 96,21 |
| ENSMUSG00000030088 | Aldh1l1 | aldehyde dehydrogenase 1 family, member L1 [Source:MGI Symbol;Acc:MGI:1340024] | 1460,7 | 96,05 |
| ENSMUSG00000071172 | Srsf3 | serine/arginine-rich splicing factor 3 [Source:MGI Symbol;Acc:MGI:98285] | 1460,2 | 83,75 |
| ENSMUSG00000059119 | Nap1l4 | nucleosome assembly protein 1-like 4 [Source:MGI Symbol;Acc:MGI:1316687] | 1458,8 | 98,55 |
| ENSMUSG00000063856 | Gpx1 | glutathione peroxidase 1 [Source:MGI Symbol;Acc:MGI:104887] | 1457,5 | 95,49 |
| ENSMUSG00000033307 | Mif | macrophage migration inhibitory factor [Source:MGI Symbol;Acc:MGI:96982] | 1457,3 | 100,00 |
| ENSMUSG00000053062 | Jam2 | junction adhesion molecule 2 [Source:MGI Symbol;Acc:MGI:1933820] | 1457,1 | 84,29 |
| ENSMUSG00000022698 | Naa50 | N(alpha)-acetyltransferase 50, NatE catalytic subunit [Source:MGI Symbol;Acc:MGI:1919367] | 1454,4 | 96,88 |
| ENSMUSG00000033186 | Mzt1 | mitotic spindle organizing protein 1 [Source:MGI Symbol;Acc:MGI:1924039] | 1452,4 | 97,96 |
| ENSMUSG00000021764 | Ndufs4 | NADH dehydrogenase (ubiquinone) Fe-S protein 4 [Source:MGI Symbol;Acc:MGI:1343135] | 1451,3 | 99,70 |
| ENSMUSG00000042444 | Fam63b | family with sequence similarity 63, member B [Source:MGI Symbol;Acc:MGI:2443086] | 1451,2 | 98,65 |
| ENSMUSG00000019494 | Cops6 | COP9 signalosome subunit 6 [Source:MGI Symbol;Acc:MGI:1349439] | 1449,2 | 96,27 |
| ENSMUSG00000024109 | Nrxn1 | neurexin I [Source:MGI Symbol;Acc:MGI:1096391] | 1448,6 | 91,53 |
| ENSMUSG00000037361 | Sf3b6 | splicing factor 3B, subunit 6 [Source:MGI Symbol;Acc:MGI:1913305] | 1448,2 | 99,76 |
| ENSMUSG00000030890 | Ilk | integrin linked kinase [Source:MGI Symbol;Acc:MGI:1195267] | 1448,1 | 93,20 |
| ENSMUSG00000074637 | Sox2 | SRY (sex determining region Y)-box 2 [Source:MGI Symbol;Acc:MGI:98364] | 1447,9 | 100,00 |
| ENSMUSG00000075486 | Commd6 | COMM domain containing 6 [Source:MGI Symbol;Acc:MGI:1913450] | 1447,5 | 100,00 |
| ENSMUSG00000039682 | Lap3 | leucine aminopeptidase 3 [Source:MGI Symbol;Acc:MGI:1914238] | 1447,4 | 99,44 |
| ENSMUSG00000035237 | Lcat | lecithin cholesterol acyltransferase [Source:MGI Symbol;Acc:MGI:96755] | 1447,3 | 100,00 |
| ENSMUSG00000022964 | Tmem50b | transmembrane protein 50B [Source:MGI Symbol;Acc:MGI:1925225] | 1447,0 | 95,87 |
| ENSMUSG00000036955 | Kif1bp | KIF1 binding protein [Source:MGI Symbol;Acc:MGI:1919570] | 1445,6 | 89,66 |
| ENSMUSG00000037712 | Fermt2 | fermitin family member 2 [Source:MGI Symbol;Acc:MGI:2385001] | 1445,4 | 91,84 |
| ENSMUSG00000020954 | Strn3 | striatin, calmodulin binding protein 3 [Source:MGI Symbol;Acc:MGI:2151064] | 1444,5 | 98,62 |
| ENSMUSG00000027879 | Sec22b | SEC22 homolog B, vesicle trafficking protein [Source:MGI Symbol;Acc:MGI:1338759] | 1443,3 | 76,92 |
| ENSMUSG00000055850 | Rnf181 | ring finger protein 181 [Source:MGI Symbol;Acc:MGI:1913760] | 1440,3 | 82,89 |
| ENSMUSG00000007656 | Arpp19 | cAMP-regulated phosphoprotein 19 [Source:MGI Symbol;Acc:MGI:1891691] | 1439,2 | 96,55 |
| ENSMUSG00000052681 | Rap1b | RAS related protein 1b [Source:MGI Symbol;Acc:MGI:894315] | 1437,6 | 100,00 |
| ENSMUSG00000034120 | Srsf2 | serine/arginine-rich splicing factor 2 [Source:MGI Symbol;Acc:MGI:98284] | 1436,5 | 100,00 |
| ENSMUSG00000077450 | Rab11b | RAB11B, member RAS oncogene family [Source:MGI Symbol;Acc:MGI:99425] | 1435,8 | 88,21 |
| ENSMUSG00000033916 | Chmp2a | charged multivesicular body protein 2A [Source:MGI Symbol;Acc:MGI:1916203] | 1435,2 | 100,00 |
| ENSMUSG00000028923 | Necap2 | NECAP endocytosis associated 2 [Source:MGI Symbol;Acc:MGI:1913397] | 1432,4 | 99,91 |
| ENSMUSG00000046160 | Olig1 | oligodendrocyte transcription factor 1 [Source:MGI Symbol;Acc:MGI:1355334] | 1430,6 | 100,00 |
| ENSMUSG00000028849 | Map7d1 | MAP7 domain containing 1 [Source:MGI Symbol;Acc:MGI:2384297] | 1430,5 | 99,38 |
| ENSMUSG00000064289 | Tank | TRAF family member-associated Nf-kappa B activator [Source:MGI Symbol;Acc:MGI:107676] | 1428,2 | 93,52 |
| ENSMUSG00000030884 | Uqcrc2 | ubiquinol cytochrome c reductase core protein 2 [Source:MGI Symbol;Acc:MGI:1914253] | 1427,4 | 98,63 |
| ENSMUSG00000045160 | Bola3 | bolA-like 3 (E. coli) [Source:MGI Symbol;Acc:MGI:1925903] | 1427,2 | 91,63 |
| ENSMUSG00000038552 | Fndc4 | fibronectin type III domain containing 4 [Source:MGI Symbol;Acc:MGI:1917195] | 1426,4 | 98,92 |
| ENSMUSG00000063015 | Ccni | cyclin I [Source:MGI Symbol;Acc:MGI:1341077] | 1425,7 | 83,90 |
| ENSMUSG00000103217 | Gm38287 | predicted gene, 38287 [Source:MGI Symbol;Acc:MGI:5611515] | 1421,6 | 100,00 |
| ENSMUSG00000039988 | Ankrd13c | ankyrin repeat domain 13c [Source:MGI Symbol;Acc:MGI:2139746] | 1420,4 | 99,87 |
| ENSMUSG00000020132 | Rab21 | RAB21, member RAS oncogene family [Source:MGI Symbol;Acc:MGI:894308] | 1416,2 | 98,90 |
| ENSMUSG00000032184 | Lysmd2 | LysM, putative peptidoglycan-binding, domain containing 2 [Source:MGI Symbol;Acc:MGI:1917332] | 1415,7 | 100,00 |
| ENSMUSG00000021453 | Gadd45g | growth arrest and DNA-damage-inducible 45 gamma [Source:MGI Symbol;Acc:MGI:1346325] | 1412,0 | 100,00 |
| ENSMUSG00000030087 | Klf15 | Kruppel-like factor 15 [Source:MGI Symbol;Acc:MGI:1929988] | 1410,8 | 91,39 |
| ENSMUSG00000028833 | Ncdn | neurochondrin [Source:MGI Symbol;Acc:MGI:1347351] | 1410,6 | 99,75 |
| ENSMUSG00000007411 | Mark3 | MAP/microtubule affinity regulating kinase 3 [Source:MGI Symbol;Acc:MGI:1341865] | 1410,2 | 99,91 |
| ENSMUSG00000036192 | Rorb | RAR-related orphan receptor beta [Source:MGI Symbol;Acc:MGI:1343464] | 1409,4 | 92,74 |
| ENSMUSG00000006728 | Cdk4 | cyclin-dependent kinase 4 [Source:MGI Symbol;Acc:MGI:88357] | 1409,1 | 87,98 |
| ENSMUSG00000022337 | Emc2 | ER membrane protein complex subunit 2 [Source:MGI Symbol;Acc:MGI:1913986] | 1408,4 | 99,89 |
| ENSMUSG00000055553 | Kxd1 | KxDL motif containing 1 [Source:MGI Symbol;Acc:MGI:1922870] | 1403,6 | 81,73 |
| ENSMUSG00000029446 | Psph | phosphoserine phosphatase [Source:MGI Symbol;Acc:MGI:97788] | 1402,7 | 96,60 |
| ENSMUSG00000032757 | Bet1 | Bet1 golgi vesicular membrane trafficking protein [Source:MGI Symbol;Acc:MGI:1343104] | 1401,6 | 99,89 |
| ENSMUSG00000010064 | Slc38a3 | solute carrier family 38, member 3 [Source:MGI Symbol;Acc:MGI:1923507] | 1401,0 | 84,47 |
| ENSMUSG00000001323 | Srr | serine racemase [Source:MGI Symbol;Acc:MGI:1351636] | 1399,1 | 98,23 |
| ENSMUSG00000027706 | Sec62 | SEC62 homolog (S. cerevisiae) [Source:MGI Symbol;Acc:MGI:1916526] | 1396,8 | 96,10 |
| ENSMUSG00000033475 | Tomm6 | translocase of outer mitochondrial membrane 6 homolog (yeast) [Source:MGI Symbol;Acc:MGI:1913369] | 1396,1 | 96,39 |
| ENSMUSG00000022472 | Desi1 | desumoylating isopeptidase 1 [Source:MGI Symbol;Acc:MGI:106313] | 1395,0 | 99,95 |
| ENSMUSG00000059447 | Hadhb | enase/3-ketoacyl-Coenzyme A thiolase/enoyl-Coenzyme A hydratase (trifunctional protein), beta subunit [Sour | 1393,1 | 71,71 |
| ENSMUSG00000027332 | Ivd | isovaleryl coenzyme A dehydrogenase [Source:MGI Symbol;Acc:MGI:1929242] | 1391,7 | 99,10 |
| ENSMUSG00000021241 | Isca2 | iron-sulfur cluster assembly 2 [Source:MGI Symbol;Acc:MGI:1921566] | 1390,8 | 99,92 |
| ENSMUSG00000022708 | Zbtb20 | zinc finger and BTB domain containing 20 [Source:MGI Symbol;Acc:MGI:1929213] | 1389,6 | 91,50 |
| ENSMUSG00000019689 | 1110001J03Rik | RIKEN cDNA 1110001J03 gene [Source:MGI Symbol;Acc:MGI:1913367] | 1388,5 | 100,00 |
| ENSMUSG00000024588 | Fech | ferrochelatase [Source:MGI Symbol;Acc:MGI:95513] | 1386,2 | 100,00 |
| ENSMUSG00000049760 | 2410015M20Rik | RIKEN cDNA 2410015M20 gene [Source:MGI Symbol;Acc:MGI:2442174] | 1384,2 | 100,00 |
| ENSMUSG00000052298 | Cdc42se2 | CDC42 small effector 2 [Source:MGI Symbol;Acc:MGI:1919979] | 1383,2 | 97,40 |
| ENSMUSG00000005103 | Wdr1 | WD repeat domain 1 [Source:MGI Symbol;Acc:MGI:1337100] | 1381,2 | 99,40 |
| ENSMUSG00000060373 | Hnrnpc | heterogeneous nuclear ribonucleoprotein C [Source:MGI Symbol;Acc:MGI:107795] | 1380,6 | 97,72 |
| ENSMUSG00000032076 | Cadm1 | cell adhesion molecule 1 [Source:MGI Symbol;Acc:MGI:1889272] | 1379,8 | 91,31 |
| ENSMUSG00000059824 | Dbp | D site albumin promoter binding protein [Source:MGI Symbol;Acc:MGI:94866] | 1378,4 | 99,46 |
| ENSMUSG00000046329 | Slc25a23 | te carrier family 25 (mitochondrial carrier; phosphate carrier), member 23 [Source:MGI Symbol;Acc:MGI:1914 | 1377,2 | 99,75 |
| ENSMUSG00000039630 | Hnrnpu | heterogeneous nuclear ribonucleoprotein U [Source:MGI Symbol;Acc:MGI:1858195] | 1375,1 | 96,50 |
| ENSMUSG00000040550 | Otud6b | OTU domain containing 6B [Source:MGI Symbol;Acc:MGI:1919451] | 1374,8 | 90,78 |
| ENSMUSG00000027187 | Cat | catalase [Source:MGI Symbol;Acc:MGI:88271] | 1374,1 | 95,99 |
| ENSMUSG00000029772 | Ahcyl2 | S-adenosylhomocysteine hydrolase-like 2 [Source:MGI Symbol;Acc:MGI:1921590] | 1372,5 | 82,09 |
| ENSMUSG00000053644 | Aldh7a1 | aldehyde dehydrogenase family 7, member A1 [Source:MGI Symbol;Acc:MGI:108186] | 1371,4 | 84,82 |
| ENSMUSG00000023020 | Cox14 | cytochrome c oxidase assembly protein 14 [Source:MGI Symbol;Acc:MGI:1913629] | 1370,7 | 100,00 |
| ENSMUSG00000034994 | Eef2 | eukaryotic translation elongation factor 2 [Source:MGI Symbol;Acc:MGI:95288] | 1370,5 | 99,49 |
| ENSMUSG00000036560 | Lgi4 | leucine-rich repeat LGI family, member 4 [Source:MGI Symbol;Acc:MGI:2180197] | 1369,9 | 97,12 |
| ENSMUSG00000019883 | Echdc1 | enoyl Coenzyme A hydratase domain containing 1 [Source:MGI Symbol;Acc:MGI:1277169] | 1369,5 | 98,82 |
| ENSMUSG00000025016 | Tm9sf3 | transmembrane 9 superfamily member 3 [Source:MGI Symbol;Acc:MGI:1914262] | 1369,1 | 99,29 |
| ENSMUSG00000048307 | Ankrd46 | ankyrin repeat domain 46 [Source:MGI Symbol;Acc:MGI:1916089] | 1368,7 | 99,97 |
| ENSMUSG00000022295 | Atp6v1c1 | ATPase, H+ transporting, lysosomal V1 subunit C1 [Source:MGI Symbol;Acc:MGI:1913585] | 1367,6 | 100,00 |
| ENSMUSG00000020402 | Vdac1 | voltage-dependent anion channel 1 [Source:MGI Symbol;Acc:MGI:106919] | 1367,3 | 98,02 |
| ENSMUSG00000020386 | Sar1b | SAR1 gene homolog B (S. cerevisiae) [Source:MGI Symbol;Acc:MGI:1913647] | 1366,5 | 100,00 |
| ENSMUSG00000028669 | Pithd1 | erminal proteasome-interacting domain of thioredoxin-like) domain containing 1 [Source:MGI Symbol;Acc:MGI | 1366,3 | 92,52 |
| ENSMUSG00000031197 | Vbp1 | von Hippel-Lindau binding protein 1 [Source:MGI Symbol;Acc:MGI:1333804] | 1365,5 | 91,42 |
| ENSMUSG00000042363 | Lgalsl | lectin, galactoside binding-like [Source:MGI Symbol;Acc:MGI:1916114] | 1364,8 | 99,99 |
| ENSMUSG00000006736 | Tspan31 | tetraspanin 31 [Source:MGI Symbol;Acc:MGI:1914375] | 1361,0 | 100,00 |
| ENSMUSG00000022553 | Maf1 | MAF1 homolog (S. cerevisiae) [Source:MGI Symbol;Acc:MGI:1916127] | 1358,3 | 92,24 |
| ENSMUSG00000028098 | Rnf115 | ring finger protein 115 [Source:MGI Symbol;Acc:MGI:1915095] | 1356,8 | 99,97 |
| ENSMUSG00000044117 | 2900011O08Rik | RIKEN cDNA 2900011O08 gene [Source:MGI Symbol;Acc:MGI:1914504] | 1355,8 | 98,69 |
| ENSMUSG00000051234 | Rnf7 | ring finger protein 7 [Source:MGI Symbol;Acc:MGI:1337096] | 1355,6 | 98,46 |
| ENSMUSG00000032328 | Tmem30a | transmembrane protein 30A [Source:MGI Symbol;Acc:MGI:106402] | 1353,5 | 97,70 |
| ENSMUSG00000051154 | Commd3 | COMM domain containing 3 [Source:MGI Symbol;Acc:MGI:88218] | 1353,0 | 97,50 |
| ENSMUSG00000020044 | Timp3 | tissue inhibitor of metalloproteinase 3 [Source:MGI Symbol;Acc:MGI:98754] | 1352,2 | 93,03 |
| ENSMUSG00000038412 | Higd1a | HIG1 domain family, member 1A [Source:MGI Symbol;Acc:MGI:1930666] | 1350,4 | 98,08 |
| ENSMUSG00000028127 | Abcd3 | ATP-binding cassette, sub-family D (ALD), member 3 [Source:MGI Symbol;Acc:MGI:1349216] | 1350,3 | 96,98 |

| | | | | |
|---|---|---|---|---|
| ENSMUSG00000030307 | Slc6a11 | solute carrier family 6 (neurotransmitter transporter, GABA), member 11 [Source:MGI Symbol;Acc:MGI:95630 | 1347,4 | 99,89 |
| ENSMUSG00000073676 | Hspe1 | heat shock protein 1 (chaperonin 10) [Source:MGI Symbol;Acc:MGI:104680] | 1347,0 | 97,98 |
| ENSMUSG00000019818 | Cd164 | CD164 antigen [Source:MGI Symbol;Acc:MGI:1859568] | 1346,2 | 97,20 |
| ENSMUSG00000065990 | Aurkaip1 | aurora kinase A interacting protein 1 [Source:MGI Symbol;Acc:MGI:1913327] | 1346,1 | 93,06 |
| ENSMUSG00000026696 | Vamp4 | vesicle-associated membrane protein 4 [Source:MGI Symbol;Acc:MGI:1858730] | 1345,7 | 90,76 |
| ENSMUSG00000033429 | Mcee | methylmalonyl CoA epimerase [Source:MGI Symbol;Acc:MGI:1920974] | 1344,4 | 97,14 |
| ENSMUSG00000004364 | Cul3 | cullin 3 [Source:MGI Symbol;Acc:MGI:1347360] | 1340,8 | 99,19 |
| ENSMUSG00000067847 | Romo1 | reactive oxygen species modulator 1 [Source:MGI Symbol;Acc:MGI:1914317] | 1340,5 | 100,00 |
| ENSMUSG00000000740 | Rpl13 | ribosomal protein L13 [Source:MGI Symbol;Acc:MGI:105922] | 1340,0 | 84,53 |
| ENSMUSG00000001127 | Araf | Araf proto-oncogene, serine/threonine kinase [Source:MGI Symbol;Acc:MGI:88065] | 1336,9 | 76,99 |
| ENSMUSG00000024163 | Mapk8ip3 | mitogen-activated protein kinase 8 interacting protein 3 [Source:MGI Symbol;Acc:MGI:1353598] | 1336,0 | 99,11 |
| ENSMUSG00000016541 | Atxn10 | ataxin 10 [Source:MGI Symbol;Acc:MGI:1859293] | 1335,1 | 99,97 |
| ENSMUSG00000028998 | Tomm7 | translocase of outer mitochondrial membrane 7 homolog (yeast) [Source:MGI Symbol;Acc:MGI:1913419] | 1333,9 | 95,41 |
| ENSMUSG00000028788 | Ptp4a2 | protein tyrosine phosphatase 4a2 [Source:MGI Symbol;Acc:MGI:1277117] | 1330,5 | 98,50 |
| ENSMUSG00000019872 | Smpdl3a | sphingomyelin phosphodiesterase, acid-like 3A [Source:MGI Symbol;Acc:MGI:1931437] | 1329,6 | 88,30 |
| ENSMUSG00000030335 | Mrpl51 | mitochondrial ribosomal protein L51 [Source:MGI Symbol;Acc:MGI:1913743] | 1328,3 | 98,79 |
| ENSMUSG00000047547 | Cltb | clathrin, light polypeptide (Lcb) [Source:MGI Symbol;Acc:MGI:1921575] | 1316,6 | 100,00 |
| ENSMUSG00000036949 | Slc39a12 | solute carrier family 39 (zinc transporter), member 12 [Source:MGI Symbol;Acc:MGI:2139274] | 1315,0 | 96,08 |
| ENSMUSG00000020048 | Hsp90b1 | heat shock protein 90, beta (Grp94), member 1 [Source:MGI Symbol;Acc:MGI:98817] | 1312,1 | 99,74 |
| ENSMUSG00000030663 | 1110004F10Rik | RIKEN cDNA 1110004F10 gene [Source:MGI Symbol;Acc:MGI:1929274] | 1311,7 | 71,07 |
| ENSMUSG00000018548 | Trim37 | tripartite motif-containing 37 [Source:MGI Symbol;Acc:MGI:2153072] | 1308,7 | 99,62 |
| ENSMUSG00000061130 | Ppm1b | protein phosphatase 1B, magnesium dependent, beta isoform [Source:MGI Symbol;Acc:MGI:101841] | 1306,8 | 78,74 |
| ENSMUSG00000026239 | Pde6d | phosphodiesterase 6D, cGMP-specific, rod, delta [Source:MGI Symbol;Acc:MGI:1270843] | 1306,6 | 78,06 |
| ENSMUSG00000038619 | Ensa | endosulfine alpha [Source:MGI Symbol;Acc:MGI:1891189] | 1306,5 | 82,65 |
| ENSMUSG00000031703 | Itfg1 | integrin alpha FG-GAP repeat containing 1 [Source:MGI Symbol;Acc:MGI:106419] | 1306,3 | 99,72 |
| ENSMUSG00000014077 | Chp1 | calcineurin-like EF hand protein 1 [Source:MGI Symbol;Acc:MGI:1927185] | 1305,1 | 79,60 |
| ENSMUSG00000025736 | Jmjd8 | jumonji domain containing 8 [Source:MGI Symbol;Acc:MGI:1919356] | 1304,8 | 93,34 |
| ENSMUSG00000019437 | Tlcd1 | TLC domain containing 1 [Source:MGI Symbol;Acc:MGI:1915572] | 1303,0 | 73,99 |
| ENSMUSG00000002660 | Clpp | caseinolytic mitochondrial matrix peptidase proteolytic subunit [Source:MGI Symbol;Acc:MGI:1858213] | 1300,6 | 95,03 |
| ENSMUSG00000034868 | Myl12b | myosin, light chain 12B, regulatory [Source:MGI Symbol;Acc:MGI:107494] | 1296,8 | 93,06 |
| ENSMUSG00000021286 | Zfyve21 | zinc finger, FYVE domain containing 21 [Source:MGI Symbol;Acc:MGI:1915770] | 1296,5 | 99,57 |
| ENSMUSG00000022112 | Gpc5 | glypican 5 [Source:MGI Symbol;Acc:MGI:1194894] | 1295,0 | 98,97 |
| ENSMUSG00000032047 | Acat1 | acetyl-Coenzyme A acetyltransferase 1 [Source:MGI Symbol;Acc:MGI:87870] | 1293,5 | 88,73 |
| ENSMUSG00000022139 | Mbnl2 | muscleblind-like 2 [Source:MGI Symbol;Acc:MGI:2145597] | 1291,7 | 99,96 |
| ENSMUSG00000027620 | Rbm39 | RNA binding motif protein 39 [Source:MGI Symbol;Acc:MGI:2157953] | 1291,6 | 93,64 |
| ENSMUSG00000075012 | Fjx1 | four jointed box 1 (Drosophila) [Source:MGI Symbol;Acc:MGI:1341907] | 1291,1 | 80,62 |
| ENSMUSG00000039601 | Rcan2 | regulator of calcineurin 2 [Source:MGI Symbol;Acc:MGI:1858219] | 1290,6 | 97,43 |
| ENSMUSG00000028530 | Jak1 | Janus kinase 1 [Source:MGI Symbol;Acc:MGI:96628] | 1289,7 | 98,57 |
| ENSMUSG00000024513 | Mbd2 | methyl-CpG binding domain protein 2 [Source:MGI Symbol;Acc:MGI:1333813] | 1288,0 | 75,99 |
| ENSMUSG00000020811 | Wscd1 | WSC domain containing 1 [Source:MGI Symbol;Acc:MGI:2448493] | 1287,8 | 91,27 |
| ENSMUSG00000000826 | Dnajc5 | DnaJ heat shock protein family (Hsp40) member C5 [Source:MGI Symbol;Acc:MGI:892995] | 1285,1 | 81,27 |
| ENSMUSG00000021917 | Spcs1 | signal peptidase complex subunit 1 homolog (S. cerevisiae) [Source:MGI Symbol;Acc:MGI:1916269] | 1285,0 | 79,26 |
| ENSMUSG00000030265 | Kras | Kirsten rat sarcoma viral oncogene homolog [Source:MGI Symbol;Acc:MGI:96680] | 1279,5 | 95,80 |
| ENSMUSG00000035227 | Spcs2 | signal peptidase complex subunit 2 homolog (S. cerevisiae) [Source:MGI Symbol;Acc:MGI:1913874] | 1277,5 | 74,78 |
| ENSMUSG00000042541 | Shfm1 | split hand/foot malformation (ectrodactyly) type 1 [Source:MGI Symbol;Acc:MGI:109238] | 1277,0 | 94,95 |
| ENSMUSG00000020664 | Dld | dihydrolipoamide dehydrogenase [Source:MGI Symbol;Acc:MGI:107450] | 1276,0 | 100,00 |
| ENSMUSG00000021156 | Zmynd11 | zinc finger, MYND domain containing 11 [Source:MGI Symbol;Acc:MGI:1913755] | 1275,3 | 93,59 |
| ENSMUSG00000037010 | Apln | apelin [Source:MGI Symbol;Acc:MGI:1353624] | 1273,6 | 100,00 |
| ENSMUSG00000034463 | Scara3 | scavenger receptor class A, member 3 [Source:MGI Symbol;Acc:MGI:2444418] | 1272,9 | 100,00 |
| ENSMUSG00000024750 | Zfand5 | zinc finger, AN1-type domain 5 [Source:MGI Symbol;Acc:MGI:1278334] | 1270,5 | 96,01 |
| ENSMUSG00000004285 | Atp6v1f | ATPase, H+ transporting, lysosomal V1 subunit F [Source:MGI Symbol;Acc:MGI:1913394] | 1270,4 | 70,08 |
| ENSMUSG00000024500 | Ppp2r2b | protein phosphatase 2, regulatory subunit B, beta [Source:MGI Symbol;Acc:MGI:1920180] | 1270,2 | 95,27 |
| ENSMUSG00000027984 | Hadh | hydroxyacyl-Coenzyme A dehydrogenase [Source:MGI Symbol;Acc:MGI:96009] | 1268,7 | 98,68 |
| ENSMUSG00000003464 | Pex19 | peroxisomal biogenesis factor 19 [Source:MGI Symbol;Acc:MGI:1334458] | 1264,5 | 99,79 |
| ENSMUSG00000054428 | Atpif1 | ATPase inhibitory factor 1 [Source:MGI Symbol;Acc:MGI:1196457] | 1263,0 | 96,89 |
| ENSMUSG00000006476 | Nsmf | MDA receptor synaptonuclear signaling and neuronal migration factor [Source:MGI Symbol;Acc:MGI:186175 | 1262,6 | 98,65 |
| ENSMUSG00000006740 | Kif5b | kinesin family member 5B [Source:MGI Symbol;Acc:MGI:1098268] | 1262,4 | 95,03 |
| ENSMUSG00000055943 | Emc7 | ER membrane protein complex subunit 7 [Source:MGI Symbol;Acc:MGI:1920274] | 1259,3 | 96,50 |
| ENSMUSG00000087260 | Lamtor5 | late endosomal/lysosomal adaptor, MAPK and MTOR activator 5 [Source:MGI Symbol;Acc:MGI:1915826] | 1255,8 | 90,01 |
| ENSMUSG00000020629 | Adi1 | acireductone dioxygenase 1 [Source:MGI Symbol;Acc:MGI:2144929] | 1254,7 | 98,58 |
| ENSMUSG00000026864 | Hspa5 | heat shock protein 5 [Source:MGI Symbol;Acc:MGI:95835] | 1253,9 | 71,26 |
| ENSMUSG00000029432 | Gbas | glioblastoma amplified sequence [Source:MGI Symbol;Acc:MGI:1278343] | 1253,5 | 99,62 |
| ENSMUSG00000027599 | Armc1 | armadillo repeat containing 1 [Source:MGI Symbol;Acc:MGI:1921502] | 1252,8 | 84,05 |
| ENSMUSG00000029472 | Anapc5 | anaphase-promoting complex subunit 5 [Source:MGI Symbol;Acc:MGI:1929722] | 1251,5 | 97,56 |
| ENSMUSG00000022634 | Yaf2 | YY1 associated factor 2 [Source:MGI Symbol;Acc:MGI:1914307] | 1251,4 | 100,00 |
| ENSMUSG00000006932 | Ctnnb1 | catenin (cadherin associated protein), beta 1 [Source:MGI Symbol;Acc:MGI:88276] | 1250,7 | 84,65 |
| ENSMUSG00000030286 | Emc3 | ER membrane protein complex subunit 3 [Source:MGI Symbol;Acc:MGI:1913337] | 1248,1 | 99,71 |
| ENSMUSG00000030042 | Pole4 | polymerase (DNA-directed), epsilon 4 (p12 subunit) [Source:MGI Symbol;Acc:MGI:1914229] | 1246,9 | 85,67 |
| ENSMUSG00000003131 | Pafah1b2 | platelet-activating factor acetylhydrolase, isoform 1b, subunit 2 [Source:MGI Symbol;Acc:MGI:108415] | 1246,3 | 99,82 |
| ENSMUSG00000036781 | Rps27l | ribosomal protein S27-like [Source:MGI Symbol;Acc:MGI:1915191] | 1245,3 | 100,00 |
| ENSMUSG00000062929 | Cfl2 | cofilin 2, muscle [Source:MGI Symbol;Acc:MGI:101763] | 1245,1 | 99,80 |
| ENSMUSG00000034892 | Rps29 | ribosomal protein S29 [Source:MGI Symbol;Acc:MGI:107681] | 1244,4 | 93,50 |
| ENSMUSG00000027739 | Rab33b | RAB33B, member RAS oncogene family [Source:MGI Symbol;Acc:MGI:1330805] | 1241,5 | 91,17 |
| ENSMUSG00000022680 | Pdxdc1 | pyridoxal-dependent decarboxylase domain containing 1 [Source:MGI Symbol;Acc:MGI:1920909] | 1240,3 | 74,52 |
| ENSMUSG00000040370 | Lyrm5 | LYR motif containing 5 [Source:MGI Symbol;Acc:MGI:1914886] | 1238,5 | 99,39 |
| ENSMUSG00000026500 | Cox20 | COX20 Cox2 chaperone [Source:MGI Symbol;Acc:MGI:1913609] | 1237,8 | 100,00 |
| ENSMUSG00000027574 | Nkain4 | Na+/K+ transporting ATPase interacting 4 [Source:MGI Symbol;Acc:MGI:1915372] | 1237,7 | 100,00 |
| ENSMUSG00000017221 | Psmd3 | proteasome (prosome, macropain) 26S subunit, non-ATPase, 3 [Source:MGI Symbol;Acc:MGI:98858] | 1237,3 | 96,64 |
| ENSMUSG00000022677 | Fopnl | Fgfr1op N-terminal like [Source:MGI Symbol;Acc:MGI:1913336] | 1236,4 | 99,77 |
| ENSMUSG00000026259 | Ngef | neuronal guanine nucleotide exchange factor [Source:MGI Symbol;Acc:MGI:1858414] | 1235,4 | 82,91 |
| ENSMUSG00000029084 | Cd38 | CD38 antigen [Source:MGI Symbol;Acc:MGI:107474] | 1233,9 | 96,32 |
| ENSMUSG00000003380 | Rabac1 | Rab acceptor 1 (prenylated) [Source:MGI Symbol;Acc:MGI:1201692] | 1231,7 | 95,38 |
| ENSMUSG00000092395 | Gm20463 | predicted gene 20463 [Source:MGI Symbol;Acc:MGI:5141928] | 1229,1 | 95,72 |
| ENSMUSG00000053398 | Phgdh | 3-phosphoglycerate dehydrogenase [Source:MGI Symbol;Acc:MGI:1355330] | 1227,9 | 86,98 |
| ENSMUSG00000030879 | Mrpl17 | mitochondrial ribosomal protein L17 [Source:MGI Symbol;Acc:MGI:1351608] | 1227,6 | 96,81 |
| ENSMUSG00000028367 | Txn1 | thioredoxin 1 [Source:MGI Symbol;Acc:MGI:98874] | 1227,1 | 89,09 |
| ENSMUSG00000023452 | Pisd | phosphatidylserine decarboxylase [Source:MGI Symbol;Acc:MGI:2445114] | 1226,5 | 93,41 |
| ENSMUSG00000028138 | Adh5 | alcohol dehydrogenase 5 (class III), chi polypeptide [Source:MGI Symbol;Acc:MGI:87929] | 1226,4 | 98,61 |
| ENSMUSG00000021981 | Cab39l | calcium binding protein 39-like [Source:MGI Symbol;Acc:MGI:1914081] | 1226,2 | 99,99 |
| ENSMUSG00000048960 | Prex2 | osphatidylinositol-3,4,5-trisphosphate-dependent Rac exchange factor 2 [Source:MGI Symbol;Acc:MGI:19233 | 1225,7 | 99,90 |
| ENSMUSG00000038880 | Mrps34 | mitochondrial ribosomal protein S34 [Source:MGI Symbol;Acc:MGI:1930188] | 1224,9 | 99,93 |
| ENSMUSG00000029020 | Mfn2 | mitofusin 2 [Source:MGI Symbol;Acc:MGI:2442230] | 1224,8 | 89,29 |
| ENSMUSG00000031782 | Coq9 | coenzyme Q9 [Source:MGI Symbol;Acc:MGI:1915164] | 1224,2 | 98,36 |
| ENSMUSG00000039461 | Tcta | T cell leukemia translocation altered gene [Source:MGI Symbol;Acc:MGI:1918829] | 1222,4 | 96,99 |
| ENSMUSG00000054364 | Rhob | ras homolog gene family, member B [Source:MGI Symbol;Acc:MGI:107949] | 1222,2 | 99,97 |
| ENSMUSG00000014606 | Slc25a11 | e carrier family 25 (mitochondrial carrier oxoglutarate carrier), member 11 [Source:MGI Symbol;Acc:MGI:1915 | 1220,6 | 96,86 |
| ENSMUSG00000018585 | Atox1 | antioxidant 1 copper chaperone [Source:MGI Symbol;Acc:MGI:1333855] | 1220,0 | 98,40 |
| ENSMUSG00000025794 | Rpl14 | ribosomal protein L14 [Source:MGI Symbol;Acc:MGI:1914365] | 1218,9 | 96,36 |
| ENSMUSG00000001729 | Akt1 | thymoma viral proto-oncogene 1 [Source:MGI Symbol;Acc:MGI:87986] | 1218,2 | 86,79 |
| ENSMUSG00000027637 | 1110008F13Rik | RIKEN cDNA 1110008F13 gene [Source:MGI Symbol;Acc:MGI:1914638] | 1218,0 | 100,00 |
| ENSMUSG00000039770 | Ypel5 | yippee-like 5 (Drosophila) [Source:MGI Symbol;Acc:MGI:1916937] | 1217,8 | 91,51 |
| ENSMUSG00000042675 | Ypel3 | yippee-like 3 (Drosophila) [Source:MGI Symbol;Acc:MGI:1913340] | 1217,5 | 73,52 |
| ENSMUSG00000042682 | Selk | selenoprotein K [Source:MGI Symbol;Acc:MGI:1931466] | 1217,4 | 97,55 |
| ENSMUSG00000010277 | 2610507B11Rik | RIKEN cDNA 2610507B11 gene [Source:MGI Symbol;Acc:MGI:1919753] | 1216,2 | 99,87 |
| ENSMUSG00000029669 | Tspan12 | tetraspanin 12 [Source:MGI Symbol;Acc:MGI:1889818] | 1211,7 | 88,14 |
| ENSMUSG00000021477 | Ctsl | cathepsin L [Source:MGI Symbol;Acc:MGI:88564] | 1211,7 | 94,62 |
| ENSMUSG00000017466 | Timp2 | tissue inhibitor of metalloproteinase 2 [Source:MGI Symbol;Acc:MGI:98753] | 1211,6 | 93,11 |
| ENSMUSG00000029146 | Snx17 | sorting nexin 17 [Source:MGI Symbol;Acc:MGI:2387801] | 1209,5 | 99,79 |
| ENSMUSG00000001240 | Ramp2 | receptor (calcitonin) activity modifying protein 2 [Source:MGI Symbol;Acc:MGI:1859650] | 1209,3 | 85,78 |
| ENSMUSG00000006373 | Pgrmc1 | progesterone receptor membrane component 1 [Source:MGI Symbol;Acc:MGI:1858305] | 1208,8 | 98,65 |
| ENSMUSG00000004936 | Map2k1 | mitogen-activated protein kinase kinase 1 [Source:MGI Symbol;Acc:MGI:1346866] | 1208,1 | 99,74 |
| ENSMUSG00000031913 | Vps4a | vacuolar protein sorting 4a (yeast) [Source:MGI Symbol;Acc:MGI:1890520] | 1208,0 | 98,61 |
| ENSMUSG00000028821 | Syf2 | SYF2 homolog, RNA splicing factor (S. cerevisiae) [Source:MGI Symbol;Acc:MGI:1915842] | 1205,3 | 99,32 |
| ENSMUSG00000031696 | Vps35 | vacuolar protein sorting 35 [Source:MGI Symbol;Acc:MGI:1890467] | 1202,5 | 100,00 |
| ENSMUSG00000057236 | Rbbp4 | retinoblastoma binding protein 4 [Source:MGI Symbol;Acc:MGI:1194912] | 1202,3 | 96,42 |
| ENSMUSG00000020708 | Psmc5 | protease (prosome, macropain) 26S subunit, ATPase 5 [Source:MGI Symbol;Acc:MGI:105047] | 1201,7 | 100,00 |
| ENSMUSG00000004319 | Clcn3 | chloride channel, voltage-sensitive 3 [Source:MGI Symbol;Acc:MGI:103555] | 1201,2 | 94,84 |

| | | | | |
|---|---|---|---|---|
| ENSMUSG00000000420 | Galnt1 | acetyl-alpha-D-galactosamine:polypeptide N-acetylgalactosaminyltransferase 1 [Source:MGI Symbol;Acc:MGI: | 1198,8 | 89,63 |
| ENSMUSG00000030744 | Rps3 | ribosomal protein S3 [Source:MGI Symbol;Acc:MGI:1350917] | 1198,7 | 95,25 |
| ENSMUSG00000001017 | Chtop | chromatin target of PRMT1 [Source:MGI Symbol;Acc:MGI:1913761] | 1195,0 | 91,53 |
| ENSMUSG00000031954 | Cfdp1 | craniofacial development protein 1 [Source:MGI Symbol;Acc:MGI:1344403] | 1193,5 | 100,00 |
| ENSMUSG00000042396 | Rbm7 | RNA binding motif protein 7 [Source:MGI Symbol;Acc:MGI:1914260] | 1193,1 | 100,00 |
| ENSMUSG00000030103 | Bhlhe40 | basic helix-loop-helix family, member e40 [Source:MGI Symbol;Acc:MGI:1097714] | 1192,8 | 96,08 |
| ENSMUSG00000026755 | Arpc5l | actin related protein 2/3 complex, subunit 5-like [Source:MGI Symbol;Acc:MGI:1921442] | 1191,8 | 99,95 |
| ENSMUSG00000024454 | Hdac3 | histone deacetylase 3 [Source:MGI Symbol;Acc:MGI:1343091] | 1190,8 | 99,00 |
| ENSMUSG00000068732 | Tmem167b | transmembrane protein 167B [Source:MGI Symbol;Acc:MGI:1914745] | 1189,5 | 99,07 |
| ENSMUSG00000058756 | Thra | thyroid hormone receptor alpha [Source:MGI Symbol;Acc:MGI:98742] | 1189,1 | 97,14 |
| ENSMUSG00000021684 | Pde8b | phosphodiesterase 8B [Source:MGI Symbol;Acc:MGI:2443999] | 1189,0 | 92,48 |
| ENSMUSG00000007682 | Dio2 | deiodinase, iodothyronine, type II [Source:MGI Symbol;Acc:MGI:1338833] | 1185,2 | 99,39 |
| ENSMUSG00000030842 | Lamtor1 | late endosomal/lysosomal adaptor, MAPK and MTOR activator 1 [Source:MGI Symbol;Acc:MGI:1913758] | 1184,3 | 81,92 |
| ENSMUSG00000022681 | Ntan1 | N-terminal Asn amidase [Source:MGI Symbol;Acc:MGI:108471] | 1184,1 | 91,89 |
| ENSMUSG00000020358 | Hnrnpab | heterogeneous nuclear ribonucleoprotein A/B [Source:MGI Symbol;Acc:MGI:1330294] | 1183,4 | 91,64 |
| ENSMUSG00000028646 | Rragc | Ras-related GTP binding C [Source:MGI Symbol;Acc:MGI:1858751] | 1182,6 | 99,61 |
| ENSMUSG00000021537 | Cetn3 | centrin 3 [Source:MGI Symbol;Acc:MGI:1097706] | 1181,6 | 100,00 |
| ENSMUSG00000019868 | Vta1 | Vps20-associated 1 homolog (S. cerevisiae) [Source:MGI Symbol;Acc:MGI:1913451] | 1181,1 | 97,72 |
| ENSMUSG00000039105 | Atp6v1g1 | ATPase, H+ transporting, lysosomal V1 subunit G1 [Source:MGI Symbol;Acc:MGI:1913540] | 1180,9 | 88,60 |
| ENSMUSG00000049086 | Bmyc | brain expressed myelocytomatosis oncogene [Source:MGI Symbol;Acc:MGI:88184] | 1178,5 | 100,00 |
| ENSMUSG00000022105 | Rb1 | retinoblastoma 1 [Source:MGI Symbol;Acc:MGI:97874] | 1178,5 | 98,22 |
| ENSMUSG00000028013 | Ppa2 | pyrophosphatase (inorganic) 2 [Source:MGI Symbol;Acc:MGI:1922026] | 1176,3 | 98,26 |
| ENSMUSG00000047714 | Ppp1r2 | protein phosphatase 1, regulatory (inhibitor) subunit 2 [Source:MGI Symbol;Acc:MGI:1914099] | 1175,3 | 86,40 |
| ENSMUSG00000028759 | Hp1bp3 | heterochromatin protein 1, binding protein 3 [Source:MGI Symbol;Acc:MGI:109369] | 1174,9 | 70,18 |
| ENSMUSG00000003279 | Dlgap1 | discs, large (Drosophila) homolog-associated protein 1 [Source:MGI Symbol;Acc:MGI:1346065] | 1172,7 | 94,93 |
| ENSMUSG00000005656 | Snx6 | sorting nexin 6 [Source:MGI Symbol;Acc:MGI:1919433] | 1172,0 | 99,72 |
| ENSMUSG00000035376 | Hacd2 | 3-hydroxyacyl-CoA dehydratase 2 [Source:MGI Symbol;Acc:MGI:1918007] | 1171,8 | 99,20 |
| ENSMUSG00000040289 | Hey1 | hairy/enhancer-of-split related with YRPW motif 1 [Source:MGI Symbol;Acc:MGI:1341800] | 1171,0 | 93,56 |
| ENSMUSG00000047260 | Emc6 | ER membrane protein complex subunit 6 [Source:MGI Symbol;Acc:MGI:1913298] | 1170,5 | 97,41 |
| ENSMUSG00000090841 | Myl6 | myosin, light polypeptide 6, alkali, smooth muscle and non-muscle [Source:MGI Symbol;Acc:MGI:109318] | 1168,7 | 96,14 |
| ENSMUSG00000038332 | Sesn1 | sestrin 1 [Source:MGI Symbol;Acc:MGI:2155278] | 1168,5 | 98,97 |
| ENSMUSG00000038871 | Bpgm | 2,3-bisphosphoglycerate mutase [Source:MGI Symbol;Acc:MGI:1098242] | 1168,5 | 81,70 |
| ENSMUSG00000028927 | Padi2 | peptidyl arginine deiminase, type II [Source:MGI Symbol;Acc:MGI:1338892] | 1166,8 | 99,87 |
| ENSMUSG00000068882 | Ssb | Sjogren syndrome antigen B [Source:MGI Symbol;Acc:MGI:98423] | 1166,4 | 98,79 |
| ENSMUSG00000051695 | Pcbp1 | poly(rC) binding protein 1 [Source:MGI Symbol;Acc:MGI:1345635] | 1166,4 | 90,71 |
| ENSMUSG00000038633 | Degs1 | delta(4)-desaturase, sphingolipid 1 [Source:MGI Symbol;Acc:MGI:1097711] | 1166,0 | 94,07 |
| ENSMUSG00000029128 | Rab28 | RAB28, member RAS oncogene family [Source:MGI Symbol;Acc:MGI:1917285] | 1165,0 | 100,00 |
| ENSMUSG00000040374 | Pex2 | peroxisomal biogenesis factor 2 [Source:MGI Symbol;Acc:MGI:107486] | 1165,0 | 98,28 |
| ENSMUSG00000002524 | Puf60 | poly-U binding splicing factor 60 [Source:MGI Symbol;Acc:MGI:1915209] | 1164,1 | 99,74 |
| ENSMUSG00000022403 | St13 | suppression of tumorigenicity 13 [Source:MGI Symbol;Acc:MGI:1917606] | 1163,7 | 97,69 |
| ENSMUSG00000022043 | Trim35 | tripartite motif-containing 35 [Source:MGI Symbol;Acc:MGI:1914104] | 1163,6 | 91,41 |
| ENSMUSG00000037916 | Ndufv1 | NADH dehydrogenase (ubiquinone) flavoprotein 1 [Source:MGI Symbol;Acc:MGI:107851] | 1160,5 | 98,17 |
| ENSMUSG00000056698 | Elmod3 | ELMO/CED-12 domain containing 3 [Source:MGI Symbol;Acc:MGI:2445168] | 1160,4 | 93,30 |
| ENSMUSG00000003813 | Rad23a | RAD23a homolog (S. cerevisiae) [Source:MGI Symbol;Acc:MGI:105126] | 1158,6 | 97,09 |
| ENSMUSG00000008475 | Arpc5 | actin related protein 2/3 complex, subunit 5 [Source:MGI Symbol;Acc:MGI:1915021] | 1158,2 | 98,83 |
| ENSMUSG00000038388 | Mpp6 | membrane protein, palmitoylated 6 (MAGUK p55 subfamily member 6) [Source:MGI Symbol;Acc:MGI:192734( | 1157,5 | 98,98 |
| ENSMUSG00000071076 | Jund | jun D proto-oncogene [Source:MGI Symbol;Acc:MGI:96648] | 1157,4 | 99,44 |
| ENSMUSG00000022744 | Cldnd1 | claudin domain containing 1 [Source:MGI Symbol;Acc:MGI:2447860] | 1157,2 | 96,60 |
| ENSMUSG00000027803 | Wwtr1 | WW domain containing transcription regulator 1 [Source:MGI Symbol;Acc:MGI:1917649] | 1156,0 | 98,95 |
| ENSMUSG00000030613 | Ccdc90b | coiled-coil domain containing 90B [Source:MGI Symbol;Acc:MGI:1913615] | 1155,9 | 99,83 |
| ENSMUSG00000024914 | Drap1 | Dr1 associated protein 1 (negative cofactor 2 alpha) [Source:MGI Symbol;Acc:MGI:1913806] | 1155,7 | 93,41 |
| ENSMUSG00000003429 | Rps11 | ribosomal protein S11 [Source:MGI Symbol;Acc:MGI:1351329] | 1154,4 | 83,39 |
| ENSMUSG00000071180 | Smim15 | small integral membrane protein 15 [Source:MGI Symbol;Acc:MGI:1922866] | 1153,6 | 100,00 |
| ENSMUSG00000036880 | Acaa2 | oenzyme A acyltransferase 2 (mitochondrial 3-oxoacyl-Coenzyme A thiolase) [Source:MGI Symbol;Acc:MGI:1 | 1153,4 | 100,00 |
| ENSMUSG00000038696 | Mapkap1 | mitogen-activated protein kinase associated protein 1 [Source:MGI Symbol;Acc:MGI:2444554] | 1151,8 | 96,81 |
| ENSMUSG00000031601 | Cnot7 | CCR4-NOT transcription complex, subunit 7 [Source:MGI Symbol;Acc:MGI:1298230] | 1151,8 | 77,93 |
| ENSMUSG00000031701 | Dnaja2 | DnaJ heat shock protein family (Hsp40) member A2 [Source:MGI Symbol;Acc:MGI:1931882] | 1151,1 | 99,94 |
| ENSMUSG00000020719 | Ddx5 | DEAD (Asp-Glu-Ala-Asp) box polypeptide 5 [Source:MGI Symbol;Acc:MGI:105037] | 1151,0 | 97,62 |
| ENSMUSG00000062908 | Acadm | acyl-Coenzyme A dehydrogenase, medium chain [Source:MGI Symbol;Acc:MGI:87867] | 1151,0 | 94,62 |
| ENSMUSG00000010095 | Slc3a2 | arrier family 3 (activators of dibasic and neutral amino acid transport), member 2 [Source:MGI Symbol;Acc:MG | 1150,7 | 72,16 |
| ENSMUSG00000018501 | Ncor1 | nuclear receptor co-repressor 1 [Source:MGI Symbol;Acc:MGI:1349717] | 1150,2 | 91,68 |
| ENSMUSG00000013160 | Atp6v0d1 | ATPase, H+ transporting, lysosomal V0 subunit D1 [Source:MGI Symbol;Acc:MGI:1201778] | 1149,6 | 99,61 |
| ENSMUSG00000005510 | Ndufs3 | NADH dehydrogenase (ubiquinone) Fe-S protein 3 [Source:MGI Symbol;Acc:MGI:1915599] | 1149,4 | 89,10 |
| ENSMUSG00000019478 | Rab4a | RAB4A, member RAS oncogene family [Source:MGI Symbol;Acc:MGI:105069] | 1148,8 | 93,59 |
| ENSMUSG00000016257 | Slmo2 | slowmo homolog 2 (Drosophila) [Source:MGI Symbol;Acc:MGI:1913640] | 1147,7 | 96,22 |
| ENSMUSG00000043716 | Rpl7 | ribosomal protein L7 [Source:MGI Symbol;Acc:MGI:98073] | 1146,3 | 71,27 |
| ENSMUSG00000107002 | 0610012G03Rik | RIKEN cDNA 0610012G03 gene [Source:MGI Symbol;Acc:MGI:1913301] | 1145,5 | 99,68 |
| ENSMUSG00000004843 | Chmp2b | charged multivesicular body protein 2B [Source:MGI Symbol;Acc:MGI:1916192] | 1143,8 | 100,00 |
| ENSMUSG00000041453 | Rpl21 | ribosomal protein L21 [Source:MGI Symbol;Acc:MGI:1278340] | 1141,7 | 95,48 |
| ENSMUSG00000000168 | Dlat | amide S-acetyltransferase (E2 component of pyruvate dehydrogenase complex) [Source:MGI Symbol;Acc:MG | 1141,5 | 96,65 |
| ENSMUSG00000018965 | Ywhah | monooxygenase/tryptophan 5-monooxygenase activation protein, eta polypeptide [Source:MGI Symbol;Acc:M | 1141,5 | 77,72 |
| ENSMUSG00000021775 | Nr1d2 | nuclear receptor subfamily 1, group D, member 2 [Source:MGI Symbol;Acc:MGI:2449205] | 1136,4 | 97,08 |
| ENSMUSG00000029198 | Grpel1 | GrpE-like 1, mitochondrial [Source:MGI Symbol;Acc:MGI:1334417] | 1134,7 | 93,89 |
| ENSMUSG00000071655 | Ubxn1 | UBX domain protein 1 [Source:MGI Symbol;Acc:MGI:1289301] | 1134,2 | 100,00 |
| ENSMUSG00000027011 | Ube2e3 | ubiquitin-conjugating enzyme E2E 3 [Source:MGI Symbol;Acc:MGI:107412] | 1133,2 | 72,97 |
| ENSMUSG00000034345 | Gtf2h5 | general transcription factor IIH, polypeptide 5 [Source:MGI Symbol;Acc:MGI:107227] | 1133,1 | 97,87 |
| ENSMUSG00000000538 | Tom1l2 | target of myb1-like 2 (chicken) [Source:MGI Symbol;Acc:MGI:2443306] | 1132,1 | 84,22 |
| ENSMUSG00000035297 | Cops4 | COP9 signalosome subunit 4 [Source:MGI Symbol;Acc:MGI:1349414] | 1132,0 | 88,55 |
| ENSMUSG00000053119 | Chmp3 | charged multivesicular body protein 3 [Source:MGI Symbol;Acc:MGI:1913950] | 1131,4 | 95,39 |
| ENSMUSG00000063694 | Cycs | cytochrome c, somatic [Source:MGI Symbol;Acc:MGI:88578] | 1130,5 | 74,03 |
| ENSMUSG00000030538 | Cib1 | calcium and integrin binding 1 (calmyrin) [Source:MGI Symbol;Acc:MGI:1344418] | 1129,0 | 88,77 |
| ENSMUSG00000027618 | Nfs1 | nitrogen fixation gene 1 (S. cerevisiae) [Source:MGI Symbol;Acc:MGI:1316706] | 1128,9 | 98,61 |
| ENSMUSG00000032046 | Abhd12 | abhydrolase domain containing 12 [Source:MGI Symbol;Acc:MGI:1923442] | 1128,4 | 99,35 |
| ENSMUSG00000035027 | Map2k2 | mitogen-activated protein kinase kinase 2 [Source:MGI Symbol;Acc:MGI:1346867] | 1126,0 | 98,19 |
| ENSMUSG00000023092 | Fhl1 | four and a half LIM domains 1 [Source:MGI Symbol;Acc:MGI:1298387] | 1125,9 | 78,02 |
| ENSMUSG00000028657 | Ppt1 | palmitoyl-protein thioesterase 1 [Source:MGI Symbol;Acc:MGI:1298204] | 1124,8 | 74,92 |
| ENSMUSG00000017188 | Coa3 | cytochrome C oxidase assembly factor 3 [Source:MGI Symbol;Acc:MGI:1098757] | 1122,9 | 85,82 |
| ENSMUSG00000034432 | Cops8 | COP9 signalosome subunit 8 [Source:MGI Symbol;Acc:MGI:1915363] | 1122,8 | 78,22 |
| ENSMUSG00000029076 | Sdf4 | stromal cell derived factor 4 [Source:MGI Symbol;Acc:MGI:108079] | 1122,8 | 94,09 |
| ENSMUSG00000043496 | Tril | TLR4 interactor with leucine-rich repeats [Source:MGI Symbol;Acc:MGI:1914123] | 1120,4 | 99,98 |
| ENSMUSG00000037971 | 1110032A03Rik | RIKEN cDNA 1110032A03 gene [Source:MGI Symbol;Acc:MGI:1915971] | 1120,0 | 75,65 |
| ENSMUSG00000046432 | Ngfrap1 | nerve growth factor receptor (TNFRSF16) associated protein 1 [Source:MGI Symbol;Acc:MGI:1338016] | 1117,8 | 91,71 |
| ENSMUSG00000020456 | Ogdh | oxoglutarate (alpha-ketoglutarate) dehydrogenase (lipoamide) [Source:MGI Symbol;Acc:MGI:1098267] | 1117,0 | 94,62 |
| ENSMUSG00000025903 | Lypla1 | lysophospholipase 1 [Source:MGI Symbol;Acc:MGI:1344588] | 1116,7 | 94,11 |
| ENSMUSG00000003346 | Abhd17a | abhydrolase domain containing 17A [Source:MGI Symbol;Acc:MGI:106388] | 1115,9 | 95,28 |
| ENSMUSG00000044502 | Bod1 | biorientation of chromosomes in cell division 1 [Source:MGI Symbol;Acc:MGI:1916806] | 1115,7 | 97,47 |
| ENSMUSG00000006717 | Acot13 | acyl-CoA thioesterase 13 [Source:MGI Symbol;Acc:MGI:1914084] | 1113,9 | 99,95 |
| ENSMUSG00000027777 | Schip1 | schwannomin interacting protein 1 [Source:MGI Symbol;Acc:MGI:1353557] | 1113,4 | 82,97 |
| ENSMUSG00000027104 | Atf2 | activating transcription factor 2 [Source:MGI Symbol;Acc:MGI:109349] | 1113,1 | 98,64 |
| ENSMUSG00000031988 | Vps26b | vacuolar protein sorting 26 homolog B (yeast) [Source:MGI Symbol;Acc:MGI:1917656] | 1112,8 | 95,93 |
| ENSMUSG00000026853 | Crat | carnitine acetyltransferase [Source:MGI Symbol;Acc:MGI:109501] | 1112,1 | 80,32 |
| ENSMUSG00000025139 | Tollip | toll interacting protein [Source:MGI Symbol;Acc:MGI:1891808] | 1111,8 | 88,43 |
| ENSMUSG00000006273 | Atp6v1b2 | ATPase, H+ transporting, lysosomal V1 subunit B2 [Source:MGI Symbol;Acc:MGI:109618] | 1111,2 | 96,58 |
| ENSMUSG00000003199 | Mpnd | MPN domain containing [Source:MGI Symbol;Acc:MGI:1915297] | 1110,0 | 94,03 |
| ENSMUSG00000022201 | Zfr | zinc finger RNA binding protein [Source:MGI Symbol;Acc:MGI:1341890] | 1109,4 | 94,42 |
| ENSMUSG00000005982 | Naa60 | N(alpha)-acetyltransferase 60, NatF catalytic subunit [Source:MGI Symbol;Acc:MGI:1922013] | 1109,2 | 86,22 |
| ENSMUSG00000078919 | Dpm1 | dolichol-phosphate (beta-D) mannosyltransferase 1 [Source:MGI Symbol;Acc:MGI:1330239] | 1109,1 | 91,83 |
| ENSMUSG00000021824 | Ap3m1 | adaptor-related protein complex 3, mu 1 subunit [Source:MGI Symbol;Acc:MGI:1929212] | 1108,8 | 92,88 |
| ENSMUSG00000026112 | Coa5 | cytochrome C oxidase assembly factor 5 [Source:MGI Symbol;Acc:MGI:1923428] | 1108,4 | 98,96 |
| ENSMUSG00000028484 | Psip1 | PC4 and SFRS1 interacting protein 1 [Source:MGI Symbol;Acc:MGI:2142116] | 1106,4 | 94,06 |
| ENSMUSG00000038206 | Fbxo8 | F-box protein 8 [Source:MGI Symbol;Acc:MGI:1354696] | 1105,8 | 99,82 |
| ENSMUSG00000056770 | Setd3 | SET domain containing 3 [Source:MGI Symbol;Acc:MGI:1289184] | 1105,3 | 71,25 |
| ENSMUSG00000019795 | Pcmt1 | protein-L-isoaspartate (D-aspartate) O-methyltransferase 1 [Source:MGI Symbol;Acc:MGI:97502] | 1104,9 | 96,30 |
| ENSMUSG00000035953 | Tmem55b | transmembrane protein 55b [Source:MGI Symbol;Acc:MGI:2448501] | 1103,4 | 88,10 |
| ENSMUSG00000053310 | Nrgn | neurogranin [Source:MGI Symbol;Acc:MGI:1927184] | 1102,7 | 97,22 |
| ENSMUSG00000028851 | Nudc | nuclear distribution gene C homolog (Aspergillus) [Source:MGI Symbol;Acc:MGI:106014] | 1099,7 | 100,00 |
| ENSMUSG00000022884 | Eif4a2 | eukaryotic translation initiation factor 4A2 [Source:MGI Symbol;Acc:MGI:106906] | 1099,6 | 77,36 |

| | | | | |
|---|---|---|---|---|
| ENSMUSG00000028879 | Stx12 | syntaxin 12 [Source:MGI Symbol;Acc:MGI:1931027] | 1098,7 | 99,76 |
| ENSMUSG00000055681 | Cope | coatomer protein complex, subunit epsilon [Source:MGI Symbol;Acc:MGI:1891702] | 1098,5 | 93,30 |
| ENSMUSG00000021963 | Sap18 | Sin3-associated polypeptide 18 [Source:MGI Symbol;Acc:MGI:1277978] | 1096,3 | 88,03 |
| ENSMUSG00000002107 | Celf2 | CUGBP, Elav-like family member 2 [Source:MGI Symbol;Acc:MGI:1338822] | 1094,8 | 74,15 |
| ENSMUSG00000032393 | Dpp8 | dipeptidylpeptidase 8 [Source:MGI Symbol;Acc:MGI:1921638] | 1093,8 | 98,72 |
| ENSMUSG00000025410 | Dctn2 | dynactin 2 [Source:MGI Symbol;Acc:MGI:107733] | 1092,8 | 99,46 |
| ENSMUSG00000044600 | Smim7 | small integral membrane protein 7 [Source:MGI Symbol;Acc:MGI:1914068] | 1091,3 | 97,28 |
| ENSMUSG00000028719 | Cmpk1 | cytidine monophosphate (UMP-CMP) kinase 1 [Source:MGI Symbol;Acc:MGI:1913838] | 1091,3 | 76,98 |
| ENSMUSG00000029467 | Atp2a2 | ATPase, Ca++ transporting, cardiac muscle, slow twitch 2 [Source:MGI Symbol;Acc:MGI:88110] | 1090,6 | 99,46 |
| ENSMUSG00000022051 | Bnip3l | BCL2/adenovirus E1B interacting protein 3-like [Source:MGI Symbol;Acc:MGI:1332659] | 1090,4 | 89,08 |
| ENSMUSG00000030706 | Mrpl48 | mitochondrial ribosomal protein L48 [Source:MGI Symbol;Acc:MGI:1289321] | 1087,1 | 93,87 |
| ENSMUSG00000029413 | Naaa | N-acylethanolamine acid amidase [Source:MGI Symbol;Acc:MGI:1914361] | 1084,6 | 94,48 |
| ENSMUSG00000017615 | Tnfaip1 | tumor necrosis factor, alpha-induced protein 1 (endothelial) [Source:MGI Symbol;Acc:MGI:104961] | 1084,2 | 99,23 |
| ENSMUSG00000063888 | Rpl7l1 | ribosomal protein L7-like 1 [Source:MGI Symbol;Acc:MGI:1913479] | 1079,5 | 99,48 |
| ENSMUSG00000017734 | Dbndd2 | dysbindin (dystrobrevin binding protein 1) domain containing 2 [Source:MGI Symbol;Acc:MGI:106562] | 1079,1 | 80,00 |
| ENSMUSG00000035478 | Mbd3 | methyl-CpG binding domain protein 3 [Source:MGI Symbol;Acc:MGI:1333812] | 1078,1 | 90,56 |
| ENSMUSG00000039067 | Psmd7 | proteasome (prosome, macropain) 26S subunit, non-ATPase, 7 [Source:MGI Symbol;Acc:MGI:1351511] | 1076,0 | 98,98 |
| ENSMUSG00000062797 | l7Rn6 | lethal, Chr 7, Rinchik 6 [Source:MGI Symbol;Acc:MGI:96738] | 1075,0 | 81,91 |
| ENSMUSG00000078652 | Psme3 | teaseome (prosome, macropain) activator subunit 3 (PA28 gamma, Ki) [Source:MGI Symbol;Acc:MGI:10963 | 1074,8 | 96,59 |
| ENSMUSG00000038034 | Igsf8 | immunoglobulin superfamily, member 8 [Source:MGI Symbol;Acc:MGI:2154090] | 1074,4 | 76,16 |
| ENSMUSG00000020444 | Guk1 | guanylate kinase 1 [Source:MGI Symbol;Acc:MGI:95871] | 1073,4 | 92,51 |
| ENSMUSG00000014504 | Srp19 | signal recognition particle 19 [Source:MGI Symbol;Acc:MGI:1913634] | 1073,2 | 99,77 |
| ENSMUSG00000033111 | 3830406C13Rik | RIKEN cDNA 3830406C13 gene [Source:MGI Symbol;Acc:MGI:1917937] | 1073,1 | 99,85 |
| ENSMUSG00000029430 | Ran | RAN, member RAS oncogene family [Source:MGI Symbol;Acc:MGI:1333112] | 1072,6 | 83,78 |
| ENSMUSG00000027248 | Pdia3 | protein disulfide isomerase associated 3 [Source:MGI Symbol;Acc:MGI:95834] | 1071,7 | 86,65 |
| ENSMUSG00000031347 | Cetn2 | centrin 2 [Source:MGI Symbol;Acc:MGI:1347085] | 1071,1 | 86,51 |
| ENSMUSG00000023909 | Paqr4 | progestin and adipoQ receptor family member IV [Source:MGI Symbol;Acc:MGI:1923748] | 1070,4 | 99,96 |
| ENSMUSG00000021134 | Srsf5 | serine/arginine-rich splicing factor 5 [Source:MGI Symbol;Acc:MGI:98287] | 1069,5 | 93,24 |
| ENSMUSG00000073988 | Ttpa | tocopherol (alpha) transfer protein [Source:MGI Symbol;Acc:MGI:1354168] | 1069,2 | 95,49 |
| ENSMUSG00000027774 | Gfm1 | G elongation factor, mitochondrial 1 [Source:MGI Symbol;Acc:MGI:107339] | 1068,8 | 96,61 |
| ENSMUSG00000040652 | Oaz2 | ornithine decarboxylase antizyme 2 [Source:MGI Symbol;Acc:MGI:109492] | 1066,7 | 76,25 |
| ENSMUSG00000005779 | Psmb4 | proteasome (prosome, macropain) subunit, beta type 4 [Source:MGI Symbol;Acc:MGI:1098257] | 1066,5 | 85,52 |
| ENSMUSG00000059336 | Slc14a1 | solute carrier family 14 (urea transporter), member 1 [Source:MGI Symbol;Acc:MGI:1351654] | 1066,1 | 95,88 |
| ENSMUSG00000028436 | Dcaf12 | DDB1 and CUL4 associated factor 12 [Source:MGI Symbol;Acc:MGI:1916220] | 1063,8 | 99,68 |
| ENSMUSG00000053929 | Cyhr1 | cysteine and histidine rich 1 [Source:MGI Symbol;Acc:MGI:1859320] | 1063,4 | 91,32 |
| ENSMUSG00000028822 | Tmem50a | transmembrane protein 50A [Source:MGI Symbol;Acc:MGI:1919067] | 1063,4 | 99,74 |
| ENSMUSG00000032745 | Gpbp1 | GC-rich promoter binding protein 1 [Source:MGI Symbol;Acc:MGI:1920524] | 1062,9 | 93,48 |
| ENSMUSG00000037196 | Pacrg | PARK2 co-regulated [Source:MGI Symbol;Acc:MGI:1916560] | 1062,9 | 97,24 |
| ENSMUSG00000042032 | Mat2b | methionine adenosyltransferase II, beta [Source:MGI Symbol;Acc:MGI:1913667] | 1062,1 | 87,38 |
| ENSMUSG00000002274 | Metrn | meteorin, glial cell differentiation regulator [Source:MGI Symbol;Acc:MGI:1917333] | 1061,3 | 98,05 |
| ENSMUSG00000041548 | Hspb8 | heat shock protein 8 [Source:MGI Symbol;Acc:MGI:2135756] | 1059,1 | 99,87 |
| ENSMUSG00000023460 | Rab12 | RAB12, member RAS oncogene family [Source:MGI Symbol;Acc:MGI:894284] | 1059,0 | 99,98 |
| ENSMUSG00000013622 | Atraid | all-trans retinoic acid induced differentiation factor [Source:MGI Symbol;Acc:MGI:1918918] | 1057,6 | 80,22 |
| ENSMUSG00000102234 | Gm37885 | predicted gene, 37885 [Source:MGI Symbol;Acc:MGI:5611113] | 1057,5 | 99,95 |
| ENSMUSG00000026753 | Ppp6c | protein phosphatase 6, catalytic subunit [Source:MGI Symbol;Acc:MGI:1915107] | 1057,3 | 97,46 |
| ENSMUSG00000025791 | Pgm2 | phosphoglucomutase 2 [Source:MGI Symbol;Acc:MGI:97565] | 1056,9 | 80,76 |
| ENSMUSG00000021248 | Tmed10 | transmembrane emp24-like trafficking protein 10 (yeast) [Source:MGI Symbol;Acc:MGI:1915831] | 1056,8 | 91,49 |
| ENSMUSG00000060036 | Rpl3 | ribosomal protein L3 [Source:MGI Symbol;Acc:MGI:1351605] | 1055,9 | 81,75 |
| ENSMUSG00000030061 | Uba3 | ubiquitin-like modifier activating enzyme 3 [Source:MGI Symbol;Acc:MGI:1341217] | 1055,9 | 96,63 |
| ENSMUSG00000027180 | Fbxo3 | F-box protein 3 [Source:MGI Symbol;Acc:MGI:1929084] | 1055,5 | 78,74 |
| ENSMUSG00000020366 | Mapk9 | mitogen-activated protein kinase 9 [Source:MGI Symbol;Acc:MGI:1346862] | 1051,5 | 100,00 |
| ENSMUSG00000027552 | E2f5 | E2F transcription factor 5 [Source:MGI Symbol;Acc:MGI:105091] | 1051,3 | 99,94 |
| ENSMUSG00000020541 | Tom1l1 | target of myb1-like 1 (chicken) [Source:MGI Symbol;Acc:MGI:1919193] | 1050,9 | 75,33 |
| ENSMUSG00000056820 | Tsnax | translin-associated factor X [Source:MGI Symbol;Acc:MGI:1855672] | 1050,5 | 98,90 |
| ENSMUSG00000008036 | Ap2s1 | adaptor-related protein complex 2, sigma 1 subunit [Source:MGI Symbol;Acc:MGI:2141861] | 1048,8 | 99,96 |
| ENSMUSG00000033577 | Myo6 | myosin VI [Source:MGI Symbol;Acc:MGI:104785] | 1048,7 | 99,03 |
| ENSMUSG00000085587 | Gm14493 | predicted gene 14493 [Source:MGI Symbol;Acc:MGI:3705313] | 1048,5 | 100,00 |
| ENSMUSG00000085425 | Gm15648 | predicted gene 15648 [Source:MGI Symbol;Acc:MGI:3783092] | 1047,7 | 100,00 |
| ENSMUSG00000006010 | BC003331 | cDNA sequence BC003331 [Source:MGI Symbol;Acc:MGI:2385108] | 1047,6 | 97,37 |
| ENSMUSG00000003949 | Hlf | hepatic leukemia factor [Source:MGI Symbol;Acc:MGI:96108] | 1046,6 | 95,47 |
| ENSMUSG00000027746 | Ufm1 | ubiquitin-fold modifier 1 [Source:MGI Symbol;Acc:MGI:1915140] | 1046,2 | 94,10 |
| ENSMUSG00000022307 | Oxr1 | oxidation resistance 1 [Source:MGI Symbol;Acc:MGI:2179326] | 1044,2 | 92,90 |
| ENSMUSG00000020388 | Pdlim4 | PDZ and LIM domain 4 [Source:MGI Symbol;Acc:MGI:1353470] | 1043,8 | 92,63 |
| ENSMUSG00000040234 | Tm7sf3 | transmembrane 7 superfamily member 3 [Source:MGI Symbol;Acc:MGI:1914873] | 1043,4 | 99,58 |
| ENSMUSG00000050423 | Ppp1r3g | protein phosphatase 1, regulatory (inhibitor) subunit 3G [Source:MGI Symbol;Acc:MGI:1923737] | 1043,3 | 99,76 |
| ENSMUSG00000039515 | Ppp2r4 | protein phosphatase 2A activator, regulatory subunit B [Source:MGI Symbol;Acc:MGI:1346006] | 1043,3 | 92,25 |
| ENSMUSG00000022427 | Tomm22 | translocase of outer mitochondrial membrane 22 homolog (yeast) [Source:MGI Symbol;Acc:MGI:2450248] | 1042,9 | 99,94 |
| ENSMUSG00000031453 | Rasa3 | RAS p21 protein activator 3 [Source:MGI Symbol;Acc:MGI:1197013] | 1042,4 | 96,98 |
| ENSMUSG00000031683 | Lsm6 | LSM6 homolog, U6 small nuclear RNA associated (S. cerevisiae) [Source:MGI Symbol;Acc:MGI:1925901] | 1042,1 | 98,96 |
| ENSMUSG00000034165 | Ccnd3 | cyclin D3 [Source:MGI Symbol;Acc:MGI:88315] | 1042,0 | 84,01 |
| ENSMUSG00000015095 | Fbxw5 | F-box and WD-40 domain protein 5 [Source:MGI Symbol;Acc:MGI:1354731] | 1041,2 | 99,13 |
| ENSMUSG00000026021 | Sumo1 | small ubiquitin-like modifier 1 [Source:MGI Symbol;Acc:MGI:1197010] | 1041,2 | 78,30 |
| ENSMUSG00000048747 | E130114P18Rik | RIKEN cDNA E130114P18 gene [Source:MGI Symbol;Acc:MGI:2442873] | 1040,8 | 96,98 |
| ENSMUSG00000062373 | Tmem65 | transmembrane protein 65 [Source:MGI Symbol;Acc:MGI:1922118] | 1040,6 | 99,42 |
| ENSMUSG00000068039 | Tcp1 | t-complex protein 1 [Source:MGI Symbol;Acc:MGI:98535] | 1040,4 | 93,60 |
| ENSMUSG00000008575 | Nfib | nuclear factor I/B [Source:MGI Symbol;Acc:MGI:103188] | 1040,0 | 91,63 |
| ENSMUSG00000009894 | Snap47 | synaptosomal-associated protein, 47 [Source:MGI Symbol;Acc:MGI:1915076] | 1039,5 | 78,07 |
| ENSMUSG00000036112 | Metap2 | methionine aminopeptidase 2 [Source:MGI Symbol;Acc:MGI:1929701] | 1038,8 | 85,44 |
| ENSMUSG00000102422 | Schip1 | schwannomin interacting protein 1 [Source:MGI Symbol;Acc:MGI:1353557] | 1038,6 | 82,95 |
| ENSMUSG00000051515 | Fam181b | family with sequence similarity 181, member B [Source:MGI Symbol;Acc:MGI:1930951] | 1038,5 | 92,05 |
| ENSMUSG00000026575 | Nme7 | NME/NM23 family member 7 [Source:MGI Symbol;Acc:MGI:2449121] | 1037,8 | 97,71 |
| ENSMUSG00000030495 | Slc7a10 | e carrier family 7 (cationic amino acid transporter, y+ system), member 10 [Source:MGI Symbol;Acc:MGI:1858 | 1037,6 | 94,99 |
| ENSMUSG00000008859 | Rala | v-ral simian leukemia viral oncogene homolog A (ras related) [Source:MGI Symbol;Acc:MGI:1927243] | 1036,3 | 99,74 |
| ENSMUSG00000093989 | Rnasek | ribonuclease, RNase K [Source:MGI Symbol;Acc:MGI:106369] | 1035,2 | 93,59 |
| ENSMUSG00000032014 | Oaf | OAF homolog (Drosophila) [Source:MGI Symbol;Acc:MGI:94852] | 1034,9 | 98,62 |
| ENSMUSG00000019232 | Etnppl | ethanolamine phosphate phospholyase [Source:MGI Symbol;Acc:MGI:1919010] | 1033,1 | 97,24 |
| ENSMUSG00000032381 | Fam96a | family with sequence similarity 96, member A [Source:MGI Symbol;Acc:MGI:1915500] | 1031,1 | 99,87 |
| ENSMUSG00000026782 | Abi2 | abl-interactor 2 [Source:MGI Symbol;Acc:MGI:106913] | 1030,6 | 99,36 |
| ENSMUSG00000029152 | Ociad1 | OCIA domain containing 1 [Source:MGI Symbol;Acc:MGI:1915345] | 1030,0 | 80,35 |
| ENSMUSG00000021687 | Scamp1 | secretory carrier membrane protein 1 [Source:MGI Symbol;Acc:MGI:1349480] | 1029,4 | 86,54 |
| ENSMUSG00000025979 | Mob4 | MOB family member 4, phocein [Source:MGI Symbol;Acc:MGI:104899] | 1027,9 | 92,63 |
| ENSMUSG00000025486 | Sirt3 | sirtuin 3 [Source:MGI Symbol;Acc:MGI:1927665] | 1027,7 | 99,79 |
| ENSMUSG00000013663 | Pten | phosphatase and tensin homolog [Source:MGI Symbol;Acc:MGI:109583] | 1027,4 | 84,10 |
| ENSMUSG00000003099 | Ppp5c | protein phosphatase 5, catalytic subunit [Source:MGI Symbol;Acc:MGI:102666] | 1026,7 | 99,42 |
| ENSMUSG00000005034 | Prkacb | protein kinase, cAMP dependent, catalytic, beta [Source:MGI Symbol;Acc:MGI:97594] | 1026,6 | 87,87 |
| ENSMUSG00000016637 | Ift27 | intraflagellar transport 27 [Source:MGI Symbol;Acc:MGI:1914292] | 1026,2 | 100,00 |
| ENSMUSG00000003814 | Calr | calreticulin [Source:MGI Symbol;Acc:MGI:88252] | 1026,1 | 87,70 |
| ENSMUSG00000028447 | Dctn3 | dynactin 3 [Source:MGI Symbol;Acc:MGI:1859251] | 1025,5 | 89,24 |
| ENSMUSG00000021071 | Trim9 | tripartite motif-containing 9 [Source:MGI Symbol;Acc:MGI:2137354] | 1024,7 | 84,34 |
| ENSMUSG00000025337 | Sbds | Shwachman-Bodian-Diamond syndrome homolog (human) [Source:MGI Symbol;Acc:MGI:1913961] | 1023,8 | 78,73 |
| ENSMUSG00000025950 | Idh1 | isocitrate dehydrogenase 1 (NADP+), soluble [Source:MGI Symbol;Acc:MGI:96413] | 1023,8 | 93,28 |
| ENSMUSG00000020647 | Ncoa1 | nuclear receptor coactivator 1 [Source:MGI Symbol;Acc:MGI:1276523] | 1023,7 | 99,40 |
| ENSMUSG00000032418 | Me1 | malic enzyme 1, NADP(+)-dependent, cytosolic [Source:MGI Symbol;Acc:MGI:97043] | 1023,6 | 75,57 |
| ENSMUSG00000027309 | 4930402H24Rik | RIKEN cDNA 4930402H24 gene [Source:MGI Symbol;Acc:MGI:1923029] | 1021,0 | 98,02 |
| ENSMUSG00000047242 | Taf9b | 9B RNA polymerase II, TATA box binding protein (TBP)-associated factor [Source:MGI Symbol;Acc:MGI:3039 | 1020,8 | 83,10 |
| ENSMUSG00000037235 | Mxd4 | Max dimerization protein 4 [Source:MGI Symbol;Acc:MGI:104991] | 1020,5 | 99,18 |
| ENSMUSG00000021076 | Actr10 | ARP10 actin-related protein 10 [Source:MGI Symbol;Acc:MGI:1891654] | 1019,7 | 93,32 |
| ENSMUSG00000026821 | Ralgds | ral guanine nucleotide dissociation stimulator [Source:MGI Symbol;Acc:MGI:107485] | 1017,6 | 95,02 |
| ENSMUSG00000026154 | Sdhaf4 | succinate dehydrogenase complex assembly factor 4 [Source:MGI Symbol;Acc:MGI:1915252] | 1014,5 | 99,66 |
| ENSMUSG00000048148 | Nwd1 | NACHT and WD repeat domain containing 1 [Source:MGI Symbol;Acc:MGI:2442268] | 1013,9 | 92,39 |
| ENSMUSG00000027993 | Trim2 | tripartite motif-containing 2 [Source:MGI Symbol;Acc:MGI:1933163] | 1012,8 | 91,02 |
| ENSMUSG00000022261 | Sdc2 | syndecan 2 [Source:MGI Symbol;Acc:MGI:1349165] | 1012,6 | 99,08 |
| ENSMUSG00000030868 | Dctn5 | dynactin 5 [Source:MGI Symbol;Acc:MGI:1891689] | 1012,4 | 100,00 |
| ENSMUSG00000026356 | Dars | aspartyl-tRNA synthetase [Source:MGI Symbol;Acc:MGI:2442544] | 1012,1 | 88,74 |
| ENSMUSG00000000441 | Raf1 | v-raf-leukemia viral oncogene 1 [Source:MGI Symbol;Acc:MGI:97847] | 1010,1 | 94,94 |
| ENSMUSG00000031392 | Irak1 | interleukin-1 receptor-associated kinase 1 [Source:MGI Symbol;Acc:MGI:107420] | 1008,9 | 99,60 |
| ENSMUSG00000078566 | Bnip3 | BCL2/adenovirus E1B interacting protein 3 [Source:MGI Symbol;Acc:MGI:109326] | 1008,8 | 88,00 |

| | | | | |
|---|---|---|---|---|
| ENSMUSG00000020536 | Llgl1 | lethal giant larvae homolog 1 (Drosophila) [Source:MGI Symbol;Acc:MGI:102682] | 1008,7 | 99,84 |
| ENSMUSG00000018574 | Acadvl | acyl-Coenzyme A dehydrogenase, very long chain [Source:MGI Symbol;Acc:MGI:895149] | 1008,5 | 99,88 |
| ENSMUSG00000056429 | Tgoln1 | trans-golgi network protein [Source:MGI Symbol;Acc:MGI:105080] | 1006,7 | 96,13 |
| ENSMUSG00000074129 | Rpl13a | ribosomal protein L13A [Source:MGI Symbol;Acc:MGI:1351455] | 1005,4 | 100,00 |
| ENSMUSG00000023904 | Hcfc1r1 | host cell factor C1 regulator 1 (XPO1-dependent) [Source:MGI Symbol;Acc:MGI:2663619] | 1004,4 | 100,00 |
| ENSMUSG00000025228 | Actr1a | ARP1 actin-related protein 1A, centractin alpha [Source:MGI Symbol;Acc:MGI:1858964] | 1004,1 | 100,00 |
| ENSMUSG00000033423 | Eri3 | exoribonuclease 3 [Source:MGI Symbol;Acc:MGI:2153887] | 1003,7 | 71,45 |
| ENSMUSG00000023353 | Agap3 | ArfGAP with GTPase domain, ankyrin repeat and PH domain 3 [Source:MGI Symbol;Acc:MGI:2183446] | 1001,7 | 82,17 |
| ENSMUSG00000040415 | Dtx3 | deltex 3, E3 ubiquitin ligase [Source:MGI Symbol;Acc:MGI:2135752] | 1000,5 | 97,43 |
| ENSMUSG00000021536 | Adcy2 | adenylate cyclase 2 [Source:MGI Symbol;Acc:MGI:99676] | 1000,2 | 99,09 |
| ENSMUSG00000020857 | Nme2 | NME/NM23 nucleoside diphosphate kinase 2 [Source:MGI Symbol;Acc:MGI:97356] | 999,5 | 83,80 |
| ENSMUSG00000015224 | Cyp2j9 | cytochrome P450, family 2, subfamily j, polypeptide 9 [Source:MGI Symbol;Acc:MGI:1921769] | 999,4 | 100,00 |
| ENSMUSG00000025066 | Sfr1 | SWI5 dependent recombination repair 1 [Source:MGI Symbol;Acc:MGI:1915038] | 998,9 | 99,60 |
| ENSMUSG00000061111 | Fam195b | family with sequence similarity 195, member B [Source:MGI Symbol;Acc:MGI:2384752] | 998,4 | 99,86 |
| ENSMUSG00000045128 | Rpl18a | ribosomal protein L18A [Source:MGI Symbol;Acc:MGI:1924058] | 998,3 | 75,99 |
| ENSMUSG00000053329 | D10Jhu81e | DNA segment, Chr 10, Johns Hopkins University 81 expressed [Source:MGI Symbol;Acc:MGI:1351861] | 998,2 | 100,00 |
| ENSMUSG00000019803 | Nr2e1 | nuclear receptor subfamily 2, group E, member 1 [Source:MGI Symbol;Acc:MGI:1100526] | 997,3 | 91,57 |
| ENSMUSG00000024143 | Rhoq | ras homolog gene family, member Q [Source:MGI Symbol;Acc:MGI:1931553] | 995,5 | 95,49 |
| ENSMUSG00000037720 | Tmem33 | transmembrane protein 33 [Source:MGI Symbol;Acc:MGI:1915128] | 995,2 | 94,47 |
| ENSMUSG00000011179 | Odc1 | ornithine decarboxylase, structural 1 [Source:MGI Symbol;Acc:MGI:97402] | 993,6 | 91,41 |
| ENSMUSG00000028565 | Nfia | nuclear factor I/A [Source:MGI Symbol;Acc:MGI:108056] | 993,3 | 97,89 |
| ENSMUSG00000007892 | Rplp1 | ribosomal protein, large, P1 [Source:MGI Symbol;Acc:MGI:1927099] | 992,1 | 100,00 |
| ENSMUSG00000017707 | Serinc3 | serine incorporator 3 [Source:MGI Symbol;Acc:MGI:1349457] | 991,8 | 98,21 |
| ENSMUSG00000091556 | Gm14569 | predicted gene 14569 [Source:MGI Symbol;Acc:MGI:3705243] | 991,8 | 78,11 |
| ENSMUSG00000022332 | Khdrbs3 | (H domain containing, RNA binding, signal transduction associated 3 [Source:MGI Symbol;Acc:MGI:1313312 | 991,3 | 98,60 |
| ENSMUSG00000029578 | Wipi2 | WD repeat domain, phosphoinositide interacting 2 [Source:MGI Symbol;Acc:MGI:1923831] | 989,8 | 83,80 |
| ENSMUSG00000031785 | Adgrg1 | adhesion G protein-coupled receptor G1 [Source:MGI Symbol;Acc:MGI:1340051] | 985,3 | 93,10 |
| ENSMUSG00000001313 | Rnd2 | Rho family GTPase 2 [Source:MGI Symbol;Acc:MGI:1338755] | 985,2 | 98,90 |
| ENSMUSG00000032038 | St3gal4 | ST3 beta-galactoside alpha-2,3-sialyltransferase 4 [Source:MGI Symbol;Acc:MGI:1316743] | 984,0 | 99,97 |
| ENSMUSG00000031950 | Gabarapl2 | amma-aminobutyric acid (GABA) A receptor-associated protein-like 2 [Source:MGI Symbol;Acc:MGI:1890602 | 983,8 | 75,95 |
| ENSMUSG00000018398 | Sept8 | septin 8 [Source:MGI Symbol;Acc:MGI:894310] | 983,7 | 96,65 |
| ENSMUSG00000024208 | Uqcc2 | ubiquinol-cytochrome c reductase complex assembly factor 2 [Source:MGI Symbol;Acc:MGI:1914517] | 982,4 | 81,37 |
| ENSMUSG00000002981 | Clptm1 | cleft lip and palate associated transmembrane protein 1 [Source:MGI Symbol;Acc:MGI:1927155] | 981,9 | 88,07 |
| ENSMUSG00000003528 | Slc25a1 | ite carrier family 25 (mitochondrial carrier, citrate transporter), member 1 [Source:MGI Symbol;Acc:MGI:13452 | 981,1 | 99,40 |
| ENSMUSG00000052428 | Tmco1 | transmembrane and coiled-coil domains 1 [Source:MGI Symbol;Acc:MGI:1921173] | 980,8 | 89,32 |
| ENSMUSG00000003970 | Rpl8 | ribosomal protein L8 [Source:MGI Symbol;Acc:MGI:1350927] | 979,0 | 99,54 |
| ENSMUSG00000002102 | Psmc3 | proteasome (prosome, macropain) 26S subunit, ATPase 3 [Source:MGI Symbol;Acc:MGI:1098754] | 978,9 | 86,64 |
| ENSMUSG00000060904 | Arl1 | ADP-ribosylation factor-like 1 [Source:MGI Symbol;Acc:MGI:99436] | 978,7 | 70,92 |
| ENSMUSG00000029048 | Rer1 | RER1 retention in endoplasmic reticulum 1 homolog (S. cerevisiae) [Source:MGI Symbol;Acc:MGI:1915080] | 977,6 | 99,87 |
| ENSMUSG00000019087 | Atp6ap1 | ATPase, H+ transporting, lysosomal accessory protein 1 [Source:MGI Symbol;Acc:MGI:109629] | 977,5 | 95,90 |
| ENSMUSG00000060073 | Psma3 | proteasome (prosome, macropain) subunit, alpha type 3 [Source:MGI Symbol;Acc:MGI:104883] | 977,5 | 85,42 |
| ENSMUSG00000025503 | Taldo1 | transaldolase 1 [Source:MGI Symbol;Acc:MGI:1274789] | 976,1 | 100,00 |
| ENSMUSG00000001383 | Zmat2 | zinc finger, matrin type 2 [Source:MGI Symbol;Acc:MGI:1913742] | 976,0 | 86,15 |
| ENSMUSG00000007721 | Ccdc124 | coiled-coil domain containing 124 [Source:MGI Symbol;Acc:MGI:1916403] | 975,4 | 100,00 |
| ENSMUSG00000048175 | Asb8 | ankyrin repeat and SOCS box-containing 8 [Source:MGI Symbol;Acc:MGI:1925791] | 974,1 | 90,33 |
| ENSMUSG00000039680 | Mrps6 | mitochondrial ribosomal protein S6 [Source:MGI Symbol;Acc:MGI:2153111] | 973,7 | 97,87 |
| ENSMUSG00000030007 | Cct7 | chaperonin containing Tcp1, subunit 7 (eta) [Source:MGI Symbol;Acc:MGI:107184] | 973,7 | 99,00 |
| ENSMUSG00000038722 | Bud31 | BUD31 homolog [Source:MGI Symbol;Acc:MGI:2141291] | 973,5 | 100,00 |
| ENSMUSG00000039194 | Rlbp1 | retinaldehyde binding protein 1 [Source:MGI Symbol;Acc:MGI:97930] | 973,0 | 89,06 |
| ENSMUSG00000021748 | Pdhb | pyruvate dehydrogenase (lipoamide) beta [Source:MGI Symbol;Acc:MGI:1915513] | 972,7 | 99,03 |
| ENSMUSG00000041959 | S100a10 | S100 calcium binding protein A10 (calpactin) [Source:MGI Symbol;Acc:MGI:1339468] | 972,7 | 85,41 |
| ENSMUSG00000028959 | Fastk | Fas-activated serine/threonine kinase [Source:MGI Symbol;Acc:MGI:1913837] | 971,6 | 88,12 |
| ENSMUSG00000015766 | Eps8 | epidermal growth factor receptor pathway substrate 8 [Source:MGI Symbol;Acc:MGI:104684] | 971,5 | 80,05 |
| ENSMUSG00000030447 | Cyfip1 | cytoplasmic FMR1 interacting protein 1 [Source:MGI Symbol;Acc:MGI:1338801] | 970,2 | 90,55 |
| ENSMUSG00000060240 | Cend1 | cell cycle exit and neuronal differentiation 1 [Source:MGI Symbol;Acc:MGI:1929898] | 969,3 | 99,19 |
| ENSMUSG00000046756 | Mrps7 | mitochondrial ribosomal protein S7 [Source:MGI Symbol;Acc:MGI:1354367] | 968,4 | 89,23 |
| ENSMUSG00000027221 | Chst1 | carbohydrate (keratan sulfate Gal-6) sulfotransferase 1 [Source:MGI Symbol;Acc:MGI:1924219] | 968,1 | 100,00 |
| ENSMUSG00000031353 | Rbbp7 | retinoblastoma binding protein 7 [Source:MGI Symbol;Acc:MGI:1194910] | 968,1 | 74,01 |
| ENSMUSG00000057469 | E2f6 | E2F transcription factor 6 [Source:MGI Symbol;Acc:MGI:1354159] | 968,0 | 100,00 |
| ENSMUSG00000036644 | Tbc1d9b | TBC1 domain family, member 9B [Source:MGI Symbol;Acc:MGI:1924045] | 967,9 | 99,63 |
| ENSMUSG00000048429 | 1810026J23Rik | RIKEN cDNA 1810026J23 gene [Source:MGI Symbol;Acc:MGI:1917023] | 967,7 | 97,26 |
| ENSMUSG00000021178 | Psmc1 | protease (prosome, macropain) 26S subunit, ATPase 1 [Source:MGI Symbol;Acc:MGI:106054] | 967,6 | 100,00 |
| ENSMUSG00000029613 | Eif2ak1 | eukaryotic translation initiation factor 2 alpha kinase 1 [Source:MGI Symbol;Acc:MGI:1353448] | 966,9 | 98,98 |
| ENSMUSG00000024780 | Cdc37l1 | cell division cycle 37-like 1 [Source:MGI Symbol;Acc:MGI:1914322] | 965,6 | 100,00 |
| ENSMUSG00000042978 | Sbk1 | SH3-binding kinase 1 [Source:MGI Symbol;Acc:MGI:2135937] | 964,8 | 98,83 |
| ENSMUSG00000024039 | Cbs | cystathionine beta-synthase [Source:MGI Symbol;Acc:MGI:88285] | 964,3 | 91,87 |
| ENSMUSG00000029463 | Fam216a | family with sequence similarity 216, member A [Source:MGI Symbol;Acc:MGI:1916198] | 964,2 | 99,88 |
| ENSMUSG00000029810 | Tmem176b | transmembrane protein 176B [Source:MGI Symbol;Acc:MGI:1916348] | 963,7 | 75,31 |
| ENSMUSG00000056999 | Ide | insulin degrading enzyme [Source:MGI Symbol;Acc:MGI:96412] | 963,6 | 99,90 |
| ENSMUSG00000052833 | Sae1 | SUMO1 activating enzyme subunit 1 [Source:MGI Symbol;Acc:MGI:1929264] | 962,7 | 100,00 |
| ENSMUSG00000090129 | Olfr287 | olfactory receptor 287 [Source:MGI Symbol;Acc:MGI:3030121] | 961,2 | 71,02 |
| ENSMUSG00000075701 | Vimp | VCP-interacting membrane protein [Source:MGI Symbol;Acc:MGI:95994] | 961,1 | 93,78 |
| ENSMUSG00000029103 | Lrpap1 | low density lipoprotein receptor-related protein associated protein 1 [Source:MGI Symbol;Acc:MGI:96829] | 960,2 | 98,97 |
| ENSMUSG00000106918 | Mrpl33 | mitochondrial ribosomal protein L33 [Source:MGI Symbol;Acc:MGI:2137225] | 960,1 | 99,37 |
| ENSMUSG00000029066 | Mrpl20 | mitochondrial ribosomal protein L20 [Source:MGI Symbol;Acc:MGI:2137221] | 959,9 | 80,92 |
| ENSMUSG00000023106 | Denr | density-regulated protein [Source:MGI Symbol;Acc:MGI:1915434] | 959,8 | 91,41 |
| ENSMUSG00000021796 | Bmpr1a | bone morphogenetic protein receptor, type 1A [Source:MGI Symbol;Acc:MGI:1338938] | 959,6 | 92,38 |
| ENSMUSG00000063457 | Rps15 | ribosomal protein S15 [Source:MGI Symbol;Acc:MGI:98117] | 959,3 | 90,16 |
| ENSMUSG00000022124 | Fbxl3 | F-box and leucine-rich repeat protein 3 [Source:MGI Symbol;Acc:MGI:1354702] | 959,3 | 88,42 |
| ENSMUSG00000024072 | Yipf4 | Yip1 domain family, member 4 [Source:MGI Symbol;Acc:MGI:1915114] | 958,1 | 100,00 |
| ENSMUSG00000028955 | Vamp3 | vesicle-associated membrane protein 3 [Source:MGI Symbol;Acc:MGI:1321389] | 957,9 | 99,73 |
| ENSMUSG00000024158 | Hagh | hydroxyacyl glutathione hydrolase [Source:MGI Symbol;Acc:MGI:95745] | 956,9 | 70,89 |
| ENSMUSG00000029577 | Ube3b | ubiquitin protein ligase E3B [Source:MGI Symbol;Acc:MGI:1891295] | 956,5 | 98,74 |
| ENSMUSG00000000581 | C1d | C1D nuclear receptor co-repressor [Source:MGI Symbol;Acc:MGI:1927354] | 953,7 | 79,39 |
| ENSMUSG00000092360 | Gm20441 | predicted gene 20441 [Source:MGI Symbol;Acc:MGI:5141906] | 952,7 | 100,00 |
| ENSMUSG00000026511 | Srp9 | signal recognition particle 9 [Source:MGI Symbol;Acc:MGI:1350930] | 951,4 | 88,62 |
| ENSMUSG00000029038 | Ssu72 | Ssu72 RNA polymerase II CTD phosphatase homolog (yeast) [Source:MGI Symbol;Acc:MGI:1916241] | 951,2 | 78,71 |
| ENSMUSG00000028295 | Smim8 | small integral membrane protein 8 [Source:MGI Symbol;Acc:MGI:1913541] | 951,1 | 96,70 |
| ENSMUSG00000028207 | Asph | aspartate-beta-hydroxylase [Source:MGI Symbol;Acc:MGI:1914186] | 949,7 | 87,73 |
| ENSMUSG00000030751 | Psma1 | proteasome (prosome, macropain) subunit, alpha type 1 [Source:MGI Symbol;Acc:MGI:1347005] | 949,7 | 98,86 |
| ENSMUSG00000032348 | Gsta4 | glutathione S-transferase, alpha 4 [Source:MGI Symbol;Acc:MGI:1309515] | 949,6 | 100,00 |
| ENSMUSG00000020440 | Arf5 | ADP-ribosylation factor 5 [Source:MGI Symbol;Acc:MGI:99434] | 949,5 | 99,94 |
| ENSMUSG00000024359 | Hspa9 | heat shock protein 9 [Source:MGI Symbol;Acc:MGI:96245] | 948,6 | 99,57 |
| ENSMUSG00000029778 | Adcyap1r1 | adenylate cyclase activating polypeptide 1 receptor 1 [Source:MGI Symbol;Acc:MGI:108449] | 948,5 | 96,22 |
| ENSMUSG00000028772 | Zcchc17 | zinc finger, CCHC domain containing 17 [Source:MGI Symbol;Acc:MGI:1919955] | 948,3 | 96,77 |
| ENSMUSG00000024978 | Gpam | glycerol-3-phosphate acyltransferase, mitochondrial [Source:MGI Symbol;Acc:MGI:109162] | 947,9 | 99,36 |
| ENSMUSG00000022898 | Dscr3 | Down syndrome critical region gene 3 [Source:MGI Symbol;Acc:MGI:1206040] | 947,3 | 96,81 |
| ENSMUSG00000069763 | Tmem100 | transmembrane protein 100 [Source:MGI Symbol;Acc:MGI:1915138] | 947,0 | 84,53 |
| ENSMUSG00000002602 | Axl | AXL receptor tyrosine kinase [Source:MGI Symbol;Acc:MGI:1347244] | 945,3 | 99,83 |
| ENSMUSG00000025940 | Tmem70 | transmembrane protein 70 [Source:MGI Symbol;Acc:MGI:1915068] | 945,2 | 97,28 |
| ENSMUSG00000034781 | Gna11 | guanine nucleotide binding protein, alpha 11 [Source:MGI Symbol;Acc:MGI:95766] | 944,8 | 99,53 |
| ENSMUSG00000023156 | Rpp14 | ribonuclease P 14 subunit [Source:MGI Symbol;Acc:MGI:1914303] | 943,2 | 100,00 |
| ENSMUSG00000028070 | Apoa1bp | apolipoprotein A-I binding protein [Source:MGI Symbol;Acc:MGI:2180167] | 943,1 | 100,00 |
| ENSMUSG00000021681 | Aggf1 | angiogenic factor with G patch and FHA domains 1 [Source:MGI Symbol;Acc:MGI:1913799] | 941,7 | 100,00 |
| ENSMUSG00000000827 | Tpd52l2 | tumor protein D52-like 2 [Source:MGI Symbol;Acc:MGI:1913564] | 941,5 | 86,74 |
| ENSMUSG00000033793 | Atp6v1h | ATPase, H+ transporting, lysosomal V1 subunit H [Source:MGI Symbol;Acc:MGI:1914864] | 941,4 | 98,15 |
| ENSMUSG00000011884 | Gltp | glycolipid transfer protein [Source:MGI Symbol;Acc:MGI:1929253] | 939,8 | 99,70 |
| ENSMUSG00000052459 | Atp6v1a | ATPase, H+ transporting, lysosomal V1 subunit A [Source:MGI Symbol;Acc:MGI:1201780] | 939,0 | 90,94 |
| ENSMUSG00000078429 | Ctdsp2 | )oxy-terminal domain, RNA polymerase II, polypeptide A) small phosphatase 2 [Source:MGI Symbol;Acc:MGI: | 938,7 | 73,81 |
| ENSMUSG00000026687 | Aldh9a1 | aldehyde dehydrogenase 9, subfamily A1 [Source:MGI Symbol;Acc:MGI:1861622] | 938,3 | 99,35 |
| ENSMUSG00000010054 | Tusc2 | tumor suppressor candidate 2 [Source:MGI Symbol;Acc:MGI:1931086] | 938,1 | 99,98 |
| ENSMUSG00000032112 | Trappc4 | trafficking protein particle complex 4 [Source:MGI Symbol;Acc:MGI:1926211] | 936,5 | 100,00 |
| ENSMUSG00000021987 | Mtmr6 | myotubularin related protein 6 [Source:MGI Symbol;Acc:MGI:2145637] | 936,4 | 99,08 |
| ENSMUSG00000061461 | Smim20 | small integral membrane protein 20 [Source:MGI Symbol;Acc:MGI:1913528] | 935,9 | 100,00 |
| ENSMUSG00000017404 | Rpl19 | ribosomal protein L19 [Source:MGI Symbol;Acc:MGI:98020] | 933,3 | 94,36 |
| ENSMUSG00000072214 | Sept5 | septin 5 [Source:MGI Symbol;Acc:MGI:1195461] | 931,9 | 99,23 |
| ENSMUSG00000027642 | Rpn2 | ribophorin II [Source:MGI Symbol;Acc:MGI:98085] | 931,3 | 98,86 |

| | | | | |
|---|---|---|---|---|
| ENSMUSG00000037351 | Actr1b | ARP1 actin-related protein 1B, centractin beta [Source:MGI Symbol;Acc:MGI:1917446] | 931,1 | 90,26 |
| ENSMUSG00000032878 | Ccdc85a | coiled-coil domain containing 85A [Source:MGI Symbol;Acc:MGI:2445069] | 931,1 | 98,03 |
| ENSMUSG00000015759 | Cnih1 | cornichon family AMPA receptor auxiliary protein 1 [Source:MGI Symbol;Acc:MGI:1277202] | 930,5 | 99,17 |
| ENSMUSG00000025968 | Ndufs1 | NADH dehydrogenase (ubiquinone) Fe-S protein 1 [Source:MGI Symbol;Acc:MGI:2443241] | 929,3 | 96,89 |
| ENSMUSG00000009630 | Ppp2cb | protein phosphatase 2 (formerly 2A), catalytic subunit, beta isoform [Source:MGI Symbol;Acc:MGI:1321161] | 927,8 | 99,05 |
| ENSMUSG00000056579 | Tug1 | taurine upregulated gene 1 [Source:MGI Symbol;Acc:MGI:2144114] | 927,5 | 75,92 |
| ENSMUSG00000022774 | Ncbp2 | nuclear cap binding protein subunit 2 [Source:MGI Symbol;Acc:MGI:1915342] | 927,0 | 99,57 |
| ENSMUSG00000032350 | Gclc | glutamate-cysteine ligase, catalytic subunit [Source:MGI Symbol;Acc:MGI:104990] | 926,5 | 99,81 |
| ENSMUSG00000000326 | Comt | catechol-O-methyltransferase [Source:MGI Symbol;Acc:MGI:88470] | 926,3 | 82,44 |
| ENSMUSG00000039953 | Clstn1 | calsyntenin 1 [Source:MGI Symbol;Acc:MGI:1929895] | 925,9 | 99,44 |
| ENSMUSG00000004264 | Phb2 | prohibitin 2 [Source:MGI Symbol;Acc:MGI:102520] | 925,6 | 98,14 |
| ENSMUSG00000051469 | Zfp24 | zinc finger protein 24 [Source:MGI Symbol;Acc:MGI:1929704] | 925,6 | 98,68 |
| ENSMUSG00000057092 | Fxyd3 | FXYD domain-containing ion transport regulator 3 [Source:MGI Symbol;Acc:MGI:107497] | 925,5 | 78,07 |
| ENSMUSG00000035597 | Prpf39 | PRP39 pre-mRNA processing factor 39 homolog (yeast) [Source:MGI Symbol;Acc:MGI:104602] | 924,6 | 99,78 |
| ENSMUSG00000029397 | Rchy1 | ring finger and CHY zinc finger domain containing 1 [Source:MGI Symbol;Acc:MGI:1915348] | 924,3 | 99,93 |
| ENSMUSG00000025173 | Wdr45b | WD repeat domain 45B [Source:MGI Symbol;Acc:MGI:1914090] | 923,8 | 97,30 |
| ENSMUSG00000014195 | Dnajc7 | DnaJ heat shock protein family (Hsp40) member C7 [Source:MGI Symbol;Acc:MGI:1928373] | 923,6 | 90,93 |
| ENSMUSG00000042305 | Tmem183a | transmembrane protein 183A [Source:MGI Symbol;Acc:MGI:1914729] | 922,6 | 97,97 |
| ENSMUSG00000028455 | Stoml2 | stomatin (Epb7.2)-like 2 [Source:MGI Symbol;Acc:MGI:1913842] | 921,8 | 83,05 |
| ENSMUSG00000028145 | Them4 | thioesterase superfamily member 4 [Source:MGI Symbol;Acc:MGI:1923028] | 921,8 | 98,37 |
| ENSMUSG00000001911 | Nfix | nuclear factor I/X [Source:MGI Symbol;Acc:MGI:97311] | 921,6 | 85,40 |
| ENSMUSG00000037031 | Tspan15 | tetraspanin 15 [Source:MGI Symbol;Acc:MGI:1917673] | 921,5 | 97,28 |
| ENSMUSG00000020978 | Klhdc2 | kelch domain containing 2 [Source:MGI Symbol;Acc:MGI:1916804] | 920,8 | 100,00 |
| ENSMUSG00000015002 | Efr3a | EFR3 homolog A [Source:MGI Symbol;Acc:MGI:1923990] | 919,5 | 83,61 |
| ENSMUSG00000006423 | C330007P06Rik | RIKEN cDNA C330007P06 gene [Source:MGI Symbol;Acc:MGI:1924894] | 918,9 | 75,09 |
| ENSMUSG00000028180 | Zranb2 | zinc finger, RAN-binding domain containing 2 [Source:MGI Symbol;Acc:MGI:1858211] | 916,7 | 95,26 |
| ENSMUSG00000030603 | Psmc4 | proteasome (prosome, macropain) 26S subunit, ATPase, 4 [Source:MGI Symbol;Acc:MGI:1346093] | 916,1 | 99,72 |
| ENSMUSG00000020794 | Ube2g1 | ubiquitin-conjugating enzyme E2G 1 [Source:MGI Symbol;Acc:MGI:1914378] | 916,0 | 99,58 |
| ENSMUSG00000068742 | Cry2 | cryptochrome 2 (photolyase-like) [Source:MGI Symbol;Acc:MGI:1270859] | 915,0 | 91,55 |
| ENSMUSG00000024194 | Cuta | cutA divalent cation tolerance homolog [Source:MGI Symbol;Acc:MGI:1914925] | 914,9 | 100,00 |
| ENSMUSG00000041577 | Prelp | proline arginine-rich end leucine-rich repeat [Source:MGI Symbol;Acc:MGI:2151110] | 914,7 | 100,00 |
| ENSMUSG00000035048 | Anapc13 | anaphase promoting complex subunit 13 [Source:MGI Symbol;Acc:MGI:1916260] | 913,9 | 95,39 |
| ENSMUSG00000062963 | Ufc1 | ubiquitin-fold modifier conjugating enzyme 1 [Source:MGI Symbol;Acc:MGI:1913405] | 912,8 | 98,52 |
| ENSMUSG00000005575 | Ube2m | ubiquitin-conjugating enzyme E2M [Source:MGI Symbol;Acc:MGI:108278] | 911,2 | 89,97 |
| ENSMUSG00000073481 | Marc2 | mitochondrial amidoxime reducing component 2 [Source:MGI Symbol;Acc:MGI:1914497] | 910,5 | 89,88 |
| ENSMUSG00000020463 | Smek2 | SMEK homolog 2, suppressor of mek1 (Dictyostelium) [Source:MGI Symbol;Acc:MGI:2144474] | 910,5 | 97,02 |
| ENSMUSG00000074781 | Ube2n | ubiquitin-conjugating enzyme E2N [Source:MGI Symbol;Acc:MGI:1934835] | 910,3 | 86,52 |
| ENSMUSG00000027488 | Snta1 | syntrophin, acidic 1 [Source:MGI Symbol;Acc:MGI:101772] | 910,2 | 99,24 |
| ENSMUSG00000028218 | Fam92a | family with sequence similarity 92, member A [Source:MGI Symbol;Acc:MGI:1915349] | 909,5 | 91,08 |
| ENSMUSG00000032171 | Pin1 | protein (peptidyl-prolyl cis/trans isomerase) NIMA-interacting 1 [Source:MGI Symbol;Acc:MGI:1346036] | 908,9 | 79,86 |
| ENSMUSG00000025162 | Csnk1d | casein kinase 1, delta [Source:MGI Symbol;Acc:MGI:1355272] | 908,5 | 97,66 |
| ENSMUSG00000005469 | Prkaca | protein kinase, cAMP dependent, catalytic, alpha [Source:MGI Symbol;Acc:MGI:97592] | 908,0 | 99,93 |
| ENSMUSG00000032398 | Snapc5 | small nuclear RNA activating complex, polypeptide 5 [Source:MGI Symbol;Acc:MGI:1914282] | 907,9 | 99,82 |
| ENSMUSG00000027184 | Caprin1 | cell cycle associated protein 1 [Source:MGI Symbol;Acc:MGI:1858234] | 907,0 | 88,95 |
| ENSMUSG00000044221 | Grsf1 | G-rich RNA sequence binding factor 1 [Source:MGI Symbol;Acc:MGI:106479] | 906,9 | 97,10 |
| ENSMUSG00000028223 | Decr1 | 2,4-dienoyl CoA reductase 1, mitochondrial [Source:MGI Symbol;Acc:MGI:1914710] | 905,7 | 96,87 |
| ENSMUSG00000031565 | Fgfr1 | fibroblast growth factor receptor 1 [Source:MGI Symbol;Acc:MGI:95522] | 905,4 | 90,65 |
| ENSMUSG00000000184 | Ccnd2 | cyclin D2 [Source:MGI Symbol;Acc:MGI:88314] | 905,0 | 94,55 |
| ENSMUSG00000004937 | Sgta | small glutamine-rich tetratricopeptide repeat (TPR)-containing, alpha [Source:MGI Symbol;Acc:MGI:1098703] | 904,8 | 100,00 |
| ENSMUSG00000021901 | Bap1 | Brca1 associated protein 1 [Source:MGI Symbol;Acc:MGI:1206586] | 904,6 | 99,16 |
| ENSMUSG00000067274 | Rplp0 | ribosomal protein, large, P0 [Source:MGI Symbol;Acc:MGI:1927636] | 904,4 | 77,99 |
| ENSMUSG00000020955 | Ap4s1 | adaptor-related protein complex AP-4, sigma 1 [Source:MGI Symbol;Acc:MGI:1337065] | 903,2 | 97,75 |
| ENSMUSG00000026490 | Cdc42bpa | CDC42 binding protein kinase alpha [Source:MGI Symbol;Acc:MGI:2441841] | 902,6 | 93,89 |
| ENSMUSG00000003037 | Rab8a | RAB8A, member RAS oncogene family [Source:MGI Symbol;Acc:MGI:96960] | 902,5 | 75,91 |
| ENSMUSG00000034390 | Cmip | c-Maf inducing protein [Source:MGI Symbol;Acc:MGI:1921690] | 902,3 | 84,91 |
| ENSMUSG00000020634 | Ubxn2a | UBX domain protein 2A [Source:MGI Symbol;Acc:MGI:2442310] | 901,7 | 93,55 |
| ENSMUSG00000002968 | Med25 | mediator complex subunit 25 [Source:MGI Symbol;Acc:MGI:1922863] | 901,6 | 82,56 |
| ENSMUSG00000031969 | Acad8 | acyl-Coenzyme A dehydrogenase family, member 8 [Source:MGI Symbol;Acc:MGI:1914198] | 900,4 | 99,32 |
| ENSMUSG00000063576 | Klhdc3 | kelch domain containing 3 [Source:MGI Symbol;Acc:MGI:2651568] | 900,2 | 99,09 |
| ENSMUSG00000029319 | Coq2 | coenzyme Q2 4-hydroxybenzoate polyprenyltransferase [Source:MGI Symbol;Acc:MGI:1919133] | 899,8 | 74,63 |
| ENSMUSG00000071072 | Ptges3 | prostaglandin E synthase 3 (cytosolic) [Source:MGI Symbol;Acc:MGI:1929282] | 899,8 | 84,90 |
| ENSMUSG00000030983 | Bccip | BRCA2 and CDKN1A interacting protein [Source:MGI Symbol;Acc:MGI:1913415] | 898,9 | 100,00 |
| ENSMUSG00000041935 | AW549877 | expressed sequence AW549877 [Source:MGI Symbol;Acc:MGI:2146232] | 898,3 | 99,85 |
| ENSMUSG00000032606 | Nicn1 | nicolin 1 [Source:MGI Symbol;Acc:MGI:1913507] | 898,2 | 99,95 |
| ENSMUSG00000005881 | Ergic3 | ERGIC and golgi 3 [Source:MGI Symbol;Acc:MGI:1913616] | 895,5 | 96,57 |
| ENSMUSG00000068299 | 1700019G17Rik | RIKEN cDNA 1700019G17 gene [Source:MGI Symbol;Acc:MGI:1922791] | 895,1 | 90,63 |
| ENSMUSG00000029538 | Srsf9 | serine/arginine-rich splicing factor 9 [Source:MGI Symbol;Acc:MGI:104896] | 894,9 | 100,00 |
| ENSMUSG00000031788 | Kifc3 | kinesin family member C3 [Source:MGI Symbol;Acc:MGI:109202] | 894,7 | 96,06 |
| ENSMUSG00000064115 | Cadm2 | cell adhesion molecule 2 [Source:MGI Symbol;Acc:MGI:2442722] | 894,6 | 94,01 |
| ENSMUSG00000061559 | Wdr61 | WD repeat domain 61 [Source:MGI Symbol;Acc:MGI:1917493] | 893,8 | 78,04 |
| ENSMUSG00000034210 | Efcab14 | EF-hand calcium binding domain 14 [Source:MGI Symbol;Acc:MGI:2442397] | 893,1 | 91,06 |
| ENSMUSG00000037049 | Smpd1 | sphingomyelin phosphodiesterase 1, acid lysosomal [Source:MGI Symbol;Acc:MGI:98325] | 891,5 | 99,28 |
| ENSMUSG00000029922 | Mkrn1 | makorin, ring finger protein, 1 [Source:MGI Symbol;Acc:MGI:1859353] | 890,8 | 91,62 |
| ENSMUSG00000031647 | Mfap3l | microfibrillar-associated protein 3-like [Source:MGI Symbol;Acc:MGI:1918556] | 890,5 | 93,11 |
| ENSMUSG00000003526 | Prodh | proline dehydrogenase [Source:MGI Symbol;Acc:MGI:97770] | 889,4 | 94,86 |
| ENSMUSG00000036093 | Arl5a | ADP-ribosylation factor-like 5A [Source:MGI Symbol;Acc:MGI:1922673] | 888,9 | 81,67 |
| ENSMUSG00000026889 | Rbm18 | RNA binding motif protein 18 [Source:MGI Symbol;Acc:MGI:1915139] | 888,0 | 97,87 |
| ENSMUSG00000004069 | Dnaja3 | DnaJ heat shock protein family (Hsp40) member A3 [Source:MGI Symbol;Acc:MGI:1933786] | 887,8 | 99,16 |
| ENSMUSG00000020614 | Fam20a | family with sequence similarity 20, member A [Source:MGI Symbol;Acc:MGI:2388266] | 887,6 | 100,00 |
| ENSMUSG00000006998 | Psmd2 | proteasome (prosome, macropain) 26S subunit, non-ATPase, 2 [Source:MGI Symbol;Acc:MGI:1096584] | 887,4 | 97,24 |
| ENSMUSG00000014959 | Gorasp2 | golgi reassembly stacking protein 2 [Source:MGI Symbol;Acc:MGI:2135962] | 887,3 | 77,92 |
| ENSMUSG00000028798 | Eif3i | eukaryotic translation initiation factor 3, subunit I [Source:MGI Symbol;Acc:MGI:1860763] | 886,4 | 99,26 |
| ENSMUSG00000029004 | Kmt2e | lysine (K)-specific methyltransferase 2E [Source:MGI Symbol;Acc:MGI:1924825] | 885,1 | 71,95 |
| ENSMUSG00000015937 | H2afy | H2A histone family, member Y [Source:MGI Symbol;Acc:MGI:1349392] | 883,2 | 96,49 |
| ENSMUSG00000030876 | Mettl9 | methyltransferase like 9 [Source:MGI Symbol;Acc:MGI:1914862] | 882,9 | 92,20 |
| ENSMUSG00000056121 | Fez2 | fasciculation and elongation protein zeta 2 (zygin II) [Source:MGI Symbol;Acc:MGI:2675856] | 882,5 | 99,54 |
| ENSMUSG00000044986 | Tst | thiosulfate sulfurtransferase, mitochondrial [Source:MGI Symbol;Acc:MGI:98852] | 881,3 | 99,24 |
| ENSMUSG00000020803 | Txndc17 | thioredoxin domain containing 17 [Source:MGI Symbol;Acc:MGI:1289248] | 880,8 | 99,86 |
| ENSMUSG00000029110 | Rnf4 | ring finger protein 4 [Source:MGI Symbol;Acc:MGI:1201691] | 879,7 | 90,22 |
| ENSMUSG00000075702 | Selm | selenoprotein M [Source:MGI Symbol;Acc:MGI:2149786] | 879,3 | 100,00 |
| ENSMUSG00000027286 | Lrrc57 | leucine rich repeat containing 57 [Source:MGI Symbol;Acc:MGI:1913856] | 878,2 | 99,95 |
| ENSMUSG00000032570 | Atp2c1 | ATPase, Ca++-sequestering [Source:MGI Symbol;Acc:MGI:1889008] | 877,4 | 79,88 |
| ENSMUSG00000042429 | Adora1 | adenosine A1 receptor [Source:MGI Symbol;Acc:MGI:99401] | 877,2 | 96,08 |
| ENSMUSG00000040430 | Pitpnc1 | phosphatidylinositol transfer protein, cytoplasmic 1 [Source:MGI Symbol;Acc:MGI:1919045] | 877,2 | 98,10 |
| ENSMUSG00000040331 | Nsmce4a | non-SMC element 4 homolog A (S. cerevisiae) [Source:MGI Symbol;Acc:MGI:1915122] | 877,2 | 100,00 |
| ENSMUSG00000003299 | Mrpl4 | mitochondrial ribosomal protein L4 [Source:MGI Symbol;Acc:MGI:2137210] | 877,1 | 99,66 |
| ENSMUSG00000024777 | Ppp2r5b | protein phosphatase 2, regulatory subunit B', beta [Source:MGI Symbol;Acc:MGI:2388480] | 876,9 | 99,47 |
| ENSMUSG00000043391 | 2510009E07Rik | RIKEN cDNA 2510009E07 gene [Source:MGI Symbol;Acc:MGI:1919440] | 876,7 | 99,61 |
| ENSMUSG00000030894 | Tpp1 | tripeptidyl peptidase I [Source:MGI Symbol;Acc:MGI:1336194] | 876,6 | 100,00 |
| ENSMUSG00000039128 | Cdc123 | cell division cycle 123 [Source:MGI Symbol;Acc:MGI:2138811] | 875,5 | 91,69 |
| ENSMUSG00000063077 | Kif1b | kinesin family member 1B [Source:MGI Symbol;Acc:MGI:108426] | 875,0 | 97,90 |
| ENSMUSG00000031916 | Cog8 | component of oligomeric golgi complex 8 [Source:MGI Symbol;Acc:MGI:2142885] | 874,8 | 85,39 |
| ENSMUSG00000004667 | Polr2e | polymerase (RNA) II (DNA directed) polypeptide E [Source:MGI Symbol;Acc:MGI:1913670] | 874,7 | 96,92 |
| ENSMUSG00000073725 | Lmbrd1 | LMBR1 domain containing 1 [Source:MGI Symbol;Acc:MGI:1915671] | 874,5 | 94,83 |
| ENSMUSG00000028919 | Arhgef19 | Rho guanine nucleotide exchange factor (GEF) 19 [Source:MGI Symbol;Acc:MGI:1925912] | 873,6 | 97,94 |
| ENSMUSG00000041607 | Mbp | myelin basic protein [Source:MGI Symbol;Acc:MGI:96925] | 873,4 | 73,72 |
| ENSMUSG00000024668 | Sdhaf2 | succinate dehydrogenase complex assembly factor 2 [Source:MGI Symbol;Acc:MGI:1913322] | 872,9 | 75,51 |
| ENSMUSG00000039640 | Mrpl12 | mitochondrial ribosomal protein L12 [Source:MGI Symbol;Acc:MGI:1926273] | 872,8 | 89,64 |
| ENSMUSG00000020029 | Nudt4 | nudix (nucleoside diphosphate linked moiety X)-type motif 4 [Source:MGI Symbol;Acc:MGI:1918457] | 871,8 | 99,97 |
| ENSMUSG00000022789 | Dnm1l | dynamin 1-like [Source:MGI Symbol;Acc:MGI:1921256] | 871,4 | 99,53 |
| ENSMUSG00000061032 | Rrp1 | ribosomal RNA processing 1 homolog (S. cerevisiae) [Source:MGI Symbol;Acc:MGI:1203500] | 870,6 | 98,49 |
| ENSMUSG00000048332 | Lhfp | lipoma HMGIC fusion partner [Source:MGI Symbol;Acc:MGI:1920048] | 870,4 | 99,70 |
| ENSMUSG00000035877 | Zhx3 | zinc fingers and homeoboxes 3 [Source:MGI Symbol;Acc:MGI:2444772] | 870,3 | 92,21 |
| ENSMUSG00000044018 | Mrpl50 | mitochondrial ribosomal protein L50 [Source:MGI Symbol;Acc:MGI:107329] | 869,4 | 99,95 |
| ENSMUSG00000021548 | Ccnh | cyclin H [Source:MGI Symbol;Acc:MGI:1913921] | 868,9 | 79,84 |
| ENSMUSG00000020328 | Nudcd2 | NudC domain containing 2 [Source:MGI Symbol;Acc:MGI:1277103] | 868,9 | 99,98 |
| ENSMUSG00000008305 | Tle1 | transducin-like enhancer of split 1, homolog of Drosophila E(spl) [Source:MGI Symbol;Acc:MGI:104636] | 868,3 | 94,64 |

| | | | | |
|---|---|---|---|---|
| ENSMUSG00000000605 | Clcn4 | chloride channel, voltage-sensitive 4 [Source:MGI Symbol;Acc:MGI:104571] | 868,0 | 99,79 |
| ENSMUSG00000008140 | Emc10 | ER membrane protein complex subunit 10 [Source:MGI Symbol;Acc:MGI:1916933] | 866,2 | 96,16 |
| ENSMUSG00000057130 | Txnl4a | thioredoxin-like 4A [Source:MGI Symbol;Acc:MGI:1351613] | 865,7 | 97,44 |
| ENSMUSG00000027193 | Api5 | apoptosis inhibitor 5 [Source:MGI Symbol;Acc:MGI:1888993] | 865,6 | 89,79 |
| ENSMUSG00000034484 | Snx2 | sorting nexin 2 [Source:MGI Symbol;Acc:MGI:1915054] | 864,5 | 98,38 |
| ENSMUSG00000042712 | Wbp5 | WW domain binding protein 5 [Source:MGI Symbol;Acc:MGI:109567] | 864,3 | 98,39 |
| ENSMUSG00000057388 | Mrpl18 | mitochondrial ribosomal protein L18 [Source:MGI Symbol;Acc:MGI:1914931] | 863,9 | 86,35 |
| ENSMUSG00000026234 | Ncl | nucleolin [Source:MGI Symbol;Acc:MGI:97286] | 863,8 | 76,48 |
| ENSMUSG00000036046 | 5031439G07Rik | RIKEN cDNA 5031439G07 gene [Source:MGI Symbol;Acc:MGI:2444899] | 863,8 | 99,08 |
| ENSMUSG00000030802 | Bckdk | branched chain ketoacid dehydrogenase kinase [Source:MGI Symbol;Acc:MGI:1276121] | 863,7 | 94,67 |
| ENSMUSG00000022536 | Glyr1 | glyoxylate reductase 1 homolog (Arabidopsis) [Source:MGI Symbol;Acc:MGI:1921272] | 863,5 | 99,17 |
| ENSMUSG00000054728 | Phactr1 | phosphatase and actin regulator 1 [Source:MGI Symbol;Acc:MGI:2659021] | 863,2 | 89,71 |
| ENSMUSG00000028062 | Lamtor2 | late endosomal/lysosomal adaptor, MAPK and MTOR activator 2 [Source:MGI Symbol;Acc:MGI:1932697] | 862,3 | 98,77 |
| ENSMUSG00000045636 | Mtus1 | mitochondrial tumor suppressor 1 [Source:MGI Symbol;Acc:MGI:2142572] | 862,3 | 80,03 |
| ENSMUSG00000042613 | Pbxip1 | pre B cell leukemia transcription factor interacting protein 1 [Source:MGI Symbol;Acc:MGI:2441670] | 862,0 | 99,89 |
| ENSMUSG00000034187 | Nsf | N-ethylmaleimide sensitive fusion protein [Source:MGI Symbol;Acc:MGI:104560] | 861,4 | 97,13 |
| ENSMUSG00000033379 | Atp6v0b | ATPase, H+ transporting, lysosomal V0 subunit B [Source:MGI Symbol;Acc:MGI:1890510] | 860,3 | 74,70 |
| ENSMUSG00000023921 | Mut | methylmalonyl-Coenzyme A mutase [Source:MGI Symbol;Acc:MGI:97239] | 860,2 | 99,39 |
| ENSMUSG00000026482 | Rgl1 | ral guanine nucleotide dissociation stimulator,-like 1 [Source:MGI Symbol;Acc:MGI:107484] | 859,9 | 98,38 |
| ENSMUSG00000078656 | Vps25 | vacuolar protein sorting 25 (yeast) [Source:MGI Symbol;Acc:MGI:106354] | 859,4 | 97,02 |
| ENSMUSG00000016344 | Ppdpf | ell differentiation and proliferation factor homolog (zebrafish)RIKEN cDNA 2700038C09 gene [Source:MGI Syr | 858,1 | 95,66 |
| ENSMUSG00000006095 | Tbcb | tubulin folding cofactor B [Source:MGI Symbol;Acc:MGI:1913661] | 857,8 | 100,00 |
| ENSMUSG00000021484 | Lman2 | lectin, mannose-binding 2 [Source:MGI Symbol;Acc:MGI:1914140] | 857,8 | 99,85 |
| ENSMUSG00000023307 | March5 | membrane-associated ring finger (C3HC4) 5 [Source:MGI Symbol;Acc:MGI:1915207] | 857,0 | 92,75 |
| ENSMUSG00000017119 | Nbr1 | neighbor of Brca1 gene 1 [Source:MGI Symbol;Acc:MGI:108498] | 854,8 | 86,13 |
| ENSMUSG00000090258 | Churc1 | churchill domain containing 1 [Source:MGI Symbol;Acc:MGI:1923684] | 854,7 | 93,00 |
| ENSMUSG00000030513 | Pcsk6 | proprotein convertase subtilisin/kexin type 6 [Source:MGI Symbol;Acc:MGI:102897] | 854,5 | 97,24 |
| ENSMUSG00000078970 | Wdr92 | WD repeat domain 92 [Source:MGI Symbol;Acc:MGI:2144224] | 853,2 | 96,12 |
| ENSMUSG00000001794 | Capns1 | calpain, small subunit 1 [Source:MGI Symbol;Acc:MGI:88266] | 852,9 | 88,57 |
| ENSMUSG00000022763 | Aifm3 | apoptosis-inducing factor, mitochondrion-associated 3 [Source:MGI Symbol;Acc:MGI:1919418] | 852,3 | 97,86 |
| ENSMUSG00000020817 | Rabep1 | rabaptin, RAB GTPase binding effector protein 1 [Source:MGI Symbol;Acc:MGI:1860236] | 850,0 | 92,28 |
| ENSMUSG00000014867 | Surf4 | surfeit gene 4 [Source:MGI Symbol;Acc:MGI:98445] | 850,0 | 87,23 |
| ENSMUSG00000025439 | Clns1a | chloride channel, nucleotide-sensitive, 1A [Source:MGI Symbol;Acc:MGI:109638] | 849,9 | 98,60 |
| ENSMUSG00000029767 | Calu | calumenin [Source:MGI Symbol;Acc:MGI:1097158] | 849,9 | 74,30 |
| ENSMUSG00000022817 | Itgb5 | integrin beta 5 [Source:MGI Symbol;Acc:MGI:96614] | 849,7 | 94,96 |
| ENSMUSG00000030770 | Parva | parvin, alpha [Source:MGI Symbol;Acc:MGI:1931144] | 848,8 | 93,83 |
| ENSMUSG00000033020 | Polr2f | polymerase (RNA) II (DNA directed) polypeptide F [Source:MGI Symbol;Acc:MGI:1349393] | 848,7 | 100,00 |
| ENSMUSG00000051177 | Plcb1 | phospholipase C, beta 1 [Source:MGI Symbol;Acc:MGI:97613] | 848,7 | 94,41 |
| ENSMUSG00000021957 | Tkt | transketolase [Source:MGI Symbol;Acc:MGI:105992] | 847,7 | 81,25 |
| ENSMUSG00000000001 | Gnai3 | guanine nucleotide binding protein (G protein), alpha inhibiting 3 [Source:MGI Symbol;Acc:MGI:95773] | 846,7 | 99,86 |
| ENSMUSG00000025579 | Gaa | glucosidase, alpha, acid [Source:MGI Symbol;Acc:MGI:95609] | 846,7 | 76,73 |
| ENSMUSG00000008429 | Herpud2 | HERPUD family member 2 [Source:MGI Symbol;Acc:MGI:1915393] | 846,3 | 100,00 |
| ENSMUSG00000040028 | Elavl1 | ELAV (embryonic lethal, abnormal vision)-like 1 (Hu antigen R) [Source:MGI Symbol;Acc:MGI:1100851] | 846,1 | 92,23 |
| ENSMUSG00000032549 | Rab6b | RAB6B, member RAS oncogene family [Source:MGI Symbol;Acc:MGI:107283] | 844,9 | 99,69 |
| ENSMUSG00000004535 | Tax1bp1 | Tax1 (human T cell leukemia virus type I) binding protein 1 [Source:MGI Symbol;Acc:MGI:1289308] | 844,9 | 92,62 |
| ENSMUSG00000021288 | Klc1 | kinesin light chain 1 [Source:MGI Symbol;Acc:MGI:107978] | 844,0 | 96,49 |
| ENSMUSG00000059970 | Hspa2 | heat shock protein 2 [Source:MGI Symbol;Acc:MGI:96243] | 843,7 | 95,61 |
| ENSMUSG00000024587 | Nars | asparaginyl-tRNA synthetase [Source:MGI Symbol;Acc:MGI:1917473] | 843,6 | 100,00 |
| ENSMUSG00000027937 | Jtb | jumping translocation breakpoint [Source:MGI Symbol;Acc:MGI:1346082] | 843,3 | 98,88 |
| ENSMUSG00000021012 | Zc3h14 | zinc finger CCCH type containing 14 [Source:MGI Symbol;Acc:MGI:1919824] | 842,7 | 84,74 |
| ENSMUSG00000057858 | Fam204a | family with sequence similarity 204, member A [Source:MGI Symbol;Acc:MGI:1289174] | 842,7 | 95,04 |
| ENSMUSG00000004849 | Ap1s1 | adaptor protein complex AP-1, sigma 1 [Source:MGI Symbol;Acc:MGI:1098244] | 842,5 | 73,74 |
| ENSMUSG00000051391 | Ywhag | nooxygenase/tryptophan 5-monooxygenase activation protein, gamma polypeptide [Source:MGI Symbol;Acc: | 842,3 | 99,88 |
| ENSMUSG00000038301 | Snx10 | sorting nexin 10 [Source:MGI Symbol;Acc:MGI:1919232] | 841,5 | 94,17 |
| ENSMUSG00000062762 | Ei24 | etoposide induced 2.4 mRNA [Source:MGI Symbol;Acc:MGI:108090] | 841,4 | 95,22 |
| ENSMUSG00000021578 | Ccdc127 | coiled-coil domain containing 127 [Source:MGI Symbol;Acc:MGI:1914683] | 841,4 | 87,20 |
| ENSMUSG00000010097 | Nxf1 | nuclear RNA export factor 1 [Source:MGI Symbol;Acc:MGI:1858330] | 841,1 | 74,59 |
| ENSMUSG00000030374 | Strn4 | striatin, calmodulin binding protein 4 [Source:MGI Symbol;Acc:MGI:2142346] | 841,1 | 83,18 |
| ENSMUSG00000025728 | Pigq | phosphatidylinositol glycan anchor biosynthesis, class Q [Source:MGI Symbol;Acc:MGI:1333114] | 840,4 | 75,80 |
| ENSMUSG00000024853 | Sf3b2 | splicing factor 3b, subunit 2 [Source:MGI Symbol;Acc:MGI:2441856] | 840,3 | 99,93 |
| ENSMUSG00000051359 | Ncald | neurocalcin delta [Source:MGI Symbol;Acc:MGI:1196326] | 839,5 | 83,62 |
| ENSMUSG00000001289 | Pfdn5 | prefoldin 5 [Source:MGI Symbol;Acc:MGI:1928753] | 839,0 | 83,55 |
| ENSMUSG00000053453 | Thoc7 | THO complex 7 homolog (Drosophila) [Source:MGI Symbol;Acc:MGI:1913481] | 838,6 | 100,00 |
| ENSMUSG00000030417 | Pdcd5 | programmed cell death 5 [Source:MGI Symbol;Acc:MGI:1913538] | 838,5 | 79,24 |
| ENSMUSG00000026466 | Tor1aip1 | torsin A interacting protein 1 [Source:MGI Symbol;Acc:MGI:3582693] | 838,4 | 82,77 |
| ENSMUSG00000057897 | Camk2b | calcium/calmodulin-dependent protein kinase II, beta [Source:MGI Symbol;Acc:MGI:88257] | 838,4 | 89,93 |
| ENSMUSG00000022257 | Laptm4b | lysosomal-associated protein transmembrane 4B [Source:MGI Symbol;Acc:MGI:1890494] | 837,2 | 96,22 |
| ENSMUSG00000020225 | Tmbim4 | transmembrane BAX inhibitor motif containing 4 [Source:MGI Symbol;Acc:MGI:1915462] | 837,1 | 82,59 |
| ENSMUSG00000021660 | Btf3 | basic transcription factor 3 [Source:MGI Symbol;Acc:MGI:1202875] | 836,8 | 70,23 |
| ENSMUSG00000024583 | Txnl1 | thioredoxin-like 1 [Source:MGI Symbol;Acc:MGI:1860078] | 836,7 | 100,00 |
| ENSMUSG00000022428 | Cby1 | chibby homolog 1 (Drosophila) [Source:MGI Symbol;Acc:MGI:1920989] | 836,1 | 100,00 |
| ENSMUSG00000090639 | Gm20425 | predicted gene 20425 [Source:MGI Symbol;Acc:MGI:5141890] | 835,9 | 98,25 |
| ENSMUSG00000050565 | Tor1aip2 | torsin A interacting protein 2 [Source:MGI Symbol;Acc:MGI:3582695] | 835,6 | 85,30 |
| ENSMUSG00000037071 | Scd1 | stearoyl-Coenzyme A desaturase 1 [Source:MGI Symbol;Acc:MGI:98239] | 834,0 | 97,66 |
| ENSMUSG00000001260 | Gabrg1 | gamma-aminobutyric acid (GABA) A receptor, subunit gamma 1 [Source:MGI Symbol;Acc:MGI:103156] | 833,1 | 87,83 |
| ENSMUSG00000026737 | Pip4k2a | phosphatidylinositol-5-phosphate 4-kinase, type II, alpha [Source:MGI Symbol;Acc:MGI:1298206] | 832,8 | 99,95 |
| ENSMUSG00000024140 | Epas1 | endothelial PAS domain protein 1 [Source:MGI Symbol;Acc:MGI:109169] | 832,8 | 99,14 |
| ENSMUSG00000063787 | Chchd1 | coiled-coil-helix-coiled-coil-helix domain containing 1 [Source:MGI Symbol;Acc:MGI:1913371] | 832,7 | 99,60 |
| ENSMUSG00000026750 | Psmb7 | proteasome (prosome, macropain) subunit, beta type 7 [Source:MGI Symbol;Acc:MGI:107637] | 832,7 | 87,53 |
| ENSMUSG00000025613 | Cct8 | chaperonin containing Tcp1, subunit 8 (theta) [Source:MGI Symbol;Acc:MGI:107183] | 832,7 | 95,62 |
| ENSMUSG00000024423 | Impact | impact, RWD domain protein [Source:MGI Symbol;Acc:MGI:1098233] | 831,8 | 99,98 |
| ENSMUSG00000051285 | Pcmtd1 | ein-L-isoaspartate (D-aspartate) O-methyltransferase domain containing 1 [Source:MGI Symbol;Acc:MGI:2441 | 831,2 | 83,79 |
| ENSMUSG00000050721 | Plekho2 | pleckstrin homology domain containing, family O member 2 [Source:MGI Symbol;Acc:MGI:2143132] | 830,7 | 100,00 |
| ENSMUSG00000038459 | Abhd17c | abhydrolase domain containing 17C [Source:MGI Symbol;Acc:MGI:1917428] | 829,8 | 100,00 |
| ENSMUSG00000022272 | Myo10 | myosin X [Source:MGI Symbol;Acc:MGI:107716] | 829,7 | 71,29 |
| ENSMUSG00000021065 | Fut8 | fucosyltransferase 8 [Source:MGI Symbol;Acc:MGI:1858901] | 829,5 | 94,83 |
| ENSMUSG00000029636 | Wasf3 | WAS protein family, member 3 [Source:MGI Symbol;Acc:MGI:2658986] | 829,1 | 94,85 |
| ENSMUSG00000025545 | Clybl | citrate lyase beta like [Source:MGI Symbol;Acc:MGI:1916884] | 828,9 | 99,89 |
| ENSMUSG00000060548 | Tnfrsf19 | tumor necrosis factor receptor superfamily, member 19 [Source:MGI Symbol;Acc:MGI:1352474] | 827,3 | 93,82 |
| ENSMUSG00000047126 | Cltc | clathrin, heavy polypeptide (Hc) [Source:MGI Symbol;Acc:MGI:2388633] | 826,6 | 98,73 |
| ENSMUSG00000026470 | Stx6 | syntaxin 6 [Source:MGI Symbol;Acc:MGI:1926235] | 826,4 | 76,59 |
| ENSMUSG00000034235 | Usp54 | ubiquitin specific peptidase 54 [Source:MGI Symbol;Acc:MGI:1926037] | 826,0 | 99,95 |
| ENSMUSG00000038324 | Trpc4ap | receptor potential cation channel, subfamily C, member 4 associated protein [Source:MGI Symbol;Acc:MGI:1 | 825,2 | 98,92 |
| ENSMUSG00000006575 | Rundc3a | RUN domain containing 3A [Source:MGI Symbol;Acc:MGI:1858752] | 824,6 | 98,59 |
| ENSMUSG00000039217 | Il18 | interleukin 18 [Source:MGI Symbol;Acc:MGI:107936] | 823,3 | 98,88 |
| ENSMUSG00000028847 | Trappc3 | trafficking protein particle complex 3 [Source:MGI Symbol;Acc:MGI:1351486] | 823,3 | 98,48 |
| ENSMUSG00000038374 | Rbm8a | RNA binding motif protein 8a [Source:MGI Symbol;Acc:MGI:1913129] | 822,4 | 94,56 |
| ENSMUSG00000063200 | Nol7 | nucleolar protein 7 [Source:MGI Symbol;Acc:MGI:1917328] | 821,9 | 100,00 |
| ENSMUSG00000018565 | Elp5 | elongator acetyltransferase complex subunit 5 [Source:MGI Symbol;Acc:MGI:1859017] | 821,7 | 94,10 |
| ENSMUSG00000051319 | 1500011K16Rik | RIKEN cDNA 1500011K16 gene [Source:MGI Symbol;Acc:MGI:1915135] | 821,7 | 100,00 |
| ENSMUSG00000029781 | Fkbp9 | FK506 binding protein 9 [Source:MGI Symbol;Acc:MGI:1350921] | 821,3 | 98,59 |
| ENSMUSG00000027519 | Rab22a | RAB22A, member RAS oncogene family [Source:MGI Symbol;Acc:MGI:105072] | 818,8 | 96,91 |
| ENSMUSG00000032301 | Psma4 | proteasome (prosome, macropain) subunit, alpha type 4 [Source:MGI Symbol;Acc:MGI:1347060] | 818,4 | 88,97 |
| ENSMUSG00000003235 | Eif2b5 | eukaryotic translation initiation factor 2B, subunit 5 epsilon [Source:MGI Symbol;Acc:MGI:2446176] | 817,7 | 83,90 |
| ENSMUSG00000060923 | Acyp2 | acylphosphatase 2, muscle type [Source:MGI Symbol;Acc:MGI:1922822] | 817,7 | 99,96 |
| ENSMUSG00000024993 | Fam45a | family with sequence similarity 45, member A [Source:MGI Symbol;Acc:MGI:1915144] | 817,1 | 99,92 |
| ENSMUSG00000018379 | Srsf1 | serine/arginine-rich splicing factor 1 [Source:MGI Symbol;Acc:MGI:98283] | 816,9 | 96,01 |
| ENSMUSG00000101678 | Gm29609 | predicted gene 29609 [Source:MGI Symbol;Acc:MGI:5580315] | 816,2 | 84,30 |
| ENSMUSG00000027404 | Snrpb | small nuclear ribonucleoprotein B [Source:MGI Symbol;Acc:MGI:98342] | 815,9 | 99,02 |
| ENSMUSG00000040591 | 1110051M20Rik | RIKEN cDNA 1110051M20 gene [Source:MGI Symbol;Acc:MGI:1915079] | 815,3 | 76,63 |
| ENSMUSG00000032238 | Rora | RAR-related orphan receptor alpha [Source:MGI Symbol;Acc:MGI:104661] | 815,2 | 86,52 |
| ENSMUSG00000025034 | Trim8 | tripartite motif-containing 8 [Source:MGI Symbol;Acc:MGI:1933302] | 814,9 | 99,75 |
| ENSMUSG00000024097 | Srsf7 | serine/arginine-rich splicing factor 7 [Source:MGI Symbol;Acc:MGI:1926232] | 814,9 | 100,00 |
| ENSMUSG00000029499 | Pxmp2 | peroxisomal membrane protein 2 [Source:MGI Symbol;Acc:MGI:107487] | 814,3 | 92,51 |
| ENSMUSG00000009927 | Rps25 | ribosomal protein S25 [Source:MGI Symbol;Acc:MGI:1922867] | 814,2 | 89,07 |
| ENSMUSG00000035642 | Aamdc | adipogenesis associated Mth938 domain containing [Source:MGI Symbol;Acc:MGI:1913523] | 812,9 | 78,89 |
| ENSMUSG00000060477 | Irak2 | interleukin-1 receptor-associated kinase 2 [Source:MGI Symbol;Acc:MGI:2429603] | 812,7 | 99,63 |

| | | | | |
|---|---|---|---|---|
| ENSMUSG00000038335 | Tsr1 | TSR1 20S rRNA accumulation [Source:MGI Symbol;Acc:MGI:2144566] | 812,1 | 98,27 |
| ENSMUSG00000005417 | Mprip | myosin phosphatase Rho interacting protein [Source:MGI Symbol;Acc:MGI:1349438] | 811,9 | 83,92 |
| ENSMUSG00000102349 | Gm37376 | predicted gene, 37376 [Source:MGI Symbol;Acc:MGI:5610604] | 811,0 | 100,00 |
| ENSMUSG00000032388 | Spg21 | spastic paraplegia 21 homolog (human) [Source:MGI Symbol;Acc:MGI:106403] | 809,8 | 87,04 |
| ENSMUSG00000021518 | Ptdss1 | phosphatidylserine synthase 1 [Source:MGI Symbol;Acc:MGI:1276575] | 809,5 | 91,42 |
| ENSMUSG00000002733 | Plekha3 | mology domain-containing, family A (phosphoinositide binding specific) member 3 [Source:MGI Symbol;Acc:M | 809,3 | 98,93 |
| ENSMUSG00000035093 | Secisbp2l | SECIS binding protein 2-like [Source:MGI Symbol;Acc:MGI:1917604] | 809,0 | 89,08 |
| ENSMUSG00000021273 | Fdft1 | farnesyl diphosphate farnesyl transferase 1 [Source:MGI Symbol;Acc:MGI:102706] | 808,5 | 81,81 |
| ENSMUSG00000020993 | Trappc6b | trafficking protein particle complex 6B [Source:MGI Symbol;Acc:MGI:1925482] | 808,3 | 95,53 |
| ENSMUSG00000039001 | Rps21 | ribosomal protein S21 [Source:MGI Symbol;Acc:MGI:1913731] | 808,1 | 99,75 |
| ENSMUSG00000022808 | Snx4 | sorting nexin 4 [Source:MGI Symbol;Acc:MGI:1916400] | 807,1 | 100,00 |
| ENSMUSG00000013236 | Ptprs | protein tyrosine phosphatase, receptor type, S [Source:MGI Symbol;Acc:MGI:97815] | 806,9 | 98,13 |
| ENSMUSG00000032905 | Atg12 | autophagy related 12 [Source:MGI Symbol;Acc:MGI:1914776] | 806,8 | 99,76 |
| ENSMUSG00000026203 | Dnajb2 | DnaJ heat shock protein family (Hsp40) member B2 [Source:MGI Symbol;Acc:MGI:1928739] | 806,2 | 98,95 |
| ENSMUSG00000021486 | Prelid1 | PRELI domain containing 1 [Source:MGI Symbol;Acc:MGI:1913744] | 806,0 | 92,49 |
| ENSMUSG00000003039 | Fam32a | family with sequence similarity 32, member A [Source:MGI Symbol;Acc:MGI:1915172] | 805,9 | 100,00 |
| ENSMUSG00000039159 | Ube2h | ubiquitin-conjugating enzyme E2H [Source:MGI Symbol;Acc:MGI:104632] | 805,4 | 79,41 |
| ENSMUSG00000028861 | Mrps15 | mitochondrial ribosomal protein S15 [Source:MGI Symbol;Acc:MGI:1913657] | 804,6 | 100,00 |
| ENSMUSG00000007777 | 0610009B22Rik | RIKEN cDNA 0610009B22 gene [Source:MGI Symbol;Acc:MGI:1913300] | 803,2 | 88,51 |
| ENSMUSG00000009073 | Nf2 | neurofibromatosis 2 [Source:MGI Symbol;Acc:MGI:97307] | 803,1 | 88,05 |
| ENSMUSG00000026179 | Pnkd | paroxysmal nonkinesiogenic dyskinesia [Source:MGI Symbol;Acc:MGI:1930773] | 802,9 | 93,89 |
| ENSMUSG00000046688 | Tifa | TRAF-interacting protein with forkhead-associated domain [Source:MGI Symbol;Acc:MGI:2182965] | 802,4 | 96,98 |
| ENSMUSG00000024921 | Smarca2 | ed, matrix associated, actin dependent regulator of chromatin, subfamily a, member 2 [Source:MGI Symbol;Ac | 802,1 | 78,83 |
| ENSMUSG00000032002 | Dcun1d5 | N1, defective in cullin neddylation 1, domain containing 5 (S. cerevisiae) [Source:MGI Symbol;Acc:MGI:19241 | 801,9 | 100,00 |
| ENSMUSG00000032096 | Arcn1 | archain 1 [Source:MGI Symbol;Acc:MGI:2387591] | 801,8 | 97,44 |
| ENSMUSG00000027007 | Ssfa2 | sperm specific antigen 2 [Source:MGI Symbol;Acc:MGI:1917849] | 801,8 | 98,59 |
| ENSMUSG00000064138 | Fam172a | family with sequence similarity 172, member A [Source:MGI Symbol;Acc:MGI:1915925] | 801,1 | 98,61 |
| ENSMUSG00000027288 | Zfp106 | zinc finger protein 106 [Source:MGI Symbol;Acc:MGI:1270153] | 798,6 | 95,08 |
| ENSMUSG00000020950 | Foxg1 | forkhead box G1 [Source:MGI Symbol;Acc:MGI:1347464] | 797,8 | 85,14 |
| ENSMUSG00000031556 | Tm2d2 | TM2 domain containing 2 [Source:MGI Symbol;Acc:MGI:1916992] | 797,2 | 99,48 |
| ENSMUSG00000030960 | Mettl10 | methyltransferase like 10 [Source:MGI Symbol;Acc:MGI:1919346] | 797,2 | 99,58 |
| ENSMUSG00000028252 | Ccnc | cyclin C [Source:MGI Symbol;Acc:MGI:1858199] | 796,4 | 97,05 |
| ENSMUSG00000055022 | Cntn1 | contactin 1 [Source:MGI Symbol;Acc:MGI:105980] | 795,9 | 99,01 |
| ENSMUSG00000030291 | Med21 | mediator complex subunit 21 [Source:MGI Symbol;Acc:MGI:1347064] | 795,8 | 72,01 |
| ENSMUSG00000032382 | Snx1 | sorting nexin 1 [Source:MGI Symbol;Acc:MGI:1928395] | 794,6 | 98,60 |
| ENSMUSG00000025260 | Hsd17b10 | hydroxysteroid (17-beta) dehydrogenase 10 [Source:MGI Symbol;Acc:MGI:1333871] | 794,5 | 95,77 |
| ENSMUSG00000037999 | Arap2 | ArfGAP with RhoGAP domain, ankyrin repeat and PH domain 2 [Source:MGI Symbol;Acc:MGI:2684416] | 794,5 | 88,88 |
| ENSMUSG00000039195 | 1110008P14Rik | RIKEN cDNA 1110008P14 gene [Source:MGI Symbol;Acc:MGI:1920987] | 793,5 | 98,52 |
| ENSMUSG00000032816 | Igdcc4 | immunoglobulin superfamily, DCC subclass, member 4 [Source:MGI Symbol;Acc:MGI:1858497] | 793,4 | 98,45 |
| ENSMUSG00000071253 | Slc25a16 | rrier family 25 (mitochondrial carrier, Graves disease autoantigen), member 16 [Source:MGI Symbol;Acc:MGI:1 | 793,1 | 99,24 |
| ENSMUSG00000027239 | Mdk | midkine [Source:MGI Symbol;Acc:MGI:96949] | 793,0 | 84,94 |
| ENSMUSG00000021737 | Psmd6 | proteasome (prosome, macropain) 26S subunit, non-ATPase, 6 [Source:MGI Symbol;Acc:MGI:1913663] | 792,8 | 98,67 |
| ENSMUSG00000039183 | Nubp2 | nucleotide binding protein 2 [Source:MGI Symbol;Acc:MGI:1347072] | 792,2 | 99,77 |
| ENSMUSG00000031536 | Polb | polymerase (DNA directed), beta [Source:MGI Symbol;Acc:MGI:97740] | 792,2 | 99,76 |
| ENSMUSG00000027679 | Dnajc19 | DnaJ heat shock protein family (Hsp40) member C19 [Source:MGI Symbol;Acc:MGI:1914963] | 791,6 | 71,53 |
| ENSMUSG00000068748 | Ptprz1 | protein tyrosine phosphatase, receptor type Z, polypeptide 1 [Source:MGI Symbol;Acc:MGI:97816] | 790,5 | 98,38 |
| ENSMUSG00000019124 | Scrn1 | secernin 1 [Source:MGI Symbol;Acc:MGI:1917188] | 789,7 | 94,00 |
| ENSMUSG00000045503 | Sys1 | YS1 Golgi-localized integral membrane protein homolog (S. cerevisiae) [Source:MGI Symbol;Acc:MGI:191371 | 789,5 | 95,10 |
| ENSMUSG00000024283 | Wac | WW domain containing adaptor with coiled-coil [Source:MGI Symbol;Acc:MGI:2387357] | 789,1 | 88,66 |
| ENSMUSG00000097919 | Gm27021 | predicted gene, 27021 [Source:MGI Symbol;Acc:MGI:5504136] | 788,9 | 100,00 |
| ENSMUSG00000090266 | Mettl23 | methyltransferase like 23 [Source:MGI Symbol;Acc:MGI:1921569] | 788,4 | 97,16 |
| ENSMUSG00000032536 | Trak1 | trafficking protein, kinesin binding 1 [Source:MGI Symbol;Acc:MGI:1914345] | 787,9 | 98,02 |
| ENSMUSG00000054863 | Fam19a5 | family with sequence similarity 19, member A5 [Source:MGI Symbol;Acc:MGI:2146182] | 787,3 | 100,00 |
| ENSMUSG00000020589 | Fam49a | family with sequence similarity 49, member A [Source:MGI Symbol;Acc:MGI:1261783] | 787,3 | 99,83 |
| ENSMUSG00000028932 | Psmc2 | proteasome (prosome, macropain) 26S subunit, ATPase 2 [Source:MGI Symbol;Acc:MGI:109555] | 787,3 | 85,53 |
| ENSMUSG00000020277 | Pfkl | phosphofructokinase, liver, B-type [Source:MGI Symbol;Acc:MGI:97547] | 786,9 | 95,27 |
| ENSMUSG00000032740 | Ccdc88a | coiled coil domain containing 88A [Source:MGI Symbol;Acc:MGI:1925177] | 786,7 | 89,82 |
| ENSMUSG00000028136 | Snx27 | sorting nexin family member 27 [Source:MGI Symbol;Acc:MGI:1923992] | 786,5 | 97,11 |
| ENSMUSG00000094483 | Purb | purine rich element binding protein B [Source:MGI Symbol;Acc:MGI:1338779] | 786,3 | 99,11 |
| ENSMUSG00000022663 | Atg3 | autophagy related 3 [Source:MGI Symbol;Acc:MGI:1915091] | 786,1 | 95,19 |
| ENSMUSG00000013822 | Elof1 | ELF1 homolog, elongation factor 1 [Source:MGI Symbol;Acc:MGI:1913376] | 784,6 | 79,37 |
| ENSMUSG00000024944 | Arl2 | ADP-ribosylation factor-like 2 [Source:MGI Symbol;Acc:MGI:1928393] | 784,5 | 91,00 |
| ENSMUSG00000025555 | Farp1 | I, RhoGEF (Arhgef) and pleckstrin domain protein 1 (chondrocyte-derived) [Source:MGI Symbol;Acc:MGI:244 | 784,2 | 96,21 |
| ENSMUSG00000074649 | BC029722 | cDNA sequence BC029722 [Source:MGI Symbol;Acc:MGI:3584273] | 784,2 | 98,89 |
| ENSMUSG00000028278 | Rragd | Ras-related GTP binding D [Source:MGI Symbol;Acc:MGI:1098604] | 783,8 | 93,98 |
| ENSMUSG00000005054 | Cstb | cystatin B [Source:MGI Symbol;Acc:MGI:109514] | 783,6 | 100,00 |
| ENSMUSG00000020734 | Grin2c | glutamate receptor, ionotropic, NMDA2C (epsilon 3) [Source:MGI Symbol;Acc:MGI:95822] | 783,3 | 95,16 |
| ENSMUSG00000026341 | Actr3 | ARP3 actin-related protein 3 [Source:MGI Symbol;Acc:MGI:1921367] | 781,5 | 88,57 |
| ENSMUSG00000002205 | Vrk3 | vaccinia related kinase 3 [Source:MGI Symbol;Acc:MGI:2182465] | 781,4 | 77,32 |
| ENSMUSG00000024695 | Zfp91 | zinc finger protein 91 [Source:MGI Symbol;Acc:MGI:104854] | 780,3 | 86,83 |
| ENSMUSG00000015094 | Npdc1 | neural proliferation, differentiation and control 1 [Source:MGI Symbol;Acc:MGI:1099802] | 779,6 | 97,37 |
| ENSMUSG00000074247 | Dda1 | DET1 and DDB1 associated 1 [Source:MGI Symbol;Acc:MGI:1913748] | 779,5 | 99,93 |
| ENSMUSG00000030235 | Slco1c1 | solute carrier organic anion transporter family, member 1c1 [Source:MGI Symbol;Acc:MGI:1889679] | 779,4 | 94,38 |
| ENSMUSG00000025153 | Fasn | fatty acid synthase [Source:MGI Symbol;Acc:MGI:95485] | 778,9 | 84,19 |
| ENSMUSG00000057113 | Npm1 | nucleophosmin 1 [Source:MGI Symbol;Acc:MGI:106184] | 778,3 | 92,12 |
| ENSMUSG00000024259 | Slc25a46 | solute carrier family 25, member 46 [Source:MGI Symbol;Acc:MGI:1914703] | 777,6 | 85,66 |
| ENSMUSG00000003269 | Cyth2 | cytohesin 2 [Source:MGI Symbol;Acc:MGI:1334255] | 776,6 | 78,92 |
| ENSMUSG00000015839 | Nfe2l2 | nuclear factor, erythroid derived 2, like 2 [Source:MGI Symbol;Acc:MGI:108420] | 776,2 | 98,59 |
| ENSMUSG00000062691 | Cebpzos | CCAAT/enhancer binding protein (C/EBP), zeta, opposite strand [Source:MGI Symbol;Acc:MGI:1915804] | 775,5 | 99,20 |
| ENSMUSG00000032010 | Usp2 | ubiquitin specific peptidase 2 [Source:MGI Symbol;Acc:MGI:1858178] | 775,0 | 95,43 |
| ENSMUSG00000057522 | Spop | speckle-type POZ protein [Source:MGI Symbol;Acc:MGI:1343085] | 774,9 | 98,17 |
| ENSMUSG00000050043 | Tmx2 | thioredoxin-related transmembrane protein 2 [Source:MGI Symbol;Acc:MGI:1914208] | 774,4 | 70,80 |
| ENSMUSG00000020250 | Txnrd1 | thioredoxin reductase 1 [Source:MGI Symbol;Acc:MGI:1354175] | 774,0 | 99,76 |
| ENSMUSG00000032965 | Ift57 | intraflagellar transport 57 [Source:MGI Symbol;Acc:MGI:1921166] | 773,5 | 76,97 |
| ENSMUSG00000061244 | Exoc5 | exocyst complex component 5 [Source:MGI Symbol;Acc:MGI:2145645] | 773,5 | 98,72 |
| ENSMUSG00000046447 | Camk2n1 | calcium/calmodulin-dependent protein kinase II inhibitor 1 [Source:MGI Symbol;Acc:MGI:1913509] | 773,5 | 90,20 |
| ENSMUSG00000033540 | Idua | iduronidase, alpha-L- [Source:MGI Symbol;Acc:MGI:96418] | 773,2 | 80,78 |
| ENSMUSG00000052337 | Immt | inner membrane protein, mitochondrial [Source:MGI Symbol;Acc:MGI:1923864] | 772,7 | 96,75 |
| ENSMUSG00000015290 | Ubl4a | ubiquitin-like 4A [Source:MGI Symbol;Acc:MGI:95049] | 771,4 | 93,15 |
| ENSMUSG00000024560 | Cxxc1 | CXXC finger 1 (PHD domain) [Source:MGI Symbol;Acc:MGI:1921572] | 771,0 | 99,47 |
| ENSMUSG00000022338 | Eny2 | enhancer of yellow 2 homolog (Drosophila) [Source:MGI Symbol;Acc:MGI:1919286] | 770,7 | 100,00 |
| ENSMUSG00000037242 | Clic4 | chloride intracellular channel 4 (mitochondrial) [Source:MGI Symbol;Acc:MGI:1352754] | 769,8 | 99,00 |
| ENSMUSG00000031068 | Glrx3 | glutaredoxin 3 [Source:MGI Symbol;Acc:MGI:1353653] | 769,1 | 83,10 |
| ENSMUSG00000031198 | Fundc2 | FUN14 domain containing 2 [Source:MGI Symbol;Acc:MGI:1914641] | 768,7 | 99,43 |
| ENSMUSG00000029571 | Tmem106b | transmembrane protein 106B [Source:MGI Symbol;Acc:MGI:1919150] | 767,4 | 97,90 |
| ENSMUSG00000037669 | Ldah | lipid droplet associated hydrolase [Source:MGI Symbol;Acc:MGI:1916082] | 767,1 | 99,53 |
| ENSMUSG00000017639 | Rab11fip4 | RAB11 family interacting protein 4 (class II) [Source:MGI Symbol;Acc:MGI:2442920] | 766,2 | 97,99 |
| ENSMUSG00000024346 | Pfdn1 | prefoldin 1 [Source:MGI Symbol;Acc:MGI:1914449] | 765,2 | 94,68 |
| ENSMUSG00000024639 | Gnaq | guanine nucleotide binding protein, alpha q polypeptide [Source:MGI Symbol;Acc:MGI:95776] | 764,5 | 98,68 |
| ENSMUSG00000018287 | Spag7 | sperm associated antigen 7 [Source:MGI Symbol;Acc:MGI:107380] | 764,1 | 98,62 |
| ENSMUSG00000032594 | Ip6k1 | inositol hexaphosphate kinase 1 [Source:MGI Symbol;Acc:MGI:1351633] | 763,4 | 73,63 |
| ENSMUSG00000032413 | Rasa2 | RAS p21 protein activator 2 [Source:MGI Symbol;Acc:MGI:2149960] | 763,4 | 99,32 |
| ENSMUSG00000068921 | Dap3 | death associated protein 3 [Source:MGI Symbol;Acc:MGI:1929538] | 762,2 | 76,48 |
| ENSMUSG00000060032 | H2afj | H2A histone family, member J [Source:MGI Symbol;Acc:MGI:3606192] | 762,2 | 99,54 |
| ENSMUSG00000045302 | Preb | prolactin regulatory element binding [Source:MGI Symbol;Acc:MGI:1355326] | 762,1 | 80,69 |
| ENSMUSG00000021476 | Habp4 | hyaluronic acid binding protein 4 [Source:MGI Symbol;Acc:MGI:1891713] | 762,0 | 100,00 |
| ENSMUSG00000038023 | Atp6v0a2 | ATPase, H+ transporting, lysosomal V0 subunit A2 [Source:MGI Symbol;Acc:MGI:104855] | 761,1 | 99,62 |
| ENSMUSG00000025537 | Phkg1 | phosphorylase kinase gamma 1 [Source:MGI Symbol;Acc:MGI:97579] | 760,6 | 87,53 |
| ENSMUSG00000032554 | Trf | transferrin [Source:MGI Symbol;Acc:MGI:98821] | 760,6 | 87,62 |
| ENSMUSG00000008690 | Ncaph2 | non-SMC condensin II complex, subunit H2 [Source:MGI Symbol;Acc:MGI:1289164] | 760,2 | 98,56 |
| ENSMUSG00000026667 | Uhmk1 | U2AF homology motif (UHM) kinase 1 [Source:MGI Symbol;Acc:MGI:1341908] | 760,0 | 96,84 |
| ENSMUSG00000026201 | Stk16 | serine/threonine kinase 16 [Source:MGI Symbol;Acc:MGI:1313271] | 759,7 | 96,45 |
| ENSMUSG00000027809 | Etfdh | electron transferring flavoprotein, dehydrogenase [Source:MGI Symbol;Acc:MGI:106100] | 759,5 | 95,50 |
| ENSMUSG00000021996 | Esd | esterase D/formylglutathione hydrolase [Source:MGI Symbol;Acc:MGI:95421] | 758,9 | 91,24 |
| ENSMUSG00000046330 | Rpl37a | ribosomal protein L37a [Source:MGI Symbol;Acc:MGI:98068] | 757,0 | 100,00 |
| ENSMUSG00000020078 | Vps26a | vacuolar protein sorting 26 homolog A (yeast) [Source:MGI Symbol;Acc:MGI:1353654] | 756,9 | 80,48 |
| ENSMUSG00000034190 | Chmp7 | charged multivesicular body protein 7 [Source:MGI Symbol;Acc:MGI:1913922] | 756,2 | 99,15 |

| | | | | |
|---|---|---|---|---|
| ENSMUSG00000061981 | Flot2 | flotillin 2 [Source:MGI Symbol;Acc:MGI:103309] | 756,0 | 99,10 |
| ENSMUSG00000021711 | Trappc13 | trafficking protein particle complex 13 [Source:MGI Symbol;Acc:MGI:1914225] | 755,9 | 99,42 |
| ENSMUSG00000090733 | Rps27 | ribosomal protein S27 [Source:MGI Symbol;Acc:MGI:1888676] | 755,8 | 91,16 |
| ENSMUSG00000018446 | C1qbp | complement component 1, q subcomponent binding protein [Source:MGI Symbol;Acc:MGI:1194505] | 755,6 | 100,00 |
| ENSMUSG00000024571 | Gm16286 | predicted gene 16286 [Source:MGI Symbol;Acc:MGI:3833940] | 754,9 | 96,94 |
| ENSMUSG00000023075 | Akirin1 | akirin 1 [Source:MGI Symbol;Acc:MGI:1915300] | 754,5 | 85,26 |
| ENSMUSG00000022378 | Fam49b | family with sequence similarity 49, member B [Source:MGI Symbol;Acc:MGI:1923520] | 754,3 | 99,78 |
| ENSMUSG00000022949 | Clic6 | chloride intracellular channel 6 [Source:MGI Symbol;Acc:MGI:2146607] | 753,3 | 95,85 |
| ENSMUSG00000036275 | 9530068E07Rik | RIKEN cDNA 9530068E07 gene [Source:MGI Symbol;Acc:MGI:2654705] | 753,2 | 100,00 |
| ENSMUSG00000036854 | Hspb6 | heat shock protein, alpha-crystallin-related, B6 [Source:MGI Symbol;Acc:MGI:2685325] | 752,7 | 99,90 |
| ENSMUSG00000078931 | Pdf | peptide deformylase (mitochondrial) [Source:MGI Symbol;Acc:MGI:1915273] | 752,5 | 100,00 |
| ENSMUSG00000002768 | Mea1 | male enhanced antigen 1 [Source:MGI Symbol;Acc:MGI:96957] | 752,4 | 100,00 |
| ENSMUSG00000074657 | Kif5a | kinesin family member 5A [Source:MGI Symbol;Acc:MGI:109564] | 752,0 | 96,88 |
| ENSMUSG00000025917 | Cops5 | COP9 signalosome subunit 5 [Source:MGI Symbol;Acc:MGI:1349415] | 751,9 | 98,33 |
| ENSMUSG00000053253 | Ndfip2 | Nedd4 family interacting protein 2 [Source:MGI Symbol;Acc:MGI:1923523] | 751,7 | 97,01 |
| ENSMUSG00000026080 | Chst10 | carbohydrate sulfotransferase 10 [Source:MGI Symbol;Acc:MGI:2138283] | 751,2 | 70,30 |
| ENSMUSG00000009575 | Cbx5 | chromobox 5 [Source:MGI Symbol;Acc:MGI:109372] | 750,8 | 98,91 |
| ENSMUSG00000021302 | Ggps1 | geranylgeranyl diphosphate synthase 1 [Source:MGI Symbol;Acc:MGI:1341724] | 750,7 | 100,00 |
| ENSMUSG00000025134 | Alyref | Aly/REF export factor [Source:MGI Symbol;Acc:MGI:1341044] | 750,1 | 96,07 |
| ENSMUSG00000030697 | Ppp4c | protein phosphatase 4, catalytic subunit [Source:MGI Symbol;Acc:MGI:1891763] | 749,1 | 89,05 |
| ENSMUSG00000027206 | Cops2 | COP9 signalosome subunit 2 [Source:MGI Symbol;Acc:MGI:1330276] | 748,0 | 97,00 |
| ENSMUSG00000029536 | Gatc | glutamyl-tRNA(Gln) amidotransferase, subunit C [Source:MGI Symbol;Acc:MGI:1923776] | 747,9 | 100,00 |
| ENSMUSG00000040242 | Fgfr1op2 | FGFR1 oncogene partner 2 [Source:MGI Symbol;Acc:MGI:1914779] | 747,3 | 96,39 |
| ENSMUSG00000032802 | Srxn1 | sulfiredoxin 1 homolog (S. cerevisiae) [Source:MGI Symbol;Acc:MGI:104971] | 746,7 | 97,62 |
| ENSMUSG00000053581 | Zfand2a | zinc finger, AN1-type domain 2A [Source:MGI Symbol;Acc:MGI:2140729] | 746,2 | 98,89 |
| ENSMUSG00000037119 | D15Ertd621e | DNA segment, Chr 15, ERATO Doi 621, expressed [Source:MGI Symbol;Acc:MGI:1277178] | 745,6 | 99,48 |
| ENSMUSG00000031985 | Gnpat | glyceronephosphate O-acyltransferase [Source:MGI Symbol;Acc:MGI:1343460] | 745,2 | 99,32 |
| ENSMUSG00000064339 | mt-Rnr2 | mitochondrially encoded 16S rRNA [Source:MGI Symbol;Acc:MGI:102492] | 745,0 | 99,77 |
| ENSMUSG00000026434 | Nucks1 | nuclear casein kinase and cyclin-dependent kinase substrate 1 [Source:MGI Symbol;Acc:MGI:1934811] | 744,6 | 99,45 |
| ENSMUSG00000078861 | Zfp931 | zinc finger protein 931 [Source:MGI Symbol;Acc:MGI:2441662] | 744,5 | 93,84 |
| ENSMUSG00000063065 | Mapk3 | mitogen-activated protein kinase 3 [Source:MGI Symbol;Acc:MGI:1346859] | 743,7 | 85,28 |
| ENSMUSG00000061046 | Haghl | hydroxyacylglutathione hydrolase-like [Source:MGI Symbol;Acc:MGI:1919877] | 743,5 | 91,85 |
| ENSMUSG00000039530 | Tusc3 | tumor suppressor candidate 3 [Source:MGI Symbol;Acc:MGI:1933134] | 742,6 | 100,00 |
| ENSMUSG00000027952 | Pmvk | phosphomevalonate kinase [Source:MGI Symbol;Acc:MGI:1915853] | 742,6 | 73,85 |
| ENSMUSG00000028020 | Glrb | glycine receptor, beta subunit [Source:MGI Symbol;Acc:MGI:95751] | 742,1 | 96,12 |
| ENSMUSG00000021589 | Rhobtb3 | Rho-related BTB domain containing 3 [Source:MGI Symbol;Acc:MGI:1920546] | 740,9 | 98,35 |
| ENSMUSG00000028277 | Ube2j1 | ubiquitin-conjugating enzyme E2J 1 [Source:MGI Symbol;Acc:MGI:1926245] | 740,7 | 99,61 |
| ENSMUSG00000027133 | Nop10 | NOP10 ribonucleoprotein [Source:MGI Symbol;Acc:MGI:1913431] | 738,9 | 97,42 |
| ENSMUSG00000026927 | Sdccag3 | serologically defined colon cancer antigen 3 [Source:MGI Symbol;Acc:MGI:1915362] | 738,7 | 99,09 |
| ENSMUSG00000000439 | Mkrn2 | makorin, ring finger protein, 2 [Source:MGI Symbol;Acc:MGI:1914277] | 738,1 | 91,08 |
| ENSMUSG00000025151 | Maged1 | melanoma antigen, family D, 1 [Source:MGI Symbol;Acc:MGI:1930187] | 737,8 | 97,20 |
| ENSMUSG00000061517 | Sox21 | SRY (sex determining region Y)-box 21 [Source:MGI Symbol;Acc:MGI:2654070] | 737,7 | 100,00 |
| ENSMUSG00000055782 | Abcd2 | ATP-binding cassette, sub-family D (ALD), member 2 [Source:MGI Symbol;Acc:MGI:1349467] | 737,6 | 99,26 |
| ENSMUSG00000037815 | Ctnna1 | catenin (cadherin associated protein), alpha 1 [Source:MGI Symbol;Acc:MGI:88274] | 737,3 | 99,91 |
| ENSMUSG00000061436 | Hipk2 | homeodomain interacting protein kinase 2 [Source:MGI Symbol;Acc:MGI:1314872] | 736,8 | 89,68 |
| ENSMUSG00000012848 | Rps5 | ribosomal protein S5 [Source:MGI Symbol;Acc:MGI:1097682] | 736,4 | 88,60 |
| ENSMUSG00000029864 | Gstk1 | glutathione S-transferase kappa 1 [Source:MGI Symbol;Acc:MGI:1923513] | 735,8 | 100,00 |
| ENSMUSG00000024966 | Stip1 | stress-induced phosphoprotein 1 [Source:MGI Symbol;Acc:MGI:109130] | 735,8 | 99,06 |
| ENSMUSG00000028779 | Pef1 | penta-EF hand domain containing 1 [Source:MGI Symbol;Acc:MGI:1915148] | 735,5 | 83,03 |
| ENSMUSG00000046312 | AI464131 | expressed sequence AI464131 [Source:MGI Symbol;Acc:MGI:2140300] | 735,0 | 93,83 |
| ENSMUSG00000018909 | Arrb1 | arrestin, beta 1 [Source:MGI Symbol;Acc:MGI:99473] | 734,7 | 95,96 |
| ENSMUSG00000020621 | Rdh14 | retinol dehydrogenase 14 (all-trans and 9-cis) [Source:MGI Symbol;Acc:MGI:1920402] | 734,6 | 100,00 |
| ENSMUSG00000024811 | Tnks2 | ankyrase, TRF1-interacting ankyrin-related ADP-ribose polymerase 2 [Source:MGI Symbol;Acc:MGI:1921743 | 734,5 | 96,91 |
| ENSMUSG00000028843 | Sh3bgrl3 | SH3 domain binding glutamic acid-rich protein-like 3 [Source:MGI Symbol;Acc:MGI:1920973] | 734,5 | 99,60 |
| ENSMUSG00000022136 | Dnajc3 | DnaJ heat shock protein family (Hsp40) member C3 [Source:MGI Symbol;Acc:MGI:107373] | 733,7 | 98,40 |
| ENSMUSG00000037287 | Tbcel | tubulin folding cofactor E-like [Source:MGI Symbol;Acc:MGI:1925543] | 733,7 | 92,76 |
| ENSMUSG00000028412 | Slc44a1 | solute carrier family 44, member 1 [Source:MGI Symbol;Acc:MGI:2140592] | 733,4 | 81,36 |
| ENSMUSG00000028673 | Fuca1 | fucosidase, alpha-L- 1, tissue [Source:MGI Symbol;Acc:MGI:95593] | 733,4 | 86,39 |
| ENSMUSG00000028857 | Tmem222 | transmembrane protein 222 [Source:MGI Symbol;Acc:MGI:1098568] | 733,2 | 100,00 |
| ENSMUSG00000027363 | Usp8 | ubiquitin specific peptidase 8 [Source:MGI Symbol;Acc:MGI:1934029] | 732,5 | 99,10 |
| ENSMUSG00000026510 | Trp53bp2 | transformation related protein 53 binding protein 2 [Source:MGI Symbol;Acc:MGI:2138319] | 732,2 | 99,93 |
| ENSMUSG00000032018 | Sc5d | ·C5-desaturase (fungal ERG3, delta-5-desaturase) homolog (S. cerevisae) [Source:MGI Symbol;Acc:MGI:135 | 730,2 | 96,58 |
| ENSMUSG00000025374 | Nabp2 | nucleic acid binding protein 2 [Source:MGI Symbol;Acc:MGI:1917167] | 729,7 | 78,31 |
| ENSMUSG00000026185 | Igfbp5 | insulin-like growth factor binding protein 5 [Source:MGI Symbol;Acc:MGI:96440] | 729,2 | 99,58 |
| ENSMUSG00000059409 | Ppp2r5d | protein phosphatase 2, regulatory subunit B', delta [Source:MGI Symbol;Acc:MGI:2388481] | 729,1 | 100,00 |
| ENSMUSG00000037032 | Apbb1 | amyloid beta (A4) precursor protein-binding, family B, member 1 [Source:MGI Symbol;Acc:MGI:107765] | 729,0 | 79,22 |
| ENSMUSG00000035559 | Mpv17l2 | MPV17 mitochondrial membrane protein-like 2 [Source:MGI Symbol;Acc:MGI:2681846] | 728,7 | 100,00 |
| ENSMUSG00000024067 | Dpy30 | dpy-30, histone methyltransferase complex regulatory subunit [Source:MGI Symbol;Acc:MGI:1913560] | 728,2 | 73,92 |
| ENSMUSG00000046668 | Cxxc5 | CXXC finger 5 [Source:MGI Symbol;Acc:MGI:1914643] | 728,1 | 99,97 |
| ENSMUSG00000055932 | Fto | fat mass and obesity associated [Source:MGI Symbol;Acc:MGI:1347093] | 728,0 | 88,01 |
| ENSMUSG00000029474 | Rnf34 | ring finger protein 34 [Source:MGI Symbol;Acc:MGI:2153340] | 727,2 | 75,82 |
| ENSMUSG00000030327 | Necap1 | NECAP endocytosis associated 1 [Source:MGI Symbol;Acc:MGI:1914852] | 727,1 | 97,31 |
| ENSMUSG00000021102 | Glrx5 | glutaredoxin 5 [Source:MGI Symbol;Acc:MGI:1920296] | 727,0 | 99,68 |
| ENSMUSG00000028988 | Ctnnbip1 | catenin beta interacting protein 1 [Source:MGI Symbol;Acc:MGI:1915756] | 726,8 | 95,76 |
| ENSMUSG00000019433 | Gipc1 | GIPC PDZ domain containing family, member 1 [Source:MGI Symbol;Acc:MGI:1926252] | 725,8 | 100,00 |
| ENSMUSG00000024197 | Plin3 | perilipin 3 [Source:MGI Symbol;Acc:MGI:1914155] | 724,2 | 99,95 |
| ENSMUSG00000025381 | Cnpy2 | canopy FGF signaling regulator 2 [Source:MGI Symbol;Acc:MGI:1928477] | 724,0 | 97,51 |
| ENSMUSG00000035443 | Thyn1 | thymocyte nuclear protein 1 [Source:MGI Symbol;Acc:MGI:1925112] | 724,0 | 100,00 |
| ENSMUSG00000024530 | Slmo1 | slowmo homolog 1 (Drosophila) [Source:MGI Symbol;Acc:MGI:2442865] | 723,5 | 99,94 |
| ENSMUSG00000003948 | Mmd | monocyte to macrophage differentiation-associated [Source:MGI Symbol;Acc:MGI:1914718] | 723,4 | 79,70 |
| ENSMUSG00000057342 | Sphk2 | sphingosine kinase 2 [Source:MGI Symbol;Acc:MGI:1861380] | 723,3 | 93,05 |
| ENSMUSG00000031827 | Cotl1 | coactosin-like 1 (Dictyostelium) [Source:MGI Symbol;Acc:MGI:1919292] | 723,2 | 99,65 |
| ENSMUSG00000040097 | Flywch1 | FLYWCH-type zinc finger 1 [Source:MGI Symbol;Acc:MGI:2442638] | 723,2 | 85,94 |
| ENSMUSG00000040848 | Sft2d2 | SFT2 domain containing 2 [Source:MGI Symbol;Acc:MGI:1917362] | 722,8 | 98,52 |
| ENSMUSG00000041733 | Coq5 | coenzyme Q5 methyltransferase [Source:MGI Symbol;Acc:MGI:1098643] | 722,5 | 99,57 |
| ENSMUSG00000029634 | Rnf6 | ring finger protein (C3H2C3 type) 6 [Source:MGI Symbol;Acc:MGI:1921382] | 722,4 | 94,69 |
| ENSMUSG00000034252 | Senp6 | SUMO/sentrin specific peptidase 6 [Source:MGI Symbol;Acc:MGI:1922075] | 722,4 | 85,86 |
| ENSMUSG00000042670 | Immp1l | IMP1 inner mitochondrial membrane peptidase-like (S. cerevisiae) [Source:MGI Symbol;Acc:MGI:1913791] | 721,9 | 81,47 |
| ENSMUSG00000041958 | Pigs | phosphatidylinositol glycan anchor biosynthesis, class S [Source:MGI Symbol;Acc:MGI:2687325] | 721,6 | 97,15 |
| ENSMUSG00000042750 | Bex2 | brain expressed X-linked 2 [Source:MGI Symbol;Acc:MGI:1338017] | 720,8 | 90,37 |
| ENSMUSG00000040128 | Pnrc1 | proline-rich nuclear receptor coactivator 1 [Source:MGI Symbol;Acc:MGI:1917838] | 720,6 | 99,88 |
| ENSMUSG00000046598 | Bdh1 | 3-hydroxybutyrate dehydrogenase, type 1 [Source:MGI Symbol;Acc:MGI:1919161] | 720,5 | 85,07 |
| ENSMUSG00000030888 | Rrp8 | ribosomal RNA processing 8, methyltransferase, homolog (yeast) [Source:MGI Symbol;Acc:MGI:1914251] | 720,4 | 91,96 |
| ENSMUSG00000014402 | Tsg101 | tumor susceptibility gene 101 [Source:MGI Symbol;Acc:MGI:106581] | 720,2 | 85,40 |
| ENSMUSG00000035847 | Ids | iduronate 2-sulfatase [Source:MGI Symbol;Acc:MGI:96417] | 719,8 | 96,62 |
| ENSMUSG00000027274 | Mkks | McKusick-Kaufman syndrome [Source:MGI Symbol;Acc:MGI:1891836] | 719,2 | 97,89 |
| ENSMUSG00000042608 | Stk40 | serine/threonine kinase 40 [Source:MGI Symbol;Acc:MGI:1921428] | 718,7 | 92,56 |
| ENSMUSG00000041278 | Ttc1 | tetratricopeptide repeat domain 1 [Source:MGI Symbol;Acc:MGI:1914077] | 718,5 | 99,63 |
| ENSMUSG00000030869 | Ndufab1 | NADH dehydrogenase (ubiquinone) 1, alpha/beta subcomplex, 1 [Source:MGI Symbol;Acc:MGI:1917566] | 717,6 | 87,20 |
| ENSMUSG00000030357 | Fkbp4 | FK506 binding protein 4 [Source:MGI Symbol;Acc:MGI:95543] | 717,4 | 96,32 |
| ENSMUSG00000011831 | Evi5 | ecotropic viral integration site 5 [Source:MGI Symbol;Acc:MGI:104736] | 717,2 | 94,18 |
| ENSMUSG00000049354 | Dcaf7 | DDB1 and CUL4 associated factor 7 [Source:MGI Symbol;Acc:MGI:1919083] | 717,1 | 99,02 |
| ENSMUSG00000025544 | Tm9sf2 | transmembrane 9 superfamily member 2 [Source:MGI Symbol;Acc:MGI:1915309] | 716,8 | 98,53 |
| ENSMUSG00000061740 | Cyp2d22 | cytochrome P450, family 2, subfamily d, polypeptide 22 [Source:MGI Symbol;Acc:MGI:1929474] | 715,9 | 99,70 |
| ENSMUSG00000052934 | Fbxo31 | F-box protein 31 [Source:MGI Symbol;Acc:MGI:1354708] | 715,5 | 93,39 |
| ENSMUSG00000024810 | Il33 | interleukin 33 [Source:MGI Symbol;Acc:MGI:1924375] | 715,0 | 87,76 |
| ENSMUSG00000022800 | Fyttd1 | forty-two-three domain containing 1 [Source:MGI Symbol;Acc:MGI:1917955] | 714,5 | 84,53 |
| ENSMUSG00000024327 | Slc39a7 | solute carrier family 39 (zinc transporter), member 7 [Source:MGI Symbol;Acc:MGI:95909] | 714,1 | 99,79 |
| ENSMUSG00000031446 | Cul4a | cullin 4A [Source:MGI Symbol;Acc:MGI:1914487] | 713,7 | 91,97 |
| ENSMUSG00000028261 | Ndufaf4 | DH dehydrogenase (ubiquinone) 1 alpha subcomplex, assembly factor 4 [Source:MGI Symbol;Acc:MGI:1915 | 713,7 | 91,02 |
| ENSMUSG00000054312 | Mrps21 | mitochondrial ribosomal protein S21 [Source:MGI Symbol;Acc:MGI:1913542] | 712,1 | 73,92 |
| ENSMUSG00000060703 | Cd302 | CD302 antigen [Source:MGI Symbol;Acc:MGI:1913455] | 711,3 | 79,67 |
| ENSMUSG00000087612 | A230005M16Rik | RIKEN cDNA A230005M16 gene [Source:MGI Symbol;Acc:MGI:3603401] | 711,2 | 89,41 |
| ENSMUSG00000034640 | Tiparp | TCDD-inducible poly(ADP-ribose) polymerase [Source:MGI Symbol;Acc:MGI:2159210] | 711,0 | 92,66 |
| ENSMUSG00000004945 | Tmem242 | transmembrane protein 242 [Source:MGI Symbol;Acc:MGI:1917794] | 711,0 | 80,89 |
| ENSMUSG00000003974 | Grm3 | glutamate receptor, metabotropic 3 [Source:MGI Symbol;Acc:MGI:1351340] | 710,8 | 100,00 |
| ENSMUSG00000045589 | Frrs1l | ferric-chelate reductase 1 like [Source:MGI Symbol;Acc:MGI:2442704] | 710,4 | 98,83 |

| | | | | |
|---|---|---|---|---|
| ENSMUSG00000004798 | Ulk2 | unc-51 like kinase 2 [Source:MGI Symbol;Acc:MGI:1352758] | 709,7 | 99,90 |
| ENSMUSG00000001445 | Mrpl10 | mitochondrial ribosomal protein L10 [Source:MGI Symbol;Acc:MGI:1333801] | 709,3 | 99,40 |
| ENSMUSG00000031357 | Syap1 | synapse associated protein 1 [Source:MGI Symbol;Acc:MGI:1914293] | 709,2 | 100,00 |
| ENSMUSG00000027195 | Hsd17b12 | hydroxysteroid (17-beta) dehydrogenase 12 [Source:MGI Symbol;Acc:MGI:1926967] | 708,9 | 99,70 |
| ENSMUSG00000063019 | Manbal | mannosidase, beta A, lysosomal-like [Source:MGI Symbol;Acc:MGI:1916411] | 708,7 | 100,00 |
| ENSMUSG00000029405 | G3bp2 | GTPase activating protein (SH3 domain) binding protein 2 [Source:MGI Symbol;Acc:MGI:2442040] | 708,6 | 89,24 |
| ENSMUSG00000022507 | 1810013L24Rik | RIKEN cDNA 1810013L24 gene [Source:MGI Symbol;Acc:MGI:1916303] | 708,4 | 99,07 |
| ENSMUSG00000041225 | Arhgap12 | Rho GTPase activating protein 12 [Source:MGI Symbol;Acc:MGI:1922665] | 708,2 | 95,86 |
| ENSMUSG00000022365 | Derl1 | Der1-like domain family, member 1 [Source:MGI Symbol;Acc:MGI:1915069] | 707,9 | 87,81 |
| ENSMUSG00000021986 | Amer2 | APC membrane recruitment 2 [Source:MGI Symbol;Acc:MGI:1919375] | 707,9 | 99,98 |
| ENSMUSG00000022013 | Dnajc15 | DnaJ heat shock protein family (Hsp40) member C15 [Source:MGI Symbol;Acc:MGI:1913398] | 707,9 | 100,00 |
| ENSMUSG00000020823 | Sec14l1 | SEC14-like lipid binding 1 [Source:MGI Symbol;Acc:MGI:1921386] | 707,7 | 99,17 |
| ENSMUSG00000027257 | Pacsin3 | protein kinase C and casein kinase substrate in neurons 3 [Source:MGI Symbol;Acc:MGI:1891410] | 706,9 | 73,91 |
| ENSMUSG00000006418 | Rnf114 | ring finger protein 114 [Source:MGI Symbol;Acc:MGI:1933159] | 706,8 | 81,94 |
| ENSMUSG00000032285 | Dnaja4 | DnaJ heat shock protein family (Hsp40) member A4 [Source:MGI Symbol;Acc:MGI:1927638] | 705,8 | 85,26 |
| ENSMUSG00000032526 | Deb1 | differentially expressed in B16F10 1 [Source:MGI Symbol;Acc:MGI:1349474] | 705,7 | 100,00 |
| ENSMUSG00000032383 | Ppib | peptidylprolyl isomerase B [Source:MGI Symbol;Acc:MGI:97750] | 705,2 | 97,15 |
| ENSMUSG00000055447 | Cd47 | CD47 antigen (Rh-related antigen, integrin-associated signal transducer) [Source:MGI Symbol;Acc:MGI:96617 | 705,1 | 76,50 |
| ENSMUSG00000045193 | Cirbp | cold inducible RNA binding protein [Source:MGI Symbol;Acc:MGI:893588] | 704,9 | 87,95 |
| ENSMUSG00000021025 | Nfkbia | ar factor of kappa light polypeptide gene enhancer in B cells inhibitor, alpha [Source:MGI Symbol;Acc:MGI:10 | 704,4 | 97,02 |
| ENSMUSG00000026914 | Psmd14 | proteasome (prosome, macropain) 26S subunit, non-ATPase, 14 [Source:MGI Symbol;Acc:MGI:1913284] | 703,6 | 98,90 |
| ENSMUSG00000013089 | Etv5 | ets variant 5 [Source:MGI Symbol;Acc:MGI:1096867] | 703,6 | 84,29 |
| ENSMUSG00000062591 | Tubb4a | tubulin, beta 4A class IVA [Source:MGI Symbol;Acc:MGI:107848] | 703,2 | 94,17 |
| ENSMUSG00000034412 | Tbc1d10a | TBC1 domain family, member 10a [Source:MGI Symbol;Acc:MGI:2144164] | 702,4 | 99,61 |
| ENSMUSG00000018858 | Ict1 | immature colon carcinoma transcript 1 [Source:MGI Symbol;Acc:MGI:1915822] | 702,0 | 77,48 |
| ENSMUSG00000024360 | Etf1 | eukaryotic translation termination factor 1 [Source:MGI Symbol;Acc:MGI:2385071] | 701,0 | 95,51 |
| ENSMUSG00000005625 | Psmd4 | proteasome (prosome, macropain) 26S subunit, non-ATPase, 4 [Source:MGI Symbol;Acc:MGI:1201670] | 701,0 | 71,36 |
| ENSMUSG00000020184 | Mdm2 | transformed mouse 3T3 cell double minute 2 [Source:MGI Symbol;Acc:MGI:96952] | 700,9 | 97,72 |
| ENSMUSG00000038387 | Rras | Harvey rat sarcoma oncogene, subgroup R [Source:MGI Symbol;Acc:MGI:98179] | 700,7 | 99,86 |
| ENSMUSG00000001416 | Cct3 | chaperonin containing Tcp1, subunit 3 (gamma) [Source:MGI Symbol;Acc:MGI:104708] | 700,6 | 86,84 |
| ENSMUSG00000038286 | Bphl | l hydrolase-like (serine hydrolase, breast epithelial mucin-associated antigen) [Source:MGI Symbol;Acc:MGI:1 | 700,5 | 100,00 |
| ENSMUSG00000021607 | Mrpl36 | mitochondrial ribosomal protein L36 [Source:MGI Symbol;Acc:MGI:2137228] | 699,9 | 100,00 |
| ENSMUSG00000071415 | Rpl23 | ribosomal protein L23 [Source:MGI Symbol;Acc:MGI:1929455] | 699,7 | 93,26 |
| ENSMUSG00000026617 | Bpnt1 | bisphosphate 3'-nucleotidase 1 [Source:MGI Symbol;Acc:MGI:1338800] | 699,7 | 96,99 |
| ENSMUSG00000020116 | Pno1 | partner of NOB1 homolog (S. cerevisiae) [Source:MGI Symbol;Acc:MGI:1913499] | 699,4 | 99,76 |
| ENSMUSG00000042650 | Alkbh5 | alkB homolog 5, RNA demethylase [Source:MGI Symbol;Acc:MGI:2144489] | 699,1 | 72,83 |
| ENSMUSG00000034880 | Mrpl34 | mitochondrial ribosomal protein L34 [Source:MGI Symbol;Acc:MGI:2137227] | 699,0 | 99,95 |
| ENSMUSG00000038628 | Polr3k | polymerase (RNA) III (DNA directed) polypeptide K [Source:MGI Symbol;Acc:MGI:1914255] | 698,8 | 95,68 |
| ENSMUSG00000066900 | Suds3 | suppressor of defective silencing 3 homolog (S. cerevisiae) [Source:MGI Symbol;Acc:MGI:1919204] | 698,7 | 94,49 |
| ENSMUSG00000001366 | Fbxo9 | f-box protein 9 [Source:MGI Symbol;Acc:MGI:1918788] | 698,1 | 90,82 |
| ENSMUSG00000027472 | Pdrg1 | p53 and DNA damage regulated 1 [Source:MGI Symbol;Acc:MGI:1915809] | 698,0 | 97,38 |
| ENSMUSG00000028032 | Papss1 | 3'-phosphoadenosine 5'-phosphosulfate synthase 1 [Source:MGI Symbol;Acc:MGI:1330587] | 697,9 | 95,47 |
| ENSMUSG00000026005 | Rpe | ribulose-5-phosphate-3-epimerase [Source:MGI Symbol;Acc:MGI:1913896] | 697,4 | 98,80 |
| ENSMUSG00000027962 | Vcam1 | vascular cell adhesion molecule 1 [Source:MGI Symbol;Acc:MGI:98926] | 695,7 | 76,03 |
| ENSMUSG00000022982 | Sod1 | superoxide dismutase 1, soluble [Source:MGI Symbol;Acc:MGI:98351] | 695,4 | 100,00 |
| ENSMUSG00000020003 | Pex7 | peroxisomal biogenesis factor 7 [Source:MGI Symbol;Acc:MGI:1321392] | 695,0 | 99,48 |
| ENSMUSG00000024844 | Banf1 | barrier to autointegration factor 1 [Source:MGI Symbol;Acc:MGI:1346330] | 694,4 | 100,00 |
| ENSMUSG00000020477 | Mrps24 | mitochondrial ribosomal protein S24 [Source:MGI Symbol;Acc:MGI:1928142] | 693,8 | 81,21 |
| ENSMUSG00000040006 | Ginm1 | glycoprotein integral membrane 1 [Source:MGI Symbol;Acc:MGI:2384905] | 693,5 | 99,07 |
| ENSMUSG00000022437 | Samm50 | orting and assembly machinery component 50 homolog (S. cerevisiae) [Source:MGI Symbol;Acc:MGI:191590 | 693,4 | 100,00 |
| ENSMUSG00000065979 | Cpped1 | calcineurin-like phosphoesterase domain containing 1 [Source:MGI Symbol;Acc:MGI:2443300] | 693,2 | 96,72 |
| ENSMUSG00000025702 | March8 | membrane-associated ring finger (C3HC4) 8 [Source:MGI Symbol;Acc:MGI:1919029] | 692,6 | 90,01 |
| ENSMUSG00000040712 | Camta2 | calmodulin binding transcription activator 2 [Source:MGI Symbol;Acc:MGI:2135957] | 688,8 | 96,83 |
| ENSMUSG00000022490 | Ppp1r1a | protein phosphatase 1, regulatory (inhibitor) subunit 1A [Source:MGI Symbol;Acc:MGI:1889595] | 688,3 | 100,00 |
| ENSMUSG00000024429 | Gnl1 | guanine nucleotide binding protein-like 1 [Source:MGI Symbol;Acc:MGI:95764] | 688,2 | 94,45 |
| ENSMUSG00000028797 | Tmem234 | transmembrane protein 234 [Source:MGI Symbol;Acc:MGI:1924049] | 688,1 | 100,00 |
| ENSMUSG00000036095 | Dgkb | diacylglycerol kinase, beta [Source:MGI Symbol;Acc:MGI:2442474] | 687,5 | 100,00 |
| ENSMUSG00000014748 | Tex261 | testis expressed gene 261 [Source:MGI Symbol;Acc:MGI:1096575] | 687,4 | 100,00 |
| ENSMUSG00000026051 | 1500015O10Rik | RIKEN cDNA 1500015O10 gene [Source:MGI Symbol;Acc:MGI:1926146] | 686,5 | 100,00 |
| ENSMUSG00000058690 | Ccser2 | coiled-coil serine rich 2 [Source:MGI Symbol;Acc:MGI:101859] | 686,3 | 96,99 |
| ENSMUSG00000025935 | Tram1 | translocating chain-associating membrane protein 1 [Source:MGI Symbol;Acc:MGI:1919515] | 685,3 | 99,85 |
| ENSMUSG00000033352 | Map2k4 | mitogen-activated protein kinase kinase 4 [Source:MGI Symbol;Acc:MGI:1346869] | 685,2 | 99,37 |
| ENSMUSG00000032412 | Atp1b3 | ATPase, Na+/K+ transporting, beta 3 polypeptide [Source:MGI Symbol;Acc:MGI:107788] | 684,6 | 94,80 |
| ENSMUSG00000050144 | Slc25a44 | solute carrier family 25, member 44 [Source:MGI Symbol;Acc:MGI:2444391] | 684,1 | 99,92 |
| ENSMUSG00000028266 | Lmo4 | LIM domain only 4 [Source:MGI Symbol;Acc:MGI:109360] | 683,2 | 84,00 |
| ENSMUSG00000058833 | 2810428I15Rik | RIKEN cDNA 2810428I15 gene [Source:MGI Symbol;Acc:MGI:1913712] | 683,1 | 100,00 |
| ENSMUSG00000031605 | Klhl2 | kelch-like 2, Mayven [Source:MGI Symbol;Acc:MGI:1924363] | 682,6 | 100,00 |
| ENSMUSG00000038991 | Txndc5 | thioredoxin domain containing 5 [Source:MGI Symbol;Acc:MGI:2145316] | 682,4 | 92,07 |
| ENSMUSG00000004789 | Dlst | lipoamide S-succinyltransferase (E2 component of 2-oxo-glutarate complex) [Source:MGI Symbol;Acc:MGI:1 | 682,3 | 99,41 |
| ENSMUSG00000035585 | Tsen34 | tRNA splicing endonuclease 34 homolog (S. cerevisiae) [Source:MGI Symbol;Acc:MGI:1913328] | 682,1 | 89,44 |
| ENSMUSG00000039960 | Rhou | ras homolog gene family, member U [Source:MGI Symbol;Acc:MGI:1916831] | 682,1 | 99,37 |
| ENSMUSG00000044340 | Phlpp1 | PH domain and leucine rich repeat protein phosphatase 1 [Source:MGI Symbol;Acc:MGI:2138327] | 681,4 | 98,28 |
| ENSMUSG00000042043 | Tbca | tubulin cofactor A [Source:MGI Symbol;Acc:MGI:107549] | 680,2 | 100,00 |
| ENSMUSG00000028729 | Ebna1bp2 | EBNA1 binding protein 2 [Source:MGI Symbol;Acc:MGI:1916322] | 679,3 | 97,62 |
| ENSMUSG00000025894 | Aasdhppt | oadipate-semialdehyde dehydrogenase-phosphopantetheinyl transferase [Source:MGI Symbol;Acc:MGI:1914 | 679,2 | 99,40 |
| ENSMUSG00000022516 | Nudt16l1 | nudix (nucleoside diphosphate linked moiety X)-type motif 16-like 1 [Source:MGI Symbol;Acc:MGI:1914161] | 679,1 | 99,92 |
| ENSMUSG00000024515 | Smad4 | SMAD family member 4 [Source:MGI Symbol;Acc:MGI:894293] | 676,4 | 94,67 |
| ENSMUSG00000020720 | Psmd12 | proteasome (prosome, macropain) 26S subunit, non-ATPase, 12 [Source:MGI Symbol;Acc:MGI:1914247] | 675,8 | 71,08 |
| ENSMUSG00000078851 | Hist3h2a | histone cluster 3, H2a [Source:MGI Symbol;Acc:MGI:2448458] | 675,2 | 97,93 |
| ENSMUSG00000027665 | Pik3ca | phosphatidylinositol 3-kinase, catalytic, alpha polypeptide [Source:MGI Symbol;Acc:MGI:1206581] | 674,2 | 78,78 |
| ENSMUSG00000024414 | Mrpl27 | mitochondrial ribosomal protein L27 [Source:MGI Symbol;Acc:MGI:2137224] | 673,9 | 98,59 |
| ENSMUSG00000021832 | Psmc6 | proteasome (prosome, macropain) 26S subunit, ATPase, 6 [Source:MGI Symbol;Acc:MGI:1914339] | 673,9 | 100,00 |
| ENSMUSG00000025326 | Ube3a | ubiquitin protein ligase E3A [Source:MGI Symbol;Acc:MGI:105098] | 673,7 | 79,51 |
| ENSMUSG00000022451 | Twf1 | twinfilin, actin-binding protein, homolog 1 (Drosophila) [Source:MGI Symbol;Acc:MGI:1100520] | 673,3 | 98,23 |
| ENSMUSG00000052387 | Trpm3 | transient receptor potential cation channel, subfamily M, member 3 [Source:MGI Symbol;Acc:MGI:2443101] | 673,1 | 94,19 |
| ENSMUSG00000017760 | Ctsa | cathepsin A [Source:MGI Symbol;Acc:MGI:97748] | 672,4 | 73,57 |
| ENSMUSG00000028121 | Bcar3 | breast cancer anti-estrogen resistance 3 [Source:MGI Symbol;Acc:MGI:1352501] | 671,8 | 85,28 |
| ENSMUSG00000025130 | P4hb | prolyl 4-hydroxylase, beta polypeptide [Source:MGI Symbol;Acc:MGI:97464] | 671,0 | 95,99 |
| ENSMUSG00000059070 | Rpl18 | ribosomal protein L18 [Source:MGI Symbol;Acc:MGI:98003] | 669,5 | 95,09 |
| ENSMUSG00000039485 | Tspyl4 | TSPY-like 4 [Source:MGI Symbol;Acc:MGI:106393] | 669,5 | 99,86 |
| ENSMUSG00000035569 | Ankrd11 | ankyrin repeat domain 11 [Source:MGI Symbol;Acc:MGI:1924337] | 669,3 | 96,84 |
| ENSMUSG00000000751 | Rpa1 | replication protein A1 [Source:MGI Symbol;Acc:MGI:1915525] | 669,3 | 70,27 |
| ENSMUSG00000021427 | Ssr1 | signal sequence receptor, alpha [Source:MGI Symbol;Acc:MGI:105082] | 668,7 | 88,57 |
| ENSMUSG00000046562 | Unc119b | unc-119 homolog B (C. elegans) [Source:MGI Symbol;Acc:MGI:2147162] | 668,7 | 98,84 |
| ENSMUSG00000025221 | Kcnip2 | Kv channel-interacting protein 2 [Source:MGI Symbol;Acc:MGI:2135916] | 668,2 | 85,89 |
| ENSMUSG00000060376 | Bckdha | branched chain ketoacid dehydrogenase E1, alpha polypeptide [Source:MGI Symbol;Acc:MGI:107701] | 668,2 | 94,75 |
| ENSMUSG00000036565 | Ttyh3 | tweety homolog 3 (Drosophila) [Source:MGI Symbol;Acc:MGI:1925589] | 668,1 | 97,39 |
| ENSMUSG00000024900 | Cpt1a | carnitine palmitoyltransferase 1a, liver [Source:MGI Symbol;Acc:MGI:1098296] | 667,9 | 99,44 |
| ENSMUSG00000064215 | Ifi27 | interferon, alpha-inducible protein 27 [Source:MGI Symbol;Acc:MGI:1277180] | 666,7 | 99,74 |
| ENSMUSG00000045948 | Mrps12 | mitochondrial ribosomal protein S12 [Source:MGI Symbol;Acc:MGI:1346333] | 666,5 | 88,85 |
| ENSMUSG00000022340 | Sybu | syntabulin (syntaxin-interacting) [Source:MGI Symbol;Acc:MGI:2442392] | 666,5 | 92,49 |
| ENSMUSG00000060601 | Nr1h2 | nuclear receptor subfamily 1, group H, member 2 [Source:MGI Symbol;Acc:MGI:1352463] | 666,5 | 79,57 |
| ENSMUSG00000025321 | Itgb8 | integrin beta 8 [Source:MGI Symbol;Acc:MGI:1338035] | 666,3 | 99,87 |
| ENSMUSG00000073987 | Ggh | gamma-glutamyl hydrolase [Source:MGI Symbol;Acc:MGI:1329035] | 666,1 | 94,01 |
| ENSMUSG00000031505 | Carkd | carbohydrate kinase domain containing [Source:MGI Symbol;Acc:MGI:1913353] | 666,0 | 100,00 |
| ENSMUSG00000058488 | Kl | klotho [Source:MGI Symbol;Acc:MGI:1101771] | 665,6 | 98,34 |
| ENSMUSG00000020457 | Drg1 | developmentally regulated GTP binding protein 1 [Source:MGI Symbol;Acc:MGI:1343297] | 664,6 | 100,00 |
| ENSMUSG00000079659 | Tmem243 | transmembrane protein 243, mitochondrial [Source:MGI Symbol;Acc:MGI:3606159] | 664,5 | 86,45 |
| ENSMUSG00000020230 | Prmt2 | protein arginine N-methyltransferase 2 [Source:MGI Symbol;Acc:MGI:1316652] | 664,0 | 97,00 |
| ENSMUSG00000032517 | Mobp | myelin-associated oligodendrocytic basic protein [Source:MGI Symbol;Acc:MGI:108511] | 664,0 | 95,42 |
| ENSMUSG00000032397 | Tipin | timeless interacting protein [Source:MGI Symbol;Acc:MGI:1921571] | 662,7 | 100,00 |
| ENSMUSG00000023967 | Mrps18a | mitochondrial ribosomal protein S18A [Source:MGI Symbol;Acc:MGI:1915815] | 662,4 | 85,21 |
| ENSMUSG00000000876 | Pxmp4 | peroxisomal membrane protein 4 [Source:MGI Symbol;Acc:MGI:1891701] | 662,2 | 95,07 |
| ENSMUSG00000025041 | Nt5c2 | 5'-nucleotidase, cytosolic II [Source:MGI Symbol;Acc:MGI:2178563] | 661,7 | 97,35 |
| ENSMUSG00000016256 | Ctsz | cathepsin Z [Source:MGI Symbol;Acc:MGI:1891190] | 661,5 | 97,57 |
| ENSMUSG00000050017 | Pitpnb | phosphatidylinositol transfer protein, beta [Source:MGI Symbol;Acc:MGI:1927542] | 661,2 | 100,00 |
| ENSMUSG00000028608 | 0610037L13Rik | RIKEN cDNA 0610037L13 gene [Source:MGI Symbol;Acc:MGI:1921348] | 660,8 | 82,07 |

| | | | | |
|---|---|---|---|---|
| ENSMUSG00000038497 | Tmco3 | transmembrane and coiled-coil domains 3 [Source:MGI Symbol;Acc:MGI:2444946] | 660,7 | 82,67 |
| ENSMUSG00000038612 | Mcl1 | myeloid cell leukemia sequence 1 [Source:MGI Symbol;Acc:MGI:101769] | 660,4 | 99,96 |
| ENSMUSG00000041084 | Ostc | oligosaccharyltransferase complex subunit [Source:MGI Symbol;Acc:MGI:1913607] | 659,7 | 99,56 |
| ENSMUSG00000079179 | Rab10os | RAB10, member RAS oncogene family, opposite strand [Source:MGI Symbol;Acc:MGI:1921423] | 659,6 | 77,36 |
| ENSMUSG00000029461 | Fam168a | family with sequence similarity 168, member A [Source:MGI Symbol;Acc:MGI:2442372] | 659,4 | 98,69 |
| ENSMUSG00000030682 | Cdipt | acylglycerol--inositol 3-phosphatidyltransferase (phosphatidylinositol synthase) [Source:MGI Symbol;Acc:MGI:1 | 658,9 | 83,19 |
| ENSMUSG00000014177 | Tvp23b | trans-golgi network vesicle protein 23B [Source:MGI Symbol;Acc:MGI:1914760] | 658,9 | 100,00 |
| ENSMUSG00000031485 | Prosc | proline synthetase co-transcribed [Source:MGI Symbol;Acc:MGI:1891207] | 658,8 | 70,34 |
| ENSMUSG00000028443 | Nudt2 | nudix (nucleoside diphosphate linked moiety X)-type motif 2 [Source:MGI Symbol;Acc:MGI:1913651] | 658,0 | 100,00 |
| ENSMUSG00000057181 | 5730455P16Rik | RIKEN cDNA 5730455P16 gene [Source:MGI Symbol;Acc:MGI:1917841] | 657,4 | 96,33 |
| ENSMUSG00000030410 | Dmwd | dystrophia myotonica-containing WD repeat motif [Source:MGI Symbol;Acc:MGI:94907] | 657,0 | 99,81 |
| ENSMUSG00000031791 | Tmem38a | transmembrane protein 38A [Source:MGI Symbol;Acc:MGI:1921416] | 656,7 | 99,97 |
| ENSMUSG00000079436 | Kcnj13 | potassium inwardly-rectifying channel, subfamily J, member 13 [Source:MGI Symbol;Acc:MGI:3781032] | 655,6 | 99,95 |
| ENSMUSG00000002741 | Ykt6 | YKT6 v-SNARE homolog (S. cerevisiae) [Source:MGI Symbol;Acc:MGI:1927550] | 654,9 | 99,02 |
| ENSMUSG00000030207 | Fam234b | family with sequence similarity 234, member B [Source:MGI Symbol;Acc:MGI:1921775] | 653,8 | 78,81 |
| ENSMUSG00000022040 | Ephx2 | epoxide hydrolase 2, cytoplasmic [Source:MGI Symbol;Acc:MGI:99500] | 653,6 | 100,00 |
| ENSMUSG00000019978 | Epb41l2 | erythrocyte membrane protein band 4.1 like 2 [Source:MGI Symbol;Acc:MGI:103009] | 653,6 | 91,73 |
| ENSMUSG00000005804 | Bloc1s6 | biogenesis of lysosomal organelles complex-1, subunit 6, pallidin [Source:MGI Symbol;Acc:MGI:1927580] | 653,5 | 77,62 |
| ENSMUSG00000032547 | Ryk | receptor-like tyrosine kinase [Source:MGI Symbol;Acc:MGI:101766] | 653,5 | 99,61 |
| ENSMUSG00000045410 | Akr1e1 | aldo-keto reductase family 1, member E1 [Source:MGI Symbol;Acc:MGI:1914758] | 653,1 | 96,40 |
| ENSMUSG00000024767 | Otub1 | OTU domain, ubiquitin aldehyde binding 1 [Source:MGI Symbol;Acc:MGI:2147616] | 653,0 | 88,17 |
| ENSMUSG00000031885 | Cbfb | core binding factor beta [Source:MGI Symbol;Acc:MGI:99851] | 652,7 | 97,68 |
| ENSMUSG00000031820 | Babam1 | BRISC and BRCA1 A complex member 1 [Source:MGI Symbol;Acc:MGI:1915501] | 652,3 | 94,18 |
| ENSMUSG00000035268 | Pkig | protein kinase inhibitor, gamma [Source:MGI Symbol;Acc:MGI:1343086] | 652,1 | 86,98 |
| ENSMUSG00000022234 | Cct5 | chaperonin containing Tcp1, subunit 5 (epsilon) [Source:MGI Symbol;Acc:MGI:107185] | 652,0 | 95,81 |
| ENSMUSG00000017843 | Ppp2r5c | protein phosphatase 2, regulatory subunit B', gamma [Source:MGI Symbol;Acc:MGI:1349475] | 651,8 | 99,87 |
| ENSMUSG00000026173 | Plcd4 | phospholipase C, delta 4 [Source:MGI Symbol;Acc:MGI:107469] | 651,1 | 87,92 |
| ENSMUSG00000060147 | Serpinb6a | serine (or cysteine) peptidase inhibitor, clade B, member 6a [Source:MGI Symbol;Acc:MGI:103123] | 651,0 | 88,61 |
| ENSMUSG00000033845 | Mrpl15 | mitochondrial ribosomal protein L15 [Source:MGI Symbol;Acc:MGI:1351639] | 650,9 | 79,59 |
| ENSMUSG00000044629 | Cnrip1 | cannabinoid receptor interacting protein 1 [Source:MGI Symbol;Acc:MGI:1917505] | 650,9 | 81,43 |
| ENSMUSG00000024949 | Sf1 | splicing factor 1 [Source:MGI Symbol;Acc:MGI:1095403] | 650,4 | 88,35 |
| ENSMUSG00000037656 | Slc20a2 | solute carrier family 20, member 2 [Source:MGI Symbol;Acc:MGI:97851] | 649,8 | 98,83 |
| ENSMUSG00000075415 | Fnbp1 | formin binding protein 1 [Source:MGI Symbol;Acc:MGI:109606] | 649,8 | 88,13 |
| ENSMUSG00000031865 | Dctn1 | dynactin 1 [Source:MGI Symbol;Acc:MGI:107745] | 649,6 | 89,47 |
| ENSMUSG00000025980 | Hspd1 | heat shock protein 1 (chaperonin) [Source:MGI Symbol;Acc:MGI:96242] | 649,4 | 91,16 |
| ENSMUSG00000026893 | Gca | grancalcin [Source:MGI Symbol;Acc:MGI:1918521] | 649,1 | 99,74 |
| ENSMUSG00000031007 | Atp6ap2 | ATPase, H+ transporting, lysosomal accessory protein 2 [Source:MGI Symbol;Acc:MGI:1917745] | 649,0 | 99,54 |
| ENSMUSG00000038615 | Nfe2l1 | nuclear factor, erythroid derived 2,-like 1 [Source:MGI Symbol;Acc:MGI:99421] | 649,0 | 96,66 |
| ENSMUSG00000024911 | Fibp | fibroblast growth factor (acidic) intracellular binding protein [Source:MGI Symbol;Acc:MGI:1926233] | 648,4 | 99,76 |
| ENSMUSG00000025932 | Eya1 | EYA transcriptional coactivator and phosphatase 1 [Source:MGI Symbol;Acc:MGI:109344] | 648,3 | 96,98 |
| ENSMUSG00000019782 | Rwdd1 | RWD domain containing 1 [Source:MGI Symbol;Acc:MGI:1913771] | 648,2 | 81,96 |
| ENSMUSG00000020037 | Rfx4 | regulatory factor X, 4 (influences HLA class II expression) [Source:MGI Symbol;Acc:MGI:1918387] | 648,1 | 93,61 |
| ENSMUSG00000004040 | Stat3 | signal transducer and activator of transcription 3 [Source:MGI Symbol;Acc:MGI:103038] | 647,9 | 72,57 |
| ENSMUSG00000020283 | Pex13 | peroxisomal biogenesis factor 13 [Source:MGI Symbol;Acc:MGI:1919379] | 647,7 | 92,46 |
| ENSMUSG00000026727 | Rsu1 | Ras suppressor protein 1 [Source:MGI Symbol;Acc:MGI:103040] | 647,6 | 90,41 |
| ENSMUSG00000079215 | Zfp664 | zinc finger protein 664 [Source:MGI Symbol;Acc:MGI:2442505] | 647,5 | 99,76 |
| ENSMUSG00000031367 | Ap1s2 | adaptor-related protein complex 1, sigma 2 subunit [Source:MGI Symbol;Acc:MGI:1889383] | 647,2 | 89,78 |
| ENSMUSG00000024403 | Atp6v1g2 | ATPase, H+ transporting, lysosomal V1 subunit G2 [Source:MGI Symbol;Acc:MGI:1913487] | 647,1 | 98,50 |
| ENSMUSG00000025967 | Eef1b2 | eukaryotic translation elongation factor 1 beta 2 [Source:MGI Symbol;Acc:MGI:1929520] | 647,0 | 81,06 |
| ENSMUSG00000021039 | Snw1 | SNW domain containing 1 [Source:MGI Symbol;Acc:MGI:1913604] | 646,8 | 100,00 |
| ENSMUSG00000029156 | Sgcb | sarcoglycan, beta (dystrophin-associated glycoprotein) [Source:MGI Symbol;Acc:MGI:1346523] | 646,8 | 99,08 |
| ENSMUSG00000055762 | Eef1d | ic translation elongation factor 1 delta (guanine nucleotide exchange protein) [Source:MGI Symbol;Acc:MGI:1 | 646,7 | 79,77 |
| ENSMUSG00000022463 | Srebf2 | sterol regulatory element binding factor 2 [Source:MGI Symbol;Acc:MGI:107585] | 646,6 | 98,40 |
| ENSMUSG00000021094 | Dhrs7 | dehydrogenase/reductase (SDR family) member 7 [Source:MGI Symbol;Acc:MGI:1913625] | 646,4 | 100,00 |
| ENSMUSG00000027763 | Mbnl1 | muscleblind-like 1 (Drosophila) [Source:MGI Symbol;Acc:MGI:1928482] | 646,3 | 93,19 |
| ENSMUSG00000027189 | Trim44 | tripartite motif-containing 44 [Source:MGI Symbol;Acc:MGI:1931835] | 646,0 | 99,20 |
| ENSMUSG00000021610 | Clptm1l | CLPTM1-like [Source:MGI Symbol;Acc:MGI:2442892] | 645,9 | 99,43 |
| ENSMUSG00000006315 | Tmem147 | transmembrane protein 147 [Source:MGI Symbol;Acc:MGI:1915011] | 645,6 | 95,66 |
| ENSMUSG00000024507 | Hsd17b4 | hydroxysteroid (17-beta) dehydrogenase 4 [Source:MGI Symbol;Acc:MGI:105089] | 645,3 | 98,05 |
| ENSMUSG00000001924 | Uba1 | ubiquitin-like modifier activating enzyme 1 [Source:MGI Symbol;Acc:MGI:98890] | 645,2 | 95,46 |
| ENSMUSG00000032437 | Stt3b | subunit of the oligosaccharyltransferase complex, homolog B (S. cerevisiae) [Source:MGI Symbol;Acc:MGI:19 | 645,0 | 99,45 |
| ENSMUSG00000026926 | Pmpca | peptidase (mitochondrial processing) alpha [Source:MGI Symbol;Acc:MGI:1918568] | 644,8 | 95,77 |
| ENSMUSG00000037348 | Paqr7 | progestin and adipoQ receptor family member VII [Source:MGI Symbol;Acc:MGI:1919154] | 644,3 | 89,53 |
| ENSMUSG00000064246 | Chil1 | chitinase-like 1 [Source:MGI Symbol;Acc:MGI:1340899] | 644,2 | 93,08 |
| ENSMUSG00000038150 | Ormdl3 | ORM1-like 3 (S. cerevisiae) [Source:MGI Symbol;Acc:MGI:1913862] | 644,2 | 95,92 |
| ENSMUSG00000022951 | Rcan1 | regulator of calcineurin 1 [Source:MGI Symbol;Acc:MGI:1890564] | 643,6 | 99,91 |
| ENSMUSG00000014551 | Mrps25 | mitochondrial ribosomal protein S25 [Source:MGI Symbol;Acc:MGI:1928140] | 643,6 | 89,70 |
| ENSMUSG00000032540 | Abhd5 | abhydrolase domain containing 5 [Source:MGI Symbol;Acc:MGI:1914719] | 643,5 | 93,73 |
| ENSMUSG00000022540 | Rogdi | rogdi homolog (Drosophila) [Source:MGI Symbol;Acc:MGI:1913299] | 643,4 | 91,33 |
| ENSMUSG00000020744 | Slc25a19 | carrier family 25 (mitochondrial thiamine pyrophosphate carrier), member 19 [Source:MGI Symbol;Acc:MGI:19 | 643,3 | 90,01 |
| ENSMUSG00000022456 | Sept3 | septin 3 [Source:MGI Symbol;Acc:MGI:1345148] | 642,6 | 97,84 |
| ENSMUSG00000035521 | Gnptg | N-acetylglucosamine-1-phosphotransferase, gamma subunit [Source:MGI Symbol;Acc:MGI:2147006] | 642,6 | 70,89 |
| ENSMUSG00000039542 | Ncam1 | neural cell adhesion molecule 1 [Source:MGI Symbol;Acc:MGI:97281] | 642,0 | 76,36 |
| ENSMUSG00000052889 | Prkcb | protein kinase C, beta [Source:MGI Symbol;Acc:MGI:97596] | 642,0 | 99,90 |
| ENSMUSG00000043323 | Fbrsl1 | fibrosin-like 1 [Source:MGI Symbol;Acc:MGI:1920907] | 641,7 | 97,86 |
| ENSMUSG00000021087 | Rtn1 | reticulon 1 [Source:MGI Symbol;Acc:MGI:1933947] | 641,6 | 96,98 |
| ENSMUSG00000037253 | Mex3c | mex3 homolog C (C. elegans) [Source:MGI Symbol;Acc:MGI:2652843] | 640,5 | 98,61 |
| ENSMUSG00000024083 | Pja2 | praja 2, RING-H2 motif containing [Source:MGI Symbol;Acc:MGI:2159342] | 640,4 | 98,99 |
| ENSMUSG00000059689 | Zfp637 | zinc finger protein 637 [Source:MGI Symbol;Acc:MGI:2448537] | 640,3 | 96,40 |
| ENSMUSG00000020590 | Snx13 | sorting nexin 13 [Source:MGI Symbol;Acc:MGI:2661416] | 639,9 | 84,07 |
| ENSMUSG00000049299 | Trappc1 | trafficking protein particle complex 1 [Source:MGI Symbol;Acc:MGI:1098727] | 639,8 | 100,00 |
| ENSMUSG00000034361 | Cpne2 | copine II [Source:MGI Symbol;Acc:MGI:2387578] | 639,5 | 99,36 |
| ENSMUSG00000025786 | Zdhhc3 | zinc finger, DHHC domain containing 3 [Source:MGI Symbol;Acc:MGI:1926134] | 639,2 | 90,89 |
| ENSMUSG00000029209 | Gnpda2 | glucosamine-6-phosphate deaminase 2 [Source:MGI Symbol;Acc:MGI:1915230] | 638,6 | 71,13 |
| ENSMUSG00000002550 | Uck1 | uridine-cytidine kinase 1 [Source:MGI Symbol;Acc:MGI:98904] | 638,2 | 74,75 |
| ENSMUSG00000031840 | Rab3a | RAB3A, member RAS oncogene family [Source:MGI Symbol;Acc:MGI:97843] | 637,9 | 83,20 |
| ENSMUSG00000040138 | Ndp | Norrie disease (pseudoglioma) (human) [Source:MGI Symbol;Acc:MGI:102570] | 637,9 | 99,78 |
| ENSMUSG00000032557 | Uba5 | ubiquitin-like modifier activating enzyme 5 [Source:MGI Symbol;Acc:MGI:1913913] | 637,8 | 99,94 |
| ENSMUSG00000041895 | Wipi1 | WD repeat domain, phosphoinositide interacting 1 [Source:MGI Symbol;Acc:MGI:1261864] | 637,2 | 87,93 |
| ENSMUSG00000036398 | Ppp1r11 | protein phosphatase 1, regulatory (inhibitor) subunit 11 [Source:MGI Symbol;Acc:MGI:1923747] | 636,6 | 81,86 |
| ENSMUSG00000071662 | Polr2g | polymerase (RNA) II (DNA directed) polypeptide G [Source:MGI Symbol;Acc:MGI:1914960] | 636,4 | 100,00 |
| ENSMUSG00000026229 | Psmd1 | proteasome (prosome, macropain) 26S subunit, non-ATPase, 1 [Source:MGI Symbol;Acc:MGI:1917497] | 636,4 | 99,64 |
| ENSMUSG00000091512 | Lamtor3 | late endosomal/lysosomal adaptor, MAPK and MTOR activator 3 [Source:MGI Symbol;Acc:MGI:1929467] | 636,2 | 97,40 |
| ENSMUSG00000062753 | AI413582 | expressed sequence AI413582 [Source:MGI Symbol;Acc:MGI:2146839] | 635,8 | 76,71 |
| ENSMUSG00000022265 | Ank | progressive ankylosis [Source:MGI Symbol;Acc:MGI:3045421] | 635,6 | 99,64 |
| ENSMUSG00000020376 | Rnf130 | ring finger protein 130 [Source:MGI Symbol;Acc:MGI:1891717] | 635,2 | 99,69 |
| ENSMUSG00000042745 | Id1 | inhibitor of DNA binding 1 [Source:MGI Symbol;Acc:MGI:96396] | 635,1 | 81,01 |
| ENSMUSG00000048000 | Gigyf2 | GRB10 interacting GYF protein 2 [Source:MGI Symbol;Acc:MGI:2138584] | 635,1 | 85,87 |
| ENSMUSG00000001025 | S100a6 | S100 calcium binding protein A6 (calcyclin) [Source:MGI Symbol;Acc:MGI:1339467] | 635,0 | 90,36 |
| ENSMUSG00000043259 | Fam13c | family with sequence similarity 13, member C [Source:MGI Symbol;Acc:MGI:1918971] | 634,4 | 99,56 |
| ENSMUSG00000031848 | Lsm4 | LSM4 homolog, U6 small nuclear RNA associated (S. cerevisiae) [Source:MGI Symbol;Acc:MGI:1354692] | 633,8 | 99,89 |
| ENSMUSG00000045767 | B230219D22Rik | RIKEN cDNA B230219D22 gene [Source:MGI Symbol;Acc:MGI:1925771] | 633,4 | 99,79 |
| ENSMUSG00000039323 | Igfbp2 | insulin-like growth factor binding protein 2 [Source:MGI Symbol;Acc:MGI:96437] | 633,2 | 96,64 |
| ENSMUSG00000025220 | Mgea5 | meningioma expressed antigen 5 (hyaluronidase) [Source:MGI Symbol;Acc:MGI:1932139] | 633,1 | 99,31 |
| ENSMUSG00000022892 | App | amyloid beta (A4) precursor protein [Source:MGI Symbol;Acc:MGI:88059] | 633,0 | 100,00 |
| ENSMUSG00000025208 | Mrpl43 | mitochondrial ribosomal protein L43 [Source:MGI Symbol;Acc:MGI:2137229] | 632,6 | 100,00 |
| ENSMUSG00000029686 | Cul1 | cullin 1 [Source:MGI Symbol;Acc:MGI:1349658] | 632,5 | 98,09 |
| ENSMUSG00000020359 | Phykpl | 5-phosphohydroxy-L-lysine phospholyase [Source:MGI Symbol;Acc:MGI:1920197] | 632,3 | 76,80 |
| ENSMUSG00000069744 | Psmb3 | proteasome (prosome, macropain) subunit, beta type 3 [Source:MGI Symbol;Acc:MGI:1347014] | 631,9 | 98,09 |
| ENSMUSG00000024431 | Nr3c1 | nuclear receptor subfamily 3, group C, member 1 [Source:MGI Symbol;Acc:MGI:95824] | 631,6 | 93,81 |
| ENSMUSG00000039263 | Npepl1 | aminopeptidase-like 1 [Source:MGI Symbol;Acc:MGI:2448523] | 631,1 | 84,84 |
| ENSMUSG00000047368 | Abhd17b | abhydrolase domain containing 17B [Source:MGI Symbol;Acc:MGI:1917816] | 630,2 | 99,95 |
| ENSMUSG00000018377 | Vezf1 | vascular endothelial zinc finger 1 [Source:MGI Symbol;Acc:MGI:1313291] | 630,2 | 99,76 |
| ENSMUSG00000020936 | Nmt1 | N-myristoyltransferase 1 [Source:MGI Symbol;Acc:MGI:102579] | 629,9 | 97,52 |
| ENSMUSG00000022469 | Rapgef3 | Rap guanine nucleotide exchange factor (GEF) 3 [Source:MGI Symbol;Acc:MGI:2441741] | 629,8 | 95,18 |
| ENSMUSG00000031600 | Vps37a | vacuolar protein sorting 37A (yeast) [Source:MGI Symbol;Acc:MGI:1261835] | 629,8 | 99,42 |
| ENSMUSG00000007338 | Mrpl49 | mitochondrial ribosomal protein L49 [Source:MGI Symbol;Acc:MGI:108180] | 629,6 | 100,00 |

| | | | | |
|---|---|---|---|---|
| ENSMUSG00000027589 | Pcmtd2 | ein-L-isoaspartate (D-aspartate) O-methyltransferase domain containing 2 [Source:MGI Symbol;Acc:MGI:1923 | 629,4 | 91,15 |
| ENSMUSG00000078941 | Ak6 | adenylate kinase 6 [Source:MGI Symbol;Acc:MGI:5510732] | 629,4 | 71,41 |
| ENSMUSG00000020571 | Pdia6 | protein disulfide isomerase associated 6 [Source:MGI Symbol;Acc:MGI:1919103] | 629,0 | 99,20 |
| ENSMUSG00000031980 | Agt | angiotensinogen (serpin peptidase inhibitor, clade A, member 8) [Source:MGI Symbol;Acc:MGI:87963] | 628,4 | 97,21 |
| ENSMUSG00000030774 | Pak1 | p21 protein (Cdc42/Rac)-activated kinase 1 [Source:MGI Symbol;Acc:MGI:1339975] | 628,2 | 94,44 |
| ENSMUSG00000038582 | Pptc7 | PTC7 protein phosphatase homolog (S. cerevisiae) [Source:MGI Symbol;Acc:MGI:2444593] | 628,1 | 98,19 |
| ENSMUSG00000027502 | Rtfdc1 | replication termination factor 2 domain containing 1 [Source:MGI Symbol;Acc:MGI:1913654] | 627,9 | 99,60 |
| ENSMUSG00000006050 | Sra1 | steroid receptor RNA activator 1 [Source:MGI Symbol;Acc:MGI:1344414] | 627,8 | 81,12 |
| ENSMUSG00000027160 | Ccdc34 | coiled-coil domain containing 34 [Source:MGI Symbol;Acc:MGI:1915451] | 626,9 | 94,36 |
| ENSMUSG00000027775 | Mfsd1 | major facilitator superfamily domain containing 1 [Source:MGI Symbol;Acc:MGI:1914118] | 626,8 | 99,07 |
| ENSMUSG00000022722 | Arl6 | ADP-ribosylation factor-like 6 [Source:MGI Symbol;Acc:MGI:1927136] | 626,7 | 88,78 |
| ENSMUSG00000012405 | Rpl15 | ribosomal protein L15 [Source:MGI Symbol;Acc:MGI:1913730] | 626,6 | 81,60 |
| ENSMUSG00000038014 | Fam120a | family with sequence similarity 120, member A [Source:MGI Symbol;Acc:MGI:2446163] | 626,2 | 97,49 |
| ENSMUSG00000020869 | Lrrc59 | leucine rich repeat containing 59 [Source:MGI Symbol;Acc:MGI:2138133] | 625,9 | 100,00 |
| ENSMUSG00000020133 | 2310011J03Rik | RIKEN cDNA 2310011J03 gene [Source:MGI Symbol;Acc:MGI:1913624] | 625,7 | 99,92 |
| ENSMUSG00000102627 | Snurf | SNRPN upstream reading frame [Source:MGI Symbol;Acc:MGI:1891236] | 625,1 | 100,00 |
| ENSMUSG00000028757 | Ddost | dolichyl-di-phosphooligosaccharide-protein glycotransferase [Source:MGI Symbol;Acc:MGI:1194508] | 625,0 | 98,24 |
| ENSMUSG00000024516 | Sec11c | SEC11 homolog C, signal peptidase complex subunit [Source:MGI Symbol;Acc:MGI:1913536] | 624,3 | 98,15 |
| ENSMUSG00000044147 | Arf6 | ADP-ribosylation factor 6 [Source:MGI Symbol;Acc:MGI:99435] | 624,3 | 89,14 |
| ENSMUSG00000019857 | Asf1a | anti-silencing function 1A histone chaperone [Source:MGI Symbol;Acc:MGI:1913653] | 624,2 | 99,97 |
| ENSMUSG00000020460 | Rps27a | ribosomal protein S27A [Source:MGI Symbol;Acc:MGI:1925544] | 623,7 | 72,54 |
| ENSMUSG00000020111 | Micu1 | mitochondrial calcium uptake 1 [Source:MGI Symbol;Acc:MGI:2384909] | 623,6 | 81,15 |
| ENSMUSG00000058325 | Dock1 | dedicator of cytokinesis 1 [Source:MGI Symbol;Acc:MGI:2429765] | 623,5 | 99,99 |
| ENSMUSG00000034168 | Irf2bpl | interferon regulatory factor 2 binding protein-like [Source:MGI Symbol;Acc:MGI:2442463] | 623,2 | 98,37 |
| ENSMUSG00000000838 | Fmr1 | fragile X mental retardation syndrome 1 [Source:MGI Symbol;Acc:MGI:95564] | 623,1 | 98,27 |
| ENSMUSG00000042747 | Krtcap2 | keratinocyte associated protein 2 [Source:MGI Symbol;Acc:MGI:1913309] | 623,0 | 74,64 |
| ENSMUSG00000031987 | Egln1 | egl-9 family hypoxia-inducible factor 1 [Source:MGI Symbol;Acc:MGI:1932286] | 622,1 | 98,36 |
| ENSMUSG00000059482 | 2610301B20Rik | RIKEN cDNA 2610301B20 gene [Source:MGI Symbol;Acc:MGI:1914407] | 621,7 | 99,37 |
| ENSMUSG00000096740 | Lbhd1 | LBH domain containing 1 [Source:MGI Symbol;Acc:MGI:5516029] | 621,3 | 75,55 |
| ENSMUSG00000021807 | 2700060E02Rik | RIKEN cDNA 2700060E02 gene [Source:MGI Symbol;Acc:MGI:1915295] | 621,2 | 90,73 |
| ENSMUSG00000068798 | Rap1a | RAS-related protein-1a [Source:MGI Symbol;Acc:MGI:97852] | 620,3 | 88,28 |
| ENSMUSG00000067713 | Prkag1 | protein kinase, AMP-activated, gamma 1 non-catalytic subunit [Source:MGI Symbol;Acc:MGI:108411] | 620,1 | 99,84 |
| ENSMUSG00000025423 | Pias2 | protein inhibitor of activated STAT 2 [Source:MGI Symbol;Acc:MGI:1096566] | 619,6 | 98,09 |
| ENSMUSG00000025264 | Tsr2 | TSR2 20S rRNA accumulation [Source:MGI Symbol;Acc:MGI:1916749] | 619,6 | 99,08 |
| ENSMUSG00000034902 | Pip5k1c | phosphatidylinositol-4-phosphate 5-kinase, type 1 gamma [Source:MGI Symbol;Acc:MGI:1298224] | 619,5 | 98,62 |
| ENSMUSG00000027605 | Acss2 | acyl-CoA synthetase short-chain family member 2 [Source:MGI Symbol;Acc:MGI:1890410] | 618,6 | 75,99 |
| ENSMUSG00000022706 | Mrpl40 | mitochondrial ribosomal protein L40 [Source:MGI Symbol;Acc:MGI:1332635] | 617,5 | 96,27 |
| ENSMUSG00000008892 | Vdac3 | voltage-dependent anion channel 3 [Source:MGI Symbol;Acc:MGI:106922] | 617,0 | 80,56 |
| ENSMUSG00000025766 | D3Ertd751e | DNA segment, Chr 3, ERATO Doi 751, expressed [Source:MGI Symbol;Acc:MGI:1289213] | 616,3 | 82,24 |
| ENSMUSG00000031352 | Hccs | holocytochrome c synthetase [Source:MGI Symbol;Acc:MGI:106911] | 616,3 | 99,88 |
| ENSMUSG00000021709 | Erbb2ip | Erbb2 interacting protein [Source:MGI Symbol;Acc:MGI:1890169] | 616,2 | 99,80 |
| ENSMUSG00000029684 | Wasl | Wiskott-Aldrich syndrome-like (human) [Source:MGI Symbol;Acc:MGI:1920428] | 616,0 | 87,15 |
| ENSMUSG00000023960 | Enpp5 | ectonucleotide pyrophosphatase/phosphodiesterase 5 [Source:MGI Symbol;Acc:MGI:1933830] | 615,6 | 99,70 |
| ENSMUSG00000074656 | Eif2s2 | eukaryotic translation initiation factor 2, subunit 2 (beta) [Source:MGI Symbol;Acc:MGI:1914454] | 615,5 | 83,61 |
| ENSMUSG00000059820 | AU019823 | expressed sequence AU019823 [Source:MGI Symbol;Acc:MGI:2143205] | 615,4 | 71,35 |
| ENSMUSG00000054302 | Eapp | E2F-associated phosphoprotein [Source:MGI Symbol;Acc:MGI:1913516] | 615,1 | 70,73 |
| ENSMUSG00000021905 | Dph3 | diphthamine biosynthesis 3 [Source:MGI Symbol;Acc:MGI:1922658] | 614,6 | 93,36 |
| ENSMUSG00000024258 | Polr2d | polymerase (RNA) II (DNA directed) polypeptide D [Source:MGI Symbol;Acc:MGI:1916491] | 614,6 | 100,00 |
| ENSMUSG00000020486 | Sept4 | septin 4 [Source:MGI Symbol;Acc:MGI:1270156] | 614,6 | 92,60 |
| ENSMUSG00000030126 | Tmcc1 | transmembrane and coiled coil domains 1 [Source:MGI Symbol;Acc:MGI:2442368] | 614,0 | 86,28 |
| ENSMUSG00000079478 | Sssca1 | Sjogren's syndrome/scleroderma autoantigen 1 homolog (human) [Source:MGI Symbol;Acc:MGI:1913482] | 613,9 | 92,57 |
| ENSMUSG00000054766 | Set | SET nuclear oncogene [Source:MGI Symbol;Acc:MGI:1860267] | 613,8 | 85,16 |
| ENSMUSG00000041645 | Ddx24 | DEAD (Asp-Glu-Ala-Asp) box polypeptide 24 [Source:MGI Symbol;Acc:MGI:1351337] | 613,7 | 97,82 |
| ENSMUSG00000017286 | Glod4 | glyoxalase domain containing 4 [Source:MGI Symbol;Acc:MGI:1914451] | 613,4 | 85,06 |
| ENSMUSG00000032299 | Commd4 | COMM domain containing 4 [Source:MGI Symbol;Acc:MGI:1913449] | 613,3 | 100,00 |
| ENSMUSG00000032563 | Mrpl3 | mitochondrial ribosomal protein L3 [Source:MGI Symbol;Acc:MGI:2137204] | 613,1 | 91,78 |
| ENSMUSG00000001105 | Ift20 | intraflagellar transport 20 [Source:MGI Symbol;Acc:MGI:1915585] | 612,7 | 99,05 |
| ENSMUSG00000006442 | Srm | spermidine synthase [Source:MGI Symbol;Acc:MGI:102690] | 612,7 | 98,60 |
| ENSMUSG00000020362 | Cnot6 | CCR4-NOT transcription complex, subunit 6 [Source:MGI Symbol;Acc:MGI:2144529] | 612,7 | 98,61 |
| ENSMUSG00000099881 | 2810013P06Rik | RIKEN cDNA 2810013P06 gene [Source:MGI Symbol;Acc:MGI:1914456] | 612,5 | 99,56 |
| ENSMUSG00000024294 | Mib1 | mindbomb homolog 1 (Drosophila) [Source:MGI Symbol;Acc:MGI:2443157] | 612,4 | 95,92 |
| ENSMUSG00000020515 | Cnot8 | CCR4-NOT transcription complex, subunit 8 [Source:MGI Symbol;Acc:MGI:1916375] | 612,2 | 95,58 |
| ENSMUSG00000032867 | Fbxw8 | F-box and WD-40 domain protein 8 [Source:MGI Symbol;Acc:MGI:1923041] | 611,5 | 93,47 |
| ENSMUSG00000053483 | Usp21 | ubiquitin specific peptidase 21 [Source:MGI Symbol;Acc:MGI:1353665] | 611,3 | 93,12 |
| ENSMUSG00000000295 | Hddc2 | HD domain containing 2 [Source:MGI Symbol;Acc:MGI:1916942] | 611,2 | 100,00 |
| ENSMUSG00000090000 | Ier3ip1 | immediate early response 3 interacting protein 1 [Source:MGI Symbol;Acc:MGI:1913441] | 611,1 | 99,88 |
| ENSMUSG00000048240 | Gng7 | guanine nucleotide binding protein (G protein), gamma 7 [Source:MGI Symbol;Acc:MGI:95787] | 611,0 | 95,86 |
| ENSMUSG00000023572 | Ccndbp1 | cyclin D-type binding-protein 1 [Source:MGI Symbol;Acc:MGI:109595] | 610,9 | 93,52 |
| ENSMUSG00000024875 | Yif1a | Yip1 interacting factor homolog A (S. cerevisiae) [Source:MGI Symbol;Acc:MGI:1915340] | 610,7 | 100,00 |
| ENSMUSG00000036352 | Ubac1 | ubiquitin associated domain containing 1 [Source:MGI Symbol;Acc:MGI:1920995] | 610,3 | 100,00 |
| ENSMUSG00000066568 | Lsm14a | LSM14 homolog A (SCD6, S. cerevisiae) [Source:MGI Symbol;Acc:MGI:1914320] | 610,1 | 93,22 |
| ENSMUSG00000023495 | Pcbp4 | poly(rC) binding protein 4 [Source:MGI Symbol;Acc:MGI:1890471] | 609,6 | 78,68 |
| ENSMUSG00000035770 | Dync1li2 | dynein, cytoplasmic 1 light intermediate chain 2 [Source:MGI Symbol;Acc:MGI:107738] | 608,9 | 97,68 |
| ENSMUSG00000037257 | Aagab | alpha- and gamma-adaptin binding protein [Source:MGI Symbol;Acc:MGI:1914189] | 608,9 | 100,00 |
| ENSMUSG00000029148 | Nrbp1 | nuclear receptor binding protein 1 [Source:MGI Symbol;Acc:MGI:2183436] | 608,8 | 99,51 |
| ENSMUSG00000037514 | Pank2 | pantothenate kinase 2 [Source:MGI Symbol;Acc:MGI:1921700] | 608,6 | 99,93 |
| ENSMUSG00000025743 | Sdc3 | syndecan 3 [Source:MGI Symbol;Acc:MGI:1349163] | 608,5 | 97,43 |
| ENSMUSG00000066150 | Slc31a1 | solute carrier family 31, member 1 [Source:MGI Symbol;Acc:MGI:1333843] | 608,3 | 87,30 |
| ENSMUSG00000028291 | Akirin2 | akirin 2 [Source:MGI Symbol;Acc:MGI:1889364] | 608,2 | 100,00 |
| ENSMUSG00000036860 | Mrpl55 | mitochondrial ribosomal protein L55 [Source:MGI Symbol;Acc:MGI:1914462] | 608,2 | 99,19 |
| ENSMUSG00000093574 | Gm20671 | predicted gene 20671 [Source:MGI Symbol;Acc:MGI:5313118] | 607,9 | 80,53 |
| ENSMUSG00000031534 | Smim19 | small integral membrane protein 19 [Source:MGI Symbol;Acc:MGI:2142501] | 607,5 | 95,29 |
| ENSMUSG00000041353 | Tmem29 | transmembrane protein 29 [Source:MGI Symbol;Acc:MGI:1923420] | 607,4 | 73,18 |
| ENSMUSG00000071652 | Ints5 | integrator complex subunit 5 [Source:MGI Symbol;Acc:MGI:1923578] | 607,3 | 96,25 |
| ENSMUSG00000070319 | Eif3g | eukaryotic translation initiation factor 3, subunit G [Source:MGI Symbol;Acc:MGI:1858258] | 607,1 | 100,00 |
| ENSMUSG00000004070 | Hmox2 | heme oxygenase 2 [Source:MGI Symbol;Acc:MGI:109373] | 606,9 | 85,08 |
| ENSMUSG00000023094 | Msrb2 | methionine sulfoxide reductase B2 [Source:MGI Symbol;Acc:MGI:1923717] | 606,9 | 100,00 |
| ENSMUSG00000020180 | Snrpd3 | small nuclear ribonucleoprotein D3 [Source:MGI Symbol;Acc:MGI:1914582] | 606,8 | 90,92 |
| ENSMUSG00000031568 | Rwdd4a | RWD domain containing 4A [Source:MGI Symbol;Acc:MGI:2681000] | 606,7 | 99,94 |
| ENSMUSG00000035781 | R3hdm4 | R3H domain containing 4 [Source:MGI Symbol;Acc:MGI:1924814] | 606,6 | 98,63 |
| ENSMUSG00000005846 | Rsl1d1 | ribosomal L1 domain containing 1 [Source:MGI Symbol;Acc:MGI:1913659] | 606,5 | 98,59 |
| ENSMUSG00000048450 | Msx1 | msh homeobox 1 [Source:MGI Symbol;Acc:MGI:97168] | 605,8 | 99,14 |
| ENSMUSG00000030982 | 9030624J02Rik | RIKEN cDNA 9030624J02 gene [Source:MGI Symbol;Acc:MGI:1918767] | 605,8 | 79,92 |
| ENSMUSG00000032615 | Nt5m | 5',3'-nucleotidase, mitochondrial [Source:MGI Symbol;Acc:MGI:1917127] | 605,7 | 100,00 |
| ENSMUSG00000052962 | Mrpl35 | mitochondrial ribosomal protein L35 [Source:MGI Symbol;Acc:MGI:1913473] | 605,6 | 96,62 |
| ENSMUSG00000007987 | Ift22 | intraflagellar transport 22 [Source:MGI Symbol;Acc:MGI:1914536] | 605,1 | 98,30 |
| ENSMUSG00000061650 | Med9 | mediator complex subunit 9 [Source:MGI Symbol;Acc:MGI:2183151] | 604,9 | 99,83 |
| ENSMUSG00000042508 | Dmtf1 | cyclin D binding myb-like transcription factor 1 [Source:MGI Symbol;Acc:MGI:1344415] | 603,2 | 95,10 |
| ENSMUSG00000056851 | Pcbp2 | poly(rC) binding protein 2 [Source:MGI Symbol;Acc:MGI:108202] | 602,7 | 95,23 |
| ENSMUSG00000028159 | Dapp1 | dual adaptor for phosphotyrosine and 3-phosphoinositides 1 [Source:MGI Symbol;Acc:MGI:1347063] | 602,7 | 99,68 |
| ENSMUSG00000030058 | Copg1 | coatomer protein complex, subunit gamma 1 [Source:MGI Symbol;Acc:MGI:1858696] | 602,5 | 98,50 |
| ENSMUSG00000021377 | Dek | DEK oncogene (DNA binding) [Source:MGI Symbol;Acc:MGI:1926209] | 601,5 | 97,01 |
| ENSMUSG00000061787 | Rps17 | ribosomal protein S17 [Source:MGI Symbol;Acc:MGI:1309526] | 601,5 | 96,29 |
| ENSMUSG00000046818 | Ddit4l | DNA-damage-inducible transcript 4-like [Source:MGI Symbol;Acc:MGI:1920534] | 601,4 | 97,57 |
| ENSMUSG00000064036 | Mro | maestro [Source:MGI Symbol;Acc:MGI:2152817] | 601,2 | 98,82 |
| ENSMUSG00000024941 | Scyl1 | SCY1-like 1 (S. cerevisiae) [Source:MGI Symbol;Acc:MGI:1931787] | 601,1 | 100,00 |
| ENSMUSG00000031320 | Rps4x | ribosomal protein S4, X-linked [Source:MGI Symbol;Acc:MGI:98158] | 601,0 | 91,73 |
| ENSMUSG00000022191 | Drosha | drosha, ribonuclease type III [Source:MGI Symbol;Acc:MGI:1261425] | 600,8 | 98,58 |
| ENSMUSG00000083282 | Ctsf | cathepsin F [Source:MGI Symbol;Acc:MGI:1861434] | 600,8 | 99,93 |
| ENSMUSG00000047205 | Dusp18 | dual specificity phosphatase 18 [Source:MGI Symbol;Acc:MGI:1922469] | 600,2 | 99,86 |
| ENSMUSG00000004394 | Tmed4 | transmembrane emp24 protein transport domain containing 4 [Source:MGI Symbol;Acc:MGI:1915070] | 600,1 | 98,40 |
| ENSMUSG00000041120 | Nbl1 | neuroblastoma, suppression of tumorigenicity 1 [Source:MGI Symbol;Acc:MGI:104591] | 599,9 | 99,70 |
| ENSMUSG00000038160 | Atg5 | autophagy related 5 [Source:MGI Symbol;Acc:MGI:1277186] | 599,8 | 99,74 |
| ENSMUSG00000037736 | Limch1 | LIM and calponin homology domains 1 [Source:MGI Symbol;Acc:MGI:1924819] | 599,7 | 95,05 |
| ENSMUSG00000025738 | Fbxl16 | F-box and leucine-rich repeat protein 16 [Source:MGI Symbol;Acc:MGI:2448488] | 599,5 | 99,84 |
| ENSMUSG00000078765 | U2af1l4 | U2 small nuclear RNA auxiliary factor 1-like 4 [Source:MGI Symbol;Acc:MGI:2678374] | 599,3 | 99,96 |
| ENSMUSG00000024096 | Ralbp1 | ralA binding protein 1 [Source:MGI Symbol;Acc:MGI:108466] | 598,8 | 94,78 |

| | | | | |
|---|---|---|---|---|
| ENSMUSG00000053012 | Krcc1 | lysine-rich coiled-coil 1 [Source:MGI Symbol;Acc:MGI:1889377] | 598,4 | 88,05 |
| ENSMUSG00000024335 | Brd2 | bromodomain containing 2 [Source:MGI Symbol;Acc:MGI:99495] | 598,3 | 75,42 |
| ENSMUSG00000048578 | Mlec | malectin [Source:MGI Symbol;Acc:MGI:1924015] | 597,8 | 99,84 |
| ENSMUSG00000030615 | Tmem126a | transmembrane protein 126A [Source:MGI Symbol;Acc:MGI:1913521] | 597,7 | 98,66 |
| ENSMUSG00000028524 | Sgip1 | SH3-domain GRB2-like (endophilin) interacting protein 1 [Source:MGI Symbol;Acc:MGI:1920344] | 597,6 | 86,95 |
| ENSMUSG00000001467 | Cyp51 | cytochrome P450, family 51 [Source:MGI Symbol;Acc:MGI:106040] | 596,7 | 98,37 |
| ENSMUSG00000031622 | Sin3b | transcriptional regulator, SIN3B (yeast) [Source:MGI Symbol;Acc:MGI:107158] | 596,6 | 96,77 |
| ENSMUSG00000054894 | Atp5s | ATP synthase, H+ transporting, mitochondrial F0 complex, subunit S [Source:MGI Symbol;Acc:MGI:1915305 | 596,0 | 100,00 |
| ENSMUSG00000024081 | Cebpz | CCAAT/enhancer binding protein zeta [Source:MGI Symbol;Acc:MGI:109386] | 595,8 | 91,72 |
| ENSMUSG00000041203 | 2310036O22Rik | RIKEN cDNA 2310036O22 gene [Source:MGI Symbol;Acc:MGI:1922833] | 595,5 | 100,00 |
| ENSMUSG00000059456 | Ptk2b | PTK2 protein tyrosine kinase 2 beta [Source:MGI Symbol;Acc:MGI:104908] | 595,3 | 98,93 |
| ENSMUSG00000030733 | Sh2b1 | SH2B adaptor protein 1 [Source:MGI Symbol;Acc:MGI:1201407] | 595,2 | 93,25 |
| ENSMUSG00000000202 | Btbd17 | BTB (POZ) domain containing 17 [Source:MGI Symbol;Acc:MGI:1919264] | 595,1 | 99,94 |
| ENSMUSG00000043131 | Mob1a | MOB kinase activator 1A [Source:MGI Symbol;Acc:MGI:2442631] | 594,8 | 98,10 |
| ENSMUSG00000039218 | Srrm2 | serine/arginine repetitive matrix 2 [Source:MGI Symbol;Acc:MGI:1923206] | 594,3 | 93,65 |
| ENSMUSG00000036748 | Cuedc2 | CUE domain containing 2 [Source:MGI Symbol;Acc:MGI:1914366] | 594,2 | 81,44 |
| ENSMUSG00000062604 | Srpk2 | serine/arginine-rich protein specific kinase 2 [Source:MGI Symbol;Acc:MGI:1201408] | 594,1 | 88,96 |
| ENSMUSG00000037531 | Mrpl47 | mitochondrial ribosomal protein L47 [Source:MGI Symbol;Acc:MGI:1921850] | 593,6 | 86,44 |
| ENSMUSG00000037197 | Rbm17 | RNA binding motif protein 17 [Source:MGI Symbol;Acc:MGI:1924188] | 593,6 | 88,67 |
| ENSMUSG00000091896 | Ube2d2a | ubiquitin-conjugating enzyme E2D 2A [Source:MGI Symbol;Acc:MGI:1930715] | 593,6 | 71,20 |
| ENSMUSG00000026728 | Vim | vimentin [Source:MGI Symbol;Acc:MGI:98932] | 593,5 | 87,92 |
| ENSMUSG00000029126 | Nsg1 | neuron specific gene family member 1 [Source:MGI Symbol;Acc:MGI:109149] | 593,4 | 86,05 |
| ENSMUSG00000040725 | Hnrnpul1 | heterogeneous nuclear ribonucleoprotein U-like 1 [Source:MGI Symbol;Acc:MGI:2443517] | 593,4 | 96,24 |
| ENSMUSG00000027339 | Rassf2 | Ras association (RalGDS/AF-6) domain family member 2 [Source:MGI Symbol;Acc:MGI:2442060] | 593,4 | 75,17 |
| ENSMUSG00000035790 | Cep19 | centrosomal protein 19 [Source:MGI Symbol;Acc:MGI:1914244] | 593,2 | 98,10 |
| ENSMUSG00000038416 | Cdc16 | CDC16 cell division cycle 16 [Source:MGI Symbol;Acc:MGI:1917207] | 593,0 | 99,86 |
| ENSMUSG00000031157 | Pqbp1 | polyglutamine binding protein 1 [Source:MGI Symbol;Acc:MGI:1859638] | 592,4 | 85,66 |
| ENSMUSG00000019920 | Lims1 | LIM and senescent cell antigen-like domains 1 [Source:MGI Symbol;Acc:MGI:1195263] | 592,0 | 93,16 |
| ENSMUSG00000018672 | Copz2 | coatomer protein complex, subunit zeta 2 [Source:MGI Symbol;Acc:MGI:1929008] | 591,6 | 100,00 |
| ENSMUSG00000036775 | Decr2 | 2-4-dienoyl-Coenzyme A reductase 2, peroxisomal [Source:MGI Symbol;Acc:MGI:1347059] | 591,5 | 99,59 |
| ENSMUSG00000043687 | 1190005I06Rik | RIKEN cDNA 1190005I06 gene [Source:MGI Symbol;Acc:MGI:1916168] | 591,2 | 83,11 |
| ENSMUSG00000059291 | Rpl11 | ribosomal protein L11 [Source:MGI Symbol;Acc:MGI:1914275] | 591,0 | 92,14 |
| ENSMUSG00000052751 | Repin1 | replication initiator 1 [Source:MGI Symbol;Acc:MGI:1889817] | 590,6 | 99,63 |
| ENSMUSG00000028639 | Ybx1 | Y box protein 1 [Source:MGI Symbol;Acc:MGI:99146] | 590,0 | 93,25 |
| ENSMUSG00000004268 | Emg1 | EMG1 N1-specific pseudouridine methyltransferase [Source:MGI Symbol;Acc:MGI:1315195] | 589,8 | 98,36 |
| ENSMUSG00000026827 | Gpd2 | glycerol phosphate dehydrogenase 2, mitochondrial [Source:MGI Symbol;Acc:MGI:99778] | 589,6 | 96,73 |
| ENSMUSG00000003585 | Sec14l2 | SEC14-like lipid binding 2 [Source:MGI Symbol;Acc:MGI:1915065] | 589,5 | 98,75 |
| ENSMUSG00000058355 | Abce1 | ATP-binding cassette, sub-family E (OABP), member 1 [Source:MGI Symbol;Acc:MGI:1195458] | 589,4 | 95,84 |
| ENSMUSG00000060938 | Rpl26 | ribosomal protein L26 [Source:MGI Symbol;Acc:MGI:106022] | 589,0 | 87,82 |
| ENSMUSG00000007739 | Cct4 | chaperonin containing Tcp1, subunit 4 (delta) [Source:MGI Symbol;Acc:MGI:104689] | 588,9 | 95,53 |
| ENSMUSG00000058709 | Egln2 | egl-9 family hypoxia-inducible factor 2 [Source:MGI Symbol;Acc:MGI:1932287] | 588,8 | 98,00 |
| ENSMUSG00000031591 | Asah1 | N-acylsphingosine amidohydrolase 1 [Source:MGI Symbol;Acc:MGI:1277124] | 588,2 | 86,65 |
| ENSMUSG00000026739 | Bmi1 | Bmi1 polycomb ring finger oncogene [Source:MGI Symbol;Acc:MGI:88174] | 588,2 | 86,75 |
| ENSMUSG00000038650 | Rnh1 | ribonuclease/angiogenin inhibitor 1 [Source:MGI Symbol;Acc:MGI:1195456] | 587,9 | 96,34 |
| ENSMUSG00000027495 | Fam210b | family with sequence similarity 210, member B [Source:MGI Symbol;Acc:MGI:1914267] | 587,7 | 99,71 |
| ENSMUSG00000058351 | Smim4 | small integral membrane protein 4 [Source:MGI Symbol;Acc:MGI:1913737] | 587,6 | 99,88 |
| ENSMUSG00000028936 | Rpl22 | ribosomal protein L22 [Source:MGI Symbol;Acc:MGI:99262] | 587,6 | 99,95 |
| ENSMUSG00000005882 | Uqcc1 | ubiquinol-cytochrome c reductase complex assembly factor 1 [Source:MGI Symbol;Acc:MGI:1929472] | 587,3 | 91,25 |
| ENSMUSG00000060227 | Casc4 | cancer susceptibility candidate 4 [Source:MGI Symbol;Acc:MGI:2443129] | 587,2 | 96,40 |
| ENSMUSG00000022899 | Slc15a2 | solute carrier family 15 (H+/peptide transporter), member 2 [Source:MGI Symbol;Acc:MGI:1890457] | 587,2 | 98,88 |
| ENSMUSG00000021033 | Gstz1 | glutathione transferase zeta 1 (maleylacetoacetate isomerase) [Source:MGI Symbol;Acc:MGI:1341859] | 587,0 | 100,00 |
| ENSMUSG00000054426 | A930005H10Rik | RIKEN cDNA A930005H10 gene [Source:MGI Symbol;Acc:MGI:1915411] | 587,0 | 100,00 |
| ENSMUSG00000044550 | Tceal3 | transcription elongation factor A (SII)-like 3 [Source:MGI Symbol;Acc:MGI:1913354] | 586,8 | 79,21 |
| ENSMUSG00000045257 | Morn2 | MORN repeat containing 2 [Source:MGI Symbol;Acc:MGI:2674071] | 586,1 | 100,00 |
| ENSMUSG00000039887 | Alg14 | asparagine-linked glycosylation 14 [Source:MGI Symbol;Acc:MGI:1914039] | 585,8 | 84,55 |
| ENSMUSG00000028149 | Rap1gds1 | RAP1, GTP-GDP dissociation stimulator 1 [Source:MGI Symbol;Acc:MGI:2385189] | 585,4 | 81,31 |
| ENSMUSG00000039114 | Nrn1 | neuritin 1 [Source:MGI Symbol;Acc:MGI:1915654] | 584,8 | 93,13 |
| ENSMUSG00000045817 | Zfp36l2 | zinc finger protein 36, C3H type-like 2 [Source:MGI Symbol;Acc:MGI:107945] | 584,1 | 82,67 |
| ENSMUSG00000031660 | Brd7 | bromodomain containing 7 [Source:MGI Symbol;Acc:MGI:1349766] | 583,9 | 87,38 |
| ENSMUSG00000020372 | Gnb2l1 | uanine nucleotide binding protein (G protein), beta polypeptide 2 like 1 [Source:MGI Symbol;Acc:MGI:10184 | 583,7 | 87,70 |
| ENSMUSG00000040479 | Dgkz | diacylglycerol kinase zeta [Source:MGI Symbol;Acc:MGI:1278339] | 583,6 | 87,27 |
| ENSMUSG00000024902 | Mrpl11 | mitochondrial ribosomal protein L11 [Source:MGI Symbol;Acc:MGI:2137215] | 583,5 | 87,80 |
| ENSMUSG00000027854 | Sike1 | suppressor of IKBKE 1 [Source:MGI Symbol;Acc:MGI:1913891] | 582,8 | 78,05 |
| ENSMUSG00000029621 | Arpc1a | actin related protein 2/3 complex, subunit 1A [Source:MGI Symbol;Acc:MGI:1928896] | 582,8 | 91,43 |
| ENSMUSG00000040746 | Rnf167 | ring finger protein 167 [Source:MGI Symbol;Acc:MGI:1917760] | 582,5 | 99,98 |
| ENSMUSG00000024404 | Riok3 | RIO kinase 3 [Source:MGI Symbol;Acc:MGI:1914128] | 582,5 | 93,82 |
| ENSMUSG00000021497 | Txndc15 | thioredoxin domain containing 15 [Source:MGI Symbol;Acc:MGI:1916922] | 582,4 | 100,00 |
| ENSMUSG00000026851 | BC005624 | cDNA sequence BC005624 [Source:MGI Symbol;Acc:MGI:2385132] | 582,3 | 95,03 |
| ENSMUSG00000067787 | Blcap | bladder cancer associated protein [Source:MGI Symbol;Acc:MGI:1858907] | 581,9 | 92,21 |
| ENSMUSG00000047037 | Nipa1 | on imprinted in Prader-Willi/Angelman syndrome 1 homolog (human) [Source:MGI Symbol;Acc:MGI:244205 | 581,8 | 86,48 |
| ENSMUSG00000022781 | Pak2 | p21 protein (Cdc42/Rac)-activated kinase 2 [Source:MGI Symbol;Acc:MGI:1339984] | 581,7 | 98,48 |
| ENSMUSG00000020089 | Ppa1 | pyrophosphatase (inorganic) 1 [Source:MGI Symbol;Acc:MGI:97831] | 581,6 | 98,44 |
| ENSMUSG00000068220 | Lgals1 | lectin, galactose binding, soluble 1 [Source:MGI Symbol;Acc:MGI:96777] | 581,6 | 95,92 |
| ENSMUSG00000040822 | 1700123O20Rik | RIKEN cDNA 1700123O20 gene [Source:MGI Symbol;Acc:MGI:1920893] | 581,5 | 96,06 |
| ENSMUSG00000038370 | Pcp4l1 | Purkinje cell protein 4-like 1 [Source:MGI Symbol;Acc:MGI:1913675] | 581,5 | 99,34 |
| ENSMUSG00000056216 | Cebpg | CCAAT/enhancer binding protein (C/EBP), gamma [Source:MGI Symbol;Acc:MGI:104982] | 581,5 | 84,85 |
| ENSMUSG00000017176 | Nt5c3b | 5'-nucleotidase, cytosolic IIIB [Source:MGI Symbol;Acc:MGI:1915356] | 581,4 | 76,47 |
| ENSMUSG00000022905 | Kpna1 | karyopherin (importin) alpha 1 [Source:MGI Symbol;Acc:MGI:103560] | 581,2 | 98,88 |
| ENSMUSG00000021364 | Elovl2 | gation of very long chain fatty acids (FEN1/Elo2, SUR4/Elo3, yeast)-like 2 [Source:MGI Symbol;Acc:MGI:1858 | 581,2 | 96,73 |
| ENSMUSG00000028743 | Akr7a5 | ldo-keto reductase family 7, member A5 (aflatoxin aldehyde reductase) [Source:MGI Symbol;Acc:MGI:10779 | 580,6 | 99,92 |
| ENSMUSG00000026553 | Copa | coatomer protein complex subunit alpha [Source:MGI Symbol;Acc:MGI:1334462] | 580,4 | 98,62 |
| ENSMUSG00000029167 | Ppargc1a | eroxisome proliferative activated receptor, gamma, coactivator 1 alpha [Source:MGI Symbol;Acc:MGI:134277 | 580,3 | 98,29 |
| ENSMUSG00000022312 | Eif3h | eukaryotic translation initiation factor 3, subunit H [Source:MGI Symbol;Acc:MGI:1915385] | 580,3 | 93,99 |
| ENSMUSG00000043969 | Emx2 | empty spiracles homeobox 2 [Source:MGI Symbol;Acc:MGI:95388] | 579,7 | 97,60 |
| ENSMUSG00000020650 | Bcap29 | B cell receptor associated protein 29 [Source:MGI Symbol;Acc:MGI:101917] | 579,3 | 95,75 |
| ENSMUSG00000039703 | Nploc4 | nuclear protein localization 4 homolog (S. cerevisiae) [Source:MGI Symbol;Acc:MGI:2679787] | 579,2 | 98,55 |
| ENSMUSG00000017686 | Rhot1 | ras homolog gene family, member T1 [Source:MGI Symbol;Acc:MGI:1926078] | 579,1 | 87,45 |
| ENSMUSG00000037266 | Rsrp1 | arginine/serine rich protein 1 [Source:MGI Symbol;Acc:MGI:106498] | 579,0 | 97,83 |
| ENSMUSG00000047866 | Lonp2 | lon peptidase 2, peroxisomal [Source:MGI Symbol;Acc:MGI:1914137] | 578,9 | 97,11 |
| ENSMUSG00000014602 | Kif1a | kinesin family member 1A [Source:MGI Symbol;Acc:MGI:108391] | 578,7 | 94,54 |
| ENSMUSG00000033335 | Dnm2 | dynamin 2 [Source:MGI Symbol;Acc:MGI:109547] | 578,1 | 98,59 |
| ENSMUSG00000033373 | Fntb | farnesyltransferase, CAAX box, beta [Source:MGI Symbol;Acc:MGI:1861305] | 577,7 | 95,98 |
| ENSMUSG00000027030 | Stk39 | serine/threonine kinase 39 [Source:MGI Symbol;Acc:MGI:1858416] | 577,3 | 92,69 |
| ENSMUSG00000034024 | Cct2 | chaperonin containing Tcp1, subunit 2 (beta) [Source:MGI Symbol;Acc:MGI:107186] | 577,2 | 99,90 |
| ENSMUSG00000039166 | Akap7 | A kinase (PRKA) anchor protein 7 [Source:MGI Symbol;Acc:MGI:1859150] | 576,5 | 78,36 |
| ENSMUSG00000098950 | Gm28036 | predicted gene, 28036 [Source:MGI Symbol;Acc:MGI:5547772] | 576,3 | 100,00 |
| ENSMUSG00000028559 | Osbpl9 | oxysterol binding protein-like 9 [Source:MGI Symbol;Acc:MGI:1923784] | 576,2 | 87,44 |
| ENSMUSG00000052214 | Opa3 | optic atrophy 3 [Source:MGI Symbol;Acc:MGI:2686271] | 575,9 | 88,42 |
| ENSMUSG00000049969 | Plekhf2 | strin homology domain containing, family F (with FYVE domain) member 2 [Source:MGI Symbol;Acc:MGI:191 | 575,8 | 96,44 |
| ENSMUSG00000028576 | Ift74 | intraflagellar transport 74 [Source:MGI Symbol;Acc:MGI:1914944] | 575,5 | 82,16 |
| ENSMUSG00000027466 | Rbck1 | RanBP-type and C3HC4-type zinc finger containing 1 [Source:MGI Symbol;Acc:MGI:1344372] | 575,4 | 90,91 |
| ENSMUSG00000104208 | Gm37401 | predicted gene, 37401 [Source:MGI Symbol;Acc:MGI:5610629] | 575,3 | 89,53 |
| ENSMUSG00000003992 | Ssbp2 | single-stranded DNA binding protein 2 [Source:MGI Symbol;Acc:MGI:1914220] | 575,3 | 96,21 |
| ENSMUSG00000037509 | Arhgef4 | Rho guanine nucleotide exchange factor (GEF) 4 [Source:MGI Symbol;Acc:MGI:2442507] | 574,9 | 98,17 |
| ENSMUSG00000025487 | Psmd13 | proteasome (prosome, macropain) 26S subunit, non-ATPase, 13 [Source:MGI Symbol;Acc:MGI:1345192] | 574,7 | 82,29 |
| ENSMUSG00000057863 | Rpl36 | ribosomal protein L36 [Source:MGI Symbol;Acc:MGI:1860603] | 574,3 | 87,87 |
| ENSMUSG00000021550 | 2210016F16Rik | RIKEN cDNA 2210016F16 gene [Source:MGI Symbol;Acc:MGI:1917403] | 573,5 | 95,59 |
| ENSMUSG00000032601 | Prkar2a | protein kinase, cAMP dependent regulatory, type II alpha [Source:MGI Symbol;Acc:MGI:108025] | 573,5 | 95,75 |
| ENSMUSG00000021811 | Dnajc9 | DnaJ heat shock protein family (Hsp40) member C9 [Source:MGI Symbol;Acc:MGI:1915326] | 572,7 | 90,46 |
| ENSMUSG00000028102 | Pex11b | peroxisomal biogenesis factor 11 beta [Source:MGI Symbol;Acc:MGI:1338882] | 572,4 | 90,01 |
| ENSMUSG00000021910 | Nisch | nischarin [Source:MGI Symbol;Acc:MGI:1928323] | 572,3 | 72,18 |
| ENSMUSG00000042406 | Atf4 | activating transcription factor 4 [Source:MGI Symbol;Acc:MGI:88096] | 572,2 | 86,56 |
| ENSMUSG00000018340 | Anxa6 | annexin A6 [Source:MGI Symbol;Acc:MGI:88255] | 572,1 | 100,00 |
| ENSMUSG00000002058 | Unc119 | unc-119 homolog (C. elegans) [Source:MGI Symbol;Acc:MGI:1328357] | 571,7 | 93,37 |
| ENSMUSG00000025354 | Dnajc14 | DnaJ heat shock protein family (Hsp40) member C14 [Source:MGI Symbol;Acc:MGI:1921580] | 571,5 | 97,47 |
| ENSMUSG00000021037 | Ahsa1 | AHA1, activator of heat shock protein ATPase 1 [Source:MGI Symbol;Acc:MGI:2387603] | 571,3 | 94,93 |
| ENSMUSG00000024873 | Cnih2 | cornichon family AMPA receptor auxiliary protein 2 [Source:MGI Symbol;Acc:MGI:1277225] | 571,3 | 100,00 |

| | | | | |
|---|---|---|---|---|
| ENSMUSG00000012535 | Tnpo3 | transportin 3 [Source:MGI Symbol;Acc:MGI:1196412] | 571,2 | 97,54 |
| ENSMUSG00000058240 | Cryzl1 | crystallin, zeta (quinone reductase)-like 1 [Source:MGI Symbol;Acc:MGI:1913859] | 571,1 | 76,06 |
| ENSMUSG00000001525 | Tubb5 | tubulin, beta 5 class I [Source:MGI Symbol;Acc:MGI:107812] | 570,7 | 87,07 |
| ENSMUSG00000064370 | mt-Cytb | mitochondrially encoded cytochrome b [Source:MGI Symbol;Acc:MGI:102501] | 570,3 | 100,00 |
| ENSMUSG00000029201 | Ugdh | UDP-glucose dehydrogenase [Source:MGI Symbol;Acc:MGI:1306785] | 570,3 | 93,33 |
| ENSMUSG00000032648 | Pygm | muscle glycogen phosphorylase [Source:MGI Symbol;Acc:MGI:97830] | 569,9 | 99,59 |
| ENSMUSG00000015709 | Arnt2 | aryl hydrocarbon receptor nuclear translocator 2 [Source:MGI Symbol;Acc:MGI:107188] | 569,4 | 89,93 |
| ENSMUSG00000040722 | Scamp5 | secretory carrier membrane protein 5 [Source:MGI Symbol;Acc:MGI:1928948] | 569,4 | 99,94 |
| ENSMUSG00000047675 | Rps8 | ribosomal protein S8 [Source:MGI Symbol;Acc:MGI:98166] | 569,4 | 91,74 |
| ENSMUSG00000000355 | Mcts1 | malignant T cell amplified sequence 1 [Source:MGI Symbol;Acc:MGI:1916245] | 569,3 | 100,00 |
| ENSMUSG00000031148 | Gpkow | G patch domain and KOW motifs [Source:MGI Symbol;Acc:MGI:1859610] | 569,2 | 98,36 |
| ENSMUSG00000021072 | Tmx1 | thioredoxin-related transmembrane protein 1 [Source:MGI Symbol;Acc:MGI:1919986] | 569,1 | 95,31 |
| ENSMUSG00000039285 | Azi2 | 5-azacytidine induced gene 2 [Source:MGI Symbol;Acc:MGI:1351332] | 568,2 | 76,38 |
| ENSMUSG00000058546 | Rpl23a | ribosomal protein L23A [Source:MGI Symbol;Acc:MGI:3040672] | 567,7 | 75,95 |
| ENSMUSG00000053317 | Sec61b | Sec61 beta subunit [Source:MGI Symbol;Acc:MGI:1913462] | 567,7 | 75,09 |
| ENSMUSG00000033022 | Cdo1 | cysteine dioxygenase 1, cytosolic [Source:MGI Symbol;Acc:MGI:105925] | 567,6 | 100,00 |
| ENSMUSG00000020069 | Hnrnph3 | heterogeneous nuclear ribonucleoprotein H3 [Source:MGI Symbol;Acc:MGI:1926462] | 567,2 | 97,62 |
| ENSMUSG00000028452 | Vcp | valosin containing protein [Source:MGI Symbol;Acc:MGI:99919] | 567,1 | 80,11 |
| ENSMUSG00000022091 | Sorbs3 | sorbin and SH3 domain containing 3 [Source:MGI Symbol;Acc:MGI:700013] | 567,1 | 97,60 |
| ENSMUSG00000042506 | Usp22 | ubiquitin specific peptidase 22 [Source:MGI Symbol;Acc:MGI:2144157] | 566,7 | 99,54 |
| ENSMUSG00000021395 | Spin1 | spindlin 1 [Source:MGI Symbol;Acc:MGI:109242] | 566,6 | 91,45 |
| ENSMUSG00000031834 | Pik3r2 | osphatidylinositol 3-kinase, regulatory subunit, polypeptide 2 (p85 beta) [Source:MGI Symbol;Acc:MGI:10987 | 566,5 | 94,88 |
| ENSMUSG00000019578 | Ubxn6 | UBX domain protein 6 [Source:MGI Symbol;Acc:MGI:1913780] | 566,2 | 82,06 |
| ENSMUSG00000091264 | Smim13 | small integral membrane protein 13 [Source:MGI Symbol;Acc:MGI:2652854] | 565,9 | 98,66 |
| ENSMUSG00000059811 | Atl2 | atlastin GTPase 2 [Source:MGI Symbol;Acc:MGI:1929492] | 565,4 | 99,10 |
| ENSMUSG00000002477 | Snrpd1 | small nuclear ribonucleoprotein D1 [Source:MGI Symbol;Acc:MGI:98344] | 565,3 | 100,00 |
| ENSMUSG00000000340 | Dbt | dihydrolipoamide branched chain transacylase E2 [Source:MGI Symbol;Acc:MGI:105386] | 565,3 | 98,38 |
| ENSMUSG00000062661 | Ncs1 | neuronal calcium sensor 1 [Source:MGI Symbol;Acc:MGI:109166] | 564,9 | 96,51 |
| ENSMUSG00000058594 | Fbxo18 | F-box protein 18 [Source:MGI Symbol;Acc:MGI:1354699] | 564,5 | 98,94 |
| ENSMUSG00000049421 | Zfp260 | zinc finger protein 260 [Source:MGI Symbol;Acc:MGI:1347071] | 564,5 | 92,70 |
| ENSMUSG00000035824 | Tk2 | thymidine kinase 2, mitochondrial [Source:MGI Symbol;Acc:MGI:1913266] | 564,2 | 99,81 |
| ENSMUSG00000015087 | Rabl6 | RAB, member RAS oncogene family-like 6 [Source:MGI Symbol;Acc:MGI:2442633] | 564,2 | 99,93 |
| ENSMUSG00000003721 | Insig2 | insulin induced gene 2 [Source:MGI Symbol;Acc:MGI:1920249] | 563,2 | 82,69 |
| ENSMUSG00000018736 | Ndel1 | nuclear distribution gene E-like homolog 1 (A. nidulans) [Source:MGI Symbol;Acc:MGI:1932915] | 562,5 | 76,81 |
| ENSMUSG00000055239 | Kcmf1 | potassium channel modulatory factor 1 [Source:MGI Symbol;Acc:MGI:1921537] | 562,2 | 92,72 |
| ENSMUSG00000025609 | Mkln1 | muskelin 1, intracellular mediator containing kelch motifs [Source:MGI Symbol;Acc:MGI:1351638] | 562,0 | 72,41 |
| ENSMUSG00000026155 | Smap1 | small ArfGAP 1 [Source:MGI Symbol;Acc:MGI:2138261] | 562,0 | 98,73 |
| ENSMUSG00000068747 | Sort1 | sortilin 1 [Source:MGI Symbol;Acc:MGI:1338015] | 561,3 | 99,11 |
| ENSMUSG00000053565 | Eif3k | eukaryotic translation initiation factor 3, subunit K [Source:MGI Symbol;Acc:MGI:1921080] | 561,1 | 84,40 |
| ENSMUSG00000030298 | Sec13 | SEC13 homolog, nuclear pore and COPII coat complex component [Source:MGI Symbol;Acc:MGI:99832] | 561,1 | 99,26 |
| ENSMUSG00000022761 | Lztr1 | leucine-zipper-like transcriptional regulator, 1 [Source:MGI Symbol;Acc:MGI:1914113] | 561,1 | 96,01 |
| ENSMUSG00000026869 | Psmd5 | proteasome (prosome, macropain) 26S subunit, non-ATPase, 5 [Source:MGI Symbol;Acc:MGI:1914248] | 560,9 | 83,11 |
| ENSMUSG00000001441 | Npepps | aminopeptidase puromycin sensitive [Source:MGI Symbol;Acc:MGI:1101358] | 560,4 | 94,46 |
| ENSMUSG00000020015 | Cdk17 | cyclin-dependent kinase 17 [Source:MGI Symbol;Acc:MGI:97517] | 560,3 | 99,52 |
| ENSMUSG00000022537 | Tmem44 | transmembrane protein 44 [Source:MGI Symbol;Acc:MGI:1924489] | 560,2 | 99,90 |
| ENSMUSG00000024856 | Cdk2ap2 | CDK2-associated protein 2 [Source:MGI Symbol;Acc:MGI:1098779] | 560,1 | 89,84 |
| ENSMUSG00000052369 | Tmem106c | transmembrane protein 106C [Source:MGI Symbol;Acc:MGI:1196384] | 559,9 | 99,61 |
| ENSMUSG00000027852 | Nras | neuroblastoma ras oncogene [Source:MGI Symbol;Acc:MGI:97376] | 559,6 | 92,71 |
| ENSMUSG00000040859 | Bsdc1 | BSD domain containing 1 [Source:MGI Symbol;Acc:MGI:1913466] | 559,6 | 93,50 |
| ENSMUSG00000041712 | Ubr7 | ubiquitin protein ligase E3 component n-recognin 7 (putative) [Source:MGI Symbol;Acc:MGI:1913872] | 559,3 | 100,00 |
| ENSMUSG00000034761 | Map4k5 | mitogen-activated protein kinase kinase kinase kinase 5 [Source:MGI Symbol;Acc:MGI:1925503] | 559,0 | 98,10 |
| ENSMUSG00000047514 | Tspyl1 | testis-specific protein, Y-encoded-like 1 [Source:MGI Symbol;Acc:MGI:1298395] | 558,8 | 88,77 |
| ENSMUSG00000020646 | Mboat2 | membrane bound O-acyltransferase domain containing 2 [Source:MGI Symbol;Acc:MGI:1914466] | 558,8 | 100,00 |
| ENSMUSG00000019831 | Wasf1 | WAS protein family, member 1 [Source:MGI Symbol;Acc:MGI:1890563] | 558,6 | 96,38 |
| ENSMUSG00000046519 | Golph3l | golgi phosphoprotein 3-like [Source:MGI Symbol;Acc:MGI:1917129] | 558,3 | 99,86 |
| ENSMUSG00000038005 | 2700029M09Rik | RIKEN cDNA 2700029M09 gene [Source:MGI Symbol;Acc:MGI:1919862] | 558,2 | 99,92 |
| ENSMUSG00000047694 | Yipf6 | Yip1 domain family, member 6 [Source:MGI Symbol;Acc:MGI:1925179] | 558,1 | 97,21 |
| ENSMUSG00000031563 | Wwc2 | WW, C2 and coiled-coil domain containing 2 [Source:MGI Symbol;Acc:MGI:1261872] | 558,0 | 97,72 |
| ENSMUSG00000025745 | Hadha | enase/3-ketoacyl-Coenzyme A thiolase/enoyl-Coenzyme A hydratase (trifunctional protein), alpha subunit [Sou | 557,4 | 79,11 |
| ENSMUSG00000039157 | Fam102a | family with sequence similarity 102, member A [Source:MGI Symbol;Acc:MGI:2138935] | 557,3 | 99,92 |
| ENSMUSG00000060261 | Gtf2i | general transcription factor II I [Source:MGI Symbol;Acc:MGI:1202722] | 557,2 | 92,49 |
| ENSMUSG00000039849 | Pcif1 | PDX1 C-terminal inhibiting factor 1 [Source:MGI Symbol;Acc:MGI:2443858] | 557,2 | 99,46 |
| ENSMUSG00000032463 | Faim | Fas apoptotic inhibitory molecule [Source:MGI Symbol;Acc:MGI:1344387] | 557,0 | 97,97 |
| ENSMUSG00000019970 | Sgk1 | serum/glucocorticoid regulated kinase 1 [Source:MGI Symbol;Acc:MGI:1340062] | 556,9 | 78,21 |
| ENSMUSG00000071653 | 1810009A15Rik | RIKEN cDNA 1810009A15 gene [Source:MGI Symbol;Acc:MGI:1913526] | 556,6 | 99,50 |
| ENSMUSG00000030301 | Ccdc91 | coiled-coil domain containing 91 [Source:MGI Symbol;Acc:MGI:1914265] | 556,5 | 99,13 |
| ENSMUSG00000051483 | Cbr1 | carbonyl reductase 1 [Source:MGI Symbol;Acc:MGI:88284] | 556,3 | 87,04 |
| ENSMUSG00000029993 | Nfu1 | NFU1 iron-sulfur cluster scaffold homolog (S. cerevisiae) [Source:MGI Symbol;Acc:MGI:1913290] | 556,0 | 80,20 |
| ENSMUSG00000001036 | Epn2 | epsin 2 [Source:MGI Symbol;Acc:MGI:1333766] | 555,8 | 96,52 |
| ENSMUSG00000021906 | Oxnad1 | oxidoreductase NAD-binding domain containing 1 [Source:MGI Symbol;Acc:MGI:1916953] | 555,8 | 99,26 |
| ENSMUSG00000052456 | Asna1 | arsA arsenite transporter, ATP-binding, homolog 1 (bacterial) [Source:MGI Symbol;Acc:MGI:1928379] | 555,5 | 99,15 |
| ENSMUSG00000015714 | Cers2 | ceramide synthase 2 [Source:MGI Symbol;Acc:MGI:1924143] | 555,4 | 90,06 |
| ENSMUSG00000027223 | Mapk8ip1 | mitogen-activated protein kinase 8 interacting protein 1 [Source:MGI Symbol;Acc:MGI:1309464] | 554,3 | 94,43 |
| ENSMUSG00000013662 | Atad1 | ATPase family, AAA domain containing 1 [Source:MGI Symbol;Acc:MGI:1915229] | 553,9 | 99,96 |
| ENSMUSG00000018761 | Mpdu1 | mannose-P-dolichol utilization defect 1 [Source:MGI Symbol;Acc:MGI:1346040] | 553,8 | 99,60 |
| ENSMUSG00000053646 | Plxnb1 | plexin B1 [Source:MGI Symbol;Acc:MGI:2154238] | 553,7 | 94,11 |
| ENSMUSG00000055436 | Srsf11 | serine/arginine-rich splicing factor 11 [Source:MGI Symbol;Acc:MGI:1916457] | 553,5 | 92,86 |
| ENSMUSG00000022296 | Baalc | brain and acute leukemia, cytoplasmic [Source:MGI Symbol;Acc:MGI:1928704] | 553,3 | 71,65 |
| ENSMUSG00000033365 | Ipo13 | importin 13 [Source:MGI Symbol;Acc:MGI:2385205] | 553,1 | 99,97 |
| ENSMUSG00000028057 | Rit1 | Ras-like without CAAX 1 [Source:MGI Symbol;Acc:MGI:108053] | 552,2 | 99,51 |
| ENSMUSG00000019837 | Gtf3c6 | general transcription factor IIIC, polypeptide 6, alpha [Source:MGI Symbol;Acc:MGI:1914621] | 552,2 | 100,00 |
| ENSMUSG00000036825 | Ssx2ip | synovial sarcoma, X breakpoint 2 interacting protein [Source:MGI Symbol;Acc:MGI:2139150] | 552,2 | 94,04 |
| ENSMUSG00000039701 | Usp53 | ubiquitin specific peptidase 53 [Source:MGI Symbol;Acc:MGI:2139607] | 552,1 | 80,58 |
| ENSMUSG00000031799 | Tpm4 | tropomyosin 4 [Source:MGI Symbol;Acc:MGI:2449202] | 551,6 | 98,28 |
| ENSMUSG00000028409 | Smu1 | smu-1 suppressor of mec-8 and unc-52 homolog (C. elegans) [Source:MGI Symbol;Acc:MGI:1915546] | 551,4 | 96,64 |
| ENSMUSG00000027285 | Haus2 | HAUS augmin-like complex, subunit 2 [Source:MGI Symbol;Acc:MGI:1913546] | 551,0 | 98,17 |
| ENSMUSG00000073131 | Vma21 | VMA21 vacuolar H+-ATPase homolog (S. cerevisiae) [Source:MGI Symbol;Acc:MGI:1914298] | 550,6 | 80,96 |
| ENSMUSG00000063931 | Pepd | peptidase D [Source:MGI Symbol;Acc:MGI:97542] | 550,3 | 98,62 |
| ENSMUSG00000035864 | Syt1 | synaptotagmin I [Source:MGI Symbol;Acc:MGI:99667] | 550,1 | 97,03 |
| ENSMUSG00000028676 | Srsf10 | serine/arginine-rich splicing factor 10 [Source:MGI Symbol;Acc:MGI:1333805] | 549,8 | 84,48 |
| ENSMUSG00000062352 | Itgb1bp1 | integrin beta 1 binding protein 1 [Source:MGI Symbol;Acc:MGI:1306802] | 549,8 | 100,00 |
| ENSMUSG00000033610 | Pank1 | pantothenate kinase 1 [Source:MGI Symbol;Acc:MGI:1922985] | 549,2 | 98,25 |
| ENSMUSG00000072694 | 1500011B03Rik | RIKEN cDNA 1500011B03 gene [Source:MGI Symbol;Acc:MGI:1913486] | 549,1 | 99,54 |
| ENSMUSG00000022505 | Emp2 | epithelial membrane protein 2 [Source:MGI Symbol;Acc:MGI:1098726] | 549,0 | 99,37 |
| ENSMUSG00000000902 | Smarcb1 | d, matrix associated, actin dependent regulator of chromatin, subfamily b, member 1 [Source:MGI Symbol;Acc | 548,8 | 99,90 |
| ENSMUSG00000001482 | Def8 | differentially expressed in FDCP 8 [Source:MGI Symbol;Acc:MGI:1346331] | 548,7 | 85,04 |
| ENSMUSG00000037697 | Ddhd1 | DDHD domain containing 1 [Source:MGI Symbol;Acc:MGI:2150302] | 548,6 | 98,94 |
| ENSMUSG00000011096 | Akt1s1 | AKT1 substrate 1 (proline-rich) [Source:MGI Symbol;Acc:MGI:1914855] | 548,2 | 91,47 |
| ENSMUSG00000027067 | Ssrp1 | structure specific recognition protein 1 [Source:MGI Symbol;Acc:MGI:107912] | 548,2 | 98,19 |
| ENSMUSG00000002341 | Ncan | neurocan [Source:MGI Symbol;Acc:MGI:104694] | 548,0 | 98,37 |
| ENSMUSG00000020152 | Actr2 | ARP2 actin-related protein 2 [Source:MGI Symbol;Acc:MGI:1913963] | 547,9 | 97,78 |
| ENSMUSG00000024603 | Dctn4 | dynactin 4 [Source:MGI Symbol;Acc:MGI:1914915] | 547,3 | 99,94 |
| ENSMUSG00000021373 | Cap2 | CAP, adenylate cyclase-associated protein, 2 (yeast) [Source:MGI Symbol;Acc:MGI:1914502] | 547,1 | 94,22 |
| ENSMUSG00000025246 | Tbl1x | transducin (beta)-like 1 X-linked [Source:MGI Symbol;Acc:MGI:1336172] | 547,1 | 99,26 |
| ENSMUSG00000019471 | Cdc37 | cell division cycle 37 [Source:MGI Symbol;Acc:MGI:109531] | 547,0 | 88,78 |
| ENSMUSG00000039221 | Rpl22l1 | ribosomal protein L22 like 1 [Source:MGI Symbol;Acc:MGI:1915278] | 546,9 | 87,82 |
| ENSMUSG00000026276 | Sept2 | septin 2 [Source:MGI Symbol;Acc:MGI:97298] | 546,6 | 79,53 |
| ENSMUSG00000009216 | Fam163b | family with sequence similarity 163, member B [Source:MGI Symbol;Acc:MGI:1926106] | 546,6 | 99,22 |
| ENSMUSG00000008090 | Fgfrl1 | fibroblast growth factor receptor-like 1 [Source:MGI Symbol;Acc:MGI:2150920] | 546,1 | 92,81 |
| ENSMUSG00000030986 | Dhx32 | DEAH (Asp-Glu-Ala-His) box polypeptide 32 [Source:MGI Symbol;Acc:MGI:2141813] | 546,1 | 84,68 |
| ENSMUSG00000020432 | Tcn2 | transcobalamin 2 [Source:MGI Symbol;Acc:MGI:98534] | 546,1 | 93,88 |
| ENSMUSG00000006412 | Pfdn2 | prefoldin 2 [Source:MGI Symbol;Acc:MGI:1276111] | 546,0 | 97,44 |
| ENSMUSG00000045854 | Lyrm2 | LYR motif containing 2 [Source:MGI Symbol;Acc:MGI:1917573] | 545,7 | 100,00 |
| ENSMUSG00000008301 | Phax | phosphorylated adaptor for RNA export [Source:MGI Symbol;Acc:MGI:1891839] | 545,3 | 91,26 |
| ENSMUSG00000062098 | Btbd3 | BTB (POZ) domain containing 3 [Source:MGI Symbol;Acc:MGI:2385155] | 545,1 | 99,04 |
| ENSMUSG00000021010 | Npas3 | neuronal PAS domain protein 3 [Source:MGI Symbol;Acc:MGI:1351610] | 544,9 | 99,79 |
| ENSMUSG00000015790 | Surf1 | surfeit gene 1 [Source:MGI Symbol;Acc:MGI:98443] | 544,9 | 99,65 |

| | | | | |
|---|---|---|---|---|
| ENSMUSG00000047989 | Ino80c | INO80 complex subunit C [Source:MGI Symbol;Acc:MGI:2443014] | 543,4 | 88,13 |
| ENSMUSG00000033751 | Gadd45gip1 | rowth arrest and DNA-damage-inducible, gamma interacting protein 1 [Source:MGI Symbol;Acc:MGI:191494 | 542,8 | 74,57 |
| ENSMUSG00000059796 | Eif4a1 | eukaryotic translation initiation factor 4A1 [Source:MGI Symbol;Acc:MGI:95303] | 542,7 | 88,46 |
| ENSMUSG00000009394 | Syn2 | synapsin II [Source:MGI Symbol;Acc:MGI:103020] | 542,3 | 98,84 |
| ENSMUSG00000026313 | Hdac4 | histone deacetylase 4 [Source:MGI Symbol;Acc:MGI:3036234] | 542,2 | 96,17 |
| ENSMUSG00000018481 | Appbp2 | amyloid beta precursor protein (cytoplasmic tail) binding protein 2 [Source:MGI Symbol;Acc:MGI:1914134] | 542,2 | 99,21 |
| ENSMUSG00000024789 | Jak2 | Janus kinase 2 [Source:MGI Symbol;Acc:MGI:96629] | 542,2 | 99,62 |
| ENSMUSG00000024843 | Chka | choline kinase alpha [Source:MGI Symbol;Acc:MGI:107760] | 541,6 | 95,64 |
| ENSMUSG00000022247 | Brix1 | BRX1, biogenesis of ribosomes [Source:MGI Symbol;Acc:MGI:1915082] | 541,4 | 95,75 |
| ENSMUSG00000024826 | Dpf2 | D4, zinc and double PHD fingers family 2 [Source:MGI Symbol;Acc:MGI:109529] | 541,3 | 81,65 |
| ENSMUSG00000036879 | Phkb | phosphorylase kinase beta [Source:MGI Symbol;Acc:MGI:97578] | 540,7 | 81,11 |
| ENSMUSG00000063802 | Hspbp1 | SPA (heat shock 70kDa) binding protein, cytoplasmic cochaperone 1 [Source:MGI Symbol;Acc:MGI:191349 | 540,7 | 93,57 |
| ENSMUSG00000074227 | Spint2 | serine protease inhibitor, Kunitz type 2 [Source:MGI Symbol;Acc:MGI:1338031] | 540,6 | 100,00 |
| ENSMUSG00000030452 | Nipa2 | on imprinted in Prader-Willi/Angelman syndrome 2 homolog (human) [Source:MGI Symbol;Acc:MGI:191391 | 540,5 | 84,71 |
| ENSMUSG00000003873 | Bax | BCL2-associated X protein [Source:MGI Symbol;Acc:MGI:99702] | 540,1 | 97,70 |
| ENSMUSG00000026095 | Asnsd1 | asparagine synthetase domain containing 1 [Source:MGI Symbol;Acc:MGI:1917646] | 540,1 | 85,81 |
| ENSMUSG00000030849 | Fgfr2 | fibroblast growth factor receptor 2 [Source:MGI Symbol;Acc:MGI:95523] | 540,0 | 96,37 |
| ENSMUSG00000109336 | Samd4b | sterile alpha motif domain containing 4B [Source:MGI Symbol;Acc:MGI:2448542] | 539,9 | 98,14 |
| ENSMUSG00000010392 | Gosr1 | golgi SNAP receptor complex member 1 [Source:MGI Symbol;Acc:MGI:1858260] | 539,8 | 99,91 |
| ENSMUSG00000005505 | Kbtbd4 | kelch repeat and BTB (POZ) domain containing 4 [Source:MGI Symbol;Acc:MGI:1914386] | 539,5 | 99,63 |
| ENSMUSG00000105204 | Gm43738 | predicted gene 43738 [Source:MGI Symbol;Acc:MGI:5663875] | 539,4 | 93,26 |
| ENSMUSG00000062647 | Rpl7a | ribosomal protein L7A [Source:MGI Symbol;Acc:MGI:1353472] | 539,1 | 93,07 |
| ENSMUSG00000032366 | Tpm1 | tropomyosin 1, alpha [Source:MGI Symbol;Acc:MGI:98809] | 538,8 | 91,88 |
| ENSMUSG00000035764 | Fbxo45 | F-box protein 45 [Source:MGI Symbol;Acc:MGI:2447775] | 538,6 | 97,73 |
| ENSMUSG00000020530 | Ggnbp2 | gametogenetin binding protein 2 [Source:MGI Symbol;Acc:MGI:2387356] | 538,6 | 90,77 |
| ENSMUSG00000066324 | Impad1 | inositol monophosphatase domain containing 1 [Source:MGI Symbol;Acc:MGI:1915720] | 538,5 | 99,47 |
| ENSMUSG00000029617 | Ccz1 | CCZ1 vacuolar protein trafficking and biogenesis associated [Source:MGI Symbol;Acc:MGI:2141070] | 538,5 | 99,83 |
| ENSMUSG00000020598 | Nrcam | neuronal cell adhesion molecule [Source:MGI Symbol;Acc:MGI:104750] | 538,0 | 85,23 |
| ENSMUSG00000087408 | Cers1 | ceramide synthase 1 [Source:MGI Symbol;Acc:MGI:2136690] | 537,9 | 100,00 |
| ENSMUSG00000052957 | Gas1 | growth arrest specific 1 [Source:MGI Symbol;Acc:MGI:95655] | 537,8 | 99,95 |
| ENSMUSG00000032009 | Sesn3 | sestrin 3 [Source:MGI Symbol;Acc:MGI:1922997] | 537,5 | 89,76 |
| ENSMUSG00000026127 | Imp4 | IMP4, U3 small nucleolar ribonucleoprotein [Source:MGI Symbol;Acc:MGI:106572] | 537,1 | 98,14 |
| ENSMUSG00000008976 | Gabpa | GA repeat binding protein, alpha [Source:MGI Symbol;Acc:MGI:95610] | 537,1 | 96,35 |
| ENSMUSG00000007458 | M6pr | mannose-6-phosphate receptor, cation dependent [Source:MGI Symbol;Acc:MGI:96904] | 537,0 | 95,92 |
| ENSMUSG00000022382 | Wnt7b | wingless-type MMTV integration site family, member 7B [Source:MGI Symbol;Acc:MGI:98962] | 537,0 | 81,63 |
| ENSMUSG00000046093 | Hpcal4 | hippocalcin-like 4 [Source:MGI Symbol;Acc:MGI:2157521] | 536,7 | 95,72 |
| ENSMUSG00000034157 | Cipc | CLOCK interacting protein, circadian [Source:MGI Symbol;Acc:MGI:1919185] | 536,3 | 91,54 |
| ENSMUSG00000014361 | Mertk | c-mer proto-oncogene tyrosine kinase [Source:MGI Symbol;Acc:MGI:96965] | 536,3 | 100,00 |
| ENSMUSG00000005262 | Ufd1l | ubiquitin fusion degradation 1 like [Source:MGI Symbol;Acc:MGI:109353] | 536,1 | 75,82 |
| ENSMUSG00000032609 | Klhdc8b | kelch domain containing 8B [Source:MGI Symbol;Acc:MGI:1925517] | 536,1 | 70,29 |
| ENSMUSG00000097675 | 1700101I11Rik | RIKEN cDNA 1700101I11 gene [Source:MGI Symbol;Acc:MGI:1915180] | 535,6 | 71,18 |
| ENSMUSG00000037149 | Ddx1 | DEAD (Asp-Glu-Ala-Asp) box polypeptide 1 [Source:MGI Symbol;Acc:MGI:2144727] | 535,6 | 99,48 |
| ENSMUSG00000021760 | Gpx8 | glutathione peroxidase 8 (putative) [Source:MGI Symbol;Acc:MGI:1916840] | 535,4 | 100,00 |
| ENSMUSG00000029603 | Dtx1 | deltex 1, E3 ubiquitin ligase [Source:MGI Symbol;Acc:MGI:1352744] | 534,9 | 93,13 |
| ENSMUSG00000052430 | Bmpr1b | bone morphogenetic protein receptor, type 1B [Source:MGI Symbol;Acc:MGI:107191] | 534,9 | 93,96 |
| ENSMUSG00000030421 | Uri1 | URI1, prefoldin-like chaperone [Source:MGI Symbol;Acc:MGI:1342294] | 534,6 | 96,20 |
| ENSMUSG00000028211 | Trp53inp1 | transformation related protein 53 inducible nuclear protein 1 [Source:MGI Symbol;Acc:MGI:1926609] | 534,2 | 99,65 |
| ENSMUSG00000068099 | 1500009C09Rik | RIKEN cDNA 1500009C09 gene [Source:MGI Symbol;Acc:MGI:1923755] | 533,9 | 78,82 |
| ENSMUSG00000020601 | Trib2 | tribbles homolog 2 (Drosophila) [Source:MGI Symbol;Acc:MGI:2145021] | 533,7 | 99,95 |
| ENSMUSG00000009075 | Cabp7 | calcium binding protein 7 [Source:MGI Symbol;Acc:MGI:2183437] | 533,3 | 99,39 |
| ENSMUSG00000021236 | Entpd5 | ectonucleoside triphosphate diphosphohydrolase 5 [Source:MGI Symbol;Acc:MGI:1321385] | 532,6 | 97,63 |
| ENSMUSG00000038145 | Snrk | SNF related kinase [Source:MGI Symbol;Acc:MGI:108104] | 532,2 | 85,17 |
| ENSMUSG00000001576 | Ergic1 | endoplasmic reticulum-golgi intermediate compartment (ERGIC) 1 [Source:MGI Symbol;Acc:MGI:1914708] | 532,2 | 95,16 |
| ENSMUSG00000024847 | Aip | aryl-hydrocarbon receptor-interacting protein [Source:MGI Symbol;Acc:MGI:109622] | 532,2 | 99,60 |
| ENSMUSG00000020340 | Cyfip2 | cytoplasmic FMR1 interacting protein 2 [Source:MGI Symbol;Acc:MGI:1924134] | 532,0 | 89,29 |
| ENSMUSG00000025495 | Ptdss2 | phosphatidylserine synthase 2 [Source:MGI Symbol;Acc:MGI:1351664] | 531,7 | 89,55 |
| ENSMUSG00000028273 | Pdlim5 | PDZ and LIM domain 5 [Source:MGI Symbol;Acc:MGI:1927489] | 531,4 | 86,78 |
| ENSMUSG00000035413 | Tmem98 | transmembrane protein 98 [Source:MGI Symbol;Acc:MGI:1923457] | 531,4 | 88,53 |
| ENSMUSG00000022562 | Oplah | 5-oxoprolinase (ATP-hydrolysing) [Source:MGI Symbol;Acc:MGI:1922725] | 531,4 | 91,11 |
| ENSMUSG00000049517 | Rps23 | ribosomal protein S23 [Source:MGI Symbol;Acc:MGI:1913725] | 531,2 | 90,08 |
| ENSMUSG00000018583 | G3bp1 | GTPase activating protein (SH3 domain) binding protein 1 [Source:MGI Symbol;Acc:MGI:1351465] | 531,2 | 86,70 |
| ENSMUSG00000025580 | Eif4a3 | eukaryotic translation initiation factor 4A3 [Source:MGI Symbol;Acc:MGI:1923731] | 530,7 | 99,78 |
| ENSMUSG00000024248 | Cox7a2l | cytochrome c oxidase subunit VIIa polypeptide 2-like [Source:MGI Symbol;Acc:MGI:106015] | 530,5 | 82,33 |
| ENSMUSG00000010290 | AI597479 | expressed sequence AI597479 [Source:MGI Symbol;Acc:MGI:2138299] | 530,5 | 99,65 |
| ENSMUSG00000018411 | Mapt | microtubule-associated protein tau [Source:MGI Symbol;Acc:MGI:97180] | 529,9 | 81,63 |
| ENSMUSG00000026526 | Fh1 | fumarate hydratase 1 [Source:MGI Symbol;Acc:MGI:95530] | 529,5 | 85,13 |
| ENSMUSG00000021591 | Glrx | glutaredoxin [Source:MGI Symbol;Acc:MGI:2135625] | 529,1 | 73,80 |
| ENSMUSG00000053141 | Ptprt | protein tyrosine phosphatase, receptor type, T [Source:MGI Symbol;Acc:MGI:1321152] | 529,0 | 78,49 |
| ENSMUSG00000034667 | Xpot | exportin, tRNA (nuclear export receptor for tRNAs) [Source:MGI Symbol;Acc:MGI:1920442] | 528,6 | 98,09 |
| ENSMUSG00000022175 | Lrp10 | low-density lipoprotein receptor-related protein 10 [Source:MGI Symbol;Acc:MGI:1929480] | 528,5 | 75,30 |
| ENSMUSG00000039917 | Rhbdd2 | rhomboid domain containing 2 [Source:MGI Symbol;Acc:MGI:1915612] | 528,5 | 96,06 |
| ENSMUSG00000041168 | Lonp1 | lon peptidase 1, mitochondrial [Source:MGI Symbol;Acc:MGI:1921392] | 527,9 | 100,00 |
| ENSMUSG00000059495 | Arhgef12 | Rho guanine nucleotide exchange factor (GEF) 12 [Source:MGI Symbol;Acc:MGI:1916882] | 527,6 | 91,94 |
| ENSMUSG00000028035 | Dnajb4 | DnaJ heat shock protein family (Hsp40) member B4 [Source:MGI Symbol;Acc:MGI:1914285] | 527,4 | 91,57 |
| ENSMUSG00000034789 | Rab24 | RAB24, member RAS oncogene family [Source:MGI Symbol;Acc:MGI:105065] | 527,0 | 90,58 |
| ENSMUSG00000005615 | Pcyt1a | phosphate cytidylyltransferase 1, choline, alpha isoform [Source:MGI Symbol;Acc:MGI:88557] | 526,9 | 92,38 |
| ENSMUSG00000038170 | Pde4dip | phosphodiesterase 4D interacting protein (myomegalin) [Source:MGI Symbol;Acc:MGI:1891434] | 526,8 | 78,82 |
| ENSMUSG00000003363 | Pld3 | phospholipase D family, member 3 [Source:MGI Symbol;Acc:MGI:1333782] | 526,7 | 92,11 |
| ENSMUSG00000039536 | Stau1 | staufen (RNA binding protein) homolog 1 (Drosophila) [Source:MGI Symbol;Acc:MGI:1338864] | 526,7 | 98,18 |
| ENSMUSG00000073436 | Eme2 | essential meiotic structure-specific endonuclease subunit 2 [Source:MGI Symbol;Acc:MGI:1919889] | 526,4 | 79,80 |
| ENSMUSG00000020397 | Med7 | mediator complex subunit 7 [Source:MGI Symbol;Acc:MGI:1913463] | 526,3 | 90,82 |
| ENSMUSG00000085838 | Chn1os1 | chimerin 1, opposite strand 1 [Source:MGI Symbol;Acc:MGI:3702041] | 526,2 | 91,98 |
| ENSMUSG00000009614 | Sardh | sarcosine dehydrogenase [Source:MGI Symbol;Acc:MGI:2183102] | 525,7 | 95,87 |
| ENSMUSG00000026761 | Orc4 | origin recognition complex, subunit 4 [Source:MGI Symbol;Acc:MGI:1347043] | 525,1 | 95,60 |
| ENSMUSG00000028437 | Ubap1 | ubiquitin-associated protein 1 [Source:MGI Symbol;Acc:MGI:2149543] | 524,9 | 77,68 |
| ENSMUSG00000070934 | Rraga | Ras-related GTP binding A [Source:MGI Symbol;Acc:MGI:1915691] | 524,6 | 100,00 |
| ENSMUSG00000029681 | Bcl7b | B cell CLL/lymphoma 7B [Source:MGI Symbol;Acc:MGI:1332238] | 524,5 | 95,56 |
| ENSMUSG00000046314 | Stxbp6 | syntaxin binding protein 6 (amisyn) [Source:MGI Symbol;Acc:MGI:2384963] | 524,5 | 95,83 |
| ENSMUSG00000040771 | Oard1 | O-acyl-ADP-ribose deacylase 1 [Source:MGI Symbol;Acc:MGI:2146818] | 524,5 | 98,70 |
| ENSMUSG00000031697 | Orc6 | origin recognition complex, subunit 6 [Source:MGI Symbol;Acc:MGI:1929285] | 524,2 | 98,51 |
| ENSMUSG00000006058 | Snf8 | SNF8, ESCRT-II complex subunit, homolog (S. cerevisiae) [Source:MGI Symbol;Acc:MGI:1343161] | 524,0 | 94,89 |
| ENSMUSG00000071856 | Mcc | mutated in colorectal cancers [Source:MGI Symbol;Acc:MGI:96930] | 524,0 | 89,15 |
| ENSMUSG00000025630 | Hprt | hypoxanthine guanine phosphoribosyl transferase [Source:MGI Symbol;Acc:MGI:96217] | 523,9 | 99,95 |
| ENSMUSG00000032050 | Rdx | radixin [Source:MGI Symbol;Acc:MGI:97887] | 523,8 | 91,37 |
| ENSMUSG00000031639 | Tlr3 | toll-like receptor 3 [Source:MGI Symbol;Acc:MGI:2156367] | 523,7 | 98,10 |
| ENSMUSG00000073418 | C4b | complement component 4B (Chido blood group) [Source:MGI Symbol;Acc:MGI:88228] | 523,6 | 89,86 |
| ENSMUSG00000005506 | Celf1 | CUGBP, Elav-like family member 1 [Source:MGI Symbol;Acc:MGI:1342295] | 523,4 | 90,93 |
| ENSMUSG00000061758 | Akr1b10 | aldo-keto reductase family 1, member B10 (aldose reductase) [Source:MGI Symbol;Acc:MGI:1915111] | 522,9 | 99,66 |
| ENSMUSG00000057177 | Gsk3a | glycogen synthase kinase 3 alpha [Source:MGI Symbol;Acc:MGI:2152453] | 522,8 | 96,67 |
| ENSMUSG00000038274 | Fau | is-Reilly murine sarcoma virus (FBR-MuSV) ubiquitously expressed (fox derived) [Source:MGI Symbol;Acc:MG | 522,7 | 83,35 |
| ENSMUSG00000057766 | Ankrd29 | ankyrin repeat domain 29 [Source:MGI Symbol;Acc:MGI:2687055] | 522,7 | 99,60 |
| ENSMUSG00000059278 | Naa38 | N(alpha)-acetyltransferase 38, NatC auxiliary subunit [Source:MGI Symbol;Acc:MGI:1925554] | 522,3 | 93,24 |
| ENSMUSG00000033918 | Parl | presenilin associated, rhomboid-like [Source:MGI Symbol;Acc:MGI:1277152] | 522,2 | 99,13 |
| ENSMUSG00000039952 | Dag1 | dystroglycan 1 [Source:MGI Symbol;Acc:MGI:101864] | 522,1 | 91,94 |
| ENSMUSG00000025369 | Smarcc2 | d, matrix associated, actin dependent regulator of chromatin, subfamily c, member 2 [Source:MGI Symbol;Ac | 522,1 | 96,90 |
| ENSMUSG00000017485 | Top2b | topoisomerase (DNA) II beta [Source:MGI Symbol;Acc:MGI:98791] | 522,0 | 99,58 |
| ENSMUSG00000030432 | Rpl28 | ribosomal protein L28 [Source:MGI Symbol;Acc:MGI:101839] | 522,0 | 71,64 |
| ENSMUSG00000040532 | Abhd11 | abhydrolase domain containing 11 [Source:MGI Symbol;Acc:MGI:1916008] | 521,8 | 87,89 |
| ENSMUSG00000030275 | Etnk1 | ethanolamine kinase 1 [Source:MGI Symbol;Acc:MGI:1922570] | 521,8 | 73,95 |
| ENSMUSG00000040351 | Ankib1 | ankyrin repeat and IBR domain containing 1 [Source:MGI Symbol;Acc:MGI:1918047] | 521,5 | 98,72 |
| ENSMUSG00000036873 | 2410004B18Rik | RIKEN cDNA 2410004B18 gene [Source:MGI Symbol;Acc:MGI:1913671] | 520,5 | 94,50 |
| ENSMUSG00000028444 | Cntfr | ciliary neurotrophic factor receptor [Source:MGI Symbol;Acc:MGI:99605] | 520,3 | 83,98 |
| ENSMUSG00000020523 | Fam114a2 | family with sequence similarity 114, member A2 [Source:MGI Symbol;Acc:MGI:1917629] | 520,0 | 98,97 |
| ENSMUSG00000032307 | Ube2q2 | ubiquitin-conjugating enzyme E2Q (putative) 2 [Source:MGI Symbol;Acc:MGI:2388672] | 519,7 | 86,42 |
| ENSMUSG00000025982 | Sf3b1 | splicing factor 3b, subunit 1 [Source:MGI Symbol;Acc:MGI:1932339] | 519,3 | 94,59 |
| ENSMUSG00000039686 | Zer1 | zyg-11 related, cell cycle regulator [Source:MGI Symbol;Acc:MGI:2442511] | 518,6 | 95,56 |
| ENSMUSG00000022607 | Ptk2 | PTK2 protein tyrosine kinase 2 [Source:MGI Symbol;Acc:MGI:95481] | 517,9 | 90,15 |

| | | | | |
|---|---|---|---|---|
| ENSMUSG00000037643 | Prkci | protein kinase C, iota [Source:MGI Symbol;Acc:MGI:99260] | 517,7 | 91,24 |
| ENSMUSG00000025059 | Gk | glycerol kinase [Source:MGI Symbol;Acc:MGI:106594] | 517,7 | 81,49 |
| ENSMUSG00000068328 | Aup1 | ancient ubiquitous protein 1 [Source:MGI Symbol;Acc:MGI:107789] | 517,4 | 92,65 |
| ENSMUSG00000034152 | Exoc3 | exocyst complex component 3 [Source:MGI Symbol;Acc:MGI:2443972] | 516,6 | 99,95 |
| ENSMUSG00000023050 | Map3k12 | mitogen-activated protein kinase kinase kinase 12 [Source:MGI Symbol;Acc:MGI:1346881] | 516,2 | 86,03 |
| ENSMUSG00000027109 | Sp3 | trans-acting transcription factor 3 [Source:MGI Symbol;Acc:MGI:1277166] | 516,0 | 98,25 |
| ENSMUSG00000028527 | Ak4 | adenylate kinase 4 [Source:MGI Symbol;Acc:MGI:87979] | 515,7 | 90,31 |
| ENSMUSG00000033781 | Asb13 | ankyrin repeat and SOCS box-containing 13 [Source:MGI Symbol;Acc:MGI:2145525] | 515,5 | 99,86 |
| ENSMUSG00000020198 | Ap3d1 | adaptor-related protein complex 3, delta 1 subunit [Source:MGI Symbol;Acc:MGI:107734] | 515,3 | 99,97 |
| ENSMUSG00000075467 | Dnlz | DNL-type zinc finger [Source:MGI Symbol;Acc:MGI:106559] | 514,9 | 99,62 |
| ENSMUSG00000031158 | Timm17b | translocase of inner mitochondrial membrane 17b [Source:MGI Symbol;Acc:MGI:1343176] | 514,8 | 89,22 |
| ENSMUSG00000015672 | Mrpl32 | mitochondrial ribosomal protein L32 [Source:MGI Symbol;Acc:MGI:2137226] | 514,4 | 100,00 |
| ENSMUSG00000048644 | Ctxn1 | cortexin 1 [Source:MGI Symbol;Acc:MGI:88566] | 514,3 | 97,47 |
| ENSMUSG00000003559 | As3mt | arsenic (+3 oxidation state) methyltransferase [Source:MGI Symbol;Acc:MGI:1929882] | 514,2 | 99,70 |
| ENSMUSG00000019960 | Dusp6 | dual specificity phosphatase 6 [Source:MGI Symbol;Acc:MGI:1914853] | 514,0 | 99,81 |
| ENSMUSG00000003420 | Fcgrt | Fc receptor, IgG, alpha chain transporter [Source:MGI Symbol;Acc:MGI:103017] | 513,9 | 99,52 |
| ENSMUSG00000071657 | Bscl2 | Berardinelli-Seip congenital lipodystrophy 2 (seipin) [Source:MGI Symbol;Acc:MGI:1298392] | 513,7 | 94,34 |
| ENSMUSG00000021646 | Mccc2 | methylcrotonoyl-Coenzyme A carboxylase 2 (beta) [Source:MGI Symbol;Acc:MGI:1925288] | 513,6 | 100,00 |
| ENSMUSG00000014592 | Camta1 | calmodulin binding transcription activator 1 [Source:MGI Symbol;Acc:MGI:2140230] | 513,6 | 77,18 |
| ENSMUSG00000024927 | Rela | v-rel reticuloendotheliosis viral oncogene homolog A (avian) [Source:MGI Symbol;Acc:MGI:103290] | 513,4 | 98,51 |
| ENSMUSG00000074513 | Arfip1 | ADP-ribosylation factor interacting protein 1 [Source:MGI Symbol;Acc:MGI:1277120] | 513,4 | 99,67 |
| ENSMUSG00000028191 | Bcl10 | B cell leukemia/lymphoma 10 [Source:MGI Symbol;Acc:MGI:1337994] | 512,8 | 99,06 |
| ENSMUSG00000056204 | Pgpep1 | pyroglutamyl-peptidase I [Source:MGI Symbol;Acc:MGI:1913772] | 512,6 | 74,49 |
| ENSMUSG00000027742 | Cog6 | component of oligomeric golgi complex 6 [Source:MGI Symbol;Acc:MGI:1914792] | 512,1 | 88,70 |
| ENSMUSG00000027347 | Rasgrp1 | RAS guanyl releasing protein 1 [Source:MGI Symbol;Acc:MGI:1314635] | 511,7 | 97,30 |
| ENSMUSG00000032309 | Fbxo22 | F-box protein 22 [Source:MGI Symbol;Acc:MGI:1926014] | 511,5 | 87,89 |
| ENSMUSG00000047617 | BC029214 | cDNA sequence BC029214 [Source:MGI Symbol;Acc:MGI:2442831] | 511,1 | 100,00 |
| ENSMUSG00000021982 | Cdadc1 | cytidine and dCMP deaminase domain containing 1 [Source:MGI Symbol;Acc:MGI:1919141] | 511,0 | 98,84 |
| ENSMUSG00000000223 | Drp2 | dystrophin related protein 2 [Source:MGI Symbol;Acc:MGI:107432] | 510,9 | 96,54 |
| ENSMUSG00000042705 | Commd10 | COMM domain containing 10 [Source:MGI Symbol;Acc:MGI:1916706] | 510,9 | 99,96 |
| ENSMUSG00000024642 | Tle4 | transducin-like enhancer of split 4, homolog of Drosophila E(spl) [Source:MGI Symbol;Acc:MGI:104633] | 510,8 | 97,78 |
| ENSMUSG00000020925 | Ccdc43 | coiled-coil domain containing 43 [Source:MGI Symbol;Acc:MGI:1289318] | 510,6 | 99,72 |
| ENSMUSG00000039660 | D2Wsu81e | DNA segment, Chr 2, Wayne State University 81, expressed [Source:MGI Symbol;Acc:MGI:106544] | 510,3 | 100,00 |
| ENSMUSG00000031584 | Gsr | glutathione reductase [Source:MGI Symbol;Acc:MGI:95804] | 510,2 | 93,93 |
| ENSMUSG00000040003 | Magi2 | nbrane associated guanylate kinase, WW and PDZ domain containing 2 [Source:MGI Symbol;Acc:MGI:1354 | 510,1 | 84,07 |
| ENSMUSG00000034290 | Nek9 | NIMA (never in mitosis gene a)-related expressed kinase 9 [Source:MGI Symbol;Acc:MGI:2387995] | 509,3 | 99,33 |
| ENSMUSG00000024835 | Coro1b | coronin, actin binding protein 1B [Source:MGI Symbol;Acc:MGI:1345963] | 509,3 | 100,00 |
| ENSMUSG00000026309 | Ilkap | integrin-linked kinase-associated serine/threonine phosphatase 2C [Source:MGI Symbol;Acc:MGI:1914694] | 509,1 | 83,74 |
| ENSMUSG00000028243 | Ubxn2b | UBX domain protein 2B [Source:MGI Symbol;Acc:MGI:1915303] | 509,0 | 91,75 |
| ENSMUSG00000001380 | Hars | histidyl-tRNA synthetase [Source:MGI Symbol;Acc:MGI:108087] | 508,9 | 98,89 |
| ENSMUSG00000087370 | Tmem170b | transmembrane protein 170B [Source:MGI Symbol;Acc:MGI:3647046] | 508,9 | 93,10 |
| ENSMUSG00000039911 | Spsb1 | splA/ryanodine receptor domain and SOCS box containing 1 [Source:MGI Symbol;Acc:MGI:1921896] | 508,7 | 74,01 |
| ENSMUSG00000056076 | Eif3b | eukaryotic translation initiation factor 3, subunit B [Source:MGI Symbol;Acc:MGI:106478] | 508,6 | 98,27 |
| ENSMUSG00000016528 | Mapkapk2 | MAP kinase-activated protein kinase 2 [Source:MGI Symbol;Acc:MGI:109298] | 507,8 | 92,67 |
| ENSMUSG00000026028 | Trak2 | trafficking protein, kinesin binding 2 [Source:MGI Symbol;Acc:MGI:1918077] | 507,8 | 99,22 |
| ENSMUSG00000036106 | Prr5 | proline rich 5 (renal) [Source:MGI Symbol;Acc:MGI:1924714] | 507,6 | 90,14 |
| ENSMUSG00000049960 | Mrps16 | mitochondrial ribosomal protein S16 [Source:MGI Symbol;Acc:MGI:1913492] | 507,6 | 100,00 |
| ENSMUSG00000028470 | Hint2 | histidine triad nucleotide binding protein 2 [Source:MGI Symbol;Acc:MGI:1916167] | 507,3 | 100,00 |
| ENSMUSG00000022055 | Nefl | neurofilament, light polypeptide [Source:MGI Symbol;Acc:MGI:97313] | 507,1 | 99,14 |
| ENSMUSG00000045515 | Pou3f3 | POU domain, class 3, transcription factor 3 [Source:MGI Symbol;Acc:MGI:102564] | 506,9 | 89,13 |
| ENSMUSG00000022635 | Zcrb1 | zinc finger CCHC-type and RNA binding motif 1 [Source:MGI Symbol;Acc:MGI:1914447] | 506,8 | 98,76 |
| ENSMUSG00000019467 | Arhgef25 | Rho guanine nucleotide exchange factor (GEF) 25 [Source:MGI Symbol;Acc:MGI:1277173] | 506,6 | 95,89 |
| ENSMUSG00000030612 | Mrpl46 | mitochondrial ribosomal protein L46 [Source:MGI Symbol;Acc:MGI:1914558] | 506,6 | 97,38 |
| ENSMUSG00000004267 | Eno2 | enolase 2, gamma neuronal [Source:MGI Symbol;Acc:MGI:95394] | 506,5 | 85,08 |
| ENSMUSG00000025178 | Pi4k2a | phosphatidylinositol 4-kinase type 2 alpha [Source:MGI Symbol;Acc:MGI:1934031] | 506,1 | 96,08 |
| ENSMUSG00000057649 | Brd9 | bromodomain containing 9 [Source:MGI Symbol;Acc:MGI:2145317] | 505,9 | 99,76 |
| ENSMUSG00000027163 | Commd9 | COMM domain containing 9 [Source:MGI Symbol;Acc:MGI:1923751] | 505,5 | 100,00 |
| ENSMUSG00000060733 | Ipmk | inositol polyphosphate multikinase [Source:MGI Symbol;Acc:MGI:1916968] | 505,1 | 90,59 |
| ENSMUSG00000021823 | Vcl | vinculin [Source:MGI Symbol;Acc:MGI:98927] | 505,0 | 99,56 |
| ENSMUSG00000026615 | Eprs | glutamyl-prolyl-tRNA synthetase [Source:MGI Symbol;Acc:MGI:97838] | 505,0 | 99,67 |
| ENSMUSG00000007039 | Ddah2 | dimethylarginine dimethylaminohydrolase 2 [Source:MGI Symbol;Acc:MGI:1859016] | 504,8 | 98,04 |
| ENSMUSG00000020076 | Ddx50 | DEAD (Asp-Glu-Ala-Asp) box polypeptide 50 [Source:MGI Symbol;Acc:MGI:2182303] | 504,8 | 100,00 |
| ENSMUSG00000020484 | Xbp1 | X-box binding protein 1 [Source:MGI Symbol;Acc:MGI:98970] | 504,8 | 76,30 |
| ENSMUSG00000049076 | Acap2 | ArfGAP with coiled-coil, ankyrin repeat and PH domains 2 [Source:MGI Symbol;Acc:MGI:1925868] | 504,6 | 98,56 |
| ENSMUSG00000026353 | Ubxn4 | UBX domain protein 4 [Source:MGI Symbol;Acc:MGI:1915062] | 504,3 | 96,05 |
| ENSMUSG00000047879 | Usp14 | ubiquitin specific peptidase 14 [Source:MGI Symbol;Acc:MGI:1928898] | 504,1 | 98,86 |
| ENSMUSG00000067873 | Htatsf1 | HIV TAT specific factor 1 [Source:MGI Symbol;Acc:MGI:1919709] | 504,1 | 99,64 |
| ENSMUSG00000032602 | Slc25a20 | arrier family 25 (mitochondrial carnitine/acylcarnitine translocase), member 20 [Source:MGI Symbol;Acc:MGI:1 | 503,7 | 100,00 |
| ENSMUSG00000031432 | Prps1 | phosphoribosyl pyrophosphate synthetase 1 [Source:MGI Symbol;Acc:MGI:97775] | 503,7 | 98,63 |
| ENSMUSG00000025240 | Sacm1l | SAC1 (suppressor of actin mutations 1, homolog)-like (S. cerevisiae) [Source:MGI Symbol;Acc:MGI:1933169] | 503,7 | 99,93 |
| ENSMUSG00000028394 | Pole3 | polymerase (DNA directed), epsilon 3 (p17 subunit) [Source:MGI Symbol;Acc:MGI:1933378] | 503,7 | 99,82 |
| ENSMUSG00000022940 | Pigp | phosphatidylinositol glycan anchor biosynthesis, class P [Source:MGI Symbol;Acc:MGI:1860433] | 503,6 | 93,46 |
| ENSMUSG00000031875 | Cmtm3 | CKLF-like MARVEL transmembrane domain containing 3 [Source:MGI Symbol;Acc:MGI:2447162] | 503,5 | 98,16 |
| ENSMUSG00000061838 | Suclg2 | succinate-Coenzyme A ligase, GDP-forming, beta subunit [Source:MGI Symbol;Acc:MGI:1306824] | 503,0 | 72,70 |
| ENSMUSG00000047084 | Ngrn | neugrin, neurite outgrowth associated [Source:MGI Symbol;Acc:MGI:1933212] | 503,0 | 99,97 |
| ENSMUSG00000025477 | Inpp5a | inositol polyphosphate-5-phosphatase A [Source:MGI Symbol;Acc:MGI:2686961] | 502,8 | 91,04 |
| ENSMUSG00000021951 | N6amt2 | N-6 adenine-specific DNA methyltransferase 2 (putative) [Source:MGI Symbol;Acc:MGI:1915293] | 502,2 | 100,00 |
| ENSMUSG00000035469 | Rcbtb1 | chromosome condensation (RCC1) and BTB (POZ) domain containing protein 1 [Source:MGI Symbol;Acc:M | 501,8 | 92,93 |
| ENSMUSG00000023951 | Vegfa | vascular endothelial growth factor A [Source:MGI Symbol;Acc:MGI:103178] | 501,7 | 98,52 |
| ENSMUSG00000020780 | Srp68 | signal recognition particle 68 [Source:MGI Symbol;Acc:MGI:1917447] | 501,5 | 96,61 |
| ENSMUSG00000027131 | Emc4 | ER membrane protein complex subunit 4 [Source:MGI Symbol;Acc:MGI:1915282] | 501,5 | 85,66 |
| ENSMUSG00000019738 | Polr2i | polymerase (RNA) II (DNA directed) polypeptide I [Source:MGI Symbol;Acc:MGI:1917170] | 501,3 | 95,85 |
| ENSMUSG00000061360 | Phf5a | PHD finger protein 5A [Source:MGI Symbol;Acc:MGI:2156864] | 501,3 | 100,00 |
| ENSMUSG00000028926 | Cdk14 | cyclin-dependent kinase 14 [Source:MGI Symbol;Acc:MGI:894318] | 500,7 | 89,83 |
| ENSMUSG00000038776 | Ephx1 | epoxide hydrolase 1, microsomal [Source:MGI Symbol;Acc:MGI:95405] | 500,4 | 87,71 |
| ENSMUSG00000050552 | Lamtor4 | late endosomal/lysosomal adaptor, MAPK and MTOR activator 4 [Source:MGI Symbol;Acc:MGI:1913346] | 500,3 | 73,51 |
| ENSMUSG00000021779 | Thrb | thyroid hormone receptor beta [Source:MGI Symbol;Acc:MGI:98743] | 500,2 | 91,36 |
| ENSMUSG00000040964 | Arhgef10l | Rho guanine nucleotide exchange factor (GEF) 10-like [Source:MGI Symbol;Acc:MGI:1920004] | 500,2 | 82,40 |
| ENSMUSG00000036138 | Acaa1a | acetyl-Coenzyme A acyltransferase 1A [Source:MGI Symbol;Acc:MGI:2148491] | 500,1 | 87,43 |
| ENSMUSG00000042605 | Atxn2 | ataxin 2 [Source:MGI Symbol;Acc:MGI:1277223] | 500,0 | 94,81 |

Table S2.4

| Id | baseMean Read | Mean read Astrocyte | Mean read PAP | log2FoldChange | padj | Associated Gene Name | Description | Mean Astrocyte Exon Coverage | Mean PAP exon Coverage |
|---|---|---|---|---|---|---|---|---|---|
| ENSMUSG00000000184 | 21813,72757 | 905,0 | 39873,2 | -5,456549541 | 2,9364E-09 | Ccnd2 | cyclin D2 [Source:MGI Symbol;Acc:MGI:88314] | 94,55 | 93,2 |
| ENSMUSG00000050708 | 25890,21632 | 1603,8 | 50106,7 | -4,972222268 | 9,99202E-09 | Ftl1 | ferritin light polypeptide 1 [Source:MGI Symbol;Acc:MGI:95589] | 90,71 | 87,1 |
| ENSMUSG00000020184 | 10069,65634 | 700,9 | 19108,8 | -4,670599456 | 9,25997E-08 | Mdm2 | transformed mouse 3T3 cell double minute 2 [Source:MGI Symbol;Acc:MGI:96952] | 97,72 | 97,5 |
| ENSMUSG00000020372 | 6053,641255 | 583,7 | 12434,4 | -4,398034559 | 2,96566E-09 | Gnb2l1 | guanine nucleotide binding protein (G protein), beta polypeptide 2 like 1 [Source:MGI Symbol;Acc:MGI:101849] | 87,70 | 82,7 |
| ENSMUSG00000028211 | 6452,519215 | 534,2 | 10260,5 | -4,248306982 | 0,000303492 | Trp53inp1 | transformation related protein 53 inducible nuclear protein 1 [Source:MGI Symbol;Acc:MGI:1926609] | 99,65 | 98,6 |
| ENSMUSG00000007892 | 9021,236542 | 992,1 | 16082,4 | -3,996641756 | 1,10834E-06 | Rplp1 | ribosomal protein, large, P1 [Source:MGI Symbol;Acc:MGI:1927099] | 100,00 | 100,0 |
| ENSMUSG00000057113 | 6483,911835 | 778,3 | 12048,4 | -3,895958317 | 1,76545E-07 | Npm1 | nucleophosmin 1 [Source:MGI Symbol;Acc:MGI:106184] | 92,12 | 80,7 |
| ENSMUSG00000037742 | 24476,69821 | 3129,0 | 46682,3 | -3,873586275 | 4,43069E-07 | Eef1a1 | eukaryotic translation elongation factor 1 alpha 1 [Source:MGI Symbol;Acc:MGI:1096881] | 80,10 | 71,1 |
| ENSMUSG00000020460 | 4762,165298 | 623,7 | 9248,6 | -3,860575047 | 5,47199E-07 | Rps27a | ribosomal protein S27A [Source:MGI Symbol;Acc:MGI:1925544] | 72,54 | 71,7 |
| ENSMUSG00000034994 | 10952,36374 | 1370,5 | 19645,5 | -3,820031026 | 4,31989E-07 | Eef2 | eukaryotic translation elongation factor 2 [Source:MGI Symbol;Acc:MGI:95288] | 99,49 | 99,5 |
| ENSMUSG00000032399 | 16817,97492 | 2220,7 | 31493,7 | -3,796835035 | 2,29172E-07 | Rpl4 | ribosomal protein L4 [Source:MGI Symbol;Acc:MGI:1915141] | 100,00 | 98,3 |
| ENSMUSG00000067274 | 6635,247618 | 904,4 | 12313,4 | -3,752169344 | 1,50113E-07 | Rplp0 | ribosomal protein, large, P0 [Source:MGI Symbol;Acc:MGI:1927636] | 77,99 | 77,0 |
| ENSMUSG00000003970 | 7106,387782 | 979,0 | 13267,5 | -3,742325153 | 2,68524E-08 | Rpl8 | ribosomal protein L8 [Source:MGI Symbol;Acc:MGI:1350927] | 99,54 | 99,0 |
| ENSMUSG00000102349 | 5422,44528 | 811,0 | 10992,7 | -3,577752685 | 0,000238304 | Gm37376 | predicted gene, 37376 [Source:MGI Symbol;Acc:MGI:5610604] | 100,00 | 100,0 |
| ENSMUSG00000071415 | 4645,282686 | 699,7 | 8521,2 | -3,560964177 | 7,22388E-06 | Rpl23 | ribosomal protein L23 [Source:MGI Symbol;Acc:MGI:1929455] | 93,26 | 88,6 |
| ENSMUSG00000074129 | 6108,031172 | 1005,4 | 11325,3 | -3,45260639 | 7,17095E-06 | Rpl13a | ribosomal protein L13A [Source:MGI Symbol;Acc:MGI:1351455] | 100,00 | 99,4 |
| ENSMUSG00000030744 | 7066,868887 | 1198,7 | 12966,5 | -3,429732311 | 1,55505E-06 | Rps3 | ribosomal protein S3 [Source:MGI Symbol;Acc:MGI:1350917] | 95,25 | 95,2 |
| ENSMUSG00000025794 | 6977,535508 | 1218,9 | 12940,7 | -3,407211413 | 5,25356E-06 | Rpl14 | ribosomal protein L14 [Source:MGI Symbol;Acc:MGI:1914365] | 96,36 | 96,5 |
| ENSMUSG00000012848 | 4655,01696 | 736,4 | 7638,9 | -3,35613699 | 3,46276E-05 | Rps5 | ribosomal protein S5 [Source:MGI Symbol;Acc:MGI:1097682] | 88,60 | 87,9 |
| ENSMUSG00000059070 | 3805,434615 | 669,5 | 6651,3 | -3,309531537 | 4,04787E-06 | Rpl18 | ribosomal protein L18 [Source:MGI Symbol;Acc:MGI:98003] | 95,09 | 88,8 |
| ENSMUSG00000043131 | 2915,651384 | 594,8 | 5588,2 | -3,18293955 | 3,02051E-06 | Mob1a | MOB kinase activator 1A [Source:MGI Symbol;Acc:MGI:2442631] | 98,10 | 96,1 |
| ENSMUSG00000057863 | 2897,925885 | 574,3 | 5232,3 | -3,170735729 | 0,000119227 | Rpl36 | ribosomal protein L36 [Source:MGI Symbol;Acc:MGI:1860603] | 87,87 | 88,0 |
| ENSMUSG00000046330 | 3650,847536 | 757,0 | 6688,8 | -3,123765101 | 0,000132751 | Rpl37a | ribosomal protein L37a [Source:MGI Symbol;Acc:MGI:98068] | 100,00 | 99,6 |
| ENSMUSG00000031320 | 2902,037563 | 601,0 | 5246,0 | -3,092556917 | 1,21468E-05 | Rps4x | ribosomal protein S4, X-linked [Source:MGI Symbol;Acc:MGI:98158] | 91,73 | 86,3 |
| ENSMUSG00000038274 | 2436,760178 | 522,7 | 4377,7 | -3,04399687 | 3,74958E-05 | Fau | l-Biskis-Reilly murine sarcoma virus (FBR-MuSV) ubiquitously expressed (fox derived) [Source:MGI Symbol;Acc:MGI:10 | 83,35 | 85,6 |
| ENSMUSG00000047675 | 2653,753854 | 569,4 | 4826,7 | -3,023021225 | 3,67112E-05 | Rps8 | ribosomal protein S8 [Source:MGI Symbol;Acc:MGI:98166] | 91,74 | 80,5 |
| ENSMUSG00000018583 | 2353,155654 | 531,2 | 4112,1 | -2,993425709 | 1,23964E-05 | G3bp1 | GTPase activating protein (SH3 domain) binding protein 1 [Source:MGI Symbol;Acc:MGI:1351465] | 86,70 | 80,1 |
| ENSMUSG00000038612 | 2908,13207 | 660,4 | 4907,4 | -2,865627693 | 7,88054E-05 | Mcl1 | myeloid cell leukemia sequence 1 [Source:MGI Symbol;Acc:MGI:101769] | 99,96 | 99,0 |
| ENSMUSG00000017404 | 3831,906176 | 933,3 | 6821,4 | -2,846447961 | 0,000164224 | Rpl19 | ribosomal protein L19 [Source:MGI Symbol;Acc:MGI:98020] | 94,36 | 85,4 |
| ENSMUSG00000061787 | 2309,568286 | 601,5 | 4096,3 | -2,757215888 | 0,000565985 | Rps17 | ribosomal protein S17 [Source:MGI Symbol;Acc:MGI:1309526] | 96,29 | 91,0 |
| ENSMUSG00000032050 | 2231,358465 | 523,8 | 3631,2 | -2,728851539 | 5,51281E-05 | Rdx | radixin [Source:MGI Symbol;Acc:MGI:97887] | 91,37 | 72,8 |
| ENSMUSG00000049517 | 2022,520846 | 531,2 | 3493,2 | -2,712308579 | 0,000154358 | Rps23 | ribosomal protein S23 [Source:MGI Symbol;Acc:MGI:1913725] | 90,08 | 86,6 |
| ENSMUSG00000009927 | 2954,88596 | 814,2 | 5100,5 | -2,63845456 | 0,000492468 | Rps25 | ribosomal protein S25 [Source:MGI Symbol;Acc:MGI:1922867] | 89,07 | 80,3 |
| ENSMUSG00000003429 | 4102,407704 | 1154,4 | 6890,2 | -2,554113086 | 0,000910617 | Rps11 | ribosomal protein S11 [Source:MGI Symbol;Acc:MGI:1351329] | 83,39 | 83,4 |
| ENSMUSG00000053565 | 1779,189802 | 561,1 | 3282,0 | -2,546552653 | 0,000362015 | Eif3k | eukaryotic translation initiation factor 3, subunit K [Source:MGI Symbol;Acc:MGI:1921080] | 84,40 | 71,6 |
| ENSMUSG00000068039 | 3732,511154 | 1040,4 | 5987,1 | -2,530588177 | 0,001128767 | Tcp1 | t-complex protein 1 [Source:MGI Symbol;Acc:MGI:98535] | 93,60 | 82,8 |
| ENSMUSG00000034892 | 4204,406769 | 1244,4 | 7113,4 | -2,481884087 | 0,002985684 | Rps29 | ribosomal protein S29 [Source:MGI Symbol;Acc:MGI:107681] | 93,50 | 79,7 |
| ENSMUSG00000062762 | 3127,65182 | 841,4 | 4645,9 | -2,467298274 | 0,000907964 | Ei24 | etoposide induced 2.4 mRNA [Source:MGI Symbol;Acc:MGI:108090] | 95,22 | 94,5 |
| ENSMUSG00000090000 | 1819,692041 | 611,1 | 3334,1 | -2,463551176 | 0,005455322 | Ier3ip1 | immediate early response 3 interacting protein 1 [Source:MGI Symbol;Acc:MGI:1913441] | 99,88 | 98,4 |
| ENSMUSG00000000740 | 4239,369353 | 1340,0 | 7298,1 | -2,439813905 | 0,000641048 | Rpl13 | ribosomal protein L13 [Source:MGI Symbol;Acc:MGI:105922] | 84,53 | 79,1 |
| ENSMUSG00000039001 | 2401,635816 | 808,1 | 4487,3 | -2,406922041 | 0,005578315 | Rps21 | ribosomal protein S21 [Source:MGI Symbol;Acc:MGI:1913731] | 99,75 | 99,2 |
| ENSMUSG00000064370 | 1269,584883 | 570,3 | 2333,8 | -2,405933538 | 0,042667317 | mt-Cytb | mitochondrially encoded cytochrome b [Source:MGI Symbol;Acc:MGI:102501] | 100,00 | 99,9 |
| ENSMUSG00000049354 | 2124,930454 | 717,1 | 4331,6 | -2,387158893 | 0,004736532 | Dcaf7 | DDB1 and CUL4 associated factor 7 [Source:MGI Symbol;Acc:MGI:1919083] | 99,02 | 86,6 |
| ENSMUSG00000026434 | 1736,777739 | 744,6 | 3922,2 | -2,375009144 | 0,00123279 | Nucks1 | nuclear casein kinase and cyclin-dependent kinase substrate 1 [Source:MGI Symbol;Acc:MGI:1934811] | 99,45 | 90,0 |
| ENSMUSG00000020362 | 1855,787994 | 612,7 | 3361,0 | -2,324602013 | 0,005326329 | Cnot6 | CCR4-NOT transcription complex, subunit 6 [Source:MGI Symbol;Acc:MGI:2144529] | 98,61 | 87,5 |
| ENSMUSG00000063457 | 2861,403994 | 959,3 | 4788,3 | -2,306708321 | 0,002039489 | Rps15 | ribosomal protein S15 [Source:MGI Symbol;Acc:MGI:98117] | 90,16 | 86,2 |
| ENSMUSG00000035268 | 2064,61097 | 652,1 | 3203,0 | -2,278002248 | 0,007481539 | Pkig | protein kinase inhibitor, gamma [Source:MGI Symbol;Acc:MGI:1343086] | 86,98 | 83,1 |
| ENSMUSG00000028639 | 1856,535722 | 590,0 | 2884,8 | -2,276330921 | 0,000164224 | Ybx1 | Y box protein 1 [Source:MGI Symbol;Acc:MGI:99146] | 93,25 | 89,1 |
| ENSMUSG00000026028 | 1163,713224 | 507,8 | 2473,6 | -2,271898162 | 0,001111771 | Trak2 | trafficking protein, kinesin binding 2 [Source:MGI Symbol;Acc:MGI:1918077] | 99,22 | 81,7 |
| ENSMUSG00000001525 | 1768,974525 | 570,7 | 2857,5 | -2,233341064 | 0,010254873 | Tubb5 | tubulin, beta 5 class I [Source:MGI Symbol;Acc:MGI:107812] | 87,07 | 87,7 |
| ENSMUSG00000023944 | 5424,421096 | 1938,0 | 8787,9 | -2,209574748 | 0,004084001 | Hsp90ab1 | heat shock protein 90 alpha (cytosolic), class B member 1 [Source:MGI Symbol;Acc:MGI:96247] | 98,17 | 95,8 |
| ENSMUSG00000031985 | 2087,854959 | 745,2 | 3634,2 | -2,207332522 | 0,001323936 | Gnpat | glyceronephosphate O-acyltransferase [Source:MGI Symbol;Acc:MGI:1343460] | 99,32 | 98,5 |
| ENSMUSG00000037503 | 4965,443605 | 1901,9 | 8599,9 | -2,19304745 | 0,003347776 | Fam168b | family with sequence similarity 168, member B [Source:MGI Symbol;Acc:MGI:2448487] | 99,04 | 96,4 |
| ENSMUSG00000029621 | 1691,156089 | 582,8 | 3059,2 | -2,180372259 | 0,018081101 | Arpc1a | actin related protein 2/3 complex, subunit 1A [Source:MGI Symbol;Acc:MGI:1928896] | 91,43 | 90,4 |
| ENSMUSG00000025130 | 1421,529502 | 671,0 | 2762,3 | -2,131298414 | 0,000935031 | P4hb | prolyl 4-hydroxylase, beta polypeptide [Source:MGI Symbol;Acc:MGI:97464] | 95,99 | 86,6 |
| ENSMUSG00000018446 | 1873,587826 | 755,6 | 3348,3 | -2,128023523 | 0,004054846 | C1qbp | complement component 1, q subcomponent binding protein [Source:MGI Symbol;Acc:MGI:1194505] | 100,00 | 98,1 |
| ENSMUSG00000007739 | 1569,08876 | 588,9 | 2302,2 | -2,011873071 | 0,004307395 | Cct4 | chaperonin containing Tcp1, subunit 4 (delta) [Source:MGI Symbol;Acc:MGI:104689] | 95,53 | 77,7 |
| ENSMUSG00000032601 | 1404,717129 | 573,5 | 2315,5 | -1,999030518 | 0,024619628 | Prkar2a | protein kinase, cAMP dependent regulatory, type II alpha [Source:MGI Symbol;Acc:MGI:108025] | 95,75 | 82,8 |
| ENSMUSG00000020857 | 2408,26337 | 999,5 | 3835,6 | -1,923658836 | 0,004092103 | Nme2 | NME/NM23 nucleoside diphosphate kinase 2 [Source:MGI Symbol;Acc:MGI:97356] | 83,80 | 72,3 |
| ENSMUSG00000017639 | 1754,738925 | 766,2 | 2901,8 | -1,908235507 | 0,01949513 | Rab11fip4 | RAB11 family interacting protein 4 (class II) [Source:MGI Symbol;Acc:MGI:2442920] | 97,99 | 88,4 |
| ENSMUSG00000022234 | 1538,296136 | 652,0 | 2621,3 | -1,899305899 | 0,021568781 | Cct5 | chaperonin containing Tcp1, subunit 5 (epsilon) [Source:MGI Symbol;Acc:MGI:107185] | 95,81 | 87,6 |
| ENSMUSG00000034024 | 1584,521891 | 577,2 | 2666,9 | -1,896572479 | 0,041917199 | Cct2 | chaperonin containing Tcp1, subunit 2 (beta) [Source:MGI Symbol;Acc:MGI:107186] | 99,90 | 94,6 |
| ENSMUSG00000090733 | 1872,028442 | 755,8 | 2792,5 | -1,885613528 | 0,015750201 | Rps27 | ribosomal protein S27 [Source:MGI Symbol;Acc:MGI:1888676] | 91,16 | 74,9 |
| ENSMUSG00000026615 | 1083,440134 | 505,0 | 2064,5 | -1,876413275 | 0,035858194 | Eprs | glutamyl-prolyl-tRNA synthetase [Source:MGI Symbol;Acc:MGI:97838] | 99,67 | 87,3 |
| ENSMUSG00000056851 | 1144,737468 | 602,7 | 2190,3 | -1,853477619 | 0,003181242 | Pcbp2 | poly(rC) binding protein 2 [Source:MGI Symbol;Acc:MGI:108202] | 95,23 | 74,0 |
| ENSMUSG00000024516 | 1456,187784 | 624,3 | 2272,9 | -1,844074025 | 0,006566597 | Sec11c | SEC11 homolog C, signal peptidase complex subunit [Source:MGI Symbol;Acc:MGI:1913536] | 98,15 | 96,2 |
| ENSMUSG00000027852 | 1382,003522 | 559,6 | 2061,9 | -1,841175577 | 0,020740575 | Nras | neuroblastoma ras oncogene [Source:MGI Symbol;Acc:MGI:97376] | 92,71 | 74,4 |
| ENSMUSG00000057522 | 1919,007695 | 774,9 | 2760,1 | -1,796794735 | 0,031022712 | Spop | speckle-type POZ protein [Source:MGI Symbol;Acc:MGI:1343085] | 98,17 | 97,7 |
| ENSMUSG00000068328 | 960,0880752 | 517,4 | 1709,7 | -1,735767076 | 0,003898346 | Aup1 | ancient ubiquitous protein 1 [Source:MGI Symbol;Acc:MGI:107789] | 92,65 | 81,9 |
| ENSMUSG00000046562 | 1082,955718 | 668,7 | 2246,3 | -1,725947919 | 0,015376826 | Unc119b | unc-119 homolog B (C. elegans) [Source:MGI Symbol;Acc:MGI:2147162] | 98,84 | 97,8 |
| ENSMUSG00000040848 | 1696,488964 | 722,8 | 2607,2 | -1,709341604 | 0,009576467 | Sft2d2 | SFT2 domain containing 2 [Source:MGI Symbol;Acc:MGI:1917362] | 98,52 | 94,5 |
| ENSMUSG00000062647 | 1176,277331 | 539,1 | 1737,7 | -1,687696787 | 0,012275369 | Rpl7a | ribosomal protein L7A [Source:MGI Symbol;Acc:MGI:1353472] | 93,07 | 88,1 |
| ENSMUSG00000029538 | 1927,565427 | 894,9 | 2802,4 | -1,650396913 | 0,019292726 | Srsf9 | serine/arginine-rich splicing factor 9 [Source:MGI Symbol;Acc:MGI:104896] | 100,00 | 99,8 |
| ENSMUSG00000020484 | 1529,384883 | 504,8 | 1725,9 | -1,590412666 | 0,048661456 | Xbp1 | X-box binding protein 1 [Source:MGI Symbol;Acc:MGI:98970] | 76,30 | 81,4 |
| ENSMUSG00000026095 | 1023,117591 | 540,1 | 1634,3 | -1,585685964 | 0,042232315 | Asnsd1 | asparagine synthetase domain containing 1 [Source:MGI Symbol;Acc:MGI:1917646] | 85,81 | 83,8 |

| | | | | | | | | | |
|---|---|---|---|---|---|---|---|---|---|
| ENSMUSG00000024248 | 1001,61802 | 530,5 | 1586,2 | -1,577515157 | 0,008448967 | Cox7a2l | cytochrome c oxidase subunit VIIa polypeptide 2-like [Source:MGI Symbol;Acc:MGI:106015] | 82,33 | 71,0 |
| ENSMUSG00000024661 | 17122,44557 | 8833,9 | 25673,0 | -1,548844492 | 0,038318939 | Fth1 | ferritin heavy polypeptide 1 [Source:MGI Symbol;Acc:MGI:95588] | 100,00 | 100,0 |
| ENSMUSG00000036781 | 2500,904047 | 1245,3 | 3537,6 | -1,503845315 | 0,020374405 | Rps27l | ribosomal protein S27-like [Source:MGI Symbol;Acc:MGI:1915191] | 100,00 | 94,5 |
| ENSMUSG00000042305 | 1401,817417 | 922,6 | 2495,1 | -1,450965537 | 0,020610821 | Tmem183a | transmembrane protein 183A [Source:MGI Symbol;Acc:MGI:1914729] | 97,97 | 96,5 |
| ENSMUSG00000025381 | 1174,393307 | 724,0 | 1981,3 | -1,444115694 | 0,012772283 | Cnpy2 | canopy FGF signaling regulator 2 [Source:MGI Symbol;Acc:MGI:1928477] | 97,51 | 96,1 |
| ENSMUSG00000052397 | 3201,255426 | 1881,3 | 4642,6 | -1,339596073 | 0,036909524 | Ezr | ezrin [Source:MGI Symbol;Acc:MGI:98931] | 92,84 | 91,7 |
| ENSMUSG00000027479 | 3264,489934 | 1737,3 | 4213,5 | -1,249305238 | 0,031661078 | Mapre1 | microtubule-associated protein, RP/EB family, member 1 [Source:MGI Symbol;Acc:MGI:891995] | 82,72 | 81,2 |

Table S2.5

| Id | baseMean Read | Mean read Astrocyte | Mean read PAP | log2FoldChange | padj | Associated Gene Name | Description | Mean Astrocyte Exon Coverage | Mean PAP exon Coverage |
|---|---|---|---|---|---|---|---|---|---|
| ENSMUSG00000017390 | 27993,96595 | 45498,3 | 9857,7 | 2,442382009 | 0,039994072 | Aldoc | aldolase C, fructose-bisphosphate [Source:MGI Symbol;Acc:MGI:101863] | 100,00 | 82,3 |
| ENSMUSG00000026701 | 25807,46035 | 44118,1 | 8096,1 | 2,52666555 | 0,005912613 | Prdx6 | peroxiredoxin 6 [Source:MGI Symbol;Acc:MGI:894320] | 99,72 | 63,2 |
| ENSMUSG00000004558 | 24355,62011 | 37500,0 | 10834,1 | 2,181282147 | 0,042457815 | Ndrg2 | N-myc downstream regulated gene 2 [Source:MGI Symbol;Acc:MGI:1352498] | 100,00 | 95,8 |
| ENSMUSG00000058135 | 15538,58151 | 27393,6 | 5406,5 | 2,481091227 | 0,014154423 | Gstm1 | glutathione S-transferase, mu 1 [Source:MGI Symbol;Acc:MGI:95860] | 100,00 | 86,7 |
| ENSMUSG00000001270 | 15673,42835 | 25914,7 | 4548,4 | 2,653357783 | 0,006924761 | Ckb | creatine kinase, brain [Source:MGI Symbol;Acc:MGI:88407] | 98,92 | 95,7 |
| ENSMUSG00000004032 | 13402,09956 | 22563,9 | 4169,9 | 2,878087256 | 0,009805025 | Gstm5 | glutathione S-transferase, mu 5 [Source:MGI Symbol;Acc:MGI:1309466] | 95,99 | 81,5 |
| ENSMUSG00000001175 | 10205,77799 | 15676,0 | 5669,9 | 1,497333076 | 0,02058664 | Calm1 | calmodulin 1 [Source:MGI Symbol;Acc:MGI:88251] | 95,53 | 78,5 |
| ENSMUSG00000026421 | 9539,536171 | 15542,5 | 2631,6 | 3,069311308 | 0,005598936 | Csrp1 | cysteine and glycine-rich protein 1 [Source:MGI Symbol;Acc:MGI:88549] | 96,06 | 87,4 |
| ENSMUSG00000031517 | 10067,10059 | 15521,9 | 3318,6 | 2,185972251 | 0,038235961 | Gpm6a | glycoprotein m6a [Source:MGI Symbol;Acc:MGI:107671] | 99,47 | 83,6 |
| ENSMUSG00000031762 | 7759,039747 | 14105,0 | 4348,8 | 1,935607756 | 0,028442082 | Mt2 | metallothionein 2 [Source:MGI Symbol;Acc:MGI:97172] | 100,00 | 89,0 |
| ENSMUSG00000020644 | 7067,298447 | 13113,0 | 1346,6 | 3,658053668 | 0,032295618 | Id2 | inhibitor of DNA binding 2 [Source:MGI Symbol;Acc:MGI:96397] | 100,00 | 98,5 |
| ENSMUSG00000039533 | 7834,091637 | 12560,3 | 3170,0 | 2,130947325 | 0,025444664 | Mmd2 | monocyte to macrophage differentiation-associated 2 [Source:MGI Symbol;Acc:MGI:1922354] | 99,95 | 65,7 |
| ENSMUSG00000021750 | 7821,690492 | 11038,8 | 3003,6 | 2,793243523 | 0,041551111 | Fam107a | family with sequence similarity 107, member A [Source:MGI Symbol;Acc:MGI:3041256] | 91,94 | 77,4 |
| ENSMUSG00000045092 | 6861,182325 | 10766,9 | 2331,6 | 2,36893765 | 0,02830088 | S1pr1 | sphingosine-1-phosphate receptor 1 [Source:MGI Symbol;Acc:MGI:1096355] | 99,84 | 95,0 |
| ENSMUSG00000030246 | 6025,373907 | 10219,7 | 1961,4 | 2,566471378 | 0,018209447 | Ldhb | lactate dehydrogenase B [Source:MGI Symbol;Acc:MGI:96763] | 89,81 | 80,7 |
| ENSMUSG00000000346 | 6265,36545 | 10160,4 | 2772,7 | 2,114820033 | 0,020067293 | Dazap2 | DAZ associated protein 2 [Source:MGI Symbol;Acc:MGI:1344344] | 100,00 | 99,8 |
| ENSMUSG00000022323 | 4453,483543 | 9257,7 | 1530,0 | 2,965120876 | 0,015512448 | Hrsp12 | heat-responsive protein 12 [Source:MGI Symbol;Acc:MGI:1095401] | 99,87 | 98,9 |
| ENSMUSG00000030695 | 6852,655302 | 9155,1 | 4096,1 | 1,156792513 | 0,028422025 | Aldoa | aldolase A, fructose-bisphosphate [Source:MGI Symbol;Acc:MGI:87994] | 78,40 | 54,6 |
| ENSMUSG00000028691 | 5370,474218 | 8949,8 | 2330,1 | 1,890915356 | 0,009128616 | Prdx1 | peroxiredoxin 1 [Source:MGI Symbol;Acc:MGI:99523] | 94,53 | 45,4 |
| ENSMUSG00000024654 | 5574,076828 | 8843,8 | 2613,1 | 2,065770252 | 0,031924952 | Asrgl1 | asparaginase like 1 [Source:MGI Symbol;Acc:MGI:1913764] | 99,91 | 81,3 |
| ENSMUSG00000026473 | 5271,908081 | 8817,2 | 1752,7 | 2,864783904 | 0,018591859 | Glul | glutamate-ammonia ligase (glutamine synthetase) [Source:MGI Symbol;Acc:MGI:95739] | 99,68 | 70,1 |
| ENSMUSG00000036309 | 5370,101594 | 8601,9 | 2451,8 | 1,842311099 | 0,013575227 | Skp1a | S-phase kinase-associated protein 1A [Source:MGI Symbol;Acc:MGI:103575] | 93,96 | 85,5 |
| ENSMUSG00000031633 | 5994,209438 | 8533,9 | 3340,3 | 1,351773778 | 0,022655966 | Slc25a4 | nily 25 (mitochondrial carrier, adenine nucleotide translocator), member 4 [Source:MGI Symbol;Ac | 88,76 | 68,1 |
| ENSMUSG00000025666 | 5058,207393 | 8454,8 | 1975,7 | 3,902021586 | 0,034693621 | Tmem47 | transmembrane protein 47 [Source:MGI Symbol;Acc:MGI:2177570] | 100,00 | 66,3 |
| ENSMUSG00000028517 | 5294,244796 | 8309,9 | 1852,3 | 2,163435082 | 0,038475475 | Plpp3 | phospholipid phosphatase 3 [Source:MGI Symbol;Acc:MGI:1915166] | 100,00 | 96,4 |
| ENSMUSG00000031431 | 3733,547352 | 8167,4 | 579,5 | 4,720736378 | 2,59185E-05 | Tsc22d3 | TSC22 domain family, member 3 [Source:MGI Symbol;Acc:MGI:1196284] | 90,35 | 57,3 |
| ENSMUSG00000030120 | 5066,190039 | 7983,9 | 2097,3 | 2,213138383 | 0,011883954 | Mlf2 | myeloid leukemia factor 2 [Source:MGI Symbol;Acc:MGI:1353554] | 98,76 | 88,4 |
| ENSMUSG00000021379 | 4031,702164 | 7929,5 | 465,2 | 4,43826935 | 0,000420757 | Id4 | inhibitor of DNA binding 4 [Source:MGI Symbol;Acc:MGI:99414] | 100,00 | 77,3 |
| ENSMUSG00000027562 | 5040,418104 | 7637,9 | 1579,5 | 2,587544728 | 0,033825651 | Car2 | carbonic anhydrase 2 [Source:MGI Symbol;Acc:MGI:88269] | 100,00 | 96,6 |
| ENSMUSG00000040997 | 4139,711407 | 7373,1 | 1172,5 | 2,898001054 | 0,010278719 | Abhd4 | abhydrolase domain containing 4 [Source:MGI Symbol;Acc:MGI:1915938] | 92,71 | 78,9 |
| ENSMUSG00000030057 | 5128,913023 | 6781,9 | 2662,1 | 1,384311052 | 0,012526629 | Cnbp | cellular nucleic acid binding protein [Source:MGI Symbol;Acc:MGI:88431] | 96,35 | 71,5 |
| ENSMUSG00000001833 | 3636,853996 | 6315,9 | 938,9 | 3,492282414 | 0,001270433 | Sept7 | septin 7 [Source:MGI Symbol;Acc:MGI:1335094] | 98,20 | 73,2 |
| ENSMUSG00000056201 | 4225,503326 | 6077,7 | 2100,2 | 1,572990399 | 0,011616912 | Cfl1 | cofilin 1, non-muscle [Source:MGI Symbol;Acc:MGI:101757] | 100,00 | 99,5 |
| ENSMUSG00000001847 | 3898,788056 | 6064,7 | 1874,0 | 1,714536245 | 0,035066767 | Rac1 | RAS-related C3 botulinum substrate 1 [Source:MGI Symbol;Acc:MGI:97845] | 76,23 | 53,6 |
| ENSMUSG00000049422 | 3675,660824 | 5927,8 | 1328,1 | 2,559704302 | 0,034635525 | Chchd10 | coiled-coil-helix-coiled-coil-helix domain containing 10 [Source:MGI Symbol;Acc:MGI:2143558] | 85,59 | 68,5 |
| ENSMUSG00000074457 | 3310,028881 | 5645,4 | 874,7 | 3,223754098 | 0,014747957 | S100a16 | S100 calcium binding protein A16 [Source:MGI Symbol;Acc:MGI:1915110] | 98,95 | 71,5 |
| ENSMUSG00000001891 | 3376,807116 | 5626,4 | 1082,7 | 2,463630929 | 0,00551827 | Ugp2 | UDP-glucose pyrophosphorylase 2 [Source:MGI Symbol;Acc:MGI:2183447] | 100,00 | 89,3 |
| ENSMUSG00000019804 | 3374,039649 | 5299,6 | 1889,2 | 1,884236659 | 0,045031522 | Snx3 | sorting nexin 3 [Source:MGI Symbol;Acc:MGI:1860188] | 99,53 | 88,1 |
| ENSMUSG00000018102 | 3090,352678 | 5158,8 | 742,9 | 3,284319456 | 0,007337311 | Hist1h2bc | histone cluster 1, H2bc [Source:MGI Symbol;Acc:MGI:1915274] | 71,78 | 65,9 |
| ENSMUSG00000006699 | 3239,697166 | 5008,4 | 1725,7 | 1,543907669 | 0,018427617 | Cdc42 | cell division cycle 42 [Source:MGI Symbol;Acc:MGI:106211] | 99,92 | 71,9 |
| ENSMUSG00000030647 | 3334,965764 | 4999,8 | 1664,9 | 1,781381849 | 0,026582956 | Ndufc2 | .DH dehydrogenase (ubiquinone) 1, subcomplex unknown, 2 [Source:MGI Symbol;Acc:MGI:13443 | 93,63 | 86,1 |
| ENSMUSG00000047187 | 2986,571069 | 4964,9 | 950,8 | 2,406560618 | 0,008355771 | Rab2a | RAB2A, member RAS oncogene family [Source:MGI Symbol;Acc:MGI:1928750] | 97,97 | 92,0 |
| ENSMUSG00000029030 | 3162,941696 | 4771,5 | 1449,4 | 2,269373971 | 0,013831112 | Tprgl | transformation related protein 63 regulated like [Source:MGI Symbol;Acc:MGI:1915058] | 100,00 | 99,8 |
| ENSMUSG00000033059 | 3110,054318 | 4740,9 | 1500,3 | 1,714160302 | 0,025988092 | Pygb | brain glycogen phosphorylase [Source:MGI Symbol;Acc:MGI:97828] | 100,00 | 97,3 |
| ENSMUSG00000020733 | 2595,653112 | 4739,3 | 385,4 | 3,981956606 | 3,84838E-05 | Slc9a3r1 | rrier family 9 (sodium/hydrogen exchanger), member 3 regulator 1 [Source:MGI Symbol;Acc:MGI: | 100,00 | 85,4 |
| ENSMUSG00000022658 | 2871,127137 | 4602,4 | 1041,8 | 2,486328788 | 0,022252444 | Tagln3 | transgelin 3 [Source:MGI Symbol;Acc:MGI:1926784] | 100,00 | 82,5 |
| ENSMUSG00000053931 | 2555,568949 | 4571,4 | 569,3 | 3,113843633 | 0,01918874 | Cnn3 | calponin 3, acidic [Source:MGI Symbol;Acc:MGI:1919244] | 91,83 | 82,0 |
| ENSMUSG00000056515 | 2959,495762 | 4392,7 | 1427,3 | 2,702277605 | 0,032958225 | Rab31 | RAB31, member RAS oncogene family [Source:MGI Symbol;Acc:MGI:1914603] | 99,44 | 69,4 |
| ENSMUSG00000041556 | 2597,615743 | 4362,4 | 923,1 | 2,510377292 | 0,027415843 | Fbxo2 | F-box protein 2 [Source:MGI Symbol;Acc:MGI:2446216] | 100,00 | 98,9 |
| ENSMUSG00000020333 | 2718,540867 | 4355,3 | 398,2 | 3,48703406 | 0,00138233 | Acsl6 | acyl-CoA synthetase long-chain family member 6 [Source:MGI Symbol;Acc:MGI:894291] | 94,11 | 22,7 |
| ENSMUSG00000073639 | 2312,973779 | 4276,3 | 697,2 | 3,048622156 | 0,005008091 | Rab18 | RAB18, member RAS oncogene family [Source:MGI Symbol;Acc:MGI:102790] | 100,00 | 74,5 |
| ENSMUSG00000007872 | 2402,980681 | 4238,2 | 567,3 | 3,163569043 | 0,035908736 | Id3 | inhibitor of DNA binding 3 [Source:MGI Symbol;Acc:MGI:96398] | 84,74 | 75,2 |
| ENSMUSG00000030310 | 2505,336743 | 4233,9 | 607,6 | 3,002877566 | 0,007479235 | Slc6a1 | carrier family 6 (neurotransmitter transporter, GABA), member 1 [Source:MGI Symbol;Acc:MGI:9 | 99,84 | 74,0 |
| ENSMUSG00000039197 | 2391,377095 | 4193,7 | 879,4 | 2,265107272 | 0,028021437 | Adk | adenosine kinase [Source:MGI Symbol;Acc:MGI:87930] | 99,68 | 95,8 |
| ENSMUSG00000024099 | 2884,777939 | 4121,3 | 1704,8 | 1,302587233 | 0,038303911 | Ndufv2 | NADH dehydrogenase (ubiquinone) flavoprotein 2 [Source:MGI Symbol;Acc:MGI:1920150] | 83,33 | 56,7 |
| ENSMUSG00000018567 | 2797,049007 | 4116,9 | 1521,0 | 1,496759153 | 0,02894751 | Gabarap | gamma-aminobutyric acid receptor associated protein [Source:MGI Symbol;Acc:MGI:1861742] | 86,43 | 63,7 |
| ENSMUSG00000075706 | 2789,889853 | 4102,1 | 1129,6 | 2,142257017 | 0,024379257 | Gpx4 | glutathione peroxidase 4 [Source:MGI Symbol;Acc:MGI:104767] | 89,05 | 50,5 |
| ENSMUSG00000027805 | 2694,994484 | 3989,0 | 1130,1 | 2,262200441 | 0,022794374 | Pfn2 | profilin 2 [Source:MGI Symbol;Acc:MGI:97550] | 71,96 | 49,6 |
| ENSMUSG00000079477 | 2551,660479 | 3970,3 | 1164,3 | 1,882824329 | 0,004893845 | Rab7 | RAB7, member RAS oncogene family [Source:MGI Symbol;Acc:MGI:105068] | 83,23 | 76,2 |
| ENSMUSG00000035133 | 2617,959338 | 3949,3 | 1097,4 | 1,978087273 | 0,043707608 | Arhgap5 | Rho GTPase activating protein 5 [Source:MGI Symbol;Acc:MGI:1332637] | 100,00 | 74,7 |
| ENSMUSG00000001687 | 2450,620557 | 3740,5 | 600,0 | 2,872811188 | 0,001753896 | Ubl3 | ubiquitin-like 3 [Source:MGI Symbol;Acc:MGI:1344373] | 99,92 | 87,9 |
| ENSMUSG00000057880 | 2713,525491 | 3680,1 | 1936,4 | 2,725275362 | 0,049616998 | Abat | 4-aminobutyrate aminotransferase [Source:MGI Symbol;Acc:MGI:2443582] | 95,36 | 57,0 |
| ENSMUSG00000025351 | 2573,716586 | 3643,7 | 807,8 | 2,620052873 | 0,046266669 | Cd63 | CD63 antigen [Source:MGI Symbol;Acc:MGI:99529] | 99,81 | 84,7 |
| ENSMUSG00000021877 | 2483,707402 | 3635,1 | 1122,5 | 1,688334094 | 0,026989554 | Arf4 | ADP-ribosylation factor 4 [Source:MGI Symbol;Acc:MGI:99433] | 99,51 | 84,6 |
| ENSMUSG00000022474 | 2290,444281 | 3622,5 | 643,3 | 2,501701865 | 0,001903062 | Pmm1 | phosphomannomutase 1 [Source:MGI Symbol;Acc:MGI:1353418] | 96,82 | 94,8 |
| ENSMUSG00000030605 | 2679,837537 | 3595,0 | 906,0 | 2,587940381 | 0,039595924 | Mfge8 | milk fat globule-EGF factor 8 protein [Source:MGI Symbol;Acc:MGI:102768] | 97,62 | 81,3 |
| ENSMUSG00000024646 | 2228,291289 | 3590,4 | 1172,4 | 1,74365462 | 0,032220951 | Cyb5a | cytochrome b5 type A (microsomal) [Source:MGI Symbol;Acc:MGI:1926952] | 72,23 | 57,7 |
| ENSMUSG00000021238 | 1986,693012 | 3555,3 | 1,9 | 11,20646142 | 3,89286E-06 | Aldh6a1 | aldehyde dehydrogenase family 6, subfamily A1 [Source:MGI Symbol;Acc:MGI:1915077] | 100,00 | 1,1 |

| | | | | | | | | | |
|---|---|---|---|---|---|---|---|---|---|
| ENSMUSG00000008035 | 2010,53251 | 3547,8 | 525,3 | 2,861309636 | 0,040314285 | Mid1ip1 | interacting protein 1 (gastrulation specific G12-like (zebrafish)) [Source:MGI Symbol;Acc:MGI:191 | 99,82 | 92,7 |
| ENSMUSG00000020894 | 1871,688246 | 3441,5 | 652,1 | 2,968334165 | 0,037808165 | Vamp2 | vesicle-associated membrane protein 2 [Source:MGI Symbol;Acc:MGI:1313277] | 93,67 | 66,3 |
| ENSMUSG00000004056 | 2029,523432 | 3419,9 | 497,4 | 3,056415922 | 0,002796786 | Akt2 | thymoma viral proto-oncogene 2 [Source:MGI Symbol;Acc:MGI:104874] | 86,94 | 35,6 |
| ENSMUSG00000027351 | 2235,347083 | 3418,9 | 746,8 | 10,96034909 | 6,54566E-05 | Spred1 | sprouty protein with EVH-1 domain 1, related sequence [Source:MGI Symbol;Acc:MGI:2150016] | 98,93 | 28,2 |
| ENSMUSG00000027273 | 2212,826448 | 3297,9 | 637,5 | 2,542920331 | 0,001885975 | Snap25 | synaptosomal-associated protein 25 [Source:MGI Symbol;Acc:MGI:98331] | 99,91 | 90,0 |
| ENSMUSG00000033174 | 2261,334726 | 3294,5 | 821,7 | 2,087457622 | 0,022729402 | Mgll | monoglyceride lipase [Source:MGI Symbol;Acc:MGI:1346042] | 88,24 | 48,8 |
| ENSMUSG00000009863 | 2311,688329 | 3260,2 | 982,6 | 1,750697322 | 0,010559107 | Sdhb | ıccinate dehydrogenase complex, subunit B, iron sulfur (Ip) [Source:MGI Symbol;Acc:MGI:191493 | 99,42 | 87,3 |
| ENSMUSG00000040759 | 2169,567804 | 3259,6 | 712,5 | 2,734264431 | 0,024671043 | Cmtm5 | CKLF-like MARVEL transmembrane domain containing 5 [Source:MGI Symbol;Acc:MGI:2447164 | 100,00 | 97,0 |
| ENSMUSG00000040048 | 2137,904751 | 3248,3 | 1034,9 | 1,702389882 | 0,017208103 | Ndufb10 | IADH dehydrogenase (ubiquinone) 1 beta subcomplex, 10 [Source:MGI Symbol;Acc:MGI:191559 | 98,76 | 87,5 |
| ENSMUSG00000015837 | 1934,48964 | 3246,5 | 934,4 | 1,807555197 | 0,0291501 | Sqstm1 | sequestosome 1 [Source:MGI Symbol;Acc:MGI:107931] | 92,89 | 75,4 |
| ENSMUSG00000004071 | 2050,891936 | 3245,1 | 730,9 | 2,164359026 | 0,000211054 | Cdip1 | cell death inducing Trp53 target 1 [Source:MGI Symbol;Acc:MGI:1913876] | 97,33 | 65,7 |
| ENSMUSG00000022477 | 2393,048746 | 3240,0 | 1001,4 | 1,750042761 | 0,016615332 | Aco2 | aconitase 2, mitochondrial [Source:MGI Symbol;Acc:MGI:87880] | 97,36 | 89,9 |
| ENSMUSG00000036402 | 2064,768853 | 3228,4 | 742,3 | 3,071097663 | 0,032017121 | Gng12 | guanine nucleotide binding protein (G protein), gamma 12 [Source:MGI Symbol;Acc:MGI:1336171 | 98,48 | 41,0 |
| ENSMUSG00000028982 | 1716,50645 | 3218,5 | 454,7 | 2,87780834 | 0,003532165 | Slc25a33 | solute carrier family 25, member 33 [Source:MGI Symbol;Acc:MGI:1917806] | 100,00 | 80,0 |
| ENSMUSG00000020149 | 2176,358434 | 3214,8 | 910,7 | 1,8555957 | 0,02655426 | Rab1a | RAB1A, member RAS oncogene family [Source:MGI Symbol;Acc:MGI:97842] | 98,51 | 76,1 |
| ENSMUSG00000031748 | 1880,105392 | 3210,9 | 662,7 | 2,490622748 | 0,01949513 | Gnao1 | guanine nucleotide binding protein, alpha O [Source:MGI Symbol;Acc:MGI:95775] | 98,13 | 58,3 |
| ENSMUSG00000004902 | 1954,037323 | 3148,9 | 339,5 | 3,368045064 | 0,001512458 | Slc25a18 | solute carrier family 25 (mitochondrial carrier), member 18 [Source:MGI Symbol;Acc:MGI:1919053 | 100,00 | 80,6 |
| ENSMUSG00000052738 | 1952,195315 | 3010,1 | 854,5 | 1,930174958 | 0,019361857 | Suclg1 | succinate-CoA ligase, GDP-forming, alpha subunit [Source:MGI Symbol;Acc:MGI:1927234] | 89,20 | 80,1 |
| ENSMUSG00000006356 | 1712,865211 | 3009,7 | 281,2 | 3,625101782 | 0,012282057 | Crip2 | cysteine rich protein 2 [Source:MGI Symbol;Acc:MGI:1915587] | 93,73 | 67,8 |
| ENSMUSG00000019923 | 2037,42836 | 3006,5 | 1072,7 | 1,535743182 | 0,012214782 | Zwint | ZW10 interactor [Source:MGI Symbol;Acc:MGI:1289227] | 70,55 | 46,6 |
| ENSMUSG00000022678 | 1718,410152 | 3006,1 | 727,3 | 2,202016842 | 0,007525357 | Nde1 | nuclear distribution gene E homolog 1 (A nidulans) [Source:MGI Symbol;Acc:MGI:1914453] | 100,00 | 85,4 |
| ENSMUSG00000028675 | 1496,90851 | 2993,6 | 372,1 | 4,37713914 | 0,03340453 | Pnrc2 | proline-rich nuclear receptor coactivator 2 [Source:MGI Symbol;Acc:MGI:106512] | 100,00 | 54,5 |
| ENSMUSG00000031924 | 2188,954268 | 2963,3 | 959,0 | 2,480193552 | 0,02185231 | Cyb5b | cytochrome b5 type B [Source:MGI Symbol;Acc:MGI:1913677] | 99,98 | 56,2 |
| ENSMUSG00000034341 | 1925,996557 | 2945,6 | 565,2 | 2,436461757 | 0,011411264 | Wbp2 | WW domain binding protein 2 [Source:MGI Symbol;Acc:MGI:104709] | 100,00 | 93,3 |
| ENSMUSG00000021520 | 2041,329657 | 2911,2 | 933,2 | 1,805019809 | 0,044783053 | Uqcrb | ubiquinol-cytochrome c reductase binding protein [Source:MGI Symbol;Acc:MGI:1914780] | 99,19 | 90,7 |
| ENSMUSG00000024091 | 1811,973348 | 2899,8 | 751,0 | 1,944708908 | 0,011034089 | Vapa | esicle-associated membrane protein, associated protein A [Source:MGI Symbol;Acc:MGI:135356 | 100,00 | 92,2 |
| ENSMUSG00000040659 | 1865,924429 | 2832,3 | 535,9 | 2,654244529 | 0,036094827 | Efhd2 | EF hand domain containing 2 [Source:MGI Symbol;Acc:MGI:106504] | 99,83 | 86,3 |
| ENSMUSG00000030161 | 1781,509257 | 2811,1 | 835,5 | 1,91998543 | 0,037496059 | Gabarapl1 | a-aminobutyric acid (GABA) A receptor-associated protein-like 1 [Source:MGI Symbol;Acc:MGI:19 | 98,97 | 90,6 |
| ENSMUSG00000017631 | 1724,899954 | 2803,6 | 792,9 | 1,913469508 | 0,032006231 | Abr | active BCR-related gene [Source:MGI Symbol;Acc:MGI:107771] | 78,13 | 51,3 |
| ENSMUSG00000026223 | 1930,130961 | 2790,6 | 854,1 | 1,863351175 | 0,041869735 | Itm2c | integral membrane protein 2C [Source:MGI Symbol;Acc:MGI:1927594] | 91,24 | 86,4 |
| ENSMUSG00000028603 | 1832,008607 | 2754,7 | 795,8 | 2,4328615 | 0,039405259 | Scp2 | sterol carrier protein 2, liver [Source:MGI Symbol;Acc:MGI:98254] | 78,74 | 29,7 |
| ENSMUSG00000030934 | 1689,644553 | 2751,1 | 252,7 | 4,443483032 | 0,004375869 | Oat | ornithine aminotransferase [Source:MGI Symbol;Acc:MGI:97394] | 99,35 | 60,4 |
| ENSMUSG00000020153 | 1894,682385 | 2725,9 | 1043,4 | 1,479969006 | 0,02491849 | Ndufs7 | NADH dehydrogenase (ubiquinone) Fe-S protein 7 [Source:MGI Symbol;Acc:MGI:1922656] | 100,00 | 77,9 |
| ENSMUSG00000022010 | 1215,684324 | 2723,5 | 359,5 | 3,309580977 | 0,004779948 | Tsc22d1 | TSC22 domain family, member 1 [Source:MGI Symbol;Acc:MGI:109127] | 70,62 | 24,3 |
| ENSMUSG00000006299 | 1719,96549 | 2712,9 | 619,4 | 2,162223385 | 0,009115328 | Aamp | angio-associated migratory protein [Source:MGI Symbol;Acc:MGI:107809] | 88,33 | 84,9 |
| ENSMUSG00000031819 | 1725,682957 | 2664,0 | 918,1 | 1,589262123 | 0,031449128 | Emc8 | ER membrane protein complex subunit 8 [Source:MGI Symbol;Acc:MGI:1343095] | 99,96 | 45,6 |
| ENSMUSG00000036564 | 1703,765749 | 2659,6 | 561,0 | 2,301540314 | 0,037604751 | Ndrg4 | N-myc downstream regulated gene 4 [Source:MGI Symbol;Acc:MGI:2384590] | 86,34 | 67,0 |
| ENSMUSG00000056486 | 1631,697437 | 2658,4 | 569,2 | 3,549330761 | 0,001198022 | Chn1 | chimerin 1 [Source:MGI Symbol;Acc:MGI:1915674] | 90,92 | 29,7 |
| ENSMUSG00000028745 | 1860,041406 | 2618,2 | 858,4 | 1,72263377 | 0,008385673 | Capzb | capping protein (actin filament) muscle Z-line, beta [Source:MGI Symbol;Acc:MGI:104652] | 79,73 | 72,5 |
| ENSMUSG00000025911 | 1586,347819 | 2611,2 | 470,9 | 4,433672572 | 0,018661386 | Adhfe1 | alcohol dehydrogenase, iron containing, 1 [Source:MGI Symbol;Acc:MGI:1923437] | 86,12 | 38,3 |
| ENSMUSG00000058267 | 1767,32929 | 2481,6 | 877,7 | 1,565661973 | 0,018346333 | Mrps14 | mitochondrial ribosomal protein S14 [Source:MGI Symbol;Acc:MGI:1928141] | 90,41 | 26,5 |
| ENSMUSG00000093930 | 1515,438221 | 2470,9 | 447,6 | 2,838680654 | 0,005058806 | Hmgcs1 | 3-hydroxy-3-methylglutaryl-Coenzyme A synthase 1 [Source:MGI Symbol;Acc:MGI:107592] | 95,81 | 63,0 |
| ENSMUSG00000019210 | 1460,060984 | 2443,7 | 554,1 | 2,306610391 | 0,043042394 | Atp6v1e1 | ATPase, H+ transporting, lysosomal V1 subunit E1 [Source:MGI Symbol;Acc:MGI:894326] | 95,31 | 77,5 |
| ENSMUSG00000015478 | 1566,798617 | 2428,2 | 621,6 | 2,378579114 | 0,01670605 | Rnf5 | ring finger protein 5 [Source:MGI Symbol;Acc:MGI:1860076] | 97,71 | 84,1 |
| ENSMUSG00000026895 | 1575,596634 | 2427,0 | 605,8 | 2,094147255 | 0,008019388 | Ndufa8 | IADH dehydrogenase (ubiquinone) 1 alpha subcomplex, 8 [Source:MGI Symbol;Acc:MGI:191562 | 99,54 | 89,4 |
| ENSMUSG00000096210 | 1685,333745 | 2419,1 | 958,6 | 1,622162446 | 0,048969442 | H1f0 | H1 histone family, member 0 [Source:MGI Symbol;Acc:MGI:95893] | 95,98 | 83,6 |
| ENSMUSG00000024870 | 1448,181509 | 2402,2 | 548,7 | 2,376310822 | 0,012503927 | Rab1b | RAB1B, member RAS oncogene family [Source:MGI Symbol;Acc:MGI:1923558] | 100,00 | 94,2 |
| ENSMUSG00000003161 | 1520,814897 | 2380,4 | 298,3 | 4,575573785 | 0,003406984 | Sri | sorcin [Source:MGI Symbol;Acc:MGI:98419] | 90,88 | 41,8 |
| ENSMUSG00000022476 | 1439,630778 | 2319,6 | 676,8 | 1,792890454 | 0,008890511 | Polr3h | polymerase (RNA) III (DNA directed) polypeptide H [Source:MGI Symbol;Acc:MGI:1926179] | 100,00 | 77,4 |
| ENSMUSG00000038059 | 1023,446181 | 2280,3 | 177,3 | 5,210595532 | 0,016782598 | Smim3 | small integral membrane protein 3 [Source:MGI Symbol;Acc:MGI:1917088] | 97,63 | 41,4 |
| ENSMUSG00000021771 | 1487,728861 | 2251,0 | 846,1 | 1,530492292 | 0,032785753 | Vdac2 | voltage-dependent anion channel 2 [Source:MGI Symbol;Acc:MGI:106915] | 74,71 | 64,5 |
| ENSMUSG00000007815 | 1527,43217 | 2229,5 | 683,7 | 1,766259746 | 0,011984883 | Rhoa | ras homolog gene family, member A [Source:MGI Symbol;Acc:MGI:1096342] | 89,57 | 68,6 |
| ENSMUSG00000032314 | 1559,219018 | 2222,0 | 783,2 | 1,651294618 | 0,026876988 | Etfa | electron transferring flavoprotein, alpha polypeptide [Source:MGI Symbol;Acc:MGI:106092] | 98,37 | 87,3 |
| ENSMUSG00000054252 | 1421,456376 | 2177,6 | 334,5 | 4,327974932 | 0,032460068 | Fgfr3 | fibroblast growth factor receptor 3 [Source:MGI Symbol;Acc:MGI:95524] | 95,18 | 34,0 |
| ENSMUSG00000015092 | 1483,773919 | 2160,2 | 836,7 | 1,503425603 | 0,033078575 | Edf1 | endothelial differentiation-related factor 1 [Source:MGI Symbol;Acc:MGI:1891227] | 100,00 | 97,4 |
| ENSMUSG00000017009 | 1296,644605 | 2148,1 | 429,7 | 3,406763036 | 0,033778987 | Sdc4 | syndecan 4 [Source:MGI Symbol;Acc:MGI:1349164] | 100,00 | 54,4 |
| ENSMUSG00000006782 | 1391,115634 | 2135,2 | 336,9 | 3,121961733 | 0,000397712 | Cnp | 2',3'-cyclic nucleotide 3' phosphodiesterase [Source:MGI Symbol;Acc:MGI:88437] | 98,85 | 49,9 |
| ENSMUSG00000072772 | 1445,586207 | 2108,5 | 769,2 | 1,548932818 | 0,040451804 | Grcc10 | gene rich cluster, C10 gene [Source:MGI Symbol;Acc:MGI:1315201] | 91,85 | 63,9 |
| ENSMUSG00000021371 | 1243,973363 | 2060,9 | 350,3 | 2,838174058 | 0,046938122 | Mcur1 | mitochondrial calcium uniporter regulator 1 [Source:MGI Symbol;Acc:MGI:1923387] | 99,98 | 43,4 |
| ENSMUSG00000028128 | 1180,710347 | 2050,6 | 286,4 | 3,439879082 | 0,029459488 | F3 | coagulation factor III [Source:MGI Symbol;Acc:MGI:88381] | 100,00 | 63,3 |
| ENSMUSG00000027533 | 1556,38122 | 2029,8 | 786,0 | 1,360619246 | 0,032399561 | Fabp5 | fatty acid binding protein 5, epidermal [Source:MGI Symbol;Acc:MGI:101790] | 91,02 | 48,7 |
| ENSMUSG00000026227 | 1185,538334 | 2028,4 | 361,1 | 5,42136497 | 0,019428823 | 2810459M11Rik | RIKEN cDNA 2810459M11 gene [Source:MGI Symbol;Acc:MGI:1920042] | 99,77 | 38,4 |
| ENSMUSG00000001700 | 1565,983308 | 2022,6 | 691,6 | 2,200625467 | 0,034742336 | Gramd3 | GRAM domain containing 3 [Source:MGI Symbol;Acc:MGI:1914815] | 99,99 | 70,1 |
| ENSMUSG00000027712 | 1247,022224 | 2006,0 | 366,1 | 2,988737599 | 0,00133698 | Anxa5 | annexin A5 [Source:MGI Symbol;Acc:MGI:106008] | 99,11 | 79,2 |
| ENSMUSG00000031021 | 1417,297086 | 1984,0 | 604,9 | 1,768879675 | 0,043819032 | Tmem9b | TMEM9 domain family, member B [Source:MGI Symbol;Acc:MGI:1915254] | 99,87 | 72,3 |
| ENSMUSG00000026568 | 1400,223172 | 1961,9 | 773,2 | 1,365040645 | 0,04419823 | Mpc2 | mitochondrial pyruvate carrier 2 [Source:MGI Symbol;Acc:MGI:1917706] | 99,77 | 73,6 |
| ENSMUSG00000044252 | 1190,735998 | 1959,8 | 86,3 | 8,197390857 | 2,63496E-05 | Osbpl1a | oxysterol binding protein-like 1A [Source:MGI Symbol;Acc:MGI:1927551] | 77,30 | 13,8 |
| ENSMUSG00000028221 | 1196,441892 | 1956,7 | 221,5 | 6,318912847 | 1,68917E-06 | Tmem55a | transmembrane protein 55A [Source:MGI Symbol;Acc:MGI:1919769] | 95,77 | 34,9 |
| ENSMUSG00000020949 | 1264,009972 | 1939,9 | 621,9 | 1,816015528 | 0,015863155 | Fkbp3 | FK506 binding protein 3 [Source:MGI Symbol;Acc:MGI:1353460] | 100,00 | 99,1 |

| | | | | | | | | | |
|---|---|---|---|---|---|---|---|---|---|
| ENSMUSG00000058672 | 1059,397743 | 1899,0 | 298,0 | 2,712962672 | 0,001000199 | Tubb2a | tubulin, beta 2A class IIA [Source:MGI Symbol;Acc:MGI:107861] | 77,27 | 64,9 |
| ENSMUSG00000027797 | 1488,855498 | 1881,1 | 643,4 | 3,185303448 | 0,030148615 | Dclk1 | doublecortin-like kinase 1 [Source:MGI Symbol;Acc:MGI:1330861] | 94,18 | 21,7 |
| ENSMUSG00000021417 | 1275,863532 | 1873,8 | 668,6 | 1,63798989 | 0,037737088 | Eci2 | enoyl-Coenzyme A delta isomerase 2 [Source:MGI Symbol;Acc:MGI:1346064] | 93,12 | 78,0 |
| ENSMUSG00000025509 | 810,3082676 | 1856,8 | 0,0 | 14,21350216 | 1,76206E-05 | Pnpla2 | patatin-like phospholipase domain containing 2 [Source:MGI Symbol;Acc:MGI:1914103] | 85,24 | 0,0 |
| ENSMUSG00000059734 | 1316,029065 | 1838,4 | 559,7 | 1,758414445 | 0,01404616 | Ndufs8 | NADH dehydrogenase (ubiquinone) Fe-S protein 8 [Source:MGI Symbol;Acc:MGI:2385079] | 100,00 | 83,0 |
| ENSMUSG00000027406 | 1230,697481 | 1815,6 | 541,8 | 1,786623633 | 0,036470737 | Idh3b | isocitrate dehydrogenase 3 (NAD+) beta [Source:MGI Symbol;Acc:MGI:2158650] | 98,66 | 92,0 |
| ENSMUSG00000002459 | 1068,107793 | 1815,1 | 262,5 | 3,011057567 | 0,036785298 | Rgs20 | regulator of G-protein signaling 20 [Source:MGI Symbol;Acc:MGI:1929866] | 95,53 | 54,1 |
| ENSMUSG00000019998 | 1127,205005 | 1811,1 | 251,7 | 2,936334293 | 0,036029848 | Stx7 | syntaxin 7 [Source:MGI Symbol;Acc:MGI:1858210] | 92,43 | 63,8 |
| ENSMUSG00000013921 | 988,2428338 | 1809,4 | 121,3 | 4,89527626 | 0,02958134 | Clip3 | CAP-GLY domain containing linker protein 3 [Source:MGI Symbol;Acc:MGI:1923936] | 99,90 | 34,4 |
| ENSMUSG00000009291 | 1239,941908 | 1784,4 | 325,4 | 2,467305244 | 0,0047196 | Pttg1ip | pituitary tumor-transforming 1 interacting protein [Source:MGI Symbol;Acc:MGI:2652132] | 92,41 | 79,7 |
| ENSMUSG00000036503 | 1121,304896 | 1778,8 | 230,9 | 4,018600589 | 0,000755551 | Rnf13 | ring finger protein 13 [Source:MGI Symbol;Acc:MGI:1346341] | 76,85 | 25,9 |
| ENSMUSG00000015484 | 1061,184467 | 1766,4 | 299,9 | 6,194183287 | 0,011731631 | Fam163a | family with sequence similarity 163, member A [Source:MGI Symbol;Acc:MGI:3618859] | 100,00 | 34,5 |
| ENSMUSG00000053768 | 1343,433037 | 1761,2 | 672,7 | 1,484451715 | 0,049495484 | Chchd3 | coiled-coil-helix-coiled-coil-helix domain containing 3 [Source:MGI Symbol;Acc:MGI:1913325] | 91,00 | 65,5 |
| ENSMUSG00000039347 | 1178,61283 | 1761,1 | 273,6 | 2,830326952 | 0,002086313 | Atp6v0e2 | ATPase, H+ transporting, lysosomal V0 subunit E2 [Source:MGI Symbol;Acc:MGI:1923502] | 99,98 | 76,6 |
| ENSMUSG00000004610 | 1377,465746 | 1750,1 | 678,4 | 1,494929256 | 0,040198466 | Etfb | electron transferring flavoprotein, beta polypeptide [Source:MGI Symbol;Acc:MGI:106098] | 98,10 | 93,6 |
| ENSMUSG00000002808 | 1109,202615 | 1724,2 | 366,7 | 2,510182537 | 0,004023292 | Epdr1 | ependymin related protein 1 (zebrafish) [Source:MGI Symbol;Acc:MGI:2145369] | 99,61 | 60,1 |
| ENSMUSG00000035242 | 1226,91357 | 1721,7 | 665,1 | 1,418093689 | 0,043293286 | Oaz1 | ornithine decarboxylase antizyme 1 [Source:MGI Symbol;Acc:MGI:109433] | 72,03 | 58,2 |
| ENSMUSG00000034853 | 1112,892301 | 1710,1 | 530,8 | 5,520561062 | 0,023510692 | Acot11 | acyl-CoA thioesterase 11 [Source:MGI Symbol;Acc:MGI:1913736] | 95,22 | 33,5 |
| ENSMUSG00000040147 | 928,3758273 | 1700,1 | 366,9 | 3,407038748 | 0,045877678 | Maob | monoamine oxidase B [Source:MGI Symbol;Acc:MGI:96916] | 100,00 | 58,1 |
| ENSMUSG00000026781 | 1045,220997 | 1693,4 | 208,4 | 3,398455386 | 0,032118278 | Acbd5 | acyl-Coenzyme A binding domain containing 5 [Source:MGI Symbol;Acc:MGI:1921409] | 94,94 | 25,0 |
| ENSMUSG00000019428 | 1202,659648 | 1689,2 | 417,0 | 2,1358753 | 0,020672134 | Fkbp8 | FK506 binding protein 8 [Source:MGI Symbol;Acc:MGI:1341070] | 87,80 | 61,9 |
| ENSMUSG00000022240 | 966,1032321 | 1685,6 | 341,3 | 4,127390369 | 0,013193418 | Ctnnd2 | catenin (cadherin associated protein), delta 2 [Source:MGI Symbol;Acc:MGI:1195966] | 98,79 | 31,3 |
| ENSMUSG00000039206 | 1025,17827 | 1682,0 | 487,7 | 1,795300187 | 0,02076277 | Daglb | diacylglycerol lipase, beta [Source:MGI Symbol;Acc:MGI:2442032] | 79,67 | 13,3 |
| ENSMUSG00000022214 | 1110,993832 | 1667,5 | 329,0 | 2,552926558 | 0,00303887 | Dcaf11 | DDB1 and CUL4 associated factor 11 [Source:MGI Symbol;Acc:MGI:90168] | 99,77 | 65,7 |
| ENSMUSG00000032470 | 1160,094472 | 1652,4 | 651,7 | 1,568896614 | 0,038333083 | Mras | muscle and microspikes RAS [Source:MGI Symbol;Acc:MGI:1100856] | 95,61 | 54,5 |
| ENSMUSG00000078202 | 962,0327184 | 1648,6 | 359,8 | 7,871820206 | 8,51191E-06 | Nrarp | Notch-regulated ankyrin repeat protein [Source:MGI Symbol;Acc:MGI:1914372] | 100,00 | 34,2 |
| ENSMUSG00000027668 | 1082,5317 | 1643,4 | 208,5 | 4,331102778 | 8,21045E-05 | Mfn1 | mitofusin 1 [Source:MGI Symbol;Acc:MGI:1914664] | 83,49 | 20,0 |
| ENSMUSG00000026032 | 1083,248404 | 1636,9 | 441,1 | 2,083151555 | 0,005912613 | Ndufb3 | NADH dehydrogenase (ubiquinone) 1 beta subcomplex 3 [Source:MGI Symbol;Acc:MGI:1913745] | 99,48 | 65,7 |
| ENSMUSG00000030127 | 975,4883976 | 1630,7 | 266,1 | 3,103750097 | 0,014694066 | Cops7a | COP9 signalosome subunit 7A [Source:MGI Symbol;Acc:MGI:1349400] | 98,88 | 60,2 |
| ENSMUSG00000029213 | 1254,891883 | 1619,4 | 599,5 | 1,493828888 | 0,035667203 | Commd8 | COMM domain containing 8 [Source:MGI Symbol;Acc:MGI:1343485] | 99,74 | 60,9 |
| ENSMUSG00000033720 | 924,4757416 | 1618,3 | 223,0 | 3,234648542 | 0,005708695 | Sfxn5 | sideroflexin 5 [Source:MGI Symbol;Acc:MGI:2137681] | 96,41 | 25,3 |
| ENSMUSG00000024899 | 750,1755373 | 1611,3 | 20,2 | 8,277173954 | 2,15802E-05 | Papss2 | 3'-phosphoadenosine 5'-phosphosulfate synthase 2 [Source:MGI Symbol;Acc:MGI:1330223] | 99,55 | 5,7 |
| ENSMUSG00000018849 | 999,007905 | 1602,2 | 347,4 | 9,62886232 | 0,000138865 | Wwc1 | WW, C2 and coiled-coil domain containing 1 [Source:MGI Symbol;Acc:MGI:2388637] | 99,93 | 31,2 |
| ENSMUSG00000020889 | 844,2585255 | 1585,4 | 180,7 | 3,392786163 | 0,008552163 | Nr1d1 | nuclear receptor subfamily 1, group D, member 1 [Source:MGI Symbol;Acc:MGI:2444210] | 98,84 | 48,6 |
| ENSMUSG00000043445 | 1063,991897 | 1573,1 | 421,1 | 2,228539158 | 0,003934342 | Pgp | phosphoglycolate phosphatase [Source:MGI Symbol;Acc:MGI:1914328] | 100,00 | 93,1 |
| ENSMUSG00000016427 | 1216,394086 | 1566,6 | 546,8 | 1,491259401 | 0,035329739 | Ndufa1 | ADH dehydrogenase (ubiquinone) 1 alpha subcomplex, 1 [Source:MGI Symbol;Acc:MGI:1929511 | 100,00 | 96,4 |
| ENSMUSG00000017670 | 990,9450104 | 1557,2 | 299,2 | 3,440091995 | 0,025410497 | Elmo2 | engulfment and cell motility 2 [Source:MGI Symbol;Acc:MGI:2153045] | 81,45 | 37,6 |
| ENSMUSG00000079316 | 862,3735515 | 1549,2 | 144,6 | 9,45235827 | 3,2957E-05 | Rab9 | RAB9, member RAS oncogene family [Source:MGI Symbol;Acc:MGI:1890695] | 74,37 | 23,9 |
| ENSMUSG00000029106 | 981,31401 | 1548,2 | 491,2 | 1,707967149 | 0,025833693 | Add1 | adducin 1 (alpha) [Source:MGI Symbol;Acc:MGI:87918] | 84,75 | 56,7 |
| ENSMUSG00000024827 | 810,761106 | 1545,1 | 321,5 | 3,269587192 | 0,021874999 | Gldc | glycine decarboxylase [Source:MGI Symbol;Acc:MGI:1341155] | 99,84 | 58,3 |
| ENSMUSG00000000399 | 1097,394795 | 1541,2 | 539,7 | 2,591507616 | 0,033632765 | Ndufa9 | ADH dehydrogenase (ubiquinone) 1 alpha subcomplex, 9 [Source:MGI Symbol;Acc:MGI:1913358 | 89,25 | 56,8 |
| ENSMUSG00000001768 | 931,1079024 | 1541,2 | 250,2 | 4,426095014 | 0,00220446 | Rin2 | Ras and Rab interactor 2 [Source:MGI Symbol;Acc:MGI:1921280] | 93,09 | 23,2 |
| ENSMUSG00000054604 | 907,272714 | 1535,8 | 271,8 | 3,164185179 | 0,047974717 | Cggbp1 | CGG triplet repeat binding protein 1 [Source:MGI Symbol;Acc:MGI:2146370] | 99,65 | 30,3 |
| ENSMUSG00000027199 | 1102,213686 | 1518,1 | 384,6 | 2,029583876 | 0,009341226 | Gatm | ne amidinotransferase (L-arginine:glycine amidinotransferase) [Source:MGI Symbol;Acc:MGI:1914 | 99,80 | 87,0 |
| ENSMUSG00000021024 | 1069,402194 | 1508,9 | 517,8 | 1,60913497 | 0,012234787 | Psma6 | proteasome (prosome, macropain) subunit, alpha type 6 [Source:MGI Symbol;Acc:MGI:1347006] | 89,26 | 84,0 |
| ENSMUSG00000005312 | 1109,756903 | 1498,5 | 413,8 | 2,617859019 | 0,009401723 | Ubqln1 | ubiquilin 1 [Source:MGI Symbol;Acc:MGI:1860276] | 99,90 | 49,7 |
| ENSMUSG00000016534 | 1071,441088 | 1494,1 | 258,7 | 2,443334079 | 0,003836524 | Lamp2 | lysosomal-associated membrane protein 2 [Source:MGI Symbol;Acc:MGI:96748] | 97,51 | 27,1 |
| ENSMUSG00000020107 | 1016,63675 | 1488,8 | 585,6 | 1,516180585 | 0,042818634 | Anapc16 | anaphase promoting complex subunit 16 [Source:MGI Symbol;Acc:MGI:1289325] | 93,03 | 53,0 |
| ENSMUSG00000027602 | 1044,470816 | 1477,6 | 476,5 | 2,097219576 | 0,039651736 | Map1lc3a | microtubule-associated protein 1 light chain 3 alpha [Source:MGI Symbol;Acc:MGI:1915661] | 98,86 | 85,0 |
| ENSMUSG00000085793 | 977,1521822 | 1476,1 | 325,4 | 2,490694723 | 0,026653364 | Lin52 | lin-52 homolog (C. elegans) [Source:MGI Symbol;Acc:MGI:3045391] | 92,65 | 68,9 |
| ENSMUSG00000028383 | 984,5809727 | 1462,6 | 421,8 | 10,15159043 | 0,001354712 | Hsdl2 | hydroxysteroid dehydrogenase like 2 [Source:MGI Symbol;Acc:MGI:1919729] | 93,73 | 29,9 |
| ENSMUSG00000003623 | 788,2001875 | 1461,8 | 146,3 | 8,183781815 | 6,8726E-07 | Crot | carnitine O-octanoyltransferase [Source:MGI Symbol;Acc:MGI:1921364] | 96,21 | 18,7 |
| ENSMUSG00000019494 | 1089,297468 | 1449,2 | 546,9 | 1,506204476 | 0,03434132 | Cops6 | COP9 signalosome subunit 6 [Source:MGI Symbol;Acc:MGI:1349439] | 96,27 | 57,4 |
| ENSMUSG00000037361 | 973,4242926 | 1448,2 | 381,2 | 2,000299186 | 0,005071598 | Sf3b6 | splicing factor 3B, subunit 6 [Source:MGI Symbol;Acc:MGI:1913305] | 99,76 | 64,2 |
| ENSMUSG00000074637 | 798,7731557 | 1447,9 | 52,9 | 4,814030343 | 0,010125358 | Sox2 | SRY (sex determining region Y)-box 2 [Source:MGI Symbol;Acc:MGI:98364] | 100,00 | 28,8 |
| ENSMUSG00000039682 | 1068,970325 | 1447,4 | 583,2 | 1,346464514 | 0,044124433 | Lap3 | leucine aminopeptidase 3 [Source:MGI Symbol;Acc:MGI:1914238] | 99,44 | 90,6 |
| ENSMUSG00000037712 | 922,2854553 | 1445,4 | 250,7 | 3,248149635 | 0,049283007 | Fermt2 | fermitin family member 2 [Source:MGI Symbol;Acc:MGI:2385001] | 91,84 | 38,2 |
| ENSMUSG00000077450 | 921,3516496 | 1435,8 | 351,6 | 2,167389697 | 0,003898346 | Rab11b | RAB11B, member RAS oncogene family [Source:MGI Symbol;Acc:MGI:99425] | 88,21 | 23,0 |
| ENSMUSG00000028923 | 922,4617878 | 1432,4 | 184,4 | 3,83310751 | 0,046935289 | Necap2 | NECAP endocytosis associated 2 [Source:MGI Symbol;Acc:MGI:1913397] | 99,91 | 56,5 |
| ENSMUSG00000103217 | 830,4869917 | 1421,6 | 316,8 | 7,667103101 | 1,3278E-05 | Gm38287 | predicted gene, 38287 [Source:MGI Symbol;Acc:MGI:5611515] | 100,00 | 34,4 |
| ENSMUSG00000021453 | 738,739459 | 1412,0 | 213,2 | 3,617881543 | 0,000663947 | Gadd45g | growth arrest and DNA-damage-inducible 45 gamma [Source:MGI Symbol;Acc:MGI:1346325] | 100,00 | 58,3 |
| ENSMUSG00000030087 | 785,9506496 | 1410,8 | 239,5 | 10,13344809 | 0,000145766 | Klf15 | Kruppel-like factor 15 [Source:MGI Symbol;Acc:MGI:1929988] | 91,39 | 26,9 |
| ENSMUSG00000007411 | 832,2872984 | 1410,2 | 264,6 | 3,114436716 | 0,04449741 | Mark3 | MAP/microtubule affinity regulating kinase 3 [Source:MGI Symbol;Acc:MGI:1341865] | 99,91 | 47,8 |
| ENSMUSG00000029446 | 950,342606 | 1402,7 | 434,0 | 2,05864391 | 0,014153278 | Psph | phosphoserine phosphatase [Source:MGI Symbol;Acc:MGI:97788] | 96,60 | 66,1 |
| ENSMUSG00000022472 | 946,9749004 | 1395,0 | 415,3 | 1,815806113 | 0,023276764 | Desi1 | desumoylating isopeptidase 1 [Source:MGI Symbol;Acc:MGI:106313] | 99,95 | 74,8 |
| ENSMUSG00000059447 | 898,8476499 | 1393,1 | 338,6 | 2,060632758 | 0,041531226 | Hadhb | e/3-ketoacyl-Coenzyme A thiolase/enoyl-Coenzyme A hydratase (trifunctional protein), beta subunit | 71,71 | 60,1 |
| ENSMUSG00000019689 | 943,0729735 | 1388,5 | 435,7 | 2,039557532 | 0,025713947 | 1110001J03Rik | RIKEN cDNA 1110001J03 gene [Source:MGI Symbol;Acc:MGI:1913367] | 100,00 | 94,4 |
| ENSMUSG00000005103 | 963,5555563 | 1381,2 | 338,4 | 2,527830655 | 0,031192751 | Wdr1 | WD repeat domain 1 [Source:MGI Symbol;Acc:MGI:1337100] | 99,40 | 50,3 |
| ENSMUSG00000059824 | 884,9900427 | 1378,4 | 280,0 | 2,618336911 | 0,010339674 | Dbp | D site albumin promoter binding protein [Source:MGI Symbol;Acc:MGI:94866] | 99,46 | 73,6 |

| | | | | | | | | | |
|---|---|---|---|---|---|---|---|---|---|
| ENSMUSG00000027187 | 794,9331423 | 1374,1 | 212,9 | 4,302970593 | 0,014342208 | Cat | catalase [Source:MGI Symbol;Acc:MGI:88271] | 95,99 | 26,5 |
| ENSMUSG00000028669 | 971,5740285 | 1366,3 | 340,9 | 2,847730765 | 0,032095287 | Pithd1 | al proteasome-interacting domain of thioredoxin-like) domain containing 1 [Source:MGI Symbol;Ac | 92,52 | 55,1 |
| ENSMUSG00000042363 | 942,6844532 | 1364,8 | 86,3 | 5,334901017 | 3,44087E-05 | Lgalsl | lectin, galactoside binding-like [Source:MGI Symbol;Acc:MGI:1916114] | 99,99 | 21,4 |
| ENSMUSG00000065990 | 913,5002483 | 1346,1 | 456,5 | 1,588464021 | 0,042232315 | Aurkaip1 | aurora kinase A interacting protein 1 [Source:MGI Symbol;Acc:MGI:1913327] | 93,06 | 53,9 |
| ENSMUSG00000036949 | 845,8175333 | 1315,0 | 237,1 | 4,083614229 | 0,007016941 | Slc39a12 | solute carrier family 39 (zinc transporter), member 12 [Source:MGI Symbol;Acc:MGI:2139274] | 96,08 | 32,8 |
| ENSMUSG00000031703 | 977,2546134 | 1306,3 | 217,1 | 3,265502326 | 0,010660604 | Itfg1 | integrin alpha FG-GAP repeat containing 1 [Source:MGI Symbol;Acc:MGI:106419] | 99,72 | 44,0 |
| ENSMUSG00000021286 | 791,0110138 | 1296,5 | 179,5 | 3,02189402 | 0,027719879 | Zfyve21 | zinc finger, FYVE domain containing 21 [Source:MGI Symbol;Acc:MGI:1915770] | 99,57 | 77,6 |
| ENSMUSG00000022112 | 760,5132483 | 1295,0 | 49,9 | 7,990192687 | 0,000121059 | Gpc5 | glypican 5 [Source:MGI Symbol;Acc:MGI:1194894] | 98,97 | 8,4 |
| ENSMUSG00000022139 | 735,7644587 | 1291,7 | 313,9 | 3,527580559 | 0,04509201 | Mbnl2 | muscleblind-like 2 [Source:MGI Symbol;Acc:MGI:2145597] | 99,96 | 41,2 |
| ENSMUSG00000034463 | 795,2566497 | 1272,9 | 320,9 | 7,21896488 | 0,00461289 | Scara3 | scavenger receptor class A, member 3 [Source:MGI Symbol;Acc:MGI:2444418] | 100,00 | 33,5 |
| ENSMUSG00000024500 | 742,2971983 | 1270,2 | 0,9 | 10,81461772 | 1,46512E-06 | Ppp2r2b | protein phosphatase 2, regulatory subunit B, beta [Source:MGI Symbol;Acc:MGI:1920180] | 95,27 | 0,7 |
| ENSMUSG00000020629 | 856,0173071 | 1254,7 | 314,0 | 3,383464105 | 0,020940193 | Adi1 | acireductone dioxygenase 1 [Source:MGI Symbol;Acc:MGI:2144929] | 98,58 | 58,1 |
| ENSMUSG00000027739 | 718,3458159 | 1241,5 | 0,9 | 10,88246443 | 7,08271E-06 | Rab33b | RAB33B, member RAS oncogene family [Source:MGI Symbol;Acc:MGI:1330805] | 91,17 | 0,5 |
| ENSMUSG00000022680 | 997,0763851 | 1240,3 | 450,0 | 2,029272382 | 0,04176854 | Pdxdc1 | pyridoxal-dependent decarboxylase domain containing 1 [Source:MGI Symbol;Acc:MGI:1920909] | 74,52 | 29,3 |
| ENSMUSG00000017221 | 831,2640235 | 1237,3 | 451,4 | 1,551167653 | 0,035445955 | Psmd3 | oteasome (prosome, macropain) 26S subunit, non-ATPase, 3 [Source:MGI Symbol;Acc:MGI:9885 | 96,64 | 89,1 |
| ENSMUSG00000026259 | 844,6142754 | 1235,4 | 261,6 | 4,405705429 | 0,043053311 | Ngef | neuronal guanine nucleotide exchange factor [Source:MGI Symbol;Acc:MGI:1858414] | 82,91 | 34,5 |
| ENSMUSG00000092395 | 783,6529183 | 1229,1 | 300,9 | 2,481219493 | 0,019372396 | Gm20463 | predicted gene 20463 [Source:MGI Symbol;Acc:MGI:5141928] | 95,72 | 86,4 |
| ENSMUSG00000023452 | 786,3439518 | 1226,5 | 253,8 | 2,92791599 | 0,031441632 | Pisd | phosphatidylserine decarboxylase [Source:MGI Symbol;Acc:MGI:2445114] | 93,41 | 42,4 |
| ENSMUSG00000031782 | 739,7624998 | 1224,2 | 215,4 | 3,917082161 | 0,048045881 | Coq9 | coenzyme Q9 [Source:MGI Symbol;Acc:MGI:1915164] | 98,36 | 57,4 |
| ENSMUSG00000054364 | 686,0014878 | 1222,2 | 312,3 | 2,388504401 | 0,021531594 | Rhob | ras homolog gene family, member B [Source:MGI Symbol;Acc:MGI:107949] | 99,97 | 72,6 |
| ENSMUSG00000039770 | 753,2811402 | 1217,8 | 178,1 | 3,036069369 | 0,023777367 | Ypel5 | yippee-like 5 (Drosophila) [Source:MGI Symbol;Acc:MGI:1916937] | 91,51 | 29,0 |
| ENSMUSG00000029669 | 803,3551436 | 1211,7 | 308,3 | 4,874627621 | 0,037161083 | Tspan12 | tetraspanin 12 [Source:MGI Symbol;Acc:MGI:1889818] | 88,14 | 26,5 |
| ENSMUSG00000031913 | 847,8854329 | 1208,0 | 257,4 | 3,414817368 | 0,022180393 | Vps4a | vacuolar protein sorting 4a (yeast) [Source:MGI Symbol;Acc:MGI:1890520] | 98,61 | 57,5 |
| ENSMUSG00000057236 | 917,2594864 | 1202,3 | 341,8 | 2,712466555 | 0,020333974 | Rbbp4 | retinoblastoma binding protein 4 [Source:MGI Symbol;Acc:MGI:1194912] | 96,42 | 26,1 |
| ENSMUSG00000030103 | 725,4548401 | 1192,8 | 98,4 | 5,113210667 | 0,037822042 | Bhlhe40 | basic helix-loop-helix family, member e40 [Source:MGI Symbol;Acc:MGI:1097714] | 96,08 | 25,3 |
| ENSMUSG00000024454 | 794,5743063 | 1190,8 | 346,3 | 2,415219963 | 0,009892318 | Hdac3 | histone deacetylase 3 [Source:MGI Symbol;Acc:MGI:1343091] | 99,00 | 67,4 |
| ENSMUSG00000021684 | 650,5198139 | 1189,0 | 149,1 | 3,180344554 | 0,003685841 | Pde8b | phosphodiesterase 8B [Source:MGI Symbol;Acc:MGI:2443999] | 92,48 | 26,8 |
| ENSMUSG00000039105 | 782,1106069 | 1180,9 | 351,8 | 1,874845356 | 0,038989198 | Atp6v1g1 | ATPase, H+ transporting, lysosomal V1 subunit G1 [Source:MGI Symbol;Acc:MGI:1913540] | 88,60 | 71,9 |
| ENSMUSG00000022105 | 770,8607163 | 1178,5 | 334,6 | 6,222251532 | 0,001730007 | Rb1 | retinoblastoma 1 [Source:MGI Symbol;Acc:MGI:97874] | 98,22 | 31,2 |
| ENSMUSG00000038332 | 562,4530342 | 1168,5 | 3,0 | 9,6868209 | 8,36606E-06 | Sesn1 | sestrin 1 [Source:MGI Symbol;Acc:MGI:2155278] | 98,97 | 1,5 |
| ENSMUSG00000002524 | 787,3787945 | 1164,1 | 303,8 | 2,221990018 | 0,009219475 | Puf60 | poly-U binding splicing factor 60 [Source:MGI Symbol;Acc:MGI:1915209] | 99,74 | 71,7 |
| ENSMUSG00000027803 | 748,0561478 | 1156,0 | 67,5 | 9,383283909 | 0,000101185 | Wwtr1 | WW domain containing transcription regulator 1 [Source:MGI Symbol;Acc:MGI:1917649] | 98,95 | 11,8 |
| ENSMUSG00000030613 | 730,0011595 | 1155,9 | 99,1 | 4,474191675 | 0,032419045 | Ccdc90b | coiled-coil domain containing 90B [Source:MGI Symbol;Acc:MGI:1913615] | 99,83 | 41,0 |
| ENSMUSG00000062908 | 807,8869061 | 1151,0 | 144,5 | 4,007039645 | 0,016615332 | Acadm | acyl-Coenzyme A dehydrogenase, medium chain [Source:MGI Symbol;Acc:MGI:87867] | 94,62 | 37,5 |
| ENSMUSG00000010095 | 805,4621326 | 1150,7 | 316,1 | 2,898416317 | 0,016311869 | Slc3a2 | amily 3 (activators of dibasic and neutral amino acid transport), member 2 [Source:MGI Symbol;A | 72,16 | 36,0 |
| ENSMUSG00000013160 | 676,2240055 | 1149,6 | 39,7 | 5,344879598 | 2,80125E-06 | Atp6v0d1 | ATPase, H+ transporting, lysosomal V0 subunit D1 [Source:MGI Symbol;Acc:MGI:1201778] | 99,61 | 35,3 |
| ENSMUSG00000004843 | 723,2696758 | 1143,8 | 36,1 | 7,919181402 | 0,000851624 | Chmp2b | charged multivesicular body protein 2B [Source:MGI Symbol;Acc:MGI:1916192] | 100,00 | 20,0 |
| ENSMUSG00000018965 | 820,592509 | 1141,5 | 331,0 | 1,920194769 | 0,008430306 | Ywhah | oxygenase/tryptophan 5-monooxygenase activation protein, eta polypeptide [Source:MGI Symbol;A | 77,72 | 63,5 |
| ENSMUSG00000000538 | 633,3079565 | 1132,1 | 104,0 | 3,767832524 | 0,018030954 | Tom1l2 | target of myb1-like 2 (chicken) [Source:MGI Symbol;Acc:MGI:2443306] | 84,22 | 17,0 |
| ENSMUSG00000063694 | 818,6256562 | 1130,5 | 367,3 | 1,734909065 | 0,02753792 | Cycs | cytochrome c, somatic [Source:MGI Symbol;Acc:MGI:88578] | 74,03 | 12,5 |
| ENSMUSG00000030538 | 656,373653 | 1129,0 | 169,2 | 4,730889407 | 0,012435143 | Cib1 | calcium and integrin binding 1 (calmyrin) [Source:MGI Symbol;Acc:MGI:1344418] | 88,77 | 44,4 |
| ENSMUSG00000023092 | 672,7176838 | 1125,9 | 23,7 | 8,003535773 | 0,000142552 | Fhl1 | four and a half LIM domains 1 [Source:MGI Symbol;Acc:MGI:1298387] | 78,02 | 5,3 |
| ENSMUSG00000043496 | 714,9791074 | 1120,4 | 114,4 | 9,311505575 | 0,000245776 | Tril | TLR4 interactor with leucine-rich repeats [Source:MGI Symbol;Acc:MGI:1914123] | 99,98 | 20,1 |
| ENSMUSG00000046432 | 786,057818 | 1117,8 | 337,9 | 1,814063785 | 0,00624442 | Ngfrap1 | ve growth factor receptor (TNFRSF16) associated protein 1 [Source:MGI Symbol;Acc:MGI:13380 | 91,71 | 78,9 |
| ENSMUSG00000027777 | 625,9823286 | 1113,4 | 173,2 | 3,516976598 | 0,010101794 | Schip1 | schwannomin interacting protein 1 [Source:MGI Symbol;Acc:MGI:1353557] | 82,97 | 28,8 |
| ENSMUSG00000038206 | 613,6710587 | 1105,8 | 116,7 | 7,979856908 | 0,000686115 | Fbxo8 | F-box protein 8 [Source:MGI Symbol;Acc:MGI:1354696] | 99,82 | 29,3 |
| ENSMUSG00000022884 | 918,567066 | 1099,6 | 336,3 | 1,935333157 | 0,029323948 | Eif4a2 | eukaryotic translation initiation factor 4A2 [Source:MGI Symbol;Acc:MGI:106906] | 77,36 | 29,2 |
| ENSMUSG00000028879 | 706,3689385 | 1098,7 | 188,1 | 5,333893475 | 0,004681539 | Stx12 | syntaxin 12 [Source:MGI Symbol;Acc:MGI:1931027] | 99,76 | 29,2 |
| ENSMUSG00000025410 | 889,7881771 | 1092,8 | 327,8 | 1,818068269 | 0,009373845 | Dctn2 | dynactin 2 [Source:MGI Symbol;Acc:MGI:107733] | 99,46 | 65,9 |
| ENSMUSG00000022051 | 744,5990077 | 1090,4 | 171,4 | 2,912796724 | 0,00065863 | Bnip3l | BCL2/adenovirus E1B interacting protein 3-like [Source:MGI Symbol;Acc:MGI:1332659] | 89,08 | 24,7 |
| ENSMUSG00000017734 | 711,9491356 | 1079,1 | 150,5 | 3,936562623 | 0,036438839 | Dbndd2 | ysbindin (dystrobrevin binding protein 1) domain containing 2 [Source:MGI Symbol;Acc:MGI:10656 | 80,00 | 18,0 |
| ENSMUSG00000039067 | 674,9342418 | 1076,0 | 137,4 | 3,363206497 | 0,021130539 | Psmd7 | teasome (prosome, macropain) 26S subunit, non-ATPase, 7 [Source:MGI Symbol;Acc:MGI:13515 | 98,98 | 60,1 |
| ENSMUSG00000062797 | 737,9690946 | 1075,0 | 380,6 | 1,617310689 | 0,019428823 | l7Rn6 | lethal, Chr 7, Rinchik 6 [Source:MGI Symbol;Acc:MGI:96738] | 81,91 | 44,9 |
| ENSMUSG00000033111 | 710,3539597 | 1073,1 | 42,4 | 8,020068689 | 4,01052E-07 | 3830406C13Rik | RIKEN cDNA 3830406C13 gene [Source:MGI Symbol;Acc:MGI:1917937] | 99,85 | 11,4 |
| ENSMUSG00000031347 | 691,0435734 | 1071,1 | 156,9 | 4,081195167 | 0,009302039 | Cetn2 | centrin 2 [Source:MGI Symbol;Acc:MGI:1347085] | 86,51 | 27,3 |
| ENSMUSG00000073988 | 608,1843833 | 1069,2 | 126,5 | 9,347708529 | 0,001002202 | Ttpa | tocopherol (alpha) transfer protein [Source:MGI Symbol;Acc:MGI:1354168] | 95,49 | 15,4 |
| ENSMUSG00000027180 | 646,0047545 | 1055,5 | 44,4 | 8,203905018 | 8,57275E-08 | Fbxo3 | F-box protein 3 [Source:MGI Symbol;Acc:MGI:1929084] | 78,74 | 5,3 |
| ENSMUSG00000020366 | 670,6595536 | 1051,5 | 228,7 | 3,660470531 | 0,016261287 | Mapk9 | mitogen-activated protein kinase 9 [Source:MGI Symbol;Acc:MGI:1346862] | 100,00 | 30,3 |
| ENSMUSG00000085587 | 607,5720966 | 1048,5 | 172,8 | 2,69998563 | 0,043140291 | Gm14493 | predicted gene 14493 [Source:MGI Symbol;Acc:MGI:3705313] | 100,00 | 97,2 |
| ENSMUSG00000050423 | 526,5832015 | 1043,3 | 49,7 | 5,234971213 | 0,012259683 | Ppp1r3g | protein phosphatase 1, regulatory (inhibitor) subunit 3G [Source:MGI Symbol;Acc:MGI:1923737] | 99,76 | 22,3 |
| ENSMUSG00000102422 | 585,8950719 | 1038,6 | 161,2 | 3,619618972 | 0,015518472 | Schip1 | schwannomin interacting protein 1 [Source:MGI Symbol;Acc:MGI:1353557] | 82,95 | 33,3 |
| ENSMUSG00000019232 | 698,7722339 | 1033,1 | 105,8 | 9,755746558 | 0,000377106 | Etnppl | ethanolamine phosphate phospholyase [Source:MGI Symbol;Acc:MGI:1919010] | 97,24 | 4,7 |
| ENSMUSG00000026782 | 824,0320584 | 1030,6 | 305,3 | 1,782752341 | 0,009850501 | Abi2 | abl-interactor 2 [Source:MGI Symbol;Acc:MGI:106913] | 99,36 | 34,3 |
| ENSMUSG00000005034 | 649,1593359 | 1026,6 | 374,8 | 2,304170887 | 0,01781242 | Prkacb | protein kinase, cAMP dependent, catalytic, beta [Source:MGI Symbol;Acc:MGI:97594] | 87,87 | 32,1 |
| ENSMUSG00000016637 | 624,8752474 | 1026,2 | 233,7 | 2,688511809 | 0,007880777 | Ift27 | intraflagellar transport 27 [Source:MGI Symbol;Acc:MGI:1914292] | 100,00 | 73,2 |
| ENSMUSG00000021071 | 567,5835207 | 1024,7 | 73,1 | 5,395387432 | 0,010167511 | Trim9 | tripartite motif-containing 9 [Source:MGI Symbol;Acc:MGI:2137354] | 84,34 | 10,8 |
| ENSMUSG00000020647 | 594,7663381 | 1023,7 | 80,2 | 3,733771615 | 0,020359596 | Ncoa1 | nuclear receptor coactivator 1 [Source:MGI Symbol;Acc:MGI:1276523] | 99,40 | 11,3 |
| ENSMUSG00000027309 | 718,0173823 | 1021,0 | 592,9 | 5,650332698 | 0,008345158 | 4930402H24Rik | RIKEN cDNA 4930402H24 gene [Source:MGI Symbol;Acc:MGI:1923029] | 98,02 | 26,0 |
| ENSMUSG00000047242 | 649,9223064 | 1020,8 | 298,1 | 7,276453746 | 0,005222124 | Taf9b | IA polymerase II, TATA box binding protein (TBP)-associated factor [Source:MGI Symbol;Acc:MG | 83,10 | 25,2 |

| | | | | | | | | | |
|---|---|---|---|---|---|---|---|---|---|
| ENSMUSG00000021076 | 708,7366497 | 1019,7 | 326,4 | 2,407450913 | 0,047498192 | Actr10 | ARP10 actin-related protein 10 [Source:MGI Symbol;Acc:MGI:1891654] | 93,32 | 38,8 |
| ENSMUSG00000026154 | 617,7917886 | 1014,5 | 265,8 | 2,4942935 | 0,038235961 | Sdhaf4 | succinate dehydrogenase complex assembly factor 4 [Source:MGI Symbol;Acc:MGI:1915252] | 99,66 | 87,8 |
| ENSMUSG00000048148 | 645,6487335 | 1013,9 | 91,9 | 4,430547884 | 0,047499556 | Nwd1 | NACHT and WD repeat domain containing 1 [Source:MGI Symbol;Acc:MGI:2442268] | 92,39 | 10,7 |
| ENSMUSG00000022261 | 569,4634829 | 1012,6 | 0,0 | 13,33260589 | 0,000625913 | Sdc2 | syndecan 2 [Source:MGI Symbol;Acc:MGI:1349165] | 99,08 | 0,0 |
| ENSMUSG00000078566 | 588,5516072 | 1008,8 | 174,5 | 2,752086181 | 0,012127993 | Bnip3 | BCL2/adenovirus E1B interacting protein 3 [Source:MGI Symbol;Acc:MGI:109326] | 88,00 | 51,2 |
| ENSMUSG00000023904 | 748,862289 | 1004,4 | 394,1 | 1,909496949 | 0,041609376 | Hcfc1r1 | host cell factor C1 regulator 1 (XPO1-dependent) [Source:MGI Symbol;Acc:MGI:2663619] | 100,00 | 74,2 |
| ENSMUSG00000040415 | 611,5031199 | 1000,5 | 127,0 | 3,421926085 | 0,008196587 | Dtx3 | deltex 3, E3 ubiquitin ligase [Source:MGI Symbol;Acc:MGI:2135752] | 97,43 | 50,6 |
| ENSMUSG00000019803 | 625,0499056 | 997,3 | 362,4 | 3,973994495 | 0,027719879 | Nr2e1 | nuclear receptor subfamily 2, group E, member 1 [Source:MGI Symbol;Acc:MGI:1100526] | 91,57 | 36,9 |
| ENSMUSG00000022332 | 653,3312119 | 991,3 | 169,2 | 3,84943369 | 0,026680113 | Khdrbs3 | omain containing, RNA binding, signal transduction associated 3 [Source:MGI Symbol;Acc:MGI:131 | 98,60 | 41,7 |
| ENSMUSG00000018398 | 566,6545399 | 983,7 | 108,2 | 3,309033335 | 0,014694066 | Sept8 | septin 8 [Source:MGI Symbol;Acc:MGI:894310] | 96,65 | 16,9 |
| ENSMUSG00000002981 | 667,7155322 | 981,9 | 185,5 | 3,391958464 | 0,025913375 | Clptm1 | cleft lip and palate associated transmembrane protein 1 [Source:MGI Symbol;Acc:MGI:1927155] | 88,07 | 29,9 |
| ENSMUSG00000039680 | 640,3152604 | 973,7 | 140,6 | 3,034833904 | 0,001260716 | Mrps6 | mitochondrial ribosomal protein S6 [Source:MGI Symbol;Acc:MGI:2153111] | 97,87 | 75,1 |
| ENSMUSG00000030447 | 725,8870621 | 970,2 | 243,9 | 3,008836977 | 0,027751101 | Cyfip1 | cytoplasmic FMR1 interacting protein 1 [Source:MGI Symbol;Acc:MGI:1338801] | 90,55 | 16,2 |
| ENSMUSG00000057469 | 855,7155963 | 968,0 | 407,9 | 4,131249221 | 0,002708293 | E2f6 | E2F transcription factor 6 [Source:MGI Symbol;Acc:MGI:1354159] | 100,00 | 37,6 |
| ENSMUSG00000029613 | 713,2738151 | 966,9 | 242,2 | 3,096960796 | 0,005587554 | Eif2ak1 | eukaryotic translation initiation factor 2 alpha kinase 1 [Source:MGI Symbol;Acc:MGI:1353448] | 98,98 | 29,8 |
| ENSMUSG00000021796 | 829,1429023 | 959,6 | 92,6 | 4,037759421 | 0,0347406 | Bmpr1a | bone morphogenetic protein receptor, type 1A [Source:MGI Symbol;Acc:MGI:1338938] | 92,38 | 6,8 |
| ENSMUSG00000028207 | 778,6014344 | 949,7 | 350,0 | 2,675242486 | 0,040270912 | Asph | aspartate-beta-hydroxylase [Source:MGI Symbol;Acc:MGI:1914186] | 87,73 | 19,3 |
| ENSMUSG00000024978 | 515,0201062 | 947,9 | 242,2 | 5,319261838 | 0,019063996 | Gpam | glycerol-3-phosphate acyltransferase, mitochondrial [Source:MGI Symbol;Acc:MGI:109162] | 99,36 | 37,8 |
| ENSMUSG00000022898 | 612,9403905 | 947,3 | 148,5 | 4,875269003 | 0,012119144 | Dscr3 | Down syndrome critical region gene 3 [Source:MGI Symbol;Acc:MGI:1206040] | 96,81 | 31,6 |
| ENSMUSG00000069763 | 571,1147391 | 947,0 | 121,6 | 9,350761094 | 0,001264052 | Tmem100 | transmembrane protein 100 [Source:MGI Symbol;Acc:MGI:1915138] | 84,53 | 25,8 |
| ENSMUSG00000023156 | 689,0810211 | 943,2 | 354,0 | 8,845991967 | 0,000235523 | Rpp14 | ribonuclease P 14 subunit [Source:MGI Symbol;Acc:MGI:1914303] | 100,00 | 32,9 |
| ENSMUSG00000032878 | 593,3703502 | 931,1 | 228,9 | 6,027160152 | 0,003823378 | Ccdc85a | coiled-coil domain containing 85A [Source:MGI Symbol;Acc:MGI:2445069] | 98,03 | 25,1 |
| ENSMUSG00000051469 | 572,4937631 | 925,6 | 96,4 | 5,489464119 | 0,001912223 | Zfp24 | zinc finger protein 24 [Source:MGI Symbol;Acc:MGI:1929704] | 98,68 | 9,4 |
| ENSMUSG00000006423 | 576,2405048 | 918,9 | 67,4 | 4,38351691 | 0,014738218 | C330007P06Rik | RIKEN cDNA C330007P06 gene [Source:MGI Symbol;Acc:MGI:1924894] | 75,09 | 10,5 |
| ENSMUSG00000028180 | 869,7005151 | 916,7 | 367,1 | 2,717079903 | 0,037067021 | Zranb2 | zinc finger, RAN-binding domain containing 2 [Source:MGI Symbol;Acc:MGI:1858211] | 95,26 | 40,7 |
| ENSMUSG00000020794 | 671,2508399 | 916,0 | 262,2 | 3,963150889 | 0,019734779 | Ube2g1 | ubiquitin-conjugating enzyme E2G 1 [Source:MGI Symbol;Acc:MGI:1914378] | 99,58 | 21,2 |
| ENSMUSG00000068742 | 588,4488923 | 915,0 | 148,2 | 5,039207022 | 0,004170187 | Cry2 | cryptochrome 2 (photolyase-like) [Source:MGI Symbol;Acc:MGI:1270859] | 91,55 | 26,6 |
| ENSMUSG00000041577 | 579,4489536 | 914,7 | 142,5 | 5,376070811 | 0,00404364 | Prelp | proline arginine-rich end leucine-rich repeat [Source:MGI Symbol;Acc:MGI:2151110] | 100,00 | 28,0 |
| ENSMUSG00000074781 | 531,3799325 | 910,3 | 122,4 | 3,231326716 | 6,21322E-06 | Ube2n | ubiquitin-conjugating enzyme E2N [Source:MGI Symbol;Acc:MGI:1934835] | 86,52 | 13,1 |
| ENSMUSG00000027488 | 541,5952148 | 910,2 | 0,0 | 13,21805002 | 0,000652685 | Snta1 | syntrophin, acidic 1 [Source:MGI Symbol;Acc:MGI:101772] | 99,24 | 0,0 |
| ENSMUSG00000031565 | 520,5452656 | 905,4 | 68,6 | 4,023382579 | 0,005279836 | Fgfr1 | fibroblast growth factor receptor 1 [Source:MGI Symbol;Acc:MGI:95522] | 90,65 | 17,5 |
| ENSMUSG00000041935 | 596,8820503 | 898,3 | 157,6 | 3,800317794 | 0,008682778 | AW549877 | expressed sequence AW549877 [Source:MGI Symbol;Acc:MGI:2146232] | 99,85 | 25,9 |
| ENSMUSG00000068299 | 495,2325802 | 895,1 | 382,9 | 7,290950822 | 0,003685841 | 1700019G17Rik | RIKEN cDNA 1700019G17 gene [Source:MGI Symbol;Acc:MGI:1922791] | 90,63 | 30,3 |
| ENSMUSG00000031788 | 545,5937213 | 894,7 | 140,1 | 3,711332977 | 0,044692 | Kifc3 | kinesin family member C3 [Source:MGI Symbol;Acc:MGI:109202] | 96,06 | 38,0 |
| ENSMUSG00000020614 | 461,8008978 | 887,6 | 103,5 | 8,966837915 | 0,000490711 | Fam20a | family with sequence similarity 20, member A [Source:MGI Symbol;Acc:MGI:2388266] | 100,00 | 31,6 |
| ENSMUSG00000003299 | 651,390733 | 877,1 | 355,5 | 1,492016094 | 0,049460093 | Mrpl4 | mitochondrial ribosomal protein L4 [Source:MGI Symbol;Acc:MGI:2137210] | 99,66 | 82,2 |
| ENSMUSG00000043391 | 581,8867587 | 876,7 | 1,0 | 10,24725866 | 1,96332E-05 | 2510009E07Rik | RIKEN cDNA 2510009E07 gene [Source:MGI Symbol;Acc:MGI:1919440] | 99,61 | 0,5 |
| ENSMUSG00000031916 | 668,286609 | 874,8 | 241,9 | 2,160611846 | 0,016739916 | Cog8 | component of oligomeric golgi complex 8 [Source:MGI Symbol;Acc:MGI:2142885] | 85,39 | 27,5 |
| ENSMUSG00000048332 | 467,280596 | 870,4 | 207,6 | 9,606714316 | 0,003107701 | Lhfp | lipoma HMGIC fusion partner [Source:MGI Symbol;Acc:MGI:1920048] | 99,70 | 33,0 |
| ENSMUSG00000044018 | 632,9854342 | 869,4 | 216,6 | 3,088304034 | 0,021130539 | Mrpl50 | mitochondrial ribosomal protein L50 [Source:MGI Symbol;Acc:MGI:107329] | 99,95 | 41,3 |
| ENSMUSG00000008305 | 490,37892 | 868,3 | 84,1 | 8,697570604 | 5,63806E-05 | Tle1 | nsducin-like enhancer of split 1, homolog of Drosophila E(spl) [Source:MGI Symbol;Acc:MGI:1046 | 94,64 | 12,1 |
| ENSMUSG00000034484 | 555,1475291 | 864,5 | 147,7 | 4,712895578 | 0,017066504 | Snx2 | sorting nexin 2 [Source:MGI Symbol;Acc:MGI:1915054] | 98,38 | 34,2 |
| ENSMUSG00000042613 | 550,3691547 | 862,0 | 222,0 | 1,957736177 | 0,049958565 | Pbxip1 | re B cell leukemia transcription factor interacting protein 1 [Source:MGI Symbol;Acc:MGI:244167( | 99,89 | 39,6 |
| ENSMUSG00000030513 | 439,5542671 | 854,5 | 3,8 | 8,211432197 | 3,22281E-06 | Pcsk6 | proprotein convertase subtilisin/kexin type 6 [Source:MGI Symbol;Acc:MGI:102897] | 97,24 | 1,2 |
| ENSMUSG00000030770 | 587,1226938 | 848,8 | 215,5 | 8,967445563 | 0,000323009 | Parva | parvin, alpha [Source:MGI Symbol;Acc:MGI:1931144] | 93,83 | 27,7 |
| ENSMUSG00000000001 | 715,7066754 | 846,7 | 108,0 | 6,832794982 | 5,39421E-06 | Gnai3 | uanine nucleotide binding protein (G protein), alpha inhibiting 3 [Source:MGI Symbol;Acc:MGI:9577 | 99,86 | 25,6 |
| ENSMUSG00000008429 | 482,923573 | 846,3 | 135,1 | 4,371340023 | 0,015819883 | Herpud2 | HERPUD family member 2 [Source:MGI Symbol;Acc:MGI:1915393] | 100,00 | 18,8 |
| ENSMUSG00000059970 | 489,4639224 | 843,7 | 21,5 | 6,241837072 | 7,50734E-05 | Hspa2 | heat shock protein 2 [Source:MGI Symbol;Acc:MGI:96243] | 95,61 | 6,8 |
| ENSMUSG00000004849 | 546,1437265 | 842,5 | 239,8 | 2,268581824 | 0,009302039 | Ap1s1 | adaptor protein complex AP-1, sigma 1 [Source:MGI Symbol;Acc:MGI:1098244] | 73,74 | 53,2 |
| ENSMUSG00000053453 | 566,9933803 | 838,6 | 238,6 | 1,978428361 | 0,016274719 | Thoc7 | THO complex 7 homolog (Drosophila) [Source:MGI Symbol;Acc:MGI:1913481] | 100,00 | 74,6 |
| ENSMUSG00000022428 | 504,7432095 | 836,1 | 83,7 | 4,169409615 | 0,006499377 | Cby1 | chibby homolog 1 (Drosophila) [Source:MGI Symbol;Acc:MGI:1920989] | 100,00 | 60,9 |
| ENSMUSG00000050565 | 559,7682589 | 835,6 | 202,0 | 2,499472278 | 0,006726105 | Tor1aip2 | torsin A interacting protein 2 [Source:MGI Symbol;Acc:MGI:3582695] | 85,30 | 13,2 |
| ENSMUSG00000024140 | 517,6015488 | 832,8 | 168,4 | 4,192429595 | 0,014602 | Epas1 | endothelial PAS domain protein 1 [Source:MGI Symbol;Acc:MGI:109169] | 99,14 | 27,1 |
| ENSMUSG00000024423 | 664,8487652 | 831,8 | 319,0 | 2,103693997 | 0,0282791 | Impact | impact, RWD domain protein [Source:MGI Symbol;Acc:MGI:1098233] | 99,98 | 43,9 |
| ENSMUSG00000050721 | 716,7367821 | 830,7 | 116,9 | 5,055084201 | 0,000674007 | Plekho2 | leckstrin homology domain containing, family O member 2 [Source:MGI Symbol;Acc:MGI:214313: | 100,00 | 28,2 |
| ENSMUSG00000022272 | 611,5575088 | 829,7 | 111,0 | 3,470886686 | 0,019063996 | Myo10 | myosin X [Source:MGI Symbol;Acc:MGI:107716] | 71,29 | 10,1 |
| ENSMUSG00000021065 | 502,168909 | 829,5 | 73,1 | 7,318091804 | 0,000378972 | Fut8 | fucosyltransferase 8 [Source:MGI Symbol;Acc:MGI:1858901] | 94,83 | 20,1 |
| ENSMUSG00000060548 | 530,8751706 | 827,3 | 118,6 | 9,332821306 | 0,000957662 | Tnfrsf19 | tumor necrosis factor receptor superfamily, member 19 [Source:MGI Symbol;Acc:MGI:1352474] | 93,82 | 13,7 |
| ENSMUSG00000034235 | 502,8089575 | 826,0 | 244,7 | 7,13709247 | 0,002117304 | Usp54 | ubiquitin specific peptidase 54 [Source:MGI Symbol;Acc:MGI:1926037] | 99,95 | 24,1 |
| ENSMUSG00000038324 | 593,4117314 | 825,2 | 298,0 | 4,228417732 | 0,015231624 | Trpc4ap | tor potential cation channel, subfamily C, member 4 associated protein [Source:MGI Symbol;Acc: | 98,92 | 37,0 |
| ENSMUSG00000039217 | 558,9442091 | 823,3 | 123,9 | 3,60657922 | 0,00921148 | Il18 | interleukin 18 [Source:MGI Symbol;Acc:MGI:107936] | 98,88 | 37,8 |
| ENSMUSG00000029781 | 537,1075951 | 821,3 | 16,6 | 8,411235727 | 1,33767E-05 | Fkbp9 | FK506 binding protein 9 [Source:MGI Symbol;Acc:MGI:1350921] | 98,59 | 2,6 |
| ENSMUSG00000024993 | 651,6130822 | 817,1 | 128,1 | 5,539054613 | 0,000154767 | Fam45a | family with sequence similarity 45, member A [Source:MGI Symbol;Acc:MGI:1915144] | 99,92 | 32,9 |
| ENSMUSG00000032238 | 432,0270189 | 815,2 | 67,3 | 4,623851816 | 0,006626268 | Rora | RAR-related orphan receptor alpha [Source:MGI Symbol;Acc:MGI:104661] | 86,52 | 5,0 |
| ENSMUSG00000032388 | 582,5763096 | 809,8 | 167,0 | 2,289529332 | 0,008088217 | Spg21 | spastic paraplegia 21 homolog (human) [Source:MGI Symbol;Acc:MGI:106403] | 87,04 | 48,4 |
| ENSMUSG00000032905 | 643,4864475 | 806,8 | 96,4 | 4,232918111 | 0,008088217 | Atg12 | autophagy related 12 [Source:MGI Symbol;Acc:MGI:1914776] | 99,76 | 30,4 |
| ENSMUSG00000007777 | 487,7046392 | 803,2 | 119,2 | 3,719021449 | 0,031598818 | 0610009B22Rik | RIKEN cDNA 0610009B22 gene [Source:MGI Symbol;Acc:MGI:1913300] | 88,51 | 50,9 |
| ENSMUSG00000046688 | 447,3657828 | 802,4 | 107,8 | 4,838028648 | 0,003602277 | Tifa | TRAF-interacting protein with forkhead-associated domain [Source:MGI Symbol;Acc:MGI:2182965 | 96,98 | 19,7 |
| ENSMUSG00000032816 | 463,6894341 | 793,4 | 22,4 | 8,783608846 | 0,000191491 | Igdcc4 | immunoglobulin superfamily, DCC subclass, member 4 [Source:MGI Symbol;Acc:MGI:1858497] | 98,45 | 4,3 |

| | | | | | | | | | |
|---|---|---|---|---|---|---|---|---|---|
| ENSMUSG00000039183 | 544,0310683 | 792,2 | 246,4 | 1,725802066 | 0,010921483 | Nubp2 | nucleotide binding protein 2 [Source:MGI Symbol;Acc:MGI:1347072] | 99,77 | 92,2 |
| ENSMUSG00000031536 | 435,0680435 | 792,2 | 137,7 | 4,637662953 | 0,009723044 | Polb | polymerase (DNA directed), beta [Source:MGI Symbol;Acc:MGI:97740] | 99,76 | 32,3 |
| ENSMUSG00000054863 | 512,7219966 | 787,3 | 132,0 | 3,165286878 | 0,023150736 | Fam19a5 | family with sequence similarity 19, member A5 [Source:MGI Symbol;Acc:MGI:2146182] | 100,00 | 33,4 |
| ENSMUSG00000025555 | 570,6495437 | 784,2 | 47,0 | 4,73606724 | 3,85469E-05 | Farp1 | oGEF (Arhgef) and pleckstrin domain protein 1 (chondrocyte-derived) [Source:MGI Symbol;Acc:MC | 96,21 | 10,8 |
| ENSMUSG00000028278 | 585,1836176 | 783,8 | 68,0 | 5,077811641 | 0,014778862 | Rragd | Ras-related GTP binding D [Source:MGI Symbol;Acc:MGI:1098604] | 93,98 | 9,7 |
| ENSMUSG00000002205 | 522,0306241 | 781,4 | 215,3 | 2,28667077 | 0,009976334 | Vrk3 | vaccinia related kinase 3 [Source:MGI Symbol;Acc:MGI:2182465] | 77,32 | 43,7 |
| ENSMUSG00000030235 | 584,8064651 | 779,4 | 227,8 | 4,729652828 | 0,004694199 | Slco1c1 | olute carrier organic anion transporter family, member 1c1 [Source:MGI Symbol;Acc:MGI:188967 | 94,38 | 25,7 |
| ENSMUSG00000025153 | 534,1794263 | 778,9 | 148,0 | 2,387598616 | 0,00303887 | Fasn | fatty acid synthase [Source:MGI Symbol;Acc:MGI:95485] | 84,19 | 10,3 |
| ENSMUSG00000024259 | 522,4021203 | 777,6 | 185,2 | 7,846447205 | 7,83545E-05 | Slc25a46 | solute carrier family 25, member 46 [Source:MGI Symbol;Acc:MGI:1914703] | 85,66 | 18,1 |
| ENSMUSG00000003269 | 510,3468141 | 776,6 | 195,6 | 2,533360847 | 0,012958678 | Cyth2 | cytohesin 2 [Source:MGI Symbol;Acc:MGI:1334255] | 78,92 | 40,2 |
| ENSMUSG00000015839 | 458,830767 | 776,2 | 131,7 | 8,326764594 | 1,15903E-05 | Nfe2l2 | nuclear factor, erythroid derived 2, like 2 [Source:MGI Symbol;Acc:MGI:108420] | 98,59 | 25,2 |
| ENSMUSG00000062691 | 515,852525 | 775,5 | 233,5 | 1,78859764 | 0,024124442 | Cebpzos | AAT/enhancer binding protein (C/EBP), zeta, opposite strand [Source:MGI Symbol;Acc:MGI:1915 | 99,20 | 54,6 |
| ENSMUSG00000032010 | 531,6443062 | 775,0 | 190,3 | 3,183145006 | 0,004192399 | Usp2 | ubiquitin specific peptidase 2 [Source:MGI Symbol;Acc:MGI:1858178] | 95,43 | 25,6 |
| ENSMUSG00000029571 | 719,5971036 | 767,4 | 20,1 | 5,832771925 | 5,39421E-06 | Tmem106b | transmembrane protein 106B [Source:MGI Symbol;Acc:MGI:1919150] | 97,90 | 2,8 |
| ENSMUSG00000018287 | 512,1549412 | 764,1 | 228,5 | 1,938276547 | 0,01416993 | Spag7 | sperm associated antigen 7 [Source:MGI Symbol;Acc:MGI:107380] | 98,62 | 79,6 |
| ENSMUSG00000032413 | 501,8604574 | 763,4 | 108,3 | 3,981208163 | 0,025391383 | Rasa2 | RAS p21 protein activator 2 [Source:MGI Symbol;Acc:MGI:2149960] | 99,32 | 9,3 |
| ENSMUSG00000025537 | 468,3290363 | 760,6 | 21,8 | 6,109328353 | 2,55052E-05 | Phkg1 | phosphorylase kinase gamma 1 [Source:MGI Symbol;Acc:MGI:97579] | 87,53 | 7,1 |
| ENSMUSG00000027809 | 464,0805089 | 759,5 | 270,7 | 4,910085638 | 0,022041571 | Etfdh | electron transferring flavoprotein, dehydrogenase [Source:MGI Symbol;Acc:MGI:106100] | 95,50 | 11,3 |
| ENSMUSG00000021711 | 424,7727081 | 755,9 | 75,5 | 6,382079098 | 0,000358526 | Trappc13 | trafficking protein particle complex 13 [Source:MGI Symbol;Acc:MGI:1914225] | 99,42 | 13,7 |
| ENSMUSG00000022949 | 682,0979044 | 753,3 | 46,9 | 8,38954608 | 0,00086999 | Clic6 | chloride intracellular channel 6 [Source:MGI Symbol;Acc:MGI:2146607] | 95,85 | 3,7 |
| ENSMUSG00000036854 | 453,7225793 | 752,7 | 66,9 | 9,159333957 | 0,000532199 | Hspb6 | heat shock protein, alpha-crystallin-related, B6 [Source:MGI Symbol;Acc:MGI:2685325] | 99,90 | 27,0 |
| ENSMUSG00000078931 | 553,1619615 | 752,5 | 191,6 | 2,37991509 | 0,010159129 | Pdf | peptide deformylase (mitochondrial) [Source:MGI Symbol;Acc:MGI:1915273] | 100,00 | 49,3 |
| ENSMUSG00000032802 | 509,3481607 | 746,7 | 260,8 | 8,597928345 | 0,000564638 | Srxn1 | sulfiredoxin 1 homolog (S. cerevisiae) [Source:MGI Symbol;Acc:MGI:104971] | 97,62 | 29,7 |
| ENSMUSG00000053581 | 430,0723381 | 746,2 | 125,2 | 6,383954851 | 0,008263317 | Zfand2a | zinc finger, AN1-type domain 2A [Source:MGI Symbol;Acc:MGI:2140729] | 98,89 | 26,6 |
| ENSMUSG00000039530 | 507,7168092 | 742,6 | 109,7 | 3,590002148 | 0,019481169 | Tusc3 | tumor suppressor candidate 3 [Source:MGI Symbol;Acc:MGI:1933134] | 100,00 | 54,2 |
| ENSMUSG00000021589 | 428,2824058 | 740,9 | 208,1 | 6,386136024 | 0,000756646 | Rhobtb3 | Rho-related BTB domain containing 3 [Source:MGI Symbol;Acc:MGI:1920546] | 98,35 | 24,9 |
| ENSMUSG00000046312 | 407,8900163 | 735,0 | 0,9 | 10,17979973 | 8,88537E-05 | AI464131 | expressed sequence AI464131 [Source:MGI Symbol;Acc:MGI:2140300] | 93,83 | 0,5 |
| ENSMUSG00000028843 | 556,9498559 | 734,5 | 273,2 | 1,510393074 | 0,033172832 | Sh3bgrl3 | SH3 domain binding glutamic acid-rich protein-like 3 [Source:MGI Symbol;Acc:MGI:1920973] | 99,60 | 90,8 |
| ENSMUSG00000028857 | 564,1920553 | 733,2 | 218,2 | 3,381106392 | 0,006684206 | Tmem222 | transmembrane protein 222 [Source:MGI Symbol;Acc:MGI:1098568] | 100,00 | 44,4 |
| ENSMUSG00000026185 | 515,4174616 | 729,2 | 105,9 | 4,521962538 | 0,013986814 | Igfbp5 | insulin-like growth factor binding protein 5 [Source:MGI Symbol;Acc:MGI:96440] | 99,58 | 11,9 |
| ENSMUSG00000059409 | 539,0084069 | 729,1 | 174,5 | 2,398224107 | 0,009132349 | Ppp2r5d | protein phosphatase 2, regulatory subunit B', delta [Source:MGI Symbol;Acc:MGI:2388481] | 100,00 | 32,9 |
| ENSMUSG00000035559 | 522,1958701 | 728,7 | 179,6 | 2,021587406 | 0,013631482 | Mpv17l2 | MPV17 mitochondrial membrane protein-like 2 [Source:MGI Symbol;Acc:MGI:2681846] | 100,00 | 93,0 |
| ENSMUSG00000055932 | 529,8185053 | 728,0 | 458,3 | 3,790825004 | 0,016316251 | Fto | fat mass and obesity associated [Source:MGI Symbol;Acc:MGI:1347093] | 88,01 | 32,5 |
| ENSMUSG00000029474 | 440,4184696 | 727,2 | 129,1 | 2,97388898 | 0,009176851 | Rnf34 | ring finger protein 34 [Source:MGI Symbol;Acc:MGI:2153340] | 75,82 | 18,2 |
| ENSMUSG00000019433 | 470,6447524 | 725,8 | 180,1 | 2,224493651 | 0,025104874 | Gipc1 | GIPC PDZ domain containing family, member 1 [Source:MGI Symbol;Acc:MGI:1926252] | 100,00 | 83,3 |
| ENSMUSG00000057342 | 487,1653518 | 723,3 | 207,2 | 1,918092421 | 0,039812398 | Sphk2 | sphingosine kinase 2 [Source:MGI Symbol;Acc:MGI:1861380] | 93,05 | 27,0 |
| ENSMUSG00000040097 | 389,7278472 | 723,2 | 97,6 | 5,577298774 | 0,002167561 | Flywch1 | FLYWCH-type zinc finger 1 [Source:MGI Symbol;Acc:MGI:2442638] | 85,94 | 24,6 |
| ENSMUSG00000042750 | 395,8143897 | 720,8 | 53,1 | 4,805836205 | 0,000190848 | Bex2 | brain expressed X-linked 2 [Source:MGI Symbol;Acc:MGI:1338017] | 90,37 | 33,1 |
| ENSMUSG00000035847 | 617,3466869 | 719,8 | 106,3 | 8,109693803 | 9,58287E-06 | Ids | iduronate 2-sulfatase [Source:MGI Symbol;Acc:MGI:96417] | 96,62 | 20,7 |
| ENSMUSG00000061740 | 423,2892587 | 715,9 | 37,5 | 6,027371583 | 0,018984452 | Cyp2d22 | cytochrome P450, family 2, subfamily d, polypeptide 22 [Source:MGI Symbol;Acc:MGI:1929474] | 99,70 | 16,0 |
| ENSMUSG00000022800 | 541,9182751 | 714,5 | 153,5 | 2,376581851 | 0,018401236 | Fyttd1 | forty-two-three domain containing 1 [Source:MGI Symbol;Acc:MGI:1917955] | 84,53 | 23,9 |
| ENSMUSG00000003974 | 438,6457561 | 710,8 | 183,1 | 7,237895722 | 0,005428678 | Grm3 | glutamate receptor, metabotropic 3 [Source:MGI Symbol;Acc:MGI:1351340] | 100,00 | 32,9 |
| ENSMUSG00000027257 | 433,8289756 | 706,9 | 84,7 | 8,689278099 | 0,001128767 | Pacsin3 | protein kinase C and casein kinase substrate in neurons 3 [Source:MGI Symbol;Acc:MGI:1891410 | 73,91 | 21,6 |
| ENSMUSG00000032285 | 515,0352402 | 705,8 | 96,5 | 5,099225382 | 0,012689677 | Dnaja4 | DnaJ heat shock protein family (Hsp40) member A4 [Source:MGI Symbol;Acc:MGI:1927638] | 85,26 | 24,7 |
| ENSMUSG00000038387 | 419,0321684 | 700,7 | 200,7 | 2,087108059 | 0,044827322 | Rras | Harvey rat sarcoma oncogene, subgroup R [Source:MGI Symbol;Acc:MGI:98179] | 99,86 | 86,1 |
| ENSMUSG00000038286 | 411,9693714 | 700,5 | 35,1 | 8,838444465 | 0,001454212 | Bphl | olase-like (serine hydrolase, breast epithelial mucin-associated antigen) [Source:MGI Symbol;Acc:M | 100,00 | 26,5 |
| ENSMUSG00000021607 | 500,8761745 | 699,9 | 170,4 | 2,520429145 | 0,017920081 | Mrpl36 | mitochondrial ribosomal protein L36 [Source:MGI Symbol;Acc:MGI:2137228] | 100,00 | 64,0 |
| ENSMUSG00000065979 | 399,0341558 | 693,2 | 0,0 | 12,82164289 | 0,000276871 | Cpped1 | calcineurin-like phosphoesterase domain containing 1 [Source:MGI Symbol;Acc:MGI:2443300] | 96,72 | 0,0 |
| ENSMUSG00000036095 | 450,561104 | 687,5 | 35,5 | 7,512888667 | 0,000296708 | Dgkb | diacylglycerol kinase, beta [Source:MGI Symbol;Acc:MGI:2442474] | 100,00 | 4,5 |
| ENSMUSG00000050144 | 429,4937676 | 684,1 | 65,9 | 5,460875972 | 0,002750301 | Slc25a44 | solute carrier family 25, member 44 [Source:MGI Symbol;Acc:MGI:2444391] | 99,92 | 21,4 |
| ENSMUSG00000035585 | 400,6338273 | 682,1 | 145,6 | 3,92325707 | 0,026432771 | Tsen34 | tRNA splicing endonuclease 34 homolog (S. cerevisiae) [Source:MGI Symbol;Acc:MGI:1913328] | 89,44 | 33,2 |
| ENSMUSG00000039960 | 338,4913592 | 682,1 | 58,5 | 4,631808303 | 0,014677393 | Rhou | ras homolog gene family, member U [Source:MGI Symbol;Acc:MGI:1916831] | 99,37 | 18,3 |
| ENSMUSG00000044340 | 455,3316389 | 681,4 | 36,6 | 7,456646219 | 0,001647634 | Phlpp1 | PH domain and leucine rich repeat protein phosphatase 1 [Source:MGI Symbol;Acc:MGI:2138327 | 98,28 | 5,7 |
| ENSMUSG00000027665 | 418,2734575 | 674,2 | 72,6 | 4,076634085 | 0,012371023 | Pik3ca | phosphatidylinositol 3-kinase, catalytic, alpha polypeptide [Source:MGI Symbol;Acc:MGI:1206581] | 78,78 | 4,2 |
| ENSMUSG00000024414 | 488,0545719 | 673,9 | 150,8 | 2,383952208 | 0,006314703 | Mrpl27 | mitochondrial ribosomal protein L27 [Source:MGI Symbol;Acc:MGI:2137224] | 98,59 | 74,8 |
| ENSMUSG00000039485 | 496,5078394 | 669,5 | 137,5 | 3,146433223 | 0,043422761 | Tspyl4 | TSPY-like 4 [Source:MGI Symbol;Acc:MGI:106393] | 99,86 | 30,4 |
| ENSMUSG00000036565 | 453,2412347 | 668,1 | 155,0 | 2,210138495 | 0,039710916 | Ttyh3 | tweety homolog 3 (Drosophila) [Source:MGI Symbol;Acc:MGI:1925589] | 97,39 | 38,8 |
| ENSMUSG00000064215 | 556,6591641 | 666,7 | 94,9 | 3,584757005 | 0,014423451 | Ifi27 | interferon, alpha-inducible protein 27 [Source:MGI Symbol;Acc:MGI:1277180] | 99,74 | 58,5 |
| ENSMUSG00000060601 | 521,663964 | 666,5 | 162,3 | 2,778218622 | 0,003206771 | Nr1h2 | nuclear receptor subfamily 1, group H, member 2 [Source:MGI Symbol;Acc:MGI:1352463] | 79,57 | 41,0 |
| ENSMUSG00000025321 | 558,6375429 | 666,3 | 17,1 | 6,659339955 | 0,004625348 | Itgb8 | integrin beta 8 [Source:MGI Symbol;Acc:MGI:1338035] | 99,87 | 2,6 |
| ENSMUSG00000073987 | 432,7127801 | 666,1 | 24,8 | 8,289129699 | 8,13172E-05 | Ggh | gamma-glutamyl hydrolase [Source:MGI Symbol;Acc:MGI:1329035] | 94,01 | 13,2 |
| ENSMUSG00000000876 | 379,8415621 | 662,2 | 62,0 | 3,842006462 | 0,005979142 | Pxmp4 | peroxisomal membrane protein 4 [Source:MGI Symbol;Acc:MGI:1891701] | 95,07 | 21,1 |
| ENSMUSG00000038497 | 338,1019725 | 660,7 | 0,0 | 12,69399095 | 0,000910617 | Tmco3 | transmembrane and coiled-coil domains 3 [Source:MGI Symbol;Acc:MGI:2444946] | 82,67 | 0,0 |
| ENSMUSG00000030682 | 509,5327067 | 658,9 | 139,8 | 3,117524926 | 0,031082957 | Cdipt | ycerol--inositol 3-phosphatidyltransferase (phosphatidylinositol synthase) [Source:MGI Symbol;Acc | 83,19 | 28,2 |
| ENSMUSG00000031485 | 455,0666049 | 658,8 | 230,9 | 4,069753848 | 0,032949239 | Prosc | proline synthetase co-transcribed [Source:MGI Symbol;Acc:MGI:1891207] | 70,34 | 16,2 |
| ENSMUSG00000028443 | 399,1859145 | 658,0 | 0,0 | 12,7445966 | 0,000323542 | Nudt2 | udix (nucleoside diphosphate linked moiety X)-type motif 2 [Source:MGI Symbol;Acc:MGI:191365 | 100,00 | 0,0 |
| ENSMUSG00000022040 | 381,0775271 | 653,6 | 0,6 | 10,35718761 | 0,001096359 | Ephx2 | epoxide hydrolase 2, cytoplasmic [Source:MGI Symbol;Acc:MGI:99500] | 100,00 | 1,2 |
| ENSMUSG00000005804 | 454,4563505 | 653,5 | 183,6 | 6,360406926 | 0,011506412 | Bloc1s6 | genesis of lysosomal organelles complex-1, subunit 6, pallidin [Source:MGI Symbol;Acc:MGI:1927 | 77,62 | 20,5 |
| ENSMUSG00000024767 | 399,3464628 | 653,0 | 138,7 | 2,821819904 | 0,0135501 | Otub1 | OTU domain, ubiquitin aldehyde binding 1 [Source:MGI Symbol;Acc:MGI:2147616] | 88,17 | 49,7 |

| | | | | | | | | | |
|---|---|---|---|---|---|---|---|---|---|
| ENSMUSG00000026173 | 401,6669988 | 651,1 | 332,7 | 4,011660005 | 0,038651503 | Plcd4 | phospholipase C, delta 4 [Source:MGI Symbol;Acc:MGI:107469] | 87,92 | 20,9 |
| ENSMUSG00000060147 | 475,972002 | 651,0 | 312,3 | 8,61978222 | 0,000270676 | Serpinb6a | serine (or cysteine) peptidase inhibitor, clade B, member 6a [Source:MGI Symbol;Acc:MGI:103123 | 88,61 | 18,6 |
| ENSMUSG00000037656 | 371,847478 | 649,8 | 47,2 | 8,790219913 | 0,001199733 | Slc20a2 | solute carrier family 20, member 2 [Source:MGI Symbol;Acc:MGI:97851] | 98,83 | 12,5 |
| ENSMUSG00000026893 | 278,080922 | 649,1 | 30,5 | 8,007145236 | 5,29242E-05 | Gca | grancalcin [Source:MGI Symbol;Acc:MGI:1918521] | 99,74 | 7,9 |
| ENSMUSG00000020037 | 412,8020798 | 648,1 | 25,9 | 9,016134343 | 0,000492468 | Rfx4 | regulatory factor X, 4 (influences HLA class II expression) [Source:MGI Symbol;Acc:MGI:1918387 | 93,61 | 5,6 |
| ENSMUSG00000026727 | 374,8116654 | 647,6 | 7,0 | 9,250409403 | 0,000693649 | Rsu1 | Ras suppressor protein 1 [Source:MGI Symbol;Acc:MGI:103040] | 90,41 | 2,8 |
| ENSMUSG00000031367 | 444,1664251 | 647,2 | 189,4 | 5,589735535 | 0,0135501 | Ap1s2 | adaptor-related protein complex 1, sigma 2 subunit [Source:MGI Symbol;Acc:MGI:1889383] | 89,78 | 8,4 |
| ENSMUSG00000021039 | 560,4026852 | 646,8 | 195,9 | 1,717055624 | 0,024493876 | Snw1 | SNW domain containing 1 [Source:MGI Symbol;Acc:MGI:1913604] | 100,00 | 55,2 |
| ENSMUSG00000024507 | 376,0846306 | 645,3 | 171,9 | 7,946786063 | 0,000507497 | Hsd17b4 | hydroxysteroid (17-beta) dehydrogenase 4 [Source:MGI Symbol;Acc:MGI:105089] | 98,05 | 31,2 |
| ENSMUSG00000032540 | 492,8816675 | 643,5 | 116,8 | 8,594956832 | 0,000358526 | Abhd5 | abhydrolase domain containing 5 [Source:MGI Symbol;Acc:MGI:1914719] | 93,73 | 22,0 |
| ENSMUSG00000022540 | 457,5144483 | 643,4 | 213,2 | 1,825661497 | 0,037073718 | Rogdi | rogdi homolog (Drosophila) [Source:MGI Symbol;Acc:MGI:1913299] | 91,33 | 67,0 |
| ENSMUSG00000037253 | 398,167823 | 640,5 | 21,3 | 8,056893724 | 2,07881E-05 | Mex3c | mex3 homolog C (C. elegans) [Source:MGI Symbol;Acc:MGI:2652843] | 98,61 | 5,2 |
| ENSMUSG00000002550 | 370,5328053 | 638,2 | 126,3 | 3,519702036 | 0,039939641 | Uck1 | uridine-cytidine kinase 1 [Source:MGI Symbol;Acc:MGI:98904] | 74,75 | 32,9 |
| ENSMUSG00000031840 | 402,1136673 | 637,9 | 105,9 | 2,567483766 | 0,000212251 | Rab3a | RAB3A, member RAS oncogene family [Source:MGI Symbol;Acc:MGI:97843] | 83,20 | 57,9 |
| ENSMUSG00000091512 | 428,8963462 | 636,2 | 185,5 | 2,21681615 | 0,036439871 | Lamtor3 | endosomal/lysosomal adaptor, MAPK and MTOR activator 3 [Source:MGI Symbol;Acc:MGI:1929 | 97,40 | 52,4 |
| ENSMUSG00000042745 | 366,7859004 | 635,1 | 110,7 | 4,441513155 | 0,033753697 | Id1 | inhibitor of DNA binding 1 [Source:MGI Symbol;Acc:MGI:96396] | 81,01 | 37,8 |
| ENSMUSG00000022469 | 351,4399669 | 629,8 | 133,4 | 5,005476843 | 0,019553887 | Rapgef3 | Rap guanine nucleotide exchange factor (GEF) 3 [Source:MGI Symbol;Acc:MGI:2441741] | 95,18 | 25,5 |
| ENSMUSG00000031600 | 427,4722554 | 629,8 | 39,3 | 8,059793014 | 0,000300739 | Vps37a | vacuolar protein sorting 37A (yeast) [Source:MGI Symbol;Acc:MGI:1261835] | 99,42 | 5,0 |
| ENSMUSG00000007338 | 448,2562701 | 629,6 | 202,4 | 1,657096996 | 0,019933848 | Mrpl49 | mitochondrial ribosomal protein L49 [Source:MGI Symbol;Acc:MGI:108180] | 100,00 | 83,4 |
| ENSMUSG00000031980 | 458,2439055 | 628,4 | 59,6 | 4,630712684 | 0,011193495 | Agt | giotensinogen (serpin peptidase inhibitor, clade A, member 8) [Source:MGI Symbol;Acc:MGI:879 | 97,21 | 33,8 |
| ENSMUSG00000030774 | 356,4816932 | 628,2 | 163,6 | 5,6054888 | 0,000581979 | Pak1 | p21 protein (Cdc42/Rac)-activated kinase 1 [Source:MGI Symbol;Acc:MGI:1339975] | 94,44 | 27,3 |
| ENSMUSG00000058325 | 344,9268633 | 623,5 | 310,3 | 6,930938041 | 0,001089664 | Dock1 | dedicator of cytokinesis 1 [Source:MGI Symbol;Acc:MGI:2429765] | 99,99 | 28,0 |
| ENSMUSG00000034168 | 325,1908621 | 623,2 | 39,1 | 4,893084073 | 0,000387247 | Irf2bpl | interferon regulatory factor 2 binding protein-like [Source:MGI Symbol;Acc:MGI:2442463] | 98,37 | 12,1 |
| ENSMUSG00000000838 | 479,110113 | 623,1 | 85,7 | 5,242950626 | 0,005424796 | Fmr1 | fragile X mental retardation syndrome 1 [Source:MGI Symbol;Acc:MGI:95564] | 98,27 | 13,3 |
| ENSMUSG00000059482 | 365,9242273 | 621,7 | 76,3 | 8,538786434 | 0,002402222 | 2610301B20Rik | RIKEN cDNA 2610301B20 gene [Source:MGI Symbol;Acc:MGI:1914407] | 99,37 | 25,5 |
| ENSMUSG00000027605 | 413,427811 | 618,6 | 33,9 | 4,458938397 | 0,009417009 | Acss2 | acyl-CoA synthetase short-chain family member 2 [Source:MGI Symbol;Acc:MGI:1890410] | 75,99 | 3,3 |
| ENSMUSG00000025766 | 523,609447 | 616,3 | 160,8 | 2,65247345 | 0,026344109 | D3Ertd751e | DNA segment, Chr 3, ERATO Doi 751, expressed [Source:MGI Symbol;Acc:MGI:1289213] | 82,24 | 26,1 |
| ENSMUSG00000029684 | 356,7173895 | 616,0 | 19,8 | 6,734072051 | 0,000212251 | Wasl | Wiskott-Aldrich syndrome-like (human) [Source:MGI Symbol;Acc:MGI:1920428] | 87,15 | 3,6 |
| ENSMUSG00000079478 | 512,834649 | 613,9 | 150,0 | 2,232782898 | 0,009806038 | Sssca1 | ren's syndrome/scleroderma autoantigen 1 homolog (human) [Source:MGI Symbol;Acc:MGI:1913 | 92,57 | 76,8 |
| ENSMUSG00000032867 | 414,2152627 | 611,5 | 365,8 | 6,323568615 | 0,002993446 | Fbxw8 | F-box and WD-40 domain protein 8 [Source:MGI Symbol;Acc:MGI:1923041] | 93,47 | 22,7 |
| ENSMUSG00000024875 | 471,9737984 | 610,7 | 195,3 | 1,832398626 | 0,035785983 | Yif1a | Yip1 interacting factor homolog A (S. cerevisiae) [Source:MGI Symbol;Acc:MGI:1915340] | 100,00 | 65,4 |
| ENSMUSG00000035770 | 551,5439822 | 608,9 | 165,9 | 2,004989817 | 0,032713879 | Dync1li2 | dynein, cytoplasmic 1 light intermediate chain 2 [Source:MGI Symbol;Acc:MGI:107738] | 97,68 | 31,8 |
| ENSMUSG00000029148 | 457,2027418 | 608,8 | 113,8 | 6,3891806 | 7,31758E-05 | Nrbp1 | nuclear receptor binding protein 1 [Source:MGI Symbol;Acc:MGI:2183436] | 99,51 | 31,2 |
| ENSMUSG00000066150 | 597,3780349 | 608,3 | 7,4 | 8,588241999 | 0,000133309 | Slc31a1 | solute carrier family 31, member 1 [Source:MGI Symbol;Acc:MGI:1333843] | 87,30 | 1,4 |
| ENSMUSG00000093574 | 399,708487 | 607,9 | 94,6 | 3,717471292 | 0,01949513 | Gm20671 | predicted gene 20671 [Source:MGI Symbol;Acc:MGI:5313118] | 80,53 | 39,8 |
| ENSMUSG00000041353 | 501,294939 | 607,4 | 120,1 | 8,086509012 | 0,000170322 | Tmem29 | transmembrane protein 29 [Source:MGI Symbol;Acc:MGI:1923420] | 73,18 | 22,4 |
| ENSMUSG00000031568 | 375,1032714 | 606,7 | 122,3 | 3,896733059 | 0,03063669 | Rwdd4a | RWD domain containing 4A [Source:MGI Symbol;Acc:MGI:2681000] | 99,94 | 29,6 |
| ENSMUSG00000048450 | 425,6923305 | 605,8 | 26,8 | 9,045793952 | 0,000942143 | Msx1 | msh homeobox 1 [Source:MGI Symbol;Acc:MGI:97168] | 99,14 | 16,3 |
| ENSMUSG00000032615 | 420,9453983 | 605,7 | 207,3 | 2,315520156 | 0,035956226 | Nt5m | 5',3'-nucleotidase, mitochondrial [Source:MGI Symbol;Acc:MGI:1917127] | 100,00 | 52,1 |
| ENSMUSG00000007987 | 391,3506562 | 605,1 | 60,6 | 4,104220322 | 0,015569151 | Ift22 | intraflagellar transport 22 [Source:MGI Symbol;Acc:MGI:1914536] | 98,30 | 15,1 |
| ENSMUSG00000028159 | 351,9704584 | 602,7 | 0,0 | 12,58330101 | 0,000575064 | Dapp1 | ual adaptor for phosphotyrosine and 3-phosphoinositides 1 [Source:MGI Symbol;Acc:MGI:134706 | 99,68 | 0,0 |
| ENSMUSG00000046818 | 327,0055694 | 601,4 | 2,0 | 9,358582851 | 0,000213819 | Ddit4l | DNA-damage-inducible transcript 4-like [Source:MGI Symbol;Acc:MGI:1920534] | 97,57 | 0,8 |
| ENSMUSG00000047205 | 361,6723135 | 600,2 | 0,0 | 12,62255679 | 0,001385088 | Dusp18 | dual specificity phosphatase 18 [Source:MGI Symbol;Acc:MGI:1922469] | 99,86 | 0,0 |
| ENSMUSG00000054894 | 343,2835097 | 596,0 | 96,2 | 5,931093464 | 0,019015056 | Atp5s | synthase, H+ transporting, mitochondrial F0 complex, subunit S [Source:MGI Symbol;Acc:MGI:191 | 100,00 | 17,4 |
| ENSMUSG00000000202 | 403,309056 | 595,1 | 110,3 | 6,362865499 | 0,005518094 | Btbd17 | BTB (POZ) domain containing 17 [Source:MGI Symbol;Acc:MGI:1919264] | 99,94 | 29,6 |
| ENSMUSG00000027339 | 309,5512202 | 593,4 | 57,1 | 6,392745356 | 0,000131252 | Rassf2 | Ras association (RalGDS/AF-6) domain family member 2 [Source:MGI Symbol;Acc:MGI:2442060 | 75,17 | 5,4 |
| ENSMUSG00000036775 | 320,1384869 | 591,5 | 139,7 | 8,566002616 | 0,000525404 | Decr2 | 2-4-dienoyl-Coenzyme A reductase 2, peroxisomal [Source:MGI Symbol;Acc:MGI:1347059] | 99,59 | 28,6 |
| ENSMUSG00000052751 | 342,57704 | 590,6 | 157,5 | 4,734354113 | 0,025306745 | Repin1 | replication initiator 1 [Source:MGI Symbol;Acc:MGI:1889817] | 99,63 | 31,6 |
| ENSMUSG00000003585 | 383,2636972 | 589,5 | 79,0 | 4,197100972 | 0,031195337 | Sec14l2 | SEC14-like lipid binding 2 [Source:MGI Symbol;Acc:MGI:1915065] | 98,75 | 28,5 |
| ENSMUSG00000031591 | 346,301213 | 588,2 | 0,0 | 12,49748319 | 0,001283295 | Asah1 | N-acylsphingosine amidohydrolase 1 [Source:MGI Symbol;Acc:MGI:1277124] | 86,65 | 0,0 |
| ENSMUSG00000027495 | 400,3709525 | 587,7 | 23,9 | 8,443161691 | 0,000114954 | Fam210b | family with sequence similarity 210, member B [Source:MGI Symbol;Acc:MGI:1914267] | 99,71 | 3,7 |
| ENSMUSG00000005882 | 376,0408137 | 587,3 | 54,8 | 4,038164113 | 0,009961272 | Uqcc1 | iquinol-cytochrome c reductase complex assembly factor 1 [Source:MGI Symbol;Acc:MGI:192947 | 91,25 | 20,0 |
| ENSMUSG00000045817 | 302,8993588 | 584,1 | 50,6 | 4,521790191 | 0,008002308 | Zfp36l2 | zinc finger protein 36, C3H type-like 2 [Source:MGI Symbol;Acc:MGI:107945] | 82,67 | 17,5 |
| ENSMUSG00000026851 | 381,7554733 | 582,3 | 90,1 | 2,894196241 | 0,003820711 | BC005624 | cDNA sequence BC005624 [Source:MGI Symbol;Acc:MGI:2385132] | 95,03 | 33,0 |
| ENSMUSG00000038370 | 466,3617072 | 581,5 | 154,7 | 8,298082965 | 0,002876863 | Pcp4l1 | Purkinje cell protein 4-like 1 [Source:MGI Symbol;Acc:MGI:1913675] | 99,34 | 28,5 |
| ENSMUSG00000021364 | 295,1918566 | 581,2 | 1,5 | 8,672692951 | 0,000104656 | Elovl2 | of very long chain fatty acids (FEN1/Elo2, SUR4/Elo3, yeast)-like 2 [Source:MGI Symbol;Acc:MG | 96,73 | 0,7 |
| ENSMUSG00000043969 | 369,7379423 | 579,7 | 215,1 | 3,485824536 | 0,037609424 | Emx2 | empty spiracles homeobox 2 [Source:MGI Symbol;Acc:MGI:95388] | 97,60 | 30,2 |
| ENSMUSG00000047866 | 330,6547435 | 578,9 | 175,5 | 8,056079936 | 0,001907102 | Lonp2 | lon peptidase 2, peroxisomal [Source:MGI Symbol;Acc:MGI:1914137] | 97,11 | 31,7 |
| ENSMUSG00000027030 | 564,9515063 | 577,3 | 9,9 | 5,841018099 | 6,8013E-05 | Stk39 | serine/threonine kinase 39 [Source:MGI Symbol;Acc:MGI:1858416] | 92,69 | 2,6 |
| ENSMUSG00000098950 | 390,4451128 | 576,3 | 33,0 | 4,697026559 | 1,21917E-05 | Gm28036 | predicted gene, 28036 [Source:MGI Symbol;Acc:MGI:5547772] | 100,00 | 0,0 |
| ENSMUSG00000028576 | 350,1474769 | 575,5 | 72,1 | 3,995197818 | 0,019258586 | Ift74 | intraflagellar transport 74 [Source:MGI Symbol;Acc:MGI:1914944] | 82,16 | 22,7 |
| ENSMUSG00000104208 | 416,0432738 | 575,3 | 202,3 | 1,605226905 | 0,029520617 | Gm37401 | predicted gene, 37401 [Source:MGI Symbol;Acc:MGI:5610629] | 89,53 | 79,0 |
| ENSMUSG00000018340 | 554,1598538 | 572,1 | 149,4 | 2,835073367 | 0,006043248 | Anxa6 | annexin A6 [Source:MGI Symbol;Acc:MGI:88255] | 100,00 | 30,1 |
| ENSMUSG00000029201 | 335,0432402 | 570,3 | 286,9 | 8,230989276 | 0,001049358 | Ugdh | UDP-glucose dehydrogenase [Source:MGI Symbol;Acc:MGI:1306785] | 93,33 | 31,1 |
| ENSMUSG00000033022 | 344,6975024 | 567,6 | 0,0 | 12,52759032 | 0,0003463 | Cdo1 | cysteine dioxygenase 1, cytosolic [Source:MGI Symbol;Acc:MGI:105925] | 100,00 | 0,0 |
| ENSMUSG00000022091 | 330,3765287 | 567,1 | 113,8 | 8,243604098 | 0,001787001 | Sorbs3 | sorbin and SH3 domain containing 3 [Source:MGI Symbol;Acc:MGI:700013] | 97,60 | 23,8 |
| ENSMUSG00000042506 | 415,6496038 | 566,7 | 78,2 | 2,88243317 | 0,001135361 | Usp22 | ubiquitin specific peptidase 22 [Source:MGI Symbol;Acc:MGI:2144157] | 99,54 | 16,8 |
| ENSMUSG00000059811 | 416,3495456 | 565,4 | 138,6 | 7,779855946 | 0,000285361 | Atl2 | atlastin GTPase 2 [Source:MGI Symbol;Acc:MGI:1929492] | 99,10 | 17,5 |
| ENSMUSG00000058594 | 403,4253431 | 564,5 | 126,4 | 3,4323884 | 0,030993462 | Fbxo18 | F-box protein 18 [Source:MGI Symbol;Acc:MGI:1354699] | 98,94 | 36,2 |

| | | | | | | | | | |
|---|---|---|---|---|---|---|---|---|---|
| ENSMUSG00000025609 | 343,2243945 | 562,0 | 57,8 | 4,578406134 | 0,01521624 | Mkln1 | muskelin 1, intracellular mediator containing kelch motifs [Source:MGI Symbol;Acc:MGI:1351638] | 72,41 | 3,5 |
| ENSMUSG00000068747 | 350,1364497 | 561,3 | 92,7 | 4,785509058 | 0,002883434 | Sort1 | sortilin 1 [Source:MGI Symbol;Acc:MGI:1338015] | 99,11 | 14,5 |
| ENSMUSG00000022537 | 438,6408429 | 560,2 | 342,5 | 8,893771657 | 0,007146733 | Tmem44 | transmembrane protein 44 [Source:MGI Symbol;Acc:MGI:1924489] | 99,90 | 28,8 |
| ENSMUSG00000046519 | 339,1168622 | 558,3 | 469,0 | 8,844541281 | 0,001463867 | Golph3l | golgi phosphoprotein 3-like [Source:MGI Symbol;Acc:MGI:1917129] | 99,86 | 32,6 |
| ENSMUSG00000047694 | 365,9985655 | 558,1 | 0,0 | 12,52148051 | 0,001908984 | Yipf6 | Yip1 domain family, member 6 [Source:MGI Symbol;Acc:MGI:1925179] | 97,21 | 0,0 |
| ENSMUSG00000019970 | 466,5309259 | 556,9 | 202,2 | 4,404179458 | 0,038497541 | Sgk1 | serum/glucocorticoid regulated kinase 1 [Source:MGI Symbol;Acc:MGI:1340062] | 78,21 | 20,9 |
| ENSMUSG00000052456 | 376,9891458 | 555,5 | 132,8 | 2,175549671 | 0,00331406 | Asna1 | sA arsenite transporter, ATP-binding, homolog 1 (bacterial) [Source:MGI Symbol;Acc:MGI:192837 | 99,15 | 76,3 |
| ENSMUSG00000053646 | 376,1360993 | 553,7 | 179,9 | 6,258723918 | 0,002856502 | Plxnb1 | plexin B1 [Source:MGI Symbol;Acc:MGI:2154238] | 94,11 | 18,3 |
| ENSMUSG00000019837 | 345,7396928 | 552,2 | 91,3 | 3,433964336 | 0,04497998 | Gtf3c6 | general transcription factor IIIC, polypeptide 6, alpha [Source:MGI Symbol;Acc:MGI:1914621] | 100,00 | 55,6 |
| ENSMUSG00000027285 | 380,2655836 | 551,0 | 65,7 | 6,492643241 | 0,000762765 | Haus2 | HAUS augmin-like complex, subunit 2 [Source:MGI Symbol;Acc:MGI:1913546] | 98,17 | 18,4 |
| ENSMUSG00000033610 | 312,383273 | 549,2 | 173,0 | 3,94370195 | 0,032722538 | Pank1 | pantothenate kinase 1 [Source:MGI Symbol;Acc:MGI:1922985] | 98,25 | 19,7 |
| ENSMUSG00000000902 | 319,7287086 | 548,8 | 88,7 | 8,202238516 | 0,000393615 | Smarcb1 | trix associated, actin dependent regulator of chromatin, subfamily b, member 1 [Source:MGI Symb | 99,90 | 25,8 |
| ENSMUSG00000037697 | 425,0357441 | 548,6 | 280,6 | 5,305866348 | 0,005026681 | Ddhd1 | DDHD domain containing 1 [Source:MGI Symbol;Acc:MGI:2150302] | 98,94 | 21,2 |
| ENSMUSG00000024603 | 380,7942734 | 547,3 | 83,7 | 3,577209313 | 0,008733864 | Dctn4 | dynactin 4 [Source:MGI Symbol;Acc:MGI:1914915] | 99,94 | 13,5 |
| ENSMUSG00000009216 | 285,2225561 | 546,6 | 8,8 | 6,28684634 | 1,92458E-05 | Fam163b | family with sequence similarity 163, member B [Source:MGI Symbol;Acc:MGI:1926106] | 99,22 | 3,4 |
| ENSMUSG00000030986 | 366,1728408 | 546,1 | 94,2 | 2,86289479 | 0,020710182 | Dhx32 | DEAH (Asp-Glu-Ala-His) box polypeptide 32 [Source:MGI Symbol;Acc:MGI:2141813] | 84,68 | 6,1 |
| ENSMUSG00000020432 | 341,3501992 | 546,1 | 22,4 | 8,69109972 | 0,002117304 | Tcn2 | transcobalamin 2 [Source:MGI Symbol;Acc:MGI:98534] | 93,88 | 10,9 |
| ENSMUSG00000021010 | 303,8941622 | 544,9 | 183,1 | 2,914481234 | 0,028723909 | Npas3 | neuronal PAS domain protein 3 [Source:MGI Symbol;Acc:MGI:1351610] | 99,79 | 22,9 |
| ENSMUSG00000033751 | 369,530322 | 542,8 | 225,4 | 1,288144208 | 0,049415649 | Gadd45gip1 | arrest and DNA-damage-inducible, gamma interacting protein 1 [Source:MGI Symbol;Acc:MGI:19 | 74,57 | 40,7 |
| ENSMUSG00000009394 | 367,3098017 | 542,3 | 100,6 | 4,040948312 | 0,018744531 | Syn2 | synapsin II [Source:MGI Symbol;Acc:MGI:103020] | 98,84 | 16,6 |
| ENSMUSG00000024789 | 330,8457761 | 542,2 | 9,8 | 6,128699089 | 0,000928068 | Jak2 | Janus kinase 2 [Source:MGI Symbol;Acc:MGI:96629] | 99,62 | 2,1 |
| ENSMUSG00000024843 | 351,6647078 | 541,6 | 98,0 | 7,869335012 | 7,23739E-05 | Chka | choline kinase alpha [Source:MGI Symbol;Acc:MGI:107760] | 95,64 | 23,7 |
| ENSMUSG00000024826 | 442,1781409 | 541,3 | 201,7 | 2,185263437 | 0,038936575 | Dpf2 | D4, zinc and double PHD fingers family 2 [Source:MGI Symbol;Acc:MGI:109529] | 81,65 | 37,5 |
| ENSMUSG00000063802 | 326,2606295 | 540,7 | 56,3 | 7,686561654 | 0,000339105 | Hspbp1 | (heat shock 70kDa) binding protein, cytoplasmic cochaperone 1 [Source:MGI Symbol;Acc:MGI:19 | 93,57 | 23,4 |
| ENSMUSG00000109336 | 327,4332195 | 539,9 | 131,9 | 7,499087225 | 0,000218895 | Samd4b | sterile alpha motif domain containing 4B [Source:MGI Symbol;Acc:MGI:2448542] | 98,14 | 23,7 |
| ENSMUSG00000005505 | 433,2811363 | 539,5 | 206,2 | 1,423373462 | 0,023822826 | Kbtbd4 | kelch repeat and BTB (POZ) domain containing 4 [Source:MGI Symbol;Acc:MGI:1914386] | 99,63 | 8,8 |
| ENSMUSG00000052957 | 323,3520274 | 537,8 | 3,7 | 8,83932191 | 0,000291495 | Gas1 | growth arrest specific 1 [Source:MGI Symbol;Acc:MGI:95655] | 99,95 | 1,9 |
| ENSMUSG00000022382 | 336,6156626 | 537,0 | 52,4 | 5,937932335 | 0,006161154 | Wnt7b | wingless-type MMTV integration site family, member 7B [Source:MGI Symbol;Acc:MGI:98962] | 81,63 | 13,1 |
| ENSMUSG00000014361 | 362,5514549 | 536,3 | 0,9 | 9,884057267 | 0,000939386 | Mertk | c-mer proto-oncogene tyrosine kinase [Source:MGI Symbol;Acc:MGI:96965] | 100,00 | 0,5 |
| ENSMUSG00000097675 | 312,5648649 | 535,6 | 111,8 | 2,478936215 | 0,009892318 | 1700101I11Rik | RIKEN cDNA 1700101I11 gene [Source:MGI Symbol;Acc:MGI:1915180] | 71,18 | 22,9 |
| ENSMUSG00000021760 | 346,8294753 | 535,4 | 77,0 | 3,702796559 | 0,036798784 | Gpx8 | glutathione peroxidase 8 (putative) [Source:MGI Symbol;Acc:MGI:1916840] | 100,00 | 52,6 |
| ENSMUSG00000068099 | 350,597587 | 533,9 | 162,2 | 8,220573823 | 0,001196977 | 1500009C09Rik | RIKEN cDNA 1500009C09 gene [Source:MGI Symbol;Acc:MGI:1923755] | 78,82 | 25,2 |
| ENSMUSG00000020601 | 303,0988371 | 533,7 | 129,7 | 8,431847373 | 0,000359804 | Trib2 | tribbles homolog 2 (Drosophila) [Source:MGI Symbol;Acc:MGI:2145021] | 99,95 | 15,7 |
| ENSMUSG00000022562 | 344,1642148 | 531,4 | 34,8 | 3,890070123 | 0,000263846 | Oplah | 5-oxoprolinase (ATP-hydrolysing) [Source:MGI Symbol;Acc:MGI:1922725] | 91,11 | 6,3 |
| ENSMUSG00000010290 | 555,863795 | 530,5 | 249,1 | 6,927258229 | 0,001631726 | AI597479 | expressed sequence AI597479 [Source:MGI Symbol;Acc:MGI:2138299] | 99,65 | 30,3 |
| ENSMUSG00000021591 | 305,501265 | 529,1 | 39,9 | 8,590388754 | 0,000652685 | Glrx | glutaredoxin [Source:MGI Symbol;Acc:MGI:2135625] | 73,80 | 18,4 |
| ENSMUSG00000053141 | 387,8599904 | 529,0 | 391,6 | 8,486607887 | 0,003510352 | Ptprt | protein tyrosine phosphatase, receptor type, T [Source:MGI Symbol;Acc:MGI:1321152] | 78,49 | 18,0 |
| ENSMUSG00000022175 | 371,4231996 | 528,5 | 56,4 | 6,572750112 | 0,00012881 | Lrp10 | low-density lipoprotein receptor-related protein 10 [Source:MGI Symbol;Acc:MGI:1929480] | 75,30 | 15,5 |
| ENSMUSG00000039917 | 378,5573085 | 528,5 | 133,5 | 6,398213727 | 0,000756646 | Rhbdd2 | rhomboid domain containing 2 [Source:MGI Symbol;Acc:MGI:1915612] | 96,06 | 18,5 |
| ENSMUSG00000028035 | 362,6037309 | 527,4 | 124,8 | 6,513107715 | 0,007303788 | Dnajb4 | DnaJ heat shock protein family (Hsp40) member B4 [Source:MGI Symbol;Acc:MGI:1914285] | 91,57 | 25,3 |
| ENSMUSG00000038170 | 280,8077427 | 526,8 | 80,8 | 4,414207482 | 0,005219865 | Pde4dip | phosphodiesterase 4D interacting protein (myomegalin) [Source:MGI Symbol;Acc:MGI:1891434] | 78,82 | 4,5 |
| ENSMUSG00000085838 | 302,9191447 | 526,2 | 79,3 | 5,691606114 | 0,001198022 | Chn1os1 | chimerin 1, opposite strand 1 [Source:MGI Symbol;Acc:MGI:3702041] | 91,98 | 34,7 |
| ENSMUSG00000070934 | 275,9281599 | 524,6 | 0,0 | 12,39166006 | 0,001239038 | Rraga | Ras-related GTP binding A [Source:MGI Symbol;Acc:MGI:1915691] | 100,00 | 0,0 |
| ENSMUSG00000031639 | 289,5384572 | 523,7 | 0,6 | 9,652978759 | 0,000253141 | Tlr3 | toll-like receptor 3 [Source:MGI Symbol;Acc:MGI:2156367] | 98,10 | 0,5 |
| ENSMUSG00000039952 | 342,2513054 | 522,1 | 46,1 | 6,357333701 | 0,006961964 | Dag1 | dystroglycan 1 [Source:MGI Symbol;Acc:MGI:101864] | 91,94 | 5,4 |
| ENSMUSG00000028444 | 319,3522002 | 520,3 | 76,0 | 4,261566469 | 0,009023159 | Cntfr | ciliary neurotrophic factor receptor [Source:MGI Symbol;Acc:MGI:99605] | 83,98 | 27,7 |
| ENSMUSG00000039686 | 282,9111432 | 518,6 | 22,2 | 7,552399183 | 1,683E-05 | Zer1 | zyg-11 related, cell cycle regulator [Source:MGI Symbol;Acc:MGI:2442511] | 95,56 | 7,6 |
| ENSMUSG00000022607 | 314,2304779 | 517,9 | 38,9 | 4,306253034 | 0,002238453 | Ptk2 | PTK2 protein tyrosine kinase 2 [Source:MGI Symbol;Acc:MGI:95481] | 90,15 | 8,9 |
| ENSMUSG00000037643 | 267,3738268 | 517,7 | 37,1 | 5,844163516 | 0,002774311 | Prkci | protein kinase C, iota [Source:MGI Symbol;Acc:MGI:99260] | 91,24 | 5,7 |
| ENSMUSG00000025059 | 325,5065604 | 517,7 | 184,5 | 4,834950967 | 0,011804736 | Gk | glycerol kinase [Source:MGI Symbol;Acc:MGI:106594] | 81,49 | 23,5 |
| ENSMUSG00000034152 | 342,9016649 | 516,6 | 9,2 | 8,287650719 | 0,000166554 | Exoc3 | exocyst complex component 3 [Source:MGI Symbol;Acc:MGI:2443972] | 99,95 | 1,9 |
| ENSMUSG00000028527 | 331,5419051 | 515,7 | 97,8 | 4,507246676 | 0,017796356 | Ak4 | adenylate kinase 4 [Source:MGI Symbol;Acc:MGI:87979] | 90,31 | 21,0 |
| ENSMUSG00000048644 | 347,2326346 | 514,3 | 191,7 | 2,07021208 | 0,041767397 | Ctxn1 | cortexin 1 [Source:MGI Symbol;Acc:MGI:88566] | 97,47 | 72,4 |
| ENSMUSG00000003559 | 333,3217482 | 514,2 | 44,2 | 8,393528798 | 0,002857421 | As3mt | arsenic (+3 oxidation state) methyltransferase [Source:MGI Symbol;Acc:MGI:1929882] | 99,70 | 23,3 |
| ENSMUSG00000019960 | 301,1396462 | 514,0 | 40,1 | 5,829645373 | 0,006792606 | Dusp6 | dual specificity phosphatase 6 [Source:MGI Symbol;Acc:MGI:1914853] | 99,81 | 10,7 |
| ENSMUSG00000003420 | 344,7461987 | 513,9 | 28,9 | 8,336766772 | 0,00123279 | Fcgrt | Fc receptor, IgG, alpha chain transporter [Source:MGI Symbol;Acc:MGI:103017] | 99,52 | 11,3 |
| ENSMUSG00000021646 | 257,9132942 | 513,6 | 75,9 | 6,635843429 | 0,00489173 | Mccc2 | methylcrotonoyl-Coenzyme A carboxylase 2 (beta) [Source:MGI Symbol;Acc:MGI:1925288] | 100,00 | 31,3 |
| ENSMUSG00000028191 | 384,1904572 | 512,8 | 175,0 | 2,582263178 | 0,043968139 | Bcl10 | B cell leukemia/lymphoma 10 [Source:MGI Symbol;Acc:MGI:1337994] | 99,06 | 41,5 |
| ENSMUSG00000000223 | 282,8210578 | 510,9 | 46,9 | 4,635150507 | 0,009687502 | Drp2 | dystrophin related protein 2 [Source:MGI Symbol;Acc:MGI:107432] | 96,54 | 4,8 |
| ENSMUSG00000042705 | 325,3099971 | 510,9 | 79,3 | 8,389229078 | 0,00337462 | Commd10 | COMM domain containing 10 [Source:MGI Symbol;Acc:MGI:1916706] | 99,96 | 30,8 |
| ENSMUSG00000024642 | 355,2898146 | 510,8 | 230,7 | 3,750901666 | 0,046232076 | Tle4 | nsducin-like enhancer of split 4, homolog of Drosophila E(spl) [Source:MGI Symbol;Acc:MGI:1046 | 97,78 | 29,7 |
| ENSMUSG00000031584 | 352,0166391 | 510,2 | 314,3 | 6,378094559 | 0,002535368 | Gsr | glutathione reductase [Source:MGI Symbol;Acc:MGI:95804] | 93,93 | 29,4 |
| ENSMUSG00000040003 | 318,1989736 | 510,1 | 115,4 | 3,367948324 | 0,016973227 | Magi2 | e associated guanylate kinase, WW and PDZ domain containing 2 [Source:MGI Symbol;Acc:MGI: | 84,07 | 14,7 |
| ENSMUSG00000039911 | 270,6228466 | 508,7 | 59,1 | 8,664307178 | 0,001901811 | Spsb1 | olA/ryanodine receptor domain and SOCS box containing 1 [Source:MGI Symbol;Acc:MGI:192189 | 74,01 | 14,7 |
| ENSMUSG00000022055 | 342,6120833 | 507,1 | 263,9 | 3,943360753 | 0,020000617 | Nefl | neurofilament, light polypeptide [Source:MGI Symbol;Acc:MGI:97313] | 99,14 | 32,0 |
| ENSMUSG00000021823 | 229,8978689 | 505,0 | 0,6 | 9,759854 | 0,0001986 | Vcl | vinculin [Source:MGI Symbol;Acc:MGI:98927] | 99,56 | 0,3 |
| ENSMUSG00000049076 | 243,200996 | 504,6 | 71,2 | 8,420934652 | 0,001743956 | Acap2 | ArfGAP with coiled-coil, ankyrin repeat and PH domains 2 [Source:MGI Symbol;Acc:MGI:1925868 | 98,56 | 6,3 |
| ENSMUSG00000031875 | 271,1529784 | 503,5 | 35,1 | 8,037232866 | 0,000262805 | Cmtm3 | CKLF-like MARVEL transmembrane domain containing 3 [Source:MGI Symbol;Acc:MGI:2447162 | 98,16 | 21,8 |
| ENSMUSG00000061838 | 376,5989317 | 503,0 | 18,1 | 5,39460277 | 0,00015953 | Suclg2 | succinate-Coenzyme A ligase, GDP-forming, beta subunit [Source:MGI Symbol;Acc:MGI:1306824 | 72,70 | 4,4 |

| | | | | | | | | | |
|---|---|---|---|---|---|---|---|---|---|
| ENSMUSG00000025477 | 454,3526982 | 502,8 | 160,8 | 3,221558502 | 0,023648568 | Inpp5a | inositol polyphosphate-5-phosphatase A [Source:MGI Symbol;Acc:MGI:2686961] | 91,04 | 36,8 |
| ENSMUSG00000035469 | 326,3044668 | 501,8 | 8,0 | 6,815395039 | 2,70315E-08 | Rcbtb1 | ıosome condensation (RCC1) and BTB (POZ) domain containing protein 1 [Source:MGI Symbol;A | 92,93 | 2,2 |
| ENSMUSG00000028926 | 322,2874965 | 500,7 | 3,0 | 7,994910712 | 1,10834E-06 | Cdk14 | cyclin-dependent kinase 14 [Source:MGI Symbol;Acc:MGI:894318] | 89,83 | 1,6 |
| ENSMUSG00000038776 | 266,5702175 | 500,4 | 36,9 | 8,159807141 | 0,000392863 | Ephx1 | epoxide hydrolase 1, microsomal [Source:MGI Symbol;Acc:MGI:95405] | 87,71 | 17,5 |
| ENSMUSG00000021779 | 329,4712778 | 500,2 | 73,2 | 8,110933763 | 0,001820656 | Thrb | thyroid hormone receptor beta [Source:MGI Symbol;Acc:MGI:98743] | 91,36 | 12,5 |
| ENSMUSG00000040964 | 296,0030253 | 500,2 | 153,6 | 6,766314641 | 0,002669967 | Arhgef10l | Rho guanine nucleotide exchange factor (GEF) 10-like [Source:MGI Symbol;Acc:MGI:1920004] | 82,40 | 21,5 |

Table S2.6

| Id | baseMean Read | Mean read Astrocyte | Mean read PAP | log2FoldChange | padj | Associated Gene Name | Description | Mean Astrocyte Exon Coverage | Mean PAP exon Coverage |
|---|---|---|---|---|---|---|---|---|---|
| ENSMUSG00000037852 | 57312,24097 | 77322,8 | 36749,6 | 1,163246478 | 0,32792602 | Cpe | carboxypeptidase E [Source:MGI Symbol;Acc:MGI:101932] | 100,00 | 100,0 |
| ENSMUSG00000061808 | 52620,12161 | 53183,3 | 25063,3 | 1,171355253 | 0,46951909 | Ttr | transthyretin [Source:MGI Symbol;Acc:MGI:98865] | 100,00 | 95,5 |
| ENSMUSG00000027447 | 53363,6157 | 86625,9 | 24711,3 | 2,092676124 | 0,0807423 | Cst3 | cystatin C [Source:MGI Symbol;Acc:MGI:102519] | 91,68 | 85,1 |
| ENSMUSG00000031765 | 39617,59049 | 62434,7 | 23512,3 | 1,602052255 | 0,0648659 | Mt1 | metallothionein 1 [Source:MGI Symbol;Acc:MGI:97171] | 100,00 | 100,0 |
| ENSMUSG00000020932 | 31548,84248 | 51709,3 | 15244,8 | 1,957259788 | 0,06600305 | Gfap | glial fibrillary acidic protein [Source:MGI Symbol;Acc:MGI:95697] | 99,40 | 86,7 |
| ENSMUSG00000025428 | 8142,776329 | 4941,1 | 11614,7 | -1,238136255 | 0,05031008 | Atp5a1 | ATP synthase, H+ transporting, mitochondrial F1 complex, alpha subunit 1 [Source:MGI Symbol;Acc:MGI:88115] | 86,89 | 84,6 |
| ENSMUSG00000031342 | 15121,5718 | 19934,3 | 11195,8 | 1,040582708 | 0,42887772 | Gpm6b | glycoprotein m6b [Source:MGI Symbol;Acc:MGI:107672] | 99,91 | 97,9 |
| ENSMUSG00000025203 | 11253,57196 | 13344,1 | 9057,2 | 0,556554773 | 0,51243503 | Scd2 | stearoyl-Coenzyme A desaturase 2 [Source:MGI Symbol;Acc:MGI:98240] | 95,31 | 92,1 |
| ENSMUSG00000032199 | 5596,595083 | 4537,2 | 7673,8 | -0,759339581 | 0,2890684 | Polr2m | polymerase (RNA) II (DNA directed) polypeptide M [Source:MGI Symbol;Acc:MGI:107282] | 100,00 | 92,9 |
| ENSMUSG00000005610 | 10862,8587 | 14563,6 | 7064,9 | 1,125189155 | 0,16143564 | Eif4g2 | eukaryotic translation initiation factor 4, gamma 2 [Source:MGI Symbol;Acc:MGI:109207] | 87,19 | 77,1 |
| ENSMUSG00000031760 | 13317,19866 | 21793,1 | 6913,5 | 2,141105348 | 0,06935742 | Mt3 | metallothionein 3 [Source:MGI Symbol;Acc:MGI:97173] | 100,00 | 81,1 |
| ENSMUSG00000053279 | 9750,086592 | 14626,8 | 6567,6 | 1,656826701 | 0,39325364 | Aldh1a1 | aldehyde dehydrogenase family 1, subfamily A1 [Source:MGI Symbol;Acc:MGI:1353450] | 99,01 | 98,3 |
| ENSMUSG00000021792 | 8385,55898 | 11709,2 | 6162,4 | 0,974402232 | 0,3953241 | Fam213a | family with sequence similarity 213, member A [Source:MGI Symbol;Acc:MGI:1917814] | 92,35 | 73,7 |
| ENSMUSG00000005360 | 12265,97181 | 19652,7 | 6135,4 | 1,644038758 | 0,13677814 | Slc1a3 | solute carrier family 1 (glial high affinity glutamate transporter), member 3 [Source:MGI Symbol;Acc:MGI:99917] | 96,68 | 92,6 |
| ENSMUSG00000054226 | 9718,047782 | 15367,4 | 6017,9 | 1,641734225 | 0,15154048 | Tprkb | Tp53rk binding protein [Source:MGI Symbol;Acc:MGI:1917036] | 99,04 | 73,3 |
| ENSMUSG00000029309 | 11281,62449 | 16389,2 | 5886,9 | 1,610284017 | 0,22872137 | Sparcl1 | SPARC-like 1 [Source:MGI Symbol;Acc:MGI:108110] | 100,00 | 98,0 |
| ENSMUSG00000033208 | 8165,8054 | 11166,7 | 5755,8 | 1,512001706 | 0,23913222 | S100b | S100 protein, beta polypeptide, neural [Source:MGI Symbol;Acc:MGI:98217] | 100,00 | 100,0 |
| ENSMUSG00000078812 | 4568,642193 | 3427,9 | 5701,5 | -0,68313744 | 0,25984335 | Eif5a | eukaryotic translation initiation factor 5A [Source:MGI Symbol;Acc:MGI:106248] | 73,58 | 70,3 |
| ENSMUSG00000035885 | 6980,095074 | 8039,9 | 5661,2 | 0,656734339 | 0,37985916 | Cox8a | cytochrome c oxidase subunit VIIIa [Source:MGI Symbol;Acc:MGI:105959] | 98,72 | 87,3 |
| ENSMUSG00000013698 | 8388,972128 | 11374,1 | 5516,3 | 1,631519205 | 0,1571542 | Pea15a | phosphoprotein enriched in astrocytes 15A [Source:MGI Symbol;Acc:MGI:104799] | 97,95 | 89,8 |
| ENSMUSG00000025393 | 7223,592489 | 8729,7 | 5395,0 | 0,691871118 | 0,32987156 | Atp5b | ATP synthase, H+ transporting mitochondrial F1 complex, beta subunit [Source:MGI Symbol;Acc:MGI:107801] | 99,39 | 92,9 |
| ENSMUSG00000031818 | 6676,901625 | 7165,2 | 5232,2 | 0,470349305 | 0,40692753 | Cox4i1 | cytochrome c oxidase subunit IV isoform 1 [Source:MGI Symbol;Acc:MGI:88473] | 100,00 | 96,1 |
| ENSMUSG00000055302 | 4022,145205 | 3032,4 | 5173,9 | -0,763555477 | 0,18454011 | Mrfap1 | Morf4 family associated protein 1 [Source:MGI Symbol;Acc:MGI:1914818] | 84,59 | 79,1 |
| ENSMUSG00000090625 | 8961,534035 | 11999,4 | 5150,8 | 1,308623444 | 0,07407347 | Gm20721 | predicted gene, 20721 [Source:MGI Symbol;Acc:MGI:5313030] | 99,33 | 99,1 |
| ENSMUSG00000018770 | 6509,648156 | 7550,4 | 5031,5 | 0,595964209 | 0,32320821 | Atp5g3 | ATP synthase, H+ transporting, mitochondrial F0 complex, subunit C3 (subunit 9) [Source:MGI Symbol;Acc:MGI:2442035] | 100,00 | 100,0 |
| ENSMUSG00000062078 | 5665,86184 | 7196,7 | 4984,2 | 0,788238764 | 0,46554274 | Qk | quaking [Source:MGI Symbol;Acc:MGI:97837] | 99,50 | 71,4 |
| ENSMUSG00000064373 | 6454,177762 | 8993,2 | 4823,9 | 1,501688525 | 0,44504481 | Sepp1 | selenoprotein P, plasma, 1 [Source:MGI Symbol;Acc:MGI:894288] | 89,64 | 76,7 |
| ENSMUSG00000091228 | 3352,900889 | 1971,9 | 4682,5 | -1,223479605 | 0,05780494 | Gm20390 | predicted gene 20390 [Source:MGI Symbol;Acc:MGI:5141855] | 98,59 | 97,4 |
| ENSMUSG00000027893 | 8630,853458 | 13181,8 | 4578,2 | 1,729966015 | 0,09086628 | Ahcyl1 | S-adenosylhomocysteine hydrolase-like 1 [Source:MGI Symbol;Acc:MGI:2385184] | 100,00 | 93,1 |
| ENSMUSG00000030428 | 8370,541755 | 11354,2 | 4528,4 | 1,638218466 | 0,13968168 | Ttyh1 | tweety homolog 1 (Drosophila) [Source:MGI Symbol;Acc:MGI:1889007] | 99,16 | 85,7 |
| ENSMUSG00000022425 | 13510,66339 | 12078,0 | 4468,5 | 1,784322903 | 0,25444618 | Enpp2 | ectonucleotide pyrophosphatase/phosphodiesterase 2 [Source:MGI Symbol;Acc:MGI:1321390] | 99,90 | 88,0 |
| ENSMUSG00000032294 | 4278,468205 | 3554,1 | 4460,3 | -0,326736839 | 0,60387969 | Pkm | pyruvate kinase, muscle [Source:MGI Symbol;Acc:MGI:97591] | 97,35 | 89,1 |
| ENSMUSG00000010663 | 5632,224101 | 6867,2 | 4405,5 | 0,841316243 | 0,48004355 | Fads1 | fatty acid desaturase 1 [Source:MGI Symbol;Acc:MGI:1923517] | 99,84 | 97,2 |
| ENSMUSG00000005089 | 8616,104327 | 12099,8 | 4289,4 | 1,588017366 | 0,16643545 | Slc1a2 | solute carrier family 1 (glial high affinity glutamate transporter), member 2 [Source:MGI Symbol;Acc:MGI:101931] | 99,20 | 73,5 |
| ENSMUSG00000025871 | 6064,769561 | 7817,7 | 4171,2 | 1,413692088 | 0,22441109 | 4833439L19Rik | RIKEN cDNA 4833439L19 gene [Source:MGI Symbol;Acc:MGI:1921162] | 98,96 | 94,9 |
| ENSMUSG00000020611 | 3312,652225 | 3193,9 | 3986,4 | -0,204384416 | 0,83361663 | Gna13 | guanine nucleotide binding protein, alpha 13 [Source:MGI Symbol;Acc:MGI:95768] | 99,91 | 91,8 |
| ENSMUSG00000022108 | 7831,240569 | 11491,4 | 3942,4 | 1,534048952 | 0,14633 | Itm2b | integral membrane protein 2B [Source:MGI Symbol;Acc:MGI:1309517] | 100,00 | 99,9 |
| ENSMUSG00000020267 | 3299,048301 | 2439,9 | 3940,5 | -0,687624279 | 0,34458216 | Hint1 | histidine triad nucleotide binding protein 1 [Source:MGI Symbol;Acc:MGI:1321133] | 72,97 | 72,7 |
| ENSMUSG00000020849 | 6450,488276 | 9375,3 | 3931,7 | 1,278443694 | 0,07314317 | Ywhae | ıe 3-monooxygenase/tryptophan 5-monooxygenase activation protein, epsilon polypeptide [Source:MGI Symbol;Acc:MGI:8[illegible] | 99,01 | 89,0 |
| ENSMUSG00000022890 | 5312,009684 | 7019,3 | 3876,8 | 0,902023233 | 0,14455809 | Atp5j | ATP synthase, H+ transporting, mitochondrial F0 complex, subunit F [Source:MGI Symbol;Acc:MGI:107777] | 100,00 | 71,9 |
| ENSMUSG00000028416 | 3330,597588 | 2741,6 | 3853,3 | -0,485545516 | 0,43997541 | Bag1 | BCL2-associated athanogene 1 [Source:MGI Symbol;Acc:MGI:108047] | 100,00 | 97,5 |
| ENSMUSG00000024213 | 4024,083251 | 3686,8 | 3613,4 | 0,020373869 | 0,97709704 | Nudt3 | nudix (nucleotide diphosphate linked moiety X)-type motif 3 [Source:MGI Symbol;Acc:MGI:1928484] | 93,75 | 89,4 |
| ENSMUSG00000030224 | 2482,003778 | 1771,9 | 3608,6 | -1,023145783 | 0,09952869 | Strap | serine/threonine kinase receptor associated protein [Source:MGI Symbol;Acc:MGI:1329037] | 98,96 | 96,1 |
| ENSMUSG00000059325 | 7735,943485 | 13770,0 | 3578,5 | 2,268131236 | 0,07407347 | Hopx | HOP homeobox [Source:MGI Symbol;Acc:MGI:1916782] | 100,00 | 85,5 |
| ENSMUSG00000020591 | 5944,417846 | 8823,2 | 3576,6 | 1,638038738 | 0,15381031 | Ntsr2 | neurotensin receptor 2 [Source:MGI Symbol;Acc:MGI:108018] | 99,87 | 97,8 |
| ENSMUSG00000041571 | 3883,141386 | 4154,7 | 3486,4 | 0,457496725 | 0,61410117 | Sepw1 | selenoprotein W, muscle 1 [Source:MGI Symbol;Acc:MGI:1100878] | 99,91 | 96,5 |
| ENSMUSG00000021282 | 2921,388153 | 2774,2 | 3384,4 | -0,299515121 | 0,68550249 | Eif5 | eukaryotic translation initiation factor 5 [Source:MGI Symbol;Acc:MGI:95309] | 92,49 | 86,2 |
| ENSMUSG00000007097 | 7619,125499 | 11364,3 | 3366,1 | 1,728714844 | 0,15028748 | Atp1a2 | ATPase, Na+/K+ transporting, alpha 2 polypeptide [Source:MGI Symbol;Acc:MGI:88106] | 98,03 | 86,9 |
| ENSMUSG00000063882 | 3237,633433 | 3500,0 | 3328,3 | 0,085444063 | 0,89162013 | Uqcrh | ubiquinol-cytochrome c reductase hinge protein [Source:MGI Symbol;Acc:MGI:1913826] | 100,00 | 98,4 |
| ENSMUSG00000024782 | 4912,505492 | 7240,6 | 3247,4 | 1,28267149 | 0,21735249 | Ak3 | adenylate kinase 3 [Source:MGI Symbol;Acc:MGI:1860835] | 100,00 | 97,2 |
| ENSMUSG00000031246 | 3027,896598 | 3800,3 | 3111,3 | 0,465560441 | 0,63356866 | Sh3bgrl | SH3-binding domain glutamic acid-rich protein like [Source:MGI Symbol;Acc:MGI:1930849] | 99,40 | 95,3 |
| ENSMUSG00000041740 | 1895,418392 | 1569,7 | 3067,1 | -0,967942214 | 0,16545199 | Rnf10 | ring finger protein 10 [Source:MGI Symbol;Acc:MGI:1859162] | 93,45 | 89,5 |
| ENSMUSG00000026424 | 6146,161661 | 9669,7 | 3059,1 | 1,690300552 | 0,1977193 | Gpr37l1 | G protein-coupled receptor 37-like 1 [Source:MGI Symbol;Acc:MGI:1928503] | 99,00 | 94,2 |
| ENSMUSG00000032883 | 3726,874898 | 5619,9 | 3049,8 | 0,862314234 | 0,52942551 | Acsl3 | acyl-CoA synthetase long-chain family member 3 [Source:MGI Symbol;Acc:MGI:1921455] | 99,72 | 95,6 |
| ENSMUSG00000026457 | 2536,627911 | 1607,0 | 3040,0 | -0,853569052 | 0,26648115 | Adipor1 | adiponectin receptor 1 [Source:MGI Symbol;Acc:MGI:1919924] | 97,42 | 70,7 |
| ENSMUSG00000022186 | 2932,068558 | 2988,6 | 3011,9 | 0,039103755 | 0,95622346 | Oxct1 | 3-oxoacid CoA transferase 1 [Source:MGI Symbol;Acc:MGI:1914291] | 99,57 | 95,9 |
| ENSMUSG00000029364 | 2439,460621 | 1524,8 | 2987,5 | -0,858076932 | 0,21153141 | Wsb2 | WD repeat and SOCS box-containing 2 [Source:MGI Symbol;Acc:MGI:2144041] | 94,24 | 90,1 |
| ENSMUSG00000038388 | 2046,14426 | 1157,5 | 2980,5 | -1,366952748 | 0,07153956 | Mpp6 | membrane protein, palmitoylated 6 (MAGUK p55 subfamily member 6) [Source:MGI Symbol;Acc:MGI:1927340] | 98,98 | 74,5 |
| ENSMUSG00000061904 | 3054,661637 | 3316,2 | 2921,1 | 0,193426172 | 0,78293268 | Slc25a3 | solute carrier family 25 (mitochondrial carrier, phosphate carrier), member 3 [Source:MGI Symbol;Acc:MGI:1353498] | 98,75 | 79,8 |
| ENSMUSG00000050953 | 5551,359317 | 8700,2 | 2885,5 | 1,809170544 | 0,07857246 | Gja1 | gap junction protein, alpha 1 [Source:MGI Symbol;Acc:MGI:95713] | 99,99 | 98,6 |
| ENSMUSG00000062014 | 3652,741862 | 4064,8 | 2857,3 | 0,658371408 | 0,43522345 | Gmfb | glia maturation factor, beta [Source:MGI Symbol;Acc:MGI:1927133] | 94,94 | 87,8 |
| ENSMUSG00000023456 | 3856,066508 | 5079,6 | 2830,2 | 0,874252054 | 0,12090592 | Tpi1 | triosephosphate isomerase 1 [Source:MGI Symbol;Acc:MGI:98797] | 92,97 | 80,2 |
| ENSMUSG00000063229 | 2870,582485 | 2272,0 | 2819,1 | -0,304908061 | 0,68287307 | Ldha | lactate dehydrogenase A [Source:MGI Symbol;Acc:MGI:96759] | 77,85 | 72,8 |
| ENSMUSG00000034566 | 3247,221157 | 3355,3 | 2790,1 | 0,256946264 | 0,70460327 | Atp5h | ATP synthase, H+ transporting, mitochondrial F0 complex, subunit D [Source:MGI Symbol;Acc:MGI:1918929] | 97,93 | 84,3 |
| ENSMUSG00000028788 | 2194,067769 | 1330,5 | 2774,9 | -1,059869157 | 0,10751057 | Ptp4a2 | protein tyrosine phosphatase 4a2 [Source:MGI Symbol;Acc:MGI:1277117] | 98,50 | 88,1 |
| ENSMUSG00000026112 | 1739,195165 | 1108,4 | 2766,8 | -1,257854754 | 0,08879438 | Coa5 | cytochrome C oxidase assembly factor 5 [Source:MGI Symbol;Acc:MGI:1923428] | 98,96 | 98,2 |
| ENSMUSG00000005683 | 2502,534694 | 3002,5 | 2729,6 | 0,248293093 | 0,72885231 | Cs | citrate synthase [Source:MGI Symbol;Acc:MGI:88529] | 99,99 | 98,6 |
| ENSMUSG00000020444 | 2140,378605 | 1073,4 | 2726,3 | -1,294184703 | 0,08530191 | Guk1 | guanylate kinase 1 [Source:MGI Symbol;Acc:MGI:95871] | 92,51 | 85,6 |

| | | | | | | | | | |
|---|---|---|---|---|---|---|---|---|---|
| ENSMUSG00000020088 | 1841,923225 | 1553,6 | 2717,1 | -0,789189331 | 0,40777322 | Sar1a | SAR1 gene homolog A (S. cerevisiae) [Source:MGI Symbol;Acc:MGI:98230] | 100,00 | 99,3 |
| ENSMUSG00000037706 | 6088,715515 | 9604,2 | 2678,5 | 1,836478176 | 0,06902254 | Cd81 | CD81 antigen [Source:MGI Symbol;Acc:MGI:1096398] | 95,12 | 89,8 |
| ENSMUSG00000074093 | 1975,021459 | 2078,9 | 2657,0 | 0,271346889 | 0,85346645 | Svip | small VCP/p97-interacting protein [Source:MGI Symbol;Acc:MGI:1922994] | 92,39 | 85,6 |
| ENSMUSG00000054452 | 2300,461062 | 2127,6 | 2653,5 | -0,308320422 | 0,62555685 | Aes | amino-terminal enhancer of split [Source:MGI Symbol;Acc:MGI:95806] | 99,98 | 99,3 |
| ENSMUSG00000026932 | 2145,596859 | 2251,8 | 2636,8 | -0,204921617 | 0,77560734 | Nacc2 | nucleus accumbens associated 2, BEN and BTB (POZ) domain containing [Source:MGI Symbol;Acc:MGI:1915241] | 98,78 | 85,0 |
| ENSMUSG00000041028 | 4648,22352 | 5937,9 | 2634,5 | 1,188612654 | 0,09914485 | Ghitm | growth hormone inducible transmembrane protein [Source:MGI Symbol;Acc:MGI:1913342] | 100,00 | 85,5 |
| ENSMUSG00000028756 | 4055,164579 | 5438,2 | 2621,2 | 1,421686201 | 0,19980326 | Pink1 | PTEN induced putative kinase 1 [Source:MGI Symbol;Acc:MGI:1916193] | 91,12 | 87,0 |
| ENSMUSG00000016252 | 2301,124822 | 2482,3 | 2608,5 | -0,008796235 | 0,99142344 | Atp5e | ATP synthase, H+ transporting, mitochondrial F1 complex, epsilon subunit [Source:MGI Symbol;Acc:MGI:1855697] | 99,76 | 98,2 |
| ENSMUSG00000026249 | 5666,608047 | 8748,6 | 2578,7 | 1,76088288 | 0,23340606 | Serpine2 | serine (or cysteine) peptidase inhibitor, clade E, member 2 [Source:MGI Symbol;Acc:MGI:101780] | 95,51 | 87,6 |
| ENSMUSG00000020321 | 3808,111951 | 5058,5 | 2560,8 | 0,978417368 | 0,08835563 | Mdh1 | malate dehydrogenase 1, NAD (soluble) [Source:MGI Symbol;Acc:MGI:97051] | 91,38 | 73,2 |
| ENSMUSG00000068823 | 1993,787585 | 1570,8 | 2553,9 | -0,678833076 | 0,38618594 | Csde1 | cold shock domain containing E1, RNA binding [Source:MGI Symbol;Acc:MGI:92356] | 85,78 | 79,3 |
| ENSMUSG00000015090 | 3038,742802 | 2472,2 | 2553,6 | 0,87130925 | 0,54238917 | Ptgds | prostaglandin D2 synthase (brain) [Source:MGI Symbol;Acc:MGI:99261] | 99,75 | 82,4 |
| ENSMUSG00000009630 | 1579,223773 | 927,8 | 2542,7 | -1,363052569 | 0,05912182 | Ppp2cb | protein phosphatase 2 (formerly 2A), catalytic subunit, beta isoform [Source:MGI Symbol;Acc:MGI:1321161] | 99,05 | 93,9 |
| ENSMUSG00000020368 | 1904,239584 | 1668,4 | 2521,5 | -0,608531484 | 0,50758975 | Canx | calnexin [Source:MGI Symbol;Acc:MGI:88261] | 97,72 | 86,8 |
| ENSMUSG00000027423 | 2789,968485 | 2810,5 | 2500,2 | 0,41544202 | 0,60028652 | Snx5 | sorting nexin 5 [Source:MGI Symbol;Acc:MGI:1916428] | 100,00 | 94,4 |
| ENSMUSG00000020163 | 2762,565566 | 3052,3 | 2475,3 | 0,357010851 | 0,57554726 | Uqcr11 | ubiquinol-cytochrome c reductase, complex III subunit XI [Source:MGI Symbol;Acc:MGI:1913844] | 100,00 | 100,0 |
| ENSMUSG00000028692 | 2941,079006 | 3327,0 | 2469,9 | 0,468617475 | 0,49502329 | Akr1a1 | aldo-keto reductase family 1, member A1 (aldehyde reductase) [Source:MGI Symbol;Acc:MGI:1929955] | 92,38 | 91,3 |
| ENSMUSG00000059534 | 2975,266089 | 3446,9 | 2461,0 | 0,539425255 | 0,37166328 | Uqcr10 | ubiquinol-cytochrome c reductase, complex III subunit X [Source:MGI Symbol;Acc:MGI:1913402] | 100,00 | 100,0 |
| ENSMUSG00000031839 | 3575,532825 | 5014,2 | 2454,4 | 1,330335888 | 0,14551712 | Hsbp1 | heat shock factor binding protein 1 [Source:MGI Symbol;Acc:MGI:1915446] | 100,00 | 99,8 |
| ENSMUSG00000014769 | 2114,332258 | 2043,2 | 2427,9 | -0,246646547 | 0,71778004 | Psmb1 | proteasome (prosome, macropain) subunit, beta type 1 [Source:MGI Symbol;Acc:MGI:104884] | 100,00 | 99,0 |
| ENSMUSG00000036751 | 3694,06882 | 4895,1 | 2425,0 | 1,123051713 | 0,09401621 | Cox6b1 | cytochrome c oxidase, subunit VIb polypeptide 1 [Source:MGI Symbol;Acc:MGI:107460] | 91,05 | 81,7 |
| ENSMUSG00000025132 | 2876,359539 | 3223,3 | 2405,9 | 0,419782743 | 0,51395515 | Arhgdia | Rho GDP dissociation inhibitor (GDI) alpha [Source:MGI Symbol;Acc:MGI:2178103] | 92,87 | 91,0 |
| ENSMUSG00000026626 | 3155,126767 | 3623,3 | 2392,4 | 0,817417754 | 0,30317024 | Ppp2r5a | protein phosphatase 2, regulatory subunit B', alpha [Source:MGI Symbol;Acc:MGI:2388479] | 99,96 | 94,0 |
| ENSMUSG00000037266 | 1372,375321 | 579,0 | 2387,2 | -1,963202981 | 0,05889842 | Rsrp1 | arginine/serine rich protein 1 [Source:MGI Symbol;Acc:MGI:106498] | 97,83 | 87,6 |
| ENSMUSG00000032320 | 3422,649944 | 4624,9 | 2360,6 | 1,203512005 | 0,58932587 | Rcn2 | reticulocalbin 2 [Source:MGI Symbol;Acc:MGI:1349765] | 99,52 | 88,9 |
| ENSMUSG00000064339 | 1568,253358 | 745,0 | 2355,1 | -2,026434748 | 0,12293047 | mt-Rnr2 | mitochondrially encoded 16S rRNA [Source:MGI Symbol;Acc:MGI:102492] | 99,77 | 99,1 |
| ENSMUSG00000014294 | 2453,538432 | 2406,1 | 2312,9 | 0,07321698 | 0,91460689 | Ndufa2 | NADH dehydrogenase (ubiquinone) 1 alpha subcomplex, 2 [Source:MGI Symbol;Acc:MGI:1343103] | 99,18 | 93,8 |
| ENSMUSG00000029472 | 2014,596278 | 1251,5 | 2287,7 | -0,814468582 | 0,27750294 | Anapc5 | anaphase-promoting complex subunit 5 [Source:MGI Symbol;Acc:MGI:1929722] | 97,56 | 89,4 |
| ENSMUSG00000022253 | 2359,492517 | 3050,2 | 2283,6 | 0,436442145 | 0,60344043 | Nadk2 | NAD kinase 2, mitochondrial [Source:MGI Symbol;Acc:MGI:1915896] | 92,80 | 75,4 |
| ENSMUSG00000000563 | 3338,598558 | 4435,6 | 2279,6 | 0,954755698 | 0,12193431 | Atp5f1 | ATP synthase, H+ transporting, mitochondrial F0 complex, subunit B1 [Source:MGI Symbol;Acc:MGI:1100495] | 95,49 | 79,0 |
| ENSMUSG00000032324 | 4995,821501 | 7052,1 | 2271,0 | 1,704437652 | 0,13035325 | Tspan3 | tetraspanin 3 [Source:MGI Symbol;Acc:MGI:1928098] | 100,00 | 98,5 |
| ENSMUSG00000020326 | 2231,47152 | 2412,0 | 2267,9 | 0,108857995 | 0,90377405 | Ccng1 | cyclin G1 [Source:MGI Symbol;Acc:MGI:102890] | 99,53 | 97,6 |
| ENSMUSG00000030007 | 1608,106507 | 973,7 | 2254,5 | -1,152557344 | 0,12395758 | Cct7 | chaperonin containing Tcp1, subunit 7 (eta) [Source:MGI Symbol;Acc:MGI:107184] | 99,00 | 75,8 |
| ENSMUSG00000027367 | 2186,326796 | 2858,3 | 2249,1 | 0,386195585 | 0,5559543 | Stard7 | START domain containing 7 [Source:MGI Symbol;Acc:MGI:2139090] | 75,25 | 77,9 |
| ENSMUSG00000006651 | 3464,671267 | 4500,2 | 2248,2 | 1,216415781 | 0,25753759 | Aplp1 | amyloid beta (A4) precursor-like protein 1 [Source:MGI Symbol;Acc:MGI:88046] | 100,00 | 97,1 |
| ENSMUSG00000037058 | 2457,660086 | 2877,3 | 2234,8 | 0,380221104 | 0,56475774 | Paip2 | polyadenylate-binding protein-interacting protein 2 [Source:MGI Symbol;Acc:MGI:1915119] | 87,25 | 79,5 |
| ENSMUSG00000025702 | 1398,387403 | 692,6 | 2229,6 | -1,501524581 | 0,10939557 | March8 | membrane-associated ring finger (C3HC4) 8 [Source:MGI Symbol;Acc:MGI:1919029] | 90,01 | 82,9 |
| ENSMUSG00000027828 | 2262,248371 | 2067,5 | 2214,6 | -0,13691218 | 0,85565633 | Ssr3 | signal sequence receptor, gamma [Source:MGI Symbol;Acc:MGI:1914687] | 97,59 | 73,5 |
| ENSMUSG00000032281 | 4375,120962 | 7056,1 | 2199,7 | 1,780004659 | 0,1391128 | Acsbg1 | acyl-CoA synthetase bubblegum family member 1 [Source:MGI Symbol;Acc:MGI:2385656] | 84,38 | 76,9 |
| ENSMUSG00000032966 | 2747,699618 | 3530,9 | 2187,3 | 0,755538101 | 0,30028156 | Fkbp1a | FK506 binding protein 1a [Source:MGI Symbol;Acc:MGI:95541] | 99,86 | 88,2 |
| ENSMUSG00000022956 | 3298,606636 | 4385,1 | 2167,0 | 1,061265066 | 0,09983077 | Atp5o | ATP synthase, H+ transporting, mitochondrial F1 complex, O subunit [Source:MGI Symbol;Acc:MGI:106341] | 97,87 | 73,2 |
| ENSMUSG00000029064 | 2491,925301 | 2838,0 | 2163,3 | 0,393527553 | 0,5275801 | Gnb1 | guanine nucleotide binding protein (G protein), beta 1 [Source:MGI Symbol;Acc:MGI:95781] | 90,69 | 86,0 |
| ENSMUSG00000063015 | 2169,592287 | 1425,7 | 2161,4 | -0,580393576 | 0,36943976 | Ccni | cyclin I [Source:MGI Symbol;Acc:MGI:1341077] | 83,90 | 73,2 |
| ENSMUSG00000041697 | 3473,788587 | 4385,9 | 2159,6 | 1,212826497 | 0,11517165 | Cox6a1 | cytochrome c oxidase subunit VIa polypeptide 1 [Source:MGI Symbol;Acc:MGI:103099] | 100,00 | 96,4 |
| ENSMUSG00000011179 | 1631,489136 | 993,6 | 2157,0 | -0,991449921 | 0,18322851 | Odc1 | ornithine decarboxylase, structural 1 [Source:MGI Symbol;Acc:MGI:97402] | 91,41 | 81,9 |
| ENSMUSG00000038462 | 2603,340821 | 3073,1 | 2153,9 | 0,514716535 | 0,43297493 | Uqcrfs1 | ubiquinol-cytochrome c reductase, Rieske iron-sulfur polypeptide 1 [Source:MGI Symbol;Acc:MGI:1913944] | 100,00 | 90,5 |
| ENSMUSG00000048388 | 2764,498685 | 3611,0 | 2144,3 | 1,183357388 | 0,44816894 | Fam171b | family with sequence similarity 171, member B [Source:MGI Symbol;Acc:MGI:2444579] | 100,00 | 70,9 |
| ENSMUSG00000060450 | 1781,613166 | 1849,7 | 2142,1 | -0,075592687 | 0,92562652 | Rnf14 | ring finger protein 14 [Source:MGI Symbol;Acc:MGI:1929668] | 99,11 | 91,2 |
| ENSMUSG00000019370 | 2534,419469 | 3298,4 | 2142,0 | 0,869351577 | 0,2518475 | Calm3 | calmodulin 3 [Source:MGI Symbol;Acc:MGI:103249] | 99,71 | 82,4 |
| ENSMUSG00000035674 | 2118,591011 | 2162,8 | 2128,2 | 0,009053454 | 0,9880838 | Ndufa3 | NADH dehydrogenase (ubiquinone) 1 alpha subcomplex, 3 [Source:MGI Symbol;Acc:MGI:1913341] | 97,39 | 74,3 |
| ENSMUSG00000063297 | 2872,327901 | 3485,7 | 2126,7 | 0,882863256 | 0,71297914 | Luzp2 | leucine zipper protein 2 [Source:MGI Symbol;Acc:MGI:1889615] | 99,99 | 96,4 |
| ENSMUSG00000006728 | 1689,476739 | 1409,1 | 2121,0 | -0,586963482 | 0,3784655 | Cdk4 | cyclin-dependent kinase 4 [Source:MGI Symbol;Acc:MGI:88357] | 87,98 | 87,7 |
| ENSMUSG00000003072 | 3142,813654 | 3885,7 | 2119,6 | 0,955110869 | 0,12668244 | Atp5d | ATP synthase, H+ transporting, mitochondrial F1 complex, delta subunit [Source:MGI Symbol;Acc:MGI:1913293] | 89,63 | 72,3 |
| ENSMUSG00000079037 | 5347,738892 | 8124,3 | 2116,7 | 2,090563007 | 0,05835903 | Prnp | prion protein [Source:MGI Symbol;Acc:MGI:97769] | 99,69 | 98,5 |
| ENSMUSG00000024359 | 1443,871843 | 948,6 | 2113,9 | -1,073973347 | 0,19500858 | Hspa9 | heat shock protein 9 [Source:MGI Symbol;Acc:MGI:96245] | 99,57 | 96,5 |
| ENSMUSG00000048022 | 2950,182367 | 3858,0 | 2111,7 | 1,189151365 | 0,61063741 | Tmem229a | transmembrane protein 229A [Source:MGI Symbol;Acc:MGI:2442812] | 99,02 | 91,1 |
| ENSMUSG00000036046 | 1381,928029 | 863,8 | 2106,2 | -1,274215579 | 0,10623381 | 5031439G07Rik | RIKEN cDNA 5031439G07 gene [Source:MGI Symbol;Acc:MGI:2444899] | 99,08 | 83,4 |
| ENSMUSG00000032330 | 3160,049227 | 3740,7 | 2096,7 | 0,849609348 | 0,14354716 | Cox7a2 | cytochrome c oxidase subunit VIIa 2 [Source:MGI Symbol;Acc:MGI:1316715] | 92,48 | 71,9 |
| ENSMUSG00000048578 | 1249,350687 | 597,8 | 2086,8 | -1,656482552 | 0,06491364 | Mlec | malectin [Source:MGI Symbol;Acc:MGI:1924015] | 99,84 | 74,5 |
| ENSMUSG00000035805 | 3961,118095 | 6156,6 | 2075,6 | 1,627176595 | 0,20777576 | Mlc1 | megalencephalic leukoencephalopathy with subcortical cysts 1 homolog (human) [Source:MGI Symbol;Acc:MGI:2157910] | 99,52 | 89,7 |
| ENSMUSG00000005779 | 1615,146149 | 1066,5 | 2073,9 | -0,905558075 | 0,19373563 | Psmb4 | proteasome (prosome, macropain) subunit, beta type 4 [Source:MGI Symbol;Acc:MGI:1098257] | 85,52 | 78,8 |
| ENSMUSG00000029632 | 3461,122917 | 4589,9 | 2065,1 | 1,303171261 | 0,06654685 | Ndufa4 | NADH dehydrogenase (ubiquinone) 1 alpha subcomplex, 4 [Source:MGI Symbol;Acc:MGI:107686] | 100,00 | 86,8 |
| ENSMUSG00000057388 | 1255,365167 | 863,9 | 2055,3 | -1,157287738 | 0,07013872 | Mrpl18 | mitochondrial ribosomal protein L18 [Source:MGI Symbol;Acc:MGI:1914931] | 86,35 | 81,5 |
| ENSMUSG00000051695 | 1613,250529 | 1166,4 | 2050,3 | -0,793896484 | 0,20933344 | Pcbp1 | poly(rC) binding protein 1 [Source:MGI Symbol;Acc:MGI:1345635] | 90,71 | 84,3 |
| ENSMUSG00000018593 | 3329,688726 | 4608,5 | 2036,7 | 1,243945697 | 0,32560908 | Sparc | secreted acidic cysteine rich glycoprotein [Source:MGI Symbol;Acc:MGI:98373] | 97,97 | 91,8 |
| ENSMUSG00000032181 | 3408,425474 | 4924,2 | 2017,5 | 1,402995176 | 0,3958695 | Scg3 | secretogranin III [Source:MGI Symbol;Acc:MGI:103032] | 100,00 | 98,0 |
| ENSMUSG00000021124 | 1693,024728 | 1790,5 | 1992,5 | -0,069702833 | 0,93391114 | Vti1b | vesicle transport through interaction with t-SNAREs 1B [Source:MGI Symbol;Acc:MGI:1855688] | 98,81 | 75,2 |
| ENSMUSG00000031667 | 2010,782507 | 1879,7 | 1991,3 | 0,087910541 | 0,91649772 | Aktip | thymoma viral proto-oncogene 1 interacting protein [Source:MGI Symbol;Acc:MGI:3693832] | 90,94 | 74,2 |
| ENSMUSG00000044080 | 4711,067344 | 7245,7 | 1986,9 | 2,439874651 | 0,05286793 | S100a1 | S100 calcium binding protein A1 [Source:MGI Symbol;Acc:MGI:1338917] | 100,00 | 89,2 |

| | | | | | | | | | |
|---|---|---|---|---|---|---|---|---|---|
| ENSMUSG00000072704 | 2090,024841 | 2296,4 | 1982,3 | 0,320363202 | 0,73740505 | Smim10l1 | small integral membrane protein 10 like 1 [Source:MGI Symbol;Acc:MGI:1914379] | 89,39 | 82,7 |
| ENSMUSG00000039016 | 1740,645304 | 1629,2 | 1981,0 | -0,277847028 | 0,66683019 | Timm8b | translocase of inner mitochondrial membrane 8B [Source:MGI Symbol;Acc:MGI:1353424] | 100,00 | 100,0 |
| ENSMUSG00000015749 | 1442,396191 | 1628,6 | 1969,9 | -0,144851994 | 0,90216868 | Anp32e | acidic (leucine-rich) nuclear phosphoprotein 32 family, member E [Source:MGI Symbol;Acc:MGI:1913721] | 95,05 | 93,2 |
| ENSMUSG00000020719 | 1333,793431 | 1151,0 | 1969,8 | -0,765229809 | 0,3376246 | Ddx5 | DEAD (Asp-Glu-Ala-Asp) box polypeptide 5 [Source:MGI Symbol;Acc:MGI:105037] | 97,62 | 85,8 |
| ENSMUSG00000006304 | 1943,866362 | 2023,1 | 1959,7 | 0,091455493 | 0,89942426 | Arpc2 | actin related protein 2/3 complex, subunit 2 [Source:MGI Symbol;Acc:MGI:1923959] | 81,55 | 72,7 |
| ENSMUSG00000026755 | 1454,854696 | 1191,8 | 1941,7 | -0,704620479 | 0,3500366 | Arpc5l | actin related protein 2/3 complex, subunit 5-like [Source:MGI Symbol;Acc:MGI:1921442] | 99,95 | 97,9 |
| ENSMUSG00000023913 | 5748,025221 | 9406,0 | 1935,8 | 2,26362525 | 0,05433802 | Pla2g7 | phospholipase A2, group VII (platelet-activating factor acetylhydrolase, plasma) [Source:MGI Symbol;Acc:MGI:1351327] | 99,84 | 85,4 |
| ENSMUSG00000014504 | 1301,781412 | 1073,2 | 1911,4 | -0,796125309 | 0,24281405 | Srp19 | signal recognition particle 19 [Source:MGI Symbol;Acc:MGI:1913634] | 99,77 | 99,0 |
| ENSMUSG00000029580 | 3426,475468 | 5087,8 | 1900,6 | 1,47664894 | 0,05216259 | Actb | actin, beta [Source:MGI Symbol;Acc:MGI:87904] | 92,66 | 85,8 |
| ENSMUSG00000021290 | 2704,396082 | 3154,7 | 1899,9 | 0,735629228 | 0,23677553 | 2010107E04Rik | RIKEN cDNA 2010107E04 gene [Source:MGI Symbol;Acc:MGI:1917507] | 100,00 | 100,0 |
| ENSMUSG00000031776 | 2900,986141 | 3573,5 | 1886,1 | 0,973742484 | 0,17298835 | Arl2bp | ADP-ribosylation factor-like 2 binding protein [Source:MGI Symbol;Acc:MGI:1349429] | 100,00 | 100,0 |
| ENSMUSG00000026511 | 1409,667344 | 951,4 | 1881,3 | -0,986266896 | 0,082204 | Srp9 | signal recognition particle 9 [Source:MGI Symbol;Acc:MGI:1350930] | 88,62 | 83,4 |
| ENSMUSG00000021474 | 1872,525901 | 1830,1 | 1873,3 | 0,007130099 | 0,99300766 | Sfxn1 | sideroflexin 1 [Source:MGI Symbol;Acc:MGI:2137677] | 99,95 | 93,2 |
| ENSMUSG00000026833 | 1626,500393 | 1615,1 | 1867,1 | -0,181592576 | 0,79932211 | Olfm1 | olfactomedin 1 [Source:MGI Symbol;Acc:MGI:1860437] | 99,51 | 90,2 |
| ENSMUSG00000019877 | 1980,179618 | 2476,6 | 1863,1 | 1,036448572 | 0,32060764 | Serinc1 | serine incorporator 1 [Source:MGI Symbol;Acc:MGI:1926228] | 99,91 | 93,5 |
| ENSMUSG00000010277 | 1364,178305 | 1216,2 | 1860,3 | -0,173229171 | 0,86621716 | 2610507B11Rik | RIKEN cDNA 2610507B11 gene [Source:MGI Symbol;Acc:MGI:1919753] | 99,87 | 71,3 |
| ENSMUSG00000024533 | 2067,464929 | 2444,4 | 1860,3 | 0,865051861 | 0,64304439 | Spire1 | spire homolog 1 (Drosophila) [Source:MGI Symbol;Acc:MGI:1915416] | 98,88 | 84,1 |
| ENSMUSG00000029128 | 1492,177674 | 1165,0 | 1860,0 | -0,675023398 | 0,23759723 | Rab28 | RAB28, member RAS oncogene family [Source:MGI Symbol;Acc:MGI:1917285] | 100,00 | 97,5 |
| ENSMUSG00000041126 | 1995,194358 | 2361,3 | 1853,2 | 0,453017781 | 0,56184282 | H2afv | H2A histone family, member V [Source:MGI Symbol;Acc:MGI:1924855] | 99,98 | 97,8 |
| ENSMUSG00000050856 | 1968,539729 | 2264,1 | 1845,3 | 0,28774986 | 0,62669006 | Atp5k | ATP synthase, H+ transporting, mitochondrial F1F0 complex, subunit E [Source:MGI Symbol;Acc:MGI:106636] | 96,32 | 91,8 |
| ENSMUSG00000000088 | 2978,76713 | 3986,5 | 1844,6 | 1,118935233 | 0,05592704 | Cox5a | cytochrome c oxidase subunit Va [Source:MGI Symbol;Acc:MGI:88474] | 100,00 | 96,9 |
| ENSMUSG00000031447 | 4123,792757 | 5978,9 | 1841,1 | 1,741258489 | 0,06706328 | Lamp1 | lysosomal-associated membrane protein 1 [Source:MGI Symbol;Acc:MGI:96745] | 99,79 | 92,3 |
| ENSMUSG00000019179 | 2131,661804 | 2590,9 | 1827,1 | 0,515896472 | 0,43979352 | Mdh2 | malate dehydrogenase 2, NAD (mitochondrial) [Source:MGI Symbol;Acc:MGI:97050] | 97,09 | 87,6 |
| ENSMUSG00000016319 | 1966,702936 | 2202,9 | 1799,0 | 0,292637443 | 0,6225431 | Slc25a5 | ıte carrier family 25 (mitochondrial carrier, adenine nucleotide translocator), member 5 [Source:MGI Symbol;Acc:MGI:1353 | 99,27 | 97,2 |
| ENSMUSG00000032562 | 3236,846203 | 4993,4 | 1791,4 | 1,529834277 | 0,14423691 | Gnai2 | guanine nucleotide binding protein (G protein), alpha inhibiting 2 [Source:MGI Symbol;Acc:MGI:95772] | 92,18 | 87,6 |
| ENSMUSG00000041329 | 4328,70784 | 6734,9 | 1770,4 | 2,022641205 | 0,06873617 | Atp1b2 | ATPase, Na+/K+ transporting, beta 2 polypeptide [Source:MGI Symbol;Acc:MGI:88109] | 99,13 | 95,3 |
| ENSMUSG00000027162 | 2042,433167 | 1988,0 | 1760,0 | 0,499046572 | 0,55478791 | Lin7c | lin-7 homolog C (C. elegans) [Source:MGI Symbol;Acc:MGI:1330839] | 99,91 | 77,3 |
| ENSMUSG00000006740 | 1246,005079 | 1262,4 | 1757,4 | -0,379230874 | 0,65807161 | Kif5b | kinesin family member 5B [Source:MGI Symbol;Acc:MGI:1098268] | 95,03 | 72,7 |
| ENSMUSG00000030357 | 1206,425388 | 717,4 | 1752,4 | -1,172240618 | 0,11237002 | Fkbp4 | FK506 binding protein 4 [Source:MGI Symbol;Acc:MGI:95543] | 96,32 | 93,9 |
| ENSMUSG00000018293 | 1825,786204 | 1904,7 | 1749,9 | 0,124748127 | 0,86033805 | Pfn1 | profilin 1 [Source:MGI Symbol;Acc:MGI:97549] | 90,16 | 83,3 |
| ENSMUSG00000026914 | 1264,73608 | 703,6 | 1745,3 | -1,166419883 | 0,17150635 | Psmd14 | proteasome (prosome, macropain) 26S subunit, non-ATPase, 14 [Source:MGI Symbol;Acc:MGI:1913284] | 98,90 | 89,8 |
| ENSMUSG00000004897 | 1914,763093 | 2025,7 | 1737,9 | 0,260464764 | 0,77046173 | Hdgf | hepatoma-derived growth factor [Source:MGI Symbol;Acc:MGI:1194494] | 99,18 | 91,0 |
| ENSMUSG00000004980 | 2258,627151 | 2462,3 | 1733,2 | 0,637353402 | 0,40959178 | Hnrnpa2b1 | heterogeneous nuclear ribonucleoprotein A2/B1 [Source:MGI Symbol;Acc:MGI:104819] | 99,90 | 83,8 |
| ENSMUSG00000004892 | 3790,157218 | 6071,2 | 1731,1 | 2,052155174 | 0,13766074 | Bcan | brevican [Source:MGI Symbol;Acc:MGI:1096385] | 99,12 | 98,2 |
| ENSMUSG00000021832 | 1252,733465 | 673,9 | 1729,4 | -1,330408555 | 0,10427232 | Psmc6 | proteasome (prosome, macropain) 26S subunit, ATPase, 6 [Source:MGI Symbol;Acc:MGI:1914339] | 100,00 | 91,1 |
| ENSMUSG00000035686 | 4172,682153 | 8458,4 | 1724,6 | 2,476856437 | 0,08216074 | Thrsp | thyroid hormone responsive [Source:MGI Symbol;Acc:MGI:109126] | 94,13 | 83,6 |
| ENSMUSG00000028798 | 1268,627281 | 886,4 | 1715,6 | -0,919964511 | 0,22974143 | Eif3i | eukaryotic translation initiation factor 3, subunit I [Source:MGI Symbol;Acc:MGI:1860763] | 99,26 | 92,6 |
| ENSMUSG00000024695 | 1108,650347 | 780,3 | 1714,7 | -1,081683483 | 0,15180159 | Zfp91 | zinc finger protein 91 [Source:MGI Symbol;Acc:MGI:104854] | 86,83 | 76,2 |
| ENSMUSG00000032328 | 1276,480463 | 1353,5 | 1713,5 | -0,186902937 | 0,87804917 | Tmem30a | transmembrane protein 30A [Source:MGI Symbol;Acc:MGI:106402] | 97,70 | 74,4 |
| ENSMUSG00000001018 | 2385,731178 | 3526,1 | 1700,8 | 1,338058553 | 0,16306656 | Snapin | SNAP-associated protein [Source:MGI Symbol;Acc:MGI:1333745] | 100,00 | 92,8 |
| ENSMUSG00000010376 | 1471,971823 | 1475,8 | 1698,9 | -0,206141318 | 0,75290949 | Nedd8 | neural precursor cell expressed, developmentally down-regulated gene 8 [Source:MGI Symbol;Acc:MGI:97301] | 100,00 | 100,0 |
| ENSMUSG00000019054 | 1722,324365 | 2080,4 | 1696,9 | 0,392949081 | 0,61142606 | Fis1 | fission, mitochondrial 1 [Source:MGI Symbol;Acc:MGI:1913687] | 100,00 | 99,0 |
| ENSMUSG00000040242 | 1242,789385 | 747,3 | 1696,5 | -1,132367189 | 0,10876782 | Fgfr1op2 | FGFR1 oncogene partner 2 [Source:MGI Symbol;Acc:MGI:1914779] | 96,39 | 84,4 |
| ENSMUSG00000036438 | 2823,00107 | 3903,9 | 1693,7 | 1,340936106 | 0,07583496 | Calm2 | calmodulin 2 [Source:MGI Symbol;Acc:MGI:103250] | 97,93 | 94,9 |
| ENSMUSG00000048076 | 2117,633497 | 2517,5 | 1667,9 | 0,59335205 | 0,39957537 | Arf1 | ADP-ribosylation factor 1 [Source:MGI Symbol;Acc:MGI:99431] | 97,81 | 86,2 |
| ENSMUSG00000032412 | 1217,626554 | 684,6 | 1666,3 | -1,146588918 | 0,17273027 | Atp1b3 | ATPase, Na+/K+ transporting, beta 3 polypeptide [Source:MGI Symbol;Acc:MGI:107788] | 94,80 | 89,9 |
| ENSMUSG00000024712 | 2262,791117 | 2977,6 | 1665,6 | 0,945472533 | 0,27750294 | Rfk | riboflavin kinase [Source:MGI Symbol;Acc:MGI:1914688] | 100,00 | 91,9 |
| ENSMUSG00000021484 | 1004,294476 | 857,8 | 1662,7 | -0,918234516 | 0,19408237 | Lman2 | lectin, mannose-binding 2 [Source:MGI Symbol;Acc:MGI:1914140] | 99,85 | 96,3 |
| ENSMUSG00000040731 | 1772,340302 | 2046,4 | 1660,7 | 0,302691859 | 0,68252213 | Eif4h | eukaryotic translation initiation factor 4H [Source:MGI Symbol;Acc:MGI:1341822] | 90,98 | 88,3 |
| ENSMUSG00000022698 | 1581,155452 | 1454,4 | 1657,0 | 0,015701916 | 0,98799944 | Naa50 | N(alpha)-acetyltransferase 50, NatE catalytic subunit [Source:MGI Symbol;Acc:MGI:1919367] | 96,88 | 78,3 |
| ENSMUSG00000031176 | 3128,472487 | 4171,3 | 1653,2 | 1,502220158 | 0,11657205 | Dynlt3 | dynein light chain Tctex-type 3 [Source:MGI Symbol;Acc:MGI:1914367] | 99,86 | 98,3 |
| ENSMUSG00000032667 | 2057,644682 | 2580,7 | 1643,6 | 0,834744629 | 0,47603982 | Pon2 | paraoxonase 2 [Source:MGI Symbol;Acc:MGI:106687] | 96,60 | 75,1 |
| ENSMUSG00000024587 | 1430,588629 | 843,6 | 1643,2 | -0,937511319 | 0,29475439 | Nars | asparaginyl-tRNA synthetase [Source:MGI Symbol;Acc:MGI:1917473] | 100,00 | 99,3 |
| ENSMUSG00000025917 | 908,323169 | 751,9 | 1640,0 | -1,033053199 | 0,12263828 | Cops5 | COP9 signalosome subunit 5 [Source:MGI Symbol;Acc:MGI:1349415] | 98,33 | 83,1 |
| ENSMUSG00000021242 | 1583,014517 | 1598,7 | 1633,7 | -0,081706824 | 0,92453806 | Npc2 | Niemann-Pick type C2 [Source:MGI Symbol;Acc:MGI:1915213] | 99,01 | 71,4 |
| ENSMUSG00000055538 | 2176,057197 | 2983,0 | 1633,5 | 1,560213324 | 0,21449811 | Zcchc24 | zinc finger, CCHC domain containing 24 [Source:MGI Symbol;Acc:MGI:1919168] | 99,22 | 73,4 |
| ENSMUSG00000023175 | 3595,363268 | 4535,2 | 1632,8 | 1,254568193 | 0,21780811 | Bsg | basigin [Source:MGI Symbol;Acc:MGI:88208] | 93,66 | 76,4 |
| ENSMUSG00000004096 | 1740,633625 | 1917,0 | 1623,2 | 0,290452682 | 0,70992063 | Cwc15 | CWC15 spliceosome-associated protein [Source:MGI Symbol;Acc:MGI:1913320] | 100,00 | 95,4 |
| ENSMUSG00000000149 | 1806,008806 | 2389,9 | 1620,5 | 1,066686201 | 0,52904521 | Gna12 | guanine nucleotide binding protein, alpha 12 [Source:MGI Symbol;Acc:MGI:95767] | 100,00 | 92,7 |
| ENSMUSG00000030654 | 3625,530935 | 4595,4 | 1619,8 | 1,648070831 | 0,09908452 | Arl6ip1 | ADP-ribosylation factor-like 6 interacting protein 1 [Source:MGI Symbol;Acc:MGI:1858943] | 98,43 | 94,0 |
| ENSMUSG00000009073 | 1087,365329 | 803,1 | 1619,2 | -1,013839931 | 0,27356117 | Nf2 | neurofibromatosis 2 [Source:MGI Symbol;Acc:MGI:97307] | 88,05 | 79,6 |
| ENSMUSG00000032060 | 2436,137212 | 3078,2 | 1615,7 | 1,403939448 | 0,25215372 | Cryab | crystallin, alpha B [Source:MGI Symbol;Acc:MGI:88516] | 95,33 | 84,1 |
| ENSMUSG00000029247 | 1812,909169 | 1701,5 | 1615,1 | 0,109435832 | 0,87474077 | Paics | ıoimidazole carboxylase, phosphoribosylaminoribosylaminoimidazole, succinocarboxamide synthetase [Source:MGI Symbol | 98,50 | 81,7 |
| ENSMUSG00000071528 | 1782,133545 | 2087,8 | 1614,6 | 0,536470607 | 0,48711242 | Usmg5 | upregulated during skeletal muscle growth 5 [Source:MGI Symbol;Acc:MGI:1891435] | 100,00 | 94,6 |
| ENSMUSG00000032381 | 1271,969926 | 1031,1 | 1606,4 | -0,433443278 | 0,60892955 | Fam96a | family with sequence similarity 96, member A [Source:MGI Symbol;Acc:MGI:1915500] | 99,87 | 91,6 |
| ENSMUSG00000081534 | 1997,571815 | 2285,5 | 1606,3 | 0,50556997 | 0,51066279 | Slc48a1 | solute carrier family 48 (heme transporter), member 1 [Source:MGI Symbol;Acc:MGI:1914989] | 99,42 | 96,3 |
| ENSMUSG00000091264 | 945,9818417 | 565,9 | 1602,5 | -1,09230104 | 0,32344156 | Smim13 | small integral membrane protein 13 [Source:MGI Symbol;Acc:MGI:2652854] | 98,66 | 78,0 |
| ENSMUSG00000015733 | 4405,502608 | 6478,9 | 1599,4 | 2,018445735 | 0,12233143 | Capza2 | capping protein (actin filament) muscle Z-line, alpha 2 [Source:MGI Symbol;Acc:MGI:106222] | 99,24 | 87,8 |
| ENSMUSG00000021218 | 1742,406185 | 1741,7 | 1592,7 | 0,183277761 | 0,7960925 | Gdi2 | guanosine diphosphate (GDP) dissociation inhibitor 2 [Source:MGI Symbol;Acc:MGI:99845] | 97,02 | 85,0 |

| | | | | | | | | | |
|---|---|---|---|---|---|---|---|---|---|
| ENSMUSG00000031392 | 1274,459766 | 1008,9 | 1581,3 | -0,624988171 | 0,34452242 | Irak1 | interleukin-1 receptor-associated kinase 1 [Source:MGI Symbol;Acc:MGI:107420] | 99,60 | 82,8 |
| ENSMUSG00000020048 | 1568,818163 | 1312,1 | 1574,8 | -0,292656154 | 0,81194946 | Hsp90b1 | heat shock protein 90, beta (Grp94), member 1 [Source:MGI Symbol;Acc:MGI:98817] | 99,74 | 95,4 |
| ENSMUSG00000033475 | 1374,423638 | 1396,1 | 1556,0 | -0,156072085 | 0,79560321 | Tomm6 | translocase of outer mitochondrial membrane 6 homolog (yeast) [Source:MGI Symbol;Acc:MGI:1913369] | 96,39 | 86,7 |
| ENSMUSG00000025980 | 1185,837064 | 649,4 | 1552,5 | -1,195409426 | 0,11816835 | Hspd1 | heat shock protein 1 (chaperonin) [Source:MGI Symbol;Acc:MGI:96242] | 91,16 | 77,8 |
| ENSMUSG00000020496 | 1879,869905 | 2392,7 | 1543,8 | 1,019265423 | 0,22772882 | Rnf187 | ring finger protein 187 [Source:MGI Symbol;Acc:MGI:1914224] | 100,00 | 98,2 |
| ENSMUSG00000026150 | 2695,83374 | 4010,3 | 1539,2 | 1,499798421 | 0,41086737 | Mff | mitochondrial fission factor [Source:MGI Symbol;Acc:MGI:1922984] | 99,02 | 82,8 |
| ENSMUSG00000006442 | 998,094809 | 612,7 | 1537,7 | -1,297066304 | 0,16475146 | Srm | spermidine synthase [Source:MGI Symbol;Acc:MGI:102690] | 98,60 | 93,9 |
| ENSMUSG00000014226 | 2022,642926 | 1690,6 | 1529,6 | 0,183676027 | 0,81323981 | Cacybp | calcyclin binding protein [Source:MGI Symbol;Acc:MGI:1270839] | 93,76 | 70,4 |
| ENSMUSG00000032026 | 1423,501139 | 1481,0 | 1525,2 | 0,01239314 | 0,98627462 | Rexo2 | REX2, RNA exonuclease 2 homolog (S. cerevisiae) [Source:MGI Symbol;Acc:MGI:1888981] | 100,00 | 100,0 |
| ENSMUSG00000070319 | 1108,237057 | 607,1 | 1520,1 | -1,295948484 | 0,08017021 | Eif3g | eukaryotic translation initiation factor 3, subunit G [Source:MGI Symbol;Acc:MGI:1858258] | 100,00 | 100,0 |
| ENSMUSG00000003131 | 1365,072666 | 1246,3 | 1515,8 | 0,121677543 | 0,89369157 | Pafah1b2 | platelet-activating factor acetylhydrolase, isoform 1b, subunit 2 [Source:MGI Symbol;Acc:MGI:108415] | 99,82 | 80,8 |
| ENSMUSG00000071655 | 1233,395393 | 1134,2 | 1508,5 | -0,37800936 | 0,68285505 | Ubxn1 | UBX domain protein 1 [Source:MGI Symbol;Acc:MGI:1289301] | 100,00 | 98,7 |
| ENSMUSG00000044627 | 1766,205966 | 1987,3 | 1500,0 | 0,450421343 | 0,50129171 | Swi5 | SWI5 recombination repair homolog (yeast) [Source:MGI Symbol;Acc:MGI:1920181] | 81,49 | 81,3 |
| ENSMUSG00000046240 | 3052,900788 | 4657,1 | 1497,0 | 2,130116691 | 0,12465238 | Hepacam | hepatocyte cell adhesion molecule [Source:MGI Symbol;Acc:MGI:1920177] | 100,00 | 83,8 |
| ENSMUSG00000020585 | 3281,791697 | 4396,1 | 1492,5 | 1,990578701 | 0,07607816 | Laptm4a | lysosomal-associated protein transmembrane 4A [Source:MGI Symbol;Acc:MGI:108017] | 90,41 | 72,5 |
| ENSMUSG00000004268 | 1048,138551 | 589,8 | 1490,5 | -1,328039357 | 0,072248 | Emg1 | EMG1 N1-specific pseudouridine methyltransferase [Source:MGI Symbol;Acc:MGI:1315195] | 98,36 | 98,4 |
| ENSMUSG00000062981 | 1919,229982 | 2047,5 | 1482,5 | 0,608276278 | 0,3885505 | Mrpl42 | mitochondrial ribosomal protein L42 [Source:MGI Symbol;Acc:MGI:1333774] | 98,85 | 96,5 |
| ENSMUSG00000035597 | 847,0866536 | 924,6 | 1482,0 | -0,419311634 | 0,6836644 | Prpf39 | PRP39 pre-mRNA processing factor 39 homolog (yeast) [Source:MGI Symbol;Acc:MGI:104602] | 99,78 | 80,9 |
| ENSMUSG00000084957 | 1728,300237 | 2427,4 | 1476,5 | 0,94641921 | 0,54803183 | Bbip1 | BBSome interacting protein 1 [Source:MGI Symbol;Acc:MGI:1913610] | 96,29 | 79,2 |
| ENSMUSG00000052298 | 1428,823419 | 1383,2 | 1473,8 | 0,019792306 | 0,98166611 | Cdc42se2 | CDC42 small effector 2 [Source:MGI Symbol;Acc:MGI:1919979] | 97,40 | 95,9 |
| ENSMUSG00000020271 | 1641,026271 | 1735,2 | 1472,6 | 0,279699541 | 0,71360766 | Fbxw11 | F-box and WD-40 domain protein 11 [Source:MGI Symbol;Acc:MGI:2144023] | 98,50 | 84,2 |
| ENSMUSG00000028419 | 1898,456426 | 2579,3 | 1470,5 | 1,241208874 | 0,51102404 | Chmp5 | charged multivesicular body protein 5 [Source:MGI Symbol;Acc:MGI:1924209] | 96,49 | 90,2 |
| ENSMUSG00000022312 | 1076,147556 | 580,3 | 1463,9 | -1,311306504 | 0,07263922 | Eif3h | eukaryotic translation initiation factor 3, subunit H [Source:MGI Symbol;Acc:MGI:1915385] | 93,99 | 91,6 |
| ENSMUSG00000020664 | 1094,416852 | 1276,0 | 1460,6 | 0,135863867 | 0,91280985 | Dld | dihydrolipoamide dehydrogenase [Source:MGI Symbol;Acc:MGI:107450] | 100,00 | 94,5 |
| ENSMUSG00000008690 | 1125,848156 | 760,2 | 1457,3 | -0,808201399 | 0,35385141 | Ncaph2 | non-SMC condensin II complex, subunit H2 [Source:MGI Symbol;Acc:MGI:1289164] | 98,56 | 74,6 |
| ENSMUSG00000032393 | 1153,12242 | 1093,8 | 1452,5 | -0,421859452 | 0,55065652 | Dpp8 | dipeptidylpeptidase 8 [Source:MGI Symbol;Acc:MGI:1921638] | 98,72 | 88,4 |
| ENSMUSG00000031059 | 2018,240235 | 2674,1 | 1448,8 | 0,956356637 | 0,18061101 | Ndufb11 | NADH dehydrogenase (ubiquinone) 1 beta subcomplex, 11 [Source:MGI Symbol;Acc:MGI:1349919] | 99,96 | 83,1 |
| ENSMUSG00000020029 | 1049,022934 | 871,8 | 1447,4 | -0,716536078 | 0,49214673 | Nudt4 | nudix (nucleoside diphosphate linked moiety X)-type motif 4 [Source:MGI Symbol;Acc:MGI:1918457] | 99,97 | 98,9 |
| ENSMUSG00000033940 | 1927,818765 | 2692,6 | 1446,5 | 0,918262014 | 0,12473549 | Brk1 | BRICK1, SCAR/WAVE actin-nucleating complex subunit [Source:MGI Symbol;Acc:MGI:1915406] | 100,00 | 99,6 |
| ENSMUSG00000048429 | 1326,50631 | 967,7 | 1446,1 | -0,581783501 | 0,49184067 | 1810026J23Rik | RIKEN cDNA 1810026J23 gene [Source:MGI Symbol;Acc:MGI:1917023] | 97,26 | 90,5 |
| ENSMUSG00000028445 | 2589,643393 | 3812,1 | 1441,7 | 2,336977001 | 0,21596264 | Enho | energy homeostasis associated [Source:MGI Symbol;Acc:MGI:1916888] | 99,53 | 74,2 |
| ENSMUSG00000015341 | 1710,999532 | 1914,3 | 1431,2 | 0,447786519 | 0,46801598 | Golga7 | golgi autoantigen, golgin subfamily a, 7 [Source:MGI Symbol;Acc:MGI:1931029] | 86,52 | 77,9 |
| ENSMUSG00000022354 | 2048,789334 | 2264,9 | 1430,8 | 0,705849965 | 0,24114149 | Ndufb9 | NADH dehydrogenase (ubiquinone) 1 beta subcomplex, 9 [Source:MGI Symbol;Acc:MGI:1913468] | 100,00 | 100,0 |
| ENSMUSG00000003380 | 1414,525699 | 1231,7 | 1426,2 | -0,228301409 | 0,76478477 | Rabac1 | Rab acceptor 1 (prenylated) [Source:MGI Symbol;Acc:MGI:1201692] | 95,38 | 79,2 |
| ENSMUSG00000000171 | 2492,600051 | 3649,6 | 1420,6 | 1,368722133 | 0,06764349 | Sdhd | succinate dehydrogenase complex, subunit D, integral membrane protein [Source:MGI Symbol;Acc:MGI:1914175] | 99,82 | 97,0 |
| ENSMUSG00000027566 | 1528,857531 | 1712,9 | 1406,2 | 0,292137689 | 0,66239025 | Psma7 | proteasome (prosome, macropain) subunit, alpha type 7 [Source:MGI Symbol;Acc:MGI:1347070] | 100,00 | 99,3 |
| ENSMUSG00000006941 | 1480,139325 | 1917,5 | 1403,8 | 0,567732733 | 0,50687526 | Eif1b | eukaryotic translation initiation factor 1B [Source:MGI Symbol;Acc:MGI:1916219] | 100,00 | 99,3 |
| ENSMUSG00000028161 | 2119,793105 | 2618,5 | 1402,5 | 0,912085318 | 0,22877975 | Ppp3ca | protein phosphatase 3, catalytic subunit, alpha isoform [Source:MGI Symbol;Acc:MGI:107164] | 99,98 | 78,0 |
| ENSMUSG00000014748 | 874,6657582 | 687,4 | 1399,0 | -1,054970594 | 0,27271278 | Tex261 | testis expressed gene 261 [Source:MGI Symbol;Acc:MGI:1096575] | 100,00 | 81,4 |
| ENSMUSG00000055839 | 1629,016094 | 1866,5 | 1387,1 | 0,496791812 | 0,42893355 | Tceb2 | transcription elongation factor B (SIII), polypeptide 2 [Source:MGI Symbol;Acc:MGI:1914923] | 99,93 | 99,5 |
| ENSMUSG00000024962 | 1329,812736 | 1627,7 | 1385,3 | 0,73209296 | 0,53015022 | Vegfb | vascular endothelial growth factor B [Source:MGI Symbol;Acc:MGI:106199] | 100,00 | 91,0 |
| ENSMUSG00000102918 | 2545,158272 | 3543,2 | 1383,8 | 1,565657721 | 0,35709513 | Pcdhgc3 | protocadherin gamma subfamily C, 3 [Source:MGI Symbol;Acc:MGI:1935201] | 100,00 | 88,7 |
| ENSMUSG00000030884 | 1434,9645 | 1427,4 | 1383,3 | 0,083779204 | 0,90573949 | Uqcrc2 | ubiquinol cytochrome c reductase core protein 2 [Source:MGI Symbol;Acc:MGI:1914253] | 98,63 | 87,8 |
| ENSMUSG00000006476 | 1288,61047 | 1262,6 | 1382,3 | 0,101933923 | 0,91849702 | Nsmf | NMDA receptor synaptonuclear signaling and neuronal migration factor [Source:MGI Symbol;Acc:MGI:1861755] | 98,65 | 97,0 |
| ENSMUSG00000059796 | 913,0223148 | 542,7 | 1378,1 | -1,346072061 | 0,06776873 | Eif4a1 | eukaryotic translation initiation factor 4A1 [Source:MGI Symbol;Acc:MGI:95303] | 88,46 | 85,3 |
| ENSMUSG00000024844 | 972,0578631 | 694,4 | 1375,2 | -0,969491633 | 0,11007827 | Banf1 | barrier to autointegration factor 1 [Source:MGI Symbol;Acc:MGI:1346330] | 100,00 | 95,2 |
| ENSMUSG00000031996 | 1570,393819 | 1625,2 | 1365,0 | 0,194950081 | 0,89383019 | Aplp2 | amyloid beta (A4) precursor-like protein 2 [Source:MGI Symbol;Acc:MGI:88047] | 99,97 | 82,3 |
| ENSMUSG00000034432 | 1186,80949 | 1122,8 | 1358,5 | -0,270714713 | 0,68613611 | Cops8 | COP9 signalosome subunit 8 [Source:MGI Symbol;Acc:MGI:1915363] | 78,22 | 72,4 |
| ENSMUSG00000026576 | 1953,463502 | 2167,9 | 1355,9 | 0,863475321 | 0,47823167 | Atp1b1 | ATPase, Na+/K+ transporting, beta 1 polypeptide [Source:MGI Symbol;Acc:MGI:88108] | 93,70 | 79,6 |
| ENSMUSG00000040385 | 1840,066084 | 2439,8 | 1353,2 | 1,026120436 | 0,2367734 | Ppp1ca | protein phosphatase 1, catalytic subunit, alpha isoform [Source:MGI Symbol;Acc:MGI:103016] | 100,00 | 99,3 |
| ENSMUSG00000024411 | 2883,812073 | 4397,6 | 1352,1 | 1,850033612 | 0,36124015 | Aqp4 | aquaporin 4 [Source:MGI Symbol;Acc:MGI:107387] | 99,90 | 79,0 |
| ENSMUSG00000004207 | 2759,795706 | 3091,9 | 1351,8 | 1,242686562 | 0,27653232 | Psap | prosaposin [Source:MGI Symbol;Acc:MGI:97783] | 95,77 | 70,5 |
| ENSMUSG00000022450 | 1687,752575 | 1993,8 | 1351,7 | 0,657372604 | 0,32949446 | Ndufa6 | NADH dehydrogenase (ubiquinone) 1 alpha subcomplex, 6 (B14) [Source:MGI Symbol;Acc:MGI:1914380] | 100,00 | 93,8 |
| ENSMUSG00000018379 | 1002,957964 | 816,9 | 1351,6 | -0,63808348 | 0,36383156 | Srsf1 | serine/arginine-rich splicing factor 1 [Source:MGI Symbol;Acc:MGI:98283] | 96,01 | 73,4 |
| ENSMUSG00000020358 | 1449,033024 | 1183,4 | 1348,1 | -0,10162664 | 0,89384678 | Hnrnpab | heterogeneous nuclear ribonucleoprotein A/B [Source:MGI Symbol;Acc:MGI:1330294] | 91,64 | 78,3 |
| ENSMUSG00000070394 | 1455,101637 | 1590,8 | 1342,1 | 0,249716484 | 0,66815007 | Tmem256 | transmembrane protein 256 [Source:MGI Symbol;Acc:MGI:1916436] | 100,00 | 97,9 |
| ENSMUSG00000032547 | 1171,77069 | 653,5 | 1339,1 | -0,705128495 | 0,47086445 | Ryk | receptor-like tyrosine kinase [Source:MGI Symbol;Acc:MGI:101766] | 99,61 | 83,4 |
| ENSMUSG00000015759 | 1114,202978 | 930,5 | 1335,2 | -0,534267835 | 0,40617456 | Cnih1 | cornichon family AMPA receptor auxiliary protein 1 [Source:MGI Symbol;Acc:MGI:1277202] | 99,17 | 93,2 |
| ENSMUSG00000033629 | 1245,553213 | 1574,6 | 1322,7 | 0,33274362 | 0,84819593 | Hacd3 | 3-hydroxyacyl-CoA dehydratase 3 [Source:MGI Symbol;Acc:MGI:1889341] | 99,91 | 98,9 |
| ENSMUSG00000008475 | 1272,376183 | 1158,2 | 1318,2 | 0,00527263 | 0,99497875 | Arpc5 | actin related protein 2/3 complex, subunit 5 [Source:MGI Symbol;Acc:MGI:1915021] | 98,83 | 92,7 |
| ENSMUSG00000031065 | 1497,206514 | 1792,0 | 1315,0 | 0,843827128 | 0,34939919 | Cdk16 | cyclin-dependent kinase 16 [Source:MGI Symbol;Acc:MGI:97516] | 89,22 | 75,5 |
| ENSMUSG00000022452 | 2107,390761 | 2803,2 | 1314,0 | 1,160353306 | 0,07429549 | Smdt1 | single-pass membrane protein with aspartate rich tail 1 [Source:MGI Symbol;Acc:MGI:1916279] | 87,16 | 77,0 |
| ENSMUSG00000035864 | 743,3491968 | 550,1 | 1311,7 | -1,25102921 | 0,36340437 | Syt1 | synaptotagmin I [Source:MGI Symbol;Acc:MGI:99667] | 97,03 | 90,3 |
| ENSMUSG00000020390 | 1525,965948 | 1994,5 | 1309,1 | 0,709229521 | 0,32584062 | Ube2b | ubiquitin-conjugating enzyme E2B [Source:MGI Symbol;Acc:MGI:102944] | 99,82 | 89,4 |
| ENSMUSG00000046722 | 1126,204936 | 1489,5 | 1308,0 | 0,243768326 | 0,74357766 | Cdc42se1 | CDC42 small effector 1 [Source:MGI Symbol;Acc:MGI:1889510] | 93,78 | 90,9 |
| ENSMUSG00000022437 | 830,3463435 | 693,4 | 1307,5 | -0,675666671 | 0,38762229 | Samm50 | sorting and assembly machinery component 50 homolog (S. cerevisiae) [Source:MGI Symbol;Acc:MGI:1915903] | 100,00 | 98,3 |
| ENSMUSG00000025651 | 1841,918085 | 2557,4 | 1300,2 | 1,104119734 | 0,17167482 | Uqcrc1 | ubiquinol-cytochrome c reductase core protein 1 [Source:MGI Symbol;Acc:MGI:107876] | 91,70 | 89,6 |
| ENSMUSG00000006205 | 3495,568854 | 5560,3 | 1300,0 | 2,349564309 | 0,05743305 | Htra1 | HtrA serine peptidase 1 [Source:MGI Symbol;Acc:MGI:1929076] | 100,00 | 95,4 |
| ENSMUSG00000030265 | 1182,752795 | 1279,5 | 1297,0 | 0,046531629 | 0,96199586 | Kras | Kirsten rat sarcoma viral oncogene homolog [Source:MGI Symbol;Acc:MGI:96680] | 95,80 | 75,1 |

| | | | | | | | | | |
|---|---|---|---|---|---|---|---|---|---|
| ENSMUSG00000031299 | 1882,82302 | 2454,6 | 1295,4 | 0,962669397 | 0,16617123 | Pdha1 | pyruvate dehydrogenase E1 alpha 1 [Source:MGI Symbol;Acc:MGI:97532] | 99,94 | 96,9 |
| ENSMUSG00000028194 | 2911,636082 | 4449,6 | 1292,0 | 2,000047649 | 0,19215663 | Ddah1 | dimethylarginine dimethylaminohydrolase 1 [Source:MGI Symbol;Acc:MGI:1916469] | 99,26 | 92,8 |
| ENSMUSG00000046671 | 2114,219931 | 2581,2 | 1290,2 | 1,538165235 | 0,10523268 | Mtfr1l | mitochondrial fission regulator 1-like [Source:MGI Symbol;Acc:MGI:1924074] | 98,05 | 81,0 |
| ENSMUSG00000033933 | 1664,961673 | 2512,7 | 1289,9 | 1,305922372 | 0,18832454 | Vhl | von Hippel-Lindau tumor suppressor [Source:MGI Symbol;Acc:MGI:103223] | 98,94 | 92,9 |
| ENSMUSG00000047260 | 1188,065124 | 1170,5 | 1288,8 | -0,005710665 | 0,99490214 | Emc6 | ER membrane protein complex subunit 6 [Source:MGI Symbol;Acc:MGI:1913298] | 97,41 | 94,9 |
| ENSMUSG00000022400 | 1737,572948 | 2181,6 | 1288,7 | 0,796443419 | 0,21063552 | Rbx1 | ring-box 1 [Source:MGI Symbol;Acc:MGI:1891829] | 99,17 | 72,0 |
| ENSMUSG00000021395 | 1008,494728 | 566,6 | 1285,0 | -1,157977457 | 0,08546735 | Spin1 | spindlin 1 [Source:MGI Symbol;Acc:MGI:109242] | 91,45 | 79,8 |
| ENSMUSG00000004264 | 1130,647689 | 925,6 | 1283,0 | -0,401521678 | 0,6430787 | Phb2 | prohibitin 2 [Source:MGI Symbol;Acc:MGI:102520] | 98,14 | 98,1 |
| ENSMUSG00000023150 | 1266,004984 | 1483,7 | 1279,2 | 0,278510164 | 0,85006742 | Ivns1abp | influenza virus NS1A binding protein [Source:MGI Symbol;Acc:MGI:2152389] | 92,04 | 72,5 |
| ENSMUSG00000022427 | 1095,396567 | 1042,9 | 1276,7 | -0,271614909 | 0,67638189 | Tomm22 | translocase of outer mitochondrial membrane 22 homolog (yeast) [Source:MGI Symbol;Acc:MGI:2450248] | 99,94 | 99,6 |
| ENSMUSG00000038150 | 1018,201381 | 644,2 | 1269,4 | -1,000041899 | 0,16178462 | Ormdl3 | ORM1-like 3 (S. cerevisiae) [Source:MGI Symbol;Acc:MGI:1913862] | 95,92 | 93,4 |
| ENSMUSG00000044600 | 1163,333667 | 1091,3 | 1264,3 | -0,115030252 | 0,88950227 | Smim7 | small integral membrane protein 7 [Source:MGI Symbol;Acc:MGI:1914068] | 97,28 | 70,6 |
| ENSMUSG00000034424 | 1598,764042 | 2236,3 | 1261,9 | 1,390495701 | 0,14643579 | Gcsh | glycine cleavage system protein H (aminomethyl carrier) [Source:MGI Symbol;Acc:MGI:1915383] | 96,95 | 84,4 |
| ENSMUSG00000093989 | 1049,164365 | 1035,2 | 1261,2 | -0,222257566 | 0,80731858 | Rnasek | ribonuclease, RNase K [Source:MGI Symbol;Acc:MGI:106369] | 93,59 | 93,4 |
| ENSMUSG00000006058 | 900,6271028 | 524,0 | 1260,7 | -1,235186948 | 0,08216117 | Snf8 | SNF8, ESCRT-II complex subunit, homolog (S. cerevisiae) [Source:MGI Symbol;Acc:MGI:1343161] | 94,89 | 88,4 |
| ENSMUSG00000024012 | 2314,100865 | 3129,5 | 1255,4 | 1,42287532 | 0,08322348 | Mtch1 | mitochondrial carrier homolog 1 (C. elegans) [Source:MGI Symbol;Acc:MGI:1929261] | 96,47 | 91,1 |
| ENSMUSG00000051391 | 1089,51326 | 842,3 | 1252,4 | -0,539028528 | 0,55039931 | Ywhag | e 3-monooxygenase/tryptophan 5-monooxygenase activation protein, gamma polypeptide [Source:MGI Symbol;Acc:MGI:1 | 99,88 | 90,2 |
| ENSMUSG00000042406 | 1085,395691 | 572,2 | 1249,7 | -1,000467594 | 0,20197657 | Atf4 | activating transcription factor 4 [Source:MGI Symbol;Acc:MGI:88096] | 86,56 | 80,2 |
| ENSMUSG00000005687 | 1459,996444 | 1597,4 | 1240,0 | 0,411626599 | 0,58503449 | Bcas2 | breast carcinoma amplified sequence 2 [Source:MGI Symbol;Acc:MGI:1915433] | 96,80 | 92,0 |
| ENSMUSG00000008859 | 1128,195259 | 1036,3 | 1239,2 | -0,233791541 | 0,75664566 | Rala | v-ral simian leukemia viral oncogene homolog A (ras related) [Source:MGI Symbol;Acc:MGI:1927243] | 99,74 | 92,9 |
| ENSMUSG00000039640 | 1264,06807 | 872,8 | 1237,6 | -0,436876226 | 0,54914228 | Mrpl12 | mitochondrial ribosomal protein L12 [Source:MGI Symbol;Acc:MGI:1926273] | 89,64 | 72,9 |
| ENSMUSG00000018326 | 2015,502189 | 2637,9 | 1236,3 | 1,087471065 | 0,20865062 | Ywhab | ne 3-monooxygenase/tryptophan 5-monooxygenase activation protein, beta polypeptide [Source:MGI Symbol;Acc:MGI:189 | 86,44 | 75,5 |
| ENSMUSG00000015994 | 1873,701584 | 2628,5 | 1232,2 | 1,334159775 | 0,14823092 | Fnta | farnesyltransferase, CAAX box, alpha [Source:MGI Symbol;Acc:MGI:104683] | 98,81 | 89,6 |
| ENSMUSG00000022551 | 1750,261779 | 2365,7 | 1230,6 | 0,960429789 | 0,22779559 | Cyc1 | cytochrome c-1 [Source:MGI Symbol;Acc:MGI:1913695] | 100,00 | 99,4 |
| ENSMUSG00000001100 | 1315,02702 | 1593,5 | 1230,1 | 0,4076777 | 0,58118322 | Poldip2 | polymerase (DNA-directed), delta interacting protein 2 [Source:MGI Symbol;Acc:MGI:1915061] | 97,76 | 96,0 |
| ENSMUSG00000025283 | 2706,570568 | 4210,7 | 1228,3 | 2,101466614 | 0,21922889 | Sat1 | spermidine/spermine N1-acetyl transferase 1 [Source:MGI Symbol;Acc:MGI:98233] | 91,91 | 77,3 |
| ENSMUSG00000033953 | 2231,615113 | 3287,1 | 1225,3 | 1,489176994 | 0,05147411 | Ppp3r1 | protein phosphatase 3, regulatory subunit B, alpha isoform (calcineurin B, type I) [Source:MGI Symbol;Acc:MGI:107172] | 99,21 | 84,5 |
| ENSMUSG00000054428 | 1175,57815 | 1263,0 | 1223,5 | 0,036413594 | 0,96308942 | Atpif1 | ATPase inhibitory factor 1 [Source:MGI Symbol;Acc:MGI:1196457] | 96,89 | 82,9 |
| ENSMUSG00000044408 | 1475,398601 | 1898,0 | 1220,6 | 0,913454996 | 0,33854133 | Sptssa | serine palmitoyltransferase, small subunit A [Source:MGI Symbol;Acc:MGI:1913399] | 100,00 | 97,4 |
| ENSMUSG00000029063 | 2597,283398 | 3882,9 | 1217,2 | 1,87106604 | 0,11209957 | Nadk | NAD kinase [Source:MGI Symbol;Acc:MGI:2183149] | 88,32 | 77,4 |
| ENSMUSG00000024853 | 823,7039827 | 840,3 | 1213,7 | -0,18399406 | 0,87317255 | Sf3b2 | splicing factor 3b, subunit 2 [Source:MGI Symbol;Acc:MGI:2441856] | 99,93 | 95,9 |
| ENSMUSG00000018286 | 1393,391949 | 1507,7 | 1211,9 | 0,31484217 | 0,62940843 | Psmb6 | proteasome (prosome, macropain) subunit, beta type 6 [Source:MGI Symbol;Acc:MGI:104880] | 100,00 | 100,0 |
| ENSMUSG00000001472 | 1712,923861 | 2289,5 | 1210,3 | 0,998183913 | 0,1798819 | Tcf25 | transcription factor 25 (basic helix-loop-helix) [Source:MGI Symbol;Acc:MGI:1914105] | 97,72 | 84,8 |
| ENSMUSG00000036570 | 2376,05225 | 3511,8 | 1209,0 | 1,927097108 | 0,07517589 | Fxyd1 | FXYD domain-containing ion transport regulator 1 [Source:MGI Symbol;Acc:MGI:1889273] | 93,83 | 70,2 |
| ENSMUSG00000027637 | 1197,201176 | 1218,0 | 1202,8 | 0,163674834 | 0,82808071 | 1110008F13Rik | RIKEN cDNA 1110008F13 gene [Source:MGI Symbol;Acc:MGI:1914638] | 100,00 | 96,5 |
| ENSMUSG00000020869 | 759,2631925 | 625,9 | 1198,2 | -0,791344816 | 0,46122737 | Lrrc59 | leucine rich repeat containing 59 [Source:MGI Symbol;Acc:MGI:2138133] | 100,00 | 95,8 |
| ENSMUSG00000078652 | 1180,746418 | 1074,8 | 1197,5 | -0,018151481 | 0,99006039 | Psme3 | proteaseome (prosome, macropain) activator subunit 3 (PA28 gamma, Ki) [Source:MGI Symbol;Acc:MGI:1096366] | 96,59 | 91,5 |
| ENSMUSG00000020650 | 809,5938595 | 579,3 | 1196,5 | -0,919322966 | 0,26626167 | Bcap29 | B cell receptor associated protein 29 [Source:MGI Symbol;Acc:MGI:101917] | 95,75 | 86,8 |
| ENSMUSG00000027002 | 2467,458511 | 3301,3 | 1193,6 | 1,665056464 | 0,16206816 | Nckap1 | NCK-associated protein 1 [Source:MGI Symbol;Acc:MGI:1355333] | 98,68 | 71,6 |
| ENSMUSG00000028618 | 2895,301692 | 4502,1 | 1189,7 | 2,061349116 | 0,19744724 | Tmem59 | transmembrane protein 59 [Source:MGI Symbol;Acc:MGI:1929278] | 92,75 | 83,6 |
| ENSMUSG00000038014 | 778,1644882 | 626,2 | 1188,6 | -0,961393459 | 0,47051952 | Fam120a | family with sequence similarity 120, member A [Source:MGI Symbol;Acc:MGI:2446163] | 97,49 | 76,8 |
| ENSMUSG00000022679 | 1464,89476 | 2246,5 | 1182,2 | 1,516171424 | 0,4522173 | Mpv17l | Mpv17 transgene, kidney disease mutant-like [Source:MGI Symbol;Acc:MGI:2135951] | 93,29 | 73,6 |
| ENSMUSG00000021156 | 999,9492112 | 1275,3 | 1175,6 | 0,265562965 | 0,75603383 | Zmynd11 | zinc finger, MYND domain containing 11 [Source:MGI Symbol;Acc:MGI:1913755] | 93,59 | 77,9 |
| ENSMUSG00000044502 | 1119,18583 | 1115,7 | 1174,1 | 0,033070732 | 0,96627774 | Bod1 | biorientation of chromosomes in cell division 1 [Source:MGI Symbol;Acc:MGI:1916806] | 97,47 | 80,8 |
| ENSMUSG00000054766 | 864,6927403 | 613,8 | 1173,6 | -0,923645136 | 0,20306532 | Set | SET nuclear oncogene [Source:MGI Symbol;Acc:MGI:1860267] | 85,16 | 72,8 |
| ENSMUSG00000005732 | 1560,79374 | 1933,8 | 1170,7 | 0,747876953 | 0,23271014 | Ranbp1 | RAN binding protein 1 [Source:MGI Symbol;Acc:MGI:96269] | 94,76 | 90,6 |
| ENSMUSG00000015087 | 754,3061721 | 564,2 | 1169,1 | -0,847175627 | 0,28342373 | Rabl6 | RAB, member RAS oncogene family-like 6 [Source:MGI Symbol;Acc:MGI:2442633] | 99,93 | 91,6 |
| ENSMUSG00000046329 | 1333,362763 | 1377,2 | 1150,6 | 0,403622472 | 0,57289748 | Slc25a23 | solute carrier family 25 (mitochondrial carrier; phosphate carrier), member 23 [Source:MGI Symbol;Acc:MGI:1914222] | 99,75 | 91,5 |
| ENSMUSG00000020803 | 991,6406303 | 880,8 | 1147,2 | -0,362763557 | 0,57187693 | Txndc17 | thioredoxin domain containing 17 [Source:MGI Symbol;Acc:MGI:1289248] | 99,86 | 84,3 |
| ENSMUSG00000022635 | 738,8174246 | 506,8 | 1145,7 | -1,112559843 | 0,15636717 | Zcrb1 | zinc finger CCHC-type and RNA binding motif 1 [Source:MGI Symbol;Acc:MGI:1914447] | 98,76 | 97,7 |
| ENSMUSG00000023307 | 838,9238734 | 857,0 | 1144,4 | -0,20132352 | 0,851916 | March5 | membrane-associated ring finger (C3HC4) 5 [Source:MGI Symbol;Acc:MGI:1915207] | 92,75 | 79,0 |
| ENSMUSG00000066456 | 1605,643094 | 2229,7 | 1141,2 | 1,219825275 | 0,4544235 | Hmgn3 | high mobility group nucleosomal binding domain 3 [Source:MGI Symbol;Acc:MGI:2138069] | 100,00 | 91,2 |
| ENSMUSG00000034290 | 664,158074 | 509,3 | 1138,0 | -1,022967577 | 0,28901311 | Nek9 | NIMA (never in mitosis gene a)-related expressed kinase 9 [Source:MGI Symbol;Acc:MGI:2387995] | 99,33 | 71,1 |
| ENSMUSG00000037242 | 1242,352137 | 769,8 | 1132,5 | -0,240856911 | 0,79899669 | Clic4 | chloride intracellular channel 4 (mitochondrial) [Source:MGI Symbol;Acc:MGI:1352754] | 99,00 | 75,3 |
| ENSMUSG00000020946 | 1472,978769 | 1651,2 | 1132,0 | 0,602647303 | 0,3630348 | Gosr2 | golgi SNAP receptor complex member 2 [Source:MGI Symbol;Acc:MGI:1927204] | 92,89 | 82,5 |
| ENSMUSG00000047459 | 1821,080274 | 2677,0 | 1129,0 | 1,604067914 | 0,13968168 | Dynlrb1 | dynein light chain roadblock-type 1 [Source:MGI Symbol;Acc:MGI:1914318] | 99,50 | 75,7 |
| ENSMUSG00000010054 | 925,3146695 | 938,1 | 1123,9 | -0,254824312 | 0,72531 | Tusc2 | tumor suppressor candidate 2 [Source:MGI Symbol;Acc:MGI:1931086] | 99,98 | 99,2 |
| ENSMUSG00000049760 | 1219,62755 | 1384,2 | 1122,5 | 0,37798751 | 0,60821857 | 2410015M20Rik | RIKEN cDNA 2410015M20 gene [Source:MGI Symbol;Acc:MGI:2442174] | 100,00 | 88,7 |
| ENSMUSG00000022110 | 1172,394547 | 1472,0 | 1122,4 | 0,496833784 | 0,58115321 | Sucla2 | succinate-Coenzyme A ligase, ADP-forming, beta subunit [Source:MGI Symbol;Acc:MGI:1306775] | 87,49 | 71,2 |
| ENSMUSG00000001105 | 739,9946584 | 612,7 | 1121,7 | -0,663635736 | 0,51274338 | Ift20 | intraflagellar transport 20 [Source:MGI Symbol;Acc:MGI:1915585] | 99,05 | 92,0 |
| ENSMUSG00000003955 | 1407,002493 | 1830,4 | 1121,2 | 0,999401084 | 0,25742252 | Fam162a | family with sequence similarity 162, member A [Source:MGI Symbol;Acc:MGI:1917436] | 100,00 | 99,2 |
| ENSMUSG00000027937 | 947,661833 | 843,3 | 1116,5 | -0,161601303 | 0,86628369 | Jtb | jumping translocation breakpoint [Source:MGI Symbol;Acc:MGI:1346082] | 98,88 | 90,4 |
| ENSMUSG00000033938 | 1679,964855 | 2314,6 | 1113,2 | 1,09659484 | 0,08726414 | Ndufb7 | NADH dehydrogenase (ubiquinone) 1 beta subcomplex, 7 [Source:MGI Symbol;Acc:MGI:1914166] | 90,96 | 91,0 |
| ENSMUSG00000008393 | 1899,664427 | 2646,0 | 1107,9 | 1,456022843 | 0,14987693 | Carhsp1 | calcium regulated heat stable protein 1 [Source:MGI Symbol;Acc:MGI:1196368] | 96,86 | 90,7 |
| ENSMUSG00000060938 | 865,3337908 | 589,0 | 1107,0 | -0,909929871 | 0,12974867 | Rpl26 | ribosomal protein L26 [Source:MGI Symbol;Acc:MGI:106022] | 87,82 | 91,8 |
| ENSMUSG00000016257 | 1049,860601 | 1147,7 | 1103,2 | 0,0959117 | 0,90052552 | Slmo2 | slowmo homolog 2 (Drosophila) [Source:MGI Symbol;Acc:MGI:1913640] | 96,22 | 85,0 |
| ENSMUSG00000067847 | 1147,992957 | 1340,5 | 1100,3 | 0,304661618 | 0,64696107 | Romo1 | reactive oxygen species modulator 1 [Source:MGI Symbol;Acc:MGI:1914317] | 100,00 | 91,4 |
| ENSMUSG00000023075 | 834,5625767 | 754,5 | 1099,9 | -0,519036764 | 0,42056432 | Akirin1 | akirin 1 [Source:MGI Symbol;Acc:MGI:1915300] | 85,26 | 75,1 |
| ENSMUSG00000002477 | 751,3856486 | 565,3 | 1093,9 | -0,87428961 | 0,16001488 | Snrpd1 | small nuclear ribonucleoprotein D1 [Source:MGI Symbol;Acc:MGI:98344] | 100,00 | 79,2 |

| | | | | | | | | | |
|---|---|---|---|---|---|---|---|---|---|
| ENSMUSG00000039515 | 950,7134074 | 1043,3 | 1092,1 | 0,169005689 | 0,84783654 | Ppp2r4 | protein phosphatase 2A activator, regulatory subunit B [Source:MGI Symbol;Acc:MGI:1346006] | 92,25 | 81,4 |
| ENSMUSG00000033065 | 1842,894308 | 2805,8 | 1091,3 | 1,432508004 | 0,16628347 | Pfkm | phosphofructokinase, muscle [Source:MGI Symbol;Acc:MGI:97548] | 96,92 | 94,1 |
| ENSMUSG00000021111 | 1211,96642 | 1481,6 | 1088,5 | 0,524091363 | 0,45415997 | Papola | poly (A) polymerase alpha [Source:MGI Symbol;Acc:MGI:109301] | 97,56 | 76,1 |
| ENSMUSG00000025439 | 850,8362996 | 849,9 | 1086,4 | -0,296927099 | 0,73569703 | Clns1a | chloride channel, nucleotide-sensitive, 1A [Source:MGI Symbol;Acc:MGI:109638] | 98,60 | 96,5 |
| ENSMUSG00000030788 | 1707,592527 | 2223,1 | 1086,2 | 1,164965735 | 0,48533148 | Rnf141 | ring finger protein 141 [Source:MGI Symbol;Acc:MGI:1914400] | 91,46 | 76,1 |
| ENSMUSG00000025728 | 917,2328699 | 840,4 | 1086,0 | -0,270111602 | 0,780948 | Pigq | phosphatidylinositol glycan anchor biosynthesis, class Q [Source:MGI Symbol;Acc:MGI:1333114] | 75,80 | 75,7 |
| ENSMUSG00000020402 | 1311,668502 | 1367,3 | 1085,6 | 0,615757626 | 0,55480071 | Vdac1 | voltage-dependent anion channel 1 [Source:MGI Symbol;Acc:MGI:106919] | 98,02 | 85,6 |
| ENSMUSG00000034868 | 1430,416141 | 1296,8 | 1085,4 | 0,325348569 | 0,67533253 | Myl12b | myosin, light chain 12B, regulatory [Source:MGI Symbol;Acc:MGI:107494] | 93,06 | 78,4 |
| ENSMUSG00000026664 | 3237,338153 | 4945,9 | 1084,9 | 2,190549845 | 0,1262489 | Phyh | phytanoyl-CoA hydroxylase [Source:MGI Symbol;Acc:MGI:891978] | 100,00 | 99,5 |
| ENSMUSG00000024953 | 1897,636662 | 2623,2 | 1083,2 | 1,395612725 | 0,09815305 | Prdx5 | peroxiredoxin 5 [Source:MGI Symbol;Acc:MGI:1859821] | 92,86 | 78,8 |
| ENSMUSG00000022295 | 1440,964248 | 1367,6 | 1081,6 | 0,413495817 | 0,59662969 | Atp6v1c1 | ATPase, H+ transporting, lysosomal V1 subunit C1 [Source:MGI Symbol;Acc:MGI:1913585] | 100,00 | 96,3 |
| ENSMUSG00000053907 | 1687,541041 | 1862,5 | 1081,5 | 1,130999991 | 0,30176517 | Mat2a | methionine adenosyltransferase II, alpha [Source:MGI Symbol;Acc:MGI:2443731] | 97,55 | 88,6 |
| ENSMUSG00000007891 | 2638,524874 | 3328,6 | 1081,1 | 1,792748007 | 0,23272615 | Ctsd | cathepsin D [Source:MGI Symbol;Acc:MGI:88562] | 97,86 | 84,6 |
| ENSMUSG00000006315 | 780,1781591 | 645,6 | 1077,9 | -0,786369946 | 0,17194965 | Tmem147 | transmembrane protein 147 [Source:MGI Symbol;Acc:MGI:1915011] | 95,66 | 92,9 |
| ENSMUSG00000019362 | 1191,900028 | 1486,0 | 1071,6 | 0,644567491 | 0,42498674 | D8Ertd738e | DNA segment, Chr 8, ERATO Doi 738, expressed [Source:MGI Symbol;Acc:MGI:1289231] | 99,04 | 93,3 |
| ENSMUSG00000060992 | 1807,855864 | 2168,6 | 1062,8 | 1,050663691 | 0,06028468 | Copz1 | coatomer protein complex, subunit zeta 1 [Source:MGI Symbol;Acc:MGI:1929063] | 100,00 | 99,6 |
| ENSMUSG00000033272 | 1275,134566 | 1522,8 | 1061,5 | 0,852139381 | 0,31938974 | Slc35a4 | solute carrier family 35, member A4 [Source:MGI Symbol;Acc:MGI:1915093] | 94,75 | 70,8 |
| ENSMUSG00000037149 | 911,6491431 | 535,6 | 1061,2 | -0,360262652 | 0,75330992 | Ddx1 | DEAD (Asp-Glu-Ala-Asp) box polypeptide 1 [Source:MGI Symbol;Acc:MGI:2144727] | 99,48 | 74,7 |
| ENSMUSG00000036352 | 928,9862155 | 610,3 | 1053,1 | -0,491560194 | 0,60192242 | Ubac1 | ubiquitin associated domain containing 1 [Source:MGI Symbol;Acc:MGI:1920995] | 100,00 | 94,0 |
| ENSMUSG00000028861 | 903,2977638 | 804,6 | 1053,0 | -0,389517458 | 0,54605751 | Mrps15 | mitochondrial ribosomal protein S15 [Source:MGI Symbol;Acc:MGI:1913657] | 100,00 | 93,2 |
| ENSMUSG00000024208 | 1039,90724 | 982,4 | 1051,3 | -0,060603993 | 0,93811024 | Uqcc2 | ubiquinol-cytochrome c reductase complex assembly factor 2 [Source:MGI Symbol;Acc:MGI:1914517] | 81,37 | 80,5 |
| ENSMUSG00000019471 | 802,8788463 | 547,0 | 1049,9 | -0,791404512 | 0,31742344 | Cdc37 | cell division cycle 37 [Source:MGI Symbol;Acc:MGI:109531] | 88,78 | 80,2 |
| ENSMUSG00000021102 | 850,0317068 | 727,0 | 1048,7 | -0,507542636 | 0,49213122 | Glrx5 | glutaredoxin 5 [Source:MGI Symbol;Acc:MGI:1920296] | 99,68 | 97,1 |
| ENSMUSG00000034120 | 1088,99733 | 1436,5 | 1046,8 | 0,495175394 | 0,73348647 | Srsf2 | serine/arginine-rich splicing factor 2 [Source:MGI Symbol;Acc:MGI:98284] | 100,00 | 96,0 |
| ENSMUSG00000053317 | 688,3716157 | 567,7 | 1045,7 | -0,854777626 | 0,11741973 | Sec61b | Sec61 beta subunit [Source:MGI Symbol;Acc:MGI:1913462] | 75,09 | 71,3 |
| ENSMUSG00000039615 | 1394,097235 | 1703,0 | 1040,5 | 0,919119919 | 0,39477825 | Stub1 | STIP1 homology and U-Box containing protein 1 [Source:MGI Symbol;Acc:MGI:1891731] | 99,34 | 79,8 |
| ENSMUSG00000032002 | 841,8195743 | 801,9 | 1039,6 | -0,197361581 | 0,82735169 | Dcun1d5 | DCN1, defective in cullin neddylation 1, domain containing 5 (S. cerevisiae) [Source:MGI Symbol;Acc:MGI:1924113] | 100,00 | 98,7 |
| ENSMUSG00000024146 | 1365,093025 | 1744,1 | 1036,8 | 0,955900307 | 0,27191916 | Cript | cysteine-rich PDZ-binding protein [Source:MGI Symbol;Acc:MGI:1929655] | 99,75 | 97,3 |
| ENSMUSG00000024665 | 1825,559646 | 2454,7 | 1032,5 | 1,453770922 | 0,40222168 | Fads2 | fatty acid desaturase 2 [Source:MGI Symbol;Acc:MGI:1930079] | 100,00 | 100,0 |
| ENSMUSG00000051234 | 1272,519219 | 1355,6 | 1027,5 | 0,506454052 | 0,50793562 | Rnf7 | ring finger protein 7 [Source:MGI Symbol;Acc:MGI:1337096] | 98,46 | 93,3 |
| ENSMUSG00000003814 | 1115,456501 | 1026,1 | 1027,0 | -0,110189446 | 0,91428066 | Calr | calreticulin [Source:MGI Symbol;Acc:MGI:88252] | 87,70 | 84,2 |
| ENSMUSG00000013593 | 1521,232303 | 1921,5 | 1026,1 | 0,928529242 | 0,3565762 | Ndufs2 | NADH dehydrogenase (ubiquinone) Fe-S protein 2 [Source:MGI Symbol;Acc:MGI:2385112] | 93,63 | 89,1 |
| ENSMUSG00000063888 | 1004,065426 | 1079,5 | 1022,3 | 0,07987727 | 0,91379655 | Rpl7l1 | ribosomal protein L7-like 1 [Source:MGI Symbol;Acc:MGI:1913479] | 99,48 | 87,3 |
| ENSMUSG00000027067 | 795,8724819 | 548,2 | 1017,1 | -0,705155091 | 0,43820954 | Ssrp1 | structure specific recognition protein 1 [Source:MGI Symbol;Acc:MGI:107912] | 98,19 | 92,8 |
| ENSMUSG00000006024 | 1252,499486 | 1663,1 | 1016,9 | 0,7243755 | 0,26829317 | Napa | N-ethylmaleimide sensitive fusion protein attachment protein alpha [Source:MGI Symbol;Acc:MGI:104563] | 97,17 | 82,2 |
| ENSMUSG00000041881 | 1803,680088 | 2499,3 | 1014,0 | 1,415807798 | 0,05227268 | Ndufa7 | NADH dehydrogenase (ubiquinone) 1 alpha subcomplex, 7 (B14.5a) [Source:MGI Symbol;Acc:MGI:1913666] | 99,80 | 96,9 |
| ENSMUSG00000032383 | 942,4766571 | 705,2 | 1012,5 | -0,566001495 | 0,4167711 | Ppib | peptidylprolyl isomerase B [Source:MGI Symbol;Acc:MGI:97750] | 97,15 | 92,4 |
| ENSMUSG00000020671 | 3102,490728 | 5077,5 | 1010,7 | 2,635507437 | 0,09945031 | Rab10 | RAB10, member RAS oncogene family [Source:MGI Symbol;Acc:MGI:105066] | 96,17 | 85,0 |
| ENSMUSG00000038965 | 1491,290921 | 2244,9 | 1010,6 | 1,286834666 | 0,37021052 | Ube2l3 | ubiquitin-conjugating enzyme E2L 3 [Source:MGI Symbol;Acc:MGI:109240] | 83,31 | 72,1 |
| ENSMUSG00000021486 | 1095,546258 | 806,0 | 1003,3 | -0,307267918 | 0,67924109 | Prelid1 | PRELI domain containing 1 [Source:MGI Symbol;Acc:MGI:1913744] | 92,49 | 86,1 |
| ENSMUSG00000015149 | 1924,077495 | 2945,4 | 1002,6 | 1,981758532 | 0,25956871 | Sirt2 | sirtuin 2 [Source:MGI Symbol;Acc:MGI:1927664] | 83,33 | 73,5 |
| ENSMUSG00000049612 | 1052,749335 | 1472,1 | 1000,5 | 0,8091026 | 0,59615443 | Omg | oligodendrocyte myelin glycoprotein [Source:MGI Symbol;Acc:MGI:106586] | 99,72 | 79,6 |
| ENSMUSG00000009549 | 1635,365147 | 2226,4 | 999,9 | 1,278507179 | 0,11671384 | Srp14 | signal recognition particle 14 [Source:MGI Symbol;Acc:MGI:107169] | 76,92 | 70,9 |
| ENSMUSG00000027620 | 1161,502559 | 1291,6 | 998,5 | 0,42141764 | 0,57187693 | Rbm39 | RNA binding motif protein 39 [Source:MGI Symbol;Acc:MGI:2157953] | 93,64 | 70,5 |
| ENSMUSG00000021134 | 1209,839842 | 1069,5 | 998,0 | 0,284815573 | 0,74993736 | Srsf5 | serine/arginine-rich splicing factor 5 [Source:MGI Symbol;Acc:MGI:98287] | 93,24 | 90,2 |
| ENSMUSG00000029776 | 2363,489897 | 3720,8 | 997,6 | 2,138005426 | 0,19435257 | Hibadh | 3-hydroxyisobutyrate dehydrogenase [Source:MGI Symbol;Acc:MGI:1889802] | 99,45 | 92,0 |
| ENSMUSG00000067786 | 1806,932746 | 3078,5 | 995,1 | 2,140586015 | 0,07844276 | Nnat | neuronatin [Source:MGI Symbol;Acc:MGI:104716] | 93,02 | 84,5 |
| ENSMUSG00000037710 | 1301,355511 | 1590,2 | 993,5 | 0,758731562 | 0,2826457 | Cisd1 | CDGSH iron sulfur domain 1 [Source:MGI Symbol;Acc:MGI:1261855] | 97,02 | 80,9 |
| ENSMUSG00000025613 | 1222,189082 | 832,7 | 988,9 | 0,080241881 | 0,93579852 | Cct8 | chaperonin containing Tcp1, subunit 8 (theta) [Source:MGI Symbol;Acc:MGI:107183] | 95,62 | 74,1 |
| ENSMUSG00000015671 | 1308,484297 | 1733,4 | 988,7 | 0,968014714 | 0,18109705 | Psma2 | proteasome (prosome, macropain) subunit, alpha type 2 [Source:MGI Symbol;Acc:MGI:104885] | 100,00 | 96,4 |
| ENSMUSG00000017843 | 944,1930075 | 651,8 | 985,5 | -0,430152864 | 0,66682845 | Ppp2r5c | protein phosphatase 2, regulatory subunit B', gamma [Source:MGI Symbol;Acc:MGI:1349475] | 99,87 | 73,5 |
| ENSMUSG00000022199 | 1498,801543 | 1640,2 | 983,5 | 0,692285431 | 0,70254489 | Slc22a17 | solute carrier family 22 (organic cation transporter), member 17 [Source:MGI Symbol;Acc:MGI:1926225] | 99,71 | 96,5 |
| ENSMUSG00000024997 | 1372,228406 | 1963,7 | 975,9 | 1,067123007 | 0,27307563 | Prdx3 | peroxiredoxin 3 [Source:MGI Symbol;Acc:MGI:88034] | 99,05 | 92,3 |
| ENSMUSG00000021794 | 1804,286846 | 2340,0 | 967,1 | 1,273139451 | 0,08020609 | Glud1 | glutamate dehydrogenase 1 [Source:MGI Symbol;Acc:MGI:95753] | 99,42 | 75,0 |
| ENSMUSG00000068882 | 867,6216279 | 1166,4 | 963,2 | 0,289697227 | 0,82319348 | Ssb | Sjogren syndrome antigen B [Source:MGI Symbol;Acc:MGI:98423] | 98,79 | 81,9 |
| ENSMUSG00000047557 | 2739,919044 | 4546,5 | 958,1 | 2,57287966 | 0,06828041 | Lxn | latexin [Source:MGI Symbol;Acc:MGI:107633] | 100,00 | 96,4 |
| ENSMUSG00000001576 | 685,1794944 | 532,2 | 953,0 | -0,836379967 | 0,40229687 | Ergic1 | endoplasmic reticulum-golgi intermediate compartment (ERGIC) 1 [Source:MGI Symbol;Acc:MGI:1914708] | 95,16 | 76,9 |
| ENSMUSG00000032349 | 1361,38788 | 1837,7 | 951,8 | 1,066052858 | 0,34835492 | Elovl5 | ELOVL family member 5, elongation of long chain fatty acids (yeast) [Source:MGI Symbol;Acc:MGI:1916051] | 94,03 | 82,8 |
| ENSMUSG00000041084 | 777,7094345 | 659,7 | 951,8 | -0,505627188 | 0,43226501 | Ostc | oligosaccharyltransferase complex subunit [Source:MGI Symbol;Acc:MGI:1913607] | 99,56 | 96,3 |
| ENSMUSG00000015291 | 1333,201134 | 1956,7 | 949,7 | 1,114152441 | 0,21579779 | Gdi1 | guanosine diphosphate (GDP) dissociation inhibitor 1 [Source:MGI Symbol;Acc:MGI:99846] | 91,21 | 85,0 |
| ENSMUSG00000036299 | 1144,726605 | 1709,7 | 949,6 | 1,131798258 | 0,39212989 | BC031181 | cDNA sequence BC031181 [Source:MGI Symbol;Acc:MGI:3039614] | 98,09 | 97,6 |
| ENSMUSG00000023020 | 1097,315147 | 1370,7 | 949,1 | 0,554831812 | 0,41920723 | Cox14 | cytochrome c oxidase assembly protein 14 [Source:MGI Symbol;Acc:MGI:1913629] | 100,00 | 94,2 |
| ENSMUSG00000025935 | 735,7866345 | 685,3 | 948,7 | -0,493819749 | 0,54982757 | Tram1 | translocating chain-associating membrane protein 1 [Source:MGI Symbol;Acc:MGI:1919515] | 99,85 | 90,3 |
| ENSMUSG00000052681 | 1164,332066 | 1437,6 | 947,0 | 0,654324959 | 0,57373196 | Rap1b | RAS related protein 1b [Source:MGI Symbol;Acc:MGI:894315] | 100,00 | 95,0 |
| ENSMUSG00000024143 | 1103,913638 | 995,5 | 943,2 | 0,197347646 | 0,85582634 | Rhoq | ras homolog gene family, member Q [Source:MGI Symbol;Acc:MGI:1931553] | 95,49 | 71,3 |
| ENSMUSG00000002475 | 1780,143393 | 2700,4 | 937,8 | 1,783953218 | 0,09876958 | Abhd3 | abhydrolase domain containing 3 [Source:MGI Symbol;Acc:MGI:2147183] | 99,95 | 94,6 |
| ENSMUSG00000071014 | 1687,194817 | 2452,4 | 935,5 | 1,564716687 | 0,05061445 | Ndufb6 | NADH dehydrogenase (ubiquinone) 1 beta subcomplex, 6 [Source:MGI Symbol;Acc:MGI:2684983] | 100,00 | 86,3 |
| ENSMUSG00000060802 | 1483,743396 | 1990,0 | 934,4 | 0,985256683 | 0,25469056 | B2m | beta-2 microglobulin [Source:MGI Symbol;Acc:MGI:88127] | 100,00 | 98,7 |
| ENSMUSG00000029066 | 829,070818 | 959,9 | 930,9 | 0,232542546 | 0,75775724 | Mrpl20 | mitochondrial ribosomal protein L20 [Source:MGI Symbol;Acc:MGI:2137221] | 80,92 | 78,2 |

| | | | | | | | | | |
|---|---|---|---|---|---|---|---|---|---|
| ENSMUSG00000039703 | 897,373202 | 579,2 | 930,4 | -0,555240277 | 0,56837751 | Nploc4 | nuclear protein localization 4 homolog (S. cerevisiae) [Source:MGI Symbol;Acc:MGI:2679787] | 98,55 | 91,0 |
| ENSMUSG00000024181 | 1224,202179 | 1471,3 | 929,5 | 0,706914619 | 0,26977312 | Mrpl28 | mitochondrial ribosomal protein L28 [Source:MGI Symbol;Acc:MGI:1915861] | 100,00 | 99,7 |
| ENSMUSG00000002010 | 1303,748871 | 1494,9 | 926,6 | 0,966847762 | 0,25614213 | Idh3g | isocitrate dehydrogenase 3 (NAD+), gamma [Source:MGI Symbol;Acc:MGI:1099463] | 100,00 | 97,4 |
| ENSMUSG00000033307 | 1168,438261 | 1457,3 | 921,3 | 0,663923976 | 0,25771368 | Mif | macrophage migration inhibitory factor [Source:MGI Symbol;Acc:MGI:96982] | 100,00 | 96,3 |
| ENSMUSG00000021737 | 804,7695807 | 792,8 | 919,4 | -0,222841377 | 0,76912632 | Psmd6 | proteasome (prosome, macropain) 26S subunit, non-ATPase, 6 [Source:MGI Symbol;Acc:MGI:1913663] | 98,67 | 93,7 |
| ENSMUSG00000024571 | 809,5332837 | 754,9 | 918,9 | -0,236890394 | 0,73915645 | Gm16286 | predicted gene 16286 [Source:MGI Symbol;Acc:MGI:3833940] | 96,94 | 90,8 |
| ENSMUSG00000059119 | 1167,257681 | 1458,8 | 915,7 | 0,777219066 | 0,39317711 | Nap1l4 | nucleosome assembly protein 1-like 4 [Source:MGI Symbol;Acc:MGI:1316687] | 98,55 | 89,3 |
| ENSMUSG00000033917 | 1455,20578 | 1855,1 | 914,1 | 1,023608629 | 0,5026755 | Gde1 | glycerophosphodiester phosphodiesterase 1 [Source:MGI Symbol;Acc:MGI:1891827] | 96,85 | 92,5 |
| ENSMUSG00000053898 | 1223,796248 | 1743,8 | 913,6 | 1,135341474 | 0,4846002 | Ech1 | enoyl coenzyme A hydratase 1, peroxisomal [Source:MGI Symbol;Acc:MGI:1858208] | 96,69 | 92,6 |
| ENSMUSG00000066900 | 818,4151565 | 698,7 | 912,5 | -0,269946215 | 0,72923704 | Suds3 | suppressor of defective silencing 3 homolog (S. cerevisiae) [Source:MGI Symbol;Acc:MGI:1919204] | 94,49 | 85,4 |
| ENSMUSG00000032554 | 923,6747899 | 760,6 | 911,8 | -0,168612064 | 0,91056121 | Trf | transferrin [Source:MGI Symbol;Acc:MGI:98821] | 87,62 | 77,4 |
| ENSMUSG00000002274 | 1230,953804 | 1061,3 | 911,6 | 0,249456889 | 0,73417041 | Metrn | meteorin, glial cell differentiation regulator [Source:MGI Symbol;Acc:MGI:1917333] | 98,05 | 90,2 |
| ENSMUSG00000027303 | 1376,772641 | 1923,7 | 910,8 | 1,334168526 | 0,44816894 | Ptpra | protein tyrosine phosphatase, receptor type, A [Source:MGI Symbol;Acc:MGI:97808] | 96,76 | 81,4 |
| ENSMUSG00000029840 | 2426,25063 | 3793,9 | 910,6 | 2,286686601 | 0,0712134 | Mtpn | myotrophin [Source:MGI Symbol;Acc:MGI:99445] | 99,65 | 74,4 |
| ENSMUSG00000016256 | 735,1734977 | 661,5 | 909,2 | -0,364569012 | 0,69762682 | Ctsz | cathepsin Z [Source:MGI Symbol;Acc:MGI:1891190] | 97,57 | 92,8 |
| ENSMUSG00000025978 | 1318,554474 | 1965,6 | 905,9 | 1,475335969 | 0,43828489 | Rftn2 | raftlin family member 2 [Source:MGI Symbol;Acc:MGI:1921263] | 87,43 | 75,6 |
| ENSMUSG00000022744 | 910,2207863 | 1157,2 | 901,3 | 0,730250347 | 0,65420769 | Cldnd1 | claudin domain containing 1 [Source:MGI Symbol;Acc:MGI:2447860] | 96,60 | 86,5 |
| ENSMUSG00000031604 | 1335,711725 | 2088,3 | 899,8 | 1,447567898 | 0,31775604 | Msmo1 | methylsterol monoxygenase 1 [Source:MGI Symbol;Acc:MGI:1913484] | 87,03 | 76,6 |
| ENSMUSG00000071662 | 716,3441132 | 636,4 | 899,2 | -0,486837486 | 0,55793788 | Polr2g | polymerase (RNA) II (DNA directed) polypeptide G [Source:MGI Symbol;Acc:MGI:1914960] | 100,00 | 99,7 |
| ENSMUSG00000090639 | 996,3681534 | 835,9 | 899,2 | -0,035054097 | 0,97709704 | Gm20425 | predicted gene 20425 [Source:MGI Symbol;Acc:MGI:5141890] | 98,25 | 79,8 |
| ENSMUSG00000027984 | 1052,938455 | 1268,7 | 897,9 | 0,615014144 | 0,48303991 | Hadh | hydroxyacyl-Coenzyme A dehydrogenase [Source:MGI Symbol;Acc:MGI:96009] | 98,68 | 95,0 |
| ENSMUSG00000020440 | 975,880091 | 949,5 | 895,7 | 0,084081121 | 0,91384494 | Arf5 | ADP-ribosylation factor 5 [Source:MGI Symbol;Acc:MGI:99434] | 99,94 | 93,0 |
| ENSMUSG00000024132 | 1397,792358 | 2094,5 | 893,3 | 1,457084802 | 0,11405528 | Eci1 | enoyl-Coenzyme A delta isomerase 1 [Source:MGI Symbol;Acc:MGI:94871] | 100,00 | 99,1 |
| ENSMUSG00000029146 | 1021,902236 | 1209,5 | 892,7 | 0,653407651 | 0,46938535 | Snx17 | sorting nexin 17 [Source:MGI Symbol;Acc:MGI:2387801] | 99,79 | 94,0 |
| ENSMUSG00000026229 | 695,8995615 | 636,4 | 889,1 | -0,292394968 | 0,75242712 | Psmd1 | proteasome (prosome, macropain) 26S subunit, non-ATPase, 1 [Source:MGI Symbol;Acc:MGI:1917497] | 99,64 | 85,9 |
| ENSMUSG00000031617 | 1271,639875 | 1683,6 | 886,8 | 1,271036502 | 0,2501066 | Tmem184c | transmembrane protein 184C [Source:MGI Symbol;Acc:MGI:2384562] | 100,00 | 86,4 |
| ENSMUSG00000056820 | 950,8421868 | 1050,5 | 883,9 | 0,245485652 | 0,81564567 | Tsnax | translin-associated factor X [Source:MGI Symbol;Acc:MGI:1855672] | 98,90 | 94,0 |
| ENSMUSG00000057858 | 797,724348 | 842,7 | 883,5 | 0,120977504 | 0,93240742 | Fam204a | family with sequence similarity 204, member A [Source:MGI Symbol;Acc:MGI:1289174] | 95,04 | 77,3 |
| ENSMUSG00000024429 | 623,4429095 | 688,2 | 872,9 | -0,253328917 | 0,75134769 | Gnl1 | guanine nucleotide binding protein-like 1 [Source:MGI Symbol;Acc:MGI:95764] | 94,45 | 76,7 |
| ENSMUSG00000029432 | 904,092197 | 1253,5 | 871,9 | 1,791324756 | 0,12691383 | Gbas | glioblastoma amplified sequence [Source:MGI Symbol;Acc:MGI:1278343] | 99,62 | 75,4 |
| ENSMUSG00000026341 | 800,1141534 | 781,5 | 869,6 | -0,090778142 | 0,92801708 | Actr3 | ARP3 actin-related protein 3 [Source:MGI Symbol;Acc:MGI:1921367] | 88,57 | 78,5 |
| ENSMUSG00000035953 | 894,8177084 | 1103,4 | 868,0 | 0,350257067 | 0,63575872 | Tmem55b | transmembrane protein 55b [Source:MGI Symbol;Acc:MGI:2448501] | 88,10 | 72,9 |
| ENSMUSG00000032397 | 837,3683441 | 662,7 | 864,7 | 0,133237853 | 0,89936894 | Tipin | timeless interacting protein [Source:MGI Symbol;Acc:MGI:1921571] | 100,00 | 95,4 |
| ENSMUSG00000007950 | 1208,718038 | 1640,6 | 863,0 | 0,926636536 | 0,56519849 | Abhd8 | abhydrolase domain containing 8 [Source:MGI Symbol;Acc:MGI:1918946] | 99,85 | 99,3 |
| ENSMUSG00000040055 | 2709,029295 | 4828,8 | 862,6 | 2,627236048 | 0,13988341 | Gjb6 | gap junction protein, beta 6 [Source:MGI Symbol;Acc:MGI:107588] | 100,00 | 92,2 |
| ENSMUSG00000031987 | 783,5666907 | 622,1 | 861,6 | -0,440890829 | 0,64384232 | Egln1 | egl-9 family hypoxia-inducible factor 1 [Source:MGI Symbol;Acc:MGI:1932286] | 98,36 | 86,7 |
| ENSMUSG00000053398 | 1184,192491 | 1227,9 | 860,5 | 0,501847488 | 0,46619662 | Phgdh | 3-phosphoglycerate dehydrogenase [Source:MGI Symbol;Acc:MGI:1355330] | 86,98 | 73,6 |
| ENSMUSG00000025006 | 1405,186557 | 2218,6 | 860,5 | 1,467414939 | 0,14772198 | Sorbs1 | sorbin and SH3 domain containing 1 [Source:MGI Symbol;Acc:MGI:700014] | 98,91 | 78,3 |
| ENSMUSG00000066568 | 815,165032 | 610,1 | 858,9 | -0,441019251 | 0,73312976 | Lsm14a | LSM14 homolog A (SCD6, S. cerevisiae) [Source:MGI Symbol;Acc:MGI:1914320] | 93,22 | 77,4 |
| ENSMUSG00000016946 | 1293,498192 | 1829,7 | 857,7 | 1,182484872 | 0,3950818 | Kctd5 | potassium channel tetramerisation domain containing 5 [Source:MGI Symbol;Acc:MGI:1916509] | 99,65 | 94,1 |
| ENSMUSG00000032067 | 1368,295601 | 1913,1 | 854,8 | 1,407862845 | 0,38744793 | Pts | 6-pyruvoyl-tetrahydropterin synthase [Source:MGI Symbol;Acc:MGI:1338783] | 100,00 | 97,4 |
| ENSMUSG00000028822 | 838,3958478 | 1063,4 | 851,0 | 0,319293584 | 0,72102051 | Tmem50a | transmembrane protein 50A [Source:MGI Symbol;Acc:MGI:1919067] | 99,74 | 96,8 |
| ENSMUSG00000107283 | 1143,110132 | 1711,7 | 849,6 | 1,010450625 | 0,44671787 | Mpv17 | MpV17 mitochondrial inner membrane protein [Source:MGI Symbol;Acc:MGI:97138] | 96,04 | 74,6 |
| ENSMUSG00000022892 | 851,1550077 | 633,0 | 848,1 | -0,254684417 | 0,81127255 | App | amyloid beta (A4) precursor protein [Source:MGI Symbol;Acc:MGI:88059] | 100,00 | 87,6 |
| ENSMUSG00000020152 | 634,4023311 | 547,9 | 846,6 | -0,220936848 | 0,82631484 | Actr2 | ARP2 actin-related protein 2 [Source:MGI Symbol;Acc:MGI:1913963] | 97,78 | 70,3 |
| ENSMUSG00000028837 | 1282,479705 | 1780,2 | 846,2 | 1,128863843 | 0,19702765 | Psmb2 | proteasome (prosome, macropain) subunit, beta type 2 [Source:MGI Symbol;Acc:MGI:1347045] | 100,00 | 95,6 |
| ENSMUSG00000020708 | 1095,212997 | 1201,7 | 844,5 | 0,538411185 | 0,35972104 | Psmc5 | protease (prosome, macropain) 26S subunit, ATPase 5 [Source:MGI Symbol;Acc:MGI:105047] | 100,00 | 99,3 |
| ENSMUSG00000015806 | 1230,172131 | 1494,0 | 837,6 | 0,964959277 | 0,19965149 | Qdpr | quinoid dihydropteridine reductase [Source:MGI Symbol;Acc:MGI:97836] | 86,48 | 75,6 |
| ENSMUSG00000028277 | 809,2865015 | 740,7 | 836,3 | -0,099166042 | 0,91242238 | Ube2j1 | ubiquitin-conjugating enzyme E2J 1 [Source:MGI Symbol;Acc:MGI:1926245] | 99,61 | 70,2 |
| ENSMUSG00000051853 | 1618,215241 | 2259,2 | 834,6 | 1,514323475 | 0,33628593 | Arf3 | ADP-ribosylation factor 3 [Source:MGI Symbol;Acc:MGI:99432] | 94,79 | 81,7 |
| ENSMUSG00000025162 | 902,83561 | 908,5 | 834,4 | 0,122519645 | 0,87229828 | Csnk1d | casein kinase 1, delta [Source:MGI Symbol;Acc:MGI:1355272] | 97,66 | 90,9 |
| ENSMUSG00000029048 | 929,3419855 | 977,6 | 834,3 | 0,372137614 | 0,69939646 | Rer1 | RER1 retention in endoplasmic reticulum 1 homolog (S. cerevisiae) [Source:MGI Symbol;Acc:MGI:1915080] | 99,87 | 86,8 |
| ENSMUSG00000028851 | 954,5281995 | 1099,7 | 831,4 | 0,492586571 | 0,52768046 | Nudc | nuclear distribution gene C homolog (Aspergillus) [Source:MGI Symbol;Acc:MGI:106014] | 100,00 | 97,4 |
| ENSMUSG00000024914 | 992,1688875 | 1155,7 | 830,5 | 0,497915252 | 0,40405259 | Drap1 | Dr1 associated protein 1 (negative cofactor 2 alpha) [Source:MGI Symbol;Acc:MGI:1913806] | 93,41 | 80,9 |
| ENSMUSG00000022553 | 1016,616211 | 1358,3 | 829,6 | 0,908500214 | 0,21893496 | Maf1 | MAF1 homolog (S. cerevisiae) [Source:MGI Symbol;Acc:MGI:1916127] | 92,24 | 92,0 |
| ENSMUSG00000025151 | 720,9834305 | 737,8 | 829,6 | -0,208072656 | 0,90227664 | Maged1 | melanoma antigen, family D, 1 [Source:MGI Symbol;Acc:MGI:1930187] | 97,20 | 93,9 |
| ENSMUSG00000035376 | 786,3835761 | 1171,8 | 823,7 | 0,755496422 | 0,41263705 | Hacd2 | 3-hydroxyacyl-CoA dehydratase 2 [Source:MGI Symbol;Acc:MGI:1918007] | 99,20 | 70,0 |
| ENSMUSG00000002332 | 1088,426185 | 1531,3 | 823,2 | 0,951749037 | 0,23721484 | Dhrs1 | dehydrogenase/reductase (SDR family) member 1 [Source:MGI Symbol;Acc:MGI:1196314] | 97,79 | 95,6 |
| ENSMUSG00000022122 | 1443,446932 | 1861,1 | 823,0 | 1,396948273 | 0,3661564 | Ednrb | endothelin receptor type B [Source:MGI Symbol;Acc:MGI:102720] | 100,00 | 80,4 |
| ENSMUSG00000007721 | 835,5039707 | 975,4 | 816,8 | 0,336870777 | 0,66300896 | Ccdc124 | coiled-coil domain containing 124 [Source:MGI Symbol;Acc:MGI:1916403] | 100,00 | 99,9 |
| ENSMUSG00000037060 | 1737,247342 | 2782,9 | 813,4 | 2,497993533 | 0,1541188 | Prkcdbp | protein kinase C, delta binding protein [Source:MGI Symbol;Acc:MGI:1923422] | 100,00 | 81,6 |
| ENSMUSG00000038615 | 717,8557585 | 649,0 | 812,9 | -0,272137795 | 0,81348521 | Nfe2l1 | nuclear factor, erythroid derived 2,-like 1 [Source:MGI Symbol;Acc:MGI:99421] | 96,66 | 80,4 |
| ENSMUSG00000061032 | 750,9793044 | 870,6 | 808,9 | 0,268130708 | 0,8771957 | Rrp1 | ribosomal RNA processing 1 homolog (S. cerevisiae) [Source:MGI Symbol;Acc:MGI:1203500] | 98,49 | 95,0 |
| ENSMUSG00000038936 | 1272,14602 | 2078,2 | 807,8 | 1,372691404 | 0,43042209 | Sccpdh | saccharopine dehydrogenase (putative) [Source:MGI Symbol;Acc:MGI:1924486] | 92,71 | 80,9 |
| ENSMUSG00000029103 | 834,3562474 | 960,2 | 804,7 | 0,35725986 | 0,85345701 | Lrpap1 | low density lipoprotein receptor-related protein associated protein 1 [Source:MGI Symbol;Acc:MGI:96829] | 98,97 | 87,8 |
| ENSMUSG00000022564 | 1535,34632 | 2465,7 | 804,4 | 1,78640418 | 0,05464049 | Grina | nate receptor, ionotropic, N-methyl D-aspartate-associated protein 1 (glutamate binding) [Source:MGI Symbol;Acc:MGI:191 | 100,00 | 94,2 |
| ENSMUSG00000059336 | 742,7972032 | 1066,1 | 803,1 | 0,6079018 | 0,77560734 | Slc14a1 | solute carrier family 14 (urea transporter), member 1 [Source:MGI Symbol;Acc:MGI:1351654] | 95,88 | 83,2 |
| ENSMUSG00000001666 | 1083,912848 | 1478,4 | 802,9 | 0,902639993 | 0,18380568 | Ddt | D-dopachrome tautomerase [Source:MGI Symbol;Acc:MGI:1298381] | 95,86 | 81,6 |
| ENSMUSG00000038459 | 860,1931551 | 829,8 | 801,8 | 0,12755565 | 0,9359459 | Abhd17c | abhydrolase domain containing 17C [Source:MGI Symbol;Acc:MGI:1917428] | 100,00 | 90,8 |

| | | | | | | | | | |
|---|---|---|---|---|---|---|---|---|---|
| ENSMUSG00000003346 | 1078,572004 | 1115,9 | 799,2 | 0,742127465 | 0,33993732 | Abhd17a | abhydrolase domain containing 17A [Source:MGI Symbol;Acc:MGI:106388] | 95,28 | 86,8 |
| ENSMUSG00000024072 | 744,6102538 | 958,1 | 798,2 | 0,311875542 | 0,80756215 | Yipf4 | Yip1 domain family, member 4 [Source:MGI Symbol;Acc:MGI:1915114] | 100,00 | 76,9 |
| ENSMUSG00000020646 | 492,5807232 | 558,8 | 797,0 | -0,422506662 | 0,81720334 | Mboat2 | membrane bound O-acyltransferase domain containing 2 [Source:MGI Symbol;Acc:MGI:1914466] | 100,00 | 88,0 |
| ENSMUSG00000042043 | 709,6949598 | 680,2 | 796,3 | -0,162334507 | 0,87333017 | Tbca | tubulin cofactor A [Source:MGI Symbol;Acc:MGI:107549] | 100,00 | 100,0 |
| ENSMUSG00000016528 | 862,3900987 | 507,8 | 794,7 | -0,459457181 | 0,60850517 | Mapkapk2 | MAP kinase-activated protein kinase 2 [Source:MGI Symbol;Acc:MGI:109298] | 92,67 | 83,5 |
| ENSMUSG00000024780 | 723,6260775 | 965,6 | 794,2 | 0,580218229 | 0,58162868 | Cdc37l1 | cell division cycle 37-like 1 [Source:MGI Symbol;Acc:MGI:1914322] | 100,00 | 72,4 |
| ENSMUSG00000013662 | 588,453685 | 553,9 | 793,1 | -0,272241613 | 0,77558601 | Atad1 | ATPase family, AAA domain containing 1 [Source:MGI Symbol;Acc:MGI:1915229] | 99,96 | 79,4 |
| ENSMUSG00000036106 | 729,2383552 | 507,6 | 793,0 | -0,478681444 | 0,58414223 | Prr5 | proline rich 5 (renal) [Source:MGI Symbol;Acc:MGI:1924714] | 90,14 | 89,5 |
| ENSMUSG00000015937 | 827,8822547 | 883,2 | 786,4 | 0,206277316 | 0,77737549 | H2afy | H2A histone family, member Y [Source:MGI Symbol;Acc:MGI:1349392] | 96,49 | 88,2 |
| ENSMUSG00000021508 | 1705,462246 | 2766,7 | 785,2 | 1,89069001 | 0,12373244 | Cxcl14 | chemokine (C-X-C motif) ligand 14 [Source:MGI Symbol;Acc:MGI:1888514] | 99,87 | 90,4 |
| ENSMUSG00000035478 | 935,8652188 | 1078,1 | 784,8 | 0,494014799 | 0,4858251 | Mbd3 | methyl-CpG binding domain protein 3 [Source:MGI Symbol;Acc:MGI:1333812] | 90,56 | 79,9 |
| ENSMUSG00000091337 | 1164,953271 | 1576,9 | 782,2 | 1,081656085 | 0,26375864 | Eid1 | EP300 interacting inhibitor of differentiation 1 [Source:MGI Symbol;Acc:MGI:1889651] | 100,00 | 95,4 |
| ENSMUSG00000038696 | 835,4883331 | 1151,8 | 777,0 | 0,598128141 | 0,66300896 | Mapkap1 | mitogen-activated protein kinase associated protein 1 [Source:MGI Symbol;Acc:MGI:2444554] | 96,81 | 88,8 |
| ENSMUSG00000030842 | 951,9938407 | 1184,3 | 775,3 | 0,808176234 | 0,32103739 | Lamtor1 | late endosomal/lysosomal adaptor, MAPK and MTOR activator 1 [Source:MGI Symbol;Acc:MGI:1913758] | 81,92 | 78,0 |
| ENSMUSG00000030890 | 1112,022015 | 1448,1 | 775,1 | 0,879711672 | 0,17408053 | Ilk | integrin linked kinase [Source:MGI Symbol;Acc:MGI:1195267] | 93,20 | 74,3 |
| ENSMUSG00000002580 | 1229,297054 | 1701,1 | 773,7 | 1,190036797 | 0,10371464 | Mien1 | migration and invasion enhancer 1 [Source:MGI Symbol;Acc:MGI:1913678] | 99,51 | 97,1 |
| ENSMUSG00000019978 | 696,198894 | 653,6 | 773,7 | -0,114053122 | 0,89403347 | Epb41l2 | erythrocyte membrane protein band 4.1 like 2 [Source:MGI Symbol;Acc:MGI:103009] | 91,73 | 79,5 |
| ENSMUSG00000068732 | 907,6443652 | 1189,5 | 773,3 | 0,833013893 | 0,63224541 | Tmem167b | transmembrane protein 167B [Source:MGI Symbol;Acc:MGI:1914745] | 99,07 | 78,9 |
| ENSMUSG00000011884 | 803,2296258 | 939,8 | 770,0 | 0,655836488 | 0,70917746 | Gltp | glycolipid transfer protein [Source:MGI Symbol;Acc:MGI:1929253] | 99,70 | 91,6 |
| ENSMUSG00000031848 | 685,0610955 | 633,8 | 768,6 | -0,279791416 | 0,69092144 | Lsm4 | LSM4 homolog, U6 small nuclear RNA associated (S. cerevisiae) [Source:MGI Symbol;Acc:MGI:1354692] | 99,89 | 90,5 |
| ENSMUSG00000023048 | 1209,559864 | 1468,2 | 766,9 | 1,155785267 | 0,14823092 | Prr13 | proline rich 13 [Source:MGI Symbol;Acc:MGI:1913401] | 96,52 | 74,3 |
| ENSMUSG00000055850 | 1114,922189 | 1440,3 | 765,7 | 0,910829515 | 0,11777558 | Rnf181 | ring finger protein 181 [Source:MGI Symbol;Acc:MGI:1913760] | 82,89 | 76,4 |
| ENSMUSG00000030704 | 2224,530862 | 3768,5 | 764,7 | 2,310174685 | 0,07820041 | Rab6a | RAB6A, member RAS oncogene family [Source:MGI Symbol;Acc:MGI:894313] | 97,09 | 87,5 |
| ENSMUSG00000003813 | 862,3489401 | 1158,6 | 763,6 | 0,65278152 | 0,35984334 | Rad23a | RAD23a homolog (S. cerevisiae) [Source:MGI Symbol;Acc:MGI:105126] | 97,09 | 79,4 |
| ENSMUSG00000005846 | 687,1644536 | 606,5 | 763,2 | 0,354304661 | 0,84397612 | Rsl1d1 | ribosomal L1 domain containing 1 [Source:MGI Symbol;Acc:MGI:1913659] | 98,59 | 82,0 |
| ENSMUSG00000032350 | 592,3428169 | 926,5 | 762,6 | 0,386580767 | 0,78227477 | Gclc | glutamate-cysteine ligase, catalytic subunit [Source:MGI Symbol;Acc:MGI:104990] | 99,81 | 89,0 |
| ENSMUSG00000025173 | 909,740902 | 923,8 | 762,6 | 0,525451917 | 0,51368066 | Wdr45b | WD repeat domain 45B [Source:MGI Symbol;Acc:MGI:1914090] | 97,30 | 88,8 |
| ENSMUSG00000024327 | 732,5019815 | 714,1 | 762,1 | -0,07153781 | 0,93169628 | Slc39a7 | solute carrier family 39 (zinc transporter), member 7 [Source:MGI Symbol;Acc:MGI:95909] | 99,79 | 96,5 |
| ENSMUSG00000015932 | 1717,683141 | 2527,6 | 761,7 | 1,739717102 | 0,11902307 | Dstn | destrin [Source:MGI Symbol;Acc:MGI:1929270] | 82,50 | 72,5 |
| ENSMUSG00000024588 | 976,9853133 | 1386,2 | 759,2 | 1,008233786 | 0,20090917 | Fech | ferrochelatase [Source:MGI Symbol;Acc:MGI:95513] | 100,00 | 84,0 |
| ENSMUSG00000030876 | 894,7958077 | 882,9 | 757,3 | 0,25031761 | 0,78448968 | Mettl9 | methyltransferase like 9 [Source:MGI Symbol;Acc:MGI:1914862] | 92,20 | 83,2 |
| ENSMUSG00000024194 | 828,5645212 | 914,9 | 753,7 | 0,27677139 | 0,67261414 | Cuta | cutA divalent cation tolerance homolog [Source:MGI Symbol;Acc:MGI:1914925] | 100,00 | 99,4 |
| ENSMUSG00000022265 | 754,358687 | 635,6 | 751,8 | -0,24026696 | 0,80866253 | Ank | progressive ankylosis [Source:MGI Symbol;Acc:MGI:3045421] | 99,64 | 87,4 |
| ENSMUSG00000090247 | 1423,019661 | 1974,3 | 743,8 | 1,719072058 | 0,09983077 | Bloc1s1 | biogenesis of lysosomal organelles complex-1, subunit 1 [Source:MGI Symbol;Acc:MGI:1195276] | 88,20 | 70,8 |
| ENSMUSG00000038412 | 1031,166307 | 1350,4 | 743,5 | 0,93279225 | 0,15807559 | Higd1a | HIG1 domain family, member 1A [Source:MGI Symbol;Acc:MGI:1930666] | 98,08 | 86,5 |
| ENSMUSG00000021681 | 823,07781 | 941,7 | 739,3 | 0,429500231 | 0,6591569 | Aggf1 | angiogenic factor with G patch and FHA domains 1 [Source:MGI Symbol;Acc:MGI:1913799] | 100,00 | 94,1 |
| ENSMUSG00000051154 | 1012,79529 | 1353,0 | 738,1 | 1,029274785 | 0,21449811 | Commd3 | COMM domain containing 3 [Source:MGI Symbol;Acc:MGI:88218] | 97,50 | 95,0 |
| ENSMUSG00000062963 | 810,2945623 | 912,8 | 737,1 | 0,40343172 | 0,66322016 | Ufc1 | ubiquitin-fold modifier conjugating enzyme 1 [Source:MGI Symbol;Acc:MGI:1913405] | 98,52 | 87,5 |
| ENSMUSG00000028833 | 939,9879327 | 1410,6 | 737,1 | 0,897639918 | 0,50858103 | Ncdn | neurochondrin [Source:MGI Symbol;Acc:MGI:1347351] | 99,75 | 87,5 |
| ENSMUSG00000041733 | 560,7696482 | 722,5 | 737,0 | 0,284916479 | 0,86223773 | Coq5 | coenzyme Q5 methyltransferase [Source:MGI Symbol;Acc:MGI:1098643] | 99,57 | 87,2 |
| ENSMUSG00000022634 | 1029,255979 | 1251,4 | 735,3 | 0,769084623 | 0,16240307 | Yaf2 | YY1 associated factor 2 [Source:MGI Symbol;Acc:MGI:1914307] | 100,00 | 94,6 |
| ENSMUSG00000030327 | 637,9730537 | 727,1 | 734,2 | 0,120071868 | 0,90252666 | Necap1 | NECAP endocytosis associated 1 [Source:MGI Symbol;Acc:MGI:1914852] | 97,31 | 90,1 |
| ENSMUSG00000037119 | 743,3294598 | 745,6 | 734,1 | 0,033856459 | 0,98186729 | D15Ertd621e | DNA segment, Chr 15, ERATO Doi 621, expressed [Source:MGI Symbol;Acc:MGI:1277178] | 99,48 | 78,0 |
| ENSMUSG00000027221 | 776,7850458 | 968,1 | 733,8 | 0,416042624 | 0,54983143 | Chst1 | carbohydrate (keratan sulfate Gal-6) sulfotransferase 1 [Source:MGI Symbol;Acc:MGI:1924219] | 100,00 | 93,9 |
| ENSMUSG00000028821 | 880,3133045 | 1205,3 | 733,5 | 0,794942481 | 0,32770533 | Syf2 | SYF2 homolog, RNA splicing factor (S. cerevisiae) [Source:MGI Symbol;Acc:MGI:1915842] | 99,32 | 91,7 |
| ENSMUSG00000037916 | 1049,73841 | 1160,5 | 727,1 | 0,741271043 | 0,29384829 | Ndufv1 | NADH dehydrogenase (ubiquinone) flavoprotein 1 [Source:MGI Symbol;Acc:MGI:107851] | 98,17 | 85,8 |
| ENSMUSG00000002102 | 946,649786 | 978,9 | 726,3 | 0,516186519 | 0,51048399 | Psmc3 | proteasome (prosome, macropain) 26S subunit, ATPase 3 [Source:MGI Symbol;Acc:MGI:1098754] | 86,64 | 77,0 |
| ENSMUSG00000052934 | 496,3878016 | 715,5 | 725,5 | -0,058321333 | 0,94606362 | Fbxo31 | F-box protein 31 [Source:MGI Symbol;Acc:MGI:1354708] | 93,39 | 80,2 |
| ENSMUSG00000020076 | 711,631763 | 504,8 | 722,1 | -0,533431131 | 0,37522079 | Ddx50 | DEAD (Asp-Glu-Ala-Asp) box polypeptide 50 [Source:MGI Symbol;Acc:MGI:2182303] | 100,00 | 97,0 |
| ENSMUSG00000031610 | 1425,967324 | 1880,2 | 719,7 | 1,961262984 | 0,25746908 | Scrg1 | scrapie responsive gene 1 [Source:MGI Symbol;Acc:MGI:1328308] | 100,00 | 93,6 |
| ENSMUSG00000063856 | 1097,813494 | 1457,5 | 719,3 | 1,076476873 | 0,23064482 | Gpx1 | glutathione peroxidase 1 [Source:MGI Symbol;Acc:MGI:104887] | 95,49 | 80,1 |
| ENSMUSG00000022789 | 638,7061612 | 871,4 | 718,5 | 0,273164794 | 0,8010503 | Dnm1l | dynamin 1-like [Source:MGI Symbol;Acc:MGI:1921256] | 99,53 | 88,8 |
| ENSMUSG00000030960 | 654,4870015 | 797,2 | 718,5 | 0,201808845 | 0,834944 | Mettl10 | methyltransferase like 10 [Source:MGI Symbol;Acc:MGI:1919346] | 99,58 | 94,6 |
| ENSMUSG00000021981 | 1246,716869 | 1226,2 | 712,6 | 1,390905299 | 0,37689946 | Cab39l | calcium binding protein 39-like [Source:MGI Symbol;Acc:MGI:1914081] | 99,99 | 73,9 |
| ENSMUSG00000063576 | 692,8404691 | 900,2 | 711,5 | 0,553550601 | 0,52780138 | Klhdc3 | kelch domain containing 3 [Source:MGI Symbol;Acc:MGI:2651568] | 99,09 | 92,8 |
| ENSMUSG00000040374 | 865,6338869 | 1165,0 | 709,0 | 1,014525087 | 0,54714902 | Pex2 | peroxisomal biogenesis factor 2 [Source:MGI Symbol;Acc:MGI:107486] | 98,28 | 72,1 |
| ENSMUSG00000031532 | 1726,347693 | 2414,3 | 707,6 | 1,786191326 | 0,10389727 | Saraf | store-operated calcium entry-associated regulatory factor [Source:MGI Symbol;Acc:MGI:1915137] | 99,58 | 87,4 |
| ENSMUSG00000031708 | 1272,793778 | 1634,0 | 707,0 | 1,387489693 | 0,10184166 | Tecr | trans-2,3-enoyl-CoA reductase [Source:MGI Symbol;Acc:MGI:1915408] | 88,58 | 84,4 |
| ENSMUSG00000031197 | 1331,42688 | 1365,5 | 704,1 | 1,195085278 | 0,32537762 | Vbp1 | von Hippel-Lindau binding protein 1 [Source:MGI Symbol;Acc:MGI:1333804] | 91,42 | 70,4 |
| ENSMUSG00000003721 | 778,686514 | 563,2 | 697,1 | -0,312094644 | 0,72815937 | Insig2 | insulin induced gene 2 [Source:MGI Symbol;Acc:MGI:1920249] | 82,69 | 74,6 |
| ENSMUSG00000022982 | 605,9676683 | 695,4 | 695,1 | 0,099937796 | 0,91480252 | Sod1 | superoxide dismutase 1, soluble [Source:MGI Symbol;Acc:MGI:98351] | 100,00 | 97,7 |
| ENSMUSG00000019738 | 529,1859337 | 501,3 | 692,9 | -0,408675243 | 0,55754405 | Polr2i | polymerase (RNA) II (DNA directed) polypeptide I [Source:MGI Symbol;Acc:MGI:1917170] | 95,85 | 90,1 |
| ENSMUSG00000041236 | 1030,476193 | 1540,2 | 692,6 | 1,195286701 | 0,5987965 | Vps41 | vacuolar protein sorting 41 (yeast) [Source:MGI Symbol;Acc:MGI:1929215] | 99,91 | 86,2 |
| ENSMUSG00000020571 | 832,0558957 | 629,0 | 689,3 | -0,192348461 | 0,78589239 | Pdia6 | protein disulfide isomerase associated 6 [Source:MGI Symbol;Acc:MGI:1919103] | 99,20 | 77,3 |
| ENSMUSG00000106918 | 880,7589684 | 960,1 | 686,9 | 0,516907392 | 0,3974293 | Mrpl33 | mitochondrial ribosomal protein L33 [Source:MGI Symbol;Acc:MGI:2137225] | 99,37 | 93,5 |
| ENSMUSG00000021025 | 487,1492237 | 704,4 | 686,1 | 0,243778801 | 0,84309128 | Nfkbia | nuclear factor of kappa light polypeptide gene enhancer in B cells inhibitor, alpha [Source:MGI Symbol;Acc:MGI:104741] | 97,02 | 82,8 |
| ENSMUSG00000037197 | 814,4204808 | 593,6 | 686,0 | -0,071847738 | 0,93242073 | Rbm17 | RNA binding motif protein 17 [Source:MGI Symbol;Acc:MGI:1924188] | 88,67 | 82,7 |
| ENSMUSG00000017615 | 926,433003 | 1084,2 | 682,4 | 1,156001676 | 0,19040611 | Tnfaip1 | tumor necrosis factor, alpha-induced protein 1 (endothelial) [Source:MGI Symbol;Acc:MGI:104961] | 99,23 | 74,1 |
| ENSMUSG00000006736 | 930,6344374 | 1361,0 | 680,3 | 0,931875243 | 0,58546406 | Tspan31 | tetraspanin 31 [Source:MGI Symbol;Acc:MGI:1914375] | 100,00 | 99,6 |

| | | | | | | | | | |
|---|---|---|---|---|---|---|---|---|---|
| ENSMUSG00000028797 | 655,0215881 | 688,1 | 679,1 | 0,102875282 | 0,90856201 | Tmem234 | transmembrane protein 234 [Source:MGI Symbol;Acc:MGI:1924049] | 100,00 | 84,5 |
| ENSMUSG00000023495 | 532,5944691 | 609,6 | 678,9 | -0,113716883 | 0,8946164 | Pcbp4 | poly(rC) binding protein 4 [Source:MGI Symbol;Acc:MGI:1890471] | 78,68 | 72,8 |
| ENSMUSG00000021577 | 2130,211689 | 3327,0 | 677,7 | 2,464535103 | 0,08769736 | Sdha | succinate dehydrogenase complex, subunit A, flavoprotein (Fp) [Source:MGI Symbol;Acc:MGI:1914195] | 99,17 | 88,8 |
| ENSMUSG00000025260 | 715,3242593 | 794,5 | 677,4 | 0,251318599 | 0,73016629 | Hsd17b10 | hydroxysteroid (17-beta) dehydrogenase 10 [Source:MGI Symbol;Acc:MGI:1333871] | 95,77 | 92,4 |
| ENSMUSG00000024835 | 469,4410354 | 509,3 | 677,2 | -0,421516482 | 0,51395515 | Coro1b | coronin, actin binding protein 1B [Source:MGI Symbol;Acc:MGI:1345963] | 100,00 | 98,9 |
| ENSMUSG00000026500 | 918,3381515 | 1237,8 | 674,4 | 1,225816511 | 0,12834316 | Cox20 | COX20 Cox2 chaperone [Source:MGI Symbol;Acc:MGI:1913609] | 100,00 | 92,5 |
| ENSMUSG00000028772 | 741,0638026 | 948,3 | 673,2 | 0,593786805 | 0,42766218 | Zcchc17 | zinc finger, CCHC domain containing 17 [Source:MGI Symbol;Acc:MGI:1919955] | 96,77 | 72,1 |
| ENSMUSG00000021606 | 1196,009876 | 1462,5 | 672,1 | 1,194928299 | 0,06178216 | Ndufs6 | NADH dehydrogenase (ubiquinone) Fe-S protein 6 [Source:MGI Symbol;Acc:MGI:107932] | 97,01 | 86,4 |
| ENSMUSG00000025465 | 1448,458677 | 2378,4 | 668,6 | 2,082114176 | 0,13642338 | Echs1 | enoyl Coenzyme A hydratase, short chain, 1, mitochondrial [Source:MGI Symbol;Acc:MGI:2136460] | 99,36 | 93,8 |
| ENSMUSG00000069744 | 715,3488607 | 631,9 | 668,4 | -0,058656093 | 0,94175231 | Psmb3 | proteasome (prosome, macropain) subunit, beta type 3 [Source:MGI Symbol;Acc:MGI:1347014] | 98,09 | 92,7 |
| ENSMUSG00000022257 | 653,8728732 | 837,2 | 667,0 | 0,645620833 | 0,77688819 | Laptm4b | lysosomal-associated protein transmembrane 4B [Source:MGI Symbol;Acc:MGI:1890494] | 96,22 | 83,1 |
| ENSMUSG00000032112 | 763,8960814 | 936,5 | 666,8 | 0,774479045 | 0,34568212 | Trappc4 | trafficking protein particle complex 4 [Source:MGI Symbol;Acc:MGI:1926211] | 100,00 | 73,4 |
| ENSMUSG00000035297 | 915,2674106 | 1132,0 | 666,6 | 0,8101754 | 0,41960231 | Cops4 | COP9 signalosome subunit 4 [Source:MGI Symbol;Acc:MGI:1349414] | 88,55 | 81,6 |
| ENSMUSG00000058709 | 630,8021693 | 588,8 | 666,3 | 0,17628465 | 0,89936894 | Egln2 | egl-9 family hypoxia-inducible factor 2 [Source:MGI Symbol;Acc:MGI:1932287] | 98,00 | 77,8 |
| ENSMUSG00000008036 | 812,9623764 | 1048,8 | 666,2 | 0,666537119 | 0,34889171 | Ap2s1 | adaptor-related protein complex 2, sigma 1 subunit [Source:MGI Symbol;Acc:MGI:2141861] | 99,96 | 95,8 |
| ENSMUSG00000030541 | 1286,015386 | 1809,3 | 665,9 | 1,420689514 | 0,0553745 | Idh2 | isocitrate dehydrogenase 2 (NADP+), mitochondrial [Source:MGI Symbol;Acc:MGI:96414] | 98,19 | 95,7 |
| ENSMUSG00000030603 | 714,7546582 | 916,1 | 663,4 | 0,781140034 | 0,44419963 | Psmc4 | proteasome (prosome, macropain) 26S subunit, ATPase, 4 [Source:MGI Symbol;Acc:MGI:1346093] | 99,72 | 91,4 |
| ENSMUSG00000047084 | 396,4696694 | 503,0 | 662,7 | -0,053203746 | 0,98014335 | Ngrn | neugrin, neurite outgrowth associated [Source:MGI Symbol;Acc:MGI:1933212] | 99,97 | 98,2 |
| ENSMUSG00000000743 | 1100,530983 | 1463,6 | 662,3 | 1,194387326 | 0,12078464 | Chmp1a | charged multivesicular body protein 1A [Source:MGI Symbol;Acc:MGI:1920159] | 100,00 | 92,6 |
| ENSMUSG00000038552 | 1145,814614 | 1426,4 | 662,1 | 1,561922516 | 0,19196428 | Fndc4 | fibronectin type III domain containing 4 [Source:MGI Symbol;Acc:MGI:1917195] | 98,92 | 74,4 |
| ENSMUSG00000004394 | 557,7421111 | 600,1 | 662,0 | 0,046533938 | 0,98366007 | Tmed4 | transmembrane emp24 protein transport domain containing 4 [Source:MGI Symbol;Acc:MGI:1915070] | 98,40 | 88,8 |
| ENSMUSG00000024197 | 614,8583026 | 724,2 | 661,2 | 0,279087084 | 0,77233446 | Plin3 | perilipin 3 [Source:MGI Symbol;Acc:MGI:1914155] | 99,95 | 91,0 |
| ENSMUSG00000020744 | 698,0520763 | 643,3 | 659,4 | 0,181499225 | 0,85636392 | Slc25a19 | solute carrier family 25 (mitochondrial thiamine pyrophosphate carrier), member 19 [Source:MGI Symbol;Acc:MGI:1914533 | 90,01 | 75,9 |
| ENSMUSG00000022043 | 840,2414799 | 1163,6 | 658,8 | 0,848252551 | 0,4256461 | Trim35 | tripartite motif-containing 35 [Source:MGI Symbol;Acc:MGI:1914104] | 91,41 | 74,3 |
| ENSMUSG00000028757 | 679,2133728 | 625,0 | 656,6 | -0,0574352 | 0,96560075 | Ddost | dolichyl-di-phosphooligosaccharide-protein glycotransferase [Source:MGI Symbol;Acc:MGI:1194508] | 98,24 | 88,1 |
| ENSMUSG00000036850 | 985,7222575 | 1910,1 | 655,5 | 1,517069799 | 0,29379074 | Mrpl41 | mitochondrial ribosomal protein L41 [Source:MGI Symbol;Acc:MGI:1333816] | 100,00 | 97,4 |
| ENSMUSG00000038607 | 1208,3787 | 1845,4 | 654,1 | 1,848236308 | 0,1052698 | Gng10 | guanine nucleotide binding protein (G protein), gamma 10 [Source:MGI Symbol;Acc:MGI:1336169] | 96,90 | 86,0 |
| ENSMUSG00000020536 | 732,1204265 | 1008,7 | 652,1 | 0,930960876 | 0,69017368 | Llgl1 | lethal giant larvae homolog 1 (Drosophila) [Source:MGI Symbol;Acc:MGI:102682] | 99,84 | 73,9 |
| ENSMUSG00000038967 | 2147,769418 | 4005,2 | 651,7 | 2,901867856 | 0,06273351 | Pdk2 | pyruvate dehydrogenase kinase, isoenzyme 2 [Source:MGI Symbol;Acc:MGI:1343087] | 100,00 | 89,7 |
| ENSMUSG00000025208 | 705,0407019 | 632,6 | 649,9 | 0,031755192 | 0,96996707 | Mrpl43 | mitochondrial ribosomal protein L43 [Source:MGI Symbol;Acc:MGI:2137229] | 100,00 | 92,0 |
| ENSMUSG00000040722 | 451,2079207 | 569,4 | 648,5 | -0,051900705 | 0,97401114 | Scamp5 | secretory carrier membrane protein 5 [Source:MGI Symbol;Acc:MGI:1928948] | 99,94 | 93,3 |
| ENSMUSG00000092345 | 1185,196753 | 1798,7 | 646,8 | 1,825247257 | 0,10069837 | Gm20503 | predicted gene 20503 [Source:MGI Symbol;Acc:MGI:5141968] | 98,48 | 77,3 |
| ENSMUSG00000047514 | 470,7771108 | 558,8 | 646,6 | -0,050143836 | 0,98014335 | Tspyl1 | testis-specific protein, Y-encoded-like 1 [Source:MGI Symbol;Acc:MGI:1298395] | 88,77 | 81,9 |
| ENSMUSG00000032463 | 532,2494264 | 557,0 | 645,8 | -0,052923606 | 0,97175597 | Faim | Fas apoptotic inhibitory molecule [Source:MGI Symbol;Acc:MGI:1344387] | 97,97 | 89,5 |
| ENSMUSG00000022463 | 694,3534556 | 646,6 | 644,5 | 0,170621209 | 0,85088352 | Srebf2 | sterol regulatory element binding factor 2 [Source:MGI Symbol;Acc:MGI:107585] | 98,40 | 73,9 |
| ENSMUSG00000024403 | 534,7083258 | 647,1 | 643,6 | 0,292004486 | 0,83253327 | Atp6v1g2 | ATPase, H+ transporting, lysosomal V1 subunit G2 [Source:MGI Symbol;Acc:MGI:1913487] | 98,50 | 79,5 |
| ENSMUSG00000026309 | 701,0808549 | 509,1 | 643,6 | -0,321848391 | 0,69191811 | Ilkap | integrin-linked kinase-associated serine/threonine phosphatase 2C [Source:MGI Symbol;Acc:MGI:1914694] | 83,74 | 76,6 |
| ENSMUSG00000003199 | 985,2002284 | 1110,0 | 642,3 | 1,162677989 | 0,15777801 | Mpnd | MPN domain containing [Source:MGI Symbol;Acc:MGI:1915297] | 94,03 | 76,4 |
| ENSMUSG00000036752 | 1032,223383 | 1543,9 | 640,0 | 1,335907518 | 0,0577686 | Tubb4b | tubulin, beta 4B class IVB [Source:MGI Symbol;Acc:MGI:1915472] | 86,05 | 80,8 |
| ENSMUSG00000033863 | 1238,897495 | 2353,1 | 638,1 | 2,371896283 | 0,10365894 | Klf9 | Kruppel-like factor 9 [Source:MGI Symbol;Acc:MGI:1333856] | 100,00 | 73,3 |
| ENSMUSG00000040234 | 693,5950602 | 1043,4 | 637,6 | 1,077059885 | 0,51855632 | Tm7sf3 | transmembrane 7 superfamily member 3 [Source:MGI Symbol;Acc:MGI:1914873] | 99,58 | 71,5 |
| ENSMUSG00000061461 | 723,6767843 | 935,9 | 637,1 | 0,752423443 | 0,38226563 | Smim20 | small integral membrane protein 20 [Source:MGI Symbol;Acc:MGI:1913528] | 100,00 | 97,0 |
| ENSMUSG00000020993 | 762,86169 | 808,3 | 636,2 | 0,537689328 | 0,53099132 | Trappc6b | trafficking protein particle complex 6B [Source:MGI Symbol;Acc:MGI:1925482] | 95,53 | 87,5 |
| ENSMUSG00000028436 | 951,6574168 | 1063,8 | 633,1 | 0,793420863 | 0,30522282 | Dcaf12 | DDB1 and CUL4 associated factor 12 [Source:MGI Symbol;Acc:MGI:1916220] | 99,68 | 79,8 |
| ENSMUSG00000025337 | 739,9016732 | 1023,8 | 633,0 | 1,048699111 | 0,50700229 | Sbds | Shwachman-Bodian-Diamond syndrome homolog (human) [Source:MGI Symbol;Acc:MGI:1913961] | 78,73 | 73,6 |
| ENSMUSG00000047547 | 916,4920581 | 1316,6 | 632,6 | 1,222608453 | 0,18029514 | Cltb | clathrin, light polypeptide (Lcb) [Source:MGI Symbol;Acc:MGI:1921575] | 100,00 | 86,7 |
| ENSMUSG00000024873 | 529,6453828 | 571,3 | 629,7 | -0,007198066 | 0,99300766 | Cnih2 | cornichon family AMPA receptor auxiliary protein 2 [Source:MGI Symbol;Acc:MGI:1277225] | 100,00 | 99,2 |
| ENSMUSG00000052833 | 813,0837 | 962,7 | 626,0 | 0,619095962 | 0,62999835 | Sae1 | SUMO1 activating enzyme subunit 1 [Source:MGI Symbol;Acc:MGI:1929264] | 100,00 | 98,5 |
| ENSMUSG00000055681 | 841,1812737 | 1098,5 | 622,0 | 0,934782396 | 0,27387513 | Cope | coatomer protein complex, subunit epsilon [Source:MGI Symbol;Acc:MGI:1891702] | 93,30 | 83,4 |
| ENSMUSG00000001729 | 1095,883506 | 1218,2 | 621,8 | 1,059480244 | 0,14906418 | Akt1 | thymoma viral proto-oncogene 1 [Source:MGI Symbol;Acc:MGI:87986] | 86,79 | 77,9 |
| ENSMUSG00000026509 | 1664,425944 | 2807,6 | 620,2 | 2,244723775 | 0,13045627 | Capn2 | calpain 2 [Source:MGI Symbol;Acc:MGI:88264] | 100,00 | 79,4 |
| ENSMUSG00000061360 | 571,1928217 | 501,3 | 613,9 | -0,075845421 | 0,93424065 | Phf5a | PHD finger protein 5A [Source:MGI Symbol;Acc:MGI:2156864] | 100,00 | 97,3 |
| ENSMUSG00000020457 | 614,1088141 | 664,6 | 613,4 | 0,462911415 | 0,79641879 | Drg1 | developmentally regulated GTP binding protein 1 [Source:MGI Symbol;Acc:MGI:1343297] | 100,00 | 91,3 |
| ENSMUSG00000033763 | 1166,257293 | 1682,9 | 611,9 | 1,572899243 | 0,37324749 | Mtss1l | metastasis suppressor 1-like [Source:MGI Symbol;Acc:MGI:3039591] | 98,73 | 72,9 |
| ENSMUSG00000032046 | 945,2440031 | 1128,4 | 611,4 | 1,149909798 | 0,23630403 | Abhd12 | abhydrolase domain containing 12 [Source:MGI Symbol;Acc:MGI:1923442] | 99,35 | 83,1 |
| ENSMUSG00000020225 | 774,5009998 | 837,1 | 610,5 | 0,581758452 | 0,56950257 | Tmbim4 | transmembrane BAX inhibitor motif containing 4 [Source:MGI Symbol;Acc:MGI:1915462] | 82,59 | 71,5 |
| ENSMUSG00000030307 | 968,3411847 | 1347,4 | 609,6 | 1,255147007 | 0,31310081 | Slc6a11 | solute carrier family 6 (neurotransmitter transporter, GABA), member 11 [Source:MGI Symbol;Acc:MGI:95630] | 99,89 | 80,4 |
| ENSMUSG00000033998 | 1633,538081 | 2759,4 | 609,5 | 2,319346313 | 0,14139827 | Kcnk1 | potassium channel, subfamily K, member 1 [Source:MGI Symbol;Acc:MGI:109322] | 100,00 | 92,7 |
| ENSMUSG00000022677 | 868,0810453 | 1236,4 | 609,1 | 1,184829422 | 0,43857069 | Fopnl | Fgfr1op N-terminal like [Source:MGI Symbol;Acc:MGI:1913336] | 99,77 | 85,2 |
| ENSMUSG00000087260 | 906,612748 | 1255,8 | 607,9 | 1,072941501 | 0,14589725 | Lamtor5 | late endosomal/lysosomal adaptor, MAPK and MTOR activator 5 [Source:MGI Symbol;Acc:MGI:1915826] | 90,01 | 82,1 |
| ENSMUSG00000013822 | 703,9087446 | 784,6 | 607,6 | 0,52311033 | 0,45740211 | Elof1 | ELF1 homolog, elongation factor 1 [Source:MGI Symbol;Acc:MGI:1913376] | 79,37 | 78,9 |
| ENSMUSG00000020250 | 707,7463412 | 774,0 | 602,9 | 0,661126902 | 0,70836664 | Txnrd1 | thioredoxin reductase 1 [Source:MGI Symbol;Acc:MGI:1354175] | 99,76 | 70,0 |
| ENSMUSG00000005510 | 869,7514042 | 1149,4 | 602,6 | 0,996503204 | 0,20930183 | Ndufs3 | NADH dehydrogenase (ubiquinone) Fe-S protein 3 [Source:MGI Symbol;Acc:MGI:1915599] | 89,10 | 73,6 |
| ENSMUSG00000031007 | 708,3861067 | 649,0 | 598,5 | 0,049525287 | 0,96773906 | Atp6ap2 | ATPase, H+ transporting, lysosomal accessory protein 2 [Source:MGI Symbol;Acc:MGI:1917745] | 99,54 | 76,4 |
| ENSMUSG00000025868 | 1234,221851 | 1734,5 | 592,8 | 1,773874314 | 0,09068917 | Higd2a | HIG1 domain family, member 2A [Source:MGI Symbol;Acc:MGI:1914294] | 99,38 | 98,6 |
| ENSMUSG00000054405 | 1008,760925 | 1571,1 | 592,3 | 1,419739685 | 0,19618181 | Dnajc8 | DnaJ heat shock protein family (Hsp40) member C8 [Source:MGI Symbol;Acc:MGI:1915848] | 92,69 | 85,8 |
| ENSMUSG00000021178 | 767,3084058 | 967,6 | 590,2 | 0,84796036 | 0,26216087 | Psmc1 | protease (prosome, macropain) 26S subunit, ATPase 1 [Source:MGI Symbol;Acc:MGI:106054] | 100,00 | 95,5 |
| ENSMUSG00000028955 | 709,5534304 | 957,9 | 588,9 | 0,926686411 | 0,60525041 | Vamp3 | vesicle-associated membrane protein 3 [Source:MGI Symbol;Acc:MGI:1321389] | 99,73 | 89,0 |
| ENSMUSG00000026356 | 790,0296128 | 1012,1 | 588,8 | 0,829101305 | 0,32515295 | Dars | aspartyl-tRNA synthetase [Source:MGI Symbol;Acc:MGI:2442544] | 88,74 | 82,7 |

| | | | | | | | | | |
|---|---|---|---|---|---|---|---|---|---|
| ENSMUSG00000025630 | 527,3269349 | 523,9 | 587,5 | -0,023974757 | 0,98649897 | Hprt | hypoxanthine guanine phosphoribosyl transferase [Source:MGI Symbol;Acc:MGI:96217] | 99,95 | 92,8 |
| ENSMUSG00000026426 | 1158,150225 | 1849,8 | 583,5 | 1,679120985 | 0,30194106 | Arl8a | ADP-ribosylation factor-like 8A [Source:MGI Symbol;Acc:MGI:1915974] | 99,72 | 91,4 |
| ENSMUSG00000042396 | 884,8011275 | 1193,1 | 582,3 | 1,056730298 | 0,40725498 | Rbm7 | RNA binding motif protein 7 [Source:MGI Symbol;Acc:MGI:1914260] | 100,00 | 95,6 |
| ENSMUSG00000052889 | 587,7744697 | 642,0 | 581,2 | 0,080779172 | 0,95184066 | Prkcb | protein kinase C, beta [Source:MGI Symbol;Acc:MGI:97596] | 99,90 | 81,9 |
| ENSMUSG00000001313 | 830,81182 | 985,2 | 581,1 | 0,876705555 | 0,35505203 | Rnd2 | Rho family GTPase 2 [Source:MGI Symbol;Acc:MGI:1338755] | 98,90 | 89,6 |
| ENSMUSG00000025228 | 796,8035785 | 1004,1 | 579,8 | 0,946751719 | 0,25053486 | Actr1a | ARP1 actin-related protein 1A, centractin alpha [Source:MGI Symbol;Acc:MGI:1858964] | 100,00 | 86,1 |
| ENSMUSG00000079659 | 612,4660911 | 664,5 | 576,0 | 0,25614775 | 0,74963322 | Tmem243 | transmembrane protein 243, mitochondrial [Source:MGI Symbol;Acc:MGI:3606159] | 86,45 | 83,1 |
| ENSMUSG00000032299 | 568,4705183 | 613,3 | 575,2 | 0,091769113 | 0,90535824 | Commd4 | COMM domain containing 4 [Source:MGI Symbol;Acc:MGI:1913449] | 100,00 | 100,0 |
| ENSMUSG00000021241 | 1004,968267 | 1390,8 | 574,9 | 1,450027689 | 0,10147572 | Isca2 | iron-sulfur cluster assembly 2 [Source:MGI Symbol;Acc:MGI:1921566] | 99,92 | 97,5 |
| ENSMUSG00000026203 | 594,1104154 | 806,2 | 572,2 | 0,519599334 | 0,58951522 | Dnajb2 | DnaJ heat shock protein family (Hsp40) member B2 [Source:MGI Symbol;Acc:MGI:1928739] | 98,95 | 71,7 |
| ENSMUSG00000027404 | 626,200628 | 815,9 | 572,2 | 0,944453267 | 0,34367811 | Snrpb | small nuclear ribonucleoprotein B [Source:MGI Symbol;Acc:MGI:98342] | 99,02 | 93,3 |
| ENSMUSG00000003363 | 535,0442987 | 526,7 | 571,4 | 0,075741294 | 0,95557263 | Pld3 | phospholipase D family, member 3 [Source:MGI Symbol;Acc:MGI:1333782] | 92,11 | 77,0 |
| ENSMUSG00000063787 | 665,8876808 | 832,7 | 571,3 | 1,020031103 | 0,28162657 | Chchd1 | coiled-coil-helix-coiled-coil-helix domain containing 1 [Source:MGI Symbol;Acc:MGI:1913371] | 99,60 | 95,5 |
| ENSMUSG00000078656 | 692,247027 | 859,4 | 566,3 | 0,637571556 | 0,63598748 | Vps25 | vacuolar protein sorting 25 (yeast) [Source:MGI Symbol;Acc:MGI:106354] | 97,02 | 74,8 |
| ENSMUSG00000018585 | 904,8127129 | 1220,0 | 566,2 | 1,214858764 | 0,2121972 | Atox1 | antioxidant 1 copper chaperone [Source:MGI Symbol;Acc:MGI:1333855] | 98,40 | 89,2 |
| ENSMUSG00000020132 | 936,1918314 | 1416,2 | 563,2 | 1,464074951 | 0,10635458 | Rab21 | RAB21, member RAS oncogene family [Source:MGI Symbol;Acc:MGI:894308] | 98,90 | 87,6 |
| ENSMUSG00000037257 | 531,5723727 | 608,9 | 560,9 | 0,675014361 | 0,51912671 | Aagab | alpha- and gamma-adaptin binding protein [Source:MGI Symbol;Acc:MGI:1914189] | 100,00 | 72,3 |
| ENSMUSG00000038722 | 675,889953 | 973,5 | 559,9 | 0,907959619 | 0,23777049 | Bud31 | BUD31 homolog [Source:MGI Symbol;Acc:MGI:2141291] | 100,00 | 95,2 |
| ENSMUSG00000028932 | 677,0902185 | 787,3 | 557,8 | 0,597994929 | 0,3830816 | Psmc2 | proteasome (prosome, macropain) 26S subunit, ATPase 2 [Source:MGI Symbol;Acc:MGI:109555] | 85,53 | 79,8 |
| ENSMUSG00000015095 | 707,4310207 | 1041,2 | 556,1 | 1,147804722 | 0,44658293 | Fbxw5 | F-box and WD-40 domain protein 5 [Source:MGI Symbol;Acc:MGI:1354731] | 99,13 | 88,1 |
| ENSMUSG00000090841 | 931,5964755 | 1168,7 | 555,4 | 1,123544899 | 0,06121894 | Myl6 | myosin, light polypeptide 6, alkali, smooth muscle and non-muscle [Source:MGI Symbol;Acc:MGI:109318] | 96,14 | 76,4 |
| ENSMUSG00000048307 | 1028,106195 | 1368,7 | 550,1 | 1,560511993 | 0,05178385 | Ankrd46 | ankyrin repeat domain 46 [Source:MGI Symbol;Acc:MGI:1916089] | 99,97 | 88,5 |
| ENSMUSG00000033020 | 609,0661659 | 848,7 | 549,6 | 0,785330569 | 0,38757395 | Polr2f | polymerase (RNA) II (DNA directed) polypeptide F [Source:MGI Symbol;Acc:MGI:1349393] | 100,00 | 99,0 |
| ENSMUSG00000029577 | 734,9510791 | 956,5 | 548,3 | 1,026705783 | 0,20906365 | Ube3b | ubiquitin protein ligase E3B [Source:MGI Symbol;Acc:MGI:1891295] | 98,74 | 73,1 |
| ENSMUSG00000022536 | 571,250139 | 863,5 | 546,6 | 0,664600266 | 0,72815937 | Glyr1 | glyoxylate reductase 1 homolog (Arabidopsis) [Source:MGI Symbol;Acc:MGI:1921272] | 99,17 | 85,4 |
| ENSMUSG00000007564 | 1039,956283 | 1664,9 | 541,7 | 1,62316673 | 0,20859346 | Ppp2r1a | protein phosphatase 2, regulatory subunit A, alpha [Source:MGI Symbol;Acc:MGI:1926334] | 96,67 | 90,4 |
| ENSMUSG00000021476 | 707,8176408 | 762,0 | 538,6 | 1,388318842 | 0,21265562 | Habp4 | hyaluronic acid binding protein 4 [Source:MGI Symbol;Acc:MGI:1891713] | 100,00 | 73,8 |
| ENSMUSG00000032038 | 706,8879243 | 984,0 | 538,3 | 0,918379326 | 0,66477451 | St3gal4 | ST3 beta-galactoside alpha-2,3-sialyltransferase 4 [Source:MGI Symbol;Acc:MGI:1316743] | 99,97 | 91,4 |
| ENSMUSG00000049299 | 541,6630225 | 639,8 | 536,7 | 0,535965743 | 0,57590342 | Trappc1 | trafficking protein particle complex 1 [Source:MGI Symbol;Acc:MGI:1098727] | 100,00 | 89,5 |
| ENSMUSG00000074227 | 590,701095 | 540,6 | 534,2 | -0,489633487 | 0,70773007 | Spint2 | serine protease inhibitor, Kunitz type 2 [Source:MGI Symbol;Acc:MGI:1338031] | 100,00 | 86,1 |
| ENSMUSG00000052296 | 908,0168157 | 1474,2 | 533,5 | 1,555343199 | 0,2998204 | Ppp6r1 | protein phosphatase 6, regulatory subunit 1 [Source:MGI Symbol;Acc:MGI:2442163] | 99,22 | 83,8 |
| ENSMUSG00000027502 | 539,8776715 | 627,9 | 533,3 | 0,325146342 | 0,83393207 | Rtfdc1 | replication termination factor 2 domain containing 1 [Source:MGI Symbol;Acc:MGI:1913654] | 99,60 | 92,4 |
| ENSMUSG00000049960 | 475,8313761 | 507,6 | 533,3 | -0,050985443 | 0,95246807 | Mrps16 | mitochondrial ribosomal protein S16 [Source:MGI Symbol;Acc:MGI:1913492] | 100,00 | 99,2 |
| ENSMUSG00000031954 | 927,0509509 | 1193,5 | 531,6 | 1,727300755 | 0,0582813 | Cfdp1 | craniofacial development protein 1 [Source:MGI Symbol;Acc:MGI:1344403] | 100,00 | 81,3 |
| ENSMUSG00000032526 | 583,3667302 | 705,7 | 531,6 | 0,503668981 | 0,50715321 | Deb1 | differentially expressed in B16F10 1 [Source:MGI Symbol;Acc:MGI:1349474] | 100,00 | 99,9 |
| ENSMUSG00000028013 | 841,1090583 | 1176,3 | 530,0 | 1,302064429 | 0,0686533 | Ppa2 | pyrophosphatase (inorganic) 2 [Source:MGI Symbol;Acc:MGI:1922026] | 98,26 | 81,9 |
| ENSMUSG00000072694 | 446,8259207 | 549,1 | 528,3 | 0,265239441 | 0,85916323 | 1500011B03Rik | RIKEN cDNA 1500011B03 gene [Source:MGI Symbol;Acc:MGI:1913486] | 99,54 | 87,6 |
| ENSMUSG00000039660 | 446,2668509 | 510,3 | 527,5 | -0,001902336 | 0,99794692 | D2Wsu81e | DNA segment, Chr 2, Wayne State University 81, expressed [Source:MGI Symbol;Acc:MGI:106544] | 100,00 | 96,8 |
| ENSMUSG00000022681 | 991,972749 | 1184,1 | 526,7 | 1,599588663 | 0,06479287 | Ntan1 | N-terminal Asn amidase [Source:MGI Symbol;Acc:MGI:108471] | 91,89 | 78,8 |
| ENSMUSG00000030615 | 530,8247685 | 597,7 | 524,0 | 1,36032677 | 0,2091015 | Tmem126a | transmembrane protein 126A [Source:MGI Symbol;Acc:MGI:1913521] | 98,66 | 78,5 |
| ENSMUSG00000025035 | 1036,647337 | 1635,7 | 522,5 | 1,79297426 | 0,10393512 | Arl3 | ADP-ribosylation factor-like 3 [Source:MGI Symbol;Acc:MGI:1929699] | 100,00 | 96,6 |
| ENSMUSG00000022200 | 1452,881031 | 2362,7 | 520,6 | 2,198542261 | 0,09549981 | Golph3 | golgi phosphoprotein 3 [Source:MGI Symbol;Acc:MGI:1913879] | 100,00 | 89,8 |
| ENSMUSG00000020780 | 570,2360766 | 501,5 | 517,9 | 0,236775482 | 0,87283104 | Srp68 | signal recognition particle 68 [Source:MGI Symbol;Acc:MGI:1917447] | 96,61 | 79,4 |
| ENSMUSG00000006575 | 665,6610963 | 824,6 | 517,2 | 0,718312365 | 0,64304439 | Rundc3a | RUN domain containing 3A [Source:MGI Symbol;Acc:MGI:1858752] | 98,59 | 80,2 |
| ENSMUSG00000028394 | 528,6646787 | 503,7 | 514,5 | 0,187258714 | 0,82822972 | Pole3 | polymerase (DNA directed), epsilon 3 (p17 subunit) [Source:MGI Symbol;Acc:MGI:1933378] | 99,82 | 91,4 |
| ENSMUSG00000024911 | 533,8695108 | 648,4 | 513,9 | 0,513510578 | 0,60730301 | Fibp | fibroblast growth factor (acidic) intracellular binding protein [Source:MGI Symbol;Acc:MGI:1926233] | 99,76 | 95,9 |
| ENSMUSG00000061111 | 808,394451 | 998,4 | 513,3 | 1,085806989 | 0,1541188 | Fam195b | family with sequence similarity 195, member B [Source:MGI Symbol;Acc:MGI:2384752] | 99,86 | 96,5 |
| ENSMUSG00000020089 | 559,250112 | 581,6 | 510,5 | 0,854762114 | 0,44333449 | Ppa1 | pyrophosphatase (inorganic) 1 [Source:MGI Symbol;Acc:MGI:97831] | 98,44 | 83,0 |
| ENSMUSG00000031983 | 1068,757956 | 1721,0 | 509,4 | 1,750823736 | 0,05486295 | 2310022B05Rik | RIKEN cDNA 2310022B05 gene [Source:MGI Symbol;Acc:MGI:1916801] | 97,57 | 87,9 |
| ENSMUSG00000021114 | 965,2601584 | 1519,7 | 509,2 | 1,580971575 | 0,05686833 | Atp6v1d | ATPase, H+ transporting, lysosomal V1 subunit D [Source:MGI Symbol;Acc:MGI:1921084] | 98,95 | 91,4 |
| ENSMUSG00000058833 | 563,524934 | 683,1 | 508,4 | 0,536340136 | 0,45621176 | 2810428I15Rik | RIKEN cDNA 2810428I15 gene [Source:MGI Symbol;Acc:MGI:1913712] | 100,00 | 92,9 |
| ENSMUSG00000002660 | 958,8031551 | 1300,6 | 508,4 | 1,543410422 | 0,08437679 | Clpp | caseinolytic mitochondrial matrix peptidase proteolytic subunit [Source:MGI Symbol;Acc:MGI:1858213] | 95,03 | 78,9 |
| ENSMUSG00000005881 | 759,7580324 | 895,5 | 508,1 | 1,444141926 | 0,11201444 | Ergic3 | ERGIC and golgi 3 [Source:MGI Symbol;Acc:MGI:1913616] | 96,57 | 76,7 |
| ENSMUSG00000057649 | 443,0451611 | 505,9 | 506,8 | 0,355342226 | 0,74202346 | Brd9 | bromodomain containing 9 [Source:MGI Symbol;Acc:MGI:2145317] | 99,76 | 82,2 |
| ENSMUSG00000022951 | 485,3669667 | 643,6 | 505,2 | 0,573523972 | 0,74161886 | Rcan1 | regulator of calcineurin 1 [Source:MGI Symbol;Acc:MGI:1890564] | 99,91 | 84,9 |
| ENSMUSG00000025066 | 851,2825215 | 998,9 | 503,6 | 1,142871687 | 0,23621324 | Sfr1 | SWI5 dependent recombination repair 1 [Source:MGI Symbol;Acc:MGI:1915038] | 99,60 | 78,7 |
| ENSMUSG00000041203 | 553,0638638 | 595,5 | 501,5 | 0,410358864 | 0,62045182 | 2310036O22Rik | RIKEN cDNA 2310036O22 gene [Source:MGI Symbol;Acc:MGI:1922833] | 100,00 | 96,7 |
| ENSMUSG00000014606 | 896,7645382 | 1220,6 | 500,7 | 1,391803746 | 0,10857141 | Slc25a11 | solute carrier family 25 (mitochondrial carrier oxoglutarate carrier), member 11 [Source:MGI Symbol;Acc:MGI:1915113] | 96,86 | 81,5 |

Table S2.7

| Id | baseMean Read | log2FoldChange (Whole astrocyte/PAP) | padj | Associated Gene Name | Description | Exon rank | Exon count per transcript |
| --- | --- | --- | --- | --- | --- | --- | --- |
| **ENSMUSE00000486160** | **15988,91806** | **1,665273322** | **0,0003** | **Gfap alpha** | **acidic protein [Source:MGI Symbol;Ac** | **9** | **9** |
| ENSMUSE00000908554 | 5004,257074 | 1,607847563 | 0,0006 | Gfap | acidic protein [Source:MGI Symbol;Acc | 1 | 8 |
| ENSMUSE00000513876 | 5002,826012 | 1,60737515 | 0,0006 | Gfap | acidic protein [Source:MGI Symbol;Acc | 1 | 9 |
| ENSMUSE00000576526 | 4150,587202 | 1,70266312 | 0,0006 | Gfap | acidic protein [Source:MGI Symbol;Acc | 6 | 8 |
| ENSMUSE00000363222 | 2868,240771 | 1,70950039 | 0,0006 | Gfap | acidic protein [Source:MGI Symbol;Acc | 4 | 8 |
| **ENSMUSE00000647296** | **2526,938454** | **2,912321883** | **0,0014** | **Gfap delta** | **acidic protein [Source:MGI Symbol;Ac** | **8** | **8** |
| ENSMUSE00000113329 | 2261,223046 | 1,758704902 | 0,0005 | Gfap | acidic protein [Source:MGI Symbol;Acc | 5 | 8 |
| ENSMUSE00001246067 | 2011,87202 | 1,638915818 | 0,0007 | Gfap | acidic protein [Source:MGI Symbol;Acc | 3 | 8 |
| ENSMUSE00000113335 | 1954,930859 | 1,690484808 | 0,0006 | Gfap | acidic protein [Source:MGI Symbol;Acc | 8 | 9 |
| ENSMUSE00000113334 | 1884,880225 | 1,700502479 | 0,0006 | Gfap | acidic protein [Source:MGI Symbol;Acc | 7 | 8 |
| ENSMUSE00001268811 | 1581,194699 | 1,733062847 | 0,0004 | Gfap | acidic protein [Source:MGI Symbol;Acc | 2 | 8 |

Table S3

| Gene name | Astro | | | | | PAP | | | | | p-value | significance |
|---|---|---|---|---|---|---|---|---|---|---|---|---|
| | Astro 1 | Astro 2 | Astro 3 | Mean | SEM | PAP 1 | PAP 2 | PAP 3 | Mean | SEM | | |
| Fth1 | 4,55 | 1,29 | 3,45 | 3,10 | 0,96 | 63,67 | 81,66 | 19,86 | 55,06 | 18,35 | 0,05 | * |
| Ftl1 | 0,13 | 0,05 | 0,11 | 0,10 | 0,02 | 95,14 | 75,54 | 15,76 | 62,15 | 23,87 | 0,05 | * |
| Ezr | 0,06 | 0,04 | 0,2 | 0,10 | 0,05 | 11,78 | 15,6 | 3,15 | 10,18 | 3,68 | 0,05 | * |
| Gnb2l1 | 0,45 | 0,58 | 0,97 | 0,67 | 0,16 | 125,7 | 87,13 | 42,08 | 84,97 | 24,16 | 0,05 | * |
| Rpl8 | 1,54 | 0,94 | 1,72 | 1,40 | 0,24 | 168,06 | 102,36 | 48,8 | 106,41 | 34,49 | 0,05 | * |
| Eef1a1 | 4,08 | 3,76 | 5,19 | 4,34 | 0,43 | 26667,84 | 309,58 | 13384,88 | 13454,10 | 7609,05 | 0,05 | * |
| Eef2 | 0,92 | 0,64 | 1,51 | 1,02 | 0,26 | 120,34 | 120,7 | 89,12 | 110,05 | 10,47 | 0,05 | * |
| Rdx | 0 | 0,03 | 0,05 | 0,03 | 0,01 | 7,02 | 9,29 | 2,51 | 6,27 | 1,99 | 0,05 | * |
| Rplp1 | 2,85 | 3,3 | 6,61 | 4,25 | 1,19 | 299,1 | 296,49 | 72,7 | 222,76 | 75,04 | 0,05 | * |
| Rpl4 | 0,82 | 0,92 | 1,58 | 1,11 | 0,24 | 257,02 | 56,67 | 300 | 204,56 | 74,98 | 0,05 | * |
| Rps14 | 0,08 | 0,06 | 0,18 | 0,11 | 0,04 | 3,6 | 2,93 | 1,8 | 2,78 | 0,53 | 0,05 | * |
| Rps3 | 0,9 | 0,87 | 1,76 | 1,18 | 0,29 | 220,49 | 178,38 | 38,69 | 145,85 | 54,94 | 0,05 | * |
| Ccnd2 | 0,08 | 0,4 | 0,4 | 0,29 | 0,11 | 74,3 | 71,71 | 37,73 | 61,25 | 11,78 | 0,05 | * |
| Hsp90 | 1,08 | 0,75 | 0,98 | 0,94 | 0,10 | 70,97 | 48,28 | 19,9 | 46,38 | 14,77 | 0,05 | * |
| Mdm2 | 0,02 | 0,02 | 0,01 | 0,02 | 0,00 | 24,36 | 17,61 | 9,05 | 17,01 | 4,43 | 0,05 | * |
| Ncl | 0,05 | 0,15 | 0,15 | 0,12 | 0,03 | 106,9 | 92,14 | 22,74 | 73,93 | 25,95 | 0,05 | * |
| Npm1 | 0,6 | 0,14 | 0,26 | 0,33 | 0,14 | 95,97 | 87,93 | 31,31 | 71,74 | 20,35 | 0,05 | * |

Table S4

| Gene name | PvAP | | | | | PAP | | | | | p-value | significance | Enrichment |
|---|---|---|---|---|---|---|---|---|---|---|---|---|---|
| | PvAP 1 | PvAP 2 | PvAP 3 | Mean | SEM | PAP 1 | PAP 2 | PAP 3 | Mean | SEM | | | |
| Ezr | 1,42 | 1,61 | 1,29 | 1,44 | 0,09 | 3,64 | 3,00 | 3,15 | 3,26 | 0,19 | 0,05 | * | PAP |
| Fth1 | 4,33 | 4,11 | 6,14 | 4,86 | 0,64 | 8,33 | 7,67 | 18,40 | 11,47 | 3,47 | 0,05 | * | PAP |
| Rpl1 | 6,18 | 4,94 | 4,78 | 5,30 | 0,44 | 7,56 | 10,79 | 14,38 | 10,91 | 1,97 | 0,05 | * | PAP |
| Ftl1 | 1,45 | 1,20 | 0,64 | 1,10 | 0,24 | 0,64 | 0,99 | 3,36 | 1,66 | 0,85 | 0,5 | ns | None |
| Gnb2l1 | 2,06 | 1,92 | 0,90 | 1,63 | 0,37 | 1,18 | 0,76 | 3,96 | 1,97 | 1,00 | 0,5 | ns | None |
| Hsp90 | 6,17 | 4,54 | 4,56 | 5,09 | 0,54 | 1,51 | 0,43 | 9,32 | 3,75 | 2,80 | 0,35 | ns | None |
| Ncl | 0,62 | 1,35 | 1,01 | 0,99 | 0,21 | 0,47 | 0,31 | 2,86 | 1,21 | 0,82 | 0,35 | ns | None |
| Npm | 2,25 | 7,22 | 1,96 | 3,81 | 1,71 | 3,06 | 0,98 | 5,39 | 3,14 | 1,27 | 0,5 | ns | None |
| Rdx | 2,24 | 15,17 | 3,14 | 6,85 | 4,17 | 3,00 | 1,21 | 5,80 | 3,34 | 1,34 | 0,35 | ns | None |
| Eef2 | 13,48 | 43,79 | 11,03 | 22,77 | 10,54 | 18,75 | 5,56 | 27,18 | 17,16 | 6,29 | 0,5 | ns | None |
| Rpl4 | 0,89 | 1,57 | 1,44 | 1,30 | 0,21 | 0,71 | 0,43 | 4,46 | 1,87 | 1,30 | 0,5 | ns | None |
| Rpl8 | 5,90 | 3,51 | 2,00 | 3,80 | 1,14 | 2,65 | 2,10 | 11,81 | 5,52 | 3,15 | 0,5 | ns | None |
| Rps14 | 0,89 | 1,57 | 1,44 | 1,30 | 0,21 | 0,71 | 0,43 | 4,46 | 1,87 | 1,30 | 0,35 | ns | None |
| Rps3 | 0,17 | 0,12 | 0,05 | 0,11 | 0,03 | 0,17 | 0,22 | 1,22 | 0,54 | 0,34 | 0,1 | ns | None |

Table S5

| Gene name | PAP Control | | | | | | | PAP Fear | | | | | | p-value | significance |
|---|---|---|---|---|---|---|---|---|---|---|---|---|---|---|---|
| | C1 | C2 | C3 | C4 | C5 | Mean | SEM | F1 | F2 | F3 | F4 | Mean | SEM | | |
| eEF1A1 | 1 | 0,9 | 0,9 | 0,9 | 1,1 | 0,96 | 0,04 | 1,2 | 1,4 | 1,6 | 1,6 | 1,45 | 0,10 | 0,0011 | ** |
| CCND2 | 2 | 2,1 | 1,8 | 1,2 | 1,3 | 1,68 | 0,18 | 1,7 | 2,1 | 1,4 | 1,3 | 1,63 | 0,18 | 0,8388 | ns |
| FTH1/FTL1 | 0,89 | 0,74 | 0,83 | 1,3 | | 0,94 | 0,12 | 0,75 | 0,62 | 0,62 | 0,41 | 0,60 | 0,07 | 0,0543 | ns |
| RACK1 | 7,77 | 8,32 | 6,39 | 4,36 | | 6,71 | 0,88 | 3,57 | 4,79 | 5,24 | 3,65 | 4,31 | 0,42 | 0,0493 | * |
| MDM2 | 0,56 | 0,44 | 0,49 | 0,44 | 0,52 | 0,49 | 0,02 | 0,81 | 0,71 | 0,92 | 0,98 | 0,86 | 0,06 | 0,0004 | *** |

Table S6

| Gene name | Density | | | | | | Distribution in Soma | | | | | | Distribution in Large Processes | | | | | | Distribution in Fine Processes | | | | | |
|---|---|---|---|---|---|---|---|---|---|---|---|---|---|---|---|---|---|---|---|---|---|---|---|---|
| | Control | | Fear | | p-value | Significance | Control | | Fear | | p-value | Significance | Control | | Fear | | p-value | Significance | Control | | Fear | | p-value | Significance |
| | Mean | SEM | Mean | SEM | | | Mean | SEM | Mean | SEM | | | Mean | SEM | Mean | SEM | | | Mean | SEM | Mean | SEM | | |
| Ccnd2 | 0,0064 | 0,0008 | 0,0070 | 0,0007 | <0,0001 | **** | 7,7220 | 0,5279 | 7,3100 | 0,6611 | 0,1500 | ns | 16,21 | 1,001 | 12,82 | 0,6692 | 0,0024 | ** | 76,07 | 1,179 | 79,87 | 0,9033 | 0,0008 | *** |
| Ftl1 | 0,0128 | 0,0011 | 0,01 | 0,0005 | 0,0282 | * | 10,61 | 0,7933 | 12,57 | 0,7321 | 0,0719 | ns | 8,83 | 0,6126 | 10,89 | 0,7023 | 0,0288 | * | 80,56 | 0,896 | 76,54 | 0,9856 | 0,0037 | ** |
| Fth1 | 0,1024 | 0,0039 | 0,0926 | 0,0067 | 0,2 | ns | 3,883 | 0,2967 | 6,98 | 0,5634 | <0,0001 | **** | 8,678 | 0,5576 | 10,47 | 0,7563 | 0,05 | ns | 87,44 | 0,7392 | 82,55 | 0,9345 | <0,0001 | **** |
| Eef1a1 | 0,0593 | 0,0031 | 0,0455 | 0,002 | 0,0007 | *** | 7,636 | 0,5853 | 7,15 | 0,6849 | 0,5911 | ns | 14,41 | 0,5538 | 13,28 | 0,5514 | 0,1621 | ns | 77,96 | 0,8681 | 79,57 | 0,9426 | 0,2158 | ns |
| Mdm2 | 0,007 | 0,0007 | 0,0028 | 0,0002 | <0,0001 | **** | 6,366 | 0,5433 | 7,743 | 0,781 | 0,1504 | ns | 11,9 | 0,9328 | 16,33 | 1,084 | 0,0024 | ** | 81,77 | 1,025 | 75,93 | 1,362 | 0,0008 | *** |
| Gnb2l1 | 0,008 | 0,0006 | 0,0066 | 0,0003 | 0,0327 | * | 8,517 | 0,6932 | 13,16 | 1,102 | 0,0006 | *** | 11,64 | 0,9398 | 15,92 | 0,9353 | 0,0017 | ** | 79,85 | 1,339 | 70,91 | 1,572 | <0,0001 | **** |

Table S7

| Gene name | PAP Control | | | | | PAP Fear | | | | | | | p-value | significance |
|---|---|---|---|---|---|---|---|---|---|---|---|---|---|---|
| | C1 | C2 | C3 | Mean | SEM | F1 | F2 | F3 | F4 | F5 | Mean | SEM | | |
| Eef1a1 | 26667,8 | 309,6 | 13384,9 | 13454,1 | 7609,1 | 27,8 | 17,1 | 6,1 | 11,1 | 9,6 | 14,3 | 3,8 | 1,00E-04 | *** |
| Ccnd2 | 74,3 | 71,7 | 37,7 | 61,2 | 11,8 | 0,9 | 0,6 | 0,3 | 0,3 | 0,2 | 0,5 | 0,1 | 0,02 | * |
| Fth1 | 63,7 | 81,7 | 19,9 | 55,1 | 18,4 | 16,8 | 17,1 | 9,7 | 17,3 | 12,4 | 14,7 | 1,5 | 0,02 | * |
| Ftl1 | 95,1 | 75,5 | 15,8 | 62,1 | 23,9 | 1,5 | 1,7 | 1,1 | 1,3 | 0,9 | 1,3 | 0,1 | 0,02 | * |
| Gnb2l1 | 125,7 | 87,1 | 42,1 | 85,0 | 24,2 | 1,5 | 2,1 | 1,1 | 2,0 | 2,6 | 1,9 | 0,3 | 0,02 | * |
| Mdm2 | 24,4 | 17,6 | 9,1 | 17,0 | 4,4 | 0,1 | 0,0 | 0,0 | 0,1 | 0,1 | 0,1 | 0,0 | 0,02 | * |

| Gene name | Astro Control | | | | | Astro Fear | | | | | | p-value | significance |
|---|---|---|---|---|---|---|---|---|---|---|---|---|---|
| | C1 | C2 | C3 | Mean | SEM | F1 | F2 | F3 | F4 | Mean | SEM | | |
| Eef1a1 | 4,1 | 3,8 | 5,2 | 4,34 | 0,43 | 2,99 | 3,18 | 2,76 | 3,12 | 3,01 | 0,11 | 0,03 | * |
| Ccnd2 | 0,08 | 0,4 | 0,4 | 0,29 | 0,11 | 0,08 | 0,12 | 0,18 | 0,20 | 0,15 | 0,03 | 0,26 | ns |
| Fth1 | 4,55 | 1,29 | 3,45 | 3,10 | 0,96 | 2,43 | 1,21 | 2,72 | 3,03 | 2,35 | 0,46 | 0,20 | ns |
| Ftl1 | 0,13 | 0,05 | 0,11 | 0,10 | 0,02 | 0,09 | 0,13 | 0,24 | 0,12 | 0,15 | 0,04 | 0,23 | ns |
| Gnb2l1 | 0,45 | 0,58 | 0,97 | 0,67 | 0,16 | 0,77 | 0,67 | 0,41 | 0,72 | 0,64 | 0,09 | 0,50 | ns |
| Mdm2 | 0,02 | 0,02 | 0,01 | 0,02 | 0,00 | 0,04 | 0,01 | 0,00 | 0,00 | 0,01 | 0,01 | 0,20 | ns |
